# Occam’s bias undermines inferences from phylogenetic linear models

**DOI:** 10.64898/2026.02.06.704358

**Authors:** Jacinta Guirguis, Luke E.B. Goodyear, Daniel Pincheira-Donoso

## Abstract

Phylogenetic modelling has consolidated as the analytical standard to address hypotheses about the patterns and dynamics of biodiversity in inter-specific contexts. These analyses are traditionally performed implementing phylogenetic linear models where single outcomes are regressed against multiple predictors without explicitly modelling the relationships amongst predictors. A prevailing, yet largely overlooked consequence of neglecting these relationships is what we introduce as ‘Occam’s bias’ – a statistical distortion arising where the model has fewer cause-effect connections than predicted by theory. Here, we propose that Occam’s bias is likely to have impacted a wide range of inferences about ecological and evolutionary processes made from phylogenetic linear models across the literature, and thus, that the adoption of approaches to address this bias are critical. We present an empirical test of the long-standing hypothesis that interspecific variation in life-history traits influences the likelihood of extinction risk across 13,949 species of terrestrial vertebrates to show the impacts of Occam’s bias in phenomenological inference. Our study calls for a re-evaluation of hypotheses tested using the traditional linear modelling structure and advocate the use and continued development of multi-response model structures that account for all causal pathways in phylogenetic analyses.

## Introduction

Phylogenetic linear modelling has consolidated as a statistical paradigm for testing hypotheses on the factors underlying evolutionary adaptation and the mechanisms behind large-scale patterns of biodiversity (1,2). These phylogenetic methods search for signals in the patterns of correlations among the characteristics of lineages both within and among clades, which indicate how characteristics of organisms have evolved in response to factors predicted to influence evolutionary change (3). To address these questions, studies have traditionally relied on multiple linear regression, where a single response is regressed on multiple predictors (single-response models, SR hereafter) (4). Such methods are widely available as packages implemented in the R software and include phylogenetic generalist least squares (*caper*), phylogenetic generalised linear model *(phylolm*) (5,6), as well as Bayesian approaches (*MCMCglmm* and *brms*) (7,8), among others. However, reliable inferences on the factors underlying the direction of evolutionary change and other phenomena, such as extinctions, require a rigorous assessment of the influences that multiple predictors can have when performing phylogenetic models. To address this issue, we propose ‘Occam’s bias’ –a statistical distortion that expresses in coefficients when a statistical model has fewer cause-effect connections than predicted by theory. We argue that overlooking Occam’s bias is likely to have impacted a potentially considerable fraction of inferences from tests of ecological and evolutionary processes appeared in the literature.

The rationale underlying the proposition that Occam’s bias impacts inferences from phylogenetic linear modelling arises from three main problems. First, predictions derived from hypotheses about ecological and evolutionary processes often include feedback cycles and bidirectional causal paths, which prevent the establishment of the clear, temporal ordering required to infer cause and effect, thus violating the assumptions of modern empirical causal inference for observational studies (9,10). Second, while no statistical model is infallible to establish cause-effect, the use of SR models can mislead interpretations when empirically testing theory (Fig 1). In the context of hypothesis testing in phylogenetics using SR models, the statistical model often contains fewer cause-effect connections than those derived from the scientific theory (Fig 1). This is because SR models cannot model the relationships among predictors, and thus, they fail to consider the influence of the immense range of factors that drive trait interactions such as (but not limited to) genetic correlations, pleiotropic effects and extragenetic inheritance. For instance, divergent natural selection towards different body size optima may result due to competition for ecological niches (11). At the same time, selection towards an optimal number of offspring may indirectly influence the evolution of body size (12), while environmental features also influence body size via natural selection for enhanced thermoregulatory performance (13,14). Consequently, the multiple paths that interact simultaneously during (e.g. phenotypic) evolutionary change (Fig 1a), can often fail to be realistically accounted for (or accounted for at all) by SR approaches (Fig 1b) as these do not model the relationships among predictors. Third, there is a misleading approach often applied in phylogenetic linear modelling in which more ‘control’ –achieved by the addition of covariates to models – is traditionally considered to improve rather than worsen inferences. Yet, no such general rule exists in statistics (15–17), and the addition of covariates may in fact worsen inferences by increasing Occam’s bias. As the number of explanatory variable increases linearly, the number of unaccounted, indirect paths grows quadratically (given by the function *y* = (*x*^2^-*x*)/2, defined for positive integers) (Fig 2), leading to biased coefficients. This is Occam’s bias, and the additional explanatory variables distort effect sizes if their relationships amongst each other are not accounted for. Consequently, we argue that the widespread implementation of phylogenetic linear models which has contributed to shape our interpretations of a range of phenomena about the evolution and extinction of biodiversity are likely to be heavily impacted by Occam’s bias.

**Fig 1.**
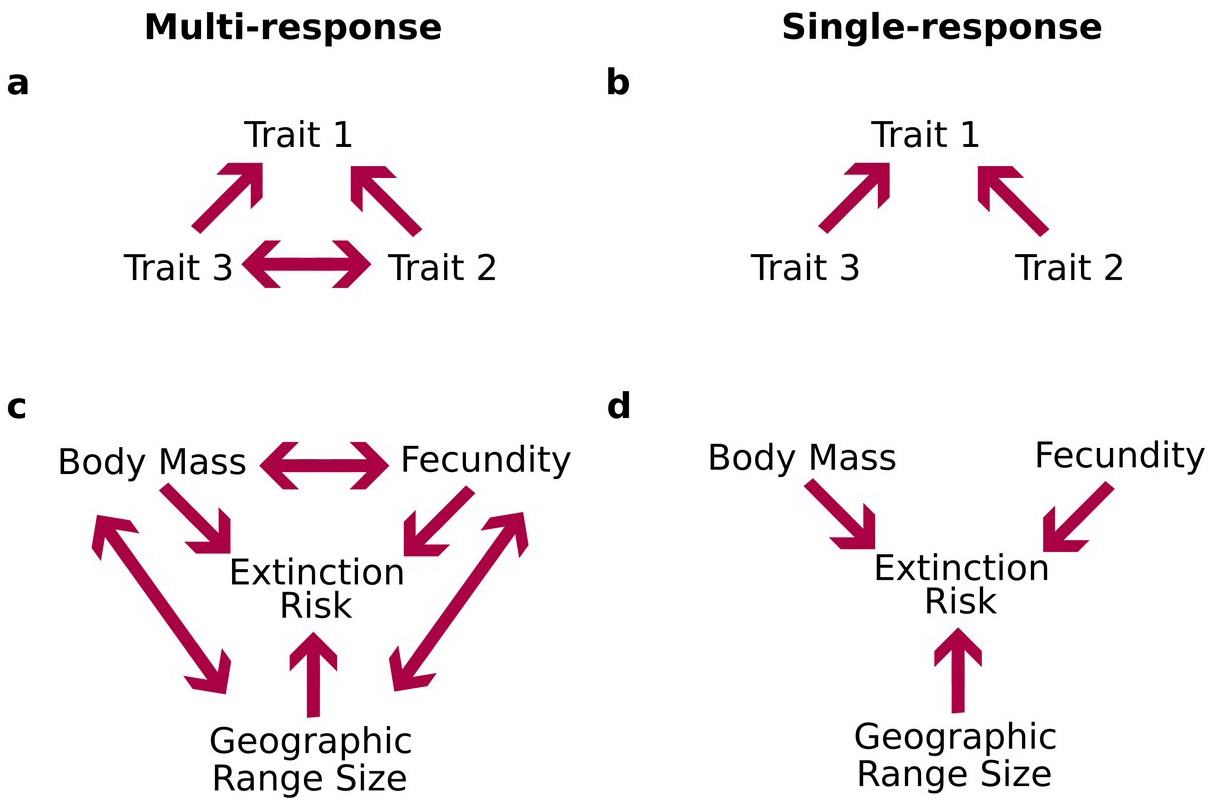
Causal relationships modelled by multi-response (MR) and single-response (SR). Graphs showing the causal relationships modelled by multi-response (MR) (**a** and **c**) and single-response (SR) (**b** and **d**) approaches. Arrows represent relationships between variables, which contribute to Occam’s bias when omitted, for example, in SR models (i.e., **b** and **d**). **(a)** and **(b)** show causal relationships among three hypothetical traits while **(c)** and **(d)** show causal relationships among life history traits and extinction risk.

**Fig 2.**
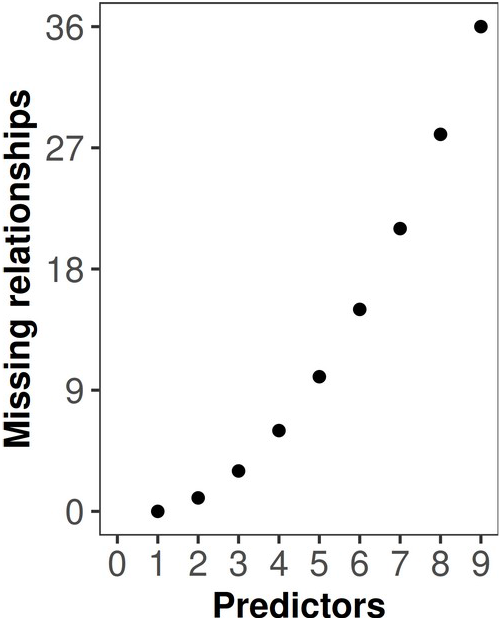
Relationship between the number of explanatory variables and indirect paths. The quadratic relationship between the number of predictors in a single-response (SR) model (x-axis) and the number of relationships which are not explicitly modelled (given by *y* = (*x* ^2^ -*x*) /2, defined for positive integers) but present in the data (y-axis).

Despite these risks, both direct and indirect causal paths on processes underpinning the evolution and extinctions of biodiversity can be accounted for in phylogenetic linear modelling. Bayesian multivariate or multiple-response (MR hereafter) models adapted to the phylogenetic framework have been available for 15 years (7,8), but have remained outside the mainstream (4). MR models use unstructured random effects to estimate the degree of phylogenetic and non-phylogenetic covariation among all possible combinations of response variables, as well as phylogenetic and non-phylogenetic variances (from which estimated phylogenetic signal is derived) for each response variable. In this way, the MR framework provides a base where the direct and indirect influences of species traits can be estimated, shaping statistical models into better structures that offer the analytical capabilities to identify more accurately the signals underlying the biological processes they are designed to search for.

Here, we elaborate on the need to re-focus hypothesis-testing approaches in comparative biology by considering Occam’s bias in phylogenetic linear modelling. We discuss the impacts of omitting relationships among predictors on the significance and directionality of coefficients. To formalise these discussions, we apply a general approach that is applicable to both phylogenetic and non-phylogenetic linear modelling contexts. Specifically, we identify the numerical conditions that make Occam’s bias more likely to cause false significant results and changes to directionality in coefficients, and show that those numerical conditions are biologically relevant using empirical data. To illustrate this rationale empirically, we employ both SR and MR approaches on a global-scale database of 13,949 species of tetrapods (amphibians, reptiles represented by lizards, mammals and birds) to revisit a core hypothesis that predicts the influence of life history traits (body mass, fecundity and geographic range size) on extinction risk (Figs 1c and 1d).

## Methods

### Data

We assembled a novel database spanning extinction risk (IUCN Red List conservation status) and life histories (body mass, fecundity, geographic range size) for 13,949 species from across all four classes of tetrapod vertebrates – amphibians, ‘reptiles’ (restricted to lizards), mammals and birds. These data were obtained from the scientific literature (field guides, journal articles, books), existing class-specific databases (see details below), as well as field and museum records collected by the authors. Firstly, we incorporated extinction risk into our models as the response variable using a binary proxy (the probability of being threatened) based on threat categories from the IUCN Red List, which we downloaded using IUCN API 2024-1, accessed with the R package *rredlist* (18). We assigned ’1’ to species if their threat status was ‘Vulnerable’, ‘Endangered’ and ‘Critically Engendered’ and ‘0’ to species if their threat status was ‘Least Concern’ and ‘Near Threatened’ while excluding ‘Data Deficient’ species. Secondly, we collated a database of body mass of tetrapods. For amphibians, we calculated species body mass from snout-vent length (SVL) for frogs and salamanders, and from total body length for caecilians. We then converted maximum SVL and maximum total body length into body mass using order-specific allometric formulas (19). Body mass data for lizards was gathered from Meiri et al. (20). For mammals, we used body mass from Cox et al. (21), defined as mean adult body mass. For birds, we obtained body masses from the AVONET database (22), where it was defined as average species body mass. Thirdly, we collected data for clutch and litter sizes (fecundity) from published databases. For amphibians, we used the minimum and maximum number of eggs per reproductive episode recorded for each given species to estimate clutch size range. From this, we obtained the midpoint clutch size that was used as a single value for fecundity per species in our analyses (23,24). We obtained clutch sizes for lizards from the dataset employed by Meiri et al. (20). This dataset used mean clutch size where possible, but took the midpoint between the smallest and largest clutch size reported for species which no means were available. For mammals, we used litter size from Cox et al.(21), defined as mean number of offspring per reproductive effort. For birds, we obtained clutch sizes from Jetz et al. (25), defined as the geometric mean of the typical minimum and maximum clutch size per species. Fourthly, geographic range size (i.e., geographic surface over which a species is known to occur in km^2^) data were extracted from species distribution polygons from the IUCN archive (www.iucnredlist.org) for amphibians and mammals, from GARD (www.gardinitiative.org) for lizards, and from Birdlife International (www.datazone.birdlife.org) for birds. We consider geographic range size to be a proxy life history trait because it captures the life history adaptations that allow species to achieve such a distribution. We also note that among tetrapods, geographic range size carries an extremely right-skewed distribution, reflecting that of other life history traits. Finally, the taxonomic nomenclature used in our database follows a range of primary repositories of tetrapod taxonomy. For amphibians, we follow Frost version 6.2 (26). For lizards, Reptile Database (March 2024) (27). For mammals, we follow version 1.12.1 of the Mammal Diversity Database species checklist (28). Finally, for birds, we follow version 8.1 (January 2024) of the HBW Birdlife taxonomic checklist (29), maintained by Birdlife International. Mismatches in names between our data sources and these taxonomies were dealt with individually, depending on context. All data are available in Dataset 1.

### Phylogenies

For phylogenetic analyses, we used consensus trees. For amphibians, we used Jetz and Pyron’s phylogenetic supertree (30). For lizards, mammals and birds, we summarised the posterior distributions of time-calibrated trees available from Tonini et al. (31), the Phylacine 1.2 database (32), and the Birdtree database (33) respectively, using median node heights with the TreeAnnotator app of Beast 2.5 (34). For birds, we selected the Hackett backbone.

### Overview of statistical approach

We implemented Bayesian phylogenetic generalised linear mixed modelling using the R package *MCMCglmm* version 2.36 (7) in R version 4.5.2 (35). We used MCMC to sample the posterior distribution of multivariate mixed models and incorporated the phylogeny (represented by the inverse of the phylogenetic covariance matrix for both tips and internal nodes produced by the function ’inverseA()’ into the random effects. We implemented both single-response (SR hereafter) and multiple-response (MR hereafter) model structures. For MR models, fixed effects were restricted to fixed intercepts for each response variable. For random effects, we utilised an unstructured, multidimensional variance-covariance matrix, which allows for the estimation of the phylogenetic and residual variances of each response variable as well as the phylogenetic and residual covariances of each pairwise combination of response variables. Consequently, evidence for a prediction is derived from observing significant phylogenetic or residual covariances, with the former indicating a significant relationship with clustering by clades while the latter indicates a significant relationship without clustering by clades. In contrast, SR models regress a single response variables on multiple predictors as fixed effect. In these models, the phylogenetic and residual variances are estimated only for the response, and the relationships among the predictors are not accounted for. Thus, evidence for a prediction is derived from observing significance in the fixed effects.

### Simulations

For the simulated analysis, we implemented a three trait system (Trait 1, Trait 2 and Trait 3), where we held the phylogenetic correlation of Trait 1 and Trait 2 (r_12_) close to zero while systematically varying the correlations between Trait 1 and Trait 3 (r_13_) as well as Trait 2 and Trait 3 (r_23_) in order to determine which values of r_13_ and r_23_ would produce false significant results. These systematically varying correlations (i.e., r_13_ and r_23_), were organised into a matrix of possible correlations which ranged from -0.7 to 0.7 in steps of 0.1. For each combination of trait correlations, we simulated fresh multivariate, normal, phylogenetically correlated data to run the model. For this purpose, we generated purely random phylogenies using the function ‘rtree()’ from the package *ape (36)*, and then rescaled branch lengths so that root-to-tip distance was 1. To generate trait data, we took the Kronecker product of the phylogenetic correlation matrix and the matrix that represents the trait correlation structure, and used the outputted matrix in the ‘mvrnorm()’ function from the R package *MASS (37)*. This function creates multivariate, normal, phylogenetically correlated data by generating random samples from a multivariate normal distribution given a mean vector and a covariance matrix as inputs.

We used SR and MR model structures utilising Gaussian link functions. For the SR model structure, the response was Trait 1 while the predictors were Trait 2 and Trait 3. The phylogenetic and residual variances were estimated for the response. For the MR model structure, we modelled three response variables (Trait 1, Trait 2 and Trait 3). Fixed effects included only the fixed intercept for each response. Random and residual effects each implemented an unstructured, multi-dimensional matrix for the covariances of each pairwise combination of responses.

We repeated this process using three different sample sizes (*n* = 100, *n* = 1,500 and *n* = 5,000) resulting in 1,350 Bayesian models. While we ensured all parameters of each of these models had an effective sample size of at least 1,000, we did not check retrospectively for convergence in each of these outputs. Instead, prior to analysis, we ran pilot tests using selected combinations of correlations and checked those outputs for convergence by examining trace plots in order to get a feel for the general complexity of the parameter space created by a three-trait system with a random phylogeny. We then implemented the specifications relating to successful convergence in the pilot analyses to the main, simulated analyses. The purpose of these simulations was to visualise the existence of a pattern that is fundamental to linear modelling, rather than to test a prediction using empirical data. Consequently, we emphasise that this approach to convergence is only adequate for this specific context. We do not recommend that this approach to convergence is applied to other contexts. Prior and sampling specifications are listed in Table S3.

### Empirical analyses

To test our hypothesis on life history drivers of extinction risk among tetrapods, and that Occam’s bias impacts these analyses, we ran SR and MR models. For both SR and MR models, extinction risk (binary) was modelled using a threshold (probit) link function. In other words, we assume extinction risk is expressed by a latent, normally distributed continuous process which contains a threshold value at which species are separated into the binary outcome. We chose this approach because directly modelling the threshold or ‘tipping point’ more accurately represents the mechanism of demographic decline to extinction (38–40). All continuous variables were transformed using natural logarithms, then *z*-transformed and modelled using a Gaussian link function. We utilised four different SR model structures, with each regressing extinction risk on various combinations of body mass, fecundity and geographic range size. Firstly, model ‘B’ incorporated body mass as the sole predictor thus excluding covariates. Secondly, model ‘BF’ used body mass and fecundity as predictors. Thirdly, model ’BG’used body mass and geographic range size as predictors. Finally, model ‘BFG’ used body mass, fecundity and geographic range size as predictors. We modelled the phylogenetic variance of extinction risk as a random effect, while the residual variance was fixed to 1. We utilised a single MR model structure. This included all variables (i.e., extinction risk, body mass, fecundity and geographic range size) as response variables. Fixed effects included only the fixed intercept for each response. Random and residual effects each implemented an unstructured, multi-dimensional matrix for the covariances of each pairwise combination of responses. However, the residual variance was fixed to 1 for extinction risk, but estimated for the continuous response variables.

In all models, and for each parameter, convergence was determined by examination of trace plots and we ensured that effective sample size was greater than 1,000. Prior and sampling specifications are listed in Table S3.

## Results

### Type I and Type II errors increase with increasing sample size and mildly correlated predictors

To present Occam’s bias, we consider a hypothetical situation where the prediction that Trait 2 influences Trait 1 is tested, while modelling the effect that Trait 3 also exerts onto Trait 1 (i.e. a model structure such that Trait 1 ∼ Trait 2 + Trait 3). In this scenario, theory predicts that Trait 3 and Trait 2 may also influence one another, and thus the correct model to capture these interactions is given by Fig 1a. However, we test the prediction that Trait 2 influences Trait 1 using the SR model structure (Fig 1b) and therefore, Trait 3 would be the bias-inducing variable whose inclusion causes spurious results. We set the bivariate phylogenetic correlation between Trait 1 and Trait 2 to be approximately zero and we investigate under what numerical conditions a spurious, statistically significant relationship could be induced between them. For this purpose, we derive the ‘Occam’s bias inequality’. This inequality is based on the partial correlation formula, and may be used as a broad approximation of impacts relating to directionality and significance (under the same significance threshold) of coefficients (but not precise coefficient estimates) across a variety of SR model contexts. The Occam’s bias inequality is given by eq. 1,

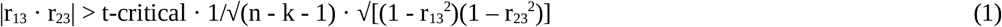

where r_13_ is the correlation between Trait 1 and Trait 3, r_23_ is the correlation between Trait 2 and Trait 3, n is sample size and k is number of predictors controlled for, and which is only valid where r_12_ (the correlation between Trait 1 and Trait 2) is 0. When the inequality is true, the inclusion of the covariate Trait 3 in SR model structure causes a spurious, significant relationship between Trait 1 and Trait 2, but not under MR models. The right-hand side of the Occam’s bias inequality comprises of two terms– the significance threshold 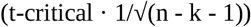 and a scalar 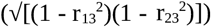. Thus, we can write the Occam’s bias inequality as

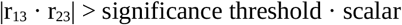

where the significance threshold is influenced by correlations of the other variables in the model (r_13_ and r_23_) via the scalar. We use the Occam’s bias inequality to predict which values of r_13_ and r_23_ would cause spurious results in Trait 2 where Trait 1 is regressed on Trait 2 and Trait 3. Unsurprisingly, we found predictions from the Occam’s bias inequality on false significance are broadly consistent with Bayesian phylogenetic generalised linear mixed models using a Gaussian link function with simulated, systematically correlated data and random phylogenies (Fig 3). In contrast, MR models revealed only apparently random patterns of false significance, likely resulting from multiple testing (Fig 3). The Occam’s bias inequality and simulations reveal three important findings on false significance (see below).

**Fig 3.**
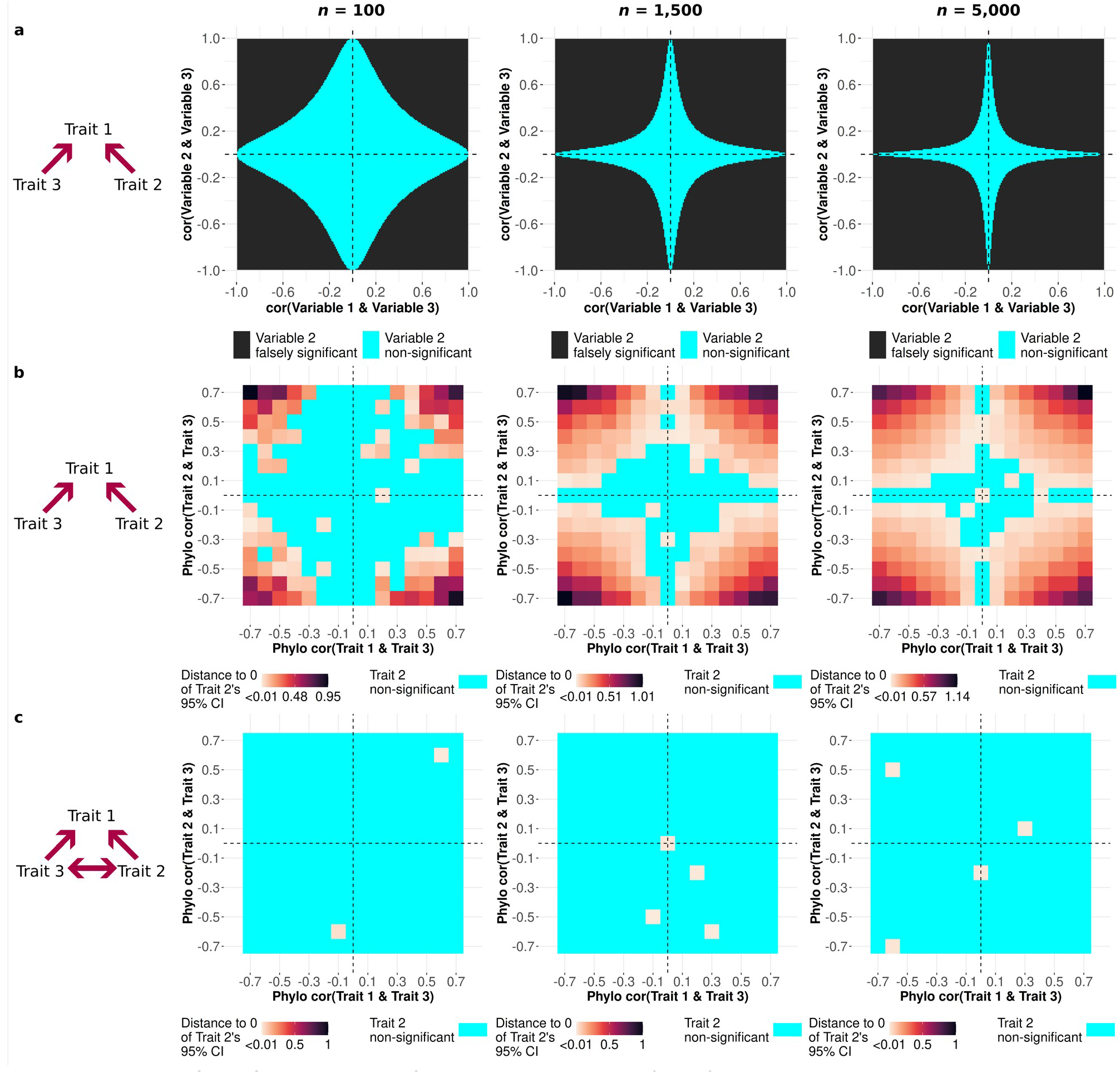
Response surface of statistical significance. The response surface of a three variable system, showing the correlation between Trait 1 and Trait 3 (x-axes), the correlation between Trait 2 and Trait 3 (y-axes), and impact of these on the significance of Trait 2 in predicting Trait 1 (represented by the colour in the tile). Trait 1 and Trait 2 do not have a relationship (or have a bivariate correlation of approximately zero) while the correlation of Trait 1 and Trait 3 and the correlation of Trait 2 and Trait 3 are allowed to vary. Graphs show, at three different sample sizes (*n* = 100, *n* = 1,500 and *n* = 5,000), **(a)** the impact on the significance of Trait 2 when Trait 1 is regressed on Trait 2 and Trait 3 (i.e., single-response or SR model structure), as predicted by the Occam’s bias inequality; **(b)** the impact on the significance of Trait 2 when Trait 1 is regressed on Trait 2 and Trait 3 (i.e., single-response or SR model structure), as revealed by Bayesian phylogenetic mixed models of simulated, phylogenetically correlated data; and **(c)** the impact on the significance of Trait 2 in predicting Trait 1 when all possible causal paths among Trait 1, Trait 2 and Trait 3 are considered (i.e., multi-response or MR model structure), as revealed by Bayesian phylogenetic mixed models of simulated, phylogenetically correlated data.

First, we find that the significance of spurious relationships increases as the strength of the other pairwise correlations among variables increases in magnitude (i.e., r_13_ or r_23_) (Figs 3 and 4). This is because as either r_13_ or r_23_ increases in magnitude, the left-hand side of the Occam’s bias inequality increases while the right-hand side (where the significance threshold is located) decreases (Fig 4). The decrease in the right-hand side is because larger decimal values given by r_13_ or r_23_ subtracted from 1 result in a smaller decimal scalar for the significance threshold, creating an increasingly larger difference between the sides of the inequality. Consequently, as r_13_ or r_23_ increases in strength, the significance threshold becomes progressively lower as the value being compared becomes progressively larger (Fig 4).

**Fig 4.**
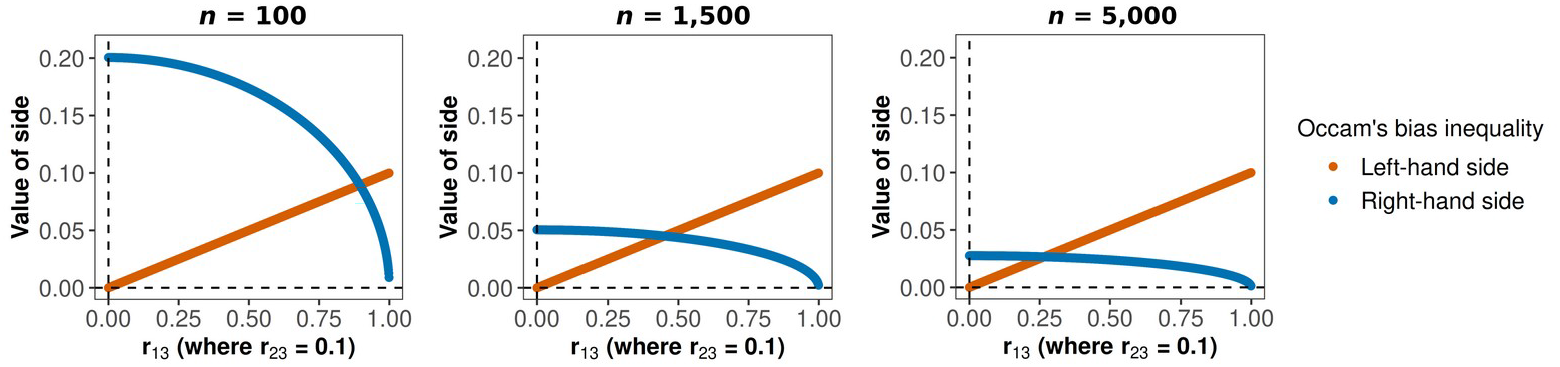
Occam’s bias inequality. Graphs depicting the left-hand and right-hand sides of the Occam’s bias inequality at three different sample sizes (*n* = 100, *n* = 1,500 and *n* = 5,000) for the three-variable system where Trait 1 is regressed on Trait 2 and Trait 3. The value for the sides of the Occam’s bias inequality appear on the y-axes with each side corresponding to a colour, while the value for r_13_ (the correlation between Trait 1 and Trait 3) appears on the x-axes, and the value for r_23_ (the correlation between Trait 2 and Trait 3) is fixed to 0.1. Note that due to equal weights of r_13_ and r_23_ in the Occam’s bias inequality, the same pattern would emerge if r_23_ appeared on the x-axes while r_13_ was fixed to 0.1. The Occam’s bias inequality is non-significant until the lines cross. Beyond the crossing point, the distance between the lines increase progressively, reflecting the increases in significance as measured in our simulations by the distance of the 95% credible interval from zero.

Second, we found that as sample size increases, the omitted trait correlations (i.e., r_23_) needed to induce spurious, significant results become progressively and considerably reduced, indicating that large-scale studies are most likely to be impacted by Occam’s bias (Figs 3 and 4). This is because of the impact of sample size and degrees of freedom on components used to construct the Occam’s bias inequality. Larger sample sizes result in decreases in the standard error of the partial correlation formula due to its location in the denominator (Supplementary Methods), which in turn decreases the significance threshold determined by the t-critical value. At the same time, larger degrees of freedom (i.e., achieved by larger sample size) results in thinner tails in the distribution used to calculate the t-critical value, which also decreases the significance threshold.

Third, we found that the stronger the relationship between the response (Trait 1) and the covariate (Trait 3), the weaker the omitted relationship (i.e., that between Trait 2 and Trait 3) has to be to induce a falsely significant result (Fig 3). This is because the in the left-hand side of the Occam’s bias inequality, r_13_ and r_23_ have equal weight such that if one increases, lower magnitudes of the other are needed to achieve the same effect (Fig 3). Therefore, including a covariate because it is known to be a particularly strong predictor of the outcome would ironically bias the coefficient of the variable of interest even more than it would have had the covariate been a weaker predictor of the outcome.

Note that while we focused on Type I error (false positives) in these simulations, if we had used simulated data with different marginal correlations, the same partial correlation mechanism could apply to result in Type II error (false negatives) as a significant coefficient can be dragged towards zero due to Occam’s bias.

### Impacts of Occam’s bias on the magnitude and directionality of coefficients

While still considering the specific situation in which Trait 1 is regressed on Trait 2 and Trait 3, where Trait 2 becomes spuriously significant when we condition on Trait 3, we use the partial correlation formula (eq. 2) to understand the impact of the bias on the coefficient of Trait 2 (regardless of significance).

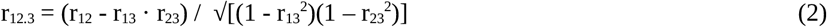

We observe that the biased coefficient in the SR model can be either positive or negative, depending on the sign of the correlations of the other variables in the data (r_13_ and r_23_) (Fig 5). A positive product of r_13_ and r_23_ results in a negative posterior mean for Trait 2, while a negative product of r_13_ and r_23_ results in a positive coefficient for Trait 2 (Fig 5). These sign changes in coefficients of SR models are determined in the numerator of the partial correlation formula (r_12_ - r_13_ · r_23_). While in the Occam’s bias inequality we assume r_12_ equals precisely zero (Supplementary Methods), in empirical datasets correlations cannot be precisely zero and instead express as a value very close to zero, either positive or negative. Therefore, where r_13_ and r_23_ have the same sign, the biased coefficient (r_12_) will be negative, because a value close to zero minus a positive value further from zero will result in a negative value (Fig 5). In contrast, where r_13_ and r_23_ differ in sign, the biased coefficient (r_12_) will be positive, because a value close to zero minus a negative value further from zero will result in a positive value (Fig 5).

**Fig 5.**
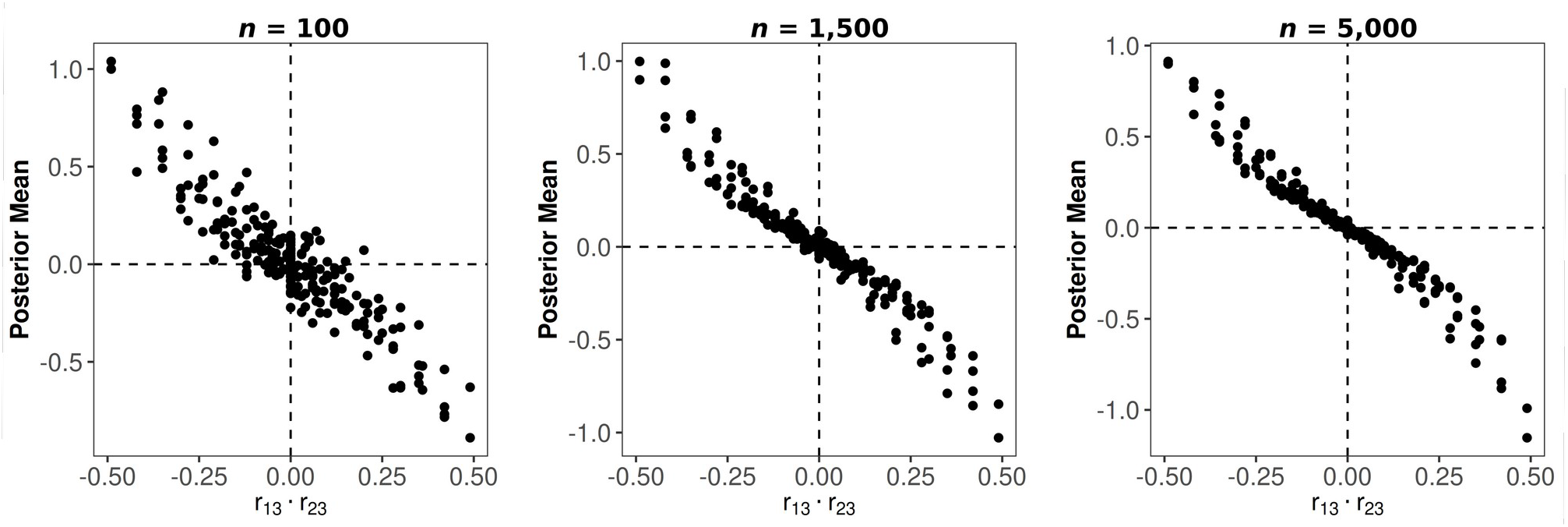
Response surface of posterior means. Scatter plots showing the response surface of coefficients (posterior means) of the biased coefficient (Trait 2) from simulated Bayesian phylogenetic models where Trait 1 is regressed on Trait 2 and Trait 3 at three different sample sizes (*n* = 100, *n* = 1,500 and *n* = 5,000). The graphs how the product of the correlation between r_13_ and r_23_ (x-axes) influences the posterior mean of the Trait 2 (y-axes), whose bivariate correlation with Trait 1 is approximately zero. A positive product of r_13_ and r_23_ results in a negative posterior mean for Trait 2, while a negative product of r_13_ and r_23_ results in a positive coefficient for Trait 2.

### Empirical case study: Occam’s bias impacts analyses of life history evolution and extinction risk

Our analyses addressing the hypothesis that life history traits influence variation in extinction risk (binary, based on IUCN Red List threat categories) across species reveal that modelling the effect of life history traits without accounting for the evolutionary relationships among them obscures their predicted effects by returning spurious results.

Consistent with life history theory, MR models reveal significant relationships in life history traits among tetrapods, at sufficiently high magnitudes to cause Occam’s bias (Fig 6; Table S1). The observation of these relationships supports the use of MR models over SR models when the effects of more than one trait (life histories in the case of our analysis) on any other variable are modelled. Specifically, we find that fecundity significantly increases with body mass among ectotherms, whereas it decreases among endotherms (Fig 6; Table S1). Regarding geographic range size, while it increases with fecundity among tetrapods in general, the relationship between geographic range size and body mass is positive among ectotherms but non-significant among endotherms (Fig 6; Table S1). The relationships among our three life history traits –if omitted from models that seek to determine the effect of life history traits on some other outcome– all contribute to false coefficients because the Occam’s bias inequality for four variables (i.e., extinction risk, body mass, fecundity and geographic range size) would be based on second-order partial correlations.

**Fig 6.**
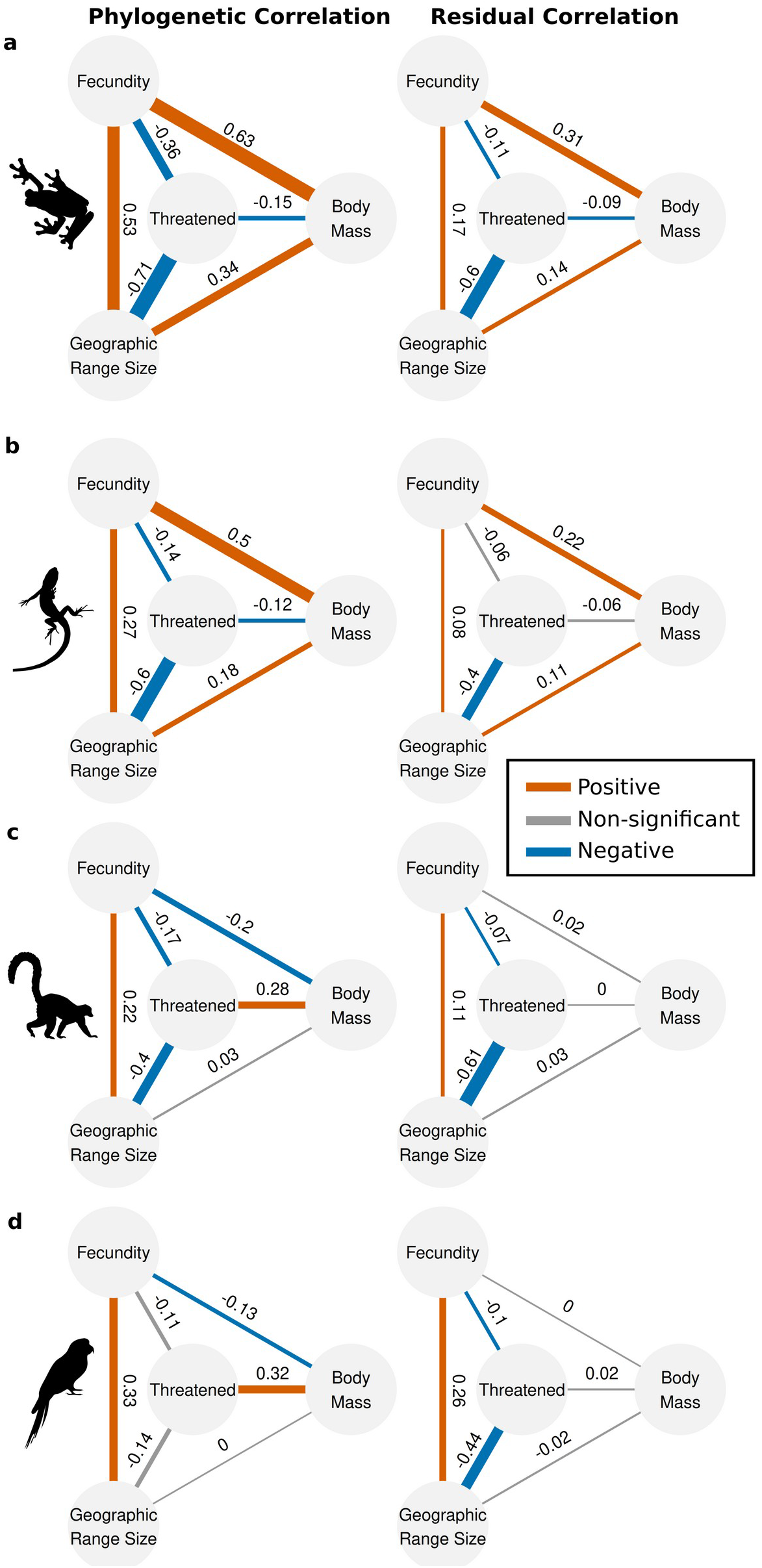
Relationships of life history traits and extinction risk. Paths depicting the relationships between life history traits and extinction risk (binary, the probability of being threatened based on IUCN conservation categories) from MR models. For ease of interpretability, the posterior of each covariance was converted to the correlation scale, before the means were taken and displayed in the figure (see Table S1 for original covariance estimates). Thickness of the line represents the absolute value of the mean, while the colour represents the significance taken from the 95% credible intervals of the covariances, which we considered to be significant if they did not include zero. Both phylogenetic (between-clade) and residual (within-clade) covariation are shown for amphibians, lizards mammals and birds.

Having estimated covariation among life history traits by use of MR models, findings reveal the effect of life history traits on extinction risk are aligned with theory (in contrast to previous work and analyses below using SR models) (Fig 6; Table S1). Among tetrapods, increased extinction risk is consistently significantly associated with decreasing fecundity and decreasing geographic range size (Fig 6; Table S1). However, as expected, the role of body mass in influencing extinction risk differs between ectotherms and endotherms. Significant, negative relationships are observed among ectotherms while significant, positive relationships are observed among endotherms (Fig 6; Table S1). These findings are predicted by theory (41–43) and corroborate the idea that ecophysiological differences between ectotherms and endotherms need to be accounted for.

Specific findings from SR and MR models differ in a number of ways when it comes to quantifying the predictive role of life history traits in extinction risk. First, SR models consistently reveal that fecundity plays no significant role in influencing variation in extinction risk (Fig 7; Table S2). Second, while SR models reveal a significant, positive association between body mass and extinction risk in endotherms, the reverse relationship is not observed among ectotherms (Fig 7; Table S2). Instead, larger body mass increases extinction risk among amphibians, while a positive, non-significant relationship between body mass and extinction risk is observed among lizards (Fig 7; Table S2). Third, we found that the addition of related predictors (i.e., fecundity, geographic range size) progressively causes further bias in coefficients (Fig 7; Table S2), but this effect may be contextual (i.e. it may depend on the specific combination of correlations observed in our dataset rather than constituting a generalisable rule). We note that the deviance information criterion (*DIC*) improves by the addition of related predictors (Table S2) and subsequent biasing of coefficients, highlighting that model comparison metrics –while helpful in predictive modelling– may not serve modelling for causal inference. In fact, among ectotherms, the SR models that offer the most biologically relevant inferences (i.e., those in which extinction risk is predicted only by body mass) have the worst *DIC* (Table S1).

**Fig 7.**
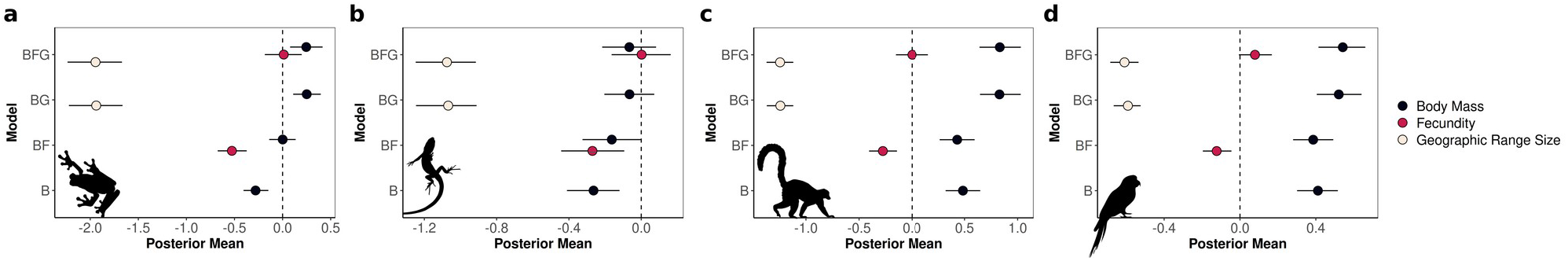
Coefficients of single-response life history models. Posterior means from single-response models of life history predictors of extinction risk (binary, based on IUCN threat categories) ran using R package *MCMCglmm*. Four models were ran. Model ‘B’ includes body mass as the single predictor. Model ‘BF’ includes body mass and adds the covariate fecundity. Model ‘BG’ includes body mass and adds the covariate geographic range size. Model ‘BFG’ includes body mass, fecundity and geographic range size as predictors. The bias in the posterior mean of body mass (and fecundity) worsens as predictors as added, in that they deviate from what would be expected based on their relationships in the multi-response models (see Fig 6). Posterior means are represented by dots while 95% credible intervals are represented by whiskers. Effects are considered significant when the 95% credible interval does not include zero, which is represented in the figure by the dashed black lines.

Similar to coefficient changes in simulated findings, we observed that the impacts of Occam’s bias may be understood in terms of the numerator of the partial correlation formula (eq. 2 above). We consider how the correlations among the variables extinction risk (E), body mass (B), fecundity (F) and geographic range size (G) impact the estimated coefficient of body mass among ectotherms and endotherms.

Among ectotherms, taking the SR models of

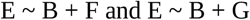

the numerator of the estimated relationship between body mass and extinction risk in both models would be

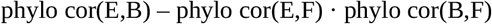

and

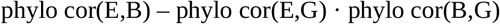

These correlations among ectotherms are

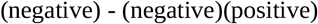

(Fig 6; Table S1) resulting in a negative value subtracted from a negative value, thus making the effect size of the relationship of body mass and extinction risk more positive (Fig 7; Table S2). This results in moving the estimate of body mass closer to zero (e.g., lizards with fecundity; lizards with geographic range size), making it non-significant (e.g., amphibians with fecundity; lizards with geographic range size), or even significantly positive (e.g., amphibians with geographic range size) (Fig 7; Table S2).

Among endotherms, Occam’s bias in coefficients is present but less apparent due to differences in relationships of body size, fecundity and geographic range size. In contrast to ectotherms, among endotherms the relationships between body size and fecundity are negative and relatively weak Fig 6; Table S1). Thus, the estimated coefficient for the relationship between body mass and extinction risk from the SR model

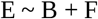

is

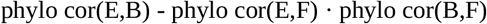

resulting in

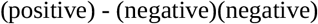

which manifests as a slightly less positive coefficient (Fig 7; Table S2). In addition, while no significant relationships were observed between body size and geographic range size among endotherms (Fig 6; Table S1), the inclusion of geographic range size in SR models nevertheless causes Occam’s bias in the coefficient of body mass due to the sign and strength of their correlation. For instance, in the SR mode

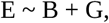

the coefficient for body mass would be

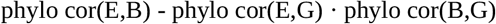

resulting in

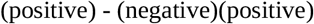

causing the inflation of the already positive coefficient for body mass (Fig 7; Table S2).

We note that, while the MR approach decomposes each variable’s effect into phylogenetic (between-clade) and residual (within-clade) covariation, the non-linear link functions in SR model produces predictor coefficients that are a weighted average of the within- and between-clade variation. Thus –irrespective of Occam’s bias– the SR coefficients reflect the effect of the predictor in the phylogenetic space which concentrates the greatest variation in the outcome, rather than a clear, biologically meaningful within- or between-clade effect. Therefore, in relation to Occam’s bias, the observed predictor coefficient may (in some cases) be better understood when considering the between-clade estimates of MR models (as done here) or (in other cases) the within-clade estimates.

## Discussion

Our study proposes Occam’s bias and provides both simulation and empirical evidence of the impact that this form of bias exerts on inference from phylogenetic analyses. These empirical analyses in particular align with our proposition that a large expanse of studies are likely to be impacted by Occam’s bias – our results support the hypothesis that the numerical conditions which lead to Occam’s bias (e.g. the magnitude of correlation between predictors; the increase in bias with increasing same sizes) are biologically relevant and impact findings when correlations are at levels below multicollinearity thresholds(i.e. 0.7 (44)). Importantly, however, the identification of Occam’s bias in phylogenetic analyses serves not merely to retrospectively highlight the impact that Occam’s bias has had on past hypothesis testing, but as a catalyst for reconsidering how statistical models for inferences on observational data are employed going forward. To this end, we open the discussion on the appropriateness of SR models in phylogenetic analyses.

### Reconciliation of model with theory

SR and MR modelling performed on the same dataset are subject to the same ‘covariation budget’, such that the total covariation present in the data is equivalent to the sum of covariation captured by the model and the residuals. In SR models, while the relationships between predictors are not *explicitly* estimated as parameters, their covariation is *implicitly* captured by the model (i.e. taken as given), expressing in the beta coefficients via the partial correlation mechanism in which the relationship between the predictors is subtracted from the estimates of predictors. The implicit allocation of covariation to the model avoids correlation in residuals, but also results in the statistical manifestations of what can be interpreted as Occam’s bias. In contrast, the structure of MR model used here is ‘fully saturated’ (i.e. the number of estimated parameters in the model equals the total number of pieces of information available in the data), thus all covariation in the data is captured explicitly under distinct coefficients.

Can the reallocation of covariation between predictors to predictor coefficients in SR models be justified for inferences restricted to interspecific, observational data. One perspective is that the reallocation is justified if no causal pathway is hypothesised to exist between our predictors (45). Alternatively, another perspective is that a fully saturated MR model should be preferred because the coefficients are more interpretable biologically. Even when predictors are causally independent in theory but statistically correlated, their covariation should be treated as incidental and therefore should be separated from the other parameters by estimating it explicitly rather than mixing it with the effects of the predictors via the partial correlation mechanism. The mixing of the covariation among predictors into the coefficients of SR models is often referred to as ‘partialling out’ or ‘controlling for’, but this language is easily misinterpreted in biological contexts. Technically, SR models do ‘control for’ the relationship between predictors in that the covariation is captured by the model rather than the residuals, but ultimately the covariation is conserved under ‘labels’ which are less useful for inferences. In a predictive context, the partial correlation mechanism is advantageous as it improves predictive ability by accounting for variance explained indirectly by the covariation among predictors without explicitly modelling it. Whereas the biological meaning of the partial correlation mechanism of SR approach is not clear, the MR approach offers estimates that are more interpretable biologically. By estimating all relationships simultaneously, MR models allow us to isolate the unaltered relationships between each variable.

The approach of explicitly estimating hypothesised null relationships in a saturated MR model serves as a diagnostic tool which can be used to decide whether there is empirical evidence for the proposed causal structure. Specifically, this tool addresses the issue that hypothesised null relationships may nevertheless express as significant in the fitted model. Thus, the map of all possible relationships estimated from the saturated MR model does not need to represent the proposed causal theory (provided no relationships are missing), but instead presents us with information which better allows for the evaluation of empirical support for the proposed causal theory because covariation has been appropriately categorised while accounting for all back-door paths (mitigating Occam’s bias). Thus, in relation to any hypothesis, a significant relationship observed in a fully saturated MR model may have causal meaning (forming a causal path within the mechanism presented by the hypothesis) or may be incidental (in which its ‘cause’ is superficial or more proximate than the mechanism presented by the hypothesis), and the decision for which it is derives solely from scientific theory. Therefore, while in other statistical contexts the fully saturated MR model often isn’t considered useful for hypothesis testing (46), we propose that due to the violations of assumptions (9,10) presented by the hypothesised relationships among species traits, MR approaches offer a more biologically-relevant test for empirical support of hypotheses in phylogenetic analyses than SR.

## Conclusions

Our study shows that the omission of causal pathways among variables is a major cause of Occam’s bias in phylogenetic linear modelling. Our findings derive in six important implications:

1. The addition of mildly correlated predictors and larger sample sizes increases Occam’s bias. Studies using small datasets or those with no covariates are less likely to be implicated by our findings.
2. These findings challenge the standard method of incorporating a covariate into a multiple linear regression because it is known to be a strong predictor of the outcome. Our findings identify that the stronger the predictor of the outcome is the lower the level of correlation among predictors is required to cause false results.
3. Occam’s bias emerges from correlations among predictors at magnitudes below multicollinearity thresholds (i.e. 0.7 (44)). In other words, Occam’s bias interferes with reliable inference to the extent of changing significance and direction of coefficients before the impacts of multicollinearity need to be considered.
4. We highlight that model performance metrics serve limited purpose in scientific inference (47), as our findings revealed that models with better performance metrics were nevertheless the least biologically relevant.
5. We emphasise that our study identifies that inference using more than two variables can be made more reliable by the use of models which account for all possible pathways among variables, such as MR models. Therefore, we call for a re-evaluation of scientific inferences made using SR model structures, and advocate the use of MR model structures or others that account for all causal pathways.
6. We call for continued development and expansion of phylogenetic MR approaches.

## Acknowledgements

JG is indebted to the Natural Environment Research Council (NERC) DTP QUADRAT for full PhD funding.

