## Supplementary material for "Occam’s bias undermines inferences from phylogenetic linear models": Dataset 1

| taxon | scientific_name | phylo_name | threatened | body_mass | fecundity | range_size |
|---|---|---|---|---|---|---|
| amphibians | "Cochranella"_riveroi | Cochranella_riveroi | 1 | 1.17 | 21 | 59.43 |
| amphibians | "Litoria"_castanea | Litoria_castanea | 1 | 79.58 | 2889 | 54538.32 |
| amphibians | Acanthixalus_sonjae | Acanthixalus_sonjae | 1 | 4.93 | 9 | 6030.02 |
| amphibians | Acanthixalus_spinosus | Acanthixalus_spinosus | 0 | 4.93 | 9 | 1213834.98 |
| amphibians | Acris_blanchardi | Acris_blanchardi | 0 | 4.57 | 275 | 2345138.95 |
| amphibians | Acris_crepitans | Acris_crepitans | 0 | 4.54 | 270 | 970974.66 |
| amphibians | Acris_gryllus | Acris_gryllus | 0 | 2.87 | 207.5 | 667451.52 |
| amphibians | Adelophryne_baturitensis | Adelophryne_baturitensis | 0 | 0.33 | 3.5 | 1866.43 |
| amphibians | Adelophryne_gutturosa | Adelophryne_gutturosa | 0 | 0.28 | 1 | 161196.36 |
| amphibians | Adelophryne_maranguapensis | Adelophryne_maranguapensis | 1 | 0.36 | 5.5 | 1.29 |
| amphibians | Adelotus_brevis | Adelotus_brevis | 0 | 7.84 | 440.5 | 151591.63 |
| amphibians | Adelphobates_castaneoticus | Adelphobates_castaneoticus | 0 | 1.17 | 7 | 630666.8 |
| amphibians | Adelphobates_galactonotus | Adelphobates_galactonotus | 0 | 5.58 | 10 | 996582.34 |
| amphibians | Adelphobates_quinquevittatus | Adelphobates_quinquevittatus | 0 | 0.59 | 6.5 | 777019.41 |
| amphibians | Adenomera_andreae | Adenomera_andreae | 0 | 1.87 | 16 | 5953365.83 |
| amphibians | Adenomera_araucaria | Adenomera_araucaria | 0 | 0.57 | 9.5 | 57646.59 |
| amphibians | Adenomera_bokermanni | Adenomera_bokermanni | 0 | 1.18 | 18 | 10978.04 |
| amphibians | Adenomera_diptyx | Adenomera_diptyx | 0 | 1.17 | 81.5 | 568792.85 |
| amphibians | Adenomera_hylaedactyla | Adenomera_hylaedactyla | 0 | 2.11 | 18.5 | 8309254.3 |
| amphibians | Adenomera_lutzi | Adenomera_lutzi | 1 | 3.02 | 6.5 | 2985.45 |
| amphibians | Adenomera_marmorata | Adenomera_marmorata | 0 | 2.11 | 8 | 52409.52 |
| amphibians | Adenomera_martinezi | Adenomera_martinezi | 0 | 1.17 | 13 | 5534.84 |
| amphibians | Adenomus_kelaartii | Adenomus_kelaartii | 1 | 11.04 | 975 | 11262.45 |
| amphibians | Afrixalus_aureus | Afrixalus_aureus | 0 | 1.02 | 49.5 | 100277.56 |
| amphibians | Afrixalus_delicatus | Afrixalus_delicatus | 0 | 1.5 | 55 | 748759.8 |
| amphibians | Afrixalus_dorsalis | Afrixalus_dorsalis | 0 | 4.54 | 60.5 | 609648.18 |
| amphibians | Afrixalus_fornasini | Afrixalus_fornasini | 0 | 5.36 | 50.5 | 944389.11 |
| amphibians | Afrixalus_fulvovittatus | Afrixalus_fulvovittatus | 0 | 1.69 | 45 | 95612.23 |
| amphibians | Afrixalus_knysnae | Afrixalus_knysnae | 1 | 1.33 | 29.5 | 1597.21 |
| amphibians | Afrixalus_laevis | Afrixalus_laevis | 0 | 1.69 | 5 | 1211518.69 |
| amphibians | Afrixalus_paradorsalis | Afrixalus_paradorsalis | 0 | 3.47 | 49 | 450758.52 |
| amphibians | Afrixalus_quadrivittatus | Afrixalus_quadrivittatus | 0 | 2.6 | 50 | 5187441.86 |
| amphibians | Afrixalus_spinifrons | Afrixalus_spinifrons | 0 | 1.17 | 30 | 58901 |

| | | | | | | |
|---|---|---|---|---|---|---|
| amphibians | Afrixalus_uluguruensis | Afrixalus_uluguruensis | 1 | 2.35 | 13 | 593.27 |
| amphibians | Afrixalus_vittiger | Afrixalus_vittiger | 0 | 1.69 | 100 | 1528799.61 |
| amphibians | Afrixalus_weidholzi | Afrixalus_weidholzi | 0 | 1.17 | 140 | 2158264.35 |
| amphibians | Agalychnis_annae | Agalychnis_annae | 1 | 59.73 | 697.5 | 2708.23 |
| amphibians | Agalychnis_callidryas | Agalychnis_callidryas | 0 | 45.08 | 138 | 501190.03 |
| amphibians | Agalychnis_dacnicolor | Pachymedusa_dacnicolor | 0 | 116.93 | 283.5 | 150009.61 |
| amphibians | Agalychnis_lemur | Agalychnis_lemur | 1 | 13.33 | 30.5 | 16484.3 |
| amphibians | Agalychnis_moreletii | Agalychnis_moreletii | 0 | 61.58 | 115 | 59737.54 |
| amphibians | Agalychnis_saltator | Agalychnis_saltator | 0 | 27.13 | 168 | 11425.7 |
| amphibians | Agalychnis_spurrelli | Agalychnis_spurrelli | 0 | 82.42 | 138.5 | 132657.12 |
| amphibians | Aglyptodactylus_laticeps | Aglyptodactylus_laticeps | 1 | 35.97 | 3798.5 | 2021.95 |
| amphibians | Aglyptodactylus_madagascariensis | Aglyptodactylus_madagascariensis | 0 | 31.33 | 1811 | 110547.76 |
| amphibians | Aglyptodactylus_securifer | Aglyptodactylus_securifer | 0 | 11.04 | 2686 | 117095 |
| amphibians | Alcalus_baluensis | Ingerana_baluensis | 0 | 2.11 | 8 | 240419.75 |
| amphibians | Allobates_brunneus | Allobates_brunneus | 1 | 1.02 | 17 | 28521.75 |
| amphibians | Allobates_femoralis | Allobates_femoralis | 0 | 3.81 | 28 | 4353103.97 |
| amphibians | Allobates_granti | Allobates_granti | 0 | 0.43 | 9 | 33358.99 |
| amphibians | Allobates_grillisimilis | Allobates_grillisimilis | 0 | 0.28 | 9.5 | 17629.35 |
| amphibians | Allobates_insperatus | Allobates_insperatus | 0 | 0.43 | 17 | 74685.49 |
| amphibians | Allobates_kingsburyi | Allobates_kingsburyi | 0 | 1.36 | 17.5 | 15817.21 |
| amphibians | Allobates_marchesianus | Allobates_marchesianus | 0 | 0.57 | 17 | 2390953.86 |
| amphibians | Allobates_nidicola | Allobates_nidicola | 0 | 0.71 | 4 | 239832.4 |
| amphibians | Allobates_olfersioides | Allobates_olfersioides | 0 | 0.48 | 8 | 92525.88 |
| amphibians | Allobates_paleovarzensis | Allobates_paleovarzensis | 0 | 0.82 | 28 | 205782.9 |
| amphibians | Allobates_subfolionidificans | Allobates_subfolionidificans | 0 | 0.46 | 9 | 152731.78 |
| amphibians | Allobates_talamancae | Allobates_talamancae | 0 | 1.17 | 29 | 117497.08 |
| amphibians | Allobates_trilineatus | Allobates_trilineatus | 0 | 0.51 | 14.5 | 1039919.36 |
| amphibians | Allobates_zaparo | Allobates_zaparo | 0 | 2.35 | 34.5 | 47502.85 |
| amphibians | Allophryne_ruthveni | Allophryne_ruthveni | 0 | 2.35 | 300 | 3055083.07 |
| amphibians | Alsodes_gargola | Alsodes_gargola | 0 | 11.04 | 35 | 2443.34 |
| amphibians | Alsodes_nodosus | Alsodes_nodosus | 0 | 41.05 | 55 | 40420.34 |
| amphibians | Altiphrynoides_malcolmi | Altiphrynoides_malcolmi | 1 | 2.6 | 21 | 629.85 |
| amphibians | Altiphrynoides_osgoodi | Altiphrynoides_osgoodi | 1 | 22.16 | 307 | 55.8 |
| amphibians | Alytes_cisternasii | Alytes_cisternasii | 0 | 11.04 | 110 | 168715.8 |

| | | | | | | |
|---|---|---|---|---|---|---|
| amphibians | Alytes_dickhilleni | Alytes_dickhilleni | 1 | 16.87 | 93.5 | 31457.44 |
| amphibians | Alytes_maurus | Alytes_maurus | 1 | 16.87 | 42.5 | 5263.82 |
| amphibians | Alytes_muletensis | Alytes_muletensis | 1 | 5.36 | 14 | 228.51 |
| amphibians | Alytes_obstetricans | Alytes_obstetricans | 0 | 15.03 | 95.5 | 957570.84 |
| amphibians | Amazophrynella_minuta | Amazophrynella_minuta | 0 | 2.18 | 160 | 992304.1 |
| amphibians | Ambystoma_annulatum | Ambystoma_annulatum | 0 | 21.88 | 232.5 | 140918.31 |
| amphibians | Ambystoma_barbouri | Ambystoma_barbouri | 0 | 13.9 | 290.5 | 76990.05 |
| amphibians | Ambystoma_bishopi | Ambystoma_bishopi | 1 | 3.49 | 162.5 | 28444.29 |
| amphibians | Ambystoma_californiense | Ambystoma_californiense | 1 | 56.87 | 671 | 38873.7 |
| amphibians | Ambystoma_cingulatum | Ambystoma_cingulatum | 1 | 3.49 | 113 | 89204.97 |
| amphibians | Ambystoma_gracile | Ambystoma_gracile | 0 | 35.46 | 150 | 350902.19 |
| amphibians | Ambystoma_jeffersonianum | Ambystoma_jeffersonianum | 0 | 16.14 | 210 | 470696.23 |
| amphibians | Ambystoma_laterale | Ambystoma_laterale | 0 | 6.73 | 186.5 | 1767652.42 |
| amphibians | Ambystoma_mabeei | Ambystoma_mabeei | 0 | 2.6 | 4 | 112024.47 |
| amphibians | Ambystoma_macrodactylum | Ambystoma_macrodactylum | 0 | 11.13 | 237.5 | 1443398.78 |
| amphibians | Ambystoma_maculatum | Ambystoma_maculatum | 0 | 11.28 | 247.5 | 2992980.17 |
| amphibians | Ambystoma_mavortium | Ambystoma_mavortium | 0 | 23.22 | 120 | 3659783.25 |
| amphibians | Ambystoma_mexicanum | Ambystoma_mexicanum | 1 | 32.02 | 550 | 32.18 |
| amphibians | Ambystoma_opacum | Ambystoma_opacum | 0 | 6.73 | 131 | 1683648.27 |
| amphibians | Ambystoma_ordinarium | Ambystoma_ordinarium | 1 | 10.06 | 109 | 3394.33 |
| amphibians | Ambystoma_rosaceum | Ambystoma_rosaceum | 0 | 7.69 | 30 | 163030.96 |
| amphibians | Ambystoma_talpoideum | Ambystoma_talpoideum | 0 | 3.32 | 449.5 | 678160.02 |
| amphibians | Ambystoma_texanum | Ambystoma_texanum | 0 | 8.76 | 481.5 | 1389181.65 |
| amphibians | Ambystoma_tigrinum | Ambystoma_tigrinum | 0 | 68.34 | 3875.5 | 3989359.36 |
| amphibians | Ameerega_altamazonica | Ameerega_altamazonica | 0 | 1.17 | 20 | 35895.82 |
| amphibians | Ameerega_bassleri | Ameerega_bassleri | 1 | 7.84 | 21 | 17729.53 |
| amphibians | Ameerega_bilinguis | Ameerega_bilinguis | 0 | 0.85 | 10 | 69393.41 |
| amphibians | Ameerega_boliviana | Ameerega_boliviana | 0 | 1.43 | 10 | 12764.91 |
| amphibians | Ameerega_braccata | Ameerega_braccata | 0 | 0.75 | 7 | 51772.54 |
| amphibians | Ameerega_cainarachi | Ameerega_cainarachi | 1 | 2.42 | 20 | 2256.92 |
| amphibians | Ameerega_flavopicta | Ameerega_flavopicta | 0 | 2.22 | 24.5 | 375628.25 |
| amphibians | Ameerega_hahneli | Ameerega_hahneli | 0 | 1.17 | 19.5 | 3909906.31 |
| amphibians | Ameerega_parvula | Ameerega_parvula | 0 | 1.02 | 9.5 | 192116.22 |
| amphibians | Ameerega_pepperi | Ameerega_pepperi | 1 | 3.29 | 44 | 2962.19 |

| | | | | | | |
|---|---|---|---|---|---|---|
| amphibians | Ameerega_petersi | Ameerega_petersi | 0 | 2.35 | 9 | 126813.34 |
| amphibians | Ameerega_picta | Ameerega_picta | 0 | 1.17 | 17.5 | 3960562.05 |
| amphibians | Ameerega_rubriventris | Ameerega_rubriventris | 1 | 1.09 | 12.5 | 753.3 |
| amphibians | Ameerega_silverstonei | Ameerega_silverstonei | 1 | 7.84 | 25 | 2764.05 |
| amphibians | Ameerega_trivittata | Ameerega_trivittata | 0 | 15.03 | 34.5 | 3357773.1 |
| amphibians | Amerana_aurora | Rana_aurora | 0 | 250.1 | 2750 | 218209.31 |
| amphibians | Amerana_boylii | Rana_boylii | 0 | 54.82 | 750 | 236132.63 |
| amphibians | Amerana_cascadae | Rana_cascadae | 0 | 42.85 | 550 | 113888.53 |
| amphibians | Amerana_draytonii | Rana_draytonii | 0 | 282.37 | 3150 | 141156.35 |
| amphibians | Amerana_luteiventris | Rana_luteiventris | 0 | 107.68 | 1575 | 1322515.34 |
| amphibians | Amerana_muscosa | Rana_muscosa | 1 | 71.48 | 226 | 11894.09 |
| amphibians | Amerana_pretiosa | Rana_pretiosa | 1 | 131.8 | 1124.5 | 31015.42 |
| amphibians | Amerana_sierrae | Rana_sierrae | 1 | 61.58 | 450 | 24073.9 |
| amphibians | Amietia_angolensis | Amietia_angolensis | 0 | 141.99 | 450 | 343127.22 |
| amphibians | Amietia_delalandii | Amietia_quecketti | 0 | 74.11 | 150 | 1731220.2 |
| amphibians | Amietia_fuscigula | Amietia_fuscigula | 0 | 214.85 | 7950 | 333657.09 |
| amphibians | Amietia_hymenopus | Strongylopus_hymenopus | 0 | 25.82 | 350 | 6280.44 |
| amphibians | Amietia_inyangae | Amietia_inyangae | 1 | 18.87 | 60 | 591.99 |
| amphibians | Amolops_chunganensis | Amolops_chunganensis | 0 | 14.42 | 452.5 | 197777.05 |
| amphibians | Amolops_cremnobatus | Amolops_cremnobatus | 0 | 5.62 | 20 | 121959.74 |
| amphibians | Amolops_loloensis | Amolops_loloensis | 1 | 45.66 | 153.5 | 61845.41 |
| amphibians | Amolops_ricketti | Amolops_ricketti | 0 | 21.02 | 990.5 | 555082.37 |
| amphibians | Amolops_wuyiensis | Amolops_wuyiensis | 0 | 10.34 | 600.5 | 102692.34 |
| amphibians | Amphiuma_means | Amphiuma_means | 0 | 3026.27 | 190 | 491481.78 |
| amphibians | Amphiuma_pholeter | Amphiuma_pholeter | 0 | 146.79 | 11.5 | 56376.44 |
| amphibians | Amphiuma_tridactylum | Amphiuma_tridactylum | 0 | 4114.98 | 196 | 477336.12 |
| amphibians | Anaxyrus_americanus | Anaxyrus_americanus | 0 | 431.35 | 11150 | 5652731.68 |
| amphibians | Anaxyrus_boreas | Anaxyrus_boreas | 0 | 214.85 | 9600 | 2865276.61 |
| amphibians | Anaxyrus_californicus | Anaxyrus_californicus | 1 | 52.68 | 6190.5 | 37565.51 |
| amphibians | Anaxyrus_canorus | Anaxyrus_canorus | 1 | 39.31 | 3250 | 10686.36 |
| amphibians | Anaxyrus_cognatus | Anaxyrus_cognatus | 0 | 163.99 | 23177 | 2795125.82 |
| amphibians | Anaxyrus_compactilis | Anaxyrus_compactilis | 0 | 76.81 | 5000 | 314894.54 |
| amphibians | Anaxyrus_debilis | Anaxyrus_debilis | 0 | 14.16 | 1650 | 1460610.64 |
| amphibians | Anaxyrus_exsul | Anaxyrus_exsul | 1 | 48.58 | 8200 | 31.95 |

| | | | | | | |
|---|---|---|---|---|---|---|
| amphibians | Anaxyrus_fowleri | Anaxyrus_fowleri | 0 | 88.3 | 8999 | 1991769.67 |
| amphibians | Anaxyrus_hemiophrys | Anaxyrus_hemiophrys | 0 | 76.81 | 5177 | 1401163.64 |
| amphibians | Anaxyrus_houstonensis | Anaxyrus_houstonensis | 1 | 50.6 | 3356 | 30066.84 |
| amphibians | Anaxyrus_mexicanus | Anaxyrus_mexicanus | 0 | 32.83 | 4000 | 155243.16 |
| amphibians | Anaxyrus_microscaphus | Anaxyrus_microscaphus | 0 | 76.81 | 4075 | 242190.78 |
| amphibians | Anaxyrus_nelsoni | Anaxyrus_nelsoni | 1 | 198.59 | 5200 | 76.39 |
| amphibians | Anaxyrus_punctatus | Anaxyrus_punctatus | 0 | 43.22 | 2515 | 2436573.85 |
| amphibians | Anaxyrus_quercicus | Anaxyrus_quercicus | 0 | 3.47 | 3305 | 450929.87 |
| amphibians | Anaxyrus_retiformis | Anaxyrus_retiformis | 0 | 34.37 | 162.5 | 55041.86 |
| amphibians | Anaxyrus_terrestris | Anaxyrus_terrestris | 0 | 170.07 | 9120.5 | 545693.55 |
| amphibians | Anaxyrus_woodhousii | Anaxyrus_woodhousii | 0 | 226.19 | 18500 | 3242461.38 |
| amphibians | Andinobates_bombetes | Andinobates_bombetes | 1 | 0.57 | 1.5 | 569.99 |
| amphibians | Andinobates_claudiae | Andinobates_claudiae | 1 | 0.61 | 3.5 | 199.7 |
| amphibians | Andinobates_fulguritus | Andinobates_fulguritus | 0 | 0.3 | 3 | 39940.74 |
| amphibians | Andinobates_minutus | Andinobates_minutus | 0 | 0.25 | 2.5 | 55926.22 |
| amphibians | Andinobates_opisthomelas | Andinobates_opisthomelas | 1 | 0.57 | 3 | 7404.47 |
| amphibians | Andinobates_tolimense | Andinobates_tolimensis | 1 | 0.47 | 3 | 28.71 |
| amphibians | Andinobates_virolinensis | Andinobates_virolinensis | 1 | 0.47 | 2 | 3158.12 |
| amphibians | Andrias_davidianus | Andrias_davidianus | 1 | 3845.76 | 900 | 330599.84 |
| amphibians | Andrias_japonicus | Andrias_japonicus | 1 | 10792.73 | 450 | 57086.66 |
| amphibians | Aneides_aeneus | Aneides_aeneus | 0 | 2.47 | 21 | 169792.05 |
| amphibians | Aneides_ferreus | Aneides_ferreus | 0 | 9.6 | 13 | 81592.5 |
| amphibians | Aneides_flavipunctatus | Aneides_flavipunctatus | 0 | 13.48 | 17.5 | 15597.08 |
| amphibians | Aneides_hardii | Aneides_hardii | 0 | 4.13 | 10 | 16690.5 |
| amphibians | Aneides_lugubris | Aneides_lugubris | 0 | 16.14 | 14.5 | 105848.84 |
| amphibians | Aneides_vagrans | Aneides_vagrans | 0 | 7 | 7.5 | 52040.65 |
| amphibians | Anhydrophryne_hewitti | Anhydrophryne_hewitti | 0 | 3.81 | 27 | 43053.9 |
| amphibians | Anhydrophryne_ngongoniensis | Anhydrophryne_ngongoniensis | 1 | 0.77 | 12.5 | 412.41 |
| amphibians | Anhydrophryne_rattrayi | Anhydrophryne_rattrayi | 1 | 0.89 | 15.5 | 2285.03 |
| amphibians | Anodonthyla_boulengerii | Anodonthyla_boulengerii | 0 | 0.89 | 26.5 | 80954.07 |
| amphibians | Anodonthyla_hutchisoni | Anodonthyla_hutchisoni | 1 | 1.08 | 26 | 429.58 |
| amphibians | Anodonthyla_rouxae | Anodonthyla_rouxae | 1 | 3.16 | 31.5 | 103.59 |
| amphibians | Anomaloglossus_beebei | Anomaloglossus_beebei | 1 | 0.46 | 4 | 2022.89 |
| amphibians | Anomaloglossus_roraima | Anomaloglossus_roraima | 1 | 0.51 | 5 | 248.57 |

| | | | | | | |
|---|---|---|---|---|---|---|
| amphibians | Anomaloglossus_stepheni | Anomaloglossus_stepheni | 0 | 0.42 | 4.5 | 388182.01 |
| amphibians | Anomaloglossus_verbeeksnyderorum | Anomaloglossus_verbeeksnyderorum | 0 | 0.81 | 5 | 11066.93 |
| amphibians | Ansonia_hanitschi | Ansonia_hanitschi | 0 | 4.16 | 59 | 21080.18 |
| amphibians | Ansonia_malayana | Ansonia_malayana | 0 | 1.89 | 300 | 8564.77 |
| amphibians | Anstisia_alba | Geocrinia_alba | 1 | 1.17 | 10 | 215.72 |
| amphibians | Anstisia_lutea | Geocrinia_lutea | 0 | 1.17 | 20.5 | 263.57 |
| amphibians | Anstisia_rosea | Geocrinia_rosea | 0 | 1.17 | 21.5 | 2401.76 |
| amphibians | Anstisia_vitellina | Geocrinia_vitellina | 1 | 1.17 | 36 | 22.28 |
| amphibians | Aphantophryne_pansa | Aphantophryne_pansa | 0 | 2.5 | 20 | 848.61 |
| amphibians | Aphantophryne_sabini | Aphantophryne_sabini | 0 | 1.04 | 15.5 | 14.45 |
| amphibians | Aplastodiscus_arildae | Aplastodiscus_arildae | 0 | 6.08 | 307 | 166613.89 |
| amphibians | Aplastodiscus_cochranae | Aplastodiscus_cochranae | 0 | 11.25 | 333 | 31715.66 |
| amphibians | Aplastodiscus_eugenioi | Aplastodiscus_eugenioi | 0 | 4.93 | 75 | 16103.43 |
| amphibians | Aplastodiscus_leucopygius | Aplastodiscus_leucopygius | 0 | 9.03 | 224.5 | 116912.51 |
| amphibians | Aplastodiscus_perviridis | Aplastodiscus_perviridis | 0 | 9.03 | 194.5 | 234555.88 |
| amphibians | Aquarana_catesbeiana | Rana_catesbeiana | 0 | 1033.8 | 24400 | 7019850.75 |
| amphibians | Aquarana_clamitans | Rana_clamitans | 0 | 122.13 | 3632.5 | 3829381.03 |
| amphibians | Aquarana_grylio | Rana_grylio | 0 | 528.2 | 19500 | 365850.88 |
| amphibians | Aquarana_heckscheri | Rana_heckscheri | 0 | 431.35 | 6500 | 253556.06 |
| amphibians | Aquarana_okaloosae | Rana_okaloosae | 1 | 10.89 | 248.5 | 492.62 |
| amphibians | Aquarana_septentrionalis | Rana_septentrionalis | 0 | 42.85 | 2250 | 1981074.12 |
| amphibians | Aquarana_virgatipes | Rana_virgatipes | 0 | 37.61 | 400 | 166516.82 |
| amphibians | Arcovomer_passarellii | Arcovomer_passarellii | 0 | 1.33 | 136.5 | 42462.21 |
| amphibians | Arenophryne_rotunda | Arenophryne_rotunda | 0 | 4.93 | 7.5 | 1871.97 |
| amphibians | Arlequinus_krebsi | Arlequinus_krebsi | 1 | 2.35 | 15.5 | 70.37 |
| amphibians | Arthroleptella_bicolor | Arthroleptella_bicolor | 0 | 0.77 | 9 | 2040.15 |
| amphibians | Arthroleptella_drewesii | Arthroleptella_drewesii | 0 | 0.66 | 10 | 44.04 |
| amphibians | Arthroleptella_lightfooti | Arthroleptella_lightfooti | 0 | 0.77 | 8.5 | 253.97 |
| amphibians | Arthroleptella_subvoce | Arthroleptella_subvoce | 1 | 0.18 | 9 | 41.22 |
| amphibians | Arthroleptella_villiersi | Arthroleptella_villiersi | 0 | 0.77 | 11 | 5675.61 |
| amphibians | Arthroleptides_yakusini | Arthroleptides_yakusini | 1 | 37.61 | 105 | 3113.2 |
| amphibians | Arthroleptis_crusculum | Arthroleptis_crusculum | 0 | 0.66 | 15 | 1521.83 |
| amphibians | Arthroleptis_langeri | Arthroleptis_langeri | 1 | 0.51 | 7 | 264.23 |
| amphibians | Arthroleptis_poecilonotus | Arthroleptis_poecilonotus | 0 | 3.16 | 20 | 2565497.12 |

| | | | | | | |
|---|---|---|---|---|---|---|
| amphibians | Arthroleptis_schubotzi | Arthroleptis_schubotzi | 0 | 0.89 | 14.5 | 27346.81 |
| amphibians | Arthroleptis_stenodactylus | Arthroleptis_stenodactylus | 0 | 7.84 | 55 | 2386920.61 |
| amphibians | Arthroleptis_wahlbergii | Arthroleptis_wahlbergii | 0 | 3.35 | 20.5 | 50856.12 |
| amphibians | Arthroleptis_xenochirus | Arthroleptis_xenochirus | 0 | 1.02 | 9 | 1044689.79 |
| amphibians | Arthroleptis_xenodactyloides | Arthroleptis_xenodactyloides | 0 | 1.17 | 21.5 | 1109401.59 |
| amphibians | Ascaphus_montanus | Ascaphus_montanus | 0 | 15.93 | 60 | 239008.97 |
| amphibians | Ascaphus_truei | Ascaphus_truei | 0 | 11.77 | 57.5 | 304263.38 |
| amphibians | Assa_darlingtoni | Assa_darlingtoni | 1 | 2.11 | 13 | 6062.97 |
| amphibians | Atelopus_bomolochos | Atelopus_bomolochos | 1 | 11.77 | 251.19 | 3922.28 |
| amphibians | Atelopus_carbonerensis | Atelopus_carbonerensis | 1 | 14.68 | 279.71 | 142.87 |
| amphibians | Atelopus_cruciger | Atelopus_cruciger | 1 | 11.04 | 270.5 | 16489.97 |
| amphibians | Atelopus_elegans | Atelopus_elegans | 1 | 3.71 | 370 | 8362.61 |
| amphibians | Atelopus_exiguus | Atelopus_exiguus | 1 | 3.61 | 1200 | 410.87 |
| amphibians | Atelopus_franciscus | Atelopus_franciscus | 0 | 1.41 | 250 | 35332.26 |
| amphibians | Atelopus_ignescens | Atelopus_ignescens | 1 | 9.8 | 340.5 | 10227.64 |
| amphibians | Atelopus_mucubajiensis | Atelopus_mucubajiensis | 1 | 7.29 | 251.19 | 10.7 |
| amphibians | Atelopus_muisca | Atelopus_muisca | 1 | 6.47 | 69 | 87.36 |
| amphibians | Atelopus_onorei | Atelopus_onorei | 1 | 9.6 | 133 | 11.15 |
| amphibians | Atelopus_patazensis | Atelopus_patazensis | 1 | 10.2 | 348 | 27.74 |
| amphibians | Atelopus_pulcher | Atelopus_pulcher | 1 | 5.36 | 600 | 9255.54 |
| amphibians | Atelopus_sernai | Atelopus_sernai | 1 | 2.96 | 219.5 | 318.58 |
| amphibians | Atelopus_sorianoi | Atelopus_sorianoi | 1 | 11.4 | 251.19 | 5.89 |
| amphibians | Atelopus_spumarius | Atelopus_spumarius | 0 | 5.36 | 341.5 | 90169.92 |
| amphibians | Atelopus_subornatus | Atelopus_subornatus | 1 | 4.93 | 189.39 | 617.18 |
| amphibians | Atelopus_varius | Atelopus_varius | 1 | 19.92 | 485 | 10807.36 |
| amphibians | Atelopus_zeteki | Atelopus_zeteki | 1 | 23.34 | 411.5 | 580.72 |
| amphibians | Atlantihyla_spinipollex | Ptychohyla_spinipollex | 1 | 8.48 | 200 | 445.95 |
| amphibians | Atympanophrys_gigantica | Xenophrys_gigantica | 0 | 165.84 | 1192 | 5315.76 |
| amphibians | Austrochaperina_adelphe | Austrochaperina_adelphe | 0 | 0.77 | 12 | 82170.1 |
| amphibians | Austrochaperina_fryi | Austrochaperina_fryi | 0 | 3.47 | 10 | 4191.13 |
| amphibians | Austrochaperina_gracilipes | Austrochaperina_gracilipes | 0 | 0.89 | 8 | 98893.76 |
| amphibians | Austrochaperina_novaebritanniae | Austrochaperina_novaebritanniae | 1 | 0.66 | 5.5 | 35333.54 |
| amphibians | Austrochaperina_palmipes | Austrochaperina_palmipes | 0 | 10.34 | 15.5 | 29095.84 |
| amphibians | Austrochaperina_pluvialis | Austrochaperina_pluvialis | 0 | 1.89 | 11 | 10030.96 |

| | | | | | | |
|---|---|---|---|---|---|---|
| amphibians | Austrochaperina_robusta | Austrochaperina_robusta | 0 | 2.87 | 12 | 6472.09 |
| amphibians | Babina_holsti | Babina_holsti | 1 | 209.33 | 900 | 250.71 |
| amphibians | Babina_subaspera | Babina_subaspera | 1 | 310.17 | 1150 | 784.55 |
| amphibians | Barbarophryne_brongersmai | Barbarophryne_brongersmai | 0 | 12.69 | 690 | 131702.78 |
| amphibians | Barbourula_busuangensis | Barbourula_busuangensis | 0 | 62.29 | 78 | 1842.91 |
| amphibians | Barycholos_pulcher | Barycholos_pulcher | 0 | 2.35 | 43 | 58746.12 |
| amphibians | Batrachoseps_attenuatus | Batrachoseps_attenuatus | 0 | 1.93 | 11 | 72715.59 |
| amphibians | Batrachoseps_gregarius | Batrachoseps_gregarius | 0 | 1.6 | 19.5 | 20831.06 |
| amphibians | Batrachoseps_luciae | Batrachoseps_luciae | 0 | 1.6 | 12.5 | 1703.9 |
| amphibians | Batrachoseps_pacificus | Batrachoseps_pacificus | 0 | 5.49 | 16.5 | 513.58 |
| amphibians | Batrachoseps_wrighti | Batrachoseps_wrighti | 0 | 3.67 | 7 | 15905.03 |
| amphibians | Batrachuperus_karlschmidti | Batrachuperus_karlschmidti | 1 | 10.18 | 9.5 | 100637.11 |
| amphibians | Batrachuperus_pinchonii | Batrachuperus_pinchonii | 1 | 18.17 | 31 | 107908.57 |
| amphibians | Batrachuperus_tibetanus | Batrachuperus_tibetanus | 1 | 13.94 | 40 | 171618.72 |
| amphibians | Batrachuperus_yenyuanensis | Batrachuperus_yenyuanensis | 1 | 9 | 31 | 14635.19 |
| amphibians | Batrachyla_antartandica | Batrachyla_antartandica | 0 | 11.04 | 65 | 95777.05 |
| amphibians | Batrachyla_leptopus | Batrachyla_leptopus | 0 | 7.84 | 90 | 172540.62 |
| amphibians | Batrachyla_taeniata | Batrachyla_taeniata | 0 | 7.84 | 185 | 145739.87 |
| amphibians | Beduka_amboli | Xanthophryne_tigerina | 1 | 3.57 | 165 | 35.53 |
| amphibians | Beduka_koynayensis | Xanthophryne_koynayensis | 1 | 2.87 | 165 | 70.62 |
| amphibians | Blommersia_domerguei | Blommersia_domerguei | 0 | 0.59 | 66.5 | 51578.29 |
| amphibians | Blommersia_wittei | Blommersia_wittei | 0 | 1.5 | 70 | 69134.08 |
| amphibians | Boana_aguilari | Hypsiboas_aguilari | 0 | 11.04 | 440 | 1040.68 |
| amphibians | Boana_albomarginata | Hypsiboas_albomarginatus | 0 | 22.16 | 1247.5 | 504714.8 |
| amphibians | Boana_albopunctata | Hypsiboas_albopunctatus | 0 | 41.05 | 774.5 | 4885110.93 |
| amphibians | Boana_atlantica | Hypsiboas_atlanticus | 0 | 7.03 | 233.5 | 42358.19 |
| amphibians | Boana_balzani | Hypsiboas_balzani | 0 | 12.77 | 300 | 24585.41 |
| amphibians | Boana_bischoffi | Hypsiboas_bischoffi | 0 | 31.33 | 995.5 | 164695.23 |
| amphibians | Boana_boans | Hypsiboas_boans | 0 | 256.34 | 2980 | 7123580.79 |
| amphibians | Boana_caipora | Hypsiboas_caipora | 0 | 7.46 | 68 | 3405.13 |
| amphibians | Boana_calcarata | Hypsiboas_calcaratus | 0 | 22.16 | 893 | 4707380.25 |
| amphibians | Boana_cinerascens | Hypsiboas_cinerascens | 0 | 5.89 | 502.5 | 5780987.86 |
| amphibians | Boana_cipoensis | Hypsiboas_cipoensis | 0 | 4.35 | 161.5 | 33311.84 |
| amphibians | Boana_crepitans | Hypsiboas_crepitans | 0 | 41.05 | 1597.5 | 935349.72 |

| | | | | | | |
|---|---|---|---|---|---|---|
| amphibians | Boana_curupi | Hypsiboas_curupi | 0 | 9.03 | 263 | 76642.57 |
| amphibians | Boana_faber | Hypsiboas_faber | 0 | 118.4 | 2862 | 1288354.88 |
| amphibians | Boana_fasciata | Hypsiboas_fasciatus | 0 | 14.85 | 1057.5 | 17312.19 |
| amphibians | Boana_freicanecae | Hypsiboas_freicanecae | 1 | 6.37 | 30 | 2005.18 |
| amphibians | Boana_geographica | Hypsiboas_geographicus | 0 | 61.58 | 2243 | 6749152.75 |
| amphibians | Boana_goiana | Hypsiboas_goianus | 0 | 4.65 | 176 | 51882.46 |
| amphibians | Boana_guentheri | Hypsiboas_guentheri | 0 | 9.03 | 1000 | 23644.05 |
| amphibians | Boana_heilprini | Hypsiboas_heilprini | 1 | 14.42 | 332 | 37933.01 |
| amphibians | Boana_joaquini | Hypsiboas_joaquini | 0 | 16.3 | 300 | 11319.7 |
| amphibians | Boana_lanciformis | Hypsiboas_lanciformis | 0 | 104.27 | 1293 | 4670364.27 |
| amphibians | Boana_lemai | Hypsiboas_lemai | 0 | 5.36 | 177 | 38398.08 |
| amphibians | Boana_leptolineata | Hypsiboas_leptolineatus | 0 | 4.93 | 157.5 | 114377.64 |
| amphibians | Boana_marginata | Hypsiboas_marginatus | 0 | 16.87 | 200 | 12500.97 |
| amphibians | Boana_marianitae | Hypsiboas_marianitae | 0 | 16.68 | 300 | 70845.72 |
| amphibians | Boana_multifasciata | Hypsiboas_multifasciatus | 0 | 57.01 | 1343 | 164135.1 |
| amphibians | Boana_nympha | Hypsiboas_nympha | 0 | 3.81 | 117 | 416259.32 |
| amphibians | Boana_pellucens | Hypsiboas_pellucens | 0 | 21.7 | 1420 | 76138.72 |
| amphibians | Boana_polytaenia | Hypsiboas_polytaenius | 0 | 6.27 | 230 | 209403.86 |
| amphibians | Boana_prasina | Hypsiboas_prasinus | 0 | 15.03 | 995.5 | 80899.99 |
| amphibians | Boana_pugnax | Hypsiboas_pugnax | 0 | 50.6 | 1916.5 | 286727 |
| amphibians | Boana_pulchella | Hypsiboas_pulchellus | 0 | 11.04 | 750 | 1267157.23 |
| amphibians | Boana_punctata | Hypsiboas_punctatus | 0 | 6.13 | 298.5 | 8320786.59 |
| amphibians | Boana_raniceps | Hypsiboas_raniceps | 0 | 54.82 | 1656.5 | 7490502.56 |
| amphibians | Boana_riojana | Hypsiboas_riojanus | 0 | 22.98 | 300 | 271454.51 |
| amphibians | Boana_rosenbergi | Hypsiboas_rosenbergi | 0 | 88.3 | 1682.5 | 164358.87 |
| amphibians | Boana_rufitela | Hypsiboas_rufitelus | 0 | 15.03 | 265 | 39575.39 |
| amphibians | Boana_secedens | Hypsiboas_secedens | 1 | 16.87 | 1771 | 3504.46 |
| amphibians | Boana_semilineata | Hypsiboas_semilineatus | 0 | 15.93 | 891 | 444831.27 |
| amphibians | Boana_sibleszi | Hypsiboas_sibleszi | 0 | 3.71 | 61 | 145943.68 |
| amphibians | Boana_stellae | Hypsiboas_stellae | 0 | 15.93 | 200 | 10397.95 |
| amphibians | Bokermannohyla_alvarengai | Bokermannohyla_alvarengai | 0 | 316.68 | 1471.5 | 60203.62 |
| amphibians | Bokermannohyla_circumdata | Bokermannohyla_circumdata | 0 | 34.37 | 342.5 | 948421.36 |
| amphibians | Bokermannohyla_ibitiguara | Bokermannohyla_ibitiguara | 0 | 9.93 | 237.5 | 12112.51 |
| amphibians | Bokermannohyla_itapoty | Bokermannohyla_itapoty | 0 | 11.19 | 315.5 | 818.39 |

| | | | | | | |
|---|---|---|---|---|---|---|
| amphibians | Bokermannohyla_luctuosa | Bokermannohyla_luctuosa | 0 | 32.83 | 504.5 | 42440.04 |
| amphibians | Bokermannohyla_nanuzae | Bokermannohyla_nanuzae | 0 | 7.29 | 105 | 27757.77 |
| amphibians | Bokermannohyla_oxente | Bokermannohyla_oxente | 0 | 10.34 | 302.5 | 9126.44 |
| amphibians | Bokermannohyla_saxicola | Bokermannohyla_saxicola | 0 | 15.03 | 200 | 74214.51 |
| amphibians | Bolitoglossa_adspersa | Bolitoglossa_adspersa | 0 | 2.66 | 10 | 22488.82 |
| amphibians | Bolitoglossa_altamazonica | Bolitoglossa_altamazonica | 0 | 2.94 | 9.5 | 1058929.16 |
| amphibians | Bolitoglossa_mexicana | Bolitoglossa_mexicana | 0 | 8.75 | 63 | 205820.18 |
| amphibians | Bolitoglossa_orestes | Bolitoglossa_orestes | 1 | 1.24 | 2545 | 1026.56 |
| amphibians | Bolitoglossa_palmata | Bolitoglossa_palmata | 0 | 2.47 | 2053 | 5733.58 |
| amphibians | Bolitoglossa_pesrubra | Bolitoglossa_pesrubra | 0 | 4.83 | 25.5 | 629.99 |
| amphibians | Bolitoglossa_rostrata | Bolitoglossa_rostrata | 0 | 3.66 | 37 | 961.32 |
| amphibians | Bolitoglossa_subpalmata | Bolitoglossa_subpalmata | 0 | 3.49 | 23 | 1237.4 |
| amphibians | Bombina_bombina | Bombina_bombina | 0 | 19.92 | 503 | 3052238.79 |
| amphibians | Bombina_orientalis | Bombina_orientalis | 0 | 25.82 | 166.5 | 942069.95 |
| amphibians | Bombina_variegata | Bombina_variegata | 0 | 15.93 | 107.5 | 1205595 |
| amphibians | Boophis_albilabris | Boophis_albilabris | 0 | 94.79 | 725 | 82597.19 |
| amphibians | Boophis_doulioti | Boophis_doulioti | 0 | 12.85 | 3351 | 243894.94 |
| amphibians | Boophis_goudotii | Boophis_goudotii | 0 | 105.97 | 1000 | 52329.98 |
| amphibians | Boophis_luteus | Boophis_luteus | 0 | 19.92 | 560 | 40589.22 |
| amphibians | Boophis_madagascariensis | Boophis_madagascariensis | 0 | 114.75 | 402.5 | 84306.25 |
| amphibians | Boophis_marojezensis | Boophis_marojezensis | 0 | 2.87 | 133 | 43462.67 |
| amphibians | Boophis_miniatus | Boophis_miniatus | 1 | 4.54 | 248 | 7853.75 |
| amphibians | Boophis_occidentalis | Boophis_occidentalis | 0 | 12.85 | 1342 | 575.25 |
| amphibians | Boophis_picturatus | Boophis_picturatus | 0 | 2.96 | 50 | 28021.68 |
| amphibians | Boophis_rappiodes | Boophis_rappiodes | 0 | 3.16 | 625 | 31810.04 |
| amphibians | Boophis_tampoka | Boophis_tampoka | 0 | 8.42 | 240 | 42.88 |
| amphibians | Boophis_tephraeomystax | Boophis_tephraeomystax | 0 | 32.83 | 1000 | 112253.61 |
| amphibians | Boophis_tsilomaro | Boophis_tsilomaro | 1 | 61.58 | 1342 | 89.93 |
| amphibians | Boophis_viridis | Boophis_viridis | 0 | 3.47 | 154 | 64241.13 |
| amphibians | Boophis_xerophilus | Boophis_xerophilus | 0 | 10.34 | 2617 | 63546.65 |
| amphibians | Boreorana_sylvatica | Rana_sylvatica | 0 | 57.01 | 2520 | 9135971.74 |
| amphibians | Boulengerula_boulengeri | Boulengerula_boulengeri | 0 | 10.95 | 3 | 1853.53 |
| amphibians | Boulengerula_taitana | Boulengerula_taitana | 1 | 27.94 | 8.5 | 340.74 |
| amphibians | Boulenophrys_boettgeri | Xenophrys_boettgeri | 0 | 8.91 | 200 | 367630.19 |

| | | | | | | |
|---|---|---|---|---|---|---|
| amphibians | Boulenophrys_brachykolos | Xenophrys_brachykolos | 1 | 9.67 | 305 | 261.01 |
| amphibians | Boulenophrys_jingdongensis | Xenophrys_jingdongensis | 0 | 23.94 | 215 | 58231.51 |
| amphibians | Boulenophrys_kuatunensis | Xenophrys_kuatunensis | 0 | 4.31 | 54 | 231307.51 |
| amphibians | Boulenophrys_omeimontis | Xenophrys_omeimontis | 0 | 36.78 | 3141 | 25114.39 |
| amphibians | Boulenophrys_sangzhiensis | Xenophrys_sangzhiensis | 1 | 14.76 | 255 | 10.67 |
| amphibians | Brachycephalus_ephippium | Brachycephalus_ephippium | 0 | 0.57 | 5 | 57893.63 |
| amphibians | Brachycephalus_ferruginus | Brachycephalus_ferruginus | 0 | 0.2 | 3 | 113.61 |
| amphibians | Brachytarsophrys_feae | Brachytarsophrys_feae | 0 | 157.16 | 579 | 343348.74 |
| amphibians | Breviceps_acutirostris | Breviceps_acutirostris | 0 | 5.36 | 24 | 6682.48 |
| amphibians | Breviceps_adspersus | Breviceps_adspersus | 0 | 19.92 | 39.5 | 2329054.01 |
| amphibians | Breviceps_fuscus | Breviceps_fuscus | 0 | 11.77 | 43.5 | 7155.28 |
| amphibians | Breviceps_gibbosus | Breviceps_gibbosus | 0 | 50.6 | 26 | 3287.49 |
| amphibians | Breviceps_macrops | Breviceps_macrops | 0 | 14.16 | 43 | 5555.54 |
| amphibians | Breviceps_mossambicus | Breviceps_mossambicus | 0 | 17.85 | 25 | 1579734.52 |
| amphibians | Breviceps_sylvestris | Breviceps_sylvestris | 0 | 13.33 | 53 | 5575.65 |
| amphibians | Breviceps_verrucosus | Breviceps_verrucosus | 0 | 13.33 | 35 | 106155.59 |
| amphibians | Bromeliohyla_bromeliacia | Bromeliohyla_bromeliacia | 0 | 3.71 | 12.5 | 6976.63 |
| amphibians | Bryophryne_cophites | Bryophryne_cophites | 1 | 3.74 | 20 | 42.48 |
| amphibians | Buergeria_buergeri | Buergeria_buergeri | 0 | 42.85 | 418 | 284874.19 |
| amphibians | Buergeria_japonica | Buergeria_japonica | 0 | 4.16 | 319.5 | 2643.66 |
| amphibians | Buergeria_robusta | Buergeria_robusta | 0 | 28.62 | 475 | 25850.21 |
| amphibians | Bufo_aspinius | Bufo_aspinius | 1 | 114.75 | 2250 | 1323.48 |
| amphibians | Bufo_bankorensis | Bufo_bankorensis | 0 | 152.72 | 4362.5 | 21413.55 |
| amphibians | Bufo_bufo | Bufo_bufo | 0 | 1341.53 | 7500 | 9388939.92 |
| amphibians | Bufo_gargarizans | Bufo_gargarizans | 0 | 269.14 | 7600 | 5233722.26 |
| amphibians | Bufo_praetextatus | Bufo_japonicus | 0 | 651.05 | 10000 | 295039.75 |
| amphibians | Bufo_spinosus | Bufo_spinosus | 0 | 1341.53 | 6500 | 1155874.72 |
| amphibians | Bufo_stejnegeri | Bufo_stejnegeri | 0 | 22.16 | 830 | 150839.18 |
| amphibians | Bufo_torrenticola | Bufo_torrenticola | 0 | 559.96 | 3250 | 21579.83 |
| amphibians | Bufo_verrucosissimus | Bufo_verrucosissimus | 0 | 834.28 | 5345 | 134366.24 |
| amphibians | Bufotes_boulengeri | Bufotes_boulengeri | 0 | 129.82 | 20000 | 1966248.49 |
| amphibians | Bufotes_pewzowi | Bufotes_pewzowi | 0 | 54.6 | 3250 | 1286335.42 |
| amphibians | Bufotes_viridis | Bufotes_viridis | 0 | 310.17 | 19475 | 10026528.87 |
| amphibians | Cacosternum_boettgeri | Cacosternum_boettgeri | 0 | 1.33 | 140 | 2352879.44 |

| | | | | | | |
|---|---|---|---|---|---|---|
| amphibians | Cacosternum_capense | Cacosternum_capense | 0 | 4.93 | 228.5 | 6661.78 |
| amphibians | Cacosternum_karooicum | Cacosternum_karooicum | 0 | 2.35 | 323 | 47401.51 |
| amphibians | Cacosternum_namaquense | Cacosternum_namaquense | 0 | 1.87 | 56 | 45611.11 |
| amphibians | Cacosternum_nanum | Cacosternum_nanum | 0 | 1.17 | 16.5 | 229198.21 |
| amphibians | Cacosternum_plimptoni | Cacosternum_plimptoni | 0 | 1.17 | 250 | 44886.59 |
| amphibians | Caecilia_orientalis | Caecilia_orientalis | 0 | 116.06 | 75 | 1511.45 |
| amphibians | Callimedusa_atelopoides | Phyllomedusa_atelopoides | 0 | 10.34 | 21.5 | 815329.15 |
| amphibians | Callimedusa_perinesos | Phyllomedusa_perinesos | 1 | 26.08 | 63 | 533.44 |
| amphibians | Callimedusa_tomopterna | Phyllomedusa_tomopterna | 0 | 53.1 | 105.5 | 5426896.3 |
| amphibians | Callulina_dawida | Callulina_dawida | 1 | 11.04 | 40 | 249.82 |
| amphibians | Callulina_kreffti | Callulina_kreffti | 0 | 9.03 | 35 | 5605.07 |
| amphibians | Callulops_robustus | Callulops_robustus | 0 | 46.62 | 17 | 582358 |
| amphibians | Calotriton_arnoldi | Calotriton_arnoldi | 1 | 3.32 | 20 | 289.55 |
| amphibians | Calotriton_asper | Calotriton_asper | 0 | 9.72 | 43.5 | 27428.33 |
| amphibians | Calyptocephalella_gayi | Calyptocephalella_gayi | 1 | 4516.87 | 5400 | 137474.05 |
| amphibians | Capensibufo_rosei | Capensibufo_rosei | 1 | 4.93 | 140 | 228.79 |
| amphibians | Capensibufo_tradouwi | Capensibufo_tradouwi | 0 | 9.67 | 75 | 27722.47 |
| amphibians | Cardioglossa_leucomystax | Cardioglossa_leucomystax | 0 | 5.23 | 25 | 1490484.84 |
| amphibians | Cardioglossa_occidentalis | Cardioglossa_occidentalis | 0 | 3.41 | 33 | 209290.82 |
| amphibians | Celsiella_revocata | Celsiella_revocata | 1 | 1.2 | 40 | 684.41 |
| amphibians | Celsiella_vozmedianoi | Celsiella_vozmedianoi | 1 | 1.77 | 20 | 47.21 |
| amphibians | Centrolene_altitudinalis | Centrolene_altitudinale | 1 | 1.09 | 20 | 322.73 |
| amphibians | Centrolene_ballux | Centrolene_ballux | 1 | 0.93 | 16.5 | 481.54 |
| amphibians | Centrolene_buckleyi | Centrolene_buckleyi | 1 | 3.38 | 18 | 74234 |
| amphibians | Centrolene_geckoidea | Centrolene_geckoideum | 1 | 52.05 | 100.5 | 8570.5 |
| amphibians | Centrolene_heloderma | Centrolene_heloderma | 1 | 2.68 | 28.5 | 4737.58 |
| amphibians | Centrolene_lynchi | Centrolene_lynchi | 1 | 1.39 | 22 | 672.68 |
| amphibians | Centrolene_notosticta | Centrolene_notostictum | 0 | 1.15 | 16.5 | 5600.18 |
| amphibians | Centrolene_peristicta | Centrolene_peristictum | 0 | 0.66 | 23.5 | 10885.61 |
| amphibians | Centrolene_pipilata | Centrolene_pipilatum | 1 | 0.9 | 18 | 35.33 |
| amphibians | Centrolene_sabini | Centrolene_sabini | 1 | 2.35 | 40 | 9.78 |
| amphibians | Centrolene_sanchezi | Centrolene_sanchezi | 1 | 0.63 | 16 | 337.68 |
| amphibians | Centrolene_savagei | Centrolene_savagei | 0 | 1.02 | 19 | 6382.97 |
| amphibians | Ceratophrys_calcarata | Ceratophrys_calcarata | 0 | 74.11 | 380 | 134165.12 |

| | | | | | | |
|---|---|---|---|---|---|---|
| amphibians | Ceratophrys_cornuta | Ceratophrys_cornuta | 0 | 175.82 | 1285 | 5479643.7 |
| amphibians | Ceratophrys_cranwelli | Ceratophrys_cranwelli | 0 | 458.99 | 3696 | 1124371.25 |
| amphibians | Ceratophrys_joazeirensis | Ceratophrys_joazeirensis | 0 | 122.13 | 2676 | 580713.41 |
| amphibians | Ceratophrys_ornata | Ceratophrys_ornata | 0 | 528.2 | 1250 | 513220.27 |
| amphibians | Ceratophrys_stolzmanni | Ceratophrys_stolzmanni | 1 | 54.82 | 1150 | 11075.09 |
| amphibians | Chaltenobatrachus_grandisonae | Chaltenobatrachus_grandisonae | 0 | 8.42 | 29 | 29846.94 |
| amphibians | Chaperina_fusca | Chaperina_fusca | 0 | 1.33 | 8 | 169007.35 |
| amphibians | Charadrahyla_nephila | Charadrahyla_nephila | 1 | 52.68 | 1541.5 | 1925.89 |
| amphibians | Charadrahyla_taeniopus | Charadrahyla_taeniopus | 1 | 32.83 | 265 | 6562.93 |
| amphibians | Chiasmocleis_antenori | Syncope_antenori | 0 | 0.17 | 5 | 609221.62 |
| amphibians | Chiasmocleis_atlantica | Chiasmocleis_atlantica | 0 | 2.6 | 730 | 11602.14 |
| amphibians | Chiasmocleis_avilapiresae | Chiasmocleis_avilapiresae | 0 | 4.46 | 1920 | 1182660.33 |
| amphibians | Chiasmocleis_bassleri | Syncope_bassleri | 0 | 2.09 | 175 | 3212224.95 |
| amphibians | Chiasmocleis_carvalhoi | Chiasmocleis_carvalhoi | 0 | 1.08 | 200 | 122088.12 |
| amphibians | Chiasmocleis_leucosticta | Chiasmocleis_leucosticta | 0 | 1.33 | 201.5 | 167239.02 |
| amphibians | Chiasmocleis_magnova | Syncope_magnova | 0 | 0.43 | 10 | 2542.85 |
| amphibians | Chiasmocleis_mehelyi | Chiasmocleis_mehelyi | 0 | 1.09 | 216.5 | 45686.35 |
| amphibians | Chiasmocleis_shudikarensis | Chiasmocleis_shudikarensis | 0 | 2.6 | 398 | 2076572.05 |
| amphibians | Chiasmocleis_tridactyla | Syncope_tridactyla | 0 | 0.12 | 3 | 349221.27 |
| amphibians | Chiasmocleis_ventrimaculata | Chiasmocleis_ventrimaculata | 0 | 3.16 | 281 | 1805618.13 |
| amphibians | Chioglossa_lusitanica | Chioglossa_lusitanica | 0 | 2.14 | 22 | 54174.61 |
| amphibians | Chirixalus_doriae | Chiromantis_doriae | 0 | 1.57 | 585 | 1893691.98 |
| amphibians | Chirixalus_dudhwaensis | Chiromantis_dudhwaensis | 1 | 5.36 | 216 | 47.83 |
| amphibians | Chirixalus_nongkhorensis | Chiromantis_nongkhorensis | 0 | 3.81 | 200 | 1344686.89 |
| amphibians | Chirixalus_simus | Chiromantis_simus | 0 | 0.77 | 193.5 | 169100.98 |
| amphibians | Chiromantis_petersii | Chiromantis_petersii | 0 | 25.82 | 325 | 732830.4 |
| amphibians | Chiromantis_rufescens | Chiromantis_rufescens | 0 | 25.82 | 157 | 2415856.56 |
| amphibians | Chiromantis_xerampelina | Chiromantis_xerampelina | 0 | 74.11 | 688 | 2783181.51 |
| amphibians | Chthonerpeton_indistinctum | Chthonerpeton_indistinctum | 0 | 32.16 | 7 | 113033.66 |
| amphibians | Chthonerpeton_noctinectes | Chthonerpeton_noctinectes | 0 | 15.43 | 8 | 2598.91 |
| amphibians | Churamiti_maridadi | Churamiti_maridadi | 1 | 16.3 | 195 | 94.8 |
| amphibians | Cochranella_euknemos | Cochranella_euknemos | 0 | 2.6 | 51 | 25146.67 |
| amphibians | Cochranella_granulosa | Cochranella_granulosa | 0 | 2.6 | 39 | 94988.32 |
| amphibians | Cochranella_mache | Cochranella_mache | 0 | 2.99 | 30 | 29709.15 |

| | | | | | | |
|---|---|---|---|---|---|---|
| amphibians | Cochranella_resplendens | Cochranella_resplendens | 0 | 1.55 | 74 | 147868.57 |
| amphibians | Colostethus_panamansis | Colostethus_panamansis | 0 | 1.93 | 27.5 | 28472.76 |
| amphibians | Conraua_crassipes | Conraua_crassipes | 0 | 52.68 | 281.5 | 557445.38 |
| amphibians | Conraua_goliath | Conraua_goliath | 1 | 5497.22 | 100 | 69715.98 |
| amphibians | Cophixalus_aenigma | Cophixalus_aenigma | 1 | 0.89 | 14.5 | 360.47 |
| amphibians | Cophixalus_australis | Cophixalus_australis | 0 | 1.89 | 13.5 | 5598.8 |
| amphibians | Cophixalus_bombiens | Cophixalus_bombiens | 0 | 0.33 | 6.5 | 2471.1 |
| amphibians | Cophixalus_concinnus | Cophixalus_concinnus | 1 | 1.41 | 17 | 11.83 |
| amphibians | Cophixalus_exiguus | Cophixalus_exiguus | 0 | 0.48 | 9 | 99.51 |
| amphibians | Cophixalus_hinchinbrookensis | Cophixalus_hinchinbrookensis | 0 | 1.33 | 9 | 102.78 |
| amphibians | Cophixalus_hosmeri | Cophixalus_hosmeri | 1 | 0.33 | 5.5 | 224.65 |
| amphibians | Cophixalus_infacetus | Cophixalus_infacetus | 0 | 0.4 | 11 | 4731.32 |
| amphibians | Cophixalus_kulakula | Cophixalus_kulakula | 0 | 9.67 | 47 | 63.48 |
| amphibians | Cophixalus_mcdonaldi | Cophixalus_mcdonaldi | 1 | 1.33 | 17 | 11.62 |
| amphibians | Cophixalus_monticola | Cophixalus_monticola | 1 | 1.02 | 10.5 | 11.81 |
| amphibians | Cophixalus_neglectus | Cophixalus_neglectus | 1 | 1.89 | 16.5 | 66.52 |
| amphibians | Cophixalus_ornatus | Cophixalus_ornatus | 0 | 1.89 | 12.5 | 2804.62 |
| amphibians | Cophixalus_parkeri | Cophixalus_parkeri | 0 | 2.9 | 18 | 14025.52 |
| amphibians | Cophixalus_saxatilis | Cophixalus_saxatilis | 0 | 9.03 | 13 | 14.1 |
| amphibians | Cophixalus_sphagnicola | Cophixalus_sphagnicola | 1 | 0.62 | 20 | 496.65 |
| amphibians | Cophyla_phyllodactyla | Cophyla_phyllodactyla | 0 | 2.11 | 50 | 9136.55 |
| amphibians | Cornufer_guentheri | Ceratobatrachus_guentheri | 0 | 104.27 | 20 | 29925.08 |
| amphibians | Cornufer_pelewensis | Platymantis_pelewensis | 0 | 19.92 | 30 | 467.03 |
| amphibians | Cornufer_vitianus | Platymantis_vitianus | 0 | 141.99 | 50 | 9910.78 |
| amphibians | Cornufer_vitiensis | Platymantis_vitiensis | 0 | 19.92 | 30 | 16546.65 |
| amphibians | Corythomantis_greeningi | Corythomantis_greeningi | 0 | 66.4 | 700 | 1230069.47 |
| amphibians | Craugastor_angelicus | Craugastor_angelicus | 1 | 41.05 | 64 | 1469.47 |
| amphibians | Craugastor_augusti | Craugastor_augusti | 0 | 88.3 | 43 | 934920.21 |
| amphibians | Craugastor_aurilegulus | Craugastor_aurilegulus | 1 | 51.63 | 80 | 1673.91 |
| amphibians | Craugastor_bransfordii | Craugastor_bransfordii | 0 | 2.11 | 6 | 62031.14 |
| amphibians | Craugastor_crassidigitus | Craugastor_crassidigitus | 0 | 9.67 | 35 | 84725.12 |
| amphibians | Craugastor_fitzingeri | Craugastor_fitzingeri | 0 | 13.33 | 58 | 282360.7 |
| amphibians | Craugastor_psephosypharus | Craugastor_psephosypharus | 0 | 26.21 | 81 | 5869.67 |
| amphibians | Crinia_bilingua | Crinia_bilingua | 0 | 0.89 | 180 | 602369.9 |

| | | | | | | |
|---|---|---|---|---|---|---|
| amphibians | Crinia_deserticola | Crinia_deserticola | 0 | 0.57 | 142.5 | 2649360.65 |
| amphibians | Crinia_georgiana | Crinia_georgiana | 0 | 7.84 | 175 | 115254.34 |
| amphibians | Crinia_glauerti | Crinia_glauerti | 0 | 1.02 | 60 | 47392.03 |
| amphibians | Crinia_insignifera | Crinia_insignifera | 0 | 2.11 | 145 | 12384.67 |
| amphibians | Crinia_nimbus | Crinia_nimbus | 0 | 2.11 | 10 | 3508.74 |
| amphibians | Crinia_parinsignifera | Crinia_parinsignifera | 0 | 0.89 | 136.5 | 1121430.76 |
| amphibians | Crinia_pseudinsignifera | Crinia_pseudinsignifera | 0 | 2.11 | 115 | 285007.4 |
| amphibians | Crinia_riparia | Crinia_riparia | 1 | 1.17 | 94 | 1463.14 |
| amphibians | Crinia_signifera | Crinia_signifera | 0 | 3.47 | 247 | 1079554.52 |
| amphibians | Crinia_tasmaniensis | Crinia_tasmaniensis | 0 | 2.11 | 57.5 | 59170 |
| amphibians | Crinia_tinnula | Crinia_tinnula | 0 | 2.11 | 75 | 47281.54 |
| amphibians | Crossodactylus_gaudichaudii | Crossodactylus_gaudichaudii | 0 | 3.51 | 144 | 49389.17 |
| amphibians | Cruziohyla_calcarifer | Cruziohyla_calcarifer | 0 | 70.96 | 45 | 27099.78 |
| amphibians | Cruziohyla_craspedopus | Cruziohyla_craspedopus | 0 | 41.95 | 13 | 2096426.8 |
| amphibians | Cryptobatrachus_boulengeri | Cryptobatrachus_boulengeri | 1 | 3.47 | 48 | 10173.83 |
| amphibians | Cryptobatrachus_fuhrmanni | Cryptobatrachus_fuhrmanni | 0 | 37.61 | 24.5 | 8373.47 |
| amphibians | Cryptobatrachus_pedroruizi | Cryptobatrachus_pedroruizi | 1 | 37.11 | 29.5 | 4803.16 |
| amphibians | Cryptobatrachus_ruthveni | Cryptobatrachus_ruthveni | 1 | 24.19 | 35 | 411.43 |
| amphibians | Cryptobranchus_alleganiensis | Cryptobranchus_alleganiensis | 1 | 923.27 | 300.5 | 435321.18 |
| amphibians | Cryptothylax_greshoffii | Cryptothylax_greshoffii | 0 | 19.92 | 15 | 1418395.85 |
| amphibians | Cryptotriton_sierraminensis | Cryptotriton_sierraminensis | 1 | 0.74 | 29.5 | 53.54 |
| amphibians | Ctenophryne_aterrima | Ctenophryne_aterrima | 0 | 32.83 | 75 | 153830.15 |
| amphibians | Ctenophryne_barbatula | Ctenophryne_barbatula | 1 | 1.5 | 22 | 9.95 |
| amphibians | Ctenophryne_carpish | Ctenophryne_carpish | 1 | 3.32 | 83 | 61.49 |
| amphibians | Ctenophryne_geayi | Ctenophryne_geayi | 0 | 15.12 | 685 | 6199340.37 |
| amphibians | Cycloramphus_boraceiensis | Cycloramphus_boraceiensis | 0 | 18.87 | 89.5 | 6189.13 |
| amphibians | Cycloramphus_carvalhoi | Zachaenus_carvalhoi | 0 | 4.16 | 37 | 23506.04 |
| amphibians | Cycloramphus_dubius | Cycloramphus_dubius | 0 | 20.25 | 55 | 4376.42 |
| amphibians | Cycloramphus_faustoi | Cycloramphus_faustoi | 1 | 7.29 | 25.5 | 1.8 |
| amphibians | Cycloramphus_organensis | Cycloramphus_organensis | 1 | 2.96 | 21 | 75.87 |
| amphibians | Cycloramphus_stejnegeri | Cycloramphus_stejnegeri | 1 | 15.93 | 30 | 7.67 |
| amphibians | Cynops_pyrrhogaster | Cynops_pyrrhogaster | 0 | 6.73 | 128 | 287453.99 |
| amphibians | Dendrobates_auratus | Dendrobates_auratus | 0 | 6.27 | 8.5 | 116892.07 |
| amphibians | Dendrobates_leucomelas | Dendrobates_leucomelas | 0 | 11.04 | 7 | 844723.62 |

| | | | | | | |
|---|---|---|---|---|---|---|
| amphibians | Dendrobates_tinctorius | Dendrobates_tinctorius | 0 | 11.04 | 10 | 528727.86 |
| amphibians | Dendrobates_truncatus | Dendrobates_truncatus | 0 | 2.35 | 5 | 66001.85 |
| amphibians | Dendrophryniscus_brevipollicatus | Dendrophryniscus_brevipollicatus | 0 | 1.17 | 37 | 3288.22 |
| amphibians | Dendropsophus_berthalutzae | Dendropsophus_berthalutzae | 0 | 1.02 | 56.5 | 122224.43 |
| amphibians | Dendropsophus_bifurcus | Dendropsophus_bifurcus | 0 | 3.47 | 203 | 1059615.48 |
| amphibians | Dendropsophus_bokermanni | Dendropsophus_bokermanni | 0 | 1.69 | 128 | 1374636.81 |
| amphibians | Dendropsophus_brevifrons | Dendropsophus_brevifrons | 0 | 1.17 | 74.5 | 1471694.42 |
| amphibians | Dendropsophus_carnifex | Dendropsophus_carnifex | 0 | 2.87 | 100 | 3308.19 |
| amphibians | Dendropsophus_ebraccatus | Dendropsophus_ebraccatus | 0 | 4.16 | 155.5 | 234634.69 |
| amphibians | Dendropsophus_elegans | Dendropsophus_elegans | 0 | 3.71 | 355.5 | 545786.36 |
| amphibians | Dendropsophus_giesleri | Dendropsophus_giesleri | 0 | 3.77 | 190 | 327238.31 |
| amphibians | Dendropsophus_haddadi | Dendropsophus_haddadi | 0 | 1.02 | 45 | 112513.18 |
| amphibians | Dendropsophus_haraldschultzi | Dendropsophus_haraldschultzi | 0 | 1.17 | 216 | 979393.96 |
| amphibians | Dendropsophus_joannae | Dendropsophus_joannae | 0 | 0.62 | 207 | 354126.84 |
| amphibians | Dendropsophus_leali | Dendropsophus_leali | 0 | 1.69 | 688 | 1207985.64 |
| amphibians | Dendropsophus_leucophyllatus | Dendropsophus_leucophyllatus | 0 | 11.04 | 557 | 2574716.36 |
| amphibians | Dendropsophus_marmoratus | Dendropsophus_marmoratus | 0 | 15.93 | 883 | 4020292.39 |
| amphibians | Dendropsophus_microcephalus | Dendropsophus_microcephalus | 0 | 2.6 | 217 | 4742856.85 |
| amphibians | Dendropsophus_microps | Dendropsophus_microps | 0 | 2.87 | 325 | 350316.31 |
| amphibians | Dendropsophus_minusculus | Dendropsophus_minusculus | 0 | 1.02 | 225 | 1240670.19 |
| amphibians | Dendropsophus_minutus | Dendropsophus_minutus | 0 | 1.69 | 287 | 10714770.45 |
| amphibians | Dendropsophus_miyatai | Dendropsophus_miyatai | 0 | 1.17 | 231 | 1186964.8 |
| amphibians | Dendropsophus_nanus | Dendropsophus_nanus | 0 | 1 | 325.5 | 8670974.69 |
| amphibians | Dendropsophus_parviceps | Dendropsophus_parviceps | 0 | 1.5 | 237 | 3216087.93 |
| amphibians | Dendropsophus_rhodopeplus | Dendropsophus_rhodopeplus | 0 | 1.89 | 333 | 2517105.76 |
| amphibians | Dendropsophus_riveroi | Dendropsophus_riveroi | 0 | 0.89 | 394 | 1957343.5 |
| amphibians | Dendropsophus_robertmertensi | Dendropsophus_robertmertensi | 0 | 1.71 | 265 | 19651.82 |
| amphibians | Dendropsophus_rossalleni | Dendropsophus_rossalleni | 0 | 2.11 | 117.5 | 2083437.49 |
| amphibians | Dendropsophus_ruschii | Dendropsophus_ruschii | 0 | 1.89 | 15 | 26721.87 |
| amphibians | Dendropsophus_sanborni | Dendropsophus_sanborni | 0 | 0.67 | 90 | 1187849.23 |
| amphibians | Dendropsophus_sarayacuensis | Dendropsophus_sarayacuensis | 0 | 7.29 | 118.5 | 3370986.48 |
| amphibians | Dendropsophus_sartori | Dendropsophus_sartori | 0 | 1.81 | 265 | 55772.47 |
| amphibians | Dendropsophus_schubarti | Dendropsophus_schubarti | 0 | 1.25 | 229 | 1871193.77 |
| amphibians | Dendropsophus_shiwiarum | Dendropsophus_shiwiarum | 0 | 0.46 | 121 | 64358.61 |

| | | | | | | |
|---|---|---|---|---|---|---|
| amphibians | Dendropsophus_soaresi | Dendropsophus_soaresi | 0 | 3.47 | 393.5 | 1428862.35 |
| amphibians | Dendropsophus_studerae | Dendropsophus_studerae | 1 | 2.02 | 415 | 96.4 |
| amphibians | Dendropsophus_timbeba | Dendropsophus_timbeba | 0 | 1.36 | 136 | 486391.55 |
| amphibians | Dendropsophus_triangulum | Dendropsophus_triangulum | 0 | 6.27 | 384.5 | 2397126.38 |
| amphibians | Dendropsophus_walfordi | Dendropsophus_walfordi | 0 | 0.57 | 181 | 1199450.83 |
| amphibians | Dendropsophus_werneri | Dendropsophus_werneri | 0 | 0.89 | 249 | 58187.86 |
| amphibians | Dermatonotus_muelleri | Dermatonotus_muelleri | 0 | 39.57 | 7343.5 | 3775072.17 |
| amphibians | Dermophis_mexicanus | Dermophis_mexicanus | 0 | 82.05 | 11 | 78747.83 |
| amphibians | Desmognathus_aeneus | Desmognathus_aeneus | 0 | 0.19 | 11.5 | 45549.23 |
| amphibians | Desmognathus_auriculatus | Desmognathus_auriculatus | 0 | 2.55 | 23.5 | 325799.61 |
| amphibians | Desmognathus_brimleyorum | Desmognathus_brimleyorum | 0 | 7.13 | 25.5 | 58242.36 |
| amphibians | Desmognathus_carolinensis | Desmognathus_carolinensis | 0 | 1.93 | 21 | 11431.37 |
| amphibians | Desmognathus_conanti | Desmognathus_conanti | 0 | 1.47 | 21.5 | 622621.85 |
| amphibians | Desmognathus_folkertsi | Desmognathus_folkertsi | 0 | 8.44 | 34 | 6364.52 |
| amphibians | Desmognathus_fuscus | Desmognathus_fuscus | 0 | 3.04 | 26.5 | 1007633.86 |
| amphibians | Desmognathus_imitator | Desmognathus_imitator | 0 | 1.6 | 21.5 | 3698.89 |
| amphibians | Desmognathus_marmoratus | Desmognathus_marmoratus | 0 | 8.87 | 42.5 | 7803.44 |
| amphibians | Desmognathus_monticola | Desmognathus_monticola | 0 | 3.81 | 25 | 324006.87 |
| amphibians | Desmognathus_ochrophaeus | Desmognathus_ochrophaeus | 0 | 1.09 | 20 | 287424.36 |
| amphibians | Desmognathus_ocoee | Desmognathus_ocoee | 0 | 1.37 | 17 | 42012.81 |
| amphibians | Desmognathus_orestes | Desmognathus_orestes | 0 | 0.98 | 13.5 | 22135.66 |
| amphibians | Desmognathus_santeetlah | Desmognathus_santeetlah | 0 | 1.52 | 22.5 | 6447.54 |
| amphibians | Desmognathus_welteri | Desmognathus_welteri | 0 | 6.22 | 37.5 | 57404.94 |
| amphibians | Desmognathus_wrighti | Desmognathus_wrighti | 0 | 0.42 | 9 | 8875.85 |
| amphibians | Diasporus_diastema | Diasporus_diastema | 0 | 1.17 | 14 | 109484.48 |
| amphibians | Diasporus_vocator | Diasporus_vocator | 0 | 0.57 | 5 | 23566.49 |
| amphibians | Dicamptodon_aterrimus | Dicamptodon_aterrimus | 0 | 24.87 | 135 | 53294.47 |
| amphibians | Dicamptodon_copei | Dicamptodon_copei | 0 | 26.8 | 69 | 34398.56 |
| amphibians | Dicamptodon_ensatus | Dicamptodon_ensatus | 0 | 78.55 | 104 | 16015.83 |
| amphibians | Dicamptodon_tenebrosus | Dicamptodon_tenebrosus | 0 | 62.43 | 141.5 | 169824.69 |
| amphibians | Didynamipus_sjostedti | Didynamipus_sjostedti | 1 | 0.57 | 18 | 7639.94 |
| amphibians | Discoglossus_galganoi | Discoglossus_galganoi | 0 | 50.6 | 910 | 488944.42 |
| amphibians | Discoglossus_montalentii | Discoglossus_montalentii | 0 | 25.82 | 1000 | 4154.89 |
| amphibians | Discoglossus_pictus | Discoglossus_pictus | 0 | 50.6 | 2828 | 356005.89 |

| | | | | | | |
|---|---|---|---|---|---|---|
| amphibians | Discoglossus_sardus | Discoglossus_sardus | 0 | 32.83 | 1000 | 32995.85 |
| amphibians | Discoglossus_scovazzi | Discoglossus_scovazzi | 0 | 50.6 | 550 | 181223.14 |
| amphibians | Dryophytes_andersonii | Hyla_andersonii | 0 | 11.77 | 509 | 67727.63 |
| amphibians | Dryophytes_arenicolor | Hyla_arenicolor | 0 | 16.97 | 232 | 1286044.86 |
| amphibians | Dryophytes_avivoca | Hyla_avivoca | 0 | 13.33 | 409 | 446462.52 |
| amphibians | Dryophytes_chrysoscelis | Hyla_chrysoscelis | 0 | 22.16 | 2493 | 3693368.95 |
| amphibians | Dryophytes_cinereus | Hyla_cinerea | 0 | 27.13 | 1595.5 | 1290978.87 |
| amphibians | Dryophytes_euphorbiaceus | Hyla_euphorbiacea | 0 | 6.98 | 265 | 10694.19 |
| amphibians | Dryophytes_eximius | Hyla_eximia | 0 | 15.93 | 265 | 357544.21 |
| amphibians | Dryophytes_femoralis | Hyla_femoralis | 0 | 7.29 | 1093 | 482056.2 |
| amphibians | Dryophytes_gratiosus | Hyla_gratiosa | 0 | 32.83 | 2295.5 | 593264.04 |
| amphibians | Dryophytes_immaculatus | Hyla_immaculata | 0 | 5.8 | 220 | 226228.58 |
| amphibians | Dryophytes_japonicus | Hyla_japonica | 0 | 9.67 | 832.5 | 2185028.15 |
| amphibians | Dryophytes_plicatus | Hyla_plicata | 0 | 9.28 | 265 | 25310.34 |
| amphibians | Dryophytes_squirellus | Hyla_squirella | 0 | 7.84 | 1156.5 | 747619.54 |
| amphibians | Dryophytes_versicolor | Hyla_versicolor | 0 | 19.92 | 1800 | 2240480.06 |
| amphibians | Dryophytes_walkeri | Hyla_walkeri | 1 | 4.46 | 1000 | 7092.26 |
| amphibians | Dryophytes_wrightorum | Hyla_wrightorum | 0 | 9.67 | 232 | 528650.58 |
| amphibians | Duellmanohyla_legleri | Ptychohyla_legleri | 1 | 4.93 | 92 | 3631.58 |
| amphibians | Duellmanohyla_rufioculis | Duellmanohyla_rufioculis | 0 | 5.36 | 173.5 | 20510.27 |
| amphibians | Duellmanohyla_salvavida | Duellmanohyla_salvavida | 1 | 4.16 | 100 | 1794.91 |
| amphibians | Duellmanohyla_schmidtorum | Duellmanohyla_schmidtorum | 0 | 4.65 | 191 | 11388.83 |
| amphibians | Duellmanohyla_uranochroa | Duellmanohyla_uranochroa | 1 | 5.36 | 69 | 7897.41 |
| amphibians | Duttaphrynus_melanostictus | Duttaphrynus_melanostictus | 0 | 310.17 | 20500 | 7410138.94 |
| amphibians | Dyscophus_antongilii | Dyscophus_antongilii | 0 | 188.23 | 1250 | 37130.04 |
| amphibians | Dyscophus_guineti | Dyscophus_guineti | 0 | 152.28 | 5000 | 6095.4 |
| amphibians | Echinotriton_andersoni | Echinotriton_andersoni | 1 | 15.68 | 58.5 | 2193.92 |
| amphibians | Echinotriton_chinhaiensis | Echinotriton_chinhaiensis | 1 | 6.71 | 113 | 114.65 |
| amphibians | Ecnomiohyla_miliaria | Ecnomiohyla_miliaria | 0 | 154.93 | 265 | 13433.21 |
| amphibians | Edalorhina_perezi | Edalorhina_perezi | 0 | 5.36 | 142 | 1481341.99 |
| amphibians | Elachistocleis_bicolor | Elachistocleis_bicolor | 0 | 11.04 | 840 | 2588357.63 |
| amphibians | Elachistocleis_cesarii | Elachistocleis_cesarii | 0 | 6.62 | 893 | 2162380.35 |
| amphibians | Elachistocleis_erythrogaster | Elachistocleis_erythrogaster | 1 | 4.54 | 700 | 5054.73 |
| amphibians | Elachistocleis_ovalis | Elachistocleis_ovalis | 0 | 7.19 | 385 | 1366726.17 |

| | | | | | | |
|---|---|---|---|---|---|---|
| amphibians | Elachistocleis_skotogaster | Elachistocleis_skotogaster | 0 | 3.29 | 1072 | 8322.22 |
| amphibians | Eleutherodactylus_adelus | Eleutherodactylus_adelus | 1 | 0.22 | 3.5 | 302.55 |
| amphibians | Eleutherodactylus_alticola | Eleutherodactylus_alticola | 1 | 1.33 | 25.5 | 60.05 |
| amphibians | Eleutherodactylus_amadeus | Eleutherodactylus_amadeus | 1 | 1.17 | 19 | 500.76 |
| amphibians | Eleutherodactylus_amplinympha | Eleutherodactylus_amplinympha | 1 | 11.04 | 18 | 217.37 |
| amphibians | Eleutherodactylus_andrewsi | Eleutherodactylus_andrewsi | 1 | 1.33 | 21 | 213.25 |
| amphibians | Eleutherodactylus_antillensis | Eleutherodactylus_antillensis | 0 | 9.03 | 33 | 9408.59 |
| amphibians | Eleutherodactylus_aporostegus | Eleutherodactylus_aporostegus | 1 | 9.67 | 18 | 4627.86 |
| amphibians | Eleutherodactylus_armstrongi | Eleutherodactylus_armstrongi | 1 | 6.77 | 40 | 985.3 |
| amphibians | Eleutherodactylus_atkinsi | Eleutherodactylus_atkinsi | 0 | 6.77 | 68 | 22235.38 |
| amphibians | Eleutherodactylus_audanti | Eleutherodactylus_audanti | 1 | 1.17 | 22 | 2879.6 |
| amphibians | Eleutherodactylus_auriculatus | Eleutherodactylus_auriculatus | 0 | 1.02 | 28 | 16894.64 |
| amphibians | Eleutherodactylus_cochranae | Eleutherodactylus_cochranae | 0 | 0.89 | 7.5 | 3963.48 |
| amphibians | Eleutherodactylus_cooki | Eleutherodactylus_cooki | 1 | 14.16 | 27 | 238.77 |
| amphibians | Eleutherodactylus_coqui | Eleutherodactylus_coqui | 0 | 23.34 | 24 | 13154.86 |
| amphibians | Eleutherodactylus_corona | Eleutherodactylus_corona | 1 | 0.49 | 3 | 31.54 |
| amphibians | Eleutherodactylus_cubanus | Eleutherodactylus_cubanus | 1 | 0.18 | 1 | 667.25 |
| amphibians | Eleutherodactylus_cundalli | Eleutherodactylus_cundalli | 1 | 19.92 | 49.5 | 6292.65 |
| amphibians | Eleutherodactylus_cuneatus | Eleutherodactylus_cuneatus | 0 | 20.25 | 35 | 10968.08 |
| amphibians | Eleutherodactylus_cystignathoides | Eleutherodactylus_cystignathoides | 0 | 1.29 | 9 | 107464.67 |
| amphibians | Eleutherodactylus_dilatus | Eleutherodactylus_dilatus | 0 | 2.11 | 14 | 511.23 |
| amphibians | Eleutherodactylus_dimidiatus | Eleutherodactylus_dimidiatus | 0 | 17.85 | 42 | 18105.17 |
| amphibians | Eleutherodactylus_eneidae | Eleutherodactylus_eneidae | 1 | 2.11 | 11.5 | 1921.11 |
| amphibians | Eleutherodactylus_fowleri | Eleutherodactylus_fowleri | 1 | 9.67 | 12.5 | 236.06 |
| amphibians | Eleutherodactylus_glamyrus | Eleutherodactylus_glamyrus | 1 | 1.54 | 8.5 | 860.28 |
| amphibians | Eleutherodactylus_glaucoreius | Eleutherodactylus_glaucoreius | 1 | 4.54 | 28.5 | 1339.15 |
| amphibians | Eleutherodactylus_gossei | Eleutherodactylus_gossei | 1 | 3.16 | 56 | 8661.19 |
| amphibians | Eleutherodactylus_gryllus | Eleutherodactylus_gryllus | 1 | 0.4 | 5 | 1092.05 |
| amphibians | Eleutherodactylus_guanahacabibes | Eleutherodactylus_guanahacabibes | 0 | 0.89 | 8 | 951.79 |
| amphibians | Eleutherodactylus_guantanamera | Eleutherodactylus_guantanamera | 1 | 4.27 | 9 | 2108.34 |
| amphibians | Eleutherodactylus_guttilatus | Eleutherodactylus_guttilatus | 0 | 2.6 | 12.5 | 72979.53 |
| amphibians | Eleutherodactylus_hedricki | Eleutherodactylus_hedricki | 1 | 3.47 | 23 | 137.72 |
| amphibians | Eleutherodactylus_iberia | Eleutherodactylus_iberia | 1 | 0.08 | 1 | 1867.96 |
| amphibians | Eleutherodactylus_inoptatus | Eleutherodactylus_inoptatus | 0 | 68.91 | 27.5 | 50054.38 |

| | | | | | | |
|---|---|---|---|---|---|---|
| amphibians | Eleutherodactylus_jamaicensis | Eleutherodactylus_jamaicensis | 1 | 2.11 | 20 | 3409.6 |
| amphibians | Eleutherodactylus_jasperi | Eleutherodactylus_jasperi | 1 | 1.17 | 3.5 | 12.64 |
| amphibians | Eleutherodactylus_jaumei | Eleutherodactylus_jaumei | 1 | 0.13 | 2 | 26.88 |
| amphibians | Eleutherodactylus_johnstonei | Eleutherodactylus_johnstonei | 0 | 3.47 | 20 | 89710.27 |
| amphibians | Eleutherodactylus_juanariveroi | Eleutherodactylus_juanariveroi | 1 | 0.35 | 3 | 2.47 |
| amphibians | Eleutherodactylus_leberi | Eleutherodactylus_leberi | 1 | 2.87 | 27 | 85.19 |
| amphibians | Eleutherodactylus_limbatus | Eleutherodactylus_limbatus | 0 | 0.33 | 1 | 8614.96 |
| amphibians | Eleutherodactylus_locustus | Eleutherodactylus_locustus | 1 | 1.02 | 2.5 | 115.28 |
| amphibians | Eleutherodactylus_marnockii | Eleutherodactylus_marnockii | 0 | 5.14 | 13 | 178842.93 |
| amphibians | Eleutherodactylus_martinicensis | Eleutherodactylus_martinicensis | 0 | 9.03 | 34.5 | 3655.2 |
| amphibians | Eleutherodactylus_melacara | Eleutherodactylus_melacara | 1 | 3.81 | 5 | 496.71 |
| amphibians | Eleutherodactylus_monensis | Eleutherodactylus_monensis | 1 | 10.61 | 15 | 50.57 |
| amphibians | Eleutherodactylus_nubicola | Eleutherodactylus_nubicola | 1 | 10.34 | 57 | 195.16 |
| amphibians | Eleutherodactylus_orientalis | Eleutherodactylus_orientalis | 1 | 0.15 | 1 | 3.93 |
| amphibians | Eleutherodactylus_pantoni | Eleutherodactylus_pantoni | 1 | 7.29 | 76.5 | 4011.53 |
| amphibians | Eleutherodactylus_patriciae | Eleutherodactylus_patriciae | 1 | 3.47 | 22 | 714.13 |
| amphibians | Eleutherodactylus_pituinus | Eleutherodactylus_pituinus | 1 | 1.89 | 19.5 | 98.11 |
| amphibians | Eleutherodactylus_planirostris | Eleutherodactylus_planirostris | 0 | 11.04 | 14 | 330361.6 |
| amphibians | Eleutherodactylus_portoricensis | Eleutherodactylus_portoricensis | 1 | 6.27 | 15.5 | 338.55 |
| amphibians | Eleutherodactylus_richmondi | Eleutherodactylus_richmondi | 1 | 11.04 | 24.5 | 218.4 |
| amphibians | Eleutherodactylus_ricordii | Eleutherodactylus_ricordii | 1 | 5.36 | 22 | 4518.7 |
| amphibians | Eleutherodactylus_rivularis | Eleutherodactylus_rivularis | 1 | 2.35 | 42 | 56.93 |
| amphibians | Eleutherodactylus_rufescens | Eleutherodactylus_nivicolimae | 1 | 0.96 | 12 | 5468.11 |
| amphibians | Eleutherodactylus_sisyphodemus | Eleutherodactylus_sisyphodemus | 1 | 0.4 | 8 | 55.18 |
| amphibians | Eleutherodactylus_tetajulia | Eleutherodactylus_tetajulia | 1 | 0.18 | 6 | 444.17 |
| amphibians | Eleutherodactylus_thorectes | Eleutherodactylus_thorectes | 1 | 0.22 | 3 | 42.19 |
| amphibians | Eleutherodactylus_turquinensis | Eleutherodactylus_turquinensis | 1 | 14.16 | 24 | 269.13 |
| amphibians | Eleutherodactylus_varians | Eleutherodactylus_varians | 0 | 5.36 | 21 | 4268.58 |
| amphibians | Eleutherodactylus_varleyi | Eleutherodactylus_varleyi | 0 | 0.44 | 4 | 66329.94 |
| amphibians | Eleutherodactylus_wetmorei | Eleutherodactylus_wetmorei | 1 | 5.8 | 10 | 7458.75 |
| amphibians | Eleutherodactylus_wightmanae | Eleutherodactylus_wightmanae | 1 | 0.89 | 17 | 1994.82 |
| amphibians | Engystomops_coloradorum | Engystomops_coloradorum | 1 | 1.28 | 130 | 2919.12 |
| amphibians | Engystomops_petersi | Engystomops_petersi | 0 | 4.98 | 431.5 | 2015286.4 |
| amphibians | Engystomops_pustulatus | Engystomops_pustulatus | 0 | 3.98 | 320 | 23877.51 |

| | | | | | | |
|---|---|---|---|---|---|---|
| amphibians | Engystomops_pustulosus | Engystomops_pustulosus | 0 | 9.8 | 265 | 894865.28 |
| amphibians | Engystomops_randi | Engystomops_randi | 0 | 1.02 | 110 | 8331.1 |
| amphibians | Ensatina_eschscholtzii | Ensatina_eschscholtzii | 0 | 8.44 | 14 | 325428.35 |
| amphibians | Epidalea_calamita | Epidalea_calamita | 0 | 104.27 | 6250 | 2234800.72 |
| amphibians | Epipedobates_anthonyi | Epipedobates_anthonyi | 0 | 1.5 | 24 | 20770.84 |
| amphibians | Epipedobates_boulengeri | Epipedobates_boulengeri | 0 | 0.75 | 20 | 55338.23 |
| amphibians | Epipedobates_machalilla | Epipedobates_machalilla | 0 | 0.37 | 15 | 37054.53 |
| amphibians | Epipedobates_tricolor | Epipedobates_tricolor | 1 | 1.41 | 16 | 4671.22 |
| amphibians | Espadarana_andina | Espadarana_andina | 0 | 1.61 | 55 | 34642.72 |
| amphibians | Espadarana_callistomma | Espadarana_callistomma | 0 | 2.55 | 35 | 30974.08 |
| amphibians | Espadarana_prosoblepon | Espadarana_prosoblepon | 0 | 2.35 | 66 | 346069.79 |
| amphibians | Euphlyctis_cyanophlyctis | Euphlyctis_cyanophlyctis | 0 | 79.58 | 844 | 4408170.13 |
| amphibians | Euphlyctis_hexadactyla | Euphlyctis_hexadactylus | 0 | 347.52 | 2000 | 330377.45 |
| amphibians | Euproctus_montanus | Euproctus_montanus | 0 | 3.16 | 40 | 7955.86 |
| amphibians | Euproctus_platycephalus | Euproctus_platycephalus | 1 | 5.54 | 220 | 5723.5 |
| amphibians | Eupsophus_altor | Eupsophus_altor | 1 | 4.02 | 23.5 | 312.67 |
| amphibians | Eupsophus_emiliopugini | Eupsophus_emiliopugini | 0 | 11.96 | 88 | 64006.96 |
| amphibians | Eurycea_bislineata | Eurycea_bislineata | 0 | 1.09 | 63.5 | 1198568.49 |
| amphibians | Eurycea_chamberlaini | Eurycea_chamberlaini | 0 | 0.57 | 49.5 | 79016.26 |
| amphibians | Eurycea_cirrigera | Eurycea_cirrigera | 0 | 1.7 | 61 | 1080486.82 |
| amphibians | Eurycea_guttolineata | Eurycea_guttolineata | 0 | 3.67 | 11 | 738309.3 |
| amphibians | Eurycea_junaluska | Eurycea_junaluska | 1 | 1.36 | 49 | 4834.61 |
| amphibians | Eurycea_latitans | Eurycea_tridentifera | 1 | 1.6 | 12.5 | 6213.75 |
| amphibians | Eurycea_longicauda | Eurycea_longicauda | 0 | 3.34 | 83 | 848916.9 |
| amphibians | Eurycea_lucifuga | Eurycea_lucifuga | 0 | 3.51 | 84.5 | 449641.61 |
| amphibians | Eurycea_multiplicata | Eurycea_multiplicata | 0 | 1.55 | 11.5 | 136465.11 |
| amphibians | Eurycea_nana | Eurycea_nana | 1 | 0.35 | 37.5 | 0.09 |
| amphibians | Eurycea_neotenes | Eurycea_neotenes | 1 | 2.16 | 28 | 774.62 |
| amphibians | Eurycea_pterophila | Eurycea_pterophila | 1 | 0.63 | 12.5 | 3412.38 |
| amphibians | Eurycea_quadridigitata | Eurycea_quadridigitata | 0 | 0.65 | 34.5 | 251090.11 |
| amphibians | Eurycea_rathbuni | Eurycea_rathbuni | 1 | 7.55 | 39 | 56.69 |
| amphibians | Eurycea_sosorum | Eurycea_sosorum | 1 | 0.42 | 23.5 | 1011.62 |
| amphibians | Eurycea_spelaea | Eurycea_spelaea | 0 | 5.49 | 16 | 32788.25 |
| amphibians | Eurycea_tynerensis | Eurycea_tynerensis | 0 | 1.75 | 6.5 | 22705.87 |

| | | | | | | |
|---|---|---|---|---|---|---|
| amphibians | Eurycea_wilderae | Eurycea_wilderae | 0 | 1.93 | 54 | 76390.17 |
| amphibians | Excidobates_mysteriosus | Excidobates_mysteriosus | 1 | 2.11 | 10.5 | 1306.16 |
| amphibians | Exerodonta_melanomma | Exerodonta_melanomma | 1 | 2.47 | 265 | 1328.47 |
| amphibians | Exerodonta_sumichrasti | Exerodonta_sumichrasti | 0 | 2.9 | 157.5 | 40451.19 |
| amphibians | Feihyla_hansenae | Chiromantis_hansenae | 0 | 1.17 | 235 | 458688.58 |
| amphibians | Feihyla_vittata | Chiromantis_vittatus | 0 | 2.6 | 217 | 1369629.74 |
| amphibians | Feihyla_vittiger | Philautus_vittiger | 0 | 2.11 | 51.5 | 2907.52 |
| amphibians | Fejervarya_cancrivora | Fejervarya_cancrivora | 0 | 188.23 | 1045 | 817338.52 |
| amphibians | Fejervarya_limnocharis | Fejervarya_limnocharis | 0 | 42.85 | 855.5 | 6160012.44 |
| amphibians | Fejervarya_sakishimensis | Fejervarya_sakishimensis | 0 | 31.63 | 3550 | 790.04 |
| amphibians | Fejervarya_vittigera | Fejervarya_vittigera | 0 | 51.84 | 1500 | 288730.35 |
| amphibians | Firouzophrynus_stomaticus | Duttaphrynus_stomaticus | 0 | 74.11 | 10000 | 4282362.48 |
| amphibians | Flectonotus_fitzgeraldi | Flectonotus_fitzgeraldi | 0 | 1.02 | 4.5 | 4192.85 |
| amphibians | Flectonotus_pygmaeus | Flectonotus_pygmaeus | 0 | 2.71 | 9 | 41606.16 |
| amphibians | Fritziana_fissilis | Fritziana_fissilis | 0 | 4.81 | 11 | 1963.94 |
| amphibians | Fritziana_goeldii | Fritziana_goeldii | 0 | 4.16 | 15.5 | 31471.82 |
| amphibians | Fritziana_ohausi | Fritziana_ohausi | 0 | 3.38 | 15 | 38985.79 |
| amphibians | Gastrophryne_carolinensis | Gastrophryne_carolinensis | 0 | 4.93 | 911 | 1576052.91 |
| amphibians | Gastrophryne_olivacea | Gastrophryne_olivacea | 0 | 6.27 | 1300 | 1462157.64 |
| amphibians | Gastrotheca_albolineata | Gastrotheca_albolineata | 0 | 19.92 | 19 | 18226.12 |
| amphibians | Gastrotheca_andaquiensis | Gastrotheca_andaquiensis | 0 | 44.71 | 12 | 9079.87 |
| amphibians | Gastrotheca_argenteovirens | Gastrotheca_argenteovirens | 0 | 25.82 | 37.5 | 4237.16 |
| amphibians | Gastrotheca_aureomaculata | Gastrotheca_aureomaculata | 1 | 42.85 | 93 | 1186.19 |
| amphibians | Gastrotheca_christiani | Gastrotheca_christiani | 1 | 5.36 | 9 | 56.55 |
| amphibians | Gastrotheca_chrysosticta | Gastrotheca_chrysosticta | 1 | 3.88 | 33 | 210.6 |
| amphibians | Gastrotheca_cornuta | Gastrotheca_cornuta | 1 | 52.89 | 10 | 45654.31 |
| amphibians | Gastrotheca_dendronastes | Gastrotheca_dendronastes | 1 | 49.58 | 11 | 3405.58 |
| amphibians | Gastrotheca_dunni | Gastrotheca_dunni | 0 | 15.93 | 46 | 2109.02 |
| amphibians | Gastrotheca_ernestoi | Gastrotheca_ernestoi | 0 | 41.05 | 22 | 55978.95 |
| amphibians | Gastrotheca_excubitor | Gastrotheca_excubitor | 1 | 9.22 | 21 | 5227.98 |
| amphibians | Gastrotheca_fissipes | Gastrotheca_fissipes | 0 | 76 | 20 | 45618.32 |
| amphibians | Gastrotheca_gracilis | Gastrotheca_gracilis | 1 | 9.67 | 66 | 1279.31 |
| amphibians | Gastrotheca_griswoldi | Gastrotheca_griswoldi | 0 | 7.29 | 21 | 6390.73 |
| amphibians | Gastrotheca_helenae | Gastrotheca_helenae | 1 | 27.13 | 12 | 446.87 |

| | | | | | | |
|---|---|---|---|---|---|---|
| amphibians | Gastrotheca_lateonota | Gastrotheca_lateonota | 1 | 24.19 | 108 | 2352.04 |
| amphibians | Gastrotheca_litonedis | Gastrotheca_litonedis | 1 | 22.62 | 69 | 81 |
| amphibians | Gastrotheca_longipes | Gastrotheca_longipes | 0 | 69.67 | 16 | 54620.77 |
| amphibians | Gastrotheca_marsupiata | Gastrotheca_marsupiata | 0 | 10.75 | 115 | 245721.67 |
| amphibians | Gastrotheca_megacephala | Gastrotheca_megacephala | 0 | 141.99 | 17 | 72659.37 |
| amphibians | Gastrotheca_microdiscus | Gastrotheca_microdiscus | 0 | 28.35 | 4.5 | 69344.74 |
| amphibians | Gastrotheca_monticola | Gastrotheca_monticola | 0 | 44.71 | 117.5 | 30970.55 |
| amphibians | Gastrotheca_nebulanastes | Gastrotheca_nebulanastes | 1 | 5.67 | 12.5 | 120.48 |
| amphibians | Gastrotheca_nicefori | Gastrotheca_nicefori | 0 | 54.82 | 38 | 113015.13 |
| amphibians | Gastrotheca_ochoai | Gastrotheca_ochoai | 1 | 4.5 | 16 | 454.64 |
| amphibians | Gastrotheca_orophylax | Gastrotheca_orophylax | 1 | 39.31 | 24.5 | 3165.92 |
| amphibians | Gastrotheca_ovifera | Gastrotheca_ovifera | 1 | 57.01 | 27 | 1413.66 |
| amphibians | Gastrotheca_pacchamama | Gastrotheca_pacchamama | 1 | 4.54 | 6.5 | 224.06 |
| amphibians | Gastrotheca_phelloderma | Gastrotheca_phelloderma | 1 | 17.55 | 45 | 161.87 |
| amphibians | Gastrotheca_plumbea | Gastrotheca_plumbea | 1 | 37.61 | 24 | 7778.5 |
| amphibians | Gastrotheca_pseustes | Gastrotheca_pseustes | 0 | 24.19 | 123 | 34203.84 |
| amphibians | Gastrotheca_pulchra | Gastrotheca_pulchra | 0 | 3.22 | 10 | 76462.29 |
| amphibians | Gastrotheca_rebeccae | Gastrotheca_rebeccae | 1 | 8.25 | 22 | 310 |
| amphibians | Gastrotheca_recava | Gastrotheca_recava | 0 | 61.35 | 20 | 34642.8 |
| amphibians | Gastrotheca_riobambae | Gastrotheca_riobambae | 1 | 27.67 | 115 | 9041.94 |
| amphibians | Gastrotheca_ruizi | Gastrotheca_ruizi | 0 | 46.62 | 126.5 | 145.46 |
| amphibians | Gastrotheca_stictopleura | Gastrotheca_stictopleura | 1 | 30.32 | 82 | 167.69 |
| amphibians | Gastrotheca_testudinea | Gastrotheca_testudinea | 0 | 56.13 | 49.5 | 204436.36 |
| amphibians | Gastrotheca_trachyceps | Gastrotheca_trachyceps | 1 | 29.89 | 30 | 99.92 |
| amphibians | Gastrotheca_walkeri | Gastrotheca_walkeri | 1 | 50.6 | 17 | 1181.87 |
| amphibians | Gastrotheca_weinlandii | Gastrotheca_weinlandii | 0 | 63.96 | 14 | 25298.42 |
| amphibians | Gegeneophis_ramaswamii | Gegeneophis_ramaswamii | 0 | 14.76 | 11.5 | 4867.72 |
| amphibians | Geocrinia_laevis | Geocrinia_laevis | 0 | 3.47 | 210 | 76935.88 |
| amphibians | Geocrinia_leai | Geocrinia_leai | 0 | 1.33 | 74 | 36314.16 |
| amphibians | Geocrinia_victoriana | Geocrinia_victoriana | 0 | 3.47 | 132.5 | 104489.33 |
| amphibians | Geotrypetes_seraphini | Geotrypetes_seraphini | 0 | 24.12 | 2.5 | 1139615.21 |
| amphibians | Gephyromantis_eiselti | Gephyromantis_eiselti | 1 | 0.89 | 4 | 585.74 |
| amphibians | Gephyromantis_horridus | Gephyromantis_horridus | 1 | 3.47 | 80 | 1003.38 |
| amphibians | Gephyromantis_malagasius | Gephyromantis_malagasius | 0 | 1.33 | 232 | 83503.16 |

| | | | | | | |
|---|---|---|---|---|---|---|
| amphibians | Gephyromantis_pseudoasper | Gephyromantis_pseudoasper | 0 | 3.16 | 12 | 20436.01 |
| amphibians | Gephyromantis_webbi | Gephyromantis_webbi | 1 | 2.87 | 7 | 2489.09 |
| amphibians | Glandirana_emeljanovi | Rugosa_emelianjovi | 0 | 15.39 | 783 | 439660.28 |
| amphibians | Glandirana_minima | Glandirana_minima | 1 | 1.85 | 269.5 | 3406.45 |
| amphibians | Glandirana_rugosa | Rugosa_rugosa | 0 | 19.92 | 1305 | 310817.99 |
| amphibians | Glandirana_tientaiensis | Rugosa_tiantaiensis | 0 | 2.41 | 975 | 59479.47 |
| amphibians | Glyphoglossus_molossus | Glyphoglossus_molossus | 0 | 63.96 | 250 | 871247.26 |
| amphibians | Gracixalus_gracilipes | Gracixalus_gracilipes | 0 | 2 | 5 | 192172.5 |
| amphibians | Gracixalus_quangi | Gracixalus_quangi | 0 | 1.55 | 12.5 | 134844.51 |
| amphibians | Gracixalus_supercornutus | Gracixalus_supercornutus | 0 | 2.63 | 10.5 | 2976.64 |
| amphibians | Grandisonia_sechellensis | Grandisonia_sechellensis | 0 | 2.55 | 12 | 217.11 |
| amphibians | Guibemantis_bicalcaratus | Guibemantis_bicalcaratus | 0 | 1.89 | 25 | 112874.11 |
| amphibians | Guibemantis_depressiceps | Guibemantis_depressiceps | 0 | 7.84 | 66 | 134877.68 |
| amphibians | Guibemantis_liber | Guibemantis_liber | 0 | 1.89 | 60 | 143701.42 |
| amphibians | Guibemantis_tornieri | Guibemantis_tornieri | 0 | 18.25 | 70 | 61565.49 |
| amphibians | Gymnopis_multiplicata | Gymnopis_multiplicata | 0 | 41.82 | 16 | 182860.26 |
| amphibians | Gyrinophilus_porphyriticus | Gyrinophilus_porphyriticus | 0 | 26.8 | 73.5 | 764568.27 |
| amphibians | Haddadus_binotatus | Haddadus_binotatus | 0 | 24.56 | 31.5 | 496585.75 |
| amphibians | Hadromophryne_natalensis | Hadromophryne_natalensis | 0 | 25.82 | 125 | 121695.1 |
| amphibians | Hamptophryne_boliviana | Hamptophryne_boliviana | 0 | 7.84 | 220 | 4439329.92 |
| amphibians | Heleioporus_albopunctatus | Heleioporus_albopunctatus | 0 | 104.27 | 459 | 203573.95 |
| amphibians | Heleioporus_australiacus | Heleioporus_australiacus | 1 | 94.47 | 820 | 42343.84 |
| amphibians | Heleioporus_barycragus | Heleioporus_barycragus | 0 | 63.96 | 384 | 15500.39 |
| amphibians | Heleioporus_eyrei | Heleioporus_eyrei | 0 | 27.13 | 285 | 115826.89 |
| amphibians | Heleioporus_inornatus | Heleioporus_inornatus | 0 | 37.61 | 170 | 34319.42 |
| amphibians | Heleioporus_psammophilus | Heleioporus_psammophilus | 0 | 22.16 | 185 | 154841.83 |
| amphibians | Heleophryne_hewitti | Heleophryne_hewitti | 1 | 11.04 | 146.5 | 140.52 |
| amphibians | Heleophryne_orientalis | Heleophryne_orientalis | 0 | 8.42 | 150.5 | 1191.27 |
| amphibians | Heleophryne_purcelli | Heleophryne_purcelli | 0 | 15.93 | 129 | 14223.9 |
| amphibians | Heleophryne_regis | Heleophryne_regis | 0 | 10.34 | 157 | 6393.13 |
| amphibians | Hemidactylium_scutatum | Hemidactylium_scutatum | 0 | 2.29 | 74.5 | 1511888.54 |
| amphibians | Hemiphractus_fasciatus | Hemiphractus_fasciatus | 1 | 30.9 | 12.5 | 57066.6 |
| amphibians | Hemiphractus_helioi | Hemiphractus_helioi | 0 | 41.05 | 17.5 | 400478.01 |
| amphibians | Hemiphractus_johnsoni | Hemiphractus_johnsoni | 1 | 45.08 | 18 | 267.74 |

| | | | | | | |
|---|---|---|---|---|---|---|
| amphibians | Hemiphractus_proboscideus | Hemiphractus_proboscideus | 0 | 28.49 | 23 | 338926.37 |
| amphibians | Hemiphractus_scutatus | Hemiphractus_scutatus | 0 | 52.68 | 11 | 2046879.87 |
| amphibians | Hemisus_guttatus | Hemisus_guttatus | 0 | 50.6 | 1098.5 | 8652.86 |
| amphibians | Hemisus_marmoratus | Hemisus_marmoratus | 0 | 19.92 | 165 | 9852655.86 |
| amphibians | Herpele_squalostoma | Herpele_squalostoma | 0 | 68.43 | 9 | 914810.4 |
| amphibians | Heterixalus_alboguttatus | Heterixalus_alboguttatus | 0 | 2.87 | 500 | 30825.68 |
| amphibians | Heterixalus_betsileo | Heterixalus_betsileo | 0 | 1.89 | 71 | 106400.59 |
| amphibians | Hildebrandtia_ornata | Hildebrandtia_ornata | 0 | 32.83 | 1336 | 6748590.7 |
| amphibians | Holoaden_bradei | Holoaden_bradei | 1 | 9.67 | 8 | 3.26 |
| amphibians | Holoaden_luederwaldti | Holoaden_luederwaldti | 0 | 9.67 | 38.5 | 2364.03 |
| amphibians | Hoplobatrachus_chinensis | Hoplobatrachus_rugulosus | 0 | 262.68 | 2300 | 3004293.14 |
| amphibians | Hoplobatrachus_occipitalis | Hoplobatrachus_occipitalis | 0 | 478.08 | 2042.5 | 8574915.61 |
| amphibians | Hoplobatrachus_tigerinus | Hoplobatrachus_tigerinus | 0 | 675.32 | 5750 | 3535067.89 |
| amphibians | Hoplophryne_rogersi | Hoplophryne_rogersi | 1 | 2.6 | 6 | 1002.54 |
| amphibians | Hoplophryne_uluguruensis | Hoplophryne_uluguruensis | 1 | 1.17 | 28 | 902.69 |
| amphibians | Hyalinobatrachium_aureoguttatum | Hyalinobatrachium_aureoguttatum | 0 | 1.02 | 33.5 | 25282.31 |
| amphibians | Hyalinobatrachium_cappellei | Hyalinobatrachium_cappellei | 0 | 1.15 | 18.5 | 2321705.75 |
| amphibians | Hyalinobatrachium_carlesvilai | Hyalinobatrachium_carlesvilai | 0 | 1.01 | 32 | 204335.6 |
| amphibians | Hyalinobatrachium_chirripoi | Hyalinobatrachium_chirripoi | 0 | 2.11 | 72.5 | 57455.95 |
| amphibians | Hyalinobatrachium_colymbiphyllum | Hyalinobatrachium_colymbiphyllum | 0 | 1.89 | 102.5 | 113097.16 |
| amphibians | Hyalinobatrachium_duranti | Hyalinobatrachium_duranti | 1 | 1.11 | 24.5 | 689.4 |
| amphibians | Hyalinobatrachium_fleischmanni | Hyalinobatrachium_fleischmanni | 0 | 2.6 | 30 | 37182.31 |
| amphibians | Hyalinobatrachium_fragile | Hyalinobatrachium_fragile | 0 | 1.09 | 22 | 7370.83 |
| amphibians | Hyalinobatrachium_iaspidiense | Hyalinobatrachium_iaspidiense | 0 | 0.85 | 22 | 2097241.12 |
| amphibians | Hyalinobatrachium_kawense | Hyalinobatrachium_kawense | 0 | 0.57 | 33 | 57741.24 |
| amphibians | Hyalinobatrachium_mondolfii | Hyalinobatrachium_mondolfii | 0 | 0.83 | 24.5 | 1129956.56 |
| amphibians | Hyalinobatrachium_munozorum | Hyalinobatrachium_munozorum | 0 | 0.97 | 18 | 39779.41 |
| amphibians | Hyalinobatrachium_orientale | Hyalinobatrachium_orientale | 1 | 0.97 | 26 | 2704.26 |
| amphibians | Hyalinobatrachium_orocostale | Hyalinobatrachium_orocostale | 1 | 0.61 | 45 | 460.01 |
| amphibians | Hyalinobatrachium_talamancae | Hyalinobatrachium_talamancae | 0 | 1.5 | 28.5 | 519.06 |
| amphibians | Hyalinobatrachium_tatayoi | Hyalinobatrachium_tatayoi | 0 | 0.82 | 40 | 232033.58 |
| amphibians | Hyalinobatrachium_taylori | Hyalinobatrachium_taylori | 0 | 0.77 | 32 | 463793.3 |
| amphibians | Hyalinobatrachium_tricolor | Hyalinobatrachium_tricolor | 0 | 0.66 | 22 | 37061.53 |
| amphibians | Hyalinobatrachium_valerioi | Hyalinobatrachium_valerioi | 0 | 1.33 | 38.5 | 69861.61 |

| | | | | | | |
|---|---|---|---|---|---|---|
| amphibians | Hydromantes_platycephalus | Hydromantes_platycephalus | 0 | 4.83 | 10 | 48834.54 |
| amphibians | Hydromantes_shastae | Hydromantes_shastae | 0 | 4.14 | 10.5 | 747.51 |
| amphibians | Hyla_annectans | Hyla_annectans | 0 | 7.03 | 946 | 1002137.49 |
| amphibians | Hyla_arborea | Hyla_arborea | 0 | 12.53 | 1100 | 1771404.5 |
| amphibians | Hyla_chinensis | Hyla_chinensis | 0 | 4.65 | 459 | 640031.7 |
| amphibians | Hyla_intermedia | Hyla_intermedia | 0 | 19.92 | 1200 | 230703.13 |
| amphibians | Hyla_meridionalis | Hyla_meridionalis | 0 | 25.82 | 2000 | 643813.43 |
| amphibians | Hyla_orientalis | Hyla_orientalis | 0 | 13.33 | 120 | 1852620.44 |
| amphibians | Hyla_sanchiangensis | Hyla_sanchiangensis | 0 | 4.35 | 1075 | 442581.78 |
| amphibians | Hyla_sarda | Hyla_sarda | 0 | 11.04 | 625 | 31503.52 |
| amphibians | Hyla_savignyi | Hyla_savignyi | 0 | 10.34 | 600 | 694154.18 |
| amphibians | Hyla_tsinlingensis | Hyla_tsinlingensis | 0 | 9.67 | 1185 | 89298.64 |
| amphibians | Hylambates_boulengeri | Phlyctimantis_boulengeri | 0 | 18.87 | 225 | 384199.39 |
| amphibians | Hylambates_maculatus | Kassina_maculata | 0 | 35.97 | 172.5 | 915307.44 |
| amphibians | Hylarana_chalconota | Hylarana_chalconota | 0 | 48.58 | 2100 | 318282.29 |
| amphibians | Hylarana_erythraea | Hylarana_erythraea | 0 | 46.62 | 1550 | 2299374.73 |
| amphibians | Hylarana_galamensis | Hylarana_galamensis | 0 | 63.96 | 3067.5 | 5152211.2 |
| amphibians | Hylarana_guentheri | Hylarana_guentheri | 0 | 50.6 | 1995 | 1858234.22 |
| amphibians | Hylarana_labialis | Hylarana_labialis | 0 | 18.87 | 1443.5 | 131956.59 |
| amphibians | Hylarana_latouchii | Hylarana_latouchii | 0 | 10.65 | 655 | 875372.51 |
| amphibians | Hylarana_macrodactyla | Hylarana_macrodactyla | 0 | 7.29 | 300 | 1762927.83 |
| amphibians | Hylarana_megalonesa | Hylarana_megalonesa | 0 | 26.6 | 1048.5 | 22774.67 |
| amphibians | Hylarana_nigrovittata | Hylarana_nigrovittata | 0 | 32.83 | 281 | 783557.81 |
| amphibians | Hylarana_raniceps | Hylarana_raniceps | 0 | 34.06 | 2158 | 115107.9 |
| amphibians | Hylarana_taipehensis | Hylarana_taipehensis | 0 | 7.84 | 194.5 | 1177402.89 |
| amphibians | Hylarana_temporalis | Hylarana_temporalis | 1 | 54.82 | 1548 | 17658.53 |
| amphibians | Hylodes_asper | Hylodes_asper | 0 | 11.04 | 85.5 | 23535.62 |
| amphibians | Hylodes_dactylocinus | Hylodes_dactylocinus | 1 | 2.35 | 50 | 7122.31 |
| amphibians | Hylodes_heyeri | Hylodes_heyeri | 0 | 8.25 | 132 | 45673.36 |
| amphibians | Hylodes_phyllodes | Hylodes_phyllodes | 0 | 2.65 | 40.5 | 22660.21 |
| amphibians | Hylodes_sazimai | Hylodes_sazimai | 1 | 2.18 | 109 | 5953.21 |
| amphibians | Hylomantis_aspera | Hylomantis_aspera | 0 | 9.67 | 60 | 37306.79 |
| amphibians | Hylomantis_granulosa | Hylomantis_granulosa | 0 | 9.67 | 43 | 110840.52 |
| amphibians | Hylorina_sylvatica | Hylorina_sylvatica | 0 | 41.05 | 475 | 91167.03 |

| | | | | | | |
|---|---|---|---|---|---|---|
| amphibians | Hyloscirtus_armatus | Hyloscirtus_armatus | 0 | 41.05 | 300 | 57355.31 |
| amphibians | Hyloscirtus_caucanus | Hyloscirtus_caucanus | 1 | 25.06 | 120 | 749.03 |
| amphibians | Hyloscirtus_platydactylus | Hyloscirtus_platydactylus | 1 | 6.42 | 30 | 5255.64 |
| amphibians | Hyloscirtus_tapichalaca | Hyloscirtus_tapichalaca | 1 | 46.81 | 428 | 84.1 |
| amphibians | Hyloxalus_awa | Hyloxalus_awa | 0 | 1.31 | 12.5 | 31202.07 |
| amphibians | Hyloxalus_azureiventris | Hyloxalus_azureiventris | 1 | 1.59 | 14 | 301.04 |
| amphibians | Hyloxalus_bocagei | Hyloxalus_bocagei | 1 | 2 | 39 | 3003.42 |
| amphibians | Hyloxalus_elachyhistus | Hyloxalus_elachyhistus | 0 | 1.28 | 13.5 | 24992.2 |
| amphibians | Hyloxalus_insulatus | Hyloxalus_insulatus | 1 | 1.33 | 217 | 15135.12 |
| amphibians | Hyloxalus_italoi | Hyloxalus_italoi | 0 | 2.13 | 22.5 | 36406.04 |
| amphibians | Hyloxalus_maculosus | Hyloxalus_maculosus | 1 | 1.89 | 34 | 2130.19 |
| amphibians | Hyloxalus_nexipus | Hyloxalus_nexipus | 0 | 3.47 | 79 | 31137.26 |
| amphibians | Hyloxalus_sauli | Hyloxalus_sauli | 0 | 1.83 | 10 | 56197.95 |
| amphibians | Hyloxalus_subpunctatus | Hyloxalus_subpunctatus | 0 | 1.07 | 36.5 | 11032.55 |
| amphibians | Hyloxalus_sylvaticus | Hyloxalus_sylvaticus | 1 | 2.11 | 265 | 784.38 |
| amphibians | Hyloxalus_toachi | Hyloxalus_toachi | 1 | 1.73 | 10 | 8072.7 |
| amphibians | Hyloxalus_vertebralis | Hyloxalus_vertebralis | 1 | 0.59 | 9 | 13409.5 |
| amphibians | Hymenochirus_boettgeri | Hymenochirus_boettgeri | 0 | 5.8 | 563.5 | 2034734.85 |
| amphibians | Hymenochirus_curtipes | Hymenochirus_curtipes | 0 | 5.36 | 1500 | 663528.66 |
| amphibians | Hynobius_abei | Hynobius_abei | 1 | 5.73 | 67.5 | 2256.78 |
| amphibians | Hynobius_amjiensis | Hynobius_amjiensis | 1 | 10.24 | 123.5 | 102.79 |
| amphibians | Hynobius_arisanensis | Hynobius_arisanensis | 1 | 5.27 | 12 | 2348.37 |
| amphibians | Hynobius_boulengeri | Hynobius_boulengeri | 1 | 19.13 | 36 | 2837.24 |
| amphibians | Hynobius_dunni | Hynobius_dunni | 1 | 8.13 | 119 | 4935.01 |
| amphibians | Hynobius_formosanus | Hynobius_formosanus | 1 | 3.79 | 28 | 1985.85 |
| amphibians | Hynobius_fucus | Hynobius_fuca | 0 | 2.67 | 19 | 1562.91 |
| amphibians | Hynobius_glacialis | Hynobius_glacialis | 1 | 4.92 | 19 | 99.63 |
| amphibians | Hynobius_guabangshanensis | Hynobius_guabangshanensis | 1 | 10.59 | 155 | 13.34 |
| amphibians | Hynobius_hidamontanus | Hynobius_hidamontanus | 1 | 3.16 | 91 | 2501.29 |
| amphibians | Hynobius_katoi | Hynobius_katoi | 1 | 4.64 | 11 | 892.69 |
| amphibians | Hynobius_kimurae | Hynobius_kimurae | 0 | 14.33 | 32 | 33130.53 |
| amphibians | Hynobius_leechii | Hynobius_leechii | 0 | 4.57 | 155 | 542724.12 |
| amphibians | Hynobius_lichenatus | Hynobius_lichenatus | 0 | 5.49 | 78 | 68172.07 |
| amphibians | Hynobius_naevius | Hynobius_naevius | 1 | 13.36 | 26.5 | 2850.59 |

| | | | | | | |
|---|---|---|---|---|---|---|
| amphibians | Hynobius_nebulosus | Hynobius_nebulosus | 0 | 5.49 | 119 | 10929.53 |
| amphibians | Hynobius_nigrescens | Hynobius_nigrescens | 0 | 9.39 | 90 | 82972.24 |
| amphibians | Hynobius_okiensis | Hynobius_okiensis | 1 | 6.22 | 40 | 249.54 |
| amphibians | Hynobius_quelpaertensis | Hynobius_quelpaertensis | 1 | 5.78 | 105 | 7743.62 |
| amphibians | Hynobius_retardatus | Hynobius_retardatus | 0 | 9.72 | 101 | 77446.26 |
| amphibians | Hynobius_sonani | Hynobius_sonani | 1 | 4.77 | 32 | 3090.83 |
| amphibians | Hynobius_stejnegeri | Hynobius_stejnegeri | 0 | 9.72 | 36.5 | 9545.7 |
| amphibians | Hynobius_takedai | Hynobius_takedai | 1 | 4.83 | 87 | 2286.87 |
| amphibians | Hynobius_tokyoensis | Hynobius_tokyoensis | 1 | 9.39 | 103.5 | 8430.91 |
| amphibians | Hynobius_tsuensis | Hynobius_tsuensis | 0 | 6.73 | 50 | 702.75 |
| amphibians | Hynobius_yangi | Hynobius_yangi | 1 | 3.73 | 76 | 341.25 |
| amphibians | Hynobius_yiwuensis | Hynobius_yiwuensis | 0 | 3.58 | 90.5 | 31199.3 |
| amphibians | Hyperolius_argus | Hyperolius_argus | 0 | 5.36 | 115 | 710482.93 |
| amphibians | Hyperolius_bopeleti | Hyperolius_bopeleti | 1 | 0.77 | 100 | 6241.71 |
| amphibians | Hyperolius_castaneus | Hyperolius_castaneus | 0 | 3.81 | 125 | 17288.83 |
| amphibians | Hyperolius_cinnamomeoventris | Hyperolius_cinnamomeoventris | 0 | 2.87 | 132 | 4028644.91 |
| amphibians | Hyperolius_concolor | Hyperolius_concolor | 0 | 6.27 | 250 | 1119901.09 |
| amphibians | Hyperolius_cystocandicans | Hyperolius_cystocandicans | 1 | 3.81 | 180 | 7790.33 |
| amphibians | Hyperolius_frontalis | Hyperolius_frontalis | 0 | 3.47 | 23.5 | 9352.36 |
| amphibians | Hyperolius_glandicolor | Hyperolius_sheldricki | 0 | 3.47 | 175 | 1059258.74 |
| amphibians | Hyperolius_guttulatus | Hyperolius_guttulatus | 0 | 4.16 | 250 | 615759.43 |
| amphibians | Hyperolius_horstockii | Hyperolius_horstockii | 0 | 6.77 | 20 | 18583.63 |
| amphibians | Hyperolius_hypsiphonus | Alexteroon_hypsiphonus | 0 | 1.69 | 70 | 528920.81 |
| amphibians | Hyperolius_igbettensis | Hyperolius_igbettensis | 0 | 1.17 | 270 | 1652727.2 |
| amphibians | Hyperolius_kivuensis | Hyperolius_kivuensis | 0 | 4.93 | 216.5 | 2983645.54 |
| amphibians | Hyperolius_koehleri | Chlorolius_koehleri | 0 | 2.11 | 60 | 203103.26 |
| amphibians | Hyperolius_langi | Hyperolius_langi | 0 | 2.35 | 56 | 90318.33 |
| amphibians | Hyperolius_lateralis | Hyperolius_lateralis | 0 | 2.11 | 136.5 | 302571.6 |
| amphibians | Hyperolius_marginatus | Hyperolius_marginatus | 0 | 3.16 | 452.5 | 1725323.76 |
| amphibians | Hyperolius_mariae | Hyperolius_mariae | 0 | 1.69 | 330 | 78115.65 |
| amphibians | Hyperolius_marmoratus | Hyperolius_marmoratus | 0 | 6.77 | 400 | 1214080.42 |
| amphibians | Hyperolius_microps | Hyperolius_acuticeps | 0 | 1.33 | 176 | 64270.6 |
| amphibians | Hyperolius_mitchelli | Hyperolius_mitchelli | 0 | 2.6 | 125 | 809668.97 |
| amphibians | Hyperolius_molleri | Hyperolius_molleri | 0 | 2.87 | 90 | 1003.48 |

| | | | | | | |
|---|---|---|---|---|---|---|
| amphibians | Hyperolius_mosaicus | Hyperolius_mosaicus | 0 | 1.5 | 20 | 145843.81 |
| amphibians | Hyperolius_nasutus | Hyperolius_nasutus | 0 | 1.33 | 176 | 1164689.56 |
| amphibians | Hyperolius_nitidulus | Hyperolius_nitidulus | 0 | 3.47 | 472 | 3147892.17 |
| amphibians | Hyperolius_obstetricans | Alexteroon_obstetricans | 0 | 2.35 | 45 | 284849.88 |
| amphibians | Hyperolius_parallelus | Hyperolius_parallelus | 0 | 3.47 | 200 | 2511290.57 |
| amphibians | Hyperolius_parkeri | Hyperolius_parkeri | 0 | 1.69 | 73 | 237865.01 |
| amphibians | Hyperolius_pickersgilli | Hyperolius_pickersgilli | 1 | 2.11 | 50.5 | 3039.01 |
| amphibians | Hyperolius_pictus | Hyperolius_pictus | 0 | 4.54 | 80 | 33516.33 |
| amphibians | Hyperolius_poweri | Hyperolius_poweri | 0 | 1.17 | 245 | 24887.74 |
| amphibians | Hyperolius_puncticulatus | Hyperolius_puncticulatus | 1 | 6.77 | 102.5 | 920.56 |
| amphibians | Hyperolius_pusillus | Hyperolius_pusillus | 0 | 1.17 | 260 | 802133.78 |
| amphibians | Hyperolius_rubrovermiculatus | Hyperolius_rubrovermiculatus | 1 | 2.6 | 75 | 421.83 |
| amphibians | Hyperolius_rwandae | Hyperolius_rwandae | 0 | 0.77 | 80 | 25205.71 |
| amphibians | Hyperolius_semidiscus | Hyperolius_semidiscus | 0 | 3.81 | 115 | 142267.32 |
| amphibians | Hyperolius_spinigularis | Hyperolius_spinigularis | 1 | 2.35 | 175 | 1022.35 |
| amphibians | Hyperolius_substriatus | Hyperolius_substriatus | 0 | 6.77 | 37.5 | 579853 |
| amphibians | Hyperolius_thomensis | Hyperolius_thomensis | 1 | 9.03 | 42.5 | 586.43 |
| amphibians | Hyperolius_tuberilinguis | Hyperolius_tuberilinguis | 0 | 5.36 | 318 | 1134975.17 |
| amphibians | Hyperolius_veithi | Hyperolius_veithi | 0 | 4.16 | 50 | 7.07 |
| amphibians | Hyperolius_viridiflavus | Hyperolius_viridiflavus | 0 | 6.77 | 331 | 1960115.39 |
| amphibians | Hypogeophis_rostratus | Hypogeophis_rostratus | 0 | 18.29 | 12.5 | 237.67 |
| amphibians | Hypopachus_variolosus | Hypopachus_variolosus | 0 | 13.33 | 700 | 759251.16 |
| amphibians | Hypselotriton_cyanurus | Cynops_cyanurus | 1 | 1.8 | 334.5 | 9756.69 |
| amphibians | Hypselotriton_orientalis | Cynops_orientalis | 0 | 1.06 | 181.5 | 392443.52 |
| amphibians | Ichthyophis_beddomei | Ichthyophis_beddomei | 0 | 7.78 | 17.5 | 17913.69 |
| amphibians | Ichthyophis_glutinosus | Ichthyophis_glutinosus | 1 | 34.42 | 40 | 8162.63 |
| amphibians | Ichthyophis_nguyenorum | Ichthyophis_nguyenorum | 0 | 11.17 | 15 | 62546.98 |
| amphibians | Ichthyophis_sikkimensis | Ichthyophis_sikkimensis | 0 | 12.29 | 32.5 | 347.19 |
| amphibians | Ichthyosaura_alpestris | Ichthyosaura_alpestris | 0 | 5.49 | 290 | 1366049.96 |
| amphibians | Ikakogi_tayrona | Ikakogi_tayrona | 1 | 3.47 | 55.5 | 4097.01 |
| amphibians | Incilius_alvarius | Incilius_alvarius | 0 | 848.59 | 7750 | 358467.99 |
| amphibians | Incilius_campbelli | Incilius_campbelli | 0 | 74.11 | 5000 | 4812.75 |
| amphibians | Incilius_canaliferus | Incilius_canaliferus | 0 | 12.53 | 5000 | 13767.33 |
| amphibians | Incilius_coccifer | Incilius_coccifer | 0 | 100.93 | 4000 | 139093.15 |

| | | | | | | |
|---|---|---|---|---|---|---|
| amphibians | Incilius_coniferus | Incilius_coniferus | 0 | 88.3 | 3000 | 120109.56 |
| amphibians | Incilius_fastidiosus | Incilius_fastidiosus | 1 | 22.16 | 85 | 252.38 |
| amphibians | Incilius_holdridgei | Incilius_holdridgei | 1 | 13.33 | 91 | 7.62 |
| amphibians | Incilius_ibarrai | Incilius_ibarrai | 0 | 85.33 | 3000 | 8185.36 |
| amphibians | Incilius_luetkenii | Incilius_luetkenii | 0 | 129.82 | 3300 | 107380.94 |
| amphibians | Incilius_nebulifer | Incilius_nebulifer | 0 | 243.97 | 10010.5 | 668550.66 |
| amphibians | Incilius_valliceps | Incilius_valliceps | 0 | 243.97 | 27 | 565201.89 |
| amphibians | Indirana_semipalmata | Indirana_semipalmata | 0 | 3.02 | 243 | 21471.76 |
| amphibians | Ischnocnema_guentheri | Ischnocnema_guentheri | 0 | 8.42 | 30 | 110849.12 |
| amphibians | Ischnocnema_henselii | Ischnocnema_henselii | 0 | 4.93 | 25 | 223011.77 |
| amphibians | Ischnocnema_holti | Ischnocnema_holti | 0 | 2.52 | 15 | 779.61 |
| amphibians | Ischnocnema_juipoca | Ischnocnema_juipoca | 0 | 1.37 | 15 | 443008.05 |
| amphibians | Ischnocnema_melanopygia | Ischnocnema_melanopygia | 1 | 0.93 | 12 | 9.94 |
| amphibians | Ischnocnema_parva | Ischnocnema_parva | 0 | 1.25 | 19 | 298074.01 |
| amphibians | Ischnocnema_randorum | Ischnocnema_randorum | 0 | 0.42 | 7 | 166.57 |
| amphibians | Isthmohyla_calypsa | Isthmohyla_calypsa | 1 | 5.8 | 23 | 323.32 |
| amphibians | Isthmohyla_insolita | Isthmohyla_insolita | 1 | 4.54 | 33.5 | 66.36 |
| amphibians | Isthmohyla_lancasteri | Isthmohyla_lancasteri | 0 | 5.85 | 50 | 5171.36 |
| amphibians | Isthmohyla_pictipes | Isthmohyla_pictipes | 1 | 7.9 | 124.5 | 2795.17 |
| amphibians | Isthmohyla_pseudopuma | Isthmohyla_pseudopuma | 0 | 12.53 | 2150 | 5381.23 |
| amphibians | Isthmohyla_rivularis | Isthmohyla_rivularis | 1 | 6.77 | 90 | 3827.42 |
| amphibians | Isthmohyla_tica | Isthmohyla_tica | 1 | 6.77 | 227.5 | 5714.19 |
| amphibians | Isthmohyla_zeteki | Isthmohyla_zeteki | 1 | 3.54 | 144 | 1242.58 |
| amphibians | Itapotihyla_langsdorffii | Itapotihyla_langsdorffii | 0 | 150.53 | 4326.5 | 729247.13 |
| amphibians | Julianus_pinimus | Scinax_pinima | 0 | 1.89 | 300 | 263114.51 |
| amphibians | Julianus_uruguayus | Scinax_uruguayus | 0 | 2.11 | 399 | 122136.43 |
| amphibians | Kalophrynus_minusculus | Kalophrynus_minusculus | 0 | 3.47 | 40 | 6442.96 |
| amphibians | Kalophrynus_pleurostigma | Kalophrynus_pleurostigma | 0 | 19.92 | 3334 | 687690.04 |
| amphibians | Kaloula_conjuncta | Kaloula_conjuncta | 0 | 9.03 | 1650 | 27864.97 |
| amphibians | Kaloula_picta | Kaloula_picta | 0 | 16.4 | 2264.5 | 272030.78 |
| amphibians | Kaloula_pulchra | Kaloula_pulchra | 0 | 57.01 | 4126 | 2300321.31 |
| amphibians | Kassina_cassinoides | Kassina_cassinoides | 0 | 11.04 | 185 | 1590492.98 |
| amphibians | Kassina_cochranae | Kassina_cochranae | 0 | 6.27 | 180 | 174153.54 |
| amphibians | Kassina_fusca | Kassina_fusca | 0 | 5.8 | 550 | 819826.69 |

| | | | | | | |
|---|---|---|---|---|---|---|
| amphibians | Kassina_kuvangensis | Kassina_kuvangensis | 0 | 11.77 | 129.5 | 561513.54 |
| amphibians | Kassina_lamottei | Kassina_lamottei | 0 | 9.03 | 37 | 72132.14 |
| amphibians | Kassina_schioetzi | Kassina_schioetzi | 0 | 5.36 | 185 | 181030.7 |
| amphibians | Kassina_senegalensis | Kassina_senegalensis | 0 | 12.53 | 350 | 12196935.29 |
| amphibians | Kassina_somalica | Kassina_somalica | 0 | 5.8 | 330 | 612349.84 |
| amphibians | Kurixalus_eiffingeri | Kurixalus_eiffingeri | 0 | 7.29 | 44.5 | 12611.51 |
| amphibians | Kurixalus_idiootocus | Kurixalus_idiootocus | 0 | 7.84 | 185 | 26688.63 |
| amphibians | Laotriton_laoensis | Laotriton_laoensis | 1 | 23.04 | 15 | 4559.12 |
| amphibians | Leiopelma_archeyi | Leiopelma_archeyi | 1 | 4.54 | 9.5 | 1376.63 |
| amphibians | Leiopelma_hamiltoni | Leiopelma_hamiltoni | 1 | 12.53 | 10 | 9.65 |
| amphibians | Leiopelma_hochstetteri | Leiopelma_hochstetteri | 0 | 9.03 | 90.5 | 10109.77 |
| amphibians | Lepidobatrachus_laevis | Lepidobatrachus_laevis | 0 | 190.79 | 2539 | 465963.74 |
| amphibians | Lepidobatrachus_llanensis | Lepidobatrachus_llanensis | 0 | 104.27 | 2190 | 302269.2 |
| amphibians | Leptobrachella_alpina | Leptolalax_alpinus | 0 | 2.73 | 129 | 4407.05 |
| amphibians | Leptobrachella_liui | Leptolalax_liui | 0 | 1.83 | 83 | 328958.9 |
| amphibians | Leptobrachium_ailaonicum | Leptobrachium_ailaonicum | 0 | 67.9 | 333 | 15284.7 |
| amphibians | Leptobrachium_boringii | Leptobrachium_boringii | 1 | 82.42 | 312.5 | 65807.47 |
| amphibians | Leptobrachium_leishanense | Leptobrachium_leishanense | 1 | 90.43 | 324 | 2876.86 |
| amphibians | Leptobrachium_liui | Leptobrachium_liui | 0 | 75.46 | 352 | 269304.84 |
| amphibians | Leptobrachium_montanum | Leptobrachium_montanum | 0 | 66.4 | 217 | 26539.32 |
| amphibians | Leptodactylus_albilabris | Leptodactylus_albilabris | 0 | 10.34 | 137.5 | 10649.91 |
| amphibians | Leptodactylus_bolivianus | Leptodactylus_bolivianus | 0 | 310.17 | 1325 | 2780429.65 |
| amphibians | Leptodactylus_bufonius | Leptodactylus_bufonius | 0 | 25.82 | 512.5 | 1569066.46 |
| amphibians | Leptodactylus_didymus | Leptodactylus_didymus | 0 | 18.46 | 326 | 304466.13 |
| amphibians | Leptodactylus_discodactylus | Leptodactylus_discodactylus | 0 | 3.47 | 284 | 1332439.43 |
| amphibians | Leptodactylus_fallax | Leptodactylus_fallax | 1 | 1153.83 | 34 | 152.03 |
| amphibians | Leptodactylus_fragilis | Leptodactylus_fragilis | 0 | 7.29 | 137.5 | 1557081.89 |
| amphibians | Leptodactylus_furnarius | Leptodactylus_furnarius | 0 | 7.29 | 73 | 1429867.37 |
| amphibians | Leptodactylus_fuscus | Leptodactylus_fuscus | 0 | 15.03 | 149 | 12048757.37 |
| amphibians | Leptodactylus_gracilis | Leptodactylus_gracilis | 0 | 11.04 | 175 | 2227779.98 |
| amphibians | Leptodactylus_jolyi | Leptodactylus_jolyi | 0 | 14.16 | 100 | 1113014.95 |
| amphibians | Leptodactylus_knudseni | Leptodactylus_knudseni | 0 | 581.84 | 1000 | 5858442.05 |
| amphibians | Leptodactylus_labyrinthicus | Leptodactylus_labyrinthicus | 0 | 765.22 | 2124.5 | 3926794.41 |
| amphibians | Leptodactylus_laticeps | Leptodactylus_laticeps | 0 | 188.23 | 3976 | 433225.6 |

| | | | | | | |
|---|---|---|---|---|---|---|
| amphibians | Leptodactylus_latinasus | Leptodactylus_latinasus | 0 | 5.36 | 481 | 1908729.09 |
| amphibians | Leptodactylus_latrans | Leptodactylus_latrans | 0 | 310.17 | 2782.5 | 179747.56 |
| amphibians | Leptodactylus_leptodactyloides | Leptodactylus_leptodactyloides | 0 | 16.12 | 888.5 | 6143679.61 |
| amphibians | Leptodactylus_macrosternum | Leptodactylus_chaquensis | 0 | 61.58 | 8225.5 | 9590666.54 |
| amphibians | Leptodactylus_melanonotus | Leptodactylus_melanonotus | 0 | 17.85 | 1009 | 833155.68 |
| amphibians | Leptodactylus_mystaceus | Leptodactylus_mystaceus | 0 | 19.92 | 251 | 6802464.72 |
| amphibians | Leptodactylus_mystacinus | Leptodactylus_mystacinus | 0 | 27.26 | 286 | 4307146.39 |
| amphibians | Leptodactylus_natalensis | Leptodactylus_natalensis | 0 | 12.53 | 1356.5 | 701823.02 |
| amphibians | Leptodactylus_pentadactylus | Leptodactylus_pentadactylus | 0 | 907.54 | 900.5 | 6672526.95 |
| amphibians | Leptodactylus_plaumanni | Leptodactylus_plaumanni | 0 | 8.42 | 225 | 211860.66 |
| amphibians | Leptodactylus_podicipinus | Leptodactylus_podicipinus | 0 | 14.16 | 2330.5 | 6340303.57 |
| amphibians | Leptodactylus_rhodomystax | Leptodactylus_rhodomystax | 0 | 74.11 | 268.5 | 4578521 |
| amphibians | Leptodactylus_rhodonotus | Leptodactylus_rhodonotus | 0 | 74.11 | 201 | 886872.68 |
| amphibians | Leptodactylus_riveroi | Leptodactylus_riveroi | 0 | 39.31 | 1000 | 924607.47 |
| amphibians | Leptodactylus_savagei | Leptodactylus_savagei | 0 | 765.22 | 1000 | 255617.77 |
| amphibians | Leptodactylus_syphax | Leptodactylus_syphax | 0 | 54.82 | 526 | 3088798.52 |
| amphibians | Leptodactylus_troglodytes | Leptodactylus_troglodytes | 0 | 13.82 | 263.5 | 1873310.41 |
| amphibians | Leptodactylus_wagneri | Leptodactylus_wagneri | 0 | 54.17 | 2075 | 1058311.35 |
| amphibians | Leptopelis_flavomaculatus | Leptopelis_flavomaculatus | 0 | 32.83 | 141 | 858622.21 |
| amphibians | Leptopelis_natalensis | Leptopelis_natalensis | 0 | 25.82 | 192 | 36599.57 |
| amphibians | Leptopelis_spiritusnoctis | Leptopelis_spiritusnoctis | 0 | 12.53 | 95 | 521402.46 |
| amphibians | Leptopelis_viridis | Leptopelis_viridis | 0 | 11.04 | 203.5 | 3428870.05 |
| amphibians | Leptophryne_borbonica | Leptophryne_borbonica | 0 | 11.04 | 250 | 269261.2 |
| amphibians | Leucostethus_fraterdanieli | Colostethus_fraterdanieli | 0 | 1.5 | 12 | 42537.71 |
| amphibians | Limnodynastes_convexiusculus | Limnodynastes_convexiusculus | 0 | 21.02 | 668 | 692576.27 |
| amphibians | Limnodynastes_dorsalis | Limnodynastes_dorsalis | 0 | 37.61 | 1000 | 259159.3 |
| amphibians | Limnodynastes_dumerilii | Limnodynastes_dumerilii | 0 | 57.01 | 2605 | 921457.09 |
| amphibians | Limnodynastes_fletcheri | Limnodynastes_fletcheri | 0 | 15.03 | 300 | 1212018.53 |
| amphibians | Limnodynastes_interioris | Limnodynastes_interioris | 0 | 74.11 | 962.5 | 273943.64 |
| amphibians | Limnodynastes_lignarius | Limnodynastes_lignarius | 0 | 22.16 | 350 | 318842.49 |
| amphibians | Limnodynastes_peronii | Limnodynastes_peronii | 0 | 37.61 | 1153.5 | 939844.67 |
| amphibians | Limnodynastes_salmini | Limnodynastes_salmini | 0 | 42.85 | 2030 | 521308.28 |
| amphibians | Limnodynastes_tasmaniensis | Limnodynastes_tasmaniensis | 0 | 11.4 | 820 | 2878716.72 |
| amphibians | Limnodynastes_terraereginae | Limnodynastes_terraereginae | 0 | 48.58 | 768.5 | 706376.63 |

| | | | | | | |
|---|---|---|---|---|---|---|
| amphibians | Limnomedusa_macroglossa | Limnomedusa_macroglossa | 0 | 25.82 | 398 | 398365.42 |
| amphibians | Limnonectes_arathooni | Limnonectes_arathooni | 1 | 5.18 | 19.5 | 3387.39 |
| amphibians | Limnonectes_blythii | Limnonectes_blythii | 0 | 2304.97 | 1839 | 416463.01 |
| amphibians | Limnonectes_finchi | Limnonectes_finchi | 0 | 8.02 | 42.5 | 16721.93 |
| amphibians | Limnonectes_fujianensis | Limnonectes_fujianensis | 0 | 14.08 | 119.5 | 446729.46 |
| amphibians | Limnonectes_grunniens | Limnonectes_grunniens | 0 | 581.84 | 1460 | 330320.89 |
| amphibians | Limnonectes_ibanorum | Limnonectes_ibanorum | 0 | 275.7 | 1122 | 105854.46 |
| amphibians | Limnonectes_khasianus | Limnonectes_khasianus | 0 | 13.33 | 142.5 | 246140.86 |
| amphibians | Limnonectes_kuhlii | Limnonectes_kuhlii | 0 | 66.4 | 53 | 2038423.89 |
| amphibians | Limnonectes_leporinus | Limnonectes_leporinus | 0 | 639.14 | 1852 | 346338.31 |
| amphibians | Limnonectes_leytensis | Limnonectes_leytensis | 0 | 17.46 | 46 | 144027.56 |
| amphibians | Limnonectes_limborgi | Limnonectes_limborgi | 0 | 4.54 | 12 | 432176.21 |
| amphibians | Limnonectes_macrodon | Limnonectes_macrodon | 0 | 387.87 | 1144.5 | 66067.28 |
| amphibians | Limnonectes_magnus | Limnonectes_magnus | 0 | 522 | 178 | 18642.24 |
| amphibians | Limnonectes_microdiscus | Limnonectes_microdiscus | 0 | 32.83 | 45 | 142980.11 |
| amphibians | Limnonectes_palavanensis | Limnonectes_palavanensis | 0 | 5.36 | 42.5 | 126845.17 |
| amphibians | Limnonectes_parvus | Limnonectes_parvus | 0 | 2.11 | 9.5 | 6349.28 |
| amphibians | Limnonectes_visayanus | Limnonectes_visayanus | 0 | 85.92 | 178 | 5846.23 |
| amphibians | Lissotriton_boscai | Lissotriton_boscai | 0 | 1.5 | 175 | 201777.08 |
| amphibians | Lissotriton_helveticus | Lissotriton_helveticus | 0 | 1.7 | 352.5 | 1084367.76 |
| amphibians | Lissotriton_italicus | Lissotriton_italicus | 0 | 1.5 | 350 | 75200.41 |
| amphibians | Lissotriton_montandoni | Lissotriton_montandoni | 0 | 2.85 | 247.5 | 123049.57 |
| amphibians | Lissotriton_vulgaris | Lissotriton_vulgaris | 0 | 3.16 | 300 | 898593.44 |
| amphibians | Lithobates_areolatus | Rana_areolata | 0 | 198.59 | 6000 | 417915.34 |
| amphibians | Lithobates_berlandieri | Rana_berlandieri | 0 | 188.23 | 500 | 888950.83 |
| amphibians | Lithobates_blairi | Rana_blairi | 0 | 159.41 | 5000 | 1423691.13 |
| amphibians | Lithobates_capito | Rana_capito | 1 | 154.93 | 3994 | 241845.51 |
| amphibians | Lithobates_chiricahuensis | Rana_chiricahuensis | 1 | 289.15 | 892.5 | 228519.32 |
| amphibians | Lithobates_forreri | Rana_forreri | 0 | 159.41 | 550 | 227810.28 |
| amphibians | Lithobates_onca | Rana_onca | 1 | 71.48 | 225 | 19570.15 |
| amphibians | Lithobates_palmipes | Rana_palmipes | 0 | 220.47 | 3468 | 9806600.5 |
| amphibians | Lithobates_palustris | Rana_palustris | 0 | 66.4 | 2116 | 2479424.19 |
| amphibians | Lithobates_pipiens | Rana_pipiens | 0 | 146.22 | 4300 | 6012762.57 |
| amphibians | Lithobates_sevosus | Rana_sevosa | 1 | 122.13 | 3750 | 21134.38 |

| | | | | | | |
|---|---|---|---|---|---|---|
| amphibians | Lithobates_sphenocephalus | Rana_sphenocephala | 0 | 226.19 | 4449 | 1727205.35 |
| amphibians | Lithobates_tarahumarae | Rana_tarahumarae | 1 | 159.41 | 1363.5 | 95619.47 |
| amphibians | Lithobates_taylori | Rana_taylori | 0 | 68.91 | 1000 | 28326.39 |
| amphibians | Lithodytes_lineatus | Lithodytes_lineatus | 0 | 15.93 | 195 | 7015371.83 |
| amphibians | Litoria_adelaidensis | Litoria_adelaidensis | 0 | 19.92 | 110 | 115762.05 |
| amphibians | Litoria_aurifera | Litoria_aurifera | 0 | 0.77 | 85.5 | 12301.73 |
| amphibians | Litoria_bicolor | Litoria_bicolor | 0 | 2.11 | 518 | 841293.87 |
| amphibians | Litoria_burrowsi | Litoria_burrowsi | 0 | 19.92 | 95 | 18271.89 |
| amphibians | Litoria_coplandi | Litoria_coplandi | 0 | 6.77 | 216 | 736072.34 |
| amphibians | Litoria_dentata | Litoria_dentata | 0 | 8.13 | 1646 | 69176.2 |
| amphibians | Litoria_ewingii | Litoria_ewingii | 0 | 11.04 | 600 | 614726.57 |
| amphibians | Litoria_fallax | Litoria_fallax | 0 | 2.6 | 813 | 591734.54 |
| amphibians | Litoria_freycineti | Litoria_freycineti | 0 | 10.34 | 477.5 | 125683.59 |
| amphibians | Litoria_havina | Litoria_havina | 0 | 4.54 | 19.5 | 1938.33 |
| amphibians | Litoria_hilli | Litoria_hilli | 0 | 14.76 | 46 | 211.07 |
| amphibians | Litoria_inermis | Litoria_inermis | 0 | 4.16 | 213 | 2018405.39 |
| amphibians | Litoria_iris | Litoria_iris | 0 | 6.77 | 20.5 | 31474.28 |
| amphibians | Litoria_jervisiensis | Litoria_jervisiensis | 0 | 19.92 | 920 | 78207.59 |
| amphibians | Litoria_latopalmata | Litoria_latopalmata | 0 | 6.27 | 268 | 1925902 |
| amphibians | Litoria_littlejohni | Litoria_littlejohni | 1 | 35.97 | 132 | 6476.17 |
| amphibians | Litoria_longirostris | Litoria_longirostris | 0 | 1.5 | 42.5 | 525.44 |
| amphibians | Litoria_meiriana | Litoria_meiriana | 0 | 0.89 | 72.5 | 367981 |
| amphibians | Litoria_microbelos | Litoria_microbelos | 0 | 0.57 | 474.5 | 223558.04 |
| amphibians | Litoria_nasuta | Litoria_nasuta | 0 | 15.93 | 84.5 | 1189277.51 |
| amphibians | Litoria_nigrofrenata | Litoria_nigrofrenata | 0 | 11.04 | 447.5 | 116520.11 |
| amphibians | Litoria_olongburensis | Litoria_olongburensis | 1 | 3.19 | 117.5 | 9065.69 |
| amphibians | Litoria_pallida | Litoria_pallida | 0 | 5.8 | 485.5 | 1442166.84 |
| amphibians | Litoria_peronii | Litoria_peronii | 0 | 32.83 | 3783.5 | 1260639.65 |
| amphibians | Litoria_revelata | Litoria_revelata | 0 | 3.81 | 574.5 | 66672.92 |
| amphibians | Litoria_rothii | Litoria_rothii | 0 | 16.87 | 596 | 1441739.84 |
| amphibians | Litoria_rubella | Litoria_rubella | 0 | 6.77 | 274 | 4908530.23 |
| amphibians | Litoria_tornieri | Litoria_tornieri | 0 | 3.81 | 217.5 | 547460.07 |
| amphibians | Litoria_verreauxii | Litoria_verreauxii | 0 | 3.81 | 723.5 | 362379.98 |
| amphibians | Litoria_vocivincens | Litoria_vocivincens | 0 | 1.41 | 300 | 28902.42 |

| | | | | | | |
|---|---|---|---|---|---|---|
| amphibians | Litoria_watjulumensis | Litoria_watjulumensis | 0 | 41.05 | 167.5 | 293581.11 |
| amphibians | Liua_shihi | Liua_shihi | 0 | 6.6 | 30.5 | 48801.94 |
| amphibians | Liua_tsinpaensis | Liua_tsinpaensis | 1 | 5.73 | 14 | 432.81 |
| amphibians | Liuhurana_shuchinae | Rana_shuchinae | 0 | 7.57 | 180 | 35189.11 |
| amphibians | Lyciasalamandra_antalyana | Lyciasalamandra_antalyana | 1 | 4.42 | 2 | 588.34 |
| amphibians | Lyciasalamandra_atifi | Lyciasalamandra_atifi | 1 | 11.13 | 2 | 2610.12 |
| amphibians | Lyciasalamandra_fazilae | Lyciasalamandra_fazilae | 1 | 5.73 | 2 | 552.42 |
| amphibians | Lyciasalamandra_flavimembris | Lyciasalamandra_flavimembris | 1 | 43.05 | 2 | 595.12 |
| amphibians | Lyciasalamandra_luschani | Lyciasalamandra_luschani | 1 | 4.11 | 2 | 1801.91 |
| amphibians | Lysapsus_bolivianus | Lysapsus_boliviana | 0 | 0.55 | 19.5 | 2141528.53 |
| amphibians | Lysapsus_limellum | Lysapsus_limellum | 0 | 1.14 | 113.5 | 1061961.66 |
| amphibians | Macrogenioglottus_alipioi | Macrogenioglottus_alipioi | 0 | 158.51 | 3650 | 557849.16 |
| amphibians | Madecassophryne_truebae | Madecassophryne_truebae | 1 | 0.89 | 18.5 | 1447.66 |
| amphibians | Mannophryne_collaris | Mannophryne_collaris | 1 | 2.6 | 14 | 1687.61 |
| amphibians | Mannophryne_herminae | Mannophryne_herminae | 0 | 1.89 | 9.5 | 599.18 |
| amphibians | Mannophryne_lamarcai | Mannophryne_lamarcai | 1 | 2.3 | 10 | 824.85 |
| amphibians | Mannophryne_olmonae | Mannophryne_olmonae | 1 | 0.73 | 12 | 159.17 |
| amphibians | Mannophryne_trinitatis | Mannophryne_trinitatis | 0 | 1.89 | 7 | 1339.01 |
| amphibians | Mantella_aurantiaca | Mantella_aurantiaca | 1 | 2.35 | 126 | 10.39 |
| amphibians | Mantella_baroni | Mantella_baroni | 0 | 2.11 | 130 | 32997.28 |
| amphibians | Mantella_betsileo | Mantella_betsileo | 0 | 1.33 | 35 | 84844.81 |
| amphibians | Mantella_cowanii | Mantella_cowanii | 1 | 2.35 | 39.5 | 252.56 |
| amphibians | Mantella_crocea | Mantella_crocea | 1 | 1.02 | 53.5 | 535.94 |
| amphibians | Mantella_haraldmeieri | Mantella_haraldmeieri | 1 | 2.11 | 64.5 | 1976.39 |
| amphibians | Mantella_laevigata | Mantella_laevigata | 0 | 2.11 | 6.5 | 17602.27 |
| amphibians | Mantella_madagascariensis | Mantella_madagascariensis | 1 | 2.35 | 26.5 | 5208.68 |
| amphibians | Mantidactylus_argenteus | Mantidactylus_argenteus | 0 | 2.35 | 12 | 25689.66 |
| amphibians | Mantidactylus_brevipalmatus | Mantidactylus_brevipalmatus | 0 | 7.84 | 30 | 39368.29 |
| amphibians | Mantidactylus_majori | Mantidactylus_majori | 0 | 9.03 | 15 | 34406.4 |
| amphibians | Mantidactylus_ulcerosus | Mantidactylus_ulcerosus | 0 | 7.84 | 60 | 50502.91 |
| amphibians | Melanophryniscus_admirabilis | Melanophryniscus_admirabilis | 1 | 5.8 | 20 | 2402.14 |
| amphibians | Melanophryniscus_atroluteus | Melanophryniscus_atroluteus | 0 | 1.69 | 125 | 203412.14 |
| amphibians | Melanophryniscus_cambaraensis | Melanophryniscus_cambaraensis | 1 | 4.54 | 100 | 633.8 |
| amphibians | Melanophryniscus_montevidensis | Melanophryniscus_montevidensis | 0 | 1.89 | 181 | 123386.17 |

| | | | | | | |
|---|---|---|---|---|---|---|
| amphibians | Melanophryniscus_moreirae | Melanophryniscus_moreirae | 0 | 2.11 | 78 | 3184.71 |
| amphibians | Melanophryniscus_setiba | Melanophryniscus_setiba | 1 | 0.28 | 8 | 144.62 |
| amphibians | Melanophryniscus_stelzneri | Melanophryniscus_stelzneri | 0 | 2.11 | 351 | 35188.74 |
| amphibians | Melanophryniscus_vilavelhensis | Melanophryniscus_vilavelhensis | 1 | 0.35 | 8 | 402.47 |
| amphibians | Mercurana_myristicapalustris | Mercurana_myristicapalustris | 1 | 25.82 | 125 | 141.57 |
| amphibians | Meristogenys_amoropalamus | Meristogenys_amoropalamus | 0 | 29.04 | 232 | 12721.21 |
| amphibians | Meristogenys_jerboa | Meristogenys_jerboa | 1 | 70.7 | 1531 | 7986.39 |
| amphibians | Meristogenys_orphnocnemis | Meristogenys_orphnocnemis | 0 | 32.83 | 800 | 42640.06 |
| amphibians | Mertensiella_caucasica | Mertensiella_caucasica | 0 | 7.16 | 51.5 | 14693.64 |
| amphibians | Mertensophryne_anotis | Mertensophryne_anotis | 0 | 8.42 | 81 | 15134.69 |
| amphibians | Mertensophryne_howelli | Mertensophryne_howelli | 1 | 7.84 | 57.5 | 50.33 |
| amphibians | Mertensophryne_lindneri | Mertensophryne_lindneri | 0 | 3.16 | 81 | 387516.44 |
| amphibians | Mertensophryne_lonnbergi | Mertensophryne_lonnbergi | 1 | 7.29 | 125 | 9756.86 |
| amphibians | Mertensophryne_loveridgei | Mertensophryne_loveridgei | 0 | 4.93 | 131 | 57222.56 |
| amphibians | Mertensophryne_melanopleura | Mertensophryne_melanopleura | 0 | 1.5 | 34 | 299350.79 |
| amphibians | Mertensophryne_micranotis | Mertensophryne_micranotis | 0 | 1.02 | 39 | 124039.54 |
| amphibians | Mertensophryne_nyikae | Mertensophryne_nyikae | 0 | 7.29 | 125 | 1429.99 |
| amphibians | Mertensophryne_taitana | Mertensophryne_taitana | 0 | 3.81 | 237.5 | 1144764.24 |
| amphibians | Mertensophryne_usambarae | Mertensophryne_usambarae | 1 | 7.84 | 57.5 | 42.54 |
| amphibians | Mertensophryne_uzunguensis | Mertensophryne_uzunguensis | 1 | 2.11 | 188 | 14754.37 |
| amphibians | Metacrinia_nichollsi | Metacrinia_nichollsi | 0 | 1.69 | 21 | 16909.97 |
| amphibians | Metaphrynella_sundana | Metaphrynella_sundana | 0 | 2.11 | 689.5 | 85353.7 |
| amphibians | Microbatrachella_capensis | Microbatrachella_capensis | 1 | 0.4 | 19.5 | 120.92 |
| amphibians | Microcaecilia_dermatophaga | Microcaecilia_dermatophaga | 0 | 2.27 | 5 | 1334.74 |
| amphibians | Microhyla_achatina | Microhyla_achatina | 0 | 1.69 | 20 | 94800.65 |
| amphibians | Microhyla_butleri | Microhyla_butleri | 0 | 1.33 | 9 | 2590639.86 |
| amphibians | Microhyla_heymonsi | Microhyla_heymonsi | 0 | 2.2 | 253.5 | 3505075.57 |
| amphibians | Microhyla_mixtura | Microhyla_mixtura | 0 | 1.5 | 528.5 | 85898.21 |
| amphibians | Microhyla_okinavensis | Microhyla_okinavensis | 0 | 2.35 | 515 | 2725.68 |
| amphibians | Microhyla_ornata | Microhyla_ornata | 0 | 2.6 | 693.5 | 790904.3 |
| amphibians | Microhyla_pulchra | Microhyla_pulchra | 0 | 3.98 | 633 | 2011158.5 |
| amphibians | Microkayla_boettgeri | Psychrophrynella_boettgeri | 1 | 0.43 | 6.5 | 10.09 |
| amphibians | Microkayla_illampu | Psychrophrynella_illampu | 1 | 1.01 | 8 | 10.17 |
| amphibians | Microkayla_wettsteini | Psychrophrynella_wettsteini | 1 | 2.99 | 12.5 | 63.97 |

| | | | | | | |
|---|---|---|---|---|---|---|
| amphibians | Minervarya_keralensis | Fejervarya_keralensis | 1 | 21.02 | 162.5 | 63835.38 |
| amphibians | Minervarya_syhadrensis | Fejervarya_syhadrensis | 0 | 6.52 | 53.5 | 2634149.58 |
| amphibians | Minyobates_steyermarki | Minyobates_steyermarki | 1 | 0.48 | 6 | 9.42 |
| amphibians | Mixophyes_balbus | Mixophyes_balbus | 1 | 104.27 | 577.5 | 76843.43 |
| amphibians | Mixophyes_carbinensis | Mixophyes_carbinensis | 0 | 46.62 | 227.5 | 746.86 |
| amphibians | Mixophyes_coggeri | Mixophyes_coggeri | 0 | 118.4 | 500 | 14191.38 |
| amphibians | Mixophyes_fasciolatus | Mixophyes_fasciolatus | 0 | 107.68 | 1475 | 184187.49 |
| amphibians | Mixophyes_fleayi | Mixophyes_fleayi | 1 | 71.48 | 935 | 4870.91 |
| amphibians | Mixophyes_iteratus | Mixophyes_iteratus | 1 | 163.99 | 2763.5 | 23086.44 |
| amphibians | Mixophyes_schevilli | Mixophyes_schevilli | 0 | 79.58 | 427 | 17561.59 |
| amphibians | Morerella_cyanophthalma | Morerella_cyanophthalma | 1 | 3.47 | 87 | 375.39 |
| amphibians | Myersiella_microps | Myersiella_microps | 0 | 5.36 | 22 | 171653.06 |
| amphibians | Myersiohyla_chamaeleo | Myersiohyla_chamaeleo | 0 | 16.78 | 108 | 405.56 |
| amphibians | Myersiohyla_neblinaria | Myersiohyla_neblinaria | 0 | 21.7 | 238 | 757.06 |
| amphibians | Myobatrachus_gouldii | Myobatrachus_gouldii | 0 | 19.92 | 24.5 | 197427.13 |
| amphibians | Nannophryne_variegata | Nannophryne_variegata | 0 | 11.04 | 103.5 | 135856.65 |
| amphibians | Nannophrys_ceylonensis | Nannophrys_ceylonensis | 1 | 14.16 | 27.5 | 8067.27 |
| amphibians | Nanorana_annandalii | Nanorana_annandalii | 0 | 12.15 | 159.5 | 37343.07 |
| amphibians | Nanorana_blanfordii | Nanorana_blanfordii | 0 | 9.67 | 85.5 | 168900.02 |
| amphibians | Nanorana_ercepeae | Nanorana_ercepeae | 0 | 76.81 | 187.5 | 49998.97 |
| amphibians | Nanorana_kangxianensis | Nanorana_kangxianensis | 1 | 108.03 | 782 | 912.14 |
| amphibians | Nanorana_liebigii | Nanorana_liebigii | 0 | 570.83 | 104.5 | 261384.53 |
| amphibians | Nanorana_minica | Nanorana_minica | 0 | 5.8 | 72.5 | 101689.4 |
| amphibians | Nanorana_parkeri | Nanorana_parkeri | 0 | 17.85 | 251 | 384079.59 |
| amphibians | Nanorana_pleskei | Nanorana_pleskei | 0 | 6.27 | 50 | 142696.5 |
| amphibians | Nanorana_polunini | Nanorana_polunini | 0 | 15.93 | 85.5 | 121833.7 |
| amphibians | Nanorana_rostandi | Nanorana_rostandi | 0 | 25.82 | 443.5 | 76398.18 |
| amphibians | Nanorana_taihangnica | Nanorana_taihangnica | 0 | 116.2 | 456.5 | 70363.44 |
| amphibians | Nanorana_unculuanus | Nanorana_unculuanus | 1 | 50.6 | 97 | 47670.64 |
| amphibians | Nanorana_ventripunctata | Nanorana_ventripunctata | 0 | 10.61 | 311 | 29325.26 |
| amphibians | Nanorana_yunnanensis | Nanorana_yunnanensis | 1 | 137.85 | 811.5 | 654146.94 |
| amphibians | Nasikabatrachus_sahyadrensis | Nasikabatrachus_sahyadrensis | 0 | 74.11 | 2350 | 11357.47 |
| amphibians | Natalobatrachus_bonebergi | Natalobatrachus_bonebergi | 1 | 4.16 | 92.5 | 5826.82 |
| amphibians | Nectophryne_afra | Nectophryne_afra | 0 | 4.54 | 31 | 951359.52 |

| | | | | | | |
|---|---|---|---|---|---|---|
| amphibians | Nectophryne_batesii | Nectophryne_batesii | 0 | 1.17 | 37.5 | 946911.72 |
| amphibians | Nectophrynoides_cryptus | Nectophrynoides_cryptus | 1 | 3.16 | 20.5 | 83.01 |
| amphibians | Nectophrynoides_laticeps | Nectophrynoides_laticeps | 1 | 1.02 | 45 | 17.18 |
| amphibians | Nectophrynoides_minutus | Nectophrynoides_minutus | 1 | 0.77 | 25.5 | 438.56 |
| amphibians | Nectophrynoides_paulae | Nectophrynoides_paulae | 1 | 1.21 | 20 | 17.95 |
| amphibians | Nectophrynoides_poyntoni | Nectophrynoides_poyntoni | 1 | 3.16 | 9 | 3.09 |
| amphibians | Nectophrynoides_tornieri | Nectophrynoides_tornieri | 0 | 3.16 | 23 | 2879.87 |
| amphibians | Nectophrynoides_vestergaardi | Nectophrynoides_vestergaardi | 1 | 1.02 | 32 | 773.25 |
| amphibians | Nectophrynoides_viviparus | Nectophrynoides_viviparus | 0 | 19.92 | 137 | 2397.71 |
| amphibians | Necturus_beyeri | Necturus_beyeri | 0 | 94.15 | 40 | 303770.78 |
| amphibians | Necturus_lewisi | Necturus_lewisi | 0 | 41.28 | 27.5 | 21188.52 |
| amphibians | Necturus_maculosus | Necturus_maculosus | 0 | 533.85 | 94.5 | 1636355.17 |
| amphibians | Necturus_punctatus | Necturus_punctatus | 0 | 31 | 35 | 177790.95 |
| amphibians | Neobatrachus_albipes | Neobatrachus_albipes | 0 | 11.04 | 966 | 282547.96 |
| amphibians | Neobatrachus_aquilonius | Neobatrachus_aquilonius | 0 | 19.92 | 1426 | 288535.42 |
| amphibians | Neobatrachus_kunapalari | Neobatrachus_kunapalari | 0 | 22.39 | 1303 | 412663.84 |
| amphibians | Neobatrachus_pelobatoides | Neobatrachus_pelobatoides | 0 | 11.04 | 681.5 | 361136.88 |
| amphibians | Neobatrachus_pictus | Neobatrachus_pictus | 0 | 23.34 | 600 | 434433.25 |
| amphibians | Neobatrachus_sudelli | Neobatrachus_sudelli | 0 | 15.03 | 3161.5 | 3558999.63 |
| amphibians | Neobatrachus_sutor | Neobatrachus_sutor | 0 | 11.77 | 615 | 1478854.25 |
| amphibians | Neobatrachus_wilsmorei | Neobatrachus_wilsmorei | 0 | 23.34 | 2000 | 275721.79 |
| amphibians | Nesorohyla_kanaima | Myersiohyla_kanaima | 1 | 10.41 | 38 | 3318.43 |
| amphibians | Neurergus_crocatus | Neurergus_crocatus | 1 | 8.62 | 250 | 11081.37 |
| amphibians | Neurergus_kaiseri | Neurergus_kaiseri | 1 | 8.56 | 160 | 9198.41 |
| amphibians | Neurergus_strauchii | Neurergus_strauchii | 0 | 8.02 | 177.5 | 20397.41 |
| amphibians | Niceforonia_nigrovittata | Hypodactylus_nigrovittatus | 0 | 2.35 | 9.5 | 939846.04 |
| amphibians | Nidirana_adenopleura | Babina_adenopleura | 0 | 25.82 | 185 | 538862.92 |
| amphibians | Nidirana_okinavana | Babina_okinavana | 1 | 10.34 | 499.5 | 512.17 |
| amphibians | Nidirana_pleuraden | Babina_pleuraden | 0 | 16.87 | 606 | 357284.95 |
| amphibians | Nimbaphrynoides_occidentalis | Nimbaphrynoides_occidentalis | 1 | 2.87 | 19.5 | 6.46 |
| amphibians | Notaden_bennettii | Notaden_bennettii | 0 | 29.89 | 1863.5 | 648800.84 |
| amphibians | Notaden_melanoscaphus | Notaden_melanoscaphus | 0 | 15.03 | 850 | 759801.96 |
| amphibians | Notaden_nichollsi | Notaden_nichollsi | 0 | 28.49 | 1000 | 2358108.74 |
| amphibians | Nothophryne_broadleyi | Nothophryne_broadleyi | 1 | 1.69 | 30 | 383.86 |

| | | | | | | |
|---|---|---|---|---|---|---|
| amphibians | Notophthalmus_meridionalis | Notophthalmus_meridionalis | 1 | 11.88 | 300 | 108071.61 |
| amphibians | Notophthalmus_viridescens | Notophthalmus_viridescens | 0 | 2.56 | 265 | 3440358.26 |
| amphibians | Nototriton_abscondens | Nototriton_abscondens | 0 | 0.56 | 17 | 902.95 |
| amphibians | Nototriton_barbouri | Nototriton_barbouri | 1 | 1.05 | 12 | 36 |
| amphibians | Nototriton_guanacaste | Nototriton_guanacaste | 0 | 0.66 | 6 | 35.79 |
| amphibians | Nyctibatrachus_aliciae | Nyctibatrachus_aliciae | 1 | 2.6 | 17 | 5553.37 |
| amphibians | Nyctibatrachus_danieli | Nyctibatrachus_danieli | 1 | 3.51 | 40 | 2548.56 |
| amphibians | Nyctibatrachus_humayuni | Nyctibatrachus_humayuni | 0 | 12.53 | 33 | 23268.76 |
| amphibians | Nyctibatrachus_jog | Nyctibatrachus_jog | 1 | 4.93 | 35.5 | 10910.5 |
| amphibians | Nyctibatrachus_karnatakaensis | Nyctibatrachus_karnatakaensis | 1 | 61.58 | 25 | 852.75 |
| amphibians | Nyctibatrachus_kempholeyensis | Nyctibatrachus_kempholeyensis | 0 | 1.08 | 10 | 35495.47 |
| amphibians | Nyctibatrachus_major | Nyctibatrachus_major | 1 | 14.33 | 17 | 2635.67 |
| amphibians | Nyctibatrachus_petraeus | Nyctibatrachus_petraeus | 0 | 9.41 | 24 | 6905.85 |
| amphibians | Nyctimantis_arapapa | Aparasphenodon_arapapa | 1 | 19.92 | 34.5 | 4273.65 |
| amphibians | Nyctimantis_rugiceps | Nyctimantis_rugiceps | 0 | 29.89 | 519 | 45141.11 |
| amphibians | Nyctimantis_siemersi | Argenteohyla_siemersi | 0 | 57.01 | 5750 | 91004.38 |
| amphibians | Nyctimystes_brevipalmatus | Litoria_brevipalmata | 0 | 9.03 | 484 | 75777.4 |
| amphibians | Nyctimystes_infrafrenatus | Litoria_infrafrenata | 0 | 275.7 | 2193.5 | 774851.39 |
| amphibians | Nyctixalus_margaritifer | Nyctixalus_margaritifer | 0 | 3.47 | 14 | 17371.87 |
| amphibians | Nyctixalus_pictus | Nyctixalus_pictus | 0 | 4.54 | 11 | 231431.81 |
| amphibians | Nyctixalus_spinosus | Nyctixalus_spinosus | 0 | 4.16 | 35 | 12654.06 |
| amphibians | Nymphargus_buenaventura | Nymphargus_buenaventura | 1 | 0.96 | 38 | 1883.01 |
| amphibians | Nymphargus_grandisonae | Nymphargus_grandisonae | 0 | 2.77 | 50.5 | 21229.61 |
| amphibians | Nymphargus_griffithsi | Nymphargus_griffithsi | 0 | 2.52 | 29.5 | 45955.22 |
| amphibians | Nymphargus_ignotus | Nymphargus_ignotus | 0 | 1.23 | 19 | 3565.85 |
| amphibians | Nymphargus_lasgralarias | Nymphargus_lasgralarias | 1 | 1.52 | 24 | 760.77 |
| amphibians | Nymphargus_rosada | Nymphargus_rosada | 1 | 1.79 | 33.5 | 994.49 |
| amphibians | Nymphargus_wileyi | Nymphargus_wileyi | 1 | 1.52 | 23.5 | 10.86 |
| amphibians | Occidozyga_laevis | Occidozyga_laevis | 0 | 22.51 | 123.5 | 356736.47 |
| amphibians | Occidozyga_lima | Occidozyga_lima | 0 | 11.04 | 379 | 2814348.17 |
| amphibians | Occidozyga_martensii | Occidozyga_martensii | 0 | 2.87 | 250 | 1761564.64 |
| amphibians | Occidozyga_sumatrana | Occidozyga_sumatrana | 0 | 11.04 | 24 | 137030.46 |
| amphibians | Odontophrynus_americanus | Odontophrynus_americanus | 0 | 32.83 | 6500 | 2881157.37 |
| amphibians | Odontophrynus_lavillai | Odontophrynus_lavillai | 0 | 36.46 | 9208 | 808950.55 |

| | | | | | | |
|---|---|---|---|---|---|---|
| amphibians | Odontophrynus_maisuma | Odontophrynus_maisuma | 0 | 7.08 | 2980 | 33564.85 |
| amphibians | Odorrana_amamiensis | Odorrana_amamiensis | 1 | 107.68 | 1500 | 956.79 |
| amphibians | Odorrana_andersonii | Odorrana_andersonii | 0 | 129.82 | 990 | 633784.85 |
| amphibians | Odorrana_banaorum | Odorrana_banaorum | 0 | 136.63 | 2966 | 121051 |
| amphibians | Odorrana_grahami | Odorrana_grahami | 1 | 129.82 | 2446 | 306670.97 |
| amphibians | Odorrana_hosii | Odorrana_hosii | 0 | 104.27 | 1410 | 1116465.62 |
| amphibians | Odorrana_ishikawae | Odorrana_ishikawae | 1 | 173.41 | 1000 | 311.63 |
| amphibians | Odorrana_jingdongensis | Odorrana_jingdongensis | 1 | 133.8 | 367.5 | 111826.01 |
| amphibians | Odorrana_livida | Odorrana_livida | 0 | 105.56 | 2162 | 41226.89 |
| amphibians | Odorrana_margaretae | Odorrana_margaretae | 0 | 154.93 | 705 | 614109.52 |
| amphibians | Odorrana_narina | Odorrana_narina | 1 | 100.93 | 200 | 311.63 |
| amphibians | Odorrana_schmackeri | Odorrana_schmackeri | 0 | 50.6 | 992.5 | 1286455.01 |
| amphibians | Odorrana_supranarina | Odorrana_supranarina | 1 | 114.75 | 1100 | 512.17 |
| amphibians | Odorrana_swinhoana | Odorrana_swinhoana | 0 | 60.65 | 45 | 17356.98 |
| amphibians | Odorrana_tormota | Odorrana_tormota | 0 | 19.92 | 589.5 | 28319.16 |
| amphibians | Odorrana_utsunomiyaorum | Odorrana_utsunomiyaorum | 1 | 17.85 | 100 | 512.17 |
| amphibians | Odorrana_versabilis | Odorrana_versabilis | 0 | 22.51 | 350.5 | 188567.38 |
| amphibians | Odorrana_yentuensis | Odorrana_yentuensis | 1 | 23.46 | 350 | 2007.16 |
| amphibians | Ololygon_albicans | Scinax_albicans | 0 | 8.99 | 631.5 | 2601.54 |
| amphibians | Ololygon_angrensis | Scinax_angrensis | 0 | 5.02 | 556 | 2483.06 |
| amphibians | Ololygon_argyreornata | Scinax_argyreornatus | 0 | 1.29 | 121 | 266745.59 |
| amphibians | Ololygon_aromothyella | Scinax_aromothyella | 0 | 1.5 | 500 | 377908.12 |
| amphibians | Ololygon_catharinae | Scinax_catharinae | 0 | 9.03 | 450 | 1123002.09 |
| amphibians | Ololygon_centralis | Scinax_centralis | 0 | 1.32 | 64 | 312402.9 |
| amphibians | Ololygon_hiemalis | Scinax_hiemalis | 0 | 3.47 | 270 | 44065.08 |
| amphibians | Ololygon_littoralis | Scinax_littoralis | 0 | 5.31 | 549.5 | 22250.25 |
| amphibians | Ololygon_littorea | Scinax_littoreus | 1 | 1.25 | 9 | 1771.68 |
| amphibians | Ololygon_perpusilla | Scinax_perpusillus | 0 | 1.17 | 9 | 15916.16 |
| amphibians | Ololygon_rizibilis | Scinax_rizibilis | 0 | 3.81 | 1200 | 202433.45 |
| amphibians | Ololygon_tripui | Scinax_tripui | 0 | 5.02 | 332 | 6323.52 |
| amphibians | Ololygon_v-signata | Scinax_v-signatus | 0 | 1.5 | 9 | 5662.04 |
| amphibians | Ombrana_sikimensis | Ombrana_sikimensis | 0 | 68.91 | 233 | 75449.88 |
| amphibians | Ommatotriton_ophryticus | Ommatotriton_ophryticus | 0 | 10.77 | 124.5 | 118354.75 |
| amphibians | Ommatotriton_vittatus | Ommatotriton_vittatus | 0 | 3.54 | 43 | 57209.04 |

| | | | | | | |
|---|---|---|---|---|---|---|
| amphibians | Onychodactylus_fischeri | Onychodactylus_fischeri | 0 | 9.23 | 46 | 31338.51 |
| amphibians | Onychodactylus_japonicus | Onychodactylus_japonicus | 0 | 11.5 | 21.5 | 78734.7 |
| amphibians | Onychodactylus_koreanus | Onychodactylus_koreanus | 0 | 7.72 | 15 | 69906.07 |
| amphibians | Onychodactylus_tsukubaensis | Onychodactylus_tsukubaensis | 1 | 5.38 | 16.5 | 38.25 |
| amphibians | Onychodactylus_zhaoermii | Onychodactylus_zhaoermii | 0 | 8.5 | 13.5 | 21631.12 |
| amphibians | Oophaga_granulifera | Oophaga_granulifera | 1 | 1.17 | 3 | 6349.83 |
| amphibians | Oophaga_pumilio | Oophaga_pumilio | 0 | 1.17 | 10 | 61975.53 |
| amphibians | Oophaga_sylvatica | Oophaga_sylvatica | 0 | 4.54 | 5 | 48578.3 |
| amphibians | Oophaga_vicentei | Oophaga_vicentei | 1 | 0.66 | 4 | 2942.17 |
| amphibians | Ophryophryne_microstoma | Ophryophryne_microstoma | 0 | 10 | 436 | 451283.37 |
| amphibians | Ophryophryne_synoria | Ophryophryne_synoria | 1 | 34.37 | 800.5 | 7557.54 |
| amphibians | Opisthothylax_immaculatus | Opisthothylax_immaculatus | 0 | 3.47 | 8 | 816120.04 |
| amphibians | Oreobates_cruralis | Oreobates_cruralis | 0 | 3.04 | 13 | 146293.04 |
| amphibians | Oreobates_quixensis | Oreobates_quixensis | 0 | 23.34 | 33 | 1916308.79 |
| amphibians | Oreolalax_chuanbeiensis | Oreolalax_chuanbeiensis | 1 | 18.87 | 168 | 3829.55 |
| amphibians | Oreolalax_granulosus | Oreolalax_granulosus | 0 | 21.02 | 105 | 16633.88 |
| amphibians | Oreolalax_jingdongensis | Oreolalax_jingdongensis | 1 | 20.25 | 163 | 16481.87 |
| amphibians | Oreolalax_liangbeiensis | Oreolalax_liangbeiensis | 1 | 26.73 | 255 | 6.75 |
| amphibians | Oreolalax_lichuanensis | Oreolalax_lichuanensis | 0 | 25.57 | 215 | 94754.56 |
| amphibians | Oreolalax_multipunctatus | Oreolalax_multipunctatus | 1 | 10.89 | 77.5 | 4143.1 |
| amphibians | Oreolalax_omeimontis | Oreolalax_omeimontis | 1 | 18.25 | 183.5 | 2901.35 |
| amphibians | Oreolalax_pingii | Oreolalax_pingii | 1 | 14.5 | 293 | 2325.75 |
| amphibians | Oreolalax_popei | Oreolalax_popei | 0 | 31.33 | 393 | 34208.9 |
| amphibians | Oreolalax_puxiongensis | Oreolalax_puxiongensis | 1 | 11.04 | 126.5 | 1248.61 |
| amphibians | Oreolalax_rugosus | Oreolalax_rugosus | 0 | 15.3 | 436 | 40739.72 |
| amphibians | Oreolalax_schmidti | Oreolalax_schmidti | 0 | 14.16 | 119 | 25470.73 |
| amphibians | Oreophryne_anthonyi | Oreophryne_anthonyi | 0 | 9.03 | 24.5 | 2504.16 |
| amphibians | Oreophryne_idenburgensis | Oreophryne_idenburgensis | 0 | 9.03 | 15 | 9.88 |
| amphibians | Oreophryne_oviprotector | Oreophryne_oviprotector | 0 | 1.52 | 8 | 53939.18 |
| amphibians | Oreophrynella_quelchii | Oreophrynella_quelchii | 1 | 2.11 | 12 | 40.52 |
| amphibians | Oscaecilia_ochrocephala | Oscaecilia_ochrocephala | 0 | 90.59 | 9.5 | 34953.85 |
| amphibians | Osornophryne_cofanorum | Osornophryne_cofanorum | 1 | 3.29 | 7 | 46.84 |
| amphibians | Osornophryne_guacamayo | Osornophryne_guacamayo | 1 | 5.64 | 42.5 | 1891.79 |
| amphibians | Osornophryne_puruanta | Osornophryne_puruanta | 1 | 9.09 | 30.5 | 40.44 |

| | | | | | | |
|---|---|---|---|---|---|---|
| amphibians | Osteocephalus_alboguttatus | Osteocephalus_alboguttatus | 0 | 25.82 | 544 | 17609.51 |
| amphibians | Osteocephalus_buckleyi | Osteocephalus_buckleyi | 0 | 31.33 | 1090 | 816911.23 |
| amphibians | Osteocephalus_cabrerai | Osteocephalus_cabrerai | 0 | 35.01 | 989.5 | 2206491.4 |
| amphibians | Osteocephalus_festae | Osteocephalus_festae | 0 | 61.35 | 1234 | 26630.89 |
| amphibians | Osteocephalus_leprieurii | Osteocephalus_leprieurii | 0 | 41.05 | 1681 | 4980892.49 |
| amphibians | Osteocephalus_mutabor | Osteocephalus_mutabor | 0 | 42.31 | 35 | 58502.97 |
| amphibians | Osteocephalus_oophagus | Osteocephalus_oophagus | 0 | 22.98 | 395 | 1269609.69 |
| amphibians | Osteocephalus_taurinus | Osteocephalus_taurinus | 0 | 118.03 | 1275 | 7474189.45 |
| amphibians | Osteopilus_crucialis | Osteopilus_crucialis | 1 | 198.59 | 332 | 3741.94 |
| amphibians | Osteopilus_dominicensis | Osteopilus_dominicensis | 0 | 100.93 | 436 | 74765.19 |
| amphibians | Osteopilus_marianae | Osteopilus_marianae | 1 | 5.36 | 332 | 1934.52 |
| amphibians | Osteopilus_ocellatus | Osteopilus_ocellatus | 0 | 42.85 | 278 | 8550.17 |
| amphibians | Osteopilus_septentrionalis | Osteopilus_septentrionalis | 0 | 528.2 | 2045.5 | 279117.6 |
| amphibians | Osteopilus_vastus | Osteopilus_vastus | 1 | 324.76 | 332 | 23538.76 |
| amphibians | Otophryne_steyermarki | Otophryne_steyermarki | 0 | 9.67 | 8 | 23873.37 |
| amphibians | Pachytriton_brevipes | Pachytriton_brevipes | 0 | 15.56 | 45 | 78142.06 |
| amphibians | Pachytriton_feii | Pachytriton_feii | 1 | 8.62 | 30 | 16412.31 |
| amphibians | Pachytriton_granulosus | Pachytriton_granulosus | 0 | 8.62 | 40 | 114712.76 |
| amphibians | Paracrinia_haswelli | Paracrinia_haswelli | 0 | 3.47 | 504 | 74554.22 |
| amphibians | Paradactylodon_mustersi | Paradactylodon_mustersi | 1 | 6.68 | 19 | 1119.72 |
| amphibians | Paradoxophyla_palmata | Paradoxophyla_palmata | 0 | 1.33 | 300 | 33457.19 |
| amphibians | Paramesotriton_caudopunctatus | Paramesotriton_caudopunctatus | 0 | 5.18 | 67.5 | 28929.92 |
| amphibians | Paramesotriton_chinensis | Paramesotriton_chinensis | 0 | 10.41 | 199.5 | 107136.06 |
| amphibians | Paramesotriton_hongkongensis | Paramesotriton_hongkongensis | 0 | 6.02 | 117.5 | 6650.47 |
| amphibians | Paramesotriton_labiatus | Paramesotriton_labiatus | 1 | 18.87 | 40 | 6.87 |
| amphibians | Parapelophryne_scalpta | Parapelophryne_scalpta | 1 | 1.52 | 39 | 8696.38 |
| amphibians | Paratelmatobius_cardosoi | Paratelmatobius_cardosoi | 0 | 0.41 | 20 | 2229.42 |
| amphibians | Paratelmatobius_gaigeae | Paratelmatobius_gaigeae | 1 | 0.57 | 6 | 21.55 |
| amphibians | Paratelmatobius_lutzii | Paratelmatobius_lutzii | 1 | 1.02 | 27 | 10.75 |
| amphibians | Paratelmatobius_mantiqueira | Paratelmatobius_mantiqueira | 1 | 0.51 | 40 | 324.64 |
| amphibians | Paratelmatobius_poecilogaster | Paratelmatobius_poecilogaster | 0 | 2.13 | 9 | 5433.5 |
| amphibians | Paruwrobates_andinus | Ameerega_andina | 1 | 0.77 | 3.5 | 41.78 |
| amphibians | Paruwrobates_erythromos | Ameerega_erythromos | 1 | 1.2 | 3 | 18.7 |
| amphibians | Pelobates_cultripes | Pelobates_cultripes | 1 | 214.85 | 4750 | 493091.33 |

| | | | | | | |
|---|---|---|---|---|---|---|
| amphibians | Pelobates_fuscus | Pelobates_fuscus | 0 | 50.6 | 1772.5 | 2386089.75 |
| amphibians | Pelobates_syriacus | Pelobates_syriacus | 0 | 74.11 | 4000 | 372668.29 |
| amphibians | Pelobates_varaldii | Pelobates_varaldii | 1 | 32.83 | 1500 | 6672.95 |
| amphibians | Pelobatrachus_nasutus | Megophrys_nasuta | 0 | 478.08 | 1753.5 | 988325.78 |
| amphibians | Pelodytes_caucasicus | Pelodytes_caucasicus | 0 | 15.03 | 415.5 | 111848.59 |
| amphibians | Pelodytes_ibericus | Pelodytes_ibericus | 0 | 9.67 | 509 | 102533.41 |
| amphibians | Pelodytes_punctatus | Pelodytes_punctatus | 0 | 11.77 | 820 | 707169.23 |
| amphibians | Pelophryne_brevipes | Pelophryne_brevipes | 0 | 0.61 | 15.5 | 41109.31 |
| amphibians | Pelophryne_misera | Pelophryne_misera | 0 | 0.89 | 10.5 | 2654.7 |
| amphibians | Pelophylax_bedriagae | Pelophylax_bedriagae | 0 | 104.27 | 6172 | 1027992.46 |
| amphibians | Pelophylax_cerigensis | Pelophylax_cerigensis | 1 | 32.83 | 15000 | 4.05 |
| amphibians | Pelophylax_cretensis | Pelophylax_cretensis | 1 | 50.6 | 15000 | 2535.9 |
| amphibians | Pelophylax_epeiroticus | Pelophylax_epeiroticus | 0 | 100.93 | 2737.5 | 12315.98 |
| amphibians | Pelophylax_fukienensis | Pelophylax_fukienensis | 0 | 26.34 | 1011 | 423666.2 |
| amphibians | Pelophylax_hubeiensis | Pelophylax_hubeiensis | 0 | 11.04 | 1425.5 | 260899.53 |
| amphibians | Pelophylax_kurtmuelleri | Pelophylax_kurtmuelleri | 0 | 104.27 | 10000 | 140722.78 |
| amphibians | Pelophylax_lessonae | Pelophylax_lessonae | 0 | 188.23 | 3210 | 3431615.93 |
| amphibians | Pelophylax_nigromaculatus | Pelophylax_nigromaculatus | 0 | 85.33 | 2800 | 4306604.69 |
| amphibians | Pelophylax_perezi | Pelophylax_perezi | 0 | 141.99 | 5400 | 650025.7 |
| amphibians | Pelophylax_plancyi | Pelophylax_plancyi | 0 | 20.49 | 929 | 814241.55 |
| amphibians | Pelophylax_porosus | Pelophylax_porosus | 0 | 66.4 | 1900 | 96735.43 |
| amphibians | Pelophylax_ridibundus | Pelophylax_ridibundus | 0 | 700.22 | 8335 | 7792587.74 |
| amphibians | Pelophylax_saharicus | Pelophylax_saharicus | 0 | 310.17 | 5650 | 1403222.43 |
| amphibians | Pelophylax_shqipericus | Pelophylax_shqipericus | 1 | 41.05 | 2594.5 | 5620.94 |
| amphibians | Peltophryne_lemur | Peltophryne_lemur | 1 | 188.23 | 14983.9 | 407.91 |
| amphibians | Petropedetes_perreti | Petropedetes_perreti | 1 | 10.34 | 82 | 550.32 |
| amphibians | Petropedetes_vulpiae | Petropedetes_vulpiae | 0 | 9.03 | 10 | 190194.3 |
| amphibians | Phaeognathus_hubrichti | Phaeognathus_hubrichti | 1 | 20.25 | 26 | 1166.99 |
| amphibians | Phasmahyla_cochranae | Phasmahyla_cochranae | 0 | 8.42 | 261.5 | 63311.81 |
| amphibians | Phasmahyla_cruzi | Phasmahyla_cruzi | 0 | 6.08 | 53 | 9656.59 |
| amphibians | Phasmahyla_guttata | Phasmahyla_guttata | 0 | 7.84 | 40 | 9871.99 |
| amphibians | Phasmahyla_jandaia | Phasmahyla_jandaia | 0 | 3.81 | 24 | 18654.37 |
| amphibians | Phasmahyla_spectabilis | Phasmahyla_spectabilis | 0 | 10.07 | 40 | 2688.49 |
| amphibians | Philautus_amoenus | Philautus_amoenus | 0 | 3.38 | 15 | 290.49 |

| | | | | | | |
|---|---|---|---|---|---|---|
| amphibians | Philautus_aurifasciatus | Philautus_aurifasciatus | 0 | 2.87 | 13 | 164758.33 |
| amphibians | Philautus_bunitus | Philautus_bunitus | 0 | 8.6 | 17 | 4837.7 |
| amphibians | Philautus_hosii | Philautus_hosii | 0 | 23.34 | 36 | 60897.22 |
| amphibians | Philautus_kerangae | Philautus_kerangae | 1 | 6.77 | 7 | 583.03 |
| amphibians | Philautus_mjobergi | Philautus_mjobergi | 0 | 2.6 | 7 | 15067.32 |
| amphibians | Philautus_saueri | Philautus_saueri | 0 | 0.71 | 13 | 22.28 |
| amphibians | Philautus_schmackeri | Philautus_schmackeri | 1 | 2.18 | 10.5 | 1308.13 |
| amphibians | Philautus_surdus | Philautus_surdus | 0 | 4.69 | 12 | 211208.8 |
| amphibians | Philoria_frosti | Philoria_frosti | 1 | 15.03 | 117.5 | 113.32 |
| amphibians | Philoria_kundagungan | Philoria_kundagungan | 1 | 2.11 | 41.5 | 607.11 |
| amphibians | Philoria_loveridgei | Philoria_loveridgei | 1 | 2.6 | 28.5 | 682.21 |
| amphibians | Philoria_pughi | Philoria_pughi | 1 | 2.11 | 41 | 2017.5 |
| amphibians | Philoria_richmondensis | Philoria_richmondensis | 1 | 1.69 | 43 | 669.57 |
| amphibians | Philoria_sphagnicola | Philoria_sphagnicolus | 1 | 4.16 | 60.5 | 10171.24 |
| amphibians | Phrynobatrachus_acridoides | Phrynobatrachus_acridoides | 0 | 2.35 | 105 | 1759744.51 |
| amphibians | Phrynobatrachus_alleni | Phrynobatrachus_alleni | 0 | 1.69 | 400 | 389259.35 |
| amphibians | Phrynobatrachus_auritus | Phrynobatrachus_auritus | 0 | 5.36 | 200 | 1793524.13 |
| amphibians | Phrynobatrachus_calcaratus | Phrynobatrachus_calcaratus | 0 | 1.17 | 187.5 | 1531152.42 |
| amphibians | Phrynobatrachus_dendrobates | Phrynobatrachus_dendrobates | 0 | 4.16 | 23.5 | 112117.23 |
| amphibians | Phrynobatrachus_francisci | Phrynobatrachus_francisci | 0 | 0.77 | 1750 | 1825263.55 |
| amphibians | Phrynobatrachus_guineensis | Phrynobatrachus_guineensis | 0 | 0.77 | 21 | 187114.79 |
| amphibians | Phrynobatrachus_gutturosus | Phrynobatrachus_gutturosus | 0 | 1.02 | 350 | 733802.55 |
| amphibians | Phrynobatrachus_krefftii | Phrynobatrachus_krefftii | 1 | 11.77 | 22.5 | 2119.88 |
| amphibians | Phrynobatrachus_latifrons | Phrynobatrachus_latifrons | 0 | 2.11 | 800 | 2626407.09 |
| amphibians | Phrynobatrachus_mababiensis | Phrynobatrachus_mababiensis | 0 | 0.77 | 775 | 4355476.06 |
| amphibians | Phrynobatrachus_natalensis | Phrynobatrachus_natalensis | 0 | 5.36 | 901 | 10523648.24 |
| amphibians | Phrynobatrachus_phyllophilus | Phrynobatrachus_phyllophilus | 0 | 0.89 | 32.5 | 258033.29 |
| amphibians | Phrynobatrachus_plicatus | Phrynobatrachus_plicatus | 0 | 6.27 | 1550 | 285268.63 |
| amphibians | Phrynobatrachus_sandersoni | Phrynobatrachus_sandersoni | 0 | 1.17 | 14.5 | 64991.23 |
| amphibians | Phrynobatrachus_tokba | Phrynobatrachus_tokba | 0 | 0.61 | 13.5 | 187123.53 |
| amphibians | Phrynoidis_asper | Phrynoidis_aspera | 0 | 310.17 | 12750 | 1614710.7 |
| amphibians | Phrynomantis_annectens | Phrynomantis_annectens | 0 | 5.36 | 90 | 528204.27 |
| amphibians | Phrynomantis_bifasciatus | Phrynomantis_bifasciatus | 0 | 50.6 | 900 | 4714507.51 |
| amphibians | Phrynomantis_microps | Phrynomantis_microps | 0 | 23.34 | 765.5 | 3023303.7 |

| | | | | | | |
|---|---|---|---|---|---|---|
| amphibians | Phrynomedusa_appendiculata | Phrynomedusa_appendiculata | 1 | 4.31 | 50 | 1061.42 |
| amphibians | Phrynopus_kauneorum | Phrynopus_kauneorum | 1 | 16.3 | 56 | 306.91 |
| amphibians | Phyllobates_aurotaenia | Phyllobates_aurotaenia | 0 | 3.16 | 23.5 | 21407.51 |
| amphibians | Phyllobates_bicolor | Phyllobates_bicolor | 1 | 6.77 | 20 | 2879.3 |
| amphibians | Phyllobates_lugubris | Phyllobates_lugubris | 0 | 1.02 | 17.5 | 19092 |
| amphibians | Phyllobates_terribilis | Phyllobates_terribilis | 1 | 9.03 | 14 | 1288.44 |
| amphibians | Phyllobates_vittatus | Phyllobates_vittatus | 1 | 2.35 | 14 | 4258.39 |
| amphibians | Phyllodytes_luteolus | Phyllodytes_luteolus | 0 | 1.02 | 13 | 93868.56 |
| amphibians | Phyllomedusa_bicolor | Phyllomedusa_bicolor | 0 | 275.7 | 1011 | 5032516.9 |
| amphibians | Phyllomedusa_burmeisteri | Phyllomedusa_burmeisteri | 0 | 48.58 | 315.5 | 555418.7 |
| amphibians | Phyllomedusa_camba | Phyllomedusa_camba | 0 | 59.27 | 312.5 | 1129163.05 |
| amphibians | Phyllomedusa_distincta | Phyllomedusa_distincta | 0 | 34.37 | 215 | 120035.96 |
| amphibians | Phyllomedusa_iheringii | Phyllomedusa_iheringii | 0 | 41.05 | 160 | 157479.79 |
| amphibians | Phyllomedusa_sauvagii | Phyllomedusa_sauvagii | 0 | 93.21 | 466 | 1025952.82 |
| amphibians | Phyllomedusa_tarsius | Phyllomedusa_tarsius | 0 | 150.53 | 418 | 2240122.38 |
| amphibians | Phyllomedusa_tetraploidea | Phyllomedusa_tetraploidea | 0 | 31.93 | 231 | 292985.4 |
| amphibians | Phyllomedusa_trinitatis | Phyllomedusa_trinitatis | 0 | 89.82 | 40 | 56575.58 |
| amphibians | Phyllomedusa_vaillantii | Phyllomedusa_vaillantii | 0 | 59.27 | 415.5 | 5960632.72 |
| amphibians | Physalaemus_aguirrei | Physalaemus_aguirrei | 0 | 1.73 | 474 | 82434.95 |
| amphibians | Physalaemus_albonotatus | Physalaemus_albonotatus | 0 | 3.19 | 785.5 | 1492280.79 |
| amphibians | Physalaemus_atlanticus | Physalaemus_atlanticus | 0 | 1.46 | 90.5 | 2359.02 |
| amphibians | Physalaemus_barrioi | Physalaemus_barrioi | 0 | 1.89 | 193 | 71.25 |
| amphibians | Physalaemus_biligonigerus | Physalaemus_biligonigerus | 0 | 5.36 | 4953 | 2026421.1 |
| amphibians | Physalaemus_bokermanni | Physalaemus_bokermanni | 0 | 0.52 | 55 | 1569.17 |
| amphibians | Physalaemus_centralis | Physalaemus_centralis | 0 | 5.36 | 1903 | 2405327.65 |
| amphibians | Physalaemus_crombiei | Physalaemus_crombiei | 0 | 1.2 | 170.5 | 122259.19 |
| amphibians | Physalaemus_cuqui | Physalaemus_cuqui | 0 | 2.11 | 1033 | 102136.3 |
| amphibians | Physalaemus_cuvieri | Physalaemus_cuvieri | 0 | 3.98 | 1107 | 7110840.08 |
| amphibians | Physalaemus_ephippifer | Physalaemus_ephippifer | 0 | 2.87 | 414 | 1445710.77 |
| amphibians | Physalaemus_evangelistai | Physalaemus_evangelistai | 0 | 0.89 | 133 | 15052.15 |
| amphibians | Physalaemus_fernandezae | Physalaemus_fernandezae | 0 | 1.33 | 225 | 124620.94 |
| amphibians | Physalaemus_gracilis | Physalaemus_gracilis | 0 | 5.36 | 450 | 505595.37 |
| amphibians | Physalaemus_henselii | Physalaemus_henselii | 0 | 1.02 | 275 | 407683.17 |
| amphibians | Physalaemus_kroyeri | Physalaemus_kroyeri | 0 | 3.47 | 1458.5 | 492507.06 |

| | | | | | | |
|---|---|---|---|---|---|---|
| amphibians | Physalaemus_lisei | Physalaemus_lisei | 0 | 2.87 | 500 | 33751.74 |
| amphibians | Physalaemus_maculiventris | Physalaemus_maculiventris | 0 | 1.17 | 129 | 85909.93 |
| amphibians | Physalaemus_marmoratus | Physalaemus_marmoratus | 0 | 4.93 | 2007 | 1349605.19 |
| amphibians | Physalaemus_nanus | Physalaemus_nanus | 0 | 0.77 | 75 | 165830.7 |
| amphibians | Physalaemus_nattereri | Eupemphix_nattereri | 0 | 16.3 | 3032.5 | 1921704.34 |
| amphibians | Physalaemus_olfersii | Physalaemus_olfersii | 0 | 5.8 | 700.5 | 124940.92 |
| amphibians | Physalaemus_riograndensis | Physalaemus_riograndensis | 0 | 1.17 | 225 | 470374.73 |
| amphibians | Physalaemus_rupestris | Physalaemus_rupestris | 1 | 0.39 | 34 | 66.62 |
| amphibians | Physalaemus_santafecinus | Physalaemus_santafecinus | 0 | 3.16 | 500 | 87322.56 |
| amphibians | Physalaemus_signifer | Physalaemus_signifer | 0 | 1.57 | 273 | 873.48 |
| amphibians | Physalaemus_spiniger | Physalaemus_spiniger | 0 | 0.85 | 85.5 | 15997.53 |
| amphibians | Pipa_arrabali | Pipa_arrabali | 0 | 16.87 | 15 | 2129877.41 |
| amphibians | Pipa_aspera | Pipa_aspera | 0 | 16.87 | 19 | 75201.04 |
| amphibians | Pipa_carvalhoi | Pipa_carvalhoi | 0 | 29.89 | 220 | 238909.69 |
| amphibians | Pipa_myersi | Pipa_myersi | 1 | 7.4 | 12 | 168.2 |
| amphibians | Pipa_parva | Pipa_parva | 0 | 7.29 | 186 | 83384.84 |
| amphibians | Pipa_pipa | Pipa_pipa | 0 | 985.12 | 95 | 7019956.39 |
| amphibians | Pipa_snethlageae | Pipa_snethlageae | 0 | 79.58 | 12 | 1568740.87 |
| amphibians | Pithecopus_ayeaye | Phyllomedusa_ayeaye | 0 | 8.48 | 25.5 | 87959.35 |
| amphibians | Pithecopus_azureus | Phyllomedusa_azurea | 0 | 7.51 | 96 | 1220516.48 |
| amphibians | Pithecopus_hypochondrialis | Phyllomedusa_hypochondrialis | 0 | 11.77 | 162.5 | 4002445.6 |
| amphibians | Pithecopus_megacephalus | Phyllomedusa_megacephala | 0 | 20.14 | 28.5 | 43611.34 |
| amphibians | Pithecopus_oreades | Phyllomedusa_oreades | 0 | 10.82 | 29 | 108460.6 |
| amphibians | Pithecopus_palliatus | Phyllomedusa_palliata | 0 | 22.51 | 112 | 1120855.58 |
| amphibians | Platymantis_dorsalis | Platymantis_dorsalis | 0 | 7.84 | 33.5 | 62585.8 |
| amphibians | Platymantis_guentheri | Platymantis_guentheri | 0 | 4.27 | 8 | 16594.98 |
| amphibians | Platymantis_hazelae | Platymantis_hazelae | 1 | 3.88 | 7 | 2345.98 |
| amphibians | Platymantis_negrosensis | Platymantis_negrosensis | 0 | 11.18 | 27.5 | 2035.7 |
| amphibians | Platymantis_panayensis | Platymantis_panayensis | 1 | 2.87 | 7.5 | 1778.32 |
| amphibians | Platypelis_grandis | Platypelis_grandis | 0 | 122.13 | 95 | 129098.93 |
| amphibians | Platyplectrum_fletcheri | Lechriodus_fletcheri | 0 | 14.16 | 471.5 | 63071.21 |
| amphibians | Platyplectrum_ornatum | Platyplectrum_ornatum | 0 | 7.84 | 874 | 2936283.59 |
| amphibians | Platyplectrum_spenceri | Platyplectrum_spenceri | 0 | 11.04 | 1128.5 | 3182540.99 |
| amphibians | Plectrohyla_glandulosa | Plectrohyla_glandulosa | 1 | 11.04 | 265 | 778.44 |

| | | | | | | |
|---|---|---|---|---|---|---|
| amphibians | Plectrohyla_guatemalensis | Plectrohyla_guatemalensis | 0 | 43.04 | 196 | 36439.68 |
| amphibians | Plectrohyla_hartwegi | Plectrohyla_hartwegi | 1 | 44.71 | 271.5 | 8321.27 |
| amphibians | Plectrohyla_ixil | Plectrohyla_ixil | 1 | 9.03 | 114 | 2924.83 |
| amphibians | Plectrohyla_matudai | Plectrohyla_matudai | 0 | 10.34 | 122.5 | 7723.69 |
| amphibians | Plectrohyla_quecchi | Plectrohyla_quecchi | 1 | 9.03 | 94.5 | 3677.77 |
| amphibians | Plectrohyla_sagorum | Plectrohyla_sagorum | 1 | 12.53 | 141 | 3741.32 |
| amphibians | Plectrohyla_tecunumani | Plectrohyla_tecunumani | 1 | 22.16 | 113 | 58.04 |
| amphibians | Plectrohyla_teuchestes | Plectrohyla_teuchestes | 1 | 43.04 | 384.5 | 54.14 |
| amphibians | Plethodon_albagula | Plethodon_albagula | 0 | 3.46 | 11.5 | 323541 |
| amphibians | Plethodon_angusticlavius | Plethodon_angusticlavius | 0 | 1.11 | 6 | 65433.76 |
| amphibians | Plethodon_asupak | Plethodon_asupak | 1 | 4.87 | 9 | 553.81 |
| amphibians | Plethodon_caddoensis | Plethodon_caddoensis | 0 | 2.29 | 7.5 | 961.74 |
| amphibians | Plethodon_cinereus | Plethodon_cinereus | 0 | 1.36 | 7.5 | 1830423.81 |
| amphibians | Plethodon_dorsalis | Plethodon_dorsalis | 0 | 1.6 | 5.5 | 287031.84 |
| amphibians | Plethodon_dunni | Plethodon_dunni | 0 | 7 | 11.5 | 73586.87 |
| amphibians | Plethodon_elongatus | Plethodon_elongatus | 0 | 7 | 7 | 24407.8 |
| amphibians | Plethodon_glutinosus | Plethodon_glutinosus | 0 | 4.61 | 19.5 | 1260100.04 |
| amphibians | Plethodon_hoffmani | Plethodon_hoffmani | 0 | 1.71 | 5.5 | 59988.37 |
| amphibians | Plethodon_hubrichti | Plethodon_hubrichti | 1 | 1.5 | 10 | 49.26 |
| amphibians | Plethodon_idahoensis | Plethodon_idahoensis | 0 | 4.62 | 8 | 82818.05 |
| amphibians | Plethodon_kentucki | Plethodon_kentucki | 0 | 2.94 | 10.5 | 29180.69 |
| amphibians | Plethodon_larselli | Plethodon_larselli | 0 | 2.56 | 7 | 24133.26 |
| amphibians | Plethodon_neomexicanus | Plethodon_neomexicanus | 1 | 4.83 | 8.5 | 626.41 |
| amphibians | Plethodon_nettingi | Plethodon_nettingi | 0 | 1.7 | 10.5 | 3468.12 |
| amphibians | Plethodon_ouachitae | Plethodon_ouachitae | 0 | 3.24 | 18 | 3272.45 |
| amphibians | Plethodon_punctatus | Plethodon_punctatus | 0 | 4.72 | 11.5 | 6730.99 |
| amphibians | Plethodon_richmondi | Plethodon_richmondi | 0 | 2.05 | 10 | 100792.82 |
| amphibians | Plethodon_serratus | Plethodon_serratus | 0 | 0.95 | 6 | 192070.27 |
| amphibians | Plethodon_shenandoah | Plethodon_shenandoah | 1 | 1.69 | 13 | 27.84 |
| amphibians | Plethodon_stormi | Plethodon_stormi | 1 | 4.01 | 10 | 2274.87 |
| amphibians | Plethodon_vandykei | Plethodon_vandykei | 0 | 4.03 | 12.5 | 16759.71 |
| amphibians | Plethodon_vehiculum | Plethodon_vehiculum | 0 | 4.03 | 13 | 166117.49 |
| amphibians | Plethodon_ventralis | Plethodon_ventralis | 0 | 1.15 | 3.5 | 17275.36 |
| amphibians | Plethodon_websteri | Plethodon_websteri | 0 | 0.73 | 5.5 | 23302.24 |

| | | | | | | |
|---|---|---|---|---|---|---|
| amphibians | Plethodon_wehrlei | Plethodon_wehrlei | 0 | 2.86 | 15 | 96744.46 |
| amphibians | Plethodon_welleri | Plethodon_welleri | 1 | 1.45 | 7.5 | 3540.24 |
| amphibians | Plethodon_yonahlossee | Plethodon_yonahlossee | 0 | 5.97 | 23 | 12313.33 |
| amphibians | Plethodontohyla_mihanika | Plethodontohyla_mihanika | 0 | 3.29 | 75.5 | 34747.56 |
| amphibians | Plethodontohyla_notosticta | Plethodontohyla_notosticta | 0 | 6.27 | 81.5 | 102707.46 |
| amphibians | Plethodontohyla_tuberata | Plethodontohyla_tuberata | 0 | 7.84 | 36.5 | 21100.6 |
| amphibians | Pleurodeles_nebulosus | Pleurodeles_nebulosus | 0 | 95.67 | 775 | 59414.74 |
| amphibians | Pleurodeles_waltl | Pleurodeles_waltl | 0 | 28.81 | 1075 | 377028.15 |
| amphibians | Pleurodema_bibroni | Pleurodema_bibroni | 0 | 15.03 | 450 | 99712.45 |
| amphibians | Pleurodema_brachyops | Pleurodema_brachyops | 0 | 11.77 | 1075 | 890578.52 |
| amphibians | Pleurodema_bufoninum | Pleurodema_bufoninum | 0 | 15.93 | 25 | 650772.48 |
| amphibians | Pleurodema_cordobae | Pleurodema_cordobae | 0 | 6.72 | 144.5 | 2851.59 |
| amphibians | Pleurodema_diplolister | Pleurodema_diplolister | 0 | 3.32 | 650 | 969995.98 |
| amphibians | Pleurodema_thaul | Pleurodema_thaul | 0 | 17.85 | 436 | 274397.66 |
| amphibians | Polypedates_braueri | Polypedates_braueri | 0 | 41.05 | 350 | 2138740.74 |
| amphibians | Polypedates_leucomystax | Polypedates_leucomystax | 0 | 61.58 | 500 | 3798236.02 |
| amphibians | Polypedates_maculatus | Polypedates_maculatus | 0 | 71.48 | 575 | 2581270.75 |
| amphibians | Polypedates_megacephalus | Polypedates_megacephalus | 0 | 37.61 | 396 | 1324419.43 |
| amphibians | Poyntonophrynus_dombensis | Poyntonophrynus_dombensis | 0 | 5.36 | 900 | 286233.16 |
| amphibians | Poyntonophrynus_fenoulheti | Poyntonophrynus_fenoulheti | 0 | 7.84 | 1122.5 | 670931.94 |
| amphibians | Poyntonophrynus_hoeschi | Poyntonophrynus_hoeschi | 0 | 4.16 | 400 | 120435.96 |
| amphibians | Pristimantis_achatinus | Pristimantis_achatinus | 0 | 8.48 | 35.5 | 187731.78 |
| amphibians | Pristimantis_acuminatus | Pristimantis_acuminatus | 0 | 3.16 | 16.5 | 96581.52 |
| amphibians | Pristimantis_albertus | Pristimantis_albertus | 1 | 0.63 | 16 | 80.88 |
| amphibians | Pristimantis_altamazonicus | Pristimantis_altamazonicus | 0 | 3.13 | 18.5 | 1034298.6 |
| amphibians | Pristimantis_andinognomus | Pristimantis_andinognomus | 0 | 0.4 | 10 | 1554.26 |
| amphibians | Pristimantis_aureolineatus | Pristimantis_aureolineatus | 0 | 2.22 | 17 | 205337.42 |
| amphibians | Pristimantis_bambu | Pristimantis_bambu | 1 | 1.39 | 30 | 119.53 |
| amphibians | Pristimantis_bicantus | Pristimantis_bicantus | 1 | 0.74 | 16.5 | 6792.43 |
| amphibians | Pristimantis_bipunctatus | Pristimantis_bipunctatus | 0 | 6.03 | 63 | 572.44 |
| amphibians | Pristimantis_buccinator | Pristimantis_buccinator | 0 | 5.36 | 43 | 146985.81 |
| amphibians | Pristimantis_calcarulatus | Pristimantis_calcarulatus | 1 | 1.87 | 20 | 3865.01 |
| amphibians | Pristimantis_colodactylus | Pristimantis_colodactylus | 0 | 1.29 | 8 | 5159.43 |
| amphibians | Pristimantis_conspicillatus | Pristimantis_conspicillatus | 0 | 10.34 | 44.5 | 2709642.99 |

| | | | | | | |
|---|---|---|---|---|---|---|
| amphibians | Pristimantis_croceoinguinis | Pristimantis_croceoinguinis | 0 | 0.89 | 94 | 149848.42 |
| amphibians | Pristimantis_diadematus | Pristimantis_diadematus | 0 | 7.57 | 56.5 | 548888.37 |
| amphibians | Pristimantis_esmeraldas | Pristimantis_esmeraldas | 0 | 0.79 | 8.5 | 29959.85 |
| amphibians | Pristimantis_fenestratus | Pristimantis_fenestratus | 0 | 17.07 | 34 | 2603658.33 |
| amphibians | Pristimantis_imitatrix | Pristimantis_imitatrix | 0 | 0.59 | 5.5 | 54873.07 |
| amphibians | Pristimantis_lacrimosus | Pristimantis_lacrimosus | 0 | 3.16 | 11.5 | 1144976.57 |
| amphibians | Pristimantis_lanthanites | Pristimantis_lanthanites | 0 | 8.42 | 176 | 692688.53 |
| amphibians | Pristimantis_latidiscus | Pristimantis_latidiscus | 0 | 13.66 | 40 | 90430.56 |
| amphibians | Pristimantis_martiae | Pristimantis_martiae | 0 | 1.17 | 8.5 | 968361.27 |
| amphibians | Pristimantis_mindo | Pristimantis_mindo | 1 | 5.71 | 50 | 1224.37 |
| amphibians | Pristimantis_minimus | Pristimantis_minimus | 1 | 0.47 | 3 | 296.77 |
| amphibians | Pristimantis_ockendeni | Pristimantis_ockendeni | 0 | 2.47 | 16.5 | 1624566.16 |
| amphibians | Pristimantis_parvillus | Pristimantis_parvillus | 1 | 1.31 | 36.5 | 37549.88 |
| amphibians | Pristimantis_paululus | Pristimantis_paululus | 0 | 0.66 | 5 | 54443.27 |
| amphibians | Pristimantis_peruvianus | Pristimantis_peruvianus | 0 | 8.66 | 34.5 | 1084951.52 |
| amphibians | Pristimantis_philipi | Pristimantis_philipi | 0 | 3.07 | 27 | 169.02 |
| amphibians | Pristimantis_pseudoacuminatus | Pristimantis_pseudoacuminatus | 0 | 0.89 | 7.5 | 39677.58 |
| amphibians | Pristimantis_quaquaversus | Pristimantis_quaquaversus | 0 | 2.42 | 22 | 90970.77 |
| amphibians | Pristimantis_ramagii | Pristimantis_ramagii | 0 | 2.11 | 10 | 71229.48 |
| amphibians | Pristimantis_rosadoi | Pristimantis_rosadoi | 1 | 1.28 | 52 | 15668.02 |
| amphibians | Pristimantis_skydmainos | Pristimantis_skydmainos | 0 | 2.71 | 18 | 581754.51 |
| amphibians | Pristimantis_sulculus | Pristimantis_sulculus | 1 | 14.16 | 32 | 5.47 |
| amphibians | Pristimantis_toftae | Pristimantis_toftae | 0 | 1.57 | 16.5 | 233565.13 |
| amphibians | Pristimantis_tungurahua | Pristimantis_tungurahua | 1 | 1.67 | 8 | 57.81 |
| amphibians | Pristimantis_variabilis | Pristimantis_variabilis | 0 | 1.46 | 6.5 | 605438.53 |
| amphibians | Pristimantis_vertebralis | Pristimantis_vertebralis | 1 | 7.96 | 67 | 4072.46 |
| amphibians | Pristimantis_wagteri | Pristimantis_wagteri | 1 | 7.29 | 14 | 529.24 |
| amphibians | Probreviceps_loveridgei | Probreviceps_loveridgei | 1 | 8.42 | 35 | 2121.85 |
| amphibians | Probreviceps_macrodactylus | Probreviceps_macrodactylus | 1 | 25.82 | 42.5 | 864.45 |
| amphibians | Probreviceps_rhodesianus | Probreviceps_rhodesianus | 1 | 10.34 | 19.5 | 1323.65 |
| amphibians | Probreviceps_uluguruensis | Probreviceps_uluguruensis | 1 | 6.77 | 20 | 443.67 |
| amphibians | Proceratophrys_appendiculata | Proceratophrys_appendiculata | 0 | 23.55 | 598.5 | 6804.88 |
| amphibians | Proceratophrys_boiei | Proceratophrys_boiei | 0 | 39.82 | 1296 | 385929.03 |
| amphibians | Proceratophrys_melanopogon | Proceratophrys_melanopogon | 0 | 26.87 | 664 | 18819.42 |

| | | | | | | |
|---|---|---|---|---|---|---|
| amphibians | Proceratophrys_tupinamba | Proceratophrys_tupinambis | 0 | 37.28 | 837.5 | 184.39 |
| amphibians | Proteus_anguinus | Proteus_anguinus | 1 | 74.48 | 52.5 | 16162.8 |
| amphibians | Pseudacris_brachyphona | Pseudacris_brachyphona | 0 | 4.54 | 4005 | 190497.81 |
| amphibians | Pseudacris_brimleyi | Pseudacris_brimleyi | 0 | 3.47 | 247 | 135121.32 |
| amphibians | Pseudacris_cadaverina | Pseudacris_cadaverina | 0 | 11.77 | 265 | 75369.28 |
| amphibians | Pseudacris_clarkii | Pseudacris_clarkii | 0 | 2.6 | 575 | 556285.47 |
| amphibians | Pseudacris_crucifer | Pseudacris_crucifer | 0 | 4.16 | 917 | 4040940.03 |
| amphibians | Pseudacris_feriarum | Pseudacris_feriarum | 0 | 4.54 | 509 | 759360.33 |
| amphibians | Pseudacris_fouquettei | Pseudacris_fouquettei | 0 | 2.11 | 912.5 | 604902.11 |
| amphibians | Pseudacris_illinoensis | Pseudacris_illinoensis | 0 | 9.67 | 580 | 29551.18 |
| amphibians | Pseudacris_kalmi | Pseudacris_kalmi | 0 | 3.81 | 648.5 | 34400.23 |
| amphibians | Pseudacris_maculata | Pseudacris_maculata | 0 | 12.53 | 798 | 5606940.29 |
| amphibians | Pseudacris_nigrita | Pseudacris_nigrita | 0 | 2.87 | 429.5 | 384813.58 |
| amphibians | Pseudacris_ocularis | Pseudacris_ocularis | 0 | 0.37 | 550 | 341591.46 |
| amphibians | Pseudacris_ornata | Pseudacris_ornata | 0 | 5.36 | 505 | 347129.29 |
| amphibians | Pseudacris_regilla | Pseudacris_regilla | 0 | 11.04 | 882.5 | 1621880.11 |
| amphibians | Pseudacris_streckeri | Pseudacris_streckeri | 0 | 9.67 | 450 | 519936.45 |
| amphibians | Pseudacris_triseriata | Pseudacris_triseriata | 0 | 5.14 | 800 | 409563.54 |
| amphibians | Pseudis_bolbodactyla | Pseudis_bolbodactyla | 0 | 17.85 | 230 | 592141.85 |
| amphibians | Pseudis_cardosoi | Pseudis_cardosoi | 0 | 15.93 | 135 | 47382.88 |
| amphibians | Pseudis_minuta | Pseudis_minuta | 0 | 11.84 | 486 | 453415.64 |
| amphibians | Pseudis_paradoxa | Pseudis_paradoxa | 0 | 104.27 | 2574.5 | 5040866.18 |
| amphibians | Pseudis_platensis | Pseudis_platensis | 0 | 17.85 | 1025.5 | 1605316.21 |
| amphibians | Pseudobranchus_axanthus | Pseudobranchus_axanthus | 0 | 10.95 | 39 | 99563.71 |
| amphibians | Pseudobranchus_striatus | Pseudobranchus_striatus | 0 | 11.55 | 6 | 102698.04 |
| amphibians | Pseudoeurycea_goebeli | Pseudoeurycea_goebeli | 1 | 3.32 | 30 | 616.47 |
| amphibians | Pseudoeurycea_leprosa | Pseudoeurycea_leprosa | 0 | 3.18 | 1722 | 12025.38 |
| amphibians | Pseudoeurycea_rex | Pseudoeurycea_rex | 1 | 1.93 | 20 | 1123.25 |
| amphibians | Pseudohynobius_flavomaculatus | Pseudohynobius_flavomaculatus | 1 | 9.76 | 42 | 1587.05 |
| amphibians | Pseudohynobius_puxiongensis | Pseudohynobius_puxiongensis | 1 | 5.82 | 34 | 10.79 |
| amphibians | Pseudopaludicola_boliviana | Pseudopaludicola_boliviana | 0 | 0.22 | 184.5 | 3016227.57 |
| amphibians | Pseudopaludicola_falcipes | Pseudopaludicola_falcipes | 0 | 0.57 | 300 | 729436.77 |
| amphibians | Pseudopaludicola_murundu | Pseudopaludicola_murundu | 0 | 0.48 | 32 | 203289.43 |
| amphibians | Pseudopaludicola_mystacalis | Pseudopaludicola_mystacalis | 0 | 0.47 | 119 | 3075446.31 |

| | | | | | | |
|---|---|---|---|---|---|---|
| amphibians | Pseudopaludicola_saltica | Pseudopaludicola_saltica | 0 | 0.77 | 92 | 1526283.82 |
| amphibians | Pseudopaludicola_ternetzi | Pseudopaludicola_ternetzi | 0 | 0.57 | 46.5 | 1102585.07 |
| amphibians | Pseudophilautus_popularis | Pseudophilautus_popularis | 1 | 1.26 | 11 | 7545.45 |
| amphibians | Pseudophilautus_regius | Pseudophilautus_regius | 0 | 0.74 | 17 | 34013.8 |
| amphibians | Pseudophilautus_wynaadensis | Pseudophilautus_wynaadensis | 0 | 1.75 | 31 | 56408.82 |
| amphibians | Pseudophryne_australis | Pseudophryne_australis | 0 | 2.11 | 22 | 23013.34 |
| amphibians | Pseudophryne_bibronii | Pseudophryne_bibronii | 0 | 3.81 | 185 | 630882.11 |
| amphibians | Pseudophryne_coriacea | Pseudophryne_coriacea | 0 | 3.47 | 87.5 | 99816.2 |
| amphibians | Pseudophryne_corroboree | Pseudophryne_corroboree | 1 | 2.35 | 26.5 | 724.62 |
| amphibians | Pseudophryne_covacevichae | Pseudophryne_covacevichae | 1 | 2.11 | 44 | 368.19 |
| amphibians | Pseudophryne_dendyi | Pseudophryne_dendyi | 0 | 3.47 | 109 | 73886.95 |
| amphibians | Pseudophryne_douglasi | Pseudophryne_douglasi | 0 | 2.6 | 84.5 | 117377.92 |
| amphibians | Pseudophryne_guentheri | Pseudophryne_guentheri | 0 | 2.87 | 150.5 | 384203.41 |
| amphibians | Pseudophryne_pengilleyi | Pseudophryne_pengilleyi | 1 | 2.11 | 25 | 980.52 |
| amphibians | Pseudophryne_semimarmorata | Pseudophryne_semimarmorata | 0 | 3.47 | 242.5 | 172714.54 |
| amphibians | Pseudorana_weiningensis | Rana_weiningensis | 0 | 6.77 | 313 | 57098.11 |
| amphibians | Pseudotriton_montanus | Pseudotriton_montanus | 0 | 6.72 | 109.5 | 758984.36 |
| amphibians | Pseudotriton_ruber | Pseudotriton_ruber | 0 | 8.98 | 79.5 | 1068841.55 |
| amphibians | Ptychadena_aequiplicata | Ptychadena_aequiplicata | 0 | 24.56 | 499.5 | 1235543.07 |
| amphibians | Ptychadena_anchietae | Ptychadena_anchietae | 0 | 22.16 | 250 | 6632225.54 |
| amphibians | Ptychadena_bibroni | Ptychadena_bibroni | 0 | 24.56 | 1150 | 3611728.65 |
| amphibians | Ptychadena_mascareniensis | Ptychadena_mascareniensis | 0 | 35.97 | 1089.5 | 6801836.23 |
| amphibians | Ptychadena_nana | Ptychadena_nana | 1 | 2.11 | 335.5 | 18.55 |
| amphibians | Ptychadena_nilotica | Ptychadena_nilotica | 0 | 5.36 | 5621.5 | 1968905.85 |
| amphibians | Ptychadena_oxyrhynchus | Ptychadena_oxyrhynchus | 0 | 61.58 | 2675 | 8134935.63 |
| amphibians | Ptychadena_schillukorum | Ptychadena_schillukorum | 0 | 15.03 | 1089 | 5068749.54 |
| amphibians | Ptychadena_subpunctata | Ptychadena_subpunctata | 0 | 29.89 | 1000 | 1572767.26 |
| amphibians | Ptychadena_taenioscelis | Ptychadena_taenioscelis | 0 | 5.36 | 6 | 3975471.58 |
| amphibians | Ptychadena_tellinii | Ptychadena_tellinii | 0 | 16.87 | 1255.5 | 2476861.67 |
| amphibians | Ptychohyla_euthysanota | Ptychohyla_euthysanota | 0 | 13.58 | 210 | 13194.68 |
| amphibians | Ptychohyla_leonhardschultzei | Ptychohyla_leonhardschultzei | 0 | 6.98 | 265 | 3683.9 |
| amphibians | Pyxicephalus_adspersus | Pyxicephalus_adspersus | 0 | 1901.29 | 3500 | 3658402.38 |
| amphibians | Pyxicephalus_edulis | Pyxicephalus_edulis | 0 | 347.52 | 2200 | 2945288.23 |
| amphibians | Quasipaa_boulengeri | Quasipaa_boulengeri | 1 | 269.14 | 218 | 990447.79 |

| | | | | | | |
|---|---|---|---|---|---|---|
| amphibians | Quasipaa_exilispinosa | Quasipaa_exilispinosa | 0 | 25.69 | 7.5 | 323280.94 |
| amphibians | Quasipaa_shini | Quasipaa_shini | 1 | 99.61 | 10.5 | 102297.28 |
| amphibians | Quasipaa_spinosa | Quasipaa_spinosa | 1 | 107.68 | 338 | 765191.44 |
| amphibians | Quasipaa_verrucospinosa | Quasipaa_verrucospinosa | 0 | 359.31 | 3168 | 136064.68 |
| amphibians | Quasipaa_yei | Quasipaa_yei | 1 | 41.05 | 265 | 21970.92 |
| amphibians | Quilticohyla_sanctaecrucis | Ptychohyla_sanctaecrucis | 1 | 11.77 | 146 | 170.35 |
| amphibians | Rana_amurensis | Rana_amurensis | 0 | 32.83 | 2225 | 7752968.84 |
| amphibians | Rana_arvalis | Rana_arvalis | 0 | 50.6 | 2250 | 9971161.18 |
| amphibians | Rana_asiatica | Rana_asiatica | 1 | 13.99 | 1150 | 159462.91 |
| amphibians | Rana_chaochiaoensis | Rana_chaochiaoensis | 0 | 21.02 | 1500 | 359124.02 |
| amphibians | Rana_chensinensis | Rana_chensinensis | 0 | 27.13 | 1082.5 | 2084914.76 |
| amphibians | Rana_coreana | Rana_coreana | 0 | 7.73 | 918 | 149953.33 |
| amphibians | Rana_dalmatina | Rana_dalmatina | 0 | 88.3 | 1150 | 1950863.52 |
| amphibians | Rana_dybowskii | Rana_dybowskii | 0 | 59.27 | 1716.5 | 2023583.6 |
| amphibians | Rana_graeca | Rana_graeca | 0 | 50.6 | 1100 | 236208.94 |
| amphibians | Rana_huanrensis | Rana_huanrensis | 0 | 10.34 | 450 | 240875.48 |
| amphibians | Rana_iberica | Rana_iberica | 1 | 32.83 | 325 | 106637.61 |
| amphibians | Rana_italica | Rana_italica | 0 | 25.82 | 1675 | 115730.94 |
| amphibians | Rana_japonica | Rana_japonica | 0 | 35.97 | 1750 | 236424.42 |
| amphibians | Rana_johnsi | Rana_johnsi | 0 | 8.72 | 1550 | 672932.98 |
| amphibians | Rana_kukunoris | Rana_kukunoris | 0 | 22.04 | 1132.5 | 733978.08 |
| amphibians | Rana_latastei | Rana_latastei | 1 | 41.05 | 1560 | 33894.54 |
| amphibians | Rana_longicrus | Rana_longicrus | 1 | 11.23 | 520.5 | 6984.76 |
| amphibians | Rana_macrocnemis | Rana_macrocnemis | 0 | 89.21 | 2040 | 1083970.7 |
| amphibians | Rana_omeimontis | Rana_omeimontis | 0 | 32.38 | 1550 | 445271.27 |
| amphibians | Rana_ornativentris | Rana_ornativentris | 0 | 46.62 | 6500 | 281925.22 |
| amphibians | Rana_pirica | Rana_pirica | 0 | 35.97 | 900 | 158262.5 |
| amphibians | Rana_pyrenaica | Rana_pyrenaica | 1 | 11.04 | 142.5 | 2237.02 |
| amphibians | Rana_sakuraii | Rana_sakuraii | 0 | 19.92 | 110 | 46994.73 |
| amphibians | Rana_sauteri | Rana_sauteri | 1 | 13.17 | 667.5 | 15700.25 |
| amphibians | Rana_tagoi | Rana_tagoi | 0 | 17.95 | 1833 | 286740.38 |
| amphibians | Rana_temporaria | Rana_temporaria | 0 | 141.99 | 2382.5 | 7787374.87 |
| amphibians | Rana_tsushimensis | Rana_tsushimensis | 0 | 12.53 | 400 | 702.75 |
| amphibians | Rana_zhenhaiensis | Rana_zhenhaiensis | 0 | 9.73 | 659.5 | 836731.21 |

| | | | | | | |
|---|---|---|---|---|---|---|
| amphibians | Ranitomeya_amazonica | Ranitomeya_amazonica | 0 | 0.48 | 5.5 | 1301154.77 |
| amphibians | Ranitomeya_benedicta | Ranitomeya_benedicta | 1 | 0.66 | 5 | 10994.67 |
| amphibians | Ranitomeya_fantastica | Ranitomeya_fantastica | 1 | 0.89 | 5 | 7057.57 |
| amphibians | Ranitomeya_flavovittata | Ranitomeya_flavovittata | 0 | 0.48 | 2 | 10342.87 |
| amphibians | Ranitomeya_imitator | Ranitomeya_imitator | 0 | 0.77 | 2.5 | 9929.38 |
| amphibians | Ranitomeya_reticulata | Ranitomeya_reticulata | 0 | 0.57 | 3 | 26307.98 |
| amphibians | Ranitomeya_sirensis | Ranitomeya_sirensis | 0 | 3.13 | 2.5 | 309963.93 |
| amphibians | Ranitomeya_summersi | Ranitomeya_summersi | 1 | 0.6 | 6.5 | 242.87 |
| amphibians | Ranitomeya_toraro | Ranitomeya_toraro | 0 | 0.33 | 2 | 866697.71 |
| amphibians | Ranitomeya_uakarii | Ranitomeya_uakarii | 0 | 0.37 | 4 | 312456.02 |
| amphibians | Ranitomeya_vanzolinii | Ranitomeya_vanzolinii | 0 | 0.56 | 2.5 | 149685.29 |
| amphibians | Ranitomeya_variabilis | Ranitomeya_variabilis | 0 | 0.66 | 7.5 | 896931.18 |
| amphibians | Ranitomeya_ventrimaculata | Ranitomeya_ventrimaculata | 0 | 0.77 | 5.5 | 304522.45 |
| amphibians | Ranodon_sibiricus | Ranodon_sibiricus | 1 | 10.06 | 57 | 3060.71 |
| amphibians | Ranoidea_alboguttata | Cyclorana_alboguttata | 0 | 57.01 | 3743 | 1810747.76 |
| amphibians | Ranoidea_auae | Litoria_auae | 0 | 6.18 | 200 | 85554.68 |
| amphibians | Ranoidea_aurea | Litoria_aurea | 0 | 133.8 | 6821 | 140805.03 |
| amphibians | Ranoidea_australis | Cyclorana_australis | 0 | 122.13 | 3575 | 1353476.41 |
| amphibians | Ranoidea_barringtonensis | Litoria_barringtonensis | 0 | 4.16 | 450 | 30206.53 |
| amphibians | Ranoidea_booroolongensis | Litoria_booroolongensis | 1 | 14.16 | 1250 | 101669.58 |
| amphibians | Ranoidea_brevipes | Cyclorana_brevipes | 0 | 7.84 | 2172 | 1714335.15 |
| amphibians | Ranoidea_caerulea | Litoria_caerulea | 0 | 141.99 | 2303 | 3396988.57 |
| amphibians | Ranoidea_chloris | Litoria_chloris | 0 | 32.83 | 1431.5 | 127522.54 |
| amphibians | Ranoidea_citropa | Litoria_citropa | 0 | 32.83 | 797.5 | 54430.03 |
| amphibians | Ranoidea_cryptotis | Cyclorana_cryptotis | 0 | 8.42 | 975 | 660637.75 |
| amphibians | Ranoidea_daviesae | Litoria_daviesae | 1 | 24.56 | 425 | 9106.68 |
| amphibians | Ranoidea_dayi | Litoria_dayi | 1 | 19.92 | 228.5 | 13558.64 |
| amphibians | Ranoidea_eucnemis | Litoria_eucnemis | 0 | 61.58 | 861 | 410475.96 |
| amphibians | Ranoidea_genimaculata | Litoria_genimaculata | 0 | 61.58 | 523 | 244188.6 |
| amphibians | Ranoidea_gracilenta | Litoria_gracilenta | 0 | 7.84 | 750 | 164656.93 |
| amphibians | Ranoidea_jungguy | Litoria_jungguy | 0 | 34.37 | 1440 | 22545.06 |
| amphibians | Ranoidea_kroombitensis | Litoria_kroombitensis | 1 | 7.29 | 300 | 38.49 |
| amphibians | Ranoidea_lesueuri | Litoria_lesueurii | 0 | 32.83 | 2048.5 | 171995.93 |
| amphibians | Ranoidea_longipes | Cyclorana_longipes | 0 | 15.03 | 1369.5 | 872725.52 |

| | | | | | | |
|---|---|---|---|---|---|---|
| amphibians | Ranoidea_lorica | Litoria_lorica | 1 | 5.8 | 106.5 | 819.11 |
| amphibians | Ranoidea_myola | Litoria_myola | 1 | 31.33 | 509 | 16.61 |
| amphibians | Ranoidea_nannotis | Litoria_nannotis | 0 | 25.82 | 135 | 15721.69 |
| amphibians | Ranoidea_novaehollandiae | Cyclorana_novaehollandiae | 0 | 107.68 | 4304 | 1711516.63 |
| amphibians | Ranoidea_nudidigita | Litoria_nudidigita | 0 | 5.8 | 377 | 62221.61 |
| amphibians | Ranoidea_pearsoniana | Litoria_pearsoniana | 0 | 5.36 | 547.5 | 85282.47 |
| amphibians | Ranoidea_phyllochroa | Litoria_phyllochroa | 0 | 5.36 | 428 | 60330.59 |
| amphibians | Ranoidea_platycephala | Cyclorana_platycephala | 0 | 35.97 | 1750 | 2020180.95 |
| amphibians | Ranoidea_raniformis | Litoria_raniformis | 1 | 118.4 | 2796.5 | 731178.31 |
| amphibians | Ranoidea_rheocola | Litoria_rheocola | 0 | 11.04 | 138.5 | 14112.25 |
| amphibians | Ranoidea_serrata | Litoria_serrata | 0 | 61.58 | 741.5 | 20180.93 |
| amphibians | Ranoidea_spenceri | Litoria_spenceri | 1 | 15.03 | 558 | 9004.79 |
| amphibians | Ranoidea_splendida | Litoria_splendida | 0 | 178.26 | 7281.5 | 213142.87 |
| amphibians | Ranoidea_subglandulosa | Litoria_subglandulosa | 1 | 11.04 | 358.5 | 14117.03 |
| amphibians | Ranoidea_vagitus | Cyclorana_vagitus | 0 | 10.34 | 2715 | 55382.59 |
| amphibians | Ranoidea_wilcoxii | Litoria_wilcoxii | 0 | 32.83 | 3185 | 486704.9 |
| amphibians | Ranoidea_xanthomera | Litoria_xanthomera | 0 | 61.58 | 863 | 22195.09 |
| amphibians | Raorchestes_glandulosus | Raorchestes_glandulosus | 1 | 4.54 | 47 | 10082.3 |
| amphibians | Raorchestes_ponmudi | Raorchestes_ponmudi | 0 | 4.89 | 79.5 | 18282.62 |
| amphibians | Rentapia_hosii | Pedostibes_hosii | 0 | 122.13 | 5860 | 293952.56 |
| amphibians | Rhacophorus_annamensis | Rhacophorus_annamensis | 0 | 74.11 | 175 | 178585.83 |
| amphibians | Rhacophorus_kio | Rhacophorus_kio | 0 | 53.95 | 265 | 883418.81 |
| amphibians | Rhacophorus_malabaricus | Rhacophorus_malabaricus | 0 | 52.68 | 145.5 | 54438.94 |
| amphibians | Rhacophorus_pardalis | Rhacophorus_pardalis | 0 | 41.05 | 47 | 157763.77 |
| amphibians | Rhacophorus_reinwardtii | Rhacophorus_reinwardtii | 0 | 56.35 | 68 | 125954.88 |
| amphibians | Rhacophorus_rhodopus | Rhacophorus_rhodopus | 0 | 13.33 | 125 | 1098451.28 |
| amphibians | Rhaebo_blombergi | Rhaebo_blombergi | 1 | 2029.91 | 47500 | 18995.37 |
| amphibians | Rhaebo_ceratophrys | Rhinella_ceratophrys | 0 | 128.26 | 1250 | 953465.34 |
| amphibians | Rhaebo_glaberrimus | Rhaebo_glaberrimus | 0 | 141.99 | 6034.5 | 42940.72 |
| amphibians | Rhaebo_haematiticus | Rhaebo_haematiticus | 0 | 147.5 | 1000 | 316858.72 |
| amphibians | Rheobates_palmatus | Rheobates_palmatus | 0 | 4.93 | 23.5 | 49585.93 |
| amphibians | Rheohyla_miotympanum | Ecnomiohyla_miotympanum | 0 | 12.22 | 192.5 | 25544.43 |
| amphibians | Rhinella_achalensis | Rhinella_achalensis | 1 | 31.33 | 1624 | 2338.98 |
| amphibians | Rhinella_acrolopha | Rhinella_acrolopha | 1 | 7.84 | 290 | 900.63 |

| | | | | | | |
|---|---|---|---|---|---|---|
| amphibians | Rhinella_amabilis | Rhinella_amabilis | 1 | 93.84 | 3218 | 6.64 |
| amphibians | Rhinella_arborescandens | Rhinella_arborescandens | 1 | 4.54 | 5000 | 377.61 |
| amphibians | Rhinella_arenarum | Rhinella_arenarum | 0 | 188.23 | 19023 | 1798246.51 |
| amphibians | Rhinella_arunco | Rhinella_arunco | 0 | 163.99 | 5000 | 73509.7 |
| amphibians | Rhinella_atacamensis | Rhinella_atacamensis | 1 | 77.36 | 5000 | 13595.03 |
| amphibians | Rhinella_castaneotica | Rhinella_castaneotica | 0 | 11.4 | 224 | 2519135.75 |
| amphibians | Rhinella_chavin | Rhinella_chavin | 1 | 25.69 | 276 | 75.73 |
| amphibians | Rhinella_chrysophora | Rhinella_chrysophora | 1 | 19.5 | 915 | 1027.6 |
| amphibians | Rhinella_crucifer | Rhinella_crucifer | 0 | 107.68 | 3334.84 | 1232177.54 |
| amphibians | Rhinella_dapsilis | Rhinella_dapsilis | 0 | 81.85 | 22 | 692612.78 |
| amphibians | Rhinella_diptycha | Rhinella_schneideri | 0 | 2029.91 | 1608.5 | 6282831.93 |
| amphibians | Rhinella_dorbignyi | Rhinella_dorbignyi | 0 | 32.83 | 9000 | 873585.25 |
| amphibians | Rhinella_granulosa | Rhinella_granulosa | 0 | 74.11 | 4224 | 1204162.53 |
| amphibians | Rhinella_humboldti | Rhinella_humboldti | 0 | 23.94 | 2863 | 256711.46 |
| amphibians | Rhinella_icterica | Rhinella_icterica | 0 | 834.28 | 8840.42 | 661347.21 |
| amphibians | Rhinella_major | Rhinella_major | 0 | 15.03 | 9732 | 3999880.42 |
| amphibians | Rhinella_manu | Rhinella_manu | 0 | 6.18 | 37.5 | 9176.62 |
| amphibians | Rhinella_margaritifera | Rhinella_margaritifera | 0 | 46.62 | 1343 | 7506418.39 |
| amphibians | Rhinella_marina | Rhinella_marina | 0 | 2029.91 | 20050 | 9335781.42 |
| amphibians | Rhinella_nesiotes | Rhinella_nesiotes | 1 | 1.89 | 20 | 63.2 |
| amphibians | Rhinella_ocellata | Rhinella_ocellata | 0 | 13.33 | 1345 | 1916157.85 |
| amphibians | Rhinella_ornata | Rhinella_ornata | 0 | 59.93 | 3720.5 | 439232.47 |
| amphibians | Rhinella_proboscidea | Rhinella_proboscidea | 0 | 15.03 | 456 | 436709.57 |
| amphibians | Rhinella_pygmaea | Rhinella_pygmaea | 0 | 11.04 | 4140 | 34412.58 |
| amphibians | Rhinella_quechua | Rhinella_quechua | 1 | 22.16 | 1014 | 5461.05 |
| amphibians | Rhinella_scitula | Rhinella_scitula | 0 | 9.67 | 532.5 | 142806.32 |
| amphibians | Rhinella_spinulosa | Rhinella_spinulosa | 0 | 82.42 | 4825 | 1145779.99 |
| amphibians | Rhinella_yanachaga | Rhinella_yanachaga | 1 | 8.25 | 136 | 160.22 |
| amphibians | Rhinoderma_darwinii | Rhinoderma_darwinii | 1 | 4.35 | 22 | 88596.83 |
| amphibians | Rhinoderma_rufum | Rhinoderma_rufum | 1 | 2.87 | 18 | 23981.23 |
| amphibians | Rhinophrynus_dorsalis | Rhinophrynus_dorsalis | 0 | 74.11 | 4132.5 | 342090.37 |
| amphibians | Rhombophryne_testudo | Rhombophryne_testudo | 1 | 7.84 | 11 | 311.16 |
| amphibians | Rhyacotriton_cascadae | Rhyacotriton_cascadae | 0 | 2.2 | 8 | 11339.48 |
| amphibians | Rhyacotriton_kezeri | Rhyacotriton_kezeri | 0 | 2.7 | 12.5 | 17307.96 |

| | | | | | | |
|---|---|---|---|---|---|---|
| amphibians | Rhyacotriton_olympicus | Rhyacotriton_olympicus | 0 | 3.32 | 8.5 | 4998.62 |
| amphibians | Rhyacotriton_variegatus | Rhyacotriton_variegatus | 0 | 1.61 | 10 | 62030.56 |
| amphibians | Rulyrana_flavopunctata | Rulyrana_flavopunctata | 0 | 1.28 | 39 | 81909.66 |
| amphibians | Sachatamia_albomaculata | Sachatamia_albomaculata | 0 | 2.6 | 162.5 | 81367.95 |
| amphibians | Sachatamia_ilex | Sachatamia_ilex | 0 | 4.16 | 37.5 | 85352.74 |
| amphibians | Salamandra_algira | Salamandra_algira | 1 | 20.2 | 14.5 | 29255.66 |
| amphibians | Salamandra_atra | Salamandra_atra | 0 | 7 | 26 | 111210.66 |
| amphibians | Salamandra_corsica | Salamandra_corsica | 0 | 26.8 | 22.5 | 4417.81 |
| amphibians | Salamandra_infraimmaculata | Salamandra_infraimmaculata | 0 | 50.14 | 60 | 90468.94 |
| amphibians | Salamandra_lanzai | Salamandra_lanzai | 1 | 10.52 | 3.5 | 779.62 |
| amphibians | Salamandra_salamandra | Salamandra_salamandra | 1 | 57.14 | 39 | 2161490.75 |
| amphibians | Salamandrella_keyserlingii | Salamandrella_keyserlingii | 0 | 5.27 | 166 | 12290772.13 |
| amphibians | Salamandrina_perspicillata | Salamandrina_perspicillata | 1 | 2.04 | 47.5 | 74001.37 |
| amphibians | Salamandrina_terdigitata | Salamandrina_terdigitata | 0 | 0.84 | 53 | 28138.54 |
| amphibians | Sanguirana_everetti | Sanguirana_everetti | 0 | 65.42 | 925 | 2604.67 |
| amphibians | Sanguirana_sanguinea | Sanguirana_sanguinea | 0 | 34.53 | 180 | 15221.28 |
| amphibians | Sarcohyla_arborescandens | Plectrohyla_arborescandens | 0 | 12.22 | 436 | 13656.19 |
| amphibians | Sarcohyla_thorectes | Plectrohyla_thorectes | 1 | 4.46 | 10 | 136.61 |
| amphibians | Scaphiophryne_calcarata | Scaphiophryne_calcarata | 0 | 2.87 | 465 | 146997 |
| amphibians | Scaphiophryne_gottlebei | Scaphiophryne_gottlebei | 1 | 5.36 | 182 | 619.48 |
| amphibians | Scaphiophryne_menabensis | Scaphiophryne_menabensis | 0 | 7.84 | 560 | 41657.73 |
| amphibians | Scaphiopus_couchii | Scaphiopus_couchii | 0 | 74.11 | 1805 | 1960473.33 |
| amphibians | Scaphiopus_holbrookii | Scaphiopus_holbrookii | 0 | 46.62 | 3134 | 1122187.24 |
| amphibians | Scaphiopus_hurterii | Scaphiopus_hurterii | 0 | 57.01 | 2539.5 | 453628.45 |
| amphibians | Scarthyla_goinorum | Scarthyla_goinorum | 0 | 1.17 | 112 | 1659883.96 |
| amphibians | Schismaderma_carens | Schismaderma_carens | 0 | 79.58 | 10100 | 2960397.4 |
| amphibians | Scinax_acuminatus | Scinax_acuminatus | 0 | 9.67 | 973 | 769187.37 |
| amphibians | Scinax_alter | Scinax_alter | 0 | 2.6 | 365.5 | 207023.41 |
| amphibians | Scinax_boesemani | Scinax_boesemani | 0 | 5.36 | 600 | 4415912.24 |
| amphibians | Scinax_boulengeri | Scinax_boulengeri | 0 | 18.35 | 516 | 122080.66 |
| amphibians | Scinax_chiquitanus | Scinax_chiquitanus | 0 | 3.88 | 789 | 500762.76 |
| amphibians | Scinax_constrictus | Scinax_constrictus | 0 | 3.67 | 207 | 1236009.45 |
| amphibians | Scinax_cruentomma | Scinax_cruentommus | 0 | 2.6 | 640 | 1751608.82 |
| amphibians | Scinax_cuspidatus | Scinax_cuspidatus | 0 | 2.73 | 170.5 | 291909.04 |

| | | | | | | |
|---|---|---|---|---|---|---|
| amphibians | Scinax_duartei | Scinax_duartei | 0 | 4.16 | 385 | 3306.41 |
| amphibians | Scinax_elaeochroa | Scinax_elaeochroa | 0 | 5.49 | 530 | 46071.23 |
| amphibians | Scinax_eurydice | Scinax_eurydice | 0 | 13.99 | 1699.5 | 586523.83 |
| amphibians | Scinax_funereus | Scinax_funereus | 0 | 6.77 | 482.5 | 959399.49 |
| amphibians | Scinax_fuscomarginatus | Scinax_fuscomarginatus | 0 | 1.45 | 303 | 5108321.19 |
| amphibians | Scinax_fuscovarius | Scinax_fuscovarius | 0 | 15.44 | 2684 | 5763739.25 |
| amphibians | Scinax_garbei | Scinax_garbei | 0 | 10.41 | 662 | 4506038.85 |
| amphibians | Scinax_granulatus | Scinax_granulatus | 0 | 6.77 | 1200 | 888877.14 |
| amphibians | Scinax_hayii | Scinax_hayii | 0 | 13.33 | 1947.5 | 222916.66 |
| amphibians | Scinax_ictericus | Scinax_ictericus | 0 | 3.41 | 639 | 564135.36 |
| amphibians | Scinax_nasicus | Scinax_nasicus | 0 | 4.93 | 798.5 | 2861068.44 |
| amphibians | Scinax_nebulosus | Scinax_nebulosus | 0 | 5.36 | 527 | 6018245.05 |
| amphibians | Scinax_pedromedinae | Scinax_pedromedinae | 0 | 2.47 | 234 | 802070.35 |
| amphibians | Scinax_perereca | Scinax_perereca | 0 | 8.42 | 835 | 314816.56 |
| amphibians | Scinax_ruber | Scinax_ruber | 0 | 7.84 | 619 | 8019881.42 |
| amphibians | Scinax_squalirostris | Scinax_squalirostris | 0 | 1.89 | 237 | 1411664.87 |
| amphibians | Scinax_staufferi | Scinax_staufferi | 0 | 2.6 | 265 | 569553.53 |
| amphibians | Scinax_x-signatus | Scinax_x-signatus | 0 | 9.67 | 270 | 4512856.2 |
| amphibians | Sclerophrys_brauni | Amietophrynus_brauni | 0 | 141.99 | 9000 | 4459.78 |
| amphibians | Sclerophrys_camerunensis | Amietophrynus_camerunensis | 0 | 76.81 | 2250 | 900373.35 |
| amphibians | Sclerophrys_capensis | Amietophrynus_rangeri | 0 | 173.41 | 10380 | 731803.49 |
| amphibians | Sclerophrys_channingi | Amietophrynus_channingi | 0 | 332.23 | 4500 | 166636.39 |
| amphibians | Sclerophrys_dodsoni | Duttaphrynus_dodsoni | 0 | 24.93 | 238.5 | 863705.23 |
| amphibians | Sclerophrys_garmani | Amietophrynus_garmani | 0 | 163.99 | 16000 | 1754416.47 |
| amphibians | Sclerophrys_gutturalis | Amietophrynus_gutturalis | 0 | 188.23 | 20000 | 5933376.33 |
| amphibians | Sclerophrys_kassasii | Amietophrynus_kassasii | 0 | 4.93 | 300 | 40165.07 |
| amphibians | Sclerophrys_kisoloensis | Amietophrynus_kisoloensis | 0 | 66.4 | 2400 | 52751.39 |
| amphibians | Sclerophrys_lemairii | Amietophrynus_lemairii | 0 | 32.83 | 2500 | 1959972.77 |
| amphibians | Sclerophrys_maculata | Amietophrynus_maculatus | 0 | 61.58 | 5000 | 2329563.01 |
| amphibians | Sclerophrys_mauritanica | Amietophrynus_mauritanicus | 0 | 387.87 | 7500 | 776785.9 |
| amphibians | Sclerophrys_pantherina | Amietophrynus_pantherinus | 1 | 363.29 | 24738 | 679.82 |
| amphibians | Sclerophrys_pardalis | Amietophrynus_pardalis | 0 | 363.29 | 14000 | 27620.34 |
| amphibians | Sclerophrys_pentoni | Bufo_pentoni | 0 | 88.3 | 2600 | 3382939.44 |
| amphibians | Sclerophrys_poweri | Amietophrynus_poweri | 0 | 104.27 | 23000 | 1057179.84 |

| | | | | | | |
|---|---|---|---|---|---|---|
| amphibians | Sclerophrys_regularis | Amietophrynus_regularis | 0 | 243.97 | 12000 | 7462640 |
| amphibians | Sclerophrys_superciliaris | Amietophrynus_superciliaris | 0 | 507.74 | 4250 | 821127.39 |
| amphibians | Sclerophrys_tuberosa | Amietophrynus_tuberosus | 0 | 39.31 | 2700 | 864201.92 |
| amphibians | Sclerophrys_xeros | Amietophrynus_xeros | 0 | 94.47 | 4600 | 6286630.2 |
| amphibians | Scutiger_boulengeri | Scutiger_boulengeri | 0 | 29.46 | 685 | 704209.28 |
| amphibians | Scutiger_chintingensis | Scutiger_chintingensis | 1 | 13.17 | 147 | 5827.96 |
| amphibians | Scutiger_jiulongensis | Scutiger_jiulongensis | 1 | 53.74 | 990 | 2664.22 |
| amphibians | Scutiger_liupanensis | Scutiger_liupanensis | 1 | 19.39 | 350 | 2210.99 |
| amphibians | Scutiger_mammatus | Scutiger_mammatus | 0 | 51.84 | 577 | 288247.19 |
| amphibians | Scutiger_muliensis | Scutiger_muliensis | 1 | 50.6 | 203 | 2859.78 |
| amphibians | Scutiger_ningshanensis | Scutiger_ningshanensis | 0 | 11.77 | 149 | 12171.86 |
| amphibians | Scutiger_nyingchiensis | Scutiger_nyingchiensis | 0 | 39.65 | 1013 | 99061.44 |
| amphibians | Scutiger_pingwuensis | Scutiger_pingwuensis | 1 | 45.66 | 505 | 4280.82 |
| amphibians | Scutiger_sikimmensis | Scutiger_sikimmensis | 0 | 21.58 | 433.5 | 117217.27 |
| amphibians | Scutiger_tuberculatus | Scutiger_tuberculatus | 1 | 48.58 | 436 | 12890.18 |
| amphibians | Scutiger_wanglangensis | Scutiger_wanglangensis | 1 | 24.93 | 218 | 2970.69 |
| amphibians | Scythrophrys_sawayae | Scythrophrys_sawayae | 0 | 0.66 | 33 | 5108.92 |
| amphibians | Sechellophryne_gardineri | Sechellophryne_gardineri | 1 | 0.11 | 12 | 32.9 |
| amphibians | Semnodactylus_wealii | Semnodactylus_wealii | 0 | 7.29 | 300 | 376475.47 |
| amphibians | Silverstoneia_flotator | Silverstoneia_flotator | 0 | 0.43 | 6 | 57612.37 |
| amphibians | Silverstoneia_nubicola | Silverstoneia_nubicola | 1 | 0.89 | 6 | 67282.36 |
| amphibians | Siphonops_annulatus | Siphonops_annulatus | 0 | 59.36 | 10.5 | 4201157.22 |
| amphibians | Siphonops_paulensis | Siphonops_paulensis | 0 | 39.25 | 5 | 4627404.02 |
| amphibians | Siren_intermedia | Siren_intermedia | 0 | 205.63 | 783 | 1150933.99 |
| amphibians | Siren_lacertina | Siren_lacertina | 0 | 2111.76 | 750 | 418248.93 |
| amphibians | Smilisca_baudinii | Smilisca_baudinii | 0 | 74.11 | 1882.5 | 968474.9 |
| amphibians | Smilisca_cyanosticta | Smilisca_cyanosticta | 0 | 32.83 | 1107 | 26226.25 |
| amphibians | Smilisca_phaeota | Smilisca_phaeota | 0 | 52.68 | 1014 | 322059.06 |
| amphibians | Smilisca_puma | Smilisca_puma | 0 | 8.42 | 268 | 18050.36 |
| amphibians | Smilisca_sila | Smilisca_sila | 0 | 22.39 | 473 | 169790.18 |
| amphibians | Smilisca_sordida | Smilisca_sordida | 0 | 24.56 | 438 | 50630.82 |
| amphibians | Sooglossus_sechellensis | Sooglossus_sechellensis | 1 | 0.57 | 10.5 | 28.63 |
| amphibians | Spea_bombifrons | Spea_bombifrons | 0 | 24.56 | 2022 | 2525493.51 |
| amphibians | Spea_hammondii | Spea_hammondii | 0 | 41.05 | 1093 | 134297.68 |

| | | | | | | |
|---|---|---|---|---|---|---|
| amphibians | Spea_intermontana | Spea_intermontana | 0 | 25.82 | 593 | 1000665.47 |
| amphibians | Spea_multiplicata | Spea_multiplicata | 0 | 25.82 | 685 | 1909224.03 |
| amphibians | Speleomantes_ambrosii | Hydromantes_ambrosii | 1 | 5.04 | 11 | 393.04 |
| amphibians | Speleomantes_flavus | Hydromantes_flavus | 1 | 4.2 | 10 | 630.54 |
| amphibians | Speleomantes_genei | Hydromantes_genei | 1 | 3.91 | 10 | 1144.78 |
| amphibians | Speleomantes_imperialis | Hydromantes_imperialis | 0 | 5.85 | 8 | 2657.76 |
| amphibians | Speleomantes_italicus | Hydromantes_italicus | 1 | 5.97 | 11 | 21061.67 |
| amphibians | Speleomantes_sarrabusensis | Hydromantes_sarrabusensis | 1 | 7.38 | 10 | 103.13 |
| amphibians | Speleomantes_strinatii | Hydromantes_strinatii | 1 | 5.97 | 11.5 | 10276.41 |
| amphibians | Speleomantes_supramontis | Hydromantes_supramontis | 1 | 4.68 | 11 | 573.9 |
| amphibians | Sphaenorhynchus_carneus | Sphaenorhynchus_carneus | 0 | 0.89 | 147.5 | 1628057.29 |
| amphibians | Sphaenorhynchus_dorisae | Sphaenorhynchus_dorisae | 0 | 5.36 | 254 | 1408682.42 |
| amphibians | Sphaenorhynchus_lacteus | Sphaenorhynchus_lacteus | 0 | 9.67 | 298 | 4964958.56 |
| amphibians | Sphaerotheca_breviceps | Sphaerotheca_breviceps | 0 | 41.05 | 75 | 3248554.31 |
| amphibians | Sphaerotheca_rolandae | Sphaerotheca_rolandae | 0 | 11.04 | 743 | 707273.66 |
| amphibians | Sphenophryne_brevicrus | Oxydactyla_brevicrus | 0 | 1.54 | 14 | 2295.54 |
| amphibians | Sphenophryne_cornuta | Sphenophryne_cornuta | 0 | 6.27 | 17.5 | 470007.83 |
| amphibians | Sphenophryne_stenodactyla | Oxydactyla_stenodactyla | 0 | 2.6 | 23 | 1592.63 |
| amphibians | Spicospina_flammocaerulea | Spicospina_flammocaerulea | 1 | 3.81 | 124 | 331.13 |
| amphibians | Spinomantis_aglavei | Spinomantis_aglavei | 0 | 11.77 | 34 | 91736.52 |
| amphibians | Spinomantis_fimbriatus | Spinomantis_fimbriatus | 0 | 4.93 | 12 | 35060.69 |
| amphibians | Stefania_ayangannae | Stefania_ayangannae | 1 | 11.04 | 6.5 | 53.49 |
| amphibians | Stefania_coxi | Stefania_coxi | 1 | 32.83 | 6 | 26.6 |
| amphibians | Stefania_ginesi | Stefania_ginesi | 0 | 35.64 | 9.5 | 743.63 |
| amphibians | Stefania_goini | Stefania_goini | 0 | 82.42 | 15 | 73.49 |
| amphibians | Stefania_satelles | Stefania_satelles | 0 | 19.39 | 5 | 615.87 |
| amphibians | Stereochilus_marginatus | Stereochilus_marginatus | 0 | 3.11 | 63.5 | 150435.09 |
| amphibians | Strabomantis_anomalus | Strabomantis_anomalus | 1 | 80.71 | 69 | 86973.11 |
| amphibians | Strabomantis_biporcatus | Strabomantis_biporcatus | 0 | 104.27 | 45 | 4780.52 |
| amphibians | Strabomantis_sulcatus | Strabomantis_sulcatus | 0 | 19.6 | 33.5 | 1112766.69 |
| amphibians | Strauchbufo_raddei | Strauchbufo_raddei | 0 | 71.48 | 3500 | 4773678.42 |
| amphibians | Strongylopus_bonaespei | Strongylopus_bonaespei | 0 | 6.27 | 23 | 28830.12 |
| amphibians | Strongylopus_grayii | Strongylopus_grayii | 0 | 24.56 | 192.5 | 589930.75 |
| amphibians | Strongylopus_merumontanus | Strongylopus_merumontanus | 0 | 13.33 | 7.5 | 31759.22 |

| | | | | | | |
|---|---|---|---|---|---|---|
| amphibians | Strongylopus_wageri | Strongylopus_wageri | 0 | 10.34 | 185 | 50288.07 |
| amphibians | Stumpffia_grandis | Stumpffia_grandis | 0 | 1.17 | 20 | 23178.81 |
| amphibians | Stumpffia_pygmaea | Stumpffia_pygmaea | 1 | 0.12 | 9 | 311.21 |
| amphibians | Synapturanus_mirandaribeiroi | Synapturanus_mirandaribeiroi | 0 | 6.27 | 8.5 | 1189452.18 |
| amphibians | Synapturanus_rabus | Synapturanus_rabus | 0 | 0.48 | 3.5 | 309467.92 |
| amphibians | Synapturanus_salseri | Synapturanus_salseri | 0 | 3.47 | 6 | 411294.71 |
| amphibians | Tachycnemis_seychellensis | Tachycnemis_seychellensis | 0 | 42.85 | 300 | 180.37 |
| amphibians | Taricha_granulosa | Taricha_granulosa | 0 | 11.13 | 13.5 | 372204.67 |
| amphibians | Taricha_rivularis | Taricha_rivularis | 1 | 11.13 | 12 | 13931.34 |
| amphibians | Taricha_sierrae | Taricha_sierrae | 0 | 8.13 | 16.5 | 82830.55 |
| amphibians | Taricha_torosa | Taricha_torosa | 0 | 11.13 | 70 | 53483.46 |
| amphibians | Taruga_longinasus | Taruga_longinasus | 1 | 19.92 | 35 | 3233.5 |
| amphibians | Taudactylus_eungellensis | Taudactylus_eungellensis | 1 | 3.81 | 40 | 464.48 |
| amphibians | Taudactylus_liemi | Taudactylus_liemi | 0 | 1.89 | 56.5 | 768.54 |
| amphibians | Taudactylus_pleione | Taudactylus_pleione | 1 | 2.35 | 30.5 | 19.2 |
| amphibians | Taudactylus_rheophilus | Taudactylus_rheophilus | 1 | 2.35 | 42.5 | 724.82 |
| amphibians | Telmatobius_culeus | Telmatobius_culeus | 1 | 355.35 | 500 | 8452.4 |
| amphibians | Telmatobius_laticeps | Telmatobius_laticeps | 1 | 61.58 | 80 | 437.22 |
| amphibians | Telmatobius_niger | Telmatobius_niger | 1 | 45.85 | 446.5 | 12667.53 |
| amphibians | Telmatobufo_australis | Telmatobufo_australis | 0 | 44.71 | 100 | 27157.3 |
| amphibians | Telmatobufo_bullocki | Telmatobufo_bullocki | 1 | 57.01 | 112 | 2672.27 |
| amphibians | Telmatobufo_venustus | Telmatobufo_venustus | 1 | 32.83 | 111 | 16051.96 |
| amphibians | Teratohyla_midas | Teratohyla_midas | 0 | 1.5 | 63 | 3562726.29 |
| amphibians | Teratohyla_pulverata | Teratohyla_pulverata | 0 | 2.87 | 51.5 | 117386.03 |
| amphibians | Teratohyla_spinosa | Teratohyla_spinosa | 0 | 1.69 | 20 | 100062.16 |
| amphibians | Theloderma_laeve | Theloderma_laeve | 0 | 1.83 | 2.5 | 66346.99 |
| amphibians | Theloderma_ryabovi | Theloderma_ryabovi | 1 | 7.19 | 32 | 957.38 |
| amphibians | Thoropa_lutzi | Thoropa_lutzi | 1 | 2.11 | 149 | 1848.96 |
| amphibians | Thoropa_megatympanum | Thoropa_megatympanum | 0 | 13.33 | 67.5 | 27762.43 |
| amphibians | Thoropa_miliaris | Thoropa_miliaris | 0 | 52.68 | 516.5 | 485972.86 |
| amphibians | Thoropa_petropolitana | Thoropa_petropolitana | 1 | 1.5 | 24.46 | 1175.44 |
| amphibians | Thoropa_saxatilis | Thoropa_saxatilis | 0 | 29.89 | 33 | 12301.23 |
| amphibians | Thoropa_taophora | Thoropa_taophora | 0 | 111.53 | 793 | 7736.78 |
| amphibians | Tlalocohyla_loquax | Tlalocohyla_loquax | 0 | 9.03 | 134.5 | 348675.79 |

| | | | | | | |
|---|---|---|---|---|---|---|
| amphibians | Tlalocohyla_smithii | Tlalocohyla_smithii | 0 | 2.32 | 29 | 260450.69 |
| amphibians | Tomopterna_cryptotis | Tomopterna_cryptotis | 0 | 24.56 | 2500 | 9242511.75 |
| amphibians | Tomopterna_delalandii | Tomopterna_delalandii | 0 | 11.04 | 2500 | 216013.65 |
| amphibians | Tomopterna_krugerensis | Tomopterna_krugerensis | 0 | 17.85 | 5000 | 1212760 |
| amphibians | Tomopterna_marmorata | Tomopterna_marmorata | 0 | 15.03 | 150 | 1000987.26 |
| amphibians | Trachycephalus_coriaceus | Trachycephalus_coriaceus | 0 | 29.89 | 1325 | 3944543.13 |
| amphibians | Trachycephalus_cunauaru | Trachycephalus_cunauaru | 0 | 61.35 | 1050 | 2892478.97 |
| amphibians | Trachycephalus_mesophaeus | Trachycephalus_mesophaeus | 0 | 51.63 | 1740.5 | 327726.91 |
| amphibians | Trachycephalus_nigromaculatus | Trachycephalus_nigromaculatus | 0 | 79.58 | 270 | 936356.48 |
| amphibians | Trachycephalus_resinifictrix | Trachycephalus_resinifictrix | 0 | 104.27 | 1346 | 1369861.56 |
| amphibians | Trachycephalus_typhonius | Trachycephalus_venulosus | 0 | 159.41 | 5005.5 | 9757480.35 |
| amphibians | Triprion_petasatus | Triprion_petasatus | 0 | 41.41 | 1007.5 | 153656.14 |
| amphibians | Triprion_spatulatus | Diaglena_spatulata | 0 | 154.93 | 1500 | 73095.05 |
| amphibians | Triprion_spinosus | Anotheca_spinosa | 0 | 50.6 | 182.5 | 22780.67 |
| amphibians | Triturus_carnifex | Triturus_carnifex | 1 | 11.13 | 300 | 501475.03 |
| amphibians | Triturus_cristatus | Triturus_cristatus | 0 | 14.33 | 385 | 4456023.27 |
| amphibians | Triturus_dobrogicus | Triturus_dobrogicus | 0 | 11.5 | 225 | 271908.81 |
| amphibians | Triturus_karelinii | Triturus_karelinii | 0 | 11.5 | 200 | 242121.65 |
| amphibians | Triturus_marmoratus | Triturus_marmoratus | 1 | 9.72 | 325 | 474835.69 |
| amphibians | Triturus_pygmaeus | Triturus_pygmaeus | 0 | 3.68 | 274 | 174284.92 |
| amphibians | Tylototriton_asperrimus | Tylototriton_asperrimus | 0 | 4.67 | 41 | 111992.2 |
| amphibians | Tylototriton_hainanensis | Tylototriton_hainanensis | 1 | 3 | 74 | 438.66 |
| amphibians | Tylototriton_kweichowensis | Tylototriton_kweichowensis | 1 | 14.77 | 161 | 51079.42 |
| amphibians | Tylototriton_panhai | Tylototriton_panhai | 1 | 8.87 | 17.5 | 5385.56 |
| amphibians | Tylototriton_shanjing | Tylototriton_shanjing | 1 | 11.18 | 133.5 | 125306.09 |
| amphibians | Tylototriton_taliangensis | Tylototriton_taliangensis | 1 | 21.31 | 265 | 29860.18 |
| amphibians | Tylototriton_uyenoi | Tylototriton_uyenoi | 0 | 7.64 | 17.5 | 13094.23 |
| amphibians | Tylototriton_verrucosus | Tylototriton_verrucosus | 0 | 15.68 | 207.5 | 25986.71 |
| amphibians | Tylototriton_wenxianensis | Tylototriton_wenxianensis | 1 | 3.7 | 50.5 | 43334.53 |
| amphibians | Tylototriton_yangi | Tylototriton_yangi | 1 | 13.82 | 70 | 10452.15 |
| amphibians | Tylototriton_ziegleri | Tylototriton_ziegleri | 1 | 5.68 | 128.5 | 5257.54 |
| amphibians | Typhlonectes_compressicauda | Typhlonectes_compressicauda | 0 | 195.61 | 20 | 3464909.43 |
| amphibians | Uperodon_nagaoi | Ramanella_nagaoi | 1 | 2.47 | 20 | 3295.81 |
| amphibians | Uperoleia_altissima | Uperoleia_altissima | 0 | 1.89 | 123 | 19047.39 |

| | | | | | | |
|---|---|---|---|---|---|---|
| amphibians | Uperoleia_borealis | Uperoleia_borealis | 0 | 2.6 | 324 | 266876.47 |
| amphibians | Uperoleia_crassa | Uperoleia_crassa | 0 | 2.35 | 368 | 449339.88 |
| amphibians | Uperoleia_lithomoda | Uperoleia_lithomoda | 0 | 2.11 | 250.5 | 842374.26 |
| amphibians | Uperoleia_mimula | Uperoleia_mimula | 0 | 1.69 | 205 | 252574.46 |
| amphibians | Uperoleia_rugosa | Uperoleia_rugosa | 0 | 2.6 | 216.5 | 1344572.07 |
| amphibians | Uperoleia_talpa | Uperoleia_talpa | 0 | 6.27 | 1468.5 | 70839.76 |
| amphibians | Uraeotyphlus_bombayensis | Ichthyophis_bombayensis | 0 | 130.7 | 102 | 111814.53 |
| amphibians | Urspelerpes_brucei | Urspelerpes_brucei | 0 | 0.28 | 10 | 2944.31 |
| amphibians | Vandijkophrynus_angusticeps | Vandijkophrynus_angusticeps | 0 | 17.85 | 1825 | 76440.89 |
| amphibians | Vandijkophrynus_robinsoni | Vandijkophrynus_robinsoni | 0 | 29.89 | 2000 | 61138.68 |
| amphibians | Vitreorana_castroviejoi | Vitreorana_castroviejoi | 1 | 1.18 | 31 | 97.25 |
| amphibians | Vitreorana_gorzulae | Vitreorana_gorzulae | 0 | 0.83 | 18.5 | 23059.11 |
| amphibians | Vitreorana_ritae | Cochranella_ritae | 0 | 1.02 | 16 | 3428488.12 |
| amphibians | Vitreorana_uranoscopa | Vitreorana_uranoscopa | 0 | 1.5 | 30.5 | 613709.59 |
| amphibians | Walkerana_leptodactyla | Indirana_leptodactyla | 1 | 7.84 | 30 | 4472.59 |
| amphibians | Werneria_bambutensis | Werneria_bambutensis | 1 | 6.47 | 431.5 | 1578.36 |
| amphibians | Werneria_tandyi | Werneria_tandyi | 1 | 5.89 | 629 | 231.26 |
| amphibians | Wolterstorffina_mirei | Wolterstorffina_mirei | 1 | 4.93 | 65 | 147.26 |
| amphibians | Wolterstorffina_parvipalmata | Wolterstorffina_parvipalmata | 1 | 4.93 | 219 | 2466.48 |
| amphibians | Xenohyla_truncata | Xenohyla_truncata | 1 | 11.04 | 186 | 1925.56 |
| amphibians | Xenophrys_glandulosa | Xenophrys_glandulosa | 0 | 104.27 | 954 | 82333.78 |
| amphibians | Xenopus_gilli | Xenopus_gilli | 1 | 21.02 | 450 | 749.92 |
| amphibians | Xenopus_laevis | Xenopus_laevis | 0 | 363.29 | 13640 | 3812401.03 |
| amphibians | Xenopus_muelleri | Xenopus_muelleri | 0 | 74.11 | 436 | 3295891.82 |
| amphibians | Xenorhina_arboricola | Xenorhina_arboricola | 0 | 8.42 | 11 | 1200.28 |
| amphibians | Zhangixalus_arboreus | Rhacophorus_arboreus | 0 | 54.82 | 482.5 | 162840.26 |
| amphibians | Zhangixalus_dennysi | Rhacophorus_dennysi | 0 | 233.19 | 2625 | 922599.08 |
| amphibians | Zhangixalus_dugritei | Rhacophorus_dugritei | 0 | 27.13 | 400 | 105966.23 |
| amphibians | Zhangixalus_moltrechti | Rhacophorus_moltrechti | 0 | 10.89 | 240 | 24134.25 |
| amphibians | Zhangixalus_omeimontis | Rhacophorus_omeimontis | 0 | 50.6 | 543 | 167218.05 |
| amphibians | Zhangixalus_schlegelii | Rhacophorus_schlegelii | 0 | 23.58 | 370 | 286417.98 |
| amphibians | Zhangixalus_taipeianus | Rhacophorus_taipeianus | 1 | 15.03 | 350 | 3030.64 |
| lizards | Ablepharus_bivittatus | Ablepharus_bivittatus | 0 | 4.37 | 4 | 1124383.73 |
| lizards | Ablepharus_budaki | Ablepharus_budaki | 0 | 2 | 3.5 | 121185.75 |

| | | | | | | |
|---|---|---|---|---|---|---|
| lizards | Ablepharus_chernovi | Ablepharus_chernovi | 0 | 2.95 | 3.5 | 296009.76 |
| lizards | Ablepharus_deserti | Ablepharus_deserti | 0 | 3.89 | 4.5 | 889020.94 |
| lizards | Ablepharus_grayanus | Ablepharus_grayanus | 0 | 0.72 | 1.5 | 230787.61 |
| lizards | Ablepharus_kitaibelii | Ablepharus_kitaibelii | 0 | 4.79 | 2.8 | 713068.09 |
| lizards | Ablepharus_lindbergi | Ablepharus_lindbergi | 0 | 2.29 | 4.5 | 165909.25 |
| lizards | Ablepharus_pannonicus | Ablepharus_pannonicus | 0 | 3.09 | 4 | 2664129.08 |
| lizards | Ablepharus_rueppellii | Ablepharus_rueppellii | 0 | 2.63 | 1.8 | 46688.69 |
| lizards | Abronia_aurita | Abronia_aurita | 1 | 33.88 | 12 | 4835.25 |
| lizards | Abronia_campbelli | Abronia_campbelli | 1 | 36.31 | 8 | 124.21 |
| lizards | Abronia_deppii | Abronia_deppii | 1 | 30.2 | 6 | 4814.26 |
| lizards | Abronia_fimbriata | Abronia_fimbriata | 1 | 38.9 | 4 | 3088.5 |
| lizards | Abronia_graminea | Abronia_graminea | 1 | 22.39 | 5.03 | 8507.05 |
| lizards | Abronia_lythrochila | Abronia_lythrochila | 0 | 29.51 | 9.5 | 8425.63 |
| lizards | Abronia_martindelcampoi | Abronia_martindelcampoi | 1 | 25.7 | 4 | 1896.42 |
| lizards | Abronia_meledona | Abronia_meledona | 1 | 29.51 | 14 | 19.98 |
| lizards | Abronia_mixteca | Abronia_mixteca | 1 | 61.66 | 7 | 7624.97 |
| lizards | Abronia_oaxacae | Abronia_oaxacae | 1 | 26.92 | 1 | 6695.11 |
| lizards | Abronia_smithi | Abronia_smithi | 0 | 21.88 | 3.5 | 1998.08 |
| lizards | Abronia_taeniata | Abronia_taeniata | 1 | 47.86 | 6.4 | 19138.7 |
| lizards | Acanthocercus_adramitanus | Acanthocercus_adramitanus | 0 | 134.9 | 8 | 236389.56 |
| lizards | Acanthocercus_atricollis | Acanthocercus_atricollis | 0 | 177.83 | 11.36 | 6072021.84 |
| lizards | Acanthocercus_cyanogaster | Acanthocercus_cyanogaster | 0 | 165.96 | 11 | 2095557.48 |
| lizards | Acanthodactylus_bedriagai | Acanthodactylus_bedriagai | 0 | 10.47 | 5 | 192655.87 |
| lizards | Acanthodactylus_blanci | Acanthodactylus_blanci | 1 | 19.5 | 5 | 129571.9 |
| lizards | Acanthodactylus_blanfordii | Acanthodactylus_blanfordii | 0 | 9.77 | 4 | 934334.52 |
| lizards | Acanthodactylus_boskianus | Acanthodactylus_boskianus | 0 | 19.5 | 4.3 | 12634604.8 |
| lizards | Acanthodactylus_cantoris | Acanthodactylus_cantoris | 0 | 19.05 | 5.5 | 1416266.46 |
| lizards | Acanthodactylus_dumerilii | Acanthodactylus_dumerilii | 0 | 7.24 | 1.5 | 2618778.63 |
| lizards | Acanthodactylus_erythrurus | Acanthodactylus_erythrurus | 0 | 13.8 | 3.7 | 1030722.73 |
| lizards | Acanthodactylus_longipes | Acanthodactylus_longipes | 0 | 6.46 | 2.7 | 7831763.02 |
| lizards | Acanthodactylus_maculatus | Acanthodactylus_maculatus | 0 | 5.62 | 3.33 | 1193367.15 |
| lizards | Acanthodactylus_opheodurus | Acanthodactylus_opheodurus | 0 | 5.62 | 4 | 1708910.23 |
| lizards | Acanthodactylus_pardalis | Acanthodactylus_pardalis | 1 | 10.47 | 3.65 | 178962.19 |
| lizards | Acanthodactylus_savignyi | Acanthodactylus_savignyi | 0 | 9.77 | 5 | 515134.89 |

| | | | | | | |
|---|---|---|---|---|---|---|
| lizards | Acanthodactylus_schmidti | Acanthodactylus_schmidti | 0 | 26.3 | 3 | 2034614.22 |
| lizards | Acanthodactylus_schreiberi | Acanthodactylus_schreiberi | 1 | 18.62 | 3 | 9272.66 |
| lizards | Acanthodactylus_scutellatus | Acanthodactylus_scutellatus | 0 | 14.13 | 2.9 | 8730741.2 |
| lizards | Acanthodactylus_spinicauda | Acanthodactylus_spinicauda | 1 | 4.17 | 4 | 365.62 |
| lizards | Acanthodactylus_tristrami | Acanthodactylus_tristrami | 0 | 19.5 | 4 | 76447.33 |
| lizards | Acanthosaura_armata | Acanthosaura_armata | 0 | 95.5 | 14.5 | 194795.93 |
| lizards | Acanthosaura_capra | Acanthosaura_capra | 0 | 95.5 | 19.5 | 11351.15 |
| lizards | Acanthosaura_cardamomensis | Acanthosaura_cardamomensis | 0 | 117.49 | 20.5 | 10121.35 |
| lizards | Acanthosaura_crucigera | Acanthosaura_crucigera | 0 | 128.82 | 14 | 234223.5 |
| lizards | Acanthosaura_lepidogaster | Acanthosaura_lepidogaster | 0 | 45.71 | 13 | 2548799.74 |
| lizards | Acanthosaura_nataliae | Acanthosaura_nataliae | 0 | 138.04 | 13 | 56149.66 |
| lizards | Acontias_breviceps | Acontias_breviceps | 0 | 12.02 | 2 | 46456.68 |
| lizards | Acontias_gariepensis | Acontias_gariepensis | 0 | 5.37 | 1 | 51588.12 |
| lizards | Acontias_gracilicauda | Acontias_gracilicauda | 0 | 22.39 | 2 | 314474.12 |
| lizards | Acontias_jappi | Acontias_jappi | 0 | 10.47 | 4 | 26592.96 |
| lizards | Acontias_kgalagadi | Acontias_kgalagadi | 0 | 10.23 | 1.48 | 643796.4 |
| lizards | Acontias_lineatus | Acontias_lineatus | 0 | 6.31 | 1.57 | 241502.54 |
| lizards | Acontias_meleagris | Acontias_meleagris | 0 | 29.51 | 3 | 147662.82 |
| lizards | Acontias_namaquensis | Acontias_namaquensis | 0 | 16.6 | 2.5 | 23239.75 |
| lizards | Acontias_occidentalis | Acontias_occidentalis | 0 | 20.89 | 1.8 | 581586.74 |
| lizards | Acontias_orientalis | Acontias_orientalis | 0 | 12.88 | 4 | 54785.57 |
| lizards | Acontias_percivali | Acontias_percivali | 0 | 21.88 | 2 | 20300.67 |
| lizards | Acontias_plumbeus | Acontias_plumbeus | 0 | 100 | 8 | 316954.37 |
| lizards | Acontias_richardi | Acontias_richardi | 0 | 4.79 | 2 | 7211.42 |
| lizards | Acontias_rieppeli | Acontias_rieppeli | 1 | 10.23 | 2 | 13797.84 |
| lizards | Acratosaura_mentalis | Acratosaura_mentalis | 0 | 4.68 | 1.6 | 603401.12 |
| lizards | Acritoscincus_duperreyi | Bassiana_duperreyi | 0 | 10.47 | 5.3 | 413900.12 |
| lizards | Acritoscincus_platynotus | Bassiana_platynota | 0 | 10.47 | 5.25 | 336960.16 |
| lizards | Acritoscincus_trilineatus | Bassiana_trilineata | 0 | 10.47 | 5.9 | 319903.68 |
| lizards | Adolfus_alleni | Adolfus_alleni | 0 | 7.94 | 2 | 13638.53 |
| lizards | Adolfus_jacksoni | Adolfus_jacksoni | 0 | 16.6 | 3.5 | 389063.45 |
| lizards | Aeluroscalabotes_felinus | Aeluroscalabotes_felinus | 0 | 38.9 | 2 | 367109.59 |
| lizards | Afroedura_africana | Afroedura_africana | 0 | 5.5 | 2 | 57106.4 |
| lizards | Afroedura_amatolica | Afroedura_amatolica | 0 | 4.68 | 2 | 6918.45 |

| | | | | | | |
|---|---|---|---|---|---|---|
| lizards | Afroedura_bogerti | Afroedura_bogerti | 0 | 3.31 | 1.84 | 67804.63 |
| lizards | Afroedura_broadleyi | Afroedura_broadleyi | 0 | 3.8 | 2 | 325.44 |
| lizards | Afroedura_halli | Afroedura_halli | 0 | 5.62 | 2 | 27850.98 |
| lizards | Afroedura_hawequensis | Afroedura_hawequensis | 0 | 11.48 | 2 | 12444.67 |
| lizards | Afroedura_karroica | Afroedura_karroica | 0 | 5.5 | 2 | 21238.28 |
| lizards | Afroedura_langi | Afroedura_langi | 0 | 2.24 | 2 | 7590.26 |
| lizards | Afroedura_leoloensis | Afroedura_leoloensis | 0 | 1.58 | 2 | 880.63 |
| lizards | Afroedura_loveridgei | Afroedura_loveridgei | 0 | 4.47 | 2 | 19936.08 |
| lizards | Afroedura_major | Afroedura_major | 0 | 8.91 | 2 | 5165.43 |
| lizards | Afroedura_maripi | Afroedura_maripi | 0 | 5.37 | 2 | 2466.79 |
| lizards | Afroedura_marleyi | Afroedura_marleyi | 0 | 1.12 | 2 | 26508.94 |
| lizards | Afroedura_nivaria | Afroedura_nivaria | 0 | 6.03 | 2 | 19248.31 |
| lizards | Afroedura_pienaari | Afroedura_pienaari | 0 | 3.31 | 2 | 678.38 |
| lizards | Afroedura_pondolia | Afroedura_pondolia | 0 | 4.27 | 2 | 42402.43 |
| lizards | Afroedura_transvaalica | Afroedura_transvaalica | 0 | 7.94 | 1.5 | 359996.92 |
| lizards | Afrogecko_porphyreus | Afrogecko_porphyreus | 0 | 2.95 | 2 | 86912.03 |
| lizards | Agama_aculeata | Agama_aculeata | 0 | 56.23 | 12.75 | 4075015.48 |
| lizards | Agama_africana | Agama_africana | 0 | 58.88 | 7.55 | 1181582.68 |
| lizards | Agama_agama | Agama_agama | 0 | 117.49 | 8.8 | 3768296.52 |
| lizards | Agama_anchietae | Agama_anchietae | 0 | 95.5 | 11 | 875209.89 |
| lizards | Agama_armata | Agama_armata | 0 | 27.54 | 14.5 | 2843147.34 |
| lizards | Agama_atra | Agama_atra | 0 | 95.5 | 12.5 | 1292509.15 |
| lizards | Agama_boensis | Agama_boensis | 0 | 17.38 | 7.55 | 135764.88 |
| lizards | Agama_boueti | Agama_boueti | 0 | 44.67 | 9.3 | 2366175.35 |
| lizards | Agama_caudospinosa | Agama_caudospinosa | 0 | 158.49 | 7.5 | 47508.76 |
| lizards | Agama_doriae | Agama_doriae | 0 | 51.29 | 7.55 | 2796459.65 |
| lizards | Agama_etoshae | Agama_etoshae | 0 | 19.95 | 9.5 | 70851.95 |
| lizards | Agama_hispida | Agama_hispida | 0 | 83.18 | 13.7 | 286034.83 |
| lizards | Agama_kaimosae | Agama_kaimosae | 0 | 104.71 | 7.55 | 32480.06 |
| lizards | Agama_kirkii | Agama_kirkii | 0 | 51.29 | 10 | 948984.73 |
| lizards | Agama_lionotus | Agama_lionotus | 0 | 107.15 | 8.88 | 1013804.82 |
| lizards | Agama_montana | Agama_montana | 1 | 27.54 | 8 | 14081.05 |
| lizards | Agama_mossambica | Agama_mossambica | 0 | 58.88 | 9.5 | 658704.46 |
| lizards | Agama_mwanzae | Agama_mwanzae | 0 | 144.54 | 7.5 | 118000.27 |

| | | | | | | |
|---|---|---|---|---|---|---|
| lizards | Agama_planiceps | Agama_planiceps | 0 | 112.2 | 6.5 | 778036.55 |
| lizards | Agama_spinosa | Agama_spinosa | 0 | 69.18 | 7.55 | 1244916.3 |
| lizards | Agama_turuensis | Agama_turuensis | 0 | 95.5 | 7.5 | 22165.61 |
| lizards | Agama_weidholzi | Agama_weidholzi | 0 | 8.71 | 10 | 437786.78 |
| lizards | Agamura_persica | Agamura_persica | 0 | 9.33 | 2 | 1509180.46 |
| lizards | Ailuronyx_seychellensis | Ailuronyx_seychellensis | 0 | 28.84 | 1.5 | 232.57 |
| lizards | Ailuronyx_tachyscopaeus | Ailuronyx_tachyscopaeus | 0 | 12.3 | 1.5 | 234.77 |
| lizards | Algyroides_fitzingeri | Algyroides_fitzingeri | 0 | 2.19 | 2.5 | 32877.9 |
| lizards | Algyroides_moreoticus | Algyroides_moreoticus | 0 | 3.02 | 3.3 | 22700.81 |
| lizards | Algyroides_nigropunctatus | Algyroides_nigropunctatus | 0 | 7.94 | 3.5 | 85186.67 |
| lizards | Alinea_lanceolata | Alinea_lanceolata | 1 | 17.38 | 7 | 19.98 |
| lizards | Alopoglossus_angulatus | Alopoglossus_angulatus | 0 | 6.17 | 2 | 4274943.15 |
| lizards | Alopoglossus_atriventris | Alopoglossus_atriventris | 0 | 3.72 | 2 | 1315927.39 |
| lizards | Alopoglossus_buckleyi | Alopoglossus_buckleyi | 0 | 5.62 | 2 | 812923.34 |
| lizards | Alopoglossus_copii | Alopoglossus_copii | 0 | 12.88 | 2 | 144233.43 |
| lizards | Alsophylax_laevis | Alsophylax_laevis | 1 | 1.29 | 1.5 | 193212.44 |
| lizards | Alsophylax_loricatus | Alsophylax_loricatus | 1 | 0.87 | 1.5 | 128811.35 |
| lizards | Alsophylax_pipiens | Alsophylax_pipiens | 0 | 1.7 | 1.5 | 2852234.86 |
| lizards | Alsophylax_przewalskii | Alsophylax_przewalskii | 0 | 1.1 | 1.5 | 692562.46 |
| lizards | Alsophylax_szczerbaki | Alsophylax_szczerbaki | 1 | 0.87 | 1.5 | 64182.07 |
| lizards | Alsophylax_tadjikiensis | Alsophylax_tadjikiensis | 1 | 0.76 | 1.5 | 68343.24 |
| lizards | Altiphylax_levitoni | Altiphylax_levitoni | 0 | 2.09 | 1.5 | 25938.06 |
| lizards | Altiphylax_tokobajevi | Altiphylax_tokobajevi | 0 | 2.75 | 1.6 | 6465.81 |
| lizards | Amalosia_jacovae | Amalosia_jacovae | 0 | 4.68 | 2 | 72413.47 |
| lizards | Amalosia_lesueurii | Amalosia_lesueurii | 0 | 10.23 | 2 | 52067.63 |
| lizards | Amalosia_obscura | Amalosia_obscura | 0 | 4.79 | 2 | 28246.58 |
| lizards | Amalosia_rhombifer | Amalosia_rhombifer | 0 | 10.23 | 2 | 1984790.55 |
| lizards | Amapasaurus_tetradactylus | Amapasaurus_tetradactylus | 0 | 0.91 | 2 | 72339.45 |
| lizards | Amblyrhynchus_cristatus | Amblyrhynchus_cristatus | 1 | 7413.1 | 2.25 | 7990.36 |
| lizards | Ameiva_ameiva | Ameiva_ameiva | 0 | 302 | 5.25 | 12706264.26 |
| lizards | Ameiva_atrigularis | Ameiva_atrigularis | 0 | 218.78 | 4.72 | 15371.23 |
| lizards | Ameiva_bifrontata | Ameiva_bifrontata | 0 | 64.57 | 1.88 | 1615919.57 |
| lizards | Ameiva_concolor | Ameiva_concolor | 0 | 64.57 | 1.98 | 565.23 |
| lizards | Ameiva_parecis | Ameiva_parecis | 1 | 21.38 | 1.64 | 1505.57 |

| | | | | | | |
|---|---|---|---|---|---|---|
| lizards | Ameiva_praesignis | Ameiva_praesignis | 0 | 467.74 | 4.72 | 240603.03 |
| lizards | Ameivula_jalapensis | Ameivula_jalapensis | 0 | 5.5 | 1 | 11017.85 |
| lizards | Ameivula_mumbuca | Ameivula_mumbuca | 0 | 5.75 | 1 | 49042.12 |
| lizards | Ameivula_nativo | Ameivula_nativo | 1 | 9.55 | 2.2 | 35581.12 |
| lizards | Ameivula_ocellifera | Ameivula_ocellifera | 0 | 52.48 | 2.45 | 7006546.86 |
| lizards | Amphibolurus_burnsi | Lophognathus_burnsi | 0 | 85.11 | 7.5 | 816579.8 |
| lizards | Amphibolurus_muricatus | Amphibolurus_muricatus | 0 | 66.07 | 5.55 | 678727.05 |
| lizards | Amphibolurus_norrisi | Amphibolurus_norrisi | 0 | 54.95 | 4.8 | 264247.62 |
| lizards | Amphiglossus_astrolabi | Amphiglossus_astrolabi | 0 | 302 | 10 | 128986.63 |
| lizards | Amphiglossus_reticulatus | Amphiglossus_reticulatus | 0 | 275.42 | 3 | 75187.82 |
| lizards | Amphisbaena_alba | Amphisbaena_alba | 0 | 1023.29 | 9.25 | 13781841.24 |
| lizards | Amphisbaena_angustifrons | Amphisbaena_angustifrons | 0 | 33.11 | 2.5 | 1244295.49 |
| lizards | Amphisbaena_brasiliana | Amphisbaena_brasiliana | 0 | 23.44 | 9 | 35472.21 |
| lizards | Amphisbaena_cuiabana | Amphisbaena_cuiabana | 0 | 23.44 | 2 | 31624.4 |
| lizards | Amphisbaena_darwinii | Amphisbaena_darwini | 0 | 39.81 | 5 | 3114258.54 |
| lizards | Amphisbaena_dubia | Amphisbaena_dubia | 0 | 13.49 | 3.5 | 150218.38 |
| lizards | Amphisbaena_innocens | Amphisbaena_innocens | 1 | 12.02 | 2 | 11628.63 |
| lizards | Amphisbaena_kingii | Amphisbaena_kingii | 0 | 7.94 | 2.6 | 1567066.85 |
| lizards | Amphisbaena_mertensii | Amphisbaena_mertensii | 0 | 66.07 | 7 | 816727.3 |
| lizards | Amphisbaena_munoai | Amphisbaena_munoai | 0 | 2.04 | 1.8 | 295193.48 |
| lizards | Amphisbaena_nigricauda | Amphisbaena_nigricauda | 1 | 1.86 | 1.5 | 436.39 |
| lizards | Amphisbaena_roberti | Amphisbaena_roberti | 0 | 12.02 | 4 | 556159.68 |
| lizards | Amphisbaena_vermicularis | Amphisbaena_vermicularis | 0 | 26.3 | 4 | 1727813.56 |
| lizards | Anadia_bogotensis | Anadia_bogotensis | 0 | 7.24 | 2 | 23378.48 |
| lizards | Anadia_brevifrontalis | Anadia_brevifrontalis | 1 | 25.7 | 2 | 2502.95 |
| lizards | Anadia_hobarti | Anadia_hobarti | 0 | 16.6 | 2 | 9.99 |
| lizards | Anadia_rhombifera | Anadia_rhombifera | 0 | 8.13 | 1.5 | 148126.68 |
| lizards | Anatololacerta_anatolica | Anatololacerta_anatolica | 0 | 9.77 | 5.5 | 78409.31 |
| lizards | Anatololacerta_danfordi | Anatololacerta_danfordi | 0 | 9.77 | 5.5 | 56425.42 |
| lizards | Andinosaura_oculata | Riama_oculata | 1 | 17.38 | 2.5 | 4971.34 |
| lizards | Andinosaura_vespertina | Riama_vespertina | 1 | 7.41 | 2 | 9387.9 |
| lizards | Anepischetosia_maccoyi | Anepischetosia_maccoyi | 0 | 4.17 | 2.9 | 179855.28 |
| lizards | Anguis_cephallonica | Anguis_cephallonica | 0 | 75.86 | 10.5 | 22293.59 |
| lizards | Anguis_fragilis | Anguis_fragilis | 0 | 79.43 | 9.48 | 2517204.35 |

| | | | | | | |
|---|---|---|---|---|---|---|
| lizards | Anniella_geronimensis | Anniella_geronimensis | 1 | 7.76 | 1 | 2929.91 |
| lizards | Anniella_pulchra | Anniella_pulchra | 0 | 9.55 | 1.65 | 135562.36 |
| lizards | Anolis_acutus | Anolis_acutus | 0 | 6.31 | 1 | 193.13 |
| lizards | Anolis_aeneus | Anolis_aeneus | 0 | 10.47 | 1 | 421.6 |
| lizards | Anolis_aequatorialis | Anolis_aequatorialis | 0 | 34.67 | 1 | 22401.36 |
| lizards | Anolis_agassizi | Anolis_agassizi | 0 | 41.69 | 1 | 7.66 |
| lizards | Anolis_aliniger | Anolis_aliniger | 0 | 4.57 | 1 | 23061.52 |
| lizards | Anolis_allisoni | Anolis_allisoni | 0 | 20.42 | 1 | 64972.75 |
| lizards | Anolis_allogus | Anolis_allogus | 0 | 5.25 | 1.5 | 77077.83 |
| lizards | Anolis_altitudinalis | Anolis_altitudinalis | 0 | 2.95 | 1 | 16.61 |
| lizards | Anolis_alumina | Anolis_alumina | 0 | 1.38 | 1 | 3921.83 |
| lizards | Anolis_alutaceus | Anolis_alutaceus | 0 | 2.34 | 1 | 57368.13 |
| lizards | Anolis_angusticeps | Anolis_angusticeps | 0 | 3.16 | 1 | 100327.62 |
| lizards | Anolis_anisolepis | Anolis_anisolepis | 0 | 3.72 | 1 | 4345.91 |
| lizards | Anolis_aquaticus | Anolis_aquaticus | 0 | 9.55 | 1.13 | 11011.77 |
| lizards | Anolis_argenteolus | Anolis_argenteolus | 0 | 5.75 | 1.05 | 32510.03 |
| lizards | Anolis_armouri | Anolis_armouri | 1 | 8.51 | 1 | 4997.5 |
| lizards | Anolis_auratus | Anolis_auratus | 0 | 3.89 | 1 | 3453866.79 |
| lizards | Anolis_bahorucoensis | Anolis_bahorucoensis | 1 | 2.82 | 1.05 | 2842.23 |
| lizards | Anolis_baleatus | Anolis_baleatus | 0 | 114.82 | 1 | 23168.3 |
| lizards | Anolis_baracoae | Anolis_baracoae | 0 | 100 | 1.5 | 1052.44 |
| lizards | Anolis_barahonae | Anolis_barahonae | 1 | 100 | 1.08 | 1713.68 |
| lizards | Anolis_barbatus | Anolis_barbatus | 0 | 97.72 | 1 | 537.65 |
| lizards | Anolis_barbouri | Anolis_barbouri | 1 | 3.55 | 1 | 7797.02 |
| lizards | Anolis_barkeri | Anolis_barkeri | 1 | 20.89 | 1.5 | 36541.56 |
| lizards | Anolis_bartschi | Anolis_bartschi | 0 | 10.47 | 1.5 | 1318.99 |
| lizards | Anolis_beckeri | Anolis_beckeri | 0 | 6.61 | 1 | 180350.65 |
| lizards | Anolis_bicaorum | Anolis_bicaorum | 1 | 13.18 | 1 | 42.08 |
| lizards | Anolis_bimaculatus | Anolis_bimaculatus | 0 | 39.81 | 1 | 298.22 |
| lizards | Anolis_biporcatus | Anolis_biporcatus | 0 | 30.9 | 1 | 650952.07 |
| lizards | Anolis_bitectus | Anolis_bitectus | 1 | 3.8 | 1 | 15453.42 |
| lizards | Anolis_blanquillanus | Anolis_blanquillanus | 0 | 12.59 | 1 | 65.31 |
| lizards | Anolis_bombiceps | Anolis_bombiceps | 0 | 8.51 | 1.5 | 371682.21 |
| lizards | Anolis_bonairensis | Anolis_bonairensis | 1 | 8.71 | 1 | 244.26 |

| | | | | | | |
|---|---|---|---|---|---|---|
| lizards | Anolis_brasiliensis | Anolis_brasiliensis | 0 | 8.71 | 1 | 1239579.56 |
| lizards | Anolis_bremeri | Anolis_bremeri | 0 | 7.76 | 1 | 5260.16 |
| lizards | Anolis_brevirostris | Anolis_brevirostris | 0 | 3.16 | 1 | 15142.29 |
| lizards | Anolis_calimae | Anolis_calimae | 1 | 4.37 | 1 | 5315.41 |
| lizards | Anolis_capito | Anolis_capito | 0 | 21.88 | 1 | 278977.93 |
| lizards | Anolis_carolinensis | Anolis_carolinensis | 0 | 8.71 | 1.15 | 1275289.21 |
| lizards | Anolis_carpenteri | Anolis_carpenteri | 0 | 2.24 | 1 | 39659.49 |
| lizards | Anolis_centralis | Anolis_centralis | 0 | 2.34 | 1 | 40134.29 |
| lizards | Anolis_chamaeleonides | Anolis_chamaeleonides | 0 | 109.65 | 1 | 18057.05 |
| lizards | Anolis_chloris | Anolis_chloris | 0 | 5.01 | 1 | 99568.49 |
| lizards | Anolis_chlorocyanus | Anolis_chlorocyanus | 0 | 10.47 | 1 | 61926.91 |
| lizards | Anolis_christophei | Anolis_christophei | 0 | 2.63 | 1 | 9856.01 |
| lizards | Anolis_chrysolepis | Anolis_chrysolepis | 0 | 8.51 | 1 | 5296831.93 |
| lizards | Anolis_chrysops | Anolis_chrysops | 1 | 8.13 | 2 | 1.12 |
| lizards | Anolis_clivicola | Anolis_clivicola | 0 | 2.57 | 1 | 1153.76 |
| lizards | Anolis_concolor | Anolis_concolor | 0 | 10.47 | 1 | 51.49 |
| lizards | Anolis_conspersus | Anolis_conspersus | 0 | 9.12 | 1 | 203.03 |
| lizards | Anolis_cooki | Anolis_cooki | 1 | 8.13 | 1 | 258.4 |
| lizards | Anolis_crassulus | Anolis_crassulus | 0 | 4.37 | 1 | 51762.53 |
| lizards | Anolis_cristatellus | Anolis_cristatellus | 0 | 10.47 | 1 | 9147.17 |
| lizards | Anolis_cupreus | Anolis_cupreus | 0 | 3.89 | 1 | 150610.19 |
| lizards | Anolis_cuprinus | Anolis_cuprinus | 0 | 6.92 | 1 | 7965.57 |
| lizards | Anolis_cuvieri | Anolis_cuvieri | 0 | 114.82 | 1 | 3890.57 |
| lizards | Anolis_cybotes | Anolis_cybotes | 0 | 10.96 | 1 | 70110.56 |
| lizards | Anolis_desiradei | Anolis_desiradei | 0 | 10.47 | 2 | 23.04 |
| lizards | Anolis_distichus | Anolis_distichus | 0 | 3.16 | 1 | 8963.13 |
| lizards | Anolis_dollfusianus | Anolis_dollfusianus | 0 | 1.82 | 1 | 2134.48 |
| lizards | Anolis_dunni | Anolis_dunni | 0 | 4.17 | 1 | 26633.78 |
| lizards | Anolis_equestris | Anolis_equestris | 0 | 134.9 | 1 | 60238.86 |
| lizards | Anolis_ernestwilliamsi | Anolis_ernestwilliamsi | 1 | 11.48 | 1 | 5.89 |
| lizards | Anolis_etheridgei | Anolis_etheridgei | 1 | 1.82 | 1 | 1555.78 |
| lizards | Anolis_eugenegrahami | Anolis_eugenegrahami | 1 | 7.76 | 1 | 66.21 |
| lizards | Anolis_evermanni | Anolis_evermanni | 0 | 9.77 | 1 | 3511.44 |
| lizards | Anolis_extremus | Anolis_extremus | 0 | 12.59 | 1.5 | 439.35 |

| | | | | | | |
|---|---|---|---|---|---|---|
| lizards | Anolis_fasciatus | Anolis_fasciatus | 0 | 7.76 | 1 | 39215.85 |
| lizards | Anolis_festae | Anolis_festae | 0 | 4.79 | 1 | 46452.96 |
| lizards | Anolis_fowleri | Anolis_fowleri | 1 | 9.77 | 1 | 148.31 |
| lizards | Anolis_frenatus | Anolis_frenatus | 0 | 69.18 | 1 | 175594.62 |
| lizards | Anolis_fuscoauratus | Anolis_fuscoauratus | 0 | 3.16 | 1 | 9192893.9 |
| lizards | Anolis_gadovii | Anolis_gadovii | 0 | 10.47 | 1.5 | 3092.48 |
| lizards | Anolis_gaigei | Anolis_gaigei | 0 | 3.09 | 1 | 63716.66 |
| lizards | Anolis_garmani | Anolis_garmani | 0 | 52.48 | 1 | 9880.47 |
| lizards | Anolis_gemmosus | Anolis_gemmosus | 0 | 6.03 | 1 | 21456.83 |
| lizards | Anolis_gingivinus | Anolis_gingivinus | 0 | 7.76 | 1 | 200.33 |
| lizards | Anolis_grahami | Anolis_grahami | 0 | 8.71 | 1 | 11021.54 |
| lizards | Anolis_griseus | Anolis_griseus | 0 | 50.12 | 1.02 | 324.73 |
| lizards | Anolis_gundlachi | Anolis_gundlachi | 0 | 9.12 | 1 | 3942.5 |
| lizards | Anolis_heterodermus | Anolis_heterodermus | 0 | 13.18 | 1 | 94840.93 |
| lizards | Anolis_homolechis | Anolis_homolechis | 0 | 7.08 | 1 | 88748 |
| lizards | Anolis_humilis | Anolis_humilis | 0 | 2.63 | 1.07 | 72019.78 |
| lizards | Anolis_insolitus | Anolis_insolitus | 1 | 2.24 | 1 | 677.75 |
| lizards | Anolis_isolepis | Anolis_isolepis | 0 | 1.58 | 1 | 13567.02 |
| lizards | Anolis_jacare | Anolis_jacare | 0 | 8.51 | 1 | 31227.58 |
| lizards | Anolis_johnmeyeri | Anolis_johnmeyeri | 1 | 8.13 | 1 | 300.87 |
| lizards | Anolis_jubar | Anolis_jubar | 0 | 5.01 | 1 | 44811.35 |
| lizards | Anolis_kahouannensis | Anolis_kahouannensis | 0 | 9.12 | 2 | 0.34 |
| lizards | Anolis_kemptoni | Anolis_kemptoni | 0 | 4.57 | 1 | 743.2 |
| lizards | Anolis_krugi | Anolis_krugi | 0 | 3.55 | 1 | 5077.09 |
| lizards | Anolis_laeviventris | Anolis_laeviventris | 0 | 6.03 | 1 | 166442.11 |
| lizards | Anolis_leachii | Anolis_leachii | 0 | 37.15 | 2 | 322.47 |
| lizards | Anolis_lemurinus | Anolis_lemurinus | 0 | 10.23 | 1 | 569991.69 |
| lizards | Anolis_limifrons | Anolis_limifrons | 0 | 2.82 | 1.5 | 208940.29 |
| lizards | Anolis_lineatopus | Anolis_lineatopus | 0 | 8.13 | 1 | 10925.39 |
| lizards | Anolis_liogaster | Anolis_liogaster | 0 | 6.03 | 2 | 4209.27 |
| lizards | Anolis_lionotus | Anolis_lionotus | 0 | 9.77 | 1 | 55747.18 |
| lizards | Anolis_litoralis | Anolis_litoralis | 0 | 2.34 | 1 | 7433.59 |
| lizards | Anolis_lividus | Anolis_lividus | 0 | 11.48 | 1 | 96.57 |
| lizards | Anolis_longiceps | Anolis_longiceps | 0 | 11.75 | 1 | 4.62 |

| | | | | | | |
|---|---|---|---|---|---|---|
| lizards | Anolis_longitibialis | Anolis_longitibialis | 1 | 7.76 | 1.5 | 1259.24 |
| lizards | Anolis_loveridgei | Anolis_loveridgei | 1 | 33.11 | 1 | 3203.25 |
| lizards | Anolis_loysiana | Anolis_loysiana | 0 | 2.95 | 2 | 5524.16 |
| lizards | Anolis_luciae | Anolis_luciae | 1 | 15.49 | 1 | 617.4 |
| lizards | Anolis_lucius | Anolis_lucius | 0 | 7.08 | 1 | 36088.1 |
| lizards | Anolis_luteogularis | Anolis_luteogularis | 0 | 138.04 | 1 | 20864.65 |
| lizards | Anolis_lyra | Anolis_lyra | 0 | 9.33 | 1 | 86778.8 |
| lizards | Anolis_macrinii | Anolis_macrinii | 0 | 18.2 | 1 | 166.26 |
| lizards | Anolis_maculiventris | Anolis_maculiventris | 0 | 2.63 | 1 | 166970.54 |
| lizards | Anolis_mariarum | Anolis_mariarum | 0 | 3.72 | 1 | 34597.68 |
| lizards | Anolis_marmoratus | Anolis_marmoratus | 0 | 12.59 | 1 | 1510.93 |
| lizards | Anolis_marron | Anolis_marron | 0 | 9.12 | 1 | 354.79 |
| lizards | Anolis_matudai | Anolis_matudai | 0 | 3.55 | 1.5 | 11656.57 |
| lizards | Anolis_megapholidotus | Anolis_megapholidotus | 0 | 3.16 | 1 | 6213.57 |
| lizards | Anolis_menta | Anolis_menta | 0 | 3.72 | 1.5 | 9.99 |
| lizards | Anolis_meridionalis | Anolis_meridionalis | 0 | 4.37 | 1.2 | 3299660.69 |
| lizards | Anolis_mestrei | Anolis_mestrei | 0 | 4.57 | 1 | 3354.04 |
| lizards | Anolis_microlepidotus | Anolis_microlepidotus | 0 | 3.31 | 1.5 | 17303.27 |
| lizards | Anolis_naufragus | Anolis_naufragus | 1 | 4.57 | 2 | 8292.83 |
| lizards | Anolis_nebuloides | Anolis_nebuloides | 0 | 3.63 | 1 | 36060.5 |
| lizards | Anolis_nebulosus | Anolis_nebulosus | 0 | 4.37 | 1 | 575815.86 |
| lizards | Anolis_nelsoni | Anolis_nelsoni | 1 | 9.12 | 1.5 | 1.85 |
| lizards | Anolis_noblei | Anolis_noblei | 0 | 134.9 | 1 | 11591.65 |
| lizards | Anolis_oculatus | Anolis_oculatus | 0 | 19.05 | 1.05 | 766.69 |
| lizards | Anolis_olssoni | Anolis_olssoni | 0 | 3.16 | 1.5 | 46744.66 |
| lizards | Anolis_omiltemanus | Anolis_omiltemanus | 0 | 2.34 | 2 | 2889.2 |
| lizards | Anolis_onca | Anolis_onca | 0 | 15.85 | 1 | 25740.67 |
| lizards | Anolis_opalinus | Anolis_opalinus | 0 | 3.72 | 1 | 10293.32 |
| lizards | Anolis_ophiolepis | Anolis_ophiolepis | 0 | 1.35 | 1 | 14915.44 |
| lizards | Anolis_orcesi | Anolis_orcesi | 0 | 6.31 | 1 | 7097.06 |
| lizards | Anolis_ortonii | Anolis_ortonii | 0 | 4.57 | 1 | 5357221.04 |
| lizards | Anolis_otongae | Anolis_otongae | 1 | 6.31 | 1.5 | 214.85 |
| lizards | Anolis_oxylophus | Anolis_oxylophus | 0 | 12.59 | 1 | 118926.1 |
| lizards | Anolis_pachypus | Anolis_pachypus | 0 | 3.31 | 1 | 11879.36 |

| | | | | | | |
|---|---|---|---|---|---|---|
| lizards | Anolis_parvicirculatus | Anolis_parvicirculatus | 0 | 3.02 | 1 | 1246.58 |
| lizards | Anolis_pentaprion | Anolis_pentaprion | 0 | 10.47 | 1 | 355371.96 |
| lizards | Anolis_peraccae | Anolis_peraccae | 0 | 5.25 | 1 | 44318.89 |
| lizards | Anolis_petersii | Anolis_petersii | 0 | 54.95 | 1 | 34767.38 |
| lizards | Anolis_pigmaequestris | Anolis_pigmaequestris | 1 | 54.95 | 1 | 32.14 |
| lizards | Anolis_pinchoti | Anolis_pinchoti | 0 | 3.89 | 1 | 23.53 |
| lizards | Anolis_placidus | Anolis_placidus | 1 | 2.09 | 1 | 265.9 |
| lizards | Anolis_planiceps | Anolis_planiceps | 0 | 9.12 | 1.44 | 1206913.45 |
| lizards | Anolis_podocarpus | Anolis_podocarpus | 1 | 18.2 | 1 | 321.64 |
| lizards | Anolis_poecilopus | Anolis_poecilopus | 0 | 8.51 | 1 | 89834.99 |
| lizards | Anolis_pogus | Anolis_pogus | 0 | 4.07 | 1 | 192.76 |
| lizards | Anolis_polylepis | Anolis_polylepis | 0 | 4.37 | 1 | 12841.87 |
| lizards | Anolis_porcatus | Anolis_porcatus | 0 | 17.78 | 1 | 94983.29 |
| lizards | Anolis_porcus | Anolis_porcus | 0 | 100 | 1 | 13537.83 |
| lizards | Anolis_proboscis | Anolis_proboscis | 1 | 12.02 | 2 | 107.01 |
| lizards | Anolis_pulchellus | Anolis_pulchellus | 0 | 2.88 | 1 | 8714.06 |
| lizards | Anolis_punctatus | Anolis_punctatus | 0 | 15.85 | 1.25 | 7465476.2 |
| lizards | Anolis_quaggulus | Anolis_quaggulus | 0 | 1.78 | 1 | 130421.11 |
| lizards | Anolis_reconditus | Anolis_reconditus | 1 | 20.42 | 1 | 120.64 |
| lizards | Anolis_rejectus | Anolis_rejectus | 1 | 1.1 | 1 | 358.44 |
| lizards | Anolis_richardii | Anolis_richardii | 0 | 54.95 | 1 | 377.09 |
| lizards | Anolis_rodriguezii | Anolis_rodriguezi | 0 | 2.63 | 1 | 272090.6 |
| lizards | Anolis_roquet | Anolis_roquet | 0 | 13.18 | 1 | 1141.64 |
| lizards | Anolis_rubribarbaris | Anolis_rubribarbaris | 1 | 2.95 | 1 | 112.84 |
| lizards | Anolis_sabanus | Anolis_sabanus | 0 | 6.92 | 1 | 12.49 |
| lizards | Anolis_sagrei | Anolis_sagrei | 0 | 8.71 | 1 | 225473.67 |
| lizards | Anolis_saxatilis | Anolis_whitemani | 0 | 6.31 | 1 | 19236.28 |
| lizards | Anolis_schwartzi | Anolis_schwartzi | 0 | 2.51 | 1 | 230.59 |
| lizards | Anolis_scypheus | Anolis_scypheus | 0 | 13.18 | 1 | 843487.13 |
| lizards | Anolis_semilineatus | Anolis_semilineatus | 0 | 2.24 | 1 | 72703.32 |
| lizards | Anolis_sericeus | Anolis_sericeus | 0 | 3.16 | 1 | 655922.23 |
| lizards | Anolis_sheplani | Anolis_sheplani | 1 | 1.48 | 1 | 216.97 |
| lizards | Anolis_shrevei | Anolis_shrevei | 0 | 4.57 | 1 | 1408.13 |
| lizards | Anolis_singularis | Anolis_singularis | 1 | 2.95 | 1 | 9.99 |

| | | | | | | |
|---|---|---|---|---|---|---|
| lizards | Anolis_smallwoodi | Anolis_smallwoodi | 0 | 134.9 | 1 | 4402.32 |
| lizards | Anolis_smaragdinus | Anolis_smaragdinus | 0 | 5.5 | 1.3 | 2713.05 |
| lizards | Anolis_sminthus | Anolis_sminthus | 1 | 4.37 | 1 | 13721.82 |
| lizards | Anolis_spectrum | Anolis_spectrum | 1 | 1.58 | 1 | 11960.2 |
| lizards | Anolis_strahmi | Anolis_strahmi | 1 | 10.47 | 1 | 709.04 |
| lizards | Anolis_tandai | Anolis_tandai | 0 | 8.51 | 1 | 1400788.68 |
| lizards | Anolis_taylori | Anolis_taylori | 0 | 9.77 | 1 | 80.24 |
| lizards | Anolis_terraealtae | Anolis_terraealtae | 0 | 10.47 | 2 | 16.94 |
| lizards | Anolis_tetarii | Anolis_tetarii | 0 | 13.18 | 1 | 198.12 |
| lizards | Anolis_tolimensis | Anolis_tolimensis | 0 | 4.57 | 1.14 | 48671.29 |
| lizards | Anolis_trachyderma | Anolis_trachyderma | 0 | 4.79 | 1.5 | 1001940.69 |
| lizards | Anolis_transversalis | Anolis_transversalis | 0 | 19.05 | 1 | 2234134.98 |
| lizards | Anolis_trinitatis | Anolis_trinitatis | 0 | 10.47 | 1 | 348.24 |
| lizards | Anolis_tropidogaster | Anolis_tropidogaster | 0 | 5.25 | 1 | 422645.87 |
| lizards | Anolis_tropidolepis | Anolis_tropidolepis | 0 | 4.37 | 1 | 3965.11 |
| lizards | Anolis_tropidonotus | Anolis_tropidonotus | 0 | 5.75 | 1 | 359474.77 |
| lizards | Anolis_uniformis | Anolis_uniformis | 0 | 1.48 | 1 | 84815.2 |
| lizards | Anolis_utilensis | Anolis_utilensis | 1 | 5.01 | 1 | 42.08 |
| lizards | Anolis_valencienni | Anolis_valencienni | 0 | 13.18 | 1 | 11021.54 |
| lizards | Anolis_ventrimaculatus | Anolis_ventrimaculatus | 0 | 10.47 | 1 | 44750.81 |
| lizards | Anolis_vermiculatus | Anolis_vermiculatus | 0 | 48.98 | 1.5 | 2375.37 |
| lizards | Anolis_vittigerus | Anolis_vittigerus | 0 | 7.76 | 1 | 83506.2 |
| lizards | Anolis_wattsii | Anolis_wattsi | 0 | 4.07 | 1 | 716.38 |
| lizards | Anolis_woodi | Anolis_woodi | 0 | 20.42 | 1 | 11087.04 |
| lizards | Anomalopus_leuckartii | Anomalopus_leuckartii | 0 | 11.75 | 3.65 | 397667.28 |
| lizards | Anomalopus_swansoni | Anomalopus_swansoni | 0 | 2.88 | 2.5 | 18131.02 |
| lizards | Anomalopus_verreauxii | Anomalopus_verreauxi | 0 | 25.12 | 8 | 344288.5 |
| lizards | Anotosaura_vanzolinia | Anotosaura_vanzolinia | 0 | 2.69 | 1.94 | 45489.94 |
| lizards | Apathya_cappadocica | Apathya_cappadocica | 0 | 14.45 | 5 | 336629.59 |
| lizards | Aphaniotis_acutirostris | Aphaniotis_acutirostris | 0 | 12.02 | 2 | 63335.48 |
| lizards | Aphaniotis_fusca | Aphaniotis_fusca | 0 | 12.59 | 1 | 1322523.81 |
| lizards | Aphaniotis_ornata | Aphaniotis_ornata | 0 | 5.89 | 2 | 26163.78 |
| lizards | Aprasia_aurita | Aprasia_aurita | 0 | 6.76 | 2 | 72135.19 |
| lizards | Aprasia_haroldi | Aprasia_haroldi | 0 | 5.75 | 2 | 1629.5 |

| | | | | | | |
|---|---|---|---|---|---|---|
| lizards | Aprasia_inaurita | Aprasia_inaurita | 0 | 7.76 | 2 | 386166.53 |
| lizards | Aprasia_parapulchella | Aprasia_parapulchella | 0 | 8.32 | 2 | 152289.55 |
| lizards | Aprasia_pseudopulchella | Aprasia_pseudopulchella | 0 | 9.12 | 2 | 40407.36 |
| lizards | Aprasia_pulchella | Aprasia_pulchella | 0 | 7.41 | 2 | 65114.58 |
| lizards | Aprasia_repens | Aprasia_repens | 0 | 6.92 | 2 | 156382.46 |
| lizards | Aprasia_rostrata | Aprasia_rostrata | 1 | 6.92 | 2 | 12573.47 |
| lizards | Aprasia_striolata | Aprasia_striolata | 0 | 8.13 | 2 | 207815.33 |
| lizards | Archaeolacerta_bedriagae | Archaeolacerta_bedriagae | 0 | 14.13 | 4.5 | 12496.03 |
| lizards | Archaius_tigris | Archaius_tigris | 1 | 22.91 | 7.5 | 94.75 |
| lizards | Aristelliger_cochranae | Aristelliger_cochranae | 0 | 8.71 | 1 | 35091.31 |
| lizards | Aristelliger_expectatus | Aristelliger_expectatus | 0 | 4.07 | 1 | 32623.99 |
| lizards | Aristelliger_georgeensis | Aristelliger_georgeensis | 0 | 36.31 | 1 | 2198.57 |
| lizards | Aristelliger_hechti | Aristelliger_hechti | 1 | 17.78 | 1 | 152.34 |
| lizards | Aristelliger_lar | Aristelliger_lar | 0 | 79.43 | 1 | 44003.84 |
| lizards | Aristelliger_praesignis | Aristelliger_praesignis | 0 | 24.55 | 1 | 10815.1 |
| lizards | Aristelliger_reyesi | Aristelliger_reyesi | 1 | 6.31 | 1 | 17.71 |
| lizards | Arthrosaura_kockii | Arthrosaura_kockii | 0 | 8.51 | 2 | 780816.7 |
| lizards | Arthrosaura_reticulata | Arthrosaura_reticulata | 0 | 8.51 | 2 | 4621507.22 |
| lizards | Asaccus_caudivolvulus | Asaccus_caudivolvulus | 1 | 6.61 | 1 | 125.16 |
| lizards | Asaccus_elisae | Asaccus_elisae | 0 | 7.24 | 1.5 | 329259.55 |
| lizards | Asaccus_gallagheri | Asaccus_gallagheri | 0 | 1.74 | 1 | 14296.83 |
| lizards | Asaccus_nasrullahi | Asaccus_nasrullahi | 0 | 8.91 | 1.33 | 5601.9 |
| lizards | Asaccus_platyrhynchus | Asaccus_platyrhynchus | 0 | 6.61 | 1 | 5911.07 |
| lizards | Aspidoscelis_burti | Aspidoscelis_burti | 0 | 85.11 | 4.33 | 165629.67 |
| lizards | Aspidoscelis_calidipes | Aspidoscelis_calidipes | 0 | 14.13 | 2.33 | 8198.39 |
| lizards | Aspidoscelis_ceralbensis | Aspidoscelis_ceralbensis | 0 | 25.12 | 2.5 | 142.91 |
| lizards | Aspidoscelis_communis | Aspidoscelis_communis | 0 | 109.65 | 5.7 | 99380.57 |
| lizards | Aspidoscelis_costatus | Aspidoscelis_costata | 0 | 70.79 | 5.05 | 331734.79 |
| lizards | Aspidoscelis_cozumela | Aspidoscelis_cozumelae | 0 | 16.6 | 1.75 | 488.27 |
| lizards | Aspidoscelis_danheimae | Aspidoscelis_danheimae | 0 | 8.13 | 2.5 | 169.31 |
| lizards | Aspidoscelis_deppii | Aspidoscelis_deppei | 0 | 23.44 | 2.3 | 609448.25 |
| lizards | Aspidoscelis_exsanguis | Aspidoscelis_exsanguis | 0 | 36.31 | 3.55 | 409465.47 |
| lizards | Aspidoscelis_gularis | Aspidoscelis_gularis | 0 | 45.71 | 3.69 | 1417047.37 |
| lizards | Aspidoscelis_guttatus | Aspidoscelis_guttata | 0 | 93.33 | 4.5 | 169406.92 |

| | | | | | | |
|---|---|---|---|---|---|---|
| lizards | Aspidoscelis_hyperythrus | Aspidoscelis_hyperythra | 0 | 10.72 | 2.32 | 140729.2 |
| lizards | Aspidoscelis_inornatus | Aspidoscelis_inornata | 0 | 18.62 | 2.65 | 659625.45 |
| lizards | Aspidoscelis_labialis | Aspidoscelis_labialis | 1 | 7.08 | 2.26 | 13910.99 |
| lizards | Aspidoscelis_laredoensis | Aspidoscelis_laredoensis | 0 | 21.38 | 3.4 | 40046.35 |
| lizards | Aspidoscelis_lineattissimus | Aspidoscelis_lineattissima | 0 | 42.66 | 3.65 | 65053.4 |
| lizards | Aspidoscelis_martyris | Cnemidophorus_martyris | 1 | 14.79 | 1.25 | 1.22 |
| lizards | Aspidoscelis_motaguae | Aspidoscelis_motaguae | 0 | 93.33 | 2.5 | 42636.38 |
| lizards | Aspidoscelis_neomexicanus | Aspidoscelis_neomexicana | 0 | 18.62 | 1.95 | 91254.63 |
| lizards | Aspidoscelis_neotesselatus | Aspidoscelis_neotesselata | 0 | 36.31 | 2.94 | 124729.53 |
| lizards | Aspidoscelis_parvisocius | Aspidoscelis_parvisocia | 0 | 14.13 | 1.78 | 3556.21 |
| lizards | Aspidoscelis_pictus | Aspidoscelis_picta | 0 | 10.23 | 2.5 | 18.1 |
| lizards | Aspidoscelis_rodecki | Aspidoscelis_rodecki | 0 | 12.59 | 2.25 | 223.59 |
| lizards | Aspidoscelis_sackii | Aspidoscelis_sackii | 0 | 112.2 | 6.07 | 46103.86 |
| lizards | Aspidoscelis_sexlineatus | Aspidoscelis_sexlineata | 0 | 22.39 | 3.7 | 2757748.86 |
| lizards | Aspidoscelis_sonorae | Aspidoscelis_sonorae | 0 | 30.9 | 3.9 | 183626.98 |
| lizards | Aspidoscelis_stictogrammus | Aspidoscelis_stictogramma | 0 | 85.11 | 6.5 | 85889.54 |
| lizards | Aspidoscelis_tesselatus | Aspidoscelis_tesselata | 0 | 36.31 | 4.05 | 339352.99 |
| lizards | Aspidoscelis_tigris | Aspidoscelis_tigris | 0 | 79.43 | 2.7 | 2084184.45 |
| lizards | Aspidoscelis_uniparens | Aspidoscelis_uniparens | 0 | 18.62 | 3.02 | 218191.05 |
| lizards | Aspidoscelis_velox | Aspidoscelis_velox | 0 | 20.89 | 4 | 369953.79 |
| lizards | Aspidoscelis_xanthonotus | Aspidoscelis_xanthonota | 0 | 44.67 | 4 | 17223.34 |
| lizards | Aspronema_cochabambae | Aspronema_cochabambae | 1 | 11.48 | 4.5 | 3676.24 |
| lizards | Aspronema_dorsivittatum | Aspronema_dorsivittatum | 0 | 15.49 | 4.25 | 3779794.07 |
| lizards | Ateuchosaurus_chinensis | Ateuchosaurus_chinensis | 0 | 21.38 | 2.3 | 822446.79 |
| lizards | Ateuchosaurus_pellopleurus | Ateuchosaurus_pellopleurus | 0 | 6.46 | 4.5 | 2708.82 |
| lizards | Atlantolacerta_andreanskyi | Atlantolacerta_andreanskyi | 0 | 3.89 | 1.85 | 11816.94 |
| lizards | Aurivela_tergolaevigata | Aurivela_tergolaevigata | 0 | 7.76 | 2 | 114706.82 |
| lizards | Australolacerta_australis | Australolacerta_australis | 0 | 7.94 | 2 | 20866.44 |
| lizards | Bachia_bicolor | Bachia_bicolor | 0 | 2.95 | 2 | 156298.96 |
| lizards | Bachia_bresslaui | Bachia_bresslaui | 1 | 6.03 | 2 | 1123654.73 |
| lizards | Bachia_dorbignyi | Bachia_dorbignyi | 0 | 3.16 | 2 | 835376.68 |
| lizards | Bachia_flavescens | Bachia_flavescens | 0 | 3.16 | 1 | 3587470.01 |
| lizards | Bachia_heteropa | Bachia_heteropa | 0 | 1.82 | 2 | 83731.22 |
| lizards | Bachia_trisanale | Bachia_trisanale | 0 | 1.45 | 2 | 1489602.66 |

| | | | | | | |
|---|---|---|---|---|---|---|
| lizards | Barisia_herrerae | Barisia_herrerae | 1 | 33.88 | 6 | 743.64 |
| lizards | Barisia_imbricata | Barisia_imbricata | 0 | 77.62 | 11.45 | 485129.42 |
| lizards | Barisia_rudicollis | Barisia_rudicollis | 1 | 36.31 | 5 | 1072.25 |
| lizards | Basiliscus_basiliscus | Basiliscus_basiliscus | 0 | 549.54 | 9.2 | 432691.01 |
| lizards | Basiliscus_galeritus | Basiliscus_galeritus | 0 | 489.78 | 7.5 | 323161.64 |
| lizards | Basiliscus_plumifrons | Basiliscus_plumifrons | 0 | 549.54 | 11 | 107491.89 |
| lizards | Basiliscus_vittatus | Basiliscus_vittatus | 0 | 398.11 | 5.4 | 880749.99 |
| lizards | Bavayia_crassicollis | Bavayia_crassicollis | 1 | 12.88 | 2 | 1979.64 |
| lizards | Bavayia_cyclura | Bavayia_cyclura | 1 | 7.41 | 2 | 17439.17 |
| lizards | Bavayia_exsuccida | Bavayia_exsuccida | 1 | 2.04 | 2 | 1000.1 |
| lizards | Bavayia_geitaina | Bavayia_geitaina | 0 | 7.41 | 2 | 3641.36 |
| lizards | Bavayia_goroensis | Bavayia_goroensis | 1 | 2.24 | 2 | 146.48 |
| lizards | Bavayia_montana | Bavayia_montana | 1 | 8.91 | 1.5 | 10593.66 |
| lizards | Bavayia_nubila | Bavayia_nubila | 1 | 6.03 | 2 | 95.38 |
| lizards | Bavayia_ornata | Bavayia_ornata | 1 | 6.61 | 2 | 270.04 |
| lizards | Bavayia_pulchella | Bavayia_pulchella | 1 | 2.34 | 2 | 254.1 |
| lizards | Bavayia_robusta | Bavayia_robusta | 0 | 11.48 | 2 | 3095.75 |
| lizards | Bavayia_sauvagii | Bavayia_sauvagii | 1 | 4.79 | 2 | 16396.16 |
| lizards | Bavayia_septuiclavis | Bavayia_septuiclavis | 0 | 2.45 | 2 | 2941.22 |
| lizards | Bellatorias_frerei | Bellatorias_frerei | 0 | 234.42 | 3 | 460068.12 |
| lizards | Bellatorias_major | Bellatorias_major | 0 | 1778.28 | 5.35 | 72332.58 |
| lizards | Bipes_biporus | Bipes_biporus | 0 | 14.13 | 2.08 | 42660.69 |
| lizards | Bipes_canaliculatus | Bipes_canaliculatus | 0 | 14.13 | 2.8 | 9816.83 |
| lizards | Bipes_tridactylus | Bipes_tridactylus | 0 | 4.27 | 2.2 | 91.16 |
| lizards | Blaesodactylus_antongilensis | Blaesodactylus_antongilensis | 0 | 17.38 | 2 | 33912.21 |
| lizards | Blaesodactylus_boivini | Blaesodactylus_boivini | 1 | 64.57 | 2 | 9379.96 |
| lizards | Blanus_cinereus | Blanus_cinereus | 0 | 16.6 | 2 | 47115.08 |
| lizards | Blanus_mettetali | Blanus_mettetali | 0 | 9.55 | 1 | 111004.25 |
| lizards | Blanus_strauchi | Blanus_strauchi | 0 | 9.77 | 1.5 | 88278.28 |
| lizards | Brachylophus_bulabula | Brachylophus_bulabula | 1 | 676.08 | 3.5 | 15793.11 |
| lizards | Brachylophus_fasciatus | Brachylophus_fasciatus | 1 | 676.08 | 3.89 | 12789.31 |
| lizards | Brachylophus_vitiensis | Brachylophus_vitiensis | 1 | 707.95 | 4 | 1562.71 |
| lizards | Brachymeles_apus | Brachymeles_apus | 0 | 4.57 | 4 | 166.53 |
| lizards | Brachymeles_bonitae | Brachymeles_bonitae | 0 | 4.68 | 1.5 | 13683.25 |

| | | | | | | |
|---|---|---|---|---|---|---|
| lizards | Brachymeles_boulengeri | Brachymeles_boulengeri | 0 | 26.92 | 3.1 | 156558.46 |
| lizards | Brachymeles_cebuensis | Brachymeles_cebuensis | 1 | 2.63 | 2 | 785.02 |
| lizards | Brachymeles_gracilis | Brachymeles_gracilis | 0 | 31.62 | 2.5 | 122838.1 |
| lizards | Brachymeles_hilong | Brachymeles_hilong | 0 | 11.22 | 2.5 | 14796.09 |
| lizards | Brachymeles_lukbani | Brachymeles_lukbani | 0 | 1.86 | 2 | 9.99 |
| lizards | Brachymeles_muntingkamay | Brachymeles_muntingkamay | 0 | 3.24 | 2 | 8080.54 |
| lizards | Brachymeles_orientalis | Brachymeles_orientalis | 0 | 33.88 | 3 | 10861.21 |
| lizards | Brachymeles_pathfinderi | Brachymeles_pathfinderi | 0 | 2.14 | 2 | 17543.39 |
| lizards | Brachymeles_samarensis | Brachymeles_samarensis | 0 | 1.95 | 1.5 | 24628.39 |
| lizards | Brachymeles_schadenbergi | Brachymeles_schadenbergi | 0 | 38.02 | 2.5 | 118956.83 |
| lizards | Brachymeles_talinis | Brachymeles_talinis | 0 | 79.43 | 4.5 | 39075.48 |
| lizards | Brachymeles_taylori | Brachymeles_taylori | 0 | 25.12 | 2.5 | 17449 |
| lizards | Brachymeles_tridactylus | Brachymeles_tridactylus | 0 | 3.55 | 2 | 24396.93 |
| lizards | Brachyseps_frontoparietalis | Amphiglossus_frontoparietalis | 0 | 8.91 | 3 | 58116.39 |
| lizards | Brachyseps_punctatus | Amphiglossus_punctatus | 0 | 7.76 | 2 | 54731.14 |
| lizards | Bradypodion_atromontanum | Bradypodion_atromontanum | 0 | 10.72 | 7 | 792.88 |
| lizards | Bradypodion_caeruleogula | Bradypodion_caeruleogula | 1 | 7.94 | 5.5 | 119.14 |
| lizards | Bradypodion_damaranum | Bradypodion_damaranum | 0 | 13.8 | 11.5 | 2059.78 |
| lizards | Bradypodion_dracomontanum | Bradypodion_dracomontanum | 0 | 13.18 | 11 | 3664.57 |
| lizards | Bradypodion_gutturale | Bradypodion_gutturale | 0 | 14.79 | 15 | 45758.49 |
| lizards | Bradypodion_kentanicum | Bradypodion_kentanicum | 1 | 7.94 | 5 | 4262.03 |
| lizards | Bradypodion_melanocephalum | Bradypodion_melanocephalum | 0 | 10.96 | 14.5 | 21790.41 |
| lizards | Bradypodion_occidentale | Bradypodion_occidentale | 0 | 20.89 | 13 | 34988.04 |
| lizards | Bradypodion_pumilum | Bradypodion_pumilum | 0 | 25.12 | 12.5 | 27711.52 |
| lizards | Bradypodion_setaroi | Bradypodion_setaroi | 0 | 9.55 | 8.5 | 8036.76 |
| lizards | Bradypodion_taeniabronchum | Bradypodion_taeniabronchum | 0 | 9.12 | 12 | 6644.59 |
| lizards | Bradypodion_thamnobates | Bradypodion_thamnobates | 1 | 25.12 | 19 | 17251.73 |
| lizards | Bradypodion_transvaalense | Bradypodion_transvaalense | 0 | 15.49 | 12 | 45421.13 |
| lizards | Bradypodion_ventrale | Bradypodion_ventrale | 0 | 24.55 | 13 | 242128.84 |
| lizards | Brasiliscincus_agilis | Brasiliscincus_agilis | 0 | 19.05 | 4.5 | 479715.01 |
| lizards | Brasiliscincus_caissara | Brasiliscincus_caissara | 1 | 15.49 | 4.68 | 694.13 |
| lizards | Brasiliscincus_heathi | Brasiliscincus_heathi | 0 | 16.6 | 4 | 1824206.95 |
| lizards | Broadleysaurus_major | Broadleysaurus_major | 0 | 389.05 | 5 | 3762454.51 |
| lizards | Bronchocela_cristatella | Bronchocela_cristatella | 0 | 75.86 | 2 | 1963684.21 |

| | | | | | | |
|---|---|---|---|---|---|---|
| lizards | Bronchocela_hayeki | Bronchocela_hayeki | 0 | 58.88 | 2 | 2470.99 |
| lizards | Bronchocela_jubata | Bronchocela_jubata | 0 | 117.49 | 2 | 618236.18 |
| lizards | Bronchocela_marmorata | Bronchocela_marmorata | 0 | 66.07 | 2 | 276676.4 |
| lizards | Bronchocela_smaragdina | Bronchocela_smaragdina | 0 | 58.88 | 3 | 31053.28 |
| lizards | Brookesia_antakarana | Brookesia_antakarana | 0 | 6.03 | 4.5 | 45.97 |
| lizards | Brookesia_brygooi | Brookesia_brygooi | 0 | 3.98 | 3.5 | 146282.97 |
| lizards | Brookesia_decaryi | Brookesia_decaryi | 1 | 4.17 | 3.5 | 1967.45 |
| lizards | Brookesia_desperata | Brookesia_desperata | 1 | 0.91 | 2 | 50.05 |
| lizards | Brookesia_exarmata | Brookesia_exarmata | 1 | 0.66 | 2 | 1727.58 |
| lizards | Brookesia_griveaudi | Brookesia_griveaudi | 0 | 6.92 | 3.5 | 21829.16 |
| lizards | Brookesia_minima | Brookesia_minima | 1 | 0.56 | 2 | 7342.83 |
| lizards | Brookesia_stumpffi | Brookesia_stumpffi | 0 | 5.13 | 3.5 | 61920.45 |
| lizards | Brookesia_superciliaris | Brookesia_superciliaris | 0 | 4.47 | 3.5 | 90360.09 |
| lizards | Brookesia_thieli | Brookesia_thieli | 0 | 2.69 | 3 | 46977.04 |
| lizards | Brookesia_tristis | Brookesia_tristis | 1 | 0.49 | 2 | 215.3 |
| lizards | Bunopus_tuberculatus | Bunopus_tuberculatus | 0 | 5.13 | 2 | 3918133.93 |
| lizards | Cachryx_defensor | Ctenosaura_defensor | 1 | 162.18 | 5 | 45697.64 |
| lizards | Cadea_blanoides | Cadea_blanoides | 0 | 17.78 | 2 | 11054.5 |
| lizards | Caesoris_novaecaledoniae | Lioscincus_novaecaledoniae | 0 | 6.17 | 4 | 11307.16 |
| lizards | Caledoniscincus_aquilonius | Caledoniscincus_aquilonius | 0 | 2.14 | 3 | 4378.35 |
| lizards | Caledoniscincus_atropunctatus | Caledoniscincus_atropunctatus | 0 | 3.09 | 2 | 20414.73 |
| lizards | Caledoniscincus_auratus | Caledoniscincus_auratus | 1 | 2.45 | 3 | 709.31 |
| lizards | Caledoniscincus_austrocaledonicus | Caledoniscincus_austrocaledonicus | 0 | 3.47 | 3 | 18804.95 |
| lizards | Caledoniscincus_chazeaui | Caledoniscincus_chazeaui | 1 | 1.41 | 3 | 1552.38 |
| lizards | Caledoniscincus_festivus | Caledoniscincus_festivus | 0 | 7.41 | 5 | 16395.49 |
| lizards | Caledoniscincus_orestes | Caledoniscincus_orestes | 1 | 5.37 | 6.5 | 1981.23 |
| lizards | Caledoniscincus_pelletieri | Caledoniscincus_pelletieri | 1 | 3.8 | 2.33 | 46.54 |
| lizards | Caledoniscincus_renevieri | Caledoniscincus_renevieri | 1 | 2.45 | 3 | 730.44 |
| lizards | Callisaurus_draconoides | Callisaurus_draconoides | 0 | 39.81 | 4.34 | 798757.74 |
| lizards | Callopistes_maculatus | Callopistes_maculatus | 0 | 162.18 | 9 | 101772.52 |
| lizards | Calodactylodes_aureus | Calodactylodes_aureus | 0 | 13.8 | 2 | 121948.8 |
| lizards | Calotes_bachae | Calotes_bachae | 0 | 30.2 | 4.5 | 103754.13 |
| lizards | Calotes_calotes | Calotes_calotes | 0 | 95.5 | 12.5 | 153325.47 |
| lizards | Calotes_ceylonensis | Calotes_ceylonensis | 0 | 18.62 | 8 | 31880.4 |

| | | | | | | |
|---|---|---|---|---|---|---|
| lizards | Calotes_desilvai | Calotes_desilvai | 1 | 16.6 | 3 | 9.99 |
| lizards | Calotes_emma | Calotes_emma | 0 | 66.07 | 7.5 | 2135284.53 |
| lizards | Calotes_grandisquamis | Calotes_grandisquamis | 0 | 117.49 | 9 | 32181.39 |
| lizards | Calotes_jerdoni | Calotes_jerdoni | 0 | 75.86 | 15 | 613086.38 |
| lizards | Calotes_liocephalus | Calotes_liocephalus | 1 | 33.11 | 5.5 | 2402.12 |
| lizards | Calotes_liolepis | Calotes_liolepis | 0 | 33.11 | 4 | 29660.61 |
| lizards | Calotes_minor | Brachysaura_minor | 0 | 30.2 | 7 | 1862266.06 |
| lizards | Calotes_mystaceus | Calotes_mystaceus | 0 | 138.04 | 7 | 1617569.77 |
| lizards | Calotes_nemoricola | Calotes_nemoricola | 0 | 117.49 | 13.71 | 20157.3 |
| lizards | Calotes_nigrilabris | Calotes_nigrilabris | 1 | 95.5 | 3.5 | 2047.7 |
| lizards | Calotes_paulus | Oriocalotes_paulus | 1 | 12.02 | 5.5 | 49343.84 |
| lizards | Calotes_pethiyagodai | Calotes_pethiyagodai | 1 | 25.7 | 5.5 | 240.34 |
| lizards | Calotes_versicolor | Calotes_versicolor | 0 | 117.49 | 13.65 | 6740870.5 |
| lizards | Calumma_boettgeri | Calumma_boettgeri | 0 | 5.62 | 3.5 | 27341.92 |
| lizards | Calumma_brevicorne | Calumma_brevicorne | 0 | 95.5 | 17.5 | 89712.2 |
| lizards | Calumma_globifer | Calumma_globifer | 1 | 123.03 | 37.5 | 2690.77 |
| lizards | Calumma_hilleniusi | Calumma_hilleniusi | 1 | 10 | 8.5 | 820.95 |
| lizards | Calumma_malthe | Calumma_malthe | 0 | 51.29 | 34 | 50405.07 |
| lizards | Calumma_nasutum | Calumma_nasutum | 0 | 3.63 | 3.5 | 203379.2 |
| lizards | Calumma_parsonii | Calumma_parsonii | 0 | 436.52 | 38 | 63315.07 |
| lizards | Calyptommatus_leiolepis | Calyptommatus_leiolepis | 1 | 3.02 | 2 | 28595.99 |
| lizards | Calyptommatus_sinebrachiatus | Calyptommatus_sinebrachiatus | 1 | 2.34 | 2 | 106.29 |
| lizards | Calyptotis_lepidorostrum | Calyptotis_lepidorostrum | 0 | 3.09 | 3.2 | 76721.46 |
| lizards | Calyptotis_ruficauda | Calyptotis_ruficauda | 0 | 3.09 | 3.3 | 36946.69 |
| lizards | Calyptotis_scutirostrum | Calyptotis_scutirostrum | 0 | 3.89 | 3.45 | 145402.8 |
| lizards | Calyptotis_temporalis | Calyptotis_temporalis | 0 | 1.51 | 2 | 20182.33 |
| lizards | Caparaonia_itaiquara | Caparaonia_itaiquara | 0 | 4.9 | 2 | 193.44 |
| lizards | Capitellum_mariagalantae | Capitellum_mariagalantae | 1 | 9.77 | 6 | 9.99 |
| lizards | Carinascincus_coventryi | Niveoscincus_coventryi | 0 | 2.95 | 2.55 | 133421.33 |
| lizards | Carinascincus_greeni | Niveoscincus_greeni | 1 | 8.51 | 2.65 | 15462.58 |
| lizards | Carinascincus_metallicus | Niveoscincus_metallicus | 0 | 7.08 | 3.8 | 96536.61 |
| lizards | Carinascincus_microlepidotus | Niveoscincus_microlepidotus | 1 | 6.76 | 2.5 | 19919.42 |
| lizards | Carinascincus_ocellatus | Niveoscincus_ocellatus | 0 | 12.88 | 3.2 | 47312.97 |
| lizards | Carinascincus_orocryptus | Niveoscincus_orocryptus | 1 | 5.75 | 2.85 | 13931.47 |

| | | | | | | |
|---|---|---|---|---|---|---|
| lizards | Carinascincus_palfreymani | Niveoscincus_palfreymani | 1 | 18.2 | 5.5 | 0.05 |
| lizards | Carinascincus_pretiosus | Niveoscincus_pretiosus | 0 | 6.76 | 2.35 | 67928.47 |
| lizards | Carlia_ailanpalai | Carlia_ailanpalai | 0 | 5.89 | 2 | 1977.08 |
| lizards | Carlia_amax | Carlia_amax | 0 | 1.2 | 2 | 753606.38 |
| lizards | Carlia_bicarinata | Carlia_bicarinata | 0 | 2.04 | 2 | 8061.15 |
| lizards | Carlia_bomberai | Carlia_bomberai | 0 | 2.95 | 2 | 24.83 |
| lizards | Carlia_caesius | Carlia_caesius | 0 | 5.25 | 2.1 | 60131.52 |
| lizards | Carlia_eothen | Carlia_eothen | 0 | 6.31 | 2 | 7207.16 |
| lizards | Carlia_gracilis | Carlia_gracilis | 0 | 1.2 | 1.9 | 280253.17 |
| lizards | Carlia_jarnoldae | Carlia_jarnoldae | 0 | 2.14 | 2 | 239066.05 |
| lizards | Carlia_longipes | Carlia_longipes | 0 | 7.76 | 2 | 60361.7 |
| lizards | Carlia_munda | Carlia_munda | 0 | 1.51 | 2 | 2703686.6 |
| lizards | Carlia_mysi | Carlia_mysi | 0 | 4.9 | 1.86 | 71129.45 |
| lizards | Carlia_pectoralis | Carlia_pectoralis | 0 | 5.13 | 2 | 218228.79 |
| lizards | Carlia_peronii | Carlia_peronii | 0 | 1.41 | 2 | 7243.73 |
| lizards | Carlia_rhomboidalis | Carlia_rhomboidalis | 0 | 4.37 | 2 | 11002.22 |
| lizards | Carlia_rostralis | Carlia_rostralis | 0 | 6.76 | 2 | 41863.25 |
| lizards | Carlia_rubrigularis | Carlia_rubrigularis | 0 | 4.37 | 2 | 21875.9 |
| lizards | Carlia_rufilatus | Carlia_rufilatus | 0 | 1.32 | 2 | 299086.37 |
| lizards | Carlia_schmeltzii | Carlia_schmeltzii | 0 | 6.46 | 2 | 690904.07 |
| lizards | Carlia_storri | Carlia_storri | 0 | 1.74 | 2 | 147749.92 |
| lizards | Carlia_tetradactyla | Carlia_tetradactyla | 0 | 5.13 | 2 | 242145.6 |
| lizards | Carlia_triacantha | Carlia_triacantha | 0 | 2.75 | 1.85 | 1878749.83 |
| lizards | Carlia_tutela | Carlia_tutela | 0 | 2.88 | 2 | 1773.8 |
| lizards | Carlia_vivax | Carlia_vivax | 0 | 2.29 | 2 | 745884.01 |
| lizards | Carphodactylus_laevis | Carphodactylus_laevis | 0 | 33.88 | 2 | 11029.81 |
| lizards | Celatiscincus_euryotis | Celatiscincus_euryotis | 1 | 1.32 | 2.5 | 143.52 |
| lizards | Celatiscincus_similis | Celatiscincus_similis | 1 | 1.7 | 2 | 451.68 |
| lizards | Celestus_crusculus | Celestus_crusculus | 0 | 11.75 | 3.35 | 11083.75 |
| lizards | Ceratophora_aspera | Ceratophora_aspera | 1 | 2.69 | 1.93 | 10376.65 |
| lizards | Ceratophora_erdeleni | Ceratophora_erdeleni | 1 | 19.5 | 3.5 | 85.3 |
| lizards | Ceratophora_karu | Ceratophora_karu | 1 | 1.95 | 2 | 19.98 |
| lizards | Ceratophora_stoddartii | Ceratophora_stoddartii | 1 | 22.91 | 3.5 | 3144.99 |
| lizards | Ceratophora_tennentii | Ceratophora_tennentii | 1 | 23.99 | 3.5 | 751.85 |

| | | | | | | |
|---|---|---|---|---|---|---|
| lizards | Cercosaura_argulus | Cercosaura_argulus | 0 | 2.69 | 1.5 | 4709101.62 |
| lizards | Cercosaura_eigenmanni | Cercosaura_eigenmanni | 0 | 3.09 | 2 | 1229760.58 |
| lizards | Cercosaura_manicata | Cercosaura_manicata | 0 | 9.33 | 2 | 42492.95 |
| lizards | Cercosaura_ocellata | Cercosaura_ocellata | 0 | 6.46 | 2.14 | 8056581.45 |
| lizards | Cercosaura_parkeri | Cercosaura_parkeri | 0 | 2.34 | 2 | 769015.04 |
| lizards | Cercosaura_phelpsorum | Cercosaura_phelpsorum | 0 | 3.72 | 2 | 25714.66 |
| lizards | Cercosaura_schreibersii | Cercosaura_schreibersii | 0 | 3.72 | 2 | 5106870.97 |
| lizards | Chalcides_bedriagai | Chalcides_bedriagai | 0 | 19.95 | 2.44 | 364193.7 |
| lizards | Chalcides_bottegi | Chalcides_bottegi | 0 | 39.81 | 6.3 | 213760.06 |
| lizards | Chalcides_chalcides | Chalcides_chalcides | 0 | 33.88 | 7.4 | 241759.96 |
| lizards | Chalcides_colosii | Chalcides_colosii | 0 | 33.11 | 6.3 | 16857.23 |
| lizards | Chalcides_guentheri | Chalcides_guentheri | 1 | 10.72 | 4.4 | 19774 |
| lizards | Chalcides_lanzai | Chalcides_lanzai | 0 | 38.9 | 6.33 | 13339.03 |
| lizards | Chalcides_manueli | Chalcides_manueli | 1 | 8.91 | 6.3 | 24838.64 |
| lizards | Chalcides_mertensi | Chalcides_mertensi | 0 | 28.18 | 6.65 | 234082.95 |
| lizards | Chalcides_minutus | Chalcides_minutus | 1 | 7.59 | 4 | 24432.16 |
| lizards | Chalcides_mionecton | Chalcides_mionecton | 0 | 6.46 | 3 | 166737.18 |
| lizards | Chalcides_montanus | Chalcides_montanus | 0 | 20.89 | 6.3 | 64863.93 |
| lizards | Chalcides_ocellatus | Chalcides_ocellatus | 0 | 97.72 | 6.5 | 9309659.68 |
| lizards | Chalcides_parallelus | Chalcides_parallelus | 1 | 16.22 | 3.5 | 1479.52 |
| lizards | Chalcides_polylepis | Chalcides_polylepis | 0 | 97.72 | 5.5 | 169524.7 |
| lizards | Chalcides_pseudostriatus | Chalcides_pseudostriatus | 0 | 30.9 | 12.67 | 101093.96 |
| lizards | Chalcides_pulchellus | Chalcides_pulchellus | 0 | 50.12 | 6.3 | 362702.34 |
| lizards | Chalcides_ragazzii | Chalcides_ragazzii | 0 | 46.77 | 6.3 | 941595.6 |
| lizards | Chalcides_sepsoides | Chalcides_sepsoides | 0 | 7.76 | 4.35 | 893517.29 |
| lizards | Chalcides_sexlineatus | Chalcides_sexlineatus | 0 | 20.89 | 4.5 | 1538.29 |
| lizards | Chalcides_striatus | Chalcides_striatus | 0 | 33.88 | 7 | 530079.99 |
| lizards | Chalcides_thierryi | Chalcides_thierryi | 0 | 97.72 | 6.3 | 571457.26 |
| lizards | Chalcides_viridanus | Chalcides_viridanus | 0 | 48.98 | 4 | 2033.77 |
| lizards | Chalcidoseps_thwaitesi | Chalcidoseps_thwaitesi | 1 | 16.98 | 2 | 1290.76 |
| lizards | Chamaeleo_africanus | Chamaeleo_africanus | 0 | 147.91 | 34 | 4788798.97 |
| lizards | Chamaeleo_anchietae | Chamaeleo_anchietae | 0 | 26.3 | 12.5 | 763213.92 |
| lizards | Chamaeleo_calyptratus | Chamaeleo_calyptratus | 0 | 239.88 | 35 | 96601.52 |
| lizards | Chamaeleo_chamaeleon | Chamaeleo_chamaeleon | 0 | 95.5 | 23.65 | 1963765.5 |

| | | | | | | |
|---|---|---|---|---|---|---|
| lizards | Chamaeleo_dilepis | Chamaeleo_dilepis | 0 | 138.04 | 39.6 | 11094336.99 |
| lizards | Chamaeleo_gracilis | Chamaeleo_gracilis | 0 | 128.82 | 35 | 11136998.65 |
| lizards | Chamaeleo_laevigatus | Chamaeleo_laevigatus | 0 | 67.61 | 37.5 | 2382716.52 |
| lizards | Chamaeleo_monachus | Chamaeleo_monachus | 0 | 102.33 | 42 | 2898.39 |
| lizards | Chamaeleo_namaquensis | Chamaeleo_namaquensis | 0 | 81.28 | 13.2 | 875059.99 |
| lizards | Chamaeleo_senegalensis | Chamaeleo_senegalensis | 0 | 70.79 | 52.2 | 3770462.91 |
| lizards | Chamaeleo_zeylanicus | Chamaeleo_zeylanicus | 0 | 229.09 | 29.88 | 2090346.12 |
| lizards | Chamaesaura_aenea | Chamaesaura_aenea | 0 | 11.75 | 11.5 | 94654.01 |
| lizards | Chamaesaura_anguina | Chamaesaura_anguina | 0 | 15.49 | 6.41 | 334093.38 |
| lizards | Chamaesaura_macrolepis | Chamaesaura_macrolepis | 0 | 19.05 | 7 | 15302.75 |
| lizards | Chamaesaura_miopropus | Chamaesaura_miopropus | 0 | 8.51 | 4.5 | 218446.37 |
| lizards | Chamaesaura_tenuior | Chamaesaura_tenuior | 0 | 14.79 | 6 | 105658.75 |
| lizards | Chatogekko_amazonicus | Chatogekko_amazonicus | 0 | 0.62 | 1 | 5812690.7 |
| lizards | Chioninia_delalandii | Chioninia_delalandii | 0 | 16.6 | 3 | 1522.71 |
| lizards | Chioninia_vaillantii | Chioninia_vaillantii | 1 | 363.08 | 7 | 1212.24 |
| lizards | Chirindia_ewerbecki | Chirindia_ewerbecki | 0 | 1.29 | 1 | 12674.47 |
| lizards | Chirindia_swynnertoni | Chirindia_swynnertoni | 0 | 1.51 | 1 | 26728.46 |
| lizards | Chlamydosaurus_kingii | Chlamydosaurus_kingii | 0 | 912.01 | 11.99 | 1508834.78 |
| lizards | Chondrodactylus_angulifer | Chondrodactylus_angulifer | 0 | 26.92 | 1.7 | 886144.6 |
| lizards | Chondrodactylus_bibronii | Chondrodactylus_bibronii | 0 | 19.05 | 2 | 1160964.31 |
| lizards | Chondrodactylus_fitzsimonsi | Chondrodactylus_fitzsimonsi | 0 | 14.13 | 1.5 | 210931.64 |
| lizards | Chondrodactylus_laevigatus | Pachydactylus_laevigatus | 0 | 14.79 | 2 | 803157.12 |
| lizards | Chondrodactylus_turneri | Chondrodactylus_turneri | 0 | 16.6 | 2 | 2890005.35 |
| lizards | Christinus_alexanderi | Christinus_alexanderi | 0 | 7.08 | 1.82 | 97691.6 |
| lizards | Christinus_guentheri | Christinus_guentheri | 1 | 19.95 | 1.3 | 51.34 |
| lizards | Christinus_marmoratus | Christinus_marmoratus | 0 | 7.08 | 1.77 | 1169407.93 |
| lizards | Cnemaspis_affinis | Cnemaspis_affinis | 1 | 2.95 | 2 | 95.39 |
| lizards | Cnemaspis_argus | Cnemaspis_argus | 0 | 6.03 | 2 | 487.24 |
| lizards | Cnemaspis_assamensis | Cnemaspis_assamensis | 1 | 0.91 | 2 | 12358.35 |
| lizards | Cnemaspis_baueri | Cnemaspis_baueri | 0 | 6.46 | 2 | 15.58 |
| lizards | Cnemaspis_bayuensis | Cnemaspis_bayuensis | 1 | 2.24 | 2 | 9.99 |
| lizards | Cnemaspis_bidongensis | Cnemaspis_bidongensis | 0 | 4.27 | 2 | 2.2 |
| lizards | Cnemaspis_biocellata | Cnemaspis_biocellata | 0 | 1.55 | 1.5 | 181.3 |
| lizards | Cnemaspis_caudanivea | Cnemaspis_caudanivea | 1 | 2.4 | 2 | 3.07 |

| | | | | | | |
|---|---|---|---|---|---|---|
| lizards | Cnemaspis_chanardi | Cnemaspis_chanardi | 0 | 1.51 | 2 | 6519.91 |
| lizards | Cnemaspis_chanthaburiensis | Cnemaspis_chanthaburiensis | 0 | 1.78 | 2 | 12507.99 |
| lizards | Cnemaspis_flavigaster | Cnemaspis_flavigaster | 0 | 2.82 | 2 | 172.87 |
| lizards | Cnemaspis_gemunu | Cnemaspis_gemunu | 1 | 0.98 | 1.5 | 379.74 |
| lizards | Cnemaspis_gracilis | Cnemaspis_gracilis | 0 | 1.51 | 2 | 28544.64 |
| lizards | Cnemaspis_grismeri | Cnemaspis_grismeri | 0 | 2.88 | 2 | 10.95 |
| lizards | Cnemaspis_hangus | Cnemaspis_hangus | 0 | 2.88 | 2 | 9.99 |
| lizards | Cnemaspis_harimau | Cnemaspis_harimau | 1 | 1.58 | 2 | 9.99 |
| lizards | Cnemaspis_heteropholis | Cnemaspis_heteropholis | 0 | 2.14 | 2 | 14604.05 |
| lizards | Cnemaspis_huaseesom | Cnemaspis_huaseesom | 0 | 1.91 | 2 | 3078.8 |
| lizards | Cnemaspis_jerdonii | Cnemaspis_jerdonii | 1 | 1.86 | 2 | 13830.96 |
| lizards | Cnemaspis_kandiana | Cnemaspis_kandiana | 1 | 1.12 | 2 | 2342.12 |
| lizards | Cnemaspis_kendallii | Cnemaspis_kendallii | 0 | 4.9 | 2 | 45212.33 |
| lizards | Cnemaspis_kumarasinghei | Cnemaspis_kumarasinghei | 1 | 0.79 | 2 | 1042.51 |
| lizards | Cnemaspis_kumpoli | Cnemaspis_kumpoli | 0 | 5.37 | 2 | 20135.99 |
| lizards | Cnemaspis_limi | Cnemaspis_limi | 0 | 14.13 | 1.5 | 144.8 |
| lizards | Cnemaspis_mcguirei | Cnemaspis_mcguirei | 0 | 5.89 | 2 | 359.97 |
| lizards | Cnemaspis_monachorum | Cnemaspis_monachorum | 0 | 1.05 | 1 | 9.99 |
| lizards | Cnemaspis_mumpuniae | Cnemaspis_mumpuniae | 0 | 4.9 | 2 | 393.56 |
| lizards | Cnemaspis_nairi | Cnemaspis_nairi | 0 | 2 | 2 | 5454.06 |
| lizards | Cnemaspis_nigridia | Cnemaspis_nigridia | 0 | 13.8 | 2 | 390.96 |
| lizards | Cnemaspis_nuicamensis | Cnemaspis_nuicamensis | 1 | 2.51 | 2 | 173.51 |
| lizards | Cnemaspis_omari | Cnemaspis_omari | 0 | 1.66 | 2 | 19.98 |
| lizards | Cnemaspis_ornata | Cnemaspis_ornata | 0 | 3.8 | 2 | 5890.18 |
| lizards | Cnemaspis_otai | Cnemaspis_otai | 1 | 0.83 | 2 | 474.98 |
| lizards | Cnemaspis_pemanggilensis | Cnemaspis_pemanggilensis | 0 | 8.91 | 2 | 9.33 |
| lizards | Cnemaspis_peninsularis | Cnemaspis_peninsularis | 0 | 4.68 | 1.9 | 20411.1 |
| lizards | Cnemaspis_perhentianensis | Cnemaspis_perhentianensis | 0 | 2.4 | 2 | 7.85 |
| lizards | Cnemaspis_podihuna | Cnemaspis_podihuna | 1 | 0.49 | 2 | 7456.34 |
| lizards | Cnemaspis_pseudomcguirei | Cnemaspis_pseudomcguirei | 0 | 1.78 | 2 | 19.98 |
| lizards | Cnemaspis_psychedelica | Cnemaspis_psychedelica | 1 | 8.71 | 2 | 6.29 |
| lizards | Cnemaspis_punctatonuchalis | Cnemaspis_punctatonuchalis | 0 | 2.75 | 2 | 9.99 |
| lizards | Cnemaspis_rammalensis | Cnemaspis_rammalensis | 1 | 3.47 | 2 | 9.99 |
| lizards | Cnemaspis_roticanai | Cnemaspis_roticanai | 0 | 2.34 | 2 | 9.99 |

| | | | | | | |
|---|---|---|---|---|---|---|
| lizards | Cnemaspis_scalpensis | Cnemaspis_scalpensis | 1 | 1.23 | 2 | 478.36 |
| lizards | Cnemaspis_selamatkanmerapoh | Cnemaspis_selamatkanmerapoh | 1 | 1.91 | 2 | 9.99 |
| lizards | Cnemaspis_shahruli | Cnemaspis_shahruli | 0 | 1.17 | 2 | 2842.22 |
| lizards | Cnemaspis_siamensis | Cnemaspis_siamensis | 0 | 1.74 | 2 | 34842.31 |
| lizards | Cnemaspis_silvula | Cnemaspis_silvula | 0 | 1.05 | 2 | 2004.1 |
| lizards | Cnemaspis_stongensis | Cnemaspis_stongensis | 0 | 2.69 | 2 | 9.99 |
| lizards | Cnemaspis_sundainsula | Cnemaspis_sundainsula | 0 | 12.02 | 2 | 41.4 |
| lizards | Cnemaspis_temiah | Cnemaspis_temiah | 1 | 1.38 | 2 | 9.99 |
| lizards | Cnemaspis_tropidogaster | Cnemaspis_tropidogaster | 1 | 1.05 | 2 | 4356.18 |
| lizards | Cnemaspis_tucdupensis | Cnemaspis_tucdupensis | 1 | 2.95 | 2 | 13.6 |
| lizards | Cnemaspis_yercaudensis | Cnemaspis_yercaudensis | 1 | 0.69 | 2 | 311.75 |
| lizards | Cnemidophorus_arenivagus | Cnemidophorus_arenivagus | 0 | 17.78 | 1.8 | 9012.9 |
| lizards | Cnemidophorus_arubensis | Cnemidophorus_arubensis | 0 | 63.1 | 1 | 181.61 |
| lizards | Cnemidophorus_cryptus | Cnemidophorus_cryptus | 0 | 10.72 | 1.5 | 851696.15 |
| lizards | Cnemidophorus_gramivagus | Cnemidophorus_gramivagus | 0 | 46.77 | 1.61 | 446489.48 |
| lizards | Cnemidophorus_lemniscatus | Cnemidophorus_lemniscatus | 0 | 43.65 | 1.87 | 3115711.91 |
| lizards | Cnemidophorus_murinus | Cnemidophorus_murinus | 0 | 125.89 | 1.18 | 840.11 |
| lizards | Cnemidophorus_nigricolor | Cnemidophorus_nigricolor | 0 | 38.02 | 2.14 | 168.34 |
| lizards | Cnemidophorus_ruthveni | Cnemidophorus_ruthveni | 0 | 107.15 | 1.5 | 17.02 |
| lizards | Coeranoscincus_reticulatus | Coeranoscincus_reticulatus | 0 | 28.18 | 3.85 | 37562.28 |
| lizards | Coleodactylus_meridionalis | Coleodactylus_meridionalis | 0 | 0.71 | 1 | 1486866.64 |
| lizards | Coleodactylus_septentrionalis | Coleodactylus_septentrionalis | 0 | 0.81 | 1 | 399484.31 |
| lizards | Coleonyx_brevis | Coleonyx_brevis | 0 | 5.37 | 2.15 | 572250.26 |
| lizards | Coleonyx_elegans | Coleonyx_elegans | 0 | 37.15 | 2 | 504670.55 |
| lizards | Coleonyx_gypsicolus | Coleonyx_gypsicolus | 0 | 13.8 | 2 | 28.41 |
| lizards | Coleonyx_mitratus | Coleonyx_mitratus | 0 | 18.62 | 2 | 98543.9 |
| lizards | Coleonyx_reticulatus | Coleonyx_reticulatus | 0 | 16.6 | 2 | 13245.73 |
| lizards | Coleonyx_switaki | Coleonyx_switaki | 0 | 12.59 | 2 | 35273.06 |
| lizards | Coleonyx_variegatus | Coleonyx_variegatus | 0 | 8.71 | 1.97 | 624429.51 |
| lizards | Colobodactylus_dalcyanus | Colobodactylus_dalcyanus | 1 | 2.69 | 2 | 693 |
| lizards | Colobosaura_modesta | Colobosaura_modesta | 0 | 3.72 | 2 | 3349736.17 |
| lizards | Concinnia_brachysoma | Concinnia_brachyosoma | 0 | 8.91 | 3.9 | 434526.52 |
| lizards | Concinnia_martini | Concinnia_martini | 0 | 7.08 | 3.5 | 281952.63 |
| lizards | Concinnia_sokosoma | Concinnia_sokosoma | 0 | 10 | 4 | 295686.27 |

| | | | | | | |
|---|---|---|---|---|---|---|
| lizards | Concinnia_tenuis | Concinnia_tenuis | 0 | 12.88 | 4.7 | 502803.82 |
| lizards | Congolacerta_vauereselli | Congolacerta_vauereselli | 0 | 5.62 | 3 | 130221.86 |
| lizards | Conolophus_marthae | Conolophus_marthae | 1 | 4365.16 | 4.4 | 9.99 |
| lizards | Conolophus_pallidus | Conolophus_pallidus | 1 | 5248.07 | 10 | 23.18 |
| lizards | Conolophus_subcristatus | Conolophus_subcristatus | 1 | 6309.57 | 10.95 | 6397 |
| lizards | Contomastix_lacertoides | Contomastix_lacertoides | 0 | 13.18 | 3.75 | 199535.48 |
| lizards | Contomastix_vacariensis | Contomastix_vacariensis | 1 | 15.85 | 4.15 | 84279.67 |
| lizards | Copeoglossum_arajara | Copeoglossum_arajara | 0 | 33.11 | 4.8 | 60516.73 |
| lizards | Copeoglossum_aurae | Copeoglossum_aurae | 0 | 28.18 | 2 | 4117.97 |
| lizards | Copeoglossum_nigropunctatum | Copeoglossum_nigropunctatum | 0 | 31.62 | 4.83 | 10876578.19 |
| lizards | Cophosaurus_texanus | Cophosaurus_texanus | 0 | 22.91 | 4.55 | 1124004.22 |
| lizards | Cophoscincopus_durus | Cophoscincopus_durus | 0 | 3.09 | 2 | 164915.9 |
| lizards | Cophoscincopus_greeri | Cophoscincopus_greeri | 0 | 5.62 | 2.67 | 146191.84 |
| lizards | Cophoscincopus_simulans | Cophoscincopus_simulans | 0 | 4.37 | 2.16 | 357521.67 |
| lizards | Cophotis_ceylanica | Cophotis_ceylanica | 1 | 9.55 | 5 | 2093.31 |
| lizards | Cordylus_cordylus | Cordylus_cordylus | 0 | 26.92 | 2 | 286245.43 |
| lizards | Cordylus_imkeae | Cordylus_imkeae | 0 | 6.76 | 1.5 | 11.29 |
| lizards | Cordylus_jonesii | Cordylus_jonesii | 0 | 19.95 | 2.5 | 683213.39 |
| lizards | Cordylus_macropholis | Cordylus_macropholis | 0 | 10.47 | 1.5 | 24268.67 |
| lizards | Cordylus_mclachlani | Cordylus_mclachlani | 0 | 8.71 | 2 | 10908.02 |
| lizards | Cordylus_meculae | Cordylus_meculae | 0 | 21.38 | 2.5 | 27.34 |
| lizards | Cordylus_minor | Cordylus_minor | 0 | 8.32 | 1.5 | 10977.51 |
| lizards | Cordylus_niger | Cordylus_niger | 0 | 19.95 | 2.5 | 4029.4 |
| lizards | Cordylus_nyikae | Cordylus_nyikae | 0 | 22.39 | 3 | 10062.9 |
| lizards | Cordylus_oelofseni | Cordylus_oelofseni | 0 | 7.08 | 2.5 | 7056.99 |
| lizards | Cordylus_rhodesianus | Cordylus_rhodesianus | 0 | 19.05 | 2.5 | 86524.12 |
| lizards | Cordylus_tropidosternum | Cordylus_tropidosternum | 0 | 34.67 | 2.5 | 1784084.28 |
| lizards | Cordylus_vittifer | Cordylus_vittifer | 0 | 22.39 | 2.5 | 364265.97 |
| lizards | Correlophus_belepensis | Correlophus_belepensis | 1 | 20.42 | 2 | 51.75 |
| lizards | Correlophus_ciliatus | Correlophus_ciliatus | 1 | 45.71 | 2 | 2529.78 |
| lizards | Correlophus_sarasinorum | Correlophus_sarasinorum | 1 | 51.29 | 2 | 1473.56 |
| lizards | Corucia_zebrata | Corucia_zebrata | 0 | 1230.27 | 1.2 | 30522.2 |
| lizards | Coryphophylax_subcristatus | Coryphophylax_subcristatus | 0 | 46.77 | 2 | 6111.47 |
| lizards | Corytophanes_cristatus | Corytophanes_cristatus | 0 | 58.88 | 6 | 568337.17 |

| | | | | | | |
|---|---|---|---|---|---|---|
| lizards | Corytophanes_hernandesii | Corytophanes_hernandesii | 0 | 51.29 | 3.5 | 161179.62 |
| lizards | Corytophanes_percarinatus | Corytophanes_percarinatus | 0 | 38.9 | 7.1 | 45357.74 |
| lizards | Crenadactylus_ocellatus | Crenadactylus_ocellatus | 0 | 0.89 | 2 | 423447.53 |
| lizards | Cricosaura_typica | Cricosaura_typica | 0 | 1.23 | 1 | 4205.36 |
| lizards | Crocodilurus_amazonicus | Crocodilurus_amazonicus | 0 | 1096.48 | 5.59 | 3319210.35 |
| lizards | Crossobamon_eversmanni | Crossobamon_eversmanni | 0 | 7.08 | 1.5 | 1699213.71 |
| lizards | Crossobamon_orientalis | Crossobamon_orientalis | 0 | 3.98 | 3 | 300169.03 |
| lizards | Crotaphytus_antiquus | Crotaphytus_antiquus | 1 | 46.77 | 3.5 | 207.37 |
| lizards | Crotaphytus_bicinctores | Crotaphytus_bicinctores | 0 | 52.48 | 4.42 | 556345.85 |
| lizards | Crotaphytus_collaris | Crotaphytus_collaris | 0 | 61.66 | 6 | 1956785.65 |
| lizards | Crotaphytus_dickersonae | Crotaphytus_dickersonae | 0 | 51.29 | 5 | 2619.96 |
| lizards | Crotaphytus_grismeri | Crotaphytus_grismeri | 0 | 39.81 | 3 | 874.49 |
| lizards | Crotaphytus_insularis | Crotaphytus_insularis | 0 | 54.95 | 4.64 | 931.87 |
| lizards | Crotaphytus_nebrius | Crotaphytus_nebrius | 0 | 48.98 | 5 | 101563.8 |
| lizards | Crotaphytus_reticulatus | Crotaphytus_reticulatus | 1 | 66.07 | 9.8 | 95194.01 |
| lizards | Crotaphytus_vestigium | Crotaphytus_vestigium | 0 | 58.88 | 2.67 | 66129.09 |
| lizards | Cryptactites_peringueyi | Cryptactites_peringueyi | 0 | 0.59 | 2 | 3963.46 |
| lizards | Cryptoblepharus_africanus | Cryptoblepharus_africanus | 0 | 2.34 | 2 | 43203.33 |
| lizards | Cryptoblepharus_australis | Cryptoblepharus_australis | 0 | 1.78 | 1.9 | 3408736.82 |
| lizards | Cryptoblepharus_bitaeniatus | Cryptoblepharus_bitaeniatus | 0 | 1.62 | 2 | 21.72 |
| lizards | Cryptoblepharus_boutonii | Cryptoblepharus_boutonii | 0 | 1.32 | 1.97 | 2143.07 |
| lizards | Cryptoblepharus_buchananii | Cryptoblepharus_buchananii | 0 | 2.19 | 2 | 1126517.29 |
| lizards | Cryptoblepharus_burdeni | Cryptoblepharus_burdeni | 0 | 2.29 | 1 | 470.28 |
| lizards | Cryptoblepharus_cygnatus | Cryptoblepharus_cygnatus | 0 | 1.58 | 1.9 | 99121.23 |
| lizards | Cryptoblepharus_eximius | Cryptoblepharus_eximius | 0 | 1.15 | 1 | 13854.64 |
| lizards | Cryptoblepharus_litoralis | Cryptoblepharus_litoralis | 0 | 3.09 | 1.64 | 19997.87 |
| lizards | Cryptoblepharus_megastictus | Cryptoblepharus_megastictus | 0 | 1.17 | 2 | 22543.99 |
| lizards | Cryptoblepharus_metallicus | Cryptoblepharus_metallicus | 0 | 2 | 2 | 2091715.06 |
| lizards | Cryptoblepharus_nigropunctatus | Cryptoblepharus_nigropunctatus | 0 | 3.09 | 1.5 | 68.03 |
| lizards | Cryptoblepharus_novocaledonicus | Cryptoblepharus_novocaledonicus | 0 | 1.41 | 2 | 16548.09 |
| lizards | Cryptoblepharus_novohebridicus | Cryptoblepharus_novohebridicus | 0 | 0.87 | 1.5 | 10623.94 |
| lizards | Cryptoblepharus_pannosus | Cryptoblepharus_pannosus | 0 | 1.51 | 1.5 | 2570391.55 |
| lizards | Cryptoblepharus_plagiocephalus | Cryptoblepharus_plagiocephalus | 0 | 1.86 | 1.95 | 508873.63 |
| lizards | Cryptoblepharus_poecilopleurus | Cryptoblepharus_poecilopleurus | 0 | 2.63 | 1.55 | 44707.71 |

| | | | | | | |
|---|---|---|---|---|---|---|
| lizards | Cryptoblepharus_pulcher | Cryptoblepharus_pulcher | 0 | 1.29 | 1.9 | 756469.16 |
| lizards | Cryptoblepharus_renschi | Cryptoblepharus_renschi | 0 | 2.29 | 1.5 | 17157.35 |
| lizards | Cryptoblepharus_ruber | Cryptoblepharus_ruber | 0 | 1.91 | 2 | 497359.1 |
| lizards | Cryptoblepharus_virgatus | Cryptoblepharus_virgatus | 0 | 1.2 | 1.9 | 188197.61 |
| lizards | Cryptoblepharus_yulensis | Cryptoblepharus_yulensis | 0 | 1.29 | 1.6 | 2825.43 |
| lizards | Ctenophorus_adelaidensis | Ctenophorus_adelaidensis | 0 | 4.68 | 3.5 | 45133.67 |
| lizards | Ctenophorus_butlerorum | Ctenophorus_butlerorum | 0 | 2.82 | 2.5 | 3219.91 |
| lizards | Ctenophorus_caudicinctus | Ctenophorus_caudicinctus | 0 | 38.9 | 5.38 | 818951.3 |
| lizards | Ctenophorus_clayi | Ctenophorus_clayi | 0 | 6.17 | 2.45 | 2030499.97 |
| lizards | Ctenophorus_cristatus | Ctenophorus_cristatus | 0 | 47.86 | 4.55 | 816178.42 |
| lizards | Ctenophorus_decresii | Ctenophorus_decresii | 0 | 23.99 | 4.7 | 122891.22 |
| lizards | Ctenophorus_femoralis | Ctenophorus_femoralis | 0 | 5.89 | 2 | 62060.05 |
| lizards | Ctenophorus_fionni | Ctenophorus_fionni | 0 | 29.51 | 3.58 | 140174.98 |
| lizards | Ctenophorus_fordi | Ctenophorus_fordi | 0 | 6.17 | 2.3 | 1229820.21 |
| lizards | Ctenophorus_isolepis | Ctenophorus_isolepis | 0 | 18.62 | 3.7 | 3313601.45 |
| lizards | Ctenophorus_maculatus | Ctenophorus_maculatus | 0 | 9.55 | 3 | 511181 |
| lizards | Ctenophorus_maculosus | Ctenophorus_maculosus | 0 | 10.96 | 3.21 | 82125.22 |
| lizards | Ctenophorus_mirrityana | Ctenophorus_mirrityana | 0 | 24.55 | 5.5 | 2043.32 |
| lizards | Ctenophorus_nguyarna | Ctenophorus_nguyarna | 1 | 15.85 | 2.5 | 3130.81 |
| lizards | Ctenophorus_nuchalis | Ctenophorus_nuchalis | 0 | 70.79 | 4.05 | 4690120.93 |
| lizards | Ctenophorus_ornatus | Ctenophorus_ornatus | 0 | 28.18 | 3.13 | 360252.68 |
| lizards | Ctenophorus_parviceps | Ctenophorus_parviceps | 0 | 4.37 | 2.1 | 38678.17 |
| lizards | Ctenophorus_pictus | Ctenophorus_pictus | 0 | 13.49 | 3.2 | 3023219.88 |
| lizards | Ctenophorus_reticulatus | Ctenophorus_reticulatus | 0 | 42.66 | 4.3 | 2333990.52 |
| lizards | Ctenophorus_salinarum | Ctenophorus_salinarum | 0 | 15.49 | 3.5 | 607776.37 |
| lizards | Ctenophorus_scutulatus | Ctenophorus_scutulatus | 0 | 51.29 | 5.17 | 755547.53 |
| lizards | Ctenophorus_vadnappa | Ctenophorus_vadnappa | 0 | 23.99 | 5 | 31990.41 |
| lizards | Ctenosaura_acanthura | Ctenosaura_acanthura | 0 | 3890.45 | 20.67 | 63252.75 |
| lizards | Ctenosaura_bakeri | Ctenosaura_bakeri | 1 | 1348.96 | 9.8 | 42.08 |
| lizards | Ctenosaura_clarki | Ctenosaura_clarki | 1 | 676.08 | 7 | 9608.12 |
| lizards | Ctenosaura_flavidorsalis | Ctenosaura_flavidorsalis | 0 | 275.42 | 9.5 | 12090.93 |
| lizards | Ctenosaura_hemilopha | Ctenosaura_hemilopha | 0 | 2754.23 | 24 | 141254.58 |
| lizards | Ctenosaura_macrolopha | Ctenosaura_macrolopha | 0 | 416.87 | 10.5 | 42785.87 |
| lizards | Ctenosaura_melanosterna | Ctenosaura_melanosterna | 1 | 1445.44 | 17 | 580.58 |

| | | | | | | |
|---|---|---|---|---|---|---|
| lizards | Ctenosaura_oedirhina | Ctenosaura_oedirhina | 1 | 1348.96 | 8 | 160.3 |
| lizards | Ctenosaura_palearis | Ctenosaura_palearis | 1 | 1288.25 | 9 | 940.36 |
| lizards | Ctenosaura_pectinata | Ctenosaura_pectinata | 0 | 2691.53 | 28.05 | 358773.12 |
| lizards | Ctenosaura_similis | Ctenosaura_similis | 0 | 5011.87 | 32.7 | 893356.47 |
| lizards | Ctenotus_alacer | Ctenotus_alacer | 0 | 6.03 | 2 | 722622.96 |
| lizards | Ctenotus_alleni | Ctenotus_alleni | 0 | 16.98 | 3 | 13512.05 |
| lizards | Ctenotus_ariadnae | Ctenotus_ariadnae | 0 | 5.13 | 3.42 | 1189930.55 |
| lizards | Ctenotus_arnhemensis | Ctenotus_arnhemensis | 0 | 3.16 | 3.5 | 10830.53 |
| lizards | Ctenotus_atlas | Ctenotus_atlas | 0 | 8.51 | 1.5 | 985391.66 |
| lizards | Ctenotus_australis | Ctenotus_australis | 0 | 29.51 | 3.3 | 72199.85 |
| lizards | Ctenotus_brevipes | Ctenotus_brevipes | 0 | 4.17 | 2.95 | 252239.38 |
| lizards | Ctenotus_brooksi | Ctenotus_brooksi | 0 | 3.47 | 2.17 | 1745904.08 |
| lizards | Ctenotus_calurus | Ctenotus_calurus | 0 | 2.29 | 2.27 | 1489493.44 |
| lizards | Ctenotus_coggeri | Ctenotus_coggeri | 0 | 13.8 | 3 | 50864.81 |
| lizards | Ctenotus_colletti | Ctenotus_colletti | 0 | 1.62 | 3 | 31414.94 |
| lizards | Ctenotus_dux | Ctenotus_dux | 0 | 5.75 | 2.68 | 1196447.38 |
| lizards | Ctenotus_essingtonii | Ctenotus_essingtonii | 0 | 7.76 | 2.95 | 230922.52 |
| lizards | Ctenotus_gagudju | Ctenotus_gagudju | 0 | 2.95 | 2 | 4592.28 |
| lizards | Ctenotus_gemmula | Ctenotus_gemmula | 0 | 4.17 | 2 | 177921.29 |
| lizards | Ctenotus_grandis | Ctenotus_grandis | 0 | 40.74 | 4 | 1509619.04 |
| lizards | Ctenotus_hanloni | Ctenotus_hanloni | 0 | 7.94 | 3.6 | 1498577.69 |
| lizards | Ctenotus_hebetior | Ctenotus_hebetior | 0 | 4.17 | 4 | 580090.8 |
| lizards | Ctenotus_impar | Ctenotus_impar | 0 | 5.62 | 2 | 305520.16 |
| lizards | Ctenotus_inornatus | Ctenotus_inornatus | 0 | 18.2 | 6 | 4465089.52 |
| lizards | Ctenotus_kurnbudj | Ctenotus_kurnbudj | 1 | 3.31 | 3 | 3260.86 |
| lizards | Ctenotus_labillardieri | Ctenotus_labillardieri | 0 | 8.91 | 3.55 | 118665.43 |
| lizards | Ctenotus_lancelini | Ctenotus_lancelini | 1 | 13.8 | 3.33 | 12.1 |
| lizards | Ctenotus_leae | Ctenotus_leae | 0 | 4.57 | 3.25 | 1951798.11 |
| lizards | Ctenotus_leonhardii | Ctenotus_leonhardii | 0 | 10 | 3.8 | 4712048.74 |
| lizards | Ctenotus_mimetes | Ctenotus_mimetes | 0 | 11.22 | 6 | 182002.25 |
| lizards | Ctenotus_nasutus | Ctenotus_nasutus | 0 | 1.74 | 2.91 | 936346.32 |
| lizards | Ctenotus_olympicus | Ctenotus_olympicus | 0 | 8.51 | 3.5 | 705214.88 |
| lizards | Ctenotus_orientalis | Ctenotus_orientalis | 0 | 11.22 | 4 | 1008363.01 |
| lizards | Ctenotus_pallescens | Ctenotus_pallescens | 0 | 1.62 | 3 | 405369.89 |

| | | | | | | |
|---|---|---|---|---|---|---|
| lizards | Ctenotus_pantherinus | Ctenotus_pantherinus | 0 | 45.71 | 5.1 | 5203160.87 |
| lizards | Ctenotus_piankai | Ctenotus_piankai | 0 | 4.17 | 2.75 | 2013068.23 |
| lizards | Ctenotus_quattuordecimlineatus | Ctenotus_quattuordecimlineatus | 0 | 7.08 | 3.3 | 1773965.61 |
| lizards | Ctenotus_regius | Ctenotus_regius | 0 | 11.75 | 2.75 | 1705313.27 |
| lizards | Ctenotus_robustus | Ctenotus_robustus | 0 | 44.67 | 5 | 3394362.13 |
| lizards | Ctenotus_rufescens | Ctenotus_rufescens | 0 | 1.62 | 2 | 49511.45 |
| lizards | Ctenotus_rutilans | Ctenotus_rutilans | 0 | 3.98 | 2 | 119782.69 |
| lizards | Ctenotus_schomburgkii | Ctenotus_schomburgkii | 0 | 3.47 | 2.48 | 4312611.49 |
| lizards | Ctenotus_septenarius | Ctenotus_septenarius | 0 | 7.41 | 4 | 651429.86 |
| lizards | Ctenotus_spaldingi | Ctenotus_spaldingi | 0 | 29.51 | 6 | 2891125.55 |
| lizards | Ctenotus_storri | Ctenotus_storri | 0 | 1.12 | 2 | 117064.7 |
| lizards | Ctenotus_strauchii | Ctenotus_strauchii | 0 | 4.9 | 2.19 | 2788871.03 |
| lizards | Ctenotus_taeniolatus | Ctenotus_taeniolatus | 0 | 14.79 | 3.8 | 1203988.06 |
| lizards | Ctenotus_uber | Ctenotus_uber | 0 | 11.22 | 2 | 1062714.22 |
| lizards | Ctenotus_vertebralis | Ctenotus_vertebralis | 0 | 3.09 | 3 | 179335.69 |
| lizards | Cyclodomorphus_branchialis | Cyclodomorphus_branchialis | 0 | 38.9 | 3 | 56971.33 |
| lizards | Cyclodomorphus_casuarinae | Cyclodomorphus_casuarinae | 0 | 128.82 | 5.8 | 67107.96 |
| lizards | Cyclodomorphus_celatus | Cyclodomorphus_celatus | 0 | 39.81 | 3.8 | 72346.06 |
| lizards | Cyclodomorphus_gerrardii | Hemisphaeriodon_gerrardii | 0 | 446.68 | 21.65 | 419636.98 |
| lizards | Cyclodomorphus_maximus | Cyclodomorphus_maxima | 0 | 323.59 | 4.5 | 82497.22 |
| lizards | Cyclodomorphus_melanops | Cyclodomorphus_melanops | 0 | 52.48 | 2.7 | 3536483.12 |
| lizards | Cyclodomorphus_michaeli | Cyclodomorphus_michaeli | 0 | 128.82 | 7.65 | 37214.24 |
| lizards | Cyclodomorphus_praealtus | Cyclodomorphus_praealtus | 1 | 45.71 | 4.9 | 8620.45 |
| lizards | Cyclodomorphus_venustus | Cyclodomorphus_venustus | 0 | 22.39 | 3 | 576717.49 |
| lizards | Cyclura_carinata | Cyclura_carinata | 1 | 5623.41 | 4.3 | 153.24 |
| lizards | Cyclura_collei | Cyclura_collei | 1 | 3311.31 | 17.15 | 565.94 |
| lizards | Cyclura_cornuta | Cyclura_cornuta | 1 | 10964.78 | 14.7 | 29980.87 |
| lizards | Cyclura_cychlura | Cyclura_cychlura | 1 | 11748.98 | 6.4 | 154.76 |
| lizards | Cyclura_lewisi | Cyclura_lewisi | 1 | 7079.46 | 11.5 | 210.62 |
| lizards | Cyclura_nubila | Cyclura_nubila | 1 | 17782.79 | 10.3 | 86522.68 |
| lizards | Cyclura_pinguis | Cyclura_pinguis | 1 | 7413.1 | 12.3 | 41.59 |
| lizards | Cyclura_ricordi | Cyclura_ricordi | 1 | 4786.3 | 9.85 | 452.68 |
| lizards | Cyclura_rileyi | Cyclura_rileyi | 1 | 2630.27 | 3.75 | 32.58 |
| lizards | Cynisca_feae | Cynisca_feae | 0 | 1.86 | 2.82 | 157282.42 |

| | | | | | | |
|---|---|---|---|---|---|---|
| lizards | Cynisca_leonina | Cynisca_leonina | 1 | 1.35 | 2 | 13429.44 |
| lizards | Cynisca_leucura | Cynisca_leucura | 0 | 10.47 | 2.25 | 56138.21 |
| lizards | Cynisca_oligopholis | Cynisca_oligopholis | 1 | 1.1 | 4.5 | 4309.47 |
| lizards | Cyrtodactylus_agusanensis | Cyrtodactylus_agusanensis | 0 | 22.39 | 2 | 115002.6 |
| lizards | Cyrtodactylus_albofasciatus | Cyrtodactylus_albofasciatus | 0 | 8.91 | 2 | 22867.66 |
| lizards | Cyrtodactylus_angularis | Cyrtodactylus_angularis | 0 | 16.6 | 2 | 1568.18 |
| lizards | Cyrtodactylus_annandalei | Cyrtodactylus_annandalei | 0 | 3.72 | 2 | 12943.26 |
| lizards | Cyrtodactylus_astrum | Cyrtodactylus_astrum | 0 | 23.99 | 2 | 2047.8 |
| lizards | Cyrtodactylus_aurensis | Cyrtodactylus_aurensis | 0 | 18.62 | 2 | 15.58 |
| lizards | Cyrtodactylus_australotitiwangsaensis | Cyrtodactylus_australotitiwangsaensis | 0 | 31.62 | 2 | 1668.78 |
| lizards | Cyrtodactylus_ayeyarwadyensis | Cyrtodactylus_ayeyarwadyensis | 0 | 9.55 | 2 | 27844.63 |
| lizards | Cyrtodactylus_baluensis | Cyrtodactylus_baluensis | 0 | 18.2 | 2.05 | 39485.08 |
| lizards | Cyrtodactylus_batucolus | Cyrtodactylus_batucolus | 0 | 8.71 | 2 | 1.33 |
| lizards | Cyrtodactylus_bintangrendah | Cyrtodactylus_bintangrendah | 0 | 28.84 | 2 | 2649.87 |
| lizards | Cyrtodactylus_bintangtinggi | Cyrtodactylus_bintangtinggi | 0 | 25.7 | 2 | 134.09 |
| lizards | Cyrtodactylus_biordinis | Cyrtodactylus_biordinis | 0 | 16.22 | 2 | 5461.35 |
| lizards | Cyrtodactylus_boreoclivus | Cyrtodactylus_boreoclivus | 0 | 23.99 | 2 | 4892.42 |
| lizards | Cyrtodactylus_brevidactylus | Cyrtodactylus_brevidactylus | 1 | 13.49 | 2 | 230.57 |
| lizards | Cyrtodactylus_brevipalmatus | Cyrtodactylus_brevipalmatus | 0 | 7.94 | 2 | 98073.66 |
| lizards | Cyrtodactylus_capreoloides | Cyrtodactylus_capreoloides | 0 | 11.75 | 2 | 10629.39 |
| lizards | Cyrtodactylus_cattienensis | Cyrtodactylus_cattienensis | 0 | 6.92 | 1.5 | 10730.45 |
| lizards | Cyrtodactylus_cavernicolus | Cyrtodactylus_cavernicolus | 0 | 10.72 | 2 | 986.35 |
| lizards | Cyrtodactylus_chanhomeae | Cyrtodactylus_chanhomeae | 1 | 9.77 | 2 | 41.66 |
| lizards | Cyrtodactylus_chrysopylos | Cyrtodactylus_chrysopylos | 1 | 11.75 | 2 | 120.11 |
| lizards | Cyrtodactylus_collegalensis | Cyrtodactylus_collegalensis | 0 | 3.63 | 2 | 260686.13 |
| lizards | Cyrtodactylus_condorensis | Cyrtodactylus_condorensis | 0 | 14.79 | 3 | 2941.73 |
| lizards | Cyrtodactylus_consobrinus | Cyrtodactylus_consobrinus | 0 | 35.48 | 2 | 285419.7 |
| lizards | Cyrtodactylus_cryptus | Cyrtodactylus_cryptus | 0 | 15.49 | 2 | 956.47 |
| lizards | Cyrtodactylus_deccanensis | Cyrtodactylus_deccanensis | 0 | 12.3 | 2 | 71091.35 |
| lizards | Cyrtodactylus_dumnuii | Cyrtodactylus_dumnuii | 0 | 11.75 | 2 | 10965.49 |
| lizards | Cyrtodactylus_elok | Cyrtodactylus_elok | 0 | 10.96 | 2 | 4193.56 |
| lizards | Cyrtodactylus_epiroticus | Cyrtodactylus_epiroticus | 0 | 42.66 | 2 | 28614.09 |
| lizards | Cyrtodactylus_erythrops | Cyrtodactylus_erythrops | 0 | 10.23 | 2 | 70.23 |
| lizards | Cyrtodactylus_fraenatus | Cyrtodactylus_fraenatus | 1 | 19.05 | 2 | 728.33 |

| | | | | | | |
|---|---|---|---|---|---|---|
| lizards | Cyrtodactylus_gansi | Cyrtodactylus_gansi | 0 | 5.37 | 2 | 436.27 |
| lizards | Cyrtodactylus_huynhi | Cyrtodactylus_huynhi | 1 | 10.23 | 1 | 19.98 |
| lizards | Cyrtodactylus_ingeri | Cyrtodactylus_ingeri | 0 | 10.47 | 2 | 24505.71 |
| lizards | Cyrtodactylus_interdigitalis | Cyrtodactylus_interdigitalis | 0 | 10.23 | 1 | 88252.15 |
| lizards | Cyrtodactylus_intermedius | Cyrtodactylus_intermedius | 0 | 15.14 | 1.91 | 42675.81 |
| lizards | Cyrtodactylus_jarakensis | Cyrtodactylus_jarakensis | 1 | 6.31 | 2 | 0.35 |
| lizards | Cyrtodactylus_jellesmae | Cyrtodactylus_jellesmae | 0 | 10 | 2 | 138356.93 |
| lizards | Cyrtodactylus_laevigatus | Cyrtodactylus_laevigatus | 0 | 2.34 | 2 | 1085 |
| lizards | Cyrtodactylus_langkawiensis | Cyrtodactylus_langkawiensis | 0 | 19.05 | 2 | 9.99 |
| lizards | Cyrtodactylus_lateralis | Cyrtodactylus_lateralis | 0 | 16.98 | 2 | 2186.89 |
| lizards | Cyrtodactylus_leegrismeri | Cyrtodactylus_leegrismeri | 0 | 15.14 | 2 | 4.04 |
| lizards | Cyrtodactylus_lekaguli | Cyrtodactylus_lekaguli | 0 | 20.89 | 1.5 | 14886.84 |
| lizards | Cyrtodactylus_loriae | Cyrtodactylus_loriae | 0 | 64.57 | 2 | 16920.85 |
| lizards | Cyrtodactylus_louisiadensis | Cyrtodactylus_louisiadensis | 0 | 41.69 | 2 | 6258.19 |
| lizards | Cyrtodactylus_macrotuberculatus | Cyrtodactylus_macrotuberculatus | 0 | 31.62 | 1.75 | 10334.43 |
| lizards | Cyrtodactylus_majulah | Cyrtodactylus_majulah | 0 | 6.76 | 2 | 19.98 |
| lizards | Cyrtodactylus_malayanus | Cyrtodactylus_malayanus | 0 | 30.2 | 1.99 | 13679.76 |
| lizards | Cyrtodactylus_marmoratus | Cyrtodactylus_marmoratus | 0 | 12.3 | 2 | 7798 |
| lizards | Cyrtodactylus_mcdonaldi | Cyrtodactylus_mcdonaldi | 0 | 21.88 | 2 | 17782.5 |
| lizards | Cyrtodactylus_mimikanus | Cyrtodactylus_mimikanus | 0 | 20.89 | 2 | 16888.43 |
| lizards | Cyrtodactylus_oldhami | Cyrtodactylus_oldhami | 0 | 9.33 | 2 | 186494.62 |
| lizards | Cyrtodactylus_pageli | Cyrtodactylus_pageli | 0 | 13.8 | 1.5 | 24.34 |
| lizards | Cyrtodactylus_pantiensis | Cyrtodactylus_pantiensis | 1 | 9.33 | 2 | 13096.4 |
| lizards | Cyrtodactylus_papilionoides | Cyrtodactylus_papilionoides | 0 | 15.49 | 1.5 | 6130.93 |
| lizards | Cyrtodactylus_papuensis | Cyrtodactylus_papuensis | 0 | 7.08 | 2 | 122366.53 |
| lizards | Cyrtodactylus_peguensis | Cyrtodactylus_peguensis | 0 | 12.3 | 1.5 | 50780.14 |
| lizards | Cyrtodactylus_philippinicus | Cyrtodactylus_philippinicus | 0 | 17.78 | 2 | 165885.82 |
| lizards | Cyrtodactylus_phongnhakebangensis | Cyrtodactylus_phongnhakebangensis | 0 | 21.38 | 2 | 267.06 |
| lizards | Cyrtodactylus_pseudoquadrivirgatus | Cyrtodactylus_pseudoquadrivirgatus | 0 | 13.18 | 1 | 43794.9 |
| lizards | Cyrtodactylus_pubisulcus | Cyrtodactylus_pubisulcus | 0 | 9.33 | 1.97 | 54306.81 |
| lizards | Cyrtodactylus_pulchellus | Cyrtodactylus_pulchellus | 1 | 35.48 | 2 | 319.18 |
| lizards | Cyrtodactylus_redimiculus | Cyrtodactylus_redimiculus | 0 | 16.22 | 2 | 3158.92 |
| lizards | Cyrtodactylus_rubidus | Cyrtodactylus_rubidus | 0 | 10.23 | 2 | 1467.54 |
| lizards | Cyrtodactylus_sadleiri | Cyrtodactylus_sadleiri | 1 | 13.49 | 2 | 137.18 |

| | | | | | | |
|---|---|---|---|---|---|---|
| lizards | Cyrtodactylus_samroiyot | Cyrtodactylus_samroiyot | 0 | 6.31 | 2 | 17.89 |
| lizards | Cyrtodactylus_semenanjungensis | Cyrtodactylus_semenanjungensis | 0 | 7.41 | 2 | 277.95 |
| lizards | Cyrtodactylus_semiadii | Cyrtodactylus_semiadii | 0 | 2.4 | 1.5 | 873.76 |
| lizards | Cyrtodactylus_seribuatensis | Cyrtodactylus_seribuatensis | 1 | 8.71 | 2 | 28.4 |
| lizards | Cyrtodactylus_sermowaiensis | Cyrtodactylus_sermowaiensis | 0 | 21.88 | 1.5 | 102712.68 |
| lizards | Cyrtodactylus_soba | Cyrtodactylus_soba | 1 | 22.39 | 2 | 83.87 |
| lizards | Cyrtodactylus_sumonthai | Cyrtodactylus_sumonthai | 0 | 10.47 | 1.5 | 9.99 |
| lizards | Cyrtodactylus_sworderi | Cyrtodactylus_sworderi | 1 | 10.23 | 2 | 116326.58 |
| lizards | Cyrtodactylus_takouensis | Cyrtodactylus_takouensis | 1 | 10.72 | 2 | 9.99 |
| lizards | Cyrtodactylus_tebuensis | Cyrtodactylus_tebuensis | 0 | 12.3 | 2 | 14.93 |
| lizards | Cyrtodactylus_tigroides | Cyrtodactylus_tigroides | 0 | 12.3 | 2 | 19.98 |
| lizards | Cyrtodactylus_timur | Cyrtodactylus_timur | 0 | 31.62 | 2 | 14.93 |
| lizards | Cyrtodactylus_tiomanensis | Cyrtodactylus_tiomanensis | 0 | 12.88 | 2 | 96.76 |
| lizards | Cyrtodactylus_triedrus | Cyrtodactylus_triedrus | 0 | 5.13 | 2 | 23243.06 |
| lizards | Cyrtodactylus_tuberculatus | Cyrtodactylus_tuberculatus | 0 | 47.86 | 2 | 20487.1 |
| lizards | Cyrtodactylus_wakeorum | Cyrtodactylus_wakeorum | 1 | 5.5 | 2 | 5150.32 |
| lizards | Cyrtodactylus_wayakonei | Cyrtodactylus_wayakonei | 0 | 15.49 | 2 | 19.98 |
| lizards | Cyrtodactylus_yakhuna | Cyrtodactylus_yakhuna | 1 | 3.16 | 3 | 21228.66 |
| lizards | Cyrtodactylus_yangbayensis | Cyrtodactylus_yangbayensis | 0 | 15.14 | 2 | 64.75 |
| lizards | Cyrtodactylus_yoshii | Cyrtodactylus_yoshii | 0 | 20.89 | 1.99 | 10162.52 |
| lizards | Cyrtopodion_agamuroides | Cyrtopodion_agamuroides | 0 | 2.82 | 2 | 546163.08 |
| lizards | Cyrtopodion_fortmunroi | Cyrtopodion_fortmunroi | 0 | 2.88 | 2 | 15 |
| lizards | Cyrtopodion_indusoani | Cyrtopodion_indusoani | 0 | 3.47 | 2 | 4704.02 |
| lizards | Cyrtopodion_kachhense | Cyrtopodion_kachhense | 0 | 3.63 | 1.5 | 317223.42 |
| lizards | Cyrtopodion_kohsulaimanai | Cyrtopodion_kohsulaimanai | 0 | 4.47 | 2.5 | 65691.78 |
| lizards | Cyrtopodion_potoharense | Cyrtopodion_potoharense | 0 | 3.16 | 1.5 | 76284.19 |
| lizards | Cyrtopodion_rohtasfortai | Cyrtopodion_rohtasfortai | 0 | 4.07 | 2 | 6935.43 |
| lizards | Cyrtopodion_scabrum | Cyrtopodion_scabrum | 0 | 5.75 | 1.96 | 4041175.66 |
| lizards | Cyrtopodion_watsoni | Cyrtopodion_watsoni | 0 | 3.47 | 2 | 52928.68 |
| lizards | Dactylocnemis_pacificus | Dactylocnemis_pacificus | 0 | 20.42 | 2 | 84526.14 |
| lizards | Dalmatolacerta_oxycephala | Dalmatolacerta_oxycephala | 0 | 8.13 | 3.4 | 30034.6 |
| lizards | Dalophia_longicauda | Dalophia_longicauda | 0 | 173.78 | 3 | 120933.92 |
| lizards | Dalophia_pistillum | Dalophia_pistillum | 0 | 229.09 | 4 | 1307673.48 |
| lizards | Darevskia_alpina | Darevskia_alpina | 1 | 7.24 | 4 | 6118.55 |

| | | | | | | |
|---|---|---|---|---|---|---|
| lizards | Darevskia_armeniaca | Darevskia_armeniaca | 0 | 9.12 | 4 | 57387.05 |
| lizards | Darevskia_bendimahiensis | Darevskia_bendimahiensis | 1 | 6.17 | 2 | 4282.67 |
| lizards | Darevskia_brauneri | Darevskia_brauneri | 0 | 8.32 | 4 | 47237.89 |
| lizards | Darevskia_caucasica | Darevskia_caucasica | 0 | 7.08 | 4 | 142840.96 |
| lizards | Darevskia_chlorogaster | Darevskia_chlorogaster | 0 | 8.71 | 6.5 | 54120.58 |
| lizards | Darevskia_daghestanica | Darevskia_daghestanica | 0 | 4.57 | 3 | 27783.68 |
| lizards | Darevskia_dahli | Darevskia_dahli | 0 | 6.17 | 3.5 | 22647.9 |
| lizards | Darevskia_defilippii | Darevskia_defilippii | 0 | 4.57 | 5.2 | 76748.36 |
| lizards | Darevskia_derjugini | Darevskia_derjugini | 0 | 6.46 | 6.2 | 51277.37 |
| lizards | Darevskia_lindholmi | Darevskia_lindholmi | 0 | 9.77 | 4 | 25224.53 |
| lizards | Darevskia_mixta | Darevskia_mixta | 0 | 5.89 | 3 | 6446.27 |
| lizards | Darevskia_parvula | Darevskia_parvula | 0 | 5.01 | 3 | 39377.42 |
| lizards | Darevskia_portschinskii | Darevskia_portschinskii | 0 | 7.08 | 3.5 | 24669.69 |
| lizards | Darevskia_praticola | Darevskia_praticola | 0 | 6.76 | 4.3 | 548280.82 |
| lizards | Darevskia_raddei | Darevskia_raddei | 0 | 10.23 | 4 | 198744.71 |
| lizards | Darevskia_rostombekowi | Darevskia_rostombekovi | 1 | 5.01 | 2.5 | 6860.75 |
| lizards | Darevskia_rudis | Darevskia_rudis | 0 | 15.85 | 5 | 238514.98 |
| lizards | Darevskia_sapphirina | Darevskia_sapphirina | 0 | 4.79 | 3.5 | 53.09 |
| lizards | Darevskia_saxicola | Darevskia_saxicola | 0 | 15.85 | 4 | 19671.5 |
| lizards | Darevskia_unisexualis | Darevskia_unisexualis | 0 | 9.12 | 5 | 79925.59 |
| lizards | Darevskia_valentini | Darevskia_valentini | 0 | 11.75 | 5.6 | 259966.14 |
| lizards | Dasia_grisea | Dasia_grisea | 0 | 50.12 | 3.3 | 166998.03 |
| lizards | Dasia_haliana | Dasia_haliana | 1 | 15.49 | 2 | 34469.85 |
| lizards | Dasia_olivacea | Dasia_olivacea | 0 | 79.43 | 6.17 | 1953300.13 |
| lizards | Dasia_vittata | Dasia_vittata | 0 | 19.05 | 3.05 | 434667.82 |
| lizards | Delma_australis | Delma_australis | 0 | 4.57 | 2 | 1472695.69 |
| lizards | Delma_borea | Delma_borea | 0 | 4.9 | 1.5 | 2244028.15 |
| lizards | Delma_butleri | Delma_butleri | 0 | 5.75 | 2 | 3244974.97 |
| lizards | Delma_concinna | Delma_concinna | 0 | 5.89 | 2 | 12765.48 |
| lizards | Delma_fraseri | Delma_fraseri | 0 | 7.94 | 2 | 591977.39 |
| lizards | Delma_grayii | Delma_grayii | 0 | 6.92 | 2 | 115357.6 |
| lizards | Delma_impar | Delma_impar | 1 | 6.17 | 2 | 205008.69 |
| lizards | Delma_inornata | Delma_inornata | 0 | 7.41 | 2 | 656642.12 |
| lizards | Delma_labialis | Delma_labialis | 0 | 6.92 | 1 | 62756.53 |

| | | | | | | |
|---|---|---|---|---|---|---|
| lizards | Delma_molleri | Delma_molleri | 0 | 5.89 | 2 | 52247.84 |
| lizards | Delma_nasuta | Delma_nasuta | 0 | 5.89 | 2 | 3222078.03 |
| lizards | Delma_petersoni | Delma_petersoni | 0 | 7.08 | 2 | 373397.28 |
| lizards | Delma_plebeia | Delma_plebeia | 0 | 6.61 | 2 | 281255.73 |
| lizards | Delma_tincta | Delma_tincta | 0 | 4.47 | 2 | 4730812.56 |
| lizards | Delma_torquata | Delma_torquata | 0 | 2.69 | 2 | 188978.5 |
| lizards | Dendragama_boulengeri | Dendragama_boulengeri | 0 | 16.22 | 4 | 12700.22 |
| lizards | Dibamus_novaeguineae | Dibamus_novaeguineae | 0 | 7.76 | 1 | 15933.86 |
| lizards | Dierogekko_inexpectatus | Dierogekko_inexpectatus | 1 | 1.2 | 2 | 29.76 |
| lizards | Dierogekko_insularis | Dierogekko_insularis | 1 | 1.38 | 2 | 79.37 |
| lizards | Dierogekko_kaalaensis | Dierogekko_kaalaensis | 1 | 1.82 | 2 | 54.65 |
| lizards | Dierogekko_koniambo | Dierogekko_koniambo | 1 | 1.55 | 2 | 232.69 |
| lizards | Dierogekko_nehoueensis | Dierogekko_nehoueensis | 1 | 1.26 | 2 | 179.51 |
| lizards | Dierogekko_poumensis | Dierogekko_poumensis | 1 | 1.1 | 2 | 30.99 |
| lizards | Dierogekko_thomaswhitei | Dierogekko_thomaswhitei | 1 | 1.78 | 2 | 62.17 |
| lizards | Dierogekko_validiclavis | Dierogekko_validiclavis | 1 | 1.78 | 2 | 615.45 |
| lizards | Dinarolacerta_mosorensis | Dinarolacerta_mosorensis | 1 | 11.75 | 4.75 | 17451.53 |
| lizards | Diplodactylus_calcicolus | Diplodactylus_calcicolus | 0 | 3.89 | 2 | 346267.47 |
| lizards | Diplodactylus_capensis | Diplodactylus_capensis | 0 | 5.89 | 2 | 1436.86 |
| lizards | Diplodactylus_conspicillatus | Diplodactylus_conspicillatus | 0 | 5.5 | 1.99 | 1922008.31 |
| lizards | Diplodactylus_galaxias | Diplodactylus_galaxias | 0 | 1.91 | 2 | 62995.36 |
| lizards | Diplodactylus_galeatus | Diplodactylus_galeatus | 0 | 4.27 | 2 | 154489.23 |
| lizards | Diplodactylus_granariensis | Diplodactylus_granariensis | 0 | 7.41 | 2 | 578743.02 |
| lizards | Diplodactylus_klugei | Diplodactylus_klugei | 0 | 3.89 | 2 | 48287.79 |
| lizards | Diplodactylus_lateroides | Diplodactylus_lateroides | 0 | 2.63 | 2 | 16169.89 |
| lizards | Diplodactylus_mitchelli | Diplodactylus_mitchelli | 0 | 9.12 | 2 | 47363.12 |
| lizards | Diplodactylus_ornatus | Diplodactylus_ornatus | 0 | 3.89 | 2 | 63632.7 |
| lizards | Diplodactylus_pulcher | Diplodactylus_pulcher | 0 | 4.79 | 1.9 | 1324956.3 |
| lizards | Diplodactylus_savagei | Diplodactylus_savagei | 0 | 2.45 | 2 | 137750.77 |
| lizards | Diplodactylus_tessellatus | Diplodactylus_tessellatus | 0 | 4.07 | 2 | 2252041.59 |
| lizards | Diplodactylus_vittatus | Diplodactylus_vittatus | 0 | 4.17 | 2 | 1495846.67 |
| lizards | Diploglossus_delasagra | Diploglossus_delasagra | 0 | 30.2 | 4 | 16122.33 |
| lizards | Diploglossus_fasciatus | Diploglossus_fasciatus | 0 | 100 | 4 | 751798.1 |
| lizards | Diploglossus_garridoi | Diploglossus_garridoi | 0 | 17.38 | 4 | 84.79 |

| | | | | | | |
|---|---|---|---|---|---|---|
| lizards | Diploglossus_lessonae | Diploglossus_lessonae | 0 | 83.18 | 4.67 | 323282.89 |
| lizards | Diploglossus_monotropis | Diploglossus_monotropis | 0 | 223.87 | 5.5 | 329718.24 |
| lizards | Diploglossus_nigropunctatus | Diploglossus_nigropunctatus | 0 | 30.2 | 4.5 | 8067.39 |
| lizards | Diploglossus_pleii | Diploglossus_pleii | 0 | 81.28 | 3 | 4609.98 |
| lizards | Diplolaemus_bibronii | Diplolaemus_bibronii | 0 | 56.23 | 5 | 295294.09 |
| lizards | Diporiphora_albilabris | Diporiphora_albilabris | 0 | 6.61 | 3.7 | 532682.43 |
| lizards | Diporiphora_amphiboluroides | Diporiphora_amphiboluroides | 0 | 32.36 | 12 | 920616.43 |
| lizards | Diporiphora_australis | Diporiphora_australis | 0 | 14.79 | 4.5 | 935824.65 |
| lizards | Diporiphora_bennettii | Diporiphora_bennettii | 0 | 16.6 | 4.3 | 406968.26 |
| lizards | Diporiphora_bilineata | Diporiphora_bilineata | 0 | 8.32 | 5.5 | 2132931.32 |
| lizards | Diporiphora_nobbi | Diporiphora_nobbi | 0 | 19.5 | 6 | 1964903.24 |
| lizards | Diporiphora_paraconvergens | Diporiphora_paraconvergens | 0 | 6.92 | 3 | 852658 |
| lizards | Diporiphora_sobria | Diporiphora_arnhemica | 0 | 10.47 | 4.3 | 754640.1 |
| lizards | Diporiphora_superba | Diporiphora_superba | 0 | 28.18 | 6.5 | 70055.11 |
| lizards | Diporiphora_winneckei | Diporiphora_winneckei | 0 | 10.96 | 3 | 653992.83 |
| lizards | Dipsosaurus_dorsalis | Dipsosaurus_dorsalis | 0 | 158.49 | 4.5 | 456965.14 |
| lizards | Dixonius_hangseesom | Dixonius_hangseesom | 0 | 1.74 | 2 | 6426.52 |
| lizards | Dixonius_siamensis | Dixonius_siamensis | 0 | 4.07 | 1.93 | 788329.94 |
| lizards | Dixonius_vietnamensis | Dixonius_vietnamensis | 0 | 2.82 | 2 | 103532.32 |
| lizards | Dopasia_gracilis | Dopasia_gracilis | 0 | 134.9 | 5 | 1725322.86 |
| lizards | Dopasia_hainanensis | Dopasia_hainanensis | 0 | 75.86 | 21 | 5958.47 |
| lizards | Dopasia_harti | Dopasia_harti | 0 | 75.86 | 6 | 1636253.62 |
| lizards | Dopasia_sokolovi | Dopasia_sokolovi | 0 | 18.62 | 9 | 8994.15 |
| lizards | Dracaena_guianensis | Dracaena_guianensis | 0 | 2454.71 | 8.5 | 3151888.23 |
| lizards | Draco_beccarii | Draco_beccarii | 0 | 5.37 | 2.89 | 31194.77 |
| lizards | Draco_biaro | Draco_biaro | 0 | 4.57 | 2 | 26.31 |
| lizards | Draco_bimaculatus | Draco_bimaculatus | 0 | 4.07 | 2 | 62043.04 |
| lizards | Draco_blanfordii | Draco_blanfordii | 0 | 20.89 | 5.1 | 1393371.65 |
| lizards | Draco_boschmai | Draco_boschmai | 0 | 7.24 | 3.5 | 45435.85 |
| lizards | Draco_cornutus | Draco_cornutus | 0 | 7.59 | 3 | 734529.44 |
| lizards | Draco_cyanopterus | Draco_cyanopterus | 0 | 8.32 | 3.8 | 85923.13 |
| lizards | Draco_dussumieri | Draco_dussumieri | 0 | 9.33 | 3.5 | 178190.89 |
| lizards | Draco_fimbriatus | Draco_fimbriatus | 0 | 19.5 | 2.5 | 378214.49 |
| lizards | Draco_formosus | Draco_formosus | 0 | 12.59 | 3.5 | 99721.48 |

| | | | | | | |
|---|---|---|---|---|---|---|
| lizards | Draco_haematopogon | Draco_haematopogon | 0 | 10.47 | 2.5 | 1810765.36 |
| lizards | Draco_indochinensis | Draco_indochinensis | 0 | 14.13 | 3 | 56391.81 |
| lizards | Draco_lineatus | Draco_lineatus | 0 | 7.41 | 2.5 | 16010.16 |
| lizards | Draco_maculatus | Draco_maculatus | 0 | 8.51 | 3.6 | 2591726.25 |
| lizards | Draco_maximus | Draco_maximus | 0 | 28.84 | 4.5 | 427565.29 |
| lizards | Draco_melanopogon | Draco_melanopogon | 0 | 7.76 | 1.95 | 473941.95 |
| lizards | Draco_mindanensis | Draco_mindanensis | 0 | 12.02 | 2 | 112729.23 |
| lizards | Draco_modiglianii | Draco_modiglianii | 0 | 4.27 | 1.5 | 400.06 |
| lizards | Draco_obscurus | Draco_obscurus | 0 | 13.18 | 3 | 248262.76 |
| lizards | Draco_palawanensis | Draco_palawanensis | 0 | 6.17 | 3 | 11489.37 |
| lizards | Draco_quinquefasciatus | Draco_quinquefasciatus | 0 | 12.02 | 3.35 | 725920.55 |
| lizards | Draco_reticulatus | Draco_reticulatus | 0 | 19.5 | 2.5 | 23324.44 |
| lizards | Draco_rhytisma | Draco_rhytisma | 0 | 6.17 | 1.5 | 557.48 |
| lizards | Draco_spilonotus | Draco_spilonotus | 0 | 5.25 | 2.5 | 25739.59 |
| lizards | Draco_spilopterus | Draco_spilopterus | 0 | 10.23 | 4.5 | 142930.73 |
| lizards | Draco_sumatranus | Draco_sumatranus | 0 | 7.41 | 3.35 | 1021895.8 |
| lizards | Draco_taeniopterus | Draco_taeniopterus | 0 | 7.08 | 3 | 402687.72 |
| lizards | Draco_timoriensis | Draco_timorensis | 0 | 7.76 | 2.5 | 35281.88 |
| lizards | Draco_volans | Draco_volans | 0 | 8.51 | 4.5 | 115530.68 |
| lizards | Dravidogecko_anamallensis | Hemidactylus_anamallensis | 0 | 8.91 | 1.5 | 11511.34 |
| lizards | Dryadosaura_nordestina | Dryadosaura_nordestina | 0 | 4.17 | 2.5 | 108269.62 |
| lizards | Ebenavia_inunguis | Ebenavia_inunguis | 0 | 1.23 | 1.46 | 2096.98 |
| lizards | Ebenavia_maintimainty | Ebenavia_maintimainty | 1 | 0.37 | 1 | 2613.74 |
| lizards | Echinosaura_horrida | Echinosaura_horrida | 0 | 16.22 | 2 | 178484.41 |
| lizards | Egernia_cunninghami | Egernia_cunninghami | 0 | 446.68 | 4.4 | 439015.89 |
| lizards | Egernia_depressa | Egernia_depressa | 0 | 35.48 | 2 | 450583.58 |
| lizards | Egernia_formosa | Egernia_formosa | 0 | 26.92 | 3 | 316321.24 |
| lizards | Egernia_hosmeri | Egernia_hosmeri | 0 | 144.54 | 2 | 215412 |
| lizards | Egernia_kingii | Egernia_kingii | 0 | 416.87 | 5 | 96409.9 |
| lizards | Egernia_mcpheei | Egernia_mcpheei | 0 | 67.61 | 2 | 74857.14 |
| lizards | Egernia_napoleonis | Egernia_napoleonis | 0 | 53.7 | 2.5 | 206538.1 |
| lizards | Egernia_rugosa | Egernia_rugosa | 0 | 323.59 | 2.25 | 789989.54 |
| lizards | Egernia_saxatilis | Egernia_saxatilis | 0 | 56.23 | 2 | 250861.57 |
| lizards | Egernia_stokesii | Egernia_stokesii | 0 | 223.87 | 4.4 | 1248794.26 |

| | | | | | | |
|---|---|---|---|---|---|---|
| lizards | Egernia_striolata | Egernia_striolata | 0 | 38.02 | 3.8 | 1971703.67 |
| lizards | Elasmodactylus_tetensis | Elasmodactylus_tetensis | 0 | 19.05 | 2 | 86757.05 |
| lizards | Elasmodactylus_tuberculosus | Elasmodactylus_tuberculosus | 0 | 10 | 2 | 862531.25 |
| lizards | Elgaria_coerulea | Elgaria_coerulea | 0 | 45.71 | 5.4 | 551499.26 |
| lizards | Elgaria_kingii | Elgaria_kingii | 0 | 50.12 | 12 | 305414.67 |
| lizards | Elgaria_multicarinata | Elgaria_multicarinata | 0 | 117.49 | 10.9 | 217760.63 |
| lizards | Elgaria_panamintina | Elgaria_panamintina | 1 | 67.61 | 8 | 8268.71 |
| lizards | Elgaria_paucicarinata | Elgaria_paucicarinata | 0 | 29.51 | 7 | 9952.26 |
| lizards | Emoia_adspersa | Emoia_adspersa | 1 | 16.98 | 1.9 | 3024.59 |
| lizards | Emoia_aenea | Emoia_aenea | 0 | 7.41 | 2 | 48412.52 |
| lizards | Emoia_aneityumensis | Emoia_aneityumensis | 1 | 18.62 | 4.5 | 715.67 |
| lizards | Emoia_arnoensis | Emoia_arnoensis | 0 | 22.39 | 2 | 1195.76 |
| lizards | Emoia_atrocostata | Emoia_atrocostata | 0 | 21.38 | 1.95 | 2079573.15 |
| lizards | Emoia_aurulenta | Emoia_aurulenta | 0 | 2.24 | 2 | 69136.42 |
| lizards | Emoia_baudini | Emoia_baudini | 0 | 6.17 | 2 | 3102.97 |
| lizards | Emoia_bismarckensis | Emoia_bismarckensis | 0 | 7.41 | 2 | 19861.71 |
| lizards | Emoia_boettgeri | Emoia_boettgeri | 1 | 9.33 | 2 | 734.39 |
| lizards | Emoia_bogerti | Emoia_bogerti | 0 | 3.89 | 2 | 44810.71 |
| lizards | Emoia_brongersmai | Emoia_brongersmai | 0 | 6.76 | 2 | 3036.33 |
| lizards | Emoia_caeruleocauda | Emoia_caeruleocauda | 0 | 5.62 | 1.83 | 1089921.57 |
| lizards | Emoia_callisticta | Emoia_callisticta | 0 | 3.31 | 2 | 28982.58 |
| lizards | Emoia_campbelli | Emoia_campbelli | 1 | 19.95 | 2 | 9.99 |
| lizards | Emoia_concolor | Emoia_concolor | 0 | 18.2 | 2 | 9087.88 |
| lizards | Emoia_cyanogaster | Emoia_cyanogaster | 0 | 20.89 | 2 | 43311.81 |
| lizards | Emoia_cyanura | Emoia_cyanura | 0 | 5.37 | 1.98 | 93259.44 |
| lizards | Emoia_guttata | Emoia_guttata | 0 | 7.76 | 2 | 405.66 |
| lizards | Emoia_impar | Emoia_impar | 0 | 2.29 | 1.5 | 53986.92 |
| lizards | Emoia_irianensis | Emoia_irianensis | 0 | 5.01 | 2 | 9090.14 |
| lizards | Emoia_isolata | Emoia_isolata | 0 | 4.17 | 2 | 20.61 |
| lizards | Emoia_jakati | Emoia_jakati | 0 | 2.82 | 1.9 | 479940.33 |
| lizards | Emoia_kordoana | Emoia_kordoana | 0 | 4.37 | 2 | 398786.02 |
| lizards | Emoia_longicauda | Emoia_longicauda | 0 | 21.38 | 2 | 612690.27 |
| lizards | Emoia_loveridgei | Emoia_loveridgei | 0 | 1.74 | 2 | 36219.5 |
| lizards | Emoia_loyaltiensis | Emoia_loyaltiensis | 1 | 12.02 | 3.5 | 1845.67 |

| | | | | | | |
|---|---|---|---|---|---|---|
| lizards | Emoia_maculata | Emoia_maculata | 0 | 6.76 | 2 | 3100.33 |
| lizards | Emoia_maxima | Emoia_maxima | 0 | 1.86 | 2 | 29032.9 |
| lizards | Emoia_montana | Emoia_montana | 0 | 6.92 | 2 | 989.53 |
| lizards | Emoia_nigra | Emoia_nigra | 0 | 47.86 | 2.61 | 64085.69 |
| lizards | Emoia_nigromarginata | Emoia_nigromarginata | 0 | 10.47 | 2.5 | 8045.27 |
| lizards | Emoia_obscura | Emoia_obscura | 0 | 5.01 | 2 | 272171.72 |
| lizards | Emoia_oribata | Emoia_oribata | 0 | 9.33 | 2 | 30357.23 |
| lizards | Emoia_pallidiceps | Emoia_pallidiceps | 0 | 4.47 | 1.95 | 331355.8 |
| lizards | Emoia_parkeri | Emoia_parkeri | 1 | 2.88 | 2 | 11496.22 |
| lizards | Emoia_physicae | Emoia_physicae | 0 | 9.55 | 2 | 717251.46 |
| lizards | Emoia_physicina | Emoia_physicina | 0 | 2.4 | 2 | 109472.06 |
| lizards | Emoia_ponapea | Emoia_ponapea | 1 | 2.4 | 2 | 342.82 |
| lizards | Emoia_popei | Emoia_popei | 0 | 5.37 | 2 | 38743.34 |
| lizards | Emoia_pseudocyanura | Emoia_pseudocyanura | 0 | 6.76 | 2 | 21644.99 |
| lizards | Emoia_pseudopallidiceps | Emoia_pseudopallidiceps | 0 | 5.13 | 2 | 44259.14 |
| lizards | Emoia_reimschiisseli | Emoia_reimschiisseli | 0 | 9.55 | 2 | 192719.71 |
| lizards | Emoia_rennellensis | Emoia_rennellensis | 0 | 1.62 | 2 | 675.7 |
| lizards | Emoia_ruficauda | Emoia_ruficauda | 0 | 2.95 | 2 | 9745.73 |
| lizards | Emoia_samoensis | Emoia_samoensis | 1 | 37.15 | 5.3 | 2871.55 |
| lizards | Emoia_sanfordi | Emoia_sanfordi | 0 | 34.67 | 5 | 9879.17 |
| lizards | Emoia_schmidti | Emoia_schmidti | 0 | 4.9 | 2 | 2051.97 |
| lizards | Emoia_similis | Emoia_similis | 0 | 1.62 | 2 | 12873.41 |
| lizards | Emoia_slevini | Emoia_slevini | 1 | 12.88 | 2 | 956.86 |
| lizards | Emoia_sorex | Emoia_sorex | 0 | 3.98 | 2 | 192282.17 |
| lizards | Emoia_submetallica | Emoia_submetallica | 0 | 5.25 | 2 | 14091.82 |
| lizards | Emoia_tongana | Emoia_tongana | 0 | 8.51 | 2 | 2142.27 |
| lizards | Emoia_tropidolepis | Emoia_tropidolepis | 0 | 7.94 | 2 | 94680.79 |
| lizards | Emoia_trossula | Emoia_trossula | 1 | 28.18 | 3.5 | 13927.56 |
| lizards | Emoia_veracunda | Emoia_veracunda | 0 | 2.69 | 2 | 138722.05 |
| lizards | Enyalioides_cofanorum | Enyalioides_cofanorum | 0 | 85.11 | 3 | 358695.67 |
| lizards | Enyalioides_heterolepis | Enyalioides_heterolepis | 0 | 81.28 | 4 | 165895.82 |
| lizards | Enyalioides_laticeps | Enyalioides_laticeps | 0 | 123.03 | 6.25 | 1134082.42 |
| lizards | Enyalioides_palpebralis | Enyalioides_palpebralis | 0 | 53.7 | 3 | 243180.16 |
| lizards | Enyalioides_praestabilis | Enyalioides_praestabilis | 0 | 63.1 | 5 | 137697.66 |

| | | | | | | |
|---|---|---|---|---|---|---|
| lizards | Enyalioides_rubrigularis | Enyalioides_rubrigularis | 1 | 66.07 | 3 | 1002.19 |
| lizards | Enyalius_bibronii | Enyalius_bibronii | 0 | 56.23 | 7.24 | 982903.85 |
| lizards | Enyalius_bilineatus | Enyalius_bilineatus | 0 | 35.48 | 4.4 | 548759.19 |
| lizards | Enyalius_brasiliensis | Enyalius_brasiliensis | 0 | 51.29 | 9.5 | 44732.73 |
| lizards | Enyalius_catenatus | Enyalius_catenatus | 0 | 61.66 | 10 | 133671.26 |
| lizards | Enyalius_iheringii | Enyalius_iheringii | 0 | 63.1 | 13.65 | 221445.83 |
| lizards | Enyalius_leechii | Enyalius_leechii | 0 | 48.98 | 12.65 | 1438882.26 |
| lizards | Enyalius_perditus | Enyalius_perditus | 0 | 30.2 | 8.65 | 450100.05 |
| lizards | Epibator_greeri | Lioscincus_greeri | 1 | 4.37 | 2 | 77.05 |
| lizards | Epibator_nigrofasciolatus | Lioscincus_nigrofasciolatum | 0 | 30.9 | 7.5 | 18806.72 |
| lizards | Eremias_acutirostris | Eremias_acutirostris | 0 | 11.75 | 4.5 | 213043.53 |
| lizards | Eremias_argus | Eremias_argus | 0 | 7.24 | 3.25 | 4271177.41 |
| lizards | Eremias_arguta | Eremias_arguta | 0 | 22.91 | 3.98 | 4578499.4 |
| lizards | Eremias_brenchleyi | Eremias_brenchleyi | 0 | 7.59 | 4 | 600869.29 |
| lizards | Eremias_buechneri | Eremias_buechneri | 0 | 6.76 | 2.5 | 14695.4 |
| lizards | Eremias_grammica | Eremias_grammica | 0 | 22.91 | 3.5 | 1823617.28 |
| lizards | Eremias_intermedia | Eremias_intermedia | 0 | 7.59 | 2 | 2024194.59 |
| lizards | Eremias_lineolata | Eremias_lineolata | 0 | 3.98 | 2 | 1173018.61 |
| lizards | Eremias_multiocellata | Eremias_multiocellata | 0 | 10.96 | 2.85 | 4762088.33 |
| lizards | Eremias_nigrocellata | Eremias_nigrocellata | 0 | 13.18 | 5.5 | 322777.75 |
| lizards | Eremias_nikolskii | Eremias_nikolskii | 0 | 9.77 | 3.5 | 115485.94 |
| lizards | Eremias_persica | Eremias_persica | 0 | 21.38 | 5 | 1686829.41 |
| lizards | Eremias_pleskei | Eremias_pleskei | 1 | 5.01 | 4 | 47164.58 |
| lizards | Eremias_przewalskii | Eremias_przewalskii | 0 | 21.38 | 3.55 | 2448607.52 |
| lizards | Eremias_regeli | Eremias_regeli | 0 | 7.94 | 2.5 | 375839.71 |
| lizards | Eremias_scripta | Eremias_scripta | 0 | 6.76 | 1.5 | 1835545.75 |
| lizards | Eremias_strauchi | Eremias_strauchi | 0 | 11.75 | 8 | 309010.54 |
| lizards | Eremias_stummeri | Eremias_stummeri | 0 | 7.08 | 3.5 | 17321.55 |
| lizards | Eremias_suphani | Eremias_suphani | 0 | 8.32 | 3.5 | 10976.03 |
| lizards | Eremias_velox | Eremias_velox | 0 | 19.05 | 4 | 4796912.23 |
| lizards | Eremias_vermiculata | Eremias_vermiculata | 0 | 8.51 | 1.5 | 2114242.45 |
| lizards | Eremias_yarkandensis | Eremias_yarkandensis | 0 | 5.25 | 4 | 173393.29 |
| lizards | Eremiascincus_douglasi | Eremiascincus_douglasi | 0 | 11.22 | 4.3 | 203711.03 |
| lizards | Eremiascincus_emigrans | Eremiascincus_emigrans | 0 | 6.31 | 1 | 2849.39 |

| | | | | | | |
|---|---|---|---|---|---|---|
| lizards | Eremiascincus_fasciolatus | Eremiascincus_fasciolatus | 0 | 42.66 | 3.9 | 435748.86 |
| lizards | Eremiascincus_intermedius | Eremiascincus_intermedius | 0 | 14.45 | 4.5 | 778700.4 |
| lizards | Eremiascincus_isolepis | Eremiascincus_isolepis | 0 | 8.51 | 4.99 | 1417339.73 |
| lizards | Eremiascincus_musivus | Eremiascincus_musivus | 0 | 3.98 | 3 | 155315.14 |
| lizards | Eremiascincus_pallidus | Eremiascincus_pallidus | 0 | 9.33 | 1 | 1858227.79 |
| lizards | Eremiascincus_pardalis | Eremiascincus_pardalis | 0 | 8.51 | 4.4 | 226382.77 |
| lizards | Eremiascincus_phantasmus | Eremiascincus_phantasmus | 0 | 18.2 | 3.3 | 950164.67 |
| lizards | Eremiascincus_richardsonii | Eremiascincus_richardsonii | 0 | 46.77 | 5 | 5310086.51 |
| lizards | Eroticoscincus_graciloides | Eroticoscincus_graciloides | 0 | 0.72 | 4 | 26757.3 |
| lizards | Eublepharis_fuscus | Eublepharis_fuscus | 0 | 416.87 | 2 | 76785.14 |
| lizards | Eublepharis_hardwickii | Eublepharis_hardwickii | 0 | 29.51 | 1.5 | 284089.62 |
| lizards | Eublepharis_macularius | Eublepharis_macularius | 0 | 114.82 | 2.38 | 1631437.08 |
| lizards | Eublepharis_turcmenicus | Eublepharis_turcmenicus | 0 | 64.57 | 2 | 70844.55 |
| lizards | Eugongylus_albofasciolatus | Eugongylus_albofasciolatus | 0 | 245.47 | 3.5 | 88884.31 |
| lizards | Eugongylus_rufescens | Eugongylus_rufescens | 0 | 117.49 | 3.3 | 830810.75 |
| lizards | Eulamprus_heatwolei | Eulamprus_heatwolei | 0 | 22.39 | 3.55 | 283848.25 |
| lizards | Eulamprus_kosciuskoi | Eulamprus_kosciuskoi | 0 | 12.88 | 3.1 | 34679.7 |
| lizards | Eulamprus_leuraensis | Eulamprus_leuraensis | 1 | 12.88 | 2.67 | 1217.18 |
| lizards | Eulamprus_quoyii | Eulamprus_quoyii | 0 | 46.77 | 5.35 | 1004641.98 |
| lizards | Eulamprus_tympanum | Eulamprus_tympanum | 0 | 21.38 | 3.28 | 243512.56 |
| lizards | Euleptes_europaea | Euleptes_europaea | 0 | 2.09 | 1.75 | 33808.59 |
| lizards | Eumeces_algeriensis | Eumeces_algeriensis | 0 | 234.42 | 11 | 294432.77 |
| lizards | Eumeces_blythianus | Eumeces_blythianus | 0 | 30.2 | 6.3 | 1068638.79 |
| lizards | Eumeces_schneiderii | Eumeces_schneideri | 0 | 158.49 | 11 | 2484338.48 |
| lizards | Eumecia_anchietae | Eumecia_anchietae | 0 | 81.28 | 6.5 | 1405431.33 |
| lizards | Eurolophosaurus_amathites | Eurolophosaurus_amathites | 1 | 12.59 | 2 | 1499.31 |
| lizards | Eurolophosaurus_nanuzae | Eurolophosaurus_nanuzae | 0 | 9.77 | 2.04 | 49868.49 |
| lizards | Eurydactylodes_agricolae | Eurydactylodes_agricolae | 0 | 4.27 | 1.92 | 3524.97 |
| lizards | Eurydactylodes_occidentalis | Eurydactylodes_occidentalis | 1 | 3.02 | 2 | 279.85 |
| lizards | Eurydactylodes_symmetricus | Eurydactylodes_symmetricus | 0 | 4.37 | 2 | 3621.01 |
| lizards | Eurydactylodes_vieillardi | Eurydactylodes_vieillardi | 0 | 4.37 | 2 | 12846.36 |
| lizards | Eurylepis_taeniolata | Eurylepis_taeniolata | 0 | 138.04 | 5.5 | 1826116.58 |
| lizards | Euspondylus_acutirostris | Euspondylus_acutirostris | 0 | 3.89 | 1.5 | 5017.04 |
| lizards | Eutropis_beddomei | Eutropis_beddomii | 0 | 4.17 | 2.5 | 269543.69 |

| | | | | | | |
|---|---|---|---|---|---|---|
| lizards | Eutropis_carinata | Eutropis_carinata | 0 | 97.72 | 5.25 | 3483962.35 |
| lizards | Eutropis_clivicola | Eutropis_clivicola | 1 | 4.9 | 3 | 5640.2 |
| lizards | Eutropis_cumingi | Eutropis_cumingi | 0 | 2.95 | 2 | 89632 |
| lizards | Eutropis_indeprensa | Eutropis_indeprensa | 0 | 5.89 | 2.5 | 169835.35 |
| lizards | Eutropis_longicaudata | Eutropis_longicaudata | 0 | 63.1 | 6.68 | 1965017.29 |
| lizards | Eutropis_macularia | Eutropis_macularia | 0 | 9.33 | 2.15 | 4896988.01 |
| lizards | Eutropis_multicarinata | Eutropis_multicarinata | 0 | 19.5 | 2.42 | 247749.89 |
| lizards | Eutropis_multifasciata | Eutropis_multifasciata | 0 | 63.1 | 4.9 | 6105593.02 |
| lizards | Eutropis_quadricarinata | Eutropis_quadricarinata | 0 | 2.4 | 3 | 33567.75 |
| lizards | Eutropis_rudis | Eutropis_rudis | 0 | 60.26 | 2.49 | 571636.04 |
| lizards | Eutropis_tammanna | Eutropis_tammanna | 0 | 3.47 | 2 | 8845.86 |
| lizards | Feylinia_elegans | Feylinia_elegans | 0 | 5.5 | 2 | 817762.91 |
| lizards | Feylinia_grandisquamis | Feylinia_grandisquamis | 0 | 5.37 | 2.75 | 586811.36 |
| lizards | Flexiseps_melanurus | Amphiglossus_melanurus | 0 | 30.9 | 6 | 35290.44 |
| lizards | Furcifer_antimena | Furcifer_antimena | 1 | 95.5 | 15.5 | 6350.71 |
| lizards | Furcifer_campani | Furcifer_campani | 1 | 8.91 | 15 | 20927.47 |
| lizards | Furcifer_cephalolepis | Furcifer_cephalolepis | 0 | 11.48 | 5.5 | 1031.55 |
| lizards | Furcifer_labordi | Furcifer_labordi | 1 | 64.57 | 9 | 16997.63 |
| lizards | Furcifer_lateralis | Furcifer_lateralis | 0 | 56.23 | 13.5 | 467150.96 |
| lizards | Furcifer_major | Furcifer_major | 0 | 35.48 | 13.5 | 122786.39 |
| lizards | Furcifer_minor | Furcifer_minor | 1 | 22.91 | 12 | 7309.42 |
| lizards | Furcifer_oustaleti | Furcifer_oustaleti | 0 | 436.52 | 42 | 591142.81 |
| lizards | Furcifer_pardalis | Furcifer_pardalis | 0 | 269.15 | 30 | 109567.08 |
| lizards | Furcifer_petteri | Furcifer_petteri | 1 | 17.38 | 13.5 | 11024.04 |
| lizards | Furcifer_polleni | Furcifer_polleni | 0 | 25.7 | 9 | 384.22 |
| lizards | Furcifer_timoni | Furcifer_timoni | 0 | 20.89 | 12.33 | 1226.49 |
| lizards | Furcifer_verrucosus | Furcifer_verrucosus | 0 | 316.23 | 34.5 | 352674.7 |
| lizards | Furcifer_willsii | Furcifer_willsii | 0 | 13.49 | 11.5 | 100349.27 |
| lizards | Gallotia_atlantica | Gallotia_atlantica | 0 | 28.84 | 2.2 | 2586.7 |
| lizards | Gallotia_bravoana | Gallotia_bravoana | 1 | 208.93 | 5 | 108.5 |
| lizards | Gallotia_caesaris | Gallotia_caesaris | 0 | 30.9 | 3.15 | 660.38 |
| lizards | Gallotia_galloti | Gallotia_galloti | 0 | 69.18 | 4.1 | 2744.69 |
| lizards | Gallotia_intermedia | Gallotia_intermedia | 1 | 117.49 | 7.4 | 161.34 |
| lizards | Gallotia_simonyi | Gallotia_simonyi | 1 | 2691.53 | 10.15 | 190.64 |

| | | | | | | |
|---|---|---|---|---|---|---|
| lizards | Gallotia_stehlini | Gallotia_stehlini | 0 | 1096.48 | 10.05 | 1538.29 |
| lizards | Gambelia_copeii | Gambelia_copeii | 0 | 69.18 | 5 | 101483.76 |
| lizards | Gambelia_sila | Gambelia_sila | 1 | 57.54 | 3.15 | 37816.3 |
| lizards | Gambelia_wislizenii | Gambelia_wislizenii | 0 | 74.13 | 5.14 | 1387973.49 |
| lizards | Garthia_gaudichaudii | Garthia_gaudichaudii | 0 | 2.75 | 1.5 | 69245.95 |
| lizards | Gastropholis_prasina | Gastropholis_prasina | 0 | 30.2 | 5 | 10592.74 |
| lizards | Geckolepis_maculata | Geckolepis_maculata | 0 | 10 | 1.5 | 314848.15 |
| lizards | Geckolepis_typica | Geckolepis_typica | 0 | 5.37 | 2 | 186421.54 |
| lizards | Gehyra_australis | Gehyra_australis | 0 | 10.72 | 2 | 926589.68 |
| lizards | Gehyra_baliola | Gehyra_baliola | 0 | 21.88 | 2 | 197757.63 |
| lizards | Gehyra_borroloola | Gehyra_borroloola | 0 | 6.92 | 2 | 79273.85 |
| lizards | Gehyra_brevipalmata | Gehyra_brevipalmata | 0 | 13.8 | 1.5 | 454.79 |
| lizards | Gehyra_catenata | Gehyra_catenata | 0 | 3.63 | 2 | 355339.28 |
| lizards | Gehyra_dubia | Gehyra_dubia | 0 | 10.23 | 2 | 1500034.12 |
| lizards | Gehyra_georgpotthasti | Gehyra_georgpotthasti | 1 | 50.12 | 2 | 13053.85 |
| lizards | Gehyra_kimberleyi | Gehyra_kimberleyi | 0 | 4.9 | 1 | 381137.2 |
| lizards | Gehyra_koira | Gehyra_koira | 0 | 16.98 | 2 | 314185.04 |
| lizards | Gehyra_lacerata | Gehyra_lacerata | 0 | 4.47 | 2 | 14777.37 |
| lizards | Gehyra_marginata | Gehyra_marginata | 0 | 50.12 | 2 | 22924.93 |
| lizards | Gehyra_minuta | Gehyra_minuta | 0 | 2.09 | 1 | 119008.31 |
| lizards | Gehyra_montium | Gehyra_montium | 0 | 2.95 | 1.5 | 245462.19 |
| lizards | Gehyra_moritzi | Gehyra_moritzi | 0 | 2.63 | 2 | 187148.16 |
| lizards | Gehyra_multiporosa | Gehyra_multiporosa | 0 | 3.63 | 1 | 23838.19 |
| lizards | Gehyra_mutilata | Gehyra_mutilata | 0 | 5.5 | 1.8 | 3740611.78 |
| lizards | Gehyra_nana | Gehyra_nana | 0 | 3.47 | 1.5 | 570543.74 |
| lizards | Gehyra_oceanica | Gehyra_oceanica | 0 | 19.95 | 1.8 | 84829.7 |
| lizards | Gehyra_pilbara | Gehyra_pilbara | 0 | 6.03 | 1 | 1782077.77 |
| lizards | Gehyra_punctata | Gehyra_punctata | 0 | 5.75 | 1.5 | 600961.03 |
| lizards | Gehyra_purpurascens | Gehyra_purpurascens | 0 | 6.31 | 1.15 | 3319362.41 |
| lizards | Gehyra_robusta | Gehyra_robusta | 0 | 8.71 | 2 | 163324.91 |
| lizards | Gehyra_spheniscus | Gehyra_spheniscus | 0 | 2.24 | 1 | 29812.78 |
| lizards | Gehyra_variegata | Gehyra_variegata | 0 | 7.41 | 1 | 1717652.02 |
| lizards | Gehyra_versicolor | Gehyra_versicolor | 0 | 4.27 | 1 | 2543791.14 |
| lizards | Gehyra_vorax | Gehyra_vorax | 0 | 109.65 | 2 | 17009.28 |

| | | | | | | |
|---|---|---|---|---|---|---|
| lizards | Gehyra_xenopus | Gehyra_xenopus | 0 | 10 | 2 | 35702.29 |
| lizards | Gekko_badenii | Gekko_badenii | 1 | 47.86 | 2 | 25425.09 |
| lizards | Gekko_canaensis | Gekko_canaensis | 0 | 28.18 | 2 | 5.69 |
| lizards | Gekko_chinensis | Gekko_chinensis | 0 | 10.23 | 2 | 842949.14 |
| lizards | Gekko_ernstkelleri | Gekko_ernstkelleri | 0 | 15.14 | 2 | 540.44 |
| lizards | Gekko_gecko | Gekko_gecko | 0 | 169.82 | 2 | 4656309.15 |
| lizards | Gekko_hokouensis | Gekko_hokouensis | 0 | 12.3 | 2 | 651203.4 |
| lizards | Gekko_horsfieldii | Ptychozoon_horsfieldii | 0 | 16.98 | 2 | 38301.3 |
| lizards | Gekko_intermedium | Ptychozoon_intermedium | 0 | 19.05 | 2 | 54309.98 |
| lizards | Gekko_japonicus | Gekko_japonicus | 0 | 12.02 | 1.95 | 2304586.03 |
| lizards | Gekko_kikuchii | Gekko_kikuchii | 0 | 10.23 | 2 | 9799.19 |
| lizards | Gekko_kuhli | Ptychozoon_kuhli | 0 | 23.44 | 2 | 1423689.16 |
| lizards | Gekko_lionotum | Ptychozoon_lionotum | 0 | 20.89 | 2 | 1406078.46 |
| lizards | Gekko_mindorensis | Gekko_mindorensis | 0 | 13.49 | 2 | 55609.92 |
| lizards | Gekko_monarchus | Gekko_monarchus | 0 | 19.95 | 1.95 | 1740240.71 |
| lizards | Gekko_nicobarensis | Ptychozoon_nicobarensis | 1 | 19.05 | 2 | 620.15 |
| lizards | Gekko_nutaphandi | Gekko_nutaphandi | 0 | 29.51 | 2 | 19.98 |
| lizards | Gekko_palmatus | Gekko_palmatus | 0 | 10.23 | 1.5 | 69777.32 |
| lizards | Gekko_petricolus | Gekko_petricolus | 0 | 20.89 | 2 | 16575.65 |
| lizards | Gekko_russelltraini | Gekko_russelltraini | 1 | 11.48 | 2 | 23.25 |
| lizards | Gekko_scientiadventura | Gekko_scientiadventura | 0 | 7.94 | 2 | 2736.16 |
| lizards | Gekko_shibatai | Gekko_shibatai | 1 | 8.51 | 2 | 6.48 |
| lizards | Gekko_siamensis | Gekko_siamensis | 0 | 58.88 | 2 | 5157.35 |
| lizards | Gekko_smithii | Gekko_smithii | 0 | 114.82 | 2 | 1477995.91 |
| lizards | Gekko_subpalmatus | Gekko_subpalmatus | 0 | 10.96 | 2 | 1126594.91 |
| lizards | Gekko_swinhonis | Gekko_swinhonis | 1 | 8.32 | 2 | 1375479.65 |
| lizards | Gekko_takouensis | Gekko_takouensis | 1 | 22.91 | 2 | 9.99 |
| lizards | Gekko_vittatus | Gekko_vittatus | 0 | 47.86 | 2 | 729666.63 |
| lizards | Gekko_yakuensis | Gekko_yakuensis | 0 | 7.76 | 1.5 | 8949.28 |
| lizards | Gelanesaurus_cochranae | Potamites_cochranae | 0 | 19.5 | 2 | 66322.65 |
| lizards | Geomyersia_coggeri | Geomyersia_coggeri | 0 | 0.66 | 1 | 1819.49 |
| lizards | Geomyersia_glabra | Geomyersia_glabra | 0 | 0.79 | 1 | 8778.26 |
| lizards | Gerrhonotus_infernalis | Gerrhonotus_infernalis | 0 | 186.21 | 18.8 | 333221.59 |
| lizards | Gerrhonotus_liocephalus | Gerrhonotus_liocephalus | 0 | 186.21 | 10.7 | 365465.4 |

| | | | | | | |
|---|---|---|---|---|---|---|
| lizards | Gerrhonotus_lugoi | Gerrhonotus_lugoi | 0 | 17.78 | 4 | 1486.11 |
| lizards | Gerrhonotus_ophiurus | Gerrhonotus_ophiurus | 0 | 173.78 | 10.7 | 84300.56 |
| lizards | Gerrhonotus_parvus | Gerrhonotus_parvus | 1 | 8.71 | 7 | 6276.42 |
| lizards | Gerrhosaurus_auritus | Gerrhosaurus_auritus | 0 | 257.04 | 12 | 612172.21 |
| lizards | Gerrhosaurus_bulsi | Gerrhosaurus_bulsi | 0 | 208.93 | 6.5 | 839027.36 |
| lizards | Gerrhosaurus_flavigularis | Gerrhosaurus_flavigularis | 0 | 79.43 | 4.5 | 2867547.1 |
| lizards | Gerrhosaurus_multilineatus | Gerrhosaurus_multilineatus | 0 | 257.04 | 8 | 879482.26 |
| lizards | Gerrhosaurus_nigrolineatus | Gerrhosaurus_nigrolineatus | 0 | 158.49 | 6.5 | 4359388.44 |
| lizards | Gerrhosaurus_skoogi | Gerrhosaurus_skoogi | 0 | 104.71 | 2 | 33736.49 |
| lizards | Gerrhosaurus_typicus | Gerrhosaurus_typicus | 0 | 69.18 | 4 | 122707.1 |
| lizards | Glaphyromorphus_fuscicaudis | Glaphyromorphus_fuscicaudis | 0 | 15.85 | 3 | 18803.06 |
| lizards | Glaphyromorphus_nyanchupinta | Glaphyromorphus_nyanchupinta | 0 | 2.88 | 2 | 247.78 |
| lizards | Glaphyromorphus_pumilus | Glaphyromorphus_pumilus | 0 | 3.09 | 3 | 137312 |
| lizards | Glaphyromorphus_punctulatus | Glaphyromorphus_punctulatus | 0 | 6.76 | 3 | 201931.6 |
| lizards | Glaucomastix_abaetensis | Ameivula_abaetensis | 1 | 10.72 | 1.4 | 249.56 |
| lizards | Glaucomastix_littoralis | Ameivula_littoralis | 1 | 15.85 | 5.2 | 5242.46 |
| lizards | Goggia_braacki | Goggia_braacki | 0 | 1.02 | 2 | 1929.78 |
| lizards | Goggia_gemmula | Goggia_gemmula | 0 | 0.69 | 2 | 2921.46 |
| lizards | Goggia_hewitti | Goggia_hewitti | 0 | 1.26 | 2 | 10782.19 |
| lizards | Goggia_lineata | Goggia_lineata | 0 | 0.81 | 2 | 58711.37 |
| lizards | Gonatodes_albogularis | Gonatodes_albogularis | 0 | 2.75 | 1 | 873793.04 |
| lizards | Gonatodes_alexandermendesi | Gonatodes_alexandermendesi | 0 | 2.88 | 2 | 40483.84 |
| lizards | Gonatodes_annularis | Gonatodes_annularis | 0 | 4.07 | 2 | 1140558.82 |
| lizards | Gonatodes_antillensis | Gonatodes_antillensis | 0 | 1.35 | 1 | 99.47 |
| lizards | Gonatodes_caudiscutatus | Gonatodes_caudiscutatus | 0 | 2.29 | 1 | 106052.22 |
| lizards | Gonatodes_ceciliae | Gonatodes_ceciliae | 0 | 7.41 | 1 | 5164.75 |
| lizards | Gonatodes_concinnatus | Gonatodes_concinnatus | 0 | 3.55 | 1 | 851434.06 |
| lizards | Gonatodes_daudini | Gonatodes_daudini | 1 | 0.76 | 1 | 10.87 |
| lizards | Gonatodes_hasemani | Gonatodes_hasemani | 0 | 2.4 | 1 | 2613126.8 |
| lizards | Gonatodes_humeralis | Gonatodes_humeralis | 0 | 2.57 | 1.34 | 7917317.28 |
| lizards | Gonatodes_ocellatus | Gonatodes_ocellatus | 0 | 3.09 | 1 | 86.28 |
| lizards | Gonatodes_petersi | Gonatodes_petersi | 0 | 1.86 | 1 | 3100.38 |
| lizards | Gonatodes_superciliaris | Gonatodes_superciliaris | 0 | 2.88 | 1 | 14.31 |
| lizards | Gonatodes_taniae | Gonatodes_taniae | 0 | 3.47 | 1 | 840.39 |

| | | | | | | |
|---|---|---|---|---|---|---|
| lizards | Gonatodes_vittatus | Gonatodes_vittatus | 0 | 1.07 | 1.5 | 278124.21 |
| lizards | Gongylomorphus_bojerii | Gongylomorphus_bojerii | 1 | 7.41 | 2 | 1872.51 |
| lizards | Goniurosaurus_araneus | Goniurosaurus_araneus | 1 | 47.86 | 2 | 588.19 |
| lizards | Goniurosaurus_catbaensis | Goniurosaurus_catbaensis | 1 | 47.86 | 2 | 167.88 |
| lizards | Goniurosaurus_hainanensis | Goniurosaurus_hainanensis | 0 | 19.5 | 2 | 8319.64 |
| lizards | Goniurosaurus_kuroiwae | Goniurosaurus_kuroiwae | 1 | 20.42 | 2 | 1255.45 |
| lizards | Goniurosaurus_luii | Goniurosaurus_luii | 1 | 39.81 | 1.82 | 10484.79 |
| lizards | Goniurosaurus_orientalis | Goniurosaurus_orientalis | 1 | 16.98 | 2 | 3.48 |
| lizards | Goniurosaurus_splendens | Goniurosaurus_splendens | 1 | 8.71 | 2 | 248.88 |
| lizards | Goniurosaurus_toyamai | Goniurosaurus_toyamai | 1 | 11.75 | 2 | 18.93 |
| lizards | Goniurosaurus_yamashinae | Goniurosaurus_yamashinae | 1 | 13.49 | 2 | 58.12 |
| lizards | Goniurosaurus_yingdeensis | Goniurosaurus_yingdeensis | 1 | 17.78 | 2 | 9.99 |
| lizards | Gonocephalus_bellii | Gonocephalus_bellii | 0 | 123.03 | 4 | 62519.17 |
| lizards | Gonocephalus_bornensis | Gonocephalus_bornensis | 0 | 91.2 | 4 | 308153.19 |
| lizards | Gonocephalus_chamaeleontinus | Gonocephalus_chamaeleontinus | 0 | 173.78 | 4 | 233857.85 |
| lizards | Gonocephalus_doriae | Gonocephalus_doriae | 0 | 151.36 | 2 | 31728.3 |
| lizards | Gonocephalus_grandis | Gonocephalus_grandis | 0 | 144.54 | 3.59 | 1315206.16 |
| lizards | Gonocephalus_kuhlii | Gonocephalus_kuhlii | 1 | 223.87 | 4 | 12094.16 |
| lizards | Gonocephalus_lacunosus | Gonocephalus_lacunosus | 0 | 117.49 | 7.5 | 9.99 |
| lizards | Gonocephalus_liogaster | Gonocephalus_liogaster | 0 | 107.15 | 3.1 | 658393.54 |
| lizards | Gonocephalus_semperi | Gonocephalus_semperi | 0 | 50.12 | 3 | 273776.48 |
| lizards | Gonocephalus_sophiae | Gonocephalus_sophiae | 0 | 117.49 | 3.09 | 274059.4 |
| lizards | Gowidon_longirostris | Lophognathus_longirostris | 0 | 50.12 | 4.7 | 2537718.58 |
| lizards | Gymnodactylus_amarali | Gymnodactylus_amarali | 0 | 4.37 | 1.7 | 1124711.46 |
| lizards | Gymnodactylus_darwinii | Gymnodactylus_darwinii | 0 | 5.5 | 2 | 399674.46 |
| lizards | Gymnodactylus_geckoides | Gymnodactylus_geckoides | 0 | 4.37 | 1.13 | 539398.38 |
| lizards | Gymnophthalmus_speciosus | Gymnophthalmus_speciosus | 0 | 2.34 | 1.92 | 1096550.61 |
| lizards | Gymnophthalmus_underwoodi | Gymnophthalmus_underwoodi | 0 | 2.34 | 1.97 | 1215624.72 |
| lizards | Gymnophthalmus_vanzoi | Gymnophthalmus_vanzoi | 0 | 1.66 | 2 | 27581.73 |
| lizards | Haemodracon_riebeckii | Haemodracon_riebeckii | 0 | 69.18 | 1.82 | 3465.9 |
| lizards | Haemodracon_trachyrhinus | Haemodracon_trachyrhinus | 0 | 4.37 | 1 | 3228.85 |
| lizards | Harrisoniascincus_zia | Harrisoniascincus_zia | 0 | 3.89 | 5.45 | 13695.32 |
| lizards | Heliobolus_lugubris | Heliobolus_lugubris | 0 | 6.46 | 3.63 | 2200737.13 |
| lizards | Heliobolus_spekii | Heliobolus_spekii | 0 | 5.01 | 5 | 2240391.23 |

| | | | | | | |
|---|---|---|---|---|---|---|
| lizards | Hellenolacerta_graeca | Hellenolacerta_graeca | 0 | 14.13 | 3.5 | 21404.32 |
| lizards | Heloderma_horridum | Heloderma_horridum | 0 | 3019.95 | 8 | 171205.41 |
| lizards | Heloderma_suspectum | Heloderma_suspectum | 0 | 977.24 | 5.35 | 345298.44 |
| lizards | Hemicordylus_capensis | Hemicordylus_capensis | 0 | 36.31 | 2.25 | 72370.85 |
| lizards | Hemicordylus_nebulosus | Hemicordylus_nebulosus | 1 | 12.59 | 1.5 | 10.55 |
| lizards | Hemidactylus_albofasciatus | Hemidactylus_albofasciatus | 1 | 1.12 | 2 | 1814.63 |
| lizards | Hemidactylus_albopunctatus | Hemidactylus_albopunctatus | 0 | 3.72 | 2 | 8513.39 |
| lizards | Hemidactylus_angulatus | Hemidactylus_angulatus | 0 | 7.41 | 2 | 9223834.72 |
| lizards | Hemidactylus_aquilonius | Hemidactylus_aquilonius | 0 | 3.31 | 2 | 1034050.98 |
| lizards | Hemidactylus_beninensis | Hemidactylus_beninensis | 0 | 10.23 | 2 | 19.98 |
| lizards | Hemidactylus_bowringii | Hemidactylus_bowringii | 0 | 10.72 | 1.9 | 402160.74 |
| lizards | Hemidactylus_brasilianus | Hemidactylus_brasilianus | 0 | 5.5 | 1.89 | 925038.73 |
| lizards | Hemidactylus_brookii | Hemidactylus_brookii | 0 | 12.3 | 2 | 6273555.68 |
| lizards | Hemidactylus_coalescens | Hemidactylus_coalescens | 0 | 9.55 | 2 | 910124.81 |
| lizards | Hemidactylus_craspedotus | Hemidactylus_craspedotus | 0 | 5.75 | 2 | 125107.14 |
| lizards | Hemidactylus_depressus | Hemidactylus_depressus | 0 | 12.3 | 2 | 56020.99 |
| lizards | Hemidactylus_dracaenacolus | Hemidactylus_dracaenacolus | 1 | 8.71 | 2 | 94.14 |
| lizards | Hemidactylus_echinus | Hemidactylus_echinus | 0 | 7.08 | 2 | 29843.05 |
| lizards | Hemidactylus_fasciatus | Hemidactylus_fasciatus | 0 | 16.6 | 2 | 482648.75 |
| lizards | Hemidactylus_flaviviridis | Hemidactylus_flaviviridis | 0 | 16.6 | 2 | 3803737.44 |
| lizards | Hemidactylus_frenatus | Hemidactylus_frenatus | 0 | 6.31 | 1.95 | 7347932.06 |
| lizards | Hemidactylus_garnotii | Hemidactylus_garnotii | 0 | 6.61 | 2 | 4533420.36 |
| lizards | Hemidactylus_hajarensis | Hemidactylus_hajarensis | 0 | 6.31 | 2 | 12343.19 |
| lizards | Hemidactylus_homoeolepis | Hemidactylus_homoeolepis | 0 | 2.69 | 1.5 | 97776.82 |
| lizards | Hemidactylus_hunae | Hemidactylus_hunae | 1 | 38.02 | 2 | 7821.03 |
| lizards | Hemidactylus_imbricatus | Hemidactylus_imbricatus | 0 | 3.8 | 2 | 211884.84 |
| lizards | Hemidactylus_inexpectatus | Hemidactylus_inexpectatus | 0 | 2 | 2 | 15.96 |
| lizards | Hemidactylus_inintellectus | Hemidactylus_inintellectus | 0 | 5.5 | 2 | 2704.17 |
| lizards | Hemidactylus_isolepis | Hemidactylus_isolepis | 0 | 2.95 | 2 | 527262.52 |
| lizards | Hemidactylus_ituriensis | Hemidactylus_ituriensis | 0 | 13.8 | 2 | 1453333.8 |
| lizards | Hemidactylus_kamdemtohami | Hemidactylus_kamdemtohami | 0 | 7.41 | 2 | 132645.3 |
| lizards | Hemidactylus_leschenaultii | Hemidactylus_leschenaultii | 0 | 15.14 | 2 | 1578465.79 |
| lizards | Hemidactylus_longicephalus | Hemidactylus_longicephalus | 0 | 9.55 | 2 | 1525636.15 |
| lizards | Hemidactylus_mabouia | Hemidactylus_mabouia | 0 | 14.13 | 1.86 | 9561088.71 |

| | | | | | | |
|---|---|---|---|---|---|---|
| lizards | Hemidactylus_macropholis | Hemidactylus_macropholis | 0 | 14.79 | 2 | 955988.77 |
| lizards | Hemidactylus_maculatus | Hemidactylus_maculatus | 0 | 33.11 | 2 | 438618.32 |
| lizards | Hemidactylus_makolowodei | Hemidactylus_makolowodei | 1 | 19.05 | 2 | 19.98 |
| lizards | Hemidactylus_mercatorius | Hemidactylus_mercatorius | 0 | 4.47 | 2 | 647004.54 |
| lizards | Hemidactylus_muriceus | Hemidactylus_muriceus | 0 | 4.27 | 2 | 1805438.06 |
| lizards | Hemidactylus_palaichthus | Hemidactylus_palaichthus | 0 | 7.41 | 2 | 1432246.75 |
| lizards | Hemidactylus_parvimaculatus | Hemidactylus_parvimaculatus | 0 | 3.16 | 2 | 215161.38 |
| lizards | Hemidactylus_persicus | Hemidactylus_persicus | 0 | 9.12 | 2 | 1174736.64 |
| lizards | Hemidactylus_platycephalus | Hemidactylus_platycephalus | 0 | 16.22 | 2 | 2633418.31 |
| lizards | Hemidactylus_platyurus | Hemidactylus_platyurus | 0 | 6.92 | 2 | 6316105.58 |
| lizards | Hemidactylus_richardsonii | Hemidactylus_richardsonii | 0 | 10.23 | 2 | 530792.47 |
| lizards | Hemidactylus_robustus | Hemidactylus_robustus | 0 | 4.68 | 1.5 | 2651796.37 |
| lizards | Hemidactylus_ruspolii | Hemidactylus_ruspolii | 0 | 3.47 | 2 | 574666.46 |
| lizards | Hemidactylus_sataraensis | Hemidactylus_sataraensis | 1 | 2.29 | 2 | 59.65 |
| lizards | Hemidactylus_scabriceps | Hemidactylus_scabriceps | 1 | 2.4 | 2 | 49056 |
| lizards | Hemidactylus_squamulatus | Hemidactylus_squamulatus | 0 | 2.51 | 2 | 543958.5 |
| lizards | Hemidactylus_stejnegeri | Hemidactylus_stejnegeri | 0 | 4.57 | 1.5 | 19726.88 |
| lizards | Hemidactylus_tasmani | Hemidactylus_tasmani | 0 | 9.55 | 2 | 193903.73 |
| lizards | Hemidactylus_treutleri | Hemidactylus_treutleri | 0 | 7.24 | 2 | 419611.51 |
| lizards | Hemidactylus_triedrus | Hemidactylus_triedrus | 0 | 16.22 | 2 | 253557.23 |
| lizards | Hemidactylus_tropidolepis | Hemidactylus_tropidolepis | 0 | 3.24 | 2 | 322141.25 |
| lizards | Hemidactylus_turcicus | Hemidactylus_turcicus | 0 | 5.75 | 2 | 2226414 |
| lizards | Hemiergis_decresiensis | Hemiergis_decresiensis | 0 | 3.02 | 4.5 | 141899 |
| lizards | Hemiergis_gracilipes | Hemiergis_gracilipes | 0 | 14.79 | 3.2 | 30507.05 |
| lizards | Hemiergis_initialis | Hemiergis_initialis | 0 | 0.98 | 1.9 | 418696.12 |
| lizards | Hemiergis_millewae | Hemiergis_millewae | 0 | 3.72 | 2 | 279036.18 |
| lizards | Hemiergis_peronii | Hemiergis_peronii | 0 | 3.02 | 2.68 | 643819.97 |
| lizards | Hemiergis_talbingoensis | Hemiergis_talbingoensis | 0 | 1.86 | 3 | 282908.79 |
| lizards | Hemiphyllodactylus_aurantiacus | Hemiphyllodactylus_aurantiacus | 1 | 1.62 | 2 | 68026.08 |
| lizards | Hemiphyllodactylus_banaensis | Hemiphyllodactylus_banaensis | 0 | 2.95 | 2 | 9.99 |
| lizards | Hemiphyllodactylus_chiangmaiensis | Hemiphyllodactylus_chiangmaiensis | 0 | 1.66 | 2 | 19.98 |
| lizards | Hemiphyllodactylus_ganoklonis | Hemiphyllodactylus_ganoklonis | 0 | 0.98 | 1.5 | 424.34 |
| lizards | Hemiphyllodactylus_kiziriani | Hemiphyllodactylus_kiziriani | 0 | 1.62 | 2 | 12.51 |
| lizards | Hemiphyllodactylus_tehtarik | Hemiphyllodactylus_tehtarik | 0 | 1.55 | 2 | 9.99 |

| | | | | | | |
|---|---|---|---|---|---|---|
| lizards | Hemiphyllodactylus_typus | Hemiphyllodactylus_typus | 0 | 4.68 | 2 | 2660225.56 |
| lizards | Hemitheconyx_caudicinctus | Hemitheconyx_caudicinctus | 0 | 93.33 | 2 | 1712974.33 |
| lizards | Hemitheconyx_taylori | Hemitheconyx_taylori | 0 | 56.23 | 2 | 79546.17 |
| lizards | Heremites_auratus | Trachylepis_aurata | 0 | 33.88 | 5 | 554428.11 |
| lizards | Heremites_septemtaeniatus | Trachylepis_septemtaeniata | 0 | 25.12 | 4 | 1364784.6 |
| lizards | Heremites_vittatus | Trachylepis_vittata | 0 | 20.89 | 4.85 | 1163294.2 |
| lizards | Hesperoedura_reticulata | Hesperoedura_reticulata | 0 | 7.24 | 2.06 | 291832.37 |
| lizards | Heterodactylus_imbricatus | Heterodactylus_imbricatus | 0 | 36.31 | 1.5 | 262870.04 |
| lizards | Heteronotia_atra | Heteronotia_atra | 0 | 5.25 | 2 | 2236.83 |
| lizards | Heteronotia_binoei | Heteronotia_binoei | 0 | 3.63 | 1.85 | 6898166.2 |
| lizards | Heteronotia_planiceps | Heteronotia_planiceps | 0 | 3.63 | 2 | 226325.88 |
| lizards | Heteronotia_spelea | Heteronotia_spelea | 0 | 3.8 | 2 | 196701.97 |
| lizards | Holaspis_guentheri | Holaspis_guentheri | 0 | 3.55 | 2 | 2080975.37 |
| lizards | Holaspis_laevis | Holaspis_laevis | 0 | 2.75 | 2 | 546758.09 |
| lizards | Holbrookia_elegans | Holbrookia_elegans | 0 | 14.45 | 5 | 188396.54 |
| lizards | Holbrookia_lacerata | Holbrookia_lacerata | 0 | 12.59 | 8 | 267337.58 |
| lizards | Holbrookia_maculata | Holbrookia_maculata | 0 | 14.45 | 7.15 | 1870599.63 |
| lizards | Holbrookia_propinqua | Holbrookia_propinqua | 0 | 8.71 | 3.7 | 124685.75 |
| lizards | Holcosus_festivus | Holcosus_festivus | 0 | 93.33 | 2.75 | 455421.27 |
| lizards | Holcosus_niceforoi | Holcosus_niceforoi | 0 | 15.85 | 2.6 | 4698.21 |
| lizards | Holcosus_quadrilineatus | Holcosus_quadrilineatus | 0 | 19.95 | 2.05 | 41564.4 |
| lizards | Holcosus_undulatus | Holcosus_undulatus | 0 | 81.28 | 4.3 | 909409.79 |
| lizards | Holodactylus_africanus | Holodactylus_africanus | 0 | 12.02 | 2 | 325908.44 |
| lizards | Homonota_borellii | Homonota_borellii | 0 | 2 | 1 | 475244.79 |
| lizards | Homonota_darwinii | Homonota_darwinii | 0 | 4.37 | 1 | 656478.51 |
| lizards | Homonota_taragui | Homonota_taragui | 1 | 1.95 | 1 | 101.46 |
| lizards | Homonota_underwoodi | Homonota_underwoodi | 0 | 4.27 | 1 | 395255.54 |
| lizards | Homonota_uruguayensis | Homonota_uruguayensis | 1 | 3.31 | 1.5 | 167811.59 |
| lizards | Homonota_whitii | Homonota_whitii | 0 | 2.14 | 1 | 84524.98 |
| lizards | Homopholis_fasciata | Homopholis_fasciata | 0 | 10.96 | 2 | 1188680.91 |
| lizards | Homopholis_mulleri | Homopholis_mulleri | 0 | 10.72 | 1.5 | 15811.91 |
| lizards | Homopholis_walbergii | Homopholis_walbergii | 0 | 33.88 | 2 | 330104.02 |
| lizards | Hoplocercus_spinosus | Hoplocercus_spinosus | 0 | 33.11 | 4 | 2139549.62 |
| lizards | Hoplodactylus_duvaucelii | Hoplodactylus_duvaucelii | 0 | 93.33 | 2 | 86164.5 |

| | | | | | | |
|---|---|---|---|---|---|---|
| lizards | Hydrosaurus_amboinensis | Hydrosaurus_amboinensis | 0 | 1621.81 | 7.5 | 165540.29 |
| lizards | Hydrosaurus_pustulatus | Hydrosaurus_pustulatus | 0 | 812.83 | 7.5 | 160747.09 |
| lizards | Hypsilurus_longi | Hypsilurus_longi | 0 | 478.63 | 1.5 | 54905.9 |
| lizards | Hypsilurus_macrolepis | Hypsilurus_macrolepis | 0 | 95.5 | 2 | 15623.52 |
| lizards | Hypsilurus_magnus | Hypsilurus_magnus | 0 | 457.09 | 1 | 355591.22 |
| lizards | Hypsilurus_modestus | Hypsilurus_modestus | 0 | 41.69 | 1 | 653605.49 |
| lizards | Hypsilurus_papuensis | Hypsilurus_papuensis | 0 | 436.52 | 1 | 77835.64 |
| lizards | Hypsilurus_schoedei | Hypsilurus_schoedei | 0 | 72.44 | 1 | 451.14 |
| lizards | Iberolacerta_aranica | Iberolacerta_aranica | 1 | 6.92 | 2.99 | 59.95 |
| lizards | Iberolacerta_aurelioi | Iberolacerta_aurelioi | 1 | 6.46 | 2.52 | 122.45 |
| lizards | Iberolacerta_bonnali | Iberolacerta_bonnali | 0 | 6.46 | 3.03 | 1583.09 |
| lizards | Iberolacerta_cyreni | Iberolacerta_cyreni | 1 | 17.38 | 6.3 | 6606.26 |
| lizards | Iberolacerta_galani | Iberolacerta_galani | 0 | 14.13 | 4.5 | 858.91 |
| lizards | Iberolacerta_horvathi | Iberolacerta_horvathi | 0 | 6.46 | 3.51 | 27101.73 |
| lizards | Iberolacerta_martinezricai | Iberolacerta_martinezricai | 1 | 7.76 | 4 | 248.03 |
| lizards | Iberolacerta_monticola | Iberolacerta_monticola | 1 | 13.8 | 6.3 | 31589.4 |
| lizards | Ichnotropis_capensis | Ichnotropis_capensis | 0 | 7.24 | 6.14 | 4125084.82 |
| lizards | Ichnotropis_grandiceps | Ichnotropis_grandiceps | 0 | 7.94 | 7.5 | 93129.93 |
| lizards | Iguana_delicatissima | Iguana_delicatissima | 1 | 3467.37 | 17 | 4509.86 |
| lizards | Iguana_iguana | Iguana_iguana | 0 | 8128.31 | 28.8 | 12236032.36 |
| lizards | Insulasaurus_arborens | Insulasaurus_arborens | 0 | 5.75 | 1.5 | 28606.12 |
| lizards | Insulasaurus_traanorum | Insulasaurus_traanorum | 0 | 2.82 | 1 | 14.36 |
| lizards | Intellagama_lesueurii | Intellagama_lesueurii | 0 | 912.01 | 9.68 | 834282.1 |
| lizards | Iphisa_elegans | Iphisa_elegans | 0 | 5.75 | 2 | 6181116.34 |
| lizards | Isopachys_gyldenstolpei | Isopachys_gyldenstolpei | 0 | 15.14 | 5 | 8617.87 |
| lizards | Janetaescincus_braueri | Janetaescincus_braueri | 1 | 1.12 | 2 | 32.2 |
| lizards | Japalura_andersoniana | Japalura_andersoniana | 0 | 13.8 | 7 | 66509.27 |
| lizards | Japalura_kumaonensis | Japalura_kumaonensis | 0 | 11.22 | 10.4 | 136935.08 |
| lizards | Japalura_major | Japalura_major | 0 | 27.54 | 10 | 102211.67 |
| lizards | Japalura_variegata | Japalura_variegata | 0 | 61.66 | 16.33 | 39870.61 |
| lizards | Kanakysaurus_viviparus | Kanakysaurus_viviparus | 1 | 11.75 | 3 | 296.52 |
| lizards | Karusasaurus_jordani | Karusasaurus_jordani | 0 | 63.1 | 2.3 | 213618.21 |
| lizards | Karusasaurus_polyzonus | Karusasaurus_polyzonus | 0 | 45.71 | 2.43 | 754298.67 |
| lizards | Kentropyx_altamazonica | Kentropyx_altamazonica | 0 | 44.67 | 6.17 | 3922337.69 |

| | | | | | | |
|---|---|---|---|---|---|---|
| lizards | Kentropyx_borckiana | Kentropyx_borckiana | 0 | 30.9 | 6 | 211722.65 |
| lizards | Kentropyx_calcarata | Kentropyx_calcarata | 0 | 51.29 | 4.85 | 6454200.01 |
| lizards | Kentropyx_lagartija | Kentropyx_lagartija | 0 | 19.05 | 5.5 | 27136.29 |
| lizards | Kentropyx_paulensis | Kentropyx_paulensis | 0 | 13.18 | 3.85 | 1157731.09 |
| lizards | Kentropyx_pelviceps | Kentropyx_pelviceps | 0 | 67.61 | 4.53 | 2576988.52 |
| lizards | Kentropyx_striata | Kentropyx_striata | 0 | 63.1 | 4.06 | 2096781.34 |
| lizards | Kentropyx_vanzoi | Kentropyx_vanzoi | 0 | 17.78 | 2.31 | 1399763.63 |
| lizards | Kentropyx_viridistriga | Kentropyx_viridistriga | 0 | 36.31 | 5.27 | 1073309 |
| lizards | Kinyongia_adolfifriderici | Kinyongia_adolfifriderici | 0 | 7.59 | 3 | 124018.62 |
| lizards | Kinyongia_boehmei | Kinyongia_boehmei | 0 | 21.38 | 8 | 438.39 |
| lizards | Kinyongia_excubitor | Kinyongia_excubitor | 1 | 11.48 | 15.5 | 5340.72 |
| lizards | Kinyongia_fischeri | Kinyongia_fischeri | 0 | 147.91 | 21 | 256.66 |
| lizards | Kinyongia_matschiei | Kinyongia_matschiei | 1 | 77.62 | 20 | 717.96 |
| lizards | Kinyongia_multituberculata | Kinyongia_multituberculata | 1 | 53.7 | 15.5 | 1296.2 |
| lizards | Kinyongia_oxyrhina | Kinyongia_oxyrhina | 0 | 9.55 | 13 | 12394.66 |
| lizards | Kinyongia_tavetana | Kinyongia_tavetana | 0 | 21.38 | 11 | 10491.71 |
| lizards | Kinyongia_tenuis | Kinyongia_tenuis | 1 | 8.91 | 4 | 28982.47 |
| lizards | Kinyongia_uluguruensis | Kinyongia_uluguruensis | 0 | 19.05 | 9 | 878.74 |
| lizards | Kinyongia_uthmoelleri | Kinyongia_uthmoelleri | 0 | 19.05 | 9.5 | 23853.5 |
| lizards | Kinyongia_vosseleri | Kinyongia_vosseleri | 1 | 37.15 | 12 | 442.34 |
| lizards | Kinyongia_xenorhina | Kinyongia_xenorhina | 0 | 25.12 | 6 | 7845.26 |
| lizards | Lacerta_agilis | Lacerta_agilis | 0 | 33.88 | 8.4 | 11148919.36 |
| lizards | Lacerta_bilineata | Lacerta_bilineata | 0 | 50.12 | 11.05 | 856006.71 |
| lizards | Lacerta_media | Lacerta_media | 0 | 91.2 | 11.2 | 927506.6 |
| lizards | Lacerta_pamphylica | Lacerta_pamphylica | 0 | 38.9 | 12.5 | 9661.3 |
| lizards | Lacerta_schreiberi | Lacerta_schreiberi | 0 | 75.86 | 12.35 | 145400.97 |
| lizards | Lacerta_strigata | Lacerta_strigata | 0 | 91.2 | 10.17 | 415809.47 |
| lizards | Lacerta_trilineata | Lacerta_trilineata | 0 | 117.49 | 10.39 | 596692.93 |
| lizards | Lacerta_viridis | Lacerta_viridis | 0 | 91.2 | 9.01 | 1369557.62 |
| lizards | Lacertaspis_lepesmei | Lacertaspis_lepesmei | 1 | 3.72 | 2 | 146.5 |
| lizards | Lacertoides_pardalis | Lacertoides_pardalis | 1 | 54.95 | 4.5 | 375.07 |
| lizards | Laemanctus_longipes | Laemanctus_longipes | 0 | 104.71 | 4 | 177694.71 |
| lizards | Laemanctus_serratus | Laemanctus_serratus | 0 | 229.09 | 7 | 208774.87 |
| lizards | Lamprolepis_smaragdina | Lamprolepis_smaragdina | 0 | 40.74 | 2.05 | 892245.13 |

| | | | | | | |
|---|---|---|---|---|---|---|
| lizards | Lampropholis_adonis | Lampropholis_adonis | 0 | 2.75 | 3 | 43556.93 |
| lizards | Lampropholis_amicula | Lampropholis_amicula | 0 | 0.72 | 2 | 105352.54 |
| lizards | Lampropholis_caligula | Lampropholis_caligula | 0 | 2.95 | 2 | 3501.02 |
| lizards | Lampropholis_coggeri | Lampropholis_coggeri | 0 | 1.62 | 2.39 | 5206.97 |
| lizards | Lampropholis_delicata | Lampropholis_delicata | 0 | 3.09 | 3.35 | 636433.85 |
| lizards | Lampropholis_guichenoti | Lampropholis_guichenoti | 0 | 2.63 | 2.8 | 688423.8 |
| lizards | Lampropholis_mirabilis | Lampropholis_mirabilis | 0 | 2.63 | 3.03 | 9.99 |
| lizards | Lampropholis_robertsi | Lampropholis_robertsi | 0 | 3.09 | 2.37 | 5902.83 |
| lizards | Lankascincus_deignani | Lankascincus_deignani | 1 | 3.72 | 2 | 13434.62 |
| lizards | Lankascincus_fallax | Lankascincus_fallax | 0 | 4.17 | 1.9 | 65499.28 |
| lizards | Lankascincus_gansi | Lankascincus_gansi | 0 | 3.09 | 1 | 20573.04 |
| lizards | Lankascincus_sripadensis | Lankascincus_sripadensis | 1 | 3.8 | 2 | 9.99 |
| lizards | Lankascincus_taprobanensis | Lankascincus_taprobanensis | 1 | 3.72 | 2 | 13064.49 |
| lizards | Lankascincus_taylori | Lankascincus_taylori | 1 | 2.63 | 2 | 19983.83 |
| lizards | Lanthanotus_borneensis | Lanthanotus_borneensis | 1 | 204.17 | 5.3 | 45789.23 |
| lizards | Larutia_larutensis | Larutia_larutensis | 1 | 26.92 | 3 | 19.98 |
| lizards | Larutia_miodactyla | Larutia_miodactyla | 1 | 15.14 | 1 | 6793 |
| lizards | Larutia_trifasciata | Larutia_trifasciata | 0 | 52.48 | 2.5 | 3203.05 |
| lizards | Latastia_johnstonii | Latastia_johnstonii | 0 | 6.17 | 7 | 411744.38 |
| lizards | Latastia_longicaudata | Latastia_longicaudata | 0 | 30.2 | 6.8 | 6995048.78 |
| lizards | Laudakia_agrorensis | Laudakia_agrorensis | 0 | 48.98 | 7 | 377167.68 |
| lizards | Laudakia_melanura | Laudakia_melanura | 0 | 107.15 | 12.5 | 893887.52 |
| lizards | Laudakia_nupta | Laudakia_nupta | 0 | 181.97 | 10 | 1524079.16 |
| lizards | Laudakia_tuberculata | Laudakia_tuberculata | 0 | 117.49 | 9.4 | 459162.02 |
| lizards | Leiocephalus_barahonensis | Leiocephalus_barahonensis | 0 | 16.22 | 2.06 | 4140.88 |
| lizards | Leiocephalus_carinatus | Leiocephalus_carinatus | 0 | 72.44 | 4.05 | 124368.49 |
| lizards | Leiocephalus_cubensis | Leiocephalus_cubensis | 0 | 52.48 | 2.33 | 55970.77 |
| lizards | Leiocephalus_inaguae | Leiocephalus_inaguae | 0 | 19.95 | 4 | 1581.96 |
| lizards | Leiocephalus_lunatus | Leiocephalus_lunatus | 0 | 7.76 | 1.6 | 943.97 |
| lizards | Leiocephalus_macropus | Leiocephalus_macropus | 0 | 23.99 | 2.38 | 69312.2 |
| lizards | Leiocephalus_melanochlorus | Leiocephalus_melanochlorus | 0 | 66.07 | 4 | 6510.15 |
| lizards | Leiocephalus_onaneyi | Leiocephalus_onaneyi | 1 | 10.23 | 2 | 14.7 |
| lizards | Leiocephalus_personatus | Leiocephalus_personatus | 0 | 36.31 | 2 | 50792.4 |
| lizards | Leiocephalus_psammodromus | Leiocephalus_psammodromus | 1 | 38.9 | 1.72 | 305.16 |

| | | | | | | |
|---|---|---|---|---|---|---|
| lizards | Leiocephalus_punctatus | Leiocephalus_punctatus | 0 | 13.8 | 8 | 615.09 |
| lizards | Leiocephalus_raviceps | Leiocephalus_raviceps | 0 | 10.72 | 1.67 | 12074.38 |
| lizards | Leiocephalus_schreibersii | Leiocephalus_schreibersii | 0 | 35.48 | 2.47 | 29792.5 |
| lizards | Leiocephalus_semilineatus | Leiocephalus_semilineatus | 0 | 3.63 | 1.78 | 6069.81 |
| lizards | Leiocephalus_stictigaster | Leiocephalus_stictigaster | 0 | 28.18 | 1.33 | 44480.29 |
| lizards | Leiolepis_belliana | Leiolepis_belliana | 0 | 199.53 | 4.65 | 819785.38 |
| lizards | Leiolepis_guentherpetersi | Leiolepis_guentherpetersi | 1 | 131.83 | 2 | 4446.03 |
| lizards | Leiolepis_reevesii | Leiolepis_reevesii | 0 | 245.47 | 3.58 | 446691.99 |
| lizards | Leiosaurus_bellii | Leiosaurus_bellii | 0 | 41.69 | 15 | 628194.61 |
| lizards | Leiosaurus_catamarcensis | Leiosaurus_catamarcensis | 0 | 56.23 | 8.5 | 179019.49 |
| lizards | Lepidoblepharis_heyerorum | Lepidoblepharis_heyerorum | 0 | 1.07 | 1 | 2547613.74 |
| lizards | Lepidoblepharis_montecanoensis | Lepidoblepharis_montecanoensis | 0 | 0.23 | 1 | 2499.61 |
| lizards | Lepidoblepharis_sanctaemartae | Lepidoblepharis_sanctaemartae | 0 | 1.07 | 1 | 103968.01 |
| lizards | Lepidoblepharis_xanthostigma | Lepidoblepharis_xanthostigma | 0 | 2.24 | 1 | 192457.93 |
| lizards | Lepidodactylus_euaensis | Lepidodactylus_euaensis | 1 | 2.82 | 2 | 88.55 |
| lizards | Lepidodactylus_gardineri | Lepidodactylus_gardineri | 1 | 3.39 | 2 | 42.43 |
| lizards | Lepidodactylus_guppyi | Lepidodactylus_guppyi | 0 | 3.55 | 2 | 36047.37 |
| lizards | Lepidodactylus_lugubris | Lepidodactylus_lugubris | 0 | 2.95 | 1.5 | 7666582.84 |
| lizards | Lepidodactylus_magnus | Lepidodactylus_magnus | 0 | 7.41 | 2 | 30113.2 |
| lizards | Lepidodactylus_manni | Lepidodactylus_manni | 1 | 2.51 | 2 | 16606.33 |
| lizards | Lepidodactylus_moestus | Lepidodactylus_moestus | 0 | 2.09 | 1.84 | 893.46 |
| lizards | Lepidodactylus_orientalis | Lepidodactylus_orientalis | 0 | 1.86 | 1.8 | 5443.47 |
| lizards | Lepidodactylus_pumilus | Lepidodactylus_pumilus | 0 | 2.51 | 2 | 17402.98 |
| lizards | Lepidodactylus_vanuatuensis | Lepidodactylus_vanuatuensis | 0 | 2.4 | 2 | 4769.35 |
| lizards | Lepidodactylus_yami | Lepidodactylus_yami | 0 | 1.74 | 2 | 49.72 |
| lizards | Lepidophyma_flavimaculatum | Lepidophyma_flavimaculatum | 0 | 72.44 | 5 | 311582.87 |
| lizards | Lepidophyma_gaigeae | Lepidophyma_gaigeae | 1 | 5.75 | 2.6 | 6382.13 |
| lizards | Lepidophyma_micropholis | Lepidophyma_micropholis | 1 | 27.54 | 5.3 | 4050.18 |
| lizards | Lepidophyma_pajapanensis | Lepidophyma_pajapanensis | 0 | 16.98 | 5.5 | 1855.66 |
| lizards | Lepidophyma_smithii | Lepidophyma_smithii | 0 | 30.9 | 8 | 52342.6 |
| lizards | Lepidophyma_sylvaticum | Lepidophyma_sylvaticum | 0 | 28.84 | 5.6 | 58168.45 |
| lizards | Leposoma_baturitensis | Leposoma_baturitensis | 1 | 0.91 | 2 | 5620.96 |
| lizards | Leposoma_scincoides | Leposoma_scincoides | 0 | 2.04 | 1.5 | 110657.69 |
| lizards | Leposternon_infraorbitale | Amphisbaena_infraorbitale | 0 | 380.19 | 4 | 557716.69 |

| | | | | | | |
|---|---|---|---|---|---|---|
| lizards | Leposternon_microcephalum | Amphisbaena_microcephalum | 0 | 213.8 | 5 | 2205828.58 |
| lizards | Leposternon_polystegum | Amphisbaena_polystegum | 0 | 114.82 | 6 | 1986716.11 |
| lizards | Leptosiaphos_graueri | Leptosiaphos_graueri | 1 | 2.69 | 2 | 80181.43 |
| lizards | Leptosiaphos_ianthinoxantha | Leptosiaphos_ianthinoxantha | 1 | 4.9 | 4 | 2183.79 |
| lizards | Leptosiaphos_kilimensis | Leptosiaphos_kilimensis | 0 | 7.76 | 3 | 225182.31 |
| lizards | Leptosiaphos_meleagris | Leptosiaphos_meleagris | 1 | 2.69 | 2 | 27533.29 |
| lizards | Lerista_allanae | Lerista_allanae | 1 | 4.47 | 2.3 | 597.07 |
| lizards | Lerista_arenicola | Lerista_arenicola | 0 | 5.62 | 3 | 27969.54 |
| lizards | Lerista_baynesi | Lerista_baynesi | 0 | 4.27 | 3 | 230.76 |
| lizards | Lerista_bipes | Lerista_bipes | 0 | 2.04 | 2.01 | 1374937.74 |
| lizards | Lerista_borealis | Lerista_borealis | 0 | 1.74 | 3 | 34070.92 |
| lizards | Lerista_bougainvillii | Lerista_bougainvillii | 0 | 8.13 | 2.62 | 694518.31 |
| lizards | Lerista_christinae | Lerista_christinae | 1 | 0.56 | 2.15 | 14329.85 |
| lizards | Lerista_cinerea | Lerista_cinerea | 0 | 2.4 | 2.4 | 9926.02 |
| lizards | Lerista_colliveri | Lerista_colliveri | 0 | 4.17 | 2 | 25986.39 |
| lizards | Lerista_connivens | Lerista_connivens | 0 | 3.72 | 6 | 54227.56 |
| lizards | Lerista_desertorum | Lerista_desertorum | 0 | 4.68 | 3 | 1491472.97 |
| lizards | Lerista_distinguenda | Lerista_distinguenda | 0 | 1.02 | 1.72 | 405698.68 |
| lizards | Lerista_edwardsae | Lerista_edwardsae | 0 | 4.79 | 3 | 174485.98 |
| lizards | Lerista_elegans | Lerista_elegans | 0 | 0.76 | 2.5 | 199087.29 |
| lizards | Lerista_emmotti | Lerista_emmotti | 0 | 5.89 | 3 | 259402.33 |
| lizards | Lerista_fragilis | Lerista_fragilis | 0 | 1.55 | 2 | 520820.91 |
| lizards | Lerista_frosti | Lerista_frosti | 0 | 2.09 | 2.3 | 65790.15 |
| lizards | Lerista_gerrardii | Lerista_gerrardii | 0 | 4.27 | 2 | 174869.24 |
| lizards | Lerista_griffini | Lerista_griffini | 0 | 2.04 | 2.3 | 175113.68 |
| lizards | Lerista_humphriesi | Lerista_humphriesi | 0 | 0.91 | 2 | 3354.45 |
| lizards | Lerista_ips | Lerista_ips | 0 | 2.4 | 2 | 434663.31 |
| lizards | Lerista_karlschmidti | Lerista_karlschmidti | 0 | 2.34 | 1.6 | 125442.09 |
| lizards | Lerista_labialis | Lerista_labialis | 0 | 1.55 | 2 | 2260861.2 |
| lizards | Lerista_lineata | Lerista_lineata | 1 | 1.66 | 2.72 | 25103.44 |
| lizards | Lerista_lineopunctulata | Lerista_lineopunctulata | 0 | 7.41 | 3.9 | 75800.4 |
| lizards | Lerista_microtis | Lerista_microtis | 0 | 4.37 | 2.3 | 102837.82 |
| lizards | Lerista_muelleri | Lerista_muelleri | 0 | 0.98 | 1.9 | 2180748.39 |
| lizards | Lerista_petersoni | Lerista_petersoni | 0 | 2.34 | 6 | 53512.1 |

| | | | | | | |
|---|---|---|---|---|---|---|
| lizards | Lerista_picturata | Lerista_picturata | 0 | 4.47 | 3 | 172756.65 |
| lizards | Lerista_planiventralis | Lerista_planiventralis | 0 | 2.4 | 3 | 111772.36 |
| lizards | Lerista_praepedita | Lerista_praepedita | 0 | 0.95 | 2.3 | 106472.83 |
| lizards | Lerista_punctatovittata | Lerista_punctatovittata | 0 | 6.17 | 2.15 | 1426418.07 |
| lizards | Lerista_storri | Lerista_storri | 0 | 2.51 | 2 | 14805.46 |
| lizards | Lerista_stylis | Lerista_stylis | 0 | 1.32 | 2.2 | 11149.96 |
| lizards | Lerista_uniduo | Lerista_uniduo | 0 | 1.58 | 2.3 | 137380.85 |
| lizards | Lerista_vittata | Lerista_vittata | 1 | 2.75 | 2.55 | 518.34 |
| lizards | Lerista_walkeri | Lerista_walkeri | 0 | 2.19 | 2 | 23437.03 |
| lizards | Lerista_wilkinsi | Lerista_wilkinsi | 0 | 2.69 | 1.95 | 8726.06 |
| lizards | Lerista_xanthura | Lerista_xanthura | 0 | 1.41 | 1.35 | 491754.57 |
| lizards | Lialis_burtonis | Lialis_burtonis | 0 | 22.91 | 1.85 | 7207219.23 |
| lizards | Lialis_jicari | Lialis_jicari | 0 | 23.99 | 2 | 186471.1 |
| lizards | Liburnascincus_mundivensis | Liburnascincus_mundivensis | 0 | 4.17 | 2 | 195037.92 |
| lizards | Liburnascincus_scirtetis | Liburnascincus_scirtetis | 0 | 6.76 | 2 | 26.04 |
| lizards | Liolaemus_abdalai | Liolaemus_abdalai | 0 | 3.89 | 1.85 | 5916.85 |
| lizards | Liolaemus_alticolor | Liolaemus_alticolor | 0 | 5.25 | 3.05 | 45889.03 |
| lizards | Liolaemus_andinus | Liolaemus_andinus | 0 | 24.55 | 5.8 | 63469.14 |
| lizards | Liolaemus_anomalus | Liolaemus_anomalus | 0 | 28.18 | 7.5 | 44677.07 |
| lizards | Liolaemus_arambarensis | Liolaemus_arambarensis | 1 | 6.76 | 2 | 1239.66 |
| lizards | Liolaemus_atacamensis | Liolaemus_atacamensis | 0 | 14.45 | 5 | 119428.95 |
| lizards | Liolaemus_audituvelatus | Liolaemus_audituvelatus | 1 | 6.92 | 2.5 | 39887.98 |
| lizards | Liolaemus_baguali | Liolaemus_baguali | 0 | 33.88 | 3.5 | 6012.63 |
| lizards | Liolaemus_bellii | Liolaemus_bellii | 0 | 26.92 | 4.22 | 10201.5 |
| lizards | Liolaemus_bibronii | Liolaemus_bibronii | 0 | 10.96 | 2.75 | 605195.2 |
| lizards | Liolaemus_bitaeniatus | Liolaemus_bitaeniatus | 0 | 7.41 | 5.91 | 47641.75 |
| lizards | Liolaemus_boulengeri | Liolaemus_boulengeri | 0 | 15.14 | 4.08 | 557576.76 |
| lizards | Liolaemus_cazianiae | Liolaemus_cazianiae | 0 | 14.13 | 3 | 4758.89 |
| lizards | Liolaemus_chacoensis | Liolaemus_chacoensis | 0 | 5.13 | 6.07 | 484670.63 |
| lizards | Liolaemus_chaltin | Liolaemus_chaltin | 0 | 6.31 | 4.5 | 11870.57 |
| lizards | Liolaemus_chavin | Liolaemus_chavin | 0 | 9.33 | 4 | 5142.54 |
| lizards | Liolaemus_chiliensis | Liolaemus_chiliensis | 0 | 38.9 | 14.4 | 215500.98 |
| lizards | Liolaemus_chillanensis | Liolaemus_chillanensis | 0 | 22.91 | 3 | 11442.45 |
| lizards | Liolaemus_coeruleus | Liolaemus_coeruleus | 0 | 10.96 | 4.5 | 12446.76 |

| | | | | | | |
|---|---|---|---|---|---|---|
| lizards | Liolaemus_constanzae | Liolaemus_constanzae | 0 | 14.45 | 3.64 | 33922.41 |
| lizards | Liolaemus_crepuscularis | Liolaemus_crepuscularis | 0 | 8.32 | 4.5 | 1298.03 |
| lizards | Liolaemus_curis | Liolaemus_curis | 1 | 36.31 | 5.5 | 105.6 |
| lizards | Liolaemus_cyanogaster | Liolaemus_cyanogaster | 0 | 11.48 | 7.5 | 140369.79 |
| lizards | Liolaemus_darwinii | Liolaemus_darwinii | 0 | 10.47 | 5.05 | 779703.75 |
| lizards | Liolaemus_dorbignyi | Liolaemus_dorbignyi | 0 | 34.67 | 4.5 | 64763.06 |
| lizards | Liolaemus_elongatus | Liolaemus_elongatus | 0 | 27.54 | 3.9 | 270073.05 |
| lizards | Liolaemus_espinozai | Liolaemus_espinozai | 0 | 7.76 | 2.5 | 1216.31 |
| lizards | Liolaemus_fabiani | Liolaemus_fabiani | 1 | 23.99 | 2.5 | 1918.35 |
| lizards | Liolaemus_fitzgeraldi | Liolaemus_fitzgeraldi | 0 | 8.71 | 2.25 | 54255.72 |
| lizards | Liolaemus_foxi | Liolaemus_foxi | 0 | 18.2 | 4 | 771.09 |
| lizards | Liolaemus_fuscus | Liolaemus_fuscus | 0 | 4.68 | 2 | 2464.97 |
| lizards | Liolaemus_gallardoi | Liolaemus_gallardoi | 0 | 25.12 | 3 | 34575.94 |
| lizards | Liolaemus_goetschi | Liolaemus_goetschi | 0 | 13.18 | 5 | 122943.8 |
| lizards | Liolaemus_gracilis | Liolaemus_gracilis | 0 | 6.17 | 4.23 | 577965.52 |
| lizards | Liolaemus_gravenhorstii | Liolaemus_gravenhorstii | 1 | 10.47 | 6 | 6065.4 |
| lizards | Liolaemus_halonastes | Liolaemus_halonastes | 1 | 11.75 | 3.5 | 69.14 |
| lizards | Liolaemus_hatcheri | Liolaemus_hatcheri | 0 | 12.3 | 3 | 16719.84 |
| lizards | Liolaemus_huacahuasicus | Liolaemus_huacahuasicus | 0 | 14.13 | 4.77 | 841.88 |
| lizards | Liolaemus_incaicus | Liolaemus_incaicus | 0 | 6.03 | 5 | 1122.17 |
| lizards | Liolaemus_irregularis | Liolaemus_irregularis | 0 | 23.44 | 5.38 | 7925.07 |
| lizards | Liolaemus_jamesi | Liolaemus_jamesi | 0 | 32.36 | 5.6 | 24376.31 |
| lizards | Liolaemus_kingii | Liolaemus_kingii | 0 | 33.11 | 3.5 | 273140.79 |
| lizards | Liolaemus_kolengh | Liolaemus_kolengh | 0 | 7.59 | 3.2 | 1023.72 |
| lizards | Liolaemus_koslowskyi | Liolaemus_koslowskyi | 0 | 17.78 | 4.89 | 33902.59 |
| lizards | Liolaemus_kriegi | Liolaemus_kriegi | 0 | 50.12 | 4 | 112291.95 |
| lizards | Liolaemus_lemniscatus | Liolaemus_lemniscatus | 0 | 5.5 | 4.5 | 176579.8 |
| lizards | Liolaemus_leopardinus | Liolaemus_leopardinus | 1 | 34.67 | 3 | 644.2 |
| lizards | Liolaemus_lineomaculatus | Liolaemus_lineomaculatus | 0 | 10 | 3.56 | 435633.65 |
| lizards | Liolaemus_lutzae | Liolaemus_lutzae | 1 | 22.91 | 2.64 | 1707.58 |
| lizards | Liolaemus_magellanicus | Liolaemus_magellanicus | 0 | 11.48 | 4 | 94721.8 |
| lizards | Liolaemus_melanops | Liolaemus_melanops | 0 | 31.62 | 8.5 | 127566.14 |
| lizards | Liolaemus_millcayac | Liolaemus_millcayac | 0 | 15.49 | 7.5 | 21008.11 |
| lizards | Liolaemus_monticola | Liolaemus_monticola | 0 | 22.91 | 3.75 | 31123.62 |

| | | | | | | |
|---|---|---|---|---|---|---|
| lizards | Liolaemus_morenoi | Liolaemus_morenoi | 1 | 22.39 | 5 | 22101.86 |
| lizards | Liolaemus_multicolor | Liolaemus_multicolor | 0 | 19.05 | 7 | 38154.88 |
| lizards | Liolaemus_multimaculatus | Liolaemus_multimaculatus | 1 | 12.02 | 4.2 | 6252.34 |
| lizards | Liolaemus_nigromaculatus | Liolaemus_nigromaculatus | 0 | 26.3 | 3.09 | 80035.07 |
| lizards | Liolaemus_nigroviridis | Liolaemus_nigroviridis | 0 | 22.39 | 4.66 | 17486.22 |
| lizards | Liolaemus_nitidus | Liolaemus_nitidus | 0 | 38.9 | 8 | 100923.32 |
| lizards | Liolaemus_olongasta | Liolaemus_olongasta | 0 | 9.55 | 3.5 | 52531.16 |
| lizards | Liolaemus_ornatus | Liolaemus_ornatus | 0 | 11.48 | 7 | 130150.23 |
| lizards | Liolaemus_pachacutec | Liolaemus_pachacutec | 0 | 4.37 | 3 | 9627.03 |
| lizards | Liolaemus_pagaburoi | Liolaemus_pagaburoi | 0 | 5.75 | 5 | 4627.17 |
| lizards | Liolaemus_parvus | Liolaemus_parvus | 0 | 11.48 | 2.44 | 19702.16 |
| lizards | Liolaemus_patriciaiturrae | Liolaemus_patriciaiturrae | 0 | 29.51 | 3.5 | 4065.76 |
| lizards | Liolaemus_paulinae | Liolaemus_paulinae | 1 | 7.08 | 2 | 215.26 |
| lizards | Liolaemus_petrophilus | Liolaemus_petrophilus | 0 | 33.11 | 3.6 | 187888.49 |
| lizards | Liolaemus_pictus | Liolaemus_pictus | 0 | 37.15 | 3.95 | 172561.91 |
| lizards | Liolaemus_platei | Liolaemus_platei | 0 | 11.48 | 3 | 99102.12 |
| lizards | Liolaemus_pleopholis | Liolaemus_pleopholis | 0 | 16.98 | 4.4 | 4961.1 |
| lizards | Liolaemus_poecilochromus | Liolaemus_poecilochromus | 0 | 14.13 | 5.8 | 5440.98 |
| lizards | Liolaemus_pseudoanomalus | Liolaemus_pseudoanomalus | 0 | 10 | 6.8 | 59606.59 |
| lizards | Liolaemus_puna | Liolaemus_puna | 0 | 5.5 | 3 | 278719.85 |
| lizards | Liolaemus_puritamensis | Liolaemus_puritamensis | 0 | 33.88 | 4 | 24529.92 |
| lizards | Liolaemus_ramirezae | Liolaemus_ramirezae | 0 | 5.89 | 5 | 19191.9 |
| lizards | Liolaemus_riojanus | Liolaemus_riojanus | 0 | 7.76 | 4.2 | 20256.94 |
| lizards | Liolaemus_robertoi | Liolaemus_robertoi | 1 | 11.75 | 4.5 | 778.31 |
| lizards | Liolaemus_rosenmanni | Liolaemus_rosenmanni | 0 | 15.14 | 3 | 10786.24 |
| lizards | Liolaemus_rothi | Liolaemus_rothi | 0 | 37.15 | 6.5 | 138928.6 |
| lizards | Liolaemus_ruibali | Liolaemus_ruibali | 0 | 9.33 | 2.97 | 19123.26 |
| lizards | Liolaemus_sagei | Liolaemus_sagei | 0 | 33.11 | 4.5 | 15129.84 |
| lizards | Liolaemus_sanjuanensis | Liolaemus_sanjuanensis | 0 | 8.71 | 4.5 | 4084.61 |
| lizards | Liolaemus_sarmientoi | Liolaemus_sarmientoi | 0 | 30.9 | 3.81 | 82944.7 |
| lizards | Liolaemus_saxatilis | Liolaemus_saxatilis | 0 | 6.76 | 5.6 | 17738.86 |
| lizards | Liolaemus_scapularis | Liolaemus_scapularis | 1 | 14.45 | 4.33 | 13758.41 |
| lizards | Liolaemus_schroederi | Liolaemus_schroederi | 0 | 9.77 | 8 | 77024.67 |
| lizards | Liolaemus_septentrionalis | Liolaemus_septentrionalis | 0 | 6.61 | 4.2 | 3081.93 |

| | | | | | | |
|---|---|---|---|---|---|---|
| lizards | Liolaemus_signifer | Liolaemus_signifer | 0 | 41.69 | 6.1 | 373214.76 |
| lizards | Liolaemus_silvanae | Liolaemus_silvanae | 0 | 15.14 | 3 | 3113.52 |
| lizards | Liolaemus_stolzmanni | Liolaemus_stolzmanni | 1 | 7.24 | 2 | 1268.65 |
| lizards | Liolaemus_tandiliensis | Liolaemus_tandiliensis | 1 | 6.17 | 3.19 | 11868.18 |
| lizards | Liolaemus_tari | Liolaemus_tari | 0 | 34.67 | 3.5 | 4127.89 |
| lizards | Liolaemus_tenuis | Liolaemus_tenuis | 0 | 8.51 | 5.05 | 226075.72 |
| lizards | Liolaemus_torresi | Liolaemus_torresi | 1 | 8.71 | 2 | 3344.32 |
| lizards | Liolaemus_tristis | Liolaemus_tristis | 0 | 20.42 | 3.5 | 4459.62 |
| lizards | Liolaemus_uspallatensis | Liolaemus_uspallatensis | 0 | 8.71 | 3 | 18567.89 |
| lizards | Liolaemus_valdesianus | Liolaemus_valdesianus | 0 | 28.18 | 2.5 | 481.53 |
| lizards | Liolaemus_vallecurensis | Liolaemus_vallecurensis | 0 | 14.79 | 4 | 71303.66 |
| lizards | Liolaemus_velosoi | Liolaemus_velosoi | 0 | 6.46 | 1 | 153.13 |
| lizards | Liolaemus_wari | Liolaemus_wari | 0 | 7.24 | 3 | 9364.51 |
| lizards | Liolaemus_wiegmannii | Liolaemus_wiegmannii | 0 | 9.12 | 4.45 | 1481143.35 |
| lizards | Liolaemus_xanthoviridis | Liolaemus_xanthoviridis | 0 | 33.88 | 7 | 15719.97 |
| lizards | Liolaemus_yanalcu | Liolaemus_yanalcu | 0 | 7.24 | 3.5 | 1339.48 |
| lizards | Liolaemus_zapallarensis | Liolaemus_zapallarensis | 0 | 43.65 | 6.91 | 1996.25 |
| lizards | Liolaemus_zullyae | Liolaemus_zullyae | 0 | 22.91 | 2.5 | 2053.64 |
| lizards | Liopholis_guthega | Liopholis_guthega | 1 | 32.36 | 2.28 | 1108.35 |
| lizards | Liopholis_inornata | Liopholis_inornata | 0 | 12.88 | 2.5 | 2541556.98 |
| lizards | Liopholis_kintorei | Liopholis_kintorei | 1 | 223.87 | 6 | 1336754.79 |
| lizards | Liopholis_modesta | Liopholis_modesta | 0 | 30.9 | 2.7 | 299350.97 |
| lizards | Liopholis_montana | Liopholis_montana | 0 | 30.2 | 4 | 40855.75 |
| lizards | Liopholis_multiscutata | Liopholis_multiscutata | 0 | 19.05 | 2 | 568847.56 |
| lizards | Liopholis_pulchra | Liopholis_pulchra | 0 | 29.51 | 3 | 82244.66 |
| lizards | Liopholis_slateri | Liopholis_slateri | 1 | 19.5 | 3.3 | 73334.73 |
| lizards | Liopholis_striata | Liopholis_striata | 0 | 35.48 | 2.8 | 2261385.72 |
| lizards | Liopholis_whitii | Liopholis_whitii | 0 | 31.62 | 2.9 | 831367.91 |
| lizards | Lioscincus_steindachneri | Lioscincus_steindachneri | 1 | 31.62 | 6 | 1594.23 |
| lizards | Lipinia_auriculata | Lipinia_auriculata | 0 | 2.45 | 1.5 | 40497 |
| lizards | Lipinia_inexpectata | Lipinia_inexpectata | 0 | 1.17 | 2 | 62144.22 |
| lizards | Lipinia_leptosoma | Lipinia_leptosoma | 0 | 1.51 | 1 | 455.64 |
| lizards | Lipinia_longiceps | Lipinia_longiceps | 0 | 1.41 | 2 | 23559.67 |
| lizards | Lipinia_pulchella | Lipinia_pulchella | 0 | 2.29 | 1.8 | 186999.27 |

| | | | | | | |
|---|---|---|---|---|---|---|
| lizards | Lipinia_quadrivittata | Lipinia_quadrivittata | 0 | 1.74 | 2 | 38346.88 |
| lizards | Lipinia_relicta | Lipinia_relicta | 0 | 3.31 | 2 | 138.18 |
| lizards | Lipinia_rouxi | Lipinia_rouxi | 0 | 1.2 | 1 | 6878.3 |
| lizards | Lipinia_septentrionalis | Lipinia_septentrionalis | 0 | 1.41 | 2 | 68919.02 |
| lizards | Lipinia_vittigera | Lipinia_vittigera | 0 | 1.62 | 2.25 | 457593.08 |
| lizards | Lissolepis_coventryi | Lissolepis_coventryi | 1 | 79.43 | 2.75 | 109117.35 |
| lizards | Lobulia_brongersmai | Lobulia_brongersmai | 0 | 3.09 | 3 | 3306.87 |
| lizards | Lobulia_elegans | Lobulia_elegans | 0 | 5.62 | 3 | 92798.49 |
| lizards | Lophognathus_gilberti | Lophognathus_gilberti | 0 | 85.11 | 5.4 | 258890.61 |
| lizards | Lophosaurus_boydii | Hypsilurus_boydii | 0 | 190.55 | 3 | 28110.48 |
| lizards | Lophosaurus_dilophus | Hypsilurus_dilophus | 0 | 389.05 | 2 | 772102.21 |
| lizards | Lophosaurus_spinipes | Hypsilurus_spinipes | 0 | 87.1 | 5 | 78462.36 |
| lizards | Loveridgea_ionidesii | Loveridgea_ionidesii | 0 | 2.69 | 1.5 | 119919.94 |
| lizards | Loxopholis_guianense | Leposoma_guianense | 0 | 1.2 | 2 | 1480769.96 |
| lizards | Loxopholis_hexalepis | Leposoma_hexalepis | 0 | 1.41 | 2 | 67952.09 |
| lizards | Loxopholis_osvaldoi | Leposoma_osvaldoi | 0 | 1 | 1.75 | 1602331.61 |
| lizards | Loxopholis_parietalis | Leposoma_parietale | 0 | 1.91 | 2 | 632209.62 |
| lizards | Loxopholis_percarinatum | Leposoma_percarinatum | 0 | 1 | 1.89 | 5730663.91 |
| lizards | Loxopholis_rugiceps | Leposoma_rugiceps | 0 | 1.78 | 2.09 | 465129.47 |
| lizards | Loxopholis_snethlageae | Leposoma_snethlageae | 0 | 0.69 | 2 | 675574.21 |
| lizards | Loxopholis_southi | Leposoma_southi | 0 | 3.72 | 2 | 303496.73 |
| lizards | Lubuya_ivensii | Trachylepis_ivensii | 0 | 75.86 | 4.67 | 479094.03 |
| lizards | Lucasium_alboguttatum | Lucasium_alboguttatum | 0 | 3.63 | 2 | 80258.93 |
| lizards | Lucasium_bungabinna | Lucasium_bungabinna | 0 | 3.89 | 2 | 515832.85 |
| lizards | Lucasium_byrnei | Lucasium_byrnei | 0 | 3.55 | 2 | 873971.17 |
| lizards | Lucasium_damaeum | Lucasium_damaeum | 0 | 4.27 | 2 | 2068398.97 |
| lizards | Lucasium_maini | Lucasium_maini | 0 | 3.09 | 2 | 350641.05 |
| lizards | Lucasium_occultum | Lucasium_occultum | 1 | 1.35 | 2 | 2381.58 |
| lizards | Lucasium_squarrosum | Lucasium_squarrosum | 0 | 3.63 | 2 | 505496.95 |
| lizards | Lucasium_steindachneri | Lucasium_steindachneri | 0 | 4.07 | 2 | 1570837.4 |
| lizards | Lucasium_wombeyi | Lucasium_wombeyi | 0 | 3.09 | 2 | 114715.14 |
| lizards | Lygisaurus_foliorum | Lygisaurus_foliorum | 0 | 1.51 | 1.95 | 842999.21 |
| lizards | Lygisaurus_laevis | Lygisaurus_laevis | 0 | 0.87 | 2 | 11367.9 |
| lizards | Lygisaurus_macfarlani | Lygisaurus_macfarlani | 0 | 1.07 | 2 | 94494.78 |

| | | | | | | |
|---|---|---|---|---|---|---|
| lizards | Lygisaurus_rococo | Lygisaurus_rococo | 0 | 1.62 | 2 | 950.59 |
| lizards | Lygisaurus_zuma | Lygisaurus_zuma | 0 | 0.71 | 2 | 14787.68 |
| lizards | Lygodactylus_angolensis | Lygodactylus_angolensis | 0 | 1.12 | 2 | 1958377.78 |
| lizards | Lygodactylus_angularis | Lygodactylus_angularis | 0 | 2.24 | 2 | 441775.81 |
| lizards | Lygodactylus_bernardi | Lygodactylus_bernardi | 0 | 1.51 | 2 | 8187.9 |
| lizards | Lygodactylus_bivittis | Lygodactylus_bivittis | 1 | 1.15 | 2 | 20306.38 |
| lizards | Lygodactylus_bradfieldi | Lygodactylus_bradfieldi | 0 | 0.69 | 2 | 1028863.94 |
| lizards | Lygodactylus_capensis | Lygodactylus_capensis | 0 | 1.86 | 1.97 | 4100565.08 |
| lizards | Lygodactylus_chobiensis | Lygodactylus_chobiensis | 0 | 2 | 2 | 591173.94 |
| lizards | Lygodactylus_conraui | Lygodactylus_conraui | 0 | 1.51 | 2 | 560180.62 |
| lizards | Lygodactylus_depressus | Lygodactylus_depressus | 0 | 1.32 | 2 | 645324.89 |
| lizards | Lygodactylus_gravis | Lygodactylus_gravis | 1 | 1.74 | 2 | 12320.68 |
| lizards | Lygodactylus_grotei | Lygodactylus_grotei | 0 | 0.83 | 2 | 379608.54 |
| lizards | Lygodactylus_gutturalis | Lygodactylus_gutturalis | 0 | 2.09 | 2 | 6305237.66 |
| lizards | Lygodactylus_insularis | Lygodactylus_insularis | 1 | 0.69 | 2 | 3.41 |
| lizards | Lygodactylus_keniensis | Lygodactylus_keniensis | 0 | 1.74 | 2 | 262812.18 |
| lizards | Lygodactylus_kimhowelli | Lygodactylus_kimhowelli | 0 | 1.1 | 2 | 8703.89 |
| lizards | Lygodactylus_klugei | Lygodactylus_klugei | 0 | 0.98 | 1.91 | 247442.92 |
| lizards | Lygodactylus_lawrencei | Lygodactylus_lawrencei | 0 | 0.98 | 2 | 102734.88 |
| lizards | Lygodactylus_manni | Lygodactylus_manni | 0 | 1.74 | 2 | 46936.31 |
| lizards | Lygodactylus_methueni | Lygodactylus_methueni | 1 | 1.74 | 2 | 19999.02 |
| lizards | Lygodactylus_mirabilis | Lygodactylus_mirabilis | 1 | 0.65 | 1 | 23.15 |
| lizards | Lygodactylus_mombasicus | Lygodactylus_mombasicus | 0 | 1.41 | 2 | 7241.41 |
| lizards | Lygodactylus_nigropunctatus | Lygodactylus_nigropunctatus | 0 | 1.32 | 2 | 57542.95 |
| lizards | Lygodactylus_ocellatus | Lygodactylus_ocellatus | 0 | 1.32 | 2 | 56517.61 |
| lizards | Lygodactylus_picturatus | Lygodactylus_picturatus | 0 | 1.86 | 2 | 1218978.75 |
| lizards | Lygodactylus_pictus | Lygodactylus_pictus | 0 | 1.62 | 2 | 29084.42 |
| lizards | Lygodactylus_somalicus | Lygodactylus_somalicus | 0 | 0.87 | 2 | 153499.59 |
| lizards | Lygodactylus_thomensis | Lygodactylus_thomensis | 0 | 1.2 | 2 | 108.19 |
| lizards | Lygodactylus_tuberosus | Lygodactylus_tuberosus | 0 | 1.35 | 1.5 | 50959.9 |
| lizards | Lygodactylus_verticillatus | Lygodactylus_verticillatus | 0 | 1.05 | 1.5 | 66079.37 |
| lizards | Lygodactylus_williamsi | Lygodactylus_williamsi | 1 | 0.98 | 2 | 50.94 |
| lizards | Lygosoma_angeli | Lygosoma_angeli | 0 | 79.43 | 3.5 | 3681.57 |
| lizards | Lygosoma_corpulentum | Lygosoma_corpulentum | 0 | 186.21 | 4.5 | 75138.06 |

| | | | | | | |
|---|---|---|---|---|---|---|
| lizards | Lygosoma_koratense | Lygosoma_koratense | 0 | 38.9 | 3 | 2802.46 |
| lizards | Lygosoma_quadrupes | Lygosoma_quadrupes | 0 | 4.9 | 2.5 | 110.91 |
| lizards | Lyriocephalus_scutatus | Lyriocephalus_scutatus | 1 | 223.87 | 3.67 | 13983.56 |
| lizards | Mabuya_dominicana | Mabuya_dominicana | 0 | 22.39 | 3.15 | 624.11 |
| lizards | Mabuya_guadeloupae | Mabuya_guadeloupae | 1 | 25.7 | 5 | 9.99 |
| lizards | Mabuya_mabouya | Mabuya_mabouya | 1 | 22.39 | 2 | 1142.26 |
| lizards | Mabuya_montserratae | Mabuya_montserratae | 1 | 19.95 | 5 | 9.99 |
| lizards | Macropholidus_annectens | Macropholidus_annectens | 1 | 4.9 | 2 | 3607.91 |
| lizards | Macropholidus_ruthveni | Macropholidus_ruthveni | 0 | 2.04 | 1.83 | 24877 |
| lizards | Madascincus_melanopleura | Madascincus_melanopleura | 0 | 3.47 | 4.5 | 141035.31 |
| lizards | Madascincus_mouroundavae | Madascincus_mouroundavae | 0 | 7.41 | 2.5 | 69093.27 |
| lizards | Madascincus_nanus | Madascincus_nanus | 1 | 0.63 | 1 | 15683.42 |
| lizards | Madascincus_stumpffi | Madascincus_stumpffi | 0 | 33.11 | 4 | 15269.16 |
| lizards | Malayodracon_robinsonii | Gonocephalus_robinsonii | 0 | 138.04 | 7 | 6669.15 |
| lizards | Manciola_guaporicola | Manciola_guaporicola | 0 | 19.95 | 4.25 | 1493212.38 |
| lizards | Mantheyus_phuwuanensis | Mantheyus_phuwuanensis | 0 | 23.99 | 4 | 1887.29 |
| lizards | Maracaiba_meridensis | Maracaiba_meridensis | 0 | 9.12 | 4.5 | 11816.64 |
| lizards | Marisora_alliacea | Marisora_alliacea | 0 | 14.13 | 4.1 | 7785.38 |
| lizards | Marisora_brachypoda | Marisora_brachypoda | 0 | 15.49 | 5.5 | 865336.97 |
| lizards | Marisora_pergravis | Alinea_pergravis | 0 | 15.85 | 4.1 | 23.53 |
| lizards | Marisora_unimarginata | Marisora_unimarginata | 0 | 16.6 | 4.67 | 938239.04 |
| lizards | Marmorosphax_boulinda | Marmorosphax_boulinda | 1 | 2.75 | 2 | 99.94 |
| lizards | Marmorosphax_taom | Marmorosphax_taom | 1 | 3.02 | 2 | 155.74 |
| lizards | Marmorosphax_tricolor | Marmorosphax_tricolor | 0 | 5.62 | 3 | 14859.61 |
| lizards | Matobosaurus_maltzahni | Matobosaurus_maltzahni | 0 | 407.38 | 5 | 324748.28 |
| lizards | Matobosaurus_validus | Matobosaurus_validus | 0 | 616.6 | 4 | 820594.42 |
| lizards | Mediodactylus_amictopholis | Mediodactylus_amictopholis | 1 | 1.12 | 1 | 253.57 |
| lizards | Mediodactylus_heterocercus | Mediodactylus_heterocercus | 0 | 3.16 | 2 | 163494.2 |
| lizards | Mediodactylus_kotschyi | Mediodactylus_kotschyi | 0 | 3.8 | 1.63 | 122055.92 |
| lizards | Mediodactylus_russowii | Mediodactylus_russowii | 0 | 3.31 | 1.5 | 2115396.52 |
| lizards | Mediodactylus_spinicauda | Mediodactylus_spinicauda | 0 | 2.51 | 1.5 | 70327.02 |
| lizards | Melanoseps_ater | Melanoseps_ater | 0 | 10.23 | 3 | 85203.57 |
| lizards | Melanoseps_emmrichi | Melanoseps_emmrichi | 1 | 9.12 | 3 | 282.95 |
| lizards | Melanoseps_loveridgei | Melanoseps_loveridgei | 0 | 5.75 | 4.25 | 350996.05 |

| | | | | | | |
|---|---|---|---|---|---|---|
| lizards | Melanoseps_pygmaeus | Melanoseps_pygmaeus | 0 | 1.78 | 2 | 8008.98 |
| lizards | Menetia_alanae | Menetia_alanae | 0 | 0.44 | 1.65 | 138055.47 |
| lizards | Menetia_greyii | Menetia_greyii | 0 | 1.12 | 1.55 | 7074442.81 |
| lizards | Menetia_maini | Menetia_maini | 0 | 0.55 | 1.62 | 1319458.93 |
| lizards | Meroles_anchietae | Meroles_anchietae | 0 | 4.79 | 1.3 | 200330.97 |
| lizards | Meroles_ctenodactylus | Meroles_ctenodactylus | 0 | 20.89 | 6 | 58210.97 |
| lizards | Meroles_cuneirostris | Meroles_cuneirostris | 0 | 4.57 | 2.9 | 138082.05 |
| lizards | Meroles_knoxii | Meroles_knoxii | 0 | 7.24 | 4 | 165150.61 |
| lizards | Meroles_reticulatus | Meroles_reticulatus | 0 | 3.89 | 4 | 187250.75 |
| lizards | Meroles_squamulosus | Meroles_squamulosa | 0 | 11.48 | 4 | 3461354.19 |
| lizards | Meroles_suborbitalis | Meroles_suborbitalis | 0 | 8.32 | 4.6 | 701307.96 |
| lizards | Mesalina_adramitana | Mesalina_adramitana | 0 | 2.29 | 3 | 742653.62 |
| lizards | Mesalina_bahaeldini | Mesalina_bahaeldini | 0 | 4.17 | 4.25 | 647041.2 |
| lizards | Mesalina_balfouri | Mesalina_balfouri | 0 | 4.57 | 3 | 3631.2 |
| lizards | Mesalina_brevirostris | Mesalina_brevirostris | 0 | 5.01 | 3.9 | 1193015.6 |
| lizards | Mesalina_guttulata | Mesalina_guttulata | 0 | 5.01 | 4.6 | 9617162.62 |
| lizards | Mesalina_martini | Mesalina_martini | 0 | 2.95 | 3 | 100772.92 |
| lizards | Mesalina_olivieri | Mesalina_olivieri | 0 | 3.31 | 3.7 | 2601525.96 |
| lizards | Mesalina_pasteuri | Mesalina_pasteuri | 0 | 2.95 | 3 | 5154182.54 |
| lizards | Mesalina_rubropunctata | Mesalina_rubropunctata | 0 | 7.08 | 4.5 | 5592089.71 |
| lizards | Mesalina_simoni | Mesalina_simoni | 0 | 2.95 | 3 | 55851.97 |
| lizards | Mesalina_watsonana | Mesalina_watsonana | 0 | 5.01 | 2.5 | 2510586.89 |
| lizards | Micrablepharus_atticolus | Micrablepharus_atticolus | 0 | 2.34 | 1.88 | 1787612.21 |
| lizards | Micrablepharus_maximiliani | Micrablepharus_maximiliani | 0 | 1.86 | 2.25 | 4221760.12 |
| lizards | Microauris_aurantolabium | Calotes_aurantolabium | 1 | 16.22 | 4 | 621.17 |
| lizards | Microgecko_persicus | Microgecko_persicus | 0 | 1.12 | 1 | 182508.7 |
| lizards | Microlophus_albemarlensis | Microlophus_albemarlensis | 0 | 66.07 | 2.25 | 5106.21 |
| lizards | Microlophus_atacamensis | Microlophus_atacamensis | 0 | 72.44 | 4.5 | 18794.23 |
| lizards | Microlophus_delanonis | Microlophus_delanonis | 0 | 117.49 | 3.47 | 65.79 |
| lizards | Microlophus_habelii | Microlophus_habelii | 0 | 52.48 | 2.5 | 119.42 |
| lizards | Microlophus_koepckeorum | Microlophus_koepckeorum | 0 | 20.42 | 3.1 | 6021.48 |
| lizards | Microlophus_occipitalis | Microlophus_occipitalis | 0 | 19.5 | 4 | 97167.95 |
| lizards | Microlophus_pacificus | Microlophus_pacificus | 0 | 40.74 | 3 | 60.4 |
| lizards | Microlophus_peruvianus | Microlophus_peruvianus | 0 | 89.13 | 3.5 | 60949.56 |

| | | | | | | |
|---|---|---|---|---|---|---|
| lizards | Microlophus_quadrivittatus | Microlophus_quadrivittatus | 0 | 72.44 | 3.16 | 34776.74 |
| lizards | Microlophus_theresiae | Microlophus_theresiae | 0 | 42.66 | 3.4 | 11277.63 |
| lizards | Microlophus_theresioides | Microlophus_theresioides | 0 | 52.48 | 2.8 | 48894.59 |
| lizards | Microlophus_thoracicus | Microlophus_thoracicus | 0 | 30.2 | 2.8 | 54452.81 |
| lizards | Microlophus_tigris | Microlophus_tigris | 0 | 40.74 | 3.4 | 51891.66 |
| lizards | Mniarogekko_chahoua | Mniarogekko_chahoua | 1 | 66.07 | 2 | 3821.27 |
| lizards | Mniarogekko_jalu | Mniarogekko_jalu | 1 | 56.23 | 2 | 391.57 |
| lizards | Mochlus_brevicaudis | Mochlus_brevicaudis | 0 | 9.55 | 2.2 | 358660.48 |
| lizards | Mochlus_fernandi | Lepidothyris_fernandi | 0 | 144.54 | 10 | 2980336.33 |
| lizards | Mochlus_guineensis | Mochlus_guineensis | 0 | 10.47 | 2.48 | 2959270.67 |
| lizards | Mochlus_hinkeli | Lepidothyris_hinkeli | 0 | 72.44 | 11.5 | 828947.62 |
| lizards | Mochlus_striatus | Lepidothyris_striatus | 0 | 112.2 | 12 | 425436.3 |
| lizards | Mochlus_sundevallii | Mochlus_sundevalli | 0 | 63.1 | 3.87 | 7876204.79 |
| lizards | Mochlus_tanae | Mochlus_tanae | 0 | 5.13 | 2.5 | 340.59 |
| lizards | Mochlus_vinciguerrae | Mochlus_vinciguerrae | 0 | 5.13 | 3 | 19164.16 |
| lizards | Mokopirirakau_cryptozoicus | Mokopirirakau_cryptozoicus | 1 | 13.49 | 2 | 3963.55 |
| lizards | Mokopirirakau_granulatus | Mokopirirakau_granulatus | 1 | 19.05 | 2 | 118586.85 |
| lizards | Mokopirirakau_kahutarae | Mokopirirakau_kahutarae | 1 | 15.14 | 2 | 5639.71 |
| lizards | Mokopirirakau_nebulosus | Mokopirirakau_nebulosus | 0 | 14.79 | 2 | 995.26 |
| lizards | Moloch_horridus | Moloch_horridus | 0 | 61.66 | 7.25 | 3199451.13 |
| lizards | Monilesaurus_ellioti | Calotes_ellioti | 0 | 14.45 | 4 | 85602.92 |
| lizards | Monopeltis_adercae | Monopeltis_adercae | 0 | 158.49 | 5 | 5104.77 |
| lizards | Monopeltis_anchietae | Monopeltis_anchietae | 0 | 33.11 | 2.4 | 780879.93 |
| lizards | Monopeltis_capensis | Monopeltis_capensis | 0 | 31.62 | 2 | 821772.49 |
| lizards | Monopeltis_remaclei | Monopeltis_remaclei | 0 | 269.15 | 8 | 28696.72 |
| lizards | Morethia_adelaidensis | Morethia_adelaidensis | 0 | 4.17 | 3.3 | 1707088.54 |
| lizards | Morethia_boulengeri | Morethia_boulengeri | 0 | 3.47 | 3 | 3302096.32 |
| lizards | Morethia_butleri | Morethia_butleri | 0 | 3.47 | 3.7 | 1275905.51 |
| lizards | Morethia_lineoocellata | Morethia_lineoocellata | 0 | 3.47 | 2.75 | 259510.21 |
| lizards | Morethia_obscura | Morethia_obscura | 0 | 3.31 | 4 | 1568218.05 |
| lizards | Morethia_ruficauda | Morethia_ruficauda | 0 | 1.74 | 2 | 3133787.4 |
| lizards | Morethia_storri | Morethia_storri | 0 | 0.95 | 2.3 | 910508.76 |
| lizards | Morethia_taeniopleura | Morethia_taeniopleura | 0 | 1.51 | 2.35 | 739225.91 |
| lizards | Nactus_eboracensis | Nactus_eboracensis | 0 | 4.27 | 2 | 48422.38 |

| | | | | | | |
|---|---|---|---|---|---|---|
| lizards | Nactus_multicarinatus | Nactus_multicarinatus | 0 | 5.37 | 2 | 92954.72 |
| lizards | Nactus_pelagicus | Nactus_pelagicus | 0 | 10.23 | 1.9 | 266115.25 |
| lizards | Nactus_serpensinsula | Nactus_serpensinsula | 1 | 5.75 | 1 | 2.35 |
| lizards | Nactus_sphaerodactylodes | Nactus_sphaerodactylodes | 0 | 0.46 | 1 | 1921 |
| lizards | Nadzikambia_mlanjensis | Nadzikambia_mlanjensis | 1 | 14.45 | 7.5 | 199.02 |
| lizards | Namazonurus_campbelli | Namazonurus_campbelli | 0 | 11.48 | 2 | 6755.86 |
| lizards | Namazonurus_lawrenci | Namazonurus_lawrenci | 0 | 9.55 | 2.5 | 5891.69 |
| lizards | Namazonurus_namaquensis | Namazonurus_namaquensis | 0 | 13.18 | 2.5 | 5499.79 |
| lizards | Namazonurus_peersi | Namazonurus_peersi | 0 | 15.14 | 2.5 | 24004.95 |
| lizards | Namazonurus_pustulatus | Namazonurus_pustulatus | 0 | 13.18 | 1.33 | 17393.54 |
| lizards | Nannoscincus_garrulus | Nannoscincus_garrulus | 1 | 2.63 | 2 | 135.61 |
| lizards | Nannoscincus_gracilis | Nannoscincus_gracilis | 1 | 0.93 | 2 | 6301.31 |
| lizards | Nannoscincus_mariei | Nannoscincus_mariei | 1 | 1.74 | 1.5 | 3111.79 |
| lizards | Narudasia_festiva | Narudasia_festiva | 0 | 0.76 | 2 | 184589.51 |
| lizards | Naultinus_elegans | Naultinus_elegans | 1 | 6.92 | 2 | 42897.18 |
| lizards | Naultinus_gemmeus | Naultinus_gemmeus | 1 | 10.23 | 1.94 | 72027.27 |
| lizards | Naultinus_grayii | Naultinus_grayii | 0 | 17.38 | 2 | 1952.65 |
| lizards | Naultinus_manukanus | Naultinus_manukanus | 1 | 10.72 | 1.9 | 2842.96 |
| lizards | Naultinus_punctatus | Naultinus_punctatus | 1 | 17.38 | 1.5 | 56072.89 |
| lizards | Naultinus_rudis | Naultinus_rudis | 1 | 7.41 | 2 | 4131.88 |
| lizards | Naultinus_stellatus | Naultinus_stellatus | 1 | 10.23 | 2 | 11292.1 |
| lizards | Naultinus_tuberculatus | Naultinus_tuberculatus | 1 | 12.3 | 2 | 5594.21 |
| lizards | Nephrurus_amyae | Nephrurus_amyae | 0 | 39.81 | 2 | 111723.02 |
| lizards | Nephrurus_asper | Nephrurus_asper | 0 | 26.3 | 2 | 878623.09 |
| lizards | Nephrurus_deleani | Nephrurus_deleani | 0 | 18.2 | 2 | 8983.91 |
| lizards | Nephrurus_laevissimus | Nephrurus_laevissimus | 0 | 18.2 | 1.98 | 1629449.61 |
| lizards | Nephrurus_levis | Nephrurus_levis | 0 | 23.99 | 2 | 3537661.71 |
| lizards | Nephrurus_sheai | Nephrurus_sheai | 0 | 28.84 | 2 | 320983.27 |
| lizards | Nephrurus_stellatus | Nephrurus_stellatus | 0 | 16.6 | 2 | 180306.52 |
| lizards | Nephrurus_vertebralis | Nephrurus_vertebralis | 0 | 15.49 | 2 | 555857.82 |
| lizards | Nephrurus_wheeleri | Nephrurus_wheeleri | 0 | 18.62 | 2 | 508326.24 |
| lizards | Nessia_bipes | Nessia_bipes | 1 | 1.48 | 3 | 716.52 |
| lizards | Nessia_burtonii | Nessia_burtonii | 0 | 2.75 | 2 | 12090.07 |
| lizards | Nessia_layardi | Nessia_layardi | 1 | 2.14 | 2 | 3836.19 |

| | | | | | | |
|---|---|---|---|---|---|---|
| lizards | Nessia_monodactyla | Nessia_monodactyla | 1 | 2 | 1.5 | 13848.97 |
| lizards | Neusticurus_bicarinatus | Neusticurus_bicarinatus | 0 | 44.67 | 2 | 4356982.2 |
| lizards | Neusticurus_rudis | Neusticurus_rudis | 0 | 21.88 | 2 | 809214.03 |
| lizards | Neusticurus_tatei | Neusticurus_tatei | 0 | 30.2 | 2 | 317553.84 |
| lizards | Ninurta_coeruleopunctatus | Ninurta_coeruleopunctatus | 0 | 13.18 | 3.5 | 15088.56 |
| lizards | Nothobachia_ablephara | Nothobachia_ablephara | 0 | 1.7 | 2 | 59192.36 |
| lizards | Notomabuya_frenata | Notomabuya_frenata | 0 | 15.85 | 4.3 | 4419369.54 |
| lizards | Notoscincus_ornatus | Notoscincus_ornatus | 0 | 1.17 | 2.62 | 3561381.47 |
| lizards | Nucras_boulengeri | Nucras_boulengeri | 0 | 6.46 | 4 | 523266.74 |
| lizards | Nucras_holubi | Nucras_holubi | 0 | 3.16 | 4.7 | 791737.59 |
| lizards | Nucras_intertexta | Nucras_intertexta | 0 | 19.05 | 5 | 1010673.72 |
| lizards | Nucras_lalandii | Nucras_lalandii | 0 | 38.9 | 7 | 303549.88 |
| lizards | Nucras_livida | Nucras_livida | 0 | 14.13 | 4 | 135077.6 |
| lizards | Nucras_ornata | Nucras_ornata | 0 | 30.9 | 5.5 | 1027910.45 |
| lizards | Nucras_taeniolata | Nucras_taeniolata | 0 | 30.9 | 4.79 | 58948.73 |
| lizards | Nucras_tessellata | Nucras_tessellata | 0 | 19.05 | 3.96 | 697346.95 |
| lizards | Oedodera_marmorata | Oedodera_marmorata | 1 | 5.5 | 2 | 121.1 |
| lizards | Oedura_castelnaui | Oedura_castelnaui | 0 | 18.62 | 2 | 383086.6 |
| lizards | Oedura_coggeri | Oedura_coggeri | 0 | 10.72 | 2 | 98443.02 |
| lizards | Oedura_filicipoda | Oedura_filicipoda | 0 | 24.55 | 2 | 21336.45 |
| lizards | Oedura_gemmata | Oedura_gemmata | 0 | 27.54 | 2 | 38325.09 |
| lizards | Oedura_gracilis | Oedura_gracilis | 0 | 12.3 | 2 | 189248.19 |
| lizards | Oedura_jowalbinna | Oedura_jowalbinna | 0 | 6.61 | 2 | 66.23 |
| lizards | Oedura_marmorata | Oedura_marmorata | 0 | 27.54 | 2 | 221682.2 |
| lizards | Oedura_monilis | Oedura_monilis | 0 | 19.05 | 2 | 574184.24 |
| lizards | Oedura_murrumanu | Oedura_murrumanu | 0 | 22.39 | 2 | 174.62 |
| lizards | Oedura_tryoni | Oedura_tryoni | 0 | 22.39 | 2 | 250672.48 |
| lizards | Oligosoma_acrinasum | Oligosoma_acrinasum | 0 | 14.13 | 4 | 1579.71 |
| lizards | Oligosoma_aeneum | Oligosoma_aeneum | 0 | 8.91 | 2.2 | 100139.8 |
| lizards | Oligosoma_alani | Oligosoma_alani | 1 | 79.43 | 3 | 103.06 |
| lizards | Oligosoma_burganae | Oligosoma_burganae | 1 | 6.76 | 5.5 | 19.98 |
| lizards | Oligosoma_chloronoton | Oligosoma_chloronoton | 1 | 44.67 | 2.5 | 37946.17 |
| lizards | Oligosoma_fallai | Oligosoma_fallai | 1 | 70.79 | 4.5 | 4.57 |
| lizards | Oligosoma_grande | Oligosoma_grande | 1 | 37.15 | 2.13 | 9040.37 |

| | | | | | | |
|---|---|---|---|---|---|---|
| lizards | Oligosoma_hardyi | Oligosoma_hardyi | 1 | 4.57 | 5 | 2.01 |
| lizards | Oligosoma_homalonotum | Oligosoma_homalonotum | 1 | 72.44 | 7.5 | 313.85 |
| lizards | Oligosoma_inconspicuum | Oligosoma_inconspicuum | 1 | 8.51 | 3 | 42485.12 |
| lizards | Oligosoma_infrapunctatum | Oligosoma_infrapunctatum | 1 | 33.88 | 4 | 44858.16 |
| lizards | Oligosoma_judgei | Oligosoma_judgei | 1 | 15.85 | 4 | 795.61 |
| lizards | Oligosoma_levidensum | Oligosoma_levidensum | 1 | 2.4 | 5 | 509.57 |
| lizards | Oligosoma_lineoocellatum | Oligosoma_lineoocellatum | 1 | 30.2 | 2.5 | 10139.38 |
| lizards | Oligosoma_longipes | Oligosoma_longipes | 1 | 8.51 | 4 | 30550.55 |
| lizards | Oligosoma_maccanni | Oligosoma_maccanni | 0 | 9.55 | 3.25 | 57177.21 |
| lizards | Oligosoma_macgregori | Oligosoma_macgregori | 1 | 35.48 | 2 | 20.34 |
| lizards | Oligosoma_microlepis | Oligosoma_microlepis | 1 | 7.76 | 2.5 | 7616.18 |
| lizards | Oligosoma_moco | Oligosoma_moco | 0 | 10.96 | 4 | 4289.08 |
| lizards | Oligosoma_nigriplantare | Oligosoma_nigriplantare | 0 | 19.95 | 4.47 | 74.15 |
| lizards | Oligosoma_notosaurus | Oligosoma_notosaurus | 0 | 15.49 | 2.5 | 1288.43 |
| lizards | Oligosoma_oliveri | Oligosoma_oliveri | 0 | 34.67 | 3.3 | 47.97 |
| lizards | Oligosoma_ornatum | Oligosoma_ornatum | 0 | 13.8 | 5 | 94721.43 |
| lizards | Oligosoma_otagense | Oligosoma_otagense | 1 | 117.49 | 2.17 | 8968.58 |
| lizards | Oligosoma_polychroma | Oligosoma_polychroma | 0 | 10.47 | 4.8 | 177591.86 |
| lizards | Oligosoma_smithi | Oligosoma_smithi | 0 | 11.22 | 2.6 | 14624.13 |
| lizards | Oligosoma_stenotis | Oligosoma_stenotis | 1 | 8.51 | 3 | 820.07 |
| lizards | Oligosoma_striatum | Oligosoma_striatum | 1 | 8.91 | 5.5 | 23867.62 |
| lizards | Oligosoma_suteri | Oligosoma_suteri | 0 | 45.71 | 3.65 | 13451.26 |
| lizards | Oligosoma_whitakeri | Oligosoma_whitakeri | 1 | 31.62 | 2 | 136.68 |
| lizards | Oligosoma_zelandicum | Oligosoma_zelandicum | 0 | 8.51 | 5.15 | 13845.24 |
| lizards | Omanosaura_cyanura | Omanosaura_cyanura | 0 | 5.01 | 2.83 | 23800.1 |
| lizards | Omanosaura_jayakari | Omanosaura_jayakari | 0 | 177.83 | 6 | 35859.7 |
| lizards | Ophiodes_fragilis | Ophiodes_fragilis | 0 | 81.28 | 5.55 | 1668843.42 |
| lizards | Ophiodes_intermedius | Ophiodes_intermedius | 0 | 63.1 | 8.7 | 1342019.39 |
| lizards | Ophiodes_striatus | Ophiodes_striatus | 0 | 102.33 | 7.3 | 978405.1 |
| lizards | Ophiodes_vertebralis | Ophiodes_vertebralis | 0 | 33.88 | 11.5 | 243664.34 |
| lizards | Ophiomorus_nuchalis | Ophiomorus_nuchalis | 0 | 7.41 | 4 | 10407.95 |
| lizards | Ophiomorus_punctatissimus | Ophiomorus_punctatissimus | 0 | 2.45 | 3 | 19823.91 |
| lizards | Ophiomorus_tridactylus | Ophiomorus_tridactylus | 0 | 6.17 | 4 | 385438.01 |
| lizards | Ophioscincus_cooloolensis | Ophioscincus_cooloolensis | 0 | 1.1 | 3 | 18332.64 |

| | | | | | | |
|---|---|---|---|---|---|---|
| lizards | Ophioscincus_ophioscincus | Ophioscincus_ophioscincus | 0 | 2.29 | 2.5 | 41069.91 |
| lizards | Ophioscincus_truncatus | Ophioscincus_truncatus | 0 | 1.45 | 2.5 | 37032.49 |
| lizards | Ophisaurus_attenuatus | Ophisaurus_attenuatus | 0 | 147.91 | 9.74 | 1936754.71 |
| lizards | Ophisaurus_compressus | Ophisaurus_compressus | 0 | 25.12 | 11 | 149110.7 |
| lizards | Ophisaurus_mimicus | Ophisaurus_mimicus | 0 | 20.89 | 6 | 105948.08 |
| lizards | Ophisaurus_ventralis | Ophisaurus_ventralis | 0 | 91.2 | 11 | 472335.08 |
| lizards | Ophisops_elegans | Ophisops_elegans | 0 | 7.94 | 3.57 | 2393067.59 |
| lizards | Ophisops_jerdonii | Ophisops_jerdonii | 0 | 2.75 | 4.5 | 2891076.11 |
| lizards | Ophisops_leschenaultii | Ophisops_leschenaultii | 0 | 4.37 | 6 | 1300623.34 |
| lizards | Ophisops_microlepis | Ophisops_microlepis | 0 | 6.46 | 7 | 664093.06 |
| lizards | Ophisops_occidentalis | Ophisops_occidentalis | 0 | 2.63 | 3.5 | 333927.52 |
| lizards | Oplurus_cuvieri | Oplurus_cuvieri | 0 | 177.83 | 3.5 | 207380.26 |
| lizards | Oplurus_cyclurus | Oplurus_cyclurus | 0 | 177.83 | 4 | 230746.92 |
| lizards | Oplurus_fierinensis | Oplurus_fierinensis | 0 | 27.54 | 2.5 | 9875.94 |
| lizards | Oplurus_grandidieri | Oplurus_grandidieri | 0 | 52.48 | 2 | 16124.11 |
| lizards | Oplurus_quadrimaculatus | Oplurus_quadrimaculatus | 0 | 120.23 | 7 | 149236.18 |
| lizards | Oplurus_saxicola | Oplurus_saxicola | 0 | 38.9 | 2 | 67455.68 |
| lizards | Oreosaurus_achlyens | Riama_achlyens | 0 | 17.38 | 2 | 4024.02 |
| lizards | Oreosaurus_rhodogaster | Riama_rhodogaster | 1 | 2.24 | 2 | 9.99 |
| lizards | Oreosaurus_shrevei | Riama_shrevei | 0 | 2.69 | 2 | 277.89 |
| lizards | Orraya_occultus | Orraya_occultus | 1 | 21.88 | 2 | 617.73 |
| lizards | Otocryptis_nigristigma | Otocryptis_nigristigma | 0 | 16.6 | 4 | 26990.7 |
| lizards | Otocryptis_wiegmanni | Otocryptis_wiegmanni | 0 | 16.6 | 4 | 17038.49 |
| lizards | Ouroborus_cataphractus | Ouroborus_cataphractus | 0 | 58.88 | 1 | 95329.77 |
| lizards | Pachydactylus_acuminatus | Pachydactylus_acuminatus | 0 | 2.82 | 1 | 25198.33 |
| lizards | Pachydactylus_affinis | Pachydactylus_affinis | 0 | 6.61 | 1 | 132821.15 |
| lizards | Pachydactylus_amoenus | Pachydactylus_amoenus | 0 | 1.12 | 2 | 3091.41 |
| lizards | Pachydactylus_atorquatus | Pachydactylus_atorquatus | 0 | 3.55 | 2 | 21400.86 |
| lizards | Pachydactylus_austeni | Pachydactylus_austeni | 0 | 2.34 | 2 | 42139.86 |
| lizards | Pachydactylus_barnardi | Pachydactylus_barnardi | 0 | 5.25 | 1 | 34338.92 |
| lizards | Pachydactylus_bicolor | Pachydactylus_bicolor | 0 | 1.86 | 2 | 231661.68 |
| lizards | Pachydactylus_boehmei | Pachydactylus_boehmei | 0 | 2.04 | 2 | 2040.26 |
| lizards | Pachydactylus_capensis | Pachydactylus_capensis | 0 | 6.61 | 1.97 | 1796090.87 |
| lizards | Pachydactylus_carinatus | Pachydactylus_carinatus | 0 | 2.19 | 2 | 315481.56 |

| | | | | | | |
|---|---|---|---|---|---|---|
| lizards | Pachydactylus_fasciatus | Pachydactylus_fasciatus | 0 | 3.8 | 1.5 | 124432.49 |
| lizards | Pachydactylus_formosus | Pachydactylus_formosus | 0 | 4.68 | 1 | 62618.72 |
| lizards | Pachydactylus_geitje | Pachydactylus_geitje | 0 | 2.09 | 2 | 132809.23 |
| lizards | Pachydactylus_griffini | Pachydactylus_griffini | 0 | 1.45 | 2 | 22217.51 |
| lizards | Pachydactylus_haackei | Pachydactylus_haackei | 0 | 12.3 | 1.5 | 99726.08 |
| lizards | Pachydactylus_katanganus | Pachydactylus_katanganus | 0 | 4.07 | 1 | 54523.88 |
| lizards | Pachydactylus_kobosensis | Pachydactylus_kobosensis | 0 | 3.39 | 2 | 9742.77 |
| lizards | Pachydactylus_labialis | Pachydactylus_labialis | 0 | 2.24 | 1 | 68381.98 |
| lizards | Pachydactylus_maculatus | Pachydactylus_maculatus | 0 | 4.27 | 2 | 338309.26 |
| lizards | Pachydactylus_maraisi | Pachydactylus_maraisi | 0 | 1.62 | 2 | 1202.65 |
| lizards | Pachydactylus_mariquensis | Pachydactylus_mariquensis | 0 | 4.27 | 2 | 351788.39 |
| lizards | Pachydactylus_mclachlani | Pachydactylus_mclachlani | 0 | 2.63 | 2 | 21243.02 |
| lizards | Pachydactylus_monicae | Pachydactylus_monicae | 0 | 2.88 | 2 | 10223.84 |
| lizards | Pachydactylus_montanus | Pachydactylus_montanus | 0 | 1.91 | 2 | 107345.2 |
| lizards | Pachydactylus_namaquensis | Pachydactylus_namaquensis | 0 | 12.3 | 2 | 27679.32 |
| lizards | Pachydactylus_oculatus | Pachydactylus_oculatus | 0 | 3.31 | 2 | 122997.76 |
| lizards | Pachydactylus_otaviensis | Pachydactylus_otaviensis | 0 | 1.86 | 2 | 19.98 |
| lizards | Pachydactylus_parascutatus | Pachydactylus_parascutatus | 0 | 1.35 | 2 | 9094.75 |
| lizards | Pachydactylus_punctatus | Pachydactylus_punctatus | 0 | 1.74 | 2 | 3247237.48 |
| lizards | Pachydactylus_purcelli | Pachydactylus_purcelli | 0 | 2.95 | 2 | 217357.64 |
| lizards | Pachydactylus_rangei | Pachydactylus_rangei | 0 | 10.23 | 2 | 229410.87 |
| lizards | Pachydactylus_reconditus | Pachydactylus_reconditus | 0 | 2.04 | 2 | 87652.43 |
| lizards | Pachydactylus_robertsi | Pachydactylus_robertsi | 0 | 1.74 | 2 | 19.98 |
| lizards | Pachydactylus_rugosus | Pachydactylus_rugosus | 0 | 6.03 | 2 | 476375.14 |
| lizards | Pachydactylus_sansteynae | Pachydactylus_sansteynae | 0 | 2.51 | 2 | 47434.14 |
| lizards | Pachydactylus_scutatus | Pachydactylus_scutatus | 0 | 2.34 | 2 | 217542.83 |
| lizards | Pachydactylus_serval | Pachydactylus_serval | 0 | 3.39 | 2 | 199116.19 |
| lizards | Pachydactylus_tigrinus | Pachydactylus_tigrinus | 0 | 3.31 | 1.5 | 273512.4 |
| lizards | Pachydactylus_tsodiloensis | Pachydactylus_tsodiloensis | 0 | 4.68 | 1.95 | 9.99 |
| lizards | Pachydactylus_vansoni | Pachydactylus_vansoni | 0 | 4.47 | 2 | 346112.43 |
| lizards | Pachydactylus_visseri | Pachydactylus_visseri | 0 | 2.57 | 2 | 10976.91 |
| lizards | Pachydactylus_wahlbergii | Colopus_wahlbergii | 0 | 4.9 | 2 | 834161.5 |
| lizards | Pachydactylus_weberi | Pachydactylus_weberi | 0 | 2.82 | 2 | 264086.27 |
| lizards | Pachydactylus_werneri | Pachydactylus_werneri | 0 | 3.24 | 1 | 23199.68 |

| | | | | | | |
|---|---|---|---|---|---|---|
| lizards | Palleon_lolontany | Palleon_lolontany | 0 | 1.1 | 3 | 800.24 |
| lizards | Pamelaescincus_gardineri | Pamelaescincus_gardineri | 0 | 11.22 | 2 | 237.62 |
| lizards | Panaspis_breviceps | Panaspis_breviceps | 0 | 6.76 | 2.5 | 930288.51 |
| lizards | Panaspis_helleri | Panaspis_helleri | 0 | 4.57 | 3 | 29.97 |
| lizards | Panaspis_maculicollis | Afroablepharus_maculicollis | 0 | 1.62 | 3 | 863948.32 |
| lizards | Panaspis_togoensis | Panaspis_togoensis | 0 | 3.09 | 1.92 | 2676662.93 |
| lizards | Panaspis_tristaoi | Panaspis_nimbaensis | 0 | 6.17 | 2.51 | 533551.18 |
| lizards | Paniegekko_madjo | Paniegekko_madjo | 1 | 8.51 | 2 | 388.54 |
| lizards | Papuascincus_morokanus | Papuascincus_morokanus | 0 | 2.63 | 2 | 112736.52 |
| lizards | Papuascincus_stanleyanus | Papuascincus_stanleyanus | 0 | 4.17 | 2 | 184604.11 |
| lizards | Paracontias_minimus | Paracontias_minimus | 1 | 1.26 | 4 | 89.34 |
| lizards | Paradelma_orientalis | Paradelma_orientalis | 0 | 12.88 | 2 | 407132.73 |
| lizards | Paragehyra_gabriellae | Paragehyra_gabriellae | 1 | 8.71 | 2 | 1348.88 |
| lizards | Paralaudakia_bochariensis | Paralaudakia_bochariensis | 0 | 58.88 | 4.5 | 100406.9 |
| lizards | Paralaudakia_caucasia | Paralaudakia_caucasia | 0 | 186.21 | 9.45 | 1711153.78 |
| lizards | Paralaudakia_erythrogaster | Paralaudakia_erythrogaster | 0 | 128.82 | 12.5 | 237044.87 |
| lizards | Paralaudakia_himalayana | Paralaudakia_himalayana | 0 | 107.15 | 3.5 | 549740.63 |
| lizards | Paralaudakia_lehmanni | Paralaudakia_lehmanni | 0 | 117.49 | 10 | 386332.21 |
| lizards | Paralaudakia_stoliczkana | Paralaudakia_stoliczkana | 0 | 208.93 | 7.5 | 508336.61 |
| lizards | Paroedura_androyensis | Paroedura_androyensis | 1 | 2.34 | 2 | 19244.04 |
| lizards | Paroedura_bastardi | Paroedura_bastardi | 0 | 11.75 | 2 | 172114.55 |
| lizards | Paroedura_gracilis | Paroedura_gracilis | 0 | 7.76 | 2 | 38038.34 |
| lizards | Paroedura_ibityensis | Paroedura_ibityensis | 0 | 4.9 | 2 | 2999.55 |
| lizards | Paroedura_karstophila | Paroedura_karstophila | 0 | 3.63 | 2 | 4776.62 |
| lizards | Paroedura_lohatsara | Paroedura_lohatsara | 1 | 10.47 | 2 | 288.1 |
| lizards | Paroedura_masobe | Paroedura_masobe | 1 | 25.7 | 2 | 121.67 |
| lizards | Paroedura_oviceps | Paroedura_oviceps | 0 | 7.41 | 2 | 30337.09 |
| lizards | Paroedura_picta | Paroedura_picta | 0 | 25.7 | 1.89 | 136497.28 |
| lizards | Paroedura_stellata | Paroedura_stellata | 1 | 5.01 | 2 | 332.63 |
| lizards | Paroedura_stumpffi | Paroedura_stumpffi | 0 | 8.51 | 2 | 59105.43 |
| lizards | Paroedura_tanjaka | Paroedura_tanjaka | 1 | 19.95 | 2 | 1302.97 |
| lizards | Paroedura_vahiny | Paroedura_vahiny | 0 | 1.74 | 1 | 55037.06 |
| lizards | Paroedura_vazimba | Paroedura_vazimba | 1 | 2.82 | 2 | 8442.64 |
| lizards | Parvilacerta_fraasii | Parvilacerta_fraasii | 1 | 5.01 | 5.8 | 1418.9 |

| | | | | | | |
|---|---|---|---|---|---|---|
| lizards | Parvilacerta_parva | Parvilacerta_parva | 0 | 5.62 | 3 | 293339.17 |
| lizards | Parvoscincus_beyeri | Parvoscincus_beyeri | 1 | 7.76 | 2 | 17317.82 |
| lizards | Parvoscincus_kitangladensis | Parvoscincus_kitangladensis | 0 | 3.47 | 1.5 | 3697.32 |
| lizards | Parvoscincus_palawanensis | Parvoscincus_palawanensis | 1 | 0.72 | 1 | 19.98 |
| lizards | Parvoscincus_sisoni | Parvoscincus_sisoni | 1 | 0.83 | 1 | 9.99 |
| lizards | Parvoscincus_steerei | Parvoscincus_steerei | 0 | 1.32 | 1.05 | 268427.77 |
| lizards | Parvoscincus_tagapayo | Parvoscincus_tagapayo | 0 | 0.55 | 2 | 2455.43 |
| lizards | Pedioplanis_breviceps | Pedioplanis_breviceps | 0 | 2.29 | 3 | 149069.04 |
| lizards | Pedioplanis_burchelli | Pedioplanis_burchelli | 0 | 5.75 | 4.5 | 371177.28 |
| lizards | Pedioplanis_husabensis | Pedioplanis_husabensis | 0 | 5.37 | 1.5 | 19159.9 |
| lizards | Pedioplanis_inornata | Pedioplanis_inornata | 0 | 3.89 | 5.5 | 295832.05 |
| lizards | Pedioplanis_lineoocellata | Pedioplanis_lineoocellata | 0 | 7.08 | 6.88 | 1985439.31 |
| lizards | Pedioplanis_namaquensis | Pedioplanis_namaquensis | 0 | 4.17 | 3.9 | 1464925.1 |
| lizards | Pedioplanis_rubens | Pedioplanis_rubens | 0 | 2.95 | 4 | 879.55 |
| lizards | Pedioplanis_undata | Pedioplanis_undata | 0 | 5.62 | 2.5 | 611403.52 |
| lizards | Perochirus_ateles | Perochirus_ateles | 1 | 14.13 | 2.05 | 1550.83 |
| lizards | Perochirus_scutellatus | Perochirus_scutellatus | 0 | 43.65 | 2 | 2.25 |
| lizards | Petracola_ventrimaculatus | Petracola_ventrimaculatus | 1 | 6.46 | 2 | 19058.38 |
| lizards | Petrosaurus_mearnsi | Petrosaurus_mearnsi | 0 | 37.15 | 4 | 24622.52 |
| lizards | Petrosaurus_thalassinus | Petrosaurus_thalassinus | 0 | 141.25 | 13.8 | 7545.62 |
| lizards | Phasmasaurus_maruia | Lioscincus_maruia | 1 | 4.37 | 2.5 | 905.26 |
| lizards | Phasmasaurus_tillieri | Lioscincus_tillieri | 0 | 5.13 | 2.5 | 3316.01 |
| lizards | Phelsuma_abbotti | Phelsuma_abbotti | 0 | 5.75 | 1.5 | 33813.53 |
| lizards | Phelsuma_andamanensis | Phelsuma_andamanense | 0 | 5.5 | 2 | 4566.09 |
| lizards | Phelsuma_antanosy | Phelsuma_antanosy | 1 | 2.51 | 2 | 524.92 |
| lizards | Phelsuma_astriata | Phelsuma_astriata | 0 | 4.68 | 2 | 242.92 |
| lizards | Phelsuma_barbouri | Phelsuma_barbouri | 0 | 5.5 | 1.5 | 573.17 |
| lizards | Phelsuma_berghofi | Phelsuma_berghofi | 0 | 4.27 | 1.5 | 1982.53 |
| lizards | Phelsuma_borbonica | Phelsuma_borbonica | 1 | 7.94 | 2 | 34.38 |
| lizards | Phelsuma_breviceps | Phelsuma_breviceps | 1 | 3.47 | 2 | 9278.48 |
| lizards | Phelsuma_cepediana | Phelsuma_cepediana | 0 | 4.68 | 2 | 1872.58 |
| lizards | Phelsuma_comorensis | Phelsuma_comorensis | 0 | 3.8 | 2 | 1031.55 |
| lizards | Phelsuma_dorsivittata | Phelsuma_dorsivittata | 0 | 4.07 | 2 | 975.47 |
| lizards | Phelsuma_dubia | Phelsuma_dubia | 0 | 7.08 | 2 | 25456.11 |

| | | | | | | |
|---|---|---|---|---|---|---|
| lizards | Phelsuma_grandis | Phelsuma_grandis | 0 | 30.2 | 2 | 9173.23 |
| lizards | Phelsuma_guentheri | Phelsuma_guentheri | 1 | 69.18 | 1.5 | 1872.61 |
| lizards | Phelsuma_guimbeaui | Phelsuma_guimbeaui | 1 | 7.08 | 2 | 1877.19 |
| lizards | Phelsuma_guttata | Phelsuma_guttata | 0 | 6.61 | 1.5 | 33638.22 |
| lizards | Phelsuma_inexpectata | Phelsuma_inexpectata | 1 | 4.27 | 1.5 | 383.25 |
| lizards | Phelsuma_klemmeri | Phelsuma_klemmeri | 1 | 1.86 | 1.5 | 958.94 |
| lizards | Phelsuma_kochi | Phelsuma_kochi | 0 | 28.18 | 1.5 | 66294.7 |
| lizards | Phelsuma_laticauda | Phelsuma_laticauda | 0 | 5.75 | 2 | 39094.21 |
| lizards | Phelsuma_lineata | Phelsuma_lineata | 0 | 5.75 | 1.88 | 224353.23 |
| lizards | Phelsuma_madagascariensis | Phelsuma_madagascariensis | 0 | 31.62 | 1.88 | 79710.52 |
| lizards | Phelsuma_malamakibo | Phelsuma_malamakibo | 0 | 4.9 | 2 | 836.71 |
| lizards | Phelsuma_masohoala | Phelsuma_masohoala | 1 | 2.34 | 2.5 | 841.31 |
| lizards | Phelsuma_modesta | Phelsuma_modesta | 0 | 2.24 | 1.5 | 25546.69 |
| lizards | Phelsuma_mutabilis | Phelsuma_mutabilis | 0 | 2.82 | 1.5 | 103490.07 |
| lizards | Phelsuma_nigristriata | Phelsuma_nigristriata | 1 | 2.09 | 1.5 | 386.17 |
| lizards | Phelsuma_ornata | Phelsuma_ornata | 0 | 4.47 | 2 | 882.07 |
| lizards | Phelsuma_parkeri | Phelsuma_parkeri | 0 | 6.92 | 2 | 908.25 |
| lizards | Phelsuma_parva | Phelsuma_parva | 0 | 1.12 | 2 | 39161.32 |
| lizards | Phelsuma_pronki | Phelsuma_pronki | 1 | 2.82 | 2 | 3452.38 |
| lizards | Phelsuma_pusilla | Phelsuma_pusilla | 0 | 1.23 | 2 | 75550.24 |
| lizards | Phelsuma_quadriocellata | Phelsuma_quadriocellata | 0 | 4.9 | 2 | 96380.24 |
| lizards | Phelsuma_robertmertensi | Phelsuma_robertmertensi | 0 | 2.82 | 1.5 | 386.17 |
| lizards | Phelsuma_rosagularis | Phelsuma_rosagularis | 1 | 6.92 | 2 | 1872.58 |
| lizards | Phelsuma_seippi | Phelsuma_seippi | 1 | 3.63 | 1.5 | 3751.59 |
| lizards | Phelsuma_serraticauda | Phelsuma_serraticauda | 1 | 6.03 | 1.5 | 6783.58 |
| lizards | Phelsuma_standingi | Phelsuma_standingi | 1 | 15.49 | 1.5 | 17234.11 |
| lizards | Phelsuma_sundbergi | Phelsuma_sundbergi | 0 | 24.55 | 1.5 | 263.25 |
| lizards | Phelsuma_v-nigra | Phelsuma_v-nigra | 0 | 3.31 | 2 | 2039.61 |
| lizards | Phoenicolacerta_kulzeri | Phoenicolacerta_kulzeri | 1 | 15.14 | 5.5 | 14652.33 |
| lizards | Phoenicolacerta_laevis | Phoenicolacerta_laevis | 0 | 15.14 | 5.3 | 70636.46 |
| lizards | Phoenicolacerta_troodica | Phoenicolacerta_troodica | 0 | 7.94 | 5 | 9272.65 |
| lizards | Pholidobolus_affinis | Pholidobolus_affinis | 0 | 6.46 | 2 | 1731.87 |
| lizards | Pholidobolus_macbrydei | Pholidobolus_macbrydei | 0 | 4.9 | 2 | 29668.99 |
| lizards | Pholidobolus_montium | Pholidobolus_montium | 0 | 6.76 | 2 | 5905.26 |

| | | | | | | |
|---|---|---|---|---|---|---|
| lizards | Pholidobolus_prefrontalis | Pholidobolus_prefrontalis | 0 | 5.75 | 2 | 1269.34 |
| lizards | Pholidobolus_vertebralis | Cercosaura_vertebralis | 0 | 7.41 | 2 | 149316.92 |
| lizards | Pholidoscelis_auberi | Ameiva_auberi | 0 | 77.62 | 1 | 111513.76 |
| lizards | Pholidoscelis_chrysolaemus | Ameiva_chrysolaema | 0 | 128.82 | 5 | 52162.88 |
| lizards | Pholidoscelis_exsul | Ameiva_exsul | 0 | 263.03 | 2.68 | 9254.63 |
| lizards | Pholidoscelis_fuscatus | Ameiva_fuscata | 0 | 512.86 | 4 | 660.07 |
| lizards | Pholidoscelis_plei | Ameiva_plei | 0 | 186.21 | 2.6 | 199.36 |
| lizards | Pholidoscelis_taeniurus | Ameiva_taeniura | 0 | 34.67 | 4 | 62690.91 |
| lizards | Pholidoscelis_wetmorei | Ameiva_wetmorei | 1 | 3.98 | 1 | 495.87 |
| lizards | Phrynocephalus_arabicus | Phrynocephalus_arabicus | 0 | 6.92 | 3.5 | 1958315.28 |
| lizards | Phrynocephalus_axillaris | Phrynocephalus_axillaris | 0 | 8.13 | 2.6 | 1029671.95 |
| lizards | Phrynocephalus_clarkorum | Phrynocephalus_clarkorum | 0 | 2.82 | 4 | 130981.09 |
| lizards | Phrynocephalus_erythrurus | Phrynocephalus_erythrurus | 0 | 5.62 | 2.21 | 371161.08 |
| lizards | Phrynocephalus_forsythii | Phrynocephalus_forsythii | 0 | 7.08 | 4.8 | 343495.01 |
| lizards | Phrynocephalus_golubewii | Phrynocephalus_golubewii | 1 | 10.96 | 2.5 | 193.25 |
| lizards | Phrynocephalus_guttatus | Phrynocephalus_guttatus | 0 | 8.71 | 3.23 | 2125606.43 |
| lizards | Phrynocephalus_helioscopus | Phrynocephalus_helioscopus | 0 | 10.96 | 3.7 | 2586056.74 |
| lizards | Phrynocephalus_interscapularis | Phrynocephalus_interscapularis | 0 | 3.24 | 1 | 1167045.41 |
| lizards | Phrynocephalus_luteoguttatus | Phrynocephalus_luteoguttatus | 0 | 3.24 | 3 | 291725.81 |
| lizards | Phrynocephalus_maculatus | Phrynocephalus_maculatus | 0 | 25.12 | 2.5 | 2828625.89 |
| lizards | Phrynocephalus_mystaceus | Phrynocephalus_mystaceus | 0 | 63.1 | 5 | 2820240.64 |
| lizards | Phrynocephalus_ornatus | Phrynocephalus_ornatus | 0 | 3.24 | 3.5 | 364906.62 |
| lizards | Phrynocephalus_persicus | Phrynocephalus_persicus | 1 | 6.92 | 4.5 | 355797.13 |
| lizards | Phrynocephalus_przewalskii | Phrynocephalus_przewalskii | 0 | 23.99 | 2.85 | 3274656.58 |
| lizards | Phrynocephalus_putjatai | Phrynocephalus_putjatai | 0 | 19.5 | 3.1 | 11425.2 |
| lizards | Phrynocephalus_raddei | Phrynocephalus_raddei | 0 | 6.17 | 4 | 315414.94 |
| lizards | Phrynocephalus_rossikowi | Phrynocephalus_rossikowi | 1 | 3.89 | 4 | 88378.17 |
| lizards | Phrynocephalus_scutellatus | Phrynocephalus_scutellatus | 0 | 5.5 | 4 | 1614986.16 |
| lizards | Phrynocephalus_strauchi | Phrynocephalus_strauchi | 1 | 5.25 | 3.5 | 24633.99 |
| lizards | Phrynocephalus_theobaldi | Phrynocephalus_theobaldi | 0 | 8.13 | 2.2 | 1631165.54 |
| lizards | Phrynocephalus_versicolor | Phrynocephalus_versicolor | 0 | 9.55 | 3.3 | 4002944.43 |
| lizards | Phrynocephalus_vlangalii | Phrynocephalus_vlangalii | 0 | 18.62 | 3.36 | 1144045.77 |
| lizards | Phrynosoma_asio | Phrynosoma_asio | 0 | 56.23 | 16.9 | 115543.61 |
| lizards | Phrynosoma_blainvillii | Phrynosoma_blainvillii | 0 | 44.67 | 15.5 | 216760.1 |

| | | | | | | |
|---|---|---|---|---|---|---|
| lizards | Phrynosoma_braconnieri | Phrynosoma_braconnieri | 0 | 16.6 | 7.7 | 12632.63 |
| lizards | Phrynosoma_cornutum | Phrynosoma_cornutum | 0 | 63.1 | 27.9 | 1798114.31 |
| lizards | Phrynosoma_coronatum | Phrynosoma_coronatum | 0 | 44.67 | 11.8 | 287997.75 |
| lizards | Phrynosoma_douglasii | Phrynosoma_douglasii | 0 | 17.38 | 15 | 209725.16 |
| lizards | Phrynosoma_goodei | Phrynosoma_goodei | 0 | 23.99 | 9 | 72654.2 |
| lizards | Phrynosoma_hernandesi | Phrynosoma_hernandesi | 0 | 57.54 | 18 | 1210130.55 |
| lizards | Phrynosoma_mcallii | Phrynosoma_mcallii | 0 | 39.81 | 4.95 | 31422.87 |
| lizards | Phrynosoma_modestum | Phrynosoma_modestum | 0 | 12.59 | 11 | 811535.93 |
| lizards | Phrynosoma_orbiculare | Phrynosoma_orbiculare | 0 | 28.84 | 12.65 | 448940.88 |
| lizards | Phrynosoma_platyrhinos | Phrynosoma_platyrhinos | 0 | 27.54 | 5.52 | 836125.64 |
| lizards | Phrynosoma_solare | Phrynosoma_solare | 0 | 47.86 | 18.9 | 330439.41 |
| lizards | Phrynosoma_taurus | Phrynosoma_taurus | 0 | 23.99 | 13 | 26989.03 |
| lizards | Phyllodactylus_angustidigitus | Phyllodactylus_angustidigitus | 1 | 4.9 | 1 | 1017.36 |
| lizards | Phyllodactylus_bordai | Phyllodactylus_bordai | 0 | 5.62 | 1.5 | 21631.01 |
| lizards | Phyllodactylus_bugastrolepis | Phyllodactylus_bugastrolepis | 0 | 6.61 | 1.5 | 27.73 |
| lizards | Phyllodactylus_davisi | Phyllodactylus_davisi | 0 | 8.51 | 1.5 | 1794.77 |
| lizards | Phyllodactylus_delcampoi | Phyllodactylus_delcampoi | 0 | 22.91 | 1.5 | 100.48 |
| lizards | Phyllodactylus_delsolari | Phyllodactylus_delsolari | 1 | 18.2 | 2 | 214.7 |
| lizards | Phyllodactylus_duellmani | Phyllodactylus_duellmani | 0 | 2.14 | 1.5 | 1646.19 |
| lizards | Phyllodactylus_galapagensis | Phyllodactylus_galapagensis | 0 | 2.57 | 1 | 4717.6 |
| lizards | Phyllodactylus_gerrhopygus | Phyllodactylus_gerrhopygus | 0 | 4.68 | 1.5 | 125801.39 |
| lizards | Phyllodactylus_inaequalis | Phyllodactylus_inaequalis | 0 | 2 | 1 | 65967.18 |
| lizards | Phyllodactylus_insularis | Phyllodactylus_insularis | 1 | 8.51 | 2 | 11.47 |
| lizards | Phyllodactylus_interandinus | Phyllodactylus_interandinus | 0 | 3.16 | 1 | 23866.1 |
| lizards | Phyllodactylus_johnwrighti | Phyllodactylus_johnwrighti | 1 | 3.31 | 1 | 983.14 |
| lizards | Phyllodactylus_kofordi | Phyllodactylus_kofordi | 0 | 2.57 | 1 | 43204.7 |
| lizards | Phyllodactylus_lanei | Phyllodactylus_lanei | 0 | 13.8 | 2 | 124168.38 |
| lizards | Phyllodactylus_leei | Phyllodactylus_leei | 0 | 2.45 | 1.5 | 447.41 |
| lizards | Phyllodactylus_leoni | Phyllodactylus_leoni | 1 | 4.37 | 1.5 | 320.57 |
| lizards | Phyllodactylus_lepidopygus | Phyllodactylus_lepidopygus | 1 | 4.37 | 1 | 44015.77 |
| lizards | Phyllodactylus_microphyllus | Phyllodactylus_microphyllus | 0 | 5.13 | 1 | 56050.39 |
| lizards | Phyllodactylus_muralis | Phyllodactylus_muralis | 0 | 6.92 | 1.8 | 9377.79 |
| lizards | Phyllodactylus_nocticolus | Phyllodactylus_nocticolus | 0 | 6.61 | 2 | 92187 |
| lizards | Phyllodactylus_palmeus | Phyllodactylus_palmeus | 0 | 12.88 | 2 | 246.88 |

| | | | | | | |
|---|---|---|---|---|---|---|
| lizards | Phyllodactylus_pulcher | Phyllodactylus_pulcher | 1 | 7.94 | 1 | 179.1 |
| lizards | Phyllodactylus_reissii | Phyllodactylus_reissii | 0 | 13.18 | 1.85 | 228951.32 |
| lizards | Phyllodactylus_thompsoni | Phyllodactylus_thompsoni | 1 | 2.57 | 1.25 | 898.53 |
| lizards | Phyllodactylus_tuberculosus | Phyllodactylus_tuberculosus | 0 | 25.7 | 1.86 | 340776.32 |
| lizards | Phyllodactylus_unctus | Phyllodactylus_unctus | 0 | 5.62 | 2 | 8776.55 |
| lizards | Phyllodactylus_wirshingi | Phyllodactylus_wirshingi | 0 | 7.59 | 2 | 40.44 |
| lizards | Phyllodactylus_xanti | Phyllodactylus_xanti | 0 | 11.48 | 2 | 126932.84 |
| lizards | Phyllopezus_maranjonensis | Phyllopezus_maranjonensis | 0 | 52.48 | 2 | 34376.79 |
| lizards | Phyllopezus_periosus | Phyllopezus_periosus | 0 | 51.29 | 2 | 278407.36 |
| lizards | Phyllopezus_pollicaris | Phyllopezus_pollicaris | 0 | 21.88 | 1.95 | 1857384.69 |
| lizards | Phyllurus_amnicola | Phyllurus_amnicola | 0 | 24.55 | 2 | 224.05 |
| lizards | Phyllurus_caudiannulatus | Phyllurus_caudiannulatus | 0 | 23.99 | 2 | 239.07 |
| lizards | Phyllurus_championae | Phyllurus_championae | 0 | 10.96 | 1.5 | 192.74 |
| lizards | Phyllurus_gulbaru | Phyllurus_gulbaru | 1 | 17.38 | 2 | 513.71 |
| lizards | Phyllurus_nepthys | Phyllurus_nepthys | 0 | 19.5 | 2 | 840.32 |
| lizards | Phyllurus_platurus | Phyllurus_platurus | 0 | 22.91 | 2 | 28511.11 |
| lizards | Phymaturus_aguanegra | Phymaturus_aguanegra | 0 | 46.77 | 1.5 | 10.06 |
| lizards | Phymaturus_antofagastensis | Phymaturus_antofagastensis | 0 | 40.74 | 2 | 1725.1 |
| lizards | Phymaturus_camilae | Phymaturus_camilae | 0 | 42.66 | 2 | 8558.51 |
| lizards | Phymaturus_castillensis | Phymaturus_castillensis | 0 | 33.11 | 1.5 | 50.66 |
| lizards | Phymaturus_darwini | Phymaturus_darwini | 1 | 48.98 | 2 | 370.1 |
| lizards | Phymaturus_delheyi | Phymaturus_delheyi | 0 | 33.88 | 2 | 15.47 |
| lizards | Phymaturus_nevadoi | Phymaturus_nevadoi | 0 | 32.36 | 2 | 4473.49 |
| lizards | Phymaturus_palluma | Phymaturus_palluma | 0 | 56.23 | 2.75 | 180821.43 |
| lizards | Phymaturus_patagonicus | Phymaturus_patagonicus | 0 | 53.7 | 2 | 67493.56 |
| lizards | Phymaturus_payuniae | Phymaturus_payuniae | 0 | 28.18 | 2 | 4022.97 |
| lizards | Phymaturus_punae | Phymaturus_punae | 0 | 63.1 | 1.5 | 840.2 |
| lizards | Phymaturus_roigorum | Phymaturus_roigorum | 0 | 51.29 | 2 | 5174.84 |
| lizards | Phymaturus_sinervoi | Phymaturus_sinervoi | 0 | 45.71 | 1.5 | 86.28 |
| lizards | Phymaturus_somuncurensis | Phymaturus_somuncurensis | 0 | 40.74 | 3 | 21525.14 |
| lizards | Phymaturus_tenebrosus | Phymaturus_tenebrosus | 1 | 56.23 | 2 | 2691.94 |
| lizards | Phymaturus_videlai | Phymaturus_videlai | 0 | 33.88 | 2 | 124.73 |
| lizards | Phymaturus_vociferator | Phymaturus_vociferator | 1 | 44.67 | 1.5 | 14135.8 |
| lizards | Phymaturus_williamsi | Phymaturus_williamsi | 0 | 48.98 | 1.9 | 570.91 |

| | | | | | | |
|---|---|---|---|---|---|---|
| lizards | Phymaturus_zapalensis | Phymaturus_zapalensis | 0 | 48.98 | 1.5 | 10818.32 |
| lizards | Physignathus_cocincinus | Physignathus_cocincinus | 1 | 575.44 | 10.36 | 1243408.5 |
| lizards | Pinoyscincus_coxi | Pinoyscincus_coxi | 0 | 15.49 | 1.67 | 134924.06 |
| lizards | Pinoyscincus_jagori | Pinoyscincus_jagori | 0 | 29.51 | 1.9 | 273098.44 |
| lizards | Pinoyscincus_mindanensis | Pinoyscincus_mindanensis | 0 | 5.25 | 2 | 29535.74 |
| lizards | Placosoma_glabellum | Placosoma_glabellum | 0 | 3.31 | 1.5 | 36443.93 |
| lizards | Platysaurus_broadleyi | Platysaurus_broadleyi | 0 | 15.85 | 2 | 5396.49 |
| lizards | Platysaurus_capensis | Platysaurus_capensis | 0 | 16.98 | 2 | 54943.57 |
| lizards | Platysaurus_guttatus | Platysaurus_guttatus | 0 | 41.69 | 2 | 19528.73 |
| lizards | Platysaurus_imperator | Platysaurus_imperator | 0 | 104.71 | 2 | 34203.09 |
| lizards | Platysaurus_intermedius | Platysaurus_intermedius | 0 | 67.61 | 2 | 405952.4 |
| lizards | Platysaurus_lebomboensis | Platysaurus_lebomboensis | 0 | 9.55 | 1.5 | 8411.92 |
| lizards | Platysaurus_maculatus | Platysaurus_maculatus | 0 | 13.18 | 2 | 93.23 |
| lizards | Platysaurus_minor | Platysaurus_minor | 0 | 8.71 | 2 | 19496.33 |
| lizards | Platysaurus_monotropis | Platysaurus_monotropis | 0 | 10.47 | 2 | 2219.34 |
| lizards | Platysaurus_ocellatus | Platysaurus_ocellatus | 0 | 21.38 | 2 | 23479.51 |
| lizards | Platysaurus_orientalis | Platysaurus_orientalis | 0 | 19.05 | 2 | 28276.45 |
| lizards | Platysaurus_pungweensis | Platysaurus_pungweensis | 0 | 19.05 | 2 | 38507.7 |
| lizards | Platysaurus_torquatus | Platysaurus_torquatus | 0 | 10 | 2 | 30309.74 |
| lizards | Plestiodon_anthracinus | Plestiodon_anthracinus | 0 | 6.76 | 8.5 | 1099280.69 |
| lizards | Plestiodon_brevirostris | Plestiodon_brevirostris | 0 | 7.08 | 3 | 88662.24 |
| lizards | Plestiodon_callicephalus | Plestiodon_callicephalus | 0 | 8.91 | 5 | 174367.88 |
| lizards | Plestiodon_capito | Plestiodon_capito | 0 | 10.47 | 6 | 1059338.26 |
| lizards | Plestiodon_chinensis | Plestiodon_chinensis | 0 | 54.95 | 16.4 | 1767825.96 |
| lizards | Plestiodon_copei | Plestiodon_copei | 0 | 8.91 | 3.62 | 9117.4 |
| lizards | Plestiodon_dugesii | Plestiodon_dugesii | 1 | 6.46 | 4 | 19119.85 |
| lizards | Plestiodon_egregius | Plestiodon_egregius | 0 | 5.13 | 4.8 | 251532.98 |
| lizards | Plestiodon_elegans | Plestiodon_elegans | 0 | 25.12 | 5.95 | 2780942.55 |
| lizards | Plestiodon_fasciatus | Plestiodon_fasciatus | 0 | 13.8 | 7.7 | 2540293.28 |
| lizards | Plestiodon_gilberti | Plestiodon_gilberti | 0 | 35.48 | 6.3 | 217185.41 |
| lizards | Plestiodon_inexpectatus | Plestiodon_inexpectatus | 0 | 14.79 | 7.5 | 862871.06 |
| lizards | Plestiodon_japonicus | Plestiodon_japonicus | 0 | 19.05 | 10.5 | 106243.36 |
| lizards | Plestiodon_kishinouyei | Plestiodon_kishinouyei | 1 | 123.03 | 3 | 796.55 |
| lizards | Plestiodon_lagunensis | Plestiodon_lagunensis | 0 | 4.17 | 5.4 | 10333.94 |

| | | | | | | |
|---|---|---|---|---|---|---|
| lizards | Plestiodon_laticeps | Plestiodon_laticeps | 0 | 67.61 | 11.58 | 1777565.65 |
| lizards | Plestiodon_latiscutatus | Plestiodon_latiscutatus | 0 | 19.05 | 7.95 | 2989.86 |
| lizards | Plestiodon_lynxe | Plestiodon_lynxe | 0 | 8.51 | 4.35 | 252363.33 |
| lizards | Plestiodon_multivirgatus | Plestiodon_multivirgatus | 0 | 8.91 | 8.65 | 609662.31 |
| lizards | Plestiodon_obsoletus | Plestiodon_obsoletus | 0 | 67.61 | 12.65 | 1800483.91 |
| lizards | Plestiodon_quadrilineatus | Plestiodon_quadrilineatus | 0 | 30.2 | 2 | 412691.73 |
| lizards | Plestiodon_reynoldsi | Plestiodon_reynoldsi | 1 | 1.86 | 2 | 8146.16 |
| lizards | Plestiodon_septentrionalis | Plestiodon_septentrionalis | 0 | 15.49 | 10.65 | 1851681.79 |
| lizards | Plestiodon_skiltonianus | Plestiodon_skiltonianus | 0 | 13.18 | 4.9 | 1164304.83 |
| lizards | Plestiodon_stimpsonii | Plestiodon_stimpsonii | 0 | 10.47 | 9 | 554.76 |
| lizards | Plestiodon_sumichrasti | Plestiodon_sumichrasti | 0 | 21.38 | 11.5 | 135378.52 |
| lizards | Plestiodon_tetragrammus | Plestiodon_tetragrammus | 0 | 8.91 | 8.5 | 584850.08 |
| lizards | Pletholax_gracilis | Pletholax_gracilis | 0 | 4.37 | 2 | 40485.45 |
| lizards | Plica_plica | Plica_plica | 0 | 169.82 | 3.08 | 6864445.87 |
| lizards | Plica_umbra | Plica_umbra | 0 | 35.48 | 2.02 | 5802933.83 |
| lizards | Podarcis_bocagei | Podarcis_bocagei | 0 | 7.94 | 3.4 | 59240.89 |
| lizards | Podarcis_carbonelli | Podarcis_carbonelli | 1 | 6.46 | 2.15 | 15441.88 |
| lizards | Podarcis_cretensis | Podarcis_cretensis | 1 | 8.51 | 3.5 | 6298.08 |
| lizards | Podarcis_erhardii | Podarcis_erhardii | 0 | 13.8 | 2.53 | 153937.83 |
| lizards | Podarcis_filfolensis | Podarcis_filfolensis | 0 | 14.79 | 2.5 | 338.33 |
| lizards | Podarcis_gaigeae | Podarcis_gaigeae | 1 | 22.91 | 2.47 | 227.05 |
| lizards | Podarcis_hispanicus | Podarcis_hispanicus | 0 | 9.33 | 2.85 | 62282.82 |
| lizards | Podarcis_lilfordi | Podarcis_lilfordi | 1 | 12.3 | 2.5 | 4346.5 |
| lizards | Podarcis_melisellensis | Podarcis_melisellensis | 0 | 9.33 | 4.25 | 31611.34 |
| lizards | Podarcis_milensis | Podarcis_milensis | 1 | 9.77 | 1.73 | 242.86 |
| lizards | Podarcis_muralis | Podarcis_muralis | 0 | 10.96 | 4.93 | 1837171.64 |
| lizards | Podarcis_peloponnesiacus | Podarcis_peloponnesiacus | 0 | 14.13 | 3.48 | 21531.4 |
| lizards | Podarcis_pityusensis | Podarcis_pityusensis | 0 | 12.59 | 2.65 | 660.63 |
| lizards | Podarcis_raffonei | Podarcis_raffoneae | 1 | 14.13 | 6 | 71.51 |
| lizards | Podarcis_siculus | Podarcis_siculus | 0 | 16.6 | 5.7 | 276524.86 |
| lizards | Podarcis_tauricus | Podarcis_tauricus | 0 | 17.38 | 4.79 | 543683.3 |
| lizards | Podarcis_tiliguerta | Podarcis_tiliguerta | 0 | 15.14 | 9 | 32891.8 |
| lizards | Podarcis_vaucheri | Podarcis_vaucheri | 0 | 5.01 | 3 | 390655.23 |
| lizards | Podarcis_waglerianus | Podarcis_waglerianus | 0 | 10.23 | 4 | 23913.3 |

| | | | | | | |
|---|---|---|---|---|---|---|
| lizards | Pogona_barbata | Pogona_barbata | 0 | 691.83 | 16.75 | 1641296.36 |
| lizards | Pogona_henrylawsoni | Pogona_henrylawsoni | 0 | 123.03 | 12 | 167036.48 |
| lizards | Pogona_microlepidota | Pogona_microlepidota | 0 | 208.93 | 18 | 38139.89 |
| lizards | Pogona_minor | Pogona_minor | 0 | 173.78 | 6.5 | 3027728.34 |
| lizards | Pogona_nullarbor | Pogona_nullarbor | 0 | 97.72 | 14 | 131690.64 |
| lizards | Pogona_vitticeps | Pogona_vitticeps | 0 | 575.44 | 20.75 | 2466684.34 |
| lizards | Polychrus_acutirostris | Polychrus_acutirostris | 0 | 104.71 | 13.25 | 6733865.57 |
| lizards | Polychrus_femoralis | Polychrus_femoralis | 0 | 36.31 | 12 | 56600.86 |
| lizards | Polychrus_gutturosus | Polychrus_gutturosus | 0 | 165.96 | 7 | 422163.6 |
| lizards | Polychrus_liogaster | Polychrus_liogaster | 0 | 109.65 | 10 | 963491.22 |
| lizards | Polychrus_marmoratus | Polychrus_marmoratus | 0 | 102.33 | 8.1 | 8528534.28 |
| lizards | Polychrus_peruvianus | Polychrus_peruvianus | 1 | 109.65 | 6.6 | 63098.51 |
| lizards | Potamites_ecpleopus | Potamites_ecpleopus | 0 | 15.14 | 2 | 2258840.43 |
| lizards | Potamites_juruazensis | Potamites_juruazensis | 0 | 4.37 | 2 | 47076.45 |
| lizards | Prasinohaema_flavipes | Prasinohaema_flavipes | 0 | 23.44 | 6 | 49779.93 |
| lizards | Prasinohaema_semoni | Prasinohaema_semoni | 0 | 8.13 | 2 | 66690.41 |
| lizards | Prasinohaema_virens | Prasinohaema_virens | 0 | 5.37 | 2 | 103574.88 |
| lizards | Pristidactylus_achalensis | Pristidactylus_achalensis | 0 | 56.23 | 3 | 1357.23 |
| lizards | Pristidactylus_scapulatus | Pristidactylus_scapulatus | 0 | 41.69 | 6 | 102259.23 |
| lizards | Pristidactylus_torquatus | Pristidactylus_torquatus | 1 | 41.69 | 5.5 | 111288.84 |
| lizards | Pristidactylus_volcanensis | Pristidactylus_volcanensis | 1 | 27.54 | 6.5 | 749.24 |
| lizards | Pristurus_abdelkuri | Pristurus_abdelkuri | 0 | 1.26 | 1 | 1987.75 |
| lizards | Pristurus_carteri | Pristurus_carteri | 0 | 12.3 | 1.33 | 217855.55 |
| lizards | Pristurus_celerrimus | Pristurus_celerrimus | 0 | 1.58 | 1 | 26382.49 |
| lizards | Pristurus_collaris | Pristurus_collaris | 0 | 5.25 | 1.5 | 77478.83 |
| lizards | Pristurus_crucifer | Pristurus_crucifer | 0 | 1.58 | 1.11 | 774631.52 |
| lizards | Pristurus_flavipunctatus | Pristurus_flavipunctatus | 0 | 1.58 | 1 | 2017062.79 |
| lizards | Pristurus_gallagheri | Pristurus_gallagheri | 0 | 1.58 | 1 | 6567.88 |
| lizards | Pristurus_guichardi | Pristurus_guichardi | 0 | 1.29 | 1 | 347.6 |
| lizards | Pristurus_insignis | Pristurus_insignis | 0 | 5.25 | 1 | 2940.83 |
| lizards | Pristurus_insignoides | Pristurus_insignoides | 0 | 3.63 | 1 | 66.73 |
| lizards | Pristurus_minimus | Pristurus_minimus | 0 | 0.68 | 1 | 702202.61 |
| lizards | Pristurus_obsti | Pristurus_obsti | 0 | 1.32 | 1 | 1033.7 |
| lizards | Pristurus_ornithocephalus | Pristurus_ornithocephalus | 0 | 5.25 | 1 | 55646.25 |

| | | | | | | |
|---|---|---|---|---|---|---|
| lizards | Pristurus_phillipsii | Pristurus_phillipsii | 0 | 0.68 | 1 | 80172.13 |
| lizards | Pristurus_popovi | Pristurus_popovi | 0 | 1.32 | 1 | 28560.03 |
| lizards | Pristurus_rupestris | Pristurus_rupestris | 0 | 0.81 | 1 | 884378.49 |
| lizards | Pristurus_saada | Pristurus_saada | 0 | 1.7 | 1.67 | 27459.21 |
| lizards | Pristurus_samhaensis | Pristurus_samhaensis | 0 | 1.41 | 1 | 49.46 |
| lizards | Pristurus_sokotranus | Pristurus_sokotranus | 0 | 1.58 | 1 | 3551.63 |
| lizards | Pristurus_somalicus | Pristurus_somalicus | 0 | 1.48 | 1 | 3640.31 |
| lizards | Proablepharus_reginae | Proablepharus_reginae | 0 | 1.2 | 2 | 2311047.71 |
| lizards | Proablepharus_tenuis | Proablepharus_tenuis | 0 | 0.55 | 1.5 | 1534087.38 |
| lizards | Proctoporus_bolivianus | Proctoporus_bolivianus | 0 | 6.17 | 2 | 11067.29 |
| lizards | Proctoporus_chasqui | Proctoporus_chasqui | 0 | 9.77 | 2 | 21044.16 |
| lizards | Proctoporus_guentheri | Proctoporus_guentheri | 0 | 2.19 | 1.5 | 188762.74 |
| lizards | Proctoporus_pachyurus | Proctoporus_pachyurus | 0 | 4.37 | 2 | 2292.65 |
| lizards | Proctoporus_sucullucu | Proctoporus_sucullucu | 0 | 3.89 | 2 | 30309.6 |
| lizards | Proctoporus_unsaacae | Proctoporus_unsaacae | 0 | 2.95 | 2 | 11651.6 |
| lizards | Proctoporus_xestus | Proctoporus_xestus | 0 | 4.37 | 1.5 | 350333.03 |
| lizards | Proscelotes_arnoldi | Proscelotes_arnoldi | 0 | 4.79 | 5 | 49591.76 |
| lizards | Proscelotes_eggeli | Proscelotes_eggeli | 0 | 5.75 | 3 | 5991.46 |
| lizards | Psammodromus_algirus | Psammodromus_algirus | 0 | 19.5 | 5.35 | 1118062.97 |
| lizards | Psammodromus_blanci | Psammodromus_blanci | 0 | 2.45 | 3 | 155249.47 |
| lizards | Psammodromus_hispanicus | Psammodromus_hispanicus | 0 | 4.17 | 3.25 | 149004.2 |
| lizards | Psammophilus_blanfordanus | Psammophilus_blanfordanus | 0 | 38.02 | 5 | 889843.53 |
| lizards | Psammophilus_dorsalis | Psammophilus_dorsalis | 0 | 95.5 | 8.5 | 369569.72 |
| lizards | Pseudemoia_baudini | Pseudemoia_baudini | 0 | 2.75 | 3.5 | 89633.06 |
| lizards | Pseudemoia_cryodroma | Pseudemoia_cryodroma | 1 | 4.17 | 3.05 | 22949.37 |
| lizards | Pseudemoia_entrecasteauxii | Pseudemoia_entrecasteauxii | 0 | 5.37 | 4.05 | 581993.39 |
| lizards | Pseudemoia_pagenstecheri | Pseudemoia_pagenstecheri | 0 | 6.46 | 6.55 | 250162.81 |
| lizards | Pseudemoia_spenceri | Pseudemoia_spenceri | 0 | 5.37 | 1.95 | 191785.55 |
| lizards | Pseudocalotes_brevipes | Pseudocalotes_brevipes | 0 | 15.14 | 3 | 216284.45 |
| lizards | Pseudocalotes_cybelidermus | Pseudocalotes_cybelidermus | 0 | 27.54 | 2.5 | 268.06 |
| lizards | Pseudocalotes_flavigula | Pseudocalotes_flavigula | 1 | 16.98 | 3 | 9.99 |
| lizards | Pseudocalotes_floweri | Pseudocalotes_floweri | 1 | 32.36 | 4 | 115266.66 |
| lizards | Pseudocalotes_guttalineatus | Pseudocalotes_guttalineatus | 0 | 28.84 | 1.5 | 256 |
| lizards | Pseudocalotes_kakhienensis | Pseudocalotes_kakhienensis | 0 | 70.79 | 5 | 123991.86 |

| | | | | | | |
|---|---|---|---|---|---|---|
| lizards | Pseudocalotes_larutensis | Pseudocalotes_larutensis | 1 | 15.14 | 3 | 9.99 |
| lizards | Pseudocalotes_tympanistriga | Pseudocalotes_tympanistriga | 0 | 17.38 | 2 | 5078.07 |
| lizards | Pseudoceramodactylus_khobarensis | Pseudoceramodactylus_khobarensis | 0 | 6.92 | 1.5 | 99923.85 |
| lizards | Pseudocordylus_langi | Pseudocordylus_langi | 0 | 33.11 | 2 | 30620.21 |
| lizards | Pseudocordylus_melanotus | Pseudocordylus_melanotus | 0 | 117.49 | 3.45 | 165377 |
| lizards | Pseudocordylus_microlepidotus | Pseudocordylus_microlepidotus | 0 | 162.18 | 4 | 216570.52 |
| lizards | Pseudocordylus_spinosus | Pseudocordylus_spinosus | 0 | 17.78 | 2.5 | 15084.26 |
| lizards | Pseudocordylus_transvaalensis | Pseudocordylus_transvaalensis | 0 | 117.49 | 4.5 | 25028.99 |
| lizards | Pseudogekko_brevipes | Pseudogekko_brevipes | 1 | 3.47 | 2 | 29759.21 |
| lizards | Pseudogekko_compresicorpus | Pseudogekko_compressicorpus | 0 | 5.13 | 2 | 86089.02 |
| lizards | Pseudogekko_smaragdinus | Pseudogekko_smaragdinus | 0 | 5.62 | 2 | 920.81 |
| lizards | Pseudogonatodes_guianensis | Pseudogonatodes_guianensis | 0 | 0.68 | 1 | 3849398.49 |
| lizards | Pseudogonatodes_lunulatus | Pseudogonatodes_lunulatus | 0 | 0.66 | 1 | 48421.53 |
| lizards | Pseudopus_apodus | Pseudopus_apodus | 0 | 588.84 | 8.65 | 1920903.7 |
| lizards | Pseudotrapelus_sinaitus | Pseudotrapelus_sinaitus | 0 | 44.67 | 4.45 | 255754.01 |
| lizards | Psilops_paeminosus | Psilophthalmus_paeminosus | 0 | 1.29 | 1.86 | 1706.72 |
| lizards | Psychosaura_agmosticha | Psychosaura_agmosticha | 0 | 19.05 | 3.47 | 24199.95 |
| lizards | Psychosaura_macrorhyncha | Psychosaura_macrorhyncha | 0 | 12.88 | 3.33 | 889124.18 |
| lizards | Ptenopus_carpi | Ptenopus_carpi | 0 | 4.79 | 2 | 38353.56 |
| lizards | Ptenopus_garrulus | Ptenopus_garrulus | 0 | 5.37 | 1.5 | 1314010.36 |
| lizards | Ptyctolaemus_gularis | Ptyctolaemus_gularis | 0 | 21.88 | 10 | 173499.41 |
| lizards | Ptyodactylus_guttatus | Ptyodactylus_guttatus | 0 | 21.38 | 2 | 525182.43 |
| lizards | Ptyodactylus_hasselquistii | Ptyodactylus_hasselquistii | 0 | 29.51 | 2 | 2220688.31 |
| lizards | Ptyodactylus_oudrii | Ptyodactylus_oudrii | 0 | 6.31 | 1.5 | 509271.98 |
| lizards | Ptyodactylus_puiseuxi | Ptyodactylus_puiseuxi | 0 | 18.62 | 1.9 | 145812.08 |
| lizards | Ptyodactylus_ragazzii | Ptyodactylus_ragazzii | 0 | 25.7 | 2 | 4056885.1 |
| lizards | Pygmaeascincus_sadlieri | Pygmaeascincus_sadlieri | 0 | 0.44 | 2 | 54.05 |
| lizards | Pygmaeascincus_timlowi | Pygmaeascincus_timlowi | 0 | 0.4 | 1.67 | 530697.43 |
| lizards | Pygomeles_trivittatus | Androngo_trivittatus | 0 | 14.13 | 3 | 18662.43 |
| lizards | Pygopus_lepidopodus | Pygopus_lepidopodus | 0 | 19.95 | 2 | 3927238.31 |
| lizards | Pygopus_nigriceps | Pygopus_nigriceps | 0 | 15.49 | 1.75 | 3851747 |
| lizards | Pygopus_schraderi | Pygopus_schraderi | 0 | 12.88 | 2 | 2696954.24 |
| lizards | Pygopus_steelescotti | Pygopus_steelescotti | 0 | 11.75 | 2 | 877195.4 |
| lizards | Quedenfeldtia_moerens | Quedenfeldtia_moerens | 0 | 1.58 | 1 | 60974.66 |

| | | | | | | |
|---|---|---|---|---|---|---|
| lizards | Quedenfeldtia_trachyblepharus | Quedenfeldtia_trachyblepharus | 0 | 2.45 | 1 | 9521.66 |
| lizards | Ramigekko_swartbergensis | Ramigekko_swartbergensis | 0 | 9.33 | 2 | 9427.1 |
| lizards | Rankinia_diemensis | Rankinia_diemensis | 0 | 19.5 | 5.4 | 361951.85 |
| lizards | Rhacodactylus_auriculatus | Rhacodactylus_auriculatus | 0 | 40.74 | 2 | 9897.77 |
| lizards | Rhacodactylus_leachianus | Rhacodactylus_leachianus | 0 | 478.63 | 2 | 12858.56 |
| lizards | Rhampholeon_acuminatus | Rhampholeon_acuminatus | 1 | 5.13 | 3 | 33.51 |
| lizards | Rhampholeon_boulengeri | Rhampholeon_boulengeri | 0 | 6.46 | 2 | 465606.98 |
| lizards | Rhampholeon_chapmanorum | Rhampholeon_chapmanorum | 1 | 3.98 | 8 | 10.58 |
| lizards | Rhampholeon_gorongosae | Rhampholeon_gorongosae | 1 | 9.12 | 11 | 112.55 |
| lizards | Rhampholeon_marshalli | Rhampholeon_marshalli | 1 | 10 | 14 | 7879.21 |
| lizards | Rhampholeon_maspictus | Rhampholeon_maspictus | 0 | 7.59 | 7 | 13.97 |
| lizards | Rhampholeon_moyeri | Rhampholeon_moyeri | 0 | 3.89 | 4.5 | 7517.58 |
| lizards | Rhampholeon_nchisiensis | Rhampholeon_nchisiensis | 0 | 7.94 | 8 | 18302.51 |
| lizards | Rhampholeon_nebulauctor | Rhampholeon_nebulauctor | 1 | 3.39 | 3 | 9.99 |
| lizards | Rhampholeon_platyceps | Rhampholeon_platyceps | 1 | 8.91 | 9.5 | 2018.29 |
| lizards | Rhampholeon_spectrum | Rhampholeon_spectrum | 0 | 6.17 | 2 | 1599030.12 |
| lizards | Rhampholeon_spinosus | Rhampholeon_spinosus | 1 | 3.8 | 3.5 | 3516.75 |
| lizards | Rhampholeon_temporalis | Rhampholeon_temporalis | 1 | 3.63 | 5 | 731.37 |
| lizards | Rhampholeon_tilburyi | Rhampholeon_tilburyi | 1 | 8.71 | 7 | 61.01 |
| lizards | Rhampholeon_uluguruensis | Rhampholeon_uluguruensis | 0 | 3.63 | 3.5 | 412.66 |
| lizards | Rhampholeon_viridis | Rhampholeon_viridis | 1 | 3.02 | 6 | 1838.77 |
| lizards | Rhineura_floridana | Rhineura_floridana | 0 | 53.7 | 2 | 63057.85 |
| lizards | Rhoptropella_ocellata | Rhoptropella_ocellata | 0 | 2.24 | 2 | 37666.96 |
| lizards | Rhoptropus_afer | Rhoptropus_afer | 0 | 3.63 | 2 | 148366.28 |
| lizards | Rhoptropus_barnardi | Rhoptropus_barnardi | 0 | 2.63 | 2 | 289977.89 |
| lizards | Rhoptropus_boultoni | Rhoptropus_boultoni | 0 | 8.32 | 2 | 248492.67 |
| lizards | Rhoptropus_bradfieldi | Rhoptropus_bradfieldi | 0 | 7.08 | 2 | 71747.54 |
| lizards | Rhynchoedura_angusta | Rhynchoedura_angusta | 0 | 2.95 | 2 | 173564.14 |
| lizards | Rhynchoedura_ormsbyi | Rhynchoedura_ormsbyi | 0 | 2.95 | 2 | 723260.83 |
| lizards | Rhynchoedura_ornata | Rhynchoedura_ornata | 0 | 3.63 | 1.99 | 2671543.63 |
| lizards | Rhynchoedura_sexapora | Rhynchoedura_sexapora | 0 | 2.24 | 2 | 229008.35 |
| lizards | Riama_cashcaensis | Riama_cashcaensis | 1 | 9.33 | 2 | 456.72 |
| lizards | Riama_columbiana | Riama_columbiana | 1 | 13.8 | 2 | 15553.12 |
| lizards | Riama_orcesi | Riama_orcesi | 1 | 5.75 | 2 | 1159.41 |

| | | | | | | |
|---|---|---|---|---|---|---|
| lizards | Riama_striata | Riama_striata | 0 | 7.08 | 1.5 | 61897.74 |
| lizards | Rieppeleon_brachyurus | Rieppeleon_brachyurus | 0 | 3.63 | 10 | 542075.66 |
| lizards | Rieppeleon_brevicaudatus | Rieppeleon_brevicaudatus | 0 | 10.72 | 5 | 178303.61 |
| lizards | Rieppeleon_kerstenii | Rieppeleon_kerstenii | 0 | 9.12 | 7.5 | 1280227.06 |
| lizards | Ristella_beddomii | Ristella_beddomii | 0 | 1.12 | 3 | 11463.76 |
| lizards | Ristella_rurkii | Ristella_rurkii | 0 | 1.86 | 2 | 10912.55 |
| lizards | Saara_hardwickii | Saara_hardwickii | 1 | 575.44 | 12.5 | 1111718.16 |
| lizards | Saara_loricata | Saara_loricata | 0 | 912.01 | 14 | 139510.09 |
| lizards | Saiphos_equalis | Saiphos_equalis | 0 | 4.47 | 2.9 | 245681.42 |
| lizards | Salea_anamallayana | Salea_anamallayana | 0 | 45.71 | 4 | 6379.91 |
| lizards | Salea_horsfieldii | Salea_horsfieldii | 0 | 28.18 | 3.5 | 4619.34 |
| lizards | Saltuarius_cornutus | Saltuarius_cornutus | 0 | 54.95 | 2 | 19859.74 |
| lizards | Saltuarius_kateae | Saltuarius_kateae | 0 | 22.91 | 2 | 257.69 |
| lizards | Saltuarius_moritzi | Saltuarius_moritzi | 0 | 22.39 | 2 | 30847.59 |
| lizards | Saltuarius_salebrosus | Saltuarius_salebrosus | 0 | 42.66 | 2 | 89491.31 |
| lizards | Saltuarius_swaini | Saltuarius_swaini | 0 | 37.15 | 2 | 11797.3 |
| lizards | Saltuarius_wyberba | Saltuarius_wyberba | 0 | 22.39 | 2 | 6886.31 |
| lizards | Salvator_merianae | Salvator_merianae | 0 | 4466.84 | 36.5 | 8025045.46 |
| lizards | Salvator_rufescens | Salvator_rufescens | 0 | 8317.64 | 21.4 | 1371652.75 |
| lizards | Saproscincus_basiliscus | Saproscincus_basiliscus | 0 | 2.29 | 2 | 22520.23 |
| lizards | Saproscincus_challengeri | Saproscincus_challengeri | 0 | 3.89 | 3.5 | 19134.33 |
| lizards | Saproscincus_czechurai | Saproscincus_czechurai | 0 | 1.12 | 2 | 13189.24 |
| lizards | Saproscincus_mustelinus | Saproscincus_mustelinus | 0 | 5.13 | 5.05 | 212712.42 |
| lizards | Saproscincus_oriarus | Saproscincus_oriarus | 0 | 1.41 | 2 | 4899.02 |
| lizards | Saproscincus_rosei | Saproscincus_rosei | 0 | 5.37 | 2 | 65680.53 |
| lizards | Saproscincus_spectabilis | Saproscincus_spectabilis | 0 | 4.17 | 2 | 66641.8 |
| lizards | Saproscincus_tetradactylus | Saproscincus_tetradactylus | 0 | 0.6 | 2 | 14781.59 |
| lizards | Saurodactylus_fasciatus | Saurodactylus_fasciatus | 1 | 1.35 | 2 | 42698.67 |
| lizards | Saurodactylus_mauritanicus | Saurodactylus_mauritanicus | 0 | 0.98 | 1 | 235990.43 |
| lizards | Sauromalus_ater | Sauromalus_ater | 0 | 524.81 | 7.75 | 387687.4 |
| lizards | Sauromalus_hispidus | Sauromalus_hispidus | 1 | 1380.38 | 21.75 | 991.44 |
| lizards | Sauromalus_varius | Sauromalus_varius | 1 | 1659.59 | 20.85 | 41.2 |
| lizards | Scelarcis_perspicillata | Scelarcis_perspicillata | 0 | 7.59 | 3 | 133305.08 |
| lizards | Sceloporus_acanthinus | Sceloporus_acanthinus | 0 | 30.9 | 6 | 13458.61 |

| | | | | | | |
|---|---|---|---|---|---|---|
| lizards | Sceloporus_adleri | Sceloporus_adleri | 0 | 16.6 | 5.19 | 4633.16 |
| lizards | Sceloporus_aeneus | Sceloporus_aeneus | 0 | 9.12 | 6.66 | 52636.26 |
| lizards | Sceloporus_angustus | Sceloporus_angustus | 0 | 32.36 | 5.5 | 45.05 |
| lizards | Sceloporus_arenicolus | Sceloporus_arenicolus | 1 | 12.02 | 5 | 5622.15 |
| lizards | Sceloporus_bicanthalis | Sceloporus_bicanthalis | 0 | 8.71 | 6.69 | 24407.25 |
| lizards | Sceloporus_bulleri | Sceloporus_bulleri | 0 | 46.77 | 7 | 29304.51 |
| lizards | Sceloporus_carinatus | Sceloporus_carinatus | 0 | 17.38 | 10 | 18402.32 |
| lizards | Sceloporus_chaneyi | Sceloporus_chaneyi | 1 | 7.08 | 5 | 3018.86 |
| lizards | Sceloporus_chrysostictus | Sceloporus_chrysostictus | 0 | 8.71 | 2.45 | 156256.05 |
| lizards | Sceloporus_clarkii | Sceloporus_clarkii | 0 | 83.18 | 13.8 | 457350.44 |
| lizards | Sceloporus_couchii | Sceloporus_couchii | 0 | 9.55 | 4 | 48041.72 |
| lizards | Sceloporus_cozumelae | Sceloporus_cozumelae | 0 | 8.13 | 1.8 | 20751.44 |
| lizards | Sceloporus_cryptus | Sceloporus_cryptus | 0 | 15.49 | 9 | 1680.9 |
| lizards | Sceloporus_cyanostictus | Sceloporus_cyanostictus | 1 | 17.78 | 7.07 | 3325.92 |
| lizards | Sceloporus_dugesii | Sceloporus_dugesii | 0 | 22.39 | 4.4 | 147176.86 |
| lizards | Sceloporus_edwardtaylori | Sceloporus_edwardtaylori | 0 | 38.02 | 8.5 | 18437.35 |
| lizards | Sceloporus_exsul | Sceloporus_exsul | 1 | 10 | 10 | 323.54 |
| lizards | Sceloporus_formosus | Sceloporus_formosus | 0 | 22.39 | 7.12 | 97765.32 |
| lizards | Sceloporus_gadoviae | Sceloporus_gadoviae | 0 | 15.14 | 3.75 | 22582.32 |
| lizards | Sceloporus_goldmani | Sceloporus_goldmani | 1 | 8.13 | 7.5 | 9208.61 |
| lizards | Sceloporus_graciosus | Sceloporus_graciosus | 0 | 22.91 | 5.4 | 1358846.8 |
| lizards | Sceloporus_grammicus | Sceloporus_grammicus | 0 | 17.78 | 4.76 | 981930.66 |
| lizards | Sceloporus_grandaevus | Sceloporus_grandaevus | 0 | 17.78 | 6.5 | 142.91 |
| lizards | Sceloporus_horridus | Sceloporus_horridus | 0 | 48.98 | 11.9 | 330779.99 |
| lizards | Sceloporus_hunsakeri | Sceloporus_hunsakeri | 0 | 21.88 | 7.5 | 9946.62 |
| lizards | Sceloporus_insignis | Sceloporus_insignis | 0 | 30.9 | 9 | 8956.53 |
| lizards | Sceloporus_internasalis | Sceloporus_internasalis | 0 | 31.62 | 6 | 34714.51 |
| lizards | Sceloporus_jalapae | Sceloporus_jalapae | 0 | 8.71 | 5.62 | 34104.5 |
| lizards | Sceloporus_jarrovii | Sceloporus_jarrovii | 0 | 37.15 | 6.7 | 532172.57 |
| lizards | Sceloporus_licki | Sceloporus_licki | 0 | 26.92 | 6 | 4628.56 |
| lizards | Sceloporus_lineatulus | Sceloporus_lineatulus | 0 | 45.71 | 9.4 | 27.73 |
| lizards | Sceloporus_macdougalli | Sceloporus_macdougalli | 0 | 27.54 | 3.09 | 8909.22 |
| lizards | Sceloporus_magister | Sceloporus_magister | 0 | 81.28 | 9.3 | 1395390.78 |
| lizards | Sceloporus_malachiticus | Sceloporus_malachiticus | 0 | 30.2 | 5.25 | 50271.06 |

| | | | | | | |
|---|---|---|---|---|---|---|
| lizards | Sceloporus_megalepidurus | Sceloporus_megalepidurus | 1 | 6.31 | 2.82 | 10818.61 |
| lizards | Sceloporus_melanorhinus | Sceloporus_melanorhinus | 0 | 36.31 | 7.7 | 164283.9 |
| lizards | Sceloporus_merriami | Sceloporus_merriami | 0 | 10.47 | 4.11 | 208660.18 |
| lizards | Sceloporus_minor | Sceloporus_minor | 0 | 21.88 | 6.58 | 133511.21 |
| lizards | Sceloporus_mucronatus | Sceloporus_mucronatus | 0 | 44.67 | 4.69 | 98787.59 |
| lizards | Sceloporus_nelsoni | Sceloporus_nelsoni | 0 | 11.22 | 6.25 | 117808.96 |
| lizards | Sceloporus_occidentalis | Sceloporus_occidentalis | 0 | 30.9 | 10.1 | 1010683.23 |
| lizards | Sceloporus_ochoterenae | Sceloporus_ochoterenae | 0 | 8.71 | 6.25 | 14081.84 |
| lizards | Sceloporus_olivaceus | Sceloporus_olivaceus | 0 | 56.23 | 17.9 | 646821 |
| lizards | Sceloporus_orcutti | Sceloporus_orcutti | 0 | 47.86 | 10.1 | 111356.74 |
| lizards | Sceloporus_ornatus | Sceloporus_ornatus | 0 | 30.2 | 3.3 | 56330.39 |
| lizards | Sceloporus_palaciosi | Sceloporus_palaciosi | 0 | 14.45 | 4 | 985.53 |
| lizards | Sceloporus_parvus | Sceloporus_parvus | 0 | 5.25 | 3.4 | 163557.95 |
| lizards | Sceloporus_poinsettii | Sceloporus_poinsettii | 0 | 74.13 | 10.95 | 842198.09 |
| lizards | Sceloporus_pyrocephalus | Sceloporus_pyrocephalus | 0 | 14.45 | 5.23 | 62214.86 |
| lizards | Sceloporus_samcolemani | Sceloporus_samcolemani | 0 | 8.32 | 8.8 | 25507.14 |
| lizards | Sceloporus_scalaris | Sceloporus_scalaris | 0 | 16.22 | 8.19 | 227283.4 |
| lizards | Sceloporus_serrifer | Sceloporus_serrifer | 0 | 43.65 | 12.9 | 225665.13 |
| lizards | Sceloporus_siniferus | Sceloporus_siniferus | 0 | 12.59 | 5.45 | 156926.04 |
| lizards | Sceloporus_slevini | Sceloporus_slevini | 0 | 12.02 | 8.5 | 151398.7 |
| lizards | Sceloporus_smaragdinus | Sceloporus_smaragdinus | 0 | 20.89 | 4.2 | 24225.81 |
| lizards | Sceloporus_smithi | Sceloporus_smithi | 0 | 12.59 | 4 | 20200.24 |
| lizards | Sceloporus_spinosus | Sceloporus_spinosus | 0 | 58.88 | 13.85 | 304248 |
| lizards | Sceloporus_squamosus | Sceloporus_squamosus | 0 | 7.76 | 4.5 | 95046.17 |
| lizards | Sceloporus_sugillatus | Sceloporus_sugillatus | 0 | 21.88 | 7.07 | 1224.52 |
| lizards | Sceloporus_taeniocnemis | Sceloporus_taeniocnemis | 0 | 18.62 | 6 | 19174.85 |
| lizards | Sceloporus_teapensis | Sceloporus_teapensis | 0 | 12.02 | 2.33 | 179968.89 |
| lizards | Sceloporus_torquatus | Sceloporus_torquatus | 0 | 79.43 | 8.05 | 321352.38 |
| lizards | Sceloporus_undulatus | Sceloporus_undulatus | 0 | 24.55 | 9.13 | 4098325.29 |
| lizards | Sceloporus_utiformis | Sceloporus_utiformis | 0 | 19.95 | 7.02 | 162522.64 |
| lizards | Sceloporus_variabilis | Sceloporus_variabilis | 0 | 15.85 | 3.9 | 953656.3 |
| lizards | Sceloporus_virgatus | Sceloporus_virgatus | 0 | 12.59 | 9.83 | 78813.04 |
| lizards | Sceloporus_woodi | Sceloporus_woodi | 0 | 10 | 4.81 | 13827.79 |
| lizards | Sceloporus_zosteromus | Sceloporus_zosteromus | 0 | 64.57 | 6 | 126799.42 |

| | | | | | | |
|---|---|---|---|---|---|---|
| lizards | Scelotes_bidigittatus | Scelotes_bidigittatus | 0 | 3.47 | 1.5 | 48632.48 |
| lizards | Scelotes_bipes | Scelotes_bipes | 0 | 4.27 | 1.5 | 60186 |
| lizards | Scelotes_caffer | Scelotes_caffer | 0 | 1.23 | 1.5 | 26702.14 |
| lizards | Scelotes_duttoni | Scelotes_duttoni | 0 | 1.82 | 2.5 | 26.78 |
| lizards | Scelotes_fitzsimonsi | Scelotes_fitzsimonsi | 0 | 0.98 | 2 | 12593.08 |
| lizards | Scelotes_gronovii | Scelotes_gronovii | 0 | 2.24 | 1.5 | 25154.96 |
| lizards | Scelotes_limpopoensis | Scelotes_limpopoensis | 0 | 3.63 | 2 | 29130.27 |
| lizards | Scelotes_mirus | Scelotes_mirus | 0 | 12.88 | 4 | 100610.75 |
| lizards | Scelotes_mossambicus | Scelotes_mossambicus | 0 | 2.69 | 2.5 | 99204.91 |
| lizards | Scelotes_uluguruensis | Scelotes_uluguruensis | 1 | 3.98 | 4 | 14340.6 |
| lizards | Scincella_boettgeri | Scincella_boettgeri | 0 | 3.31 | 7.5 | 798.67 |
| lizards | Scincella_cherriei | Scincella_cherriei | 0 | 6.17 | 2.5 | 482005.37 |
| lizards | Scincella_gemmingeri | Scincella_gemmingeri | 0 | 5.37 | 3.5 | 146449.24 |
| lizards | Scincella_huanrenensis | Scincella_huanrenensis | 1 | 4.57 | 4.56 | 15069.21 |
| lizards | Scincella_lateralis | Scincella_lateralis | 0 | 3.89 | 3.08 | 2107754.7 |
| lizards | Scincella_melanosticta | Scincella_melanosticta | 0 | 5.37 | 3 | 483325.97 |
| lizards | Scincella_modesta | Scincella_modesta | 0 | 5.75 | 7.55 | 1279283.81 |
| lizards | Scincella_monticola | Scincella_monticola | 0 | 3.89 | 2.5 | 908302.35 |
| lizards | Scincella_ochracea | Scincella_ochracea | 0 | 2.45 | 4 | 39206.7 |
| lizards | Scincella_reevesii | Scincella_reevesii | 0 | 4.17 | 4.8 | 2762379.81 |
| lizards | Scincella_silvicola | Scincella_silvicola | 0 | 3.72 | 5 | 18296.27 |
| lizards | Scincella_tsinlingensis | Scincella_tsinlingensis | 0 | 6.76 | 3 | 282019.32 |
| lizards | Scincella_vandenburghi | Scincella_vandenburghi | 0 | 2.88 | 7 | 57898.86 |
| lizards | Scincella_victoriana | Scincella_victoriana | 0 | 9.12 | 2 | 5227.43 |
| lizards | Scincus_albifasciatus | Scincus_albifasciatus | 0 | 51.29 | 2.75 | 2073161.13 |
| lizards | Scincus_mitranus | Scincus_mitranus | 0 | 54.95 | 5 | 1455446.25 |
| lizards | Scincus_scincus | Scincus_scincus | 0 | 74.13 | 5 | 10611472.07 |
| lizards | Sepsina_alberti | Sepsina_alberti | 0 | 1.23 | 5 | 41674.02 |
| lizards | Sepsina_tetradactyla | Sepsina_tetradactyla | 0 | 4.47 | 3 | 440539.85 |
| lizards | Shinisaurus_crocodilurus | Shinisaurus_crocodilurus | 1 | 83.18 | 6.3 | 90732.08 |
| lizards | Sigaloseps_deplanchei | Sigaloseps_deplanchei | 0 | 1.74 | 2.17 | 4672.6 |
| lizards | Sigaloseps_ruficauda | Sigaloseps_ruficauda | 1 | 4.17 | 3 | 193.79 |
| lizards | Silvascincus_murrayi | Silvascincus_murrayi | 0 | 27.54 | 3.6 | 75202.7 |
| lizards | Silvascincus_tryoni | Silvascincus_tryoni | 0 | 24.55 | 5 | 629.05 |

| | | | | | | |
|---|---|---|---|---|---|---|
| lizards | Simiscincus_aurantiacus | Simiscincus_aurantiacus | 1 | 12.88 | 3 | 1293.42 |
| lizards | Sitana_fusca | Sitana_fusca | 1 | 3.39 | 7.2 | 19.98 |
| lizards | Sitana_ponticeriana | Sitana_ponticeriana | 0 | 17.38 | 14.3 | 173089.06 |
| lizards | Sitana_sivalensis | Sitana_sivalensis | 0 | 2.69 | 6.86 | 14229.84 |
| lizards | Smaug_barbertonensis | Smaug_barbertonensis | 0 | 95.5 | 5 | 10140.02 |
| lizards | Smaug_breyeri | Smaug_breyeri | 0 | 102.33 | 4 | 17344.44 |
| lizards | Smaug_depressus | Smaug_depressus | 0 | 53.7 | 4 | 7008.35 |
| lizards | Smaug_giganteus | Smaug_giganteus | 1 | 457.09 | 2.1 | 119790.5 |
| lizards | Smaug_mossambicus | Smaug_mossambicus | 0 | 61.66 | 5.5 | 21926.19 |
| lizards | Smaug_regius | Smaug_regius | 0 | 102.33 | 3.5 | 26058.51 |
| lizards | Smaug_vandami | Smaug_vandami | 0 | 102.33 | 4 | 43320.67 |
| lizards | Smaug_warreni | Smaug_warreni | 0 | 128.82 | 5 | 6984.53 |
| lizards | Sphaerodactylus_altavelensis | Sphaerodactylus_altavelensis | 0 | 0.62 | 1 | 30077.36 |
| lizards | Sphaerodactylus_argivus | Sphaerodactylus_argivus | 0 | 0.62 | 1 | 265.85 |
| lizards | Sphaerodactylus_argus | Sphaerodactylus_argus | 0 | 0.89 | 1 | 55662.42 |
| lizards | Sphaerodactylus_armasi | Sphaerodactylus_armasi | 1 | 0.81 | 1 | 1708.32 |
| lizards | Sphaerodactylus_caicosensis | Sphaerodactylus_caicosensis | 0 | 0.81 | 1 | 342.6 |
| lizards | Sphaerodactylus_callocricus | Sphaerodactylus_callocricus | 1 | 0.55 | 1 | 2180.61 |
| lizards | Sphaerodactylus_cinereus | Sphaerodactylus_cinereus | 1 | 1.26 | 1 | 5532.65 |
| lizards | Sphaerodactylus_clenchi | Sphaerodactylus_clenchi | 0 | 0.98 | 1 | 1846.56 |
| lizards | Sphaerodactylus_copei | Sphaerodactylus_copei | 0 | 1.7 | 1 | 10732.24 |
| lizards | Sphaerodactylus_corticola | Sphaerodactylus_corticola | 0 | 1.48 | 1 | 1120.04 |
| lizards | Sphaerodactylus_darlingtoni | Sphaerodactylus_darlingtoni | 0 | 0.62 | 1 | 24299.55 |
| lizards | Sphaerodactylus_difficilis | Sphaerodactylus_difficilis | 0 | 0.98 | 1 | 49802.78 |
| lizards | Sphaerodactylus_elegans | Sphaerodactylus_elegans | 0 | 1.55 | 1 | 127682.27 |
| lizards | Sphaerodactylus_epiurus | Sphaerodactylus_epiurus | 1 | 0.39 | 1 | 19.98 |
| lizards | Sphaerodactylus_fantasticus | Sphaerodactylus_fantasticus | 0 | 0.62 | 1 | 1509.48 |
| lizards | Sphaerodactylus_glaucus | Sphaerodactylus_glaucus | 0 | 0.74 | 1 | 322192.51 |
| lizards | Sphaerodactylus_goniorhynchus | Sphaerodactylus_goniorhynchus | 0 | 0.81 | 1 | 8634.72 |
| lizards | Sphaerodactylus_homolepis | Sphaerodactylus_homolepis | 0 | 0.89 | 1 | 15779.13 |
| lizards | Sphaerodactylus_inaguae | Sphaerodactylus_inaguae | 1 | 0.62 | 1 | 436.1 |
| lizards | Sphaerodactylus_kirbyi | Sphaerodactylus_kirbyi | 1 | 0.62 | 1 | 78.83 |
| lizards | Sphaerodactylus_klauberi | Sphaerodactylus_klauberi | 0 | 1.26 | 1 | 4438.21 |
| lizards | Sphaerodactylus_leucaster | Sphaerodactylus_leucaster | 0 | 0.74 | 1 | 2105.33 |

| | | | | | | |
|---|---|---|---|---|---|---|
| lizards | Sphaerodactylus_levinsi | Sphaerodactylus_levinsi | 0 | 0.66 | 1 | 1.3 |
| lizards | Sphaerodactylus_macrolepis | Sphaerodactylus_macrolepis | 0 | 0.52 | 1 | 9157.78 |
| lizards | Sphaerodactylus_millepunctatus | Sphaerodactylus_millepunctatus | 0 | 1.1 | 1.5 | 251986.7 |
| lizards | Sphaerodactylus_molei | Sphaerodactylus_molei | 0 | 0.55 | 2 | 501975.05 |
| lizards | Sphaerodactylus_nicholsi | Sphaerodactylus_nicholsi | 0 | 0.39 | 1 | 5917.33 |
| lizards | Sphaerodactylus_nigropunctatus | Sphaerodactylus_nigropunctatus | 0 | 1.58 | 1 | 44934.38 |
| lizards | Sphaerodactylus_notatus | Sphaerodactylus_notatus | 0 | 0.98 | 1 | 127588.64 |
| lizards | Sphaerodactylus_oliveri | Sphaerodactylus_oliveri | 1 | 0.98 | 1 | 2033.26 |
| lizards | Sphaerodactylus_oxyrhinus | Sphaerodactylus_oxyrhinus | 1 | 0.98 | 1 | 975.54 |
| lizards | Sphaerodactylus_pacificus | Sphaerodactylus_pacificus | 0 | 2.88 | 1.5 | 24.78 |
| lizards | Sphaerodactylus_parkeri | Sphaerodactylus_parkeri | 1 | 1.07 | 1 | 664.81 |
| lizards | Sphaerodactylus_parvus | Sphaerodactylus_parvus | 0 | 0.35 | 1 | 195.94 |
| lizards | Sphaerodactylus_pimienta | Sphaerodactylus_pimienta | 1 | 1.35 | 1 | 1466.37 |
| lizards | Sphaerodactylus_richardsonii | Sphaerodactylus_richardsonii | 1 | 1.58 | 1 | 569.39 |
| lizards | Sphaerodactylus_roosevelti | Sphaerodactylus_roosevelti | 0 | 1.48 | 1 | 168.56 |
| lizards | Sphaerodactylus_ruibali | Sphaerodactylus_ruibali | 1 | 0.81 | 1 | 190.06 |
| lizards | Sphaerodactylus_sabanus | Sphaerodactylus_sabanus | 0 | 0.68 | 1 | 310.32 |
| lizards | Sphaerodactylus_savagei | Sphaerodactylus_savagei | 0 | 0.89 | 1 | 2896.71 |
| lizards | Sphaerodactylus_scaber | Sphaerodactylus_scaber | 0 | 0.98 | 1 | 13689 |
| lizards | Sphaerodactylus_siboney | Sphaerodactylus_siboney | 0 | 0.83 | 1 | 29.68 |
| lizards | Sphaerodactylus_sputator | Sphaerodactylus_sputator | 0 | 1.58 | 1 | 324.72 |
| lizards | Sphaerodactylus_streptophorus | Sphaerodactylus_streptophorus | 1 | 0.44 | 1 | 7151.28 |
| lizards | Sphaerodactylus_torrei | Sphaerodactylus_torrei | 1 | 1.48 | 1 | 791 |
| lizards | Sphaerodactylus_townsendi | Sphaerodactylus_townsendi | 0 | 0.68 | 1 | 1539.52 |
| lizards | Sphaerodactylus_vincenti | Sphaerodactylus_vincenti | 0 | 1.58 | 1 | 2876.61 |
| lizards | Sphaerodactylus_zygaena | Sphaerodactylus_zygaena | 1 | 0.81 | 1 | 587.49 |
| lizards | Sphenomorphus_aignanus | Sphenomorphus_aignanus | 0 | 10.47 | 1 | 29903.33 |
| lizards | Sphenomorphus_bignelli | Sphenomorphus_bignelli | 0 | 0.72 | 1 | 12970.58 |
| lizards | Sphenomorphus_brunneus | Sphenomorphus_brunneus | 0 | 13.8 | 5.1 | 7957.21 |
| lizards | Sphenomorphus_buenloicus | Sphenomorphus_buenloicus | 0 | 3.47 | 2.5 | 14831.68 |
| lizards | Sphenomorphus_concinnatus | Sphenomorphus_concinnatus | 0 | 6.76 | 2 | 34463.95 |
| lizards | Sphenomorphus_cranei | Sphenomorphus_cranei | 0 | 10 | 2.53 | 28487.91 |
| lizards | Sphenomorphus_cyanolaemus | Sphenomorphus_cyanolaemus | 0 | 4.17 | 2.15 | 232642.68 |
| lizards | Sphenomorphus_darlingtoni | Sphenomorphus_darlingtoni | 0 | 5.13 | 2 | 37540.51 |

| | | | | | | |
|---|---|---|---|---|---|---|
| lizards | Sphenomorphus_dussumieri | Sphenomorphus_dussumieri | 0 | 5.13 | 3.5 | 121173.56 |
| lizards | Sphenomorphus_fasciatus | Sphenomorphus_fasciatus | 0 | 39.81 | 5.35 | 118957.09 |
| lizards | Sphenomorphus_forbesi | Sphenomorphus_forbesi | 0 | 3.89 | 1.6 | 302.13 |
| lizards | Sphenomorphus_fragilis | Sphenomorphus_fragilis | 0 | 2.95 | 2.7 | 5816.94 |
| lizards | Sphenomorphus_fuscolineatus | Sphenomorphus_fuscolineatus | 0 | 3.89 | 2 | 133368.08 |
| lizards | Sphenomorphus_incognitus | Sphenomorphus_incognitus | 0 | 28.84 | 4.6 | 1013128.3 |
| lizards | Sphenomorphus_indicus | Sphenomorphus_indicus | 0 | 25.12 | 7.27 | 5514148.77 |
| lizards | Sphenomorphus_jobiensis | Sphenomorphus_jobiensis | 0 | 34.67 | 3 | 322943.89 |
| lizards | Sphenomorphus_leptofasciatus | Sphenomorphus_leptofasciatus | 0 | 13.18 | 3.25 | 29425.48 |
| lizards | Sphenomorphus_longicaudatus | Sphenomorphus_longicaudatus | 0 | 17.78 | 2.6 | 26764.7 |
| lizards | Sphenomorphus_louisiadensis | Sphenomorphus_louisiadensis | 0 | 2.14 | 1 | 294.49 |
| lizards | Sphenomorphus_maculatus | Sphenomorphus_maculatus | 0 | 6.76 | 3.85 | 2700117.52 |
| lizards | Sphenomorphus_maindroni | Sphenomorphus_maindroni | 0 | 9.33 | 2 | 72003.77 |
| lizards | Sphenomorphus_melanopogon | Sphenomorphus_melanopogon | 0 | 15.49 | 2 | 32867.41 |
| lizards | Sphenomorphus_minutus | Sphenomorphus_minutus | 0 | 1.2 | 1 | 78131.17 |
| lizards | Sphenomorphus_multisquamatus | Sphenomorphus_multisquamatus | 0 | 7.76 | 3 | 174704.78 |
| lizards | Sphenomorphus_nigriventris | Sphenomorphus_nigriventris | 0 | 15.49 | 3.7 | 42917.4 |
| lizards | Sphenomorphus_nigrolineatus | Sphenomorphus_nigrolineatus | 0 | 8.51 | 3.8 | 13453.9 |
| lizards | Sphenomorphus_oligolepis | Sphenomorphus_oligolepis | 0 | 3.09 | 2 | 35212.03 |
| lizards | Sphenomorphus_orientalis | Sphenomorphus_orientale | 0 | 2.95 | 2 | 814.41 |
| lizards | Sphenomorphus_papuae | Sphenomorphus_papuae | 0 | 10.47 | 3.37 | 12742.54 |
| lizards | Sphenomorphus_praesignis | Sphenomorphus_praesignis | 0 | 39.81 | 7.25 | 14193.99 |
| lizards | Sphenomorphus_pratti | Sphenomorphus_pratti | 0 | 18.62 | 3 | 252218.77 |
| lizards | Sphenomorphus_sabanus | Sphenomorphus_sabanus | 0 | 4.17 | 1.9 | 36610.25 |
| lizards | Sphenomorphus_schlegeli | Sphenomorphus_schlegeli | 0 | 0.6 | 6 | 333.65 |
| lizards | Sphenomorphus_schultzei | Sphenomorphus_schultzei | 0 | 1.86 | 1 | 102107.23 |
| lizards | Sphenomorphus_scotophilus | Sphenomorphus_scotophilus | 0 | 6.76 | 1.8 | 17431.78 |
| lizards | Sphenomorphus_scutatus | Sphenomorphus_scutatus | 0 | 1.51 | 2 | 415.88 |
| lizards | Sphenomorphus_simus | Sphenomorphus_simus | 0 | 3.72 | 2 | 693689.12 |
| lizards | Sphenomorphus_solomonis | Sphenomorphus_solomonis | 0 | 10 | 2.75 | 203096.63 |
| lizards | Sphenomorphus_striolatus | Sphenomorphus_striolatus | 0 | 3.09 | 1 | 25730.07 |
| lizards | Sphenomorphus_taiwanensis | Sphenomorphus_taiwanensis | 0 | 6.46 | 5.22 | 8542.11 |
| lizards | Sphenomorphus_tanneri | Sphenomorphus_tanneri | 0 | 2.63 | 2 | 19615.93 |
| lizards | Sphenomorphus_tonkinensis | Sphenomorphus_tonkinensis | 0 | 2.69 | 3.5 | 312.98 |

| | | | | | | |
|---|---|---|---|---|---|---|
| lizards | Sphenomorphus_undulatus | Sphenomorphus_undulatus | 0 | 6.17 | 2 | 51777.4 |
| lizards | Sphenomorphus_vanheurni | Sphenomorphus_vanheurni | 0 | 5.13 | 2 | 19.98 |
| lizards | Spondylurus_anegadae | Spondylurus_anegadae | 1 | 6.92 | 1.7 | 41.59 |
| lizards | Spondylurus_caicosae | Spondylurus_caicosae | 0 | 9.55 | 2.2 | 513.49 |
| lizards | Spondylurus_culebrae | Spondylurus_culebrae | 1 | 19.95 | 3.55 | 30.25 |
| lizards | Spondylurus_fulgidus | Spondylurus_fulgidus | 1 | 12.59 | 4 | 1215.82 |
| lizards | Spondylurus_haitiae | Spondylurus_haitiae | 1 | 12.88 | 4 | 9.99 |
| lizards | Spondylurus_lineolatus | Spondylurus_lineolatus | 1 | 5.01 | 2 | 19028.59 |
| lizards | Spondylurus_monae | Spondylurus_monae | 1 | 13.8 | 3 | 58.34 |
| lizards | Spondylurus_nitidus | Spondylurus_nitidus | 1 | 18.62 | 2 | 752.93 |
| lizards | Spondylurus_powelli | Spondylurus_powelli | 1 | 7.41 | 3 | 104.3 |
| lizards | Spondylurus_sloanii | Spondylurus_sloanii | 1 | 20.89 | 3 | 244.26 |
| lizards | Spondylurus_turksae | Spondylurus_turksae | 1 | 10.23 | 3 | 31.99 |
| lizards | Stenocercus_angel | Stenocercus_angel | 0 | 24.55 | 2 | 5639.94 |
| lizards | Stenocercus_angulifer | Stenocercus_angulifer | 0 | 32.36 | 2 | 6657.32 |
| lizards | Stenocercus_azureus | Stenocercus_azureus | 0 | 21.38 | 5.5 | 219015.23 |
| lizards | Stenocercus_cadlei | Stenocercus_cadlei | 0 | 22.91 | 2 | 5181.8 |
| lizards | Stenocercus_caducus | Stenocercus_caducus | 0 | 29.51 | 2.75 | 1582548.45 |
| lizards | Stenocercus_chinchaoensis | Stenocercus_chinchaoensis | 1 | 23.99 | 2 | 19.98 |
| lizards | Stenocercus_chota | Stenocercus_chota | 0 | 33.11 | 2 | 303.08 |
| lizards | Stenocercus_chrysopygus | Stenocercus_chrysopygus | 0 | 19.5 | 5.5 | 15603.93 |
| lizards | Stenocercus_cupreus | Stenocercus_cupreus | 0 | 18.2 | 3.5 | 3610.49 |
| lizards | Stenocercus_doellojuradoi | Stenocercus_doellojuradoi | 0 | 19.5 | 6.5 | 251234.79 |
| lizards | Stenocercus_dumerilii | Stenocercus_dumerilii | 1 | 48.98 | 4 | 268590.04 |
| lizards | Stenocercus_empetrus | Stenocercus_empetrus | 0 | 38.9 | 6 | 2159.23 |
| lizards | Stenocercus_eunetopsis | Stenocercus_eunetopsis | 0 | 21.38 | 2 | 9.99 |
| lizards | Stenocercus_fimbriatus | Stenocercus_fimbriatus | 0 | 27.54 | 3.5 | 366909 |
| lizards | Stenocercus_guentheri | Stenocercus_guentheri | 0 | 32.36 | 2 | 24528.27 |
| lizards | Stenocercus_huancabambae | Stenocercus_huancabambae | 0 | 34.67 | 2 | 2558.81 |
| lizards | Stenocercus_humeralis | Stenocercus_humeralis | 0 | 48.98 | 4 | 452.3 |
| lizards | Stenocercus_imitator | Stenocercus_imitator | 0 | 35.48 | 4.5 | 9913.4 |
| lizards | Stenocercus_lache | Stenocercus_lache | 0 | 25.12 | 2 | 2883.55 |
| lizards | Stenocercus_melanopygus | Stenocercus_melanopygus | 0 | 22.91 | 6 | 3280.27 |
| lizards | Stenocercus_ornatissimus | Stenocercus_ornatissimus | 0 | 16.22 | 2 | 1571.73 |

| | | | | | | |
|---|---|---|---|---|---|---|
| lizards | Stenocercus_ornatus | Stenocercus_ornatus | 1 | 22.91 | 2 | 2006.07 |
| lizards | Stenocercus_percultus | Stenocercus_percultus | 0 | 42.66 | 2 | 3864.38 |
| lizards | Stenocercus_prionotus | Stenocercus_prionotus | 0 | 29.51 | 2 | 289499.7 |
| lizards | Stenocercus_puyango | Stenocercus_puyango | 0 | 52.48 | 2 | 17869.23 |
| lizards | Stenocercus_roseiventris | Stenocercus_roseiventris | 0 | 37.15 | 2 | 1215734.9 |
| lizards | Stenocercus_santander | Stenocercus_santander | 0 | 32.36 | 4 | 10727.6 |
| lizards | Stenocercus_torquatus | Stenocercus_torquatus | 1 | 22.39 | 2 | 6273.05 |
| lizards | Stenocercus_trachycephalus | Stenocercus_trachycephalus | 0 | 26.92 | 2 | 29581.24 |
| lizards | Stenocercus_varius | Stenocercus_varius | 1 | 22.91 | 2 | 998.81 |
| lizards | Stenodactylus_doriae | Stenodactylus_doriae | 0 | 11.48 | 1.81 | 2966808.13 |
| lizards | Stenodactylus_grandiceps | Stenodactylus_grandiceps | 0 | 4.9 | 2 | 468036.75 |
| lizards | Stenodactylus_petrii | Stenodactylus_petrii | 0 | 7.41 | 2 | 7334237.65 |
| lizards | Stenodactylus_slevini | Stenodactylus_slevini | 0 | 13.49 | 2 | 1192838.56 |
| lizards | Strobilurus_torquatus | Strobilurus_torquatus | 0 | 42.66 | 2.5 | 261835.17 |
| lizards | Strophurus_assimilis | Strophurus_assimilis | 0 | 9.55 | 2 | 320714.06 |
| lizards | Strophurus_ciliaris | Strophurus_ciliaris | 0 | 14.79 | 2 | 3961905.44 |
| lizards | Strophurus_elderi | Strophurus_elderi | 0 | 2.63 | 1.92 | 2487114.84 |
| lizards | Strophurus_intermedius | Strophurus_intermedius | 0 | 10.23 | 2 | 2115405.26 |
| lizards | Strophurus_jeanae | Strophurus_jeanae | 0 | 2.45 | 2 | 1368479.81 |
| lizards | Strophurus_krisalys | Strophurus_krisalys | 0 | 8.91 | 2 | 615003.32 |
| lizards | Strophurus_mcmillani | Strophurus_mcmillani | 0 | 2.95 | 2 | 59894.77 |
| lizards | Strophurus_michaelseni | Strophurus_michaelseni | 0 | 5.75 | 2 | 36254.14 |
| lizards | Strophurus_rankini | Strophurus_rankini | 0 | 5.01 | 2 | 14827.05 |
| lizards | Strophurus_spinigerus | Strophurus_spinigerus | 0 | 10 | 2 | 322149.95 |
| lizards | Strophurus_strophurus | Strophurus_strophurus | 0 | 8.91 | 2 | 919280.07 |
| lizards | Strophurus_taeniatus | Strophurus_taeniatus | 0 | 2.45 | 2 | 229984.8 |
| lizards | Strophurus_taenicauda | Strophurus_taenicauda | 0 | 7.94 | 2 | 183820.19 |
| lizards | Strophurus_williamsi | Strophurus_williamsi | 0 | 6.92 | 2 | 1142513.22 |
| lizards | Strophurus_wilsoni | Strophurus_wilsoni | 0 | 4.27 | 2 | 31963.98 |
| lizards | Takydromus_amurensis | Takydromus_amurensis | 0 | 11.75 | 5 | 1082455.11 |
| lizards | Takydromus_dorsalis | Takydromus_dorsalis | 1 | 7.94 | 1.4 | 528.95 |
| lizards | Takydromus_formosanus | Takydromus_formosanus | 0 | 6.17 | 2 | 17564.48 |
| lizards | Takydromus_hsuehshanensis | Takydromus_hsuehshanensis | 0 | 8.71 | 2.78 | 3517.44 |
| lizards | Takydromus_khasiensis | Takydromus_khasiensis | 0 | 9.77 | 2 | 189038.79 |

| | | | | | | |
|---|---|---|---|---|---|---|
| lizards | Takydromus_kuehnei | Takydromus_kuehnei | 0 | 6.61 | 1.8 | 924777.09 |
| lizards | Takydromus_luyeanus | Takydromus_luyeanus | 0 | 3.72 | 2.5 | 4584.89 |
| lizards | Takydromus_sauteri | Takydromus_sauteri | 0 | 10.47 | 2 | 6749.74 |
| lizards | Takydromus_septentrionalis | Takydromus_septentrionalis | 0 | 11.75 | 3.3 | 2465519.62 |
| lizards | Takydromus_sexlineatus | Takydromus_sexlineatus | 0 | 7.94 | 2.2 | 4405213.07 |
| lizards | Takydromus_smaragdinus | Takydromus_smaragdinus | 0 | 6.46 | 2 | 2664.38 |
| lizards | Takydromus_stejnegeri | Takydromus_stejnegeri | 0 | 5.62 | 2.2 | 19041.83 |
| lizards | Takydromus_sylvaticus | Takydromus_sylvaticus | 0 | 4.57 | 3 | 11695.31 |
| lizards | Takydromus_tachydromoides | Takydromus_tachydromoides | 0 | 7.94 | 4 | 368570.01 |
| lizards | Takydromus_toyamai | Takydromus_toyamai | 1 | 3.89 | 2 | 193.47 |
| lizards | Takydromus_viridipunctatus | Takydromus_viridipunctatus | 0 | 4.9 | 2.5 | 5773.19 |
| lizards | Takydromus_wolteri | Takydromus_wolteri | 0 | 6.76 | 3.75 | 1893523.43 |
| lizards | Tarentola_americana | Tarentola_americana | 0 | 43.65 | 2 | 114780.05 |
| lizards | Tarentola_angustimentalis | Tarentola_angustimentalis | 0 | 13.18 | 1.92 | 2521.2 |
| lizards | Tarentola_annularis | Tarentola_annularis | 0 | 69.18 | 1.5 | 5606088.15 |
| lizards | Tarentola_boavistensis | Tarentola_boavistensis | 1 | 12.88 | 1.5 | 630.9 |
| lizards | Tarentola_boehmei | Tarentola_boehmei | 0 | 14.13 | 1.5 | 74734.55 |
| lizards | Tarentola_boettgeri | Tarentola_boettgeri | 0 | 10.72 | 1.5 | 1755.32 |
| lizards | Tarentola_chazaliae | Tarentola_chazaliae | 1 | 10.47 | 1.5 | 31281.17 |
| lizards | Tarentola_crombiei | Tarentola_crombiei | 0 | 5.13 | 1 | 212.84 |
| lizards | Tarentola_darwini | Tarentola_darwini | 0 | 13.18 | 1 | 993.53 |
| lizards | Tarentola_delalandii | Tarentola_delalandii | 0 | 15.49 | 1.5 | 2744.12 |
| lizards | Tarentola_deserti | Tarentola_deserti | 0 | 28.18 | 2 | 353807.28 |
| lizards | Tarentola_ephippiata | Tarentola_ephippiata | 0 | 54.95 | 1.9 | 7434256.94 |
| lizards | Tarentola_gigas | Tarentola_gigas | 1 | 93.33 | 2 | 10.72 |
| lizards | Tarentola_gomerensis | Tarentola_gomerensis | 0 | 11.48 | 1.5 | 365.53 |
| lizards | Tarentola_maioensis | Tarentola_maioensis | 0 | 9.33 | 1.5 | 275.33 |
| lizards | Tarentola_mauritanica | Tarentola_mauritanica | 0 | 16.6 | 2 | 1852529.32 |
| lizards | Tarentola_mindiae | Tarentola_mindiae | 0 | 13.8 | 2 | 72616.34 |
| lizards | Tarentola_neglecta | Tarentola_neglecta | 0 | 10.23 | 1 | 1321275.59 |
| lizards | Tarentola_protogigas | Tarentola_protogigas | 0 | 24.55 | 1.5 | 205.25 |
| lizards | Techmarscincus_jigurru | Techmarscincus_jigurru | 1 | 10.47 | 3 | 30.3 |
| lizards | Teira_dugesii | Teira_dugesii | 0 | 12.3 | 2.5 | 815.08 |
| lizards | Teius_oculatus | Teius_oculatus | 0 | 52.48 | 4.85 | 1753401.04 |

| | | | | | | |
|---|---|---|---|---|---|---|
| lizards | Teius_suquiensis | Teius_suquiensis | 0 | 58.88 | 3.95 | 74034.56 |
| lizards | Teius_teyou | Teius_teyou | 0 | 104.71 | 4.8 | 1622772.27 |
| lizards | Tenuidactylus_bogdanovi | Tenuidactylus_bogdanovi | 0 | 7.76 | 1.5 | 40289.05 |
| lizards | Tenuidactylus_caspius | Tenuidactylus_caspius | 0 | 10.23 | 2 | 2125031.92 |
| lizards | Tenuidactylus_elongatus | Tenuidactylus_elongatus | 0 | 3.98 | 2 | 823802.87 |
| lizards | Tenuidactylus_fedtschenkoi | Tenuidactylus_fedtschenkoi | 0 | 9.33 | 1.78 | 292971.55 |
| lizards | Tenuidactylus_longipes | Tenuidactylus_longipes | 0 | 6.76 | 1.5 | 343865.49 |
| lizards | Tenuidactylus_turcmenicus | Tenuidactylus_turcmenicus | 0 | 10.23 | 2 | 127397.07 |
| lizards | Teratoscincus_bedriagai | Teratoscincus_bedriagai | 0 | 9.55 | 1.5 | 309286.92 |
| lizards | Teratoscincus_microlepis | Teratoscincus_microlepis | 0 | 18.2 | 2 | 365555.14 |
| lizards | Teratoscincus_przewalskii | Teratoscincus_przewalskii | 0 | 21.88 | 2 | 1506810.98 |
| lizards | Teratoscincus_roborowskii | Teratoscincus_roborowskii | 0 | 20.89 | 2 | 33584.64 |
| lizards | Teratoscincus_scincus | Teratoscincus_scincus | 0 | 41.69 | 2 | 1871823.42 |
| lizards | Tetradactylus_africanus | Tetradactylus_africanus | 0 | 3.89 | 3.5 | 62541.75 |
| lizards | Tetradactylus_breyeri | Tetradactylus_breyeri | 0 | 2.4 | 1.5 | 18726.46 |
| lizards | Tetradactylus_ellenbergeri | Tetradactylus_ellenbergeri | 0 | 2.4 | 2 | 470440.28 |
| lizards | Tetradactylus_seps | Tetradactylus_seps | 0 | 7.41 | 2.5 | 78876.68 |
| lizards | Tetradactylus_tetradactylus | Tetradactylus_tetradactylus | 0 | 2.51 | 4 | 84034.52 |
| lizards | Tetradactylus_udzungwensis | Tetradactylus_udzungwensis | 1 | 2 | 2 | 19.98 |
| lizards | Thecadactylus_rapicauda | Thecadactylus_rapicauda | 0 | 50.12 | 1 | 7382390.2 |
| lizards | Thecadactylus_solimoensis | Thecadactylus_solimoensis | 0 | 50.12 | 1 | 2751783.83 |
| lizards | Tiliqua_adelaidensis | Tiliqua_adelaidensis | 1 | 26.92 | 2.41 | 11850.05 |
| lizards | Tiliqua_gigas | Tiliqua_gigas | 0 | 1148.15 | 8.5 | 441063.24 |
| lizards | Tiliqua_multifasciata | Tiliqua_multifasciata | 0 | 741.31 | 4.2 | 3512589.31 |
| lizards | Tiliqua_nigrolutea | Tiliqua_nigrolutea | 0 | 1445.44 | 7 | 362450.69 |
| lizards | Tiliqua_occipitalis | Tiliqua_occipitalis | 0 | 912.01 | 5 | 2510711.02 |
| lizards | Tiliqua_rugosa | Tiliqua_rugosa | 0 | 1230.27 | 2.05 | 3235976.76 |
| lizards | Tiliqua_scincoides | Tiliqua_scincoides | 0 | 1621.81 | 11 | 3476688.89 |
| lizards | Timon_lepidus | Timon_lepidus | 0 | 380.19 | 13.7 | 715237.91 |
| lizards | Timon_pater | Timon_pater | 0 | 109.65 | 11 | 235602.93 |
| lizards | Timon_princeps | Timon_princeps | 0 | 72.44 | 7 | 64199.22 |
| lizards | Timon_tangitanus | Timon_tangitanus | 0 | 57.54 | 10 | 341580.13 |
| lizards | Toropuku_stephensi | Toropuku_stephensi | 1 | 12.3 | 1.93 | 294.54 |
| lizards | Tracheloptychus_madagascariensis | Tracheloptychus_madagascariensis | 0 | 17.78 | 3 | 97239.92 |

| | | | | | | |
|---|---|---|---|---|---|---|
| lizards | Trachydactylus_spatalurus | Bunopus_spatalurus | 0 | 7.08 | 1.5 | 446738.22 |
| lizards | Trachylepis_acutilabris | Trachylepis_acutilabris | 0 | 4.17 | 4.5 | 370708.95 |
| lizards | Trachylepis_affinis | Trachylepis_affinis | 0 | 11.75 | 2.72 | 5568536.91 |
| lizards | Trachylepis_albilabris | Trachylepis_albilabris | 0 | 8.51 | 3.5 | 484550.49 |
| lizards | Trachylepis_bayonii | Trachylepis_bayonii | 0 | 11.22 | 5.5 | 3822391.94 |
| lizards | Trachylepis_bocagii | Trachylepis_bocagii | 0 | 7.76 | 5.2 | 237104.93 |
| lizards | Trachylepis_boettgeri | Trachylepis_boettgeri | 0 | 3.09 | 2 | 43162.17 |
| lizards | Trachylepis_boulengeri | Trachylepis_boulengeri | 0 | 30.9 | 5.5 | 766840.78 |
| lizards | Trachylepis_brauni | Trachylepis_brauni | 0 | 11.22 | 9 | 3578.39 |
| lizards | Trachylepis_brevicollis | Trachylepis_brevicollis | 0 | 144.54 | 3 | 3076857.86 |
| lizards | Trachylepis_buettneri | Trachylepis_buettneri | 0 | 15.49 | 8.7 | 780397.39 |
| lizards | Trachylepis_capensis | Trachylepis_capensis | 0 | 56.23 | 11.62 | 1821762.4 |
| lizards | Trachylepis_comorensis | Trachylepis_comorensis | 0 | 30.9 | 5 | 1964.52 |
| lizards | Trachylepis_depressa | Trachylepis_depressa | 0 | 13.8 | 7 | 170914.58 |
| lizards | Trachylepis_dichroma | Trachylepis_dichroma | 0 | 158.49 | 3 | 105146.46 |
| lizards | Trachylepis_gravenhorstii | Trachylepis_gravenhorstii | 0 | 17.78 | 3.5 | 583710.13 |
| lizards | Trachylepis_homalocephala | Trachylepis_homalocephala | 0 | 19.95 | 6 | 246655.21 |
| lizards | Trachylepis_irregularis | Trachylepis_irregularis | 0 | 8.51 | 5 | 59335.26 |
| lizards | Trachylepis_lacertiformis | Trachylepis_lacertiformis | 0 | 2.95 | 4.5 | 231841.34 |
| lizards | Trachylepis_lavarambo | Trachylepis_lavarambo | 1 | 4.37 | 3 | 291.32 |
| lizards | Trachylepis_maculilabris | Trachylepis_maculilabris | 0 | 19.95 | 7.04 | 6788234.54 |
| lizards | Trachylepis_madagascariensis | Trachylepis_madagascariensis | 0 | 9.55 | 5 | 32941.71 |
| lizards | Trachylepis_makolowodei | Trachylepis_makolowodei | 0 | 44.67 | 6.5 | 559359.22 |
| lizards | Trachylepis_margaritifera | Trachylepis_margaritifera | 0 | 38.9 | 6.1 | 1806100.15 |
| lizards | Trachylepis_megalura | Trachylepis_megalura | 0 | 12.88 | 4.8 | 1992826.73 |
| lizards | Trachylepis_nganghae | Trachylepis_nganghae | 1 | 4.17 | 4 | 19.98 |
| lizards | Trachylepis_occidentalis | Trachylepis_occidentalis | 0 | 33.88 | 5.35 | 939225.65 |
| lizards | Trachylepis_perrotetii | Trachylepis_perrotetii | 0 | 144.54 | 20.8 | 520829.89 |
| lizards | Trachylepis_planifrons | Trachylepis_planifrons | 0 | 34.67 | 8.5 | 1456491.39 |
| lizards | Trachylepis_polytropis | Trachylepis_polytropis | 0 | 33.11 | 3.5 | 1592079.71 |
| lizards | Trachylepis_pulcherrima | Trachylepis_pulcherrima | 0 | 39.81 | 7 | 18494.22 |
| lizards | Trachylepis_punctatissima | Trachylepis_punctatissima | 0 | 26.92 | 4.8 | 1021857.2 |
| lizards | Trachylepis_sechellensis | Trachylepis_sechellensis | 0 | 26.92 | 4 | 246.59 |
| lizards | Trachylepis_sparsa | Trachylepis_sparsa | 0 | 27.54 | 5.15 | 180145.35 |

| | | | | | | |
|---|---|---|---|---|---|---|
| lizards | Trachylepis_spilogaster | Trachylepis_spilogaster | 0 | 16.98 | 4.86 | 892599.19 |
| lizards | Trachylepis_striata | Trachylepis_striata | 0 | 31.62 | 6.7 | 5399481.89 |
| lizards | Trachylepis_sulcata | Trachylepis_sulcata | 0 | 29.51 | 3.75 | 1242359.54 |
| lizards | Trachylepis_tavaratra | Trachylepis_tavaratra | 1 | 4.57 | 5 | 10529.29 |
| lizards | Trachylepis_tessellata | Trachylepis_tessellata | 0 | 21.38 | 2.5 | 356472.45 |
| lizards | Trachylepis_varia | Trachylepis_varia | 0 | 7.08 | 7.36 | 7946224.69 |
| lizards | Trachylepis_variegata | Trachylepis_variegata | 0 | 3.47 | 2.32 | 906913.78 |
| lizards | Trachylepis_vato | Trachylepis_vato | 0 | 3.09 | 8 | 93023.69 |
| lizards | Trachylepis_wrightii | Trachylepis_wrightii | 1 | 97.72 | 4.5 | 3.63 |
| lizards | Trapelus_agilis | Trapelus_agilis | 0 | 52.48 | 7 | 3146170.54 |
| lizards | Trapelus_agnetae | Trapelus_agnetae | 0 | 26.92 | 9.7 | 1353248.28 |
| lizards | Trapelus_flavimaculatus | Trapelus_flavimaculatus | 0 | 75.86 | 5 | 2148031.31 |
| lizards | Trapelus_megalonyx | Trapelus_megalonyx | 0 | 13.8 | 5 | 808903.3 |
| lizards | Trapelus_mutabilis | Trapelus_mutabilis | 0 | 32.36 | 8.5 | 4067499.63 |
| lizards | Trapelus_ruderatus | Trapelus_ruderatus | 0 | 85.11 | 8 | 2535454.79 |
| lizards | Trapelus_sanguinolentus | Trapelus_sanguinolentus | 0 | 239.88 | 7.5 | 2563982.65 |
| lizards | Trapelus_savignii | Trapelus_savignii | 1 | 63.1 | 5.15 | 35892.02 |
| lizards | Trapelus_tournevillei | Trapelus_tournevillei | 0 | 33.11 | 6 | 487687.7 |
| lizards | Tribolonotus_blanchardi | Tribolonotus_blanchardi | 0 | 1.12 | 1 | 8572.39 |
| lizards | Tribolonotus_gracilis | Tribolonotus_gracilis | 0 | 23.44 | 1 | 117110.41 |
| lizards | Tribolonotus_novaeguineae | Tribolonotus_novaeguineae | 0 | 23.44 | 1 | 137845.66 |
| lizards | Tribolonotus_pseudoponceleti | Tribolonotus_pseudoponceleti | 0 | 8.91 | 1 | 12171.08 |
| lizards | Tribolonotus_schmidti | Tribolonotus_schmidti | 0 | 1.2 | 1 | 5461.35 |
| lizards | Trioceros_affinis | Trioceros_affinis | 0 | 13.18 | 16 | 343971.01 |
| lizards | Trioceros_balebicornutus | Trioceros_balebicornutus | 0 | 10.96 | 16 | 1291.2 |
| lizards | Trioceros_bitaeniatus | Trioceros_bitaeniatus | 0 | 18.62 | 13.9 | 1350654.57 |
| lizards | Trioceros_camerunensis | Trioceros_camerunensis | 0 | 17.38 | 8.5 | 23332.99 |
| lizards | Trioceros_cristatus | Trioceros_cristatus | 0 | 81.28 | 24 | 1322406.39 |
| lizards | Trioceros_deremensis | Trioceros_deremensis | 0 | 89.13 | 24 | 6206.16 |
| lizards | Trioceros_ellioti | Trioceros_ellioti | 0 | 21.38 | 10.1 | 1007953.11 |
| lizards | Trioceros_feae | Trioceros_feae | 0 | 25.7 | 4 | 924.63 |
| lizards | Trioceros_fuelleborni | Trioceros_fuelleborni | 0 | 26.3 | 15 | 500.82 |
| lizards | Trioceros_goetzei | Trioceros_goetzei | 0 | 19.95 | 8 | 42434.8 |
| lizards | Trioceros_harennae | Trioceros_harennae | 0 | 10.47 | 10 | 5204.66 |

| | | | | | | |
|---|---|---|---|---|---|---|
| lizards | Trioceros_hoehnelii | Trioceros_hoehnelii | 0 | 29.51 | 11.7 | 56737.32 |
| lizards | Trioceros_incornutus | Trioceros_incornutus | 0 | 19.05 | 13.5 | 15064.91 |
| lizards | Trioceros_ituriensis | Trioceros_ituriensis | 0 | 46.77 | 9 | 459565.93 |
| lizards | Trioceros_jacksonii | Trioceros_jacksonii | 0 | 91.2 | 16.95 | 31172.36 |
| lizards | Trioceros_johnstoni | Trioceros_johnstoni | 0 | 46.77 | 13.5 | 265597.46 |
| lizards | Trioceros_laterispinis | Trioceros_laterispinis | 1 | 12.3 | 15.5 | 11538.54 |
| lizards | Trioceros_melleri | Trioceros_melleri | 0 | 389.05 | 47 | 379655.57 |
| lizards | Trioceros_montium | Trioceros_montium | 0 | 38.9 | 6.5 | 21680.7 |
| lizards | Trioceros_oweni | Trioceros_oweni | 0 | 66.07 | 16 | 2129943.35 |
| lizards | Trioceros_perreti | Trioceros_perreti | 1 | 15.49 | 9.5 | 658.86 |
| lizards | Trioceros_pfefferi | Trioceros_pfefferi | 1 | 19.95 | 9 | 10519.04 |
| lizards | Trioceros_quadricornis | Trioceros_quadricornis | 1 | 93.33 | 13 | 13912.41 |
| lizards | Trioceros_rudis | Trioceros_rudis | 0 | 10.72 | 11.5 | 327813.06 |
| lizards | Trioceros_schubotzi | Trioceros_schubotzi | 0 | 14.13 | 8.5 | 516.09 |
| lizards | Trioceros_serratus | Trioceros_serratus | 1 | 18.62 | 8.67 | 6423.02 |
| lizards | Trioceros_sternfeldi | Trioceros_sternfeldi | 0 | 16.22 | 10.5 | 22624.66 |
| lizards | Trioceros_tempeli | Trioceros_tempeli | 0 | 32.36 | 21.5 | 7826.97 |
| lizards | Trioceros_werneri | Trioceros_werneri | 0 | 25.12 | 19 | 26795.45 |
| lizards | Trogonophis_wiegmanni | Trogonophis_wiegmanni | 0 | 14.13 | 5 | 505845.32 |
| lizards | Tropicagama_temporalis | Lophognathus_temporalis | 0 | 123.03 | 4.5 | 501763.59 |
| lizards | Tropidophorus_baviensis | Tropidophorus_baviensis | 0 | 16.6 | 5.5 | 120341.72 |
| lizards | Tropidophorus_beccarii | Tropidophorus_beccarii | 0 | 19.95 | 4 | 298445.11 |
| lizards | Tropidophorus_berdmorei | Tropidophorus_berdmorei | 0 | 19.5 | 11.5 | 348334.77 |
| lizards | Tropidophorus_brookei | Tropidophorus_brookei | 0 | 29.51 | 3.85 | 338300.75 |
| lizards | Tropidophorus_cocincinensis | Tropidophorus_cocincinensis | 0 | 22.39 | 6.5 | 92976.81 |
| lizards | Tropidophorus_davaoensis | Tropidophorus_davaoensis | 0 | 52.48 | 4.5 | 49600.77 |
| lizards | Tropidophorus_grayi | Tropidophorus_grayi | 0 | 38.02 | 3.8 | 71193.13 |
| lizards | Tropidophorus_hainanus | Tropidophorus_hainanus | 0 | 3.63 | 4 | 287009.47 |
| lizards | Tropidophorus_laotus | Tropidophorus_laotus | 1 | 15.49 | 4.6 | 41996.53 |
| lizards | Tropidophorus_microlepis | Tropidophorus_microlepis | 0 | 11.75 | 8 | 72503.97 |
| lizards | Tropidophorus_misaminius | Tropidophorus_misaminius | 0 | 30.9 | 3 | 52148.56 |
| lizards | Tropidophorus_murphyi | Tropidophorus_murphyi | 0 | 19.05 | 4 | 19.32 |
| lizards | Tropidophorus_noggei | Tropidophorus_noggei | 0 | 29.51 | 4.5 | 1222.77 |
| lizards | Tropidophorus_partelloi | Tropidophorus_partelloi | 0 | 46.77 | 5 | 94145.24 |

| | | | | | | |
|---|---|---|---|---|---|---|
| lizards | Tropidophorus_robinsoni | Tropidophorus_robinsoni | 0 | 8.51 | 4.5 | 48939.74 |
| lizards | Tropidophorus_sinicus | Tropidophorus_sinicus | 0 | 7.08 | 6 | 281145.25 |
| lizards | Tropidosaura_cottrelli | Tropidosaura_cottrelli | 0 | 8.71 | 4 | 5427.2 |
| lizards | Tropidosaura_essexi | Tropidosaura_essexi | 0 | 3.31 | 2 | 28140.74 |
| lizards | Tropidosaura_gularis | Tropidosaura_gularis | 0 | 5.62 | 6 | 72028.11 |
| lizards | Tropidosaura_montana | Tropidosaura_montana | 0 | 6.76 | 4.5 | 257256.86 |
| lizards | Tropidoscincus_boreus | Tropidoscincus_boreus | 0 | 18.2 | 2.5 | 11545.47 |
| lizards | Tropidoscincus_variabilis | Tropidoscincus_variabilis | 0 | 18.2 | 2.8 | 4548.8 |
| lizards | Tropidurus_bogerti | Tropidurus_bogerti | 0 | 28.84 | 1 | 1525.26 |
| lizards | Tropidurus_callathelys | Tropidurus_callathelys | 0 | 33.88 | 2 | 19.98 |
| lizards | Tropidurus_catalanensis | Tropidurus_catalanensis | 0 | 77.62 | 6.15 | 544715.38 |
| lizards | Tropidurus_chromatops | Tropidurus_chromatops | 0 | 45.71 | 4.5 | 5886.91 |
| lizards | Tropidurus_cocorobensis | Tropidurus_cocorobensis | 0 | 16.22 | 3.05 | 72805.58 |
| lizards | Tropidurus_erythrocephalus | Tropidurus_erythrocephalus | 1 | 22.39 | 2 | 26701.21 |
| lizards | Tropidurus_etheridgei | Tropidurus_etheridgei | 0 | 52.48 | 5.29 | 1548687.11 |
| lizards | Tropidurus_guarani | Tropidurus_guarani | 0 | 52.48 | 2 | 438039.89 |
| lizards | Tropidurus_hispidus | Tropidurus_hispidus | 0 | 95.5 | 6.85 | 3583749.07 |
| lizards | Tropidurus_hygomi | Tropidurus_hygomi | 1 | 19.5 | 2 | 15763.29 |
| lizards | Tropidurus_insulanus | Tropidurus_insulanus | 0 | 23.99 | 3.8 | 65542.84 |
| lizards | Tropidurus_itambere | Tropidurus_itambere | 0 | 30.9 | 3.74 | 1650871.85 |
| lizards | Tropidurus_melanopleurus | Tropidurus_melanopleurus | 0 | 45.71 | 2 | 5037.44 |
| lizards | Tropidurus_montanus | Tropidurus_montanus | 0 | 34.67 | 3.49 | 133739.51 |
| lizards | Tropidurus_oreadicus | Tropidurus_oreadicus | 0 | 54.95 | 4.37 | 4046111.74 |
| lizards | Tropidurus_pinima | Tropidurus_pinima | 0 | 22.91 | 2 | 302.65 |
| lizards | Tropidurus_psammonastes | Tropidurus_psammonastes | 1 | 34.67 | 4 | 5722.81 |
| lizards | Tropidurus_semitaeniatus | Tropidurus_semitaeniatus | 0 | 35.48 | 2.01 | 964683.25 |
| lizards | Tropidurus_spinulosus | Tropidurus_spinulosus | 0 | 89.13 | 4.9 | 1027962.33 |
| lizards | Tropidurus_torquatus | Tropidurus_torquatus | 0 | 79.43 | 4.71 | 4145465.99 |
| lizards | Tropidurus_xanthochilus | Tropidurus_xanthochilus | 1 | 66.07 | 5 | 9.99 |
| lizards | Tropiocolotes_algericus | Tropiocolotes_algericus | 0 | 1.23 | 1 | 455120.05 |
| lizards | Tropiocolotes_nattereri | Tropiocolotes_nattereri | 0 | 0.69 | 1 | 84669.69 |
| lizards | Tropiocolotes_somalicus | Tropiocolotes_somalicus | 0 | 0.56 | 1 | 8022.59 |
| lizards | Tropiocolotes_steudneri | Tropiocolotes_steudneri | 0 | 1.02 | 1 | 2758118.77 |
| lizards | Tropiocolotes_tripolitanus | Tropiocolotes_tripolitanus | 0 | 1.23 | 1 | 5063135.15 |

| | | | | | | |
|---|---|---|---|---|---|---|
| lizards | Tukutuku_rakiurae | Tukutuku_rakiurae | 1 | 7.24 | 1.5 | 1692.31 |
| lizards | Tupinambis_teguixin | Tupinambis_teguixin | 0 | 4466.84 | 11.95 | 9385193.53 |
| lizards | Tympanocryptis_centralis | Tympanocryptis_centralis | 0 | 7.24 | 7 | 552386.11 |
| lizards | Tympanocryptis_intima | Tympanocryptis_intima | 0 | 7.24 | 8.7 | 1159092.28 |
| lizards | Tympanocryptis_lineata | Tympanocryptis_lineata | 0 | 12.02 | 6.5 | 981181.57 |
| lizards | Tympanocryptis_pinguicolla | Tympanocryptis_pinguicolla | 1 | 8.71 | 4.5 | 8173.17 |
| lizards | Tympanocryptis_tetraporophora | Tympanocryptis_tetraporophora | 0 | 13.18 | 8.1 | 2308571.2 |
| lizards | Typhlacontias_brevipes | Typhlacontias_brevipes | 0 | 4.79 | 3 | 57199.89 |
| lizards | Typhlacontias_rohani | Typhlacontias_rohani | 0 | 1.95 | 3.5 | 595828.28 |
| lizards | Typhlosaurus_caecus | Typhlosaurus_caecus | 0 | 14.13 | 2 | 8003.5 |
| lizards | Typhlosaurus_vermis | Typhlosaurus_vermis | 0 | 25.7 | 2.5 | 19453.7 |
| lizards | Tytthoscincus_aesculeticola | Tytthoscincus_aesculeticola | 0 | 1.41 | 2 | 1433.87 |
| lizards | Tytthoscincus_bukitensis | Sphenomorphus_bukitensis | 0 | 1.51 | 1 | 866.58 |
| lizards | Tytthoscincus_hallieri | Tytthoscincus_hallieri | 0 | 3.47 | 2 | 156661.18 |
| lizards | Tytthoscincus_temmincki | Sphenomorphus_temmincki | 0 | 3.31 | 3.4 | 12431.98 |
| lizards | Uma_exsul | Uma_exsul | 1 | 31.62 | 2.75 | 10958.17 |
| lizards | Uma_inornata | Uma_inornata | 1 | 56.23 | 2.45 | 734.75 |
| lizards | Uma_notata | Uma_notata | 0 | 53.7 | 2.5 | 20568.98 |
| lizards | Uma_paraphygas | Uma_paraphygas | 0 | 22.91 | 8 | 1945.98 |
| lizards | Uma_scoparia | Uma_scoparia | 0 | 46.77 | 2.86 | 39209.63 |
| lizards | Underwoodisaurus_milii | Underwoodisaurus_milii | 0 | 22.91 | 2 | 2778149.51 |
| lizards | Underwoodisaurus_seorsus | Underwoodisaurus_seorsus | 0 | 18.2 | 2 | 14234.85 |
| lizards | Uracentron_azureum | Uracentron_azureum | 0 | 25.12 | 2 | 3804464.19 |
| lizards | Uracentron_flaviceps | Uracentron_flaviceps | 0 | 72.44 | 2 | 1189332.15 |
| lizards | Uranoscodon_superciliosus | Uranoscodon_superciliosus | 0 | 141.25 | 8.65 | 4847371.98 |
| lizards | Urocotyledon_inexpectata | Urocotyledon_inexpectata | 0 | 1.51 | 2 | 129.7 |
| lizards | Uromastyx_acanthinura | Uromastyx_acanthinura | 0 | 2454.71 | 13 | 1472842.99 |
| lizards | Uromastyx_aegyptia | Uromastyx_aegyptia | 1 | 2818.38 | 20.8 | 3447574.29 |
| lizards | Uromastyx_alfredschmidti | Uromastyx_alfredschmidti | 0 | 446.68 | 11.5 | 49623.26 |
| lizards | Uromastyx_benti | Uromastyx_benti | 0 | 269.15 | 8.5 | 293988.35 |
| lizards | Uromastyx_dispar | Uromastyx_dispar | 0 | 457.09 | 17 | 4715456.51 |
| lizards | Uromastyx_geyri | Uromastyx_geyri | 0 | 275.42 | 14.5 | 920201.6 |
| lizards | Uromastyx_macfadyeni | Uromastyx_macfadyeni | 0 | 61.66 | 8 | 115699.72 |
| lizards | Uromastyx_ocellata | Uromastyx_ocellata | 0 | 190.55 | 12 | 1527514.51 |

| | | | | | | |
|---|---|---|---|---|---|---|
| lizards | Uromastyx_ornata | Uromastyx_ornata | 0 | 331.13 | 9 | 261938.17 |
| lizards | Uromastyx_thomasi | Uromastyx_thomasi | 1 | 245.47 | 12 | 99513.08 |
| lizards | Uroplatus_alluaudi | Uroplatus_alluaudi | 0 | 10 | 2 | 2811.33 |
| lizards | Uroplatus_ebenaui | Uroplatus_ebenaui | 1 | 7.24 | 1.5 | 28776.82 |
| lizards | Uroplatus_fimbriatus | Uroplatus_fimbriatus | 0 | 120.23 | 1.5 | 94198.14 |
| lizards | Uroplatus_guentheri | Uroplatus_guentheri | 1 | 10 | 1.5 | 5471.97 |
| lizards | Uroplatus_henkeli | Uroplatus_henkeli | 1 | 69.18 | 2.5 | 12389.28 |
| lizards | Uroplatus_lineatus | Uroplatus_lineatus | 0 | 83.18 | 2 | 43159.48 |
| lizards | Uroplatus_malahelo | Uroplatus_malahelo | 1 | 10 | 2 | 1088.33 |
| lizards | Uroplatus_phantasticus | Uroplatus_phantasticus | 0 | 12.59 | 2 | 44946.07 |
| lizards | Uroplatus_pietschmanni | Uroplatus_pietschmanni | 1 | 10.72 | 2 | 1264.58 |
| lizards | Uroplatus_sameiti | Uroplatus_sameiti | 0 | 28.18 | 2 | 52955.61 |
| lizards | Uroplatus_sikorae | Uroplatus_sikorae | 0 | 33.88 | 1.5 | 88289.28 |
| lizards | Urosaurus_bicarinatus | Urosaurus_bicarinatus | 0 | 10 | 6.6 | 364544.77 |
| lizards | Urosaurus_graciosus | Urosaurus_graciosus | 0 | 11.22 | 4.45 | 183159.46 |
| lizards | Urosaurus_ornatus | Urosaurus_ornatus | 0 | 11.75 | 7.4 | 1685548.26 |
| lizards | Urostrophus_vautieri | Urostrophus_vautieri | 0 | 39.81 | 9.6 | 506846.52 |
| lizards | Uta_encantadae | Uta_encantadae | 1 | 11.75 | 3.55 | 0.62 |
| lizards | Uta_nolascensis | Uta_nolascensis | 0 | 6.31 | 4 | 2.97 |
| lizards | Uta_palmeri | Uta_palmeri | 1 | 19.05 | 2.8 | 1.22 |
| lizards | Uta_squamata | Uta_squamata | 0 | 7.08 | 3.5 | 27.73 |
| lizards | Uta_stansburiana | Uta_stansburiana | 0 | 15.85 | 3.8 | 2345521.99 |
| lizards | Uvidicolus_sphyrurus | Uvidicolus_sphyrurus | 0 | 9.33 | 2 | 59393.84 |
| lizards | Vanzosaura_multiscutata | Vanzosaura_multiscutatus | 0 | 1.1 | 2 | 761798.83 |
| lizards | Vanzosaura_rubricauda | Vanzosaura_rubricauda | 0 | 1.91 | 1.98 | 5109646.33 |
| lizards | Vanzosaura_savanicola | Vanzosaura_savanicola | 0 | 0.87 | 2 | 152213.44 |
| lizards | Varanus_acanthurus | Varanus_acanthurus | 0 | 288.4 | 9.3 | 3077096.87 |
| lizards | Varanus_albigularis | Varanus_albigularis | 0 | 15135.61 | 32 | 8253607.15 |
| lizards | Varanus_auffenbergi | Varanus_auffenbergi | 1 | 151.36 | 5 | 1221.12 |
| lizards | Varanus_bengalensis | Varanus_bengalensis | 0 | 18197.01 | 20.2 | 4839323.3 |
| lizards | Varanus_bogerti | Varanus_bogerti | 0 | 446.68 | 4.1 | 1686.13 |
| lizards | Varanus_brevicauda | Varanus_brevicauda | 0 | 30.9 | 2.92 | 2296478.74 |
| lizards | Varanus_caerulivirens | Varanus_caerulivirens | 0 | 1318.26 | 2 | 14590.35 |
| lizards | Varanus_caudolineatus | Varanus_caudolineatus | 0 | 37.15 | 3.7 | 1179363.03 |

| | | | | | | |
|---|---|---|---|---|---|---|
| lizards | Varanus_cumingi | Varanus_cumingi | 0 | 4897.79 | 7 | 96858.03 |
| lizards | Varanus_doreanus | Varanus_doreanus | 0 | 14125.38 | 9.09 | 647811.21 |
| lizards | Varanus_eremius | Varanus_eremius | 0 | 144.54 | 3.85 | 2912674.98 |
| lizards | Varanus_exanthematicus | Varanus_exanthematicus | 0 | 10000 | 19.05 | 7554393.89 |
| lizards | Varanus_finschi | Varanus_finschi | 0 | 1023.29 | 9.09 | 42667.28 |
| lizards | Varanus_flavescens | Varanus_flavescens | 1 | 2951.21 | 12.25 | 883802.28 |
| lizards | Varanus_giganteus | Varanus_giganteus | 0 | 21379.62 | 9.05 | 2464635.03 |
| lizards | Varanus_gilleni | Varanus_gilleni | 0 | 125.89 | 4.28 | 2155331.71 |
| lizards | Varanus_glauerti | Varanus_glauerti | 0 | 288.4 | 4.5 | 153277.13 |
| lizards | Varanus_glebopalma | Varanus_glebopalma | 0 | 1288.25 | 6.5 | 656187.11 |
| lizards | Varanus_gouldii | Varanus_gouldii | 0 | 6918.31 | 7.81 | 6329091.52 |
| lizards | Varanus_griseus | Varanus_griseus | 0 | 5495.41 | 15.75 | 17979964.4 |
| lizards | Varanus_indicus | Varanus_indicus | 0 | 4365.16 | 7.85 | 651554.2 |
| lizards | Varanus_jobiensis | Varanus_jobiensis | 0 | 1949.84 | 3.63 | 494063 |
| lizards | Varanus_juxtindicus | Varanus_juxtindicus | 0 | 2754.23 | 4 | 675.7 |
| lizards | Varanus_keithhornei | Varanus_keithhornei | 0 | 436.52 | 3.1 | 10996.05 |
| lizards | Varanus_kingorum | Varanus_kingorum | 0 | 26.92 | 4.5 | 40086.06 |
| lizards | Varanus_komodoensis | Varanus_komodoensis | 1 | 109647.82 | 18.4 | 1113.1 |
| lizards | Varanus_lirungensis | Varanus_lirungensis | 1 | 891.25 | 2.5 | 93.23 |
| lizards | Varanus_mabitang | Varanus_mabitang | 1 | 7943.28 | 9 | 336.89 |
| lizards | Varanus_macraei | Varanus_macraei | 1 | 933.25 | 4.03 | 453.71 |
| lizards | Varanus_marmoratus | Varanus_marmoratus | 0 | 11748.98 | 11 | 89087.08 |
| lizards | Varanus_melinus | Varanus_melinus | 1 | 2884.03 | 7.4 | 4741.45 |
| lizards | Varanus_mertensi | Varanus_mertensi | 1 | 3890.45 | 9.7 | 1201720.63 |
| lizards | Varanus_mitchelli | Varanus_mitchelli | 1 | 812.83 | 11.85 | 449926.03 |
| lizards | Varanus_niloticus | Varanus_niloticus | 0 | 23988.33 | 23.38 | 15915291.19 |
| lizards | Varanus_nuchalis | Varanus_nuchalis | 0 | 5011.87 | 10.5 | 34950.57 |
| lizards | Varanus_olivaceus | Varanus_olivaceus | 1 | 9120.11 | 7.1 | 26652.27 |
| lizards | Varanus_panoptes | Varanus_panoptes | 0 | 9549.93 | 8.8 | 3341093.98 |
| lizards | Varanus_pilbarensis | Varanus_pilbarensis | 0 | 107.15 | 3.25 | 31460.33 |
| lizards | Varanus_prasinus | Varanus_prasinus | 0 | 575.44 | 4.35 | 632368.53 |
| lizards | Varanus_primordius | Varanus_primordius | 0 | 26.92 | 3.45 | 102040.89 |
| lizards | Varanus_rosenbergi | Varanus_rosenbergi | 0 | 3019.95 | 8.25 | 413035.44 |
| lizards | Varanus_salvadorii | Varanus_salvadorii | 0 | 15135.61 | 6.8 | 505211.35 |

| | | | | | | |
|---|---|---|---|---|---|---|
| lizards | Varanus_salvator | Varanus_salvator | 0 | 41686.94 | 10.65 | 4677934.79 |
| lizards | Varanus_scalaris | Varanus_scalaris | 0 | 354.81 | 5.85 | 1203625.36 |
| lizards | Varanus_semiremex | Varanus_semiremex | 0 | 426.58 | 5.6 | 202421.38 |
| lizards | Varanus_spenceri | Varanus_spenceri | 0 | 4365.16 | 19.75 | 700991.1 |
| lizards | Varanus_spinulosus | Varanus_spinulosus | 0 | 630.96 | 9.09 | 1594.65 |
| lizards | Varanus_storri | Varanus_storri | 0 | 42.66 | 4.15 | 518428.56 |
| lizards | Varanus_timorensis | Varanus_timorensis | 0 | 436.52 | 7.95 | 28857.86 |
| lizards | Varanus_togianus | Varanus_togianus | 0 | 3981.07 | 12.7 | 184110.68 |
| lizards | Varanus_tristis | Varanus_tristis | 0 | 549.54 | 9.1 | 5950251.61 |
| lizards | Varanus_varius | Varanus_varius | 0 | 10715.19 | 7.55 | 1219347.69 |
| lizards | Varzea_bistriata | Varzea_bistriata | 0 | 28.18 | 4.7 | 8162647.76 |
| lizards | Vhembelacerta_rupicola | Vhembelacerta_rupicola | 0 | 3.24 | 3 | 15435.29 |
| lizards | Voeltzkowia_mira | Voeltzkowia_mira | 1 | 1.48 | 3 | 4891.83 |
| lizards | Woodworthia_brunnea | Woodworthia_brunneus | 1 | 10.23 | 2 | 9177.54 |
| lizards | Woodworthia_chrysosiretica | Woodworthia_chrysosireticus | 0 | 10.23 | 2 | 6515.38 |
| lizards | Woodworthia_maculata | Woodworthia_maculatus | 0 | 14.45 | 1.85 | 7.11 |
| lizards | Xantusia_arizonae | Xantusia_arizonae | 0 | 4.17 | 1.69 | 783.96 |
| lizards | Xantusia_bezyi | Xantusia_bezyi | 0 | 6.76 | 1.6 | 4891.34 |
| lizards | Xantusia_extorris | Xantusia_extorris | 0 | 1.51 | 1.4 | 1353.53 |
| lizards | Xantusia_gracilis | Xantusia_gracilis | 1 | 7.08 | 2 | 44.94 |
| lizards | Xantusia_henshawi | Xantusia_henshawi | 0 | 7.08 | 1.73 | 27942.41 |
| lizards | Xantusia_riversiana | Xantusia_riversiana | 0 | 32.36 | 4.4 | 208.08 |
| lizards | Xantusia_sanchezi | Xantusia_sanchezi | 0 | 3.24 | 3 | 1707.92 |
| lizards | Xantusia_vigilis | Xantusia_vigilis | 0 | 6.76 | 1.84 | 261352.53 |
| lizards | Xenagama_batillifera | Xenagama_batillifera | 0 | 33.88 | 8 | 40067.03 |
| lizards | Xenagama_taylori | Xenagama_taylori | 0 | 20.89 | 5 | 51741.44 |
| lizards | Xenagama_wilmsi | Xenagama_wilmsi | 0 | 30.2 | 6.5 | 154908.41 |
| lizards | Xenosaurus_grandis | Xenosaurus_grandis | 1 | 56.23 | 3.9 | 5386.36 |
| lizards | Xenosaurus_newmanorum | Xenosaurus_newmanorum | 1 | 54.95 | 2.5 | 3894.73 |
| lizards | Xenosaurus_penai | Xenosaurus_penai | 0 | 30.2 | 4 | 107.41 |
| lizards | Xenosaurus_platyceps | Xenosaurus_platyceps | 1 | 42.66 | 2.35 | 4161.09 |
| lizards | Xenosaurus_rectocollaris | Xenosaurus_rectocollaris | 0 | 29.51 | 2.62 | 1263.72 |
| lizards | Zonosaurus_anelanelany | Zonosaurus_anelanelany | 1 | 19.5 | 7 | 6448.39 |
| lizards | Zonosaurus_bemaraha | Zonosaurus_bemaraha | 0 | 10 | 7 | 1604.4 |

| | | | | | | |
|---|---|---|---|---|---|---|
| lizards | Zonosaurus_haraldmeieri | Zonosaurus_haraldmeieri | 0 | 69.18 | 5 | 1956.63 |
| lizards | Zonosaurus_karsteni | Zonosaurus_karsteni | 0 | 58.88 | 4 | 28718.01 |
| lizards | Zonosaurus_rufipes | Zonosaurus_rufipes | 0 | 16.6 | 2.5 | 23315.28 |
| lizards | Zootoca_vivipara | Zootoca_vivipara | 0 | 10.72 | 6.7 | 17595247.88 |
| lizards | Zygaspis_quadrifrons | Zygaspis_quadrifrons | 0 | 8.51 | 3.3 | 3374594.09 |
| lizards | Zygaspis_vandami | Zygaspis_vandami | 0 | 2.75 | 2.5 | 156371.16 |
| mammals | Abeomelomys_sevia | Abeomelomys_sevia | 0 | 52 | 1 | 9683.81 |
| mammals | Abrocoma_bennettii | Abrocoma_bennettii | 0 | 250 | 3.93 | 124895.99 |
| mammals | Abrocoma_cinerea | Abrocoma_cinerea | 0 | 250 | 2.09 | 529152.75 |
| mammals | Abrocoma_schistacea | Abrocoma_shistacea | 0 | 276.3 | 2 | 5496.48 |
| mammals | Abrothrix_andina | Abrothrix_andinus | 0 | 18 | 6 | 612688.89 |
| mammals | Abrothrix_longipilis | Abrothrix_longipilis | 0 | 37.6 | 4.27 | 717949.9 |
| mammals | Abrothrix_olivacea | Abrothrix_olivaceus | 0 | 27 | 5.38 | 1083345.06 |
| mammals | Acerodon_celebensis | Acerodon_celebensis | 1 | 390 | 0.98 | 176922.81 |
| mammals | Acerodon_jubatus | Acerodon_jubatus | 1 | 1075 | 1 | 155342.19 |
| mammals | Acerodon_macklotii | Acerodon_mackloti | 1 | 470.3 | 0.98 | 74622.81 |
| mammals | Acinonyx_jubatus | Acinonyx_jubatus | 1 | 46700 | 3.16 | 32853534.64 |
| mammals | Acomys_cahirinus | Acomys_cahirinus | 0 | 52.5 | 2.45 | 4017285.73 |
| mammals | Acomys_chudeaui | Acomys_airensis | 0 | 48.3 | 2.47 | 1825638.34 |
| mammals | Acomys_cineraceus | Acomys_cineraceus | 0 | 20.3 | 2.47 | 1049545.75 |
| mammals | Acomys_dimidiatus | Acomys_dimidiatus | 0 | 37.5 | 2.47 | 1558726.5 |
| mammals | Acomys_ignitus | Acomys_ignitus | 0 | 24 | 2.47 | 105257.36 |
| mammals | Acomys_johannis | Acomys_johannis | 0 | 18.5 | 2.47 | 426085.27 |
| mammals | Acomys_kempi | Acomys_kempi | 0 | 22 | 2.47 | 569643.34 |
| mammals | Acomys_mullah | Acomys_mullah | 0 | 28.4 | 2.47 | 160420.64 |
| mammals | Acomys_percivali | Acomys_percivali | 0 | 22 | 1.5 | 222958.3 |
| mammals | Acomys_russatus | Acomys_russatus | 0 | 49.8 | 2.5 | 702024.71 |
| mammals | Acomys_seurati | Acomys_seurati | 0 | 71.2 | 2.47 | 298887.04 |
| mammals | Acomys_spinosissimus | Acomys_spinosissimus | 0 | 26.5 | 2.71 | 101250.46 |
| mammals | Acomys_subspinosus | Acomys_subspinosus | 0 | 21 | 3 | 94357.7 |
| mammals | Acomys_wilsoni | Acomys_wilsoni | 0 | 18.5 | 2.29 | 1372812.73 |
| mammals | Aconaemys_fuscus | Aconaemys_fuscus | 0 | 133.5 | 3.23 | 31935.75 |
| mammals | Acrobates_pygmaeus | Acrobates_pygmaeus | 0 | 12 | 2.9 | 1200801.2 |
| mammals | Addax_nasomaculatus | Addax_nasomaculatus | 1 | 70000.3 | 1 | 49488.37 |

| | | | | | | |
|---|---|---|---|---|---|---|
| mammals | Aepeomys_lugens | Aepeomys_lugens | 0 | 37 | 2 | 9497.27 |
| mammals | Aepyceros_melampus | Aepyceros_melampus | 0 | 52500.1 | 0.99 | 3498111.54 |
| mammals | Aepyprymnus_rufescens | Aepyprymnus_rufescens | 0 | 3250 | 1.02 | 566664.64 |
| mammals | Aethalops_aequalis | Aethalops_aequalis | 0 | 11.6 | 1 | 93008.85 |
| mammals | Aethalops_alecto | Aethalops_alecto | 0 | 15 | 0.99 | 79531.47 |
| mammals | Aethomys_chrysophilus | Aethomys_chrysophilus | 0 | 74.9 | 3.22 | 4154330.28 |
| mammals | Aethomys_hindei | Aethomys_hindei | 0 | 103.5 | 2.48 | 2482352.21 |
| mammals | Aethomys_ineptus | Aethomys_ineptus | 0 | 74.9 | 3.35 | 592259.59 |
| mammals | Aethomys_kaiseri | Aethomys_kaiseri | 0 | 90 | 2.95 | 1693329.18 |
| mammals | Aethomys_nyikae | Aethomys_nyikae | 0 | 77 | 3.56 | 897965.41 |
| mammals | Aethomys_thomasi | Aethomys_thomasi | 0 | 107.6 | 2.9 | 270711.03 |
| mammals | Afronycteris_nanus | Neoromicia_nana | 0 | 3.9 | 1.86 | 10443728.67 |
| mammals | Ailuropoda_melanoleuca | Ailuropoda_melanoleuca | 1 | 108400 | 1.58 | 34603.98 |
| mammals | Ailurops_melanotis | Ailurops_melanotis | 1 | 6669 | 1.5 | 274.51 |
| mammals | Ailurops_ursinus | Ailurops_ursinus | 1 | 10000 | 1.5 | 179858.77 |
| mammals | Ailurus_fulgens | Ailurus_fulgens | 1 | 4900 | 1.85 | 338830.99 |
| mammals | Akodon_aerosus | Akodon_aerosus | 0 | 60 | 5 | 114165.46 |
| mammals | Akodon_azarae | Akodon_azarae | 0 | 25 | 4.46 | 1227616 |
| mammals | Akodon_boliviensis | Akodon_boliviensis | 0 | 27.5 | 4.52 | 217533.18 |
| mammals | Akodon_cursor | Akodon_cursor | 0 | 39.9 | 3 | 824145.09 |
| mammals | Akodon_dayi | Akodon_dayi | 0 | 32.5 | 3 | 289331.63 |
| mammals | Akodon_dolores | Akodon_dolores | 0 | 50.5 | 3.89 | 885883.78 |
| mammals | Akodon_iniscatus | Akodon_iniscatus | 0 | 28.7 | 3 | 573427.58 |
| mammals | Akodon_lutescens | Akodon_lutescens | 0 | 20.7 | 4 | 118603.69 |
| mammals | Akodon_mimus | Akodon_mimus | 0 | 24 | 2 | 57439.28 |
| mammals | Akodon_mollis | Akodon_mollis | 0 | 30.4 | 3 | 265351.81 |
| mammals | Akodon_montensis | Akodon_montensis | 0 | 44.1 | 3 | 1050409.37 |
| mammals | Akodon_simulator | Akodon_simulator | 0 | 42.5 | 3 | 156808.59 |
| mammals | Akodon_spegazzinii | Akodon_spegazzinii | 0 | 28.6 | 4.18 | 219496.23 |
| mammals | Akodon_subfuscus | Akodon_subfuscus | 0 | 30.4 | 4.18 | 171092.78 |
| mammals | Akodon_sylvanus | Akodon_sylvanus | 0 | 39 | 3 | 39317.41 |
| mammals | Akodon_toba | Akodon_toba | 0 | 51.2 | 3 | 443710.58 |
| mammals | Alcelaphus_buselaphus | Alcelaphus_buselaphus | 0 | 171001.5 | 1 | 6856359 |
| mammals | Alces_alces | Alces_alces | 0 | 356998 | 1.27 | 23096272.63 |

| | | | | | | |
|---|---|---|---|---|---|---|
| mammals | Alexandromys_fortis | Microtus_fortis | 0 | 63 | 5.26 | 1970776.53 |
| mammals | Alexandromys_kikuchii | Microtus_kikuchii | 0 | 21.7 | 1.97 | 2656.87 |
| mammals | Alexandromys_limnophilus | Microtus_limnophilus | 0 | 19.1 | 5.76 | 933449.66 |
| mammals | Alexandromys_maximowiczii | Microtus_maximowiczii | 0 | 41.7 | 6.8 | 1644298.68 |
| mammals | Alexandromys_middendorffii | Microtus_middendorffii | 0 | 56.6 | 5.61 | 3237474.78 |
| mammals | Alexandromys_mongolicus | Microtus_mongolicus | 0 | 27.5 | 5.1 | 1040262.61 |
| mammals | Alexandromys_montebelli | Microtus_montebelli | 0 | 39.9 | 4.96 | 235353.97 |
| mammals | Alexandromys_oeconomus | Microtus_oeconomus | 0 | 34.4 | 5.69 | 18612352.02 |
| mammals | Alexandromys_sachalinensis | Microtus_sachalinensis | 0 | 54.1 | 7.89 | 45125.98 |
| mammals | Alionycteris_paucidentata | Alionycteris_paucidentata | 0 | 16.2 | 1 | 6417.84 |
| mammals | Allactaga_major | Allactaga_major | 0 | 350 | 3.62 | 4297326.14 |
| mammals | Allactaga_severtzovi | Allactaga_severtzovi | 0 | 159 | 3.93 | 706422.24 |
| mammals | Allactodipus_bobrinskii | Allactodipus_bobrinskii | 0 | 70 | 5.15 | 275856.83 |
| mammals | Allenopithecus_nigroviridis | Allenopithecus_nigroviridis | 0 | 4750 | 1 | 526623 |
| mammals | Allocebus_trichotis | Allocebus_trichotis | 1 | 77.5 | 1 | 7303.54 |
| mammals | Allochrocebus_lhoesti | Allochrocebus_lhoesti | 1 | 5375 | 1 | 216021.54 |
| mammals | Allochrocebus_preussi | Allochrocebus_preussi | 1 | 5250 | 1.01 | 17526.76 |
| mammals | Allochrocebus_solatus | Allochrocebus_solatus | 0 | 6500 | 1.01 | 18941.85 |
| mammals | Allocricetulus_curtatus | Allocricetulus_curtatus | 0 | 47.8 | 5.19 | 1332299.37 |
| mammals | Allocricetulus_eversmanni | Allocricetulus_eversmanni | 0 | 115.6 | 4.99 | 1776745.67 |
| mammals | Alouatta_arctoidea | Alouatta_arctoidea | 0 | 5390 | 1.31 | 406536.15 |
| mammals | Alouatta_belzebul | Alouatta_belzebul | 1 | 6400 | 1 | 864468.57 |
| mammals | Alouatta_caraya | Alouatta_caraya | 0 | 5862.5 | 1 | 3064214.8 |
| mammals | Alouatta_discolor | Alouatta_discolor | 1 | 6262.8 | 1 | 375751.38 |
| mammals | Alouatta_guariba | Alouatta_guariba | 1 | 5589.2 | 1 | 940355.9 |
| mammals | Alouatta_macconnelli | Alouatta_macconnelli | 0 | 7554.8 | 1.2 | 1755234.56 |
| mammals | Alouatta_palliata | Alouatta_palliata | 1 | 7275 | 1.01 | 549356.89 |
| mammals | Alouatta_pigra | Alouatta_pigra | 1 | 7000 | 1.01 | 246704.22 |
| mammals | Alouatta_sara | Alouatta_sara | 0 | 6611 | 1.2 | 341403.37 |
| mammals | Alticola_argentatus | Alticola_argentatus | 0 | 37.7 | 4.5 | 1380073.11 |
| mammals | Alticola_lemminus | Alticola_lemminus | 0 | 15.5 | 4.75 | 2068169.33 |
| mammals | Alticola_macrotis | Alticola_macrotis | 0 | 36 | 5.5 | 379222.68 |
| mammals | Alticola_montosus | Alticola_montosa | 1 | 19.7 | 4.87 | 19533.93 |
| mammals | Alticola_olchonensis | Alticola_olchonensis | 1 | 33.5 | 4.43 | 1348.37 |

| | | | | | | |
|---|---|---|---|---|---|---|
| mammals | Alticola_roylei | Alticola_roylei | 0 | 37 | 5.16 | 38435.41 |
| mammals | Alticola_semicanus | Alticola_semicanus | 0 | 53.2 | 4.87 | 671213.97 |
| mammals | Alticola_stoliczkanus | Alticola_stoliczkanus | 0 | 27.5 | 7.77 | 2306359.63 |
| mammals | Alticola_strelzovi | Alticola_strelzowi | 0 | 50 | 7.1 | 441285.29 |
| mammals | Alticola_tuvinicus | Alticola_tuvinicus | 0 | 41 | 4.5 | 134541.19 |
| mammals | Ametrida_centurio | Ametrida_centurio | 0 | 10.3 | 0.99 | 5766469.12 |
| mammals | Ammodorcas_clarkei | Ammodorcas_clarkei | 1 | 28049.8 | 1 | 93566.48 |
| mammals | Ammospermophilus_harrisii | Ammospermophilus_harrisii | 0 | 122 | 6.49 | 269488.41 |
| mammals | Ammospermophilus_interpres | Ammospermophilus_interpres | 0 | 110.3 | 9.43 | 312449.22 |
| mammals | Ammospermophilus_leucurus | Ammospermophilus_leucurus | 0 | 120.5 | 8 | 990844.1 |
| mammals | Ammospermophilus_nelsoni | Ammospermophilus_nelsoni | 1 | 160.4 | 8.9 | 16430.67 |
| mammals | Ammotragus_lervia | Ammotragus_lervia | 1 | 48000 | 1.22 | 1167864.69 |
| mammals | Amorphochilus_schnablii | Amorphochilus_schnablii | 1 | 3.3 | 0.99 | 178408.34 |
| mammals | Anathana_ellioti | Anathana_ellioti | 0 | 160 | 4.84 | 1366073.82 |
| mammals | Andalgalomys_olrogi | Andalgalomys_olrogi | 0 | 32.9 | 5 | 57114.93 |
| mammals | Andinomys_edax | Andinomys_edax | 0 | 69.7 | 2.91 | 228613.14 |
| mammals | Anomalurus_beecrofti | Anomalurus_beecrofti | 0 | 389 | 0.98 | 3245229.52 |
| mammals | Anomalurus_derbianus | Anomalurus_derbianus | 0 | 650 | 1.08 | 5752727.17 |
| mammals | Anomalurus_pelii | Anomalurus_pelii | 0 | 1770 | 2.5 | 104894.75 |
| mammals | Anotomys_leander | Anotomys_leander | 1 | 66.4 | 0.97 | 12508.22 |
| mammals | Anoura_caudifer | Anoura_caudifer | 0 | 10.1 | 0.99 | 5940468.24 |
| mammals | Anoura_cultrata | Anoura_cultrata | 0 | 16.9 | 1.22 | 757500.04 |
| mammals | Anoura_geoffroyi | Anoura_geoffroyi | 0 | 14.6 | 1 | 8203646.83 |
| mammals | Anourosorex_assamensis | Anourosorex_assamensis | 0 | 25.2 | 2.82 | 17603.52 |
| mammals | Anourosorex_squamipes | Anourosorex_squamipes | 0 | 20 | 2.82 | 1189018.89 |
| mammals | Anourosorex_yamashinai | Anourosorex_yamashinai | 0 | 11.4 | 2.82 | 34157.02 |
| mammals | Antechinomys_laniger | Antechinomys_laniger | 0 | 25 | 6 | 2438423.42 |
| mammals | Antechinomys_longicaudatus | Sminthopsis_longicaudata | 0 | 18 | 3.93 | 747801.2 |
| mammals | Antechinus_adustus | Antechinus_adustus | 0 | 28.2 | 6 | 4920.27 |
| mammals | Antechinus_agilis | Antechinus_agilis | 0 | 24 | 8.75 | 163654.97 |
| mammals | Antechinus_bellus | Antechinus_bellus | 1 | 44.5 | 10.08 | 20577.83 |
| mammals | Antechinus_flavipes | Antechinus_flavipes | 0 | 45 | 9.27 | 1008801.55 |
| mammals | Antechinus_godmani | Antechinus_godmani | 0 | 76.5 | 5.29 | 3599.52 |
| mammals | Antechinus_leo | Antechinus_leo | 0 | 74 | 8.64 | 4393.22 |

| | | | | | | |
|---|---|---|---|---|---|---|
| mammals | Antechinus_minimus | Antechinus_minimus | 0 | 53.5 | 7.19 | 76275.71 |
| mammals | Antechinus_stuartii | Antechinus_stuartii | 0 | 27.5 | 6.92 | 118336.85 |
| mammals | Antechinus_subtropicus | Antechinus_subtropicus | 0 | 33.1 | 7.5 | 9618.05 |
| mammals | Antechinus_swainsonii | Antechinus_swainsonii | 0 | 53 | 8.18 | 314283.03 |
| mammals | Anthops_ornatus | Anthops_ornatus | 1 | 8 | 1 | 21294.85 |
| mammals | Antidorcas_marsupialis | Antidorcas_marsupialis | 0 | 31500 | 1 | 1950012.85 |
| mammals | Antilocapra_americana | Antilocapra_americana | 0 | 46082.9 | 1.91 | 2007736.6 |
| mammals | Antilope_cervicapra | Antilope_cervicapra | 0 | 37500 | 1.11 | 1754050.91 |
| mammals | Antrozous_pallidus | Antrozous_pallidus | 0 | 20.8 | 1.89 | 4367598.17 |
| mammals | Aotus_azarae | Aotus_azarae | 0 | 963 | 1 | 3156514.71 |
| mammals | Aotus_brumbacki | Aotus_brumbacki | 1 | 800 | 1 | 123963.78 |
| mammals | Aotus_griseimembra | Aotus_griseimembra | 1 | 966 | 1 | 234087.45 |
| mammals | Aotus_lemurinus | Aotus_lemurinus | 1 | 873 | 1 | 177191.55 |
| mammals | Aotus_miconax | Aotus_miconax | 1 | 800 | 1 | 27618.65 |
| mammals | Aotus_nancymai | Aotus_nancymaae | 1 | 788 | 1.02 | 509597.72 |
| mammals | Aotus_nigriceps | Aotus_nigriceps | 0 | 1060 | 1 | 1569879.02 |
| mammals | Aotus_trivirgatus | Aotus_trivirgatus | 0 | 900 | 1.06 | 752044.96 |
| mammals | Aotus_vociferans | Aotus_vociferans | 0 | 873 | 1.02 | 1054332.87 |
| mammals | Aotus_zonalis | Aotus_zonalis | 0 | 902.5 | 1 | 131878.68 |
| mammals | Aplodontia_rufa | Aplodontia_rufa | 0 | 1004 | 2.63 | 245471.87 |
| mammals | Apodemus_agrarius | Apodemus_agrarius | 0 | 23.3 | 5.66 | 11531272.51 |
| mammals | Apodemus_argenteus | Apodemus_argenteus | 0 | 21 | 5.16 | 366528.07 |
| mammals | Apodemus_chevrieri | Apodemus_chevrieri | 0 | 32.9 | 5.68 | 720401.61 |
| mammals | Apodemus_draco | Apodemus_draco | 0 | 22.3 | 5.18 | 1798566.22 |
| mammals | Apodemus_epimelas | Apodemus_epimelas | 0 | 42 | 3.36 | 174107.25 |
| mammals | Apodemus_latronum | Apodemus_latronum | 0 | 35.2 | 5.22 | 536889.58 |
| mammals | Apodemus_mystacinus | Apodemus_mystacinus | 0 | 45.6 | 3.62 | 804373.39 |
| mammals | Apodemus_peninsulae | Apodemus_peninsulae | 0 | 32.9 | 4.56 | 8904776.97 |
| mammals | Apodemus_rusiges | Apodemus_rusiges | 0 | 37 | 5.19 | 112760.37 |
| mammals | Apodemus_semotus | Apodemus_semotus | 0 | 28.8 | 4.36 | 33914.9 |
| mammals | Apodemus_speciosus | Apodemus_speciosus | 0 | 33.8 | 4.47 | 368742.87 |
| mammals | Apodemus_witherbyi | Apodemus_witherbyi | 0 | 30 | 4.5 | 1581288.62 |
| mammals | Apomys_abrae | Apomys_abrae | 0 | 56.6 | 2 | 12463.63 |
| mammals | Apomys_datae | Apomys_datae | 0 | 106 | 1.66 | 11591.32 |

| | | | | | | |
|---|---|---|---|---|---|---|
| mammals | Apomys_gracilirostris | Apomys_gracilirostris | 0 | 88.1 | 3 | 571.34 |
| mammals | Apomys_microdon | Apomys_microdon | 0 | 35.1 | 2 | 61956.39 |
| mammals | Apomys_musculus | Apomys_musculus | 0 | 21.2 | 2 | 47774.03 |
| mammals | Apomys_sacobianus | Apomys_sacobianus | 0 | 82.4 | 2 | 951.48 |
| mammals | Aproteles_bulmerae | Aproteles_bulmerae | 1 | 600 | 1 | 48.16 |
| mammals | Arabitragus_jayakari | Arabitragus_jayakari | 1 | 23000 | 1 | 28012.02 |
| mammals | Arborimus_albipes | Arborimus_albipes | 0 | 23 | 2.75 | 77753.37 |
| mammals | Arborimus_longicaudus | Arborimus_longicaudus | 0 | 21.8 | 2.83 | 54110.77 |
| mammals | Arborimus_pomo | Arborimus_pomo | 0 | 32.5 | 1.88 | 22003.98 |
| mammals | Archboldomys_luzonensis | Archboldomys_luzonensis | 1 | 35.4 | 1 | 22.21 |
| mammals | Arctictis_binturong | Arctictis_binturong | 1 | 9875 | 2.5 | 3049767.96 |
| mammals | Arctocebus_aureus | Arctocebus_aureus | 0 | 235 | 1 | 667484.14 |
| mammals | Arctocebus_calabarensis | Arctocebus_calabarensis | 0 | 210 | 1 | 148440.25 |
| mammals | Arctogalidia_trivirgata | Arctogalidia_trivirgata | 0 | 2250 | 2.64 | 2539704.36 |
| mammals | Arctonyx_albogularis | Arctonyx_albogularis | 0 | 9000 | 3 | 4210609.6 |
| mammals | Arctonyx_collaris | Arctonyx_collaris | 1 | 10.5 | 3 | 1878579.03 |
| mammals | Arctonyx_hoevenii | Arctonyx_hoevenii | 0 | 7000 | 3 | 98102.53 |
| mammals | Ardops_nichollsi | Ardops_nichollsi | 0 | 17.2 | 0.99 | 4742.71 |
| mammals | Arielulus_circumdatus | Arielulus_circumdatus | 0 | 10.4 | 1 | 170888.04 |
| mammals | Artibeus_concolor | Artibeus_concolor | 0 | 19.4 | 1 | 6437731.25 |
| mammals | Artibeus_fimbriatus | Artibeus_fimbriatus | 0 | 63.9 | 1 | 1480430.38 |
| mammals | Artibeus_fraterculus | Artibeus_fraterculus | 0 | 39.5 | 0.99 | 302070.73 |
| mammals | Artibeus_hirsutus | Artibeus_hirsutus | 0 | 38.7 | 0.98 | 228810.6 |
| mammals | Artibeus_jamaicensis | Artibeus_jamaicensis | 0 | 42.2 | 1 | 1559896.27 |
| mammals | Artibeus_lituratus | Artibeus_lituratus | 0 | 60.5 | 1 | 14235952.08 |
| mammals | Artibeus_obscurus | Artibeus_obscurus | 0 | 35.2 | 0.99 | 11122956.74 |
| mammals | Artibeus_planirostris | Artibeus_planirostris | 0 | 47 | 1 | 7906983.19 |
| mammals | Arvicanthis_abyssinicus | Arvicanthis_abyssinicus | 0 | 77.1 | 5.6 | 296092.69 |
| mammals | Arvicanthis_ansorgei | Arvicanthis_ansorgei | 0 | 124.5 | 5 | 651274.69 |
| mammals | Arvicanthis_blicki | Arvicanthis_blicki | 0 | 128 | 3.2 | 17893.75 |
| mammals | Arvicanthis_nairobae | Arvicanthis_nairobae | 0 | 104 | 5.5 | 151138.94 |
| mammals | Arvicanthis_niloticus | Arvicanthis_niloticus | 0 | 106.9 | 4.93 | 6185272.89 |
| mammals | Arvicanthis_rufinus | Arvicanthis_rufinus | 0 | 144.1 | 4.86 | 1154420.17 |
| mammals | Arvicola_amphibius | Arvicola_amphibius | 0 | 225 | 4.92 | 15657066.94 |

| | | | | | | |
|---|---|---|---|---|---|---|
| mammals | Arvicola_sapidus | Arvicola_sapidus | 1 | 220 | 4.12 | 882587.18 |
| mammals | Asellia_patrizii | Asellia_patrizii | 0 | 6 | 1 | 227346.99 |
| mammals | Asellia_tridens | Asellia_tridens | 0 | 11.9 | 1 | 16925970.62 |
| mammals | Aselliscus_stoliczkanus | Aselliscus_stoliczkanus | 0 | 4.9 | 1 | 1218727.09 |
| mammals | Aselliscus_tricuspidatus | Aselliscus_tricuspidatus | 0 | 4.1 | 0.99 | 274415.14 |
| mammals | Atelerix_albiventris | Atelerix_albiventris | 0 | 600 | 4.24 | 7028364.29 |
| mammals | Atelerix_algirus | Atelerix_algirus | 0 | 1250 | 3.12 | 734429.78 |
| mammals | Atelerix_frontalis | Atelerix_frontalis | 0 | 309.5 | 5.09 | 1047880.25 |
| mammals | Atelerix_sclateri | Atelerix_sclateri | 0 | 436.5 | 4.49 | 309001.28 |
| mammals | Ateles_belzebuth | Ateles_belzebuth | 1 | 5000 | 1.01 | 815705.06 |
| mammals | Ateles_chamek | Ateles_chamek | 1 | 6000 | 1 | 2602958.43 |
| mammals | Ateles_fusciceps | Ateles_fusciceps | 1 | 9100 | 1 | 212235.69 |
| mammals | Ateles_geoffroyi | Ateles_geoffroyi | 1 | 5284.9 | 1 | 901722.85 |
| mammals | Ateles_hybridus | Ateles_hybridus | 1 | 8675 | 1 | 165736.6 |
| mammals | Ateles_marginatus | Ateles_marginatus | 1 | 6000 | 1 | 524099.56 |
| mammals | Ateles_paniscus | Ateles_paniscus | 1 | 7900.1 | 1 | 918126.83 |
| mammals | Atelocynus_microtis | Atelocynus_microtis | 0 | 7750 | 2.3 | 2852893.51 |
| mammals | Atherurus_africanus | Atherurus_africanus | 0 | 1934 | 1.5 | 3544244.62 |
| mammals | Atherurus_macrourus | Atherurus_macrourus | 0 | 2000 | 1.54 | 2672201.97 |
| mammals | Atilax_paludinosus | Atilax_paludinosus | 0 | 3300 | 2 | 14049574.29 |
| mammals | Atopogale_cubana | Solenodon_cubanus | 1 | 650 | 1.5 | 1996.73 |
| mammals | Auliscomys_boliviensis | Auliscomys_boliviensis | 0 | 65.7 | 3.4 | 210614.07 |
| mammals | Auliscomys_pictus | Auliscomys_pictus | 0 | 54.7 | 4.5 | 309598.29 |
| mammals | Auliscomys_sublimis | Auliscomys_sublimis | 0 | 38 | 5 | 393512.53 |
| mammals | Austronomus_australis | Austronomus_australis | 0 | 31.8 | 1 | 6025987.53 |
| mammals | Austronomus_kuboriensis | Austronomus_kuboriensis | 0 | 25.9 | 1 | 107966.74 |
| mammals | Avahi_laniger | Avahi_laniger | 1 | 1300 | 1.01 | 51921.14 |
| mammals | Avahi_occidentalis | Avahi_occidentalis | 1 | 850 | 1.01 | 3050.16 |
| mammals | Axis_axis | Axis_axis | 0 | 66172.8 | 1.02 | 2361371.08 |
| mammals | Axis_calamianensis | Axis_calamianensis | 1 | 40500 | 1.5 | 1345.32 |
| mammals | Axis_kuhlii | Axis_kuhlii | 1 | 55000 | 1.11 | 63.8 |
| mammals | Axis_porcinus | Axis_porcinus | 1 | 34127.8 | 1 | 362934.43 |
| mammals | Babyrousa_babyrussa | Babyrousa_babyrussa | 1 | 100000 | 1.56 | 13191.94 |
| mammals | Babyrousa_celebensis | Babyrousa_celebensis | 1 | 84327.5 | 1.4 | 135977.78 |

| | | | | | | |
|---|---|---|---|---|---|---|
| mammals | Babyrousa_togeanensis | Babyrousa_togeanensis | 1 | 113762.8 | 1.4 | 533.08 |
| mammals | Baiomys_musculus | Baiomys_musculus | 0 | 9 | 2.62 | 450251.28 |
| mammals | Baiomys_taylori | Baiomys_taylori | 0 | 8 | 2.6 | 1275181.16 |
| mammals | Balantiopteryx_infusca | Balantiopteryx_infusca | 1 | 4.4 | 1 | 2841.7 |
| mammals | Balantiopteryx_io | Balantiopteryx_io | 1 | 4.3 | 1 | 170568.45 |
| mammals | Balantiopteryx_plicata | Balantiopteryx_plicata | 0 | 6.4 | 0.99 | 825040.69 |
| mammals | Balionycteris_maculata | Balionycteris_maculata | 0 | 13.4 | 1 | 439669.6 |
| mammals | Bandicota_bengalensis | Bandicota_bengalensis | 0 | 188.5 | 7.46 | 4165486.08 |
| mammals | Barbastella_barbastellus | Barbastella_barbastellus | 0 | 8 | 1.86 | 3811256.04 |
| mammals | Barbastella_leucomelas | Barbastella_leucomelas | 0 | 15.1 | 2 | 3928085.94 |
| mammals | Bassaricyon_alleni | Bassaricyon_alleni | 0 | 1235 | 1 | 4449618.02 |
| mammals | Bassaricyon_gabbii | Bassaricyon_gabbii | 0 | 1250 | 1 | 156420.12 |
| mammals | Bassaricyon_medius | Bassaricyon_medius | 0 | 1076 | 1 | 181889.99 |
| mammals | Bassaricyon_neblina | Bassaricyon_neblina | 0 | 872 | 1 | 120399.36 |
| mammals | Bassariscus_astutus | Bassariscus_astutus | 0 | 1129.5 | 3 | 3903858.56 |
| mammals | Bassariscus_sumichrasti | Bassariscus_sumichrasti | 0 | 900 | 2 | 622168.55 |
| mammals | Batomys_salomonseni | Batomys_salomonseni | 0 | 187 | 1.5 | 16015.51 |
| mammals | Bauerus_dubiaquercus | Bauerus_dubiaquercus | 0 | 22.9 | 1.71 | 265235.24 |
| mammals | Bdeogale_crassicauda | Bdeogale_crassicauda | 0 | 1550 | 0.97 | 1497508.74 |
| mammals | Bdeogale_jacksoni | Bdeogale_jacksoni | 0 | 2500 | 1 | 63953.37 |
| mammals | Bdeogale_nigripes | Bdeogale_nigripes | 0 | 2500 | 1 | 2110590.78 |
| mammals | Bdeogale_omnivora | Bdeogale_omnivora | 1 | 1750 | 0.97 | 28493 |
| mammals | Beamys_hindei | Beamys_hindei | 0 | 75.8 | 3.68 | 337356.6 |
| mammals | Beatragus_hunteri | Beatragus_hunteri | 1 | 80000 | 1 | 11330.99 |
| mammals | Berylmys_bowersii | Berylmys_bowersi | 0 | 300 | 3.88 | 2558279.08 |
| mammals | Bettongia_gaimardi | Bettongia_gaimardi | 0 | 1660 | 1 | 29089.49 |
| mammals | Bettongia_lesueur | Bettongia_lesueur | 0 | 1500 | 1 | 979.82 |
| mammals | Bettongia_penicillata | Bettongia_penicillata | 1 | 1300 | 1 | 4058.27 |
| mammals | Bettongia_tropica | Bettongia_tropica | 1 | 1200 | 1 | 396.78 |
| mammals | Blarina_brevicauda | Blarina_brevicauda | 0 | 28 | 5.56 | 4106919.94 |
| mammals | Blarina_carolinensis | Blarina_carolinensis | 0 | 13.5 | 4.04 | 866209.84 |
| mammals | Blarina_hylophaga | Blarina_hylophaga | 0 | 14.5 | 5 | 727831.72 |
| mammals | Blastocerus_dichotomus | Blastocerus_dichotomus | 1 | 86666.2 | 0.99 | 4234152.83 |
| mammals | Boneia_bidens | Rousettus_bidens | 1 | 171 | 0.98 | 154150.12 |

| | | | | | | |
|---|---|---|---|---|---|---|
| mammals | Bos_bison | Bison_bison | 0 | 579255.3 | 0.99 | 140673.08 |
| mammals | Bos_bonasus | Bison_bonasus | 0 | 500000 | 1 | 24303.16 |
| mammals | Bos_gaurus | Bos_gaurus | 1 | 825000 | 1.2 | 1370086.78 |
| mammals | Bos_javanicus | Bos_javanicus | 1 | 700000 | 1.22 | 183622.81 |
| mammals | Bos_mutus | Bos_mutus | 1 | 546250 | 1 | 469488.49 |
| mammals | Boselaphus_tragocamelus | Boselaphus_tragocamelus | 0 | 169000 | 1.45 | 2289610.81 |
| mammals | Brachyphylla_cavernarum | Brachyphylla_cavernarum | 0 | 46.2 | 1 | 14907.7 |
| mammals | Brachyphylla_nana | Brachyphylla_nana | 0 | 35.8 | 0.99 | 195038.68 |
| mammals | Brachytarsomys_albicauda | Brachytarsomys_albicauda | 0 | 200 | 6 | 64739.8 |
| mammals | Brachytarsomys_villosus | Brachytarsomys_villosa | 1 | 220 | 6 | 10127.47 |
| mammals | Brachyteles_arachnoides | Brachyteles_arachnoides | 1 | 13499.9 | 1.01 | 73233.96 |
| mammals | Brachyteles_hypoxanthus | Brachyteles_hypoxanthus | 1 | 8865 | 1.01 | 96658.15 |
| mammals | Brachyuromys_betsileoensis | Brachyuromys_betsileoensis | 0 | 92.5 | 2 | 42770.18 |
| mammals | Brachyuromys_ramirohitra | Brachyuromys_ramirohitra | 0 | 90 | 2 | 68444.56 |
| mammals | Bradypus_torquatus | Bradypus_torquatus | 1 | 3900 | 1 | 80086.16 |
| mammals | Bradypus_tridactylus | Bradypus_tridactylus | 0 | 3850 | 1 | 1043271.32 |
| mammals | Bradypus_variegatus | Bradypus_variegatus | 0 | 4335 | 1 | 10100677.31 |
| mammals | Bubalus_arnee | Bubalus_arnee | 1 | 725000 | 1.44 | 13690.35 |
| mammals | Bubalus_depressicornis | Bubalus_depressicornis | 1 | 88000 | 1 | 107246.62 |
| mammals | Bubalus_mindorensis | Bubalus_mindorensis | 1 | 224000 | 1 | 98.75 |
| mammals | Bubalus_quarlesi | Bubalus_quarlesi | 1 | 76000 | 1 | 70025.75 |
| mammals | Budorcas_taxicolor | Budorcas_taxicolor | 1 | 302000 | 1.2 | 440420.96 |
| mammals | Bullimus_gamay | Bullimus_gamay | 1 | 368 | 2 | 252.92 |
| mammals | Bullimus_luzonicus | Bullimus_luzonicus | 0 | 445 | 2 | 38308.97 |
| mammals | Bunolagus_monticularis | Bunolagus_monticularis | 1 | 1750 | 1.17 | 45473.93 |
| mammals | Bunomys_chrysocomus | Bunomys_chrysocomus | 0 | 116 | 2 | 169337.75 |
| mammals | Burramys_parvus | Burramys_parvus | 1 | 41.5 | 3.58 | 1197.79 |
| mammals | Cabassous_chacoensis | Cabassous_chacoensis | 0 | 1490 | 1 | 478652.23 |
| mammals | Cabassous_tatouay | Cabassous_tatouay | 0 | 5350 | 1 | 2538377.79 |
| mammals | Cabassous_unicinctus | Cabassous_unicinctus | 0 | 4800 | 1 | 9761562.72 |
| mammals | Cabreramops_aequatorianus | Molossops_aequatorianus | 1 | 5 | 1.05 | 3326.78 |
| mammals | Cacajao_calvus | Cacajao_calvus | 1 | 5796 | 1 | 210952.49 |
| mammals | Cacajao_hosomi | Cacajao_hosomi | 1 | 3100 | 1 | 69745.45 |
| mammals | Caenolestes_caniventer | Caenolestes_caniventer | 0 | 40 | 2 | 19664.02 |

| | | | | | | |
|---|---|---|---|---|---|---|
| mammals | Caenolestes_fuliginosus | Caenolestes_fuliginosus | 0 | 27.8 | 3.71 | 180934.66 |
| mammals | Callicebus_barbarabrownae | Callicebus_barbarabrownae | 1 | 1228.7 | 1.01 | 129698.42 |
| mammals | Callicebus_coimbrai | Callicebus_coimbrai | 1 | 1181.7 | 1.01 | 38585.35 |
| mammals | Callicebus_melanochir | Callicebus_melanochir | 1 | 1322.4 | 1.01 | 99867.07 |
| mammals | Callicebus_nigrifrons | Callicebus_nigrifrons | 0 | 1215.1 | 1.01 | 490584.71 |
| mammals | Callicebus_personatus | Callicebus_personatus | 1 | 1350 | 1.01 | 142975.24 |
| mammals | Callimico_goeldii | Callimico_goeldii | 1 | 480 | 1.02 | 602352.68 |
| mammals | Callithrix_aurita | Callithrix_aurita | 1 | 342 | 2.12 | 159614.29 |
| mammals | Callithrix_flaviceps | Callithrix_flaviceps | 1 | 342 | 2.06 | 24732.63 |
| mammals | Callithrix_geoffroyi | Callithrix_geoffroyi | 0 | 342 | 2.24 | 124967.93 |
| mammals | Callithrix_jacchus | Callithrix_jacchus | 0 | 292 | 2.28 | 943369.79 |
| mammals | Callithrix_kuhlii | Callithrix_kuhlii | 1 | 342 | 2 | 45700.72 |
| mammals | Callithrix_penicillata | Callithrix_penicillata | 0 | 342 | 2 | 1309808.49 |
| mammals | Callosciurus_adamsi | Callosciurus_adamsi | 0 | 134.5 | 1.96 | 97681.77 |
| mammals | Callosciurus_baluensis | Callosciurus_baluensis | 0 | 356 | 3 | 51137.13 |
| mammals | Callosciurus_caniceps | Callosciurus_caniceps | 0 | 257 | 2.19 | 650673.88 |
| mammals | Callosciurus_erythraeus | Callosciurus_erythraeus | 0 | 280 | 2.05 | 4042787.1 |
| mammals | Callosciurus_finlaysonii | Callosciurus_finlaysonii | 0 | 278 | 2 | 730177.83 |
| mammals | Callosciurus_nigrovittatus | Callosciurus_nigrovittatus | 0 | 202 | 2.28 | 708578.85 |
| mammals | Callosciurus_notatus | Callosciurus_notatus | 0 | 197.5 | 2.35 | 1496326.22 |
| mammals | Callosciurus_orestes | Callosciurus_orestes | 0 | 278 | 2.64 | 50506.52 |
| mammals | Callosciurus_prevostii | Callosciurus_prevostii | 0 | 361.5 | 1.53 | 1334755.19 |
| mammals | Callosciurus_pygerythrus | Callosciurus_pygerythrus | 0 | 252 | 3.5 | 587628.61 |
| mammals | Callosciurus_quinquestriatus | Callosciurus_quinquestriatus | 0 | 318.6 | 1 | 40861 |
| mammals | Callospermophilus_lateralis | Callospermophilus_lateralis | 0 | 191 | 5.15 | 1704582.98 |
| mammals | Callospermophilus_madrensis | Callospermophilus_madrensis | 0 | 275 | 4.73 | 60013.58 |
| mammals | Callospermophilus_saturatus | Callospermophilus_saturatus | 0 | 220 | 5.12 | 74904.17 |
| mammals | Calomys_boliviae | Calomys_boliviae | 0 | 27 | 4.98 | 129638.12 |
| mammals | Calomys_callidus | Calomys_callidus | 0 | 27 | 4.98 | 55966.07 |
| mammals | Calomys_callosus | Calomys_callosus | 0 | 48 | 4.95 | 3145757.29 |
| mammals | Calomys_expulsus | Calomys_expulsus | 0 | 28.2 | 4.98 | 1360452.92 |
| mammals | Calomys_hummelincki | Calomys_hummelincki | 0 | 27 | 5 | 312459.19 |
| mammals | Calomys_laucha | Calomys_laucha | 0 | 14 | 5.14 | 1437875.39 |
| mammals | Calomys_lepidus | Calomys_lepidus | 0 | 26.6 | 4.26 | 584231.18 |

| | | | | | | |
|---|---|---|---|---|---|---|
| mammals | Calomys_musculinus | Calomys_musculinus | 0 | 20.1 | 5.81 | 2024366.28 |
| mammals | Calomys_sorella | Calomys_sorellus | 0 | 20 | 4.22 | 244104.41 |
| mammals | Calomys_tener | Calomys_tener | 0 | 13.8 | 4.97 | 1710353.17 |
| mammals | Calomys_tocantinsi | Calomys_tocantinsi | 0 | 25.3 | 7.3 | 132795.31 |
| mammals | Calomys_venustus | Calomys_venustus | 0 | 50.7 | 4.98 | 58306.81 |
| mammals | Calomyscus_bailwardi | Calomyscus_bailwardi | 0 | 20.4 | 3.66 | 185798.11 |
| mammals | Calomyscus_baluchi | Calomyscus_baluchi | 0 | 20.1 | 3.33 | 426505.71 |
| mammals | Calomyscus_elburzensis | Calomyscus_elburzensis | 0 | 17 | 3.66 | 231907.31 |
| mammals | Calomyscus_hotsoni | Calomyscus_hotsoni | 0 | 16.3 | 3.33 | 141022.26 |
| mammals | Calomyscus_mystax | Calomyscus_mystax | 0 | 17 | 4.33 | 208687.98 |
| mammals | Calomyscus_urartensis | Calomyscus_urartensis | 0 | 16.5 | 3.33 | 4812.86 |
| mammals | Caluromys_derbianus | Caluromys_derbianus | 0 | 297 | 3.1 | 792215.26 |
| mammals | Caluromys_lanatus | Caluromys_lanatus | 0 | 325 | 2.5 | 5110003.56 |
| mammals | Caluromys_philander | Caluromys_philander | 0 | 229.3 | 4.26 | 3711568.69 |
| mammals | Caluromysiops_irruptus | Caluromysiops_irrupta | 0 | 250 | 2 | 182315.31 |
| mammals | Camelus_ferus | Camelus_ferus | 1 | 690000 | 1.2 | 202139.12 |
| mammals | Canis_aureus | Canis_aureus | 0 | 10345.2 | 3.85 | 10746781.91 |
| mammals | Canis_latrans | Canis_latrans | 0 | 13406.3 | 5.66 | 14921849.18 |
| mammals | Canis_lupus | Canis_lupus | 0 | 32183.3 | 4.98 | 56006225.87 |
| mammals | Canis_rufus | Canis_rufus | 1 | 22983.2 | 4.98 | 3120.73 |
| mammals | Canis_simensis | Canis_simensis | 1 | 10000 | 4 | 6913.95 |
| mammals | Cannomys_badius | Cannomys_badius | 0 | 275 | 1.95 | 1043848.45 |
| mammals | Cansumys_canus | Cansumys_canus | 0 | 90.5 | 4.24 | 161030.75 |
| mammals | Capra_aegagrus | Capra_aegagrus | 0 | 33500 | 1.48 | 1215040.11 |
| mammals | Capra_caucasica | Capra_caucasica | 1 | 55000 | 1.09 | 7868.96 |
| mammals | Capra_cylindricornis | Capra_cylindricornis | 0 | 50000 | 1.09 | 22805.65 |
| mammals | Capra_falconeri | Capra_falconeri | 0 | 41000 | 1.86 | 25328.49 |
| mammals | Capra_ibex | Capra_ibex | 0 | 85166.5 | 1.15 | 19233.35 |
| mammals | Capra_nubiana | Capra_nubiana | 1 | 62624.8 | 1.05 | 342497.98 |
| mammals | Capra_pyrenaica | Capra_pyrenaica | 0 | 50000 | 1.5 | 92676.02 |
| mammals | Capra_sibirica | Capra_sibirica | 0 | 130000 | 1.18 | 1562755.82 |
| mammals | Capra_walie | Capra_walie | 1 | 100000 | 1.05 | 844.92 |
| mammals | Capreolus_capreolus | Capreolus_capreolus | 0 | 22500 | 1.79 | 6468697.04 |
| mammals | Capreolus_pygargus | Capreolus_pygargus | 0 | 43752.2 | 1.95 | 11998106.58 |

| | | | | | | |
|---|---|---|---|---|---|---|
| mammals | Capricornis_crispus | Capricornis_crispus | 0 | 37160 | 1 | 160081 |
| mammals | Capricornis_rubidus | Capricornis_rubidus | 1 | 87500 | 1 | 86336.31 |
| mammals | Capricornis_sumatraensis | Capricornis_sumatraensis | 1 | 87500 | 1.11 | 3511835.45 |
| mammals | Capricornis_swinhoei | Capricornis_swinhoei | 0 | 21000 | 1 | 14411.9 |
| mammals | Caprolagus_hispidus | Caprolagus_hispidus | 1 | 2497 | 3.46 | 114548.3 |
| mammals | Capromys_pilorides | Capromys_pilorides | 0 | 3090 | 2 | 111094.56 |
| mammals | Caracal_aurata | Caracal_aurata | 1 | 10650 | 2.12 | 4129643.02 |
| mammals | Caracal_caracal | Caracal_caracal | 0 | 13749.9 | 2.65 | 17205653.2 |
| mammals | Cardioderma_cor | Cardioderma_cor | 0 | 27.8 | 1 | 2246034.17 |
| mammals | Carlito_syrichta | Tarsius_syrichta | 0 | 113.5 | 1 | 120136.86 |
| mammals | Carollia_brevicauda | Carollia_brevicauda | 0 | 14.4 | 0.98 | 10749190.81 |
| mammals | Carollia_castanea | Carollia_castanea | 0 | 13.3 | 0.99 | 2501247.33 |
| mammals | Carollia_perspicillata | Carollia_perspicillata | 0 | 18 | 1 | 13796363.06 |
| mammals | Carollia_subrufa | Carollia_subrufa | 0 | 15.6 | 0.98 | 212261.47 |
| mammals | Casinycteris_argynnis | Casinycteris_argynnis | 0 | 29.9 | 0.98 | 1707684.63 |
| mammals | Castor_canadensis | Castor_canadensis | 0 | 21820 | 3.59 | 15707994.78 |
| mammals | Castor_fiber | Castor_fiber | 0 | 19000 | 3 | 3573387.44 |
| mammals | Catopuma_temminckii | Catopuma_temminckii | 0 | 11500 | 1.39 | 337441.82 |
| mammals | Cavia_aperea | Cavia_aperea | 0 | 549 | 2.31 | 6132281.51 |
| mammals | Cavia_fulgida | Cavia_fulgida | 0 | 282.5 | 1.34 | 684399.49 |
| mammals | Cavia_tschudii | Cavia_tschudii | 0 | 1000 | 1.96 | 810000.65 |
| mammals | Cebus_albifrons | Cebus_albifrons | 0 | 2629 | 1 | 2701375.07 |
| mammals | Cebus_kaapori | Cebus_kaapori | 1 | 2600 | 1 | 190775.71 |
| mammals | Centronycteris_centralis | Centronycteris_centralis | 0 | 5.5 | 0.98 | 2377722.8 |
| mammals | Centronycteris_maximiliani | Centronycteris_maximiliani | 0 | 5 | 0.98 | 5647629.78 |
| mammals | Centurio_senex | Centurio_senex | 0 | 21.2 | 0.99 | 2084372.66 |
| mammals | Cephalopachus_bancanus | Tarsius_bancanus | 1 | 94.3 | 1 | 847615.95 |
| mammals | Cephalophorus_callipygus | Cephalophus_callipygus | 0 | 18249.9 | 0.7 | 745835.02 |
| mammals | Cephalophorus_leucogaster | Cephalophus_leucogaster | 0 | 12699.9 | 1 | 1031856.96 |
| mammals | Cephalophorus_natalensis | Cephalophus_natalensis | 0 | 12000 | 1 | 1245390.44 |
| mammals | Cephalophorus_niger | Cephalophus_niger | 0 | 20400 | 1 | 761614.69 |
| mammals | Cephalophorus_ogilbyi | Cephalophus_ogilbyi | 0 | 20000 | 1 | 504344.05 |
| mammals | Cephalophorus_rufilatus | Cephalophus_rufilatus | 0 | 12000 | 1 | 2928978.08 |
| mammals | Cephalophorus_weynsi | Cephalophus_weynsi | 0 | 17000 | 1 | 1484366.45 |

| | | | | | | |
|---|---|---|---|---|---|---|
| mammals | Cephalophula_zebra | Cephalophus_zebra | 1 | 17500.1 | 1 | 59057.43 |
| mammals | Cephalophus_dorsalis | Cephalophus_dorsalis | 0 | 20000 | 0.7 | 3572525.78 |
| mammals | Cephalophus_jentinki | Cephalophus_jentinki | 1 | 70000.3 | 1 | 59693.1 |
| mammals | Cephalophus_silvicultor | Cephalophus_silvicultor | 0 | 72500.3 | 1 | 6162602.94 |
| mammals | Cephalophus_spadix | Cephalophus_spadix | 1 | 56000.3 | 1 | 6572.39 |
| mammals | Ceratotherium_simum | Ceratotherium_simum | 0 | 2949986.3 | 1 | 12627012.02 |
| mammals | Cercartetus_caudatus | Cercartetus_caudatus | 0 | 25 | 2.37 | 308509.8 |
| mammals | Cercartetus_concinnus | Cercartetus_concinnus | 0 | 13 | 4.45 | 581319.93 |
| mammals | Cercartetus_lepidus | Cercartetus_lepidus | 0 | 7 | 3.5 | 103584.92 |
| mammals | Cercartetus_nana | Cercartetus_nanus | 0 | 24 | 3.97 | 386605.43 |
| mammals | Cercocebus_agilis | Cercocebus_agilis | 0 | 8023.4 | 1 | 990299.3 |
| mammals | Cercocebus_atys | Cercocebus_atys | 1 | 7643.4 | 1.01 | 316365.52 |
| mammals | Cercocebus_chrysogaster | Cercocebus_chrysogaster | 1 | 9475.6 | 1 | 78980.88 |
| mammals | Cercocebus_galeritus | Cercocebus_galeritus | 1 | 7383 | 1.01 | 214.53 |
| mammals | Cercocebus_lunulatus | Cercocebus_lunulatus | 1 | 7275 | 1 | 164399.84 |
| mammals | Cercocebus_sanjei | Cercocebus_sanjei | 1 | 8023.4 | 1 | 188.11 |
| mammals | Cercocebus_torquatus | Cercocebus_torquatus | 1 | 8135.4 | 1 | 263787.99 |
| mammals | Cercopithecus_ascanius | Cercopithecus_ascanius | 0 | 3550 | 1 | 2619625.41 |
| mammals | Cercopithecus_campbelli | Cercopithecus_campbelli | 0 | 4100 | 1.01 | 313600.26 |
| mammals | Cercopithecus_cephus | Cercopithecus_cephus | 0 | 3485 | 1 | 835775.13 |
| mammals | Cercopithecus_diana | Cercopithecus_diana | 1 | 4175 | 1 | 221588.19 |
| mammals | Cercopithecus_erythrogaster | Cercopithecus_erythrogaster | 1 | 2500 | 1.01 | 73797.06 |
| mammals | Cercopithecus_erythrotis | Cercopithecus_erythrotis | 1 | 3918.3 | 1.01 | 54822.56 |
| mammals | Cercopithecus_hamlyni | Cercopithecus_hamlyni | 1 | 6875 | 1 | 225790.34 |
| mammals | Cercopithecus_mitis | Cercopithecus_mitis | 0 | 5000 | 1 | 2345573.67 |
| mammals | Cercopithecus_mona | Cercopithecus_mona | 0 | 4208.3 | 1 | 649990.56 |
| mammals | Cercopithecus_neglectus | Cercopithecus_neglectus | 0 | 5480 | 1.01 | 2289730.95 |
| mammals | Cercopithecus_nictitans | Cercopithecus_nictitans | 0 | 5803.3 | 1 | 1354989.94 |
| mammals | Cercopithecus_petaurista | Cercopithecus_petaurista | 0 | 2775 | 1.02 | 552996.17 |
| mammals | Cercopithecus_sclateri | Cercopithecus_sclateri | 1 | 3375 | 1.01 | 34781.6 |
| mammals | Cerdocyon_thous | Cerdocyon_thous | 0 | 5240 | 3.54 | 8433351.96 |
| mammals | Cerradomys_maracajuensis | Cerradomys_maracajuensis | 0 | 91.1 | 4.58 | 963856.02 |
| mammals | Cerradomys_marinhus | Cerradomys_marinhus | 0 | 130 | 4.58 | 31994.48 |
| mammals | Cerradomys_scotti | Cerradomys_scotti | 0 | 92 | 4.58 | 1644216.95 |

| | | | | | | |
|---|---|---|---|---|---|---|
| mammals | Cerradomys_subflavus | Cerradomys_subflavus | 0 | 96.4 | 4.48 | 1461038.82 |
| mammals | Cervus_albirostris | Cervus_albirostris | 1 | 125000 | 1 | 747388.72 |
| mammals | Cervus_canadensis | Cervus_canadensis | 0 | 131250 | 1 | 6042820.88 |
| mammals | Cervus_elaphus | Cervus_elaphus | 0 | 131250 | 1.06 | 4274919.31 |
| mammals | Cervus_nippon | Cervus_nippon | 0 | 53000 | 1 | 350964.32 |
| mammals | Chaetodipus_arenarius | Chaetodipus_arenarius | 0 | 23 | 2.91 | 88085.09 |
| mammals | Chaetodipus_baileyi | Chaetodipus_baileyi | 0 | 26.4 | 3.68 | 283199.1 |
| mammals | Chaetodipus_californicus | Chaetodipus_californicus | 0 | 22 | 4 | 106977.89 |
| mammals | Chaetodipus_eremicus | Chaetodipus_eremicus | 0 | 19 | 3.64 | 538896.38 |
| mammals | Chaetodipus_fallax | Chaetodipus_fallax | 0 | 18.7 | 3.34 | 46901.83 |
| mammals | Chaetodipus_formosus | Chaetodipus_formosus | 0 | 19.5 | 5.28 | 322064.93 |
| mammals | Chaetodipus_hispidus | Chaetodipus_hispidus | 0 | 32 | 5.23 | 2300275.4 |
| mammals | Chaetodipus_intermedius | Chaetodipus_intermedius | 0 | 16.5 | 5.27 | 510455.58 |
| mammals | Chaetodipus_nelsoni | Chaetodipus_nelsoni | 0 | 15.8 | 3.52 | 564413.41 |
| mammals | Chaetodipus_penicillatus | Chaetodipus_penicillatus | 0 | 16.9 | 3.47 | 395569.82 |
| mammals | Chaetodipus_pernix | Chaetodipus_pernix | 0 | 17 | 2 | 99068.67 |
| mammals | Chaetodipus_rudinoris | Chaetodipus_rudinoris | 0 | 26 | 3.68 | 118040.43 |
| mammals | Chaetodipus_spinatus | Chaetodipus_spinatus | 0 | 16.4 | 3.5 | 125751.34 |
| mammals | Chaetophractus_vellerosus | Chaetophractus_vellerosus | 0 | 1030 | 1.5 | 1902395.92 |
| mammals | Chaetophractus_villosus | Chaetophractus_villosus | 0 | 4540 | 1.64 | 2550820.67 |
| mammals | Chalinolobus_dwyeri | Chalinolobus_dwyeri | 1 | 9 | 1.5 | 540243.08 |
| mammals | Chalinolobus_gouldii | Chalinolobus_gouldii | 0 | 14.5 | 1.56 | 7369434.57 |
| mammals | Chalinolobus_morio | Chalinolobus_morio | 0 | 8.8 | 1.28 | 2316599.99 |
| mammals | Chalinolobus_neocaledonicus | Chalinolobus_neocaledonicus | 1 | 11.9 | 1.71 | 16190.55 |
| mammals | Chalinolobus_nigrogriseus | Chalinolobus_nigrogriseus | 0 | 8.8 | 1.5 | 1476865.1 |
| mammals | Chalinolobus_picatus | Chalinolobus_picatus | 0 | 5.1 | 2 | 1154162.95 |
| mammals | Chalinolobus_tuberculatus | Chalinolobus_tuberculatus | 1 | 8.5 | 0.98 | 160397.2 |
| mammals | Cheirogaleus_crossleyi | Cheirogaleus_crossleyi | 1 | 375 | 2.26 | 20521.66 |
| mammals | Cheirogaleus_major | Cheirogaleus_major | 1 | 450 | 2.26 | 6465.23 |
| mammals | Cheirogaleus_medius | Cheirogaleus_medius | 1 | 181 | 2.09 | 8993.79 |
| mammals | Cheirogaleus_sibreei | Cheirogaleus_sibreei | 1 | 285.5 | 2.26 | 501.85 |
| mammals | Cheiromeles_parvidens | Cheiromeles_parvidens | 0 | 114.6 | 2 | 443817.37 |
| mammals | Cheiromeles_torquatus | Cheiromeles_torquatus | 0 | 163.5 | 1 | 1330218.93 |
| mammals | Cheracebus_lucifer | Callicebus_lucifer | 0 | 1329 | 1 | 255694.18 |

| | | | | | | |
|---|---|---|---|---|---|---|
| mammals | Cheracebus_lugens | Callicebus_lugens | 0 | 1203.6 | 1 | 1160872.17 |
| mammals | Cheracebus_medemi | Callicebus_medemi | 1 | 1299.6 | 1 | 14310.13 |
| mammals | Cheracebus_regulus | Callicebus_regulus | 0 | 1273.6 | 1 | 221340.27 |
| mammals | Cheracebus_torquatus | Callicebus_torquatus | 0 | 1050 | 1.01 | 189486.26 |
| mammals | Chilonatalus_micropus | Chilonatalus_micropus | 1 | 3.5 | 1 | 14935.12 |
| mammals | Chilonatalus_tumidifrons | Chilonatalus_tumidifrons | 0 | 3.3 | 1 | 4209.14 |
| mammals | Chimarrogale_hantu | Chimarrogale_hantu | 0 | 55 | 2.5 | 12589.8 |
| mammals | Chimarrogale_platycephala | Chimarrogale_platycephalus | 0 | 32 | 4.2 | 206390.28 |
| mammals | Chinchilla_chinchilla | Chinchilla_chinchilla | 1 | 500 | 2.12 | 410196.99 |
| mammals | Chinchilla_lanigera | Chinchilla_lanigera | 1 | 417 | 1.78 | 24492.8 |
| mammals | Chionomys_gud | Chionomys_gud | 0 | 93.5 | 5.83 | 78030.91 |
| mammals | Chionomys_nivalis | Chionomys_nivalis | 0 | 54 | 3.23 | 890858.69 |
| mammals | Chionomys_roberti | Chionomys_roberti | 0 | 93.5 | 1.94 | 73076.51 |
| mammals | Chiroderma_doriae | Chiroderma_doriae | 0 | 19.9 | 1 | 2630455.51 |
| mammals | Chiroderma_salvini | Chiroderma_salvini | 0 | 25.5 | 0.99 | 3419518.38 |
| mammals | Chiroderma_trinitatum | Chiroderma_trinitatum | 0 | 13.7 | 0.98 | 9386551.23 |
| mammals | Chiroderma_villosum | Chiroderma_villosum | 0 | 23.1 | 0.99 | 10414722.69 |
| mammals | Chiromyscus_langbianis | Niviventer_langbianis | 0 | 69.8 | 3.7 | 632381.02 |
| mammals | Chironax_melanocephalus | Chironax_melanocephalus | 0 | 16.7 | 0.98 | 177749.7 |
| mammals | Chironectes_minimus | Chironectes_minimus | 0 | 946 | 3.43 | 6225494.1 |
| mammals | Chiropodomys_gliroides | Chiropodomys_gliroides | 0 | 21.5 | 1.91 | 1252844.38 |
| mammals | Chiropotes_albinasus | Chiropotes_albinasus | 1 | 2900 | 1 | 981561.87 |
| mammals | Chiropotes_chiropotes | Chiropotes_chiropotes | 0 | 3174 | 1 | 1363871.24 |
| mammals | Chiropotes_satanas | Chiropotes_satanas | 1 | 3000 | 1 | 273123.32 |
| mammals | Chiropotes_utahicki | Chiropotes_utahickae | 1 | 2784.7 | 1 | 352115.26 |
| mammals | Chiruromys_forbesi | Chiruromys_forbesi | 0 | 105.2 | 2.68 | 33438.76 |
| mammals | Chiruromys_lamia | Chiruromys_lamia | 0 | 47.2 | 2.15 | 34006.1 |
| mammals | Chiruromys_vates | Chiruromys_vates | 0 | 45.6 | 1.05 | 100855.76 |
| mammals | Chlorocebus_aethiops | Chlorocebus_aethiops | 0 | 3975 | 1 | 1136054.63 |
| mammals | Chlorocebus_cynosuros | Chlorocebus_cynosuros | 0 | 3975 | 1 | 3083230.36 |
| mammals | Chlorocebus_djamdjamensis | Chlorocebus_djamdjamensis | 1 | 3975 | 1 | 9379.71 |
| mammals | Chlorocebus_dryas | Cercopithecus_dryas | 1 | 3075 | 1.01 | 41073.21 |
| mammals | Chlorocebus_pygerythrus | Chlorocebus_pygerythrus | 0 | 3975 | 1 | 4652959.28 |
| mammals | Chlorocebus_sabaeus | Chlorocebus_sabaeus | 0 | 3975 | 1 | 1439329.71 |

| | | | | | | |
|---|---|---|---|---|---|---|
| mammals | Chlorocebus_tantalus | Chlorocebus_tantalus | 0 | 4650 | 1 | 3943411.21 |
| mammals | Choeroniscus_godmani | Choeroniscus_godmani | 0 | 7.9 | 0.98 | 2679326.39 |
| mammals | Choeroniscus_minor | Choeroniscus_minor | 0 | 8.7 | 0.98 | 5047598.73 |
| mammals | Choeronycteris_mexicana | Choeronycteris_mexicana | 0 | 16.6 | 0.99 | 1777869.42 |
| mammals | Choeropsis_liberiensis | Choeropsis_liberiensis | 1 | 235001.2 | 1 | 155982.36 |
| mammals | Choloepus_didactylus | Choloepus_didactylus | 0 | 5160 | 1 | 4166311.38 |
| mammals | Choloepus_hoffmanni | Choloepus_hoffmanni | 0 | 4750 | 1 | 2348289.73 |
| mammals | Chrotogale_owstoni | Chrotogale_owstoni | 1 | 3250 | 2 | 238124.71 |
| mammals | Chrotomys_gonzalesi | Chrotomys_gonzalesi | 0 | 140.2 | 2 | 64.41 |
| mammals | Chrotomys_silaceus | Chrotomys_silaceus | 0 | 84.1 | 2 | 4484.59 |
| mammals | Chrotomys_whiteheadi | Chrotomys_whiteheadi | 0 | 140.7 | 1.75 | 16762.79 |
| mammals | Chrotopterus_auritus | Chrotopterus_auritus | 0 | 77.9 | 0.99 | 13116792.96 |
| mammals | Chrysocyon_brachyurus | Chrysocyon_brachyurus | 0 | 23249.8 | 2.5 | 4335925.96 |
| mammals | Cistugo_lesueuri | Cistugo_lesueuri | 0 | 6.4 | 1 | 70747.33 |
| mammals | Cistugo_seabrae | Cistugo_seabrae | 0 | 5.4 | 1 | 316783.73 |
| mammals | Civettictis_civetta | Civettictis_civetta | 0 | 12000 | 2.31 | 15260382.48 |
| mammals | Clethrionomys_californicus | Myodes_californicus | 0 | 18.3 | 3.2 | 200390 |
| mammals | Clethrionomys_centralis | Myodes_centralis | 0 | 32 | 4.57 | 182570.35 |
| mammals | Clethrionomys_gapperi | Myodes_gapperi | 0 | 19 | 5.29 | 8370648.96 |
| mammals | Clethrionomys_glareolus | Myodes_glareolus | 0 | 21.2 | 4.3 | 8679125.16 |
| mammals | Clethrionomys_rutilus | Myodes_rutilus | 0 | 28.2 | 5.95 | 18530154.72 |
| mammals | Cloeotis_percivali | Cloeotis_percivali | 0 | 4.1 | 0.99 | 528464.53 |
| mammals | Clyomys_laticeps | Clyomys_laticeps | 0 | 201 | 1 | 1699627.39 |
| mammals | Cnephaeus_bottae | Eptesicus_bottae | 0 | 14.2 | 2 | 1067625.07 |
| mammals | Cnephaeus_gobiensis | Eptesicus_gobiensis | 0 | 10 | 1.25 | 4527879.71 |
| mammals | Cnephaeus_isabellinus | Eptesicus_isabellinus | 0 | 21 | 2 | 851997.5 |
| mammals | Cnephaeus_japonensis | Eptesicus_japonensis | 1 | 10.1 | 1.5 | 1460.47 |
| mammals | Cnephaeus_nilssonii | Eptesicus_nilssonii | 0 | 10.4 | 1.5 | 10473774.1 |
| mammals | Cnephaeus_serotinus | Eptesicus_serotinus | 0 | 22.9 | 1.5 | 10184906.68 |
| mammals | Coelops_frithii | Coelops_frithii | 0 | 7.4 | 0.98 | 2491060.6 |
| mammals | Coelops_robinsoni | Coelops_robinsoni | 1 | 6.5 | 1 | 137740.9 |
| mammals | Coendou_bicolor | Coendou_bicolor | 0 | 4050 | 1 | 1229394.86 |
| mammals | Coendou_insidiosus | Coendou_insidiosus | 0 | 1000 | 0.97 | 171746 |
| mammals | Coendou_mexicanus | Coendou_mexicanus | 0 | 2000 | 0.99 | 707181.02 |

| | | | | | | |
|---|---|---|---|---|---|---|
| mammals | Coendou_prehensilis | Coendou_prehensilis | 0 | 4400 | 0.99 | 12257463.45 |
| mammals | Coleura_afra | Coleura_afra | 0 | 10.7 | 1 | 2796285.82 |
| mammals | Coleura_seychellensis | Coleura_seychellensis | 1 | 10.6 | 1 | 227.69 |
| mammals | Colobus_angolensis | Colobus_angolensis | 1 | 14462.2 | 1 | 2120633.56 |
| mammals | Colobus_guereza | Colobus_guereza | 0 | 16498 | 1 | 2594832.2 |
| mammals | Colobus_polykomos | Colobus_polykomos | 1 | 9089.6 | 1 | 335688.54 |
| mammals | Colobus_satanas | Colobus_satanas | 1 | 11993.8 | 1 | 327231.88 |
| mammals | Colobus_vellerosus | Colobus_vellerosus | 1 | 11505.2 | 1.01 | 470710.48 |
| mammals | Colomys_goslingi | Colomys_goslingi | 0 | 62.5 | 1.98 | 904776.06 |
| mammals | Condylura_cristata | Condylura_cristata | 0 | 84 | 4.9 | 3486994.16 |
| mammals | Conepatus_chinga | Conepatus_chinga | 0 | 1917.5 | 3.5 | 5532259.19 |
| mammals | Conepatus_leuconotus | Conepatus_leuconotus | 0 | 3500 | 3.55 | 2498458.41 |
| mammals | Conepatus_semistriatus | Conepatus_semistriatus | 0 | 1200 | 3.74 | 2021448.58 |
| mammals | Congomys_lukolelae | Praomys_lukolelae | 0 | 41.6 | 3.5 | 208467.11 |
| mammals | Conilurus_penicillatus | Conilurus_penicillatus | 1 | 153.5 | 2 | 346343.98 |
| mammals | Connochaetes_gnou | Connochaetes_gnou | 0 | 179999 | 1 | 403590.45 |
| mammals | Connochaetes_taurinus | Connochaetes_taurinus | 0 | 179999 | 0.99 | 3103294.4 |
| mammals | Cormura_brevirostris | Cormura_brevirostris | 0 | 9.2 | 1 | 7826267.29 |
| mammals | Corynorhinus_mexicanus | Corynorhinus_mexicanus | 0 | 7.7 | 1 | 514936.65 |
| mammals | Corynorhinus_rafinesquii | Corynorhinus_rafinesquii | 0 | 9.1 | 1 | 1336202.32 |
| mammals | Corynorhinus_townsendii | Corynorhinus_townsendii | 0 | 9.9 | 1 | 4596886.83 |
| mammals | Craseomys_andersoni | Myodes_andersoni | 0 | 25.5 | 3.9 | 71603.54 |
| mammals | Craseomys_regulus | Myodes_regulus | 0 | 38.6 | 4.13 | 149439.31 |
| mammals | Craseomys_rex | Myodes_rex | 0 | 55.5 | 4.13 | 58397.48 |
| mammals | Craseomys_rufocanus | Myodes_rufocanus | 0 | 37 | 4.79 | 13399426.01 |
| mammals | Craseomys_smithii | Myodes_smithii | 0 | 35 | 4.11 | 118619.05 |
| mammals | Craseonycteris_thonglongyai | Craseonycteris_thonglongyai | 0 | 2.2 | 1 | 7954.01 |
| mammals | Crateromys_heaneyi | Crateromys_heaneyi | 1 | 994 | 1.5 | 2464.52 |
| mammals | Crateromys_schadenbergi | Crateromys_schadenbergi | 1 | 1050 | 1.5 | 2699.23 |
| mammals | Cremnomys_cutchicus | Cremnomys_cutchicus | 0 | 59.8 | 5.09 | 534108.23 |
| mammals | Cricetomys_emini | Cricetomys_emini | 0 | 1300 | 2.04 | 3732493.14 |
| mammals | Cricetomys_gambianus | Cricetomys_gambianus | 0 | 1271 | 3.22 | 4141042.44 |
| mammals | Cricetulus_barabensis | Cricetulus_barabensis | 0 | 22.9 | 5.7 | 3692081.35 |
| mammals | Cricetulus_longicaudatus | Cricetulus_longicaudatus | 0 | 22.1 | 5.71 | 2465565.12 |

| | | | | | | |
|---|---|---|---|---|---|---|
| mammals | Cricetulus_sokolovi | Cricetulus_sokolovi | 0 | 24 | 6.5 | 358309.6 |
| mammals | Cricetus_cricetus | Cricetus_cricetus | 1 | 510 | 7.14 | 4841407.64 |
| mammals | Crocidura_aleksandrisi | Crocidura_aleksandrisi | 0 | 5.25 | 4 | 42683.87 |
| mammals | Crocidura_allex | Crocidura_allex | 0 | 7 | 3.99 | 8036.62 |
| mammals | Crocidura_arabica | Crocidura_arabica | 0 | 2.8 | 4.04 | 1146.28 |
| mammals | Crocidura_attenuata | Crocidura_attenuata | 0 | 14.3 | 0.99 | 5024744.53 |
| mammals | Crocidura_attila | Crocidura_attila | 0 | 5 | 2.71 | 830715.1 |
| mammals | Crocidura_baileyi | Crocidura_baileyi | 0 | 15.3 | 2.44 | 19271.55 |
| mammals | Crocidura_baluensis | Crocidura_baluensis | 0 | 15.5 | 1.61 | 2805.71 |
| mammals | Crocidura_beata | Crocidura_beatus | 0 | 10.6 | 1 | 57076.38 |
| mammals | Crocidura_beccarii | Crocidura_beccarii | 0 | 7 | 1.61 | 91817.47 |
| mammals | Crocidura_brunnea | Crocidura_brunnea | 0 | 14.9 | 1.61 | 8474.4 |
| mammals | Crocidura_buettikoferi | Crocidura_buettikoferi | 0 | 11 | 2.99 | 47433.21 |
| mammals | Crocidura_canariensis | Crocidura_canariensis | 1 | 7.5 | 2.1 | 2520.6 |
| mammals | Crocidura_crossei | Crocidura_crossei | 0 | 6.9 | 3 | 1109826.84 |
| mammals | Crocidura_cyanea | Crocidura_cyanea | 0 | 9.4 | 3.59 | 2725072.43 |
| mammals | Crocidura_denti | Crocidura_denti | 0 | 7 | 3.49 | 1196040.5 |
| mammals | Crocidura_dolichura | Crocidura_dolichura | 0 | 5.7 | 2.5 | 1488528.97 |
| mammals | Crocidura_dsinezumi | Crocidura_dsinezumi | 0 | 10.2 | 4.23 | 292835.79 |
| mammals | Crocidura_eisentrauti | Crocidura_eisentrauti | 1 | 9.1 | 1.99 | 363.73 |
| mammals | Crocidura_flavescens | Crocidura_flavescens | 0 | 28.5 | 3.37 | 292163.8 |
| mammals | Crocidura_foxi | Crocidura_foxi | 0 | 19 | 1.99 | 1660078.47 |
| mammals | Crocidura_fuliginosa | Crocidura_fuliginosa | 0 | 12 | 1.67 | 2850808.63 |
| mammals | Crocidura_fumosa | Crocidura_fumosa | 0 | 14.5 | 3.74 | 18541.19 |
| mammals | Crocidura_fuscomurina | Crocidura_fuscomurina | 0 | 3.9 | 2.95 | 9829229.26 |
| mammals | Crocidura_grayi | Crocidura_grayi | 0 | 10.3 | 1.2 | 116385.3 |
| mammals | Crocidura_hildegardeae | Crocidura_hildegardeae | 0 | 9.5 | 2 | 2340544.71 |
| mammals | Crocidura_indochinensis | Crocidura_indochinensis | 0 | 6.3 | 3.14 | 667710.31 |
| mammals | Crocidura_jacksoni | Crocidura_jacksoni | 0 | 7 | 3.49 | 911505.58 |
| mammals | Crocidura_jenkinsi | Crocidura_jenkinsi | 1 | 12.6 | 3 | 76.38 |
| mammals | Crocidura_jouvenetae | Crocidura_jouvenetae | 0 | 9.5 | 3 | 222302.34 |
| mammals | Crocidura_lamottei | Crocidura_lamottei | 0 | 20.5 | 1 | 1559816.2 |
| mammals | Crocidura_lanosa | Crocidura_lanosa | 1 | 22.6 | 3.29 | 4797.25 |
| mammals | Crocidura_lasiura | Crocidura_lasiura | 0 | 14.8 | 8.7 | 1577925.44 |

| | | | | | | |
|---|---|---|---|---|---|---|
| mammals | Crocidura_lea | Crocidura_lea | 0 | 6.6 | 2 | 116322.61 |
| mammals | Crocidura_lepidura | Crocidura_lepidura | 0 | 18.5 | 1.61 | 232052.32 |
| mammals | Crocidura_leucodon | Crocidura_leucodon | 0 | 9.8 | 4.04 | 3927700.06 |
| mammals | Crocidura_levicula | Crocidura_levicula | 0 | 1.8 | 1.5 | 42377.71 |
| mammals | Crocidura_littoralis | Crocidura_littoralis | 0 | 16 | 2.99 | 1900103.4 |
| mammals | Crocidura_lucina | Crocidura_lucina | 1 | 12.2 | 2 | 5552.6 |
| mammals | Crocidura_luna | Crocidura_luna | 0 | 14.5 | 1.99 | 1833287.87 |
| mammals | Crocidura_macmillani | Crocidura_macmillani | 0 | 13.1 | 2.87 | 22024.4 |
| mammals | Crocidura_malayana | Crocidura_malayana | 0 | 12.3 | 1.61 | 198370.39 |
| mammals | Crocidura_maquassiensis | Crocidura_maquassiensis | 0 | 6 | 3 | 334700.91 |
| mammals | Crocidura_mariquensis | Crocidura_mariquensis | 0 | 11.3 | 3.83 | 2082090.81 |
| mammals | Crocidura_maxi | Crocidura_maxi | 0 | 7.5 | 2 | 217572.15 |
| mammals | Crocidura_montis | Crocidura_montis | 0 | 14.5 | 3.74 | 21776.07 |
| mammals | Crocidura_nicobarica | Crocidura_nicobarica | 1 | 28.9 | 3 | 90.21 |
| mammals | Crocidura_nigeriae | Crocidura_nigeriae | 0 | 23 | 2.44 | 225668.75 |
| mammals | Crocidura_nigricans | Crocidura_nigricans | 0 | 20 | 3.96 | 128601.8 |
| mammals | Crocidura_nigripes | Crocidura_nigripes | 0 | 13.2 | 1.5 | 80622.63 |
| mammals | Crocidura_nigrofusca | Crocidura_nigrofusca | 0 | 20 | 2.49 | 3815204.41 |
| mammals | Crocidura_niobe | Crocidura_niobe | 0 | 16 | 2.99 | 20530.14 |
| mammals | Crocidura_obscurior | Crocidura_obscurior | 0 | 3.6 | 2.99 | 372998.11 |
| mammals | Crocidura_olivieri | Crocidura_olivieri | 0 | 58 | 3.37 | 15840910.3 |
| mammals | Crocidura_orientalis | Crocidura_orientalis | 0 | 11.3 | 1.61 | 664.6 |
| mammals | Crocidura_orii | Crocidura_orii | 1 | 14.4 | 4.16 | 945.18 |
| mammals | Crocidura_pachyura | Crocidura_pachyura | 0 | 14.4 | 4.04 | 64302.22 |
| mammals | Crocidura_poensis | Crocidura_poensis | 0 | 16.6 | 2.44 | 1557929.72 |
| mammals | Crocidura_russula | Crocidura_russula | 0 | 11.5 | 4.04 | 1926182.44 |
| mammals | Crocidura_shantungensis | Crocidura_shantungensis | 0 | 5.5 | 4 | 3049262.93 |
| mammals | Crocidura_sibirica | Crocidura_sibirica | 0 | 73 | 4.72 | 6044227.82 |
| mammals | Crocidura_sicula | Crocidura_sicula | 0 | 6.9 | 4.38 | 26033.95 |
| mammals | Crocidura_silacea | Crocidura_silacea | 0 | 8 | 4 | 339606.76 |
| mammals | Crocidura_stenocephala | Crocidura_stenocephala | 1 | 21 | 2.99 | 1374.89 |
| mammals | Crocidura_suaveolens | Crocidura_suaveolens | 0 | 5.2 | 3.74 | 10161128.93 |
| mammals | Crocidura_tanakae | Crocidura_tanakae | 0 | 10.5 | 0.99 | 82788.98 |
| mammals | Crocidura_tansaniana | Crocidura_tansaniana | 0 | 19 | 2 | 4971.38 |

| | | | | | | |
|---|---|---|---|---|---|---|
| mammals | Crocidura_tarella | Crocidura_tarella | 1 | 16.4 | 2.75 | 2336.46 |
| mammals | Crocidura_theresae | Crocidura_theresae | 0 | 17.6 | 1.99 | 336307.09 |
| mammals | Crocidura_turba | Crocidura_turba | 0 | 14.5 | 2.38 | 4878103.99 |
| mammals | Crocidura_usambarae | Crocidura_usambarae | 1 | 9.7 | 1.99 | 5471.91 |
| mammals | Crocidura_viaria | Crocidura_viaria | 0 | 17 | 3.49 | 8205161.02 |
| mammals | Crocidura_vorax | Crocidura_vorax | 0 | 6.7 | 4.04 | 1238060.04 |
| mammals | Crocidura_watasei | Crocidura_watasei | 0 | 5.5 | 3.14 | 2460.83 |
| mammals | Crocidura_zarudnyi | Crocidura_zarudnyi | 0 | 6.7 | 5.99 | 543621.68 |
| mammals | Crocidura_zimmermanni | Crocidura_zimmermanni | 1 | 7.5 | 4.04 | 900.2 |
| mammals | Crocuta_crocuta | Crocuta_crocuta | 0 | 62999.9 | 1.96 | 14474886.61 |
| mammals | Crossarchus_alexandri | Crossarchus_alexandri | 0 | 1500 | 4 | 1412932.5 |
| mammals | Crossarchus_obscurus | Crossarchus_obscurus | 0 | 1250 | 4.14 | 492155.15 |
| mammals | Crossarchus_platycephalus | Crossarchus_platycephalus | 0 | 1000 | 4 | 515163.87 |
| mammals | Crossomys_moncktoni | Crossomys_moncktoni | 0 | 165 | 1.07 | 138694.41 |
| mammals | Cryptonanus_chacoensis | Cryptonanus_chacoensis | 0 | 15.5 | 9 | 880492.79 |
| mammals | Cryptoprocta_ferox | Cryptoprocta_ferox | 1 | 9500 | 2.99 | 442881.07 |
| mammals | Cryptotis_goldmani | Cryptotis_goldmani | 0 | 8 | 3 | 47998.51 |
| mammals | Cryptotis_goodwini | Cryptotis_goodwini | 0 | 14.5 | 2.3 | 39439.36 |
| mammals | Cryptotis_gracilis | Cryptotis_gracilis | 0 | 7 | 2.5 | 12411.96 |
| mammals | Cryptotis_magnus | Cryptotis_magna | 1 | 7 | 2.99 | 14384.64 |
| mammals | Cryptotis_mayensis | Cryptotis_mayensis | 0 | 5 | 4 | 136563.23 |
| mammals | Cryptotis_medellinius | Cryptotis_medellinia | 0 | 16.2 | 2 | 14038.82 |
| mammals | Cryptotis_meridensis | Cryptotis_meridensis | 1 | 11.6 | 3 | 13485.21 |
| mammals | Cryptotis_merriami | Cryptotis_merriami | 0 | 5.9 | 3 | 95424.31 |
| mammals | Cryptotis_mexicanus | Cryptotis_mexicana | 0 | 7 | 2.99 | 46362.73 |
| mammals | Cryptotis_montivagus | Cryptotis_montivaga | 0 | 11.3 | 2 | 29768.54 |
| mammals | Cryptotis_parvus | Cryptotis_parva | 0 | 5 | 4.55 | 4122005.69 |
| mammals | Cryptotis_phillipsii | Cryptotis_phillipsii | 1 | 8 | 3 | 5969.89 |
| mammals | Cryptotis_tamensis | Cryptotis_tamensis | 0 | 13.9 | 1.5 | 5984.29 |
| mammals | Cryptotis_thomasi | Cryptotis_thomasi | 0 | 10 | 2 | 23699.21 |
| mammals | Ctenodactylus_gundi | Ctenodactylus_gundi | 0 | 289 | 1.95 | 132731.29 |
| mammals | Ctenomys_peruanus | Ctenomys_peruanus | 0 | 490 | 3 | 5464.14 |
| mammals | Cuniculus_paca | Cuniculus_paca | 0 | 8000 | 1.01 | 13436656.96 |
| mammals | Cuniculus_taczanowskii | Cuniculus_taczanowskii | 0 | 8999.9 | 1 | 406861.26 |

| | | | | | | |
|---|---|---|---|---|---|---|
| mammals | Cuon_alpinus | Cuon_alpinus | 1 | 12760 | 4.3 | 3444014.51 |
| mammals | Cyclopes_didactylus | Cyclopes_didactylus | 0 | 329.5 | 1 | 7892585.18 |
| mammals | Cynictis_penicillata | Cynictis_penicillata | 0 | 836 | 2.08 | 2252374.39 |
| mammals | Cynocephalus_volans | Cynocephalus_volans | 0 | 1250 | 1 | 120524.27 |
| mammals | Cynogale_bennettii | Cynogale_bennettii | 1 | 4500 | 2.22 | 1090109.43 |
| mammals | Cynomops_greenhalli | Cynomops_greenhalli | 0 | 15 | 0.99 | 4472130.31 |
| mammals | Cynomops_mexicanus | Cynomops_mexicanus | 0 | 20 | 1.05 | 236547.6 |
| mammals | Cynomops_planirostris | Cynomops_planirostris | 0 | 12.4 | 1 | 11852682.31 |
| mammals | Cynomys_gunnisoni | Cynomys_gunnisoni | 0 | 925 | 4.48 | 309504.09 |
| mammals | Cynomys_leucurus | Cynomys_leucurus | 0 | 908.5 | 5.44 | 269601.97 |
| mammals | Cynomys_ludovicianus | Cynomys_ludovicianus | 0 | 1364 | 4.43 | 1820824.74 |
| mammals | Cynomys_mexicanus | Cynomys_mexicanus | 1 | 900 | 4.34 | 13330.5 |
| mammals | Cynomys_parvidens | Cynomys_parvidens | 1 | 900 | 4.88 | 31805.77 |
| mammals | Cynopterus_brachyotis | Cynopterus_brachyotis | 0 | 31.2 | 1.04 | 2699824.26 |
| mammals | Cynopterus_horsfieldii | Cynopterus_horsfieldii | 0 | 55.1 | 2 | 1485603.33 |
| mammals | Cynopterus_luzoniensis | Cynopterus_luzoniensis | 0 | 27.9 | 1 | 455804.74 |
| mammals | Cynopterus_minutus | Cynopterus_minutus | 0 | 25.8 | 1.1 | 1461294.07 |
| mammals | Cynopterus_sphinx | Cynopterus_sphinx | 0 | 43.7 | 1.3 | 6456517.37 |
| mammals | Cynopterus_titthaecheilus | Cynopterus_titthaecheilus | 0 | 56.3 | 1 | 598087.14 |
| mammals | Cyttarops_alecto | Cyttarops_alecto | 0 | 5.3 | 1 | 687747.27 |
| mammals | Dactylomys_boliviensis | Dactylomys_boliviensis | 0 | 728.8 | 2 | 336905.94 |
| mammals | Dactylomys_dactylinus | Dactylomys_dactylinus | 0 | 650 | 2 | 3306574.86 |
| mammals | Dactylopsila_megalura | Dactylopsila_megalura | 0 | 387.5 | 1.52 | 62974.32 |
| mammals | Dactylopsila_palpator | Dactylonax_palpator | 0 | 398 | 1.01 | 213289.37 |
| mammals | Dactylopsila_tatei | Dactylopsila_tatei | 1 | 252 | 1.52 | 144.87 |
| mammals | Dactylopsila_trivirgata | Dactylopsila_trivirgata | 0 | 413.5 | 1.53 | 765397.19 |
| mammals | Dama_dama | Dama_dama | 0 | 56250 | 1.03 | 4018821.4 |
| mammals | Dama_mesopotamica | Dama_mesopotamica | 1 | 72500 | 1.03 | 614.21 |
| mammals | Damaliscus_lunatus | Damaliscus_lunatus | 0 | 136000.3 | 0.99 | 1840650.42 |
| mammals | Damaliscus_pygargus | Damaliscus_pygargus | 0 | 102000 | 1.33 | 395523.62 |
| mammals | Dasycercus_blythi | Dasycercus_blythi | 0 | 83.8 | 6 | 1471984.4 |
| mammals | Dasycercus_cristicauda | Dasycercus_cristicauda | 0 | 100 | 6.38 | 114850.91 |
| mammals | Dasykaluta_rosamondae | Dasykaluta_rosamondae | 0 | 28.8 | 6.98 | 386598.87 |
| mammals | Dasymys_incomtus | Dasymys_incomtus | 0 | 90.2 | 2.63 | 6435516.37 |

| | | | | | | |
|---|---|---|---|---|---|---|
| mammals | Dasymys_montanus | Dasymys_montanus | 1 | 82 | 2.58 | 1879.35 |
| mammals | Dasymys_rufulus | Dasymys_rufulus | 0 | 85 | 2.58 | 1666893.49 |
| mammals | Dasyprocta_coibae | Dasyprocta_coibae | 0 | 2195.5 | 1.64 | 497.24 |
| mammals | Dasyprocta_fuliginosa | Dasyprocta_fuliginosa | 0 | 3500 | 2.1 | 2612229.73 |
| mammals | Dasyprocta_guamara | Dasyprocta_guamara | 0 | 2650 | 2 | 7993.76 |
| mammals | Dasyprocta_leporina | Dasyprocta_leporina | 0 | 3020 | 1.57 | 4557304.87 |
| mammals | Dasyprocta_mexicana | Dasyprocta_mexicana | 1 | 5000 | 1.5 | 107846.18 |
| mammals | Dasyprocta_prymnolopha | Dasyprocta_prymnolopha | 0 | 2900 | 1.5 | 1929753.31 |
| mammals | Dasyprocta_punctata | Dasyprocta_punctata | 0 | 2675 | 1.45 | 1306837.67 |
| mammals | Dasyprocta_ruatanica | Dasyprocta_ruatanica | 1 | 1659.9 | 1.5 | 156.61 |
| mammals | Dasypus_kappleri | Dasypus_kappleri | 0 | 9500 | 3.85 | 6028983.44 |
| mammals | Dasypus_novemcinctus | Dasypus_novemcinctus | 0 | 4203.8 | 3.98 | 18207313.67 |
| mammals | Dasypus_sabanicola | Dasypus_sabanicola | 0 | 1150 | 4 | 441453.93 |
| mammals | Dasypus_septemcinctus | Dasypus_septemcinctus | 0 | 1526.7 | 5.99 | 6904376.96 |
| mammals | Dasyuroides_byrnei | Dasyuroides_byrnei | 1 | 103.8 | 5.57 | 71511.29 |
| mammals | Dasyurus_albopunctatus | Dasyurus_albopunctatus | 0 | 637 | 6.19 | 580310.28 |
| mammals | Dasyurus_geoffroii | Dasyurus_geoffroii | 0 | 1100 | 5.6 | 188679.59 |
| mammals | Dasyurus_hallucatus | Dasyurus_hallucatus | 1 | 525 | 6.49 | 331682.51 |
| mammals | Dasyurus_maculatus | Dasyurus_maculatus | 0 | 5500 | 4.47 | 464968.77 |
| mammals | Dasyurus_spartacus | Dasyurus_spartacus | 0 | 1212.6 | 6.78 | 26551.56 |
| mammals | Dasyurus_viverrinus | Dasyurus_viverrinus | 1 | 1090 | 5.35 | 63883.7 |
| mammals | Daubentonia_madagascariensis | Daubentonia_madagascariensis | 1 | 2500 | 1 | 124674 |
| mammals | Delanymys_brooksi | Delanymys_brooksi | 1 | 5 | 3.19 | 3946.32 |
| mammals | Delomys_dorsalis | Delomys_dorsalis | 0 | 67.5 | 3.5 | 349472.7 |
| mammals | Deltamys_kempi | Deltamys_kempi | 0 | 26.4 | 3 | 141581.88 |
| mammals | Dendrogale_murina | Dendrogale_murina | 0 | 50 | 2 | 204367.54 |
| mammals | Dendrohyrax_arboreus | Dendrohyrax_arboreus | 0 | 2950 | 1.63 | 2446114.61 |
| mammals | Dendrohyrax_dorsalis | Dendrohyrax_dorsalis | 0 | 3175 | 1.29 | 3057809.09 |
| mammals | Dendrohyrax_validus | Dendrohyrax_validus | 0 | 2430 | 1.63 | 37809.88 |
| mammals | Dendrolagus_bennettianus | Dendrolagus_bennettianus | 0 | 10950 | 1 | 3562.46 |
| mammals | Dendrolagus_dorianus | Dendrolagus_dorianus | 1 | 10068.3 | 1 | 39847.25 |
| mammals | Dendrolagus_goodfellowi | Dendrolagus_goodfellowi | 1 | 8025 | 1 | 114894.77 |
| mammals | Dendrolagus_inustus | Dendrolagus_inustus | 1 | 13031.3 | 1.05 | 144442.13 |
| mammals | Dendrolagus_lumholtzi | Dendrolagus_lumholtzi | 0 | 6750 | 1 | 7383.08 |

| | | | | | | |
|---|---|---|---|---|---|---|
| mammals | Dendrolagus_matschiei | Dendrolagus_matschiei | 1 | 6960 | 1.01 | 13584.63 |
| mammals | Dendrolagus_mbaiso | Dendrolagus_mbaiso | 1 | 8750 | 1 | 5558.34 |
| mammals | Dendrolagus_notatus | Dendrolagus_notatus | 1 | 6250 | 1 | 84949.68 |
| mammals | Dendrolagus_pulcherrimus | Dendrolagus_pulcherrimus | 1 | 6536.8 | 1 | 2990.49 |
| mammals | Dendrolagus_scottae | Dendrolagus_scottae | 1 | 10375 | 1.01 | 1246.36 |
| mammals | Dendrolagus_spadix | Dendrolagus_spadix | 1 | 9100 | 1 | 49872.44 |
| mammals | Dendrolagus_stellarum | Dendrolagus_stellarum | 1 | 8666.7 | 1 | 46918.52 |
| mammals | Dendrolagus_ursinus | Dendrolagus_ursinus | 1 | 13250 | 1 | 28295.88 |
| mammals | Dendromus_insignis | Dendromus_insignis | 0 | 11.1 | 4.4 | 26896.86 |
| mammals | Dendromus_kahuziensis | Dendromus_kahuziensis | 1 | 12 | 3.49 | 75.01 |
| mammals | Dendromus_mesomelas | Dendromus_mesomelas | 0 | 11.1 | 4.17 | 588635.04 |
| mammals | Dendromus_messorius | Dendromus_messorius | 0 | 14.6 | 3.09 | 60941.42 |
| mammals | Dendromus_mystacalis | Dendromus_mystacalis | 0 | 7.7 | 3.66 | 2937067.58 |
| mammals | Dendromus_nyasae | Dendromus_kivu | 0 | 11 | 4.4 | 27532.34 |
| mammals | Dendromus_oreas | Dendromus_oreas | 0 | 10.7 | 5 | 1370.2 |
| mammals | Deomys_ferrugineus | Deomys_ferrugineus | 0 | 62 | 1.7 | 1293572.48 |
| mammals | Dephomys_defua | Dephomys_defua | 0 | 43.1 | 2.75 | 426036.66 |
| mammals | Dermanura_anderseni | Dermanura_anderseni | 0 | 138 | 0.98 | 4072157.25 |
| mammals | Dermanura_azteca | Dermanura_azteca | 0 | 20.4 | 0.99 | 606576.24 |
| mammals | Dermanura_cinerea | Dermanura_cinerea | 0 | 12.6 | 0.99 | 4322309.53 |
| mammals | Dermanura_glauca | Dermanura_glauca | 0 | 11.1 | 0.99 | 2660226.9 |
| mammals | Dermanura_gnoma | Dermanura_gnoma | 0 | 10.5 | 0.99 | 10008664.03 |
| mammals | Dermanura_phaeotis | Dermanura_phaeotis | 0 | 11.6 | 0.99 | 3737367 |
| mammals | Dermanura_tolteca | Dermanura_tolteca | 0 | 15.4 | 0.99 | 849394.91 |
| mammals | Dermanura_watsoni | Dermanura_watsoni | 0 | 12 | 1 | 554677.14 |
| mammals | Desmalopex_leucoptera | Pteropus_leucopterus | 1 | 340 | 1 | 52582.64 |
| mammals | Desmana_moschata | Desmana_moschata | 1 | 383 | 4.35 | 123164.03 |
| mammals | Desmodilliscus_braueri | Desmodilliscus_braueri | 0 | 10 | 2.58 | 3380204.83 |
| mammals | Desmodillus_auricularis | Desmodillus_auricularis | 0 | 53.3 | 2.96 | 1955073.28 |
| mammals | Desmodus_rotundus | Desmodus_rotundus | 0 | 31.2 | 1 | 17726517.72 |
| mammals | Desmomys_harringtoni | Desmomys_harringtoni | 0 | 90 | 6 | 265251.58 |
| mammals | Diaemus_youngii | Diaemus_youngi | 0 | 36.3 | 1 | 13059176.18 |
| mammals | Dicerorhinus_sumatrensis | Dicerorhinus_sumatrensis | 1 | 1266667 | 1 | 960082.54 |
| mammals | Diceros_bicornis | Diceros_bicornis | 1 | 1180510.9 | 1 | 19274959.56 |

| | | | | | | |
|---|---|---|---|---|---|---|
| mammals | Diclidurus_albus | Diclidurus_albus | 0 | 15.6 | 1 | 12498370.93 |
| mammals | Diclidurus_isabella | Diclidurus_isabella | 0 | 14 | 1 | 1061836.67 |
| mammals | Diclidurus_scutatus | Diclidurus_scutatus | 0 | 13.3 | 1 | 1540656.03 |
| mammals | Dicotyles_tajacu | Pecari_tajacu | 0 | 21266.7 | 1.69 | 15938720.07 |
| mammals | Dicrostonyx_hudsonius | Dicrostonyx_hudsonius | 0 | 57 | 3.49 | 492354.1 |
| mammals | Dicrostonyx_nunatakensis | Dicrostonyx_nunatakensis | 0 | 58.5 | 3.4 | 54570.08 |
| mammals | Didelphis_albiventris | Didelphis_albiventris | 0 | 904 | 6.9 | 6795565.82 |
| mammals | Didelphis_aurita | Didelphis_aurita | 0 | 1164 | 6.16 | 1446051.05 |
| mammals | Didelphis_imperfecta | Didelphis_imperfecta | 0 | 873.7 | 6.9 | 530382.96 |
| mammals | Didelphis_marsupialis | Didelphis_marsupialis | 0 | 1091.2 | 6.64 | 9324398.93 |
| mammals | Didelphis_pernigra | Didelphis_pernigra | 0 | 1138.5 | 6.9 | 1033727.89 |
| mammals | Didelphis_virginiana | Didelphis_virginiana | 0 | 2195.5 | 7.63 | 6033229.98 |
| mammals | Dinaromys_bogdanovi | Dinaromys_bogdanovi | 1 | 56 | 2.5 | 79580.56 |
| mammals | Dinomys_branickii | Dinomys_branickii | 0 | 12500 | 1.97 | 1390156.7 |
| mammals | Diphylla_ecaudata | Diphylla_ecaudata | 0 | 29.3 | 1 | 8092435.59 |
| mammals | Diplomys_labilis | Diplomys_labilis | 0 | 227.5 | 1.18 | 110966.52 |
| mammals | Diplothrix_legata | Diplothrix_legata | 1 | 505.9 | 7 | 799.97 |
| mammals | Dipodomys_agilis | Dipodomys_agilis | 0 | 57.9 | 2.59 | 38595.52 |
| mammals | Dipodomys_californicus | Dipodomys_californicus | 0 | 85 | 2.85 | 86209.65 |
| mammals | Dipodomys_compactus | Dipodomys_compactus | 0 | 49.3 | 2.47 | 49440.9 |
| mammals | Dipodomys_deserti | Dipodomys_deserti | 0 | 104.5 | 3.33 | 307685.28 |
| mammals | Dipodomys_elator | Dipodomys_elator | 1 | 77.5 | 2.91 | 10767.66 |
| mammals | Dipodomys_heermanni | Dipodomys_heermanni | 0 | 72 | 3.11 | 70208.59 |
| mammals | Dipodomys_ingens | Dipodomys_ingens | 1 | 133.9 | 4.26 | 6784.27 |
| mammals | Dipodomys_merriami | Dipodomys_merriami | 0 | 42 | 2.35 | 1423683.82 |
| mammals | Dipodomys_microps | Dipodomys_microps | 0 | 54.6 | 2.38 | 392308.36 |
| mammals | Dipodomys_nelsoni | Dipodomys_nelsoni | 0 | 88.6 | 1.94 | 192588.89 |
| mammals | Dipodomys_nitratoides | Dipodomys_nitratoides | 1 | 43.9 | 2.22 | 11331.6 |
| mammals | Dipodomys_ordii | Dipodomys_ordii | 0 | 60.4 | 3.05 | 3419731.41 |
| mammals | Dipodomys_panamintinus | Dipodomys_panamintinus | 0 | 74.7 | 3.64 | 60988.14 |
| mammals | Dipodomys_phillipsii | Dipodomys_phillipsii | 0 | 41 | 2.59 | 62377.86 |
| mammals | Dipodomys_simulans | Dipodomys_simulans | 0 | 57.5 | 2.5 | 97814.57 |
| mammals | Dipodomys_spectabilis | Dipodomys_spectabilis | 0 | 135.9 | 2.66 | 523883.19 |
| mammals | Dipodomys_stephensi | Dipodomys_stephensi | 1 | 69.8 | 2.65 | 9357.28 |

| | | | | | | |
|---|---|---|---|---|---|---|
| mammals | Dipodomys_venustus | Dipodomys_venustus | 0 | 72 | 1.86 | 16356.39 |
| mammals | Dipus_sagitta | Dipus_sagitta | 0 | 85.7 | 3.46 | 3156284.93 |
| mammals | Distoechurus_pennatus | Distoechurus_pennatus | 0 | 44.1 | 1.23 | 536111.97 |
| mammals | Dobsonia_anderseni | Dobsonia_anderseni | 0 | 204.1 | 0.99 | 46527 |
| mammals | Dobsonia_beauforti | Dobsonia_beauforti | 0 | 159 | 0.99 | 10000.15 |
| mammals | Dobsonia_chapmani | Dobsonia_chapmani | 1 | 134 | 1 | 29.11 |
| mammals | Dobsonia_crenulata | Dobsonia_crenulata | 0 | 272 | 1 | 209291.77 |
| mammals | Dobsonia_inermis | Dobsonia_inermis | 0 | 149.5 | 0.99 | 34750.34 |
| mammals | Dobsonia_minor | Dobsonia_minor | 0 | 78.2 | 0.99 | 614847.58 |
| mammals | Dobsonia_moluccensis | Dobsonia_moluccensis | 0 | 381.4 | 0.99 | 39785.13 |
| mammals | Dobsonia_pannietensis | Dobsonia_pannietensis | 0 | 267.7 | 0.98 | 5506.82 |
| mammals | Dobsonia_peronii | Dobsonia_peronii | 0 | 216.9 | 0.98 | 87783.69 |
| mammals | Dobsonia_praedatrix | Dobsonia_praedatrix | 0 | 188.7 | 1 | 44746.12 |
| mammals | Dolichotis_patagonum | Dolichotis_patagonum | 0 | 8000 | 1.81 | 1242817 |
| mammals | Dolichotis_salinicola | Dolichotis_salinicola | 0 | 1600 | 1.77 | 735968.05 |
| mammals | Dorcatragus_megalotis | Dorcatragus_megalotis | 1 | 10249.9 | 1 | 51752.99 |
| mammals | Dorcopsis_atrata | Dorcopsis_atrata | 1 | 5916.5 | 1.01 | 83.25 |
| mammals | Dorcopsis_hageni | Dorcopsis_hageni | 0 | 5500 | 1.01 | 152971.24 |
| mammals | Dorcopsis_luctuosa | Dorcopsis_luctuosa | 1 | 6352 | 1.01 | 159592.36 |
| mammals | Dorcopsis_muelleri | Dorcopsis_muelleri | 0 | 5500 | 1 | 125655.34 |
| mammals | Dorcopsulus_macleayi | Dorcopsulus_macleayi | 0 | 2988 | 1.22 | 25885.92 |
| mammals | Dorcopsulus_vanheurni | Dorcopsulus_vanheurni | 0 | 1785.8 | 1.18 | 179392.5 |
| mammals | Doryrhina_cyclops | Hipposideros_cyclops | 0 | 33.5 | 1 | 2331176.89 |
| mammals | Dremomys_lokriah | Dremomys_lokriah | 0 | 227 | 3.67 | 361878.67 |
| mammals | Dremomys_pernyi | Dremomys_pernyi | 0 | 198.6 | 4 | 1752951.56 |
| mammals | Dromiciops_gliroides | Dromiciops_gliroides | 0 | 22.3 | 3 | 139169.35 |
| mammals | Dryomys_nitedula | Dryomys_nitedula | 0 | 28.9 | 3.24 | 5228746.46 |
| mammals | Dyacopterus_brooksi | Dyacopterus_brooksi | 1 | 88.5 | 1.5 | 429470.12 |
| mammals | Dyacopterus_spadiceus | Dyacopterus_spadiceus | 0 | 84.2 | 1.5 | 445030.61 |
| mammals | Dymecodon_pilirostris | Dymecodon_pilirostris | 0 | 16.2 | 3 | 38974.79 |
| mammals | Echimys_chrysurus | Echimys_chrysurus | 0 | 652.5 | 1.96 | 1583343.24 |
| mammals | Echinops_telfairi | Echinops_telfairi | 0 | 180 | 5.49 | 111004.16 |
| mammals | Echinosorex_gymnura | Echinosorex_gymnura | 0 | 957 | 1.88 | 1373927.43 |
| mammals | Echiothrix_centrosa | Echiothrix_centrosa | 0 | 260.9 | 3 | 34850.35 |

| | | | | | | |
|---|---|---|---|---|---|---|
| mammals | Echymipera_clara | Echymipera_clara | 0 | 1236.5 | 2.2 | 175468.6 |
| mammals | Echymipera_kalubu | Echymipera_kalubu | 0 | 782.2 | 2.08 | 840230.18 |
| mammals | Echymipera_rufescens | Echymipera_rufescens | 0 | 1100 | 2.19 | 859314.67 |
| mammals | Ectophylla_alba | Ectophylla_alba | 0 | 5.8 | 0.99 | 124361.57 |
| mammals | Eidolon_dupreanum | Eidolon_dupreanum | 1 | 283.8 | 1.15 | 460027.21 |
| mammals | Eidolon_helvum | Eidolon_helvum | 0 | 253.3 | 1.06 | 11789001.27 |
| mammals | Eira_barbara | Eira_barbara | 0 | 3910 | 2.07 | 13375343.99 |
| mammals | Elaphodus_cephalophus | Elaphodus_cephalophus | 0 | 33500 | 1 | 1944315.29 |
| mammals | Elephantulus_brachyrhynchus | Elephantulus_brachyrhynchus | 0 | 47.5 | 1.59 | 3874147.2 |
| mammals | Elephantulus_edwardii | Elephantulus_edwardii | 0 | 48 | 1.8 | 193798.45 |
| mammals | Elephantulus_intufi | Elephantulus_intufi | 0 | 49.5 | 1.57 | 1212847.09 |
| mammals | Elephantulus_myurus | Elephantulus_myurus | 0 | 52.4 | 1.84 | 968379.6 |
| mammals | Elephantulus_rupestris | Elephantulus_rupestris | 0 | 66.4 | 1.84 | 557903.85 |
| mammals | Elephas_maximus | Elephas_maximus | 1 | 3160000 | 1.21 | 894367.58 |
| mammals | Eligmodontia_morgani | Eligmodontia_morgani | 0 | 16.5 | 5.9 | 371796.23 |
| mammals | Eligmodontia_puerulus | Eligmodontia_puerulus | 0 | 28.5 | 5.12 | 447158.57 |
| mammals | Eligmodontia_typus | Eligmodontia_typus | 0 | 17.3 | 5.68 | 922783.36 |
| mammals | Eliomys_melanurus | Eliomys_melanurus | 0 | 51.8 | 4.99 | 148691.2 |
| mammals | Eliomys_munbyanus | Eliomys_munbyanus | 0 | 52 | 4.99 | 873740.08 |
| mammals | Eliomys_quercinus | Eliomys_quercinus | 0 | 115 | 4.99 | 2362160.86 |
| mammals | Eliurus_grandidieri | Eliurus_grandidieri | 0 | 51 | 3 | 60596.3 |
| mammals | Eliurus_majori | Eliurus_majori | 0 | 100 | 5 | 88713.44 |
| mammals | Eliurus_minor | Eliurus_minor | 0 | 35 | 4 | 114231.27 |
| mammals | Eliurus_myoxinus | Eliurus_myoxinus | 0 | 62.5 | 4 | 279015.1 |
| mammals | Eliurus_tanala | Eliurus_tanala | 0 | 82.5 | 4 | 120856.31 |
| mammals | Eliurus_webbi | Eliurus_webbi | 0 | 72.5 | 5 | 133408.49 |
| mammals | Emballonura_alecto | Emballonura_alecto | 0 | 5.3 | 1 | 1212058.67 |
| mammals | Emballonura_beccarii | Emballonura_beccarii | 0 | 4.2 | 1 | 91119.17 |
| mammals | Emballonura_dianae | Emballonura_dianae | 0 | 8 | 1.5 | 90456.79 |
| mammals | Emballonura_furax | Emballonura_furax | 0 | 11.2 | 1 | 67865.15 |
| mammals | Emballonura_monticola | Emballonura_monticola | 0 | 5.7 | 1 | 1690479.34 |
| mammals | Emballonura_raffrayana | Emballonura_raffrayana | 0 | 5.6 | 1 | 216455.31 |
| mammals | Emballonura_semicaudata | Emballonura_semicaudata | 1 | 6.3 | 1 | 20505.94 |
| mammals | Emballonura_serii | Emballonura_serii | 1 | 8.8 | 1 | 11497.34 |

| | | | | | | |
|---|---|---|---|---|---|---|
| mammals | Enchisthenes_hartii | Enchisthenes_hartii | 0 | 16 | 0.99 | 3104106.01 |
| mammals | Eoglaucomys_fimbriatus | Eoglaucomys_fimbriatus | 0 | 712 | 3.7 | 188936.35 |
| mammals | Eolagurus_przewalskii | Eolagurus_przewalskii | 0 | 16 | 6.4 | 903281.89 |
| mammals | Eonycteris_major | Eonycteris_major | 0 | 72.8 | 1 | 732628.36 |
| mammals | Eonycteris_robusta | Eonycteris_robusta | 1 | 77.5 | 1 | 114506.38 |
| mammals | Eonycteris_spelaea | Eonycteris_spelaea | 0 | 58.1 | 1 | 3525628.5 |
| mammals | Eothenomys_melanogaster | Eothenomys_melanogaster | 0 | 27 | 1.8 | 1722986.44 |
| mammals | Eozapus_setchuanus | Eozapus_setchuanus | 0 | 32.6 | 5.5 | 545018.87 |
| mammals | Episoriculus_caudatus | Episoriculus_caudatus | 0 | 9.1 | 4.77 | 315602.53 |
| mammals | Epixerus_ebii | Epixerus_ebii | 0 | 592 | 2 | 540525.79 |
| mammals | Epomophorus_angolensis | Epomophorus_angolensis | 0 | 131.7 | 1 | 383579.71 |
| mammals | Epomophorus_crypturus | Epomophorus_crypturus | 0 | 111.1 | 1 | 2084376.21 |
| mammals | Epomophorus_dobsonii | Epomops_dobsonii | 0 | 120 | 1.5 | 1367022.75 |
| mammals | Epomophorus_gambianus | Epomophorus_gambianus | 0 | 128.9 | 0.99 | 3791163.45 |
| mammals | Epomophorus_labiatus | Epomophorus_labiatus | 0 | 63 | 0.99 | 1765029.78 |
| mammals | Epomophorus_minor | Epomophorus_minor | 0 | 50 | 1 | 2256487.09 |
| mammals | Epomophorus_pusillus | Micropteropus_pusillus | 0 | 26.6 | 0.99 | 5380916.58 |
| mammals | Epomophorus_wahlbergi | Epomophorus_wahlbergi | 0 | 91.6 | 1 | 4999921.48 |
| mammals | Epomops_buettikoferi | Epomops_buettikoferi | 0 | 133.8 | 0.98 | 1079556.03 |
| mammals | Epomops_franqueti | Epomops_franqueti | 0 | 118.9 | 1 | 4509721.41 |
| mammals | Eptesicus_fuscus | Eptesicus_fuscus | 0 | 17.5 | 1.89 | 12798347.84 |
| mammals | Equus_africanus | Equus_africanus | 1 | 325000 | 1 | 84849.87 |
| mammals | Equus_ferus | Equus_ferus | 1 | 250000 | 1 | 26271.98 |
| mammals | Equus_grevyi | Equus_grevyi | 1 | 407999.8 | 1 | 90832.52 |
| mammals | Equus_hemionus | Equus_hemionus | 0 | 230000 | 1 | 390569.84 |
| mammals | Equus_kiang | Equus_kiang | 0 | 275000 | 1 | 1785558.32 |
| mammals | Equus_quagga | Equus_quagga | 0 | 276000 | 1 | 2314209.98 |
| mammals | Equus_zebra | Equus_zebra | 1 | 286998.7 | 1 | 321622.17 |
| mammals | Eremitalpa_granti | Eremitalpa_granti | 0 | 26.1 | 1.27 | 50294.06 |
| mammals | Eremodipus_lichtensteini | Eremodipus_lichtensteini | 0 | 51 | 4.86 | 559492.76 |
| mammals | Erethizon_dorsatum | Erethizon_dorsatum | 0 | 7085.3 | 1 | 12063228.06 |
| mammals | Erinaceus_amurensis | Erinaceus_amurensis | 0 | 800 | 4.22 | 4254779.31 |
| mammals | Erinaceus_concolor | Erinaceus_concolor | 0 | 719 | 4.86 | 1346308.2 |
| mammals | Erinaceus_europaeus | Erinaceus_europaeus | 0 | 771 | 4.43 | 3809035.1 |

| | | | | | | |
|---|---|---|---|---|---|---|
| mammals | Erinaceus_roumanicus | Erinaceus_roumanicus | 0 | 745 | 4.5 | 6029711.84 |
| mammals | Erophylla_bombifrons | Erophylla_bombifrons | 0 | 14.3 | 1 | 82891.95 |
| mammals | Erophylla_sezekorni | Erophylla_sezekorni | 0 | 15.8 | 1 | 122945.38 |
| mammals | Erythrocebus_patas | Erythrocebus_patas | 0 | 8090 | 1 | 6604128.33 |
| mammals | Euchoreutes_naso | Euchoreutes_naso | 0 | 24.1 | 3.1 | 1674821.34 |
| mammals | Euderma_maculatum | Euderma_maculatum | 0 | 15.7 | 1 | 2104986.07 |
| mammals | Eudorcas_rufifrons | Eudorcas_rufifrons | 1 | 26999.8 | 1 | 2198020.73 |
| mammals | Eudorcas_thomsonii | Eudorcas_thomsonii | 0 | 20499.8 | 1 | 157281.78 |
| mammals | Eulemur_albifrons | Eulemur_albifrons | 1 | 2300 | 1.07 | 19643.19 |
| mammals | Eulemur_cinereiceps | Eulemur_cinereiceps | 1 | 1920 | 1.1 | 1817.02 |
| mammals | Eulemur_collaris | Eulemur_collaris | 1 | 2375 | 1.1 | 8156.85 |
| mammals | Eulemur_coronatus | Eulemur_coronatus | 1 | 1650 | 1.51 | 6521.7 |
| mammals | Eulemur_flavifrons | Eulemur_flavifrons | 1 | 1850 | 1.04 | 2450.84 |
| mammals | Eulemur_fulvus | Eulemur_fulvus | 1 | 2500 | 1.1 | 48944.48 |
| mammals | Eulemur_macaco | Eulemur_macaco | 1 | 1900 | 1.04 | 7963.78 |
| mammals | Eulemur_mongoz | Eulemur_mongoz | 1 | 1350 | 1.11 | 8051.6 |
| mammals | Eulemur_rubriventer | Eulemur_rubriventer | 1 | 2000 | 1 | 51229.16 |
| mammals | Eulemur_rufifrons | Eulemur_rufifrons | 1 | 1820 | 1 | 32696.6 |
| mammals | Eulemur_rufus | Eulemur_rufus | 1 | 2375 | 1.1 | 37372.75 |
| mammals | Eulemur_sanfordi | Eulemur_sanfordi | 1 | 2250 | 1.1 | 3830.39 |
| mammals | Eumops_auripendulus | Eumops_auripendulus | 0 | 28.5 | 0.98 | 14106769.9 |
| mammals | Eumops_bonariensis | Eumops_bonariensis | 0 | 11.9 | 1 | 1161090.37 |
| mammals | Eumops_dabbenei | Eumops_dabbenei | 0 | 104.4 | 1 | 2407709.82 |
| mammals | Eumops_floridanus | Eumops_floridanus | 1 | 38.4 | 1.03 | 999.18 |
| mammals | Eumops_glaucinus | Eumops_glaucinus | 0 | 35.4 | 1.03 | 8971227.62 |
| mammals | Eumops_hansae | Eumops_hansae | 0 | 15.1 | 1.05 | 6416223.24 |
| mammals | Eumops_patagonicus | Eumops_patagonicus | 0 | 10 | 1 | 1535787.62 |
| mammals | Eumops_perotis | Eumops_perotis | 0 | 51.1 | 1 | 11695739.96 |
| mammals | Eumops_trumbulli | Eumops_trumbulli | 0 | 36.4 | 1 | 4744456.77 |
| mammals | Eumops_underwoodi | Eumops_underwoodi | 0 | 54.7 | 1 | 955823.89 |
| mammals | Euoticus_elegantulus | Euoticus_elegantulus | 0 | 300 | 0.99 | 782197.19 |
| mammals | Euoticus_pallidus | Euoticus_pallidus | 0 | 230 | 1 | 95102.99 |
| mammals | Eupetaurus_cinereus | Eupetaurus_cinereus | 1 | 2500 | 2.5 | 36743.24 |
| mammals | Euphractus_sexcinctus | Euphractus_sexcinctus | 0 | 4782.9 | 1.74 | 7716332.83 |

| | | | | | | |
|---|---|---|---|---|---|---|
| mammals | Eupleres_goudotii | Eupleres_goudotii | 1 | 1850 | 1.2 | 105598.78 |
| mammals | Euryoryzomys_macconnelli | Euryoryzomys_macconnelli | 0 | 58 | 3 | 4294868.38 |
| mammals | Euryoryzomys_nitidus | Euryoryzomys_nitidus | 0 | 55.2 | 4.02 | 1303446.66 |
| mammals | Euryzygomatomys_spinosus | Euryzygomatomys_spinosus | 0 | 187.5 | 0.97 | 807543.66 |
| mammals | Eutamias_sibiricus | Eutamias_sibiricus | 0 | 85 | 4.92 | 14527269.11 |
| mammals | Euxerus_erythropus | Xerus_erythropus | 0 | 502.4 | 3.53 | 8816274.08 |
| mammals | Exilisciurus_concinnus | Exilisciurus_concinnus | 0 | 30.9 | 1.94 | 120457.57 |
| mammals | Exilisciurus_whiteheadi | Exilisciurus_whiteheadi | 0 | 20 | 1.94 | 155450.15 |
| mammals | Falsistrellus_mackenziei | Falsistrellus_mackenziei | 0 | 21 | 1 | 24586.8 |
| mammals | Falsistrellus_tasmaniensis | Falsistrellus_tasmaniensis | 1 | 22.7 | 1 | 386354.11 |
| mammals | Felis_bieti | Felis_bieti | 1 | 7750 | 3.69 | 940432.42 |
| mammals | Felis_chaus | Felis_chaus | 0 | 7393 | 2.97 | 7405493.52 |
| mammals | Felis_margarita | Felis_margarita | 0 | 2525 | 4.31 | 3701188.81 |
| mammals | Felis_nigripes | Felis_nigripes | 1 | 1300 | 1.84 | 1948137.93 |
| mammals | Felis_silvestris | Felis_silvestris | 0 | 5500 | 3.64 | 1931656.66 |
| mammals | Felovia_vae | Felovia_vae | 0 | 205 | 1 | 183984.78 |
| mammals | Fossa_fossana | Fossa_fossana | 1 | 1500 | 1 | 91103.59 |
| mammals | Funambulus_layardi | Funambulus_layardi | 1 | 168 | 2.45 | 1276.83 |
| mammals | Funambulus_palmarum | Funambulus_palmarum | 0 | 108.4 | 2.62 | 1263136.9 |
| mammals | Funambulus_pennantii | Funambulus_pennantii | 0 | 99.1 | 3.04 | 2908289.2 |
| mammals | Funambulus_sublineatus | Funambulus_sublineatus | 1 | 70.9 | 2 | 21379.11 |
| mammals | Funambulus_tristriatus | Funambulus_tristriatus | 0 | 139 | 2.67 | 79432.11 |
| mammals | Funisciurus_anerythrus | Funisciurus_anerythrus | 0 | 218 | 1.36 | 2279777.19 |
| mammals | Funisciurus_congicus | Funisciurus_congicus | 0 | 112 | 2.22 | 1735003.2 |
| mammals | Funisciurus_isabella | Funisciurus_isabella | 0 | 109 | 1.39 | 570175.35 |
| mammals | Funisciurus_lemniscatus | Funisciurus_lemniscatus | 0 | 141 | 1.8 | 610756.52 |
| mammals | Funisciurus_leucogenys | Funisciurus_leucogenys | 0 | 250 | 0.98 | 411996.78 |
| mammals | Funisciurus_pyrropus | Funisciurus_pyrropus | 0 | 297 | 1.47 | 2153133.03 |
| mammals | Furipterus_horrens | Furipterus_horrens | 0 | 3.2 | 1 | 9816319.15 |
| mammals | Galago_gallarum | Galago_gallarum | 0 | 250 | 1.5 | 307921.62 |
| mammals | Galago_matschiei | Galago_matschiei | 0 | 210 | 1.25 | 86725.87 |
| mammals | Galago_moholi | Galago_moholi | 0 | 160 | 1.57 | 3985248.48 |
| mammals | Galago_senegalensis | Galago_senegalensis | 0 | 227.7 | 1.5 | 7758268.69 |
| mammals | Galagoides_demidoff | Galagoides_demidoff | 0 | 60.1 | 1.2 | 4348190.7 |

| | | | | | | |
|---|---|---|---|---|---|---|
| mammals | Galagoides_thomasi | Galagoides_thomasi | 0 | 99.7 | 1.29 | 4407657 |
| mammals | Galea_flavidens | Galea_flavidens | 0 | 450 | 1 | 32759.67 |
| mammals | Galea_leucoblephara | Galea_leucoblephara | 0 | 230 | 3 | 1914449.5 |
| mammals | Galea_spixii | Galea_spixii | 0 | 326.2 | 2.3 | 1699340.1 |
| mammals | Galegeeska_rufescens | Elephantulus_rufescens | 0 | 37.5 | 1.41 | 1493648.42 |
| mammals | Galemys_pyrenaicus | Galemys_pyrenaicus | 1 | 60.2 | 3.59 | 130939.77 |
| mammals | Galeopterus_variegatus | Galeopterus_variegatus | 0 | 1012.5 | 1.2 | 1410929.24 |
| mammals | Galictis_cuja | Galictis_cuja | 0 | 1000 | 3.5 | 7672218.09 |
| mammals | Galictis_vittata | Galictis_vittata | 0 | 3200 | 2 | 10014922.31 |
| mammals | Galidia_elegans | Galidia_elegans | 0 | 800 | 1 | 92492.35 |
| mammals | Galidictis_fasciata | Galidictis_fasciata | 1 | 550 | 1 | 83601.29 |
| mammals | Gardnerycteris_crenulata | Mimon_crenulatum | 0 | 13.8 | 0.99 | 11134091.34 |
| mammals | Gazella_bennettii | Gazella_bennettii | 0 | 18916.7 | 1.22 | 2764528.24 |
| mammals | Gazella_cuvieri | Gazella_cuvieri | 1 | 25000 | 1.23 | 152444.46 |
| mammals | Gazella_dorcas | Gazella_dorcas | 1 | 23000.1 | 1.22 | 9905633.09 |
| mammals | Gazella_gazella | Gazella_gazella | 1 | 22749.9 | 1.35 | 14845.71 |
| mammals | Gazella_leptoceros | Gazella_leptoceros | 1 | 22200 | 1 | 1224653.72 |
| mammals | Gazella_marica | Gazella_marica | 1 | 20200 | 1.7 | 2613936.41 |
| mammals | Gazella_spekei | Gazella_spekei | 1 | 20000 | 1 | 180961.81 |
| mammals | Gazella_subgutturosa | Gazella_subgutturosa | 1 | 28500 | 1.39 | 7107547.32 |
| mammals | Genetta_cristata | Genetta_cristata | 1 | 2500 | 2.25 | 100907.24 |
| mammals | Genetta_fieldiana | Genetta_maculata | 0 | 2225 | 2.21 | 12856912.48 |
| mammals | Genetta_genetta | Genetta_genetta | 0 | 1800 | 2.29 | 14894292.42 |
| mammals | Genetta_johnstoni | Genetta_johnstoni | 0 | 2225 | 2 | 307033.32 |
| mammals | Genetta_pardina | Genetta_pardina | 0 | 3100 | 2.36 | 1435327.91 |
| mammals | Genetta_piscivora | Genetta_piscivora | 0 | 3000 | 0.96 | 223601.46 |
| mammals | Genetta_servalina | Genetta_servalina | 0 | 1055 | 1.5 | 3171148.46 |
| mammals | Genetta_tigrina | Genetta_tigrina | 0 | 2225 | 2.63 | 212859.61 |
| mammals | Geocapromys_brownii | Geocapromys_brownii | 1 | 1500 | 1.76 | 987.88 |
| mammals | Geogale_aurita | Geogale_aurita | 0 | 7 | 3.52 | 10985.16 |
| mammals | Geosciurus_inauris | Xerus_inauris | 0 | 529 | 2.09 | 1515178.64 |
| mammals | Geosciurus_princeps | Xerus_princeps | 0 | 636 | 2 | 238199.93 |
| mammals | Geoxus_valdivianus | Geoxus_valdivianus | 0 | 31.5 | 3.4 | 232674.62 |
| mammals | Gerbilliscus_afer | Gerbilliscus_afra | 0 | 97 | 3.92 | 100211.97 |

| | | | | | | |
|---|---|---|---|---|---|---|
| mammals | Gerbilliscus_boehmi | Gerbilliscus_boehmi | 0 | 146 | 4.06 | 1160649 |
| mammals | Gerbilliscus_brantsii | Gerbilliscus_brantsii | 0 | 114.4 | 2.76 | 2084250.52 |
| mammals | Gerbilliscus_gambianus | Gerbilliscus_gambiana | 0 | 114.5 | 4 | 281829.32 |
| mammals | Gerbilliscus_guineae | Gerbilliscus_guineae | 0 | 102.5 | 5 | 892568.33 |
| mammals | Gerbilliscus_inclusus | Gerbilliscus_inclusus | 0 | 118.5 | 2.5 | 218326.24 |
| mammals | Gerbilliscus_kempii | Gerbilliscus_kempi | 0 | 101 | 4.15 | 4610375.98 |
| mammals | Gerbilliscus_leucogaster | Gerbilliscus_leucogaster | 0 | 72.8 | 4.51 | 5765221.89 |
| mammals | Gerbilliscus_nigricaudus | Gerbilliscus_nigricaudus | 0 | 123 | 4.44 | 610487.09 |
| mammals | Gerbilliscus_paeba | Gerbillurus_paeba | 0 | 23.3 | 3.64 | 2191068.63 |
| mammals | Gerbilliscus_robustus | Gerbilliscus_robustus | 0 | 96 | 5.23 | 5165821.44 |
| mammals | Gerbilliscus_setzeri | Gerbillurus_setzeri | 0 | 38.3 | 3 | 48792.27 |
| mammals | Gerbilliscus_tytonis | Gerbillurus_tytonis | 0 | 31 | 4.4 | 56157 |
| mammals | Gerbilliscus_validus | Gerbilliscus_validus | 0 | 107.7 | 4.12 | 4626297.08 |
| mammals | Gerbilliscus_vallinus | Gerbillurus_vallinus | 0 | 35.3 | 3.87 | 286817.66 |
| mammals | Gerbillus_amoenus | Gerbillus_amoenus | 0 | 25 | 4.08 | 855071.29 |
| mammals | Gerbillus_andersoni | Gerbillus_andersoni | 0 | 24.5 | 3.6 | 117593.89 |
| mammals | Gerbillus_aquilus | Gerbillus_aquilus | 0 | 43.8 | 8 | 559283.14 |
| mammals | Gerbillus_campestris | Gerbillus_campestris | 0 | 28.4 | 4.34 | 1163165.52 |
| mammals | Gerbillus_cheesmani | Gerbillus_cheesmani | 0 | 28 | 8 | 2285825.37 |
| mammals | Gerbillus_dasyurus | Gerbillus_dasyurus | 0 | 23.5 | 4.01 | 1542832.66 |
| mammals | Gerbillus_floweri | Gerbillus_floweri | 0 | 56 | 3.3 | 103318.61 |
| mammals | Gerbillus_gerbillus | Gerbillus_gerbillus | 0 | 25 | 4.12 | 9376064.67 |
| mammals | Gerbillus_gleadowi | Gerbillus_gleadowi | 0 | 24 | 3.12 | 183769.51 |
| mammals | Gerbillus_harwoodi | Gerbillus_harwoodi | 0 | 11.1 | 4 | 205253.7 |
| mammals | Gerbillus_henleyi | Gerbillus_henleyi | 0 | 8 | 4 | 3846929.15 |
| mammals | Gerbillus_hesperinus | Gerbillus_hesperinus | 1 | 30.2 | 3.3 | 724.54 |
| mammals | Gerbillus_maghrebi | Gerbillus_maghrebi | 0 | 46 | 8 | 55225.67 |
| mammals | Gerbillus_mesopotamiae | Gerbillus_mesopotamiae | 0 | 20 | 5.1 | 219864.08 |
| mammals | Gerbillus_nanus | Gerbillus_nanus | 0 | 21.5 | 3.62 | 10213065.72 |
| mammals | Gerbillus_nigeriae | Gerbillus_nigeriae | 0 | 26.8 | 4.1 | 71424.33 |
| mammals | Gerbillus_pyramidum | Gerbillus_pyramidum | 0 | 52 | 3.3 | 3344944.49 |
| mammals | Gerbillus_simoni | Gerbillus_simoni | 0 | 17.4 | 4.68 | 259962.23 |
| mammals | Gerbillus_tarabuli | Gerbillus_tarabuli | 0 | 42.5 | 3.38 | 6332711.38 |
| mammals | Giraffa_camelopardalis | Giraffa_camelopardalis | 1 | 899994.8 | 1.21 | 1794018.08 |

| | | | | | | |
|---|---|---|---|---|---|---|
| mammals | Glaucomys_sabrinus | Glaucomys_sabrinus | 0 | 166 | 3 | 7852908.63 |
| mammals | Glaucomys_volans | Glaucomys_volans | 0 | 63.9 | 3.13 | 3848743.43 |
| mammals | Glauconycteris_variegata | Glauconycteris_variegata | 0 | 11.1 | 1.09 | 8695501.17 |
| mammals | Glironia_venusta | Glironia_venusta | 0 | 114 | 4 | 2883884.4 |
| mammals | Glirulus_japonicus | Glirulus_japonicus | 0 | 27 | 4 | 211072.73 |
| mammals | Glis_glis | Glis_glis | 0 | 125 | 4.95 | 3718880.11 |
| mammals | Glischropus_tylopus | Glischropus_tylopus | 0 | 4.4 | 1 | 1373589.24 |
| mammals | Glossophaga_commissarisi | Glossophaga_commissarisi | 0 | 8.9 | 0.99 | 3716729.94 |
| mammals | Glossophaga_leachii | Glossophaga_leachii | 0 | 10.1 | 0.99 | 480889.97 |
| mammals | Glossophaga_longirostris | Glossophaga_longirostris | 0 | 11.9 | 0.99 | 1533778.45 |
| mammals | Glossophaga_morenoi | Glossophaga_morenoi | 0 | 8.4 | 1 | 327072.97 |
| mammals | Glossophaga_soricina | Glossophaga_soricina | 0 | 9.4 | 1 | 14858417.39 |
| mammals | Glyphonycteris_daviesi | Glyphonycteris_daviesi | 0 | 18.3 | 0.98 | 4708611.59 |
| mammals | Golunda_ellioti | Golunda_ellioti | 0 | 60.9 | 4.5 | 2864575.38 |
| mammals | Gorilla_beringei | Gorilla_beringei | 1 | 130000 | 1 | 48938.49 |
| mammals | Gorilla_gorilla | Gorilla_gorilla | 1 | 120950 | 1.03 | 696516.85 |
| mammals | Gracilinanus_agilis | Gracilinanus_agilis | 0 | 22 | 11.75 | 3609141.76 |
| mammals | Gracilinanus_dryas | Gracilinanus_dryas | 0 | 18 | 3.5 | 30098.2 |
| mammals | Gracilinanus_marica | Gracilinanus_marica | 0 | 23.8 | 5.99 | 329096.25 |
| mammals | Gracilinanus_microtarsus | Gracilinanus_microtarsus | 0 | 31 | 9.7 | 631663.98 |
| mammals | Grammomys_cometes | Grammomys_cometes | 0 | 48.5 | 2.92 | 417998.04 |
| mammals | Grammomys_dolichurus | Grammomys_dolichurus | 0 | 41.2 | 3.06 | 3257251.68 |
| mammals | Grammomys_ibeanus | Grammomys_ibeanus | 0 | 41.2 | 3.5 | 215705.08 |
| mammals | Graomys_domorum | Graomys_domorum | 0 | 102 | 7.01 | 101529.24 |
| mammals | Graomys_griseoflavus | Graomys_griseoflavus | 0 | 67.5 | 6.99 | 2012497.83 |
| mammals | Graphiurus_christyi | Graphiurus_christyi | 0 | 22.7 | 4.99 | 394070.48 |
| mammals | Graphiurus_kelleni | Graphiurus_kelleni | 0 | 20.1 | 4.41 | 2773770.98 |
| mammals | Graphiurus_lorraineus | Graphiurus_lorraineus | 0 | 16.8 | 4.99 | 3472143.76 |
| mammals | Graphiurus_microtis | Graphiurus_microtis | 0 | 15.5 | 3 | 3300456.43 |
| mammals | Graphiurus_murinus | Graphiurus_murinus | 0 | 30 | 3 | 966155.44 |
| mammals | Graphiurus_nagtglasii | Graphiurus_nagtglasii | 0 | 45 | 3.11 | 882351.2 |
| mammals | Graphiurus_ocularis | Graphiurus_ocularis | 0 | 30.3 | 4.99 | 323125.65 |
| mammals | Graphiurus_platyops | Graphiurus_platyops | 0 | 46.1 | 5.17 | 340551.54 |
| mammals | Graphiurus_rupicola | Graphiurus_rupicola | 0 | 53.6 | 5.17 | 54377.7 |

| | | | | | | |
|---|---|---|---|---|---|---|
| mammals | Gulo_gulo | Gulo_gulo | 0 | 17012.6 | 2.84 | 24394335.16 |
| mammals | Gymnobelideus_leadbeateri | Gymnobelideus_leadbeateri | 1 | 127 | 1.54 | 3023.86 |
| mammals | Gymnuromys_roberti | Gymnuromys_roberti | 0 | 97.5 | 1.94 | 77428.38 |
| mammals | Habromys_ixtlani | Habromys_ixtlani | 1 | 93.4 | 1.9 | 92.15 |
| mammals | Habromys_lepturus | Habromys_lepturus | 1 | 85 | 1.9 | 86 |
| mammals | Handleyomys_alfaroi | Handleyomys_alfaroi | 0 | 33.3 | 3.66 | 838400.35 |
| mammals | Handleyomys_chapmani | Handleyomys_chapmani | 1 | 50 | 3.73 | 39012.72 |
| mammals | Handleyomys_fuscatus | Handleyomys_fuscatus | 0 | 49.5 | 2.5 | 79253.02 |
| mammals | Handleyomys_intectus | Handleyomys_intectus | 0 | 60.5 | 2 | 9776.98 |
| mammals | Handleyomys_melanotis | Handleyomys_melanotis | 0 | 50 | 3.88 | 138450.88 |
| mammals | Handleyomys_rhabdops | Handleyomys_rhabdops | 1 | 33 | 3.73 | 3686.43 |
| mammals | Handleyomys_rostratus | Handleyomys_rostratus | 0 | 43.5 | 3.88 | 548403.74 |
| mammals | Hapalemur_alaotrensis | Hapalemur_alaotrensis | 1 | 1240 | 1.2 | 216.33 |
| mammals | Hapalemur_aureus | Hapalemur_aureus | 1 | 1450 | 1.01 | 2392.79 |
| mammals | Hapalemur_griseus | Hapalemur_griseus | 1 | 900 | 1.35 | 90255.61 |
| mammals | Hapalemur_meridionalis | Hapalemur_meridionalis | 1 | 900 | 1.35 | 10993.97 |
| mammals | Hapalemur_occidentalis | Hapalemur_occidentalis | 1 | 850 | 1.2 | 26197.05 |
| mammals | Haplonycteris_fischeri | Haplonycteris_fischeri | 0 | 18 | 1 | 264779.28 |
| mammals | Harpiola_isodon | Harpiola_isodon | 0 | 5.7 | 1 | 12490.23 |
| mammals | Harpyionycteris_celebensis | Harpyionycteris_celebensis | 0 | 104 | 0.99 | 169411.15 |
| mammals | Harpyionycteris_whiteheadi | Harpyionycteris_whiteheadi | 0 | 110.5 | 1 | 171865.47 |
| mammals | Heimyscus_fumosus | Heimyscus_fumosus | 0 | 14.3 | 2.5 | 504708.99 |
| mammals | Helarctos_malayanus | Helarctos_malayanus | 1 | 46500 | 1.1 | 3561925.14 |
| mammals | Heliosciurus_gambianus | Heliosciurus_gambianus | 0 | 295 | 2.93 | 4714373.11 |
| mammals | Heliosciurus_mutabilis | Heliosciurus_mutabilis | 0 | 390 | 2.75 | 1063837.25 |
| mammals | Heliosciurus_rufobrachium | Heliosciurus_rufobrachium | 0 | 371.5 | 2.38 | 2935776.37 |
| mammals | Heliosciurus_ruwenzorii | Heliosciurus_ruwenzorii | 0 | 291 | 2.93 | 140580.95 |
| mammals | Helogale_parvula | Helogale_parvula | 0 | 300 | 3.34 | 6918503.91 |
| mammals | Hemibelideus_lemuroides | Hemibelideus_lemuroides | 0 | 952.5 | 1 | 2913.77 |
| mammals | Hemicentetes_nigriceps | Hemicentetes_nigriceps | 0 | 113 | 3.93 | 39236.49 |
| mammals | Hemicentetes_semispinosus | Hemicentetes_semispinosus | 0 | 137.8 | 5.24 | 122554.36 |
| mammals | Hemiechinus_auritus | Hemiechinus_auritus | 0 | 342 | 3.87 | 8465510.75 |
| mammals | Hemiechinus_collaris | Hemiechinus_collaris | 0 | 141 | 3.9 | 272308.57 |
| mammals | Hemigalus_derbyanus | Hemigalus_derbyanus | 0 | 2322.5 | 1.86 | 1354858.33 |

| | | | | | | |
|---|---|---|---|---|---|---|
| mammals | Hemitragus_jemlahicus | Hemitragus_jemlahicus | 0 | 35200 | 1.01 | 55836.04 |
| mammals | Herpailurus_yagouaroundi | Herpailurus_yagouaroundi | 0 | 7874.9 | 2.42 | 14850896.61 |
| mammals | Herpestes_ichneumon | Herpestes_ichneumon | 0 | 5175 | 2.98 | 14478898.03 |
| mammals | Herpestes_pulverulentus | Herpestes_pulverulentus | 0 | 797 | 2 | 770354.79 |
| mammals | Herpestes_sanguineus | Herpestes_sanguineus | 0 | 550 | 3 | 16793634.04 |
| mammals | Hesperoptenus_tickelli | Hesperoptenus_tickelli | 0 | 18 | 1 | 1051792.8 |
| mammals | Heterogeomys_cherriei | Heterogeomys_cherriei | 0 | 225 | 2.5 | 10525.05 |
| mammals | Heterogeomys_dariensis | Heterogeomys_dariensis | 0 | 438 | 2.5 | 13438.01 |
| mammals | Heterogeomys_hispidus | Orthogeomys_hispidus | 0 | 500 | 1.98 | 397264.66 |
| mammals | Heterohyrax_brucei | Heterohyrax_brucei | 0 | 2625 | 1.89 | 5937564.87 |
| mammals | Heteromys_adspersus | Heteromys_adspersus | 0 | 65 | 3.2 | 22429.24 |
| mammals | Heteromys_anomalus | Heteromys_anomalus | 0 | 70 | 2.62 | 449459.77 |
| mammals | Heteromys_australis | Heteromys_australis | 0 | 57 | 2.75 | 226886.78 |
| mammals | Heteromys_desmarestianus | Heteromys_desmarestianus | 0 | 72.5 | 3.12 | 476861.29 |
| mammals | Heteromys_gaumeri | Heteromys_gaumeri | 0 | 63.6 | 2.95 | 167156.99 |
| mammals | Heteromys_irroratus | Heteromys_irroratus | 0 | 50 | 4.16 | 606822.97 |
| mammals | Heteromys_oresterus | Heteromys_oresterus | 0 | 74.8 | 3.56 | 1327.43 |
| mammals | Heteromys_pictus | Heteromys_pictus | 0 | 40 | 3.63 | 439292.26 |
| mammals | Heteromys_salvini | Heteromys_salvini | 0 | 42 | 3.62 | 169498.09 |
| mammals | Heteromys_spectabilis | Heteromys_spectabilis | 1 | 65 | 5 | 4803.8 |
| mammals | Hippocamelus_antisensis | Hippocamelus_antisensis | 1 | 68599.4 | 1 | 424715.68 |
| mammals | Hippocamelus_bisulcus | Hippocamelus_bisulcus | 1 | 70000.3 | 1 | 129675.54 |
| mammals | Hippopotamus_amphibius | Hippopotamus_amphibius | 1 | 1417490 | 1 | 1233077.11 |
| mammals | Hipposideros_abae | Hipposideros_abae | 0 | 25.5 | 1.5 | 2324228.53 |
| mammals | Hipposideros_armiger | Hipposideros_armiger | 0 | 53 | 1 | 4499367.47 |
| mammals | Hipposideros_ater | Hipposideros_ater | 0 | 5.8 | 0.99 | 3676234.68 |
| mammals | Hipposideros_beatus | Hipposideros_beatus | 0 | 6.7 | 1 | 1636043.8 |
| mammals | Hipposideros_bicolor | Hipposideros_bicolor | 0 | 7.6 | 0.99 | 1591830.81 |
| mammals | Hipposideros_caffer | Hipposideros_caffer | 0 | 9.4 | 0.99 | 1264715.77 |
| mammals | Hipposideros_calcaratus | Hipposideros_calcaratus | 0 | 11.4 | 1.5 | 217484.07 |
| mammals | Hipposideros_cervinus | Hipposideros_cervinus | 0 | 9.3 | 1 | 1964927.01 |
| mammals | Hipposideros_cineraceus | Hipposideros_cineraceus | 0 | 4.7 | 0.98 | 2205964.58 |
| mammals | Hipposideros_corynophyllus | Hipposideros_corynophyllus | 0 | 14.5 | 1 | 1988.09 |
| mammals | Hipposideros_coxi | Hipposideros_coxi | 1 | 9.3 | 1 | 2684.57 |

| | | | | | | |
|---|---|---|---|---|---|---|
| mammals | Hipposideros_curtus | Hipposideros_curtus | 1 | 7.5 | 1 | 136020.66 |
| mammals | Hipposideros_demissus | Hipposideros_demissus | 1 | 24.9 | 1 | 3048.08 |
| mammals | Hipposideros_diadema | Hipposideros_diadema | 0 | 43.8 | 1 | 3318837.89 |
| mammals | Hipposideros_dinops | Hipposideros_dinops | 1 | 71.9 | 1 | 27145.99 |
| mammals | Hipposideros_doriae | Hipposideros_doriae | 0 | 5 | 1 | 1251661.75 |
| mammals | Hipposideros_durgadasi | Hipposideros_durgadasi | 1 | 5.6 | 0.98 | 19888.86 |
| mammals | Hipposideros_dyacorum | Hipposideros_dyacorum | 0 | 6.1 | 1 | 764978.9 |
| mammals | Hipposideros_edwardshilli | Hipposideros_edwardshilli | 1 | 11.3 | 1 | 1985.79 |
| mammals | Hipposideros_fuliginosus | Hipposideros_fuliginosus | 0 | 16.6 | 1 | 1334587.14 |
| mammals | Hipposideros_fulvus | Hipposideros_fulvus | 0 | 8.5 | 1 | 2455054.71 |
| mammals | Hipposideros_galeritus | Hipposideros_galeritus | 0 | 8.2 | 0.99 | 2775105.77 |
| mammals | Hipposideros_grandis | Hipposideros_grandis | 0 | 16.3 | 0.98 | 185987.16 |
| mammals | Hipposideros_halophyllus | Hipposideros_halophyllus | 1 | 4 | 1 | 23215.65 |
| mammals | Hipposideros_inornatus | Hipposideros_inornatus | 1 | 28.5 | 1 | 6625.21 |
| mammals | Hipposideros_jonesi | Hipposideros_jonesi | 0 | 5.5 | 1 | 1271130.07 |
| mammals | Hipposideros_khaokhouayensis | Hipposideros_khaokhouayensis | 1 | 8.7 | 1 | 17926.37 |
| mammals | Hipposideros_lamottei | Hipposideros_lamottei | 1 | 13.1 | 1 | 77.52 |
| mammals | Hipposideros_lankadiva | Hipposideros_lankadiva | 0 | 52.4 | 0.99 | 1547925.26 |
| mammals | Hipposideros_larvatus | Hipposideros_larvatus | 0 | 19.9 | 0.98 | 3295221.61 |
| mammals | Hipposideros_lekaguli | Hipposideros_lekaguli | 0 | 31.1 | 1 | 96007.54 |
| mammals | Hipposideros_lylei | Hipposideros_lylei | 0 | 40 | 1 | 1013744.15 |
| mammals | Hipposideros_madurae | Hipposideros_madurae | 0 | 13.9 | 0.98 | 66492.71 |
| mammals | Hipposideros_maggietaylorae | Hipposideros_maggietaylorae | 0 | 18.5 | 1 | 263505.96 |
| mammals | Hipposideros_marisae | Hipposideros_marisae | 1 | 6.1 | 1 | 49715.66 |
| mammals | Hipposideros_megalotis | Hipposideros_megalotis | 0 | 5.6 | 1 | 943711.9 |
| mammals | Hipposideros_muscinus | Hipposideros_muscinus | 0 | 8.6 | 1 | 116471.97 |
| mammals | Hipposideros_obscurus | Hipposideros_obscurus | 0 | 9.5 | 1 | 276180.19 |
| mammals | Hipposideros_orbiculus | Hipposideros_orbiculus | 1 | 9.8 | 1 | 1922.51 |
| mammals | Hipposideros_papua | Hipposideros_papua | 0 | 10.1 | 1 | 50777.06 |
| mammals | Hipposideros_pelingensis | Hipposideros_pelingensis | 0 | 56.2 | 1 | 177328.16 |
| mammals | Hipposideros_pomona | Hipposideros_pomona | 1 | 6.2 | 0.99 | 26967.91 |
| mammals | Hipposideros_pygmaeus | Hipposideros_pygmaeus | 0 | 3.5 | 1 | 90619.52 |
| mammals | Hipposideros_ridleyi | Hipposideros_ridleyi | 1 | 8.8 | 0.98 | 218978.28 |
| mammals | Hipposideros_rotalis | Hipposideros_rotalis | 0 | 9.3 | 1 | 29363.54 |

| | | | | | | |
|---|---|---|---|---|---|---|
| mammals | Hipposideros_ruber | Hipposideros_ruber | 0 | 10.1 | 1 | 9062130.35 |
| mammals | Hipposideros_scutinares | Hipposideros_scutinares | 1 | 40 | 1 | 34112.18 |
| mammals | Hipposideros_semoni | Hipposideros_semoni | 0 | 7.5 | 1 | 76340.62 |
| mammals | Hipposideros_sorenseni | Hipposideros_sorenseni | 1 | 20.2 | 1 | 408.88 |
| mammals | Hipposideros_speoris | Hipposideros_speoris | 0 | 10.4 | 0.99 | 1221761.43 |
| mammals | Hipposideros_stenotis | Hipposideros_stenotis | 1 | 12 | 1 | 489413.65 |
| mammals | Hipposideros_sumbae | Hipposideros_sumbae | 0 | 13.3 | 0.98 | 43581.58 |
| mammals | Hipposideros_swinhoii | Hipposideros_pratti | 0 | 56 | 1 | 2242954.88 |
| mammals | Hipposideros_turpis | Hipposideros_turpis | 1 | 33.4 | 1 | 553.87 |
| mammals | Hipposideros_wollastoni | Hipposideros_wollastoni | 0 | 6.8 | 1 | 33302.6 |
| mammals | Hippotragus_equinus | Hippotragus_equinus | 0 | 269997.7 | 0.99 | 5571099.89 |
| mammals | Hippotragus_niger | Hippotragus_niger | 0 | 227499.3 | 1 | 3013917.71 |
| mammals | Histiotus_laephotis | Histiotus_laephotis | 0 | 11.3 | 0.98 | 823944.65 |
| mammals | Histiotus_macrotus | Histiotus_macrotus | 0 | 11 | 0.98 | 462171.76 |
| mammals | Histiotus_magellanicus | Histiotus_magellanicus | 0 | 14 | 1 | 330089.68 |
| mammals | Histiotus_montanus | Histiotus_montanus | 0 | 9.1 | 1 | 3290996.15 |
| mammals | Hodomys_alleni | Hodomys_alleni | 0 | 367.6 | 1.54 | 214221.36 |
| mammals | Holochilus_brasiliensis | Holochilus_brasiliensis | 0 | 155 | 3.66 | 2878494.86 |
| mammals | Holochilus_chacarius | Holochilus_chacarius | 0 | 204 | 3.44 | 766923.36 |
| mammals | Holochilus_sciureus | Holochilus_sciureus | 0 | 171 | 4.06 | 9320092.42 |
| mammals | Hoolock_hoolock | Hoolock_hoolock | 1 | 6500 | 1 | 319902.68 |
| mammals | Hoolock_leuconedys | Hoolock_leuconedys | 1 | 7200 | 1 | 112318.95 |
| mammals | Hoplomys_gymnurus | Hoplomys_gymnurus | 0 | 240 | 2.1 | 332781.88 |
| mammals | Hsunycteris_thomasi | Lonchophylla_thomasi | 0 | 6.9 | 0.98 | 8383354.14 |
| mammals | Hyaena_hyaena | Hyaena_hyaena | 0 | 41705.1 | 2.47 | 23460484.54 |
| mammals | Hybomys_basilii | Hybomys_basilii | 1 | 53.4 | 2.4 | 588.93 |
| mammals | Hybomys_lunaris | Hybomys_lunaris | 1 | 53.4 | 2.4 | 2216.95 |
| mammals | Hybomys_univittatus | Hybomys_univittatus | 0 | 47.6 | 2.35 | 2240781.54 |
| mammals | Hydrictis_maculicollis | Hydrictis_maculicollis | 0 | 4000 | 2 | 9962749.18 |
| mammals | Hydrochoerus_hydrochaeris | Hydrochoerus_hydrochaeris | 0 | 62449.6 | 3.49 | 13395889.23 |
| mammals | Hydromys_chrysogaster | Hydromys_chrysogaster | 0 | 479.8 | 3.65 | 3705391.86 |
| mammals | Hydropotes_inermis | Hydropotes_inermis | 1 | 14000 | 3 | 181922.92 |
| mammals | Hyemoschus_aquaticus | Hyemoschus_aquaticus | 0 | 10850 | 1.23 | 2450418.94 |
| mammals | Hylaeamys_megacephalus | Hylaeamys_megacephalus | 0 | 57.8 | 3.87 | 5971593.79 |

| | | | | | | |
|---|---|---|---|---|---|---|
| mammals | Hylaeamys_perenensis | Hylaeamys_perenensis | 0 | 60.6 | 4.02 | 1991968.51 |
| mammals | Hylaeamys_yunganus | Hylaeamys_yunganus | 0 | 60.5 | 2 | 5299586.68 |
| mammals | Hylobates_agilis | Hylobates_agilis | 1 | 5650 | 1.01 | 385727.49 |
| mammals | Hylobates_albibarbis | Hylobates_albibarbis | 1 | 5650 | 1 | 199695.86 |
| mammals | Hylobates_klossii | Hylobates_klossii | 1 | 5800 | 1 | 6004.48 |
| mammals | Hylobates_lar | Hylobates_lar | 1 | 4892 | 1 | 638118.63 |
| mammals | Hylobates_moloch | Hylobates_moloch | 1 | 6500 | 1 | 39231.97 |
| mammals | Hylobates_muelleri | Hylobates_muelleri | 1 | 6000 | 1.01 | 103190.46 |
| mammals | Hylobates_pileatus | Hylobates_pileatus | 1 | 5440 | 1 | 121563.53 |
| mammals | Hylochoerus_meinertzhageni | Hylochoerus_meinertzhageni | 0 | 188500 | 4.18 | 2149889.59 |
| mammals | Hylomys_parvus | Hylomys_parvus | 1 | 38.3 | 2.36 | 9948.26 |
| mammals | Hylomys_suillus | Hylomys_suillus | 0 | 65.8 | 2.33 | 1634250.04 |
| mammals | Hylomyscus_aeta | Hylomyscus_aeta | 0 | 25.1 | 4.18 | 2618038.49 |
| mammals | Hylomyscus_alleni | Hylomyscus_alleni | 0 | 18.7 | 3 | 1510866.93 |
| mammals | Hylomyscus_denniae | Hylomyscus_denniae | 0 | 30 | 4.61 | 17471.52 |
| mammals | Hylomyscus_grandis | Hylomyscus_grandis | 1 | 33 | 4.18 | 422.34 |
| mammals | Hylomyscus_parvus | Hylomyscus_parvus | 0 | 12.7 | 2.5 | 491928.87 |
| mammals | Hylomyscus_stella | Hylomyscus_stella | 0 | 20.2 | 3.08 | 89591.25 |
| mammals | Hylomyscus_walterverheyeni | Hylomyscus_walterverheyeni | 0 | 20 | 3.6 | 253573.05 |
| mammals | Hylonycteris_underwoodi | Hylonycteris_underwoodi | 0 | 6.7 | 1 | 201148.61 |
| mammals | Hylopetes_alboniger | Hylopetes_alboniger | 0 | 255.4 | 2.28 | 1399086.79 |
| mammals | Hyomys_goliath | Hyomys_goliath | 0 | 817.5 | 1 | 110796.73 |
| mammals | Hyosciurus_heinrichi | Hyosciurus_heinrichi | 0 | 296 | 1.5 | 32128.58 |
| mammals | Hypogeomys_antimena | Hypogeomys_antimena | 1 | 1250 | 1.13 | 193.69 |
| mammals | Hypsignathus_monstrosus | Hypsignathus_monstrosus | 0 | 335.1 | 1.4 | 2860674.04 |
| mammals | Hypsiprymnodon_moschatus | Hypsiprymnodon_moschatus | 0 | 520 | 2.04 | 10503.39 |
| mammals | Hypsugo_alaschanicus | Pipistrellus_alaschanicus | 0 | 8.2 | 2 | 2990994.15 |
| mammals | Hypsugo_pulveratus | Pipistrellus_pulveratus | 0 | 5.5 | 2 | 2653163.51 |
| mammals | Hypsugo_savii | Hypsugo_savii | 0 | 6.4 | 2 | 4335433.91 |
| mammals | Hystrix_africaeaustralis | Hystrix_africaeaustralis | 0 | 14936 | 1.8 | 7623760.02 |
| mammals | Hystrix_brachyura | Hystrix_brachyura | 0 | 8450 | 1.43 | 5786235.96 |
| mammals | Hystrix_cristata | Hystrix_cristata | 0 | 16250 | 2.02 | 5343513.37 |
| mammals | Hystrix_indica | Hystrix_indica | 0 | 12435.8 | 1.86 | 7063667.05 |
| mammals | Hystrix_javanica | Hystrix_javanica | 0 | 23014.4 | 1.46 | 168922.88 |

| | | | | | | |
|---|---|---|---|---|---|---|
| mammals | Ia_io | Ia_io | 0 | 49.2 | 1 | 1506489.99 |
| mammals | Ichneumia_albicauda | Ichneumia_albicauda | 0 | 3500 | 2.34 | 14289624.18 |
| mammals | Ichthyomys_pittieri | Ichthyomys_pittieri | 0 | 69.1 | 4 | 28379.94 |
| mammals | Ictidomys_mexicanus | Ictidomys_mexicanus | 0 | 189.6 | 5.91 | 715736.91 |
| mammals | Ictidomys_tridecemlineatus | Ictidomys_tridecemlineatus | 0 | 131.7 | 8.09 | 3679544.74 |
| mammals | Ictonyx_striatus | Ictonyx_striatus | 0 | 1300 | 2.23 | 15345019.39 |
| mammals | Idionycteris_phyllotis | Idionycteris_phyllotis | 0 | 12 | 1 | 1390365.36 |
| mammals | Idiurus_macrotis | Idiurus_macrotis | 0 | 30 | 0.97 | 619575.85 |
| mammals | Idiurus_zenkeri | Idiurus_zenkeri | 0 | 100 | 1 | 277784.74 |
| mammals | Indri_indri | Indri_indri | 1 | 8000 | 1.01 | 30054.88 |
| mammals | Iomys_horsfieldii | Iomys_horsfieldii | 0 | 147.5 | 2.58 | 806688.74 |
| mammals | Iomys_sipora | Iomys_sipora | 1 | 152.3 | 2.51 | 1242.49 |
| mammals | Irenomys_tarsalis | Irenomys_tarsalis | 0 | 43.2 | 3.85 | 145039.91 |
| mammals | Isoodon_auratus | Isoodon_auratus | 1 | 425 | 2.6 | 20997.78 |
| mammals | Isoodon_macrourus | Isoodon_macrourus | 0 | 1600 | 3.35 | 1220483.4 |
| mammals | Isoodon_obesulus | Isoodon_obesulus | 0 | 775 | 2.43 | 203133.14 |
| mammals | Isothrix_bistriata | Isothrix_bistriata | 0 | 445 | 0.97 | 1821069.62 |
| mammals | Isothrix_negrensis | Isothrix_negrensis | 0 | 467.7 | 0.97 | 294631.84 |
| mammals | Isthmomys_pirrensis | Isthmomys_pirrensis | 0 | 138 | 2.67 | 5206.22 |
| mammals | Jaculus_blanfordi | Jaculus_blanfordi | 0 | 54.7 | 4.04 | 1305249.84 |
| mammals | Jaculus_jaculus | Jaculus_jaculus | 0 | 56.6 | 3.24 | 8111416.6 |
| mammals | Jaculus_orientalis | Jaculus_orientalis | 0 | 138 | 2.46 | 434055.5 |
| mammals | Juliomys_pictipes | Juliomys_pictipes | 0 | 22.9 | 3 | 437708.82 |
| mammals | Kadarsanomys_sodyi | Kadarsanomys_sodyi | 1 | 199.5 | 4 | 265.89 |
| mammals | Kannabateomys_amblyonyx | Kannabateomys_amblyonyx | 0 | 600 | 1 | 916239.9 |
| mammals | Kerivoula_flora | Kerivoula_flora | 1 | 6 | 1 | 49016.83 |
| mammals | Kerivoula_hardwickii | Kerivoula_hardwickii | 0 | 4.6 | 1 | 3435503.96 |
| mammals | Kerivoula_lanosa | Kerivoula_lanosa | 0 | 6.5 | 0.99 | 9072400.79 |
| mammals | Kerivoula_lenis | Kerivoula_lenis | 0 | 6.8 | 0.98 | 129194.29 |
| mammals | Kerivoula_papillosa | Kerivoula_papillosa | 0 | 9.8 | 0.98 | 789927.98 |
| mammals | Kerivoula_pellucida | Kerivoula_pellucida | 0 | 4.1 | 1 | 1213322.06 |
| mammals | Kerivoula_picta | Kerivoula_picta | 0 | 4.5 | 1 | 2925844.48 |
| mammals | Kerivoula_smithii | Kerivoula_smithii | 0 | 13.1 | 0.98 | 1100356.07 |
| mammals | Kerodon_rupestris | Kerodon_rupestris | 0 | 800 | 1.52 | 708728.58 |

| | | | | | | |
|---|---|---|---|---|---|---|
| mammals | Kobus_ellipsiprymnus | Kobus_ellipsiprymnus | 0 | 210000.3 | 1.02 | 8949699.06 |
| mammals | Kobus_kob | Kobus_kob | 0 | 78500.1 | 1 | 3004386.39 |
| mammals | Kobus_leche | Kobus_leche | 0 | 110749 | 1 | 202159.55 |
| mammals | Kobus_megaceros | Kobus_megaceros | 1 | 87500.4 | 1 | 110602.97 |
| mammals | Kobus_vardonii | Kobus_vardonii | 0 | 71500.7 | 1 | 376058.86 |
| mammals | Kunsia_tomentosus | Kunsia_tomentosus | 0 | 115.6 | 2 | 405854.33 |
| mammals | Laephotis_capensis | Neoromicia_capensis | 0 | 6 | 1.96 | 13760702.43 |
| mammals | Laephotis_malagasyensis | Neoromicia_malagasyensis | 1 | 6.5 | 2 | 7033.93 |
| mammals | Laephotis_matroka | Neoromicia_matroka | 0 | 6.5 | 1.22 | 33499.49 |
| mammals | Laephotis_wintoni | Laephotis_wintoni | 0 | 6 | 1 | 1918973.17 |
| mammals | Lagidium_viscacia | Lagidium_viscacia | 0 | 1540 | 1.25 | 1776083.67 |
| mammals | Lagorchestes_conspicillatus | Lagorchestes_conspicillatus | 0 | 3050 | 1 | 1884281.95 |
| mammals | Lagorchestes_hirsutus | Lagorchestes_hirsutus | 1 | 1265 | 1 | 95.23 |
| mammals | Lagostomus_maximus | Lagostomus_maximus | 0 | 4647.5 | 1.97 | 1466480.45 |
| mammals | Lagostrophus_fasciatus | Lagostrophus_fasciatus | 1 | 1700 | 0.9 | 137.02 |
| mammals | Lagothrix_flavicauda | Oreonax_flavicauda | 1 | 6800 | 1 | 24210.46 |
| mammals | Lagothrix_lagothricha | Lagothrix_lagotricha | 1 | 6788.9 | 1 | 2568088.69 |
| mammals | Lagurus_lagurus | Lagurus_lagurus | 0 | 20.3 | 4.25 | 4713486.43 |
| mammals | Lama_guanicoe | Lama_guanicoe | 0 | 120000 | 1 | 1658706.17 |
| mammals | Lama_vicugna | Vicugna_vicugna | 0 | 47499.6 | 1 | 619657.91 |
| mammals | Lampronycteris_brachyotis | Lampronycteris_brachyotis | 0 | 11 | 0.98 | 4319292.52 |
| mammals | Laonastes_aenigmamus | Laonastes_aenigmamus | 0 | 375 | 1.3 | 6488.2 |
| mammals | Lariscus_hosei | Lariscus_hosei | 0 | 170.7 | 1.96 | 52688.43 |
| mammals | Lariscus_insignis | Lariscus_insignis | 0 | 175 | 2 | 1276285.79 |
| mammals | Lariscus_obscurus | Lariscus_obscurus | 0 | 188.3 | 2 | 6004.26 |
| mammals | Lasionycteris_noctivagans | Lasionycteris_noctivagans | 0 | 10.5 | 1.83 | 10110807.92 |
| mammals | Lasiopodomys_brandtii | Lasiopodomys_brandtii | 0 | 93.8 | 6 | 819069.45 |
| mammals | Lasiorhinus_krefftii | Lasiorhinus_krefftii | 1 | 31450 | 1 | 91.21 |
| mammals | Lasiurus_blossevillii | Lasiurus_blossevillii | 0 | 13 | 3.06 | 18711014.82 |
| mammals | Lasiurus_borealis | Lasiurus_borealis | 0 | 10.7 | 3.09 | 4881497.8 |
| mammals | Lasiurus_cinereus | Lasiurus_cinereus | 0 | 25.3 | 1.98 | 21071173.08 |
| mammals | Lasiurus_degelidus | Lasiurus_degelidus | 1 | 12.8 | 3.06 | 11021.01 |
| mammals | Lasiurus_ega | Lasiurus_ega | 0 | 13.2 | 2.1 | 13205185.72 |
| mammals | Lasiurus_insularis | Lasiurus_insularis | 1 | 31.6 | 2.98 | 106633.25 |

| | | | | | | |
|---|---|---|---|---|---|---|
| mammals | Lasiurus_intermedius | Lasiurus_intermedius | 0 | 23 | 2.98 | 1748790.82 |
| mammals | Lasiurus_minor | Lasiurus_minor | 1 | 9.4 | 3.12 | 84473.91 |
| mammals | Lasiurus_pfeifferi | Lasiurus_pfeifferi | 0 | 11.1 | 3.12 | 96779.23 |
| mammals | Lasiurus_seminolus | Lasiurus_seminolus | 0 | 9.9 | 2.9 | 1193528.48 |
| mammals | Lasiurus_varius | Lasiurus_varius | 0 | 11.4 | 3.06 | 545210.34 |
| mammals | Lasiurus_xanthinus | Lasiurus_xanthinus | 0 | 10.7 | 2.07 | 1737395.75 |
| mammals | Latidens_salimalii | Latidens_salimalii | 1 | 55.2 | 1 | 18947.75 |
| mammals | Lavia_frons | Lavia_frons | 0 | 23.9 | 1 | 7569286.87 |
| mammals | Leggadina_forresti | Leggadina_forresti | 0 | 20 | 3.5 | 1981608.76 |
| mammals | Leggadina_lakedownensis | Leggadina_lakedownensis | 0 | 17.5 | 3 | 1323526.63 |
| mammals | Lemmiscus_curtatus | Lemmiscus_curtatus | 0 | 28.3 | 5.36 | 1662224.71 |
| mammals | Lemmus_amurensis | Lemmus_amurensis | 0 | 33.6 | 5.77 | 2123204.58 |
| mammals | Lemmus_lemmus | Lemmus_lemmus | 0 | 47.5 | 5.11 | 758090.79 |
| mammals | Lemniscomys_barbarus | Lemniscomys_barbarus | 0 | 49 | 4.61 | 249140.73 |
| mammals | Lemniscomys_griselda | Lemniscomys_griselda | 0 | 58 | 3.57 | 599556.14 |
| mammals | Lemniscomys_linulus | Lemniscomys_linulus | 0 | 27 | 3.46 | 266618.2 |
| mammals | Lemniscomys_macculus | Lemniscomys_macculus | 0 | 30.3 | 6 | 992343.25 |
| mammals | Lemniscomys_rosalia | Lemniscomys_rosalia | 0 | 54.2 | 3.46 | 2429390.55 |
| mammals | Lemniscomys_striatus | Lemniscomys_striatus | 0 | 41.6 | 4.59 | 4422594.81 |
| mammals | Lemniscomys_zebra | Lemniscomys_zebra | 0 | 36.1 | 4.61 | 3840518.97 |
| mammals | Lemur_catta | Lemur_catta | 1 | 2900 | 1.14 | 99812.55 |
| mammals | Lenomys_meyeri | Lenomys_meyeri | 0 | 480 | 1 | 116927.09 |
| mammals | Lenothrix_cana | Lenothrix_canus | 0 | 135 | 2.91 | 238775.3 |
| mammals | Leontopithecus_caissara | Leontopithecus_caissara | 1 | 572 | 1.95 | 205.64 |
| mammals | Leontopithecus_chrysomelas | Leontopithecus_chrysomelas | 1 | 700 | 2 | 20187.3 |
| mammals | Leontopithecus_chrysopygus | Leontopithecus_chrysopygus | 1 | 700 | 1.95 | 63670.79 |
| mammals | Leontopithecus_rosalia | Leontopithecus_rosalia | 1 | 535.5 | 1.95 | 3990.55 |
| mammals | Leopardus_colocola | Leopardus_colocolo | 0 | 3935 | 2.5 | 4786752.18 |
| mammals | Leopardus_geoffroyi | Leopardus_geoffroyi | 0 | 2230 | 2.5 | 3529650.06 |
| mammals | Leopardus_guigna | Leopardus_guigna | 1 | 3590 | 2.02 | 317797.04 |
| mammals | Leopardus_guttulus | Leopardus_guttulus | 1 | 2500 | 1 | 2146517.13 |
| mammals | Leopardus_pardalis | Leopardus_pardalis | 0 | 11900.1 | 1.66 | 14546411.35 |
| mammals | Leopardus_tigrinus | Leopardus_tigrinus | 1 | 2250 | 1.5 | 9389963.34 |
| mammals | Leopardus_wiedii | Leopardus_wiedii | 0 | 3250 | 1.5 | 12288755.81 |

| | | | | | | |
|---|---|---|---|---|---|---|
| mammals | Leopoldamys_ciliatus | Leopoldamys_ciliatus | 0 | 362.5 | 1.94 | 199687.85 |
| mammals | Leopoldamys_edwardsi | Leopoldamys_edwardsi | 0 | 300 | 1.94 | 2061787.69 |
| mammals | Leopoldamys_milleti | Leopoldamys_milleti | 0 | 410.8 | 1.94 | 5654.28 |
| mammals | Leopoldamys_sabanus | Leopoldamys_sabanus | 0 | 338.3 | 3.33 | 2538761.32 |
| mammals | Lepilemur_ankaranensis | Lepilemur_ankaranensis | 1 | 750 | 1 | 1576.6 |
| mammals | Lepilemur_dorsalis | Lepilemur_dorsalis | 1 | 855 | 1 | 4754.32 |
| mammals | Lepilemur_edwardsi | Lepilemur_edwardsi | 1 | 877.5 | 1 | 9054.27 |
| mammals | Lepilemur_leucopus | Lepilemur_leucopus | 1 | 525 | 1 | 1953.79 |
| mammals | Lepilemur_microdon | Lepilemur_microdon | 1 | 900 | 1 | 2505.77 |
| mammals | Lepilemur_mustelinus | Lepilemur_mustelinus | 1 | 1000 | 1 | 12959.45 |
| mammals | Lepilemur_ruficaudatus | Lepilemur_ruficaudatus | 1 | 750 | 1.01 | 15786.76 |
| mammals | Lepilemur_sahamalaza | Lepilemur_sahamalazensis | 1 | 789.5 | 1 | 436.88 |
| mammals | Lepilemur_septentrionalis | Lepilemur_septentrionalis | 1 | 750 | 1 | 139.31 |
| mammals | Leporillus_conditor | Leporillus_conditor | 0 | 350 | 2.58 | 705.74 |
| mammals | Leptailurus_serval | Leptailurus_serval | 0 | 12000 | 2.35 | 12952360.5 |
| mammals | Leptomys_elegans | Leptomys_elegans | 0 | 84.8 | 1.88 | 73683.01 |
| mammals | Leptomys_ernstmayri | Leptomys_ernstmayri | 0 | 42.9 | 1.88 | 49834.37 |
| mammals | Leptomys_signatus | Leptomys_signatus | 0 | 54.4 | 1.88 | 764.47 |
| mammals | Leptonycteris_curasoae | Leptonycteris_curasoae | 1 | 23.4 | 0.99 | 840356.5 |
| mammals | Leptonycteris_nivalis | Leptonycteris_nivalis | 1 | 25.6 | 1 | 949993.75 |
| mammals | Leptonycteris_yerbabuenae | Leptonycteris_yerbabuenae | 0 | 20 | 1 | 1233797.18 |
| mammals | Lepus_alleni | Lepus_alleni | 0 | 3685 | 2.02 | 233589.8 |
| mammals | Lepus_americanus | Lepus_americanus | 0 | 1710 | 3.48 | 9597254.59 |
| mammals | Lepus_arcticus | Lepus_arcticus | 0 | 4405 | 4.79 | 2556757.85 |
| mammals | Lepus_brachyurus | Lepus_brachyurus | 0 | 2350 | 2.13 | 286560.64 |
| mammals | Lepus_californicus | Lepus_californicus | 0 | 2422.5 | 2.68 | 4392320.27 |
| mammals | Lepus_callotis | Lepus_callotis | 1 | 2500 | 2.41 | 378241.51 |
| mammals | Lepus_capensis | Lepus_capensis | 0 | 3261.7 | 2.48 | 11230966.34 |
| mammals | Lepus_castroviejoi | Lepus_castroviejoi | 1 | 2750 | 1.95 | 4980.05 |
| mammals | Lepus_comus | Lepus_comus | 0 | 1460 | 2.6 | 418232.27 |
| mammals | Lepus_corsicanus | Lepus_corsicanus | 1 | 3000 | 2.44 | 114558.64 |
| mammals | Lepus_europaeus | Lepus_europaeus | 0 | 3740 | 2.29 | 14444573.95 |
| mammals | Lepus_flavigularis | Lepus_flavigularis | 1 | 3000 | 2.34 | 100 |
| mammals | Lepus_granatensis | Lepus_granatensis | 0 | 4010 | 1.95 | 533085.02 |

| | | | | | | |
|---|---|---|---|---|---|---|
| mammals | Lepus_habessinicus | Lepus_habessinicus | 0 | 1962.1 | 2.51 | 1258924.32 |
| mammals | Lepus_mandshuricus | Lepus_mandshuricus | 0 | 1800 | 2.33 | 1091802.07 |
| mammals | Lepus_microtis | Lepus_victoriae | 0 | 1750 | 1.56 | 11320167.71 |
| mammals | Lepus_nigricollis | Lepus_nigricollis | 0 | 2298.5 | 1.68 | 3430852.14 |
| mammals | Lepus_oiostolus | Lepus_oiostolus | 0 | 3000 | 3.06 | 2759213.21 |
| mammals | Lepus_othus | Lepus_othus | 0 | 4806 | 5.06 | 203837.72 |
| mammals | Lepus_peguensis | Lepus_peguensis | 0 | 2105 | 3 | 1243043.05 |
| mammals | Lepus_saxatilis | Lepus_saxatilis | 0 | 2363.3 | 1.83 | 193852.4 |
| mammals | Lepus_sinensis | Lepus_sinensis | 0 | 1510 | 3 | 957212.13 |
| mammals | Lepus_starcki | Lepus_starcki | 0 | 2750 | 2.51 | 60357.6 |
| mammals | Lepus_tibetanus | Lepus_tibetanus | 0 | 2062.5 | 2.76 | 2153563.31 |
| mammals | Lepus_timidus | Lepus_timidus | 0 | 3048 | 3.16 | 20131533.22 |
| mammals | Lepus_tolai | Lepus_tolai | 0 | 1589 | 2.84 | 9351540.87 |
| mammals | Lepus_townsendii | Lepus_townsendii | 0 | 1555 | 4.45 | 3453856.92 |
| mammals | Lepus_yarkandensis | Lepus_yarkandensis | 0 | 1500 | 3.9 | 581120.53 |
| mammals | Lestoros_inca | Lestoros_inca | 0 | 21.1 | 4 | 62779.89 |
| mammals | Leucocephalophus_adersi | Cephalophus_adersi | 1 | 9250 | 1 | 5035.2 |
| mammals | Lichonycteris_obscura | Lichonycteris_obscura | 0 | 5.6 | 0.98 | 696682.6 |
| mammals | Limnomys_bryophilus | Limnomys_bryophilus | 0 | 64.1 | 1.5 | 1479.53 |
| mammals | Lionycteris_spurrelli | Lionycteris_spurrelli | 0 | 8.4 | 0.98 | 7274395.19 |
| mammals | Litocranius_walleri | Litocranius_walleri | 0 | 37999.7 | 1 | 1418356.42 |
| mammals | Lonchophylla_concava | Lonchophylla_concava | 0 | 7.4 | 0.98 | 206491.24 |
| mammals | Lonchophylla_mordax | Lonchophylla_mordax | 0 | 9 | 0.98 | 20483.63 |
| mammals | Lonchophylla_robusta | Lonchophylla_robusta | 0 | 17.2 | 0.98 | 1116127.27 |
| mammals | Lonchorhina_aurita | Lonchorhina_aurita | 0 | 15.7 | 0.98 | 11577000.21 |
| mammals | Lonchorhina_marinkellei | Lonchorhina_marinkellei | 1 | 17 | 1 | 161297.63 |
| mammals | Lonchothrix_emiliae | Lonchothrix_emiliae | 0 | 138.2 | 2 | 378168.06 |
| mammals | Lontra_canadensis | Lontra_canadensis | 0 | 8087.4 | 2.67 | 19086449.76 |
| mammals | Lontra_longicaudis | Lontra_longicaudis | 0 | 6555 | 3.39 | 15024030.12 |
| mammals | Lontra_provocax | Lontra_provocax | 1 | 7500 | 1.5 | 125850.09 |
| mammals | Lophiomys_imhausi | Lophiomys_imhausi | 0 | 755 | 1.67 | 424237.65 |
| mammals | Lophocebus_albigena | Lophocebus_albigena | 1 | 6984.8 | 1 | 1508231.45 |
| mammals | Lophocebus_aterrimus | Lophocebus_aterrimus | 1 | 6984.8 | 1 | 706964.32 |
| mammals | Lophostoma_brasiliense | Lophostoma_brasiliense | 0 | 9.4 | 0.98 | 8095483.47 |

| | | | | | | |
|---|---|---|---|---|---|---|
| mammals | Lophostoma_evotis | Lophostoma_evotis | 0 | 19.8 | 1 | 176592.76 |
| mammals | Lophostoma_silvicola | Lophostoma_silvicolum | 0 | 29.7 | 0.99 | 7691508.73 |
| mammals | Lophuromys_brevicaudus | Lophuromys_brevicaudus | 0 | 43.1 | 2.31 | 13838.89 |
| mammals | Lophuromys_chrysopus | Lophuromys_chrysopus | 0 | 43.1 | 2.31 | 112209.39 |
| mammals | Lophuromys_eisentrauti | Lophuromys_eisentrauti | 1 | 30 | 3.25 | 55.88 |
| mammals | Lophuromys_flavopunctatus | Lophuromys_flavopunctatus | 0 | 60 | 2.3 | 1820860.79 |
| mammals | Lophuromys_luteogaster | Lophuromys_luteogaster | 0 | 34 | 1.25 | 63598.45 |
| mammals | Lophuromys_medicaudatus | Lophuromys_medicaudatus | 1 | 35 | 1.5 | 7510.56 |
| mammals | Lophuromys_melanonyx | Lophuromys_melanonyx | 1 | 102.1 | 1.9 | 10810.37 |
| mammals | Lophuromys_nudicaudus | Lophuromys_nudicaudus | 0 | 39.5 | 2.28 | 612364.74 |
| mammals | Lophuromys_rahmi | Lophuromys_rahmi | 0 | 38 | 1.94 | 4293.32 |
| mammals | Lophuromys_sikapusi | Lophuromys_sikapusi | 0 | 74.1 | 3.12 | 3320549.7 |
| mammals | Lophuromys_woosnami | Lophuromys_woosnami | 0 | 42.2 | 1.85 | 52330.48 |
| mammals | Lorentzimys_nouhuysii | Lorentzimys_nouhuysi | 0 | 14.2 | 2.75 | 251852.19 |
| mammals | Loris_lydekkerianus | Loris_lydekkerianus | 0 | 235 | 1.37 | 493888.94 |
| mammals | Loris_tardigradus | Loris_tardigradus | 1 | 227.7 | 1.41 | 6904 |
| mammals | Loxodonta_africana | Loxodonta_africana | 1 | 5300000 | 0.92 | 3213906.06 |
| mammals | Loxodontomys_micropus | Loxodontomys_micropus | 0 | 72.7 | 4.16 | 452856.9 |
| mammals | Lundomys_molitor | Lundomys_molitor | 0 | 238.5 | 2.91 | 295487.69 |
| mammals | Lupulella_adustus | Canis_adustus | 0 | 10249.9 | 4.33 | 10714383.05 |
| mammals | Lupulella_mesomelas | Canis_mesomelas | 0 | 8500 | 3.92 | 6453409.89 |
| mammals | Lutra_capensis | Aonyx_capensis | 0 | 18999.8 | 2.59 | 11817070.92 |
| mammals | Lutra_cinerea | Aonyx_cinereus | 1 | 1747.6 | 1.47 | 4793188.51 |
| mammals | Lutra_congica | Aonyx_congicus | 0 | 23800.2 | 2.59 | 2924429.89 |
| mammals | Lutra_lutra | Lutra_lutra | 0 | 8785.1 | 2.02 | 27067759.29 |
| mammals | Lutra_perspicillata | Lutrogale_perspicillata | 1 | 9483.3 | 3.68 | 4875750.85 |
| mammals | Lutreolina_crassicaudata | Lutreolina_crassicaudata | 0 | 537.3 | 7.86 | 3049187.09 |
| mammals | Lycalopex_culpaeus | Lycalopex_culpaeus | 0 | 9832.4 | 4.44 | 2667800.13 |
| mammals | Lycalopex_fulvipes | Lycalopex_fulvipes | 1 | 2837.5 | 2.5 | 15381.48 |
| mammals | Lycalopex_grisea | Lycalopex_griseus | 0 | 8280 | 3.09 | 1618133.16 |
| mammals | Lycalopex_gymnocerca | Lycalopex_gymnocercus | 0 | 4690 | 3.09 | 2776265.54 |
| mammals | Lycalopex_vetula | Lycalopex_vetulus | 0 | 5350 | 3.27 | 2052494.18 |
| mammals | Lycaon_pictus | Lycaon_pictus | 1 | 22050.1 | 7.8 | 1272642.8 |
| mammals | Lyncodon_patagonicus | Lyncodon_patagonicus | 0 | 225 | 3.5 | 1386425.69 |

| | | | | | | |
|---|---|---|---|---|---|---|
| mammals | Lynx_canadensis | Lynx_canadensis | 0 | 9373.2 | 2.96 | 8328044.88 |
| mammals | Lynx_lynx | Lynx_lynx | 0 | 17950 | 2.28 | 20817635.89 |
| mammals | Lynx_pardinus | Lynx_pardinus | 1 | 9400 | 2.5 | 1191.69 |
| mammals | Lynx_rufus | Lynx_rufus | 0 | 8904.1 | 2.64 | 9879513.02 |
| mammals | Lyroderma_lyra | Megaderma_lyra | 0 | 45.5 | 1 | 6060994.63 |
| mammals | Macaca_arctoides | Macaca_arctoides | 1 | 5000 | 1 | 1712459.6 |
| mammals | Macaca_assamensis | Macaca_assamensis | 0 | 12000 | 1 | 1808926.19 |
| mammals | Macaca_cyclopis | Macaca_cyclopis | 0 | 6000 | 1 | 26772.6 |
| mammals | Macaca_fascicularis | Macaca_fascicularis | 1 | 3991.5 | 1 | 2431069.1 |
| mammals | Macaca_fuscata | Macaca_fuscata | 0 | 10400 | 1.22 | 157042.03 |
| mammals | Macaca_hecki | Macaca_hecki | 1 | 9000 | 1 | 23994.72 |
| mammals | Macaca_leonina | Macaca_leonina | 1 | 6500 | 1 | 1548715.3 |
| mammals | Macaca_maura | Macaca_maura | 1 | 7300 | 1 | 19910.24 |
| mammals | Macaca_mulatta | Macaca_mulatta | 0 | 3713 | 1 | 7472315.4 |
| mammals | Macaca_nemestrina | Macaca_nemestrina | 1 | 6296.5 | 1 | 1343992.2 |
| mammals | Macaca_nigra | Macaca_nigra | 1 | 6400 | 1 | 7665.76 |
| mammals | Macaca_nigrescens | Macaca_nigrescens | 1 | 9500 | 1 | 9771.56 |
| mammals | Macaca_ochreata | Macaca_ochreata | 1 | 2600 | 1 | 35625.65 |
| mammals | Macaca_pagensis | Macaca_pagensis | 1 | 6280 | 1 | 2140.85 |
| mammals | Macaca_radiata | Macaca_radiata | 1 | 3690 | 1 | 620918.14 |
| mammals | Macaca_siberu | Macaca_siberu | 1 | 4526.7 | 1 | 3863.63 |
| mammals | Macaca_silenus | Macaca_silenus | 1 | 4750 | 1 | 27905.78 |
| mammals | Macaca_sinica | Macaca_sinica | 1 | 3590 | 1 | 46294.77 |
| mammals | Macaca_sylvanus | Macaca_sylvanus | 1 | 12200 | 1.26 | 25901.57 |
| mammals | Macaca_thibetana | Macaca_thibetana | 0 | 12000 | 1 | 1327321.5 |
| mammals | Macaca_tonkeana | Macaca_tonkeana | 1 | 9000 | 1 | 81339.18 |
| mammals | Macroderma_gigas | Macroderma_gigas | 1 | 126.4 | 1 | 1156786.38 |
| mammals | Macroglossus_minimus | Macroglossus_minimus | 0 | 16.4 | 0.99 | 3590246 |
| mammals | Macroglossus_sobrinus | Macroglossus_sobrinus | 0 | 22.1 | 1 | 2435640.09 |
| mammals | Macronycteris_commersonii | Hipposideros_commersoni | 0 | 99.1 | 0.99 | 366161.23 |
| mammals | Macronycteris_gigas | Hipposideros_gigas | 0 | 166.5 | 1 | 3577138.02 |
| mammals | Macronycteris_thomensis | Hipposideros_thomensis | 0 | 67 | 1 | 856.81 |
| mammals | Macronycteris_vittata | Hipposideros_vittatus | 0 | 127.9 | 1 | 4006579.67 |
| mammals | Macrophyllum_macrophyllum | Macrophyllum_macrophyllum | 0 | 8.1 | 1 | 13066617.65 |

| | | | | | | |
|---|---|---|---|---|---|---|
| mammals | Macropus_fuliginosus | Macropus_fuliginosus | 0 | 22000 | 1 | 1607008.42 |
| mammals | Macropus_giganteus | Macropus_giganteus | 0 | 25875 | 1 | 2231237.8 |
| mammals | Macroscelides_flavicaudatus | Macroscelides_flavicaudatus | 0 | 30.5 | 1.76 | 52395.5 |
| mammals | Macroscelides_micus | Macroscelides_micus | 0 | 27 | 1.5 | 947.12 |
| mammals | Macroscelides_proboscideus | Macroscelides_proboscideus | 0 | 38.2 | 1.76 | 530451.92 |
| mammals | Macrotarsomys_bastardi | Macrotarsomys_bastardi | 0 | 29.5 | 2.5 | 152421.05 |
| mammals | Macrotarsomys_ingens | Macrotarsomys_ingens | 1 | 63.5 | 2 | 2998.65 |
| mammals | Macrotis_lagotis | Macrotis_lagotis | 1 | 1350 | 1.62 | 1079998.51 |
| mammals | Macrotus_californicus | Macrotus_californicus | 0 | 11.6 | 1 | 630012.55 |
| mammals | Macrotus_waterhousii | Macrotus_waterhousii | 0 | 12.8 | 0.99 | 817691.31 |
| mammals | Madoqua_guentheri | Madoqua_guentheri | 0 | 7500 | 0.99 | 1270987.22 |
| mammals | Madoqua_kirkii | Madoqua_kirkii | 0 | 5300.1 | 0.99 | 1336497 |
| mammals | Madoqua_saltiana | Madoqua_saltiana | 0 | 3250 | 1 | 1098767.22 |
| mammals | Madromys_blanfordi | Madromys_blanfordi | 0 | 141 | 2.5 | 990642.41 |
| mammals | Makalata_didelphoides | Makalata_didelphoides | 0 | 400 | 1.08 | 3605587.13 |
| mammals | Makalata_macrura | Makalata_macrura | 0 | 439 | 1.5 | 2457872.71 |
| mammals | Malacomys_cansdalei | Malacomys_cansdalei | 0 | 71 | 2.91 | 240684.25 |
| mammals | Malacomys_edwardsi | Malacomys_edwardsi | 0 | 65 | 2.57 | 482419.36 |
| mammals | Malacomys_longipes | Malacomys_longipes | 0 | 95.6 | 2.91 | 1218174.72 |
| mammals | Malacothrix_typica | Malacothrix_typica | 0 | 13.4 | 4.06 | 1387360.96 |
| mammals | Mallomys_aroaensis | Mallomys_aroaensis | 0 | 1664 | 1.02 | 136592.08 |
| mammals | Mallomys_istapantap | Mallomys_istapantap | 0 | 1950 | 1.07 | 75500.66 |
| mammals | Mallomys_rothschildi | Mallomys_rothschildi | 0 | 1157.5 | 1.02 | 190242.22 |
| mammals | Mammelomys_lanosus | Mammelomys_lanosus | 0 | 117.2 | 2.38 | 86029.86 |
| mammals | Mammelomys_rattoides | Mammelomys_rattoides | 0 | 209.5 | 1.64 | 125631.71 |
| mammals | Mandrillus_leucophaeus | Mandrillus_leucophaeus | 1 | 15000 | 1 | 17586.98 |
| mammals | Mandrillus_sphinx | Mandrillus_sphinx | 1 | 18249.9 | 1.03 | 333380.84 |
| mammals | Manis_crassicaudata | Manis_crassicaudata | 1 | 8457.3 | 1 | 2908388.91 |
| mammals | Manis_culionensis | Manis_culionensis | 1 | 5250 | 1 | 13297.09 |
| mammals | Manis_javanica | Manis_javanica | 1 | 5150 | 1 | 2319150.58 |
| mammals | Manis_pentadactyla | Manis_pentadactyla | 1 | 4675 | 1 | 2757986.81 |
| mammals | Marmosa_alstoni | Marmosa_alstoni | 0 | 132.3 | 11 | 213003.01 |
| mammals | Marmosa_demerarae | Marmosa_demerarae | 0 | 75.8 | 6.97 | 7826251.56 |
| mammals | Marmosa_lepida | Marmosa_lepida | 0 | 14 | 5 | 4163942.39 |

| | | | | | | |
|---|---|---|---|---|---|---|
| mammals | Marmosa_mexicana | Marmosa_mexicana | 0 | 49.5 | 8.7 | 927634.96 |
| mammals | Marmosa_murina | Marmosa_murina | 0 | 26 | 6.95 | 6450138.45 |
| mammals | Marmosa_paraguayana | Marmosa_paraguayanus | 0 | 74.4 | 6.71 | 962990.45 |
| mammals | Marmosa_robinsoni | Marmosa_robinsoni | 0 | 56.7 | 9.62 | 588222.59 |
| mammals | Marmosa_xerophila | Marmosa_xerophila | 1 | 46.2 | 7.9 | 20304 |
| mammals | Marmosops_caucae | Marmosops_neblina | 0 | 44 | 9.5 | 1675827.06 |
| mammals | Marmosops_parvidens | Marmosops_parvidens | 0 | 15 | 4.05 | 1459355.69 |
| mammals | Marmosops_paulensis | Marmosops_paulensis | 0 | 40 | 5 | 74073.18 |
| mammals | Marmosops_pinheiroi | Marmosops_pinheiroi | 0 | 26 | 4.05 | 586486.63 |
| mammals | Marmota_baibacina | Marmota_baibacina | 0 | 7500 | 5.69 | 606964.31 |
| mammals | Marmota_bobak | Marmota_bobak | 0 | 5500 | 6.08 | 790385.69 |
| mammals | Marmota_broweri | Marmota_broweri | 0 | 3405 | 4.64 | 218588.16 |
| mammals | Marmota_caligata | Marmota_caligata | 0 | 7230 | 4.64 | 2006099.66 |
| mammals | Marmota_camtschatica | Marmota_camtschatica | 0 | 3500 | 4.77 | 1023357.21 |
| mammals | Marmota_caudata | Marmota_caudata | 0 | 4350 | 4.19 | 547910.24 |
| mammals | Marmota_flaviventris | Marmota_flaviventris | 0 | 3350 | 4.55 | 1718256.72 |
| mammals | Marmota_himalayana | Marmota_himalayana | 0 | 5000 | 5.39 | 2387800.78 |
| mammals | Marmota_marmota | Marmota_marmota | 0 | 2010 | 4.1 | 213044.43 |
| mammals | Marmota_menzbieri | Marmota_menzbieri | 1 | 3250 | 3.37 | 2230.59 |
| mammals | Marmota_monax | Marmota_monax | 0 | 3801.7 | 4.3 | 7611091.83 |
| mammals | Marmota_olympus | Marmota_olympus | 0 | 6300 | 4 | 5368.56 |
| mammals | Marmota_sibirica | Marmota_sibirica | 1 | 8000 | 4.58 | 1534491.67 |
| mammals | Marmota_vancouverensis | Marmota_vancouverensis | 1 | 6300 | 3.59 | 2075.02 |
| mammals | Martes_americana | Martes_americana | 0 | 1250 | 2.66 | 7637867.17 |
| mammals | Martes_flavigula | Martes_flavigula | 0 | 1842.5 | 3.02 | 6697917.89 |
| mammals | Martes_foina | Martes_foina | 0 | 1540.8 | 3.78 | 12454427.46 |
| mammals | Martes_martes | Martes_martes | 0 | 1300 | 3.64 | 9929653.83 |
| mammals | Martes_zibellina | Martes_zibellina | 0 | 1130 | 3.13 | 10081922.24 |
| mammals | Massoutiera_mzabi | Massoutiera_mzabi | 0 | 194 | 2.03 | 652446.5 |
| mammals | Mastacomys_fuscus | Mastacomys_fuscus | 0 | 122 | 1.5 | 65318.33 |
| mammals | Mastomys_coucha | Mastomys_coucha | 0 | 31 | 7.96 | 1271248.17 |
| mammals | Mastomys_erythroleucus | Mastomys_erythroleucus | 0 | 46.5 | 9.6 | 7098448.73 |
| mammals | Mastomys_huberti | Mastomys_huberti | 0 | 55.3 | 7.96 | 578164.78 |
| mammals | Mastomys_kollmannspergeri | Mastomys_kollmannspergeri | 0 | 94.7 | 9.5 | 1619743.21 |

| | | | | | | |
|---|---|---|---|---|---|---|
| mammals | Mastomys_natalensis | Mastomys_natalensis | 0 | 36.4 | 8.91 | 16655440.98 |
| mammals | Maxomys_hellwaldii | Maxomys_hellwaldii | 0 | 203 | 3 | 144666.76 |
| mammals | Maxomys_inas | Maxomys_inas | 0 | 90 | 2.91 | 12645.86 |
| mammals | Maxomys_pagensis | Maxomys_pagensis | 1 | 196 | 3.54 | 6004.26 |
| mammals | Maxomys_panglima | Maxomys_panglima | 0 | 189 | 3.54 | 13144.03 |
| mammals | Maxomys_rajah | Maxomys_rajah | 1 | 150.3 | 3.29 | 1317194.65 |
| mammals | Maxomys_surifer | Maxomys_surifer | 0 | 150.4 | 3.54 | 2819980.27 |
| mammals | Maxomys_whiteheadi | Maxomys_whiteheadi | 1 | 56 | 3 | 1321686.46 |
| mammals | Mazama_chunyi | Mazama_chunyi | 1 | 16499.9 | 1.5 | 153629.41 |
| mammals | Mazama_nanus | Mazama_nana | 1 | 16499.9 | 1 | 447853.26 |
| mammals | Mazama_rufina | Mazama_rufina | 1 | 25999.8 | 1.17 | 85280.75 |
| mammals | Megaderma_spasma | Megaderma_spasma | 0 | 25.4 | 1.21 | 4056887.36 |
| mammals | Megadontomys_cryophilus | Megadontomys_cryophilus | 1 | 32 | 3 | 390.38 |
| mammals | Megadontomys_nelsoni | Megadontomys_nelsoni | 1 | 80.1 | 2.5 | 19798.12 |
| mammals | Megadontomys_thomasi | Megadontomys_thomasi | 1 | 111 | 2.62 | 5103.34 |
| mammals | Megaloglossus_woermanni | Megaloglossus_woermanni | 0 | 15.7 | 1 | 3120610.92 |
| mammals | Melanomys_caliginosus | Melanomys_caliginosus | 0 | 41 | 1.45 | 748135.41 |
| mammals | Meles_anakuma | Meles_anakuma | 0 | 11500 | 3.11 | 203248.43 |
| mammals | Meles_leucurus | Meles_leucurus | 0 | 6250 | 3.11 | 15959053.38 |
| mammals | Meles_meles | Meles_meles | 0 | 13000 | 3.11 | 9801554.24 |
| mammals | Mellivora_capensis | Mellivora_capensis | 0 | 8500 | 2.34 | 29722162.16 |
| mammals | Melogale_everetti | Melogale_everetti | 1 | 1500 | 2.25 | 19.94 |
| mammals | Melogale_moschata | Melogale_moschata | 0 | 805.3 | 2.12 | 2960755.42 |
| mammals | Melogale_orientalis | Melogale_orientalis | 0 | 2000 | 2.75 | 131392.64 |
| mammals | Melogale_personata | Melogale_personata | 0 | 1702.5 | 2.62 | 1971107.05 |
| mammals | Melomys_aerosus | Melomys_aerosus | 1 | 97.3 | 1.5 | 1314.22 |
| mammals | Melomys_bannisteri | Melomys_bannisteri | 1 | 52.8 | 2.35 | 566.77 |
| mammals | Melomys_burtoni | Melomys_burtoni | 0 | 68.5 | 2.5 | 492110.48 |
| mammals | Melomys_capensis | Melomys_capensis | 0 | 70 | 2.2 | 31410.62 |
| mammals | Melomys_caurinus | Melomys_caurinus | 1 | 148.1 | 2.08 | 929.16 |
| mammals | Melomys_cervinipes | Melomys_cervinipes | 0 | 78 | 2.28 | 217518.12 |
| mammals | Melomys_dollmani | Melomys_dollmani | 0 | 77.7 | 2.08 | 47118.68 |
| mammals | Melomys_frigicola | Melomys_frigicola | 0 | 71 | 2.5 | 2691.3 |
| mammals | Melomys_leucogaster | Melomys_leucogaster | 0 | 108.5 | 2.11 | 243984.08 |

| | | | | | | |
|---|---|---|---|---|---|---|
| mammals | Melomys_lutillus | Melomys_lutillus | 0 | 37.4 | 2.5 | 340703.58 |
| mammals | Melomys_matambuai | Melomys_matambuai | 1 | 145 | 2.08 | 1830.07 |
| mammals | Melomys_obiensis | Melomys_obiensis | 0 | 71 | 5.88 | 2708.4 |
| mammals | Melomys_rufescens | Melomys_rufescens | 0 | 60.1 | 2.13 | 769118.53 |
| mammals | Melomys_talaudium | Melomys_talaudium | 1 | 143.1 | 2.08 | 929.16 |
| mammals | Melonycteris_melanops | Melonycteris_melanops | 0 | 49 | 1 | 45239.91 |
| mammals | Melursus_ursinus | Melursus_ursinus | 1 | 93130 | 1.54 | 1080994.28 |
| mammals | Mephitis_macroura | Mephitis_macroura | 0 | 801.3 | 4.51 | 1783431.71 |
| mammals | Mephitis_mephitis | Mephitis_mephitis | 0 | 2085 | 5.4 | 12271610.67 |
| mammals | Meriones_crassus | Meriones_crassus | 0 | 80 | 4.11 | 7571087.96 |
| mammals | Meriones_grandis | Meriones_grandis | 0 | 205 | 5.62 | 708604.47 |
| mammals | Meriones_hurrianae | Meriones_hurrianae | 0 | 70 | 4.19 | 480817.19 |
| mammals | Meriones_libycus | Meriones_libycus | 0 | 91.3 | 4.76 | 7831992.14 |
| mammals | Meriones_meridianus | Meriones_meridianus | 0 | 41 | 5.62 | 6163493.31 |
| mammals | Meriones_persicus | Meriones_persicus | 0 | 100 | 5.08 | 2222403.48 |
| mammals | Meriones_sacramenti | Meriones_sacramenti | 1 | 200 | 5.05 | 8212.93 |
| mammals | Meriones_shawii | Meriones_shawi | 0 | 90.6 | 5.73 | 620432.3 |
| mammals | Meriones_tamariscinus | Meriones_tamariscinus | 0 | 120 | 4.08 | 2568252.74 |
| mammals | Meriones_tristrami | Meriones_tristrami | 0 | 105 | 5.7 | 901513.45 |
| mammals | Meriones_unguiculatus | Meriones_unguiculatus | 0 | 53.3 | 5 | 3276858.64 |
| mammals | Meriones_vinogradovi | Meriones_vinogradovi | 0 | 150 | 7.24 | 304025.53 |
| mammals | Mesechinus_dauuricus | Mesechinus_dauuricus | 0 | 342 | 5 | 5038641.96 |
| mammals | Mesembriomys_gouldii | Mesembriomys_gouldii | 1 | 716.5 | 1.92 | 237211.18 |
| mammals | Mesembriomys_macrurus | Mesembriomys_macrurus | 0 | 267 | 2 | 34779.61 |
| mammals | Mesocapromys_melanurus | Mysateles_melanurus | 1 | 1218 | 1.75 | 31192.62 |
| mammals | Mesocricetus_auratus | Mesocricetus_auratus | 1 | 105 | 8.74 | 4704.21 |
| mammals | Mesocricetus_brandti | Mesocricetus_brandti | 0 | 198 | 8.8 | 732898.24 |
| mammals | Mesocricetus_newtoni | Mesocricetus_newtoni | 0 | 97.5 | 8.69 | 38236.1 |
| mammals | Mesocricetus_raddei | Mesocricetus_raddei | 0 | 260 | 8.89 | 141380.03 |
| mammals | Mesomys_hispidus | Mesomys_hispidus | 0 | 175 | 0.97 | 4948976.38 |
| mammals | Mesomys_occultus | Mesomys_occultus | 0 | 146.7 | 1 | 53277.64 |
| mammals | Mesomys_stimulax | Mesomys_stimulax | 0 | 139 | 0.97 | 437362.91 |
| mammals | Mesophylla_macconnelli | Mesophylla_macconnelli | 0 | 6.9 | 0.99 | 8817651.77 |
| mammals | Metachirus_nudicaudatus | Metachirus_nudicaudatus | 0 | 375 | 4.43 | 10341442.4 |

| | | | | | | |
|---|---|---|---|---|---|---|
| mammals | Micaelamys_namaquensis | Aethomys_namaquensis | 0 | 50.6 | 3.62 | 3338532.92 |
| mammals | Mico_argentatus | Mico_argentatus | 0 | 440 | 1.85 | 137206.26 |
| mammals | Mico_emiliae | Mico_emiliae | 0 | 335 | 1.87 | 151986.9 |
| mammals | Mico_humeralifer | Mico_humeralifer | 0 | 350 | 2.24 | 63580.26 |
| mammals | Mico_intermedius | Mico_intermedius | 0 | 425 | 2 | 62624.11 |
| mammals | Mico_leucippe | Mico_leucippe | 0 | 440 | 1.85 | 70830.61 |
| mammals | Mico_marcai | Mico_marcai | 1 | 295 | 1.87 | 7000.76 |
| mammals | Mico_melanurus | Mico_melanurus | 0 | 302.3 | 1.87 | 769939.42 |
| mammals | Mico_nigriceps | Mico_nigriceps | 0 | 370 | 2 | 31646.7 |
| mammals | Microcavia_australis | Microcavia_australis | 0 | 286.1 | 2.48 | 1741369.31 |
| mammals | Microcavia_niata | Microcavia_niata | 0 | 255.2 | 3.16 | 155322.59 |
| mammals | Microcavia_shiptoni | Microcavia_shiptoni | 0 | 185 | 3.16 | 39515.23 |
| mammals | Microcebus_griseorufus | Microcebus_griseorufus | 0 | 62.5 | 2.41 | 35820.44 |
| mammals | Microcebus_murinus | Microcebus_murinus | 0 | 60 | 2.2 | 90390.12 |
| mammals | Microcebus_myoxinus | Microcebus_myoxinus | 1 | 47.5 | 2.1 | 31213.8 |
| mammals | Microcebus_rufus | Microcebus_rufus | 1 | 42.5 | 2.46 | 2607.4 |
| mammals | Microdipodops_megacephalus | Microdipodops_megacephalus | 0 | 10.5 | 3.92 | 226040.9 |
| mammals | Microdipodops_pallidus | Microdipodops_pallidus | 0 | 12.5 | 3.89 | 31230.11 |
| mammals | Microgale_cowani | Microgale_cowani | 0 | 13.8 | 2.5 | 93169.76 |
| mammals | Microgale_fotsifotsy | Microgale_fotsifotsy | 0 | 7.7 | 3.5 | 111846.77 |
| mammals | Microgale_gymnorhyncha | Microgale_gymnorhyncha | 0 | 18.2 | 3 | 77232.22 |
| mammals | Microgale_mergulus | Limnogale_mergulus | 1 | 78.6 | 2.8 | 56589.48 |
| mammals | Microgale_monticola | Microgale_monticola | 1 | 13.9 | 1.5 | 759.56 |
| mammals | Microgale_nasoloi | Microgale_nasoloi | 1 | 10 | 3 | 1193.56 |
| mammals | Microgale_parvula | Microgale_parvula | 0 | 3.4 | 4 | 92421.46 |
| mammals | Microgale_principula | Microgale_principula | 0 | 10.2 | 3 | 89104.73 |
| mammals | Microgale_soricoides | Microgale_soricoides | 0 | 18.7 | 2 | 102314.28 |
| mammals | Microgale_taiva | Microgale_taiva | 0 | 12.4 | 2 | 43037.75 |
| mammals | Microgale_thomasi | Microgale_thomasi | 0 | 22.9 | 2 | 22816.33 |
| mammals | Micromys_minutus | Micromys_minutus | 0 | 6 | 5.04 | 18265246.85 |
| mammals | Micronomus_norfolkensis | Mormopterus_norfolkensis | 0 | 8 | 1.05 | 74501.37 |
| mammals | Micronycteris_hirsuta | Micronycteris_hirsuta | 0 | 12 | 0.99 | 6515654.57 |
| mammals | Micronycteris_megalotis | Micronycteris_megalotis | 0 | 6.1 | 0.98 | 12095131.87 |
| mammals | Micronycteris_microtis | Micronycteris_microtis | 0 | 5.6 | 0.99 | 4395481 |

| | | | | | | |
|---|---|---|---|---|---|---|
| mammals | Micronycteris_minuta | Micronycteris_minuta | 0 | 6.5 | 0.99 | 12093270.59 |
| mammals | Micronycteris_sanborni | Micronycteris_sanborni | 0 | 6.3 | 1 | 137406.74 |
| mammals | Microperoryctes_longicauda | Microperoryctes_longicauda | 0 | 538 | 2.02 | 232311.31 |
| mammals | Microperoryctes_papuensis | Microperoryctes_papuensis | 0 | 158 | 1.01 | 19193.6 |
| mammals | Micropotamogale_lamottei | Micropotamogale_lamottei | 1 | 69.6 | 3.9 | 2100.01 |
| mammals | Micropotamogale_ruwenzorii | Micropotamogale_ruwenzorii | 0 | 98.5 | 1.86 | 52418.56 |
| mammals | Microryzomys_minutus | Microryzomys_minutus | 0 | 13.5 | 5.5 | 638218.25 |
| mammals | Microsciurus_alfari | Microsciurus_alfari | 0 | 87.5 | 2 | 117452.51 |
| mammals | Microsciurus_flaviventer | Microsciurus_flaviventer | 0 | 92 | 2 | 2575239.41 |
| mammals | Microtus_abbreviatus | Microtus_abbreviatus | 0 | 62 | 2.89 | 1432006.91 |
| mammals | Microtus_afghanus | Blanfordimys_afghanus | 0 | 30.8 | 3.4 | 409814.27 |
| mammals | Microtus_agrestis | Microtus_agrestis | 0 | 42.5 | 4.54 | 10818739.68 |
| mammals | Microtus_arvalis | Microtus_arvalis | 0 | 28 | 5.04 | 6887242.44 |
| mammals | Microtus_brachycercus | Microtus_brachycercus | 0 | 19 | 3.4 | 70234.69 |
| mammals | Microtus_bucharensis | Blanfordimys_bucharensis | 0 | 45 | 4.5 | 116187.81 |
| mammals | Microtus_cabrerae | Microtus_cabrerae | 0 | 52.5 | 4 | 119784.07 |
| mammals | Microtus_canicaudus | Microtus_canicaudus | 0 | 28.5 | 4.67 | 8888.22 |
| mammals | Microtus_chrotorrhinus | Microtus_chrotorrhinus | 0 | 39 | 3.58 | 1384481.21 |
| mammals | Microtus_felteni | Microtus_felteni | 0 | 21 | 2 | 35699.48 |
| mammals | Microtus_ilaeus | Microtus_ilaeus | 0 | 25.6 | 4.99 | 244614.04 |
| mammals | Microtus_irani | Microtus_irani | 1 | 37.1 | 5.24 | 1637.83 |
| mammals | Microtus_longicaudus | Microtus_longicaudus | 0 | 46.7 | 4.85 | 4021651.48 |
| mammals | Microtus_majori | Microtus_majori | 0 | 24.5 | 3.6 | 142325.88 |
| mammals | Microtus_mexicanus | Microtus_mexicanus | 0 | 35 | 2.43 | 854502.22 |
| mammals | Microtus_montanus | Microtus_montanus | 0 | 36.3 | 5.36 | 1453057.65 |
| mammals | Microtus_oaxacensis | Microtus_oaxacensis | 1 | 39.2 | 1.2 | 510.44 |
| mammals | Microtus_ochrogaster | Microtus_ochrogaster | 0 | 38 | 3.84 | 3243052.72 |
| mammals | Microtus_oregoni | Microtus_oregoni | 0 | 20.4 | 3.31 | 206842.17 |
| mammals | Microtus_pennsylvanicus | Microtus_pennsylvanicus | 0 | 36.8 | 5.11 | 11962526.6 |
| mammals | Microtus_quasiater | Microtus_quasiater | 0 | 40 | 1.4 | 19611.13 |
| mammals | Microtus_richardsoni | Microtus_richardsoni | 0 | 85 | 5.9 | 859823.37 |
| mammals | Microtus_schelkovnikovi | Microtus_schelkovnikovi | 0 | 26 | 2.8 | 21598.46 |
| mammals | Microtus_townsendii | Microtus_townsendii | 0 | 64.8 | 5.12 | 213940.74 |
| mammals | Microtus_transcaspicus | Microtus_transcaspicus | 0 | 38.8 | 2.1 | 11320.19 |

| | | | | | | |
|---|---|---|---|---|---|---|
| mammals | Microtus_xanthognathus | Microtus_xanthognathus | 0 | 125.8 | 8.14 | 2263668.69 |
| mammals | Microtus_yuldaschi | Neodon_juldaschi | 0 | 44.6 | 3.52 | 311957.47 |
| mammals | Mictomys_borealis | Synaptomys_borealis | 0 | 21.3 | 4.3 | 6774122.51 |
| mammals | Mimetillus_moloneyi | Mimetillus_moloneyi | 0 | 8.8 | 0.98 | 3621345.78 |
| mammals | Mimon_bennettii | Mimon_bennettii | 0 | 25.2 | 1 | 5698708.12 |
| mammals | Mimon_cozumelae | Mimon_cozumelae | 0 | 12.4 | 0.98 | 803519.04 |
| mammals | Miniopterus_australis | Miniopterus_australis | 0 | 6.2 | 1 | 2201727.95 |
| mammals | Miniopterus_fraterculus | Miniopterus_fraterculus | 0 | 7.3 | 1 | 1668579.51 |
| mammals | Miniopterus_fuscus | Miniopterus_fuscus | 1 | 10.7 | 1 | 2987.77 |
| mammals | Miniopterus_gleni | Miniopterus_gleni | 0 | 14 | 1 | 417528.44 |
| mammals | Miniopterus_inflatus | Miniopterus_inflatus | 0 | 14.2 | 1 | 2350003 |
| mammals | Miniopterus_macrocneme | Miniopterus_macrocneme | 0 | 9.2 | 1 | 470913.2 |
| mammals | Miniopterus_magnater | Miniopterus_magnater | 0 | 14.1 | 1 | 3161072.07 |
| mammals | Miniopterus_majori | Miniopterus_majori | 0 | 8 | 1 | 249561.26 |
| mammals | Miniopterus_manavi | Miniopterus_manavi | 0 | 5 | 1 | 511543.16 |
| mammals | Miniopterus_medius | Miniopterus_medius | 0 | 9 | 1 | 1256827.57 |
| mammals | Miniopterus_natalensis | Miniopterus_natalensis | 0 | 12.5 | 1 | 3787497.56 |
| mammals | Miniopterus_paululus | Miniopterus_paululus | 0 | 6 | 1 | 278632.69 |
| mammals | Miniopterus_pusillus | Miniopterus_pusillus | 0 | 8.6 | 1 | 1106938.27 |
| mammals | Miniopterus_robustior | Miniopterus_robustior | 1 | 8.1 | 1 | 1081.82 |
| mammals | Miniopterus_schreibersii | Miniopterus_schreibersii | 1 | 11.1 | 1 | 2898979.4 |
| mammals | Miniopterus_sororculus | Miniopterus_sororculus | 0 | 8 | 1 | 91220.22 |
| mammals | Miniopterus_tristis | Miniopterus_tristis | 0 | 15.8 | 1 | 732372.81 |
| mammals | Miopithecus_ogouensis | Miopithecus_ogouensis | 0 | 1278.3 | 1.1 | 486827.11 |
| mammals | Miopithecus_talapoin | Miopithecus_talapoin | 1 | 1499.2 | 1 | 399163.1 |
| mammals | Mirza_coquereli | Mirza_coquereli | 1 | 308 | 1.71 | 16761.47 |
| mammals | Mirza_zaza | Mirza_zaza | 1 | 290 | 2.3 | 9851.88 |
| mammals | Molossops_temminckii | Molossops_temminckii | 0 | 5.9 | 0.98 | 11605510.25 |
| mammals | Molossus_aztecus | Molossus_aztecus | 0 | 14.3 | 0.99 | 20680.33 |
| mammals | Molossus_coibensis | Molossus_coibensis | 0 | 13 | 1 | 6800728.09 |
| mammals | Molossus_currentium | Molossus_currentium | 0 | 17.8 | 0.99 | 1057644.54 |
| mammals | Molossus_molossus | Molossus_molossus | 0 | 13.6 | 0.99 | 15767723.28 |
| mammals | Molossus_pretiosus | Molossus_pretiosus | 0 | 27 | 0.98 | 2034434.03 |
| mammals | Molossus_rufus | Molossus_rufus | 0 | 31.7 | 1.05 | 14800275.01 |

| | | | | | | |
|---|---|---|---|---|---|---|
| mammals | Molossus_sinaloae | Molossus_sinaloae | 0 | 22.6 | 1.02 | 2024567.25 |
| mammals | Monodelphis_adusta | Monodelphis_adusta | 0 | 35 | 4 | 1083732.5 |
| mammals | Monodelphis_brevicaudata | Monodelphis_brevicaudata | 0 | 69.6 | 7.18 | 1828323.55 |
| mammals | Monodelphis_dimidiata | Monodelphis_dimidiata | 0 | 58 | 13 | 1931861.77 |
| mammals | Monodelphis_domestica | Monodelphis_domestica | 0 | 71.4 | 8.14 | 4111650.31 |
| mammals | Monophyllus_plethodon | Monophyllus_plethodon | 0 | 14.8 | 0.99 | 5400.19 |
| mammals | Monophyllus_redmani | Monophyllus_redmani | 0 | 8.7 | 0.99 | 202239.21 |
| mammals | Montemys_delectorum | Praomys_delectorum | 0 | 36 | 3.69 | 56002.78 |
| mammals | Monticolomys_koopmani | Monticolomys_koopmani | 0 | 25 | 3 | 20443.33 |
| mammals | Mops_aloysiisabaudiae | Chaerephon_aloysiisabaudiae | 0 | 25.7 | 1.05 | 1117207.37 |
| mammals | Mops_ansorgei | Chaerephon_ansorgei | 0 | 16.1 | 1 | 5633242.36 |
| mammals | Mops_bemmeleni | Chaerephon_bemmeleni | 0 | 19 | 1.05 | 671185.43 |
| mammals | Mops_bivittatus | Chaerephon_bivittatus | 0 | 15 | 1 | 1892463.64 |
| mammals | Mops_brachypterus | Mops_brachypterus | 0 | 16 | 0.99 | 1924932.17 |
| mammals | Mops_bregullae | Chaerephon_bregullae | 1 | 19.9 | 1.05 | 20613.22 |
| mammals | Mops_chapini | Chaerephon_chapini | 0 | 10 | 1.05 | 4301386.96 |
| mammals | Mops_condylurus | Mops_condylurus | 0 | 28.4 | 1 | 9108820.05 |
| mammals | Mops_congicus | Mops_congicus | 0 | 43 | 0.99 | 528771.16 |
| mammals | Mops_demonstrator | Mops_demonstrator | 0 | 23.5 | 1 | 1886714.3 |
| mammals | Mops_jobensis | Chaerephon_jobensis | 0 | 16.9 | 1.05 | 2340174.94 |
| mammals | Mops_jobimena | Chaerephon_jobimena | 0 | 14.3 | 1.05 | 186331.77 |
| mammals | Mops_johorensis | Chaerephon_johorensis | 1 | 20 | 1 | 91529.84 |
| mammals | Mops_leucostigma | Mops_leucostigma | 0 | 17.5 | 1 | 214048.94 |
| mammals | Mops_major | Chaerephon_major | 0 | 14.8 | 1 | 1759227.75 |
| mammals | Mops_midas | Mops_midas | 0 | 45.2 | 1.11 | 415330.64 |
| mammals | Mops_mops | Mops_mops | 0 | 31 | 1.05 | 46635.74 |
| mammals | Mops_nanulus | Mops_nanulus | 0 | 9.2 | 1 | 2775168.77 |
| mammals | Mops_nigeriae | Chaerephon_nigeriae | 0 | 19.8 | 1 | 4801793.71 |
| mammals | Mops_niveiventer | Mops_niveiventer | 0 | 21.7 | 1 | 1867655.71 |
| mammals | Mops_petersoni | Mops_petersoni | 0 | 12.4 | 1.05 | 41534.64 |
| mammals | Mops_plicatus | Chaerephon_plicatus | 0 | 17.1 | 0.99 | 2462336.04 |
| mammals | Mops_pumilus | Chaerephon_pumilus | 0 | 11.3 | 1 | 6943070.97 |
| mammals | Mops_solomonis | Chaerephon_solomonis | 1 | 12.3 | 1.05 | 7070.41 |
| mammals | Mops_spurrelli | Mops_spurrelli | 0 | 8.9 | 1.05 | 1162141.9 |

| | | | | | | |
|---|---|---|---|---|---|---|
| mammals | Mops_thersites | Mops_thersites | 0 | 22 | 1 | 2248550.68 |
| mammals | Mops_tomensis | Chaerephon_tomensis | 1 | 7.1 | 1.05 | 656.64 |
| mammals | Mormoops_blainvillei | Mormoops_blainvillei | 0 | 8.6 | 1 | 200604.53 |
| mammals | Mormoops_megalophylla | Mormoops_megalophylla | 0 | 15.8 | 1 | 3738685.97 |
| mammals | Mormopterus_acetabulosus | Mormopterus_acetabulosus | 1 | 14.1 | 1.05 | 1872.58 |
| mammals | Mormopterus_jugularis | Mormopterus_jugularis | 0 | 11.5 | 1.05 | 235404.57 |
| mammals | Mormopterus_kalinowskii | Mormopterus_kalinowskii | 0 | 17.6 | 1.05 | 390233.3 |
| mammals | Mormopterus_minutus | Mormopterus_minutus | 1 | 6.7 | 1.05 | 45074.17 |
| mammals | Mormopterus_phrudus | Mormopterus_phrudus | 1 | 9.3 | 1.05 | 1298.38 |
| mammals | Moschiola_indica | Moschiola_indica | 0 | 3000 | 1.5 | 2015179.41 |
| mammals | Moschiola_meminna | Moschiola_meminna | 0 | 2445 | 1.5 | 53308.92 |
| mammals | Moschus_anhuiensis | Moschus_anhuiensis | 1 | 8400 | 1.6 | 12999.74 |
| mammals | Moschus_berezovskii | Moschus_berezovskii | 1 | 9000 | 1.33 | 1605153.64 |
| mammals | Moschus_chrysogaster | Moschus_chrysogaster | 1 | 14500 | 1.15 | 1421239.73 |
| mammals | Moschus_cupreus | Moschus_cupreus | 1 | 14500 | 1.34 | 127719.94 |
| mammals | Moschus_leucogaster | Moschus_leucogaster | 1 | 15500 | 1.34 | 225843.57 |
| mammals | Moschus_moschiferus | Moschus_moschiferus | 1 | 13000 | 1.68 | 7225246.06 |
| mammals | Mosia_nigrescens | Mosia_nigrescens | 0 | 3.3 | 0.98 | 1090668.43 |
| mammals | Mungos_mungo | Mungos_mungo | 0 | 1925 | 2.68 | 10331332 |
| mammals | Mungotictis_decemlineata | Mungotictis_decemlineata | 1 | 650 | 0.99 | 13303.64 |
| mammals | Muntiacus_atherodes | Muntiacus_atherodes | 0 | 15600 | 1 | 732628.36 |
| mammals | Muntiacus_crinifrons | Muntiacus_crinifrons | 1 | 36141 | 1 | 113889.54 |
| mammals | Muntiacus_muntjak | Muntiacus_muntjak | 0 | 15925.8 | 1 | 1494965.82 |
| mammals | Muntiacus_reevesi | Muntiacus_reevesi | 0 | 13500 | 1 | 2322121.55 |
| mammals | Muntiacus_vaginalis | Muntiacus_vaginalis | 0 | 24000 | 1 | 5872023.27 |
| mammals | Muntiacus_vuquangensis | Muntiacus_vuquangensis | 1 | 34000 | 1 | 110001.6 |
| mammals | Murexia_habbema | Murexia_habbema | 0 | 34.8 | 3.65 | 143836.69 |
| mammals | Murexia_longicaudata | Murexia_longicaudata | 0 | 335.8 | 3.8 | 515141.49 |
| mammals | Murexia_melanura | Murexia_melanurus | 0 | 47.8 | 3.62 | 330753.34 |
| mammals | Murexia_naso | Murexia_naso | 0 | 48.4 | 3.68 | 181140.58 |
| mammals | Murexia_rothschildi | Murexia_rothschildi | 0 | 43.8 | 3.69 | 14840.56 |
| mammals | Murina_cyclotis | Murina_cyclotis | 0 | 9.3 | 1.99 | 1100512.77 |
| mammals | Murina_hilgendorfi | Murina_hilgendorfi | 0 | 8.1 | 1.5 | 3719628.22 |
| mammals | Murina_suilla | Murina_suilla | 0 | 3.6 | 1 | 1442016.7 |

| | | | | | | |
|---|---|---|---|---|---|---|
| mammals | Murina_ussuriensis | Murina_ussuriensis | 0 | 5.3 | 1.5 | 773194.69 |
| mammals | Mus_booduga | Mus_booduga | 0 | 11.6 | 6.15 | 3186564.99 |
| mammals | Mus_bufo | Mus_bufo | 0 | 10.4 | 3.5 | 69837.54 |
| mammals | Mus_caroli | Mus_caroli | 0 | 11.3 | 5.8 | 1841648.17 |
| mammals | Mus_cervicolor | Mus_cervicolor | 0 | 17.7 | 4.33 | 251716.07 |
| mammals | Mus_cypriacus | Mus_cypriacus | 0 | 50.6 | 6 | 9272.65 |
| mammals | Mus_indutus | Mus_indutus | 0 | 6.2 | 4.5 | 1481732.92 |
| mammals | Mus_macedonicus | Mus_macedonicus | 0 | 15 | 6.07 | 1386803.65 |
| mammals | Mus_mahomet | Mus_mahomet | 0 | 10 | 8.6 | 418704.62 |
| mammals | Mus_mayori | Mus_mayori | 1 | 29.4 | 2.96 | 1122.55 |
| mammals | Mus_minutoides | Mus_minutoides | 0 | 6.8 | 4.18 | 4215726.74 |
| mammals | Mus_musculoides | Mus_musculoides | 0 | 5.6 | 3.84 | 5014869.29 |
| mammals | Mus_musculus | Mus_musculus | 0 | 16.1 | 5.8 | 84352137.99 |
| mammals | Mus_pahari | Mus_pahari | 0 | 25.7 | 6 | 1683214.5 |
| mammals | Mus_platythrix | Mus_platythrix | 0 | 25.2 | 6.62 | 511752.21 |
| mammals | Mus_setulosus | Mus_setulosus | 0 | 11.1 | 3.92 | 1024297.29 |
| mammals | Mus_shortridgei | Mus_shortridgei | 0 | 34.6 | 6.32 | 268868.68 |
| mammals | Mus_sorella | Mus_sorella | 0 | 13.7 | 5.2 | 52516.1 |
| mammals | Mus_spicilegus | Mus_spicilegus | 0 | 14 | 6.38 | 828198.67 |
| mammals | Mus_spretus | Mus_spretus | 0 | 12.8 | 5.48 | 963695.11 |
| mammals | Mus_tenellus | Mus_tenellus | 0 | 11 | 1.94 | 491701.51 |
| mammals | Mus_terricolor | Mus_terricolor | 0 | 9 | 8 | 2890511.68 |
| mammals | Mus_triton | Mus_triton | 0 | 12.1 | 4.8 | 1251170.51 |
| mammals | Mus_vulcani | Mus_vulcani | 0 | 20.2 | 3.66 | 184.94 |
| mammals | Muscardinus_avellanarius | Muscardinus_avellanarius | 0 | 27.5 | 4.3 | 3613069.83 |
| mammals | Mustela_altaica | Mustela_altaica | 0 | 171 | 5.61 | 5661382.61 |
| mammals | Mustela_erminea | Mustela_erminea | 0 | 119.4 | 6.75 | 41482667.38 |
| mammals | Mustela_eversmanii | Mustela_eversmanii | 0 | 1350 | 7.82 | 13299450.45 |
| mammals | Mustela_itatsi | Mustela_itatsi | 0 | 565 | 6.48 | 361776.18 |
| mammals | Mustela_lutreola | Mustela_lutreola | 1 | 440 | 4.5 | 7371841.89 |
| mammals | Mustela_lutreolina | Mustela_lutreolina | 0 | 706.6 | 6.48 | 84498.73 |
| mammals | Mustela_nigripes | Mustela_nigripes | 1 | 850 | 3.38 | 2515859.23 |
| mammals | Mustela_nivalis | Mustela_nivalis | 0 | 137.5 | 5.08 | 43300266.99 |
| mammals | Mustela_nudipes | Mustela_nudipes | 0 | 500 | 4 | 1346098.67 |

| | | | | | | |
|---|---|---|---|---|---|---|
| mammals | Mustela_putorius | Mustela_putorius | 0 | 915.4 | 8.08 | 6011048.91 |
| mammals | Mustela_sibirica | Mustela_sibirica | 0 | 405 | 6.48 | 13304653.93 |
| mammals | Mydaus_javanensis | Mydaus_javanensis | 0 | 2500 | 2.5 | 105559.28 |
| mammals | Mydaus_marchei | Mydaus_marchei | 0 | 1414.2 | 2.5 | 12447.24 |
| mammals | Mylomys_dybowskii | Mylomys_dybowskii | 0 | 114.4 | 4.04 | 1279787.08 |
| mammals | Myocastor_coypus | Myocastor_coypus | 0 | 6937.5 | 5.34 | 3902051.5 |
| mammals | Myoictis_melas | Myoictis_melas | 0 | 230 | 6 | 398695.85 |
| mammals | Myoictis_wallacii | Myoictis_wallacei | 0 | 230 | 6 | 56148.8 |
| mammals | Myomimus_roachi | Myomimus_roachi | 1 | 45.5 | 8.2 | 21889.24 |
| mammals | Myonycteris_angolensis | Lissonycteris_angolensis | 0 | 74.6 | 0.99 | 9083671.62 |
| mammals | Myonycteris_leptodon | Myonycteris_leptodon | 0 | 42 | 2 | 835056.63 |
| mammals | Myonycteris_torquata | Myonycteris_torquata | 0 | 42.2 | 1 | 3651482.6 |
| mammals | Myoprocta_acouchy | Myoprocta_acouchy | 0 | 600 | 1.97 | 993244.8 |
| mammals | Myoprocta_pratti | Myoprocta_pratti | 0 | 908 | 1.82 | 2337165.24 |
| mammals | Myopterus_whitleyi | Myopterus_whitleyi | 0 | 11.5 | 1.05 | 1914285.18 |
| mammals | Myopus_schisticolor | Myopus_schisticolor | 0 | 25 | 5.1 | 10209782.76 |
| mammals | Myosciurus_pumilio | Myosciurus_pumilio | 0 | 165 | 1.99 | 346127.67 |
| mammals | Myosorex_babaulti | Myosorex_babaulti | 0 | 17 | 2.99 | 24423.89 |
| mammals | Myosorex_cafer | Myosorex_cafer | 1 | 13.4 | 2.99 | 65136.02 |
| mammals | Myosorex_geata | Myosorex_geata | 1 | 9.4 | 4 | 463.33 |
| mammals | Myosorex_kihaulei | Myosorex_kihaulei | 1 | 9.5 | 2 | 1615.63 |
| mammals | Myosorex_varius | Myosorex_varius | 0 | 13.2 | 2.92 | 424218.63 |
| mammals | Myotis_adversus | Myotis_adversus | 0 | 10.6 | 1 | 55894.18 |
| mammals | Myotis_albescens | Myotis_albescens | 0 | 5.3 | 0.98 | 14595013.01 |
| mammals | Myotis_atacamensis | Myotis_atacamensis | 1 | 7.1 | 0.98 | 253619.58 |
| mammals | Myotis_ater | Myotis_ater | 0 | 6 | 1 | 127895.39 |
| mammals | Myotis_auriculus | Myotis_auriculus | 0 | 5 | 1 | 617481.09 |
| mammals | Myotis_austroriparius | Myotis_austroriparius | 0 | 6.8 | 1.91 | 832901.78 |
| mammals | Myotis_bechsteinii | Myotis_bechsteinii | 0 | 8.5 | 1 | 2676244.15 |
| mammals | Myotis_blythii | Myotis_blythii | 0 | 22.5 | 0.99 | 6194956.01 |
| mammals | Myotis_bocagii | Myotis_bocagii | 0 | 7.8 | 0.99 | 1819217.1 |
| mammals | Myotis_bombinus | Myotis_bombinus | 0 | 6.9 | 1 | 1653772.78 |
| mammals | Myotis_brandtii | Myotis_brandtii | 0 | 5.3 | 1 | 5471819.62 |
| mammals | Myotis_californicus | Myotis_californicus | 0 | 4.6 | 0.99 | 3985147.13 |

| | | | | | | |
|---|---|---|---|---|---|---|
| mammals | Myotis_capaccinii | Myotis_capaccinii | 1 | 7.4 | 1 | 1229394.76 |
| mammals | Myotis_chiloensis | Myotis_chiloensis | 0 | 6.8 | 0.98 | 502327.68 |
| mammals | Myotis_chinensis | Myotis_chinensis | 0 | 42 | 1.25 | 2369550.79 |
| mammals | Myotis_ciliolabrum | Myotis_ciliolabrum | 0 | 4.9 | 1 | 4652123.12 |
| mammals | Myotis_dasycneme | Myotis_dasycneme | 0 | 14.1 | 1 | 5448983.29 |
| mammals | Myotis_daubentonii | Myotis_daubentonii | 0 | 7.4 | 1.21 | 7012608.32 |
| mammals | Myotis_davidii | Myotis_davidii | 0 | 5.3 | 1 | 6721248.7 |
| mammals | Myotis_dominicensis | Myotis_dominicensis | 1 | 3.2 | 1 | 2234.99 |
| mammals | Myotis_elegans | Myotis_elegans | 0 | 4.1 | 0.98 | 587280.01 |
| mammals | Myotis_emarginatus | Myotis_emarginatus | 0 | 7.7 | 0.99 | 4646898.86 |
| mammals | Myotis_evotis | Myotis_evotis | 0 | 6.7 | 0.99 | 3163297 |
| mammals | Myotis_fimbriatus | Myotis_fimbriatus | 0 | 7.6 | 1 | 1861445.65 |
| mammals | Myotis_fortidens | Myotis_fortidens | 0 | 4.4 | 0.99 | 474091.81 |
| mammals | Myotis_frater | Myotis_frater | 0 | 7.6 | 1 | 1270255.42 |
| mammals | Myotis_grisescens | Myotis_grisescens | 1 | 10.5 | 0.99 | 853015.93 |
| mammals | Myotis_hasseltii | Myotis_hasseltii | 0 | 8.8 | 0.99 | 1330531.46 |
| mammals | Myotis_horsfieldii | Myotis_horsfieldii | 0 | 6.3 | 1.5 | 3208239.96 |
| mammals | Myotis_ikonnikovi | Myotis_ikonnikovi | 0 | 3.7 | 1 | 5170695.33 |
| mammals | Myotis_keaysi | Myotis_keaysi | 0 | 5.3 | 0.98 | 2736294.01 |
| mammals | Myotis_keenii | Myotis_keenii | 0 | 6.3 | 1 | 109436.68 |
| mammals | Myotis_leibii | Myotis_leibii | 1 | 5.1 | 1 | 1423165.26 |
| mammals | Myotis_lucifugus | Myotis_lucifugus | 1 | 7.6 | 1 | 12626918.36 |
| mammals | Myotis_macrodactylus | Myotis_macrodactylus | 0 | 7.2 | 1 | 682552.49 |
| mammals | Myotis_macropus | Myotis_macropus | 0 | 10 | 1 | 1326845.82 |
| mammals | Myotis_moluccarum | Myotis_moluccarum | 0 | 9.2 | 1 | 825396.21 |
| mammals | Myotis_muricola | Myotis_muricola | 0 | 4.1 | 1 | 4387144.01 |
| mammals | Myotis_myotis | Myotis_myotis | 0 | 34 | 1.38 | 3874644.45 |
| mammals | Myotis_mystacinus | Myotis_mystacinus | 0 | 6 | 1 | 4973961.41 |
| mammals | Myotis_nattereri | Myotis_nattereri | 0 | 7.1 | 1 | 4383587.36 |
| mammals | Myotis_nigricans | Myotis_nigricans | 0 | 4.2 | 1 | 14375820.5 |
| mammals | Myotis_nipalensis | Myotis_nipalensis | 0 | 6.3 | 1 | 4946026.26 |
| mammals | Myotis_occultus | Myotis_occultus | 0 | 8 | 1 | 996794.93 |
| mammals | Myotis_oxyotus | Myotis_oxyotus | 0 | 5.5 | 0.98 | 2601562.79 |
| mammals | Myotis_pilosus | Myotis_pilosus | 1 | 15.6 | 1 | 1162073.81 |

| | | | | | | |
|---|---|---|---|---|---|---|
| mammals | Myotis_pruinosus | Myotis_pruinosus | 1 | 2.7 | 1 | 67885.32 |
| mammals | Myotis_riparius | Myotis_riparius | 0 | 5.6 | 0.98 | 14110136.66 |
| mammals | Myotis_scotti | Myotis_scotti | 1 | 8.7 | 1 | 185917.73 |
| mammals | Myotis_septentrionalis | Myotis_septentrionalis | 0 | 7.5 | 1 | 4946390.38 |
| mammals | Myotis_sicarius | Myotis_sicarius | 1 | 13.1 | 1 | 66155.33 |
| mammals | Myotis_siligorensis | Myotis_siligorensis | 0 | 2.7 | 0.98 | 1309758.19 |
| mammals | Myotis_sodalis | Myotis_sodalis | 0 | 6.9 | 0.99 | 1564971.58 |
| mammals | Myotis_stalkeri | Myotis_stalkeri | 0 | 17 | 1 | 4156.53 |
| mammals | Myotis_thysanodes | Myotis_thysanodes | 0 | 8.1 | 1 | 3472250.62 |
| mammals | Myotis_tricolor | Myotis_tricolor | 0 | 13.9 | 0.99 | 4268392.07 |
| mammals | Myotis_velifer | Myotis_velifer | 0 | 10.1 | 0.99 | 1956978.83 |
| mammals | Myotis_vivesi | Myotis_vivesi | 1 | 25 | 1 | 6940.56 |
| mammals | Myotis_volans | Myotis_volans | 0 | 8.5 | 0.99 | 4711968.98 |
| mammals | Myotis_yumanensis | Myotis_yumanensis | 0 | 5.4 | 1 | 4063522.23 |
| mammals | Myrmecobius_fasciatus | Myrmecobius_fasciatus | 1 | 471.5 | 3.84 | 2502.5 |
| mammals | Myrmecophaga_tridactyla | Myrmecophaga_tridactyla | 1 | 22333.2 | 1 | 12924626.05 |
| mammals | Mysateles_prehensilis | Mysateles_prehensilis | 0 | 1800 | 2 | 106633.6 |
| mammals | Mystacina_tuberculata | Mystacina_tuberculata | 1 | 12.6 | 1 | 13655.02 |
| mammals | Mystromys_albicaudatus | Mystromys_albicaudatus | 1 | 86.9 | 3 | 501156.54 |
| mammals | Myzopoda_aurita | Myzopoda_aurita | 0 | 9.3 | 1 | 107284.17 |
| mammals | Myzopoda_schliemanni | Myzopoda_schliemanni | 0 | 9.1 | 1 | 36340 |
| mammals | Naemorhedus_baileyi | Naemorhedus_baileyi | 1 | 27487.5 | 1.25 | 124540.26 |
| mammals | Naemorhedus_caudatus | Naemorhedus_caudatus | 1 | 27000 | 1 | 184715.63 |
| mammals | Naemorhedus_goral | Naemorhedus_goral | 0 | 28500 | 1 | 250558.82 |
| mammals | Nandinia_binotata | Nandinia_binotata | 0 | 2000 | 1.92 | 6021747.82 |
| mammals | Nanger_dama | Nanger_dama | 1 | 72999.5 | 1 | 230062.58 |
| mammals | Nanger_granti | Nanger_granti | 0 | 54999.7 | 1 | 785168.61 |
| mammals | Nanger_soemmerringii | Nanger_soemmerringii | 1 | 40000 | 1 | 291391.03 |
| mammals | Nanonycteris_veldkampii | Nanonycteris_veldkampii | 0 | 21.4 | 0.98 | 1711765.06 |
| mammals | Napaeozapus_insignis | Napaeozapus_insignis | 0 | 22.3 | 4.48 | 2010052.08 |
| mammals | Nasalis_larvatus | Nasalis_larvatus | 1 | 9000 | 1.01 | 538836.41 |
| mammals | Nasua_narica | Nasua_narica | 0 | 4030 | 4.08 | 2698807.16 |
| mammals | Nasua_nasua | Nasua_nasua | 0 | 3793.8 | 3.75 | 10629424.44 |
| mammals | Nasua_olivacea | Nasuella_olivacea | 0 | 1340 | 4 | 266801.18 |

| | | | | | | |
|---|---|---|---|---|---|---|
| mammals | Natalus_jamaicensis | Natalus_jamaicensis | 1 | 6.7 | 1 | 11024.67 |
| mammals | Natalus_macrourus | Natalus_espiritosantensis | 0 | 6 | 1 | 2483320.53 |
| mammals | Natalus_major | Natalus_major | 0 | 7.1 | 1 | 74327.94 |
| mammals | Natalus_mexicanus | Natalus_mexicanus | 0 | 5.3 | 1 | 2430998.38 |
| mammals | Natalus_primus | Natalus_primus | 1 | 9.1 | 1 | 3130.37 |
| mammals | Natalus_stramineus | Natalus_stramineus | 0 | 5 | 1 | 4059.92 |
| mammals | Natalus_tumidirostris | Natalus_tumidirostris | 0 | 6.4 | 1 | 1390815.57 |
| mammals | Neacomys_minutus | Neacomys_minutus | 0 | 10.4 | 3 | 183731.32 |
| mammals | Neacomys_musseri | Neacomys_musseri | 0 | 12.1 | 2 | 140347.47 |
| mammals | Neacomys_tenuipes | Neacomys_tenuipes | 0 | 19 | 3.9 | 224158.5 |
| mammals | Necromys_lasiurus | Necromys_lasiurus | 0 | 39.9 | 4.42 | 6100713.85 |
| mammals | Necromys_lenguarum | Necromys_lenguarum | 0 | 29.9 | 4.17 | 592293.39 |
| mammals | Necromys_obscurus | Necromys_obscurus | 0 | 40.7 | 4.86 | 45303.4 |
| mammals | Necromys_urichi | Necromys_urichi | 0 | 34 | 5.14 | 339590.64 |
| mammals | Nectomys_apicalis | Nectomys_apicalis | 0 | 297.4 | 4.32 | 1077402.36 |
| mammals | Nectomys_palmipes | Nectomys_palmipes | 0 | 197 | 4.32 | 165524.74 |
| mammals | Nectomys_rattus | Nectomys_rattus | 0 | 248.8 | 4.5 | 7115323.82 |
| mammals | Nectomys_squamipes | Nectomys_squamipes | 0 | 190.7 | 4.41 | 2344287.57 |
| mammals | Neodon_irene | Neodon_irene | 0 | 44.6 | 3 | 635022.84 |
| mammals | Neodon_leucurus | Phaiomys_leucurus | 0 | 38 | 1.94 | 2268403.08 |
| mammals | Neoeptesicus_andinus | Eptesicus_andinus | 0 | 9.3 | 1.46 | 811200.27 |
| mammals | Neoeptesicus_brasiliensis | Eptesicus_brasiliensis | 0 | 9.5 | 1.46 | 13117424.91 |
| mammals | Neoeptesicus_chiriquinus | Eptesicus_chiriquinus | 0 | 11.3 | 1.46 | 274035.83 |
| mammals | Neoeptesicus_furinalis | Eptesicus_furinalis | 0 | 7.7 | 1.86 | 16292708.03 |
| mammals | Neofelis_diardi | Neofelis_diardi | 1 | 20500 | 2.25 | 510006.38 |
| mammals | Neofelis_nebulosa | Neofelis_nebulosa | 1 | 20500 | 2.25 | 5183470.92 |
| mammals | Neofiber_alleni | Neofiber_alleni | 0 | 266 | 2.33 | 119935.2 |
| mammals | Neogale_frenata | Mustela_frenata | 0 | 147 | 6.36 | 12272775.39 |
| mammals | Neogale_vison | Neovison_vison | 0 | 945 | 4.76 | 14573783.89 |
| mammals | Neomys_anomalus | Neomys_anomalus | 0 | 16 | 6.71 | 3282589.82 |
| mammals | Neomys_fodiens | Neomys_fodiens | 0 | 14.3 | 5.95 | 13089290.24 |
| mammals | Neomys_teres | Neomys_teres | 0 | 12.9 | 5.53 | 252150.28 |
| mammals | Neophascogale_lorentzii | Neophascogale_lorentzii | 0 | 212 | 4 | 120447.44 |
| mammals | Neoplatymops_mattogrossensis | Molossops_mattogrossensis | 0 | 6.8 | 1.05 | 6688972.69 |

| | | | | | | |
|---|---|---|---|---|---|---|
| mammals | Neopteryx_frosti | Neopteryx_frosti | 1 | 225 | 0.98 | 6744 |
| mammals | Neoromicia_anchietae | Pipistrellus_anchietae | 0 | 4.6 | 2 | 1457004.26 |
| mammals | Neoromicia_somalica | Neoromicia_somalica | 0 | 3.8 | 2 | 6070953.48 |
| mammals | Neoromicia_zuluensis | Neoromicia_zuluensis | 0 | 3.9 | 2 | 3479467.79 |
| mammals | Neotamias_alpinus | Neotamias_alpinus | 0 | 36.6 | 4.5 | 8864.5 |
| mammals | Neotamias_amoenus | Neotamias_amoenus | 0 | 50.5 | 5.09 | 1102204.16 |
| mammals | Neotamias_bulleri | Neotamias_bulleri | 1 | 100 | 2.45 | 15555.61 |
| mammals | Neotamias_canipes | Neotamias_canipes | 0 | 70.4 | 4.16 | 18918.83 |
| mammals | Neotamias_cinereicollis | Neotamias_cinereicollis | 0 | 61.7 | 4.92 | 74133.93 |
| mammals | Neotamias_dorsalis | Neotamias_dorsalis | 0 | 71.1 | 5.06 | 632158.15 |
| mammals | Neotamias_durangae | Neotamias_durangae | 0 | 85 | 2.42 | 47076.71 |
| mammals | Neotamias_merriami | Neotamias_merriami | 0 | 74.8 | 4.3 | 72203.81 |
| mammals | Neotamias_minimus | Neotamias_minimus | 0 | 135.3 | 4.95 | 5243481.99 |
| mammals | Neotamias_obscurus | Neotamias_obscurus | 0 | 73 | 3.55 | 13472.17 |
| mammals | Neotamias_ochrogenys | Neotamias_ochrogenys | 0 | 91.7 | 3.94 | 9534.33 |
| mammals | Neotamias_palmeri | Neotamias_palmeri | 1 | 69.4 | 3.93 | 303.95 |
| mammals | Neotamias_panamintinus | Neotamias_panamintinus | 0 | 52.3 | 4.07 | 37380.51 |
| mammals | Neotamias_quadrimaculatus | Neotamias_quadrimaculatus | 0 | 85.2 | 5.52 | 25543.02 |
| mammals | Neotamias_quadrivittatus | Neotamias_quadrivittatus | 0 | 62.2 | 4.03 | 296948.74 |
| mammals | Neotamias_ruficaudus | Neotamias_ruficaudus | 0 | 68 | 4.85 | 147850.63 |
| mammals | Neotamias_rufus | Neotamias_rufus | 0 | 53.6 | 4.28 | 141470.69 |
| mammals | Neotamias_senex | Neotamias_senex | 0 | 89.3 | 4.35 | 99353.86 |
| mammals | Neotamias_siskiyou | Neotamias_siskiyou | 0 | 75 | 4.23 | 22528.82 |
| mammals | Neotamias_sonomae | Neotamias_sonomae | 0 | 75 | 4 | 45779.92 |
| mammals | Neotamias_speciosus | Neotamias_speciosus | 0 | 62 | 4.14 | 67800.86 |
| mammals | Neotamias_townsendii | Neotamias_townsendii | 0 | 74.8 | 3.99 | 151287.63 |
| mammals | Neotamias_umbrinus | Neotamias_umbrinus | 0 | 63 | 5.97 | 295485.18 |
| mammals | Neotoma_albigula | Neotoma_albigula | 0 | 206 | 2.14 | 834340.75 |
| mammals | Neotoma_angustapalata | Neotoma_angustapalata | 0 | 198 | 2.35 | 3326.83 |
| mammals | Neotoma_bryanti | Neotoma_bryanti | 0 | 182 | 2.92 | 271446.01 |
| mammals | Neotoma_chrysomelas | Neotoma_chrysomelas | 0 | 169.5 | 1.37 | 30203.6 |
| mammals | Neotoma_cinerea | Neotoma_cinerea | 0 | 299.2 | 3.51 | 3608407.83 |
| mammals | Neotoma_devia | Neotoma_devia | 0 | 200 | 2.81 | 73683.61 |
| mammals | Neotoma_floridana | Neotoma_floridana | 0 | 244.7 | 2.87 | 1871086.9 |

| | | | | | | |
|---|---|---|---|---|---|---|
| mammals | Neotoma_fuscipes | Neotoma_fuscipes | 0 | 229.8 | 2.76 | 171574.87 |
| mammals | Neotoma_goldmani | Neotoma_goldmani | 0 | 198 | 1.68 | 170757.26 |
| mammals | Neotoma_lepida | Neotoma_lepida | 0 | 163.8 | 2.81 | 675858.65 |
| mammals | Neotoma_leucodon | Neotoma_leucodon | 0 | 251.1 | 2.14 | 1076957.59 |
| mammals | Neotoma_macrotis | Neotoma_macrotis | 0 | 198 | 2.72 | 160394.34 |
| mammals | Neotoma_magister | Neotoma_magister | 0 | 412.5 | 2.08 | 313436.47 |
| mammals | Neotoma_mexicana | Neotoma_mexicana | 0 | 203 | 2.47 | 1784429.13 |
| mammals | Neotoma_micropus | Neotoma_micropus | 0 | 237.5 | 2.33 | 1063795.71 |
| mammals | Neotoma_nelsoni | Neotoma_nelsoni | 1 | 198 | 2.14 | 45.83 |
| mammals | Neotoma_palatina | Neotoma_palatina | 1 | 198 | 2.14 | 16059.98 |
| mammals | Neotoma_phenax | Neotoma_phenax | 0 | 227.5 | 2 | 42633.38 |
| mammals | Neotoma_stephensi | Neotoma_stephensi | 0 | 152.5 | 1.88 | 226739.19 |
| mammals | Neotomodon_alstoni | Neotomodon_alstoni | 0 | 40 | 3.22 | 27485.6 |
| mammals | Neotragus_pygmaeus | Neotragus_pygmaeus | 0 | 4000 | 1 | 455193.27 |
| mammals | Nephelomys_albigularis | Nephelomys_albigularis | 0 | 60.5 | 3.4 | 398224.72 |
| mammals | Nephelomys_auriventer | Nephelomys_auriventer | 0 | 117.5 | 3.4 | 14415.64 |
| mammals | Nephelomys_caracolus | Nephelomys_caracolus | 0 | 93 | 3.4 | 4076.97 |
| mammals | Nephelomys_devius | Nephelomys_devius | 0 | 150 | 3.4 | 9591 |
| mammals | Nephelomys_keaysi | Nephelomys_keaysi | 0 | 163.2 | 3.4 | 67023.2 |
| mammals | Nephelomys_levipes | Nephelomys_levipes | 0 | 112.8 | 3.4 | 157127.9 |
| mammals | Nephelomys_meridensis | Nephelomys_meridensis | 0 | 99.3 | 3.4 | 14351.08 |
| mammals | Nesogale_dobsoni | Microgale_dobsoni | 0 | 38 | 2.66 | 148370.96 |
| mammals | Nesogale_talazaci | Microgale_talazaci | 0 | 48 | 2.23 | 92303.03 |
| mammals | Nesomys_audeberti | Nesomys_audeberti | 0 | 212.5 | 2 | 90723.08 |
| mammals | Nesomys_lambertoni | Nesomys_lambertoni | 1 | 155 | 1 | 2220.49 |
| mammals | Nesomys_rufus | Nesomys_rufus | 0 | 162.5 | 4 | 106783.17 |
| mammals | Nesonycteris_woodfordi | Melonycteris_woodfordi | 0 | 29.2 | 0.98 | 16782.92 |
| mammals | Nesoryzomys_swarthi | Nesoryzomys_swarthi | 1 | 105.8 | 3 | 573.6 |
| mammals | Nesotragus_batesi | Neotragus_batesi | 0 | 3000 | 1 | 802403.34 |
| mammals | Nesotragus_moschatus | Nesotragus_moschatus | 0 | 6500 | 1 | 1370859 |
| mammals | Neurotrichus_gibbsii | Neurotrichus_gibbsii | 0 | 8.9 | 2.9 | 221364.52 |
| mammals | Neusticomys_monticolus | Neusticomys_monticolus | 0 | 39.5 | 1.94 | 65828.27 |
| mammals | Nilgiritragus_hylocrius | Nilgiritragus_hylocrius | 1 | 75000 | 1.35 | 2096.84 |
| mammals | Ningaui_ridei | Ningaui_ridei | 0 | 8.5 | 5.99 | 2252519.9 |

| | | | | | | |
|---|---|---|---|---|---|---|
| mammals | Ningaui_timealeyi | Ningaui_timealeyi | 0 | 5.7 | 5.78 | 250377.15 |
| mammals | Ningaui_yvonneae | Ningaui_yvonneae | 0 | 7 | 6 | 308584.03 |
| mammals | Niviventer_cameroni | Niviventer_cameroni | 1 | 86.8 | 4.41 | 9974.68 |
| mammals | Niviventer_confucianus | Niviventer_confucianus | 0 | 65 | 3.7 | 3971381.01 |
| mammals | Niviventer_coninga | Niviventer_coninga | 0 | 142 | 2.37 | 28365.69 |
| mammals | Niviventer_cremoriventer | Niviventer_cremoriventer | 0 | 69.7 | 3.7 | 1248372.6 |
| mammals | Niviventer_culturatus | Niviventer_culturatus | 0 | 90.5 | 3 | 3736.76 |
| mammals | Niviventer_eha | Niviventer_eha | 0 | 53.6 | 4.86 | 183562.82 |
| mammals | Niviventer_fraternus | Niviventer_fraternus | 0 | 86.8 | 4.41 | 111954.44 |
| mammals | Niviventer_fulvescens | Niviventer_fulvescens | 0 | 82.5 | 4.41 | 4720379.17 |
| mammals | Niviventer_tenaster | Niviventer_tenaster | 0 | 103 | 3.7 | 266057.26 |
| mammals | Noctilio_albiventris | Noctilio_albiventris | 0 | 30.8 | 1 | 12855674.66 |
| mammals | Noctilio_leporinus | Noctilio_leporinus | 0 | 58.3 | 1 | 14625648.25 |
| mammals | Nomascus_concolor | Nomascus_concolor | 1 | 7500 | 1 | 26326.31 |
| mammals | Nomascus_gabriellae | Nomascus_gabriellae | 1 | 8000 | 1 | 39684.78 |
| mammals | Nomascus_leucogenys | Nomascus_leucogenys | 1 | 8000 | 1 | 51340.52 |
| mammals | Nomascus_nasutus | Nomascus_nasutus | 1 | 8000 | 1 | 8311.9 |
| mammals | Nomascus_siki | Nomascus_siki | 1 | 8000 | 1 | 26549.04 |
| mammals | Notamacropus_agilis | Macropus_agilis | 0 | 15000 | 1.01 | 1325773.81 |
| mammals | Notamacropus_dorsalis | Macropus_dorsalis | 0 | 11250 | 1 | 570362.34 |
| mammals | Notamacropus_eugenii | Macropus_eugenii | 0 | 6500 | 1.01 | 16450.35 |
| mammals | Notamacropus_irma | Macropus_irma | 0 | 8000 | 1 | 152879.45 |
| mammals | Notamacropus_parma | Macropus_parma | 0 | 4500 | 0.92 | 50622.9 |
| mammals | Notamacropus_parryi | Macropus_parryi | 0 | 13500 | 1 | 362773.54 |
| mammals | Notamacropus_rufogriseus | Macropus_rufogriseus | 0 | 16850 | 1.01 | 652547.91 |
| mammals | Nothocricetulus_migratorius | Cricetulus_migratorius | 0 | 38.5 | 4.92 | 7677961.28 |
| mammals | Notiosorex_crawfordi | Notiosorex_crawfordi | 0 | 4.4 | 3.72 | 2165720.11 |
| mammals | Notiosorex_evotis | Notiosorex_evotis | 0 | 5.4 | 3.76 | 89137.28 |
| mammals | Notocitellus_annulatus | Notocitellus_annulatus | 0 | 500 | 3.91 | 44731.3 |
| mammals | Notomys_alexis | Notomys_alexis | 0 | 35 | 3.96 | 2927774.01 |
| mammals | Notomys_aquilo | Notomys_aquilo | 1 | 39 | 3 | 2410.57 |
| mammals | Notomys_cervinus | Notomys_cervinus | 0 | 35 | 2.92 | 83793.89 |
| mammals | Notomys_fuscus | Notomys_fuscus | 1 | 35 | 3.03 | 105460.26 |
| mammals | Notomys_mitchellii | Notomys_mitchellii | 0 | 52 | 3.28 | 646728.27 |

| | | | | | | |
|---|---|---|---|---|---|---|
| mammals | Notopteris_macdonaldii | Notopteris_macdonaldi | 1 | 65.1 | 0.99 | 19268.55 |
| mammals | Notopteris_neocaledonicus | Notopteris_neocaledonica | 1 | 45.1 | 1 | 1254.14 |
| mammals | Nyctalus_aviator | Nyctalus_aviator | 0 | 38.3 | 1.89 | 846242.21 |
| mammals | Nyctalus_azoreum | Nyctalus_azoreum | 1 | 7.1 | 1.5 | 2149.99 |
| mammals | Nyctalus_furvus | Nyctalus_furvus | 1 | 27.6 | 1.17 | 11560.16 |
| mammals | Nyctalus_lasiopterus | Nyctalus_lasiopterus | 1 | 49.7 | 1.82 | 2917259.33 |
| mammals | Nyctalus_leisleri | Nyctalus_leisleri | 0 | 13.3 | 1.61 | 5170891.77 |
| mammals | Nyctalus_montanus | Nyctalus_montanus | 0 | 14.2 | 1.5 | 123577.43 |
| mammals | Nyctalus_noctula | Nyctalus_noctula | 0 | 28 | 1.25 | 7210092.69 |
| mammals | Nyctalus_plancyi | Nyctalus_plancyi | 0 | 24.1 | 1.17 | 2017209.65 |
| mammals | Nyctereutes_procyonoides | Nyctereutes_procyonoides | 0 | 4040 | 6.33 | 11549507.06 |
| mammals | Nycteris_arge | Nycteris_arge | 0 | 12 | 0.98 | 3980021.53 |
| mammals | Nycteris_aurita | Nycteris_aurita | 0 | 10 | 1 | 1789679.26 |
| mammals | Nycteris_gambiensis | Nycteris_gambiensis | 0 | 7 | 1 | 1861005.07 |
| mammals | Nycteris_grandis | Nycteris_grandis | 0 | 30.2 | 1 | 4404698.71 |
| mammals | Nycteris_hispida | Nycteris_hispida | 0 | 8.6 | 0.99 | 11511751.47 |
| mammals | Nycteris_intermedia | Nycteris_intermedia | 0 | 10.7 | 1 | 2619716.27 |
| mammals | Nycteris_javanica | Nycteris_javanica | 1 | 17.7 | 1 | 145230.85 |
| mammals | Nycteris_macrotis | Nycteris_macrotis | 0 | 14.5 | 0.99 | 12141283.97 |
| mammals | Nycteris_nana | Nycteris_nana | 0 | 7 | 1 | 2938142.02 |
| mammals | Nycteris_thebaica | Nycteris_thebaica | 0 | 9.8 | 1 | 14428246.95 |
| mammals | Nycteris_tragata | Nycteris_tragata | 0 | 14.4 | 1 | 1394635.24 |
| mammals | Nycteris_woodi | Nycteris_woodi | 0 | 7.6 | 1 | 620419 |
| mammals | Nycticebus_bengalensis | Nycticebus_bengalensis | 1 | 1500 | 1.03 | 2049989.22 |
| mammals | Nycticebus_coucang | Nycticebus_coucang | 1 | 803.5 | 1.08 | 441468.37 |
| mammals | Nycticebus_menagensis | Nycticebus_menagensis | 1 | 803.5 | 1 | 727974.52 |
| mammals | Nycticeinops_crassulus | Pipistrellus_crassulus | 0 | 6.1 | 1 | 2601004.41 |
| mammals | Nycticeinops_schlieffenii | Nycticeinops_schlieffeni | 0 | 5 | 2.54 | 5499366.94 |
| mammals | Nycticeius_cubanus | Nycticeius_cubanus | 0 | 5.5 | 1.47 | 29183.54 |
| mammals | Nycticeius_humeralis | Nycticeius_humeralis | 0 | 9 | 1.71 | 2883260.06 |
| mammals | Nyctiellus_lepidus | Nyctiellus_lepidus | 0 | 2.4 | 1 | 110184.1 |
| mammals | Nyctimene_aello | Nyctimene_aello | 0 | 84.3 | 0.98 | 621332.74 |
| mammals | Nyctimene_albiventer | Nyctimene_albiventer | 0 | 29.9 | 0.98 | 748029.69 |
| mammals | Nyctimene_cephalotes | Nyctimene_cephalotes | 0 | 44.5 | 0.98 | 239455.09 |

| | | | | | | |
|---|---|---|---|---|---|---|
| mammals | Nyctimene_certans | Nyctimene_certans | 0 | 42.1 | 1 | 212379.59 |
| mammals | Nyctimene_keasti | Nyctimene_keasti | 0 | 44.6 | 0.99 | 47545.66 |
| mammals | Nyctimene_rabori | Nyctimene_rabori | 1 | 67.5 | 1 | 30770.51 |
| mammals | Nyctimene_robinsoni | Nyctimene_robinsoni | 0 | 49.5 | 0.99 | 351781.04 |
| mammals | Nyctimene_vizcaccia | Nyctimene_vizcaccia | 0 | 40 | 0.98 | 70176.73 |
| mammals | Nyctinomops_aurispinosus | Nyctinomops_aurispinosus | 0 | 18 | 1 | 3566231.21 |
| mammals | Nyctinomops_femorosaccus | Nyctinomops_femorosaccus | 0 | 14.6 | 0.99 | 1251911.46 |
| mammals | Nyctinomops_laticaudatus | Nyctinomops_laticaudatus | 0 | 12.2 | 0.99 | 13101230.9 |
| mammals | Nyctinomops_macrotis | Nyctinomops_macrotis | 0 | 18.4 | 0.99 | 15743813.98 |
| mammals | Nyctomys_sumichrasti | Nyctomys_sumichrasti | 0 | 49.4 | 2 | 673131.85 |
| mammals | Nyctophilus_arnhemensis | Nyctophilus_arnhemensis | 0 | 6.7 | 2 | 704005.75 |
| mammals | Nyctophilus_bifax | Nyctophilus_bifax | 0 | 9.8 | 2 | 400008.97 |
| mammals | Nyctophilus_geoffroyi | Nyctophilus_geoffroyi | 0 | 7.3 | 2 | 7272506.25 |
| mammals | Nyctophilus_gouldi | Nyctophilus_gouldi | 0 | 10 | 1.66 | 1499403.09 |
| mammals | Nyctophilus_sherrini | Nyctophilus_sherrini | 1 | 15.5 | 2 | 48998.26 |
| mammals | Nyctophilus_walkeri | Nyctophilus_walkeri | 0 | 4.3 | 2 | 397742.81 |
| mammals | Ochotona_alpina | Ochotona_alpina | 0 | 263 | 3.25 | 575019.44 |
| mammals | Ochotona_argentata | Ochotona_argentata | 1 | 206 | 4.69 | 95.49 |
| mammals | Ochotona_cansus | Ochotona_cansus | 0 | 74.5 | 3.47 | 700340.63 |
| mammals | Ochotona_collaris | Ochotona_collaris | 0 | 129 | 3.46 | 846284.43 |
| mammals | Ochotona_curzoniae | Ochotona_curzoniae | 0 | 162.5 | 4.77 | 2250961.79 |
| mammals | Ochotona_dauurica | Ochotona_dauurica | 0 | 130 | 6.06 | 2054450.61 |
| mammals | Ochotona_erythrotis | Ochotona_erythrotis | 0 | 268 | 4.1 | 961201.75 |
| mammals | Ochotona_forresti | Ochotona_forresti | 0 | 129 | 5.7 | 38532.76 |
| mammals | Ochotona_hoffmanni | Ochotona_hoffmanni | 1 | 285 | 3.42 | 1241.7 |
| mammals | Ochotona_hyperborea | Ochotona_hyperborea | 0 | 156 | 3.04 | 8990041.4 |
| mammals | Ochotona_koslowi | Ochotona_koslowi | 1 | 165 | 1 | 44741.58 |
| mammals | Ochotona_macrotis | Ochotona_macrotis | 0 | 166 | 3.74 | 2931184.45 |
| mammals | Ochotona_mantchurica | Ochotona_mantchurica | 0 | 160 | 4 | 336661.15 |
| mammals | Ochotona_nubrica | Ochotona_nubrica | 0 | 115.5 | 8.41 | 491702.49 |
| mammals | Ochotona_opaca | Ochotona_opaca | 0 | 136.5 | 7.5 | 179949.76 |
| mammals | Ochotona_pallasii | Ochotona_pallasii | 0 | 214 | 6.99 | 941438.89 |
| mammals | Ochotona_princeps | Ochotona_princeps | 0 | 157.6 | 2.94 | 894668.9 |
| mammals | Ochotona_pusilla | Ochotona_pusilla | 0 | 142.5 | 8.54 | 1530161.5 |

| | | | | | | |
|---|---|---|---|---|---|---|
| mammals | Ochotona_roylii | Ochotona_roylei | 0 | 260 | 3 | 493849.81 |
| mammals | Ochotona_rufescens | Ochotona_rufescens | 0 | 250 | 5.97 | 946121.48 |
| mammals | Ochotona_rutila | Ochotona_rutila | 0 | 270 | 4.1 | 206939.19 |
| mammals | Ochotona_thibetana | Ochotona_thibetana | 0 | 104 | 2.93 | 729354.29 |
| mammals | Ochotona_turuchanensis | Ochotona_turuchanensis | 0 | 148.2 | 3.07 | 1642659.49 |
| mammals | Ochromyscus_brockmani | Myomyscus_brockmani | 0 | 29.5 | 6.5 | 1713270.66 |
| mammals | Ochrotomys_nuttalli | Ochrotomys_nuttalli | 0 | 22.4 | 2.63 | 1444556.94 |
| mammals | Octodon_bridgesii | Octodon_bridgesi | 1 | 150 | 2.5 | 120967.76 |
| mammals | Octodon_degus | Octodon_degus | 0 | 210 | 5.31 | 63501.35 |
| mammals | Octodontomys_gliroides | Octodontomys_gliroides | 0 | 150 | 2.04 | 487312.91 |
| mammals | Odocoileus_hemionus | Odocoileus_hemionus | 0 | 54212.6 | 1.61 | 6340068.2 |
| mammals | Odocoileus_pandora | Mazama_pandora | 1 | 21000 | 1 | 146873.17 |
| mammals | Odocoileus_virginianus | Odocoileus_virginianus | 0 | 55508.6 | 1.64 | 14410637.4 |
| mammals | Oecomys_bicolor | Oecomys_bicolor | 0 | 34 | 3.15 | 8594483.01 |
| mammals | Oecomys_flavicans | Oecomys_flavicans | 0 | 73.4 | 4.5 | 42577.44 |
| mammals | Oecomys_trinitatis | Oecomys_trinitatis | 0 | 73.4 | 3 | 4830193.88 |
| mammals | Oenomys_hypoxanthus | Oenomys_hypoxanthus | 0 | 90 | 3.09 | 4172195.57 |
| mammals | Okapia_johnstoni | Okapia_johnstoni | 1 | 230001.1 | 0.99 | 264220.22 |
| mammals | Oligoryzomys_chacoensis | Oligoryzomys_chacoensis | 0 | 23 | 3.67 | 760937.18 |
| mammals | Oligoryzomys_destructor | Oligoryzomys_destructor | 0 | 25.2 | 4.38 | 661051.39 |
| mammals | Oligoryzomys_flavescens | Oligoryzomys_flavescens | 0 | 21.3 | 4.56 | 4603205 |
| mammals | Oligoryzomys_fulvescens | Oligoryzomys_fulvescens | 0 | 25 | 3.64 | 2895296.89 |
| mammals | Oligoryzomys_longicaudatus | Oligoryzomys_longicaudatus | 0 | 27 | 4.58 | 745244.39 |
| mammals | Oligoryzomys_microtis | Oligoryzomys_microtis | 0 | 22.5 | 3.4 | 2722586.2 |
| mammals | Oligoryzomys_nigripes | Oligoryzomys_nigripes | 0 | 20.5 | 3.52 | 3221464.72 |
| mammals | Oligoryzomys_stramineus | Oligoryzomys_stramineus | 0 | 25 | 4.5 | 530369.85 |
| mammals | Oligoryzomys_vegetus | Oligoryzomys_vegetus | 0 | 15 | 3.64 | 9901.69 |
| mammals | Ondatra_zibethicus | Ondatra_zibethicus | 0 | 1065.7 | 6.48 | 35093842.14 |
| mammals | Onychogalea_frenata | Onychogalea_fraenata | 1 | 5500 | 1 | 4205.87 |
| mammals | Onychogalea_unguifera | Onychogalea_unguifera | 0 | 6650 | 1 | 951054.22 |
| mammals | Onychomys_arenicola | Onychomys_arenicola | 0 | 30 | 3.66 | 686486.36 |
| mammals | Onychomys_leucogaster | Onychomys_leucogaster | 0 | 27.9 | 3.84 | 3841451.17 |
| mammals | Onychomys_torridus | Onychomys_torridus | 0 | 25 | 3.48 | 767649.88 |
| mammals | Oreamnos_americanus | Oreamnos_americanus | 0 | 72500.3 | 1.36 | 838023.45 |

| | | | | | | |
|---|---|---|---|---|---|---|
| mammals | Oreoscaptor_mizura | Euroscaptor_mizura | 0 | 35.5 | 3 | 15898.04 |
| mammals | Oreotragus_oreotragus | Oreotragus_oreotragus | 0 | 12999.9 | 0.99 | 5220692.4 |
| mammals | Orientallactaga_balikunica | Allactaga_balikunica | 0 | 72.5 | 2.52 | 234834.4 |
| mammals | Orientallactaga_bullata | Allactaga_bullata | 0 | 87 | 2.52 | 1563011.22 |
| mammals | Orientallactaga_sibirica | Allactaga_sibirica | 0 | 97.5 | 3.44 | 5412810.8 |
| mammals | Ornithorhynchus_anatinus | Ornithorhynchus_anatinus | 0 | 1484.3 | 2 | 901638 |
| mammals | Orthogeomys_grandis | Orthogeomys_cuniculus | 0 | 500 | 2 | 140934.7 |
| mammals | Orycteropus_afer | Orycteropus_afer | 0 | 52350.4 | 1.06 | 19273002.49 |
| mammals | Oryctolagus_cuniculus | Oryctolagus_cuniculus | 1 | 1958.3 | 5.24 | 634229.13 |
| mammals | Oryx_beisa | Oryx_beisa | 1 | 170000 | 1 | 975973.87 |
| mammals | Oryx_gazella | Oryx_gazella | 0 | 169001.3 | 1 | 1766027.32 |
| mammals | Oryx_leucoryx | Oryx_leucoryx | 1 | 87700 | 1 | 20909.66 |
| mammals | Oryzomys_couesi | Oryzomys_couesi | 0 | 69.3 | 3.7 | 1491857.12 |
| mammals | Oryzomys_palustris | Oryzomys_palustris | 0 | 53.9 | 4 | 1508984.73 |
| mammals | Oryzorictes_hova | Oryzorictes_hova | 0 | 36.5 | 3.5 | 124191.68 |
| mammals | Osgoodomys_banderanus | Osgoodomys_banderanus | 0 | 50 | 1.68 | 179803.69 |
| mammals | Osphranter_antilopinus | Macropus_antilopinus | 0 | 27250 | 1 | 653127.96 |
| mammals | Osphranter_bernardus | Macropus_bernardus | 0 | 17000 | 1 | 30111.04 |
| mammals | Osphranter_robustus | Macropus_robustus | 0 | 21262.5 | 1 | 6349650.74 |
| mammals | Osphranter_rufus | Macropus_rufus | 0 | 46250 | 1 | 5360863.5 |
| mammals | Otocolobus_manul | Otocolobus_manul | 0 | 3500 | 4.48 | 8387217.25 |
| mammals | Otocyon_megalotis | Otocyon_megalotis | 0 | 4150 | 4 | 4957043.24 |
| mammals | Otolemur_crassicaudatus | Otolemur_crassicaudatus | 0 | 1500 | 1.32 | 4845511.43 |
| mammals | Otolemur_garnettii | Otolemur_garnettii | 0 | 760 | 1.14 | 568222.41 |
| mammals | Otomops_madagascariensis | Otomops_madagascariensis | 0 | 25.5 | 0.99 | 3539.5 |
| mammals | Otomops_martiensseni | Otomops_martiensseni | 0 | 34.3 | 0.99 | 5186383.98 |
| mammals | Otomys_anchietae | Otomys_anchietae | 0 | 206.5 | 1.72 | 71608.94 |
| mammals | Otomys_angoniensis | Otomys_angoniensis | 0 | 121.9 | 2.59 | 595014.77 |
| mammals | Otomys_barbouri | Otomys_barbouri | 1 | 107.4 | 1.72 | 398.1 |
| mammals | Otomys_burtoni | Otomys_burtoni | 1 | 90 | 1.78 | 383.86 |
| mammals | Otomys_cuanzensis | Otomys_cuanzensis | 0 | 90.2 | 2.86 | 115191.67 |
| mammals | Otomys_denti | Otomys_denti | 0 | 120 | 1.64 | 80971.54 |
| mammals | Otomys_irroratus | Otomys_irroratus | 0 | 118 | 1.68 | 183846.81 |
| mammals | Otomys_karoensis | Otomys_saundersiae | 0 | 103 | 2.32 | 254423.21 |

| | | | | | | |
|---|---|---|---|---|---|---|
| mammals | Otomys_lacustris | Otomys_lacustris | 0 | 92.6 | 1.72 | 34134.57 |
| mammals | Otomys_laminatus | Otomys_laminatus | 0 | 145 | 1.72 | 86794.6 |
| mammals | Otomys_sloggetti | Otomys_sloggetti | 0 | 106.3 | 1.44 | 156904.33 |
| mammals | Otomys_tropicalis | Otomys_tropicalis | 0 | 108.8 | 1.47 | 143756.98 |
| mammals | Otomys_typus | Otomys_typus | 0 | 133.4 | 1.65 | 96218.78 |
| mammals | Otomys_unisulcatus | Otomys_unisulcatus | 0 | 124.5 | 1.42 | 374301.34 |
| mammals | Otonycteris_hemprichii | Otonycteris_hemprichii | 0 | 23.8 | 1.38 | 1283936.36 |
| mammals | Otonyctomys_hatti | Otonyctomys_hatti | 0 | 36.2 | 1.88 | 127970.47 |
| mammals | Otopteropus_cartilagonodus | Otopteropus_cartilagonodus | 0 | 16.8 | 0.99 | 105071.32 |
| mammals | Otospermophilus_beecheyi | Otospermophilus_beecheyi | 0 | 578.5 | 6.71 | 467246.65 |
| mammals | Otospermophilus_variegatus | Otospermophilus_variegatus | 0 | 691.6 | 4.72 | 2304576.89 |
| mammals | Ototylomys_phyllotis | Ototylomys_phyllotis | 0 | 120 | 2.22 | 491821.32 |
| mammals | Ourebia_ourebi | Ourebia_ourebi | 0 | 17250 | 0.99 | 7497157.68 |
| mammals | Ovibos_moschatus | Ovibos_moschatus | 0 | 340501.1 | 1.01 | 3760945.12 |
| mammals | Ovis_ammon | Ovis_ammon | 0 | 180000 | 1.21 | 2998114.01 |
| mammals | Ovis_canadensis | Ovis_canadensis | 0 | 74644.9 | 1.09 | 538062.18 |
| mammals | Ovis_dalli | Ovis_dalli | 0 | 55650.6 | 1.14 | 816225.07 |
| mammals | Ovis_nivicola | Ovis_nivicola | 0 | 90000 | 1 | 1369336.66 |
| mammals | Oxymycterus_dasytrichus | Oxymycterus_dasytrichus | 0 | 72.5 | 2.77 | 1718385.51 |
| mammals | Oxymycterus_delator | Oxymycterus_delator | 0 | 81.5 | 4 | 82738.64 |
| mammals | Oxymycterus_inca | Oxymycterus_inca | 0 | 35 | 5 | 360850.75 |
| mammals | Oxymycterus_josei | Oxymycterus_josei | 0 | 66.4 | 2.77 | 22830 |
| mammals | Oxymycterus_nasutus | Oxymycterus_nasutus | 0 | 68 | 2.77 | 558988.94 |
| mammals | Oxymycterus_paramensis | Oxymycterus_paramensis | 0 | 42 | 5 | 286552.68 |
| mammals | Oxymycterus_quaestor | Oxymycterus_quaestor | 0 | 75.5 | 2.77 | 784234.92 |
| mammals | Oxymycterus_rufus | Oxymycterus_rufus | 0 | 75.4 | 2.64 | 280710.36 |
| mammals | Ozimops_beccarii | Mormopterus_beccarii | 0 | 14.9 | 1.05 | 90003.55 |
| mammals | Ozimops_planiceps | Mormopterus_planiceps | 0 | 8.8 | 0.99 | 3437648.75 |
| mammals | Ozotoceros_bezoarticus | Ozotoceros_bezoarticus | 0 | 40000 | 1 | 468419.42 |
| mammals | Pachyuromys_duprasi | Pachyuromys_duprasi | 0 | 47.5 | 3.78 | 1305337.93 |
| mammals | Paguma_larvata | Paguma_larvata | 0 | 4300 | 2.23 | 6472276.6 |
| mammals | Pan_paniscus | Pan_paniscus | 1 | 34000.1 | 1 | 415766.37 |
| mammals | Pan_troglodytes | Pan_troglodytes | 1 | 42500 | 1.06 | 2314272 |
| mammals | Panthera_leo | Panthera_leo | 1 | 161499.1 | 2.79 | 26107440.05 |

| | | | | | | |
|---|---|---|---|---|---|---|
| mammals | Panthera_onca | Panthera_onca | 0 | 100000 | 1.98 | 9025908.69 |
| mammals | Panthera_pardus | Panthera_pardus | 1 | 54999.7 | 2.14 | 34544707.15 |
| mammals | Panthera_tigris | Panthera_tigris | 1 | 162564 | 2.58 | 11805900.96 |
| mammals | Panthera_uncia | Panthera_uncia | 1 | 44167 | 2.12 | 3496431.34 |
| mammals | Pantholops_hodgsonii | Pantholops_hodgsonii | 0 | 27500 | 1 | 1090874.58 |
| mammals | Papio_anubis | Papio_anubis | 0 | 28329.2 | 1.01 | 7814659.71 |
| mammals | Papio_cynocephalus | Papio_cynocephalus | 0 | 19484 | 1 | 1667455.19 |
| mammals | Papio_hamadryas | Papio_hamadryas | 0 | 14972.2 | 1 | 610339.02 |
| mammals | Papio_papio | Papio_papio | 0 | 15500.2 | 1.01 | 494995.57 |
| mammals | Papio_ursinus | Papio_ursinus | 0 | 27939.1 | 1.01 | 3348995.38 |
| mammals | Parachoerus_wagneri | Catagonus_wagneri | 1 | 35566.4 | 2.46 | 304464.11 |
| mammals | Paracrocidura_maxima | Paracrocidura_maxima | 0 | 16 | 1 | 23382.42 |
| mammals | Paracrocidura_schoutedeni | Paracrocidura_schoutedeni | 0 | 13 | 1.7 | 671227.21 |
| mammals | Paracynictis_selousi | Paracynictis_selousi | 0 | 1640 | 2.12 | 1796820.97 |
| mammals | Paradipus_ctenodactylus | Paradipus_ctenodactylus | 0 | 109 | 3.4 | 386135.52 |
| mammals | Paradoxurus_hermaphroditus | Paradoxurus_hermaphroditus | 0 | 3156.7 | 3.29 | 7641014.96 |
| mammals | Paradoxurus_jerdoni | Paradoxurus_jerdoni | 0 | 2780.8 | 3 | 66739.96 |
| mammals | Paradoxurus_zeylonensis | Paradoxurus_zeylonensis | 0 | 2780.8 | 2.5 | 18391.66 |
| mammals | Paraechinus_aethiopicus | Paraechinus_aethiopicus | 0 | 188 | 3.49 | 3874413.08 |
| mammals | Paraechinus_hypomelas | Paraechinus_hypomelas | 0 | 213 | 3.47 | 2536454.33 |
| mammals | Paraechinus_micropus | Paraechinus_micropus | 0 | 171 | 3.78 | 414921.36 |
| mammals | Paraechinus_nudiventris | Paraechinus_nudiventris | 0 | 175.3 | 3 | 9534.75 |
| mammals | Paragalago_cocos | Galagoides_cocos | 0 | 144 | 1.57 | 9853.78 |
| mammals | Paragalago_granti | Galagoides_granti | 0 | 158.7 | 1.5 | 367704.73 |
| mammals | Paragalago_orinus | Galagoides_orinus | 1 | 97 | 1.5 | 18913.3 |
| mammals | Paragalago_rondoensis | Galagoides_rondoensis | 1 | 84.2 | 1.2 | 126.8 |
| mammals | Paragalago_zanzibaricus | Galagoides_zanzibaricus | 0 | 100.5 | 1.4 | 7077.55 |
| mammals | Parahyaena_brunnea | Parahyaena_brunnea | 0 | 32200.3 | 2.3 | 2290722.98 |
| mammals | Paramelomys_levipes | Paramelomys_levipes | 0 | 86.7 | 1.51 | 27271.38 |
| mammals | Paramelomys_lorentzii | Paramelomys_lorentzii | 0 | 150 | 1.41 | 175784.2 |
| mammals | Paramelomys_mollis | Paramelomys_mollis | 0 | 90.1 | 1.6 | 225112.95 |
| mammals | Paramelomys_moncktoni | Paramelomys_moncktoni | 0 | 95.4 | 2.11 | 79468.89 |
| mammals | Paramelomys_naso | Paramelomys_naso | 0 | 129.7 | 1.51 | 115422.82 |
| mammals | Paramelomys_platyops | Paramelomys_platyops | 0 | 104.2 | 1.96 | 496373.43 |

| | | | | | | |
|---|---|---|---|---|---|---|
| mammals | Paramelomys_rubex | Paramelomys_rubex | 0 | 46.6 | 2 | 234666.79 |
| mammals | Parantechinus_apicalis | Parantechinus_apicalis | 1 | 68.8 | 7.62 | 2838.73 |
| mammals | Paranyctimene_raptor | Paranyctimene_raptor | 0 | 24.3 | 1 | 356130.33 |
| mammals | Parastrellus_hesperus | Pipistrellus_hesperus | 0 | 3.7 | 1.86 | 2686280.32 |
| mammals | Paratriaenops_auritus | Paratriaenops_auritus | 1 | 5 | 1 | 10567.58 |
| mammals | Paratriaenops_furcula | Paratriaenops_furculus | 0 | 5.5 | 1 | 57158.77 |
| mammals | Paraxerus_alexandri | Paraxerus_alexandri | 0 | 52 | 0.98 | 259068.85 |
| mammals | Paraxerus_boehmi | Paraxerus_boehmi | 0 | 70 | 1.07 | 956025.61 |
| mammals | Paraxerus_cepapi | Paraxerus_cepapi | 0 | 180 | 2.02 | 2692246.21 |
| mammals | Paraxerus_flavovittis | Paraxerus_flavovittis | 0 | 160 | 1.27 | 666787.98 |
| mammals | Paraxerus_ochraceus | Paraxerus_ochraceus | 0 | 104.3 | 2.2 | 547064.7 |
| mammals | Paraxerus_palliatus | Paraxerus_palliatus | 0 | 309.3 | 1.6 | 1045560.15 |
| mammals | Paraxerus_poensis | Paraxerus_poensis | 0 | 124.5 | 1.49 | 1570579.53 |
| mammals | Pardofelis_marmorata | Pardofelis_marmorata | 0 | 2854.3 | 2 | 1388973.7 |
| mammals | Paremballonura_atrata | Paremballonura_atrata | 0 | 4.3 | 1 | 105682.26 |
| mammals | Paremballonura_tiavato | Paremballonura_tiavato | 0 | 3.3 | 1 | 149432.76 |
| mammals | Parotomys_brantsii | Parotomys_brantsii | 0 | 129.5 | 2.04 | 679427.62 |
| mammals | Parotomys_littledalei | Parotomys_littledalei | 0 | 127 | 2.91 | 597770.68 |
| mammals | Passalites_nemorivagus | Mazama_nemorivaga | 0 | 15000 | 1 | 7647916.97 |
| mammals | Paynomys_macronyx | Chelemys_macronyx | 0 | 73.3 | 4.37 | 322022.59 |
| mammals | Pectinator_spekei | Pectinator_spekei | 0 | 160 | 1.17 | 419154.66 |
| mammals | Pedetes_capensis | Pedetes_capensis | 0 | 3155 | 0.99 | 3817904.34 |
| mammals | Pedetes_surdaster | Pedetes_surdaster | 0 | 3500 | 1 | 457191.76 |
| mammals | Pekania_pennanti | Martes_pennanti | 0 | 4000 | 2.97 | 3721001.43 |
| mammals | Pelea_capreolus | Pelea_capreolus | 0 | 20000 | 1 | 557077.48 |
| mammals | Pelomys_fallax | Pelomys_fallax | 0 | 130 | 3.67 | 3495837.14 |
| mammals | Pelomys_minor | Pelomys_minor | 0 | 50 | 2.5 | 309824.59 |
| mammals | Pentalagus_furnessi | Pentalagus_furnessi | 1 | 2396.3 | 2.44 | 370.2 |
| mammals | Penthetor_lucasii | Penthetor_lucasi | 0 | 38.2 | 1 | 1288201.06 |
| mammals | Perameles_bougainville | Perameles_bougainville | 1 | 226 | 1.9 | 235.89 |
| mammals | Perameles_gunnii | Perameles_gunnii | 1 | 870 | 2.23 | 31989.71 |
| mammals | Perameles_nasuta | Perameles_nasuta | 0 | 975 | 2.5 | 411183.14 |
| mammals | Perimyotis_subflavus | Pipistrellus_subflavus | 1 | 5.8 | 1.97 | 4264005.66 |
| mammals | Perodicticus_edwardsi | Perodicticus_edwardsi | 0 | 1210 | 1.1 | 2432405.68 |

| | | | | | | |
|---|---|---|---|---|---|---|
| mammals | Perodicticus_potto | Perodicticus_potto | 0 | 1100 | 1.06 | 680452.65 |
| mammals | Perognathus_amplus | Perognathus_amplus | 0 | 11.7 | 4 | 122119.52 |
| mammals | Perognathus_fasciatus | Perognathus_fasciatus | 0 | 11.6 | 5.24 | 1039860.87 |
| mammals | Perognathus_flavescens | Perognathus_flavescens | 0 | 8.8 | 4.46 | 1542733.95 |
| mammals | Perognathus_flavus | Perognathus_flavus | 0 | 7.7 | 4.06 | 1540956.83 |
| mammals | Perognathus_inornatus | Perognathus_inornatus | 0 | 10.9 | 4.22 | 73759.51 |
| mammals | Perognathus_longimembris | Perognathus_longimembris | 0 | 7.6 | 4.45 | 520454.39 |
| mammals | Perognathus_merriami | Perognathus_merriami | 0 | 6.9 | 4.26 | 693402.51 |
| mammals | Perognathus_parvus | Perognathus_parvus | 0 | 21.8 | 4.83 | 921305.43 |
| mammals | Peromyscus_attwateri | Peromyscus_attwateri | 0 | 27.9 | 3.52 | 452891.85 |
| mammals | Peromyscus_aztecus | Peromyscus_aztecus | 0 | 40 | 3.34 | 218569.36 |
| mammals | Peromyscus_beatae | Peromyscus_beatae | 0 | 20.7 | 3.18 | 145828.32 |
| mammals | Peromyscus_boylii | Peromyscus_boylii | 0 | 21.4 | 3.18 | 2530151.14 |
| mammals | Peromyscus_bullatus | Peromyscus_bullatus | 1 | 40 | 3.49 | 65.19 |
| mammals | Peromyscus_californicus | Peromyscus_californicus | 0 | 43.5 | 1.97 | 132623.27 |
| mammals | Peromyscus_caniceps | Peromyscus_caniceps | 1 | 30.2 | 2.6 | 18.1 |
| mammals | Peromyscus_crinitus | Peromyscus_crinitus | 0 | 18 | 3.08 | 913321.97 |
| mammals | Peromyscus_difficilis | Peromyscus_difficilis | 0 | 28 | 3.69 | 546382.94 |
| mammals | Peromyscus_eremicus | Peromyscus_eremicus | 0 | 23.7 | 2.56 | 1217908.66 |
| mammals | Peromyscus_eva | Peromyscus_eva | 0 | 22 | 2.6 | 39058.95 |
| mammals | Peromyscus_fraterculus | Peromyscus_fraterculus | 0 | 24.3 | 2.6 | 144188.19 |
| mammals | Peromyscus_gossypinus | Peromyscus_gossypinus | 0 | 29.4 | 3.87 | 1148138.97 |
| mammals | Peromyscus_grandis | Peromyscus_grandis | 0 | 71 | 2.43 | 3981.9 |
| mammals | Peromyscus_gratus | Peromyscus_gratus | 0 | 27.4 | 3.44 | 598242.3 |
| mammals | Peromyscus_guatemalensis | Peromyscus_guatemalensis | 0 | 40 | 2.1 | 17875.07 |
| mammals | Peromyscus_gymnotis | Peromyscus_gymnotis | 0 | 40 | 2.42 | 42202.78 |
| mammals | Peromyscus_hooperi | Peromyscus_hooperi | 0 | 36 | 3.04 | 19804.78 |
| mammals | Peromyscus_hylocetes | Peromyscus_hylocetes | 0 | 42.6 | 3.19 | 37378.53 |
| mammals | Peromyscus_keeni | Peromyscus_keeni | 0 | 21.3 | 5.36 | 212135.71 |
| mammals | Peromyscus_leucopus | Peromyscus_leucopus | 0 | 21.2 | 4.31 | 5831351.77 |
| mammals | Peromyscus_levipes | Peromyscus_levipes | 0 | 23 | 3.18 | 160994.64 |
| mammals | Peromyscus_madrensis | Peromyscus_madrensis | 1 | 57 | 3.04 | 415.34 |
| mammals | Peromyscus_maniculatus | Peromyscus_maniculatus | 0 | 21.3 | 4.69 | 13030881.64 |
| mammals | Peromyscus_mayensis | Peromyscus_mayensis | 1 | 32 | 1.94 | 89.08 |

| | | | | | | |
|---|---|---|---|---|---|---|
| mammals | Peromyscus_megalops | Peromyscus_megalops | 0 | 66.2 | 1.95 | 44159.06 |
| mammals | Peromyscus_mekisturus | Peromyscus_mekisturus | 1 | 60 | 2.93 | 7008.65 |
| mammals | Peromyscus_melanocarpus | Peromyscus_melanocarpus | 1 | 59 | 2.09 | 3955.83 |
| mammals | Peromyscus_melanophrys | Peromyscus_melanophrys | 0 | 40 | 2.92 | 512462.17 |
| mammals | Peromyscus_melanotis | Peromyscus_melanotis | 0 | 39.6 | 3.91 | 494800.92 |
| mammals | Peromyscus_melanurus | Peromyscus_melanurus | 1 | 40 | 2.38 | 3783.27 |
| mammals | Peromyscus_merriami | Peromyscus_merriami | 0 | 40 | 3.01 | 197642.76 |
| mammals | Peromyscus_mexicanus | Peromyscus_mexicanus | 0 | 32.6 | 2.49 | 367383.47 |
| mammals | Peromyscus_nasutus | Peromyscus_nasutus | 0 | 28 | 3.69 | 576501.02 |
| mammals | Peromyscus_ochraventer | Peromyscus_ochraventer | 1 | 40 | 4.13 | 4659.36 |
| mammals | Peromyscus_pectoralis | Peromyscus_pectoralis | 0 | 39 | 3.68 | 832035.55 |
| mammals | Peromyscus_perfulvus | Peromyscus_perfulvus | 0 | 40 | 2.69 | 66536.1 |
| mammals | Peromyscus_polionotus | Peromyscus_polionotus | 0 | 13 | 3.6 | 376320.25 |
| mammals | Peromyscus_polius | Peromyscus_polius | 0 | 40 | 3.18 | 14189.42 |
| mammals | Peromyscus_sejugis | Peromyscus_sejugis | 1 | 21.5 | 4.62 | 13.59 |
| mammals | Peromyscus_simulus | Peromyscus_simulus | 1 | 40 | 3.18 | 11840.65 |
| mammals | Peromyscus_slevini | Peromyscus_slevini | 1 | 53.6 | 4.62 | 39.51 |
| mammals | Peromyscus_spicilegus | Peromyscus_spicilegus | 0 | 36 | 3.19 | 125505.29 |
| mammals | Peromyscus_stephani | Peromyscus_stephani | 1 | 26.1 | 3.18 | 41.2 |
| mammals | Peromyscus_stirtoni | Peromyscus_stirtoni | 0 | 29.2 | 2.38 | 26277.37 |
| mammals | Peromyscus_truei | Peromyscus_truei | 0 | 27.4 | 3.49 | 1452308.01 |
| mammals | Peromyscus_winkelmanni | Peromyscus_winkelmanni | 1 | 53.8 | 3.19 | 301.03 |
| mammals | Peromyscus_yucatanicus | Peromyscus_yucatanicus | 0 | 26.3 | 3 | 137965.81 |
| mammals | Peromyscus_zarhynchus | Peromyscus_zarhynchus | 1 | 40 | 2.1 | 10773.67 |
| mammals | Peropteryx_kappleri | Peropteryx_kappleri | 0 | 9.4 | 1 | 7287612.73 |
| mammals | Peropteryx_leucoptera | Peropteryx_leucoptera | 0 | 6 | 1 | 5089787.57 |
| mammals | Peropteryx_macrotis | Peropteryx_macrotis | 0 | 5.4 | 1 | 12697128.89 |
| mammals | Peroryctes_broadbenti | Peroryctes_broadbenti | 1 | 4800 | 1.76 | 31530.63 |
| mammals | Peroryctes_raffrayanus | Peroryctes_raffrayana | 0 | 838.5 | 1.45 | 298144.48 |
| mammals | Petaurista_alborufus | Petaurista_alborufus | 0 | 4290 | 1.25 | 943217.74 |
| mammals | Petaurista_elegans | Petaurista_elegans | 0 | 997 | 1 | 2490770.45 |
| mammals | Petaurista_leucogenys | Petaurista_leucogenys | 0 | 1178 | 1.33 | 251379.48 |
| mammals | Petaurista_magnificus | Petaurista_magnificus | 0 | 1475.5 | 0.98 | 142043.46 |
| mammals | Petaurista_nobilis | Petaurista_nobilis | 0 | 2710 | 1 | 44806.38 |

| | | | | | | |
|---|---|---|---|---|---|---|
| mammals | Petaurista_petaurista | Petaurista_petaurista | 0 | 1660.1 | 1.04 | 3195973.78 |
| mammals | Petaurista_philippensis | Petaurista_philippensis | 0 | 1676 | 1.04 | 3398886.32 |
| mammals | Petaurista_xanthotis | Petaurista_xanthotis | 0 | 1900 | 1.33 | 730211.79 |
| mammals | Petauroides_volans | Petauroides_volans | 1 | 1300 | 1 | 633491.26 |
| mammals | Petaurus_abidi | Petaurus_abidi | 1 | 278 | 1.02 | 92.81 |
| mammals | Petaurus_australis | Petaurus_australis | 0 | 575 | 1.12 | 231711.03 |
| mammals | Petaurus_biacensis | Petaurus_biacensis | 0 | 88.4 | 1.68 | 2533.47 |
| mammals | Petaurus_breviceps | Petaurus_breviceps | 0 | 106.3 | 1.91 | 2761627.5 |
| mammals | Petaurus_gracilis | Petaurus_gracilis | 1 | 353 | 1.67 | 3309.37 |
| mammals | Petaurus_norfolcensis | Petaurus_norfolcensis | 0 | 230 | 1.6 | 788590.48 |
| mammals | Petinomys_fuscocapillus | Petinomys_fuscocapillus | 0 | 794.5 | 2 | 56586.73 |
| mammals | Petinomys_genibarbis | Petinomys_genibarbis | 1 | 110 | 1.05 | 844696.57 |
| mammals | Petinomys_setosus | Petinomys_setosus | 1 | 38.4 | 1.5 | 715360.47 |
| mammals | Petinomys_vordermanni | Petinomys_vordermanni | 1 | 40.4 | 2 | 114244.84 |
| mammals | Petrodromus_tetradactylus | Petrodromus_tetradactylus | 0 | 187.5 | 1.08 | 3147854.37 |
| mammals | Petrogale_assimilis | Petrogale_assimilis | 0 | 4500 | 1 | 208649.69 |
| mammals | Petrogale_brachyotis | Petrogale_brachyotis | 0 | 4050 | 1 | 221775.6 |
| mammals | Petrogale_burbidgei | Petrogale_burbidgei | 0 | 1258 | 1 | 17334.32 |
| mammals | Petrogale_coenensis | Petrogale_coenensis | 1 | 4375 | 1 | 10779.95 |
| mammals | Petrogale_concinna | Petrogale_concinna | 1 | 1400 | 1 | 82698.69 |
| mammals | Petrogale_godmani | Petrogale_godmani | 0 | 4750 | 1 | 36293.79 |
| mammals | Petrogale_herberti | Petrogale_herberti | 0 | 5150 | 1 | 281939.81 |
| mammals | Petrogale_inornata | Petrogale_inornata | 0 | 4600 | 1 | 85125.44 |
| mammals | Petrogale_lateralis | Petrogale_lateralis | 1 | 4625 | 1 | 113472.06 |
| mammals | Petrogale_mareeba | Petrogale_mareeba | 0 | 4150 | 1 | 19944.21 |
| mammals | Petrogale_penicillata | Petrogale_penicillata | 1 | 7100 | 1 | 101650.44 |
| mammals | Petrogale_persephone | Petrogale_persephone | 1 | 6200 | 1 | 4210.63 |
| mammals | Petrogale_purpureicollis | Petrogale_purpureicollis | 0 | 4500 | 1 | 80361.16 |
| mammals | Petrogale_sharmani | Petrogale_sharmani | 1 | 4250 | 1 | 16921.71 |
| mammals | Petrogale_xanthopus | Petrogale_xanthopus | 0 | 8500 | 1.04 | 24922.64 |
| mammals | Petromus_typicus | Petromus_typicus | 0 | 224 | 1.59 | 372746.37 |
| mammals | Petromyscus_barbouri | Petromyscus_barbouri | 0 | 20.1 | 2.51 | 38176.27 |
| mammals | Petromyscus_collinus | Petromyscus_collinus | 0 | 20.1 | 2.48 | 497480.98 |
| mammals | Petromyscus_shortridgei | Petromyscus_shortridgei | 0 | 46.5 | 3 | 123751.78 |

| | | | | | | |
|---|---|---|---|---|---|---|
| mammals | Petrosaltator_rozeti | Elephantulus_rozeti | 0 | 48 | 2.32 | 730293.64 |
| mammals | Phacochoerus_aethiopicus | Phacochoerus_aethiopicus | 0 | 71000.3 | 3.28 | 318268.66 |
| mammals | Phacochoerus_africanus | Phacochoerus_africanus | 0 | 82500 | 3 | 9957729.53 |
| mammals | Phalanger_carmelitae | Phalanger_carmelitae | 0 | 2047.5 | 1.01 | 198474.77 |
| mammals | Phalanger_gymnotis | Phalanger_gymnotis | 0 | 2705 | 1 | 616455.78 |
| mammals | Phalanger_intercastellanus | Phalanger_intercastellanus | 0 | 1747.5 | 2.01 | 63331.48 |
| mammals | Phalanger_lullulae | Phalanger_lullulae | 1 | 1632.5 | 1.75 | 886.3 |
| mammals | Phalanger_matanim | Phalanger_matanim | 1 | 1550 | 1.02 | 153.14 |
| mammals | Phalanger_mimicus | Phalanger_mimicus | 0 | 1850 | 2 | 233040.72 |
| mammals | Phalanger_orientalis | Phalanger_orientalis | 0 | 2168.8 | 1.85 | 456940.12 |
| mammals | Phalanger_ornatus | Phalanger_ornatus | 0 | 1787.5 | 2 | 22280.65 |
| mammals | Phalanger_pelengensis | Strigocuscus_pelengensis | 0 | 1103 | 1 | 6411.27 |
| mammals | Phalanger_rothschildi | Phalanger_rothschildi | 0 | 1365 | 1 | 2774.76 |
| mammals | Phalanger_sericeus | Phalanger_sericeus | 0 | 2003 | 1.01 | 197226.98 |
| mammals | Phalanger_vestitus | Phalanger_vestitus | 0 | 1850 | 1.01 | 88965.09 |
| mammals | Phaner_electromontis | Phaner_electromontis | 1 | 383.6 | 1 | 1663 |
| mammals | Phaner_furcifer | Phaner_furcifer | 1 | 400 | 1 | 20954.54 |
| mammals | Phaner_pallescens | Phaner_pallescens | 1 | 327 | 1 | 47678.92 |
| mammals | Phaner_parienti | Phaner_parienti | 1 | 363.4 | 1 | 1912.24 |
| mammals | Phascogale_calura | Phascogale_calura | 0 | 51.5 | 7.5 | 46849.83 |
| mammals | Phascogale_pirata | Phascogale_pirata | 1 | 181.5 | 6.69 | 40381.03 |
| mammals | Phascogale_tapoatafa | Phascogale_tapoatafa | 0 | 193.5 | 6.69 | 272010.8 |
| mammals | Phascolarctos_cinereus | Phascolarctos_cinereus | 1 | 10250 | 1.01 | 1102194.4 |
| mammals | Phascolosorex_dorsalis | Phascolosorex_dorsalis | 0 | 138.5 | 3.5 | 174794.95 |
| mammals | Phataginus_tetradactylus | Phataginus_tetradactyla | 1 | 2750 | 1.05 | 2929072.01 |
| mammals | Phataginus_tricuspis | Phataginus_tricuspis | 1 | 2066.7 | 1 | 5954123.2 |
| mammals | Phenacomys_intermedius | Phenacomys_intermedius | 0 | 25.2 | 4.64 | 524347.74 |
| mammals | Phenacomys_ungava | Phenacomys_ungava | 0 | 32.5 | 4.76 | 5082619.31 |
| mammals | Philander_andersoni | Philander_andersoni | 0 | 325 | 3.82 | 1346166.85 |
| mammals | Philander_mcilhennyi | Philander_mcilhennyi | 0 | 513.9 | 3.82 | 443217.68 |
| mammals | Philander_opossum | Philander_opossum | 0 | 585.5 | 4.68 | 5893336.84 |
| mammals | Philander_quica | Philander_frenatus | 0 | 167.7 | 4.68 | 921860.97 |
| mammals | Philantomba_maxwellii | Philantomba_maxwellii | 0 | 12000 | 1 | 872152.03 |
| mammals | Philantomba_monticola | Philantomba_monticola | 0 | 6250 | 0.99 | 6383795.31 |

| | | | | | | |
|---|---|---|---|---|---|---|
| mammals | Philetor_brachypterus | Philetor_brachypterus | 0 | 10.9 | 1 | 1292109.23 |
| mammals | Phloeomys_cumingi | Phloeomys_cumingi | 0 | 1946.3 | 1 | 19097.58 |
| mammals | Phloeomys_pallidus | Phloeomys_pallidus | 0 | 2250 | 1.1 | 43063.29 |
| mammals | Phodopus_campbelli | Phodopus_campbelli | 0 | 23.4 | 6.97 | 2049854.9 |
| mammals | Phodopus_roborovskii | Phodopus_roborovskii | 0 | 18.8 | 6.14 | 3141864.39 |
| mammals | Phodopus_sungorus | Phodopus_sungorus | 0 | 33.8 | 5.36 | 1181966.72 |
| mammals | Phoniscus_jagorii | Phoniscus_jagorii | 0 | 4.7 | 1 | 347840.56 |
| mammals | Phoniscus_papuensis | Phoniscus_papuensis | 1 | 6.2 | 1 | 474747.76 |
| mammals | Phylloderma_stenops | Phylloderma_stenops | 0 | 53.7 | 0.98 | 10972630.64 |
| mammals | Phyllomys_blainvilii | Phyllomys_blainvillii | 0 | 243.3 | 2.5 | 332344.1 |
| mammals | Phyllomys_brasiliensis | Phyllomys_brasiliensis | 1 | 298.6 | 1 | 2858.17 |
| mammals | Phyllomys_dasythrix | Phyllomys_dasythrix | 0 | 260 | 2 | 276267.04 |
| mammals | Phyllomys_medius | Phyllomys_medius | 0 | 229.9 | 2 | 501981.64 |
| mammals | Phyllomys_pattoni | Phyllomys_pattoni | 0 | 229.9 | 2 | 302917.22 |
| mammals | Phyllonycteris_aphylla | Phyllonycteris_aphylla | 1 | 24.5 | 0.99 | 215.19 |
| mammals | Phyllonycteris_poeyi | Phyllonycteris_poeyi | 0 | 22 | 0.98 | 182992.97 |
| mammals | Phyllops_falcatus | Phyllops_falcatus | 0 | 19.5 | 0.98 | 181362.69 |
| mammals | Phyllostomus_discolor | Phyllostomus_discolor | 0 | 34.1 | 1 | 12498386.82 |
| mammals | Phyllostomus_elongatus | Phyllostomus_elongatus | 0 | 40.4 | 0.99 | 10601026.64 |
| mammals | Phyllostomus_hastatus | Phyllostomus_hastatus | 0 | 87.2 | 1 | 12629829.36 |
| mammals | Phyllotis_amicus | Phyllotis_amicus | 0 | 20.2 | 2 | 71787.38 |
| mammals | Phyllotis_andium | Phyllotis_andium | 0 | 53 | 2 | 124063.11 |
| mammals | Phyllotis_bonariensis | Phyllotis_bonariensis | 0 | 42.5 | 4.02 | 27439.82 |
| mammals | Phyllotis_caprinus | Phyllotis_caprinus | 0 | 50.8 | 4.02 | 42361.2 |
| mammals | Phyllotis_darwinii | Phyllotis_darwini | 0 | 50.8 | 4.02 | 130478.74 |
| mammals | Phyllotis_definitus | Phyllotis_definitus | 1 | 89 | 4.02 | 20652.78 |
| mammals | Phyllotis_gerbillus | Phyllotis_gerbillus | 0 | 17.4 | 3 | 33728.37 |
| mammals | Phyllotis_limatus | Phyllotis_limatus | 0 | 36.9 | 4.02 | 137878.4 |
| mammals | Phyllotis_magister | Phyllotis_magister | 0 | 68.5 | 4.02 | 58908.14 |
| mammals | Phyllotis_osilae | Phyllotis_osilae | 0 | 49 | 4.25 | 309678.58 |
| mammals | Phyllotis_xanthopygus | Phyllotis_xanthopygus | 0 | 56.3 | 4.02 | 1499341.73 |
| mammals | Piliocolobus_badius | Piliocolobus_badius | 1 | 7750.1 | 1 | 342829.97 |
| mammals | Piliocolobus_pennantii | Piliocolobus_pennantii | 1 | 10896 | 1.01 | 161.41 |
| mammals | Piliocolobus_preussi | Piliocolobus_preussi | 1 | 9750 | 1.01 | 4263.46 |

| | | | | | | |
|---|---|---|---|---|---|---|
| mammals | Piliocolobus_rufomitratus | Piliocolobus_rufomitratus | 1 | 8000 | 1.01 | 201.25 |
| mammals | Pilonycteris_celebensis | Rousettus_celebensis | 0 | 56.3 | 1 | 181192.8 |
| mammals | Pipanacoctomys_aureus | Tympanoctomys_aureus | 1 | 139.4 | 1 | 16.89 |
| mammals | Pipistrellus_abramus | Pipistrellus_abramus | 0 | 5.6 | 2.15 | 3910979.37 |
| mammals | Pipistrellus_adamsi | Pipistrellus_adamsi | 0 | 4.5 | 1.92 | 277615.88 |
| mammals | Pipistrellus_angulatus | Pipistrellus_angulatus | 0 | 3.3 | 1.92 | 344529.75 |
| mammals | Pipistrellus_ceylonicus | Pipistrellus_ceylonicus | 0 | 8.4 | 1.99 | 2718309.29 |
| mammals | Pipistrellus_collinus | Pipistrellus_collinus | 0 | 6 | 1.92 | 201503.03 |
| mammals | Pipistrellus_coromandra | Pipistrellus_coromandra | 0 | 4.6 | 1.4 | 3655673.73 |
| mammals | Pipistrellus_endoi | Pipistrellus_endoi | 0 | 4.2 | 2 | 22538.23 |
| mammals | Pipistrellus_hanaki | Pipistrellus_hanaki | 1 | 6 | 2 | 48435.96 |
| mammals | Pipistrellus_hesperidus | Pipistrellus_hesperidus | 0 | 5.7 | 1.35 | 1600945.63 |
| mammals | Pipistrellus_javanicus | Pipistrellus_javanicus | 0 | 4.8 | 2.29 | 3666505.45 |
| mammals | Pipistrellus_kuhlii | Pipistrellus_kuhlii | 0 | 5.9 | 1.28 | 10905819.43 |
| mammals | Pipistrellus_nanulus | Pipistrellus_nanulus | 0 | 2.5 | 1.96 | 1565043.12 |
| mammals | Pipistrellus_nathusii | Pipistrellus_nathusii | 0 | 7.3 | 1.8 | 5890730.92 |
| mammals | Pipistrellus_papuanus | Pipistrellus_papuanus | 0 | 4.7 | 1.92 | 406795.99 |
| mammals | Pipistrellus_paterculus | Pipistrellus_paterculus | 0 | 5.3 | 2.19 | 782192.18 |
| mammals | Pipistrellus_pipistrellus | Pipistrellus_pipistrellus | 0 | 5.7 | 1.33 | 7619643.12 |
| mammals | Pipistrellus_pygmaeus | Pipistrellus_pygmaeus | 0 | 5.7 | 1.2 | 1960533.95 |
| mammals | Pipistrellus_rusticus | Pipistrellus_rusticus | 0 | 4.6 | 1.96 | 3459632.36 |
| mammals | Pipistrellus_stenopterus | Pipistrellus_stenopterus | 0 | 15.2 | 0.98 | 641151.19 |
| mammals | Pipistrellus_wattsi | Pipistrellus_wattsi | 0 | 3.3 | 1.92 | 14693.1 |
| mammals | Pipistrellus_westralis | Pipistrellus_westralis | 0 | 3.1 | 1.92 | 189808.93 |
| mammals | Pithecheir_melanurus | Pithecheir_melanurus | 1 | 108 | 1 | 132.57 |
| mammals | Pithecia_aequatorialis | Pithecia_aequatorialis | 0 | 2250 | 1 | 45738.29 |
| mammals | Pithecia_chrysocephala | Pithecia_chrysocephala | 0 | 1900 | 1.1 | 114364.76 |
| mammals | Pithecia_pithecia | Pithecia_pithecia | 0 | 1375.5 | 1.1 | 886392.05 |
| mammals | Plagiodontia_aedium | Plagiodontia_aedium | 1 | 2630 | 1.14 | 10576.6 |
| mammals | Planigale_gilesi | Planigale_gilesi | 0 | 9.2 | 5.23 | 909605.33 |
| mammals | Planigale_ingrami | Planigale_ingrami | 0 | 4.3 | 7.36 | 617152.39 |
| mammals | Planigale_maculata | Planigale_maculata | 0 | 11 | 7.64 | 1233506.99 |
| mammals | Planigale_tenuirostris | Planigale_tenuirostris | 0 | 6.1 | 5.99 | 1617101.25 |
| mammals | Platalina_genovensium | Platalina_genovensium | 0 | 47 | 0.98 | 300140.5 |

| | | | | | | |
|---|---|---|---|---|---|---|
| mammals | Platymops_setiger | Platymops_setiger | 0 | 13 | 1 | 229068.21 |
| mammals | Platyrrhinus_albericoi | Platyrrhinus_albericoi | 0 | 55 | 1 | 608162.99 |
| mammals | Platyrrhinus_brachycephalus | Platyrrhinus_brachycephalus | 0 | 14.1 | 0.98 | 6159441.37 |
| mammals | Platyrrhinus_dorsalis | Platyrrhinus_dorsalis | 0 | 26 | 0.99 | 767245.6 |
| mammals | Platyrrhinus_helleri | Platyrrhinus_helleri | 0 | 13.5 | 0.99 | 10016789.88 |
| mammals | Platyrrhinus_infuscus | Platyrrhinus_infuscus | 0 | 50.2 | 0.98 | 1466979.14 |
| mammals | Platyrrhinus_lineatus | Platyrrhinus_lineatus | 0 | 24.4 | 1 | 5807566.65 |
| mammals | Platyrrhinus_matapalensis | Platyrrhinus_matapalensis | 0 | 18 | 1 | 80808.01 |
| mammals | Platyrrhinus_vittatus | Platyrrhinus_vittatus | 0 | 41.4 | 0.98 | 769716.34 |
| mammals | Plecotus_auritus | Plecotus_auritus | 0 | 7.8 | 1.25 | 6103621.63 |
| mammals | Plecotus_austriacus | Plecotus_austriacus | 0 | 7.3 | 1.27 | 3363262.95 |
| mammals | Plecotus_kolombatovici | Plecotus_kolombatovici | 0 | 7 | 1.27 | 201013.87 |
| mammals | Plecotus_macrobullaris | Plecotus_macrobullaris | 0 | 8.5 | 1.27 | 1010971.42 |
| mammals | Plecotus_ognevi | Plecotus_ognevi | 0 | 8.4 | 1.25 | 5958715.33 |
| mammals | Plecotus_sacrimontis | Plecotus_sacrimontis | 0 | 9.3 | 1.25 | 206693.02 |
| mammals | Plecotus_teneriffae | Plecotus_teneriffae | 1 | 8 | 1.27 | 2808.5 |
| mammals | Plecturocebus_aureipalatii | Callicebus_aureipalatii | 0 | 950 | 1 | 58371.59 |
| mammals | Plecturocebus_baptista | Callicebus_baptista | 0 | 956.5 | 1.01 | 14740.99 |
| mammals | Plecturocebus_bernhardi | Callicebus_bernhardi | 0 | 950 | 1 | 119557.52 |
| mammals | Plecturocebus_brunneus | Callicebus_brunneus | 1 | 992.4 | 1.01 | 184934.53 |
| mammals | Plecturocebus_caligatus | Callicebus_caligatus | 0 | 992.4 | 1.02 | 195914.36 |
| mammals | Plecturocebus_cinerascens | Callicebus_cinerascens | 0 | 992.4 | 1.02 | 210384.5 |
| mammals | Plecturocebus_cupreus | Callicebus_cupreus | 0 | 915 | 1.01 | 776067.49 |
| mammals | Plecturocebus_discolor | Callicebus_discolor | 0 | 917.7 | 1 | 356789.18 |
| mammals | Plecturocebus_donacophilus | Callicebus_donacophilus | 0 | 795 | 1.02 | 163767.86 |
| mammals | Plecturocebus_hoffmannsi | Callicebus_hoffmannsi | 0 | 992.4 | 1.02 | 92127.78 |
| mammals | Plecturocebus_modestus | Callicebus_modestus | 1 | 992.4 | 1.02 | 6539.93 |
| mammals | Plecturocebus_moloch | Callicebus_moloch | 0 | 854.7 | 1.01 | 944033.5 |
| mammals | Plecturocebus_oenanthe | Callicebus_oenanthe | 1 | 992.4 | 1.02 | 9106.74 |
| mammals | Plecturocebus_olallae | Callicebus_olallae | 1 | 992.4 | 1.02 | 677.03 |
| mammals | Plecturocebus_ornatus | Callicebus_ornatus | 1 | 832.6 | 1 | 23835.55 |
| mammals | Plecturocebus_pallescens | Callicebus_pallescens | 0 | 832.1 | 1.02 | 323682.68 |
| mammals | Podogymnura_truei | Podogymnura_truei | 0 | 114 | 1.5 | 5291.51 |
| mammals | Podomys_floridanus | Podomys_floridanus | 0 | 30.8 | 2.6 | 47852.64 |

| | | | | | | |
|---|---|---|---|---|---|---|
| mammals | Poecilictis_libyca | Ictonyx_libycus | 0 | 2062.5 | 2 | 5082352.29 |
| mammals | Poecilogale_albinucha | Poecilogale_albinucha | 0 | 340 | 2.22 | 5663520.45 |
| mammals | Poelagus_marjorita | Poelagus_marjorita | 0 | 2500 | 1.27 | 123506.32 |
| mammals | Poemys_melanotis | Dendromus_melanotis | 0 | 9.1 | 4.26 | 4305594.49 |
| mammals | Poemys_nyikae | Dendromus_nyikae | 0 | 15 | 4.62 | 2375792.29 |
| mammals | Pogonomelomys_mayeri | Pogonomelomys_mayeri | 0 | 109.7 | 2.15 | 102264.11 |
| mammals | Pogonomys_fergussoniensis | Pogonomys_fergussoniensis | 1 | 128.5 | 2.17 | 2050.82 |
| mammals | Pogonomys_loriae | Pogonomys_loriae | 0 | 95.1 | 2.43 | 151035.63 |
| mammals | Pogonomys_macrourus | Pogonomys_macrourus | 0 | 54 | 2.6 | 317142.64 |
| mammals | Pogonomys_sylvestris | Pogonomys_sylvestris | 0 | 43.4 | 2.4 | 117161.34 |
| mammals | Poiana_leightoni | Poiana_leightoni | 1 | 600 | 2.23 | 49336.18 |
| mammals | Poiana_richardsonii | Poiana_richardsonii | 0 | 500 | 2.23 | 2017237.12 |
| mammals | Poliocitellus_franklinii | Poliocitellus_franklinii | 0 | 363 | 8.3 | 1657975.71 |
| mammals | Pongo_abelii | Pongo_abelii | 1 | 56750 | 1.14 | 33185.92 |
| mammals | Pongo_pygmaeus | Pongo_pygmaeus | 1 | 57150 | 1.1 | 249610.11 |
| mammals | Porcula_salvania | Porcula_salvania | 1 | 7718 | 3.43 | 881.68 |
| mammals | Potamochoerus_larvatus | Potamochoerus_larvatus | 0 | 97500 | 2.87 | 5881488.57 |
| mammals | Potamochoerus_porcus | Potamochoerus_porcus | 0 | 70000.3 | 3 | 4799557.05 |
| mammals | Potamogale_velox | Potamogale_velox | 0 | 660 | 2.28 | 3803856.98 |
| mammals | Potorous_longipes | Potorous_longipes | 1 | 1900 | 1 | 4229.45 |
| mammals | Potorous_tridactylus | Potorous_tridactylus | 0 | 1100 | 1 | 125038.54 |
| mammals | Potos_flavus | Potos_flavus | 0 | 3000 | 1.05 | 12691116.84 |
| mammals | Praomys_daltoni | Praomys_daltoni | 0 | 38.5 | 5.49 | 3198976.11 |
| mammals | Praomys_degraaffi | Praomys_degraaffi | 0 | 45.5 | 3 | 12854.58 |
| mammals | Praomys_hartwigi | Praomys_hartwigi | 1 | 57.2 | 3.49 | 7809.65 |
| mammals | Praomys_jacksoni | Praomys_jacksoni | 0 | 42 | 3.58 | 3744052.13 |
| mammals | Praomys_minor | Praomys_minor | 0 | 42.8 | 3.5 | 8768.52 |
| mammals | Praomys_misonnei | Praomys_misonnei | 0 | 39.1 | 4 | 144160.07 |
| mammals | Praomys_morio | Praomys_morio | 1 | 30 | 3.29 | 1343.91 |
| mammals | Praomys_mutoni | Praomys_mutoni | 0 | 52 | 2.5 | 14876.31 |
| mammals | Praomys_obscurus | Praomys_obscurus | 1 | 47.2 | 3.49 | 3123.23 |
| mammals | Praomys_rostratus | Praomys_rostratus | 0 | 52 | 3.5 | 612377.04 |
| mammals | Praomys_tullbergi | Praomys_tullbergi | 0 | 35.8 | 3.43 | 932845.13 |
| mammals | Presbytis_chrysomelas | Presbytis_chrysomelas | 1 | 7000 | 1 | 39860.91 |

| | | | | | | |
|---|---|---|---|---|---|---|
| mammals | Presbytis_comata | Presbytis_comata | 1 | 6250 | 1.01 | 41162.76 |
| mammals | Presbytis_femoralis | Presbytis_femoralis | 1 | 7000 | 1 | 11279.71 |
| mammals | Presbytis_melalophos | Presbytis_melalophos | 1 | 6300 | 1 | 94639.09 |
| mammals | Presbytis_natunae | Presbytis_natunae | 1 | 4920 | 1 | 1643.7 |
| mammals | Presbytis_potenziani | Presbytis_potenziani | 1 | 6450 | 1.01 | 2140.85 |
| mammals | Presbytis_rubicunda | Presbytis_rubicunda | 1 | 6300 | 1.01 | 503494.9 |
| mammals | Presbytis_siamensis | Presbytis_siamensis | 0 | 7000 | 1 | 160647.49 |
| mammals | Presbytis_thomasi | Presbytis_thomasi | 1 | 6690 | 1 | 60508.92 |
| mammals | Priodontes_maximus | Priodontes_maximus | 1 | 45359.7 | 1.5 | 9614613.56 |
| mammals | Prionailurus_bengalensis | Prionailurus_bengalensis | 0 | 3300 | 2.5 | 8627600.17 |
| mammals | Prionailurus_planiceps | Prionailurus_planiceps | 1 | 6750 | 1.5 | 148872.29 |
| mammals | Prionailurus_rubiginosus | Prionailurus_rubiginosus | 0 | 1384.4 | 2.23 | 2416123.72 |
| mammals | Prionailurus_viverrinus | Prionailurus_viverrinus | 1 | 9140.3 | 2 | 235697.12 |
| mammals | Prionodon_linsang | Prionodon_linsang | 0 | 700 | 2.22 | 1447867.95 |
| mammals | Prionodon_pardicolor | Prionodon_pardicolor | 0 | 512 | 2 | 2342407.95 |
| mammals | Procapra_gutturosa | Procapra_gutturosa | 0 | 27750 | 1.37 | 715708.2 |
| mammals | Procapra_picticaudata | Procapra_picticaudata | 0 | 20000 | 1 | 2394537.77 |
| mammals | Procapra_przewalskii | Procapra_przewalskii | 1 | 27500 | 1 | 1076.3 |
| mammals | Procavia_capensis | Procavia_capensis | 0 | 3030 | 2.47 | 15715542.14 |
| mammals | Procolobus_verus | Procolobus_verus | 1 | 4000 | 1.01 | 412576.7 |
| mammals | Procyon_cancrivorus | Procyon_cancrivorus | 0 | 6949.9 | 2.68 | 12754113.76 |
| mammals | Procyon_lotor | Procyon_lotor | 0 | 6550 | 3.22 | 11684834.86 |
| mammals | Proechimys_brevicauda | Proechimys_brevicauda | 0 | 285 | 2.4 | 1460844.71 |
| mammals | Proechimys_decumanus | Proechimys_decumanus | 0 | 285 | 2.4 | 65536.94 |
| mammals | Proechimys_echinothrix | Proechimys_echinothrix | 0 | 309.6 | 2 | 277424.04 |
| mammals | Proechimys_goeldii | Proechimys_goeldii | 0 | 285 | 2.4 | 1029773.28 |
| mammals | Proechimys_guairae | Proechimys_guairae | 0 | 400 | 2.79 | 329800.19 |
| mammals | Proechimys_guyannensis | Proechimys_guyannensis | 0 | 316 | 2.22 | 1497170.03 |
| mammals | Proechimys_longicaudatus | Proechimys_longicaudatus | 0 | 205 | 2.4 | 1349084.8 |
| mammals | Proechimys_pattoni | Proechimys_pattoni | 0 | 178.2 | 2.4 | 157595.73 |
| mammals | Proechimys_quadruplicatus | Proechimys_quadruplicatus | 0 | 285 | 2.4 | 1498656.18 |
| mammals | Proechimys_roberti | Proechimys_roberti | 0 | 285 | 2.4 | 1703345.3 |
| mammals | Proechimys_semispinosus | Proechimys_semispinosus | 0 | 285 | 2.69 | 348118.05 |
| mammals | Proechimys_simonsi | Proechimys_simonsi | 0 | 285 | 2.4 | 1935012.33 |

| | | | | | | |
|---|---|---|---|---|---|---|
| mammals | Proechimys_steerei | Proechimys_steerei | 0 | 323 | 2.4 | 1677776.52 |
| mammals | Prolemur_simus | Prolemur_simus | 1 | 2350 | 1 | 1709.04 |
| mammals | Promops_centralis | Promops_centralis | 0 | 26.4 | 0.99 | 4729320.4 |
| mammals | Promops_nasutus | Promops_nasutus | 0 | 15.2 | 1 | 8029235.89 |
| mammals | Pronolagus_crassicaudatus | Pronolagus_crassicaudatus | 0 | 2089 | 1.5 | 198444.31 |
| mammals | Pronolagus_randensis | Pronolagus_randensis | 0 | 2335 | 1.18 | 675441.13 |
| mammals | Pronolagus_rupestris | Pronolagus_rupestris | 0 | 1754 | 1.88 | 461340.91 |
| mammals | Propithecus_candidus | Propithecus_candidus | 1 | 5750 | 1.01 | 1360.66 |
| mammals | Propithecus_coquereli | Propithecus_coquereli | 1 | 3900 | 1 | 7389.96 |
| mammals | Propithecus_coronatus | Propithecus_coronatus | 1 | 3900 | 1.02 | 45223.88 |
| mammals | Propithecus_deckenii | Propithecus_deckenii | 1 | 4000 | 1.02 | 25611.86 |
| mammals | Propithecus_diadema | Propithecus_diadema | 1 | 5250 | 1.01 | 16421.24 |
| mammals | Propithecus_edwardsi | Propithecus_edwardsi | 1 | 5750 | 1.01 | 908.7 |
| mammals | Propithecus_perrieri | Propithecus_perrieri | 1 | 4350 | 1.01 | 62.3 |
| mammals | Propithecus_tattersalli | Propithecus_tattersalli | 1 | 3500 | 1 | 1772.87 |
| mammals | Propithecus_verreauxi | Propithecus_verreauxi | 1 | 3250 | 1.02 | 30758.38 |
| mammals | Prosciurillus_murinus | Prosciurillus_murinus | 0 | 72.9 | 1 | 67498.16 |
| mammals | Proteles_cristatus | Proteles_cristata | 0 | 10000 | 2.64 | 5997315.15 |
| mammals | Protoxerus_stangeri | Protoxerus_stangeri | 0 | 621.5 | 1.48 | 3388918.95 |
| mammals | Psammomys_obesus | Psammomys_obesus | 0 | 369.6 | 3.5 | 2313613.02 |
| mammals | Pseudantechinus_bilarni | Pseudantechinus_bilarni | 0 | 27.5 | 4.75 | 134277.93 |
| mammals | Pseudantechinus_macdonnellensis | Pseudantechinus_macdonnellensis | 0 | 32.5 | 5.91 | 1267010.73 |
| mammals | Pseudantechinus_mimulus | Pseudantechinus_mimulus | 0 | 17 | 5.94 | 3092.03 |
| mammals | Pseudantechinus_ningbing | Pseudantechinus_ningbing | 0 | 20 | 4 | 208494.5 |
| mammals | Pseudantechinus_woolleyae | Pseudantechinus_woolleyae | 0 | 40 | 5.99 | 645661.16 |
| mammals | Pseudocheirus_occidentalis | Pseudocheirus_occidentalis | 1 | 987.5 | 1.1 | 18007.81 |
| mammals | Pseudocheirus_peregrinus | Pseudocheirus_peregrinus | 0 | 900 | 1.92 | 916198.63 |
| mammals | Pseudochirops_albertisii | Pseudochirops_albertisii | 0 | 784.9 | 1.01 | 25738.52 |
| mammals | Pseudochirops_archeri | Pseudochirops_archeri | 0 | 1091.5 | 1.02 | 5210.84 |
| mammals | Pseudochirops_corinnae | Pseudochirops_corinnae | 0 | 1123 | 1.01 | 185300.98 |
| mammals | Pseudochirops_coronatus | Pseudochirops_coronatus | 1 | 1500 | 1.01 | 7454.75 |
| mammals | Pseudochirops_cupreus | Pseudochirops_cupreus | 0 | 1724.5 | 1.01 | 197734.76 |
| mammals | Pseudochirops_dahlii | Petropseudes_dahli | 0 | 1640 | 1 | 198708.38 |
| mammals | Pseudochirulus_canescens | Pseudochirulus_canescens | 0 | 300 | 2.43 | 459129.8 |

| | | | | | | |
|---|---|---|---|---|---|---|
| mammals | Pseudochirulus_cinereus | Pseudochirulus_cinereus | 0 | 1000 | 2 | 895.81 |
| mammals | Pseudochirulus_forbesi | Pseudochirulus_forbesi | 0 | 649 | 1.26 | 47168.03 |
| mammals | Pseudochirulus_herbertensis | Pseudochirulus_herbertensis | 0 | 1110.5 | 1.92 | 3432.76 |
| mammals | Pseudochirulus_mayeri | Pseudochirulus_mayeri | 0 | 151.8 | 1.26 | 84923.68 |
| mammals | Pseudochirulus_schlegelii | Pseudochirulus_schlegeli | 1 | 274.5 | 1.26 | 10303.01 |
| mammals | Pseudois_nayaur | Pseudois_nayaur | 0 | 45000 | 1.36 | 2858794.56 |
| mammals | Pseudomys_albocinereus | Pseudomys_albocinereus | 0 | 30.5 | 3.98 | 221496.58 |
| mammals | Pseudomys_apodemoides | Pseudomys_apodemoides | 0 | 20 | 4.8 | 35937.02 |
| mammals | Pseudomys_australis | Pseudomys_australis | 1 | 65 | 3.05 | 199918.69 |
| mammals | Pseudomys_bolami | Pseudomys_bolami | 0 | 15.5 | 2.38 | 746188.58 |
| mammals | Pseudomys_chapmani | Pseudomys_chapmani | 0 | 12 | 4 | 299581.3 |
| mammals | Pseudomys_delicatulus | Pseudomys_delicatulus | 0 | 11 | 3.14 | 1842508.2 |
| mammals | Pseudomys_desertor | Pseudomys_desertor | 0 | 25 | 3 | 5158688.13 |
| mammals | Pseudomys_fumeus | Pseudomys_fumeus | 1 | 70 | 3.5 | 16696.31 |
| mammals | Pseudomys_gracilicaudatus | Pseudomys_gracilicaudatus | 0 | 74.8 | 2.93 | 337741.33 |
| mammals | Pseudomys_hermannsburgensis | Pseudomys_hermannsburgensis | 0 | 12 | 2.19 | 3846942.82 |
| mammals | Pseudomys_higginsi | Pseudomys_higginsi | 0 | 67 | 2.77 | 64548.42 |
| mammals | Pseudomys_nanus | Pseudomys_nanus | 0 | 34 | 3 | 1006949.82 |
| mammals | Pseudomys_novaehollandiae | Pseudomys_novaehollandiae | 1 | 20 | 4.24 | 89948.95 |
| mammals | Pseudomys_occidentalis | Pseudomys_occidentalis | 0 | 34 | 3 | 51491.41 |
| mammals | Pseudomys_oralis | Pseudomys_oralis | 1 | 95 | 3.05 | 26516.22 |
| mammals | Pseudomys_shortridgei | Pseudomys_shortridgei | 0 | 70 | 3 | 32075.84 |
| mammals | Pseudoromicia_brunnea | Neoromicia_brunnea | 0 | 9.9 | 1 | 757189.02 |
| mammals | Pseudoromicia_rendalli | Neoromicia_rendalli | 0 | 6.3 | 2 | 8376297.1 |
| mammals | Pseudoromicia_tenuipinnis | Neoromicia_tenuipinnis | 0 | 5.3 | 1 | 4247435.61 |
| mammals | Pseudoryx_nghetinhensis | Pseudoryx_nghetinhensis | 1 | 85000 | 1 | 13994.99 |
| mammals | Pseudosoriculus_fumidus | Episoriculus_fumidus | 0 | 6.7 | 3.42 | 12808.65 |
| mammals | Ptenochirus_jagorii | Ptenochirus_jagori | 0 | 77.3 | 1 | 271017.14 |
| mammals | Ptenochirus_minor | Ptenochirus_minor | 0 | 36.3 | 1 | 118793.76 |
| mammals | Pteralopex_anceps | Pteralopex_anceps | 1 | 560.4 | 1 | 8736.5 |
| mammals | Pteralopex_taki | Pteralopex_taki | 1 | 286.7 | 1 | 3316.8 |
| mammals | Pteromys_momonga | Pteromys_momonga | 0 | 151.8 | 3 | 168642.96 |
| mammals | Pteromys_volans | Pteromys_volans | 0 | 143.8 | 2.55 | 13737442.37 |
| mammals | Pteromyscus_pulverulentus | Pteromyscus_pulverulentus | 1 | 334.3 | 1.28 | 393205.98 |

| | | | | | | |
|---|---|---|---|---|---|---|
| mammals | Pteronotus_davyi | Pteronotus_davyi | 0 | 9.2 | 0.99 | 3461633.97 |
| mammals | Pteronotus_gymnonotus | Pteronotus_gymnonotus | 0 | 14.4 | 0.98 | 3202565.12 |
| mammals | Pteronotus_macleayii | Pteronotus_macleayii | 0 | 5.4 | 1 | 119896.67 |
| mammals | Pteronotus_paraguanensis | Pteronotus_paraguanensis | 1 | 13.9 | 0.99 | 142.81 |
| mammals | Pteronotus_parnellii | Pteronotus_parnellii | 0 | 19.7 | 0.99 | 200546.18 |
| mammals | Pteronotus_personatus | Pteronotus_personatus | 0 | 8.7 | 0.98 | 4748166.22 |
| mammals | Pteronotus_quadridens | Pteronotus_quadridens | 0 | 5.2 | 1.21 | 200546.18 |
| mammals | Pteronura_brasiliensis | Pteronura_brasiliensis | 1 | 23999.9 | 2.03 | 6631572.76 |
| mammals | Pteropus_admiralitatum | Pteropus_admiralitatum | 0 | 299.9 | 1 | 75834.56 |
| mammals | Pteropus_aldabrensis | Pteropus_aldabrensis | 1 | 305 | 1 | 152.67 |
| mammals | Pteropus_alecto | Pteropus_alecto | 0 | 643.6 | 1.05 | 1427125.68 |
| mammals | Pteropus_anetianus | Pteropus_anetianus | 1 | 426 | 0.98 | 11890.49 |
| mammals | Pteropus_capistratus | Pteropus_capistratus | 1 | 223 | 1 | 36299.77 |
| mammals | Pteropus_cognatus | Pteropus_cognatus | 1 | 302.5 | 0.99 | 3106.4 |
| mammals | Pteropus_conspicillatus | Pteropus_conspicillatus | 1 | 775.7 | 1 | 219525.84 |
| mammals | Pteropus_dasymallus | Pteropus_dasymallus | 1 | 435 | 1 | 2890.67 |
| mammals | Pteropus_faunulus | Pteropus_faunulus | 1 | 288 | 1 | 602.97 |
| mammals | Pteropus_hypomelanus | Pteropus_hypomelanus | 0 | 428.1 | 1 | 524209.43 |
| mammals | Pteropus_livingstonii | Pteropus_livingstonii | 1 | 588.5 | 1 | 265.68 |
| mammals | Pteropus_lylei | Pteropus_lylei | 1 | 316.7 | 1 | 106826.97 |
| mammals | Pteropus_macrotis | Pteropus_macrotis | 0 | 357 | 1 | 619521.29 |
| mammals | Pteropus_mariannus | Pteropus_mariannus | 1 | 453.5 | 1 | 2221.34 |
| mammals | Pteropus_melanotus | Pteropus_melanotus | 1 | 375 | 1 | 399.67 |
| mammals | Pteropus_molossinus | Pteropus_molossinus | 1 | 178 | 1 | 352.29 |
| mammals | Pteropus_neohibernicus | Pteropus_neohibernicus | 0 | 966.8 | 1 | 672585.2 |
| mammals | Pteropus_niger | Pteropus_niger | 1 | 456.3 | 1 | 4413.64 |
| mammals | Pteropus_ocularis | Pteropus_ocularis | 1 | 220 | 1 | 26749.18 |
| mammals | Pteropus_ornatus | Pteropus_ornatus | 1 | 330 | 0.98 | 17983.23 |
| mammals | Pteropus_poliocephalus | Pteropus_poliocephalus | 1 | 749.9 | 1 | 454359.02 |
| mammals | Pteropus_pselaphon | Pteropus_pselaphon | 1 | 397.2 | 0.98 | 71.81 |
| mammals | Pteropus_pumilus | Pteropus_pumilus | 0 | 183.4 | 1 | 57993.96 |
| mammals | Pteropus_rayneri | Pteropus_rayneri | 0 | 724.5 | 0.99 | 30622.52 |
| mammals | Pteropus_rennelli | Pteropus_rennelli | 1 | 296.4 | 0.99 | 544.1 |
| mammals | Pteropus_rodricensis | Pteropus_rodricensis | 1 | 253.8 | 1 | 110.93 |

| | | | | | | |
|---|---|---|---|---|---|---|
| mammals | Pteropus_rufus | Pteropus_rufus | 1 | 361 | 1 | 185368.31 |
| mammals | Pteropus_samoensis | Pteropus_samoensis | 0 | 310 | 1 | 20144.58 |
| mammals | Pteropus_scapulatus | Pteropus_scapulatus | 0 | 447.5 | 1 | 3033575.2 |
| mammals | Pteropus_seychellensis | Pteropus_seychellensis | 0 | 457 | 1 | 2730.91 |
| mammals | Pteropus_tonganus | Pteropus_tonganus | 0 | 418.3 | 1 | 57838.6 |
| mammals | Pteropus_vampyrus | Pteropus_vampyrus | 1 | 935.9 | 1 | 2043987.35 |
| mammals | Pteropus_voeltzkowi | Pteropus_voeltzkowi | 1 | 527.5 | 1 | 908.53 |
| mammals | Pteropus_woodfordi | Pteropus_woodfordi | 0 | 119.3 | 1 | 13702.96 |
| mammals | Ptilocercus_lowii | Ptilocercus_lowii | 0 | 41.9 | 2 | 356708.96 |
| mammals | Pudu_puda | Pudu_puda | 0 | 9749.9 | 1.12 | 166426.17 |
| mammals | Puma_concolor | Puma_concolor | 0 | 51600 | 2.76 | 22659138 |
| mammals | Punomys_kofordi | Punomys_kofordi | 0 | 82 | 2.5 | 1443.91 |
| mammals | Punomys_lemminus | Punomys_lemminus | 1 | 84.8 | 1.94 | 47849.82 |
| mammals | Pygathrix_cinerea | Pygathrix_cinerea | 1 | 8310 | 1 | 24681.13 |
| mammals | Pygathrix_nemaeus | Pygathrix_nemaeus | 1 | 8310 | 1 | 91649.51 |
| mammals | Pygathrix_nigripes | Pygathrix_nigripes | 1 | 8310 | 1 | 95056.8 |
| mammals | Pygeretmus_platyurus | Pygeretmus_platyurus | 0 | 26.4 | 5.38 | 926949.31 |
| mammals | Pygeretmus_pumilio | Pygeretmus_pumilio | 0 | 52 | 3.26 | 3430883.5 |
| mammals | Pygeretmus_shitkovi | Pygeretmus_zhitkovi | 0 | 56.5 | 5.49 | 48808.72 |
| mammals | Pygoderma_bilabiatum | Pygoderma_bilabiatum | 0 | 18.5 | 0.98 | 2957165.72 |
| mammals | Rangifer_tarandus | Rangifer_tarandus | 1 | 86034 | 1.55 | 18784282.67 |
| mammals | Raphicerus_campestris | Raphicerus_campestris | 0 | 10500 | 0.99 | 4047877.07 |
| mammals | Raphicerus_melanotis | Raphicerus_melanotis | 0 | 10249.9 | 1.8 | 160973.79 |
| mammals | Raphicerus_sharpei | Raphicerus_sharpei | 0 | 9499.9 | 1 | 2462567.95 |
| mammals | Rattus_andamanensis | Rattus_andamanensis | 0 | 129 | 8.31 | 1747879.16 |
| mammals | Rattus_argentiventer | Rattus_argentiventer | 0 | 133.1 | 8.39 | 2183638.25 |
| mammals | Rattus_colletti | Rattus_colletti | 0 | 61 | 6 | 86055.59 |
| mammals | Rattus_everetti | Rattus_everetti | 0 | 255 | 3.7 | 241325.77 |
| mammals | Rattus_exulans | Rattus_exulans | 0 | 47.3 | 3.68 | 4966659.44 |
| mammals | Rattus_feliceus | Rattus_feliceus | 0 | 306 | 3.48 | 17400.28 |
| mammals | Rattus_fuscipes | Rattus_fuscipes | 0 | 133 | 4.58 | 486689.53 |
| mammals | Rattus_hoffmanni | Rattus_hoffmanni | 0 | 147 | 4.5 | 168758.24 |
| mammals | Rattus_jobiensis | Rattus_jobiensis | 0 | 380 | 3.48 | 5155.67 |
| mammals | Rattus_leucopus | Rattus_leucopus | 0 | 171.7 | 3.46 | 365095.78 |

| | | | | | | |
|---|---|---|---|---|---|---|
| mammals | Rattus_losea | Rattus_losea | 0 | 77 | 7.3 | 2038643.18 |
| mammals | Rattus_lugens | Rattus_lugens | 1 | 224 | 5.95 | 6004.26 |
| mammals | Rattus_lutreola | Rattus_lutreolus | 0 | 106 | 4.68 | 260763.05 |
| mammals | Rattus_mindorensis | Rattus_mindorensis | 0 | 136 | 6.4 | 5016.04 |
| mammals | Rattus_mordax | Rattus_mordax | 0 | 230 | 2.96 | 51171.65 |
| mammals | Rattus_niobe | Rattus_niobe | 0 | 46.1 | 2.15 | 110059.91 |
| mammals | Rattus_nitidus | Rattus_nitidus | 0 | 133.5 | 3.09 | 3665292.81 |
| mammals | Rattus_norvegicus | Rattus_norvegicus | 0 | 356.7 | 8.92 | 19191093.61 |
| mammals | Rattus_novaeguineae | Rattus_novaeguineae | 0 | 133 | 5.37 | 34947.82 |
| mammals | Rattus_praetor | Rattus_praetor | 0 | 204.1 | 3.94 | 310739.95 |
| mammals | Rattus_pyctoris | Rattus_pyctoris | 0 | 136 | 4.23 | 1957430.21 |
| mammals | Rattus_rattus | Rattus_rattus | 0 | 178.3 | 5.92 | 28316841.32 |
| mammals | Rattus_richardsoni | Rattus_richardsoni | 1 | 64.3 | 2.15 | 6974.6 |
| mammals | Rattus_satarae | Rattus_satarae | 1 | 180 | 5.95 | 3126.12 |
| mammals | Rattus_sordidus | Rattus_sordidus | 0 | 157.5 | 5.99 | 258376.8 |
| mammals | Rattus_steini | Rattus_steini | 0 | 155 | 3.61 | 234808.04 |
| mammals | Rattus_tanezumi | Rattus_tanezumi | 0 | 134 | 5.95 | 9717429.76 |
| mammals | Rattus_tiomanicus | Rattus_tiomanicus | 0 | 130 | 4.88 | 1495112.48 |
| mammals | Rattus_tunneyi | Rattus_tunneyi | 0 | 124 | 5.44 | 762536.61 |
| mammals | Rattus_vandeuseni | Rattus_vandeuseni | 1 | 67.3 | 2.15 | 2014.71 |
| mammals | Rattus_verecundus | Rattus_verecundus | 0 | 108.5 | 2.15 | 152812.93 |
| mammals | Rattus_villosissimus | Rattus_villosissimus | 0 | 134 | 6 | 2155750.14 |
| mammals | Ratufa_bicolor | Ratufa_bicolor | 0 | 2050 | 1.57 | 2557024.66 |
| mammals | Ratufa_indica | Ratufa_indica | 0 | 1060 | 1 | 355292.04 |
| mammals | Ratufa_macroura | Ratufa_macroura | 0 | 1327.1 | 3.41 | 29502.65 |
| mammals | Redunca_arundinum | Redunca_arundinum | 0 | 58000.3 | 0.99 | 4764551.68 |
| mammals | Redunca_fulvorufula | Redunca_fulvorufula | 1 | 29499.9 | 1 | 1044828.85 |
| mammals | Redunca_redunca | Redunca_redunca | 0 | 44050.4 | 1 | 5901393.91 |
| mammals | Reithrodon_auritus | Reithrodon_auritus | 0 | 70.9 | 3.96 | 1191439.99 |
| mammals | Reithrodon_typicus | Reithrodon_typicus | 0 | 65.8 | 4.52 | 309299.71 |
| mammals | Reithrodontomys_brevirostris | Reithrodontomys_brevirostris | 0 | 12.9 | 3.4 | 20841.43 |
| mammals | Reithrodontomys_chrysopsis | Reithrodontomys_chrysopsis | 0 | 19 | 3.4 | 53802.41 |
| mammals | Reithrodontomys_creper | Reithrodontomys_creper | 0 | 22.8 | 2.21 | 12286.38 |
| mammals | Reithrodontomys_darienensis | Reithrodontomys_darienensis | 0 | 13 | 3.88 | 43587.41 |

| | | | | | | |
|---|---|---|---|---|---|---|
| mammals | Reithrodontomys_fulvescens | Reithrodontomys_fulvescens | 0 | 11.4 | 3.43 | 2632629.57 |
| mammals | Reithrodontomys_gracilis | Reithrodontomys_gracilis | 0 | 12.3 | 3.06 | 325050.23 |
| mammals | Reithrodontomys_hirsutus | Reithrodontomys_hirsutus | 1 | 20 | 3.88 | 9699.53 |
| mammals | Reithrodontomys_humulis | Reithrodontomys_humulis | 0 | 8.3 | 2.97 | 1296370.65 |
| mammals | Reithrodontomys_megalotis | Reithrodontomys_megalotis | 0 | 9.4 | 3.96 | 5116788.76 |
| mammals | Reithrodontomys_mexicanus | Reithrodontomys_mexicanus | 0 | 16 | 2.91 | 540022.72 |
| mammals | Reithrodontomys_montanus | Reithrodontomys_montanus | 0 | 10.9 | 3.99 | 1846897.4 |
| mammals | Reithrodontomys_raviventris | Reithrodontomys_raviventris | 1 | 11.1 | 3.87 | 971.42 |
| mammals | Reithrodontomys_sumichrasti | Reithrodontomys_sumichrasti | 0 | 19 | 3.49 | 328552.49 |
| mammals | Reithrodontomys_zacatecae | Reithrodontomys_zacatecae | 0 | 9.7 | 3.75 | 211192.08 |
| mammals | Rhabdomys_pumilio | Rhabdomys_pumilio | 0 | 41.1 | 5.64 | 134988.05 |
| mammals | Rheomys_mexicanus | Rheomys_mexicanus | 1 | 40 | 2 | 4237.41 |
| mammals | Rheomys_raptor | Rheomys_raptor | 0 | 38 | 0.97 | 12331.44 |
| mammals | Rheomys_thomasi | Rheomys_thomasi | 0 | 40 | 0.97 | 27299.26 |
| mammals | Rhinoceros_sondaicus | Rhinoceros_sondaicus | 1 | 1750000 | 1 | 125954.91 |
| mammals | Rhinoceros_unicornis | Rhinoceros_unicornis | 1 | 1602333 | 1.21 | 551653.91 |
| mammals | Rhinolophus_acuminatus | Rhinolophus_acuminatus | 0 | 12.1 | 1 | 1135409.32 |
| mammals | Rhinolophus_affinis | Rhinolophus_affinis | 0 | 13.6 | 0.99 | 5505773.1 |
| mammals | Rhinolophus_alcyone | Rhinolophus_alcyone | 0 | 18.8 | 0.98 | 2859335.95 |
| mammals | Rhinolophus_beddomei | Rhinolophus_beddomei | 0 | 14.5 | 0.99 | 186306.71 |
| mammals | Rhinolophus_blasii | Rhinolophus_blasii | 0 | 9.7 | 1 | 3410468.1 |
| mammals | Rhinolophus_bocharicus | Rhinolophus_bocharicus | 0 | 15.2 | 1 | 924906.69 |
| mammals | Rhinolophus_borneensis | Rhinolophus_borneensis | 0 | 7.9 | 1 | 1426753.69 |
| mammals | Rhinolophus_canuti | Rhinolophus_canuti | 1 | 12.7 | 1 | 10574.16 |
| mammals | Rhinolophus_capensis | Rhinolophus_capensis | 0 | 12.8 | 0.99 | 242578.76 |
| mammals | Rhinolophus_celebensis | Rhinolophus_celebensis | 0 | 6.7 | 1 | 308233.83 |
| mammals | Rhinolophus_clivosus | Rhinolophus_clivosus | 0 | 15.7 | 1 | 3659664.55 |
| mammals | Rhinolophus_coelophyllus | Rhinolophus_coelophyllus | 0 | 7.1 | 1 | 426055.56 |
| mammals | Rhinolophus_cognatus | Rhinolophus_cognatus | 1 | 5.4 | 1 | 3077.32 |
| mammals | Rhinolophus_creaghi | Rhinolophus_creaghi | 0 | 12 | 1 | 161673.08 |
| mammals | Rhinolophus_darlingi | Rhinolophus_darlingi | 0 | 9.5 | 1.96 | 1360113.63 |
| mammals | Rhinolophus_deckenii | Rhinolophus_deckenii | 0 | 16.8 | 1 | 187337.72 |
| mammals | Rhinolophus_denti | Rhinolophus_denti | 0 | 6.2 | 1 | 1704313.39 |
| mammals | Rhinolophus_eloquens | Rhinolophus_eloquens | 0 | 19.1 | 0.99 | 711922.77 |

| | | | | | | |
|---|---|---|---|---|---|---|
| mammals | Rhinolophus_euryale | Rhinolophus_euryale | 0 | 8.2 | 1 | 2991976.98 |
| mammals | Rhinolophus_euryotis | Rhinolophus_euryotis | 0 | 14.3 | 1 | 878387.64 |
| mammals | Rhinolophus_ferrumequinum | Rhinolophus_ferrumequinum | 0 | 21.1 | 0.98 | 9751019.67 |
| mammals | Rhinolophus_formosae | Rhinolophus_formosae | 0 | 20 | 0.99 | 28759.05 |
| mammals | Rhinolophus_fumigatus | Rhinolophus_fumigatus | 0 | 12.9 | 0.99 | 8364570.96 |
| mammals | Rhinolophus_guineensis | Rhinolophus_guineensis | 1 | 10.4 | 0.99 | 213322.92 |
| mammals | Rhinolophus_hildebrandtii | Rhinolophus_hildebrandtii | 0 | 25.8 | 0.99 | 3184495.86 |
| mammals | Rhinolophus_hilli | Rhinolophus_hilli | 1 | 16.5 | 0.98 | 122.03 |
| mammals | Rhinolophus_hillorum | Rhinolophus_hillorum | 1 | 14.8 | 1 | 28211.39 |
| mammals | Rhinolophus_hipposideros | Rhinolophus_hipposideros | 0 | 4.7 | 1 | 6256257.88 |
| mammals | Rhinolophus_inops | Rhinolophus_inops | 0 | 13.5 | 1 | 226653.04 |
| mammals | Rhinolophus_landeri | Rhinolophus_landeri | 0 | 8.9 | 0.99 | 12125111.27 |
| mammals | Rhinolophus_lepidus | Rhinolophus_lepidus | 0 | 5.7 | 1 | 3509104.61 |
| mammals | Rhinolophus_luctus | Rhinolophus_luctus | 0 | 33.5 | 0.99 | 5655752.2 |
| mammals | Rhinolophus_maclaudi | Rhinolophus_maclaudi | 1 | 13.1 | 0.98 | 8188.63 |
| mammals | Rhinolophus_macrotis | Rhinolophus_macrotis | 0 | 6.2 | 0.98 | 2405475.71 |
| mammals | Rhinolophus_madurensis | Rhinolophus_madurensis | 1 | 6.7 | 1 | 4927.5 |
| mammals | Rhinolophus_malayanus | Rhinolophus_malayanus | 0 | 6.7 | 1 | 1922229.02 |
| mammals | Rhinolophus_marshalli | Rhinolophus_marshalli | 0 | 5 | 1 | 714382.37 |
| mammals | Rhinolophus_megaphyllus | Rhinolophus_megaphyllus | 0 | 10.1 | 1 | 876558.9 |
| mammals | Rhinolophus_mehelyi | Rhinolophus_mehelyi | 1 | 12.3 | 2 | 1976290.66 |
| mammals | Rhinolophus_montanus | Rhinolophus_montanus | 1 | 6.7 | 1 | 71.28 |
| mammals | Rhinolophus_osgoodi | Rhinolophus_osgoodi | 0 | 7.3 | 1 | 769589.59 |
| mammals | Rhinolophus_pearsonii | Rhinolophus_pearsonii | 0 | 11.8 | 1 | 3176331.02 |
| mammals | Rhinolophus_philippinensis | Rhinolophus_philippinensis | 0 | 10.3 | 1 | 247505.55 |
| mammals | Rhinolophus_pusillus | Rhinolophus_pusillus | 0 | 7.2 | 1 | 3582812.62 |
| mammals | Rhinolophus_rex | Rhinolophus_rex | 1 | 11.3 | 1 | 2144413.65 |
| mammals | Rhinolophus_robinsoni | Rhinolophus_robinsoni | 0 | 8.5 | 1 | 47297.55 |
| mammals | Rhinolophus_rouxii | Rhinolophus_rouxii | 0 | 11.8 | 0.99 | 821256.08 |
| mammals | Rhinolophus_rufus | Rhinolophus_rufus | 0 | 30 | 1 | 34730.18 |
| mammals | Rhinolophus_ruwenzorii | Rhinolophus_ruwenzorii | 1 | 17.6 | 1 | 27330.97 |
| mammals | Rhinolophus_sedulus | Rhinolophus_sedulus | 0 | 8.8 | 1 | 805437.73 |
| mammals | Rhinolophus_shameli | Rhinolophus_shameli | 0 | 9.6 | 1 | 580977.06 |
| mammals | Rhinolophus_siamensis | Rhinolophus_siamensis | 0 | 5.1 | 0.98 | 1047159.94 |

| | | | | | | |
|---|---|---|---|---|---|---|
| mammals | Rhinolophus_simulator | Rhinolophus_simulator | 0 | 8.1 | 0.99 | 1911252.19 |
| mammals | Rhinolophus_sinicus | Rhinolophus_sinicus | 0 | 11.6 | 0.99 | 2168584.38 |
| mammals | Rhinolophus_stheno | Rhinolophus_stheno | 0 | 7.9 | 0.98 | 852346.83 |
| mammals | Rhinolophus_subbadius | Rhinolophus_subbadius | 0 | 6.6 | 1 | 76243.22 |
| mammals | Rhinolophus_swinnyi | Rhinolophus_swinnyi | 0 | 7.1 | 0.98 | 1347873.84 |
| mammals | Rhinolophus_thomasi | Rhinolophus_thomasi | 0 | 8.3 | 1 | 1408310.58 |
| mammals | Rhinolophus_trifoliatus | Rhinolophus_trifoliatus | 0 | 14.5 | 1 | 1344638.63 |
| mammals | Rhinolophus_virgo | Rhinolophus_virgo | 0 | 6 | 1 | 269684.68 |
| mammals | Rhinolophus_yunanensis | Rhinolophus_yunanensis | 0 | 19.7 | 1 | 401250.63 |
| mammals | Rhinolophus_ziama | Rhinolophus_ziama | 1 | 22.3 | 1 | 2057.05 |
| mammals | Rhinonicteris_aurantia | Rhinonicteris_aurantia | 0 | 8.8 | 1 | 747496.39 |
| mammals | Rhinophylla_alethina | Rhinophylla_alethina | 0 | 14 | 1 | 125700.98 |
| mammals | Rhinophylla_fischerae | Rhinophylla_fischerae | 0 | 10.6 | 0.98 | 4564653.78 |
| mammals | Rhinophylla_pumilio | Rhinophylla_pumilio | 0 | 9.6 | 0.98 | 8334315.24 |
| mammals | Rhinopithecus_avunculus | Rhinopithecus_avunculus | 1 | 8000 | 1 | 10209.64 |
| mammals | Rhinopithecus_bieti | Rhinopithecus_bieti | 1 | 9960 | 1 | 29673.8 |
| mammals | Rhinopithecus_brelichi | Rhinopithecus_brelichi | 1 | 11500 | 1 | 67.48 |
| mammals | Rhinopithecus_roxellana | Rhinopithecus_roxellana | 1 | 12125 | 1 | 250586.15 |
| mammals | Rhinopoma_hardwickii | Rhinopoma_hardwickii | 0 | 14.2 | 0.99 | 16931124.57 |
| mammals | Rhinopoma_microphyllum | Rhinopoma_microphyllum | 0 | 27.9 | 1 | 7151525.96 |
| mammals | Rhinopoma_muscatellum | Rhinopoma_muscatellum | 0 | 9.1 | 1 | 580313.15 |
| mammals | Rhinosciurus_laticaudatus | Rhinosciurus_laticaudatus | 0 | 228.3 | 1.39 | 911388.27 |
| mammals | Rhipidomys_austrinus | Rhipidomys_austrinus | 0 | 89 | 2 | 138614.19 |
| mammals | Rhipidomys_emiliae | Rhipidomys_emiliae | 0 | 96.5 | 3.8 | 698505.62 |
| mammals | Rhipidomys_fulviventer | Rhipidomys_fulviventer | 0 | 89 | 2 | 52004.66 |
| mammals | Rhipidomys_gardneri | Rhipidomys_gardneri | 0 | 71.1 | 3 | 152165.93 |
| mammals | Rhipidomys_latimanus | Rhipidomys_latimanus | 0 | 57.5 | 2 | 157706.17 |
| mammals | Rhipidomys_leucodactylus | Rhipidomys_leucodactylus | 0 | 80 | 2 | 4402230.2 |
| mammals | Rhipidomys_macconnelli | Rhipidomys_macconnelli | 0 | 41.6 | 3.5 | 189757.69 |
| mammals | Rhipidomys_macrurus | Rhipidomys_macrurus | 0 | 79.9 | 3.8 | 1565931.64 |
| mammals | Rhipidomys_mastacalis | Rhipidomys_mastacalis | 0 | 77.5 | 3.8 | 633285.53 |
| mammals | Rhipidomys_nitela | Rhipidomys_nitela | 0 | 56 | 2 | 1447250.06 |
| mammals | Rhipidomys_venezuelae | Rhipidomys_venezuelae | 0 | 90 | 3.8 | 172678.32 |
| mammals | Rhipidomys_venustus | Rhipidomys_venustus | 0 | 89 | 2 | 8414.42 |

| | | | | | | |
|---|---|---|---|---|---|---|
| mammals | Rhipidomys_wetzeli | Rhipidomys_wetzeli | 0 | 89 | 2 | 138914.11 |
| mammals | Rhizomys_sinensis | Rhizomys_sinensis | 0 | 865 | 3 | 2057627.57 |
| mammals | Rhizomys_sumatrensis | Rhizomys_sumatrensis | 0 | 1625 | 4 | 1372330.12 |
| mammals | Rhogeessa_aenea | Rhogeessa_aeneus | 0 | 3.4 | 1.69 | 144231.54 |
| mammals | Rhogeessa_genowaysi | Rhogeessa_genowaysi | 1 | 5 | 2 | 913.22 |
| mammals | Rhogeessa_io | Rhogeessa_io | 0 | 5 | 1.69 | 4609642.85 |
| mammals | Rhogeessa_minutilla | Rhogeessa_minutilla | 1 | 3.7 | 1.98 | 663829.69 |
| mammals | Rhogeessa_parvula | Rhogeessa_parvula | 0 | 4.4 | 1.98 | 448909.73 |
| mammals | Rhogeessa_tumida | Rhogeessa_tumida | 0 | 4.5 | 1.54 | 462224.3 |
| mammals | Rhombomys_opimus | Rhombomys_opimus | 0 | 144 | 4.8 | 4585921.88 |
| mammals | Rhynchocyon_chrysopygus | Rhynchocyon_chrysopygus | 1 | 540 | 1.11 | 3574.33 |
| mammals | Rhynchocyon_cirnei | Rhynchocyon_cirnei | 0 | 490 | 1.93 | 1130811.53 |
| mammals | Rhynchocyon_petersi | Rhynchocyon_petersi | 0 | 424 | 1.93 | 31086.28 |
| mammals | Rhynchocyon_udzungwensis | Rhynchocyon_udzungwensis | 1 | 711 | 1.5 | 320.4 |
| mammals | Rhynchogale_melleri | Rhynchogale_melleri | 0 | 2500 | 2 | 2215668.76 |
| mammals | Rhyncholestes_raphanurus | Rhyncholestes_raphanurus | 0 | 21 | 6 | 40999.52 |
| mammals | Rhynchomys_isarogensis | Rhynchomys_isarogensis | 1 | 136 | 1 | 46.2 |
| mammals | Rhynchomys_soricoides | Rhynchomys_soricoides | 0 | 156 | 1.5 | 4215.03 |
| mammals | Rhynchonycteris_naso | Rhynchonycteris_naso | 0 | 3.8 | 0.99 | 11508826.66 |
| mammals | Rhyneptesicus_nasutus | Eptesicus_nasutus | 0 | 7.9 | 1.5 | 1866190.54 |
| mammals | Romerolagus_diazi | Romerolagus_diazi | 1 | 476.7 | 2.12 | 444.36 |
| mammals | Rousettus_aegyptiacus | Rousettus_aegyptiacus | 0 | 132.2 | 1.05 | 4071413.31 |
| mammals | Rousettus_amplexicaudatus | Rousettus_amplexicaudatus | 0 | 82.5 | 0.99 | 4281498.89 |
| mammals | Rousettus_leschenaultii | Rousettus_leschenaultii | 0 | 97.1 | 1.01 | 6873172.22 |
| mammals | Rousettus_madagascariensis | Rousettus_madagascariensis | 1 | 62.1 | 1.17 | 292740.75 |
| mammals | Rousettus_obliviosus | Rousettus_obliviosus | 1 | 44.1 | 0.98 | 1667.12 |
| mammals | Rousettus_spinalatus | Rousettus_spinalatus | 1 | 90.8 | 1 | 84003.77 |
| mammals | Rubrisciurus_rubriventer | Rubrisciurus_rubriventer | 0 | 688.3 | 1 | 169893.7 |
| mammals | Rucervus_duvaucelii | Rucervus_duvaucelii | 1 | 143000 | 1 | 34612.9 |
| mammals | Rucervus_eldii | Rucervus_eldii | 1 | 73000 | 1.01 | 84180.7 |
| mammals | Rungwecebus_kipunji | Rungwecebus_kipunji | 1 | 13000 | 1 | 71.06 |
| mammals | Rupicapra_pyrenaica | Rupicapra_pyrenaica | 0 | 30000 | 1 | 15275.95 |
| mammals | Rupicapra_rupicapra | Rupicapra_rupicapra | 0 | 26100 | 1 | 209048 |
| mammals | Rusa_alfredi | Rusa_alfredi | 1 | 47500 | 1 | 1689.09 |

| | | | | | | |
|---|---|---|---|---|---|---|
| mammals | Rusa_marianna | Rusa_marianna | 1 | 50000 | 1 | 112256.43 |
| mammals | Rusa_timorensis | Rusa_timorensis | 1 | 63000 | 1 | 11319.65 |
| mammals | Rusa_unicolor | Rusa_unicolor | 1 | 180344.4 | 1 | 7570390.03 |
| mammals | Saccolaimus_flaviventris | Saccolaimus_flaviventris | 0 | 42.1 | 1 | 6052125.98 |
| mammals | Saccolaimus_mixtus | Saccolaimus_mixtus | 0 | 23 | 1 | 53569.26 |
| mammals | Saccolaimus_peli | Saccolaimus_peli | 0 | 100 | 1 | 1959912.67 |
| mammals | Saccolaimus_saccolaimus | Saccolaimus_saccolaimus | 0 | 41.8 | 1 | 1930053.01 |
| mammals | Saccopteryx_antioquensis | Saccopteryx_antioquensis | 1 | 5 | 1 | 3092.05 |
| mammals | Saccopteryx_bilineata | Saccopteryx_bilineata | 0 | 7.8 | 0.99 | 12632095.59 |
| mammals | Saccopteryx_canescens | Saccopteryx_canescens | 0 | 3.4 | 1 | 6384043.27 |
| mammals | Saccopteryx_leptura | Saccopteryx_leptura | 0 | 4.7 | 1 | 12289438.47 |
| mammals | Saccostomus_campestris | Saccostomus_campestris | 0 | 51.2 | 6.78 | 5347600.61 |
| mammals | Saccostomus_mearnsi | Saccostomus_mearnsi | 0 | 45.5 | 5.75 | 667846.57 |
| mammals | Saguinus_bicolor | Saguinus_bicolor | 1 | 465 | 1.81 | 10128.76 |
| mammals | Saguinus_fuscicollis | Saguinus_fuscicollis | 0 | 387 | 1.81 | 434351.01 |
| mammals | Saguinus_geoffroyi | Saguinus_geoffroyi | 0 | 486.5 | 1.96 | 75592.62 |
| mammals | Saguinus_imperator | Saguinus_imperator | 0 | 400 | 2 | 234318.65 |
| mammals | Saguinus_inustus | Saguinus_inustus | 0 | 410.5 | 2 | 380406.84 |
| mammals | Saguinus_labiatus | Saguinus_labiatus | 0 | 575 | 1.87 | 286447.29 |
| mammals | Saguinus_leucopus | Saguinus_leucopus | 1 | 440 | 2 | 56861.64 |
| mammals | Saguinus_martinsi | Saguinus_martinsi | 0 | 603.6 | 1.63 | 42109.16 |
| mammals | Saguinus_midas | Saguinus_midas | 0 | 540 | 2.01 | 863250.69 |
| mammals | Saguinus_mystax | Saguinus_mystax | 0 | 618 | 1.93 | 589295.47 |
| mammals | Saguinus_niger | Saguinus_niger | 1 | 591.2 | 2 | 281250.69 |
| mammals | Saguinus_nigricollis | Saguinus_nigricollis | 0 | 462 | 1.81 | 211767.62 |
| mammals | Saguinus_oedipus | Saguinus_oedipus | 1 | 430 | 1.9 | 51980.63 |
| mammals | Saguinus_tripartitus | Saguinus_tripartitus | 0 | 393.5 | 1.8 | 25699.95 |
| mammals | Saguinus_weddelli | Saguinus_melanoleucus | 0 | 491 | 1.8 | 597636.46 |
| mammals | Saiga_tatarica | Saiga_tatarica | 0 | 29000 | 1.54 | 749562.59 |
| mammals | Saimiri_boliviensis | Saimiri_boliviensis | 0 | 615 | 1.01 | 1373872.35 |
| mammals | Saimiri_oerstedii | Saimiri_oerstedii | 1 | 278.5 | 1.01 | 10599.79 |
| mammals | Saimiri_sciureus | Saimiri_sciureus | 0 | 743.2 | 1 | 956591.72 |
| mammals | Saimiri_ustus | Saimiri_ustus | 0 | 1000 | 1.02 | 876710.21 |
| mammals | Saimiri_vanzolinii | Saimiri_vanzolinii | 1 | 769.6 | 1.01 | 2930.59 |

| | | | | | | |
|---|---|---|---|---|---|---|
| mammals | Salpingotus_crassicauda | Salpingotus_crassicauda | 0 | 12 | 2.58 | 1359343.51 |
| mammals | Salpingotus_kozlovi | Salpingotus_kozlovi | 0 | 9.5 | 2.6 | 1541055.82 |
| mammals | Sapajus_apella | Sapajus_apella | 0 | 2500 | 1.05 | 5970729.54 |
| mammals | Sapajus_flavius | Sapajus_flavius | 1 | 2550 | 1 | 21664.32 |
| mammals | Sapajus_libidinosus | Sapajus_libidinosus | 0 | 3200 | 1.05 | 2612578.69 |
| mammals | Sapajus_xanthosternos | Sapajus_xanthosternos | 1 | 3200 | 1.05 | 466598.87 |
| mammals | Sarcophilus_harrisii | Sarcophilus_harrisii | 1 | 11000 | 2.91 | 64284.13 |
| mammals | Sauromys_petrophilus | Sauromys_petrophilus | 0 | 13.9 | 1.05 | 728212.46 |
| mammals | Scapteromys_aquaticus | Scapteromys_aquaticus | 0 | 116.7 | 3.63 | 260142.73 |
| mammals | Scapteromys_tumidus | Scapteromys_tumidus | 0 | 146 | 3.63 | 258565.21 |
| mammals | Scarturus_elater | Allactaga_elater | 0 | 59 | 4.48 | 3246015.7 |
| mammals | Scarturus_euphraticus | Allactaga_euphratica | 0 | 70 | 5 | 758072.4 |
| mammals | Scarturus_vinogradovi | Allactaga_vinogradovi | 0 | 75.2 | 4.5 | 52562.37 |
| mammals | Scarturus_williamsi | Allactaga_williamsi | 0 | 100 | 5 | 705703.7 |
| mammals | Sciurillus_pusillus | Sciurillus_pusillus | 0 | 39 | 1.94 | 1209632.3 |
| mammals | Sciurocheirus_alleni | Sciurocheirus_alleni | 0 | 332.5 | 1.25 | 94727.98 |
| mammals | Sciurocheirus_gabonensis | Sciurocheirus_gabonensis | 0 | 271.6 | 1.22 | 530152.24 |
| mammals | Sciurotamias_davidianus | Sciurotamias_davidianus | 0 | 433.5 | 2.5 | 1330267.36 |
| mammals | Sciurus_aberti | Sciurus_aberti | 0 | 624 | 3.23 | 270389.02 |
| mammals | Sciurus_aestuans | Sciurus_aestuans | 0 | 185 | 2.3 | 6983651.39 |
| mammals | Sciurus_alleni | Sciurus_alleni | 0 | 434.5 | 3 | 48935.03 |
| mammals | Sciurus_anomalus | Sciurus_anomalus | 0 | 256 | 4 | 614332.4 |
| mammals | Sciurus_aureogaster | Sciurus_aureogaster | 0 | 595 | 1.97 | 620614.53 |
| mammals | Sciurus_carolinensis | Sciurus_carolinensis | 0 | 506.5 | 2.99 | 4083873.2 |
| mammals | Sciurus_colliaei | Sciurus_colliaei | 0 | 498 | 2.66 | 81302.04 |
| mammals | Sciurus_deppei | Sciurus_deppei | 0 | 190 | 2.81 | 694996.56 |
| mammals | Sciurus_granatensis | Sciurus_granatensis | 0 | 250 | 1.92 | 988528.23 |
| mammals | Sciurus_griseus | Sciurus_griseus | 0 | 731 | 2.98 | 376963.85 |
| mammals | Sciurus_ignitus | Sciurus_ignitus | 0 | 190 | 2 | 2092301.56 |
| mammals | Sciurus_igniventris | Sciurus_igniventris | 0 | 700 | 2 | 2756262.84 |
| mammals | Sciurus_lis | Sciurus_lis | 0 | 264.3 | 4.43 | 214979.47 |
| mammals | Sciurus_nayaritensis | Sciurus_nayaritensis | 0 | 697 | 2.46 | 249137.65 |
| mammals | Sciurus_niger | Sciurus_niger | 0 | 761.9 | 3 | 4360630.24 |
| mammals | Sciurus_richmondi | Sciurus_richmondi | 0 | 205 | 2.61 | 22521.1 |

| | | | | | | |
|---|---|---|---|---|---|---|
| mammals | Sciurus_spadiceus | Sciurus_spadiceus | 0 | 403.3 | 3 | 3247882.95 |
| mammals | Sciurus_variegatoides | Sciurus_variegatoides | 0 | 485 | 4.99 | 392638.55 |
| mammals | Sciurus_vulgaris | Sciurus_vulgaris | 0 | 333 | 4.46 | 27053169.45 |
| mammals | Sciurus_yucatanensis | Sciurus_yucatanensis | 0 | 225 | 3 | 194232.52 |
| mammals | Scolomys_melanops | Scolomys_melanops | 0 | 26.5 | 2.5 | 100718.94 |
| mammals | Scolomys_ucayalensis | Scolomys_ucayalensis | 0 | 26.5 | 2 | 197626.02 |
| mammals | Scoteanax_rueppellii | Scoteanax_rueppellii | 0 | 26.5 | 1 | 301736.58 |
| mammals | Scotinomys_teguina | Scotinomys_teguina | 0 | 11.3 | 2.54 | 129224.41 |
| mammals | Scotinomys_xerampelinus | Scotinomys_xerampelinus | 0 | 15 | 3.07 | 5119.33 |
| mammals | Scotoecus_hirundo | Scotoecus_hirundo | 0 | 9.3 | 1.96 | 4917152.73 |
| mammals | Scotoecus_pallidus | Scotoecus_pallidus | 0 | 11.65 | 2 | 187893.76 |
| mammals | Scotonycteris_zenkeri | Scotonycteris_zenkeri | 0 | 20.8 | 0.99 | 114273.67 |
| mammals | Scotophilus_collinus | Scotophilus_collinus | 0 | 23.4 | 1.5 | 210159.19 |
| mammals | Scotophilus_dinganii | Scotophilus_dinganii | 0 | 25.1 | 1.85 | 10952348.34 |
| mammals | Scotophilus_heathii | Scotophilus_heathii | 0 | 34.6 | 1.92 | 5950831.34 |
| mammals | Scotophilus_kuhlii | Scotophilus_kuhlii | 0 | 20.7 | 1.5 | 5208270.37 |
| mammals | Scotophilus_leucogaster | Scotophilus_leucogaster | 0 | 20.2 | 1.44 | 5430385.92 |
| mammals | Scotophilus_nigrita | Scotophilus_nigrita | 0 | 101 | 1.5 | 2196022.14 |
| mammals | Scotophilus_nux | Scotophilus_nux | 0 | 30 | 1.5 | 951762.19 |
| mammals | Scotophilus_viridis | Scotophilus_viridis | 0 | 19.6 | 2 | 4650910.92 |
| mammals | Scotorepens_balstoni | Scotorepens_balstoni | 0 | 11.7 | 1.46 | 4980250.11 |
| mammals | Scotorepens_greyii | Scotorepens_greyii | 0 | 10 | 1.9 | 4064788.85 |
| mammals | Scotorepens_orion | Scotorepens_orion | 0 | 11 | 1.5 | 265076.92 |
| mammals | Scotorepens_sanborni | Scotorepens_sanborni | 0 | 8 | 1.5 | 702866.88 |
| mammals | Scotozous_dormeri | Scotozous_dormeri | 0 | 6.9 | 1.08 | 2183092.1 |
| mammals | Scutisorex_somereni | Scutisorex_somereni | 0 | 91.5 | 1.85 | 434028.62 |
| mammals | Sekeetamys_calurus | Sekeetamys_calurus | 0 | 55.3 | 2.9 | 200686.83 |
| mammals | Semnopithecus_ajax | Semnopithecus_ajax | 1 | 20150 | 1.02 | 8509.04 |
| mammals | Semnopithecus_entellus | Semnopithecus_entellus | 0 | 14500 | 1.02 | 558928.44 |
| mammals | Semnopithecus_hector | Semnopithecus_hector | 0 | 17350 | 1.02 | 101204.74 |
| mammals | Semnopithecus_hypoleucos | Semnopithecus_hypoleucos | 0 | 8950 | 1.02 | 407292.34 |
| mammals | Semnopithecus_johnii | Trachypithecus_johnii | 1 | 11921 | 1.01 | 24067.17 |
| mammals | Semnopithecus_priam | Semnopithecus_priam | 0 | 12450 | 1.02 | 190726.32 |
| mammals | Semnopithecus_schistaceus | Semnopithecus_schistaceus | 0 | 17350 | 1.02 | 311696.72 |

| | | | | | | |
|---|---|---|---|---|---|---|
| mammals | Semnopithecus_vetulus | Trachypithecus_vetulus | 1 | 8830 | 1 | 36899.92 |
| mammals | Setifer_setosus | Setifer_setosus | 0 | 294 | 3.14 | 590831.63 |
| mammals | Setonix_brachyurus | Setonix_brachyurus | 1 | 3250 | 1 | 18067.24 |
| mammals | Sicista_betulina | Sicista_betulina | 0 | 8 | 5.18 | 6815198.93 |
| mammals | Sicista_caucasica | Sicista_caucasica | 0 | 10 | 5 | 6718.83 |
| mammals | Sicista_concolor | Sicista_concolor | 0 | 6.5 | 4.5 | 527987.06 |
| mammals | Sicista_kazbegica | Sicista_kazbegica | 1 | 7 | 5.5 | 3375.93 |
| mammals | Sicista_kluchorica | Sicista_kluchorica | 0 | 7.5 | 6 | 7368.94 |
| mammals | Sicista_napaea | Sicista_napaea | 0 | 10 | 3.49 | 117911.87 |
| mammals | Sicista_strandi | Sicista_strandi | 0 | 10.3 | 5.08 | 652702.1 |
| mammals | Sicista_subtilis | Sicista_subtilis | 0 | 11.8 | 5 | 2576602.47 |
| mammals | Sicista_tianschanica | Sicista_tianshanica | 0 | 11.5 | 4.37 | 358985.78 |
| mammals | Sigmodon_alleni | Sigmodon_alleni | 1 | 179 | 5.44 | 174869.57 |
| mammals | Sigmodon_alstoni | Sigmodon_alstoni | 0 | 55.7 | 5.14 | 840163.26 |
| mammals | Sigmodon_arizonae | Sigmodon_arizonae | 0 | 198 | 5.44 | 345280.16 |
| mammals | Sigmodon_fulviventer | Sigmodon_fulviventer | 0 | 214 | 6.88 | 464112.15 |
| mammals | Sigmodon_hirsutus | Sigmodon_hirsutus | 0 | 171.5 | 5.44 | 761233.05 |
| mammals | Sigmodon_hispidus | Sigmodon_hispidus | 0 | 92.4 | 5.44 | 2615707.97 |
| mammals | Sigmodon_inopinatus | Sigmodon_inopinatus | 1 | 140.5 | 6.25 | 13693.25 |
| mammals | Sigmodon_leucotis | Sigmodon_leucotis | 0 | 135.5 | 8.5 | 282532.31 |
| mammals | Sigmodon_mascotensis | Sigmodon_mascotensis | 0 | 120 | 5.44 | 252559.65 |
| mammals | Sigmodon_ochrognathus | Sigmodon_ochrognathus | 0 | 122 | 4 | 464511.07 |
| mammals | Sigmodon_peruanus | Sigmodon_peruanus | 0 | 88.7 | 5.44 | 98145.69 |
| mammals | Sigmodon_toltecus | Sigmodon_toltecus | 0 | 50 | 5.44 | 416823.49 |
| mammals | Simias_concolor | Simias_concolor | 1 | 7980 | 1.02 | 6004.48 |
| mammals | Sminthopsis_butleri | Sminthopsis_butleri | 1 | 25 | 7 | 7067.24 |
| mammals | Sminthopsis_crassicaudata | Sminthopsis_crassicaudata | 0 | 15 | 6.76 | 4137398.35 |
| mammals | Sminthopsis_dolichura | Sminthopsis_dolichura | 0 | 13.6 | 7.2 | 551815.74 |
| mammals | Sminthopsis_douglasi | Sminthopsis_douglasi | 0 | 55 | 6.78 | 47129.4 |
| mammals | Sminthopsis_fuliginosa | Sminthopsis_griseoventer | 0 | 17.5 | 8 | 144138.8 |
| mammals | Sminthopsis_gilberti | Sminthopsis_gilberti | 0 | 19.5 | 7 | 76628.82 |
| mammals | Sminthopsis_leucopus | Sminthopsis_leucopus | 0 | 26 | 8.2 | 96976.3 |
| mammals | Sminthopsis_macroura | Sminthopsis_macroura | 0 | 20 | 7 | 4300281.36 |
| mammals | Sminthopsis_murina | Sminthopsis_murina | 0 | 17 | 8 | 1210760.96 |

| | | | | | | |
|---|---|---|---|---|---|---|
| mammals | Sminthopsis_ooldea | Sminthopsis_ooldea | 0 | 11 | 7.24 | 1133609.41 |
| mammals | Sminthopsis_psammophila | Sminthopsis_psammophila | 1 | 33.5 | 5.49 | 7619.61 |
| mammals | Sminthopsis_virginiae | Sminthopsis_virginiae | 0 | 35.3 | 3.71 | 362600.72 |
| mammals | Sminthopsis_youngsoni | Sminthopsis_youngsoni | 0 | 10.1 | 5.5 | 1373345.18 |
| mammals | Smutsia_gigantea | Smutsia_gigantea | 1 | 32999.7 | 1 | 3497994.29 |
| mammals | Smutsia_temminckii | Smutsia_temminckii | 1 | 7179.9 | 1 | 7124374.96 |
| mammals | Solenodon_paradoxus | Solenodon_paradoxus | 0 | 900 | 1.75 | 13314.15 |
| mammals | Solomys_ponceleti | Solomys_ponceleti | 1 | 1000 | 1.5 | 11785.14 |
| mammals | Solomys_salebrosus | Solomys_salebrosus | 1 | 397 | 1 | 9993.96 |
| mammals | Sorex_alpinus | Sorex_alpinus | 0 | 7.8 | 5.95 | 422525.15 |
| mammals | Sorex_antinorii | Sorex_antinorii | 0 | 11.5 | 6.56 | 189627.14 |
| mammals | Sorex_araneus | Sorex_araneus | 0 | 10 | 6.56 | 13236507.59 |
| mammals | Sorex_arcticus | Sorex_arcticus | 0 | 8.2 | 6.66 | 4533841.87 |
| mammals | Sorex_asper | Sorex_asper | 0 | 7.9 | 5.7 | 379195.76 |
| mammals | Sorex_bedfordiae | Sorex_bedfordiae | 0 | 6.6 | 4.99 | 585011.12 |
| mammals | Sorex_bendirii | Sorex_bendirii | 0 | 16.1 | 5 | 207238.1 |
| mammals | Sorex_caecutiens | Sorex_caecutiens | 0 | 5.4 | 7.03 | 17639588.44 |
| mammals | Sorex_camtschaticus | Sorex_camtschatica | 0 | 4.6 | 6.5 | 447577.75 |
| mammals | Sorex_coronatus | Sorex_coronatus | 0 | 9.6 | 5.48 | 884132.19 |
| mammals | Sorex_cylindricauda | Sorex_cylindricauda | 0 | 6.1 | 4.99 | 327292.25 |
| mammals | Sorex_daphaenodon | Sorex_daphaenodon | 0 | 6.8 | 5.99 | 8192141.48 |
| mammals | Sorex_excelsus | Sorex_excelsus | 0 | 7.5 | 6.56 | 200703.77 |
| mammals | Sorex_fumeus | Sorex_fumeus | 0 | 7.7 | 5.37 | 1508616.01 |
| mammals | Sorex_gracillimus | Sorex_gracillimus | 0 | 4.5 | 5.18 | 1136758.49 |
| mammals | Sorex_granarius | Sorex_granarius | 0 | 6.3 | 5.64 | 69946.85 |
| mammals | Sorex_haydeni | Sorex_haydeni | 0 | 3.5 | 6.78 | 1160249.7 |
| mammals | Sorex_hosonoi | Sorex_hosonoi | 0 | 3.3 | 6 | 10546.77 |
| mammals | Sorex_isodon | Sorex_isodon | 0 | 9.4 | 7.09 | 12216462.27 |
| mammals | Sorex_jacksoni | Sorex_jacksoni | 0 | 3.5 | 6.5 | 4929.27 |
| mammals | Sorex_longirostris | Sorex_longirostris | 0 | 3.6 | 4.43 | 1643808.72 |
| mammals | Sorex_maritimensis | Sorex_maritimensis | 0 | 7.3 | 6.63 | 61267.68 |
| mammals | Sorex_merriami | Sorex_merriami | 0 | 5.9 | 6 | 1800350.22 |
| mammals | Sorex_milleri | Sorex_milleri | 1 | 7 | 6 | 11692.59 |
| mammals | Sorex_minutissimus | Sorex_minutissimus | 0 | 2.5 | 8.99 | 17514810.1 |

| | | | | | | |
|---|---|---|---|---|---|---|
| mammals | Sorex_minutus | Sorex_minutus | 0 | 4.5 | 5.97 | 14288920.19 |
| mammals | Sorex_monticolus | Sorex_monticolus | 0 | 5.3 | 5.72 | 4535520.83 |
| mammals | Sorex_nanus | Sorex_nanus | 0 | 2.4 | 6.49 | 997639.34 |
| mammals | Sorex_orizabae | Sorex_orizabae | 0 | 4.8 | 5.54 | 56542.85 |
| mammals | Sorex_pacificus | Sorex_pacificus | 0 | 6.7 | 4.47 | 93854.05 |
| mammals | Sorex_palustris | Sorex_palustris | 0 | 13.5 | 5.52 | 6242380.89 |
| mammals | Sorex_planiceps | Sorex_planiceps | 0 | 10.5 | 5.9 | 27733.29 |
| mammals | Sorex_preblei | Sorex_preblei | 0 | 3.1 | 4.4 | 688410.09 |
| mammals | Sorex_raddei | Sorex_raddei | 0 | 10 | 2.69 | 245664.42 |
| mammals | Sorex_roboratus | Sorex_roboratus | 0 | 9.1 | 7.04 | 9303837.8 |
| mammals | Sorex_samniticus | Sorex_samniticus | 0 | 8.3 | 6.56 | 157801.64 |
| mammals | Sorex_satunini | Sorex_satunini | 0 | 7.5 | 6.3 | 412610.26 |
| mammals | Sorex_shinto | Sorex_shinto | 0 | 5.9 | 6.82 | 27674.38 |
| mammals | Sorex_sonomae | Sorex_sonomae | 0 | 8.3 | 4.74 | 143305.12 |
| mammals | Sorex_trowbridgii | Sorex_trowbridgii | 0 | 3.8 | 4.03 | 377575.97 |
| mammals | Sorex_tundrensis | Sorex_tundrensis | 0 | 7.7 | 7.7 | 15993094.22 |
| mammals | Sorex_ugyunak | Sorex_ugyunak | 0 | 3.6 | 6.5 | 1412342.54 |
| mammals | Sorex_unguiculatus | Sorex_unguiculatus | 0 | 9.9 | 5.24 | 1194028.93 |
| mammals | Sorex_vagrans | Sorex_vagrans | 0 | 5.9 | 5.69 | 973482.08 |
| mammals | Sorex_volnuchini | Sorex_volnuchini | 0 | 5.5 | 5.9 | 434037.43 |
| mammals | Soricomys_kalinga | Soricomys_kalinga | 0 | 25.5 | 2 | 83.15 |
| mammals | Soriculus_nigrescens | Soriculus_nigrescens | 0 | 14 | 4.89 | 136320.64 |
| mammals | Spalacopus_cyanus | Spalacopus_cyanus | 0 | 127.5 | 2.77 | 76144.18 |
| mammals | Speothos_venaticus | Speothos_venaticus | 0 | 6000 | 3.96 | 11389107.51 |
| mammals | Spermophilopsis_leptodactyla | Spermophilopsis_leptodactylus | 0 | 600 | 4.46 | 1268483.87 |
| mammals | Spermophilus_alaschanicus | Spermophilus_alashanicus | 0 | 208 | 4.82 | 487834.93 |
| mammals | Spermophilus_brevicauda | Spermophilus_brevicauda | 0 | 289.5 | 6.78 | 564431.59 |
| mammals | Spermophilus_citellus | Spermophilus_citellus | 1 | 290 | 6.24 | 432578.76 |
| mammals | Spermophilus_dauricus | Spermophilus_dauricus | 0 | 161.3 | 5.42 | 1379395.54 |
| mammals | Spermophilus_erythrogenys | Spermophilus_erythrogenys | 0 | 355 | 7 | 1727062.33 |
| mammals | Spermophilus_fulvus | Spermophilus_fulvus | 0 | 596 | 6 | 1851642.09 |
| mammals | Spermophilus_major | Spermophilus_major | 0 | 786 | 7.5 | 629933.3 |
| mammals | Spermophilus_musicus | Spermophilus_musicus | 0 | 316.9 | 7.39 | 1036.69 |
| mammals | Spermophilus_nilkaensis | Spermophilus_ralli | 0 | 347.5 | 6 | 39704.86 |

| | | | | | | |
|---|---|---|---|---|---|---|
| mammals | Spermophilus_pallidicauda | Spermophilus_pallidicauda | 0 | 439.5 | 6.24 | 699451.66 |
| mammals | Spermophilus_pygmaeus | Spermophilus_pygmaeus | 0 | 136 | 7.24 | 1810870.85 |
| mammals | Spermophilus_relictus | Spermophilus_relictus | 0 | 600 | 5.3 | 188222.91 |
| mammals | Spermophilus_suslicus | Spermophilus_suslicus | 0 | 252 | 5.74 | 804158.5 |
| mammals | Spermophilus_xanthoprymnus | Spermophilus_xanthoprymnus | 0 | 289.7 | 5.5 | 475379.4 |
| mammals | Sphaeronycteris_toxophyllum | Sphaeronycteris_toxophyllum | 0 | 16.2 | 1 | 3805535.48 |
| mammals | Spilocuscus_kraemeri | Spilocuscus_kraemeri | 0 | 2300 | 1.14 | 1892.78 |
| mammals | Spilocuscus_maculatus | Spilocuscus_maculatus | 0 | 5106.8 | 1.45 | 886129.18 |
| mammals | Spilocuscus_papuensis | Spilocuscus_papuensis | 1 | 5735.5 | 1.63 | 3142.73 |
| mammals | Spilocuscus_rufoniger | Spilocuscus_rufoniger | 1 | 6000 | 1.02 | 178086.3 |
| mammals | Spilocuscus_wilsoni | Spilocuscus_wilsoni | 1 | 3262.3 | 1 | 2524.16 |
| mammals | Spilogale_angustifrons | Spilogale_angustifrons | 0 | 386.5 | 7.5 | 777447.14 |
| mammals | Spilogale_gracilis | Spilogale_gracilis | 0 | 521.8 | 5.07 | 3825446.68 |
| mammals | Spilogale_putorius | Spilogale_putorius | 1 | 341 | 5.04 | 2979584.79 |
| mammals | Spilogale_pygmaea | Spilogale_pygmaea | 1 | 235 | 3.92 | 95518.34 |
| mammals | Steatomys_caurinus | Steatomys_caurinus | 0 | 44.8 | 4.32 | 1058295.65 |
| mammals | Steatomys_krebsii | Steatomys_krebsii | 0 | 20.7 | 4.57 | 979358.6 |
| mammals | Steatomys_opimus | Steatomys_opimus | 0 | 42.7 | 4.32 | 721011.85 |
| mammals | Steatomys_pratensis | Steatomys_pratensis | 0 | 48 | 4.14 | 3477980.21 |
| mammals | Stenocephalemys_albipes | Stenocephalemys_albipes | 0 | 59.5 | 4.41 | 360891.9 |
| mammals | Stenocephalemys_albocaudatus | Stenocephalemys_albocaudata | 0 | 144 | 3.6 | 12140.11 |
| mammals | Stenocranius_gregalis | Microtus_gregalis | 0 | 47.5 | 8.48 | 4537963.91 |
| mammals | Stenoderma_rufum | Stenoderma_rufum | 0 | 20.7 | 0.99 | 9281.03 |
| mammals | Stochomys_longicaudatus | Stochomys_longicaudatus | 0 | 71 | 2.54 | 1866744.35 |
| mammals | Sturnira_aratathomasi | Sturnira_aratathomasi | 0 | 48.6 | 0.99 | 72657.45 |
| mammals | Sturnira_bidens | Sturnira_bidens | 0 | 17.6 | 1 | 705384.62 |
| mammals | Sturnira_bogotensis | Sturnira_bogotensis | 0 | 20.8 | 0.98 | 501247.49 |
| mammals | Sturnira_erythromos | Sturnira_erythromos | 0 | 15.1 | 0.99 | 2534366.14 |
| mammals | Sturnira_lilium | Sturnira_lilium | 0 | 19.1 | 1 | 4859393.43 |
| mammals | Sturnira_ludovici | Sturnira_ludovici | 0 | 21 | 0.99 | 1216315.86 |
| mammals | Sturnira_magna | Sturnira_magna | 0 | 48.6 | 0.98 | 1567758.19 |
| mammals | Sturnira_mordax | Sturnira_mordax | 0 | 25.1 | 0.98 | 28315.49 |
| mammals | Sturnira_nana | Sturnira_nana | 1 | 10.9 | 0.98 | 2319.44 |
| mammals | Sturnira_oporaphilum | Sturnira_oporaphilum | 0 | 26.5 | 0.99 | 1015753.63 |

| | | | | | | |
|---|---|---|---|---|---|---|
| mammals | Sturnira_tildae | Sturnira_tildae | 0 | 24 | 0.98 | 10661362.1 |
| mammals | Styloctenium_mindorense | Styloctenium_mindorensis | 1 | 212 | 1 | 485.35 |
| mammals | Stylodipus_andrewsi | Stylodipus_andrewsi | 0 | 60 | 3.24 | 940743.69 |
| mammals | Stylodipus_sungorus | Stylodipus_sungorus | 0 | 76.9 | 3.5 | 61392.54 |
| mammals | Stylodipus_telum | Stylodipus_telum | 0 | 80 | 3.37 | 1340954.31 |
| mammals | Subulo_gouazoubira | Mazama_gouazoubira | 0 | 16300.1 | 1 | 5746702.26 |
| mammals | Suncus_aequatorius | Suncus_aequatorius | 1 | 8.8 | 2.99 | 491.26 |
| mammals | Suncus_etruscus | Suncus_etruscus | 0 | 2.1 | 3.98 | 3765817.87 |
| mammals | Suncus_lixa | Suncus_lixus | 0 | 4.9 | 2.99 | 1437689.01 |
| mammals | Suncus_malayanus | Suncus_malayanus | 0 | 5 | 3.96 | 171596.12 |
| mammals | Suncus_megalurus | Suncus_megalura | 0 | 4.7 | 1.89 | 4240345.55 |
| mammals | Suncus_montanus | Suncus_montanus | 1 | 18 | 3.05 | 5369.62 |
| mammals | Suncus_murinus | Suncus_murinus | 0 | 61.5 | 3.08 | 9425536.56 |
| mammals | Suncus_remyi | Suncus_remyi | 0 | 1.8 | 1 | 150166.81 |
| mammals | Suncus_varilla | Suncus_varilla | 0 | 5 | 3.44 | 2013476.9 |
| mammals | Suncus_zeylanicus | Suncus_zeylanicus | 1 | 29.8 | 3.05 | 456.49 |
| mammals | Sundamys_muelleri | Sundamys_muelleri | 0 | 275.7 | 3.8 | 1408490.37 |
| mammals | Sundasciurus_fraterculus | Sundasciurus_fraterculus | 1 | 58.6 | 2.5 | 6004.26 |
| mammals | Sundasciurus_lowii | Sundasciurus_lowii | 0 | 87.5 | 2.48 | 1335061.88 |
| mammals | Sundasciurus_tenuis | Sundasciurus_tenuis | 0 | 73.8 | 3.06 | 1210112.93 |
| mammals | Surdisorex_norae | Surdisorex_norae | 0 | 9 | 1.31 | 3057.85 |
| mammals | Suricata_suricatta | Suricata_suricatta | 0 | 725.5 | 3.86 | 1758699.4 |
| mammals | Sus_ahoenobarbus | Sus_ahoenobarbus | 0 | 56749.7 | 4 | 10580.52 |
| mammals | Sus_barbatus | Sus_barbatus | 1 | 70500 | 5.3 | 813194.66 |
| mammals | Sus_cebifrons | Sus_cebifrons | 1 | 32500 | 4.8 | 2382 |
| mammals | Sus_celebensis | Sus_celebensis | 0 | 55000 | 4.36 | 257141.82 |
| mammals | Sus_oliveri | Sus_oliveri | 1 | 44801.2 | 5.24 | 2450.55 |
| mammals | Sus_philippensis | Sus_philippensis | 1 | 44253.5 | 5.74 | 64212.43 |
| mammals | Sus_scrofa | Sus_scrofa | 0 | 101052.1 | 5.04 | 28026537.21 |
| mammals | Sus_verrucosus | Sus_verrucosus | 1 | 92500 | 5.6 | 4537.91 |
| mammals | Syconycteris_australis | Syconycteris_australis | 0 | 17.1 | 1 | 1053996.42 |
| mammals | Syconycteris_hobbit | Syconycteris_hobbit | 0 | 18.3 | 0.98 | 4176.04 |
| mammals | Sylvicapra_grimmia | Sylvicapra_grimmia | 0 | 19499.8 | 0.99 | 13576010.78 |
| mammals | Sylvilagus_aquaticus | Sylvilagus_aquaticus | 0 | 2135.8 | 3.04 | 909647.52 |

| | | | | | | |
|---|---|---|---|---|---|---|
| mammals | Sylvilagus_audubonii | Sylvilagus_audubonii | 0 | 889.6 | 3 | 3575896.79 |
| mammals | Sylvilagus_bachmani | Sylvilagus_bachmani | 0 | 643.3 | 3.42 | 341187.89 |
| mammals | Sylvilagus_brasiliensis | Sylvilagus_brasiliensis | 1 | 950 | 2.22 | 7611.78 |
| mammals | Sylvilagus_cunicularius | Sylvilagus_cunicularius | 0 | 3000 | 3.71 | 323840.65 |
| mammals | Sylvilagus_dicei | Sylvilagus_dicei | 1 | 1442.9 | 3.85 | 5963.59 |
| mammals | Sylvilagus_floridanus | Sylvilagus_floridanus | 0 | 1172.8 | 4.81 | 6880226.51 |
| mammals | Sylvilagus_idahoensis | Brachylagus_idahoensis | 0 | 421.3 | 5.86 | 480341.7 |
| mammals | Sylvilagus_nuttallii | Sylvilagus_nuttallii | 0 | 755.1 | 4.21 | 2067003.63 |
| mammals | Sylvilagus_obscurus | Sylvilagus_obscurus | 0 | 653.1 | 4 | 94346.01 |
| mammals | Sylvilagus_palustris | Sylvilagus_palustris | 0 | 1500 | 2.65 | 361966.32 |
| mammals | Sylvilagus_transitionalis | Sylvilagus_transitionalis | 1 | 656.7 | 4.59 | 10493.36 |
| mammals | Sylvisorex_camerunensis | Sylvisorex_camerunensis | 1 | 5 | 1.89 | 6332.79 |
| mammals | Sylvisorex_granti | Sylvisorex_granti | 0 | 4 | 1.83 | 29983.03 |
| mammals | Sylvisorex_howelli | Sylvisorex_howelli | 0 | 4.3 | 2 | 6764.49 |
| mammals | Sylvisorex_lunaris | Sylvisorex_lunaris | 0 | 18.5 | 2.69 | 13561.08 |
| mammals | Sylvisorex_morio | Sylvisorex_morio | 1 | 8 | 1.99 | 444.44 |
| mammals | Sylvisorex_ollula | Sylvisorex_ollula | 0 | 13.2 | 2.75 | 1407399.49 |
| mammals | Symphalangus_syndactylus | Symphalangus_syndactylus | 1 | 10000 | 1.02 | 340358.94 |
| mammals | Synaptomys_cooperi | Synaptomys_cooperi | 0 | 31.9 | 3.15 | 3223190.97 |
| mammals | Syncerus_caffer | Syncerus_caffer | 0 | 580002.7 | 1.08 | 8443088.91 |
| mammals | Tachyglossus_aculeatus | Tachyglossus_aculeatus | 0 | 3025 | 1 | 7766679.89 |
| mammals | Tadarida_aegyptiaca | Tadarida_aegyptiaca | 0 | 16.8 | 1 | 4174304.73 |
| mammals | Tadarida_brasiliensis | Tadarida_brasiliensis | 0 | 12.5 | 1.08 | 13833395.32 |
| mammals | Tadarida_fulminans | Tadarida_fulminans | 0 | 35.3 | 1 | 295187 |
| mammals | Tadarida_latouchei | Tadarida_latouchei | 1 | 22 | 1 | 186758.51 |
| mammals | Tadarida_lobata | Tadarida_lobata | 0 | 26 | 1.05 | 186860.18 |
| mammals | Tadarida_teniotis | Tadarida_teniotis | 0 | 27 | 0.99 | 3893776.11 |
| mammals | Taeromys_celebensis | Taeromys_celebensis | 0 | 323 | 2.5 | 86669.87 |
| mammals | Taeromys_dominator | Paruromys_dominator | 0 | 342 | 3.5 | 169337.75 |
| mammals | Taeromys_taerae | Taeromys_taerae | 1 | 247 | 4.37 | 4355.21 |
| mammals | Tamandua_mexicana | Tamandua_mexicana | 0 | 4210 | 1 | 1495856.05 |
| mammals | Tamandua_tetradactyla | Tamandua_tetradactyla | 0 | 5515.1 | 1 | 12732832.02 |
| mammals | Tamias_striatus | Tamias_striatus | 0 | 111.9 | 4.22 | 4197616.71 |
| mammals | Tamiasciurus_douglasii | Tamiasciurus_douglasii | 0 | 225 | 5.46 | 511249.76 |

| | | | | | | |
|---|---|---|---|---|---|---|
| mammals | Tamiasciurus_hudsonicus | Tamiasciurus_hudsonicus | 0 | 201.2 | 4.07 | 10363663.52 |
| mammals | Tamiops_maritimus | Tamiops_maritimus | 0 | 70.3 | 2.86 | 1718636.15 |
| mammals | Tamiops_mcclellandii | Tamiops_macclellandii | 0 | 55.5 | 2.3 | 1464987.46 |
| mammals | Tamiops_swinhoei | Tamiops_swinhoei | 0 | 85 | 3.36 | 869533.08 |
| mammals | Tapecomys_primus | Tapecomys_primus | 0 | 71.5 | 1 | 2330.95 |
| mammals | Tapecomys_wolffsohni | Phyllotis_wolffsohni | 0 | 42.5 | 4.02 | 83243.94 |
| mammals | Taphozous_australis | Taphozous_australis | 0 | 22.9 | 1 | 38340.92 |
| mammals | Taphozous_georgianus | Taphozous_georgianus | 0 | 27.7 | 1 | 1198755.15 |
| mammals | Taphozous_hildegardeae | Taphozous_hildegardeae | 1 | 29.6 | 1 | 44478.92 |
| mammals | Taphozous_hilli | Taphozous_hilli | 0 | 22 | 1 | 1411632.65 |
| mammals | Taphozous_kapalgensis | Taphozous_kapalgensis | 0 | 26 | 1 | 97559.25 |
| mammals | Taphozous_longimanus | Taphozous_longimanus | 0 | 26.9 | 0.99 | 4903146.15 |
| mammals | Taphozous_mauritianus | Taphozous_mauritianus | 0 | 27.3 | 1 | 10153657.37 |
| mammals | Taphozous_melanopogon | Taphozous_melanopogon | 0 | 25.9 | 1 | 5670373.5 |
| mammals | Taphozous_nudiventris | Taphozous_nudiventris | 0 | 30.1 | 1 | 1206353.71 |
| mammals | Taphozous_perforatus | Taphozous_perforatus | 0 | 27.3 | 1 | 2114774.55 |
| mammals | Taphozous_theobaldi | Taphozous_theobaldi | 0 | 39.8 | 1 | 1042380.04 |
| mammals | Taphozous_troughtoni | Taphozous_troughtoni | 0 | 32.5 | 1 | 741835.35 |
| mammals | Tapirus_bairdii | Tapirus_bairdii | 1 | 299999.1 | 1 | 673656.44 |
| mammals | Tapirus_indicus | Tapirus_indicus | 1 | 296250 | 1 | 89491.88 |
| mammals | Tapirus_pinchaque | Tapirus_pinchaque | 1 | 148949.8 | 1 | 49536.77 |
| mammals | Tapirus_terrestris | Tapirus_terrestris | 1 | 207500.9 | 1 | 13052361.74 |
| mammals | Tarsipes_rostratus | Tarsipes_rostratus | 0 | 9 | 2.95 | 236675.5 |
| mammals | Tarsius_dentatus | Tarsius_dentatus | 1 | 107 | 1 | 35043.15 |
| mammals | Tarsius_pelengensis | Tarsius_pelengensis | 1 | 170 | 1.01 | 2615.02 |
| mammals | Tarsius_pumilus | Tarsius_pumilus | 1 | 27.5 | 1 | 395.74 |
| mammals | Tarsius_sangirensis | Tarsius_sangirensis | 1 | 171.3 | 1.01 | 556.37 |
| mammals | Tarsius_tarsier | Tarsius_tarsier | 1 | 200 | 1 | 37585.8 |
| mammals | Tarsomys_apoensis | Tarsomys_apoensis | 0 | 75 | 4 | 4440.29 |
| mammals | Tatera_indica | Tatera_indica | 0 | 166.3 | 5.2 | 4786124.73 |
| mammals | Taterillus_congicus | Taterillus_congicus | 0 | 54.5 | 3.63 | 1317164.85 |
| mammals | Taterillus_emini | Taterillus_emini | 0 | 52 | 3.69 | 3860531.19 |
| mammals | Taterillus_gracilis | Taterillus_gracilis | 0 | 48 | 4.12 | 2210732.89 |
| mammals | Taterillus_lacustris | Taterillus_lacustris | 0 | 41.5 | 3.63 | 39538.93 |

| | | | | | | |
|---|---|---|---|---|---|---|
| mammals | Taterillus_petteri | Taterillus_petteri | 0 | 50.8 | 3 | 178975.98 |
| mammals | Taterillus_pygargus | Taterillus_pygargus | 0 | 60.2 | 3.97 | 365303.22 |
| mammals | Taxidea_taxus | Taxidea_taxus | 0 | 7107.6 | 2.88 | 8851818.1 |
| mammals | Tayassu_pecari | Tayassu_pecari | 1 | 32233.7 | 2 | 13750306.26 |
| mammals | Tenrec_ecaudatus | Tenrec_ecaudatus | 0 | 868 | 16.89 | 590834.3 |
| mammals | Tetracerus_quadricornis | Tetracerus_quadricornis | 1 | 19000 | 1.62 | 2063086.54 |
| mammals | Thallomys_loringi | Thallomys_loringi | 0 | 80 | 4 | 56457.55 |
| mammals | Thallomys_nigricauda | Thallomys_nigricauda | 0 | 124.7 | 1.8 | 1214564.26 |
| mammals | Thallomys_paedulcus | Thallomys_paedulcus | 0 | 80.5 | 3.15 | 5096542.57 |
| mammals | Thalpomys_lasiotis | Thalpomys_lasiotis | 0 | 24 | 2.5 | 555211.18 |
| mammals | Thamnomys_kempi | Thamnomys_kempi | 1 | 75.1 | 1.33 | 5110.79 |
| mammals | Thamnomys_kuru | Grammomys_kuru | 0 | 52 | 1 | 2284482.44 |
| mammals | Thamnomys_venustus | Thamnomys_venustus | 0 | 61.9 | 1.33 | 10760.92 |
| mammals | Thaptomys_nigrita | Thaptomys_nigrita | 0 | 19.9 | 4 | 1057629.59 |
| mammals | Theropithecus_gelada | Theropithecus_gelada | 0 | 17049.8 | 1 | 104365.32 |
| mammals | Thomasomys_aureus | Thomasomys_aureus | 0 | 88 | 2.5 | 557724.62 |
| mammals | Thomasomys_baeops | Thomasomys_baeops | 0 | 36 | 3 | 86495.21 |
| mammals | Thomasomys_cinereus | Thomasomys_cinereus | 0 | 77 | 2.5 | 26409.05 |
| mammals | Thomasomys_paramorum | Thomasomys_paramorum | 0 | 21.5 | 1 | 29904.44 |
| mammals | Thoopterus_nigrescens | Thoopterus_nigrescens | 0 | 60.4 | 0.98 | 182866.82 |
| mammals | Thrichomys_apereoides | Thrichomys_apereoides | 0 | 339 | 3.17 | 1168838.93 |
| mammals | Thrichomys_inermis | Thrichomys_inermis | 0 | 298.8 | 3.15 | 115402.88 |
| mammals | Thrichomys_pachyurus | Thrichomys_pachyurus | 0 | 266 | 3.12 | 166428.89 |
| mammals | Thryonomys_gregorianus | Thryonomys_gregorianus | 0 | 2707.1 | 2.91 | 2375922.35 |
| mammals | Thryonomys_swinderianus | Thryonomys_swinderianus | 0 | 3180 | 3.88 | 7541449.63 |
| mammals | Thylamys_elegans | Thylamys_elegans | 0 | 28.9 | 11.8 | 142691.22 |
| mammals | Thylamys_pallidior | Thylamys_pallidior | 0 | 14.9 | 9.5 | 1493250.67 |
| mammals | Thylogale_billardierii | Thylogale_billardierii | 0 | 5450 | 1 | 66448.24 |
| mammals | Thylogale_brownii | Thylogale_browni | 1 | 5478 | 1.01 | 269510.34 |
| mammals | Thylogale_brunii | Thylogale_brunii | 1 | 3620 | 1 | 108853.52 |
| mammals | Thylogale_calabyi | Thylogale_calabyi | 1 | 4326.1 | 1 | 1002.37 |
| mammals | Thylogale_stigmatica | Thylogale_stigmatica | 0 | 4600 | 1 | 221199.78 |
| mammals | Thylogale_thetis | Thylogale_thetis | 0 | 5400 | 1 | 73431.48 |
| mammals | Thyroptera_tricolor | Thyroptera_tricolor | 0 | 4.4 | 0.98 | 8555467.31 |

| | | | | | | |
|---|---|---|---|---|---|---|
| mammals | Tlacuatzin_canescens | Tlacuatzin_canescens | 0 | 60 | 9.74 | 479508.39 |
| mammals | Tokudaia_muenninki | Tokudaia_muenninki | 1 | 177.4 | 7 | 240.57 |
| mammals | Tolypeutes_matacus | Tolypeutes_matacus | 0 | 1200 | 1 | 1198272.04 |
| mammals | Tolypeutes_tricinctus | Tolypeutes_tricinctus | 1 | 1487 | 0.93 | 932819.6 |
| mammals | Tomopeas_ravum | Tomopeas_ravus | 1 | 5.6 | 1 | 112800.74 |
| mammals | Tonatia_saurophila | Tonatia_saurophila | 0 | 30 | 0.98 | 6509470.68 |
| mammals | Trachops_cirrhosus | Trachops_cirrhosus | 0 | 37.7 | 0.99 | 12642871.51 |
| mammals | Trachypithecus_auratus | Trachypithecus_auratus | 1 | 7100 | 1 | 120357.24 |
| mammals | Trachypithecus_barbei | Trachypithecus_barbei | 1 | 7500 | 1 | 3817.01 |
| mammals | Trachypithecus_cristatus | Trachypithecus_cristatus | 1 | 5250 | 1 | 1213231.28 |
| mammals | Trachypithecus_delacouri | Trachypithecus_delacouri | 1 | 8899.9 | 1.5 | 6008.4 |
| mammals | Trachypithecus_francoisi | Trachypithecus_francoisi | 1 | 7325 | 1 | 248808.28 |
| mammals | Trachypithecus_geei | Trachypithecus_geei | 1 | 8350 | 1.01 | 7413.46 |
| mammals | Trachypithecus_germaini | Trachypithecus_germaini | 1 | 7179 | 1 | 261220.84 |
| mammals | Trachypithecus_hatinhensis | Trachypithecus_hatinhensis | 1 | 6323.1 | 1 | 18902.38 |
| mammals | Trachypithecus_obscurus | Trachypithecus_obscurus | 1 | 7000 | 1 | 259473.53 |
| mammals | Trachypithecus_phayrei | Trachypithecus_phayrei | 1 | 8400 | 1 | 512798.89 |
| mammals | Trachypithecus_pileatus | Trachypithecus_pileatus | 1 | 11450 | 1 | 305059.05 |
| mammals | Trachypithecus_poliocephalus | Trachypithecus_poliocephalus | 1 | 7750 | 1 | 109.88 |
| mammals | Tragelaphus_angasii | Tragelaphus_angasii | 0 | 86600.4 | 1 | 197902.7 |
| mammals | Tragelaphus_buxtoni | Tragelaphus_buxtoni | 1 | 215000.8 | 1 | 6875.04 |
| mammals | Tragelaphus_derbianus | Tragelaphus_derbianus | 1 | 680000 | 1.1 | 392963.4 |
| mammals | Tragelaphus_eurycerus | Tragelaphus_eurycerus | 0 | 329003.1 | 1 | 2217282.34 |
| mammals | Tragelaphus_imberbis | Tragelaphus_imberbis | 0 | 81575.5 | 1 | 1488947.51 |
| mammals | Tragelaphus_oryx | Tragelaphus_oryx | 0 | 569993.6 | 1 | 6776785.86 |
| mammals | Tragelaphus_scriptus | Tragelaphus_scriptus | 0 | 43250.4 | 1 | 12064054.46 |
| mammals | Tragelaphus_spekii | Tragelaphus_spekii | 0 | 77999.2 | 1 | 4392202.52 |
| mammals | Tragelaphus_strepsiceros | Tragelaphus_strepsiceros | 0 | 213501 | 1 | 6085607.88 |
| mammals | Tragulus_kanchil | Tragulus_kanchil | 0 | 2000 | 1 | 2303447.46 |
| mammals | Tragulus_napu | Tragulus_napu | 0 | 4000 | 0.98 | 1402060.13 |
| mammals | Tragulus_nigricans | Tragulus_nigricans | 1 | 2473.8 | 1 | 506.71 |
| mammals | Transandinomys_bolivaris | Transandinomys_bolivaris | 0 | 60.5 | 3.23 | 271356.28 |
| mammals | Transandinomys_talamancae | Transandinomys_talamancae | 0 | 55 | 3.9 | 510964.25 |
| mammals | Tremarctos_ornatus | Tremarctos_ornatus | 1 | 140000.6 | 1.47 | 218405.24 |

| | | | | | | |
|---|---|---|---|---|---|---|
| mammals | Triaenops_menamena | Triaenops_rufus | 0 | 10.5 | 1 | 284167.07 |
| mammals | Triaenops_persicus | Triaenops_persicus | 0 | 12.6 | 1 | 1163737.95 |
| mammals | Trichosurus_caninus | Trichosurus_caninus | 0 | 3500 | 1.02 | 148314.2 |
| mammals | Trichosurus_cunninghami | Trichosurus_cunninghami | 0 | 3270 | 1 | 119403.45 |
| mammals | Trichosurus_vulpecula | Trichosurus_vulpecula | 0 | 2650 | 1.01 | 2704232.99 |
| mammals | Trinomys_albispinus | Trinomys_albispinus | 0 | 285 | 3 | 70209.4 |
| mammals | Trinomys_dimidiatus | Trinomys_dimidiatus | 0 | 167.6 | 2 | 63989.21 |
| mammals | Trinomys_eliasi | Trinomys_eliasi | 0 | 167.9 | 3 | 4162.41 |
| mammals | Trinomys_iheringi | Trinomys_iheringi | 0 | 203.5 | 0.97 | 702152.55 |
| mammals | Trinomys_setosus | Trinomys_setosus | 0 | 285 | 2 | 414903.9 |
| mammals | Trinycteris_nicefori | Trinycteris_nicefori | 0 | 8.2 | 1 | 4253378.91 |
| mammals | Trogopterus_xanthipes | Trogopterus_xanthipes | 0 | 744.9 | 1.64 | 1062333.59 |
| mammals | Tscherskia_triton | Tscherskia_triton | 0 | 166.5 | 4.24 | 3140943.03 |
| mammals | Tupaia_belangeri | Tupaia_belangeri | 0 | 200 | 2 | 2888122.74 |
| mammals | Tupaia_chrysogaster | Tupaia_chrysogaster | 1 | 177.7 | 2.22 | 6002.77 |
| mammals | Tupaia_everetti | Urogale_everetti | 0 | 350 | 1.77 | 95385.36 |
| mammals | Tupaia_glis | Tupaia_glis | 0 | 169.8 | 2.22 | 190363.12 |
| mammals | Tupaia_gracilis | Tupaia_gracilis | 0 | 74.6 | 2 | 643734.65 |
| mammals | Tupaia_javanica | Tupaia_javanica | 0 | 105 | 2.95 | 248043.79 |
| mammals | Tupaia_longipes | Tupaia_longipes | 0 | 196 | 1.8 | 161229.93 |
| mammals | Tupaia_minor | Tupaia_minor | 0 | 64.3 | 1.96 | 1264126.17 |
| mammals | Tupaia_montana | Tupaia_montana | 0 | 199 | 1.88 | 179713.65 |
| mammals | Tupaia_nicobarica | Tupaia_nicobarica | 1 | 170.3 | 0.96 | 1108.48 |
| mammals | Tupaia_tana | Tupaia_tana | 0 | 197.8 | 2.35 | 999970.62 |
| mammals | Tylomys_mirae | Tylomys_mirae | 0 | 183.6 | 2 | 125758.52 |
| mammals | Tylomys_nudicaudus | Tylomys_nudicaudus | 0 | 100 | 2.44 | 489186.85 |
| mammals | Tylonycteris_pachypus | Tylonycteris_pachypus | 0 | 3.6 | 1.92 | 4182645.86 |
| mammals | Tylonycteris_robustula | Tylonycteris_robustula | 0 | 7.6 | 2 | 2721209.15 |
| mammals | Typhlomys_cinereus | Typhlomys_cinereus | 0 | 24.5 | 2.5 | 1030421.04 |
| mammals | Typomys_planifrons | Hybomys_planifrons | 0 | 49.6 | 2.4 | 239879.55 |
| mammals | Typomys_trivirgatus | Hybomys_trivirgatus | 0 | 56.9 | 2.72 | 387198.51 |
| mammals | Uranomys_ruddi | Uranomys_ruddi | 0 | 40.4 | 3.95 | 4057097.01 |
| mammals | Urocitellus_armatus | Urocitellus_armatus | 0 | 313 | 5.42 | 176257.8 |
| mammals | Urocitellus_beldingi | Urocitellus_beldingi | 0 | 280.5 | 5.64 | 337683.75 |

| | | | | | | |
|---|---|---|---|---|---|---|
| mammals | Urocitellus_brunneus | Urocitellus_brunneus | 1 | 300 | 6.5 | 6763.23 |
| mammals | Urocitellus_canus | Urocitellus_canus | 0 | 154 | 8 | 217761.2 |
| mammals | Urocitellus_columbianus | Urocitellus_columbianus | 0 | 493 | 3.54 | 532364.64 |
| mammals | Urocitellus_elegans | Urocitellus_elegans | 0 | 453.6 | 5.83 | 282690.69 |
| mammals | Urocitellus_mollis | Urocitellus_mollis | 0 | 165.4 | 8.2 | 416260.77 |
| mammals | Urocitellus_parryii | Urocitellus_parryii | 0 | 760 | 6.4 | 5346615.24 |
| mammals | Urocitellus_richardsonii | Urocitellus_richardsonii | 0 | 406 | 7.41 | 900811.96 |
| mammals | Urocitellus_townsendii | Urocitellus_townsendii | 1 | 213 | 8.46 | 13910.87 |
| mammals | Urocitellus_undulatus | Urocitellus_undulatus | 0 | 840 | 7.77 | 2310791.44 |
| mammals | Urocitellus_washingtoni | Urocitellus_washingtoni | 0 | 207.5 | 7.88 | 17294.85 |
| mammals | Urocricetus_alticola | Cricetulus_alticola | 0 | 35 | 7.5 | 140657.57 |
| mammals | Urocricetus_kamensis | Cricetulus_kamensis | 0 | 30 | 7.5 | 163548.69 |
| mammals | Urocyon_cinereoargenteus | Urocyon_cinereoargenteus | 0 | 3833.7 | 3.8 | 9037640.21 |
| mammals | Urocyon_littoralis | Urocyon_littoralis | 0 | 1896 | 2.28 | 911.16 |
| mammals | Uroderma_bilobatum | Uroderma_bilobatum | 0 | 16.7 | 1 | 10739162.99 |
| mammals | Uroderma_magnirostrum | Uroderma_magnirostrum | 0 | 16.9 | 0.98 | 10831975.7 |
| mammals | Uromys_anak | Uromys_anak | 0 | 664.5 | 4.3 | 207596.3 |
| mammals | Uromys_caudimaculatus | Uromys_caudimaculatus | 0 | 606.8 | 1.61 | 880888.11 |
| mammals | Uromys_hadrourus | Uromys_hadrourus | 0 | 194 | 3 | 635.19 |
| mammals | Urotrichus_talpoides | Urotrichus_talpoides | 0 | 45 | 3.5 | 286809.07 |
| mammals | Ursus_americanus | Ursus_americanus | 0 | 99949.4 | 2.43 | 16113743.93 |
| mammals | Ursus_arctos | Ursus_arctos | 0 | 180520.4 | 2.26 | 36100052.36 |
| mammals | Ursus_maritimus | Ursus_maritimus | 1 | 388750.4 | 1.66 | 22605413.56 |
| mammals | Ursus_thibetanus | Ursus_thibetanus | 1 | 77500 | 1.58 | 8529242.11 |
| mammals | Urva_auropunctata | Herpestes_auropunctatus | 0 | 483.5 | 2.3 | 4171313.51 |
| mammals | Urva_brachyura | Herpestes_brachyurus | 0 | 1853.5 | 3.49 | 1271946.62 |
| mammals | Urva_edwardsii | Herpestes_edwardsii | 0 | 1324.2 | 2.3 | 4314164.45 |
| mammals | Urva_fusca | Herpestes_fuscus | 0 | 2700 | 2.37 | 42989.01 |
| mammals | Urva_javanica | Herpestes_javanicus | 0 | 750 | 2.21 | 2125157.91 |
| mammals | Urva_semitorquata | Herpestes_semitorquatus | 0 | 3500 | 3.49 | 1174536.96 |
| mammals | Urva_smithii | Herpestes_smithii | 0 | 1782 | 2.5 | 1895779.2 |
| mammals | Urva_urva | Herpestes_urva | 0 | 1863.2 | 3 | 4060874.84 |
| mammals | Urva_vitticollis | Herpestes_vitticollis | 0 | 2575.6 | 2.67 | 152078.88 |
| mammals | Vampyressa_melissa | Vampyressa_melissa | 1 | 16.3 | 0.98 | 445446.34 |

| | | | | | | |
|---|---|---|---|---|---|---|
| mammals | Vampyressa_thyone | Vampyressa_thyone | 0 | 8.5 | 0.98 | 4543755.92 |
| mammals | Vampyriscus_bidens | Vampyriscus_bidens | 0 | 11.6 | 0.98 | 6731906.15 |
| mammals | Vampyriscus_nymphaeus | Vampyriscus_nymphaea | 0 | 11.8 | 0.99 | 248097.11 |
| mammals | Vampyrodes_caraccioli | Vampyrodes_caraccioli | 0 | 34.7 | 0.98 | 7326122.46 |
| mammals | Vampyrodes_major | Vampyrodes_major | 0 | 38.5 | 1 | 880555.25 |
| mammals | Vampyrum_spectrum | Vampyrum_spectrum | 0 | 167.7 | 1 | 5277255.15 |
| mammals | Vandeleuria_nolthenii | Vandeleuria_nolthenii | 1 | 56.6 | 4 | 316.66 |
| mammals | Vandeleuria_oleraceus | Vandeleuria_oleracea | 0 | 10 | 3.24 | 3789829.9 |
| mammals | Vansonia_rueppellii | Pipistrellus_rueppellii | 0 | 6.9 | 1.97 | 2970000.58 |
| mammals | Varecia_rubra | Varecia_rubra | 1 | 3450 | 2.2 | 5271.84 |
| mammals | Varecia_variegata | Varecia_variegata | 1 | 3350 | 2.18 | 20717.2 |
| mammals | Vespadelus_baverstocki | Vespadelus_baverstocki | 0 | 5 | 1 | 2261700.4 |
| mammals | Vespadelus_caurinus | Vespadelus_caurinus | 0 | 4 | 1.4 | 606796.83 |
| mammals | Vespadelus_darlingtoni | Vespadelus_darlingtoni | 0 | 6 | 1 | 503722.13 |
| mammals | Vespadelus_douglasorum | Vespadelus_douglasorum | 0 | 5 | 1 | 132951.75 |
| mammals | Vespadelus_finlaysoni | Vespadelus_finlaysoni | 0 | 5 | 1.5 | 3435392.14 |
| mammals | Vespadelus_pumilus | Vespadelus_pumilus | 0 | 4.4 | 1.21 | 148605.62 |
| mammals | Vespadelus_regulus | Vespadelus_regulus | 0 | 5.4 | 1 | 1241504.74 |
| mammals | Vespadelus_troughtoni | Vespadelus_troughtoni | 0 | 5.5 | 1.5 | 895634.15 |
| mammals | Vespadelus_vulturnus | Vespadelus_vulturnus | 0 | 4.1 | 1 | 1401978.48 |
| mammals | Vespertilio_murinus | Vespertilio_murinus | 0 | 18.7 | 1.5 | 15761265.22 |
| mammals | Vespertilio_sinensis | Vespertilio_sinensis | 0 | 24.5 | 2 | 6003456.6 |
| mammals | Viverra_tangalunga | Viverra_tangalunga | 0 | 6885 | 1.92 | 1602751.77 |
| mammals | Viverra_zibetha | Viverra_zibetha | 0 | 9500 | 2.89 | 5069770.83 |
| mammals | Viverricula_indica | Viverricula_indica | 0 | 2908.4 | 3.78 | 8473961.85 |
| mammals | Voalavo_antsahabensis | Voalavo_antsahabensis | 1 | 21.8 | 2 | 299.49 |
| mammals | Voalavo_gymnocaudus | Voalavo_gymnocaudus | 0 | 21.8 | 2 | 1144.09 |
| mammals | Vombatus_ursinus | Vombatus_ursinus | 0 | 26000 | 1 | 310428.62 |
| mammals | Vormela_peregusna | Vormela_peregusna | 1 | 450.4 | 5.42 | 7929465.09 |
| mammals | Vulpes_bengalensis | Vulpes_bengalensis | 0 | 2726 | 3.49 | 3129366.45 |
| mammals | Vulpes_cana | Vulpes_cana | 0 | 936.9 | 2 | 3142580.11 |
| mammals | Vulpes_chama | Vulpes_chama | 0 | 2955 | 2.69 | 2126479.34 |
| mammals | Vulpes_corsac | Vulpes_corsac | 0 | 2400 | 5.31 | 8976804.65 |
| mammals | Vulpes_ferrilata | Vulpes_ferrilata | 0 | 5000 | 4.1 | 2231673.47 |

| | | | | | | |
|---|---|---|---|---|---|---|
| mammals | Vulpes_lagopus | Vulpes_lagopus | 0 | 4867.7 | 7.88 | 10726429 |
| mammals | Vulpes_macrotis | Vulpes_macrotis | 0 | 2050 | 4.47 | 1611407.34 |
| mammals | Vulpes_pallida | Vulpes_pallida | 0 | 2800 | 4.74 | 4152381.02 |
| mammals | Vulpes_rueppellii | Vulpes_rueppellii | 0 | 2865 | 2.56 | 12646617.27 |
| mammals | Vulpes_velox | Vulpes_velox | 0 | 2197.5 | 4.34 | 1550926.19 |
| mammals | Vulpes_vulpes | Vulpes_vulpes | 0 | 5318.2 | 4.66 | 72402469.18 |
| mammals | Vulpes_zerda | Vulpes_zerda | 0 | 1100 | 2.36 | 8675207.95 |
| mammals | Wallabia_bicolor | Wallabia_bicolor | 0 | 15000 | 1.03 | 1212921.52 |
| mammals | Wiedomys_pyrrhorinos | Wiedomys_pyrrhorhinos | 0 | 46.7 | 3.8 | 694703.09 |
| mammals | Wyulda_squamicaudata | Wyulda_squamicaudata | 0 | 1675 | 1.01 | 82875.64 |
| mammals | Xanthonycticebus_pygmaeus | Nycticebus_pygmaeus | 1 | 900 | 1.44 | 481890.25 |
| mammals | Xenogale_naso | Herpestes_naso | 0 | 3000 | 3 | 2656378.52 |
| mammals | Xenomys_nelsoni | Xenomys_nelsoni | 1 | 130 | 1.6 | 9111.06 |
| mammals | Xenuromys_barbatus | Xenuromys_barbatus | 0 | 1000 | 1.76 | 591368.22 |
| mammals | Xeromys_myoides | Xeromys_myoides | 1 | 41 | 2 | 12793.23 |
| mammals | Xerospermophilus_mohavensis | Xerospermophilus_mohavensis | 0 | 190 | 6.41 | 22711.46 |
| mammals | Xerospermophilus_perotensis | Xerospermophilus_perotensis | 1 | 140 | 6.07 | 2655.85 |
| mammals | Xerospermophilus_spilosoma | Xerospermophilus_spilosoma | 0 | 89 | 6.86 | 1712244.66 |
| mammals | Xerospermophilus_tereticaudus | Xerospermophilus_tereticaudus | 0 | 156.5 | 6.09 | 309353.22 |
| mammals | Xerus_rutilus | Xerus_rutilus | 0 | 317.5 | 1.96 | 2086690.87 |
| mammals | Zaedyus_pichiy | Zaedyus_pichiy | 0 | 1380 | 2 | 1362776.27 |
| mammals | Zaglossus_bartoni | Zaglossus_bartoni | 1 | 9413.8 | 1 | 174289.77 |
| mammals | Zaglossus_bruijnii | Zaglossus_bruijnii | 1 | 7800 | 1 | 91613.59 |
| mammals | Zapus_hudsonius | Zapus_hudsonius | 0 | 17.1 | 5.36 | 8737824.37 |
| mammals | Zapus_princeps | Zapus_princeps | 0 | 29 | 5.04 | 3278014.04 |
| mammals | Zapus_trinotatus | Zapus_trinotatus | 0 | 27.5 | 5.38 | 162624.79 |
| mammals | Zelotomys_hildegardeae | Zelotomys_hildegardeae | 0 | 60 | 5.75 | 3051583.27 |
| mammals | Zelotomys_woosnami | Zelotomys_woosnami | 0 | 54.1 | 4.91 | 674859.92 |
| mammals | Zygodontomys_brevicauda | Zygodontomys_brevicauda | 0 | 52.2 | 4.25 | 1694720.07 |
| mammals | Zygodontomys_brunneus | Zygodontomys_brunneus | 0 | 75.6 | 4.28 | 75737.7 |
| mammals | Zyzomys_argurus | Zyzomys_argurus | 0 | 36 | 2.69 | 1188571.35 |
| mammals | Zyzomys_maini | Zyzomys_maini | 1 | 94 | 2.5 | 21158.55 |
| mammals | Zyzomys_pedunculatus | Zyzomys_pedunculatus | 1 | 100 | 2.5 | 7110.1 |
| mammals | Zyzomys_woodwardi | Zyzomys_woodwardi | 0 | 136 | 2.23 | 35599.28 |

| | | | | | | |
|---|---|---|---|---|---|---|
| birds | Abeillia_abeillei | Abeillia_abeillei | 0 | 2.7 | 2 | 144610.4 |
| birds | Abroscopus_albogularis | Abroscopus_albogularis | 0 | 4.8 | 3.9 | 1704650.6 |
| birds | Abroscopus_schisticeps | Abroscopus_schisticeps | 0 | 4.7 | 4.5 | 909217.05 |
| birds | Abroscopus_superciliaris | Abroscopus_superciliaris | 0 | 6.5 | 3.2 | 2493437.47 |
| birds | Acanthagenys_rufogularis | Acanthagenys_rufogularis | 0 | 47.8 | 2.4 | 5976565.2 |
| birds | Acanthis_flammea | Carduelis_flammea | 0 | 13 | 4.9 | 42235480.52 |
| birds | Acanthiza_chrysorrhoa | Acanthiza_chrysorrhoa | 0 | 9.3 | 3.5 | 4590956.33 |
| birds | Acanthiza_ewingii | Acanthiza_ewingii | 0 | 6.9 | 3.5 | 68050.51 |
| birds | Acanthiza_inornata | Acanthiza_inornata | 0 | 7 | 4 | 137406.16 |
| birds | Acanthiza_iredalei | Acanthiza_iredalei | 0 | 6 | 3 | 987131.55 |
| birds | Acanthiza_katherina | Acanthiza_katherina | 1 | 7.2 | 2 | 6563.03 |
| birds | Acanthiza_lineata | Acanthiza_lineata | 0 | 6.9 | 3 | 1043817.35 |
| birds | Acanthiza_murina | Acanthiza_murina | 0 | 8.4 | 2 | 84339.38 |
| birds | Acanthiza_nana | Acanthiza_nana | 0 | 6.2 | 3 | 2021416.13 |
| birds | Acanthiza_pusilla | Acanthiza_pusilla | 0 | 7.6 | 2.4 | 1131818.69 |
| birds | Acanthiza_reguloides | Acanthiza_reguloides | 0 | 7.7 | 3.9 | 1843422.3 |
| birds | Acanthiza_robustirostris | Acanthiza_robustirostris | 0 | 6.1 | 2.4 | 2061848.42 |
| birds | Acanthiza_uropygialis | Acanthiza_uropygialis | 0 | 6.4 | 3 | 4788831.67 |
| birds | Acanthoptila_nipalensis | Turdoides_nipalensis | 0 | 61 | 3.5 | 28765.27 |
| birds | Acanthorhynchus_superciliosus | Acanthorhynchus_superciliosus | 0 | 9.7 | 1.4 | 182218.54 |
| birds | Acanthorhynchus_tenuirostris | Acanthorhynchus_tenuirostris | 0 | 10.4 | 2 | 763502.74 |
| birds | Acanthornis_magna | Acanthornis_magna | 0 | 10.2 | 3.5 | 45032.53 |
| birds | Accipiter_badius | Accipiter_badius | 0 | 131.2 | 2.8 | 22853794.98 |
| birds | Accipiter_bicolor | Accipiter_bicolor | 0 | 287.5 | 1.7 | 14309701.27 |
| birds | Accipiter_brevipes | Accipiter_brevipes | 0 | 186.5 | 3.9 | 7225515.31 |
| birds | Accipiter_cirrocephalus | Accipiter_cirrocephalus | 0 | 165.7 | 3.5 | 5488344.15 |
| birds | Accipiter_cooperii | Accipiter_cooperii | 0 | 429.7 | 4.2 | 10487652.99 |
| birds | Accipiter_erythropus | Accipiter_erythropus | 0 | 111.2 | 2 | 3904812.09 |
| birds | Accipiter_fasciatus | Accipiter_fasciatus | 0 | 358.7 | 3 | 8180492.01 |
| birds | Accipiter_francesiae | Accipiter_francesiae | 0 | 136.8 | 3 | 519468.81 |
| birds | Accipiter_gentilis | Accipiter_gentilis | 0 | 866 | 3.5 | 36029138.68 |
| birds | Accipiter_gularis | Accipiter_gularis | 0 | 120.7 | 3.2 | 10101618.37 |
| birds | Accipiter_gundlachi | Accipiter_gundlachi | 1 | 675 | 3.5 | 106868.73 |
| birds | Accipiter_haplochrous | Accipiter_haplochrous | 0 | 198.4 | 3 | 16389.59 |

| | | | | | | |
|---|---|---|---|---|---|---|
| birds | Accipiter_madagascariensis | Accipiter_madagascariensis | 0 | 180.4 | 3 | 476517.51 |
| birds | Accipiter_melanoleucus | Accipiter_melanoleucus | 0 | 699 | 2.8 | 9071924.49 |
| birds | Accipiter_meyerianus | Accipiter_meyerianus | 0 | 763.5 | 3 | 269144.41 |
| birds | Accipiter_minullus | Accipiter_minullus | 0 | 87.4 | 2 | 6794381.58 |
| birds | Accipiter_nisus | Accipiter_nisus | 0 | 220.8 | 4.2 | 36695930.3 |
| birds | Accipiter_novaehollandiae | Accipiter_novaehollandiae | 0 | 326.1 | 2.4 | 1807067.3 |
| birds | Accipiter_ovampensis | Accipiter_ovampensis | 0 | 195 | 3 | 7295779.86 |
| birds | Accipiter_poliocephalus | Accipiter_poliocephalus | 0 | 214.8 | 1 | 671500.51 |
| birds | Accipiter_rufitorques | Accipiter_rufitorques | 0 | 209 | 2.4 | 17581.07 |
| birds | Accipiter_rufiventris | Accipiter_rufiventris | 0 | 198 | 3 | 756057.21 |
| birds | Accipiter_soloensis | Accipiter_soloensis | 0 | 164.8 | 3.5 | 4004394.76 |
| birds | Accipiter_striatus | Accipiter_striatus | 0 | 130.6 | 4.5 | 21636790.05 |
| birds | Accipiter_superciliosus | Accipiter_superciliosus | 0 | 96.4 | 1.7 | 8560355.57 |
| birds | Accipiter_tachiro | Accipiter_tachiro | 0 | 290.2 | 2.4 | 5174215.21 |
| birds | Accipiter_trivirgatus | Accipiter_trivirgatus | 0 | 265 | 2 | 3366430.69 |
| birds | Aceros_nipalensis | Aceros_nipalensis | 1 | 2385 | 1.4 | 484788.19 |
| birds | Acridotheres_cristatellus | Acridotheres_cristatellus | 0 | 116.3 | 5.3 | 3225824.49 |
| birds | Acridotheres_fuscus | Acridotheres_fuscus | 0 | 82.8 | 4.2 | 2422718.95 |
| birds | Acridotheres_ginginianus | Acridotheres_ginginianus | 0 | 72 | 4.9 | 1641344.88 |
| birds | Acridotheres_grandis | Acridotheres_grandis | 0 | 98.9 | 4.9 | 1707140.6 |
| birds | Acridotheres_melanopterus | Sturnus_melanopterus | 1 | 89 | 3.5 | 136022.66 |
| birds | Acridotheres_tristis | Acridotheres_tristis | 0 | 116.4 | 3.5 | 10019063.91 |
| birds | Acritillas_indica | Iole_indica | 0 | 30.6 | 2.4 | 219041.39 |
| birds | Acrocephalus_agricola | Acrocephalus_agricola | 0 | 9.6 | 4.2 | 7862832.83 |
| birds | Acrocephalus_arundinaceus | Acrocephalus_arundinaceus | 0 | 30 | 4.2 | 36259434.41 |
| birds | Acrocephalus_australis | Acrocephalus_australis | 0 | 17.7 | 2.4 | 6368774.96 |
| birds | Acrocephalus_bistrigiceps | Acrocephalus_bistrigiceps | 0 | 8.7 | 4.9 | 6253044.54 |
| birds | Acrocephalus_concinens | Acrocephalus_concinens | 0 | 8.3 | 3.5 | 4244812.47 |
| birds | Acrocephalus_dumetorum | Acrocephalus_dumetorum | 0 | 11.2 | 4.2 | 6977440.85 |
| birds | Acrocephalus_gracilirostris | Acrocephalus_gracilirostris | 0 | 14.4 | 2.4 | 4809980 |
| birds | Acrocephalus_melanopogon | Acrocephalus_melanopogon | 0 | 10.9 | 3.9 | 6177192.64 |
| birds | Acrocephalus_newtoni | Acrocephalus_newtoni | 0 | 17 | 3 | 591120.51 |
| birds | Acrocephalus_paludicola | Acrocephalus_paludicola | 1 | 11.6 | 4.9 | 9180072.4 |
| birds | Acrocephalus_palustris | Acrocephalus_palustris | 0 | 11.5 | 3.9 | 18399649.06 |

| | | | | | | |
|---|---|---|---|---|---|---|
| birds | Acrocephalus_rufescens | Acrocephalus_rufescens | 0 | 22.7 | 2.4 | 1400095.07 |
| birds | Acrocephalus_schoenobaenus | Acrocephalus_schoenobaenus | 0 | 11.8 | 5.5 | 39149311.53 |
| birds | Acrocephalus_scirpaceus | Acrocephalus_scirpaceus | 0 | 12.3 | 3.9 | 41060299.19 |
| birds | Acrocephalus_stentoreus | Acrocephalus_stentoreus | 0 | 24.2 | 4.2 | 8982425.75 |
| birds | Acryllium_vulturinum | Acryllium_vulturinum | 0 | 1330 | 14 | 1226666.24 |
| birds | Actenoides_concretus | Actenoides_concretus | 0 | 73.5 | 2 | 1364096.24 |
| birds | Actitis_hypoleucos | Actitis_hypoleucos | 0 | 48 | 4 | 61178024.78 |
| birds | Actitis_macularius | Actitis_macularius | 0 | 40.4 | 4 | 34169893.46 |
| birds | Actophilornis_africanus | Actophilornis_africanus | 0 | 189.1 | 4 | 16988448.66 |
| birds | Actophilornis_albinucha | Actophilornis_albinucha | 1 | 239 | 4 | 140285.77 |
| birds | Adelomyia_melanogenys | Adelomyia_melanogenys | 0 | 4.9 | 2 | 620219.31 |
| birds | Aechmophorus_clarkii | Aechmophorus_clarkii | 0 | 1090.9 | 3.5 | 3589069.35 |
| birds | Aechmophorus_occidentalis | Aechmophorus_occidentalis | 0 | 1137 | 3.5 | 6366407.79 |
| birds | Aegithalos_caudatus | Aegithalos_caudatus | 0 | 8.6 | 9.8 | 18425691.87 |
| birds | Aegithalos_concinnus | Aegithalos_concinnus | 0 | 6.1 | 4.2 | 2797505.09 |
| birds | Aegithalos_iouschistos | Aegithalos_iouschistos | 0 | 7 | 4.5 | 68201.14 |
| birds | Aegithalos_leucogenys | Aegithalos_leucogenys | 0 | 6.7 | 6.3 | 289366.03 |
| birds | Aegithalos_niveogularis | Aegithalos_niveogularis | 0 | 7.8 | 4 | 180594.85 |
| birds | Aegithina_lafresnayei | Aegithina_lafresnayei | 0 | 12 | 3 | 1675913.71 |
| birds | Aegithina_nigrolutea | Aegithina_nigrolutea | 0 | 11.8 | 2.4 | 714297.74 |
| birds | Aegithina_tiphia | Aegithina_tiphia | 0 | 12 | 2.8 | 6356008.52 |
| birds | Aegithina_viridissima | Aegithina_viridissima | 0 | 13.4 | 2 | 1109607.12 |
| birds | Aegolius_acadicus | Aegolius_acadicus | 0 | 100.7 | 5.9 | 10821011.01 |
| birds | Aegolius_funereus | Aegolius_funereus | 0 | 137.2 | 4.2 | 23807935.25 |
| birds | Aegolius_harrisii | Aegolius_harrisii | 0 | 120 | 3 | 1098550.77 |
| birds | Aegolius_ridgwayi | Aegolius_ridgwayi | 0 | 80 | 5.5 | 43992.75 |
| birds | Aegotheles_cristatus | Aegotheles_cristatus | 0 | 42.8 | 3.5 | 7702773.06 |
| birds | Aegypius_monachus | Aegypius_monachus | 0 | 9320.6 | 1 | 17944995.14 |
| birds | Aepypodius_arfakianus | Aepypodius_arfakianus | 0 | 1463.8 | 20 | 195287.34 |
| birds | Aerodramus_francicus | Collocalia_francica | 0 | 8.9 | 2 | 4418.34 |
| birds | Aerodramus_hirundinaceus | Collocalia_hirundinacea | 0 | 10.1 | 1 | 583622.07 |
| birds | Aerodramus_salangana | Collocalia_salangana | 0 | 11.3 | 1.4 | 483061.54 |
| birds | Aerodramus_spodiopygius | Collocalia_spodiopygia | 0 | 8 | 1.4 | 136466.16 |
| birds | Aerodramus_terraereginae | Collocalia_terraereginae | 0 | 12 | 1 | 85667.06 |

| | | | | | | |
|---|---|---|---|---|---|---|
| birds | Aerodramus_vanikorensis | Collocalia_vanikorensis | 0 | 10.7 | 1.4 | 1212501.77 |
| birds | Aeronautes_saxatalis | Aeronautes_saxatalis | 0 | 32.1 | 4.2 | 4032383.43 |
| birds | Aethomyias_papuensis | Sericornis_papuensis | 0 | 11.2 | 1 | 86586.53 |
| birds | Afropavo_congensis | Afropavo_congensis | 0 | 1273.7 | 2.4 | 697505.34 |
| birds | Afrotis_afra | Eupodotis_afra | 1 | 690 | 1.4 | 173167.3 |
| birds | Afrotis_afraoides | Eupodotis_afraoides | 0 | 745 | 1.4 | 1934353.37 |
| birds | Agamia_agami | Agamia_agami | 0 | 567 | 2.8 | 8217123.75 |
| birds | Agapornis_canus | Agapornis_canus | 0 | 30 | 4.2 | 415982.93 |
| birds | Agapornis_fischeri | Agapornis_fischeri | 0 | 48.3 | 4.9 | 162335.09 |
| birds | Agapornis_lilianae | Agapornis_lilianae | 0 | 37.3 | 4.9 | 211626.34 |
| birds | Agapornis_nigrigenis | Agapornis_nigrigenis | 1 | 40.5 | 4.9 | 17094.25 |
| birds | Agapornis_personatus | Agapornis_personatus | 0 | 52.4 | 4.9 | 233268.94 |
| birds | Agapornis_pullarius | Agapornis_pullarius | 0 | 39.3 | 3.9 | 2650930.74 |
| birds | Agapornis_roseicollis | Agapornis_roseicollis | 0 | 53.8 | 4.9 | 457223.49 |
| birds | Agapornis_taranta | Agapornis_taranta | 0 | 57.5 | 5 | 407732.6 |
| birds | Agelaius_assimilis | Agelaius_assimilis | 0 | 47.7 | 2.4 | 4077.17 |
| birds | Agelaius_humeralis | Agelaius_humeralis | 0 | 36.3 | 3.5 | 110595.49 |
| birds | Agelaius_phoeniceus | Agelaius_phoeniceus | 0 | 50.8 | 3 | 14132462.28 |
| birds | Agelaius_tricolor | Agelaius_tricolor | 1 | 57.9 | 4 | 113509.64 |
| birds | Agelaius_xanthomus | Agelaius_xanthomus | 1 | 38.4 | 3 | 8726.66 |
| birds | Agelasticus_cyanopus | Agelasticus_cyanopus | 0 | 37.2 | 3 | 2871356.08 |
| birds | Agelasticus_thilius | Agelasticus_thilius | 0 | 31.5 | 3 | 2816597.82 |
| birds | Aglaeactis_cupripennis | Aglaeactis_cupripennis | 0 | 7.6 | 2 | 224852.99 |
| birds | Aglaiocercus_coelestis | Aglaiocercus_coelestis | 0 | 5.6 | 2 | 29539.31 |
| birds | Aglaiocercus_kingii | Aglaiocercus_kingi | 0 | 4.8 | 2 | 267748.93 |
| birds | Agricola_infuscatus | Bradornis_infuscatus | 0 | 32.4 | 2.4 | 1446786.72 |
| birds | Agricola_pallidus | Bradornis_pallidus | 0 | 23.6 | 2.4 | 9110671.42 |
| birds | Agriornis_lividus | Agriornis_lividus | 0 | 99.1 | 2.8 | 592911.81 |
| birds | Agriornis_micropterus | Agriornis_micropterus | 0 | 67.8 | 3.5 | 2809307.82 |
| birds | Agriornis_montanus | Agriornis_montanus | 0 | 63.3 | 2.4 | 2168659.57 |
| birds | Agriornis_murinus | Agriornis_murinus | 0 | 29.9 | 3 | 2112940.19 |
| birds | Agropsar_philippensis | Sturnus_philippensis | 0 | 28.8 | 4.6 | 568180.2 |
| birds | Agropsar_sturninus | Sturnus_sturninus | 0 | 49.4 | 4.6 | 8519690.44 |
| birds | Ailuroedus_buccoides | Ailuroedus_buccoides | 0 | 138 | 1 | 374654.83 |

| | | | | | | |
|---|---|---|---|---|---|---|
| birds | Ailuroedus_crassirostris | Ailuroedus_crassirostris | 0 | 205.3 | 1.7 | 208835.98 |
| birds | Ailuroedus_melanotis | Ailuroedus_melanotis | 0 | 176.9 | 1.4 | 167335.29 |
| birds | Aimophila_rufescens | Aimophila_rufescens | 0 | 34.3 | 2.4 | 616683.25 |
| birds | Aimophila_ruficeps | Aimophila_ruficeps | 0 | 18.7 | 4 | 1184786.97 |
| birds | Aix_galericulata | Aix_galericulata | 0 | 567 | 10.4 | 3446284.16 |
| birds | Aix_sponsa | Aix_sponsa | 0 | 657.6 | 11.6 | 7693839.61 |
| birds | Akletos_goeldii | Myrmeciza_goeldii | 0 | 42 | 1.4 | 342175.6 |
| birds | Akletos_melanoceps | Myrmeciza_melanoceps | 0 | 38.5 | 2 | 1321301.01 |
| birds | Alaemon_alaudipes | Alaemon_alaudipes | 0 | 43.7 | 2.8 | 10471834.07 |
| birds | Alauda_arvensis | Alauda_arvensis | 0 | 37.3 | 2.6 | 37787455.66 |
| birds | Alauda_gulgula | Alauda_gulgula | 0 | 26.3 | 3.2 | 11900668.99 |
| birds | Alauda_leucoptera | Melanocorypha_leucoptera | 0 | 45.3 | 4.9 | 5089458.73 |
| birds | Alaudala_raytal | Calandrella_raytal | 0 | 18.5 | 2.4 | 1260034.4 |
| birds | Alaudala_rufescens | Calandrella_cheleensis | 0 | 24.1 | 3.5 | 2316407.93 |
| birds | Alcedo_atthis | Alcedo_atthis | 0 | 31.1 | 6.5 | 28202464.97 |
| birds | Alcedo_euryzona | Alcedo_euryzona | 1 | 43.1 | 3.9 | 125365.64 |
| birds | Alcedo_hercules | Alcedo_hercules | 0 | 62.5 | 4.9 | 727664.63 |
| birds | Alcedo_meninting | Alcedo_meninting | 0 | 20.4 | 4.6 | 3655192.84 |
| birds | Alcedo_quadribrachys | Alcedo_quadribrachys | 0 | 34.9 | 4.9 | 4674328.28 |
| birds | Alcedo_semitorquata | Alcedo_semitorquata | 0 | 47.3 | 3.5 | 3463736.2 |
| birds | Alcurus_striatus | Pycnonotus_striatus | 0 | 52 | 3 | 1082921.86 |
| birds | Alectoris_barbara | Alectoris_barbara | 0 | 416.3 | 11 | 1391816.95 |
| birds | Alectoris_chukar | Alectoris_chukar | 0 | 502.1 | 9.2 | 11388432.95 |
| birds | Alectoris_graeca | Alectoris_graeca | 0 | 594.1 | 10.6 | 652935.31 |
| birds | Alectoris_melanocephala | Alectoris_melanocephala | 0 | 614.8 | 6.3 | 351773.76 |
| birds | Alectoris_philbyi | Alectoris_philbyi | 0 | 441 | 6.3 | 80634.58 |
| birds | Alectoris_rufa | Alectoris_rufa | 0 | 527.9 | 12 | 1107486.35 |
| birds | Alectroenas_madagascariensis | Alectroenas_madagascariensis | 0 | 173 | 1 | 311714.72 |
| birds | Alectrurus_risora | Alectrurus_risora | 1 | 21 | 3 | 935059.48 |
| birds | Alectura_lathami | Alectura_lathami | 0 | 2333.1 | 12 | 647248.18 |
| birds | Alethe_diademata | Alethe_diademata | 0 | 31.5 | 2 | 448296.69 |
| birds | Alexandrinus_eques | Psittacula_eques | 1 | 163 | 2.8 | 1010.36 |
| birds | Alexandrinus_krameri | Psittacula_krameri | 0 | 116.1 | 3.5 | 8372649.27 |
| birds | Alisterus_amboinensis | Alisterus_amboinensis | 0 | 153 | 2 | 148979.73 |

| | | | | | | |
|---|---|---|---|---|---|---|
| birds | Alisterus_scapularis | Alisterus_scapularis | 0 | 232.3 | 4.2 | 811455.94 |
| birds | Alophoixus_bres | Alophoixus_bres | 1 | 35.3 | 2 | 87148.92 |
| birds | Alophoixus_flaveolus | Alophoixus_flaveolus | 0 | 48.3 | 3.5 | 559779.47 |
| birds | Alophoixus_ochraceus | Alophoixus_ochraceus | 0 | 41.5 | 2 | 407080.06 |
| birds | Alophoixus_phaeocephalus | Alophoixus_phaeocephalus | 0 | 32 | 2 | 1368840.15 |
| birds | Alopochelidon_fucata | Alopochelidon_fucata | 0 | 14 | 4.5 | 4411696.77 |
| birds | Amadina_erythrocephala | Amadina_erythrocephala | 0 | 22.5 | 4.9 | 1628356.18 |
| birds | Amalocichla_incerta | Amalocichla_incerta | 0 | 32.7 | 1 | 161107.33 |
| birds | Amaurolimnas_concolor | Amaurolimnas_concolor | 0 | 133 | 4 | 1725851.55 |
| birds | Amaurornis_cinerea | Porzana_cinerea | 0 | 52.1 | 4.6 | 3636563.1 |
| birds | Amaurornis_marginalis | Aenigmatolimnas_marginalis | 0 | 51.2 | 4.5 | 4694327.06 |
| birds | Amaurornis_moluccana | Amaurornis_moluccana | 0 | 186.8 | 5.3 | 1266108.02 |
| birds | Amaurornis_phoenicurus | Amaurornis_phoenicurus | 0 | 180 | 6 | 10030319.8 |
| birds | Amaurospiza_moesta | Amaurospiza_moesta | 0 | 13.6 | 2 | 864091.67 |
| birds | Amazilia_boucardi | Amazilia_boucardi | 1 | 4.5 | 2 | 1301.76 |
| birds | Amazilia_cyanifrons | Amazilia_cyanifrons | 0 | 5 | 2 | 54825.57 |
| birds | Amazilia_franciae | Amazilia_franciae | 0 | 5 | 2 | 247257.85 |
| birds | Amazilia_rutila | Amazilia_rutila | 0 | 5.2 | 2 | 397592.64 |
| birds | Amazilia_tzacatl | Amazilia_tzacatl | 0 | 4.8 | 2 | 1019992.01 |
| birds | Amazilia_yucatanensis | Amazilia_yucatanensis | 0 | 3.9 | 2 | 582084.96 |
| birds | Amazilis_amazilia | Amazilia_amazilia | 0 | 4.3 | 2 | 169380.82 |
| birds | Amazona_aestiva | Amazona_aestiva | 0 | 451 | 3 | 5857429.4 |
| birds | Amazona_agilis | Amazona_agilis | 1 | 213 | 2.8 | 8263.64 |
| birds | Amazona_albifrons | Amazona_albifrons | 0 | 206 | 3.9 | 546295.4 |
| birds | Amazona_amazonica | Amazona_amazonica | 0 | 370 | 3.2 | 7382920.98 |
| birds | Amazona_autumnalis | Amazona_autumnalis | 0 | 416 | 3.5 | 977125.18 |
| birds | Amazona_barbadensis | Amazona_barbadensis | 0 | 270 | 3.2 | 42206.45 |
| birds | Amazona_brasiliensis | Amazona_brasiliensis | 0 | 430 | 3.5 | 4747.02 |
| birds | Amazona_collaria | Amazona_collaria | 1 | 187 | 3.5 | 7226.42 |
| birds | Amazona_dufresniana | Amazona_dufresniana | 0 | 623 | 3 | 546076.57 |
| birds | Amazona_farinosa | Amazona_farinosa | 0 | 626 | 3 | 7457891.94 |
| birds | Amazona_finschi | Amazona_finschi | 1 | 302 | 2 | 112128.49 |
| birds | Amazona_leucocephala | Amazona_leucocephala | 0 | 235.3 | 3.5 | 31494.26 |
| birds | Amazona_ochrocephala | Amazona_ochrocephala | 0 | 476.9 | 2.4 | 6635282.5 |

| | | | | | | |
|---|---|---|---|---|---|---|
| birds | Amazona_pretrei | Amazona_pretrei | 1 | 272.8 | 2.8 | 181933.24 |
| birds | Amazona_rhodocorytha | Amazona_rhodocorytha | 1 | 474.3 | 4 | 26932.41 |
| birds | Amazona_tucumana | Amazona_tucumana | 1 | 260 | 4 | 129668.06 |
| birds | Amazona_ventralis | Amazona_ventralis | 1 | 214 | 2.8 | 37516.12 |
| birds | Amazona_vinacea | Amazona_vinacea | 1 | 254 | 2.8 | 534573.2 |
| birds | Amazona_viridigenalis | Amazona_viridigenalis | 1 | 316 | 3.5 | 219413.11 |
| birds | Amazona_vittata | Amazona_vittata | 1 | 279.3 | 2.8 | 8952.43 |
| birds | Amazona_xantholora | Amazona_xantholora | 0 | 217 | 4 | 141207.23 |
| birds | Amazonetta_brasiliensis | Amazonetta_brasiliensis | 0 | 500 | 6.9 | 8747518.75 |
| birds | Ammodramus_aurifrons | Ammodramus_aurifrons | 0 | 17.8 | 2.4 | 3867543.93 |
| birds | Ammodramus_humeralis | Ammodramus_humeralis | 0 | 16 | 3.5 | 9115374.38 |
| birds | Ammodramus_savannarum | Ammodramus_savannarum | 0 | 17.6 | 4.5 | 7888248.74 |
| birds | Ammomanes_cinctura | Ammomanes_cinctura | 0 | 18.1 | 2.8 | 7020113.23 |
| birds | Ammomanes_deserti | Ammomanes_deserti | 0 | 23.8 | 2.2 | 12034050.9 |
| birds | Ammomanes_phoenicura | Ammomanes_phoenicura | 0 | 25.6 | 2.8 | 1826048.11 |
| birds | Ammomanopsis_grayi | Ammomanes_grayi | 0 | 21.4 | 2 | 184607.34 |
| birds | Ammoperdix_griseogularis | Ammoperdix_griseogularis | 0 | 201.9 | 7.3 | 2996830.79 |
| birds | Ammoperdix_heyi | Ammoperdix_heyi | 0 | 189.7 | 5.9 | 1664830.95 |
| birds | Ammospiza_caudacuta | Ammodramus_caudacutus | 1 | 19.2 | 3.9 | 215557.24 |
| birds | Ammospiza_leconteii | Ammodramus_leconteii | 0 | 13 | 3.9 | 5593741.34 |
| birds | Ammospiza_maritima | Ammodramus_maritimus | 0 | 22.2 | 4.5 | 211096.75 |
| birds | Ammospiza_nelsoni | Ammodramus_nelsoni | 0 | 15.1 | 3.9 | 2904622.44 |
| birds | Amphispiza_bilineata | Amphispiza_bilineata | 0 | 13.5 | 3.5 | 2559845.86 |
| birds | Amphispiza_quinquestriata | Aimophila_quinquestriata | 0 | 19.5 | 3.9 | 151617.22 |
| birds | Amytornis_barbatus | Amytornis_barbatus | 0 | 18.4 | 2.4 | 55946.51 |
| birds | Amytornis_dorotheae | Amytornis_dorotheae | 1 | 23.2 | 2 | 47008 |
| birds | Amytornis_goyderi | Amytornis_goyderi | 0 | 16.7 | 2.4 | 227659.69 |
| birds | Amytornis_housei | Amytornis_housei | 0 | 28.9 | 2 | 13972.82 |
| birds | Amytornis_purnelli | Amytornis_purnelli | 0 | 21 | 2.4 | 653027.47 |
| birds | Amytornis_striatus | Amytornis_striatus | 0 | 19.6 | 2.4 | 912126.85 |
| birds | Amytornis_textilis | Amytornis_textilis | 0 | 23 | 2 | 108980.14 |
| birds | Amytornis_woodwardi | Amytornis_woodwardi | 1 | 35.1 | 2 | 5520.04 |
| birds | Anabacerthia_striaticollis | Anabacerthia_striaticollis | 0 | 24.8 | 2 | 335322.02 |
| birds | Anabacerthia_variegaticeps | Anabacerthia_variegaticeps | 0 | 27.8 | 2 | 215334.52 |

| | | | | | | |
|---|---|---|---|---|---|---|
| birds | Anabathmis_reichenbachii | Nectarinia_reichenbachii | 0 | 11.4 | 1 | 2222320.41 |
| birds | Anairetes_flavirostris | Anairetes_flavirostris | 0 | 6 | 2.4 | 2046579.47 |
| birds | Anairetes_parulus | Anairetes_parulus | 0 | 6.2 | 2.4 | 2638454.77 |
| birds | Anaplectes_rubriceps | Anaplectes_rubriceps | 0 | 22.6 | 2.4 | 3258277.53 |
| birds | Anas_acuta | Anas_acuta | 0 | 944.6 | 7.9 | 52934942.78 |
| birds | Anas_aucklandica | Anas_aucklandica | 0 | 499.1 | 5.5 | 588.49 |
| birds | Anas_bahamensis | Anas_bahamensis | 0 | 490 | 7.7 | 2991584.7 |
| birds | Anas_bernieri | Anas_bernieri | 1 | 378.5 | 4.5 | 53437.43 |
| birds | Anas_capensis | Anas_capensis | 0 | 402 | 8.1 | 4317287.05 |
| birds | Anas_castanea | Anas_castanea | 0 | 624 | 8.4 | 3131751.24 |
| birds | Anas_crecca | Anas_crecca | 0 | 341.9 | 9.4 | 39459336.83 |
| birds | Anas_erythrorhyncha | Anas_erythrorhyncha | 0 | 558.8 | 7.7 | 10763443.79 |
| birds | Anas_flavirostris | Anas_flavirostris | 0 | 434 | 6.3 | 3942276.35 |
| birds | Anas_georgica | Anas_georgica | 0 | 623.8 | 6.3 | 4196176.91 |
| birds | Anas_gibberifrons | Anas_gibberifrons | 0 | 488.1 | 7.5 | 787609.11 |
| birds | Anas_luzonica | Anas_luzonica | 1 | 832 | 10 | 279937.11 |
| birds | Anas_melleri | Anas_melleri | 1 | 959.2 | 7.1 | 210476.27 |
| birds | Anas_platyrhynchos | Anas_platyrhynchos | 0 | 843.4 | 10.8 | 48513515.01 |
| birds | Anas_poecilorhyncha | Anas_poecilorhyncha | 0 | 1000 | 7.9 | 5263129.88 |
| birds | Anas_rubripes | Anas_rubripes | 0 | 1241 | 9.2 | 6750626.95 |
| birds | Anas_sparsa | Anas_sparsa | 0 | 981.5 | 5.7 | 9213622.59 |
| birds | Anas_superciliosa | Anas_superciliosa | 0 | 882.8 | 8.9 | 8845309.6 |
| birds | Anas_undulata | Anas_undulata | 0 | 1008 | 6.9 | 8476793.26 |
| birds | Anastomus_lamelligerus | Anastomus_lamelligerus | 0 | 1081 | 3.2 | 13409712.69 |
| birds | Anastomus_oscitans | Anastomus_oscitans | 0 | 1590 | 3.2 | 3412455.46 |
| birds | Andigena_laminirostris | Andigena_laminirostris | 0 | 311 | 2.4 | 8893.39 |
| birds | Andropadus_importunus | Andropadus_importunus | 0 | 27.4 | 1.7 | 1338546.67 |
| birds | Anhima_cornuta | Anhima_cornuta | 0 | 3150 | 3.9 | 10980198.81 |
| birds | Anhinga_anhinga | Anhinga_anhinga | 0 | 1235 | 3.9 | 14921101.49 |
| birds | Anhinga_melanogaster | Anhinga_melanogaster | 0 | 1480.1 | 3.9 | 6866922.35 |
| birds | Anisognathus_igniventris | Anisognathus_igniventris | 0 | 34 | 2 | 96719.41 |
| birds | Anodorhynchus_hyacinthinus | Anodorhynchus_hyacinthinus | 1 | 1331 | 2.4 | 2832918.27 |
| birds | Anodorhynchus_leari | Anodorhynchus_leari | 1 | 940 | 2 | 44706.58 |
| birds | Anorrhinus_galeritus | Anorrhinus_galeritus | 0 | 1172 | 2.4 | 1381919.99 |

| | | | | | | |
|---|---|---|---|---|---|---|
| birds | Anorrhinus_tickelli | Anorrhinus_tickelli | 0 | 792.6 | 3.2 | 160242.6 |
| birds | Anser_albifrons | Anser_albifrons | 0 | 2506.4 | 5.5 | 10495879 |
| birds | Anser_anser | Anser_anser | 0 | 3302.4 | 4.9 | 19080676.34 |
| birds | Anser_brachyrhynchus | Anser_brachyrhynchus | 0 | 2642 | 3.9 | 245976.49 |
| birds | Anser_caerulescens | Chen_caerulescens | 0 | 2636.2 | 4.5 | 9421501.27 |
| birds | Anser_canagicus | Chen_canagica | 0 | 2136.5 | 5 | 464902.48 |
| birds | Anser_cygnoid | Anser_cygnoides | 1 | 3511.9 | 5.5 | 7307006.36 |
| birds | Anser_erythropus | Anser_erythropus | 1 | 1755.5 | 4.9 | 29185526.08 |
| birds | Anser_fabalis | Anser_fabalis | 0 | 2754.7 | 4.9 | 17921622.23 |
| birds | Anser_indicus | Anser_indicus | 0 | 2212.6 | 4.9 | 4470193.83 |
| birds | Anser_rossii | Chen_rossii | 0 | 1636 | 4.5 | 2899430 |
| birds | Anseranas_semipalmata | Anseranas_semipalmata | 0 | 2393.4 | 7.4 | 1308440.74 |
| birds | Anthipes_monileger | Ficedula_monileger | 0 | 11 | 4 | 941257.11 |
| birds | Anthipes_solitaris | Ficedula_solitaris | 0 | 13.1 | 2.4 | 157229.38 |
| birds | Anthobaphes_violacea | Nectarinia_violacea | 0 | 9.4 | 1.4 | 91921.14 |
| birds | Anthochaera_carunculata | Anthochaera_carunculata | 0 | 106.1 | 2 | 2173024.9 |
| birds | Anthochaera_chrysoptera | Anthochaera_chrysoptera | 0 | 67.8 | 2 | 671226.2 |
| birds | Anthochaera_paradoxa | Anthochaera_paradoxa | 0 | 143.7 | 2.4 | 46805.04 |
| birds | Anthochaera_phrygia | Xanthomyza_phrygia | 1 | 40 | 2 | 281979.51 |
| birds | Anthoscopus_caroli | Anthoscopus_caroli | 0 | 6.9 | 4.9 | 4646761.76 |
| birds | Anthoscopus_minutus | Anthoscopus_minutus | 0 | 6.6 | 4.2 | 2370698.92 |
| birds | Anthoscopus_musculus | Anthoscopus_musculus | 0 | 5.2 | 4 | 1141427.54 |
| birds | Anthoscopus_parvulus | Anthoscopus_parvulus | 0 | 5.8 | 2 | 1993198.49 |
| birds | Anthracoceros_albirostris | Anthracoceros_albirostris | 0 | 892.9 | 1.7 | 4467706.59 |
| birds | Anthracoceros_coronatus | Anthracoceros_coronatus | 0 | 806.7 | 2.8 | 701610.86 |
| birds | Anthracoceros_malayanus | Anthracoceros_malayanus | 1 | 1050 | 2.4 | 798560.44 |
| birds | Anthracothorax_dominicus | Anthracothorax_dominicus | 0 | 5.8 | 2 | 75252.82 |
| birds | Anthracothorax_mango | Anthracothorax_mango | 0 | 8.3 | 2 | 11021.79 |
| birds | Anthracothorax_nigricollis | Anthracothorax_nigricollis | 0 | 7 | 2 | 9821751.59 |
| birds | Anthracothorax_prevostii | Anthracothorax_prevostii | 0 | 6.4 | 2 | 919148.98 |
| birds | Anthracothorax_viridigula | Anthracothorax_viridigula | 0 | 7.5 | 2 | 591465.01 |
| birds | Anthracothorax_viridis | Anthracothorax_viridis | 0 | 6.6 | 2 | 8774.42 |
| birds | Anthreptes_anchietae | Anthreptes_anchietae | 0 | 8 | 1.4 | 465325.07 |
| birds | Anthreptes_aurantius | Anthreptes_aurantium | 0 | 12.5 | 1.4 | 1740388.21 |

| | | | | | | |
|---|---|---|---|---|---|---|
| birds | Anthreptes_longuemarei | Anthreptes_longuemarei | 0 | 11.4 | 1.4 | 5262392.93 |
| birds | Anthreptes_malacensis | Anthreptes_malacensis | 0 | 11.3 | 2 | 2166604.35 |
| birds | Anthreptes_orientalis | Anthreptes_orientalis | 0 | 10.3 | 1.4 | 1935952.46 |
| birds | Anthreptes_rectirostris | Anthreptes_rectirostris | 0 | 10.6 | 2 | 448143.89 |
| birds | Anthreptes_reichenowi | Anthreptes_reichenowi | 0 | 7.3 | 2.4 | 438283.12 |
| birds | Anthreptes_rhodolaemus | Anthreptes_rhodolaemus | 0 | 13.1 | 1 | 442077.76 |
| birds | Anthreptes_rubritorques | Anthreptes_rubritorques | 1 | 10.7 | 2 | 7092.32 |
| birds | Anthreptes_seimundi | Nectarinia_seimundi | 0 | 6.6 | 2 | 2083351.49 |
| birds | Anthreptes_simplex | Anthreptes_simplex | 0 | 8.9 | 2 | 1062052.92 |
| birds | Anthropoides_paradiseus | Grus_paradisea | 1 | 4540.6 | 2 | 568469.11 |
| birds | Anthropoides_virgo | Grus_virgo | 0 | 2417 | 2 | 20925462.35 |
| birds | Anthus_berthelotii | Anthus_berthelotii | 0 | 15.9 | 3.2 | 8283.44 |
| birds | Anthus_brachyurus | Anthus_brachyurus | 0 | 15.3 | 2.4 | 1510202.74 |
| birds | Anthus_caffer | Anthus_caffer | 0 | 16.8 | 2.4 | 777646.72 |
| birds | Anthus_campestris | Anthus_campestris | 0 | 23 | 4.2 | 36400563.44 |
| birds | Anthus_cervinus | Anthus_cervinus | 0 | 20.9 | 3.7 | 15867275.1 |
| birds | Anthus_correndera | Anthus_correndera | 0 | 20.9 | 2.8 | 3686700.66 |
| birds | Anthus_crenatus | Anthus_crenatus | 0 | 30.7 | 3.5 | 343685.27 |
| birds | Anthus_godlewskii | Anthus_godlewskii | 0 | 25.3 | 3.9 | 9014937.37 |
| birds | Anthus_gustavi | Anthus_gustavi | 0 | 19.8 | 4.9 | 11759006.19 |
| birds | Anthus_hodgsoni | Anthus_hodgsoni | 0 | 21.3 | 2.4 | 18925518.72 |
| birds | Anthus_hoeschi | Anthus_hoeschi | 0 | 27 | 3.5 | 1210644.43 |
| birds | Anthus_leucophrys | Anthus_leucophrys | 0 | 25 | 2.4 | 8950742.46 |
| birds | Anthus_lineiventris | Anthus_lineiventris | 0 | 35 | 2.4 | 1635119.6 |
| birds | Anthus_lutescens | Anthus_lutescens | 0 | 12.6 | 2.8 | 7241509.05 |
| birds | Anthus_melindae | Anthus_melindae | 0 | 22.5 | 2.4 | 108383.57 |
| birds | Anthus_nattereri | Anthus_nattereri | 1 | 19.5 | 4 | 753720.22 |
| birds | Anthus_novaeseelandiae | Anthus_novaeseelandiae | 0 | 24.1 | 3.2 | 6479742.41 |
| birds | Anthus_petrosus | Anthus_petrosus | 0 | 22.4 | 4.9 | 497601.29 |
| birds | Anthus_pratensis | Anthus_pratensis | 0 | 18.4 | 3.7 | 16554757.4 |
| birds | Anthus_richardi | Anthus_richardi | 0 | 32.2 | 3.5 | 23084325.31 |
| birds | Anthus_roseatus | Anthus_roseatus | 0 | 20.6 | 3.5 | 5517844.36 |
| birds | Anthus_rubescens | Anthus_rubescens | 0 | 20.7 | 4.6 | 37048750.13 |
| birds | Anthus_rufulus | Anthus_rufulus | 0 | 20.4 | 3.5 | 7678484.07 |

| | | | | | | |
|---|---|---|---|---|---|---|
| birds | Anthus_similis | Anthus_similis | 0 | 25.3 | 2.4 | 9112709.12 |
| birds | Anthus_spinoletta | Anthus_spinoletta | 0 | 23.9 | 4.9 | 17694009.14 |
| birds | Anthus_spragueii | Anthus_spragueii | 1 | 25.5 | 4.5 | 5658382.23 |
| birds | Anthus_sylvanus | Anthus_sylvanus | 0 | 31 | 3.9 | 465229.13 |
| birds | Anthus_trivialis | Anthus_trivialis | 0 | 23.3 | 4 | 56587106.05 |
| birds | Anthus_vaalensis | Anthus_vaalensis | 0 | 27 | 2.4 | 4452532.2 |
| birds | Antilophia_galeata | Antilophia_galeata | 0 | 21.5 | 2 | 1960407.09 |
| birds | Antrostomus_badius | Caprimulgus_badius | 0 | 59.8 | 2 | 139506.92 |
| birds | Antrostomus_carolinensis | Caprimulgus_carolinensis | 0 | 109 | 2 | 3554721.02 |
| birds | Antrostomus_cubanensis | Caprimulgus_cubanensis | 0 | 63.5 | 1.4 | 109234.54 |
| birds | Antrostomus_noctitherus | Caprimulgus_noctitherus | 1 | 36.1 | 1.4 | 7754.14 |
| birds | Antrostomus_ridgwayi | Caprimulgus_ridgwayi | 0 | 48 | 2 | 631238.26 |
| birds | Antrostomus_rufus | Caprimulgus_rufus | 0 | 94.2 | 1.4 | 7445742.53 |
| birds | Antrostomus_salvini | Caprimulgus_salvini | 0 | 56 | 2 | 115696.33 |
| birds | Antrostomus_saturatus | Caprimulgus_saturatus | 0 | 52 | 1 | 5381.48 |
| birds | Antrostomus_vociferus | Caprimulgus_vociferus | 0 | 53.3 | 1.4 | 4881057.02 |
| birds | Anumbius_annumbi | Anumbius_annumbi | 0 | 41.5 | 3.9 | 2928674.73 |
| birds | Apalis_binotata | Apalis_binotata | 0 | 8.2 | 1.4 | 144782.97 |
| birds | Apalis_chariessa | Apalis_chariessa | 0 | 9.2 | 2.4 | 21433.89 |
| birds | Apalis_cinerea | Apalis_cinerea | 0 | 9.9 | 3 | 438312.46 |
| birds | Apalis_flavida | Apalis_flavida | 0 | 8.2 | 2.8 | 4880471.31 |
| birds | Apalis_jacksoni | Apalis_jacksoni | 0 | 8.7 | 2 | 541008.1 |
| birds | Apalis_melanocephala | Apalis_melanocephala | 0 | 8.8 | 2.4 | 375838.95 |
| birds | Apalis_ruddi | Apalis_ruddi | 0 | 9.7 | 2 | 196688.52 |
| birds | Apalis_sharpii | Apalis_sharpii | 0 | 8.7 | 3 | 386331.11 |
| birds | Apalis_thoracica | Apalis_thoracica | 0 | 11.1 | 3 | 1229497.4 |
| birds | Apaloderma_narina | Apaloderma_narina | 0 | 67.7 | 2 | 10932030.89 |
| birds | Apaloderma_vittatum | Apaloderma_vittatum | 0 | 55 | 2.4 | 485903.03 |
| birds | Aphelocephala_leucopsis | Aphelocephala_leucopsis | 1 | 12.3 | 3.2 | 3873177.35 |
| birds | Aphelocephala_nigricincta | Aphelocephala_nigricincta | 0 | 10.4 | 2.8 | 1619312.31 |
| birds | Aphelocephala_pectoralis | Aphelocephala_pectoralis | 0 | 9 | 3 | 225402.65 |
| birds | Aphelocoma_californica | Aphelocoma_californica | 0 | 85.7 | 3.5 | 2323260.89 |
| birds | Aphelocoma_coerulescens | Aphelocoma_coerulescens | 1 | 77.1 | 3.5 | 65032.2 |
| birds | Aphelocoma_ultramarina | Aphelocoma_ultramarina | 0 | 113.3 | 4 | 87898.14 |

| | | | | | | |
|---|---|---|---|---|---|---|
| birds | Aphelocoma_unicolor | Aphelocoma_unicolor | 0 | 124 | 3.5 | 149103.75 |
| birds | Aphrastura_spinicauda | Aphrastura_spinicauda | 0 | 11.5 | 3.5 | 491332.8 |
| birds | Aplonis_cantoroides | Aplonis_cantoroides | 0 | 55.7 | 2.4 | 822753.89 |
| birds | Aplonis_metallica | Aplonis_metallica | 0 | 56.5 | 1.7 | 762829.93 |
| birds | Aplonis_panayensis | Aplonis_panayensis | 0 | 56.2 | 3.5 | 2145170.85 |
| birds | Aplonis_tabuensis | Aplonis_tabuensis | 0 | 62.5 | 2.4 | 23355.89 |
| birds | Aplopelia_larvata | Aplopelia_larvata | 0 | 155 | 2 | 2621498.99 |
| birds | Aprosmictus_erythropterus | Aprosmictus_erythropterus | 0 | 136 | 3.9 | 2743055.21 |
| birds | Apus_affinis | Apus_affinis | 0 | 22.6 | 1.7 | 22143832.39 |
| birds | Apus_apus | Apus_apus | 0 | 37.6 | 2 | 42973934.9 |
| birds | Apus_barbatus | Apus_barbatus | 0 | 42.8 | 1.4 | 1200605.42 |
| birds | Apus_berliozi | Apus_berliozi | 0 | 38.2 | 2 | 214046.13 |
| birds | Apus_bradfieldi | Apus_bradfieldi | 0 | 42.4 | 2 | 882341.76 |
| birds | Apus_caffer | Apus_caffer | 0 | 22.1 | 1.7 | 12490681.95 |
| birds | Apus_horus | Apus_horus | 0 | 27.5 | 2 | 2043600.71 |
| birds | Apus_niansae | Apus_niansae | 0 | 32.5 | 1.7 | 381006.41 |
| birds | Apus_nipalensis | Apus_nipalensis | 0 | 29.8 | 2.2 | 5768014.18 |
| birds | Apus_pacificus | Apus_pacificus | 0 | 39.4 | 1.7 | 28562076.19 |
| birds | Apus_pallidus | Apus_pallidus | 0 | 41.9 | 2.4 | 9749752.17 |
| birds | Apus_unicolor | Apus_unicolor | 0 | 22.5 | 2 | 16132.27 |
| birds | Aquila_adalberti | Aquila_adalberti | 1 | 2958 | 2.4 | 1072953.52 |
| birds | Aquila_audax | Aquila_audax | 0 | 3449.9 | 2 | 7679648.03 |
| birds | Aquila_chrysaetos | Aquila_chrysaetos | 0 | 4248 | 2 | 43360026.83 |
| birds | Aquila_fasciata | Aquila_fasciatus | 0 | 1936.5 | 1.4 | 5912536.57 |
| birds | Aquila_heliaca | Aquila_heliaca | 1 | 3125.2 | 2.4 | 31736715.07 |
| birds | Aquila_nipalensis | Aquila_nipalensis | 1 | 2714.3 | 1.7 | 30326757.84 |
| birds | Aquila_rapax | Aquila_rapax | 1 | 2236.1 | 2 | 17572054.36 |
| birds | Aquila_spilogaster | Hieraaetus_spilogaster | 0 | 1466.2 | 2 | 12423292.01 |
| birds | Aquila_verreauxii | Aquila_verreauxii | 0 | 4195 | 2 | 4020161.71 |
| birds | Ara_ambiguus | Ara_ambiguus | 1 | 1300 | 2.4 | 151979.98 |
| birds | Ara_ararauna | Ara_ararauna | 0 | 1125 | 1.7 | 7836201.13 |
| birds | Ara_chloropterus | Ara_chloropterus | 0 | 1214 | 2.4 | 8101232.43 |
| birds | Ara_glaucogularis | Ara_glaucogularis | 1 | 784 | 2.4 | 30260.16 |
| birds | Ara_macao | Ara_macao | 0 | 1015 | 2 | 7046076.25 |

| | | | | | | |
|---|---|---|---|---|---|---|
| birds | Ara_militaris | Ara_militaris | 1 | 1134 | 2.4 | 591218.42 |
| birds | Ara_rubrogenys | Ara_rubrogenys | 1 | 468 | 1.7 | 27916.97 |
| birds | Ara_severus | Ara_severus | 0 | 343 | 2 | 5828903.93 |
| birds | Arachnothera_affinis | Arachnothera_affinis | 0 | 26.4 | 1.4 | 349980.16 |
| birds | Arachnothera_chrysogenys | Arachnothera_chrysogenys | 0 | 23.2 | 2 | 1237350.36 |
| birds | Arachnothera_longirostra | Arachnothera_longirostra | 0 | 12.5 | 2.4 | 3514761.53 |
| birds | Arachnothera_magna | Arachnothera_magna | 0 | 30.7 | 2.4 | 1984624.62 |
| birds | Arachnothera_robusta | Arachnothera_robusta | 0 | 29.3 | 2 | 630859.75 |
| birds | Aramides_axillaris | Aramides_axillaris | 0 | 292 | 5 | 468740.12 |
| birds | Aramides_cajaneus | Aramides_cajanea | 0 | 397 | 4.6 | 13038120.14 |
| birds | Aratinga_jandaya | Aratinga_jandaya | 0 | 89.4 | 3 | 1514385.78 |
| birds | Aratinga_nenday | Nandayus_nenday | 0 | 128 | 4 | 373769.01 |
| birds | Aratinga_solstitialis | Aratinga_solstitialis | 1 | 119.6 | 4 | 86113.47 |
| birds | Aratinga_weddellii | Aratinga_weddellii | 0 | 108 | 3.5 | 2260164.39 |
| birds | Arborophila_ardens | Arborophila_ardens | 1 | 266.6 | 2.4 | 13199.58 |
| birds | Arborophila_atrogularis | Arborophila_atrogularis | 0 | 237.3 | 4.5 | 355074.89 |
| birds | Arborophila_crudigularis | Arborophila_crudigularis | 0 | 256.8 | 6.9 | 22768.9 |
| birds | Arborophila_gingica | Arborophila_gingica | 0 | 253 | 5.9 | 396418.35 |
| birds | Arborophila_javanica | Arborophila_javanica | 0 | 272 | 4 | 36961.99 |
| birds | Arborophila_mandellii | Arborophila_mandellii | 0 | 268 | 4 | 74187.76 |
| birds | Arborophila_orientalis | Arborophila_orientalis | 1 | 336 | 2.8 | 10707.85 |
| birds | Arborophila_rufipectus | Arborophila_rufipectus | 1 | 400.1 | 5.5 | 7707.36 |
| birds | Arborophila_rufogularis | Arborophila_rufogularis | 0 | 344.5 | 4.5 | 1245539.71 |
| birds | Arborophila_torqueola | Arborophila_torqueola | 0 | 344.5 | 3.9 | 721402.09 |
| birds | Archilochus_alexandri | Archilochus_alexandri | 0 | 3.4 | 2 | 3841632.3 |
| birds | Archilochus_colubris | Archilochus_colubris | 0 | 3.1 | 2 | 7444664.68 |
| birds | Ardea_alba | Casmerodius_albus | 0 | 871.3 | 3.9 | 80817244.41 |
| birds | Ardea_cinerea | Ardea_cinerea | 0 | 1443 | 3.9 | 51699618.43 |
| birds | Ardea_cocoi | Ardea_cocoi | 0 | 1752.4 | 2.8 | 14742289.16 |
| birds | Ardea_goliath | Ardea_goliath | 0 | 4468 | 3 | 17921027.03 |
| birds | Ardea_herodias | Ardea_herodias | 0 | 2523.4 | 3.7 | 14906902.55 |
| birds | Ardea_intermedia | Mesophoyx_intermedia | 0 | 458.8 | 3.5 | 8898068.6 |
| birds | Ardea_melanocephala | Ardea_melanocephala | 0 | 1060 | 3.5 | 19067243.58 |
| birds | Ardea_pacifica | Ardea_pacifica | 0 | 881 | 3.5 | 6934005.01 |

| | | | | | | |
|---|---|---|---|---|---|---|
| birds | Ardea_purpurea | Ardea_purpurea | 0 | 1064.5 | 4 | 42294330.03 |
| birds | Ardea_sumatrana | Ardea_sumatrana | 0 | 2024 | 2 | 1764501.96 |
| birds | Ardeola_bacchus | Ardeola_bacchus | 0 | 304.9 | 4 | 7868535.62 |
| birds | Ardeola_grayii | Ardeola_grayii | 0 | 253 | 3.9 | 3991850.35 |
| birds | Ardeola_ralloides | Ardeola_ralloides | 0 | 287 | 3.7 | 34073703.04 |
| birds | Ardeotis_arabs | Ardeotis_arabs | 0 | 5943.5 | 1.4 | 2950367.62 |
| birds | Ardeotis_australis | Ardeotis_australis | 0 | 4182.4 | 1.4 | 6717248.95 |
| birds | Ardeotis_kori | Ardeotis_kori | 0 | 7994.2 | 2 | 3565934.25 |
| birds | Ardeotis_nigriceps | Ardeotis_nigriceps | 1 | 7235.3 | 1 | 1064221.45 |
| birds | Arenaria_interpres | Arenaria_interpres | 0 | 136 | 4 | 13072863.93 |
| birds | Arenaria_melanocephala | Arenaria_melanocephala | 0 | 126.2 | 4 | 1073634.76 |
| birds | Argusianus_argus | Argusianus_argus | 1 | 1959.9 | 2 | 1480121.15 |
| birds | Argya_affinis | Turdoides_affinis | 0 | 63.3 | 3.5 | 745811.26 |
| birds | Argya_altirostris | Turdoides_altirostris | 0 | 33 | 3.5 | 149200.16 |
| birds | Argya_aylmeri | Turdoides_aylmeri | 0 | 35.3 | 2 | 1055408.06 |
| birds | Argya_caudata | Turdoides_caudata | 0 | 39.2 | 3.5 | 3191144.09 |
| birds | Argya_cinereifrons | Garrulax_cinereifrons | 1 | 70 | 3 | 11916.82 |
| birds | Argya_earlei | Turdoides_earlei | 0 | 47 | 3.5 | 1024770.23 |
| birds | Argya_fulva | Turdoides_fulva | 0 | 61.4 | 4.5 | 3666952.83 |
| birds | Argya_gularis | Turdoides_gularis | 0 | 45.5 | 3.9 | 216441.28 |
| birds | Argya_longirostris | Turdoides_longirostris | 1 | 35 | 3.9 | 28660.3 |
| birds | Argya_malcolmi | Turdoides_malcolmi | 0 | 75.7 | 3.5 | 1922153.02 |
| birds | Argya_rubiginosa | Turdoides_rubiginosa | 0 | 40.6 | 2.4 | 1410146.95 |
| birds | Argya_rufescens | Turdoides_rufescens | 0 | 64.5 | 2.4 | 12170.71 |
| birds | Argya_squamiceps | Turdoides_squamiceps | 0 | 72.9 | 3.9 | 1273158.32 |
| birds | Argya_subrufa | Turdoides_subrufa | 0 | 66.7 | 4 | 178058.68 |
| birds | Arizelocichla_masukuensis | Andropadus_masukuensis | 0 | 26.8 | 1.4 | 163721.68 |
| birds | Arizelocichla_milanjensis | Andropadus_milanjensis | 0 | 37.4 | 2 | 42374.09 |
| birds | Arizelocichla_montana | Andropadus_montanus | 0 | 32.6 | 1.4 | 61327.27 |
| birds | Arizelocichla_tephrolaema | Andropadus_tephrolaemus | 0 | 38.3 | 2 | 59489.94 |
| birds | Arremon_aurantiirostris | Arremon_aurantiirostris | 0 | 34.5 | 2 | 693974.17 |
| birds | Arremon_brunneinucha | Arremon_brunneinucha | 0 | 44 | 2 | 618730.15 |
| birds | Arses_kaupi | Arses_kaupi | 0 | 13.5 | 2 | 27770.73 |
| birds | Artamella_viridis | Artamella_viridis | 0 | 48.8 | 3 | 415338.92 |

| | | | | | | |
|---|---|---|---|---|---|---|
| birds | Artamus_cinereus | Artamus_cinereus | 0 | 35.6 | 3.5 | 7423572.8 |
| birds | Artamus_cyanopterus | Artamus_cyanopterus | 0 | 34.6 | 3.5 | 3897118.38 |
| birds | Artamus_fuscus | Artamus_fuscus | 0 | 39.8 | 2.4 | 4701116.55 |
| birds | Artamus_leucoryn | Artamus_leucorynchus | 0 | 41.7 | 3.5 | 7650935.5 |
| birds | Artamus_mentalis | Artamus_mentalis | 0 | 41.9 | 2.4 | 17666.45 |
| birds | Artamus_minor | Artamus_minor | 0 | 16 | 3 | 6485166.89 |
| birds | Artamus_personatus | Artamus_personatus | 0 | 34.6 | 2.4 | 7500863.72 |
| birds | Artamus_superciliosus | Artamus_superciliosus | 0 | 35.5 | 2.4 | 7162672.29 |
| birds | Artemisiospiza_belli | Amphispiza_belli | 0 | 17 | 3.5 | 304827.38 |
| birds | Artisornis_metopias | Artisornis_metopias | 0 | 8.4 | 2 | 68472.65 |
| birds | Artomyias_fuliginosa | Muscicapa_infuscata | 0 | 17.7 | 2.4 | 3058848.69 |
| birds | Arundinax_aedon | Acrocephalus_aedon | 0 | 22.4 | 4.9 | 14630790.95 |
| birds | Arundinicola_leucocephala | Arundinicola_leucocephala | 0 | 13.8 | 2.8 | 9087776.96 |
| birds | Asarcornis_scutulata | Cairina_scutulata | 1 | 2937.4 | 8.8 | 999723.98 |
| birds | Ashbyia_lovensis | Ashbyia_lovensis | 0 | 17.5 | 2.8 | 699489.42 |
| birds | Asio_capensis | Asio_capensis | 0 | 334 | 2.4 | 7713158.39 |
| birds | Asio_clamator | Pseudoscops_clamator | 0 | 443.3 | 3 | 9101422.66 |
| birds | Asio_flammeus | Asio_flammeus | 0 | 322.6 | 7.1 | 60054481.5 |
| birds | Asio_otus | Asio_otus | 0 | 296.6 | 5.9 | 35423274.66 |
| birds | Asio_stygius | Asio_stygius | 0 | 565 | 2 | 5772064.13 |
| birds | Asthenes_baeri | Asthenes_baeri | 0 | 17.8 | 3.9 | 1726715.05 |
| birds | Asthenes_dorbignyi | Asthenes_dorbignyi | 0 | 15.9 | 2 | 252131.25 |
| birds | Asthenes_fuliginosa | Schizoeaca_fuliginosa | 0 | 18.3 | 2 | 96182.51 |
| birds | Asthenes_harterti | Schizoeaca_harterti | 0 | 12.9 | 2 | 23849.1 |
| birds | Asthenes_hudsoni | Asthenes_hudsoni | 0 | 19 | 3.2 | 803230.15 |
| birds | Asthenes_modesta | Asthenes_modesta | 0 | 16.8 | 2.8 | 2015310.78 |
| birds | Asthenes_pyrrholeuca | Asthenes_pyrrholeuca | 0 | 13.2 | 2.8 | 2740647.46 |
| birds | Asthenes_wyatti | Asthenes_wyatti | 0 | 15.2 | 2.4 | 467539.8 |
| birds | Athene_brama | Athene_brama | 0 | 112 | 3.2 | 5213398.06 |
| birds | Athene_cunicularia | Athene_cunicularia | 0 | 150.6 | 8.1 | 15156308.38 |
| birds | Athene_noctua | Athene_noctua | 0 | 168.9 | 4.2 | 26928713.11 |
| birds | Athene_superciliaris | Ninox_superciliaris | 0 | 231 | 3.9 | 415419.42 |
| birds | Atimastillas_flavicollis | Chlorocichla_flavicollis | 0 | 48.7 | 2 | 1683203.48 |
| birds | Atrichornis_clamosus | Atrichornis_clamosus | 1 | 42.2 | 1 | 19384.96 |

| | | | | | | |
|---|---|---|---|---|---|---|
| birds | Atrichornis_rufescens | Atrichornis_rufescens | 1 | 24 | 2 | 32957.63 |
| birds | Attagis_gayi | Attagis_gayi | 0 | 311 | 4 | 1031574.14 |
| birds | Attagis_malouinus | Attagis_malouinus | 0 | 340 | 4 | 323646.86 |
| birds | Atticora_fasciata | Atticora_fasciata | 0 | 13 | 4.5 | 6512985.46 |
| birds | Atticora_pileata | Notiochelidon_pileata | 0 | 12.2 | 4 | 75275.04 |
| birds | Attila_bolivianus | Attila_bolivianus | 0 | 39.5 | 2 | 2790241.25 |
| birds | Attila_cinnamomeus | Attila_cinnamomeus | 0 | 38.8 | 2.4 | 5275009.27 |
| birds | Attila_spadiceus | Attila_spadiceus | 0 | 39.1 | 2.8 | 8099963.71 |
| birds | Augastes_lumachella | Augastes_lumachella | 0 | 4 | 2 | 20687.46 |
| birds | Augastes_scutatus | Augastes_scutatus | 0 | 3 | 2 | 28653.28 |
| birds | Aulacorhynchus_haematopygus | Aulacorhynchus_haematopygus | 0 | 194.4 | 4 | 179236.71 |
| birds | Aulacorhynchus_prasinus | Aulacorhynchus_prasinus | 0 | 170.6 | 2.2 | 420156.22 |
| birds | Auriparus_flaviceps | Auriparus_flaviceps | 0 | 6.8 | 3.5 | 1879707.38 |
| birds | Automolus_leucophthalmus | Automolus_leucophthalmus | 0 | 34.5 | 3.5 | 1800565.07 |
| birds | Automolus_ochrolaemus | Automolus_ochrolaemus | 0 | 40.2 | 2.4 | 6344367.8 |
| birds | Aviceda_cuculoides | Aviceda_cuculoides | 0 | 277 | 2 | 10593852.38 |
| birds | Aviceda_jerdoni | Aviceda_jerdoni | 0 | 363 | 2.4 | 1597402.79 |
| birds | Aviceda_leuphotes | Aviceda_leuphotes | 0 | 194 | 2.4 | 4574587.22 |
| birds | Aviceda_subcristata | Aviceda_subcristata | 0 | 322.7 | 2.4 | 1718124.83 |
| birds | Avocettula_recurvirostris | Avocettula_recurvirostris | 0 | 4.2 | 2 | 2531616.04 |
| birds | Aythya_affinis | Aythya_affinis | 0 | 819.5 | 9.9 | 16453053.34 |
| birds | Aythya_americana | Aythya_americana | 0 | 1075.7 | 9 | 11782105.96 |
| birds | Aythya_australis | Aythya_australis | 0 | 869.4 | 10.8 | 6825013.89 |
| birds | Aythya_baeri | Aythya_baeri | 1 | 681.2 | 7.7 | 9009013.37 |
| birds | Aythya_collaris | Aythya_collaris | 0 | 704.6 | 8.9 | 14291364.29 |
| birds | Aythya_ferina | Aythya_ferina | 1 | 823 | 8.9 | 27595007.89 |
| birds | Aythya_fuligula | Aythya_fuligula | 0 | 701.2 | 9.4 | 33316870.64 |
| birds | Aythya_marila | Aythya_marila | 0 | 1005.4 | 9.4 | 16793614.24 |
| birds | Aythya_novaeseelandiae | Aythya_novaeseelandiae | 0 | 682 | 5.7 | 162640.63 |
| birds | Aythya_nyroca | Aythya_nyroca | 0 | 574 | 8.9 | 36110689.77 |
| birds | Aythya_valisineria | Aythya_valisineria | 0 | 1202 | 9.5 | 10988439.08 |
| birds | Baeolophus_atricristatus | Baeolophus_atricristatus | 0 | 17.5 | 6 | 741786.53 |
| birds | Baeolophus_bicolor | Baeolophus_bicolor | 0 | 21.6 | 5.9 | 2969712.5 |
| birds | Baeolophus_ridgwayi | Baeolophus_ridgwayi | 0 | 15.9 | 6.9 | 1056208.76 |

| | | | | | | |
|---|---|---|---|---|---|---|
| birds | Baeolophus_wollweberi | Baeolophus_wollweberi | 0 | 10.4 | 5.9 | 532101.62 |
| birds | Baeopogon_indicator | Baeopogon_indicator | 0 | 45.8 | 2 | 4280773.28 |
| birds | Balaeniceps_rex | Balaeniceps_rex | 1 | 5941 | 1.7 | 1291415.43 |
| birds | Balearica_pavonina | Balearica_pavonina | 1 | 3590 | 2.4 | 2976931.85 |
| birds | Balearica_regulorum | Balearica_regulorum | 1 | 3772 | 2 | 3528882.19 |
| birds | Bambusicola_fytchii | Bambusicola_fytchii | 0 | 313.7 | 4.9 | 550319.01 |
| birds | Bambusicola_thoracicus | Bambusicola_thoracicus | 0 | 264.8 | 4.6 | 1491544.12 |
| birds | Barnardius_zonarius | Barnardius_zonarius | 0 | 133 | 5.3 | 5305452.04 |
| birds | Basileuterus_culicivorus | Basileuterus_culicivorus | 0 | 10.5 | 2.8 | 522685.98 |
| birds | Basileuterus_melanogenys | Basileuterus_melanogenys | 0 | 11.8 | 2 | 4831.72 |
| birds | Basileuterus_rufifrons | Basileuterus_rufifrons | 0 | 10.4 | 2.4 | 967644.87 |
| birds | Basileuterus_tristriatus | Basileuterus_tristriatus | 0 | 12 | 2 | 432295.34 |
| birds | Basilinna_leucotis | Hylocharis_leucotis | 0 | 3.4 | 2 | 556989.38 |
| birds | Basilinna_xantusii | Hylocharis_xantusii | 0 | 3.4 | 2 | 41923.51 |
| birds | Batara_cinerea | Batara_cinerea | 0 | 131 | 2.4 | 600001.37 |
| birds | Bathmocercus_rufus | Bathmocercus_rufus | 0 | 15.4 | 2 | 638060.54 |
| birds | Batis_capensis | Batis_capensis | 0 | 11.7 | 2 | 417033.5 |
| birds | Batis_fratrum | Batis_fratrum | 0 | 11.9 | 2.4 | 426278.31 |
| birds | Batis_minor | Batis_minor | 0 | 12.5 | 2 | 280465.17 |
| birds | Batis_minulla | Batis_minulla | 0 | 9.9 | 2 | 474658.69 |
| birds | Batis_mixta | Batis_mixta | 0 | 12.2 | 2 | 57484.87 |
| birds | Batis_molitor | Batis_molitor | 0 | 11.6 | 2 | 5233583.28 |
| birds | Batis_orientalis | Batis_orientalis | 0 | 11.1 | 2.4 | 2679950.48 |
| birds | Batis_pririt | Batis_pririt | 0 | 9.3 | 2 | 1547981.31 |
| birds | Batis_senegalensis | Batis_senegalensis | 0 | 9.6 | 2 | 3034470.61 |
| birds | Batis_soror | Batis_soror | 0 | 9.5 | 2 | 863131.73 |
| birds | Batrachostomus_hodgsoni | Batrachostomus_hodgsoni | 0 | 51 | 1.4 | 119417.56 |
| birds | Batrachostomus_javensis | Batrachostomus_javensis | 0 | 46.3 | 1.4 | 1413593.97 |
| birds | Belocercus_longicaudus | Psittacula_longicauda | 1 | 107 | 2.4 | 877581.6 |
| birds | Bias_musicus | Bias_musicus | 0 | 21.9 | 2 | 3741138.35 |
| birds | Biziura_lobata | Biziura_lobata | 0 | 1928.5 | 2.4 | 2736871.39 |
| birds | Bleda_canicapillus | Bleda_canicapillus | 0 | 41.3 | 2 | 1094053.69 |
| birds | Bleda_eximius | Bleda_eximius | 0 | 35.7 | 2 | 339296.54 |
| birds | Bleda_syndactylus | Bleda_syndactylus | 0 | 46.1 | 2 | 3527560.89 |

| | | | | | | |
|---|---|---|---|---|---|---|
| birds | Blythipicus_pyrrhotis | Blythipicus_pyrrhotis | 0 | 132 | 2.4 | 1622803.09 |
| birds | Bocagia_minuta | Tchagra_minutus | 0 | 34.4 | 2.4 | 3369836.92 |
| birds | Boissonneaua_flavescens | Boissonneaua_flavescens | 0 | 8.2 | 2 | 128710.18 |
| birds | Boissonneaua_jardini | Boissonneaua_jardini | 0 | 9.1 | 2 | 30988.1 |
| birds | Boissonneaua_matthewsii | Boissonneaua_matthewsii | 0 | 7.7 | 2 | 127445.48 |
| birds | Bolemoreus_frenatus | Lichenostomus_frenatus | 0 | 34.1 | 2 | 58164.64 |
| birds | Bolemoreus_hindwoodi | Lichenostomus_hindwoodi | 0 | 22.9 | 2 | 1242.5 |
| birds | Bombycilla_cedrorum | Bombycilla_cedrorum | 0 | 31.6 | 3.9 | 13624903.87 |
| birds | Bombycilla_garrulus | Bombycilla_garrulus | 0 | 54.4 | 5.5 | 32415917 |
| birds | Bonasa_umbellus | Bonasa_umbellus | 0 | 530.9 | 11 | 8076355.64 |
| birds | Bostrychia_carunculata | Bostrychia_carunculata | 0 | 1500 | 2.4 | 539792.68 |
| birds | Bostrychia_hagedash | Bostrychia_hagedash | 0 | 1238 | 2.4 | 16241597.55 |
| birds | Bostrychia_rara | Bostrychia_rara | 0 | 866 | 2 | 2806304.39 |
| birds | Botaurus_lentiginosus | Botaurus_lentiginosus | 0 | 706 | 3.7 | 14235010.42 |
| birds | Botaurus_pinnatus | Botaurus_pinnatus | 0 | 812.2 | 2.4 | 2262647.75 |
| birds | Botaurus_poiciloptilus | Botaurus_poiciloptilus | 1 | 1083.7 | 4.2 | 1425382.93 |
| birds | Botaurus_stellaris | Botaurus_stellaris | 0 | 1319.5 | 4.6 | 24816284.22 |
| birds | Brachygalba_goeringi | Brachygalba_goeringi | 0 | 17 | 4 | 196562.33 |
| birds | Brachygalba_lugubris | Brachygalba_lugubris | 0 | 15.9 | 3 | 4798770.73 |
| birds | Brachypodius_atriceps | Pycnonotus_atriceps | 0 | 25.5 | 2.4 | 2968877.33 |
| birds | Brachypteryx_leucophris | Brachypteryx_leucophrys | 0 | 15.8 | 2.8 | 2289506.95 |
| birds | Brachypteryx_montana | Brachypteryx_montana | 0 | 17.7 | 2.8 | 103057.9 |
| birds | Bradornis_comitatus | Muscicapa_comitata | 0 | 14.1 | 1.4 | 3634771.07 |
| birds | Bradornis_mariquensis | Bradornis_mariquensis | 0 | 25.2 | 3 | 1651176.93 |
| birds | Bradornis_microrhynchus | Bradornis_microrhynchus | 0 | 19.5 | 2 | 2477905.95 |
| birds | Bradypterus_baboecala | Bradypterus_baboecala | 0 | 12.9 | 2.4 | 3040370.23 |
| birds | Bradypterus_barratti | Bradypterus_barratti | 0 | 18.8 | 2 | 227225.24 |
| birds | Bradypterus_cinnamomeus | Bradypterus_cinnamomeus | 0 | 17.4 | 2.4 | 513979.16 |
| birds | Bradypterus_lopezi | Bradypterus_lopezi | 0 | 19.2 | 2.4 | 424179.16 |
| birds | Bradypterus_sylvaticus | Bradypterus_sylvaticus | 1 | 21 | 2.4 | 8825.28 |
| birds | Branta_bernicla | Branta_bernicla | 0 | 1277.9 | 3.9 | 4854965.92 |
| birds | Branta_canadensis | Branta_canadensis | 0 | 2811.7 | 5.3 | 16352791.22 |
| birds | Branta_leucopsis | Branta_leucopsis | 0 | 1684 | 4.5 | 259364.55 |
| birds | Branta_ruficollis | Branta_ruficollis | 1 | 1226.5 | 6.5 | 2994790.39 |

| | | | | | | |
|---|---|---|---|---|---|---|
| birds | Branta_sandvicensis | Branta_sandvicensis | 0 | 2044.1 | 3.9 | 1590.28 |
| birds | Brotogeris_chrysoptera | Brotogeris_chrysoptera | 0 | 54.5 | 3.5 | 3112139.06 |
| birds | Brotogeris_cyanoptera | Brotogeris_cyanoptera | 0 | 56 | 5 | 2567592.63 |
| birds | Brotogeris_jugularis | Brotogeris_jugularis | 0 | 63.2 | 5.3 | 781311.17 |
| birds | Brotogeris_pyrrhoptera | Brotogeris_pyrrhoptera | 1 | 63.9 | 5.3 | 25806.38 |
| birds | Brotogeris_tirica | Brotogeris_tirica | 0 | 63 | 4 | 288572.86 |
| birds | Brotogeris_versicolurus | Brotogeris_versicolurus | 0 | 71.5 | 4.5 | 714999.05 |
| birds | Brunhilda_charmosyna | Estrilda_charmosyna | 0 | 7.4 | 4 | 748636.98 |
| birds | Brunhilda_erythronotos | Estrilda_erythronotos | 0 | 9.2 | 4.2 | 2061140.19 |
| birds | Bubalornis_albirostris | Bubalornis_albirostris | 0 | 61.9 | 2.8 | 2320967.62 |
| birds | Bubo_africanus | Bubo_africanus | 0 | 645 | 2.8 | 8655406.6 |
| birds | Bubo_ascalaphus | Bubo_ascalaphus | 0 | 2100 | 2 | 3200038.62 |
| birds | Bubo_bengalensis | Bubo_bengalensis | 0 | 1100 | 3.2 | 3750743.18 |
| birds | Bubo_bubo | Bubo_bubo | 0 | 2668.5 | 2.8 | 31974373.19 |
| birds | Bubo_capensis | Bubo_capensis | 0 | 1118.6 | 1.7 | 1905194.15 |
| birds | Bubo_coromandus | Bubo_coromandus | 0 | 1392.2 | 1.7 | 1428965.17 |
| birds | Bubo_lacteus | Bubo_lacteus | 0 | 2114.9 | 1.4 | 11298632.02 |
| birds | Bubo_nipalensis | Bubo_nipalensis | 0 | 1396.4 | 1 | 2384160.56 |
| birds | Bubo_scandiacus | Bubo_scandiaca | 1 | 2028.8 | 3.9 | 27057680.29 |
| birds | Bubo_virginianus | Bubo_virginianus | 0 | 1354.5 | 1.7 | 24427057.98 |
| birds | Bucanetes_githagineus | Bucanetes_githagineus | 0 | 19.6 | 4.9 | 7293211.13 |
| birds | Bucanetes_mongolicus | Eremopsaltria_mongolicus | 0 | 21.5 | 4.9 | 4289647.08 |
| birds | Bucephala_albeola | Bucephala_albeola | 0 | 397.5 | 7.7 | 15692279.65 |
| birds | Bucephala_clangula | Bucephala_clangula | 0 | 918.6 | 9.4 | 37078063.74 |
| birds | Bucephala_islandica | Bucephala_islandica | 0 | 921.2 | 9.4 | 3452110.56 |
| birds | Buceros_bicornis | Buceros_bicornis | 1 | 2790.8 | 2 | 2951578.25 |
| birds | Buceros_hydrocorax | Buceros_hydrocorax | 1 | 1409.6 | 2.8 | 78462.72 |
| birds | Buceros_rhinoceros | Buceros_rhinoceros | 1 | 2371.6 | 1.4 | 1455791.75 |
| birds | Bucorvus_abyssinicus | Bucorvus_abyssinicus | 1 | 4000 | 2 | 4482170.58 |
| birds | Bucorvus_leadbeateri | Bucorvus_cafer | 1 | 3743.6 | 1.7 | 4867849.88 |
| birds | Bugeranus_carunculatus | Grus_carunculatus | 1 | 8159 | 2 | 2289042.19 |
| birds | Buphagus_africanus | Buphagus_africanus | 0 | 58.5 | 2.4 | 3672712.87 |
| birds | Buphagus_erythrorynchus | Buphagus_erythrorhynchus | 0 | 50.7 | 3.2 | 2881050.23 |
| birds | Burhinus_bistriatus | Burhinus_bistriatus | 0 | 787 | 2 | 787421.54 |

| | | | | | | |
|---|---|---|---|---|---|---|
| birds | Burhinus_capensis | Burhinus_capensis | 0 | 423 | 2 | 12421028.77 |
| birds | Burhinus_grallarius | Burhinus_grallarius | 0 | 648.1 | 2 | 2586160.81 |
| birds | Burhinus_oedicnemus | Burhinus_oedicnemus | 0 | 459 | 2 | 21340481.54 |
| birds | Burhinus_senegalensis | Burhinus_senegalensis | 0 | 326 | 2 | 7037522.76 |
| birds | Burhinus_superciliaris | Burhinus_superciliaris | 0 | 480 | 2 | 79703.93 |
| birds | Burhinus_vermiculatus | Burhinus_vermiculatus | 0 | 320 | 2 | 5802844.89 |
| birds | Butastur_indicus | Butastur_indicus | 0 | 414.6 | 2.8 | 6892597.52 |
| birds | Butastur_liventer | Butastur_liventer | 0 | 338 | 2.4 | 1357262.8 |
| birds | Butastur_rufipennis | Butastur_rufipennis | 0 | 334.9 | 1.7 | 7779786.48 |
| birds | Butastur_teesa | Butastur_teesa | 0 | 325 | 3 | 3969936.2 |
| birds | Buteo_albonotatus | Buteo_albonotatus | 0 | 745.9 | 2 | 8230721.24 |
| birds | Buteo_augur | Buteo_augur | 0 | 1099.2 | 2 | 3919632.94 |
| birds | Buteo_auguralis | Buteo_auguralis | 0 | 663.9 | 2.4 | 7090619.93 |
| birds | Buteo_brachypterus | Buteo_brachypterus | 0 | 511 | 2 | 591120.44 |
| birds | Buteo_brachyurus | Buteo_brachyurus | 0 | 496 | 2 | 14372802.04 |
| birds | Buteo_buteo | Buteo_buteo | 0 | 759.1 | 2.8 | 21382495.55 |
| birds | Buteo_galapagoensis | Buteo_galapagoensis | 1 | 1137.4 | 1.7 | 7978.71 |
| birds | Buteo_hemilasius | Buteo_hemilasius | 0 | 1358.9 | 2.8 | 9027045.28 |
| birds | Buteo_jamaicensis | Buteo_jamaicensis | 0 | 1101.2 | 2.4 | 15893934.47 |
| birds | Buteo_lagopus | Buteo_lagopus | 0 | 949.8 | 3.9 | 37406946.75 |
| birds | Buteo_lineatus | Buteo_lineatus | 0 | 603.7 | 3.2 | 4090576.97 |
| birds | Buteo_nitidus | Buteo_nitidus | 0 | 519 | 1.7 | 10412658.23 |
| birds | Buteo_oreophilus | Buteo_oreophilus | 0 | 700 | 2.4 | 674707.01 |
| birds | Buteo_platypterus | Buteo_platypterus | 0 | 453.7 | 2.4 | 12158210.01 |
| birds | Buteo_regalis | Buteo_regalis | 0 | 1437.2 | 3.5 | 4975226.36 |
| birds | Buteo_ridgwayi | Buteo_ridgwayi | 1 | 315 | 2 | 35683.59 |
| birds | Buteo_rufinus | Buteo_rufinus | 0 | 1166.2 | 2.8 | 20474801.17 |
| birds | Buteo_rufofuscus | Buteo_rufofuscus | 0 | 1530 | 2 | 1533534.66 |
| birds | Buteo_solitarius | Buteo_solitarius | 0 | 500.9 | 2.4 | 6957.48 |
| birds | Buteo_swainsoni | Buteo_swainsoni | 0 | 946.6 | 2.4 | 13792497.7 |
| birds | Buteogallus_aequinoctialis | Buteogallus_aequinoctialis | 0 | 657.6 | 1.4 | 132570.12 |
| birds | Buteogallus_anthracinus | Buteogallus_anthracinus | 0 | 975.1 | 1.7 | 1898078.97 |
| birds | Buteogallus_meridionalis | Buteogallus_meridionalis | 0 | 808 | 1 | 10432463.33 |
| birds | Buteogallus_urubitinga | Buteogallus_urubitinga | 0 | 1152.9 | 1.4 | 13126253.65 |

| | | | | | | |
|---|---|---|---|---|---|---|
| birds | Bycanistes_brevis | Bycanistes_brevis | 0 | 1232.8 | 1.4 | 834397.49 |
| birds | Bycanistes_bucinator | Bycanistes_bucinator | 0 | 639.4 | 2.8 | 4073212.84 |
| birds | Bycanistes_fistulator | Bycanistes_fistulator | 0 | 508.1 | 1.7 | 811024.82 |
| birds | Bycanistes_subcylindricus | Bycanistes_subcylindricus | 0 | 1195.4 | 2 | 1147243.51 |
| birds | Cacatua_alba | Cacatua_alba | 1 | 570 | 2.4 | 62157.62 |
| birds | Cacatua_galerita | Cacatua_galerita | 0 | 720.4 | 2.4 | 3967177.31 |
| birds | Cacatua_haematuropygia | Cacatua_haematuropygia | 1 | 288 | 2.4 | 216118.7 |
| birds | Cacatua_leadbeateri | Cacatua_leadbeateri | 0 | 396.8 | 3.2 | 2876418.66 |
| birds | Cacatua_moluccensis | Cacatua_moluccensis | 1 | 835 | 2 | 18717.29 |
| birds | Cacatua_ophthalmica | Cacatua_ophthalmica | 1 | 492 | 2 | 34781.17 |
| birds | Cacatua_pastinator | Cacatua_pastinator | 0 | 631.7 | 2.4 | 200538.76 |
| birds | Cacatua_sanguinea | Cacatua_sanguinea | 0 | 523.7 | 2.4 | 5317399.11 |
| birds | Cacatua_sulphurea | Cacatua_sulphurea | 1 | 303.3 | 2.4 | 1917.54 |
| birds | Cacatua_tenuirostris | Cacatua_tenuirostris | 0 | 566.6 | 2.4 | 562525.18 |
| birds | Cairina_moschata | Cairina_moschata | 0 | 2415.8 | 11 | 14998211.9 |
| birds | Calamanthus_campestris | Calamanthus_campestris | 0 | 13.9 | 3.5 | 1452514.49 |
| birds | Calamanthus_fuliginosus | Calamanthus_fuliginosus | 0 | 20.8 | 3.5 | 197682.93 |
| birds | Calamonastes_fasciolatus | Camaroptera_fasciolata | 0 | 13.1 | 2 | 1422725.53 |
| birds | Calamonastes_simplex | Camaroptera_simplex | 0 | 12 | 3.5 | 1535343.54 |
| birds | Calamonastes_undosus | Camaroptera_undosa | 0 | 13 | 2.4 | 1613961.16 |
| birds | Calamonastides_gracilirostris | Chloropeta_gracilirostris | 1 | 10.8 | 2.4 | 93067.22 |
| birds | Calamospiza_melanocorys | Calamospiza_melanocorys | 0 | 37.6 | 4.9 | 3695893.11 |
| birds | Calandrella_acutirostris | Calandrella_acutirostris | 0 | 20.2 | 2.8 | 5326192.97 |
| birds | Calandrella_brachydactyla | Calandrella_brachydactyla | 0 | 21.1 | 3.5 | 27542314.9 |
| birds | Calandrella_cinerea | Calandrella_cinerea | 0 | 23.7 | 2.8 | 5733287.57 |
| birds | Calcarius_lapponicus | Calcarius_lapponicus | 0 | 27.8 | 5.5 | 27580104.55 |
| birds | Calcarius_ornatus | Calcarius_ornatus | 1 | 20.3 | 4.9 | 3253026.17 |
| birds | Calcarius_pictus | Calcarius_pictus | 0 | 26.7 | 4 | 6213416.01 |
| birds | Calendulauda_africanoides | Mirafra_africanoides | 0 | 23.4 | 2.8 | 2859748.76 |
| birds | Calendulauda_albescens | Certhilauda_albescens | 0 | 30.4 | 2.8 | 283792.89 |
| birds | Calendulauda_burra | Certhilauda_burra | 1 | 37.4 | 2.4 | 72170.13 |
| birds | Calendulauda_erythrochlamys | Certhilauda_erythrochlamys | 0 | 28.1 | 1.4 | 70724.64 |
| birds | Calendulauda_poecilosterna | Mirafra_poecilosterna | 0 | 24.7 | 2 | 703693.95 |
| birds | Calendulauda_sabota | Mirafra_sabota | 0 | 24.7 | 2.8 | 2588939.44 |

| | | | | | | |
|---|---|---|---|---|---|---|
| birds | Calherodius_leuconotus | Gorsachius_leuconotus | 0 | 480.3 | 3.2 | 7860889.29 |
| birds | Calicalicus_madagascariensis | Calicalicus_madagascariensis | 0 | 16.4 | 2 | 363727.6 |
| birds | Calidris_acuminata | Calidris_acuminata | 1 | 66.1 | 4 | 12623150.23 |
| birds | Calidris_alba | Calidris_alba | 0 | 51.7 | 4 | 13217165.64 |
| birds | Calidris_alpina | Calidris_alpina | 0 | 51.9 | 4 | 21936163.39 |
| birds | Calidris_bairdii | Calidris_bairdii | 0 | 41 | 4 | 12711846.24 |
| birds | Calidris_canutus | Calidris_canutus | 0 | 141.9 | 3.5 | 6898573.86 |
| birds | Calidris_falcinellus | Limicola_falcinellus | 0 | 37.1 | 4 | 3247967.41 |
| birds | Calidris_fuscicollis | Calidris_fuscicollis | 0 | 43 | 4 | 27065084.74 |
| birds | Calidris_himantopus | Calidris_himantopus | 0 | 57.2 | 4 | 21365273.09 |
| birds | Calidris_maritima | Calidris_maritima | 0 | 65.1 | 4 | 3299407.83 |
| birds | Calidris_mauri | Calidris_mauri | 0 | 27.8 | 4 | 4634457.92 |
| birds | Calidris_melanotos | Calidris_melanotos | 0 | 79.7 | 4 | 33471970.38 |
| birds | Calidris_minuta | Calidris_minuta | 0 | 21.1 | 4 | 22084747.49 |
| birds | Calidris_minutilla | Calidris_minutilla | 0 | 22.9 | 4 | 25807753.61 |
| birds | Calidris_ptilocnemis | Calidris_ptilocnemis | 0 | 81.3 | 4 | 977273.69 |
| birds | Calidris_pugnax | Philomachus_pugnax | 0 | 131.7 | 4 | 28953892.13 |
| birds | Calidris_pusilla | Calidris_pusilla | 0 | 27.5 | 4 | 16118055.1 |
| birds | Calidris_ruficollis | Calidris_ruficollis | 0 | 26.8 | 4 | 9990778.18 |
| birds | Calidris_subminuta | Calidris_subminuta | 0 | 30.2 | 4 | 3687288.12 |
| birds | Calidris_subruficollis | Tryngites_subruficollis | 0 | 62.1 | 3.5 | 11646075.89 |
| birds | Calidris_temminckii | Calidris_temminckii | 0 | 23 | 4 | 17581833.28 |
| birds | Calidris_tenuirostris | Calidris_tenuirostris | 1 | 192 | 4 | 3461961.53 |
| birds | Calidris_virgata | Aphriza_virgata | 0 | 201 | 4 | 2291134.38 |
| birds | Caligavis_chrysops | Lichenostomus_chrysops | 0 | 17.3 | 2.4 | 1457474.71 |
| birds | Caligavis_obscura | Lichenostomus_obscurus | 0 | 26.4 | 2 | 173670.69 |
| birds | Calliope_calliope | Luscinia_calliope | 0 | 18.5 | 4.9 | 23381103.54 |
| birds | Calliope_obscura | Luscinia_obscura | 1 | 19.6 | 4 | 291670.81 |
| birds | Calliope_pectoralis | Luscinia_pectoralis | 0 | 23.3 | 3.5 | 3670860.21 |
| birds | Callipepla_californica | Callipepla_californica | 0 | 165.7 | 14.9 | 1691291.35 |
| birds | Callipepla_douglasii | Callipepla_douglasii | 0 | 177.1 | 9.8 | 167016.6 |
| birds | Callipepla_gambelii | Callipepla_gambelii | 0 | 166 | 13 | 558986.97 |
| birds | Callipepla_squamata | Callipepla_squamata | 0 | 183.9 | 13 | 1432736.3 |
| birds | Calliphlox_amethystina | Calliphlox_amethystina | 0 | 2.8 | 2 | 7988128.33 |

| | | | | | | |
|---|---|---|---|---|---|---|
| birds | Callocephalon_fimbriatum | Callocephalon_fimbriatum | 1 | 256 | 2 | 214821.22 |
| birds | Callonetta_leucophrys | Callonetta_leucophrys | 0 | 362.8 | 8.5 | 3218752.75 |
| birds | Caloenas_nicobarica | Caloenas_nicobarica | 0 | 537.9 | 1 | 475640.94 |
| birds | Caloperdix_oculeus | Caloperdix_oculeus | 0 | 209 | 8.9 | 534135.67 |
| birds | Caloramphus_fuliginosus | Calorhamphus_fuliginosus | 0 | 42.6 | 2.4 | 729941.66 |
| birds | Calothorax_lucifer | Calothorax_lucifer | 0 | 3.3 | 2 | 428359.87 |
| birds | Calothorax_pulcher | Calothorax_pulcher | 0 | 2.7 | 2 | 64442.51 |
| birds | Calypte_anna | Calypte_anna | 0 | 4.2 | 2 | 1270071.92 |
| birds | Calypte_costae | Calypte_costae | 0 | 3.1 | 2 | 645531.13 |
| birds | Calyptomena_hosii | Calyptomena_hosii | 0 | 92 | 2.8 | 146528.72 |
| birds | Calyptomena_viridis | Calyptomena_viridis | 0 | 58.5 | 1.7 | 1457426.48 |
| birds | Calyptomena_whiteheadi | Calyptomena_whiteheadi | 0 | 156.1 | 1.4 | 95505.36 |
| birds | Calyptorhynchus_banksii | Calyptorhynchus_banksii | 0 | 721.9 | 1 | 2798975.2 |
| birds | Calyptorhynchus_lathami | Calyptorhynchus_lathami | 1 | 437 | 1 | 661191.56 |
| birds | Camaroptera_brachyura | Camaroptera_brachyura | 0 | 9.7 | 3 | 15929835.83 |
| birds | Camaroptera_chloronota | Camaroptera_chloronota | 0 | 11.4 | 2 | 1659026 |
| birds | Camaroptera_superciliaris | Camaroptera_superciliaris | 0 | 9.7 | 3 | 3154212.79 |
| birds | Campephaga_flava | Campephaga_flava | 0 | 32.3 | 1.7 | 5048097.56 |
| birds | Campephaga_phoenicea | Campephaga_phoenicea | 0 | 27.1 | 2 | 4733993.8 |
| birds | Campephaga_quiscalina | Campephaga_quiscalina | 0 | 34.5 | 2 | 1373914.13 |
| birds | Campephilus_guatemalensis | Campephilus_guatemalensis | 0 | 242 | 2 | 888852.35 |
| birds | Campephilus_magellanicus | Campephilus_magellanicus | 0 | 324 | 2 | 395600.76 |
| birds | Campephilus_melanoleucos | Campephilus_melanoleucos | 0 | 256 | 2.4 | 11099024.68 |
| birds | Campephilus_rubricollis | Campephilus_rubricollis | 0 | 210.7 | 1 | 6911322.08 |
| birds | Campethera_abingoni | Campethera_abingoni | 0 | 67.5 | 2.4 | 7184316.96 |
| birds | Campethera_bennettii | Campethera_bennettii | 0 | 70.1 | 3 | 4469921.88 |
| birds | Campethera_cailliautii | Campethera_cailliautii | 0 | 41.1 | 2.4 | 3823971.24 |
| birds | Campethera_notata | Campethera_notata | 0 | 62 | 2.8 | 95428.78 |
| birds | Campethera_nubica | Campethera_nubica | 0 | 59.7 | 2.4 | 3241336.04 |
| birds | Campethera_punctuligera | Campethera_punctuligera | 0 | 71.8 | 2.4 | 4070381.33 |
| birds | Campocolinus_albogularis | Francolinus_albogularis | 0 | 276 | 6 | 1104886.46 |
| birds | Campocolinus_coqui | Francolinus_coqui | 0 | 262 | 4.5 | 4180135.39 |
| birds | Campocolinus_schlegelii | Francolinus_schlegelii | 0 | 236.6 | 3.2 | 553212.32 |
| birds | Camptostoma_imberbe | Camptostoma_imberbe | 0 | 7.4 | 2.8 | 1448668.84 |

| | | | | | | |
|---|---|---|---|---|---|---|
| birds | Camptostoma_obsoletum | Camptostoma_obsoletum | 0 | 8.1 | 2.4 | 14428561.74 |
| birds | Campylopterus_ensipennis | Campylopterus_ensipennis | 0 | 9.6 | 2 | 2431.08 |
| birds | Campylopterus_largipennis | Campylopterus_largipennis | 0 | 8.3 | 2 | 7116669.4 |
| birds | Campylorhynchus_brunneicapillus | Campylorhynchus_brunneicapillus | 0 | 38.9 | 3.9 | 1847263.76 |
| birds | Campylorhynchus_chiapensis | Campylorhynchus_chiapensis | 0 | 50.9 | 3 | 9934.36 |
| birds | Campylorhynchus_fasciatus | Campylorhynchus_fasciatus | 0 | 30 | 3.5 | 121819.89 |
| birds | Campylorhynchus_griseus | Campylorhynchus_griseus | 0 | 42.4 | 3.9 | 982980.09 |
| birds | Campylorhynchus_gularis | Campylorhynchus_gularis | 0 | 30.1 | 4 | 370562.24 |
| birds | Campylorhynchus_jocosus | Campylorhynchus_jocosus | 0 | 27.6 | 3.5 | 89805.27 |
| birds | Campylorhynchus_nuchalis | Campylorhynchus_nuchalis | 0 | 23.2 | 4 | 506629.36 |
| birds | Campylorhynchus_rufinucha | Campylorhynchus_rufinucha | 0 | 31 | 4.5 | 8261.17 |
| birds | Campylorhynchus_zonatus | Campylorhynchus_zonatus | 0 | 34.6 | 3.9 | 579082.34 |
| birds | Canachites_canadensis | Dendragapus_canadensis | 0 | 473.7 | 5.5 | 6068215.69 |
| birds | Canirallus_oculeus | Canirallus_oculeus | 0 | 277 | 2 | 1616653.93 |
| birds | Cantorchilus_leucotis | Thryothorus_leucotis | 0 | 19.4 | 2.4 | 8709608.57 |
| birds | Cantorchilus_modestus | Thryothorus_modestus | 0 | 17.6 | 2.4 | 263277.44 |
| birds | Cantorchilus_nigricapillus | Thryothorus_nigricapillus | 0 | 23.9 | 3 | 300564.01 |
| birds | Cantorchilus_semibadius | Thryothorus_semibadius | 0 | 17 | 2 | 14829.63 |
| birds | Cantorchilus_superciliaris | Thryothorus_superciliaris | 0 | 22.8 | 2.4 | 96843.25 |
| birds | Cantorchilus_thoracicus | Thryothorus_thoracicus | 0 | 17.6 | 2.4 | 90302.61 |
| birds | Capito_niger | Capito_niger | 0 | 57.4 | 3.5 | 1107267.21 |
| birds | Capito_squamatus | Capito_squamatus | 0 | 59.3 | 4 | 63414.19 |
| birds | Caprimulgus_aegyptius | Caprimulgus_aegyptius | 0 | 77.3 | 1.4 | 12441335.3 |
| birds | Caprimulgus_affinis | Caprimulgus_affinis | 0 | 58.7 | 1.4 | 5312716.77 |
| birds | Caprimulgus_batesi | Caprimulgus_batesi | 0 | 100 | 1 | 1859436.15 |
| birds | Caprimulgus_clarus | Caprimulgus_clarus | 0 | 42.5 | 2 | 1364940.55 |
| birds | Caprimulgus_climacurus | Caprimulgus_climacurus | 0 | 45.8 | 2 | 8738621.75 |
| birds | Caprimulgus_donaldsoni | Caprimulgus_donaldsoni | 0 | 29 | 2 | 751577.81 |
| birds | Caprimulgus_europaeus | Caprimulgus_europaeus | 0 | 67 | 1.4 | 29394935.22 |
| birds | Caprimulgus_eximius | Caprimulgus_eximius | 0 | 66.3 | 2 | 1981466.64 |
| birds | Caprimulgus_fossii | Caprimulgus_fossii | 0 | 58.3 | 1.4 | 6491736.25 |
| birds | Caprimulgus_fraenatus | Caprimulgus_fraenatus | 0 | 59.7 | 1.4 | 543293.67 |
| birds | Caprimulgus_indicus | Caprimulgus_indicus | 0 | 91.6 | 1.4 | 1651666.55 |
| birds | Caprimulgus_inornatus | Caprimulgus_inornatus | 0 | 50.3 | 2 | 8530651.41 |

| | | | | | | |
|---|---|---|---|---|---|---|
| birds | Caprimulgus_longipennis | Macrodipteryx_longipennis | 0 | 48 | 1.4 | 7317278.34 |
| birds | Caprimulgus_macrurus | Caprimulgus_macrurus | 0 | 66.2 | 1.4 | 5990101.35 |
| birds | Caprimulgus_madagascariensis | Caprimulgus_madagascariensis | 0 | 41.1 | 2 | 591293.27 |
| birds | Caprimulgus_manillensis | Caprimulgus_manillensis | 0 | 70.3 | 1.4 | 269038.47 |
| birds | Caprimulgus_natalensis | Caprimulgus_natalensis | 0 | 77.7 | 2 | 2207800.39 |
| birds | Caprimulgus_nubicus | Caprimulgus_nubicus | 0 | 49.3 | 1.4 | 713220.64 |
| birds | Caprimulgus_pectoralis | Caprimulgus_pectoralis | 0 | 51.4 | 1.4 | 5725005.93 |
| birds | Caprimulgus_poliocephalus | Caprimulgus_poliocephalus | 0 | 41.8 | 2 | 807034.95 |
| birds | Caprimulgus_pulchellus | Caprimulgus_pulchellus | 0 | 57.5 | 1.4 | 70914.3 |
| birds | Caprimulgus_ruficollis | Caprimulgus_ruficollis | 0 | 68.5 | 1.4 | 4900398.32 |
| birds | Caprimulgus_rufigena | Caprimulgus_rufigena | 0 | 52 | 1.4 | 3243389.39 |
| birds | Caprimulgus_tristigma | Caprimulgus_tristigma | 0 | 79 | 2 | 4557630.15 |
| birds | Caprimulgus_vexillarius | Macrodipteryx_vexillarius | 0 | 72.2 | 1.4 | 6625232.43 |
| birds | Capsiempis_flaveola | Capsiempis_flaveola | 0 | 7.7 | 2 | 5724076.64 |
| birds | Caracara_plancus | Caracara_plancus | 0 | 1078.6 | 2 | 12754773.65 |
| birds | Cardellina_canadensis | Wilsonia_canadensis | 0 | 10 | 3.9 | 7865400.5 |
| birds | Cardellina_pusilla | Wilsonia_pusilla | 0 | 7 | 4.9 | 18354114.38 |
| birds | Cardellina_rubrifrons | Cardellina_rubrifrons | 0 | 8.1 | 3.5 | 1056864.13 |
| birds | Cardinalis_cardinalis | Cardinalis_cardinalis | 0 | 42.6 | 3.5 | 5854913.93 |
| birds | Cardinalis_sinuatus | Cardinalis_sinuatus | 0 | 35.2 | 2.4 | 1519386.56 |
| birds | Carduelis_carduelis | Carduelis_carduelis | 0 | 16 | 4.9 | 15546092.39 |
| birds | Carduelis_citrinella | Carduelis_citrinella | 0 | 12 | 4.5 | 241931.15 |
| birds | Cariama_cristata | Cariama_cristata | 0 | 1400 | 2.4 | 5932013.38 |
| birds | Carpococcyx_renauldi | Carpococcyx_renauldi | 1 | 400 | 2.8 | 457300.49 |
| birds | Carpodacus_erythrinus | Carpodacus_erythrinus | 0 | 24 | 4.9 | 28438262.75 |
| birds | Carpodacus_rubicilla | Carpodacus_rubicilla | 0 | 43.2 | 4.5 | 3210935.5 |
| birds | Carpodacus_synoicus | Carpodacus_synoicus | 0 | 20.8 | 4.5 | 172774.39 |
| birds | Carpodectes_antoniae | Carpodectes_antoniae | 0 | 98 | 1 | 3915.87 |
| birds | Carpodectes_nitidus | Carpodectes_nitidus | 0 | 101.6 | 1 | 103015.66 |
| birds | Carterornis_leucotis | Monarcha_leucotis | 0 | 11.4 | 2 | 312561.63 |
| birds | Casuarius_bennetti | Casuarius_bennetti | 0 | 35000 | 3.9 | 396486.38 |
| birds | Casuarius_casuarius | Casuarius_casuarius | 0 | 44000 | 3.9 | 397042.35 |
| birds | Cathartes_aura | Cathartes_aura | 0 | 1518.2 | 2 | 28211569.03 |
| birds | Catharus_aurantiirostris | Catharus_aurantiirostris | 0 | 29.8 | 2 | 854579.17 |

| | | | | | | |
|---|---|---|---|---|---|---|
| birds | Catharus_bicknelli | Catharus_bicknelli | 1 | 27.8 | 3.5 | 49188.48 |
| birds | Catharus_frantzii | Catharus_frantzii | 0 | 28.9 | 2 | 283300.46 |
| birds | Catharus_fuscater | Catharus_fuscater | 0 | 33.7 | 2 | 313318.36 |
| birds | Catharus_fuscescens | Catharus_fuscescens | 0 | 31.9 | 3.5 | 16366792.63 |
| birds | Catharus_gracilirostris | Catharus_gracilirostris | 0 | 21 | 2 | 3827.25 |
| birds | Catharus_guttatus | Catharus_guttatus | 0 | 30.1 | 3.5 | 15387664.79 |
| birds | Catharus_mexicanus | Catharus_mexicanus | 0 | 33 | 2 | 143279.41 |
| birds | Catharus_minimus | Catharus_minimus | 0 | 31.6 | 3.5 | 17837621.98 |
| birds | Catharus_occidentalis | Catharus_occidentalis | 0 | 26.2 | 2.4 | 417881.84 |
| birds | Catharus_ustulatus | Catharus_ustulatus | 0 | 30.3 | 3.5 | 11035380.86 |
| birds | Catherpes_mexicanus | Catherpes_mexicanus | 0 | 12 | 6 | 4298192.46 |
| birds | Catriscus_brevirostris | Schoenicola_brevirostris | 0 | 15.7 | 2.4 | 2804794.87 |
| birds | Ceblepyris_caesius | Coracina_caesia | 0 | 47.3 | 1.4 | 905645.63 |
| birds | Ceblepyris_cinereus | Coracina_cinerea | 0 | 47.2 | 1 | 415959.23 |
| birds | Ceblepyris_pectoralis | Coracina_pectoralis | 0 | 54.4 | 1.4 | 6110219.71 |
| birds | Cecropis_abyssinica | Hirundo_abyssinica | 0 | 13.5 | 3 | 13676430.32 |
| birds | Cecropis_cucullata | Hirundo_cucullata | 0 | 27.1 | 3 | 5535896.41 |
| birds | Cecropis_daurica | Hirundo_daurica | 0 | 22.2 | 3.7 | 21306722.75 |
| birds | Cecropis_semirufa | Hirundo_semirufa | 0 | 30.1 | 3 | 8911972.36 |
| birds | Cecropis_senegalensis | Hirundo_senegalensis | 0 | 45.4 | 2.8 | 12912848.81 |
| birds | Celeus_castaneus | Celeus_castaneus | 0 | 85.6 | 3.5 | 396905.38 |
| birds | Celeus_elegans | Celeus_elegans | 0 | 135 | 3 | 6510380.16 |
| birds | Celeus_spectabilis | Celeus_spectabilis | 0 | 114 | 1 | 528390.73 |
| birds | Centrocercus_urophasianus | Centrocercus_urophasianus | 0 | 1902.5 | 7.5 | 1315889.14 |
| birds | Centropus_andamanensis | Centropus_andamanensis | 0 | 164 | 2.4 | 7211.9 |
| birds | Centropus_ateralbus | Centropus_ateralbus | 0 | 339 | 2 | 40984.98 |
| birds | Centropus_bengalensis | Centropus_bengalensis | 0 | 148.9 | 2.8 | 6030216.39 |
| birds | Centropus_bernsteini | Centropus_bernsteini | 0 | 162.1 | 2 | 516862.9 |
| birds | Centropus_chlororhynchos | Centropus_chlororhynchus | 1 | 244.4 | 2.4 | 11188.07 |
| birds | Centropus_cupreicaudus | Centropus_cupreicaudus | 0 | 285.2 | 2.8 | 1750330.64 |
| birds | Centropus_grillii | Centropus_grillii | 0 | 122.9 | 4.2 | 8530549.35 |
| birds | Centropus_leucogaster | Centropus_leucogaster | 0 | 307.8 | 2 | 1335483.71 |
| birds | Centropus_monachus | Centropus_monachus | 0 | 201.3 | 3 | 2920157.62 |
| birds | Centropus_nigrorufus | Centropus_nigrorufus | 1 | 244.4 | 1.7 | 28067.43 |

| | | | | | | |
|---|---|---|---|---|---|---|
| birds | Centropus_phasianinus | Centropus_phasianinus | 0 | 382 | 3.9 | 2665520.2 |
| birds | Centropus_senegalensis | Centropus_senegalensis | 0 | 169 | 3.2 | 9875812.37 |
| birds | Centropus_sinensis | Centropus_sinensis | 0 | 280.7 | 2.8 | 7931391.59 |
| birds | Centropus_superciliosus | Centropus_superciliosus | 0 | 148.5 | 3.9 | 7292900.16 |
| birds | Centropus_toulou | Centropus_toulou | 0 | 166.4 | 2.4 | 590995.81 |
| birds | Centropus_violaceus | Centropus_violaceus | 0 | 500 | 3 | 38717.48 |
| birds | Centropus_viridis | Centropus_viridis | 0 | 142 | 3 | 273885.38 |
| birds | Cephalopterus_glabricollis | Cephalopterus_glabricollis | 1 | 379.5 | 1 | 26873.01 |
| birds | Cephalopterus_ornatus | Cephalopterus_ornatus | 0 | 380 | 1 | 5506246.9 |
| birds | Cephalopterus_penduliger | Cephalopterus_penduliger | 1 | 338 | 1 | 56545.39 |
| birds | Ceratogymna_atrata | Ceratogymna_atrata | 0 | 1194.8 | 1.4 | 2282542.36 |
| birds | Ceratopipra_erythrocephala | Pipra_erythrocephala | 0 | 13.4 | 2 | 3456038.86 |
| birds | Ceratopipra_mentalis | Pipra_mentalis | 0 | 15 | 2 | 558740.47 |
| birds | Ceratopipra_rubrocapilla | Pipra_rubrocapilla | 0 | 12 | 2 | 3391760.24 |
| birds | Cercomacra_melanaria | Cercomacra_melanaria | 0 | 19 | 2 | 445529.08 |
| birds | Cercomacra_nigricans | Cercomacra_nigricans | 0 | 16.6 | 2 | 546994.42 |
| birds | Cercomacroides_tyrannina | Cercomacra_tyrannina | 0 | 16.3 | 2 | 3170819.2 |
| birds | Cercotrichas_galactotes | Erythropygia_galactotes | 0 | 20.3 | 3.5 | 18679450.74 |
| birds | Cercotrichas_hartlaubi | Erythropygia_hartlaubi | 0 | 19.2 | 2.4 | 612316.42 |
| birds | Cercotrichas_leucophrys | Erythropygia_leucophrys | 0 | 16.5 | 2.8 | 8126492.58 |
| birds | Cercotrichas_paena | Erythropygia_paena | 0 | 19.7 | 2.8 | 1909273.6 |
| birds | Cercotrichas_podobe | Cercotrichas_podobe | 0 | 23.9 | 2.8 | 3942700.06 |
| birds | Cereopsis_novaehollandiae | Cereopsis_novaehollandiae | 0 | 3560 | 4.2 | 595801.47 |
| birds | Certhia_americana | Certhia_americana | 0 | 8.1 | 5.5 | 11412723.75 |
| birds | Certhia_brachydactyla | Certhia_brachydactyla | 0 | 8.2 | 5.9 | 3199552.33 |
| birds | Certhia_discolor | Certhia_discolor | 0 | 10.4 | 3.9 | 173809.74 |
| birds | Certhia_familiaris | Certhia_familiaris | 0 | 9 | 4.2 | 15914424.48 |
| birds | Certhia_himalayana | Certhia_himalayana | 0 | 8.8 | 4.9 | 1333092.38 |
| birds | Certhiaxis_cinnamomeus | Certhiaxis_cinnamomeus | 0 | 15.2 | 3.2 | 8364757.31 |
| birds | Certhilauda_curvirostris | Certhilauda_curvirostris | 0 | 40.4 | 2.4 | 63461.48 |
| birds | Certhionyx_variegatus | Certhionyx_variegatus | 0 | 26.3 | 2.4 | 5122091.98 |
| birds | Ceryle_rudis | Ceryle_rudis | 0 | 84.4 | 4.5 | 23875911.23 |
| birds | Cettia_brunnifrons | Cettia_brunnifrons | 0 | 7.3 | 3.9 | 1378233.28 |
| birds | Cettia_castaneocoronata | Tesia_castaneocoronata | 0 | 8.9 | 2 | 1980546.83 |

| | | | | | | |
|---|---|---|---|---|---|---|
| birds | Cettia_cetti | Cettia_cetti | 0 | 13.2 | 4.5 | 5830320.2 |
| birds | Ceuthmochares_aereus | Ceuthmochares_aereus | 0 | 65.8 | 2.4 | 4005421.1 |
| birds | Ceyx_azureus | Alcedo_azurea | 0 | 34.9 | 5.3 | 3398489.47 |
| birds | Ceyx_erithaca | Ceyx_erithaca | 0 | 17.8 | 4.6 | 3178476.07 |
| birds | Ceyx_lepidus | Ceyx_lepidus | 0 | 21.3 | 2 | 43842.15 |
| birds | Ceyx_pusillus | Alcedo_pusilla | 0 | 13.3 | 3.9 | 910124.54 |
| birds | Chaetops_frenatus | Chaetops_frenatus | 0 | 55 | 2 | 59624.79 |
| birds | Chaetura_brachyura | Chaetura_brachyura | 0 | 18.3 | 3.7 | 7468188.25 |
| birds | Chaetura_chapmani | Chaetura_chapmani | 0 | 23.8 | 2 | 4701276.36 |
| birds | Chaetura_cinereiventris | Chaetura_cinereiventris | 0 | 13.9 | 2.8 | 6102705.51 |
| birds | Chaetura_pelagica | Chaetura_pelagica | 1 | 23.6 | 3.7 | 11481187.63 |
| birds | Chaetura_vauxi | Chaetura_vauxi | 0 | 17.1 | 2.6 | 3704613.51 |
| birds | Chalcomitra_adelberti | Nectarinia_adelberti | 0 | 9.8 | 4 | 915043.57 |
| birds | Chalcomitra_amethystina | Nectarinia_amethystina | 0 | 13.7 | 2 | 3590561.91 |
| birds | Chalcomitra_fuliginosa | Nectarinia_fuliginosa | 0 | 11.9 | 2 | 263939.43 |
| birds | Chalcomitra_hunteri | Nectarinia_hunteri | 0 | 11.8 | 1.4 | 1422643.43 |
| birds | Chalcomitra_rubescens | Nectarinia_rubescens | 0 | 10.3 | 2 | 2910535.16 |
| birds | Chalcomitra_senegalensis | Nectarinia_senegalensis | 0 | 12.2 | 1.7 | 9424883.18 |
| birds | Chalcoparia_singalensis | Anthreptes_singalensis | 0 | 8.2 | 2 | 2829894.28 |
| birds | Chalcophaps_stephani | Chalcophaps_stephani | 0 | 129 | 2 | 866814.34 |
| birds | Chamaea_fasciata | Chamaea_fasciata | 0 | 14.7 | 3.9 | 246063.25 |
| birds | Chamaetylas_fuelleborni | Alethe_fuelleborni | 0 | 52 | 2 | 189262.11 |
| birds | Chamaetylas_poliocephala | Alethe_poliocephala | 0 | 32.4 | 1.7 | 1599692.41 |
| birds | Chamaetylas_poliophrys | Alethe_poliophrys | 0 | 35.2 | 2 | 76094.57 |
| birds | Chamaeza_campanisona | Chamaeza_campanisona | 0 | 90.6 | 2 | 1801868.24 |
| birds | Charadrius_alexandrinus | Charadrius_alexandrinus | 0 | 42.3 | 3 | 22446543.51 |
| birds | Charadrius_asiaticus | Charadrius_asiaticus | 0 | 77.1 | 3 | 18276948.61 |
| birds | Charadrius_bicinctus | Charadrius_bicinctus | 0 | 59.4 | 3 | 1655193.86 |
| birds | Charadrius_collaris | Charadrius_collaris | 0 | 28.3 | 2 | 15100843.49 |
| birds | Charadrius_dubius | Charadrius_dubius | 0 | 38.7 | 4 | 46401384.75 |
| birds | Charadrius_falklandicus | Charadrius_falklandicus | 0 | 65 | 3 | 1010983.2 |
| birds | Charadrius_hiaticula | Charadrius_hiaticula | 0 | 64 | 3.5 | 28297283.64 |
| birds | Charadrius_leschenaultii | Charadrius_leschenaultii | 0 | 74.8 | 3 | 8942028.55 |
| birds | Charadrius_marginatus | Charadrius_marginatus | 0 | 48.3 | 2 | 4442943.8 |

| | | | | | | |
|---|---|---|---|---|---|---|
| birds | Charadrius_melodus | Charadrius_melodus | 0 | 54.4 | 4 | 5351896.5 |
| birds | Charadrius_modestus | Charadrius_modestus | 0 | 78 | 2 | 1462237.28 |
| birds | Charadrius_mongolus | Charadrius_mongolus | 1 | 64 | 3 | 61116255.06 |
| birds | Charadrius_montanus | Charadrius_montanus | 0 | 95.7 | 3 | 2169361.82 |
| birds | Charadrius_obscurus | Charadrius_obscurus | 1 | 146 | 3 | 311.44 |
| birds | Charadrius_pallidus | Charadrius_pallidus | 0 | 34.8 | 2 | 480800.22 |
| birds | Charadrius_pecuarius | Charadrius_pecuarius | 0 | 34 | 2 | 16351827.57 |
| birds | Charadrius_peronii | Charadrius_peronii | 0 | 42 | 3 | 1266559.93 |
| birds | Charadrius_placidus | Charadrius_placidus | 0 | 62.7 | 4 | 5396919.96 |
| birds | Charadrius_ruficapillus | Charadrius_ruficapillus | 0 | 37.4 | 2 | 4811331.66 |
| birds | Charadrius_semipalmatus | Charadrius_semipalmatus | 0 | 46.7 | 3.5 | 17838316.67 |
| birds | Charadrius_thoracicus | Charadrius_thoracicus | 1 | 11.5 | 2 | 14469.49 |
| birds | Charadrius_tricollaris | Charadrius_tricollaris | 0 | 31.2 | 2 | 9455723.89 |
| birds | Charadrius_vociferus | Charadrius_vociferus | 0 | 96.4 | 4 | 16664790.81 |
| birds | Charadrius_wilsonia | Charadrius_wilsonia | 0 | 55.1 | 3 | 966205.6 |
| birds | Chasiempis_sandwichensis | Chasiempis_sandwichensis | 0 | 13.7 | 1.7 | 10507.22 |
| birds | Chauna_chavaria | Chauna_chavaria | 0 | 2557.6 | 3.9 | 114400.07 |
| birds | Chauna_torquata | Chauna_torquata | 0 | 4400 | 3.9 | 3828524.39 |
| birds | Chelictinia_riocourii | Chelictinia_riocourii | 1 | 110 | 4 | 6292258.57 |
| birds | Chelidoptera_tenebrosa | Chelidoptera_tenebrosa | 0 | 35.9 | 1.4 | 8690116.86 |
| birds | Chelidorhynx_hypoxanthus | Chelidorhynx_hypoxantha | 0 | 6 | 3 | 1909981.69 |
| birds | Chenonetta_jubata | Chenonetta_jubata | 0 | 807.5 | 9.4 | 5996685.1 |
| birds | Cheramoeca_leucosterna | Cheramoeca_leucosterna | 0 | 14.1 | 4.9 | 5215727.85 |
| birds | Chersomanes_albofasciata | Chersomanes_albofasciata | 0 | 24.6 | 2.4 | 2153588.69 |
| birds | Chionomesa_fimbriata | Amazilia_fimbriata | 0 | 4.9 | 2 | 10740118.89 |
| birds | Chiroxiphia_caudata | Chiroxiphia_caudata | 0 | 25.6 | 2 | 1459549.86 |
| birds | Chiroxiphia_lanceolata | Chiroxiphia_lanceolata | 0 | 18.5 | 2 | 420162.32 |
| birds | Chiroxiphia_linearis | Chiroxiphia_linearis | 0 | 17.4 | 2 | 101728.76 |
| birds | Chiroxiphia_pareola | Chiroxiphia_pareola | 0 | 16.8 | 2 | 5040298.89 |
| birds | Chlamydera_cerviniventris | Chlamydera_cerviniventris | 0 | 156.4 | 1 | 152782.31 |
| birds | Chlamydera_guttata | Chlamydera_guttata | 0 | 136 | 2 | 1491137.42 |
| birds | Chlamydera_lauterbachi | Chlamydera_lauterbachi | 0 | 123.2 | 1 | 124403.09 |
| birds | Chlamydera_maculata | Chlamydera_maculata | 0 | 140 | 2.4 | 1784471.92 |
| birds | Chlamydera_nuchalis | Chlamydera_nuchalis | 0 | 200.4 | 1.4 | 1285385.85 |

| | | | | | | |
|---|---|---|---|---|---|---|
| birds | Chlamydotis_undulata | Chlamydotis_undulata | 1 | 1501.3 | 2.4 | 4448380.49 |
| birds | Chloebia_gouldiae | Erythrura_gouldiae | 0 | 13.8 | 5.7 | 1163550.38 |
| birds | Chloephaga_hybrida | Chloephaga_hybrida | 0 | 2281.8 | 4.6 | 219432.23 |
| birds | Chloephaga_melanoptera | Chloephaga_melanoptera | 0 | 2037 | 7.1 | 1159462.3 |
| birds | Chloephaga_picta | Chloephaga_picta | 0 | 2880.4 | 6.3 | 1582798.87 |
| birds | Chloephaga_poliocephala | Chloephaga_poliocephala | 0 | 2007.1 | 4.9 | 1643991.77 |
| birds | Chloephaga_rubidiceps | Chloephaga_rubidiceps | 0 | 2086 | 6.3 | 248383.81 |
| birds | Chlorestes_candida | Amazilia_candida | 0 | 3.8 | 2 | 564871.44 |
| birds | Chlorestes_cyanus | Hylocharis_cyanus | 0 | 3.3 | 2 | 7031990.78 |
| birds | Chlorestes_eliciae | Hylocharis_eliciae | 0 | 3.6 | 2 | 306966.67 |
| birds | Chlorestes_julie | Damophila_julie | 0 | 3.3 | 2 | 289656.61 |
| birds | Chlorestes_notata | Chlorestes_notata | 0 | 4.2 | 2 | 4700543.44 |
| birds | Chloris_chloris | Carduelis_chloris | 0 | 26 | 4.9 | 11398923.39 |
| birds | Chloris_sinica | Carduelis_sinica | 0 | 18.6 | 3.2 | 6355782.17 |
| birds | Chloris_spinoides | Carduelis_spinoides | 0 | 18.6 | 3.9 | 557286.26 |
| birds | Chloroceryle_aenea | Chloroceryle_aenea | 0 | 13.8 | 3.5 | 9332280.13 |
| birds | Chloroceryle_amazona | Chloroceryle_amazona | 0 | 126.4 | 3.5 | 14157184.63 |
| birds | Chloroceryle_americana | Chloroceryle_americana | 0 | 33.7 | 4.5 | 16946715.06 |
| birds | Chloroceryle_inda | Chloroceryle_inda | 0 | 52.1 | 3.9 | 8573926.25 |
| birds | Chlorocichla_falkensteini | Chlorocichla_falkensteini | 0 | 31.7 | 2 | 978923.36 |
| birds | Chlorocichla_flaviventris | Chlorocichla_flaviventris | 0 | 39.4 | 2 | 3986239.89 |
| birds | Chlorocichla_simplex | Chlorocichla_simplex | 0 | 46.4 | 2 | 4003835.48 |
| birds | Chlorophoneus_bocagei | Telophorus_bocagei | 0 | 27 | 2 | 785394.39 |
| birds | Chlorophoneus_olivaceus | Telophorus_olivaceus | 0 | 33.6 | 2 | 332431.69 |
| birds | Chlorophoneus_sulfureopectus | Telophorus_sulfureopectus | 0 | 27.1 | 2 | 8777553.1 |
| birds | Chloropsis_aurifrons | Chloropsis_aurifrons | 0 | 32.3 | 2.4 | 3471632.85 |
| birds | Chloropsis_hardwickii | Chloropsis_hardwickii | 0 | 32.3 | 2.4 | 902456.73 |
| birds | Chlorostilbon_lucidus | Chlorostilbon_lucidus | 0 | 3.5 | 2 | 5788276.16 |
| birds | Chlorostilbon_mellisugus | Chlorostilbon_mellisugus | 0 | 2.6 | 2 | 4111804.66 |
| birds | Chlorostilbon_poortmani | Chlorostilbon_poortmani | 0 | 3.3 | 2 | 60449.07 |
| birds | Chlorostilbon_russatus | Chlorostilbon_russatus | 0 | 3.4 | 2 | 27765.98 |
| birds | Chlorostilbon_stenurus | Chlorostilbon_stenurus | 0 | 3.4 | 2 | 47007.36 |
| birds | Chondestes_grammacus | Chondestes_grammacus | 0 | 29 | 4.5 | 7065667.03 |
| birds | Chondrohierax_uncinatus | Chondrohierax_uncinatus | 0 | 286.1 | 1.4 | 9787616.08 |

| | | | | | | |
|---|---|---|---|---|---|---|
| birds | Chordeiles_acutipennis | Chordeiles_acutipennis | 0 | 48.4 | 1.4 | 10717818.68 |
| birds | Chordeiles_gundlachii | Chordeiles_gundlachii | 0 | 50 | 1.4 | 228316.66 |
| birds | Chordeiles_minor | Chordeiles_minor | 0 | 79.3 | 1.7 | 28501967.68 |
| birds | Chordeiles_nacunda | Podager_nacunda | 0 | 159 | 1.4 | 12270759.1 |
| birds | Chordeiles_pusillus | Chordeiles_pusillus | 0 | 30.6 | 1 | 4968284.38 |
| birds | Chordeiles_rupestris | Chordeiles_rupestris | 0 | 52.3 | 1.4 | 1639560.09 |
| birds | Chrysocolaptes_festivus | Chrysocolaptes_festivus | 0 | 213 | 1.7 | 1703319.4 |
| birds | Chrysocolaptes_lucidus | Chrysocolaptes_lucidus | 0 | 142 | 3.2 | 119803.15 |
| birds | Chrysocolaptes_validus | Reinwardtipicus_validus | 0 | 149 | 1.4 | 1291734.8 |
| birds | Chrysoena_luteovirens | Ptilinopus_luteovirens | 0 | 46.6 | 1 | 9129.14 |
| birds | Chrysoena_victor | Ptilinopus_victor | 0 | 46.6 | 1 | 6108.65 |
| birds | Chrysolampis_mosquitus | Chrysolampis_mosquitus | 0 | 3.9 | 2 | 7397775.67 |
| birds | Chrysolophus_amherstiae | Chrysolophus_amherstiae | 0 | 732.5 | 6.5 | 387085.42 |
| birds | Chrysolophus_pictus | Chrysolophus_pictus | 0 | 555.2 | 7.7 | 975613.44 |
| birds | Chrysomus_icterocephalus | Chrysomus_icterocephalus | 0 | 30.7 | 2.4 | 1581439.98 |
| birds | Chrysophlegma_flavinucha | Picus_flavinucha | 0 | 176.8 | 2.8 | 2932266.06 |
| birds | Chrysophlegma_mentale | Picus_mentalis | 0 | 99.7 | 2.4 | 40422.48 |
| birds | Chrysophlegma_miniaceum | Picus_mineaceus | 0 | 97.3 | 2.4 | 1291605.29 |
| birds | Chrysuronia_brevirostris | Amazilia_brevirostris | 0 | 4.3 | 2 | 584285.4 |
| birds | Chrysuronia_coeruleogularis | Lepidopyga_coeruleogularis | 0 | 4.2 | 2 | 91772.24 |
| birds | Chrysuronia_leucogaster | Amazilia_leucogaster | 0 | 4.7 | 2 | 800745 |
| birds | Chrysuronia_versicolor | Amazilia_versicolor | 0 | 4.1 | 2 | 6914568.21 |
| birds | Ciccaba_nigrolineata | Strix_nigrolineata | 0 | 457.9 | 1.4 | 991396.19 |
| birds | Ciccaba_virgata | Strix_virgata | 0 | 284 | 1.4 | 11001158.19 |
| birds | Cichladusa_arquata | Cichladusa_arquata | 0 | 34.2 | 2.4 | 2314814.62 |
| birds | Cichladusa_guttata | Cichladusa_guttata | 0 | 23.1 | 2.4 | 1373382.96 |
| birds | Cichladusa_ruficauda | Cichladusa_ruficauda | 0 | 28.5 | 2 | 699193.3 |
| birds | Ciconia_abdimii | Ciconia_abdimii | 0 | 1398 | 2.4 | 17648125.86 |
| birds | Ciconia_boyciana | Ciconia_boyciana | 1 | 4847.7 | 3.5 | 3630446.15 |
| birds | Ciconia_ciconia | Ciconia_ciconia | 0 | 3445.8 | 2.6 | 30530843.13 |
| birds | Ciconia_episcopus | Ciconia_episcopus | 0 | 2061 | 2.8 | 3697795.59 |
| birds | Ciconia_maguari | Ciconia_maguari | 0 | 3995 | 2.8 | 5190878.3 |
| birds | Ciconia_nigra | Ciconia_nigra | 0 | 2926 | 3.5 | 35630654.88 |
| birds | Cinclodes_antarcticus | Cinclodes_antarcticus | 0 | 63.2 | 2.4 | 12926.55 |

| | | | | | | |
|---|---|---|---|---|---|---|
| birds | Cinclodes_atacamensis | Cinclodes_atacamensis | 0 | 53 | 2 | 870994.25 |
| birds | Cinclodes_comechingonus | Cinclodes_comechingonus | 0 | 28.3 | 2 | 44865.65 |
| birds | Cinclodes_excelsior | Cinclodes_excelsior | 0 | 64 | 2 | 37973.19 |
| birds | Cinclodes_fuscus | Cinclodes_fuscus | 0 | 30 | 2.4 | 2507430.84 |
| birds | Cinclodes_nigrofumosus | Cinclodes_nigrofumosus | 0 | 65 | 3.5 | 74685.21 |
| birds | Cinclodes_oustaleti | Cinclodes_oustaleti | 0 | 30.5 | 2.4 | 734371.46 |
| birds | Cinclodes_patagonicus | Cinclodes_patagonicus | 0 | 30.7 | 2.4 | 661640.55 |
| birds | Cincloramphus_cruralis | Cincloramphus_cruralis | 0 | 49.2 | 3.5 | 6388749.63 |
| birds | Cincloramphus_mariae | Megalurulus_mariei | 0 | 25 | 1.7 | 16389.77 |
| birds | Cincloramphus_mathewsi | Cincloramphus_mathewsi | 0 | 29.3 | 3.5 | 6668777.97 |
| birds | Cincloramphus_rubiginosus | Megalurulus_rubiginosus | 0 | 41.2 | 2 | 33761.38 |
| birds | Cincloramphus_timoriensis | Megalurus_timoriensis | 0 | 21.6 | 2.8 | 2127266.89 |
| birds | Cinclosoma_castaneothorax | Cinclosoma_castaneothorax | 0 | 62.4 | 2.4 | 488478.66 |
| birds | Cinclosoma_castanotum | Cinclosoma_castanotum | 0 | 73.3 | 2 | 2823366.52 |
| birds | Cinclosoma_cinnamomeum | Cinclosoma_cinnamomeum | 0 | 56.1 | 2.4 | 764829.56 |
| birds | Cinclosoma_punctatum | Cinclosoma_punctatum | 0 | 113.3 | 2.4 | 567920.58 |
| birds | Cinclus_cinclus | Cinclus_cinclus | 0 | 61.5 | 4.5 | 10252132.11 |
| birds | Cinclus_leucocephalus | Cinclus_leucocephalus | 0 | 44 | 2 | 619987.76 |
| birds | Cinclus_mexicanus | Cinclus_mexicanus | 0 | 56.7 | 3.2 | 4972505.5 |
| birds | Cinclus_pallasii | Cinclus_pallasii | 0 | 76 | 4.5 | 9425465.17 |
| birds | Cinclus_schulzii | Cinclus_schulzi | 1 | 37.6 | 2 | 133757.16 |
| birds | Cinnyricinclus_leucogaster | Cinnyricinclus_leucogaster | 0 | 44 | 2.8 | 12754823.21 |
| birds | Cinnyris_afer | Nectarinia_afra | 0 | 11.1 | 2 | 365697.14 |
| birds | Cinnyris_asiaticus | Nectarinia_asiatica | 0 | 8.1 | 1.7 | 5816862.3 |
| birds | Cinnyris_batesi | Nectarinia_batesi | 0 | 6.3 | 1.4 | 854825.13 |
| birds | Cinnyris_bifasciatus | Nectarinia_bifasciata | 0 | 7.2 | 1.4 | 1927794.28 |
| birds | Cinnyris_bouvieri | Nectarinia_bouvieri | 0 | 8.7 | 2 | 360313.74 |
| birds | Cinnyris_chalybeus | Nectarinia_chalybea | 0 | 7.7 | 1.7 | 496199.25 |
| birds | Cinnyris_chloropygius | Nectarinia_chloropygia | 0 | 6.2 | 1.7 | 4297123.95 |
| birds | Cinnyris_coccinigastrus | Nectarinia_coccinigaster | 0 | 14.3 | 1.4 | 1740703.51 |
| birds | Cinnyris_cupreus | Nectarinia_cuprea | 0 | 8.9 | 1.7 | 7848077.34 |
| birds | Cinnyris_erythrocercus | Nectarinia_erythrocerca | 0 | 8.8 | 1.7 | 438707.33 |
| birds | Cinnyris_fuscus | Nectarinia_fusca | 0 | 7.8 | 2.4 | 1120628.47 |
| birds | Cinnyris_habessinicus | Nectarinia_habessinica | 0 | 9.5 | 1.4 | 1107754.72 |

| | | | | | | |
|---|---|---|---|---|---|---|
| birds | Cinnyris_johannae | Nectarinia_johannae | 0 | 13 | 1.4 | 1631281.91 |
| birds | Cinnyris_jugularis | Nectarinia_jugularis | 0 | 9 | 2.4 | 5063701.81 |
| birds | Cinnyris_lotenius | Nectarinia_lotenia | 0 | 8.8 | 2.4 | 292016.32 |
| birds | Cinnyris_manoensis | Nectarinia_manoensis | 0 | 9.5 | 1.7 | 454948.66 |
| birds | Cinnyris_mariquensis | Nectarinia_mariquensis | 0 | 11.5 | 1.7 | 2606050.21 |
| birds | Cinnyris_mediocris | Nectarinia_mediocris | 0 | 8.2 | 1.4 | 195956.74 |
| birds | Cinnyris_minullus | Nectarinia_minulla | 0 | 5.7 | 1.4 | 1198202.19 |
| birds | Cinnyris_nectarinioides | Nectarinia_nectarinioides | 0 | 4.8 | 1.4 | 496613.1 |
| birds | Cinnyris_neergaardi | Nectarinia_neergardi | 0 | 6.1 | 2 | 138996.95 |
| birds | Cinnyris_notatus | Nectarinia_notata | 0 | 14.6 | 1.4 | 590045.88 |
| birds | Cinnyris_osea | Nectarinia_osea | 0 | 7.2 | 1.7 | 568902.39 |
| birds | Cinnyris_pulchellus | Nectarinia_pulchella | 0 | 7 | 1.4 | 5431378.76 |
| birds | Cinnyris_regius | Nectarinia_regia | 0 | 6.6 | 1 | 76601.79 |
| birds | Cinnyris_reichenowi | Nectarinia_preussi | 0 | 6.8 | 1.4 | 243088.55 |
| birds | Cinnyris_shelleyi | Nectarinia_shelleyi | 0 | 9 | 1.4 | 416193.99 |
| birds | Cinnyris_sovimanga | Nectarinia_sovimanga | 0 | 7.2 | 1.7 | 591013.44 |
| birds | Cinnyris_superbus | Nectarinia_superba | 0 | 16.8 | 1.4 | 2854180.16 |
| birds | Cinnyris_talatala | Nectarinia_talatala | 0 | 8.1 | 1.4 | 2820942.59 |
| birds | Cinnyris_ursulae | Nectarinia_ursulae | 0 | 6.5 | 1.4 | 27769.05 |
| birds | Cinnyris_venustus | Nectarinia_venusta | 0 | 6.6 | 1.7 | 5912223.61 |
| birds | Circaetus_cinerascens | Circaetus_cinerascens | 0 | 1126 | 1 | 5762003 |
| birds | Circaetus_cinereus | Circaetus_cinereus | 0 | 2048 | 1 | 11541044.04 |
| birds | Circaetus_fasciolatus | Circaetus_fasciolatus | 0 | 1013.6 | 1 | 734455.97 |
| birds | Circaetus_gallicus | Circaetus_gallicus | 0 | 1699.1 | 1 | 24616860.1 |
| birds | Circus_aeruginosus | Circus_aeruginosus | 0 | 704.1 | 4.2 | 38081226.31 |
| birds | Circus_approximans | Circus_approximans | 0 | 746.2 | 3.5 | 5134943.65 |
| birds | Circus_assimilis | Circus_assimilis | 0 | 558.6 | 3 | 7817547.59 |
| birds | Circus_buffoni | Circus_buffoni | 0 | 507.4 | 3.5 | 6350653.75 |
| birds | Circus_cinereus | Circus_cinereus | 0 | 399.7 | 3.5 | 3705694.07 |
| birds | Circus_cyaneus | Circus_cyaneus | 0 | 393 | 4.2 | 36969892.74 |
| birds | Circus_macrourus | Circus_macrourus | 0 | 384.4 | 4.5 | 46915251.63 |
| birds | Circus_maurus | Circus_maurus | 1 | 509 | 3.5 | 1464708.15 |
| birds | Circus_melanoleucos | Circus_melanoleucos | 0 | 336.3 | 4.5 | 13367833.88 |
| birds | Circus_pygargus | Circus_pygargus | 0 | 310.8 | 3.9 | 30385902.22 |

| | | | | | | |
|---|---|---|---|---|---|---|
| birds | Circus_ranivorus | Circus_ranivorus | 0 | 507 | 3.5 | 4426338.22 |
| birds | Circus_spilonotus | Circus_spilonotus | 0 | 558.8 | 4.6 | 11639585.51 |
| birds | Cissomela_pectoralis | Certhionyx_pectoralis | 0 | 10.1 | 2 | 1086177.68 |
| birds | Cisticola_aberdare | Cisticola_aberdare | 1 | 24 | 2 | 1272.48 |
| birds | Cisticola_aberrans | Cisticola_aberrans | 0 | 14.2 | 2.4 | 2482236.37 |
| birds | Cisticola_anonymus | Cisticola_anonymus | 0 | 13.9 | 2.4 | 2357531.94 |
| birds | Cisticola_aridulus | Cisticola_aridulus | 0 | 9.2 | 2.8 | 4468382.38 |
| birds | Cisticola_ayresii | Cisticola_ayresii | 0 | 7 | 2.2 | 1258911.91 |
| birds | Cisticola_bodessa | Cisticola_bodessa | 0 | 17.1 | 4 | 437584.83 |
| birds | Cisticola_brachypterus | Cisticola_brachypterus | 0 | 7.8 | 3 | 7674053.25 |
| birds | Cisticola_brunnescens | Cisticola_brunnescens | 0 | 8 | 2.8 | 465384.19 |
| birds | Cisticola_bulliens | Cisticola_bulliens | 0 | 15.8 | 3 | 308376.91 |
| birds | Cisticola_cantans | Cisticola_cantans | 0 | 11.7 | 3 | 5670957.17 |
| birds | Cisticola_carruthersi | Cisticola_carruthersi | 0 | 11.2 | 2.8 | 174334.6 |
| birds | Cisticola_cherina | Cisticola_cherina | 0 | 10 | 3.9 | 591128.96 |
| birds | Cisticola_chiniana | Cisticola_chiniana | 0 | 16.1 | 3.2 | 6716168.11 |
| birds | Cisticola_chubbi | Cisticola_chubbi | 0 | 16 | 2.4 | 322235.69 |
| birds | Cisticola_cinereolus | Cisticola_cinereolus | 0 | 17 | 4 | 614939.78 |
| birds | Cisticola_dambo | Cisticola_dambo | 0 | 9 | 2 | 472758.51 |
| birds | Cisticola_erythrops | Cisticola_erythrops | 0 | 14.8 | 2.8 | 6407914.95 |
| birds | Cisticola_exilis | Cisticola_exilis | 0 | 7 | 3.2 | 7177349 |
| birds | Cisticola_eximius | Cisticola_eximius | 0 | 12 | 3.5 | 674103.66 |
| birds | Cisticola_fulvicapilla | Cisticola_fulvicapilla | 0 | 8.2 | 3.5 | 4604333.83 |
| birds | Cisticola_galactotes | Cisticola_galactotes | 0 | 13.1 | 3.5 | 287146.87 |
| birds | Cisticola_hunteri | Cisticola_hunteri | 0 | 15.5 | 2 | 156331.32 |
| birds | Cisticola_juncidis | Cisticola_juncidis | 0 | 6.8 | 3.7 | 18929166.75 |
| birds | Cisticola_lais | Cisticola_lais | 0 | 14.8 | 2 | 881974.02 |
| birds | Cisticola_lateralis | Cisticola_lateralis | 0 | 16.8 | 2.4 | 3066342.45 |
| birds | Cisticola_nana | Cisticola_nanus | 0 | 5 | 3.5 | 685105.62 |
| birds | Cisticola_natalensis | Cisticola_natalensis | 0 | 16.4 | 2.4 | 7970636.77 |
| birds | Cisticola_nigriloris | Cisticola_nigriloris | 0 | 19.6 | 2.4 | 71914.39 |
| birds | Cisticola_njombe | Cisticola_njombe | 0 | 10.9 | 2.4 | 33269.22 |
| birds | Cisticola_pipiens | Cisticola_pipiens | 0 | 13.5 | 3.5 | 1127999.59 |
| birds | Cisticola_robustus | Cisticola_robustus | 0 | 18.5 | 2.8 | 1229121.48 |

| | | | | | | |
|---|---|---|---|---|---|---|
| birds | Cisticola_ruficeps | Cisticola_ruficeps | 0 | 9.7 | 3.5 | 1253642.08 |
| birds | Cisticola_rufilatus | Cisticola_rufilatus | 0 | 11.3 | 3 | 2748861.04 |
| birds | Cisticola_subruficapilla | Cisticola_subruficapilla | 0 | 9.9 | 2.8 | 910756.54 |
| birds | Cisticola_textrix | Cisticola_textrix | 0 | 10.3 | 2.8 | 702791.56 |
| birds | Cisticola_tinniens | Cisticola_tinniens | 0 | 11.4 | 3 | 1754614.86 |
| birds | Cisticola_woosnami | Cisticola_woosnami | 0 | 13.5 | 2.4 | 1290884.31 |
| birds | Cistothorus_palustris | Cistothorus_palustris | 0 | 10.8 | 4.9 | 10354699.52 |
| birds | Cistothorus_platensis | Cistothorus_platensis | 0 | 9 | 7 | 4772915.83 |
| birds | Cladorhynchus_leucocephalus | Cladorhynchus_leucocephalus | 0 | 216 | 3.5 | 1291070.25 |
| birds | Clanga_clanga | Aquila_clanga | 1 | 2154.3 | 2 | 30259017.15 |
| birds | Clanga_pomarina | Aquila_pomarina | 0 | 1359.4 | 2 | 12328656.87 |
| birds | Clangula_hyemalis | Clangula_hyemalis | 1 | 871 | 7.3 | 14175641.32 |
| birds | Claravis_pretiosa | Claravis_pretiosa | 0 | 68.2 | 2 | 12977823.95 |
| birds | Climacteris_affinis | Climacteris_affinis | 0 | 21.7 | 1.7 | 2630036.35 |
| birds | Climacteris_erythrops | Climacteris_erythrops | 0 | 23.3 | 2 | 317210.54 |
| birds | Climacteris_melanurus | Climacteris_melanurus | 0 | 31.9 | 2 | 1122025.4 |
| birds | Climacteris_picumnus | Climacteris_picumnus | 0 | 28.9 | 3 | 2269397.3 |
| birds | Climacteris_rufus | Climacteris_rufus | 0 | 32.6 | 2 | 452307.17 |
| birds | Clytolaema_rubricauda | Clytolaema_rubricauda | 0 | 7.9 | 2 | 352748.43 |
| birds | Clytorhynchus_vitiensis | Clytorhynchus_vitiensis | 0 | 29.2 | 2 | 19242.45 |
| birds | Clytospiza_monteiri | Clytospiza_monteiri | 0 | 14.9 | 4.9 | 1193530.92 |
| birds | Cnemotriccus_fuscatus | Cnemotriccus_fuscatus | 0 | 13.6 | 3 | 12243851.43 |
| birds | Coccopygia_melanotis | Estrilda_melanotis | 0 | 6.5 | 5.2 | 285539.1 |
| birds | Coccothraustes_coccothraustes | Coccothraustes_coccothraustes | 0 | 56.6 | 4.5 | 15375684.19 |
| birds | Coccycua_minuta | Coccycua_minuta | 0 | 39.3 | 2 | 8754206.58 |
| birds | Coccyzus_longirostris | Coccyzus_longirostris | 0 | 100 | 2.4 | 75087.7 |
| birds | Coccyzus_merlini | Coccyzus_merlini | 0 | 169.6 | 2.4 | 109791.83 |
| birds | Coccyzus_pluvialis | Coccyzus_pluvialis | 0 | 163 | 2.8 | 11021.79 |
| birds | Coccyzus_rufigularis | Coccyzus_rufigularis | 1 | 128 | 2 | 17282.42 |
| birds | Coccyzus_vieilloti | Coccyzus_vieilloti | 0 | 81.9 | 2.4 | 8774.42 |
| birds | Cochlearius_cochlearius | Cochlearius_cochlearius | 0 | 643.2 | 2.8 | 11698377.4 |
| birds | Cochoa_azurea | Cochoa_azurea | 1 | 111.5 | 2 | 4003.48 |
| birds | Cochoa_purpurea | Cochoa_purpurea | 0 | 103 | 2.8 | 623216.09 |
| birds | Cochoa_viridis | Cochoa_viridis | 0 | 120 | 2.8 | 1337235.03 |

| | | | | | | |
|---|---|---|---|---|---|---|
| birds | Coeligena_coeligena | Coeligena_coeligena | 0 | 6.8 | 2 | 387643.17 |
| birds | Coeligena_lutetiae | Coeligena_lutetiae | 0 | 7.3 | 2 | 63339.37 |
| birds | Coeligena_torquata | Coeligena_torquata | 0 | 6.7 | 2 | 302874.56 |
| birds | Coeligena_wilsoni | Coeligena_wilsoni | 0 | 6.7 | 2 | 34158.96 |
| birds | Colaptes_auratus | Colaptes_auratus | 0 | 131.5 | 6 | 11444924.48 |
| birds | Colaptes_campestris | Colaptes_campestris | 0 | 158 | 4.5 | 4575239.79 |
| birds | Colaptes_chrysoides | Colaptes_chrysoides | 0 | 111 | 3.9 | 377623.21 |
| birds | Colaptes_fernandinae | Colaptes_fernandinae | 1 | 135 | 3.9 | 106634.82 |
| birds | Colaptes_melanochloros | Colaptes_melanochloros | 0 | 127.3 | 4 | 4723714.99 |
| birds | Colaptes_pitius | Colaptes_pitius | 0 | 159 | 4.9 | 428459.35 |
| birds | Colaptes_rubiginosus | Colaptes_rubiginosus | 0 | 55.8 | 2.8 | 2243128.49 |
| birds | Colaptes_rupicola | Colaptes_rupicola | 0 | 174 | 2.8 | 785415.83 |
| birds | Colibri_serrirostris | Colibri_serrirostris | 0 | 6.7 | 2 | 2370635.54 |
| birds | Colibri_thalassinus | Colibri_thalassinus | 0 | 5.9 | 2 | 792390.56 |
| birds | Colinus_cristatus | Colinus_cristatus | 0 | 134.5 | 11.3 | 1970617.04 |
| birds | Colinus_virginianus | Colinus_virginianus | 0 | 171.9 | 8 | 4364464.18 |
| birds | Colius_castanotus | Colius_castanotus | 0 | 56.6 | 3 | 289063.97 |
| birds | Colius_colius | Colius_colius | 0 | 41.4 | 3 | 1611547.39 |
| birds | Colius_leucocephalus | Colius_leucocephalus | 0 | 35.8 | 3 | 533022.16 |
| birds | Colius_striatus | Colius_striatus | 0 | 51.1 | 2.4 | 8057145.03 |
| birds | Collocalia_esculenta | Collocalia_esculenta | 0 | 6.3 | 1.4 | 2759413.59 |
| birds | Colluricincla_boweri | Colluricincla_boweri | 1 | 43.8 | 2 | 7631.07 |
| birds | Colluricincla_harmonica | Colluricincla_harmonica | 0 | 65.8 | 2.8 | 6919512.58 |
| birds | Colluricincla_megarhyncha | Colluricincla_megarhyncha | 0 | 32.3 | 2.4 | 1204726.42 |
| birds | Colluricincla_woodwardi | Colluricincla_woodwardi | 0 | 54.5 | 2.4 | 549567.85 |
| birds | Colonia_colonus | Colonia_colonus | 0 | 18.3 | 2.4 | 6767109.68 |
| birds | Colorhamphus_parvirostris | Colorhamphus_parvirostris | 0 | 10.6 | 3 | 536388.92 |
| birds | Columba_albinucha | Columba_albinucha | 0 | 325 | 1 | 142261.79 |
| birds | Columba_albitorques | Columba_albitorques | 0 | 277 | 2 | 137598.58 |
| birds | Columba_arquatrix | Columba_arquatrix | 0 | 400 | 1 | 1486805.14 |
| birds | Columba_bollii | Columba_bollii | 0 | 308.5 | 1 | 2531.08 |
| birds | Columba_delegorguei | Columba_delegorguei | 0 | 133 | 2 | 232244.08 |
| birds | Columba_elphinstonii | Columba_elphinstonii | 1 | 379 | 1 | 116434.82 |
| birds | Columba_eversmanni | Columba_eversmanni | 1 | 202 | 2 | 5250966.24 |

| | | | | | | |
|---|---|---|---|---|---|---|
| birds | Columba_guinea | Columba_guinea | 0 | 352 | 2 | 10557795.16 |
| birds | Columba_hodgsonii | Columba_hodgsonii | 0 | 308.5 | 1 | 2007983.93 |
| birds | Columba_iriditorques | Columba_iriditorques | 0 | 125.9 | 1 | 2757795.12 |
| birds | Columba_janthina | Columba_janthina | 0 | 397 | 1 | 2.4 |
| birds | Columba_leucomela | Columba_leucomela | 0 | 420 | 1 | 275007.14 |
| birds | Columba_leuconota | Columba_leuconota | 0 | 284 | 2 | 2656832.05 |
| birds | Columba_livia | Columba_livia | 0 | 354.2 | 2 | 27446747.7 |
| birds | Columba_oenas | Columba_oenas | 0 | 290.8 | 2 | 9639015.46 |
| birds | Columba_palumbus | Columba_palumbus | 0 | 490 | 2 | 12401484.34 |
| birds | Columba_pulchricollis | Columba_pulchricollis | 0 | 330 | 1.4 | 565151.19 |
| birds | Columba_punicea | Columba_punicea | 1 | 434.4 | 1 | 358510.17 |
| birds | Columba_rupestris | Columba_rupestris | 0 | 245.4 | 2 | 11908353.58 |
| birds | Columba_sjostedti | Columba_sjostedti | 0 | 308.5 | 1 | 44188.92 |
| birds | Columba_torringtoniae | Columba_torringtoniae | 1 | 308.5 | 1 | 5001.87 |
| birds | Columba_unicincta | Columba_unicincta | 0 | 389.1 | 1 | 2699737.73 |
| birds | Columba_vitiensis | Columba_vitiensis | 0 | 353 | 1 | 1128735.55 |
| birds | Columbina_cruziana | Columbina_cruziana | 0 | 50.1 | 2 | 334307.94 |
| birds | Columbina_inca | Columbina_inca | 0 | 47.5 | 2 | 2760667.85 |
| birds | Columbina_minuta | Columbina_minuta | 0 | 32.6 | 2 | 7050160.83 |
| birds | Columbina_passerina | Columbina_passerina | 0 | 35.4 | 2 | 7245312.58 |
| birds | Columbina_picui | Columbina_picui | 0 | 47 | 2 | 6427122.47 |
| birds | Columbina_squammata | Columbina_squammata | 0 | 52.9 | 2 | 4814670.86 |
| birds | Columbina_talpacoti | Columbina_talpacoti | 0 | 46 | 2 | 14617238.71 |
| birds | Conopias_albovittatus | Conopias_albovittatus | 0 | 24.4 | 2 | 198320.46 |
| birds | Conopias_parvus | Conopias_parvus | 0 | 21 | 2 | 3906792.83 |
| birds | Conopophaga_aurita | Conopophaga_aurita | 0 | 26.3 | 2 | 2800393.44 |
| birds | Conopophaga_lineata | Conopophaga_lineata | 0 | 25.2 | 2 | 1896972.3 |
| birds | Conopophaga_melanops | Conopophaga_melanops | 0 | 20.1 | 2 | 242500.41 |
| birds | Conopophila_albogularis | Conopophila_albogularis | 0 | 11.8 | 2 | 496534.41 |
| birds | Conopophila_rufogularis | Conopophila_rufogularis | 0 | 10.8 | 2.4 | 1954135.96 |
| birds | Conopophila_whitei | Conopophila_whitei | 0 | 9.8 | 2 | 1437139.32 |
| birds | Contopus_caribaeus | Contopus_caribaeus | 0 | 10.5 | 2.8 | 119645.89 |
| birds | Contopus_cinereus | Contopus_cinereus | 0 | 11.6 | 2.4 | 3837797.83 |
| birds | Contopus_cooperi | Contopus_cooperi | 0 | 32.1 | 3.9 | 20278748.22 |

| | | | | | | |
|---|---|---|---|---|---|---|
| birds | Contopus_hispaniolensis | Contopus_hispaniolensis | 0 | 11.7 | 2.8 | 74799.65 |
| birds | Contopus_pertinax | Contopus_pertinax | 0 | 27.2 | 2.8 | 855800.29 |
| birds | Contopus_sordidulus | Contopus_sordidulus | 0 | 13.1 | 2.8 | 11524307.35 |
| birds | Contopus_virens | Contopus_virens | 0 | 13.9 | 2.8 | 10575580.52 |
| birds | Copsychus_albospecularis | Copsychus_albospecularis | 0 | 22.7 | 3 | 239689.82 |
| birds | Copsychus_malabaricus | Copsychus_malabaricus | 0 | 29.2 | 3.9 | 4343996.14 |
| birds | Copsychus_saularis | Copsychus_saularis | 0 | 36 | 3.2 | 9009660.2 |
| birds | Coracias_abyssinicus | Coracias_abyssinicus | 0 | 102 | 4.2 | 7043827.45 |
| birds | Coracias_benghalensis | Coracias_benghalensis | 0 | 157.5 | 3.9 | 4430960.42 |
| birds | Coracias_caudatus | Coracias_caudatus | 0 | 110 | 2.8 | 8821260.61 |
| birds | Coracias_garrulus | Coracias_garrulus | 0 | 146 | 4.2 | 34073962.64 |
| birds | Coracias_naevius | Coracias_naevia | 0 | 168 | 2.8 | 9355490.24 |
| birds | Coracias_spatulatus | Coracias_spatulatus | 0 | 90.4 | 2.8 | 2917947.7 |
| birds | Coracina_boyeri | Coracina_boyeri | 0 | 66.4 | 1 | 604477.42 |
| birds | Coracina_caeruleogrisea | Coracina_caeruleogrisea | 0 | 138.2 | 1 | 404850.4 |
| birds | Coracina_javensis | Coracina_javensis | 0 | 94.5 | 2 | 3632403.85 |
| birds | Coracina_lineata | Coracina_lineata | 0 | 76 | 2 | 643371.3 |
| birds | Coracina_longicauda | Coracina_longicauda | 0 | 101 | 1 | 135269.83 |
| birds | Coracina_macei | Coracina_macei | 0 | 105 | 2.4 | 2249595.09 |
| birds | Coracina_maxima | Coracina_maxima | 0 | 127 | 2.4 | 6390802.08 |
| birds | Coracina_novaehollandiae | Coracina_novaehollandiae | 0 | 118 | 2.4 | 8239647.62 |
| birds | Coracina_papuensis | Coracina_papuensis | 0 | 63.5 | 2.4 | 4454361.88 |
| birds | Coracopsis_nigra | Coracopsis_nigra | 0 | 186.2 | 2.4 | 591110.01 |
| birds | Coracopsis_vasa | Coracopsis_vasa | 0 | 525 | 3 | 413210.96 |
| birds | Coragyps_atratus | Coragyps_atratus | 0 | 1881.7 | 2 | 21175692.94 |
| birds | Corapipo_leucorrhoa | Corapipo_leucorrhoa | 0 | 11.7 | 2 | 267106.87 |
| birds | Corcorax_melanorhamphos | Corcorax_melanorhamphos | 0 | 364 | 3.9 | 1979313.45 |
| birds | Cormobates_leucophaea | Cormobates_leucophaea | 0 | 22 | 2.4 | 969111.73 |
| birds | Corthylio_calendula | Regulus_calendula | 0 | 6.2 | 7.9 | 16933121.98 |
| birds | Corvus_albicollis | Corvus_albicollis | 0 | 894 | 2.4 | 1615598.28 |
| birds | Corvus_albus | Corvus_albus | 0 | 524 | 2.6 | 19544750.22 |
| birds | Corvus_bennetti | Corvus_bennetti | 0 | 389 | 4.9 | 5681813.47 |
| birds | Corvus_brachyrhynchos | Corvus_brachyrhynchos | 0 | 448.8 | 4.5 | 12113678.31 |
| birds | Corvus_capensis | Corvus_capensis | 0 | 553 | 4.2 | 4036026.48 |

| | | | | | | |
|---|---|---|---|---|---|---|
| birds | Corvus_caurinus | Corvus_caurinus | 0 | 390.8 | 4.5 | 396973.08 |
| birds | Corvus_corax | Corvus_corax | 0 | 928 | 4.6 | 52342379.53 |
| birds | Corvus_corone | Corvus_corone | 0 | 570 | 4.5 | 34119496.9 |
| birds | Corvus_coronoides | Corvus_coronoides | 0 | 644.3 | 3.9 | 4803477.78 |
| birds | Corvus_crassirostris | Corvus_crassirostris | 0 | 1135 | 4 | 759238.98 |
| birds | Corvus_cryptoleucus | Corvus_cryptoleucus | 0 | 533.5 | 5.9 | 1508239.5 |
| birds | Corvus_dauuricus | Corvus_dauuricus | 0 | 208 | 4.9 | 7425668.89 |
| birds | Corvus_enca | Corvus_enca | 0 | 251 | 2 | 1629657.06 |
| birds | Corvus_frugilegus | Corvus_frugilegus | 0 | 452.1 | 3.7 | 23167955.27 |
| birds | Corvus_imparatus | Corvus_imparatus | 0 | 221 | 4 | 114579.13 |
| birds | Corvus_jamaicensis | Corvus_jamaicensis | 0 | 338 | 3 | 11021.79 |
| birds | Corvus_macrorhynchos | Corvus_macrorhynchos | 0 | 513.1 | 4.9 | 13313192.58 |
| birds | Corvus_mellori | Corvus_mellori | 0 | 540.4 | 3.9 | 1353458.96 |
| birds | Corvus_monedula | Corvus_monedula | 0 | 246 | 4 | 15225329.42 |
| birds | Corvus_moneduloides | Corvus_moneduloides | 0 | 277.4 | 2.4 | 17048.53 |
| birds | Corvus_nasicus | Corvus_nasicus | 0 | 401 | 4 | 28429.7 |
| birds | Corvus_orru | Corvus_orru | 0 | 508 | 3.2 | 5394057.72 |
| birds | Corvus_ossifragus | Corvus_ossifragus | 0 | 284.6 | 4.5 | 970986.53 |
| birds | Corvus_palmarum | Corvus_palmarum | 0 | 289 | 4 | 70693.53 |
| birds | Corvus_pectoralis | Corvus_torquatus | 1 | 470.5 | 3.5 | 2074732.63 |
| birds | Corvus_rhipidurus | Corvus_rhipidurus | 0 | 611 | 3.5 | 2756382.95 |
| birds | Corvus_ruficollis | Corvus_ruficollis | 0 | 573.5 | 3.7 | 12915376.88 |
| birds | Corvus_sinaloae | Corvus_sinaloae | 0 | 244 | 4.5 | 95101.34 |
| birds | Corvus_splendens | Corvus_splendens | 0 | 292.6 | 4.5 | 4860994.18 |
| birds | Corvus_tasmanicus | Corvus_tasmanicus | 0 | 676.3 | 4.2 | 374479.03 |
| birds | Corvus_tristis | Corvus_tristis | 0 | 635 | 2 | 693202.25 |
| birds | Corydon_sumatranus | Corydon_sumatranus | 0 | 140 | 2.8 | 2209826.9 |
| birds | Coryphistera_alaudina | Coryphistera_alaudina | 0 | 30 | 3.9 | 1447474.66 |
| birds | Corythaeola_cristata | Corythaeola_cristata | 0 | 965 | 2 | 3896093.97 |
| birds | Corythaixoides_concolor | Corythaixoides_concolor | 0 | 258 | 3 | 3808237.76 |
| birds | Corythaixoides_personatus | Corythaixoides_personatus | 0 | 250 | 2.4 | 52976.1 |
| birds | Corythopis_delalandi | Corythopis_delalandi | 0 | 15 | 2.4 | 2526735.17 |
| birds | Corythornis_cristatus | Alcedo_cristata | 0 | 15.7 | 4.2 | 14363969.72 |
| birds | Corythornis_leucogaster | Alcedo_leucogaster | 0 | 14.5 | 2 | 2739779.01 |

| | | | | | | |
|---|---|---|---|---|---|---|
| birds | Corythornis_madagascariensis | Ceyx_madagascariensis | 0 | 17.7 | 4 | 158670.3 |
| birds | Corythornis_vintsioides | Alcedo_vintsioides | 0 | 17.9 | 4.2 | 572332.97 |
| birds | Coscoroba_coscoroba | Coscoroba_coscoroba | 0 | 4294.8 | 5.3 | 4096479.61 |
| birds | Cossypha_albicapillus | Cossypha_albicapilla | 0 | 58.5 | 2 | 1269495.3 |
| birds | Cossypha_cyanocampter | Cossypha_cyanocampter | 0 | 26.7 | 2 | 1817635.07 |
| birds | Cossypha_dichroa | Cossypha_dichroa | 0 | 45.8 | 3 | 253533.92 |
| birds | Cossypha_heuglini | Cossypha_heuglini | 0 | 34.8 | 2 | 6366999.37 |
| birds | Cossypha_natalensis | Cossypha_natalensis | 0 | 34.1 | 3 | 3643626.53 |
| birds | Cossypha_niveicapilla | Cossypha_niveicapilla | 0 | 36.7 | 2.4 | 5176063.97 |
| birds | Cossypha_semirufa | Cossypha_semirufa | 0 | 27.3 | 2.4 | 865714.31 |
| birds | Cossyphicula_isabellae | Cossypha_isabellae | 0 | 24 | 2 | 51494.15 |
| birds | Cotinga_ridgwayi | Cotinga_ridgwayi | 1 | 57.2 | 2 | 14415.57 |
| birds | Coturnicops_noveboracensis | Coturnicops_noveboracensis | 0 | 60.9 | 8 | 6754791.17 |
| birds | Coturnix_coromandelica | Coturnix_coromandelica | 0 | 73.8 | 4.9 | 2899254.36 |
| birds | Coturnix_coturnix | Coturnix_coturnix | 0 | 96.3 | 10.2 | 34416948.31 |
| birds | Coturnix_delegorguei | Coturnix_delegorguei | 0 | 76 | 5.7 | 9153237.05 |
| birds | Coturnix_japonica | Coturnix_japonica | 0 | 94.8 | 9.5 | 9041141.35 |
| birds | Coturnix_pectoralis | Coturnix_pectoralis | 0 | 101.9 | 7.5 | 6323957.08 |
| birds | Coua_caerulea | Coua_caerulea | 0 | 244 | 1 | 333697.61 |
| birds | Coua_coquereli | Coua_coquereli | 0 | 160 | 2 | 158107.38 |
| birds | Coua_cristata | Coua_cristata | 0 | 147 | 2 | 271871.29 |
| birds | Coua_cursor | Coua_cursor | 0 | 110 | 2 | 51235.25 |
| birds | Coua_gigas | Coua_gigas | 0 | 424 | 3 | 149356.19 |
| birds | Coua_reynaudii | Coua_reynaudii | 0 | 153 | 2 | 320733.34 |
| birds | Coua_ruficeps | Coua_ruficeps | 0 | 180 | 2 | 107746.82 |
| birds | Coua_serriana | Coua_serriana | 0 | 293 | 2 | 62730.5 |
| birds | Cracticus_cassicus | Cracticus_cassicus | 0 | 142 | 2.4 | 554775.59 |
| birds | Cracticus_mentalis | Cracticus_mentalis | 0 | 77 | 2.4 | 218872.78 |
| birds | Cracticus_nigrogularis | Cracticus_nigrogularis | 0 | 128 | 3.9 | 6347951.06 |
| birds | Cracticus_torquatus | Cracticus_torquatus | 0 | 82.9 | 3.9 | 6032491.25 |
| birds | Cranioleuca_pyrrhophia | Cranioleuca_pyrrhophia | 0 | 14.9 | 2.4 | 2026269.35 |
| birds | Cranioleuca_vulpina | Cranioleuca_vulpina | 0 | 15.7 | 2.4 | 5731362.69 |
| birds | Crax_alector | Crax_alector | 0 | 2872 | 2 | 2099461.66 |
| birds | Crax_blumenbachii | Crax_blumenbachii | 1 | 3500 | 2 | 148947.32 |

| | | | | | | |
|---|---|---|---|---|---|---|
| birds | Crax_daubentoni | Crax_daubentoni | 0 | 2499.9 | 2 | 342112.86 |
| birds | Crax_fasciolata | Crax_fasciolata | 1 | 2600 | 2 | 3381019.12 |
| birds | Crax_globulosa | Crax_globulosa | 1 | 2500 | 2 | 44465.66 |
| birds | Crax_rubra | Crax_rubra | 1 | 4133 | 2 | 662063.72 |
| birds | Creatophora_cinerea | Creatophora_cinerea | 0 | 73 | 3.2 | 5407365.7 |
| birds | Crex_crex | Crex_crex | 0 | 154.9 | 9.8 | 22051816.35 |
| birds | Crex_egregia | Crecopsis_egregia | 0 | 119 | 5.2 | 11681471.46 |
| birds | Crinifer_piscator | Crinifer_piscator | 0 | 370 | 2.4 | 3640485.51 |
| birds | Crinifer_zonurus | Crinifer_zonurus | 0 | 527 | 2.4 | 1014236.61 |
| birds | Criniferoides_leucogaster | Corythaixoides_leucogaster | 0 | 216.2 | 2.4 | 2293621.91 |
| birds | Criniger_barbatus | Criniger_barbatus | 0 | 42.6 | 2 | 646336.43 |
| birds | Criniger_calurus | Criniger_calurus | 0 | 37.7 | 2 | 3299941.84 |
| birds | Criniger_chloronotus | Criniger_chloronotus | 0 | 45 | 2 | 1204957.83 |
| birds | Crithagra_albogularis | Serinus_albogularis | 0 | 27.1 | 2.8 | 916328.36 |
| birds | Crithagra_atrogularis | Serinus_atrogularis | 0 | 11.4 | 2.8 | 3009542.68 |
| birds | Crithagra_citrinelloides | Serinus_citrinelloides | 0 | 13 | 2.4 | 555034.99 |
| birds | Crithagra_citrinipectus | Serinus_citrinipectus | 0 | 11.2 | 3.5 | 583770.24 |
| birds | Crithagra_donaldsoni | Serinus_donaldsoni | 0 | 24 | 3 | 604511.84 |
| birds | Crithagra_dorsostriata | Serinus_dorsostriatus | 0 | 15.1 | 2.8 | 864663.9 |
| birds | Crithagra_flaviventris | Serinus_flaviventris | 0 | 16.2 | 2.8 | 1693983.55 |
| birds | Crithagra_gularis | Serinus_gularis | 0 | 16 | 2.8 | 1507366.09 |
| birds | Crithagra_leucoptera | Serinus_leucopterus | 0 | 22.1 | 2.8 | 82476.24 |
| birds | Crithagra_leucopygia | Serinus_leucopygius | 0 | 10.8 | 2.8 | 3039637.6 |
| birds | Crithagra_mennelli | Serinus_mennelli | 0 | 15.4 | 2.8 | 1495142.19 |
| birds | Crithagra_mozambica | Serinus_mozambicus | 0 | 11.9 | 3.2 | 9549283.64 |
| birds | Crithagra_scotops | Serinus_scotops | 0 | 14.2 | 2.8 | 184665.55 |
| birds | Crithagra_striolata | Serinus_striolatus | 0 | 22.4 | 3.5 | 867600.67 |
| birds | Crithagra_sulphurata | Serinus_sulphuratus | 0 | 27.9 | 2.8 | 1869165.54 |
| birds | Crithagra_totta | Serinus_totta | 0 | 13.4 | 3.9 | 85526.4 |
| birds | Crossleyia_xanthophrys | Crossleyia_xanthophrys | 0 | 19.9 | 3 | 22717.1 |
| birds | Crossoptilon_auritum | Crossoptilon_auritum | 0 | 1768.3 | 8 | 512906.05 |
| birds | Crossoptilon_crossoptilon | Crossoptilon_crossoptilon | 0 | 2075.3 | 6 | 171921.78 |
| birds | Crossoptilon_mantchuricum | Crossoptilon_mantchuricum | 1 | 1860.9 | 7.5 | 188621.54 |
| birds | Crotophaga_ani | Crotophaga_ani | 0 | 110.1 | 2.9 | 14119059.35 |

| | | | | | | |
|---|---|---|---|---|---|---|
| birds | Crotophaga_major | Crotophaga_major | 0 | 148.3 | 2.4 | 12117795.17 |
| birds | Crotophaga_sulcirostris | Crotophaga_sulcirostris | 0 | 82 | 3.5 | 2717920.78 |
| birds | Cryptillas_victorini | Bradypterus_victorini | 0 | 10 | 2.4 | 78210.99 |
| birds | Cryptolybia_olivacea | Stactolaema_olivacea | 0 | 49.4 | 4.2 | 155265.95 |
| birds | Cryptospiza_reichenovii | Cryptospiza_reichenovii | 0 | 12.4 | 3.5 | 435889.11 |
| birds | Cryptospiza_salvadorii | Cryptospiza_salvadorii | 0 | 12.2 | 3.9 | 258138.36 |
| birds | Crypturellus_boucardi | Crypturellus_boucardi | 1 | 435.6 | 3.7 | 334426.96 |
| birds | Crypturellus_cinereus | Crypturellus_cinereus | 0 | 506.6 | 2 | 5924779.18 |
| birds | Crypturellus_cinnamomeus | Crypturellus_cinnamomeus | 0 | 422 | 3 | 695817.1 |
| birds | Crypturellus_noctivagus | Crypturellus_noctivagus | 0 | 562 | 4 | 1469564.58 |
| birds | Crypturellus_obsoletus | Crypturellus_obsoletus | 0 | 443.4 | 4.5 | 2652131.25 |
| birds | Crypturellus_parvirostris | Crypturellus_parvirostris | 0 | 199 | 4.5 | 6659535.62 |
| birds | Crypturellus_soui | Crypturellus_soui | 0 | 216.2 | 2 | 11151222.76 |
| birds | Crypturellus_tataupa | Crypturellus_tataupa | 0 | 218.8 | 4.5 | 4855861.19 |
| birds | Crypturellus_undulatus | Crypturellus_undulatus | 0 | 564.4 | 4.5 | 8533511.59 |
| birds | Crypturellus_variegatus | Crypturellus_variegatus | 0 | 378 | 1 | 5436849.92 |
| birds | Culicicapa_ceylonensis | Culicicapa_ceylonensis | 0 | 7.7 | 3.5 | 6901649.27 |
| birds | Curruca_boehmi | Sylvia_boehmi | 0 | 14 | 2.4 | 757783.64 |
| birds | Curruca_buryi | Sylvia_buryi | 0 | 22 | 3 | 62421.27 |
| birds | Curruca_communis | Sylvia_communis | 0 | 15.1 | 4.5 | 42176814.38 |
| birds | Curruca_conspicillata | Sylvia_conspicillata | 0 | 9 | 5 | 4545002.93 |
| birds | Curruca_deserticola | Sylvia_deserticola | 0 | 8.6 | 3.9 | 3004118.25 |
| birds | Curruca_layardi | Sylvia_layardi | 0 | 14.3 | 3 | 780211.28 |
| birds | Curruca_leucomelaena | Sylvia_leucomelaena | 0 | 13.9 | 2.8 | 547018.24 |
| birds | Curruca_lugens | Sylvia_lugens | 0 | 14.9 | 2 | 401532.97 |
| birds | Curruca_melanocephala | Sylvia_melanocephala | 0 | 11.7 | 4.5 | 4224670.01 |
| birds | Curruca_melanothorax | Sylvia_melanothorax | 0 | 11.4 | 4.5 | 149419.08 |
| birds | Curruca_mystacea | Sylvia_mystacea | 0 | 9.9 | 4.5 | 7953271.3 |
| birds | Curruca_nisoria | Sylvia_nisoria | 0 | 22.5 | 4.2 | 12451191.9 |
| birds | Curruca_ruppeli | Sylvia_rueppelli | 0 | 12.8 | 4.5 | 5404863.93 |
| birds | Curruca_subcoerulea | Sylvia_subcaerulea | 0 | 14.6 | 2.8 | 2960791.88 |
| birds | Curruca_undata | Sylvia_undata | 0 | 10.8 | 4 | 1420763.35 |
| birds | Cursorius_cursor | Cursorius_cursor | 0 | 138 | 2 | 10509601.04 |
| birds | Cursorius_rufus | Cursorius_rufus | 0 | 75 | 2 | 1392070.57 |

| | | | | | | |
|---|---|---|---|---|---|---|
| birds | Cyanistes_caeruleus | Parus_caeruleus | 0 | 11.1 | 5.1 | 8770881.49 |
| birds | Cyanistes_cyanus | Parus_cyanus | 0 | 14.6 | 8.8 | 8153289.29 |
| birds | Cyanistes_teneriffae | Parus_teneriffae | 0 | 10 | 5.1 | 624090.33 |
| birds | Cyanocitta_cristata | Cyanocitta_cristata | 0 | 88 | 4.5 | 6809850.43 |
| birds | Cyanocitta_stelleri | Cyanocitta_stelleri | 0 | 128 | 4 | 3560822.24 |
| birds | Cyanoliseus_patagonus | Cyanoliseus_patagonus | 0 | 277.9 | 2.8 | 1217792.54 |
| birds | Cyanomitra_alinae | Nectarinia_alinae | 0 | 12.6 | 1.4 | 114321.46 |
| birds | Cyanomitra_bannermani | Nectarinia_bannermani | 0 | 14 | 2 | 454416.96 |
| birds | Cyanomitra_cyanolaema | Nectarinia_cyanolaema | 0 | 16.8 | 2 | 3142385.64 |
| birds | Cyanomitra_olivacea | Nectarinia_olivacea | 0 | 9.7 | 1.4 | 5394958.82 |
| birds | Cyanomitra_oritis | Nectarinia_oritis | 0 | 11.2 | 2 | 65561.74 |
| birds | Cyanomitra_verreauxii | Nectarinia_veroxii | 0 | 11.5 | 1.4 | 155165.64 |
| birds | Cyanomitra_verticalis | Nectarinia_verticalis | 0 | 14.4 | 2 | 5373209.67 |
| birds | Cyanopica_cyanus | Cyanopica_cyanus | 0 | 95.9 | 5.9 | 4748269.37 |
| birds | Cyclopsitta_diophthalma | Cyclopsitta_diophthalma | 0 | 41.8 | 2 | 448013.86 |
| birds | Cygnus_atratus | Cygnus_atratus | 0 | 5623.2 | 5.5 | 7027995.14 |
| birds | Cygnus_buccinator | Cygnus_buccinator | 0 | 11071.1 | 5.7 | 1616822.38 |
| birds | Cygnus_columbianus | Cygnus_columbianus | 0 | 6378 | 3.9 | 13948485.91 |
| birds | Cygnus_cygnus | Cygnus_cygnus | 0 | 9350 | 4.5 | 19300590.68 |
| birds | Cygnus_melancoryphus | Cygnus_melancoryphus | 0 | 4711.7 | 5.7 | 3619491.76 |
| birds | Cygnus_olor | Cygnus_olor | 0 | 10682 | 5.9 | 6112278.31 |
| birds | Cymbilaimus_lineatus | Cymbilaimus_lineatus | 0 | 35.8 | 2 | 6643566.16 |
| birds | Cymbirhynchus_macrorhynchos | Cymbirhynchus_macrorhynchos | 0 | 59.4 | 2.8 | 1624283.49 |
| birds | Cynanthus_latirostris | Cynanthus_latirostris | 0 | 3.1 | 2 | 990017.28 |
| birds | Cyornis_brunneatus | Rhinomyias_brunneatus | 1 | 18.1 | 2.4 | 677461.88 |
| birds | Cyornis_colonus | Rhinomyias_colonus | 0 | 23.1 | 2.4 | 3612.36 |
| birds | Cyornis_concretus | Cyornis_concretus | 0 | 24.5 | 2 | 866448.98 |
| birds | Cyornis_olivaceus | Rhinomyias_olivaceus | 0 | 15.7 | 2.4 | 595916.66 |
| birds | Cyornis_pallidipes | Cyornis_pallipes | 0 | 19 | 4 | 52751.57 |
| birds | Cyornis_rubeculoides | Cyornis_rubeculoides | 0 | 14.2 | 4.5 | 2120972.34 |
| birds | Cyornis_ruficauda | Rhinomyias_ruficauda | 0 | 17.3 | 2.4 | 81701.8 |
| birds | Cyornis_rufigastra | Cyornis_rufigastra | 0 | 18.3 | 2 | 429957.49 |
| birds | Cyornis_tickelliae | Cyornis_tickelliae | 0 | 16.2 | 3.5 | 2366894.59 |
| birds | Cyornis_umbratilis | Rhinomyias_umbratilis | 0 | 18.2 | 2.4 | 1066649.1 |

| | | | | | | |
|---|---|---|---|---|---|---|
| birds | Cyornis_unicolor | Cyornis_unicolor | 0 | 21 | 4 | 2146298.64 |
| birds | Cyphorhinus_arada | Cyphorhinus_arada | 0 | 20.9 | 2 | 4458849.32 |
| birds | Cyphorhinus_phaeocephalus | Cyphorhinus_phaeocephalus | 0 | 24.6 | 2.4 | 318922.64 |
| birds | Cyphorhinus_thoracicus | Cyphorhinus_thoracicus | 0 | 31.3 | 2 | 58720.02 |
| birds | Cypseloides_cherriei | Cypseloides_cherriei | 0 | 22.1 | 1 | 277365.45 |
| birds | Cypseloides_cryptus | Cypseloides_cryptus | 0 | 34.5 | 1 | 413124 |
| birds | Cypseloides_fumigatus | Cypseloides_fumigatus | 0 | 42 | 1 | 2325060.02 |
| birds | Cypseloides_niger | Cypseloides_niger | 1 | 39.7 | 1 | 2153809.27 |
| birds | Cypsiurus_balasiensis | Cypsiurus_balasiensis | 0 | 9.1 | 1.7 | 6867419.3 |
| birds | Cypsiurus_parvus | Cypsiurus_parvus | 0 | 13.6 | 2.4 | 15963964.24 |
| birds | Cyrtonyx_montezumae | Cyrtonyx_montezumae | 0 | 185.4 | 11 | 754065.67 |
| birds | Dacelo_gaudichaud | Dacelo_gaudichaud | 0 | 143 | 2 | 671238.41 |
| birds | Dacelo_leachii | Dacelo_leachii | 0 | 308 | 2.4 | 2183194.4 |
| birds | Dacelo_novaeguineae | Dacelo_novaeguineae | 0 | 333.8 | 2.2 | 2766080 |
| birds | Dasyornis_brachypterus | Dasyornis_brachypterus | 1 | 41.8 | 2 | 51744.96 |
| birds | Dasyornis_broadbenti | Dasyornis_broadbenti | 0 | 67.9 | 2 | 26759.89 |
| birds | Dasyornis_longirostris | Dasyornis_longirostris | 1 | 32.7 | 2 | 52040.52 |
| birds | Deconychura_longicauda | Deconychura_longicauda | 0 | 23.8 | 2 | 1094105.42 |
| birds | Delacourella_capistrata | Nesocharis_capistrata | 0 | 11.6 | 4 | 976089.61 |
| birds | Delichon_dasypus | Delichon_dasypus | 0 | 18 | 3.5 | 11430696.77 |
| birds | Delichon_nipalense | Delichon_nipalense | 0 | 15 | 3.9 | 543149.36 |
| birds | Delichon_urbicum | Delichon_urbicum | 0 | 14.5 | 4.5 | 38136395.97 |
| birds | Dendragapus_obscurus | Dendragapus_obscurus | 0 | 1047.3 | 6.9 | 1526017.86 |
| birds | Dendrexetastes_rufigula | Dendrexetastes_rufigula | 0 | 69.6 | 2.4 | 3670234 |
| birds | Dendrocincla_anabatina | Dendrocincla_anabatina | 0 | 34.2 | 2 | 378885 |
| birds | Dendrocincla_fuliginosa | Dendrocincla_fuliginosa | 0 | 38.7 | 1.7 | 8483632.64 |
| birds | Dendrocincla_homochroa | Dendrocincla_homochroa | 0 | 39.4 | 2.4 | 492694.51 |
| birds | Dendrocincla_turdina | Dendrocincla_turdina | 0 | 39 | 2 | 1193430.6 |
| birds | Dendrocolaptes_certhia | Dendrocolaptes_certhia | 0 | 68.7 | 1 | 6659679.41 |
| birds | Dendrocolaptes_picumnus | Dendrocolaptes_picumnus | 0 | 76.6 | 2 | 5749616.39 |
| birds | Dendrocopos_assimilis | Dendrocopos_assimilis | 0 | 51.8 | 3.5 | 604188.64 |
| birds | Dendrocopos_atratus | Dendrocopos_atratus | 0 | 46.7 | 4.5 | 309610.66 |
| birds | Dendrocopos_darjellensis | Dendrocopos_darjellensis | 0 | 70.7 | 2.8 | 301933.91 |
| birds | Dendrocopos_himalayensis | Dendrocopos_himalayensis | 0 | 67.7 | 3.9 | 115066.11 |

| | | | | | | |
|---|---|---|---|---|---|---|
| birds | Dendrocopos_hyperythrus | Dendrocopos_hyperythrus | 0 | 46.5 | 4.5 | 1790193.93 |
| birds | Dendrocopos_leucopterus | Dendrocopos_leucopterus | 0 | 67 | 4.9 | 2454403.28 |
| birds | Dendrocopos_leucotos | Dendrocopos_leucotos | 0 | 116.8 | 3.9 | 11229537.31 |
| birds | Dendrocopos_macei | Dendrocopos_macei | 0 | 44.9 | 3.2 | 904703.87 |
| birds | Dendrocopos_major | Dendrocopos_major | 0 | 74.9 | 5.9 | 24413724.29 |
| birds | Dendrocopos_syriacus | Dendrocopos_syriacus | 0 | 76.7 | 4.5 | 2931365.51 |
| birds | Dendrocygna_arborea | Dendrocygna_arborea | 0 | 1150 | 8.5 | 115603.83 |
| birds | Dendrocygna_arcuata | Dendrocygna_arcuata | 0 | 796.2 | 9.5 | 4892718.61 |
| birds | Dendrocygna_autumnalis | Dendrocygna_autumnalis | 0 | 755.3 | 13.9 | 9474186.49 |
| birds | Dendrocygna_bicolor | Dendrocygna_bicolor | 0 | 756.4 | 9.8 | 18682192.41 |
| birds | Dendrocygna_eytoni | Dendrocygna_eytoni | 0 | 790 | 10.6 | 4778055.17 |
| birds | Dendrocygna_guttata | Dendrocygna_guttata | 0 | 800 | 11 | 933148.33 |
| birds | Dendrocygna_javanica | Dendrocygna_javanica | 0 | 519.6 | 9.2 | 7165332.29 |
| birds | Dendrocygna_viduata | Dendrocygna_viduata | 0 | 690 | 7.2 | 27416318.34 |
| birds | Dendroma_rufa | Philydor_rufum | 0 | 25 | 2.4 | 2187683.25 |
| birds | Dendronanthus_indicus | Dendronanthus_indicus | 0 | 16.4 | 4.5 | 7547714.46 |
| birds | Dendropicos_fuscescens | Dendropicos_fuscescens | 0 | 26 | 2.4 | 12708768.94 |
| birds | Dendropicos_goertae | Mesopicos_goertae | 0 | 44.5 | 2.8 | 7460292.4 |
| birds | Dendropicos_griseocephalus | Mesopicos_griseocephalus | 0 | 40.9 | 2.4 | 2206119.9 |
| birds | Dendropicos_namaquus | Thripias_namaquus | 0 | 81.5 | 3 | 7980417.7 |
| birds | Dendropicos_obsoletus | Dendrocopos_obsoletus | 0 | 21.4 | 2 | 5967593.22 |
| birds | Dendropicos_stierlingi | Dendropicos_stierlingi | 0 | 28 | 2 | 534905.48 |
| birds | Dendroplex_picus | Dendroplex_picus | 0 | 41.3 | 1.7 | 9856567.73 |
| birds | Dendrortyx_barbatus | Dendrortyx_barbatus | 1 | 432 | 3.5 | 8177.78 |
| birds | Dendrortyx_leucophrys | Dendrortyx_leucophrys | 0 | 367.4 | 4.5 | 117127.03 |
| birds | Dendrortyx_macroura | Dendrortyx_macroura | 0 | 431 | 4 | 136253.42 |
| birds | Deroptyus_accipitrinus | Deroptyus_accipitrinus | 0 | 246 | 2.4 | 4158548.58 |
| birds | Dessonornis_anomalus | Cossypha_anomala | 0 | 24.5 | 2 | 8084.35 |
| birds | Dessonornis_archeri | Cossypha_archeri | 0 | 22.8 | 2 | 60103.14 |
| birds | Dessonornis_caffer | Cossypha_caffra | 0 | 28.5 | 2.4 | 1773198.12 |
| birds | Dessonornis_humeralis | Cossypha_humeralis | 0 | 22.8 | 3 | 644971.31 |
| birds | Devioeca_papuana | Microeca_papuana | 0 | 14.4 | 2 | 142595.35 |
| birds | Dicaeum_agile | Dicaeum_agile | 0 | 9 | 2.8 | 4429303.46 |
| birds | Dicaeum_celebicum | Dicaeum_celebicum | 0 | 6.5 | 3 | 184624.47 |

| | | | | | | |
|---|---|---|---|---|---|---|
| birds | Dicaeum_chrysorrheum | Dicaeum_chrysorrheum | 0 | 9 | 2.4 | 3155552.17 |
| birds | Dicaeum_concolor | Dicaeum_concolor | 0 | 6.2 | 2.4 | 221339.98 |
| birds | Dicaeum_cruentatum | Dicaeum_cruentatum | 0 | 5.6 | 2.8 | 4479690.77 |
| birds | Dicaeum_erythrorhynchos | Dicaeum_erythrorhynchos | 0 | 6.3 | 1.7 | 2149808.71 |
| birds | Dicaeum_geelvinkianum | Dicaeum_geelvinkianum | 0 | 6.8 | 2.4 | 535209.77 |
| birds | Dicaeum_haematostictum | Dicaeum_haematostictum | 0 | 8.8 | 1 | 24928.06 |
| birds | Dicaeum_hirundinaceum | Dicaeum_hirundinaceum | 0 | 9 | 3.5 | 6464367.1 |
| birds | Dicaeum_ignipectus | Dicaeum_ignipectus | 0 | 5.7 | 2.4 | 4029931.67 |
| birds | Dicaeum_maugei | Dicaeum_maugei | 0 | 8.5 | 2 | 39542.57 |
| birds | Dicaeum_melanozanthum | Dicaeum_melanoxanthum | 0 | 8.5 | 3 | 1518409.57 |
| birds | Dicaeum_trigonostigma | Dicaeum_trigonostigma | 0 | 7.1 | 2.4 | 2335989.48 |
| birds | Dicaeum_vincens | Dicaeum_vincens | 0 | 8.5 | 2 | 12166.57 |
| birds | Dicrurus_adsimilis | Dicrurus_adsimilis | 0 | 40.3 | 2.4 | 14611485.32 |
| birds | Dicrurus_aeneus | Dicrurus_aeneus | 0 | 26 | 2.8 | 4639369.76 |
| birds | Dicrurus_andamanensis | Dicrurus_andamanensis | 0 | 50.5 | 2.4 | 5726.81 |
| birds | Dicrurus_annectens | Dicrurus_annectans | 0 | 44 | 3.5 | 2725019.98 |
| birds | Dicrurus_atripennis | Dicrurus_atripennis | 0 | 38.9 | 2 | 1831426.11 |
| birds | Dicrurus_bracteatus | Dicrurus_bracteatus | 0 | 85.5 | 3.9 | 1975520.32 |
| birds | Dicrurus_caerulescens | Dicrurus_caerulescens | 0 | 40 | 2.4 | 2282100.81 |
| birds | Dicrurus_forficatus | Dicrurus_forficatus | 0 | 47.1 | 2.8 | 591728.75 |
| birds | Dicrurus_hottentottus | Dicrurus_hottentottus | 0 | 79.2 | 3.2 | 7412372.41 |
| birds | Dicrurus_leucophaeus | Dicrurus_leucophaeus | 0 | 37.6 | 3.5 | 9278672.07 |
| birds | Dicrurus_ludwigii | Dicrurus_ludwigii | 0 | 29.4 | 2.4 | 2047517.6 |
| birds | Dicrurus_macrocercus | Dicrurus_macrocercus | 0 | 48.3 | 3.2 | 9665439.19 |
| birds | Dicrurus_paradiseus | Dicrurus_paradiseus | 0 | 85.8 | 3.5 | 5998619.7 |
| birds | Dicrurus_remifer | Dicrurus_remifer | 0 | 43.1 | 2.8 | 2438166.85 |
| birds | Dinemellia_dinemelli | Dinemellia_dinemelli | 0 | 63.9 | 2.8 | 2361338.49 |
| birds | Dinopium_benghalense | Dinopium_benghalense | 0 | 100 | 2.4 | 3000600.76 |
| birds | Dinopium_javanense | Dinopium_javanense | 0 | 72.8 | 2.4 | 2707154.04 |
| birds | Dinopium_shorii | Dinopium_shorii | 0 | 101 | 2.4 | 382744.4 |
| birds | Diopsittaca_nobilis | Diopsittaca_nobilis | 0 | 150.9 | 4 | 495203.46 |
| birds | Discosura_longicaudus | Discosura_longicaudus | 0 | 3.4 | 2 | 2150114.71 |
| birds | Dolichonyx_oryzivorus | Dolichonyx_oryzivorus | 0 | 31.5 | 5 | 12537285.93 |
| birds | Donacobius_atricapilla | Donacobius_atricapilla | 0 | 36.8 | 2.4 | 8620634.56 |

| | | | | | | |
|---|---|---|---|---|---|---|
| birds | Doricha_eliza | Doricha_eliza | 0 | 2.5 | 2 | 4054.81 |
| birds | Doryfera_johannae | Doryfera_johannae | 0 | 4.8 | 2 | 459646.35 |
| birds | Doryfera_ludovicae | Doryfera_ludovicae | 0 | 6 | 2 | 387149.16 |
| birds | Drepanorhynchus_reichenowi | Nectarinia_reichenowi | 0 | 14.7 | 1 | 217754.73 |
| birds | Drepanornis_albertisi | Epimachus_albertisi | 0 | 110 | 1.4 | 23525.74 |
| birds | Dromaius_novaehollandiae | Dromaius_novaehollandiae | 0 | 34093.3 | 8.7 | 6731967.08 |
| birds | Dromas_ardeola | Dromas_ardeola | 0 | 325 | 1 | 1344551.83 |
| birds | Drymodes_brunneopygia | Drymodes_brunneopygia | 0 | 33 | 1 | 916861.69 |
| birds | Drymodes_superciliaris | Drymodes_superciliaris | 0 | 45.9 | 1.4 | 12382.11 |
| birds | Drymophila_squamata | Drymophila_squamata | 0 | 10.8 | 2 | 340038.68 |
| birds | Drymornis_bridgesii | Drymornis_bridgesii | 0 | 92.6 | 2.4 | 1433240.56 |
| birds | Dryobates_cathpharius | Dendrocopos_cathpharius | 0 | 30.7 | 2.8 | 125806.92 |
| birds | Dryobates_minor | Dendrocopos_minor | 0 | 19.8 | 5.5 | 18548908.46 |
| birds | Dryobates_nuttallii | Picoides_nuttallii | 0 | 37.4 | 4.5 | 183833.95 |
| birds | Dryobates_pubescens | Picoides_pubescens | 0 | 25.6 | 4.5 | 12800223.82 |
| birds | Dryobates_scalaris | Picoides_scalaris | 0 | 31.6 | 3.9 | 2902465.04 |
| birds | Dryocopus_javensis | Dryocopus_javensis | 0 | 271.4 | 2.8 | 2693444.77 |
| birds | Dryocopus_martius | Dryocopus_martius | 0 | 321 | 3.9 | 19634445.89 |
| birds | Dryolimnas_cuvieri | Dryolimnas_cuvieri | 0 | 182.4 | 3.5 | 590995.52 |
| birds | Dryoscopus_cubla | Dryoscopus_cubla | 0 | 26.2 | 2 | 5720470.9 |
| birds | Dryoscopus_gambensis | Dryoscopus_gambensis | 0 | 31.9 | 2 | 5608614.97 |
| birds | Dryoscopus_pringlii | Dryoscopus_pringlii | 0 | 19.7 | 2 | 448813.35 |
| birds | Dryoscopus_senegalensis | Dryoscopus_senegalensis | 0 | 29.3 | 2 | 1438969 |
| birds | Ducula_aenea | Ducula_aenea | 0 | 545 | 1.4 | 4813912.02 |
| birds | Ducula_badia | Ducula_badia | 0 | 486 | 1 | 2980351.8 |
| birds | Ducula_bicolor | Ducula_bicolor | 0 | 440 | 1 | 770004.59 |
| birds | Ducula_carola | Ducula_carola | 1 | 289 | 1 | 222701.31 |
| birds | Ducula_lacernulata | Ducula_lacernulata | 0 | 150 | 1 | 68766.83 |
| birds | Ducula_latrans | Ducula_latrans | 0 | 365.1 | 1 | 18103.65 |
| birds | Ducula_melanochroa | Ducula_melanochroa | 0 | 665 | 1 | 36132.41 |
| birds | Ducula_mullerii | Ducula_mullerii | 0 | 574.9 | 1 | 281060.32 |
| birds | Ducula_pacifica | Ducula_pacifica | 0 | 394.2 | 1 | 29576.6 |
| birds | Ducula_pinon | Ducula_pinon | 0 | 756 | 1 | 639390.56 |
| birds | Ducula_pistrinaria | Ducula_pistrinaria | 0 | 515 | 1 | 85422.33 |

| | | | | | | |
|---|---|---|---|---|---|---|
| birds | Ducula_radiata | Ducula_radiata | 0 | 295 | 1 | 154747.93 |
| birds | Ducula_rubricera | Ducula_rubricera | 0 | 652 | 1 | 72417.84 |
| birds | Ducula_spilorrhoa | Ducula_spilorrhoa | 0 | 464 | 1 | 518577.49 |
| birds | Dumetella_carolinensis | Dumetella_carolinensis | 0 | 35.3 | 3.5 | 8544736.27 |
| birds | Dyaphorophyia_blissetti | Platysteira_blissetti | 0 | 11 | 2 | 473029.12 |
| birds | Dyaphorophyia_castanea | Platysteira_castanea | 0 | 14.1 | 2 | 3596708.42 |
| birds | Dyaphorophyia_concreta | Platysteira_concreta | 0 | 11.4 | 2 | 175431.5 |
| birds | Dysithamnus_mentalis | Dysithamnus_mentalis | 0 | 14.9 | 2 | 4153057.29 |
| birds | Dysithamnus_plumbeus | Dysithamnus_plumbeus | 1 | 20.5 | 2 | 144601.21 |
| birds | Dysithamnus_puncticeps | Dysithamnus_puncticeps | 0 | 14.9 | 2 | 127823.89 |
| birds | Dysithamnus_striaticeps | Dysithamnus_striaticeps | 0 | 12 | 2 | 88175.27 |
| birds | Dysithamnus_xanthopterus | Dysithamnus_xanthopterus | 0 | 16 | 2 | 46590.47 |
| birds | Eclectus_roratus | Eclectus_roratus | 0 | 480.8 | 2 | 52021.22 |
| birds | Edolisoma_ostentum | Coracina_ostenta | 1 | 68 | 2.4 | 23988.16 |
| birds | Edolisoma_tenuirostre | Coracina_tenuirostris | 0 | 60.9 | 1 | 2156165.11 |
| birds | Egretta_ardesiaca | Egretta_ardesiaca | 0 | 324 | 2.8 | 8505004.79 |
| birds | Egretta_eulophotes | Egretta_eulophotes | 1 | 461.8 | 3.2 | 860382.31 |
| birds | Egretta_garzetta | Egretta_garzetta | 0 | 312 | 3.5 | 38343874.82 |
| birds | Egretta_novaehollandiae | Egretta_novaehollandiae | 0 | 558.6 | 3.9 | 7533663.44 |
| birds | Egretta_picata | Ardea_picata | 0 | 259 | 2.8 | 1218679.45 |
| birds | Egretta_rufescens | Egretta_rufescens | 0 | 614 | 3.5 | 580944.39 |
| birds | Egretta_thula | Egretta_thula | 0 | 371 | 3.2 | 22001375.46 |
| birds | Egretta_tricolor | Egretta_tricolor | 0 | 372.3 | 3.5 | 2501648.23 |
| birds | Egretta_vinaceigula | Egretta_vinaceigula | 1 | 288 | 2.4 | 1053459.65 |
| birds | Elaenia_albiceps | Elaenia_albiceps | 0 | 15.5 | 2.4 | 13323047.2 |
| birds | Elaenia_chiriquensis | Elaenia_chiriquensis | 0 | 15.4 | 2 | 8478302.49 |
| birds | Elaenia_fallax | Elaenia_fallax | 0 | 14.4 | 2 | 6030.05 |
| birds | Elaenia_flavogaster | Elaenia_flavogaster | 0 | 24.8 | 1.7 | 10327940.88 |
| birds | Elaenia_frantzii | Elaenia_frantzii | 0 | 19.6 | 2 | 266786.47 |
| birds | Elaenia_martinica | Elaenia_martinica | 0 | 20.8 | 2.4 | 26788.39 |
| birds | Elaenia_mesoleuca | Elaenia_mesoleuca | 0 | 17.6 | 2.4 | 2216525.02 |
| birds | Elaenia_parvirostris | Elaenia_parvirostris | 0 | 13.8 | 2.4 | 10962786.12 |
| birds | Elanoides_forficatus | Elanoides_forficatus | 0 | 416.2 | 1.4 | 14031977.44 |
| birds | Elanus_axillaris | Elanus_axillaris | 0 | 270.1 | 3.5 | 7649038.18 |

| | | | | | | |
|---|---|---|---|---|---|---|
| birds | Elanus_caeruleus | Elanus_caeruleus | 0 | 259.8 | 3.5 | 26925792.46 |
| birds | Elanus_leucurus | Elanus_leucurus | 0 | 346 | 3.9 | 12522294.39 |
| birds | Elanus_scriptus | Elanus_scriptus | 0 | 316 | 4.5 | 427962.13 |
| birds | Eleoscytalopus_indigoticus | Eleoscytalopus_indigoticus | 0 | 16.2 | 2 | 455549.55 |
| birds | Eleoscytalopus_psychopompus | Eleoscytalopus_psychopompus | 1 | 17.8 | 2 | 2626.39 |
| birds | Eleothreptus_candicans | Eleothreptus_candicans | 1 | 49.3 | 2 | 1398605.14 |
| birds | Elliotomyia_chionogaster | Amazilia_chionogaster | 0 | 5 | 2 | 488801.52 |
| birds | Elliotomyia_viridicauda | Amazilia_viridicauda | 0 | 6.1 | 2 | 28878.13 |
| birds | Elminia_albicauda | Elminia_albicauda | 0 | 8.3 | 2 | 1971246.25 |
| birds | Elminia_albiventris | Elminia_albiventris | 0 | 8.9 | 2 | 224843.85 |
| birds | Elminia_albonotata | Elminia_albonotata | 0 | 9.2 | 1.7 | 905274.18 |
| birds | Elminia_longicauda | Elminia_longicauda | 0 | 9.9 | 1.4 | 3910894.16 |
| birds | Elminia_nigromitrata | Elminia_nigromitrata | 0 | 9 | 2 | 2433804.71 |
| birds | Elseyornis_melanops | Elseyornis_melanops | 0 | 33 | 3 | 7354478.04 |
| birds | Emarginata_schlegelii | Cercomela_schlegelii | 0 | 23.3 | 2.8 | 1144740.06 |
| birds | Emarginata_sinuata | Cercomela_sinuata | 0 | 18.1 | 2.4 | 793361.3 |
| birds | Emarginata_tractrac | Cercomela_tractrac | 0 | 20.8 | 2.4 | 1018879.12 |
| birds | Emberiza_aureola | Emberiza_aureola | 1 | 19.6 | 3.9 | 17060140.55 |
| birds | Emberiza_bruniceps | Emberiza_bruniceps | 0 | 24.8 | 4.5 | 5514147.76 |
| birds | Emberiza_buchanani | Emberiza_buchanani | 0 | 21.2 | 4.5 | 4008927.89 |
| birds | Emberiza_cabanisi | Emberiza_cabanisi | 0 | 24.9 | 2.4 | 2851364.02 |
| birds | Emberiza_caesia | Emberiza_caesia | 0 | 19.9 | 4.5 | 1505590.67 |
| birds | Emberiza_calandra | Miliaria_calandra | 0 | 48.5 | 4.2 | 8461055.91 |
| birds | Emberiza_capensis | Emberiza_capensis | 0 | 20.9 | 2.8 | 1320689.84 |
| birds | Emberiza_chrysophrys | Emberiza_chrysophrys | 0 | 18 | 3.9 | 3119549.04 |
| birds | Emberiza_cia | Emberiza_cia | 0 | 23.4 | 3.9 | 5760543.74 |
| birds | Emberiza_cineracea | Emberiza_cineracea | 0 | 24.3 | 4.9 | 1304311.53 |
| birds | Emberiza_cioides | Emberiza_cioides | 0 | 21 | 4.2 | 10532097.13 |
| birds | Emberiza_cirlus | Emberiza_cirlus | 0 | 25.6 | 3.2 | 2210307.26 |
| birds | Emberiza_citrinella | Emberiza_citrinella | 0 | 29.7 | 3.9 | 16427829.18 |
| birds | Emberiza_elegans | Emberiza_elegans | 0 | 16.8 | 4.9 | 3666177.19 |
| birds | Emberiza_flaviventris | Emberiza_flaviventris | 0 | 18.2 | 2.4 | 6969113.54 |
| birds | Emberiza_fucata | Emberiza_fucata | 0 | 19.8 | 3.9 | 8426706.7 |
| birds | Emberiza_godlewskii | Emberiza_godlewskii | 0 | 19 | 3.9 | 6786390.71 |

| | | | | | | |
|---|---|---|---|---|---|---|
| birds | Emberiza_hortulana | Emberiza_hortulana | 0 | 19.9 | 4.5 | 10888484.28 |
| birds | Emberiza_impetuani | Emberiza_impetuani | 0 | 15.1 | 3.5 | 1657377.13 |
| birds | Emberiza_jankowskii | Emberiza_jankowskii | 1 | 22.5 | 4.5 | 288918.89 |
| birds | Emberiza_leucocephalos | Emberiza_leucocephalos | 0 | 28.5 | 4.5 | 14053729.43 |
| birds | Emberiza_melanocephala | Emberiza_melanocephala | 0 | 28.3 | 3.9 | 2718177.53 |
| birds | Emberiza_pallasi | Emberiza_pallasi | 0 | 14 | 3.9 | 14847467.73 |
| birds | Emberiza_poliopleura | Emberiza_poliopleura | 0 | 15.8 | 2.4 | 1250196.23 |
| birds | Emberiza_pusilla | Emberiza_pusilla | 0 | 14.6 | 4.9 | 13018743.8 |
| birds | Emberiza_rustica | Emberiza_rustica | 1 | 20.5 | 4.9 | 22901225.25 |
| birds | Emberiza_rutila | Emberiza_rutila | 0 | 17.5 | 4 | 5656279.1 |
| birds | Emberiza_schoeniclus | Emberiza_schoeniclus | 0 | 18.4 | 4.5 | 36135699.04 |
| birds | Emberiza_spodocephala | Emberiza_spodocephala | 0 | 18.5 | 4.5 | 6997496.16 |
| birds | Emberiza_stewarti | Emberiza_stewarti | 0 | 16 | 3.9 | 1817857.29 |
| birds | Emberiza_striolata | Emberiza_striolata | 0 | 14.8 | 2.8 | 2669183.7 |
| birds | Emberiza_sulphurata | Emberiza_sulphurata | 0 | 14.9 | 3.9 | 542896.51 |
| birds | Emberiza_tahapisi | Emberiza_tahapisi | 0 | 13 | 2.8 | 4092995.92 |
| birds | Emberiza_tristrami | Emberiza_tristrami | 0 | 18 | 4.5 | 1229215.12 |
| birds | Emberiza_variabilis | Emberiza_variabilis | 0 | 23.8 | 3.9 | 409045.4 |
| birds | Emberiza_yessoensis | Emberiza_yessoensis | 0 | 13.5 | 3.9 | 1815524.15 |
| birds | Emberizoides_herbicola | Emberizoides_herbicola | 0 | 27.6 | 2 | 5561384.25 |
| birds | Emblema_pictum | Emblema_pictum | 0 | 10.4 | 3.5 | 1295960.93 |
| birds | Eminia_lepida | Eminia_lepida | 0 | 20.2 | 2.4 | 631640.65 |
| birds | Empidonax_albigularis | Empidonax_albigularis | 0 | 9.6 | 2.4 | 1056434.02 |
| birds | Empidonax_alnorum | Empidonax_alnorum | 0 | 12.7 | 3.5 | 13408407.31 |
| birds | Empidonax_atriceps | Empidonax_atriceps | 0 | 9 | 2 | 3661.22 |
| birds | Empidonax_difficilis | Empidonax_difficilis | 0 | 10.7 | 2.2 | 1682980.97 |
| birds | Empidonax_flavescens | Empidonax_flavescens | 0 | 12.5 | 2.4 | 167716.78 |
| birds | Empidonax_flaviventris | Empidonax_flaviventris | 0 | 11.8 | 3.9 | 8805558.82 |
| birds | Empidonax_fulvifrons | Empidonax_fulvifrons | 0 | 7.9 | 3.2 | 729421.79 |
| birds | Empidonax_hammondii | Empidonax_hammondii | 0 | 10.4 | 3.5 | 5603866.93 |
| birds | Empidonax_minimus | Empidonax_minimus | 0 | 10 | 4 | 11595266.47 |
| birds | Empidonax_oberholseri | Empidonax_oberholseri | 0 | 10.4 | 3.5 | 4614600.73 |
| birds | Empidonax_occidentalis | Empidonax_occidentalis | 0 | 11.6 | 3.5 | 2964611.73 |
| birds | Empidonax_traillii | Empidonax_traillii | 0 | 13.4 | 3.5 | 11375149 |

| | | | | | | |
|---|---|---|---|---|---|---|
| birds | Empidonax_virescens | Empidonax_virescens | 0 | 12.6 | 2 | 5003149.64 |
| birds | Empidonax_wrightii | Empidonax_wrightii | 0 | 12.3 | 3.5 | 2898385.44 |
| birds | Empidonomus_varius | Empidonomus_varius | 0 | 27.1 | 2.4 | 13118402.85 |
| birds | Empidornis_semipartitus | Empidornis_semipartitus | 0 | 22.5 | 2 | 662620.71 |
| birds | Enicognathus_ferrugineus | Enicognathus_ferrugineus | 0 | 160 | 5.7 | 572089.61 |
| birds | Enicognathus_leptorhynchus | Enicognathus_leptorhynchus | 0 | 193 | 3.5 | 107787.94 |
| birds | Enicurus_immaculatus | Enicurus_immaculatus | 0 | 25.5 | 3 | 921267.82 |
| birds | Enicurus_leschenaulti | Enicurus_leschenaulti | 0 | 33.8 | 2.8 | 6166589.04 |
| birds | Enicurus_maculatus | Enicurus_maculatus | 0 | 40.4 | 3.5 | 1640498.35 |
| birds | Enicurus_ruficapillus | Enicurus_ruficapillus | 0 | 28.2 | 2 | 1105095.6 |
| birds | Enicurus_schistaceus | Enicurus_schistaceus | 0 | 31 | 2.8 | 2441930.12 |
| birds | Enicurus_scouleri | Enicurus_scouleri | 0 | 17.8 | 2.8 | 3341040.76 |
| birds | Entomodestes_coracinus | Entomodestes_coracinus | 0 | 56 | 2 | 21017.95 |
| birds | Entomyzon_cyanotis | Entomyzon_cyanotis | 0 | 105.3 | 2 | 2004063.99 |
| birds | Eolophus_roseicapilla | Cacatua_roseicapilla | 0 | 328 | 3.5 | 7020540.5 |
| birds | Eopsaltria_australis | Eopsaltria_australis | 0 | 20.2 | 2.4 | 1174338.84 |
| birds | Eopsaltria_griseogularis | Eopsaltria_griseogularis | 0 | 19.4 | 2 | 587762.32 |
| birds | Eos_cyanogenia | Eos_cyanogenia | 0 | 167 | 2 | 2964.62 |
| birds | Eos_reticulata | Eos_reticulata | 0 | 153 | 2 | 4779.51 |
| birds | Eos_squamata | Eos_squamata | 0 | 91 | 2 | 32014.97 |
| birds | Ephippiorhynchus_asiaticus | Ephippiorhynchus_asiaticus | 0 | 4100 | 3.2 | 5889190.75 |
| birds | Ephippiorhynchus_senegalensis | Ephippiorhynchus_senegalensis | 0 | 6158.7 | 2.2 | 11071552.53 |
| birds | Epimachus_meyeri | Epimachus_meyeri | 0 | 220.5 | 1 | 135052.62 |
| birds | Epinecrophylla_fulviventris | Epinecrophylla_fulviventris | 0 | 10.4 | 2 | 281139.81 |
| birds | Epinecrophylla_gutturalis | Myrmotherula_gutturalis | 0 | 9.2 | 2 | 1137766.7 |
| birds | Epthianura_albifrons | Epthianura_albifrons | 0 | 13.3 | 3 | 2667021.56 |
| birds | Epthianura_aurifrons | Epthianura_aurifrons | 0 | 10.5 | 2.4 | 5609912.51 |
| birds | Epthianura_crocea | Epthianura_crocea | 0 | 9.3 | 3 | 2179684.97 |
| birds | Epthianura_tricolor | Epthianura_tricolor | 0 | 10.7 | 3.5 | 5486066.87 |
| birds | Eremalauda_dunni | Eremalauda_dunni | 0 | 20.5 | 2.4 | 1825439.35 |
| birds | Eremomela_atricollis | Eremomela_atricollis | 0 | 10.2 | 2 | 1063082.23 |
| birds | Eremomela_flavicrissalis | Eremomela_flavicrissalis | 0 | 4.5 | 1.4 | 893727.78 |
| birds | Eremomela_gregalis | Eremomela_gregalis | 0 | 6.8 | 3.5 | 271515.78 |
| birds | Eremomela_icteropygialis | Eremomela_icteropygialis | 0 | 7.5 | 1.7 | 8262133.25 |

| | | | | | | |
|---|---|---|---|---|---|---|
| birds | Eremomela_pusilla | Eremomela_pusilla | 0 | 6.2 | 2 | 2960274.14 |
| birds | Eremomela_scotops | Eremomela_scotops | 0 | 9 | 2 | 4568185.56 |
| birds | Eremomela_usticollis | Eremomela_usticollis | 0 | 8.4 | 2 | 1430953.15 |
| birds | Eremophila_alpestris | Eremophila_alpestris | 0 | 33.3 | 3.2 | 44034741.82 |
| birds | Eremophila_bilopha | Eremophila_bilopha | 0 | 38.5 | 2.8 | 1422315.02 |
| birds | Eremopterix_australis | Eremopterix_australis | 0 | 14.2 | 2.8 | 642788.97 |
| birds | Eremopterix_griseus | Eremopterix_griseus | 0 | 16 | 2.4 | 2966642.7 |
| birds | Eremopterix_hova | Mirafra_hova | 0 | 18.9 | 2 | 591120.51 |
| birds | Eremopterix_leucopareia | Eremopterix_leucopareia | 0 | 14.5 | 2.4 | 859362.7 |
| birds | Eremopterix_leucotis | Eremopterix_leucotis | 0 | 16.4 | 1.4 | 7621320.47 |
| birds | Eremopterix_nigriceps | Eremopterix_nigriceps | 0 | 13.9 | 2.8 | 7838188.48 |
| birds | Eremopterix_signatus | Eremopterix_signatus | 0 | 15.7 | 4 | 970819.66 |
| birds | Eremopterix_verticalis | Eremopterix_verticalis | 0 | 17 | 2.4 | 3048289.88 |
| birds | Eriocnemis_cupreoventris | Eriocnemis_cupreoventris | 0 | 5.3 | 2 | 27252.17 |
| birds | Eriocnemis_derbyi | Eriocnemis_derbyi | 0 | 5.5 | 2 | 31057.37 |
| birds | Eriocnemis_glaucopoides | Eriocnemis_glaucopoides | 0 | 3.7 | 2 | 57142.21 |
| birds | Eriocnemis_godini | Eriocnemis_godini | 1 | 5.1 | 2 | 408.05 |
| birds | Eriocnemis_luciani | Eriocnemis_luciani | 0 | 6 | 2 | 39822.97 |
| birds | Eriocnemis_mosquera | Eriocnemis_mosquera | 0 | 5.3 | 2 | 36125.45 |
| birds | Eriocnemis_nigrivestis | Eriocnemis_nigrivestis | 1 | 8.5 | 2 | 967.35 |
| birds | Eriocnemis_vestita | Eriocnemis_vestita | 0 | 4.5 | 2 | 97851.28 |
| birds | Erithacus_rubecula | Erithacus_rubecula | 0 | 17.7 | 4.9 | 13266206.91 |
| birds | Erythrocercus_livingstonei | Erythrocercus_livingstonei | 0 | 5.9 | 2.4 | 1133267.59 |
| birds | Erythrocercus_mccallii | Erythrocercus_mccallii | 0 | 7.2 | 2 | 2166425.08 |
| birds | Erythrogonys_cinctus | Erythrogonys_cinctus | 0 | 51.4 | 4 | 5111119.63 |
| birds | Erythropitta_arquata | Pitta_arcuata | 0 | 54 | 2 | 214049.06 |
| birds | Erythropitta_erythrogaster | Pitta_erythrogaster | 0 | 72.4 | 2.8 | 289376.94 |
| birds | Erythropitta_granatina | Pitta_granatina | 0 | 61.5 | 2 | 852985.14 |
| birds | Erythropitta_ussheri | Pitta_ussheri | 0 | 55.3 | 2 | 41122.35 |
| birds | Erythrotriorchis_radiatus | Erythrotriorchis_radiatus | 1 | 887.4 | 2 | 1042091.38 |
| birds | Erythrura_coloria | Erythrura_coloria | 0 | 11.3 | 1.4 | 12662.23 |
| birds | Erythrura_cyaneovirens | Erythrura_cyaneovirens | 0 | 13 | 3.5 | 14573.35 |
| birds | Erythrura_hyperythra | Erythrura_hyperythra | 0 | 11 | 4 | 294579.35 |
| birds | Erythrura_pealii | Erythrura_pealii | 0 | 12 | 3.5 | 17012.67 |

| | | | | | | |
|---|---|---|---|---|---|---|
| birds | Erythrura_prasina | Erythrura_prasina | 0 | 14.7 | 4.9 | 1585424.03 |
| birds | Erythrura_psittacea | Erythrura_psittacea | 0 | 11.5 | 4 | 17405.19 |
| birds | Erythrura_trichroa | Erythrura_trichroa | 0 | 14.4 | 3.5 | 858792.1 |
| birds | Erythrura_viridifacies | Erythrura_viridifacies | 1 | 12.6 | 3 | 106324.25 |
| birds | Esacus_magnirostris | Esacus_giganteus | 0 | 1015.9 | 1 | 2207239.52 |
| birds | Esacus_recurvirostris | Esacus_recurvirostris | 0 | 790 | 1.4 | 2969603.17 |
| birds | Estrilda_astrild | Estrilda_astrild | 0 | 8.3 | 4.9 | 6015986.23 |
| birds | Estrilda_atricapilla | Estrilda_atricapilla | 0 | 7.8 | 2 | 665104.34 |
| birds | Estrilda_melpoda | Estrilda_melpoda | 0 | 7.6 | 4.9 | 3755904.77 |
| birds | Estrilda_nonnula | Estrilda_nonnula | 0 | 7.7 | 5.5 | 1216438.93 |
| birds | Estrilda_paludicola | Estrilda_paludicola | 0 | 7.6 | 4.9 | 1983230.36 |
| birds | Estrilda_troglodytes | Estrilda_troglodytes | 0 | 7.6 | 4.5 | 2017180.16 |
| birds | Eubucco_bourcierii | Eubucco_bourcierii | 0 | 37 | 3.2 | 234502.7 |
| birds | Eucometis_penicillata | Eucometis_penicillata | 0 | 27 | 2 | 5712449.56 |
| birds | Eudocimus_albus | Eudocimus_albus | 0 | 889.7 | 2.4 | 1931978.98 |
| birds | Eudocimus_ruber | Eudocimus_ruber | 0 | 660.1 | 2 | 816290.78 |
| birds | Eudromia_elegans | Eudromia_elegans | 0 | 702.9 | 5.5 | 1493624.4 |
| birds | Eudromias_morinellus | Eudromias_morinellus | 0 | 108.2 | 3 | 4095968.09 |
| birds | Eugenes_fulgens | Eugenes_fulgens | 0 | 8 | 2 | 1476036.28 |
| birds | Eugralla_paradoxa | Eugralla_paradoxa | 0 | 44 | 2.4 | 124178 |
| birds | Eulabeornis_castaneoventris | Eulabeornis_castaneoventris | 0 | 684.5 | 4.5 | 85822.05 |
| birds | Eulampis_holosericeus | Eulampis_holosericeus | 0 | 5.7 | 2 | 8331.82 |
| birds | Eumyias_additus | Rhinomyias_additus | 0 | 22.5 | 2.4 | 4091.97 |
| birds | Eumyias_albicaudatus | Eumyias_albicaudatus | 0 | 16.4 | 3 | 32809.71 |
| birds | Eumyias_sordidus | Eumyias_sordidus | 0 | 25.1 | 2.4 | 5844.08 |
| birds | Eumyias_thalassinus | Eumyias_thalassinus | 0 | 18.1 | 3.5 | 7039599.11 |
| birds | Euodice_cantans | Lonchura_cantans | 0 | 12 | 4.2 | 3802160.03 |
| birds | Euodice_malabarica | Lonchura_malabarica | 0 | 12 | 4.6 | 3376406.35 |
| birds | Eupetomena_cirrochloris | Aphantochroa_cirrochloris | 0 | 7.7 | 2 | 1438300.99 |
| birds | Eupetomena_macroura | Eupetomena_macroura | 0 | 9 | 2 | 4594117.6 |
| birds | Euphagus_carolinus | Euphagus_carolinus | 1 | 59.6 | 4.5 | 12263114.38 |
| birds | Euphagus_cyanocephalus | Euphagus_cyanocephalus | 0 | 62.5 | 5 | 10214123.73 |
| birds | Eupherusa_cyanophrys | Eupherusa_cyanophrys | 1 | 4.8 | 2 | 4063.86 |
| birds | Eupherusa_nigriventris | Eupherusa_nigriventris | 0 | 3.5 | 2 | 14338.29 |

| | | | | | | |
|---|---|---|---|---|---|---|
| birds | Euphonia_affinis | Euphonia_affinis | 0 | 10 | 3 | 724298.2 |
| birds | Euphonia_cayennensis | Euphonia_cayennensis | 0 | 14.3 | 3.9 | 1272525.45 |
| birds | Euphonia_chlorotica | Euphonia_chlorotica | 0 | 11 | 3.9 | 9620331.07 |
| birds | Euphonia_cyanocephala | Euphonia_cyanocephala | 0 | 14 | 2.4 | 2711347.81 |
| birds | Euphonia_elegantissima | Euphonia_elegantissima | 0 | 15 | 3 | 473565.77 |
| birds | Euphonia_gouldi | Euphonia_gouldi | 0 | 14 | 2.8 | 326598.75 |
| birds | Euphonia_hirundinacea | Euphonia_hirundinacea | 0 | 14 | 5 | 642669.45 |
| birds | Euphonia_imitans | Euphonia_imitans | 0 | 14 | 3 | 14869.65 |
| birds | Euphonia_jamaica | Euphonia_jamaica | 0 | 16.6 | 3.5 | 11021.79 |
| birds | Euphonia_laniirostris | Euphonia_laniirostris | 0 | 15 | 4 | 4464958.75 |
| birds | Euphonia_luteicapilla | Euphonia_luteicapilla | 0 | 13 | 3 | 111584.47 |
| birds | Euphonia_minuta | Euphonia_minuta | 0 | 10 | 3 | 5149944.3 |
| birds | Euphonia_musica | Euphonia_musica | 0 | 14.7 | 4 | 35327.88 |
| birds | Euphonia_pectoralis | Euphonia_pectoralis | 0 | 14.4 | 3 | 1361244.45 |
| birds | Euphonia_trinitatis | Euphonia_trinitatis | 0 | 11 | 3.5 | 610172.65 |
| birds | Euphonia_violacea | Euphonia_violacea | 0 | 15 | 4 | 4921834.31 |
| birds | Euphonia_xanthogaster | Euphonia_xanthogaster | 0 | 13 | 4 | 3715684.5 |
| birds | Euplectes_afer | Euplectes_afer | 0 | 14.3 | 3.2 | 3118130.65 |
| birds | Euplectes_albonotatus | Euplectes_albonotatus | 0 | 21.1 | 3.2 | 2575317.41 |
| birds | Euplectes_ardens | Euplectes_ardens | 0 | 19 | 3.5 | 3182674.54 |
| birds | Euplectes_axillaris | Euplectes_axillaris | 0 | 25.1 | 2.8 | 2371547.67 |
| birds | Euplectes_capensis | Euplectes_capensis | 0 | 30.5 | 2.8 | 2639722.61 |
| birds | Euplectes_franciscanus | Euplectes_franciscanus | 0 | 16 | 2.8 | 3267667.67 |
| birds | Euplectes_hordeaceus | Euplectes_hordeaceus | 0 | 19.4 | 3 | 5103979.03 |
| birds | Euplectes_jacksoni | Euplectes_jacksoni | 0 | 37 | 2.4 | 107944.78 |
| birds | Euplectes_macroura | Euplectes_macroura | 0 | 21.7 | 2.4 | 3993290.03 |
| birds | Euplectes_orix | Euplectes_orix | 0 | 23 | 3.2 | 2969008.67 |
| birds | Euplectes_progne | Euplectes_progne | 0 | 35.1 | 2.8 | 697961.28 |
| birds | Eupodotis_caerulescens | Eupodotis_caerulescens | 0 | 1366 | 1.7 | 264843.99 |
| birds | Eupodotis_senegalensis | Eupodotis_senegalensis | 0 | 585 | 1.7 | 4886826.69 |
| birds | Eupsittula_aurea | Aratinga_aurea | 0 | 84.6 | 2.8 | 5764751.61 |
| birds | Eupsittula_cactorum | Aratinga_cactorum | 0 | 82.5 | 6 | 1195222.88 |
| birds | Eupsittula_canicularis | Aratinga_canicularis | 1 | 85 | 3.9 | 283885.27 |
| birds | Eupsittula_nana | Aratinga_nana | 0 | 76.9 | 3.9 | 11021.79 |

| | | | | | | |
|---|---|---|---|---|---|---|
| birds | Eupsittula_pertinax | Aratinga_pertinax | 0 | 84 | 3.7 | 1480961 |
| birds | Euptilotus_eutilotus | Pycnonotus_eutilotus | 0 | 35.3 | 2 | 932677.68 |
| birds | Eurillas_gracilis | Andropadus_gracilis | 0 | 21.6 | 2 | 3552112.86 |
| birds | Eurillas_latirostris | Andropadus_latirostris | 0 | 26.2 | 2 | 3682285.72 |
| birds | Eurillas_virens | Andropadus_virens | 0 | 26.6 | 2 | 5766996.33 |
| birds | Eurocephalus_anguitimens | Eurocephalus_anguitimens | 0 | 69 | 3.5 | 1521047.71 |
| birds | Eurocephalus_ruppelli | Eurocephalus_rueppelli | 0 | 50.6 | 2.8 | 1835004.73 |
| birds | Eurostopodus_archboldi | Eurostopodus_archboldi | 0 | 77 | 1 | 74591.66 |
| birds | Eurostopodus_argus | Eurostopodus_argus | 0 | 93.4 | 1 | 7014029.22 |
| birds | Eurostopodus_diabolicus | Eurostopodus_diabolicus | 0 | 104.4 | 1 | 18765.43 |
| birds | Eurostopodus_mystacalis | Eurostopodus_mystacalis | 0 | 134.5 | 1 | 1216308.42 |
| birds | Eurostopodus_papuensis | Eurostopodus_papuensis | 0 | 80.5 | 1 | 502860.66 |
| birds | Eurylaimus_javanicus | Eurylaimus_javanicus | 0 | 78.2 | 2.4 | 126074.83 |
| birds | Eurylaimus_ochromalus | Eurylaimus_ochromalus | 0 | 33.3 | 2.4 | 1129454.95 |
| birds | Euryptila_subcinnamomea | Euryptila_subcinnamomea | 0 | 11.5 | 2.4 | 274057.07 |
| birds | Eurypyga_helias | Eurypyga_helias | 0 | 210 | 2 | 8423208.45 |
| birds | Eurystomus_glaucurus | Eurystomus_glaucurus | 0 | 110 | 2.4 | 13880859.89 |
| birds | Eurystomus_gularis | Eurystomus_gularis | 0 | 96.3 | 2.4 | 2827968.86 |
| birds | Eurystomus_orientalis | Eurystomus_orientalis | 0 | 143 | 3.9 | 12922409.59 |
| birds | Euscarthmus_meloryphus | Euscarthmus_meloryphus | 0 | 6.8 | 2 | 6211591.05 |
| birds | Eutoxeres_aquila | Eutoxeres_aquila | 0 | 10.6 | 2 | 306610.24 |
| birds | Eutoxeres_condamini | Eutoxeres_condamini | 0 | 10.8 | 2 | 370060.39 |
| birds | Falcipennis_falcipennis | Dendragapus_falcipennis | 0 | 685.6 | 7.7 | 692147.35 |
| birds | Falco_alopex | Falco_alopex | 0 | 273.9 | 2.4 | 5286034.09 |
| birds | Falco_amurensis | Falco_amurensis | 0 | 141.9 | 3.5 | 14657717.81 |
| birds | Falco_ardosiaceus | Falco_ardosiaceus | 0 | 234.2 | 3.9 | 7345675.56 |
| birds | Falco_berigora | Falco_berigora | 0 | 587 | 2.4 | 8273511.57 |
| birds | Falco_biarmicus | Falco_biarmicus | 0 | 606.1 | 3.5 | 18096388.23 |
| birds | Falco_cenchroides | Falco_cenchroides | 0 | 170.5 | 3.9 | 8034280.03 |
| birds | Falco_cherrug | Falco_cherrug | 1 | 961.2 | 3.9 | 38026625.06 |
| birds | Falco_chicquera | Falco_chicquera | 0 | 208 | 3.5 | 3166581.2 |
| birds | Falco_columbarius | Falco_columbarius | 0 | 188.5 | 4.2 | 47918643.64 |
| birds | Falco_concolor | Falco_concolor | 1 | 250 | 2.4 | 4355294.94 |
| birds | Falco_cuvierii | Falco_cuvierii | 0 | 182.2 | 3 | 7436820.01 |

| | | | | | | |
|---|---|---|---|---|---|---|
| birds | Falco_deiroleucus | Falco_deiroleucus | 0 | 459.6 | 1.7 | 3323144.47 |
| birds | Falco_dickinsoni | Falco_dickinsoni | 0 | 209 | 3 | 3765759.75 |
| birds | Falco_eleonorae | Falco_eleonorae | 0 | 390 | 2.4 | 5141958.02 |
| birds | Falco_fasciinucha | Falco_fasciinucha | 1 | 254.7 | 2.8 | 501059.82 |
| birds | Falco_femoralis | Falco_femoralis | 0 | 335.8 | 2.4 | 12113135.74 |
| birds | Falco_hypoleucos | Falco_hypoleucos | 1 | 457.2 | 2.4 | 3627330.54 |
| birds | Falco_jugger | Falco_jugger | 0 | 755 | 3.5 | 2912240.73 |
| birds | Falco_longipennis | Falco_longipennis | 0 | 249.8 | 2.4 | 8078573.63 |
| birds | Falco_mexicanus | Falco_mexicanus | 0 | 713.1 | 3.9 | 5425089.08 |
| birds | Falco_moluccensis | Falco_moluccensis | 0 | 162 | 4 | 466435.96 |
| birds | Falco_naumanni | Falco_naumanni | 0 | 152.1 | 4.2 | 43373581.24 |
| birds | Falco_newtoni | Falco_newtoni | 0 | 123.4 | 4 | 590995.81 |
| birds | Falco_novaeseelandiae | Falco_novaeseelandiae | 0 | 418.6 | 2.8 | 242840.06 |
| birds | Falco_peregrinus | Falco_peregrinus | 0 | 760 | 2.8 | 92354726.04 |
| birds | Falco_punctatus | Falco_punctatus | 1 | 167.4 | 3 | 773.93 |
| birds | Falco_rufigularis | Falco_rufigularis | 0 | 163.6 | 2.8 | 13377698.05 |
| birds | Falco_rupicoloides | Falco_rupicoloides | 0 | 261 | 3.5 | 3191508.18 |
| birds | Falco_rusticolus | Falco_rusticolus | 0 | 1431.7 | 3.5 | 26161318.43 |
| birds | Falco_severus | Falco_severus | 0 | 200.6 | 2.8 | 4016071.03 |
| birds | Falco_sparverius | Falco_sparverius | 0 | 114.6 | 4.9 | 27776585.21 |
| birds | Falco_subbuteo | Falco_subbuteo | 0 | 208.2 | 3 | 57025622.45 |
| birds | Falco_subniger | Falco_subniger | 0 | 776 | 3.5 | 5910471.62 |
| birds | Falco_tinnunculus | Falco_tinnunculus | 0 | 183.2 | 4.2 | 58595922.7 |
| birds | Falco_vespertinus | Falco_vespertinus | 1 | 151.5 | 3.5 | 33964689.97 |
| birds | Falculea_palliata | Falculea_palliata | 0 | 111 | 4 | 301726.39 |
| birds | Falcunculus_frontatus | Falcunculus_frontatus | 0 | 27.9 | 2.4 | 981576.39 |
| birds | Ferminia_cerverai | Ferminia_cerverai | 1 | 26 | 2 | 2102.53 |
| birds | Ficedula_albicollis | Ficedula_albicollis | 0 | 12.7 | 5.9 | 11815429.54 |
| birds | Ficedula_basilanica | Ficedula_basilanica | 1 | 13.5 | 2 | 116046.89 |
| birds | Ficedula_buruensis | Ficedula_buruensis | 0 | 14.5 | 2.4 | 26661.65 |
| birds | Ficedula_dumetoria | Ficedula_dumetoria | 0 | 9.4 | 2 | 377877.31 |
| birds | Ficedula_hodgsoni | Muscicapella_hodgsoni | 0 | 10 | 4.5 | 847137.81 |
| birds | Ficedula_hyperythra | Ficedula_hyperythra | 0 | 8.2 | 2.8 | 2932874.66 |
| birds | Ficedula_hypoleuca | Ficedula_hypoleuca | 0 | 13.8 | 5.7 | 21870324.64 |

| | | | | | | |
|---|---|---|---|---|---|---|
| birds | Ficedula_mugimaki | Ficedula_mugimaki | 0 | 11.7 | 5.7 | 7731077.74 |
| birds | Ficedula_narcissina | Ficedula_narcissina | 0 | 13.1 | 3.9 | 871204.77 |
| birds | Ficedula_parva | Ficedula_parva | 0 | 10.8 | 5.5 | 21638109.76 |
| birds | Ficedula_ruficauda | Muscicapa_ruficauda | 0 | 13.3 | 3.5 | 2570084.11 |
| birds | Ficedula_rufigula | Ficedula_rufigula | 0 | 11.6 | 2.4 | 114166.09 |
| birds | Ficedula_sapphira | Ficedula_sapphira | 0 | 7.8 | 4 | 1517380.4 |
| birds | Ficedula_semitorquata | Ficedula_semitorquata | 0 | 14.4 | 5.5 | 5511481.46 |
| birds | Ficedula_strophiata | Ficedula_strophiata | 0 | 12.8 | 3.5 | 2339628.4 |
| birds | Ficedula_subrubra | Ficedula_subrubra | 1 | 10.4 | 4.5 | 18887.27 |
| birds | Ficedula_superciliaris | Ficedula_superciliaris | 0 | 8 | 4 | 3996907.95 |
| birds | Ficedula_tricolor | Ficedula_tricolor | 0 | 8.3 | 3.5 | 2148183.47 |
| birds | Ficedula_westermanni | Ficedula_westermanni | 0 | 7.8 | 2 | 4080480.38 |
| birds | Ficedula_zanthopygia | Ficedula_zanthopygia | 0 | 12.1 | 5.7 | 3306839.92 |
| birds | Florisuga_fusca | Florisuga_fusca | 0 | 8 | 2 | 675214.04 |
| birds | Fluvicola_albiventer | Fluvicola_albiventer | 0 | 11.6 | 2.4 | 8121417.6 |
| birds | Fluvicola_pica | Fluvicola_pica | 0 | 12.3 | 2.4 | 1883895.03 |
| birds | Formicarius_analis | Formicarius_analis | 0 | 62.2 | 2 | 5715479.2 |
| birds | Formicarius_colma | Formicarius_colma | 0 | 47 | 2 | 6982041.19 |
| birds | Formicarius_moniliger | Formicarius_moniliger | 0 | 61.9 | 2 | 316042.05 |
| birds | Formicarius_nigricapillus | Formicarius_nigricapillus | 0 | 56.2 | 2 | 108463.37 |
| birds | Formicivora_erythronotos | Formicivora_erythronotos | 1 | 10.3 | 2 | 105.33 |
| birds | Formicivora_grisea | Formicivora_grisea | 0 | 10.4 | 2 | 5032873.87 |
| birds | Formicivora_rufa | Formicivora_rufa | 0 | 10.8 | 2 | 4090899.87 |
| birds | Formicivora_serrana | Formicivora_serrana | 0 | 11.4 | 2 | 209781.99 |
| birds | Forpus_coelestis | Forpus_coelestis | 0 | 26.1 | 4.9 | 147204.19 |
| birds | Forpus_conspicillatus | Forpus_conspicillatus | 0 | 26.4 | 2.4 | 321378.79 |
| birds | Forpus_cyanopygius | Forpus_cyanopygius | 0 | 33.5 | 3 | 115539.41 |
| birds | Forpus_passerinus | Forpus_passerinus | 0 | 23 | 5 | 1072147.57 |
| birds | Forpus_xanthops | Forpus_xanthops | 1 | 35 | 4.2 | 2888.58 |
| birds | Forpus_xanthopterygius | Forpus_xanthopterygius | 0 | 31 | 5.3 | 5781313.45 |
| birds | Francolinus_francolinus | Francolinus_francolinus | 0 | 452.1 | 9.2 | 3027937.66 |
| birds | Francolinus_pictus | Francolinus_pictus | 0 | 286.8 | 4.9 | 1554139.43 |
| birds | Francolinus_pintadeanus | Francolinus_pintadeanus | 0 | 310 | 4.5 | 2072227.73 |
| birds | Fraseria_caerulescens | Muscicapa_caerulescens | 0 | 17.6 | 2.4 | 7513617.5 |

| | | | | | | |
|---|---|---|---|---|---|---|
| birds | Fraseria_cinerascens | Fraseria_cinerascens | 0 | 23.1 | 1.4 | 3468124.59 |
| birds | Fraseria_griseigularis | Myioparus_griseigularis | 0 | 12.6 | 2 | 2490848.7 |
| birds | Fraseria_lendu | Muscicapa_lendu | 1 | 17.7 | 2 | 37404.93 |
| birds | Fraseria_ocreata | Fraseria_ocreata | 0 | 35.5 | 2.4 | 3285812.96 |
| birds | Fraseria_plumbea | Myioparus_plumbeus | 0 | 12.8 | 2 | 7861104.01 |
| birds | Frederickena_viridis | Frederickena_viridis | 0 | 76 | 2 | 1266903.39 |
| birds | Fringilla_coelebs | Fringilla_coelebs | 0 | 23.8 | 4.5 | 18764781.26 |
| birds | Fringilla_montifringilla | Fringilla_montifringilla | 0 | 23.2 | 5.9 | 24958109.43 |
| birds | Fringilla_teydea | Fringilla_teydea | 0 | 30.3 | 2 | 349.94 |
| birds | Fulica_alai | Fulica_alai | 0 | 459 | 7.7 | 3693.54 |
| birds | Fulica_americana | Fulica_americana | 0 | 636.7 | 9.5 | 13778114.98 |
| birds | Fulica_ardesiaca | Fulica_ardesiaca | 0 | 988 | 4.5 | 594594.35 |
| birds | Fulica_armillata | Fulica_armillata | 0 | 1040 | 4 | 2733323.64 |
| birds | Fulica_atra | Fulica_atra | 0 | 717.1 | 7.7 | 40041045.87 |
| birds | Fulica_cornuta | Fulica_cornuta | 0 | 2110.4 | 3.9 | 223464.94 |
| birds | Fulica_cristata | Fulica_cristata | 0 | 826 | 5.9 | 5291448.7 |
| birds | Fulica_gigantea | Fulica_gigantea | 0 | 2442.7 | 4.6 | 534521.72 |
| birds | Fulica_leucoptera | Fulica_leucoptera | 0 | 656 | 11 | 3041454.41 |
| birds | Fulica_rufifrons | Fulica_rufifrons | 0 | 644 | 4.2 | 1255982.21 |
| birds | Furnarius_cristatus | Furnarius_cristatus | 0 | 25.5 | 4 | 679538.36 |
| birds | Furnarius_figulus | Furnarius_figulus | 0 | 28 | 2 | 1971356.28 |
| birds | Furnarius_leucopus | Furnarius_leucopus | 0 | 54.8 | 2 | 4783071.89 |
| birds | Furnarius_minor | Furnarius_minor | 0 | 25.8 | 4 | 319149.09 |
| birds | Furnarius_rufus | Furnarius_rufus | 0 | 46.4 | 3.2 | 6181562.76 |
| birds | Furnarius_torridus | Furnarius_torridus | 0 | 52.3 | 4 | 87570.32 |
| birds | Gactornis_enarratus | Caprimulgus_enarratus | 0 | 52.8 | 2 | 226375.93 |
| birds | Galbula_albirostris | Galbula_albirostris | 0 | 22.1 | 2 | 1881365.04 |
| birds | Galbula_cyanicollis | Galbula_cyanicollis | 0 | 23.2 | 2 | 2705041.54 |
| birds | Galbula_leucogastra | Galbula_leucogastra | 0 | 15.9 | 1 | 2991415.41 |
| birds | Galbula_ruficauda | Galbula_ruficauda | 0 | 26.5 | 2.8 | 7607043.14 |
| birds | Galerida_cristata | Galerida_cristata | 0 | 42.7 | 3.9 | 24389170.69 |
| birds | Galerida_deva | Galerida_deva | 0 | 19.9 | 2.8 | 832847.98 |
| birds | Galerida_magnirostris | Galerida_magnirostris | 0 | 41.6 | 2.8 | 692806.52 |
| birds | Galerida_modesta | Galerida_modesta | 0 | 20.2 | 1 | 3249298.44 |

| | | | | | | |
|---|---|---|---|---|---|---|
| birds | Galerida_theklae | Galerida_theklae | 0 | 38.2 | 3.5 | 2270448.11 |
| birds | Gallicolumba_crinigera | Gallicolumba_crinigera | 1 | 192.2 | 1 | 94609.55 |
| birds | Gallicolumba_luzonica | Gallicolumba_luzonica | 0 | 196 | 2 | 107374.52 |
| birds | Gallicolumba_platenae | Gallicolumba_platenae | 1 | 196 | 2 | 475.23 |
| birds | Gallicolumba_rufigula | Gallicolumba_rufigula | 0 | 130 | 1 | 731477.39 |
| birds | Gallicrex_cinerea | Gallicrex_cinerea | 0 | 390.4 | 4.2 | 11190032.01 |
| birds | Gallinago_gallinago | Gallinago_gallinago | 0 | 112.9 | 4 | 45058410.31 |
| birds | Gallinago_hardwickii | Gallinago_hardwickii | 0 | 151.3 | 4 | 2812320.14 |
| birds | Gallinago_macrodactyla | Gallinago_macrodactyla | 1 | 216 | 3 | 220000.71 |
| birds | Gallinago_media | Gallinago_media | 0 | 170 | 4 | 39693519.26 |
| birds | Gallinago_megala | Gallinago_megala | 0 | 121 | 4 | 7562818.05 |
| birds | Gallinago_nigripennis | Gallinago_nigripennis | 0 | 112 | 2.4 | 5609482.86 |
| birds | Gallinago_paraguaiae | Gallinago_paraguaiae | 0 | 109.4 | 2.8 | 13818663 |
| birds | Gallinago_solitaria | Gallinago_solitaria | 0 | 140.1 | 4 | 7600182.79 |
| birds | Gallinago_stenura | Gallinago_stenura | 0 | 113 | 4 | 15061284.14 |
| birds | Gallinago_stricklandii | Gallinago_stricklandii | 0 | 252 | 2 | 340969.8 |
| birds | Gallinago_undulata | Gallinago_undulata | 0 | 312.4 | 2.8 | 2585357.36 |
| birds | Gallinula_chloropus | Gallinula_chloropus | 0 | 339.6 | 5.8 | 38971687.19 |
| birds | Gallinula_tenebrosa | Gallinula_tenebrosa | 0 | 530.1 | 9.5 | 3816495.38 |
| birds | Gallirallus_australis | Gallirallus_australis | 1 | 875.7 | 2.8 | 42528.43 |
| birds | Gallirex_johnstoni | Ruwenzorornis_johnstoni | 0 | 240 | 1.4 | 16706.27 |
| birds | Gallirex_porphyreolophus | Tauraco_porphyreolophus | 0 | 259 | 2.4 | 2166499.58 |
| birds | Galloperdix_bicalcarata | Galloperdix_bicalcarata | 0 | 290.9 | 2 | 37679.97 |
| birds | Galloperdix_lunulata | Galloperdix_lunulata | 0 | 254.4 | 3.5 | 1390384.27 |
| birds | Galloperdix_spadicea | Galloperdix_spadicea | 0 | 359.1 | 3.2 | 1175974.86 |
| birds | Gallus_gallus | Gallus_gallus | 0 | 751.7 | 6 | 5219583.04 |
| birds | Gallus_lafayettii | Gallus_lafayetii | 0 | 825 | 2 | 55601.24 |
| birds | Gallus_sonneratii | Gallus_sonneratii | 0 | 840.8 | 4.5 | 1030202.67 |
| birds | Gallus_varius | Gallus_varius | 0 | 839.3 | 3.5 | 187783.16 |
| birds | Gampsonyx_swainsonii | Gampsonyx_swainsonii | 0 | 92.9 | 2.4 | 10266134.72 |
| birds | Garrulax_canorus | Garrulax_canorus | 0 | 62.8 | 3.2 | 1868020.76 |
| birds | Garrulax_leucolophus | Garrulax_leucolophus | 0 | 123.4 | 4 | 1715209.64 |
| birds | Garrulax_merulinus | Garrulax_merulinus | 0 | 78.8 | 2.4 | 498651.88 |
| birds | Garrulax_monileger | Garrulax_monileger | 0 | 83.7 | 4.5 | 2496742.03 |

| | | | | | | |
|---|---|---|---|---|---|---|
| birds | Garrulus_glandarius | Garrulus_glandarius | 0 | 159.5 | 4.6 | 19458203.9 |
| birds | Garrulus_lanceolatus | Garrulus_lanceolatus | 0 | 97.7 | 3.5 | 322480.19 |
| birds | Gavia_adamsii | Gavia_adamsii | 0 | 4944.3 | 2 | 5436574.88 |
| birds | Gavia_arctica | Gavia_arctica | 0 | 2251.1 | 1.7 | 22818831.52 |
| birds | Gavia_immer | Gavia_immer | 0 | 4956.8 | 1.7 | 12626924.28 |
| birds | Gavia_stellata | Gavia_stellata | 0 | 1486 | 1.7 | 23359436.1 |
| birds | Gavicalis_fasciogularis | Lichenostomus_fasciogularis | 0 | 26.8 | 2 | 43256.67 |
| birds | Gavicalis_versicolor | Lichenostomus_versicolor | 0 | 36.7 | 2 | 161900.97 |
| birds | Gavicalis_virescens | Lichenostomus_virescens | 0 | 23.3 | 2 | 7306702.44 |
| birds | Geococcyx_californianus | Geococcyx_californianus | 0 | 376 | 3.5 | 3008009.53 |
| birds | Geococcyx_velox | Geococcyx_velox | 0 | 179.9 | 2.4 | 517104.33 |
| birds | Geocolaptes_olivaceus | Geocolaptes_olivaceus | 0 | 120 | 3 | 773995.64 |
| birds | Geokichla_citrina | Zoothera_citrina | 0 | 53.3 | 3.5 | 4482928.59 |
| birds | Geokichla_gurneyi | Zoothera_gurneyi | 0 | 57.9 | 2.4 | 421268.02 |
| birds | Geokichla_guttata | Zoothera_guttata | 1 | 57.5 | 2.4 | 84741.26 |
| birds | Geokichla_interpres | Zoothera_interpres | 1 | 81.9 | 2.4 | 670164 |
| birds | Geokichla_piaggiae | Zoothera_piaggiae | 0 | 51.1 | 2 | 328100.35 |
| birds | Geokichla_princei | Zoothera_princei | 0 | 65.4 | 1.7 | 453479.33 |
| birds | Geokichla_sibirica | Zoothera_sibirica | 0 | 75.5 | 4.5 | 10702412.67 |
| birds | Geokichla_spiloptera | Zoothera_spiloptera | 0 | 70 | 2.4 | 11408.73 |
| birds | Geokichla_wardii | Zoothera_wardii | 0 | 58.6 | 3.5 | 194220.03 |
| birds | Geopelia_cuneata | Geopelia_cuneata | 0 | 31.6 | 2 | 5703540.33 |
| birds | Geopelia_humeralis | Geopelia_humeralis | 0 | 128.4 | 2 | 2758465.78 |
| birds | Geopelia_striata | Geopelia_striata | 0 | 56.6 | 2 | 1478473.74 |
| birds | Geophaps_plumifera | Geophaps_plumifera | 0 | 94.3 | 2 | 1860384.76 |
| birds | Geophaps_scripta | Geophaps_scripta | 0 | 221.4 | 2 | 953627.9 |
| birds | Geophaps_smithii | Geophaps_smithii | 0 | 194 | 2 | 288800.13 |
| birds | Geositta_antarctica | Geositta_antarctica | 0 | 36.9 | 3 | 548387.44 |
| birds | Geositta_cunicularia | Geositta_cunicularia | 0 | 28.5 | 2.4 | 2912290.54 |
| birds | Geositta_isabellina | Geositta_isabellina | 0 | 42.7 | 3 | 71409.45 |
| birds | Geositta_peruviana | Geositta_peruviana | 0 | 20.8 | 2 | 54662.98 |
| birds | Geositta_poeciloptera | Geositta_poeciloptera | 1 | 18 | 3 | 1456963.68 |
| birds | Geositta_punensis | Geositta_punensis | 0 | 25.8 | 2.4 | 382860.93 |
| birds | Geositta_rufipennis | Geositta_rufipennis | 0 | 32.4 | 2.4 | 757788 |

| | | | | | | |
|---|---|---|---|---|---|---|
| birds | Geositta_tenuirostris | Geositta_tenuirostris | 0 | 32.2 | 2 | 622360.29 |
| birds | Geothlypis_aequinoctialis | Geothlypis_aequinoctialis | 0 | 13.1 | 2 | 7376725.6 |
| birds | Geothlypis_auricularis | Geothlypis_auricularis | 0 | 13.8 | 2 | 196093.06 |
| birds | Geothlypis_beldingi | Geothlypis_beldingi | 1 | 15.7 | 2.8 | 1747.19 |
| birds | Geothlypis_chiriquensis | Geothlypis_chiriquensis | 0 | 13.8 | 2 | 1382.25 |
| birds | Geothlypis_formosa | Oporornis_formosus | 0 | 14 | 4.5 | 3787752.22 |
| birds | Geothlypis_philadelphia | Oporornis_philadelphia | 0 | 11.7 | 3.9 | 7176055.48 |
| birds | Geothlypis_poliocephala | Geothlypis_poliocephala | 0 | 14.6 | 2.8 | 859573.04 |
| birds | Geothlypis_rostrata | Geothlypis_rostrata | 0 | 16.1 | 2 | 8764.16 |
| birds | Geothlypis_semiflava | Geothlypis_semiflava | 0 | 17.2 | 1.4 | 115622.87 |
| birds | Geothlypis_tolmiei | Oporornis_tolmiei | 0 | 10.4 | 4.2 | 5855497.18 |
| birds | Geothlypis_trichas | Geothlypis_trichas | 0 | 9.5 | 3.9 | 15147170.25 |
| birds | Geotrygon_caniceps | Geotrygon_caniceps | 1 | 192 | 1.4 | 107566.32 |
| birds | Geotrygon_chrysia | Geotrygon_chrysia | 0 | 169 | 2 | 207769.78 |
| birds | Geotrygon_montana | Geotrygon_montana | 0 | 133.9 | 2 | 13251114.74 |
| birds | Geotrygon_mystacea | Geotrygon_mystacea | 0 | 211 | 1.4 | 14198.65 |
| birds | Geotrygon_versicolor | Geotrygon_versicolor | 0 | 225 | 2 | 11021.79 |
| birds | Geotrygon_violacea | Geotrygon_violacea | 0 | 95.2 | 2 | 1490404.57 |
| birds | Geranoaetus_albicaudatus | Buteo_albicaudatus | 0 | 885 | 2 | 9466789.11 |
| birds | Geranoaetus_polyosoma | Buteo_polyosoma | 0 | 782.6 | 1.7 | 4720061.53 |
| birds | Geronticus_eremita | Geronticus_eremita | 1 | 1202 | 2.8 | 2452918.92 |
| birds | Gerygone_chloronota | Gerygone_chloronota | 0 | 6.6 | 2.4 | 1009439.84 |
| birds | Gerygone_chrysogaster | Gerygone_chrysogaster | 0 | 8 | 3 | 557753.52 |
| birds | Gerygone_dorsalis | Gerygone_dorsalis | 0 | 7.2 | 2.4 | 7004.78 |
| birds | Gerygone_fusca | Gerygone_fusca | 0 | 5.4 | 2.4 | 3748888.93 |
| birds | Gerygone_igata | Gerygone_igata | 0 | 6.4 | 3.2 | 268886.76 |
| birds | Gerygone_inornata | Gerygone_inornata | 0 | 7.2 | 2.4 | 33197.31 |
| birds | Gerygone_levigaster | Gerygone_levigaster | 0 | 6.2 | 2.4 | 342256.68 |
| birds | Gerygone_magnirostris | Gerygone_magnirostris | 0 | 6.8 | 2.4 | 1205051.15 |
| birds | Gerygone_mouki | Gerygone_mouki | 0 | 5.2 | 2.4 | 387637.87 |
| birds | Gerygone_olivacea | Gerygone_olivacea | 0 | 6.7 | 2.4 | 2358084.06 |
| birds | Gerygone_palpebrosa | Gerygone_palpebrosa | 0 | 7.4 | 2.4 | 969224.38 |
| birds | Gerygone_sulphurea | Gerygone_sulphurea | 0 | 5.6 | 2.4 | 1733224.34 |
| birds | Gerygone_tenebrosa | Gerygone_tenebrosa | 0 | 7.7 | 2.4 | 39187.75 |

| | | | | | | |
|---|---|---|---|---|---|---|
| birds | Glareola_cinerea | Glareola_cinerea | 0 | 37.4 | 1.4 | 3439738.6 |
| birds | Glareola_lactea | Glareola_lactea | 0 | 39.8 | 2.4 | 3822399.39 |
| birds | Glareola_maldivarum | Glareola_maldivarum | 0 | 75.2 | 2.4 | 12608507.89 |
| birds | Glareola_nordmanni | Glareola_nordmanni | 0 | 97.2 | 3.5 | 8161537.81 |
| birds | Glareola_nuchalis | Glareola_nuchalis | 0 | 62.7 | 2 | 6244433.49 |
| birds | Glareola_ocularis | Glareola_ocularis | 0 | 91.9 | 2 | 1261222.13 |
| birds | Glareola_pratincola | Glareola_pratincola | 0 | 84.9 | 3 | 13127791.54 |
| birds | Glaucestrilda_coerulescens | Estrilda_caerulescens | 0 | 9.4 | 3.2 | 1693485.93 |
| birds | Glaucestrilda_perreini | Estrilda_perreini | 0 | 7.4 | 6.3 | 1035934.15 |
| birds | Glaucestrilda_thomensis | Estrilda_thomensis | 0 | 7.6 | 3.5 | 210666.08 |
| birds | Glaucidium_brasilianum | Glaucidium_brasilianum | 0 | 63.3 | 3.2 | 16036298.78 |
| birds | Glaucidium_brodiei | Glaucidium_brodiei | 0 | 59.4 | 4.2 | 2339208.31 |
| birds | Glaucidium_capense | Glaucidium_capense | 0 | 122 | 2.4 | 4071921.71 |
| birds | Glaucidium_castaneum | Glaucidium_castaneum | 0 | 100 | 2.4 | 91747.23 |
| birds | Glaucidium_cuculoides | Glaucidium_cuculoides | 0 | 198.2 | 3.9 | 4612393.63 |
| birds | Glaucidium_gnoma | Glaucidium_gnoma | 0 | 61.3 | 2.8 | 3631571.53 |
| birds | Glaucidium_griseiceps | Glaucidium_griseiceps | 0 | 50.6 | 2.8 | 270382.47 |
| birds | Glaucidium_nana | Glaucidium_nanum | 0 | 73.7 | 3.9 | 656548.54 |
| birds | Glaucidium_passerinum | Glaucidium_passerinum | 0 | 57.9 | 5.3 | 11366077.56 |
| birds | Glaucidium_perlatum | Glaucidium_perlatum | 0 | 79.2 | 3 | 11923401.87 |
| birds | Glaucidium_radiatum | Glaucidium_radiatum | 0 | 101 | 2.8 | 2186922.27 |
| birds | Glaucidium_sjostedti | Glaucidium_sjostedti | 0 | 139 | 2 | 1369630.83 |
| birds | Glaucidium_tephronotum | Glaucidium_tephronotum | 0 | 87.4 | 2.8 | 2054685.26 |
| birds | Glaucis_aeneus | Glaucis_aeneus | 0 | 4.8 | 2 | 177367.01 |
| birds | Glaucis_dohrnii | Glaucis_dohrnii | 1 | 5.8 | 2 | 60089.03 |
| birds | Glaucis_hirsutus | Glaucis_hirsutus | 0 | 6.8 | 2 | 7820159.95 |
| birds | Gliciphila_melanops | Phylidonyris_melanops | 0 | 18.2 | 2 | 1135955.29 |
| birds | Glossopsitta_concinna | Glossopsitta_concinna | 0 | 70.5 | 2 | 663167.13 |
| birds | Glossopsitta_porphyrocephala | Glossopsitta_porphyrocephala | 0 | 44.7 | 3.5 | 944646.29 |
| birds | Glossopsitta_pusilla | Glossopsitta_pusilla | 0 | 39.4 | 3.9 | 776503.49 |
| birds | Glyphorynchus_spirurus | Glyphorynchus_spirurus | 0 | 14.6 | 1.7 | 7368285.07 |
| birds | Gorsachius_goisagi | Gorsachius_goisagi | 1 | 527 | 3.5 | 823887.54 |
| birds | Gorsachius_melanolophus | Gorsachius_melanolophus | 0 | 412.3 | 3.9 | 1619076.85 |
| birds | Gracula_ptilogenys | Gracula_ptilogenys | 0 | 192 | 2 | 15783.5 |

| | | | | | | |
|---|---|---|---|---|---|---|
| birds | Gracula_religiosa | Gracula_religiosa | 0 | 192 | 1.7 | 3864550.11 |
| birds | Gracupica_contra | Sturnus_contra | 0 | 84 | 4.5 | 3064197.42 |
| birds | Gracupica_nigricollis | Sturnus_nigricollis | 0 | 156.9 | 3.9 | 2900662.51 |
| birds | Grallaria_erythrotis | Grallaria_erythrotis | 0 | 57 | 2 | 39088.66 |
| birds | Grallaria_guatimalensis | Grallaria_guatimalensis | 0 | 94.2 | 2 | 1176261.46 |
| birds | Grallaria_haplonota | Grallaria_haplonota | 0 | 85 | 2 | 87992.13 |
| birds | Grallaria_ruficapilla | Grallaria_ruficapilla | 0 | 82.7 | 2 | 239637.78 |
| birds | Grallaria_rufula | Grallaria_rufula | 0 | 35.7 | 1 | 236386.27 |
| birds | Grallaria_varia | Grallaria_varia | 0 | 119 | 2 | 3247752.05 |
| birds | Grallaricula_ferrugineipectus | Grallaricula_ferrugineipectus | 0 | 17.2 | 2 | 70002.38 |
| birds | Grallaricula_flavirostris | Grallaricula_flavirostris | 0 | 17.7 | 1.4 | 271741.93 |
| birds | Graminicola_bengalensis | Graminicola_bengalensis | 0 | 14.6 | 4 | 174532.67 |
| birds | Grammatoptila_striata | Garrulax_striatus | 0 | 92 | 2 | 317905.88 |
| birds | Granatina_granatina | Uraeginthus_granatinus | 0 | 11.8 | 3.2 | 2368190.34 |
| birds | Grandala_coelicolor | Grandala_coelicolor | 0 | 45.6 | 2 | 1886746.7 |
| birds | Grantiella_picta | Grantiella_picta | 0 | 21.5 | 2 | 1455898.86 |
| birds | Griseotyrannus_aurantioatrocristatus | Empidonomus_aurantioatrocristatus | 0 | 33 | 2.8 | 9464416.15 |
| birds | Grus_americana | Grus_americana | 1 | 5826 | 2 | 1543239.24 |
| birds | Grus_antigone | Grus_antigone | 1 | 5916.2 | 2.4 | 1882141.61 |
| birds | Grus_canadensis | Grus_canadensis | 0 | 4296.2 | 2 | 14715653.06 |
| birds | Grus_grus | Grus_grus | 0 | 5500 | 2 | 20795140 |
| birds | Grus_japonensis | Grus_japonensis | 1 | 8786 | 2 | 1639871.12 |
| birds | Grus_monacha | Grus_monacha | 1 | 3729.9 | 2 | 1728477.41 |
| birds | Grus_nigricollis | Grus_nigricollis | 0 | 6000 | 2 | 886789.65 |
| birds | Grus_rubicunda | Grus_rubicunda | 0 | 6222.8 | 2 | 3559865.45 |
| birds | Grus_vipio | Grus_vipio | 1 | 4663 | 2 | 2152136.88 |
| birds | Guaruba_guarouba | Guaruba_guarouba | 1 | 194 | 2.4 | 932156.38 |
| birds | Guttera_plumifera | Guttera_plumifera | 0 | 866 | 9.5 | 834485.49 |
| birds | Guttera_pucherani | Guttera_pucherani | 0 | 1149 | 4.5 | 519919.07 |
| birds | Gymnobucco_bonapartei | Gymnobucco_bonapartei | 0 | 63 | 3.9 | 1792433.71 |
| birds | Gymnobucco_calvus | Gymnobucco_calvus | 0 | 54.5 | 3 | 1685321.51 |
| birds | Gymnocichla_nudiceps | Gymnocichla_nudiceps | 0 | 30.5 | 2 | 355899.13 |
| birds | Gymnoderus_foetidus | Gymnoderus_foetidus | 0 | 275.5 | 1 | 5252269.51 |
| birds | Gymnogyps_californianus | Gymnogyps_californianus | 1 | 8442.7 | 1 | 388190.92 |

| | | | | | | |
|---|---|---|---|---|---|---|
| birds | Gymnophaps_albertisii | Gymnophaps_albertisii | 0 | 259 | 1 | 535761.11 |
| birds | Gymnopithys_leucaspis | Gymnopithys_leucaspis | 0 | 27.2 | 2 | 1029456.75 |
| birds | Gymnopithys_rufigula | Gymnopithys_rufigula | 0 | 29.1 | 2 | 1727736.89 |
| birds | Gymnorhinus_cyanocephalus | Gymnorhinus_cyanocephalus | 1 | 104.8 | 3.5 | 1284048.89 |
| birds | Gymnoris_dentata | Petronia_dentata | 0 | 18.4 | 3 | 4184728.08 |
| birds | Gymnoris_superciliaris | Petronia_superciliaris | 0 | 24 | 3.2 | 3946522.23 |
| birds | Gymnoris_xanthocollis | Petronia_xanthocollis | 0 | 18.1 | 3.5 | 4502531.2 |
| birds | Gypaetus_barbatus | Gypaetus_barbatus | 0 | 5695 | 1.4 | 10893249.16 |
| birds | Gypohierax_angolensis | Gypohierax_angolensis | 0 | 1600 | 1 | 11874949.01 |
| birds | Gyps_africanus | Gyps_africanus | 1 | 5433 | 1 | 15114222.69 |
| birds | Gyps_bengalensis | Gyps_bengalensis | 1 | 4385 | 1 | 5834018.97 |
| birds | Gyps_coprotheres | Gyps_coprotheres | 1 | 8177 | 1 | 3702609.83 |
| birds | Gyps_fulvus | Gyps_fulvus | 0 | 7436 | 1 | 18194036.07 |
| birds | Gyps_himalayensis | Gyps_himalayensis | 0 | 9798 | 1 | 6434312.57 |
| birds | Gyps_indicus | Gyps_indicus | 1 | 5515 | 1 | 2380044.79 |
| birds | Gyps_rueppelli | Gyps_rueppellii | 1 | 7400 | 1 | 10082232.28 |
| birds | Habia_carmioli | Chlorothraupis_carmioli | 0 | 37.6 | 2 | 93291.5 |
| birds | Habia_frenata | Chlorothraupis_frenata | 0 | 37.8 | 2 | 108990.05 |
| birds | Habia_stolzmanni | Chlorothraupis_stolzmanni | 0 | 38.8 | 1.4 | 50027.82 |
| birds | Haematopus_ostralegus | Haematopus_ostralegus | 0 | 526 | 3.2 | 15759416.9 |
| birds | Haemorhous_cassinii | Carpodacus_cassinii | 0 | 26.5 | 4.5 | 2814033.13 |
| birds | Haemorhous_mexicanus | Carpodacus_mexicanus | 0 | 21.4 | 4.5 | 10299459.91 |
| birds | Haemorhous_purpureus | Carpodacus_purpureus | 0 | 23.3 | 4.5 | 7952995.84 |
| birds | Hafferia_fortis | Myrmeciza_fortis | 0 | 45.9 | 2 | 1945728.92 |
| birds | Halcyon_albiventris | Halcyon_albiventris | 0 | 65.1 | 3.2 | 3765344.98 |
| birds | Halcyon_badia | Halcyon_badia | 0 | 57.9 | 2.4 | 3197177.13 |
| birds | Halcyon_chelicuti | Halcyon_chelicuti | 0 | 43.2 | 3.9 | 13120453.97 |
| birds | Halcyon_coromanda | Halcyon_coromanda | 0 | 77.5 | 4.9 | 4234824.05 |
| birds | Halcyon_cyanoventris | Halcyon_cyanoventris | 0 | 93 | 3.9 | 131597.72 |
| birds | Halcyon_leucocephala | Halcyon_leucocephala | 0 | 41.8 | 3.5 | 14797715.21 |
| birds | Halcyon_malimbica | Halcyon_malimbica | 0 | 91.8 | 2.8 | 4774587.94 |
| birds | Halcyon_pileata | Halcyon_pileata | 1 | 84 | 4.5 | 7076437.93 |
| birds | Halcyon_senegalensis | Halcyon_senegalensis | 0 | 73.4 | 2.8 | 14752529.26 |
| birds | Halcyon_senegaloides | Halcyon_senegaloides | 0 | 61.8 | 3 | 193724.21 |

| | | | | | | |
|---|---|---|---|---|---|---|
| birds | Halcyon_smyrnensis | Halcyon_smyrnensis | 0 | 91.4 | 5.3 | 6823580.45 |
| birds | Haliaeetus_albicilla | Haliaeetus_albicilla | 0 | 4729.3 | 1.7 | 23280177.99 |
| birds | Haliaeetus_leucocephalus | Haliaeetus_leucocephalus | 0 | 4700.6 | 2 | 15549096.96 |
| birds | Haliaeetus_leucogaster | Haliaeetus_leucogaster | 0 | 2827 | 2 | 6333774.53 |
| birds | Haliaeetus_leucoryphus | Haliaeetus_leucoryphus | 1 | 2885.9 | 2.4 | 14108255.23 |
| birds | Haliaeetus_pelagicus | Haliaeetus_pelagicus | 1 | 7757 | 1.7 | 1505403.98 |
| birds | Haliaeetus_vocifer | Haliaeetus_vocifer | 0 | 3400 | 2 | 18331524.51 |
| birds | Haliaeetus_vociferoides | Haliaeetus_vociferoides | 1 | 3000 | 2 | 52761.99 |
| birds | Haliastur_indus | Haliastur_indus | 0 | 529 | 1.4 | 8840397.21 |
| birds | Haliastur_sphenurus | Haliastur_sphenurus | 0 | 753 | 2 | 5585453.97 |
| birds | Hamirostra_melanosternon | Hamirostra_melanosternon | 0 | 1196 | 2 | 5297941.74 |
| birds | Hapalopsittaca_pyrrhops | Hapalopsittaca_pyrrhops | 1 | 103 | 2 | 3517.66 |
| birds | Hapaloptila_castanea | Hapaloptila_castanea | 0 | 80.6 | 2 | 31778.47 |
| birds | Haplophaedia_aureliae | Haplophaedia_aureliae | 0 | 5.2 | 2 | 222015.49 |
| birds | Haplophaedia_lugens | Haplophaedia_lugens | 0 | 6.9 | 2 | 6460.08 |
| birds | Harpactes_erythrocephalus | Harpactes_erythrocephalus | 0 | 80.3 | 2.8 | 2608367.68 |
| birds | Harpactes_fasciatus | Harpactes_fasciatus | 0 | 62 | 2.8 | 406882.69 |
| birds | Harpactes_oreskios | Harpactes_oreskios | 0 | 57.3 | 2.4 | 1639985.91 |
| birds | Harpagus_bidentatus | Harpagus_bidentatus | 0 | 215.3 | 2 | 8976331.27 |
| birds | Harpagus_diodon | Harpagus_diodon | 0 | 200 | 2 | 8031409.73 |
| birds | Hartertula_flavoviridis | Neomixis_flavoviridis | 0 | 9.7 | 2 | 66817.98 |
| birds | Hedydipna_collaris | Anthreptes_collaris | 0 | 7 | 2 | 9197435.7 |
| birds | Hedydipna_metallica | Anthreptes_metallicus | 0 | 7.2 | 1.7 | 1304366.52 |
| birds | Hedydipna_pallidigaster | Anthreptes_pallidigaster | 1 | 6.8 | 3 | 753.18 |
| birds | Hedydipna_platura | Anthreptes_platurus | 0 | 6.7 | 1.4 | 4094486.35 |
| birds | Heliactin_bilophus | Heliactin_bilophus | 0 | 4.4 | 2 | 3879596.58 |
| birds | Heliangelus_amethysticollis | Heliangelus_amethysticollis | 0 | 5 | 2 | 146657.78 |
| birds | Heliangelus_exortis | Heliangelus_exortis | 0 | 4.6 | 2 | 64064.87 |
| birds | Heliangelus_mavors | Heliangelus_mavors | 0 | 4.2 | 2 | 38610.25 |
| birds | Heliangelus_regalis | Heliangelus_regalis | 0 | 3.6 | 2 | 5218.9 |
| birds | Heliangelus_strophianus | Heliangelus_strophianus | 0 | 5.5 | 2 | 10032.03 |
| birds | Heliangelus_viola | Heliangelus_viola | 0 | 5.4 | 2 | 35976.8 |
| birds | Helicolestes_hamatus | Helicolestes_hamatus | 0 | 425.5 | 2 | 2374863.31 |
| birds | Heliodoxa_aurescens | Heliodoxa_aurescens | 0 | 6.4 | 2 | 3600288.83 |

| | | | | | | |
|---|---|---|---|---|---|---|
| birds | Heliodoxa_leadbeateri | Heliodoxa_leadbeateri | 0 | 7.4 | 2 | 409940.21 |
| birds | Heliodoxa_rubinoides | Heliodoxa_rubinoides | 0 | 7.8 | 2 | 203573.74 |
| birds | Heliodoxa_schreibersii | Heliodoxa_schreibersii | 0 | 9.2 | 2 | 713038.86 |
| birds | Heliodoxa_xanthogonys | Heliodoxa_xanthogonys | 0 | 7 | 2 | 222443.1 |
| birds | Heliomaster_furcifer | Heliomaster_furcifer | 0 | 5.4 | 2 | 2967951.96 |
| birds | Heliomaster_longirostris | Heliomaster_longirostris | 0 | 6.6 | 2 | 8290213.63 |
| birds | Heliomaster_squamosus | Heliomaster_squamosus | 0 | 5.9 | 2 | 1519261.77 |
| birds | Heliopais_personatus | Heliopais_personatus | 1 | 510 | 5.9 | 766693.11 |
| birds | Heliornis_fulica | Heliornis_fulica | 0 | 132 | 2.4 | 8812012.44 |
| birds | Heliothryx_auritus | Heliothryx_auritus | 0 | 5.4 | 2 | 6893450.96 |
| birds | Heliothryx_barroti | Heliothryx_barroti | 0 | 5.5 | 2 | 491040.04 |
| birds | Helmitheros_vermivorum | Helmitheros_vermivorum | 0 | 14.2 | 4.5 | 3326592.3 |
| birds | Helopsaltes_certhiola | Locustella_certhiola | 0 | 14.4 | 4.9 | 18568127.92 |
| birds | Helopsaltes_ochotensis | Locustella_ochotensis | 0 | 18.5 | 5.5 | 1896276.86 |
| birds | Helopsaltes_pleskei | Locustella_pleskei | 1 | 20 | 3.9 | 5556.54 |
| birds | Helopsaltes_pryeri | Locustella_pryeri | 0 | 14.2 | 5.5 | 227242.53 |
| birds | Hemicircus_canente | Hemicircus_canente | 0 | 43 | 2.4 | 665192.85 |
| birds | Hemimacronyx_chloris | Anthus_chloris | 1 | 25.1 | 3 | 82125.98 |
| birds | Hemiprocne_mystacea | Hemiprocne_mystacea | 0 | 70.1 | 1 | 805813.24 |
| birds | Hemipus_hirundinaceus | Hemipus_hirundinaceus | 0 | 10.2 | 2 | 1391676.19 |
| birds | Hemipus_picatus | Hemipus_picatus | 0 | 9 | 2.4 | 4982703.19 |
| birds | Hemitesia_pallidipes | Cettia_pallidipes | 0 | 10 | 3.5 | 1229162.3 |
| birds | Hemitriccus_granadensis | Hemitriccus_granadensis | 0 | 7.9 | 2 | 276758.68 |
| birds | Hemitriccus_margaritaceiventer | Hemitriccus_margaritaceiventer | 0 | 8.4 | 1.7 | 5809388.94 |
| birds | Hemitriccus_nidipendulus | Hemitriccus_nidipendulus | 0 | 7.5 | 2 | 542148.01 |
| birds | Hemitriccus_obsoletus | Hemitriccus_obsoletus | 0 | 11.6 | 2 | 262970.93 |
| birds | Hemitriccus_rufigularis | Hemitriccus_rufigularis | 0 | 9.4 | 2 | 76788.81 |
| birds | Hemitriccus_striaticollis | Hemitriccus_striaticollis | 0 | 8.6 | 2 | 4016133.45 |
| birds | Hemixos_castanonotus | Hemixos_castanonotus | 0 | 35.3 | 3 | 979934.26 |
| birds | Hemixos_flavala | Hemixos_flavala | 0 | 32.5 | 2.8 | 1098549.78 |
| birds | Henicorhina_leucophrys | Henicorhina_leucophrys | 0 | 16.2 | 2 | 722586.81 |
| birds | Henicorhina_leucosticta | Henicorhina_leucosticta | 0 | 14.2 | 2 | 2745877.94 |
| birds | Hesperiphona_vespertina | Coccothraustes_vespertinus | 1 | 57.3 | 3.5 | 9649540.8 |
| birds | Heterocercus_aurantiivertex | Heterocercus_aurantiivertex | 0 | 21.5 | 2 | 233954.07 |

| | | | | | | |
|---|---|---|---|---|---|---|
| birds | Heteromirafra_archeri | Heteromirafra_archeri | 1 | 30 | 3 | 1009.96 |
| birds | Heteromirafra_ruddi | Heteromirafra_ruddi | 1 | 26.6 | 2.4 | 21832.33 |
| birds | Heteromunia_pectoralis | Heteromunia_pectoralis | 0 | 15.2 | 4.9 | 1302318.41 |
| birds | Heterotetrax_humilis | Eupodotis_humilis | 0 | 698 | 2 | 474259.61 |
| birds | Heterotetrax_rueppelii | Eupodotis_rueppellii | 0 | 1107 | 1 | 280978.34 |
| birds | Heterotetrax_vigorsii | Eupodotis_vigorsii | 0 | 1700 | 1 | 691336.88 |
| birds | Hieraaetus_ayresii | Hieraaetus_ayresii | 0 | 806.1 | 1 | 7512090.85 |
| birds | Hieraaetus_morphnoides | Hieraaetus_morphnoides | 0 | 775.1 | 2 | 7265023.23 |
| birds | Hieraaetus_pennatus | Hieraaetus_pennatus | 0 | 823.8 | 2 | 29211510.18 |
| birds | Hieraaetus_wahlbergi | Aquila_wahlbergi | 0 | 640 | 1 | 12004193.21 |
| birds | Himalayapsitta_cyanocephala | Psittacula_cyanocephala | 0 | 66 | 4.5 | 2783564.24 |
| birds | Himalayapsitta_finschii | Psittacula_finschii | 0 | 89.6 | 4 | 1639893 |
| birds | Himalayapsitta_himalayana | Psittacula_himalayana | 0 | 108.5 | 3.9 | 149148.55 |
| birds | Himantornis_haematopus | Himantornis_haematopus | 0 | 585.7 | 3 | 2345297.5 |
| birds | Hippolais_icterina | Hippolais_icterina | 0 | 13.2 | 4.5 | 29586051.33 |
| birds | Hippolais_languida | Hippolais_languida | 0 | 12 | 4.5 | 10092888.18 |
| birds | Hippolais_olivetorum | Hippolais_olivetorum | 0 | 17.5 | 3.5 | 13353897.56 |
| birds | Hippolais_polyglotta | Hippolais_polyglotta | 0 | 11 | 3.9 | 10548115.16 |
| birds | Hirundapus_caudacutus | Hirundapus_caudacutus | 0 | 95.5 | 3.7 | 16971508.02 |
| birds | Hirundapus_giganteus | Hirundapus_giganteus | 0 | 127 | 3.9 | 2715789.43 |
| birds | Hirundinea_ferruginea | Hirundinea_ferruginea | 0 | 30.6 | 2 | 916120.42 |
| birds | Hirundo_aethiopica | Hirundo_aethiopica | 0 | 13.3 | 2.8 | 4764528.41 |
| birds | Hirundo_albigularis | Hirundo_albigularis | 0 | 21.8 | 3 | 3893507.61 |
| birds | Hirundo_angolensis | Hirundo_angolensis | 0 | 17.8 | 2.4 | 1666808.13 |
| birds | Hirundo_atrocaerulea | Hirundo_atrocaerulea | 1 | 13.8 | 2.4 | 1643896.13 |
| birds | Hirundo_dimidiata | Hirundo_dimidiata | 0 | 12.1 | 2.4 | 3681173.15 |
| birds | Hirundo_javanica | Hirundo_domicola | 0 | 17.8 | 3.5 | 3415869.81 |
| birds | Hirundo_leucosoma | Hirundo_leucosoma | 0 | 17.6 | 4 | 1390558.12 |
| birds | Hirundo_lucida | Hirundo_lucida | 0 | 13 | 3.5 | 2258983.81 |
| birds | Hirundo_neoxena | Hirundo_neoxena | 0 | 14.7 | 3.5 | 6029453.14 |
| birds | Hirundo_nigrita | Hirundo_nigrita | 0 | 18.3 | 2.4 | 4040980.58 |
| birds | Hirundo_nigrorufa | Hirundo_nigrorufa | 0 | 14.2 | 3 | 581101.75 |
| birds | Hirundo_rustica | Hirundo_rustica | 0 | 17.9 | 4.5 | 94643183.39 |
| birds | Hirundo_smithii | Hirundo_smithii | 0 | 12.4 | 3.2 | 15137552.43 |

| | | | | | | |
|---|---|---|---|---|---|---|
| birds | Hirundo_tahitica | Hirundo_tahitica | 0 | 17.8 | 2 | 1208.55 |
| birds | Histrionicus_histrionicus | Histrionicus_histrionicus | 0 | 561.6 | 5.9 | 11164771.89 |
| birds | Histurgops_ruficauda | Histurgops_ruficaudus | 0 | 69 | 3 | 98235.59 |
| birds | Hoploxypterus_cayanus | Vanellus_cayanus | 0 | 75.8 | 2 | 11467799.98 |
| birds | Horizocerus_hartlaubi | Tockus_hartlaubi | 0 | 98.5 | 4 | 1163200.91 |
| birds | Horornis_acanthizoides | Cettia_acanthizoides | 0 | 5.3 | 2.4 | 311800.42 |
| birds | Horornis_diphone | Cettia_diphone | 0 | 13.3 | 4.2 | 430218.55 |
| birds | Horornis_flavolivaceus | Cettia_flavolivacea | 0 | 8 | 3.5 | 1896851.77 |
| birds | Horornis_fortipes | Cettia_fortipes | 0 | 9.6 | 3.9 | 2834443.67 |
| birds | Horornis_ruficapilla | Cettia_ruficapilla | 0 | 12.7 | 2 | 16891.28 |
| birds | Houbaropsis_bengalensis | Houbaropsis_bengalensis | 1 | 2012.5 | 1.4 | 260519.85 |
| birds | Hydrocoloeus_minutus | Larus_minutus | 0 | 118 | 2.4 | 10822470.83 |
| birds | Hydrophasianus_chirurgus | Hydrophasianus_chirurgus | 0 | 160 | 4 | 8823045.22 |
| birds | Hydropsalis_cayennensis | Caprimulgus_cayennensis | 0 | 35.5 | 1.4 | 1530293.86 |
| birds | Hydropsalis_climacocerca | Hydropsalis_climacocerca | 0 | 46 | 2 | 5856089.27 |
| birds | Hydropsalis_maculicaudus | Caprimulgus_maculicaudus | 0 | 30.3 | 2 | 3375852.78 |
| birds | Hydropsalis_torquata | Hydropsalis_torquata | 0 | 58.5 | 2 | 7882112.56 |
| birds | Hydrornis_baudii | Pitta_baudii | 1 | 66 | 2 | 604986.6 |
| birds | Hydrornis_caeruleus | Pitta_caerulea | 0 | 202 | 2.4 | 779802.96 |
| birds | Hydrornis_cyaneus | Pitta_cyanea | 0 | 109 | 5.3 | 1019585.76 |
| birds | Hydrornis_elliotii | Pitta_elliotii | 0 | 91 | 2.8 | 605196.04 |
| birds | Hydrornis_guajanus | Pitta_guajana | 0 | 81.5 | 3.2 | 129669.16 |
| birds | Hydrornis_gurneyi | Pitta_gurneyi | 1 | 60.5 | 3.9 | 1891.85 |
| birds | Hydrornis_nipalensis | Pitta_nipalensis | 0 | 124 | 3.9 | 668283.53 |
| birds | Hydrornis_oatesi | Pitta_oatesi | 0 | 120 | 3.5 | 547817.02 |
| birds | Hydrornis_phayrei | Pitta_phayrei | 0 | 100 | 4 | 927700.65 |
| birds | Hydrornis_soror | Pitta_soror | 0 | 114 | 3 | 579014.96 |
| birds | Hylacola_cauta | Hylacola_cauta | 0 | 15.2 | 2.4 | 971888.94 |
| birds | Hylacola_pyrrhopygia | Hylacola_pyrrhopygia | 0 | 16.8 | 2.4 | 767767.19 |
| birds | Hylatomus_lineatus | Dryocopus_lineatus | 0 | 183.2 | 2.4 | 13770543.52 |
| birds | Hylatomus_pileatus | Dryocopus_pileatus | 0 | 286.6 | 2.8 | 5925872.32 |
| birds | Hylatomus_schulzii | Dryocopus_schulzi | 0 | 200 | 2 | 388434.65 |
| birds | Hylia_prasina | Hylia_prasina | 0 | 13.1 | 1 | 4144582.43 |
| birds | Hyliota_australis | Hyliota_australis | 0 | 11.2 | 3 | 1342986.67 |

| | | | | | | |
|---|---|---|---|---|---|---|
| birds | Hyliota_flavigaster | Hyliota_flavigaster | 0 | 12.5 | 2 | 4822595.38 |
| birds | Hylocharis_chrysura | Hylocharis_chrysura | 0 | 4.5 | 2 | 3128401.38 |
| birds | Hylocharis_sapphirina | Hylocharis_sapphirina | 0 | 4.4 | 2 | 5411214.46 |
| birds | Hylocichla_mustelina | Hylocichla_mustelina | 0 | 50.1 | 3.5 | 5235140.13 |
| birds | Hylopezus_perspicillatus | Hylopezus_perspicillatus | 0 | 43 | 2 | 279909.57 |
| birds | Hylophylax_naevioides | Hylophylax_naevioides | 0 | 17.8 | 2 | 372510.89 |
| birds | Hylophylax_naevius | Hylophylax_naevius | 0 | 14.2 | 1.7 | 5848723.89 |
| birds | Hylophylax_punctulatus | Hylophylax_punctulatus | 0 | 12.4 | 2 | 3128205.43 |
| birds | Hylopsar_cupreocauda | Lamprotornis_cupreocauda | 0 | 58 | 3 | 458895.76 |
| birds | Hylopsar_purpureiceps | Lamprotornis_purpureiceps | 0 | 62.8 | 3 | 1844762.06 |
| birds | Hylorchilus_sumichrasti | Hylorchilus_sumichrasti | 0 | 28.4 | 3 | 6551.84 |
| birds | Hymenops_perspicillatus | Hymenops_perspicillatus | 0 | 22.9 | 2.8 | 3807233.09 |
| birds | Hypargos_margaritatus | Hypargos_margaritatus | 0 | 12.8 | 3.5 | 155741.3 |
| birds | Hypargos_niveoguttatus | Hypargos_niveoguttatus | 0 | 14.6 | 4.2 | 1748989.03 |
| birds | Hypergerus_atriceps | Hypergerus_atriceps | 0 | 29.8 | 2.4 | 2301821.94 |
| birds | Hypnelus_ruficollis | Hypnelus_ruficollis | 0 | 49.8 | 3 | 239856.99 |
| birds | Hypocnemis_cantator | Hypocnemis_cantator | 0 | 12.3 | 2 | 968489.41 |
| birds | Hypocnemoides_melanopogon | Hypocnemoides_melanopogon | 0 | 14.1 | 2 | 3494411.77 |
| birds | Hypocolius_ampelinus | Hypocolius_ampelinus | 0 | 53.3 | 3.9 | 2239945.43 |
| birds | Hypopyrrhus_pyrohypogaster | Hypopyrrhus_pyrohypogaster | 1 | 101 | 2 | 31843.13 |
| birds | Hypotaenidia_philippensis | Gallirallus_philippensis | 0 | 205.9 | 5.7 | 2532556.07 |
| birds | Hypotaenidia_rovianae | Gallirallus_rovianae | 0 | 353.1 | 2.4 | 3354.63 |
| birds | Hypotaenidia_torquata | Gallirallus_torquatus | 0 | 280 | 3.5 | 412241.22 |
| birds | Hypothymis_azurea | Hypothymis_azurea | 0 | 11.1 | 2.8 | 7041185.04 |
| birds | Hypsipetes_amaurotis | Ixos_amaurotis | 0 | 70.9 | 3.9 | 715067.91 |
| birds | Hypsipetes_leucocephalus | Hypsipetes_leucocephalus | 0 | 51.8 | 2.8 | 4955880.13 |
| birds | Hypsipetes_madagascariensis | Hypsipetes_madagascariensis | 0 | 46 | 3 | 591386.99 |
| birds | Hypsipetes_philippinus | Ixos_philippinus | 0 | 38.6 | 2.4 | 232032.85 |
| birds | Hypsipetes_thompsoni | Hypsipetes_thompsoni | 0 | 41.8 | 2.4 | 196437.35 |
| birds | Ianthocincla_ocellata | Garrulax_ocellatus | 0 | 114 | 2 | 1005117.2 |
| birds | Ianthocincla_rufogularis | Garrulax_rufogularis | 0 | 62.8 | 3 | 260288.24 |
| birds | Ibidorhyncha_struthersii | Ibidorhyncha_struthersii | 0 | 294 | 4 | 5219019.96 |
| birds | Icteria_virens | Icteria_virens | 0 | 24.9 | 3.9 | 8893581.55 |
| birds | Icterus_bullockiorum | Icterus_bullockii | 0 | 37.9 | 4.5 | 5157597.51 |

| | | | | | | |
|---|---|---|---|---|---|---|
| birds | Icterus_cayanensis | Icterus_cayanensis | 0 | 35.4 | 2.4 | 4104473.13 |
| birds | Icterus_chrysater | Icterus_chrysater | 0 | 53.4 | 2.4 | 777224.61 |
| birds | Icterus_cucullatus | Icterus_cucullatus | 0 | 24.3 | 3.9 | 2334013.11 |
| birds | Icterus_dominicensis | Icterus_dominicensis | 0 | 35.7 | 3 | 75136.96 |
| birds | Icterus_galbula | Icterus_galbula | 0 | 32.8 | 4.5 | 7852695.64 |
| birds | Icterus_graceannae | Icterus_graceannae | 0 | 41.5 | 2 | 77582.36 |
| birds | Icterus_graduacauda | Icterus_graduacauda | 0 | 42.2 | 3.9 | 347160.28 |
| birds | Icterus_gularis | Icterus_gularis | 0 | 55.2 | 3.5 | 559678.92 |
| birds | Icterus_icterus | Icterus_icterus | 0 | 60.9 | 3 | 442639.32 |
| birds | Icterus_leucopteryx | Icterus_leucopteryx | 0 | 36.9 | 3.9 | 11260.28 |
| birds | Icterus_mesomelas | Icterus_mesomelas | 0 | 39.9 | 2.4 | 995703.98 |
| birds | Icterus_nigrogularis | Icterus_nigrogularis | 0 | 40.2 | 3 | 1368029.34 |
| birds | Icterus_parisorum | Icterus_parisorum | 0 | 36.5 | 2.8 | 2080143.58 |
| birds | Icterus_pectoralis | Icterus_pectoralis | 0 | 46.5 | 3 | 206586.99 |
| birds | Icterus_pustulatus | Icterus_pustulatus | 0 | 36.8 | 3.5 | 553521.84 |
| birds | Icterus_spurius | Icterus_spurius | 0 | 19.4 | 4 | 6755981.46 |
| birds | Icterus_wagleri | Icterus_wagleri | 0 | 41.8 | 3 | 767331.59 |
| birds | Ictinia_mississippiensis | Ictinia_mississippiensis | 0 | 276 | 2 | 4891699.34 |
| birds | Ictinia_plumbea | Ictinia_plumbea | 0 | 249.9 | 1.4 | 12792063.81 |
| birds | Iduna_caligata | Hippolais_caligata | 0 | 8.9 | 4.9 | 10100495.93 |
| birds | Iduna_natalensis | Chloropeta_natalensis | 0 | 11.7 | 2 | 3742433.69 |
| birds | Iduna_pallida | Hippolais_pallida | 0 | 9 | 3 | 19827150.12 |
| birds | Iduna_rama | Hippolais_rama | 0 | 9.3 | 4.9 | 5878685.67 |
| birds | Iduna_similis | Chloropeta_similis | 0 | 11.1 | 2 | 283650.62 |
| birds | Illadopsis_albipectus | Illadopsis_albipectus | 0 | 30.6 | 2 | 1215586.75 |
| birds | Illadopsis_cleaveri | Illadopsis_cleaveri | 0 | 30.8 | 2 | 977280.76 |
| birds | Illadopsis_fulvescens | Illadopsis_fulvescens | 0 | 27.9 | 2 | 2855381.87 |
| birds | Illadopsis_puveli | Illadopsis_puveli | 0 | 48.4 | 2 | 631343.92 |
| birds | Illadopsis_pyrrhoptera | Illadopsis_pyrrhoptera | 0 | 24.8 | 2 | 145566.3 |
| birds | Illadopsis_rufipennis | Illadopsis_rufipennis | 0 | 22.2 | 2 | 2029123.6 |
| birds | Indicator_indicator | Indicator_indicator | 0 | 50.4 | 4 | 11206655.87 |
| birds | Indicator_minor | Indicator_minor | 0 | 28.2 | 2.8 | 13121952.74 |
| birds | Indicator_variegatus | Indicator_variegatus | 0 | 48.8 | 2 | 3473509.99 |
| birds | Iodopleura_pipra | Iodopleura_pipra | 1 | 10 | 1 | 277078 |

| | | | | | | |
|---|---|---|---|---|---|---|
| birds | Iole_charlottae | Iole_olivacea | 0 | 24 | 2 | 1131173.67 |
| birds | Irania_gutturalis | Irania_gutturalis | 0 | 21.6 | 4.5 | 2646506.33 |
| birds | Irediparra_gallinacea | Irediparra_gallinacea | 0 | 98.8 | 4 | 2181164.52 |
| birds | Irena_puella | Irena_puella | 0 | 64.9 | 2.4 | 3709810.22 |
| birds | Iridophanes_pulcherrimus | Iridophanes_pulcherrimus | 0 | 15.3 | 2 | 115363.41 |
| birds | Isleria_guttata | Myrmotherula_guttata | 0 | 11 | 2 | 1694815.8 |
| birds | Isleria_hauxwelli | Myrmotherula_hauxwelli | 0 | 10.7 | 2 | 4248364.67 |
| birds | Ispidina_lecontei | Ceyx_lecontei | 0 | 9.5 | 3 | 2195930.36 |
| birds | Ispidina_picta | Ceyx_pictus | 0 | 12.7 | 4.2 | 13894765.2 |
| birds | Ithaginis_cruentus | Ithaginis_cruentus | 0 | 530.7 | 3.7 | 657902.92 |
| birds | Ixidia_cyaniventris | Pycnonotus_cyaniventris | 0 | 22 | 2.4 | 926808.78 |
| birds | Ixidia_erythropthalmos | Pycnonotus_erythropthalmos | 0 | 19.2 | 2 | 1241474.05 |
| birds | Ixidia_squamata | Pycnonotus_squamatus | 0 | 22 | 2 | 298738.78 |
| birds | Ixobrychus_cinnamomeus | Ixobrychus_cinnamomeus | 0 | 126.5 | 3.9 | 8161986.56 |
| birds | Ixobrychus_eurhythmus | Ixobrychus_eurhythmus | 0 | 139 | 4.2 | 8438029.06 |
| birds | Ixobrychus_exilis | Ixobrychus_exilis | 0 | 86.3 | 3.7 | 8171743.8 |
| birds | Ixobrychus_flavicollis | Ixobrychus_flavicollis | 0 | 321 | 4.2 | 8168174.46 |
| birds | Ixobrychus_involucris | Ixobrychus_involucris | 0 | 80.1 | 3.5 | 2399740.23 |
| birds | Ixobrychus_minutus | Ixobrychus_minutus | 0 | 99.5 | 4.2 | 27727320.23 |
| birds | Ixobrychus_sinensis | Ixobrychus_sinensis | 0 | 94.3 | 4.6 | 9358675.07 |
| birds | Ixobrychus_sturmii | Ixobrychus_sturmii | 0 | 140 | 3.5 | 12872490.53 |
| birds | Ixonotus_guttatus | Ixonotus_guttatus | 0 | 35.2 | 2 | 2790568.64 |
| birds | Ixoreus_naevius | Zoothera_naevia | 0 | 77.5 | 3 | 4018779.52 |
| birds | Ixos_malaccensis | Ixos_malaccensis | 0 | 37.3 | 2 | 976235.84 |
| birds | Ixos_mcclellandii | Hypsipetes_mcclellandii | 0 | 32.5 | 2 | 2391920.69 |
| birds | Jabiru_mycteria | Jabiru_mycteria | 0 | 5996.3 | 3.2 | 9048327.31 |
| birds | Jacana_jacana | Jacana_jacana | 0 | 106.2 | 4 | 13881637.25 |
| birds | Jacana_spinosa | Jacana_spinosa | 0 | 94 | 4 | 1048144.39 |
| birds | Junco_hyemalis | Junco_hyemalis | 0 | 19.5 | 4.5 | 16177050.96 |
| birds | Junco_phaeonotus | Junco_phaeonotus | 0 | 20.4 | 3.5 | 532188.09 |
| birds | Junco_vulcani | Junco_vulcani | 0 | 28 | 2 | 1035.18 |
| birds | Jynx_ruficollis | Jynx_ruficollis | 0 | 46.4 | 3.5 | 1609043.05 |
| birds | Jynx_torquilla | Jynx_torquilla | 0 | 35 | 9.2 | 30131695.78 |
| birds | Kaupifalco_monogrammicus | Kaupifalco_monogrammicus | 0 | 278.9 | 2 | 13079368.52 |

| | | | | | | |
|---|---|---|---|---|---|---|
| birds | Kempiella_flavovirescens | Microeca_flavovirescens | 0 | 15 | 2 | 669110.72 |
| birds | Kempiella_griseoceps | Microeca_griseoceps | 0 | 10.6 | 2 | 184726.52 |
| birds | Kenopia_striata | Kenopia_striata | 0 | 19.9 | 2 | 1176589.95 |
| birds | Klais_guimeti | Klais_guimeti | 0 | 2.6 | 2 | 472690.69 |
| birds | Knipolegus_aterrimus | Knipolegus_aterrimus | 0 | 20.2 | 2.4 | 2222826.22 |
| birds | Knipolegus_cyanirostris | Knipolegus_cyanirostris | 0 | 15.4 | 1.7 | 2153351.03 |
| birds | Knipolegus_signatus | Knipolegus_signatus | 0 | 17.8 | 1.7 | 32221.11 |
| birds | Kurochkinegramma_hypogrammica | Hypogramma_hypogrammicum | 0 | 11.4 | 2.4 | 2037212.16 |
| birds | Lacedo_pulchella | Lacedo_pulchella | 0 | 47.3 | 3.2 | 1429865.78 |
| birds | Lafresnaya_lafresnayi | Lafresnaya_lafresnayi | 0 | 5.3 | 2 | 312941.28 |
| birds | Lagonosticta_larvata | Lagonosticta_larvata | 0 | 9.9 | 3.5 | 309178.84 |
| birds | Lagonosticta_rara | Lagonosticta_rara | 0 | 10.4 | 3.5 | 2433099.84 |
| birds | Lagonosticta_rhodopareia | Lagonosticta_rhodopareia | 0 | 9.3 | 3.2 | 2706237.27 |
| birds | Lagonosticta_rubricata | Lagonosticta_rubricata | 0 | 10.2 | 3.9 | 6048001.08 |
| birds | Lagonosticta_rufopicta | Lagonosticta_rufopicta | 0 | 9.3 | 4 | 3308898.34 |
| birds | Lagonosticta_senegala | Lagonosticta_senegala | 0 | 9.2 | 3.5 | 10048316.4 |
| birds | Lagonosticta_virata | Lagonosticta_virata | 0 | 9.3 | 4 | 110944.97 |
| birds | Lagopus_lagopus | Lagopus_lagopus | 0 | 566.9 | 9.4 | 26280139.2 |
| birds | Lagopus_leucura | Lagopus_leucura | 0 | 355 | 5.5 | 1846729.07 |
| birds | Lagopus_muta | Lagopus_muta | 0 | 535.3 | 6.3 | 17597025.31 |
| birds | Lalage_fimbriata | Coracina_fimbriata | 0 | 29.3 | 2 | 1316199.59 |
| birds | Lalage_leucomela | Lalage_leucomela | 0 | 29.3 | 1 | 1258230.14 |
| birds | Lalage_leucopyga | Lalage_leucopyga | 0 | 24.3 | 2 | 34357.57 |
| birds | Lalage_maculosa | Lalage_maculosa | 0 | 30.3 | 1.4 | 30893.86 |
| birds | Lalage_melanoptera | Coracina_melanoptera | 0 | 29.4 | 2.4 | 2480148.8 |
| birds | Lalage_melaschistos | Coracina_melaschistos | 0 | 43 | 3 | 6841809.69 |
| birds | Lalage_nigra | Lalage_nigra | 0 | 28.4 | 2.4 | 1519309.39 |
| birds | Lalage_polioptera | Coracina_polioptera | 0 | 35.8 | 2.4 | 932555.61 |
| birds | Lalage_sueurii | Lalage_sueurii | 0 | 26 | 2 | 259989.37 |
| birds | Lalage_tricolor | Lalage_tricolor | 0 | 25 | 2 | 7633376.41 |
| birds | Lalage_typica | Coracina_typica | 1 | 42.9 | 2 | 1872.3 |
| birds | Lampornis_amethystinus | Lampornis_amethystinus | 0 | 5.2 | 2 | 154790.45 |
| birds | Lampornis_castaneoventris | Lampornis_castaneoventris | 0 | 5.1 | 2 | 1367.41 |
| birds | Lampornis_clemenciae | Lampornis_clemenciae | 0 | 7.6 | 2 | 547582.86 |

| | | | | | | |
|---|---|---|---|---|---|---|
| birds | Lamprolaima_rhami | Lamprolaima_rhami | 0 | 7.8 | 2 | 138239.01 |
| birds | Lamprolia_victoriae | Lamprolia_victoriae | 0 | 14.7 | 1 | 442.19 |
| birds | Lamprotornis_acuticaudus | Lamprotornis_acuticaudus | 0 | 67.6 | 3 | 1542032.33 |
| birds | Lamprotornis_albicapillus | Spreo_albicapillus | 0 | 76.8 | 5.5 | 792878.49 |
| birds | Lamprotornis_australis | Lamprotornis_australis | 0 | 101.1 | 2.8 | 1557885.95 |
| birds | Lamprotornis_bicolor | Spreo_bicolor | 0 | 104 | 3.5 | 787929.61 |
| birds | Lamprotornis_caudatus | Lamprotornis_caudatus | 0 | 121 | 3.5 | 2770669.04 |
| birds | Lamprotornis_chalcurus | Lamprotornis_chalcurus | 0 | 63 | 4 | 2454558.77 |
| birds | Lamprotornis_chalybaeus | Lamprotornis_chalybaeus | 0 | 86.2 | 3.9 | 8694433.54 |
| birds | Lamprotornis_chloropterus | Lamprotornis_chloropterus | 0 | 73.1 | 3 | 5690333.67 |
| birds | Lamprotornis_fischeri | Spreo_fischeri | 0 | 51 | 4.2 | 642683.49 |
| birds | Lamprotornis_hildebrandti | Lamprotornis_hildebrandti | 0 | 55.9 | 3.5 | 413819.2 |
| birds | Lamprotornis_mevesii | Lamprotornis_mevesii | 0 | 64.2 | 3.5 | 639807.37 |
| birds | Lamprotornis_nitens | Lamprotornis_nitens | 0 | 85.6 | 2.4 | 3006322.47 |
| birds | Lamprotornis_pulcher | Lamprotornis_pulcher | 0 | 67.3 | 3.9 | 3948428.54 |
| birds | Lamprotornis_purpureus | Lamprotornis_purpureus | 0 | 112.9 | 3 | 3464722.79 |
| birds | Lamprotornis_purpuroptera | Lamprotornis_purpuroptera | 0 | 77.4 | 3 | 1362512.11 |
| birds | Lamprotornis_regius | Cosmopsarus_regius | 0 | 53.8 | 3.5 | 1243437.56 |
| birds | Lamprotornis_shelleyi | Lamprotornis_shelleyi | 0 | 45 | 4.2 | 1132646.84 |
| birds | Lamprotornis_splendidus | Lamprotornis_splendidus | 0 | 123.1 | 2 | 3906139.83 |
| birds | Lamprotornis_superbus | Lamprotornis_superbus | 0 | 64.9 | 4 | 2087560.01 |
| birds | Lamprotornis_unicolor | Cosmopsarus_unicolor | 0 | 60.2 | 3 | 309310.94 |
| birds | Laniarius_aethiopicus | Laniarius_aethiopicus | 0 | 49.4 | 3 | 7211323.51 |
| birds | Laniarius_atrococcineus | Laniarius_atrococcineus | 0 | 48.8 | 2.4 | 1962868.62 |
| birds | Laniarius_barbarus | Laniarius_barbarus | 0 | 47.1 | 2 | 2828290.55 |
| birds | Laniarius_bicolor | Laniarius_bicolor | 0 | 54.3 | 2 | 459587.12 |
| birds | Laniarius_erythrogaster | Laniarius_erythrogaster | 0 | 48.8 | 2 | 1988843.91 |
| birds | Laniarius_ferrugineus | Laniarius_ferrugineus | 0 | 46.8 | 2 | 721209.79 |
| birds | Laniarius_fuelleborni | Laniarius_fuelleborni | 0 | 46.4 | 2 | 72493.56 |
| birds | Laniarius_funebris | Laniarius_funebris | 0 | 40.4 | 2 | 1152635.02 |
| birds | Laniarius_leucorhynchus | Laniarius_leucorhynchus | 0 | 50.1 | 2 | 2498729.01 |
| birds | Laniarius_luehderi | Laniarius_luehderi | 0 | 42.9 | 2 | 1379816.38 |
| birds | Laniarius_mufumbiri | Laniarius_mufumbiri | 0 | 41.5 | 2 | 98961.39 |
| birds | Laniarius_poensis | Laniarius_poensis | 0 | 44.9 | 2 | 65722.14 |

| | | | | | | |
|---|---|---|---|---|---|---|
| birds | Laniarius_ruficeps | Laniarius_ruficeps | 0 | 32.7 | 2.4 | 660049.39 |
| birds | Laniarius_turatii | Laniarius_turatii | 0 | 45.9 | 2 | 134854.06 |
| birds | Lanio_leucothorax | Lanio_leucothorax | 0 | 40 | 2 | 91070.96 |
| birds | Laniocera_hypopyrra | Laniocera_hypopyrra | 0 | 51 | 2 | 6630155.4 |
| birds | Lanius_bucephalus | Lanius_bucephalus | 0 | 39.4 | 5.3 | 2462379.63 |
| birds | Lanius_cabanisi | Lanius_cabanisi | 0 | 73.8 | 3.5 | 548346.11 |
| birds | Lanius_collaris | Lanius_collaris | 0 | 36.4 | 3.5 | 7220481.63 |
| birds | Lanius_collurio | Lanius_collurio | 0 | 28.4 | 5.5 | 27572252.52 |
| birds | Lanius_collurioides | Lanius_collurioides | 0 | 26 | 4.2 | 2126567.6 |
| birds | Lanius_corvinus | Corvinella_corvina | 0 | 65.4 | 3.9 | 3469655.33 |
| birds | Lanius_cristatus | Lanius_cristatus | 0 | 32.2 | 4.9 | 21999940.84 |
| birds | Lanius_dorsalis | Lanius_dorsalis | 0 | 49.9 | 3.5 | 1312304.95 |
| birds | Lanius_excubitor | Lanius_excubitor | 0 | 63.4 | 5.2 | 36121373.63 |
| birds | Lanius_excubitoroides | Lanius_excubitoroides | 0 | 53.7 | 3 | 2533065.5 |
| birds | Lanius_gubernator | Lanius_gubernator | 0 | 23.5 | 3.5 | 427904.64 |
| birds | Lanius_isabellinus | Lanius_isabellinus | 0 | 25.2 | 4.5 | 25169824.71 |
| birds | Lanius_ludovicianus | Lanius_ludovicianus | 0 | 51.6 | 4.5 | 9319415.63 |
| birds | Lanius_mackinnoni | Lanius_mackinnoni | 0 | 34.3 | 2.4 | 1193617.21 |
| birds | Lanius_melanoleucus | Urolestes_melanoleucus | 0 | 82.2 | 3.9 | 1707357.86 |
| birds | Lanius_meridionalis | Lanius_meridionalis | 1 | 60.4 | 5.5 | 678211.42 |
| birds | Lanius_minor | Lanius_minor | 0 | 46.5 | 5.5 | 10497294.93 |
| birds | Lanius_nubicus | Lanius_nubicus | 0 | 19.7 | 4.6 | 8736712.43 |
| birds | Lanius_schach | Lanius_schach | 0 | 51.5 | 3.5 | 11170512.1 |
| birds | Lanius_senator | Lanius_senator | 0 | 36 | 5.5 | 19072012.9 |
| birds | Lanius_somalicus | Lanius_somalicus | 0 | 45.3 | 3.5 | 858092.07 |
| birds | Lanius_souzae | Lanius_souzae | 0 | 26.3 | 3.5 | 1679903.36 |
| birds | Lanius_sphenocercus | Lanius_sphenocercus | 0 | 92.1 | 6.9 | 4022019.93 |
| birds | Lanius_tephronotus | Lanius_tephronotus | 0 | 46.4 | 4.9 | 3050531.35 |
| birds | Lanius_tigrinus | Lanius_tigrinus | 0 | 30.1 | 5 | 5955735.75 |
| birds | Lanius_vittatus | Lanius_vittatus | 0 | 20.8 | 4 | 4168923.16 |
| birds | Larus_armenicus | Larus_armenicus | 0 | 867 | 3 | 1322509.77 |
| birds | Larus_brunnicephalus | Larus_brunnicephalus | 0 | 569.3 | 3 | 2442213 |
| birds | Larus_bulleri | Larus_bulleri | 0 | 230.2 | 2 | 133703.02 |
| birds | Larus_cachinnans | Larus_cachinnans | 0 | 1111.9 | 2.4 | 7445647.39 |

| | | | | | | |
|---|---|---|---|---|---|---|
| birds | Larus_californicus | Larus_californicus | 0 | 683.4 | 3 | 6058232.19 |
| birds | Larus_cirrocephalus | Larus_cirrocephalus | 0 | 374.7 | 2.4 | 33691966.61 |
| birds | Larus_delawarensis | Larus_delawarensis | 0 | 516.3 | 3 | 13105607.4 |
| birds | Larus_fuscus | Larus_fuscus | 0 | 762.4 | 2.4 | 47994979.5 |
| birds | Larus_genei | Larus_genei | 0 | 281 | 2.4 | 5030312.64 |
| birds | Larus_ichthyaetus | Larus_ichthyaetus | 0 | 1378.8 | 2 | 5985921.73 |
| birds | Larus_maculipennis | Larus_maculipennis | 0 | 339 | 2.4 | 2378141.14 |
| birds | Larus_melanocephalus | Larus_melanocephalus | 0 | 256 | 2.4 | 3340143.42 |
| birds | Larus_modestus | Larus_modestus | 0 | 344 | 1.7 | 567776.38 |
| birds | Larus_novaehollandiae | Larus_novaehollandiae | 0 | 269 | 3 | 8538944.92 |
| birds | Larus_philadelphia | Larus_philadelphia | 0 | 210.7 | 3 | 15309356.61 |
| birds | Larus_pipixcan | Larus_pipixcan | 0 | 280 | 3 | 8494299.95 |
| birds | Larus_relictus | Larus_relictus | 1 | 490.2 | 2.4 | 1443958.55 |
| birds | Larus_ridibundus | Larus_ridibundus | 0 | 284 | 1.7 | 44468354.72 |
| birds | Larus_serranus | Larus_serranus | 0 | 478 | 2.4 | 1115843.78 |
| birds | Larvivora_brunnea | Luscinia_brunnea | 0 | 17.5 | 3.5 | 3708882.52 |
| birds | Larvivora_ruficeps | Luscinia_ruficeps | 1 | 18 | 4 | 46895.98 |
| birds | Laterallus_albigularis | Laterallus_albigularis | 0 | 47.3 | 3.2 | 661410.86 |
| birds | Laterallus_exilis | Laterallus_exilis | 0 | 33.5 | 3 | 6585440.31 |
| birds | Laterallus_flaviventer | Porzana_flaviventer | 0 | 24.9 | 3.9 | 2593534.55 |
| birds | Laterallus_jamaicensis | Laterallus_jamaicensis | 1 | 32.6 | 7.2 | 1137301.5 |
| birds | Laterallus_leucopyrrhus | Laterallus_leucopyrrhus | 0 | 45.5 | 2.4 | 882070.87 |
| birds | Laterallus_melanophaius | Laterallus_melanophaius | 0 | 52.1 | 2.4 | 10160298.32 |
| birds | Laterallus_ruber | Laterallus_ruber | 0 | 44.4 | 4.2 | 557484.26 |
| birds | Laterallus_spilonota | Laterallus_spilonotus | 1 | 40.2 | 4.2 | 2021.77 |
| birds | Lathamus_discolor | Lathamus_discolor | 1 | 64.7 | 3.9 | 767482.19 |
| birds | Lathrotriccus_euleri | Lathrotriccus_euleri | 0 | 11.3 | 2.4 | 10279275.78 |
| birds | Legatus_leucophaius | Legatus_leucophaius | 0 | 22.2 | 2.8 | 11408794.56 |
| birds | Leiopicus_auriceps | Dendrocopos_auriceps | 0 | 39.9 | 3.9 | 79474.36 |
| birds | Leiopicus_medius | Dendrocopos_medius | 0 | 59 | 5.5 | 3966649.98 |
| birds | Leiothlypis_celata | Vermivora_celata | 0 | 9.2 | 4.5 | 15777236.34 |
| birds | Leiothlypis_crissalis | Vermivora_crissalis | 0 | 9.3 | 4 | 557643.61 |
| birds | Leiothlypis_luciae | Vermivora_luciae | 0 | 6.6 | 4.5 | 682328.06 |
| birds | Leiothlypis_peregrina | Vermivora_peregrina | 0 | 8.9 | 6 | 12155394.32 |

| | | | | | | |
|---|---|---|---|---|---|---|
| birds | Leiothlypis_ruficapilla | Vermivora_ruficapilla | 0 | 8.1 | 4.5 | 10359678 |
| birds | Leiothlypis_virginiae | Vermivora_virginiae | 0 | 8.2 | 4 | 1692005.67 |
| birds | Leipoa_ocellata | Leipoa_ocellata | 1 | 1913.9 | 12.8 | 1718483.34 |
| birds | Leistes_bellicosus | Sturnella_bellicosa | 0 | 58 | 3.9 | 351063.41 |
| birds | Leistes_loyca | Sturnella_loyca | 0 | 113 | 4 | 1809024.92 |
| birds | Leistes_militaris | Sturnella_militaris | 0 | 48 | 2.8 | 3688152.76 |
| birds | Lepidocolaptes_affinis | Lepidocolaptes_affinis | 0 | 35 | 2.4 | 238243.15 |
| birds | Lepidocolaptes_angustirostris | Lepidocolaptes_angustirostris | 0 | 29.6 | 3.5 | 6576318.88 |
| birds | Lepidocolaptes_souleyetii | Lepidocolaptes_souleyetii | 0 | 25.7 | 2 | 1484080.31 |
| birds | Lepidopygia_nana | Lonchura_nana | 0 | 8.2 | 4.9 | 590836.07 |
| birds | Lepidothrix_coronata | Lepidothrix_coronata | 0 | 8.3 | 2 | 3221211.22 |
| birds | Lepidothrix_serena | Lepidothrix_serena | 0 | 11.4 | 2 | 583711.44 |
| birds | Leptasthenura_aegithaloides | Leptasthenura_aegithaloides | 0 | 10.7 | 2.8 | 250972.15 |
| birds | Leptasthenura_andicola | Leptasthenura_andicola | 0 | 15.5 | 2 | 256098.23 |
| birds | Leptasthenura_fuliginiceps | Leptasthenura_fuliginiceps | 0 | 12.6 | 2.4 | 369467.91 |
| birds | Leptasthenura_platensis | Leptasthenura_platensis | 0 | 10.6 | 2.8 | 2007502.95 |
| birds | Leptocoma_aspasia | Nectarinia_aspasia | 0 | 8 | 2.4 | 916101.7 |
| birds | Leptocoma_calcostetha | Nectarinia_calcostetha | 0 | 8.3 | 2 | 568344.61 |
| birds | Leptocoma_minima | Nectarinia_minima | 0 | 4.9 | 2 | 57005.42 |
| birds | Leptocoma_sperata | Nectarinia_sperata | 0 | 6.4 | 2 | 229948.47 |
| birds | Leptocoma_zeylonica | Nectarinia_zeylonica | 0 | 8.8 | 2.4 | 1817376.93 |
| birds | Leptopogon_amaurocephalus | Leptopogon_amaurocephalus | 0 | 11.7 | 2.4 | 8060136.76 |
| birds | Leptopogon_superciliaris | Leptopogon_superciliaris | 0 | 11.7 | 2 | 474222.71 |
| birds | Leptopterus_chabert | Leptopterus_chabert | 0 | 19.6 | 3 | 577947.6 |
| birds | Leptoptilos_crumenifer | Leptoptilos_crumeniferus | 0 | 6067.9 | 2 | 12361408.84 |
| birds | Leptoptilos_dubius | Leptoptilos_dubius | 0 | 9050 | 2.8 | 2906151.22 |
| birds | Leptoptilos_javanicus | Leptoptilos_javanicus | 0 | 4651 | 2.8 | 4920849.1 |
| birds | Leptotila_cassinii | Leptotila_cassini | 0 | 159 | 2 | 314249.82 |
| birds | Leptotila_jamaicensis | Leptotila_jamaicensis | 0 | 165 | 2 | 92373.41 |
| birds | Leptotila_megalura | Leptotila_megalura | 0 | 218 | 2 | 147285.79 |
| birds | Leptotila_rufaxilla | Leptotila_rufaxilla | 0 | 157 | 1.4 | 11668214.68 |
| birds | Leptotila_verreauxi | Leptotila_verreauxi | 0 | 146.9 | 2 | 15542979.81 |
| birds | Leptotrygon_veraguensis | Geotrygon_veraguensis | 0 | 155 | 2 | 127743.93 |
| birds | Lerwa_lerwa | Lerwa_lerwa | 0 | 567.3 | 3.2 | 831638.53 |

| | | | | | | |
|---|---|---|---|---|---|---|
| birds | Lesbia_nuna | Lesbia_nuna | 0 | 3.7 | 2 | 251932.1 |
| birds | Lessonia_oreas | Lessonia_oreas | 0 | 13.8 | 3.5 | 831919.21 |
| birds | Lessonia_rufa | Lessonia_rufa | 0 | 13.4 | 2.8 | 3918173.55 |
| birds | Leucippus_fallax | Leucippus_fallax | 0 | 6 | 2 | 93947.05 |
| birds | Leucochloris_albicollis | Leucochloris_albicollis | 0 | 6.3 | 2 | 2176859.29 |
| birds | Leucogeranus_leucogeranus | Grus_leucogeranus | 1 | 5913.4 | 2 | 276450.45 |
| birds | Leucolia_violiceps | Amazilia_violiceps | 0 | 5.1 | 2 | 737817.46 |
| birds | Leuconotopicus_albolarvatus | Picoides_albolarvatus | 0 | 61.1 | 4.5 | 499656.93 |
| birds | Leuconotopicus_borealis | Picoides_borealis | 0 | 48 | 3.2 | 908307.87 |
| birds | Leuconotopicus_fumigatus | Picoides_fumigatus | 0 | 36.2 | 4 | 1109542.22 |
| birds | Leuconotopicus_stricklandi | Picoides_stricklandi | 0 | 40.9 | 2.8 | 18699.26 |
| birds | Leuconotopicus_villosus | Picoides_villosus | 0 | 62.7 | 3.2 | 13332874.44 |
| birds | Leucosarcia_melanoleuca | Leucosarcia_melanoleuca | 0 | 428.6 | 2 | 503204.11 |
| birds | Leucosticte_arctoa | Leucosticte_arctoa | 0 | 30.7 | 4.5 | 3906698.22 |
| birds | Leucosticte_atrata | Leucosticte_atrata | 1 | 25.2 | 4.5 | 970789.69 |
| birds | Leucosticte_australis | Leucosticte_australis | 1 | 26.6 | 4.5 | 178378.62 |
| birds | Leucosticte_nemoricola | Leucosticte_nemoricola | 0 | 22.5 | 3.5 | 4294529.95 |
| birds | Leucosticte_tephrocotis | Leucosticte_tephrocotis | 0 | 37.8 | 4.5 | 4226170.68 |
| birds | Lewinia_striata | Gallirallus_striatus | 0 | 116 | 6.7 | 4893360.6 |
| birds | Lichenostomus_cratitius | Lichenostomus_cratitius | 0 | 20.3 | 2 | 479802.48 |
| birds | Lichenostomus_melanops | Lichenostomus_melanops | 0 | 32.3 | 2 | 732848.75 |
| birds | Limnoctites_rectirostris | Limnoctites_rectirostris | 0 | 18.5 | 2.4 | 179597.01 |
| birds | Limnoctites_sulphuriferus | Cranioleuca_sulphurifera | 0 | 13.5 | 2.8 | 711047.93 |
| birds | Limnodromus_griseus | Limnodromus_griseus | 0 | 110.5 | 4 | 4634264.81 |
| birds | Limnodromus_scolopaceus | Limnodromus_scolopaceus | 0 | 104.4 | 4 | 10469880.22 |
| birds | Limnodromus_semipalmatus | Limnodromus_semipalmatus | 0 | 212 | 2 | 3508614.02 |
| birds | Limnornis_curvirostris | Limnornis_curvirostris | 0 | 28.6 | 2 | 217435.32 |
| birds | Limnothlypis_swainsonii | Limnothlypis_swainsonii | 0 | 18.9 | 3 | 1752740.74 |
| birds | Limosa_fedoa | Limosa_fedoa | 0 | 357 | 4 | 1926237.95 |
| birds | Limosa_haemastica | Limosa_haemastica | 0 | 253.3 | 4 | 3016657.16 |
| birds | Limosa_lapponica | Limosa_lapponica | 0 | 291.7 | 4 | 9184830.81 |
| birds | Limosa_limosa | Limosa_limosa | 0 | 288.4 | 4 | 55640629.68 |
| birds | Linaria_cannabina | Carduelis_cannabina | 0 | 19.5 | 4.9 | 13585958.38 |
| birds | Linaria_flavirostris | Carduelis_flavirostris | 0 | 15.4 | 4.9 | 9158045.26 |

| | | | | | | |
|---|---|---|---|---|---|---|
| birds | Linurgus_olivaceus | Linurgus_olivaceus | 0 | 20.3 | 2.4 | 220044.21 |
| birds | Lipaugus_lanioides | Lipaugus_lanioides | 0 | 94.8 | 1 | 225795.93 |
| birds | Lipaugus_unirufus | Lipaugus_unirufus | 0 | 82.1 | 1 | 462431.54 |
| birds | Lissotis_melanogaster | Eupodotis_melanogaster | 0 | 1151.5 | 1.4 | 10623922.49 |
| birds | Loboparadisea_sericea | Loboparadisea_sericea | 0 | 67.3 | 1 | 161569.22 |
| birds | Lochmias_nematura | Lochmias_nematura | 0 | 22 | 2 | 2648897.91 |
| birds | Locustella_fluviatilis | Locustella_fluviatilis | 0 | 16.1 | 4.9 | 29677457.72 |
| birds | Locustella_lanceolata | Locustella_lanceolata | 0 | 10.6 | 3.9 | 16637022.11 |
| birds | Locustella_luscinioides | Locustella_luscinioides | 0 | 13.9 | 4.5 | 6637595.4 |
| birds | Locustella_luteoventris | Bradypterus_luteoventris | 0 | 13.2 | 3.9 | 1710173.63 |
| birds | Locustella_naevia | Locustella_naevia | 0 | 13.3 | 5.5 | 9444584 |
| birds | Locustella_thoracica | Bradypterus_thoracicus | 0 | 10.8 | 3.5 | 1323689.33 |
| birds | Lonchura_castaneothorax | Lonchura_castaneothorax | 0 | 13.6 | 4.9 | 1876586.43 |
| birds | Lonchura_flaviprymna | Lonchura_flaviprymna | 0 | 11.8 | 4.5 | 117098.74 |
| birds | Lonchura_fuscans | Lonchura_fuscans | 0 | 9.5 | 4.2 | 737265.04 |
| birds | Lonchura_grandis | Lonchura_grandis | 0 | 14.3 | 5.5 | 256441.1 |
| birds | Lonchura_kelaarti | Lonchura_kelaarti | 0 | 14.1 | 5 | 139953.15 |
| birds | Lonchura_leucogastra | Lonchura_leucogastra | 0 | 11.4 | 4.9 | 1481477.6 |
| birds | Lonchura_leucogastroides | Lonchura_leucogastroides | 0 | 11.7 | 5 | 198857.57 |
| birds | Lonchura_maja | Lonchura_maja | 0 | 12.5 | 4.6 | 768204.3 |
| birds | Lonchura_malacca | Lonchura_malacca | 0 | 12.6 | 5.5 | 636026.55 |
| birds | Lonchura_melaena | Lonchura_melaena | 0 | 13.3 | 4.2 | 12280.03 |
| birds | Lonchura_molucca | Lonchura_molucca | 0 | 12 | 4.5 | 313163.12 |
| birds | Lonchura_nevermanni | Lonchura_nevermanni | 0 | 11 | 4.5 | 74957.63 |
| birds | Lonchura_pallida | Lonchura_pallida | 0 | 12 | 4.5 | 226245.63 |
| birds | Lonchura_punctulata | Lonchura_punctulata | 0 | 13.6 | 6.3 | 7671914.83 |
| birds | Lonchura_spectabilis | Lonchura_spectabilis | 0 | 11.5 | 3.9 | 214041.33 |
| birds | Lonchura_striata | Lonchura_striata | 0 | 12.3 | 5 | 5429668.12 |
| birds | Lonchura_stygia | Lonchura_stygia | 0 | 11.2 | 4.5 | 31370.72 |
| birds | Lophaetus_occipitalis | Lophaetus_occipitalis | 0 | 1067 | 1.4 | 14031980.87 |
| birds | Lophoceros_alboterminatus | Tockus_alboterminatus | 0 | 223.7 | 3.2 | 3002290.12 |
| birds | Lophoceros_fasciatus | Tockus_fasciatus | 0 | 258.8 | 4 | 3868026.54 |
| birds | Lophoceros_hemprichii | Tockus_hemprichii | 0 | 297 | 3 | 546665.06 |
| birds | Lophoceros_nasutus | Tockus_nasutus | 0 | 179.6 | 3.2 | 9830291.23 |

| | | | | | | |
|---|---|---|---|---|---|---|
| birds | Lophoceros_pallidirostris | Tockus_pallidirostris | 0 | 233 | 4.5 | 1957669.07 |
| birds | Lophodytes_cucullatus | Lophodytes_cucullatus | 0 | 613.8 | 11 | 8843716.44 |
| birds | Lophoictinia_isura | Lophoictinia_isura | 0 | 579.4 | 3 | 5951770.47 |
| birds | Lopholaimus_antarcticus | Lopholaimus_antarcticus | 0 | 496.4 | 1 | 317014.63 |
| birds | Lophophanes_cristatus | Parus_cristatus | 0 | 11 | 6.3 | 5918314.94 |
| birds | Lophophanes_dichrous | Parus_dichrous | 0 | 13.8 | 4.5 | 968325.05 |
| birds | Lophophorus_impejanus | Lophophorus_impejanus | 0 | 2066.5 | 3.9 | 480828.14 |
| birds | Lophophorus_lhuysii | Lophophorus_lhuysii | 1 | 3008 | 3.9 | 175682.11 |
| birds | Lophophorus_sclateri | Lophophorus_sclateri | 0 | 2500 | 5 | 185023.03 |
| birds | Lophorina_intercedens | Ptiloris_intercedens | 0 | 158 | 2 | 104258.24 |
| birds | Lophorina_magnifica | Ptiloris_magnificus | 0 | 149.9 | 2 | 495422.81 |
| birds | Lophorina_paradisea | Ptiloris_paradiseus | 0 | 119.5 | 2 | 128251.12 |
| birds | Lophorina_superba | Lophorina_superba | 0 | 76.7 | 2 | 14774.58 |
| birds | Lophorina_victoriae | Ptiloris_victoriae | 1 | 95 | 2 | 11240.76 |
| birds | Lophornis_adorabilis | Lophornis_adorabilis | 0 | 2.7 | 2 | 16798 |
| birds | Lophornis_chalybeus | Lophornis_chalybeus | 0 | 3 | 2 | 192385.77 |
| birds | Lophornis_gouldii | Lophornis_gouldii | 0 | 2.4 | 2 | 1585965.49 |
| birds | Lophornis_magnificus | Lophornis_magnificus | 0 | 2.7 | 2 | 2557177.01 |
| birds | Lophornis_ornatus | Lophornis_ornatus | 0 | 2.3 | 2 | 828805.98 |
| birds | Lophornis_pavoninus | Lophornis_pavoninus | 0 | 3 | 2 | 206408.3 |
| birds | Lophotibis_cristata | Lophotibis_cristata | 0 | 583 | 3 | 535452.8 |
| birds | Lophotis_gindiana | Eupodotis_gindiana | 0 | 782.3 | 1.4 | 1777669.92 |
| birds | Lophotis_ruficrista | Eupodotis_ruficrista | 0 | 712.1 | 1.4 | 2013859.39 |
| birds | Lophotriccus_galeatus | Lophotriccus_galeatus | 0 | 6.6 | 2.4 | 2884993.06 |
| birds | Lophotriccus_pileatus | Lophotriccus_pileatus | 0 | 7.5 | 2 | 475468.12 |
| birds | Lophotriorchis_kienerii | Lophotriorchis_kienerii | 0 | 799 | 1 | 3758691.08 |
| birds | Lophura_bulweri | Lophura_bulweri | 1 | 1245.2 | 1 | 274609.44 |
| birds | Lophura_diardi | Lophura_diardi | 0 | 1088.9 | 5.7 | 510759.03 |
| birds | Lophura_edwardsi | Lophura_edwardsi | 1 | 1091 | 5.3 | 8246.95 |
| birds | Lophura_erythrophthalma | Lophura_erythrophthalma | 1 | 934.3 | 4.2 | 374809.69 |
| birds | Lophura_ignita | Lophura_ignita | 1 | 1768.1 | 5.7 | 486548.86 |
| birds | Lophura_inornata | Lophura_inornata | 0 | 685 | 2 | 71415.83 |
| birds | Lophura_leucomelanos | Lophura_leucomelanos | 0 | 868 | 7.3 | 1315909.31 |
| birds | Lophura_nycthemera | Lophura_nycthemera | 0 | 1230 | 6.5 | 1072525.93 |

| | | | | | | |
|---|---|---|---|---|---|---|
| birds | Lophura_swinhoii | Lophura_swinhoii | 0 | 1100 | 5.7 | 15278.8 |
| birds | Loriculus_aurantiifrons | Loriculus_aurantiifrons | 0 | 15.5 | 4 | 573982.76 |
| birds | Loriculus_beryllinus | Loriculus_beryllinus | 0 | 30 | 2.4 | 16049.97 |
| birds | Loriculus_galgulus | Loriculus_galgulus | 0 | 28 | 3 | 1413389.85 |
| birds | Loriculus_philippensis | Loriculus_philippensis | 0 | 34.9 | 3 | 276863 |
| birds | Loriculus_vernalis | Loriculus_vernalis | 0 | 35.7 | 2.8 | 2278295.57 |
| birds | Loxia_curvirostra | Loxia_curvirostra | 0 | 38.3 | 3.5 | 29578527.67 |
| birds | Loxia_leucoptera | Loxia_leucoptera | 0 | 28.7 | 4 | 20422272.71 |
| birds | Loxia_pytyopsittacus | Loxia_pytyopsittacus | 0 | 53 | 3.5 | 3155285.39 |
| birds | Lullula_arborea | Lullula_arborea | 0 | 26.9 | 3.5 | 8808902.55 |
| birds | Lurocalis_semitorquatus | Lurocalis_semitorquatus | 0 | 75.9 | 1 | 8957931.23 |
| birds | Luscinia_luscinia | Luscinia_luscinia | 0 | 23.8 | 4.5 | 8587306.75 |
| birds | Luscinia_megarhynchos | Luscinia_megarhynchos | 0 | 19.6 | 4.5 | 11512328.43 |
| birds | Luscinia_phaenicuroides | Hodgsonius_phaenicuroides | 0 | 21.1 | 2.8 | 1681247.71 |
| birds | Luscinia_svecica | Luscinia_svecica | 0 | 17.2 | 5.5 | 61151327.32 |
| birds | Lybius_chaplini | Lybius_chaplini | 1 | 69.5 | 2.8 | 76477.21 |
| birds | Lybius_guifsobalito | Lybius_guifsobalito | 0 | 42.2 | 3 | 1310708.12 |
| birds | Lybius_leucocephalus | Lybius_leucocephalus | 0 | 62.8 | 2.8 | 1924719.61 |
| birds | Lybius_torquatus | Lybius_torquatus | 0 | 51.7 | 2.2 | 4663900.43 |
| birds | Lybius_vieilloti | Lybius_vieilloti | 0 | 36 | 2.4 | 6618281.31 |
| birds | Lycocorax_pyrrhopterus | Lycocorax_pyrrhopterus | 0 | 283.9 | 1 | 22792.49 |
| birds | Lyncornis_macrotis | Eurostopodus_macrotis | 0 | 140.6 | 1 | 1650527.3 |
| birds | Lyncornis_temminckii | Eurostopodus_temminckii | 0 | 95.9 | 1.4 | 1296744.78 |
| birds | Lyrurus_mlokosiewiczi | Tetrao_mlokosiewiczi | 0 | 814 | 6 | 104757.72 |
| birds | Lyrurus_tetrix | Tetrao_tetrix | 0 | 1068.7 | 8 | 14007422.54 |
| birds | Machaerirhynchus_flaviventer | Machaerirhynchus_flaviventer | 0 | 10.3 | 2 | 708932.93 |
| birds | Machaerirhynchus_nigripectus | Machaerirhynchus_nigripectus | 0 | 12 | 2 | 223752.26 |
| birds | Machaeropterus_deliciosus | Machaeropterus_deliciosus | 0 | 12.7 | 2 | 40123.57 |
| birds | Machaeropterus_pyrocephalus | Machaeropterus_pyrocephalus | 0 | 9.5 | 2 | 2227740.97 |
| birds | Macheiramphus_alcinus | Macheiramphus_alcinus | 0 | 650 | 1 | 13573105.09 |
| birds | Machetornis_rixosa | Machetornis_rixosa | 0 | 29.6 | 3.5 | 6805756.6 |
| birds | Machlolophus_holsti | Parus_holsti | 0 | 16.7 | 3.5 | 12987.01 |
| birds | Machlolophus_spilonotus | Parus_spilonotus | 0 | 18.8 | 4.9 | 1250920.25 |
| birds | Machlolophus_xanthogenys | Parus_xanthogenys | 0 | 14.8 | 4.5 | 1118561.25 |

| | | | | | | |
|---|---|---|---|---|---|---|
| birds | Macronus_ptilosus | Macronous_ptilosus | 0 | 18 | 2.4 | 444346.1 |
| birds | Macronus_striaticeps | Macronous_striaticeps | 0 | 16.5 | 3 | 109828 |
| birds | Macronyx_ameliae | Macronyx_ameliae | 0 | 33.5 | 2.8 | 1902624.72 |
| birds | Macronyx_aurantiigula | Macronyx_aurantiigula | 0 | 39.4 | 2.8 | 375644.56 |
| birds | Macronyx_capensis | Macronyx_capensis | 0 | 50 | 3.2 | 815796.1 |
| birds | Macronyx_croceus | Macronyx_croceus | 0 | 47.1 | 2.8 | 6192075.07 |
| birds | Macronyx_flavicollis | Macronyx_flavicollis | 0 | 45.1 | 2.8 | 404989.73 |
| birds | Macronyx_fuelleborni | Macronyx_fuellebornii | 0 | 55.2 | 2.4 | 2083060.16 |
| birds | Macronyx_sharpei | Macronyx_sharpei | 1 | 29.9 | 2.4 | 2927.17 |
| birds | Macropygia_amboinensis | Macropygia_amboinensis | 0 | 237 | 1 | 1059941.3 |
| birds | Macropygia_emiliana | Macropygia_emiliana | 0 | 161 | 1 | 186249.02 |
| birds | Macropygia_mackinlayi | Macropygia_mackinlayi | 0 | 85.9 | 1.4 | 54491.47 |
| birds | Macropygia_nigrirostris | Macropygia_nigrirostris | 0 | 90.1 | 1 | 646983.57 |
| birds | Macropygia_phasianella | Macropygia_phasianella | 0 | 237 | 1 | 582030.42 |
| birds | Macropygia_ruficeps | Macropygia_ruficeps | 0 | 81.9 | 1 | 1783125.51 |
| birds | Macropygia_unchall | Macropygia_unchall | 0 | 168 | 1.4 | 2550794.48 |
| birds | Malacocincla_abbotti | Malacocincla_abbotti | 0 | 26.8 | 3.9 | 2072332.43 |
| birds | Malacocincla_sepiaria | Malacocincla_sepiaria | 0 | 25.5 | 2 | 969773.94 |
| birds | Malaconotus_blanchoti | Malaconotus_blanchoti | 0 | 78.7 | 3.5 | 7329839.51 |
| birds | Malaconotus_lagdeni | Malaconotus_lagdeni | 0 | 92.3 | 2 | 338592.2 |
| birds | Malacopteron_affine | Malacopteron_affine | 0 | 17.1 | 2 | 387896.02 |
| birds | Malacopteron_cinereum | Malacopteron_cinereum | 0 | 18.1 | 2 | 1526343.71 |
| birds | Malacopteron_magnirostre | Malacopteron_magnirostre | 0 | 19.3 | 2 | 1312918 |
| birds | Malacopteron_magnum | Malacopteron_magnum | 0 | 27.2 | 2 | 666939.16 |
| birds | Malacoptila_panamensis | Malacoptila_panamensis | 0 | 42.6 | 2.4 | 516560.6 |
| birds | Malacorhynchus_membranaceus | Malacorhynchus_membranaceus | 0 | 372.8 | 6.9 | 6997669.67 |
| birds | Malcorus_pectoralis | Malcorus_pectoralis | 0 | 10.1 | 3.9 | 1273750.67 |
| birds | Malimbus_malimbicus | Malimbus_malimbicus | 0 | 36.1 | 2 | 2471944.17 |
| birds | Malimbus_nitens | Malimbus_nitens | 0 | 37.8 | 2 | 2636859.51 |
| birds | Malimbus_scutatus | Malimbus_scutatus | 0 | 30 | 2 | 857359.49 |
| birds | Malurus_alboscapulatus | Malurus_alboscapulatus | 0 | 9.3 | 2.8 | 778806.38 |
| birds | Malurus_amabilis | Malurus_amabilis | 0 | 8.6 | 3 | 126910.61 |
| birds | Malurus_coronatus | Malurus_coronatus | 0 | 11.1 | 2.4 | 363760.08 |
| birds | Malurus_cyaneus | Malurus_cyaneus | 0 | 10.3 | 2.8 | 1239779.31 |

| | | | | | | |
|---|---|---|---|---|---|---|
| birds | Malurus_elegans | Malurus_elegans | 0 | 9.6 | 2.4 | 71313.86 |
| birds | Malurus_lamberti | Malurus_lamberti | 0 | 8 | 3.5 | 212534.11 |
| birds | Malurus_leucopterus | Malurus_leucopterus | 0 | 7.4 | 3.5 | 5612678.07 |
| birds | Malurus_melanocephalus | Malurus_melanocephalus | 0 | 7.9 | 3.5 | 2025747.65 |
| birds | Malurus_pulcherrimus | Malurus_pulcherrimus | 0 | 9.2 | 3 | 379685.75 |
| birds | Malurus_splendens | Malurus_splendens | 0 | 9.4 | 3 | 2710216.86 |
| birds | Manacus_aurantiacus | Manacus_aurantiacus | 0 | 15.5 | 2 | 32298.09 |
| birds | Manacus_candei | Manacus_candei | 0 | 19.8 | 2 | 283521.13 |
| birds | Manacus_vitellinus | Manacus_vitellinus | 0 | 18.2 | 2 | 118914.75 |
| birds | Mandingoa_nitidula | Mandingoa_nitidula | 0 | 9.6 | 4.2 | 1540214.12 |
| birds | Manorina_flavigula | Manorina_flavigula | 0 | 55.9 | 2.8 | 6815495.21 |
| birds | Manorina_melanocephala | Manorina_melanocephala | 0 | 60.2 | 2.8 | 2065259.24 |
| birds | Manorina_melanophrys | Manorina_melanophrys | 0 | 30.4 | 2 | 317354.76 |
| birds | Manorina_melanotis | Manorina_melanotis | 1 | 54 | 2.4 | 66850.62 |
| birds | Mareca_americana | Anas_americana | 0 | 754.6 | 7.9 | 15063927.38 |
| birds | Mareca_falcata | Anas_falcata | 0 | 645.8 | 7.3 | 12027658.54 |
| birds | Mareca_penelope | Anas_penelope | 0 | 770 | 8.5 | 29250559.93 |
| birds | Mareca_sibilatrix | Anas_sibilatrix | 0 | 923.2 | 6.3 | 3564873.37 |
| birds | Mareca_strepera | Anas_strepera | 0 | 915.6 | 9.8 | 34553002.07 |
| birds | Margarops_fuscatus | Margarops_fuscatus | 0 | 98.4 | 2.4 | 20593.03 |
| birds | Marmaronetta_angustirostris | Marmaronetta_angustirostris | 0 | 477 | 9.9 | 12485474.47 |
| birds | Masius_chrysopterus | Masius_chrysopterus | 0 | 11.5 | 2 | 176828.86 |
| birds | Mayrornis_lessoni | Mayrornis_lessoni | 0 | 11.2 | 1.4 | 17898.17 |
| birds | Megaceryle_alcyon | Megaceryle_alcyon | 0 | 148 | 6.5 | 18076084.92 |
| birds | Megaceryle_lugubris | Megaceryle_lugubris | 0 | 271 | 4.5 | 5024200.4 |
| birds | Megaceryle_maxima | Megaceryle_maxima | 0 | 325 | 3.9 | 13133154.9 |
| birds | Megaceryle_torquata | Megaceryle_torquata | 0 | 317 | 4 | 16295521.19 |
| birds | Megaloprepia_formosa | Ptilinopus_bernsteinii | 0 | 118.5 | 1 | 21523.73 |
| birds | Megaloprepia_magnifica | Ptilinopus_magnificus | 0 | 185.4 | 1 | 979740.41 |
| birds | Megalurus_palustris | Megalurus_palustris | 0 | 44.1 | 3.5 | 2258407.86 |
| birds | Megarynchus_pitangua | Megarynchus_pitangua | 0 | 69.9 | 2.4 | 13243099.5 |
| birds | Megascops_asio | Megascops_asio | 0 | 180 | 3.5 | 4938942.17 |
| birds | Megascops_choliba | Megascops_choliba | 0 | 132 | 2 | 13065137.78 |
| birds | Megascops_cooperi | Megascops_cooperi | 0 | 149.9 | 3.9 | 78738.89 |

| | | | | | | |
|---|---|---|---|---|---|---|
| birds | Megascops_guatemalae | Megascops_guatemalae | 0 | 118 | 3 | 660793.49 |
| birds | Megascops_kennicottii | Megascops_kennicottii | 0 | 141.8 | 4.9 | 3582764.44 |
| birds | Megascops_nudipes | Megascops_nudipes | 0 | 137.3 | 2 | 9396.11 |
| birds | Megascops_trichopsis | Megascops_trichopsis | 0 | 88.3 | 3.5 | 660977.77 |
| birds | Megastictus_margaritatus | Megastictus_margaritatus | 0 | 20.2 | 2 | 1937447.72 |
| birds | Meiglyptes_tristis | Meiglyptes_tristis | 1 | 31.7 | 2 | 50016.69 |
| birds | Meiglyptes_tukki | Meiglyptes_tukki | 0 | 53.1 | 2 | 772219.58 |
| birds | Melaenornis_ardesiacus | Melaenornis_ardesiacus | 0 | 29 | 2 | 60515.81 |
| birds | Melaenornis_chocolatinus | Dioptrornis_chocolatinus | 0 | 23.4 | 3 | 202824.29 |
| birds | Melaenornis_edolioides | Melaenornis_edolioides | 0 | 30.7 | 2.4 | 3825770.16 |
| birds | Melaenornis_pammelaina | Melaenornis_pammelaina | 0 | 30.2 | 2.8 | 4080335.72 |
| birds | Melanerpes_aurifrons | Melanerpes_aurifrons | 0 | 80.8 | 4.5 | 1414625.36 |
| birds | Melanerpes_candidus | Melanerpes_candidus | 0 | 108 | 3.5 | 5535970.53 |
| birds | Melanerpes_carolinus | Melanerpes_carolinus | 0 | 69.5 | 4.9 | 3074161.82 |
| birds | Melanerpes_chrysauchen | Melanerpes_chrysauchen | 0 | 52.5 | 3.5 | 15256.01 |
| birds | Melanerpes_erythrocephalus | Melanerpes_erythrocephalus | 0 | 71.6 | 5.3 | 5461683.2 |
| birds | Melanerpes_formicivorus | Melanerpes_formicivorus | 0 | 79.6 | 4.9 | 1502706.45 |
| birds | Melanerpes_hoffmannii | Melanerpes_hoffmannii | 0 | 72.2 | 2.4 | 51111.87 |
| birds | Melanerpes_lewis | Melanerpes_lewis | 0 | 105.8 | 6.5 | 3401486.03 |
| birds | Melanerpes_pucherani | Melanerpes_pucherani | 0 | 62.4 | 2.8 | 486437.57 |
| birds | Melanerpes_radiolatus | Melanerpes_radiolatus | 0 | 108 | 3.9 | 11021.79 |
| birds | Melanerpes_rubricapillus | Melanerpes_rubricapillus | 0 | 52.3 | 3.5 | 877834.62 |
| birds | Melanerpes_striatus | Melanerpes_striatus | 0 | 73.4 | 4.9 | 74161.47 |
| birds | Melanerpes_superciliaris | Melanerpes_superciliaris | 0 | 77.3 | 4.5 | 114069.09 |
| birds | Melanerpes_uropygialis | Melanerpes_uropygialis | 0 | 64.7 | 3.9 | 567324.49 |
| birds | Melaniparus_afer | Parus_afer | 0 | 20.2 | 3.5 | 427814.63 |
| birds | Melaniparus_albiventris | Parus_albiventris | 0 | 21.2 | 3.9 | 322293.63 |
| birds | Melaniparus_cinerascens | Parus_cinerascens | 0 | 20 | 4.2 | 1795200.59 |
| birds | Melaniparus_fasciiventer | Parus_fasciiventer | 0 | 15.2 | 3 | 38778.02 |
| birds | Melaniparus_fringillinus | Parus_fringillinus | 0 | 13.8 | 3 | 390583.14 |
| birds | Melaniparus_funereus | Parus_funereus | 0 | 25.4 | 3 | 2252792.06 |
| birds | Melaniparus_griseiventris | Parus_griseiventris | 0 | 15.5 | 3.9 | 1535847.66 |
| birds | Melaniparus_leucomelas | Parus_leucomelas | 0 | 16.1 | 3.9 | 2587731.88 |
| birds | Melaniparus_niger | Parus_niger | 0 | 21.5 | 4.5 | 2180649.32 |

| | | | | | | |
|---|---|---|---|---|---|---|
| birds | Melaniparus_rufiventris | Parus_rufiventris | 0 | 18.2 | 3.2 | 2428644.32 |
| birds | Melanitta_fusca | Melanitta_fusca | 1 | 1800.2 | 7.9 | 5830953.16 |
| birds | Melanitta_nigra | Melanitta_nigra | 0 | 1050 | 6.9 | 5162178.68 |
| birds | Melanitta_perspicillata | Melanitta_perspicillata | 0 | 1096.3 | 6.3 | 6744994.04 |
| birds | Melanocichla_lugubris | Garrulax_lugubris | 0 | 78.6 | 2 | 41578.04 |
| birds | Melanocorypha_bimaculata | Melanocorypha_bimaculata | 0 | 54.1 | 4.2 | 4993981.9 |
| birds | Melanocorypha_calandra | Melanocorypha_calandra | 0 | 61.6 | 4.2 | 6304656.36 |
| birds | Melanocorypha_maxima | Melanocorypha_maxima | 0 | 75 | 2.8 | 1042885.9 |
| birds | Melanocorypha_mongolica | Melanocorypha_mongolica | 0 | 53.9 | 3.9 | 4731925.5 |
| birds | Melanocorypha_yeltoniensis | Melanocorypha_yeltoniensis | 0 | 59.9 | 4 | 3219779.34 |
| birds | Melanodryas_cucullata | Melanodryas_cucullata | 0 | 21.1 | 2 | 6360149.09 |
| birds | Melanodryas_vittata | Melanodryas_vittata | 1 | 26.6 | 2.4 | 68050.36 |
| birds | Melanoptila_glabrirostris | Melanoptila_glabrirostris | 0 | 35.4 | 2 | 151379.52 |
| birds | Melanotis_caerulescens | Melanotis_caerulescens | 0 | 61.6 | 2 | 577099.95 |
| birds | Melanotis_hypoleucus | Melanotis_hypoleucus | 0 | 68 | 2.4 | 98583.12 |
| birds | Meleagris_gallopavo | Meleagris_gallopavo | 0 | 5791.4 | 11.4 | 7375475.41 |
| birds | Meleagris_ocellata | Meleagris_ocellata | 0 | 5525 | 12 | 163840.22 |
| birds | Melierax_canorus | Melierax_canorus | 0 | 663.3 | 2 | 2246754.63 |
| birds | Melierax_metabates | Melierax_metabates | 0 | 674.5 | 1.4 | 12484209.96 |
| birds | Melierax_poliopterus | Melierax_poliopterus | 0 | 629.9 | 1.4 | 2071175.93 |
| birds | Meliphaga_aruensis | Meliphaga_aruensis | 0 | 26.2 | 1.4 | 665428.4 |
| birds | Meliphaga_lewinii | Meliphaga_lewinii | 0 | 33.6 | 2 | 643140.81 |
| birds | Meliphaga_notata | Meliphaga_notata | 0 | 26.4 | 2 | 110238.59 |
| birds | Melithreptus_affinis | Melithreptus_affinis | 0 | 15 | 3 | 68030 |
| birds | Melithreptus_albogularis | Melithreptus_albogularis | 0 | 11 | 2 | 1473520.94 |
| birds | Melithreptus_brevirostris | Melithreptus_brevirostris | 0 | 12.6 | 3 | 2003053.24 |
| birds | Melithreptus_gularis | Melithreptus_gularis | 0 | 19.4 | 2.4 | 1260779.55 |
| birds | Melithreptus_lunatus | Melithreptus_lunatus | 0 | 14 | 2.4 | 1242453.6 |
| birds | Melithreptus_validirostris | Melithreptus_validirostris | 1 | 24.8 | 3 | 66891.32 |
| birds | Mellisuga_helenae | Mellisuga_helenae | 0 | 2.2 | 2 | 108875.85 |
| birds | Mellisuga_minima | Mellisuga_minima | 0 | 2.4 | 2 | 86168.43 |
| birds | Melloria_quoyi | Cracticus_quoyi | 0 | 168 | 2 | 1145717.61 |
| birds | Melocichla_mentalis | Melocichla_mentalis | 0 | 33.2 | 2.4 | 6248614.06 |
| birds | Melopsittacus_undulatus | Melopsittacus_undulatus | 0 | 29 | 4.9 | 6191813 |

| | | | | | | |
|---|---|---|---|---|---|---|
| birds | Melospiza_georgiana | Melospiza_georgiana | 0 | 16.1 | 4.5 | 10922479.13 |
| birds | Melospiza_lincolnii | Melospiza_lincolnii | 0 | 16.6 | 4.5 | 16098839.35 |
| birds | Melospiza_melodia | Melospiza_melodia | 0 | 21.9 | 3.9 | 13306731.28 |
| birds | Melozone_aberti | Pipilo_aberti | 0 | 45.9 | 3.5 | 170236.93 |
| birds | Melozone_albicollis | Pipilo_albicollis | 0 | 46.4 | 4 | 64681.63 |
| birds | Melozone_crissalis | Pipilo_crissalis | 0 | 52.8 | 3.5 | 333326.69 |
| birds | Melozone_fusca | Pipilo_fuscus | 0 | 44.4 | 3.5 | 1684944.38 |
| birds | Mentocrex_kioloides | Canirallus_kioloides | 0 | 246 | 2 | 14988.07 |
| birds | Menura_alberti | Menura_alberti | 0 | 928 | 1 | 40026.78 |
| birds | Menura_novaehollandiae | Menura_novaehollandiae | 0 | 972.6 | 1 | 325624.39 |
| birds | Merganetta_armata | Merganetta_armata | 0 | 398.8 | 3.5 | 1090659.13 |
| birds | Mergellus_albellus | Mergellus_albellus | 0 | 608.6 | 7.9 | 13903839.29 |
| birds | Mergus_merganser | Mergus_merganser | 0 | 1451 | 9.8 | 35370520.54 |
| birds | Mergus_serrator | Mergus_serrator | 0 | 1015.2 | 8.9 | 37225500.74 |
| birds | Mergus_squamatus | Mergus_squamatus | 1 | 1234 | 10 | 2240002.66 |
| birds | Merops_albicollis | Merops_albicollis | 0 | 25.9 | 5.3 | 10467706.23 |
| birds | Merops_apiaster | Merops_apiaster | 0 | 56.6 | 6.3 | 13243490.57 |
| birds | Merops_boehmi | Merops_boehmi | 0 | 16.6 | 2.8 | 700033.6 |
| birds | Merops_breweri | Merops_breweri | 0 | 49.7 | 2.4 | 1632583.2 |
| birds | Merops_bullockoides | Merops_bullockoides | 0 | 34.8 | 3.2 | 5313566.69 |
| birds | Merops_bulocki | Merops_bulocki | 0 | 23.1 | 2.8 | 2933973.61 |
| birds | Merops_gularis | Merops_gularis | 0 | 27.3 | 2 | 3039017.17 |
| birds | Merops_hirundineus | Merops_hirundineus | 0 | 21.7 | 2.8 | 8552948.69 |
| birds | Merops_leschenaulti | Merops_leschenaulti | 0 | 27.2 | 5.7 | 2822822.27 |
| birds | Merops_malimbicus | Merops_malimbicus | 0 | 50.4 | 2 | 1453484.12 |
| birds | Merops_muelleri | Merops_muelleri | 0 | 22.5 | 2 | 1597560.38 |
| birds | Merops_nubicus | Merops_nubicus | 0 | 42.4 | 3.2 | 8296122.08 |
| birds | Merops_oreobates | Merops_oreobates | 0 | 24 | 2.4 | 360934.32 |
| birds | Merops_orientalis | Merops_orientalis | 0 | 14.8 | 5.7 | 5717891 |
| birds | Merops_ornatus | Merops_ornatus | 0 | 29.5 | 4.6 | 7621196.56 |
| birds | Merops_persicus | Merops_persicus | 0 | 49.3 | 5.7 | 16160299.23 |
| birds | Merops_philippinus | Merops_philippinus | 0 | 34 | 5.9 | 6418828.75 |
| birds | Merops_pusillus | Merops_pusillus | 0 | 15.1 | 4.9 | 14410773.71 |
| birds | Merops_superciliosus | Merops_superciliosus | 0 | 38.5 | 4.9 | 4269964.77 |

| | | | | | | |
|---|---|---|---|---|---|---|
| birds | Merops_variegatus | Merops_variegatus | 0 | 22.5 | 2.4 | 3491407.89 |
| birds | Merops_viridis | Merops_viridis | 0 | 34.8 | 4.2 | 3204518.18 |
| birds | Mesitornis_unicolor | Mesitornis_unicolor | 1 | 148 | 1 | 37168.05 |
| birds | Mesitornis_variegatus | Mesitornis_variegatus | 1 | 110 | 1.7 | 39538.16 |
| birds | Metopidius_indicus | Metopidius_indicus | 0 | 155 | 4 | 2649332.71 |
| birds | Micrastur_ruficollis | Micrastur_ruficollis | 0 | 177.6 | 2.4 | 12486169.39 |
| birds | Micrastur_semitorquatus | Micrastur_semitorquatus | 0 | 621.7 | 2.4 | 13182553.54 |
| birds | Micrathene_whitneyi | Micrathene_whitneyi | 0 | 41 | 2.2 | 1713909.24 |
| birds | Microbates_cinereiventris | Microbates_cinereiventris | 0 | 11.9 | 2 | 551552.19 |
| birds | Microcarbo_africanus | Phalacrocorax_africanus | 0 | 545 | 3.5 | 17477574.05 |
| birds | Microcarbo_melanoleucos | Phalacrocorax_melanoleucos | 0 | 727.4 | 3.9 | 5853894.37 |
| birds | Microcarbo_niger | Phalacrocorax_niger | 0 | 427 | 3.9 | 4902890.62 |
| birds | Microcarbo_pygmaeus | Phalacrocorax_pygmeus | 0 | 676 | 4.9 | 3509944.42 |
| birds | Microcerculus_bambla | Microcerculus_bambla | 0 | 19.1 | 1 | 1714246.73 |
| birds | Microcerculus_marginatus | Microcerculus_marginatus | 0 | 18.2 | 2.4 | 4781168.47 |
| birds | Microchera_cupreiceps | Elvira_cupreiceps | 0 | 3 | 2 | 5223.26 |
| birds | Microeca_fascinans | Microeca_fascinans | 0 | 14.5 | 2.4 | 5302401.74 |
| birds | Microeca_flavigaster | Microeca_flavigaster | 0 | 12.1 | 1 | 1175473.95 |
| birds | Microhierax_caerulescens | Microhierax_caerulescens | 0 | 38.7 | 4.5 | 1206450.7 |
| birds | Microhierax_fringillarius | Microhierax_fringillarius | 0 | 42.9 | 4.5 | 1402352.92 |
| birds | Microligea_palustris | Microligea_palustris | 0 | 13.2 | 2 | 30480.02 |
| birds | Micromonacha_lanceolata | Micromonacha_lanceolata | 0 | 19.8 | 2 | 1651230.16 |
| birds | Micronisus_gabar | Melierax_gabar | 0 | 169.6 | 2 | 13910595.47 |
| birds | Microparra_capensis | Microparra_capensis | 0 | 41.3 | 3 | 3244082.83 |
| birds | Micropternus_brachyurus | Celeus_brachyurus | 0 | 85.9 | 2.4 | 6131273.93 |
| birds | Microptilotis_albilineatus | Meliphaga_albilineata | 0 | 23.7 | 2 | 53764.24 |
| birds | Microptilotis_albonotatus | Meliphaga_albonotata | 0 | 22.4 | 1.4 | 320661.12 |
| birds | Microptilotis_gracilis | Meliphaga_gracilis | 0 | 15 | 2 | 185643.55 |
| birds | Micropygia_schomburgkii | Micropygia_schomburgkii | 0 | 31.2 | 2 | 3010330.87 |
| birds | Microrhopias_quixensis | Microrhopias_quixensis | 0 | 7.9 | 2 | 4104536.42 |
| birds | Milvus_migrans | Milvus_migrans | 0 | 734.1 | 2.4 | 71149092.14 |
| birds | Milvus_milvus | Milvus_milvus | 0 | 1071.8 | 2 | 3974917.68 |
| birds | Mimus_gilvus | Mimus_gilvus | 0 | 52 | 2.8 | 2168760.73 |
| birds | Mimus_gundlachii | Mimus_gundlachii | 0 | 66.8 | 2 | 15471.5 |

| | | | | | | |
|---|---|---|---|---|---|---|
| birds | Mimus_parvulus | Mimus_parvulus | 0 | 53.6 | 3.5 | 6967.11 |
| birds | Mimus_polyglottos | Mimus_polyglottos | 0 | 48.5 | 3.5 | 10313735.12 |
| birds | Mimus_saturninus | Mimus_saturninus | 0 | 63.7 | 3.5 | 7198431.03 |
| birds | Mimus_thenca | Mimus_thenca | 0 | 66 | 3.5 | 229319.44 |
| birds | Mimus_triurus | Mimus_triurus | 0 | 47.2 | 4 | 3284288.52 |
| birds | Mino_anais | Mino_anais | 0 | 152 | 2 | 375373.57 |
| birds | Mino_dumontii | Mino_dumontii | 0 | 217 | 1.4 | 640179.95 |
| birds | Mionectes_macconnelli | Mionectes_macconnelli | 0 | 13.5 | 3 | 2588557.72 |
| birds | Mionectes_oleagineus | Mionectes_oleagineus | 0 | 11.2 | 3.2 | 8947569.12 |
| birds | Mionectes_olivaceus | Mionectes_olivaceus | 0 | 15.3 | 2.4 | 39754.75 |
| birds | Mionectes_rufiventris | Mionectes_rufiventris | 0 | 13.3 | 3 | 1057487.4 |
| birds | Mionectes_striaticollis | Mionectes_striaticollis | 0 | 15 | 2.4 | 318010.85 |
| birds | Mirafra_africana | Mirafra_africana | 0 | 45 | 2.8 | 5074225.03 |
| birds | Mirafra_albicauda | Mirafra_albicauda | 0 | 22.2 | 2 | 516863.45 |
| birds | Mirafra_angolensis | Mirafra_angolensis | 0 | 36.2 | 3 | 763784.63 |
| birds | Mirafra_apiata | Mirafra_apiata | 0 | 26.9 | 2.4 | 258562.29 |
| birds | Mirafra_assamica | Mirafra_assamica | 0 | 26.2 | 3.5 | 808359.62 |
| birds | Mirafra_cheniana | Mirafra_cheniana | 0 | 29.2 | 2.8 | 258493.25 |
| birds | Mirafra_collaris | Mirafra_collaris | 0 | 29.2 | 2.4 | 319160.76 |
| birds | Mirafra_erythroptera | Mirafra_erythroptera | 0 | 21.3 | 2.8 | 2147813.67 |
| birds | Mirafra_hypermetra | Mirafra_hypermetra | 0 | 61.4 | 2 | 451245.37 |
| birds | Mirafra_javanica | Mirafra_javanica | 0 | 23 | 2.8 | 11882223.56 |
| birds | Mirafra_passerina | Mirafra_passerina | 0 | 24.7 | 2.8 | 1870815.46 |
| birds | Mirafra_rufocinnamomea | Mirafra_rufocinnamomea | 0 | 26.3 | 2.4 | 9716191.66 |
| birds | Mirafra_somalica | Mirafra_somalica | 0 | 46.6 | 3.5 | 124775.11 |
| birds | Mitrephanes_phaeocercus | Mitrephanes_phaeocercus | 0 | 8.6 | 2 | 882909.56 |
| birds | Mitrospingus_cassinii | Mitrospingus_cassinii | 0 | 40.4 | 2 | 167653.28 |
| birds | Mixornis_gularis | Macronous_gularis | 0 | 11.8 | 3.5 | 3009284.29 |
| birds | Mniotilta_varia | Mniotilta_varia | 0 | 10.9 | 4.5 | 12084523.74 |
| birds | Modulatrix_stictigula | Modulatrix_stictigula | 0 | 30.5 | 2 | 70768.87 |
| birds | Monarcha_castaneiventris | Monarcha_castaneiventris | 0 | 27.3 | 3 | 27842.78 |
| birds | Monarcha_cinerascens | Monarcha_cinerascens | 0 | 29.6 | 1.4 | 148476.26 |
| birds | Monarcha_frater | Monarcha_frater | 0 | 22 | 2.4 | 226581.08 |
| birds | Monarcha_melanopsis | Monarcha_melanopsis | 0 | 22.7 | 2.4 | 571217.8 |

| | | | | | | |
|---|---|---|---|---|---|---|
| birds | Monasa_morphoeus | Monasa_morphoeus | 0 | 87.9 | 2.4 | 5210334.01 |
| birds | Monasa_nigrifrons | Monasa_nigrifrons | 0 | 80.7 | 3 | 7021917.73 |
| birds | Monias_benschi | Monias_benschi | 1 | 137.9 | 1.4 | 12321.8 |
| birds | Montecincla_cachinnans | Strophocincla_cachinnans | 1 | 40.3 | 2.4 | 1070.7 |
| birds | Monticola_angolensis | Monticola_angolensis | 0 | 44.4 | 3.5 | 2218343.4 |
| birds | Monticola_brevipes | Monticola_brevipes | 0 | 33 | 2.4 | 1043690.36 |
| birds | Monticola_cinclorhyncha | Monticola_cinclorhynchus | 0 | 36 | 3.5 | 2943767.32 |
| birds | Monticola_explorator | Monticola_explorator | 0 | 44.4 | 2.4 | 368644.24 |
| birds | Monticola_gularis | Monticola_gularis | 0 | 34.6 | 3.9 | 3922614.37 |
| birds | Monticola_imerina | Monticola_imerinus | 0 | 28.3 | 2 | 18846.5 |
| birds | Monticola_rufiventris | Monticola_rufiventris | 0 | 53.2 | 3.5 | 3015321.11 |
| birds | Monticola_rufocinereus | Monticola_rufocinereus | 0 | 24 | 3 | 583391.08 |
| birds | Monticola_rupestris | Monticola_rupestris | 0 | 61.4 | 2.4 | 557569.05 |
| birds | Monticola_saxatilis | Monticola_saxatilis | 0 | 50.3 | 3.5 | 14463681.44 |
| birds | Monticola_semirufus | Myrmecocichla_semirufa | 0 | 46.7 | 3 | 386134.07 |
| birds | Monticola_solitarius | Monticola_solitarius | 0 | 53.7 | 4.2 | 21338038.17 |
| birds | Montifringilla_nivalis | Montifringilla_nivalis | 0 | 36.9 | 4.5 | 4085451.28 |
| birds | Morococcyx_erythropygus | Morococcyx_erythropygus | 0 | 64.5 | 2 | 358833.18 |
| birds | Motacilla_aguimp | Motacilla_aguimp | 0 | 27 | 3.5 | 11321309.29 |
| birds | Motacilla_alba | Motacilla_alba | 0 | 23.9 | 4.9 | 59320722.26 |
| birds | Motacilla_capensis | Motacilla_capensis | 0 | 20.8 | 2 | 3891210.17 |
| birds | Motacilla_cinerea | Motacilla_cinerea | 0 | 17.2 | 4.6 | 32926989.11 |
| birds | Motacilla_citreola | Motacilla_citreola | 0 | 19.1 | 4.2 | 33176540.87 |
| birds | Motacilla_clara | Motacilla_clara | 0 | 17.9 | 2.4 | 3122291.69 |
| birds | Motacilla_flava | Motacilla_flava | 0 | 17.7 | 4.9 | 58182475.75 |
| birds | Motacilla_flaviventris | Motacilla_flaviventris | 0 | 22.5 | 2 | 591120.51 |
| birds | Motacilla_grandis | Motacilla_grandis | 0 | 30.2 | 5.3 | 435186.17 |
| birds | Motacilla_maderaspatensis | Motacilla_madaraspatensis | 0 | 30.5 | 3.9 | 2844265.94 |
| birds | Mulleripicus_fulvus | Mulleripicus_fulvus | 0 | 312.3 | 2.4 | 103818.08 |
| birds | Mulleripicus_pulverulentus | Mulleripicus_pulverulentus | 1 | 450.2 | 2.8 | 2625286.94 |
| birds | Muscicapa_adusta | Muscicapa_adusta | 0 | 10.4 | 2.4 | 2389660.85 |
| birds | Muscicapa_aquatica | Muscicapa_aquatica | 0 | 11.5 | 2.4 | 1932273.67 |
| birds | Muscicapa_cassini | Muscicapa_cassini | 0 | 17.3 | 1.7 | 3716280.13 |
| birds | Muscicapa_dauurica | Muscicapa_dauurica | 0 | 9.8 | 2.8 | 17791128.02 |

| | | | | | | |
|---|---|---|---|---|---|---|
| birds | Muscicapa_epulata | Muscicapa_epulata | 0 | 9.6 | 2 | 903510.24 |
| birds | Muscicapa_ferruginea | Muscicapa_ferruginea | 0 | 12 | 2.4 | 2544222 |
| birds | Muscicapa_gambagae | Muscicapa_gambagae | 0 | 13 | 2 | 669433.07 |
| birds | Muscicapa_griseisticta | Muscicapa_griseisticta | 0 | 15.9 | 3.9 | 2648878.54 |
| birds | Muscicapa_muttui | Muscicapa_muttui | 0 | 12 | 4.5 | 1292048.54 |
| birds | Muscicapa_sethsmithi | Muscicapa_sethsmithi | 0 | 9.1 | 2 | 1294819.35 |
| birds | Muscicapa_sibirica | Muscicapa_sibirica | 0 | 13.2 | 3.5 | 17841855.28 |
| birds | Muscicapa_striata | Muscicapa_striata | 0 | 15.9 | 3.5 | 47138089.09 |
| birds | Muscigralla_brevicauda | Muscigralla_brevicauda | 0 | 12.6 | 3.9 | 166922.92 |
| birds | Muscipipra_vetula | Muscipipra_vetula | 0 | 27 | 3 | 828420 |
| birds | Muscisaxicola_albilora | Muscisaxicola_albilora | 0 | 22.6 | 2.4 | 1237528.62 |
| birds | Muscisaxicola_alpinus | Muscisaxicola_alpinus | 0 | 26.1 | 4 | 42973.19 |
| birds | Muscisaxicola_capistratus | Muscisaxicola_capistratus | 0 | 26.6 | 3 | 2111097.81 |
| birds | Muscisaxicola_cinereus | Muscisaxicola_cinereus | 0 | 18.6 | 2 | 770735.32 |
| birds | Muscisaxicola_flavinucha | Muscisaxicola_flavinucha | 0 | 36.2 | 2.8 | 1456441.88 |
| birds | Muscisaxicola_fluviatilis | Muscisaxicola_fluviatilis | 0 | 14.7 | 2 | 1234514.01 |
| birds | Muscisaxicola_maclovianus | Muscisaxicola_maclovianus | 0 | 23.8 | 2.4 | 2835656.76 |
| birds | Muscisaxicola_maculirostris | Muscisaxicola_maculirostris | 0 | 14.2 | 2.8 | 1996565.98 |
| birds | Muscisaxicola_rufivertex | Muscisaxicola_rufivertex | 0 | 20 | 3 | 696474.96 |
| birds | Musophaga_rossae | Musophaga_rossae | 0 | 361.4 | 1.4 | 3118175.85 |
| birds | Musophaga_violacea | Musophaga_violacea | 0 | 360 | 1.4 | 2077717.14 |
| birds | Myadestes_elisabeth | Myadestes_elisabeth | 0 | 30 | 3 | 8505.1 |
| birds | Myadestes_genibarbis | Myadestes_genibarbis | 0 | 29.2 | 2 | 88012.58 |
| birds | Myadestes_melanops | Myadestes_melanops | 0 | 32.1 | 2.4 | 31920.03 |
| birds | Myadestes_obscurus | Myadestes_obscurus | 0 | 50.4 | 1.4 | 6883.28 |
| birds | Myadestes_occidentalis | Myadestes_occidentalis | 0 | 36.4 | 2.4 | 528718.34 |
| birds | Myadestes_ralloides | Myadestes_ralloides | 0 | 29.1 | 2 | 457703.18 |
| birds | Myadestes_townsendi | Myadestes_townsendi | 0 | 33.2 | 3.5 | 6168503.62 |
| birds | Mycteria_americana | Mycteria_americana | 0 | 2554.5 | 3.2 | 14862621.42 |
| birds | Mycteria_cinerea | Mycteria_cinerea | 1 | 2393 | 2 | 155396.26 |
| birds | Mycteria_ibis | Mycteria_ibis | 0 | 2155.6 | 2.4 | 15368408.04 |
| birds | Mycteria_leucocephala | Mycteria_leucocephala | 0 | 3180 | 3.2 | 3983384 |
| birds | Myiagra_alecto | Myiagra_alecto | 0 | 23.6 | 2.4 | 1376502.43 |
| birds | Myiagra_azureocapilla | Myiagra_azureocapilla | 0 | 12.9 | 1 | 441.31 |

| | | | | | | |
|---|---|---|---|---|---|---|
| birds | Myiagra_caledonica | Myiagra_caledonica | 0 | 10.8 | 2.4 | 31488.18 |
| birds | Myiagra_cyanoleuca | Myiagra_cyanoleuca | 0 | 17.5 | 3 | 1375898.13 |
| birds | Myiagra_hebetior | Myiagra_hebetior | 1 | 20.2 | 2 | 342.84 |
| birds | Myiagra_inquieta | Myiagra_inquieta | 0 | 20.9 | 2.8 | 2820032.99 |
| birds | Myiagra_rubecula | Myiagra_rubecula | 0 | 12.1 | 2.4 | 2268381.7 |
| birds | Myiagra_ruficollis | Myiagra_ruficollis | 0 | 10.1 | 2.4 | 580888.4 |
| birds | Myiagra_vanikorensis | Myiagra_vanikorensis | 0 | 12.3 | 1.4 | 18452.63 |
| birds | Myiarchus_antillarum | Myiarchus_antillarum | 0 | 23.4 | 3.9 | 9177.01 |
| birds | Myiarchus_barbirostris | Myiarchus_barbirostris | 0 | 13.4 | 3.5 | 11021.79 |
| birds | Myiarchus_cinerascens | Myiarchus_cinerascens | 0 | 28.2 | 4.5 | 4277784.59 |
| birds | Myiarchus_crinitus | Myiarchus_crinitus | 0 | 32.1 | 5.7 | 6748704.12 |
| birds | Myiarchus_ferox | Myiarchus_ferox | 0 | 27.5 | 2.4 | 11854121.16 |
| birds | Myiarchus_magnirostris | Myiarchus_magnirostris | 0 | 14.7 | 3.9 | 7776.32 |
| birds | Myiarchus_nuttingi | Myiarchus_nuttingi | 0 | 23 | 3.9 | 634487.92 |
| birds | Myiarchus_panamensis | Myiarchus_panamensis | 0 | 31.7 | 2.4 | 370862.95 |
| birds | Myiarchus_sagrae | Myiarchus_sagrae | 0 | 19.3 | 4 | 122416.83 |
| birds | Myiarchus_stolidus | Myiarchus_stolidus | 0 | 20.8 | 3.5 | 86228.49 |
| birds | Myiarchus_swainsoni | Myiarchus_swainsoni | 0 | 25.1 | 3.5 | 13661126.22 |
| birds | Myiarchus_tuberculifer | Myiarchus_tuberculifer | 0 | 17.7 | 3.5 | 10588401.3 |
| birds | Myiarchus_tyrannulus | Myiarchus_tyrannulus | 0 | 35.5 | 3.9 | 10591424.15 |
| birds | Myiarchus_validus | Myiarchus_validus | 0 | 40.6 | 3.9 | 11021.79 |
| birds | Myiobius_atricaudus | Myiobius_atricaudus | 0 | 10 | 2 | 4897288.43 |
| birds | Myiobius_barbatus | Myiobius_barbatus | 0 | 11 | 2 | 6231670.56 |
| birds | Myioborus_flavivertex | Myioborus_flavivertex | 0 | 15 | 2 | 3571.56 |
| birds | Myioborus_melanocephalus | Myioborus_melanocephalus | 0 | 11.3 | 2 | 185791.05 |
| birds | Myioborus_miniatus | Myioborus_miniatus | 0 | 9 | 1.7 | 1254327.4 |
| birds | Myioborus_pictus | Myioborus_pictus | 0 | 9.8 | 4.6 | 1124631.49 |
| birds | Myioborus_torquatus | Myioborus_torquatus | 0 | 10.5 | 2.4 | 5261.72 |
| birds | Myiodynastes_bairdii | Myiodynastes_bairdii | 0 | 45 | 3.9 | 96352.03 |
| birds | Myiodynastes_chrysocephalus | Myiodynastes_chrysocephalus | 0 | 38.3 | 2 | 176225.83 |
| birds | Myiodynastes_hemichrysus | Myiodynastes_hemichrysus | 0 | 43.4 | 3 | 319266.74 |
| birds | Myiodynastes_luteiventris | Myiodynastes_luteiventris | 0 | 46.9 | 2.8 | 2795270.69 |
| birds | Myiodynastes_maculatus | Myiodynastes_maculatus | 0 | 43.2 | 2.8 | 4786616.48 |
| birds | Myiomela_leucura | Cinclidium_leucurum | 0 | 26.8 | 3.2 | 2713438.9 |

| | | | | | | |
|---|---|---|---|---|---|---|
| birds | Myiopagis_cotta | Myiopagis_cotta | 0 | 11.6 | 3 | 11021.79 |
| birds | Myiopagis_viridicata | Myiopagis_viridicata | 0 | 11.5 | 2 | 7381273.67 |
| birds | Myiophobus_fasciatus | Myiophobus_fasciatus | 0 | 9.9 | 1.4 | 10017556.29 |
| birds | Myiornis_atricapillus | Myiornis_atricapillus | 0 | 5.7 | 2 | 184278.95 |
| birds | Myiothlypis_conspicillata | Basileuterus_conspicillatus | 0 | 13.7 | 3.5 | 7320.25 |
| birds | Myiothlypis_flaveola | Basileuterus_flaveolus | 0 | 13.2 | 3 | 4100375.8 |
| birds | Myiothlypis_fulvicauda | Phaeothlypis_fulvicauda | 0 | 14.9 | 2 | 2473113.96 |
| birds | Myiozetetes_cayanensis | Myiozetetes_cayanensis | 0 | 25.9 | 2.4 | 7536061.77 |
| birds | Myiozetetes_granadensis | Myiozetetes_granadensis | 0 | 29.3 | 2.8 | 3322072.64 |
| birds | Myiozetetes_similis | Myiozetetes_similis | 0 | 28 | 2.8 | 10242322.01 |
| birds | Myophonus_caeruleus | Myophonus_caeruleus | 0 | 158 | 3.5 | 7139938.2 |
| birds | Myophonus_glaucinus | Myophonus_glaucinus | 0 | 89.7 | 2 | 24651.37 |
| birds | Myophonus_horsfieldii | Myophonus_horsfieldii | 0 | 117 | 2.4 | 412482.19 |
| birds | Myophonus_robinsoni | Myophonus_robinsoni | 0 | 99.2 | 1.4 | 10816.59 |
| birds | Myopornis_boehmi | Muscicapa_boehmi | 0 | 19.3 | 4 | 863704.75 |
| birds | Myrmeciza_longipes | Myrmeciza_longipes | 0 | 27.9 | 2 | 1110901.77 |
| birds | Myrmecocichla_arnotti | Myrmecocichla_arnoti | 0 | 39.6 | 2.8 | 3102387 |
| birds | Myrmecocichla_formicivora | Myrmecocichla_formicivora | 0 | 41.6 | 3.5 | 2080546.72 |
| birds | Myrmecocichla_monticola | Oenanthe_monticola | 0 | 33.1 | 2.8 | 1391804.19 |
| birds | Myrmecocichla_nigra | Myrmecocichla_nigra | 0 | 40.3 | 3.2 | 3505287.84 |
| birds | Myrmelastes_hyperythrus | Myrmeciza_hyperythra | 0 | 41 | 2 | 1642348.58 |
| birds | Myrmia_micrura | Myrmia_micrura | 0 | 2.3 | 2 | 72174.99 |
| birds | Myrmoborus_leucophrys | Myrmoborus_leucophrys | 0 | 21.5 | 2 | 5127312.13 |
| birds | Myrmoborus_myotherinus | Myrmoborus_myotherinus | 0 | 18.8 | 2 | 4778353.33 |
| birds | Myrmoderus_ferrugineus | Myrmeciza_ferruginea | 0 | 26.1 | 2 | 1290324.22 |
| birds | Myrmoderus_loricatus | Myrmeciza_loricata | 0 | 15.5 | 2 | 177950.43 |
| birds | Myrmoderus_ruficauda | Myrmeciza_ruficauda | 1 | 26 | 2 | 129646.48 |
| birds | Myrmorchilus_strigilatus | Myrmorchilus_strigilatus | 0 | 23 | 2 | 1728828.97 |
| birds | Myrmornis_torquata | Myrmornis_torquata | 0 | 46.7 | 1 | 2996202.86 |
| birds | Myrmothera_simplex | Myrmothera_simplex | 0 | 52.4 | 2 | 113218.8 |
| birds | Myrmotherula_axillaris | Myrmotherula_axillaris | 0 | 8.1 | 2 | 7734412.17 |
| birds | Myrmotherula_schisticolor | Myrmotherula_schisticolor | 0 | 9.6 | 2 | 534175.27 |
| birds | Myrmotherula_surinamensis | Myrmotherula_surinamensis | 0 | 8.3 | 2 | 1574543.3 |
| birds | Myrtis_fanny | Myrtis_fanny | 0 | 2.3 | 2 | 256471.51 |

| | | | | | | |
|---|---|---|---|---|---|---|
| birds | Myzomela_cardinalis | Myzomela_cardinalis | 0 | 13.8 | 3.9 | 22202.1 |
| birds | Myzomela_erythrocephala | Myzomela_erythrocephala | 0 | 7.8 | 2 | 442219.32 |
| birds | Myzomela_jugularis | Myzomela_jugularis | 0 | 8.7 | 2 | 18250.74 |
| birds | Myzomela_obscura | Myzomela_obscura | 0 | 11.9 | 2 | 763713.2 |
| birds | Myzomela_rosenbergii | Myzomela_rosenbergii | 0 | 10.3 | 2 | 176793.86 |
| birds | Myzomela_sanguinolenta | Myzomela_sanguinolenta | 0 | 8.3 | 2 | 912973.16 |
| birds | Namibornis_herero | Namibornis_herero | 0 | 27.2 | 2 | 174435.57 |
| birds | Nannopterum_auritum | Phalacrocorax_auritus | 0 | 1806.5 | 3.5 | 10596564.38 |
| birds | Nannopterum_brasilianum | Phalacrocorax_brasilianus | 0 | 1239.3 | 3.5 | 18298756.62 |
| birds | Neafrapus_boehmi | Neafrapus_boehmi | 0 | 14.4 | 3 | 1674288.3 |
| birds | Necrosyrtes_monachus | Necrosyrtes_monachus | 1 | 2043 | 1 | 14512300.57 |
| birds | Nectarinia_bocagii | Nectarinia_bocagii | 0 | 14.8 | 2 | 114260.42 |
| birds | Nectarinia_famosa | Nectarinia_famosa | 0 | 14.4 | 2 | 1109036.73 |
| birds | Nectarinia_johnstoni | Nectarinia_johnstoni | 0 | 15.2 | 1.4 | 21310.45 |
| birds | Nectarinia_kilimensis | Nectarinia_kilimensis | 0 | 15.7 | 1.4 | 511131.66 |
| birds | Nectarinia_purpureiventris | Nectarinia_purpureiventris | 0 | 11.5 | 1.4 | 70154.05 |
| birds | Nectarinia_tacazze | Nectarinia_tacazze | 0 | 14.9 | 1.4 | 285225.05 |
| birds | Nemosia_pileata | Nemosia_pileata | 0 | 16 | 2 | 8991826.24 |
| birds | Neochen_jubata | Neochen_jubata | 0 | 1250 | 7.7 | 7934083.33 |
| birds | Neochmia_modesta | Neochmia_modesta | 0 | 12.7 | 5.5 | 1594602.22 |
| birds | Neochmia_phaeton | Neochmia_phaeton | 0 | 10 | 6.3 | 1056985.14 |
| birds | Neochmia_ruficauda | Neochmia_ruficauda | 0 | 11.1 | 4.2 | 1490192.67 |
| birds | Neochmia_temporalis | Neochmia_temporalis | 0 | 11.4 | 4.9 | 1100465.39 |
| birds | Neocossyphus_rufus | Neocossyphus_rufus | 0 | 65.9 | 2 | 1123680.47 |
| birds | Neocrex_erythrops | Neocrex_erythrops | 0 | 58.3 | 4.6 | 3788666.38 |
| birds | Neolestes_torquatus | Neolestes_torquatus | 0 | 22.5 | 2 | 813954.89 |
| birds | Neomixis_striatigula | Neomixis_striatigula | 0 | 8.3 | 3.9 | 162877.99 |
| birds | Neomixis_tenella | Neomixis_tenella | 0 | 6.9 | 2 | 586347.22 |
| birds | Neomorphus_geoffroyi | Neomorphus_geoffroyi | 1 | 355 | 1 | 3567739.22 |
| birds | Neophedina_cincta | Riparia_cincta | 0 | 23.4 | 3.2 | 11447865.23 |
| birds | Neophema_chrysogaster | Neophema_chrysogaster | 1 | 45.9 | 4.2 | 32591.23 |
| birds | Neophema_chrysostoma | Neophema_chrysostoma | 1 | 46.1 | 4.9 | 1211024.38 |
| birds | Neophema_elegans | Neophema_elegans | 0 | 43.4 | 4.5 | 865735.95 |
| birds | Neophema_petrophila | Neophema_petrophila | 0 | 53.2 | 4.5 | 142134.67 |

| | | | | | | |
|---|---|---|---|---|---|---|
| birds | Neophema_pulchella | Neophema_pulchella | 0 | 41.4 | 4.5 | 492396.14 |
| birds | Neophema_splendida | Neophema_splendida | 0 | 38.6 | 3.9 | 262252.22 |
| birds | Neophron_percnopterus | Neophron_percnopterus | 1 | 2082 | 2 | 28188514.18 |
| birds | Neopsephotus_bourkii | Neopsephotus_bourkii | 0 | 44.4 | 4.2 | 1717936.07 |
| birds | Neopsittacus_musschenbroekii | Neopsittacus_musschenbroekii | 0 | 50.9 | 2 | 229336.72 |
| birds | Neopsittacus_pullicauda | Neopsittacus_pullicauda | 0 | 33.2 | 2 | 114873.52 |
| birds | Neosericornis_citreogularis | Sericornis_citreogularis | 0 | 17.6 | 3 | 169010.51 |
| birds | Neotis_denhami | Neotis_denhami | 0 | 3970.2 | 1.4 | 7686065.24 |
| birds | Neotis_heuglinii | Neotis_heuglinii | 0 | 4098.8 | 2 | 989206.1 |
| birds | Neotis_ludwigii | Neotis_ludwigii | 1 | 3260.9 | 2 | 1006199.57 |
| birds | Neotis_nuba | Neotis_nuba | 0 | 5440 | 2 | 2385968.92 |
| birds | Neoxolmis_rufiventris | Neoxolmis_rufiventris | 0 | 77 | 2.4 | 1290844.79 |
| birds | Nesillas_typica | Nesillas_typica | 0 | 19.7 | 2 | 520534.89 |
| birds | Nesocharis_ansorgei | Nesocharis_ansorgei | 0 | 8.4 | 3 | 157481.36 |
| birds | Nesoctites_micromegas | Nesoctites_micromegas | 0 | 32.2 | 2.8 | 74799.65 |
| birds | Nesoenas_mayeri | Nesoenas_mayeri | 1 | 302.8 | 2 | 1873.24 |
| birds | Nesoenas_picturatus | Nesoenas_picturata | 0 | 176.8 | 2 | 598530.92 |
| birds | Nesophlox_evelynae | Calliphlox_evelynae | 0 | 2.4 | 2 | 12175.66 |
| birds | Nesoptilotis_flavicollis | Lichenostomus_flavicollis | 0 | 28.7 | 2.4 | 66098.27 |
| birds | Nesoptilotis_leucotis | Lichenostomus_leucotis | 0 | 23.9 | 2 | 1974990 |
| birds | Netta_erythrophthalma | Netta_erythrophthalma | 0 | 780.4 | 8.7 | 10612856.99 |
| birds | Netta_peposaca | Netta_peposaca | 0 | 1000 | 10 | 5216896.73 |
| birds | Netta_rufina | Netta_rufina | 0 | 1118 | 8.9 | 8920116.42 |
| birds | Nettapus_auritus | Nettapus_auritus | 0 | 266 | 8.5 | 17754493.4 |
| birds | Nettapus_coromandelianus | Nettapus_coromandelianus | 0 | 391.3 | 9.8 | 9230293.27 |
| birds | Nettapus_pulchellus | Nettapus_pulchellus | 0 | 307 | 9.8 | 1855537.36 |
| birds | Nicator_chloris | Nicator_chloris | 0 | 46.1 | 1 | 3969589.79 |
| birds | Nicator_gularis | Nicator_gularis | 0 | 45.6 | 2.4 | 940421.62 |
| birds | Nicator_vireo | Nicator_vireo | 0 | 23.6 | 2 | 1466486.16 |
| birds | Nicopsitta_columboides | Psittacula_columboides | 0 | 87 | 4.5 | 121366.21 |
| birds | Nigrita_bicolor | Nigrita_bicolor | 0 | 10.5 | 2 | 3277545.3 |
| birds | Nigrita_canicapillus | Nigrita_canicapillus | 0 | 18.7 | 4.9 | 3831512 |
| birds | Nigrita_fusconotus | Nigrita_fusconotus | 0 | 9.1 | 4.2 | 2969480.86 |
| birds | Nilaus_afer | Nilaus_afer | 0 | 19.1 | 2 | 10555427.15 |

| | | | | | | |
|---|---|---|---|---|---|---|
| birds | Niltava_grandis | Niltava_grandis | 0 | 30.3 | 4 | 1622470.96 |
| birds | Niltava_macgrigoriae | Niltava_macgrigoriae | 0 | 12 | 4 | 1821884.06 |
| birds | Niltava_sundara | Niltava_sundara | 0 | 21.1 | 4 | 3184561.71 |
| birds | Ninox_boobook | Ninox_boobook | 0 | 282.2 | 3 | 7661995.2 |
| birds | Ninox_connivens | Ninox_connivens | 0 | 578 | 2.4 | 2759461.25 |
| birds | Ninox_novaeseelandiae | Ninox_novaeseelandiae | 0 | 183.4 | 2 | 269367.33 |
| birds | Ninox_rufa | Ninox_rufa | 0 | 931.7 | 2 | 824417.84 |
| birds | Ninox_scutulata | Ninox_scutulata | 0 | 191 | 3.2 | 5215181.31 |
| birds | Ninox_strenua | Ninox_strenua | 0 | 1346 | 2 | 573814.15 |
| birds | Ninox_theomacha | Ninox_theomacha | 0 | 125 | 2 | 596413.95 |
| birds | Nipponia_nippon | Nipponia_nippon | 1 | 1900 | 3 | 1119.51 |
| birds | Northiella_haematogaster | Northiella_haematogaster | 0 | 85.9 | 5.3 | 1732521.3 |
| birds | Notharchus_pectoralis | Notharchus_pectoralis | 0 | 69.1 | 3 | 197997.21 |
| birds | Notharchus_tectus | Notharchus_tectus | 0 | 26.8 | 2 | 5067701.36 |
| birds | Nothocercus_bonapartei | Nothocercus_bonapartei | 0 | 763 | 4.5 | 214864.07 |
| birds | Nothoprocta_cinerascens | Nothoprocta_cinerascens | 0 | 523.9 | 9.4 | 1208917.27 |
| birds | Nothoprocta_ornata | Nothoprocta_ornata | 0 | 619.3 | 6 | 765344.43 |
| birds | Nothoprocta_pentlandii | Nothoprocta_pentlandii | 0 | 300.7 | 6.3 | 612683.69 |
| birds | Nothoprocta_perdicaria | Nothoprocta_perdicaria | 0 | 458 | 6.3 | 118759.01 |
| birds | Nothura_boraquira | Nothura_boraquira | 0 | 283 | 4 | 1348296.47 |
| birds | Nothura_darwinii | Nothura_darwinii | 0 | 244.6 | 4.9 | 1098896.66 |
| birds | Nothura_maculosa | Nothura_maculosa | 0 | 257.5 | 4.9 | 5645223.73 |
| birds | Notopholia_corusca | Lamprotornis_corruscus | 0 | 56.1 | 2.8 | 495997.97 |
| birds | Nucifraga_caryocatactes | Nucifraga_caryocatactes | 0 | 182.5 | 3.5 | 15486182.22 |
| birds | Nucifraga_columbiana | Nucifraga_columbiana | 0 | 129.8 | 3 | 2287629.4 |
| birds | Numenius_americanus | Numenius_americanus | 0 | 583.9 | 4 | 3937491.93 |
| birds | Numenius_arquata | Numenius_arquata | 0 | 803 | 4 | 42784794.19 |
| birds | Numenius_borealis | Numenius_borealis | 1 | 350.1 | 3.5 | 2235504.86 |
| birds | Numenius_madagascariensis | Numenius_madagascariensis | 1 | 792 | 4 | 6598384.14 |
| birds | Numenius_minutus | Numenius_minutus | 0 | 173 | 4 | 1472149.77 |
| birds | Numenius_phaeopus | Numenius_phaeopus | 0 | 364.6 | 4 | 17013389.57 |
| birds | Numenius_tahitiensis | Numenius_tahitiensis | 0 | 400.1 | 4 | 102179.88 |
| birds | Numenius_tenuirostris | Numenius_tenuirostris | 1 | 308 | 4 | 887016.77 |
| birds | Numida_meleagris | Numida_meleagris | 0 | 1299 | 8.5 | 13597695.41 |

| | | | | | | |
|---|---|---|---|---|---|---|
| birds | Nycticorax_caledonicus | Nycticorax_caledonicus | 0 | 856 | 3.2 | 7791277.4 |
| birds | Nycticorax_nycticorax | Nycticorax_nycticorax | 0 | 810 | 3.9 | 52753396.92 |
| birds | Nycticryphes_semicollaris | Rostratula_semicollaris | 0 | 76.6 | 2 | 2533948.28 |
| birds | Nyctidromus_albicollis | Nyctidromus_albicollis | 0 | 57.8 | 1.4 | 13748689.24 |
| birds | Nyctidromus_anthonyi | Caprimulgus_anthonyi | 0 | 35.9 | 1.4 | 57328.24 |
| birds | Nyctiphrynus_mcleodii | Nyctiphrynus_mcleodii | 0 | 35 | 2 | 188114.4 |
| birds | Nyctiphrynus_ocellatus | Nyctiphrynus_ocellatus | 0 | 39.6 | 2 | 3398673.41 |
| birds | Nyctipolus_nigrescens | Caprimulgus_nigrescens | 0 | 35.7 | 1 | 4123782.88 |
| birds | Nyctyornis_amictus | Nyctyornis_amictus | 0 | 71.9 | 3.9 | 1492016.66 |
| birds | Nymphicus_hollandicus | Nymphicus_hollandicus | 0 | 94.6 | 4.6 | 6462512.62 |
| birds | Nystactes_tamatia | Bucco_tamatia | 0 | 35.5 | 2 | 4349445.92 |
| birds | Nystalus_chacuru | Nystalus_chacuru | 0 | 52.3 | 2.8 | 4329404.4 |
| birds | Nystalus_maculatus | Nystalus_maculatus | 0 | 42 | 2.4 | 3643937.66 |
| birds | Nystalus_striolatus | Nystalus_striolatus | 0 | 47 | 4 | 2922975.07 |
| birds | Ochetorhynchus_andaecola | Ochetorhynchus_andaecola | 0 | 30.9 | 2 | 398226.84 |
| birds | Ochetorhynchus_melanurus | Ochetorhynchus_melanura | 0 | 40 | 3.5 | 39429.48 |
| birds | Ochetorhynchus_phoenicurus | Ochetorhynchus_phoenicurus | 0 | 30.2 | 2.8 | 525155.51 |
| birds | Ochetorhynchus_ruficaudus | Ochetorhynchus_ruficaudus | 0 | 36.3 | 2 | 1066359.63 |
| birds | Ochthoeca_leucophrys | Ochthoeca_leucophrys | 0 | 13.1 | 1 | 781892.76 |
| birds | Ochthornis_littoralis | Ochthornis_littoralis | 0 | 13.4 | 3.5 | 4562555.95 |
| birds | Ocreatus_underwoodii | Ocreatus_underwoodii | 0 | 3 | 2 | 385599.41 |
| birds | Ocyphaps_lophotes | Ocyphaps_lophotes | 0 | 204 | 2 | 7382616.53 |
| birds | Odontophorus_capueira | Odontophorus_capueira | 0 | 425.4 | 5 | 1724256.7 |
| birds | Odontophorus_columbianus | Odontophorus_columbianus | 0 | 340 | 6 | 5025.3 |
| birds | Odontophorus_gujanensis | Odontophorus_gujanensis | 0 | 314.1 | 4 | 7718738.67 |
| birds | Odontophorus_leucolaemus | Odontophorus_leucolaemus | 0 | 295 | 5 | 11852.57 |
| birds | Odontophorus_melanotis | Odontophorus_melanotis | 0 | 335 | 4 | 128032.3 |
| birds | Oedistoma_iliolophus | Toxorhamphus_iliolophus | 0 | 11 | 1 | 624407.83 |
| birds | Oena_capensis | Oena_capensis | 0 | 40.6 | 2 | 17910218.79 |
| birds | Oenanthe_albifrons | Myrmecocichla_albifrons | 0 | 20.6 | 2.4 | 3294680.42 |
| birds | Oenanthe_albonigra | Oenanthe_albonigra | 0 | 25.2 | 4.5 | 1325675.77 |
| birds | Oenanthe_bottae | Oenanthe_bottae | 0 | 19.9 | 3 | 116654.28 |
| birds | Oenanthe_cypriaca | Oenanthe_cypriaca | 0 | 16.6 | 4.9 | 1608130.43 |
| birds | Oenanthe_deserti | Oenanthe_deserti | 0 | 19.8 | 4.5 | 18380455.15 |

| | | | | | | |
|---|---|---|---|---|---|---|
| birds | Oenanthe_familiaris | Cercomela_familiaris | 0 | 19.5 | 3 | 7293437.67 |
| birds | Oenanthe_finschii | Oenanthe_finschii | 0 | 27.4 | 5 | 3778500.05 |
| birds | Oenanthe_hispanica | Oenanthe_hispanica | 0 | 15 | 4.5 | 18516204.75 |
| birds | Oenanthe_isabellina | Oenanthe_isabellina | 0 | 29.4 | 5.5 | 23687917.96 |
| birds | Oenanthe_leucopyga | Oenanthe_leucopyga | 0 | 27.9 | 3.9 | 5398249.87 |
| birds | Oenanthe_leucura | Oenanthe_leucura | 0 | 41 | 3.9 | 1288299.84 |
| birds | Oenanthe_lugens | Oenanthe_lugens | 0 | 22.4 | 4.5 | 3562658.08 |
| birds | Oenanthe_melanura | Cercomela_melanura | 0 | 15.2 | 3.5 | 2174337.34 |
| birds | Oenanthe_moesta | Oenanthe_moesta | 0 | 30.2 | 4.5 | 987108.26 |
| birds | Oenanthe_monacha | Oenanthe_monacha | 0 | 21.6 | 3 | 938292.16 |
| birds | Oenanthe_oenanthe | Oenanthe_oenanthe | 0 | 25.4 | 5.3 | 39231050.36 |
| birds | Oenanthe_phillipsi | Oenanthe_phillipsi | 0 | 19.6 | 3.5 | 557674.64 |
| birds | Oenanthe_pileata | Oenanthe_pileata | 0 | 26.8 | 3.5 | 5997462.33 |
| birds | Oenanthe_pleschanka | Oenanthe_pleschanka | 0 | 19.4 | 4.9 | 12225171.53 |
| birds | Oenanthe_xanthoprymna | Oenanthe_xanthoprymna | 0 | 22.7 | 4.9 | 3804462.03 |
| birds | Ognorhynchus_icterotis | Ognorhynchus_icterotis | 1 | 285 | 4 | 146924.81 |
| birds | Oncostoma_cinereigulare | Oncostoma_cinereigulare | 0 | 6.4 | 2 | 588153.9 |
| birds | Oncostoma_olivaceum | Oncostoma_olivaceum | 0 | 6.6 | 2 | 226322.4 |
| birds | Oneillornis_salvini | Gymnopithys_salvini | 0 | 25.9 | 2 | 1355133.04 |
| birds | Onychognathus_albirostris | Onychognathus_albirostris | 0 | 137.3 | 2.4 | 162747 |
| birds | Onychognathus_blythii | Onychognathus_blythii | 0 | 100 | 4.5 | 440734.83 |
| birds | Onychognathus_morio | Onychognathus_morio | 0 | 132.9 | 3 | 3660528.38 |
| birds | Onychognathus_nabouroup | Onychognathus_nabouroup | 0 | 102.4 | 3.2 | 984106.61 |
| birds | Onychognathus_tenuirostris | Onychognathus_tenuirostris | 0 | 118 | 2.8 | 405677.73 |
| birds | Onychognathus_tristramii | Onychognathus_tristramii | 0 | 122.7 | 2.8 | 564513.21 |
| birds | Onychognathus_walleri | Onychognathus_walleri | 0 | 93 | 1.7 | 308237.32 |
| birds | Onychorhynchus_coronatus | Onychorhynchus_coronatus | 0 | 14 | 2 | 5973721.86 |
| birds | Oporornis_agilis | Oporornis_agilis | 0 | 13.3 | 4.2 | 6641240.31 |
| birds | Orchesticus_abeillei | Orchesticus_abeillei | 0 | 31.5 | 2 | 163525.7 |
| birds | Oreoica_gutturalis | Oreoica_gutturalis | 0 | 62.8 | 3.5 | 4957954.29 |
| birds | Oreolais_pulcher | Oreolais_pulchra | 0 | 8.4 | 2.4 | 100450.2 |
| birds | Oreolais_ruwenzorii | Oreolais_ruwenzorii | 0 | 9.9 | 2 | 123324.79 |
| birds | Oreophilais_robertsi | Prinia_robertsi | 0 | 10 | 2.4 | 9712.82 |
| birds | Oreopsar_bolivianus | Oreopsar_bolivianus | 0 | 72.9 | 3 | 77402.17 |

| | | | | | | |
|---|---|---|---|---|---|---|
| birds | Oreortyx_pictus | Oreortyx_pictus | 0 | 233 | 9.5 | 386114.5 |
| birds | Oreoscoptes_montanus | Oreoscoptes_montanus | 0 | 44.2 | 4.5 | 3989086.93 |
| birds | Oreoscopus_gutturalis | Oreoscopus_gutturalis | 1 | 20 | 2 | 5869.25 |
| birds | Oreothlypis_gutturalis | Parula_gutturalis | 0 | 9.5 | 2 | 3913.07 |
| birds | Oreothlypis_superciliosa | Parula_superciliosa | 0 | 9 | 2.4 | 402973.46 |
| birds | Oreotrochilus_chimborazo | Oreotrochilus_chimborazo | 0 | 8.2 | 2 | 36894.51 |
| birds | Oreotrochilus_estella | Oreotrochilus_estella | 0 | 8.1 | 2 | 595797.12 |
| birds | Oreotrochilus_leucopleurus | Oreotrochilus_leucopleurus | 0 | 8.2 | 2 | 404047.64 |
| birds | Oreotrochilus_melanogaster | Oreotrochilus_melanogaster | 0 | 9.1 | 2 | 55336.69 |
| birds | Origma_solitaria | Origma_solitaria | 0 | 14.3 | 3 | 56963.22 |
| birds | Oriolus_auratus | Oriolus_auratus | 0 | 66.2 | 1.7 | 10077113.98 |
| birds | Oriolus_brachyrynchus | Oriolus_brachyrhynchus | 0 | 49 | 2 | 2640549.2 |
| birds | Oriolus_chinensis | Oriolus_chinensis | 0 | 82.6 | 2 | 8169313.16 |
| birds | Oriolus_flavocinctus | Oriolus_flavocinctus | 0 | 109 | 2.4 | 535100.57 |
| birds | Oriolus_larvatus | Oriolus_larvatus | 0 | 66.1 | 2 | 6081428.22 |
| birds | Oriolus_oriolus | Oriolus_oriolus | 0 | 79 | 3.5 | 33861421.14 |
| birds | Oriolus_sagittatus | Oriolus_sagittatus | 0 | 95.2 | 2.8 | 3411405.87 |
| birds | Oriolus_szalayi | Oriolus_szalayi | 0 | 95.2 | 1.4 | 680368.63 |
| birds | Oriolus_tenuirostris | Oriolus_tenuirostris | 0 | 81.4 | 2.8 | 1127097.86 |
| birds | Oriolus_traillii | Oriolus_traillii | 0 | 73.7 | 2.4 | 1982675.5 |
| birds | Oriolus_xanthonotus | Oriolus_xanthonotus | 0 | 45 | 2 | 1443272.17 |
| birds | Oriolus_xanthornus | Oriolus_xanthornus | 0 | 56.3 | 2.8 | 4355542.08 |
| birds | Oriturus_superciliosus | Oriturus_superciliosus | 0 | 41.4 | 3.5 | 295047.68 |
| birds | Orochelidon_murina | Notiochelidon_murina | 0 | 11.7 | 2.4 | 425662.43 |
| birds | Ortalis_canicollis | Ortalis_canicollis | 0 | 539 | 3.5 | 1171806.84 |
| birds | Ortalis_cinereiceps | Ortalis_cinereiceps | 0 | 493 | 3 | 236334.32 |
| birds | Ortalis_erythroptera | Ortalis_erythroptera | 1 | 632 | 3 | 74565.61 |
| birds | Ortalis_garrula | Ortalis_garrula | 0 | 532 | 3 | 109708.02 |
| birds | Ortalis_leucogastra | Ortalis_leucogastra | 0 | 495.8 | 2.4 | 57398.42 |
| birds | Ortalis_motmot | Ortalis_motmot | 0 | 520 | 3 | 2134685.16 |
| birds | Ortalis_ruficauda | Ortalis_ruficauda | 0 | 586.2 | 3.5 | 662199.34 |
| birds | Ortalis_vetula | Ortalis_vetula | 0 | 562.6 | 3 | 603428.43 |
| birds | Ortalis_wagleri | Ortalis_wagleri | 0 | 834 | 3 | 114061.43 |
| birds | Orthonyx_spaldingii | Orthonyx_spaldingii | 0 | 149.9 | 1 | 23436.79 |

| | | | | | | |
|---|---|---|---|---|---|---|
| birds | Orthonyx_temminckii | Orthonyx_temminckii | 0 | 58.1 | 1.4 | 166548.28 |
| birds | Orthopsittaca_manilatus | Orthopsittaca_manilata | 0 | 372 | 2 | 7148717.74 |
| birds | Orthorhyncus_cristatus | Orthorhyncus_cristatus | 0 | 3 | 2 | 7736.3 |
| birds | Orthotomus_atrogularis | Orthotomus_atrogularis | 0 | 7.7 | 3.2 | 2752593.84 |
| birds | Orthotomus_castaneiceps | Orthotomus_castaneiceps | 0 | 12.8 | 2.4 | 33106.02 |
| birds | Orthotomus_frontalis | Orthotomus_frontalis | 0 | 8 | 2.4 | 120551.54 |
| birds | Orthotomus_ruficeps | Orthotomus_ruficeps | 0 | 7 | 2.4 | 887360.64 |
| birds | Orthotomus_sericeus | Orthotomus_sericeus | 0 | 10.8 | 2.4 | 1068502.74 |
| birds | Orthotomus_sutorius | Orthotomus_sutorius | 0 | 7.5 | 3.5 | 6849946.15 |
| birds | Ortygornis_gularis | Francolinus_gularis | 1 | 510 | 3.5 | 362104.24 |
| birds | Ortygornis_pondicerianus | Francolinus_pondicerianus | 0 | 249.9 | 7.3 | 3601349.63 |
| birds | Ortygornis_sephaena | Francolinus_sephaena | 0 | 340.5 | 6 | 3350686.8 |
| birds | Ortygospiza_atricollis | Ortygospiza_atricollis | 0 | 10.6 | 4.9 | 3231575.56 |
| birds | Otidiphaps_nobilis | Otidiphaps_nobilis | 0 | 500 | 1 | 260128.69 |
| birds | Otis_tarda | Otis_tarda | 1 | 6759.9 | 2.4 | 10008660.6 |
| birds | Otocichla_mupinensis | Turdus_mupinensis | 0 | 62.5 | 2.8 | 1082838.83 |
| birds | Otus_bakkamoena | Otus_bakkamoena | 0 | 138.5 | 3.9 | 2947848.96 |
| birds | Otus_brucei | Otus_brucei | 0 | 84.4 | 4.9 | 1794970.43 |
| birds | Otus_lempiji | Otus_lempiji | 0 | 115 | 3.9 | 1486775.34 |
| birds | Otus_lettia | Otus_lettia | 0 | 139 | 3.9 | 4204562.9 |
| birds | Otus_rutilus | Otus_rutilus | 0 | 101 | 4.5 | 590836.08 |
| birds | Otus_sagittatus | Otus_sagittatus | 1 | 122.6 | 3.5 | 148393.3 |
| birds | Otus_scops | Otus_scops | 0 | 92 | 3.5 | 26192272.39 |
| birds | Otus_semitorques | Otus_semitorques | 0 | 121.5 | 3.9 | 1982105.07 |
| birds | Otus_senegalensis | Otus_senegalensis | 0 | 67 | 2.8 | 13485667.89 |
| birds | Otus_spilocephalus | Otus_spilocephalus | 0 | 67.5 | 2.4 | 3350281.15 |
| birds | Otus_sunia | Otus_sunia | 0 | 78.1 | 4.2 | 8012147.55 |
| birds | Oxylabes_madagascariensis | Oxylabes_madagascariensis | 0 | 23.3 | 2 | 172738.74 |
| birds | Oxyruncus_cristatus | Oxyruncus_cristatus | 0 | 42 | 2 | 1872045.95 |
| birds | Oxyura_australis | Oxyura_australis | 0 | 798 | 5.5 | 3286126.17 |
| birds | Oxyura_jamaicensis | Oxyura_jamaicensis | 0 | 608.2 | 7.7 | 12454394.74 |
| birds | Oxyura_leucocephala | Oxyura_leucocephala | 1 | 661.1 | 6.3 | 10904918.6 |
| birds | Oxyura_maccoa | Oxyura_maccoa | 1 | 674 | 5.7 | 1133393.56 |
| birds | Oxyura_vittata | Oxyura_vittata | 0 | 642.7 | 3.9 | 3138365.75 |

| | | | | | | |
|---|---|---|---|---|---|---|
| birds | Pachycare_flavogriseum | Pachycare_flavogriseum | 0 | 16.2 | 2 | 171110.44 |
| birds | Pachycephala_cinerea | Pachycephala_grisola | 0 | 21 | 2 | 1077022.85 |
| birds | Pachycephala_hyperythra | Pachycephala_hyperythra | 0 | 27.7 | 2 | 103362.37 |
| birds | Pachycephala_inornata | Pachycephala_inornata | 0 | 30.8 | 2.8 | 1593660.36 |
| birds | Pachycephala_lanioides | Pachycephala_lanioides | 0 | 38.7 | 1.4 | 255698.63 |
| birds | Pachycephala_melanura | Pachycephala_melanura | 0 | 23.8 | 2.4 | 524241.76 |
| birds | Pachycephala_olivacea | Pachycephala_olivacea | 0 | 41.2 | 2.4 | 298358.81 |
| birds | Pachycephala_pectoralis | Pachycephala_pectoralis | 0 | 30.4 | 1.4 | 1963546.27 |
| birds | Pachycephala_philippinensis | Pachycephala_philippinensis | 0 | 22.3 | 2.4 | 193262.3 |
| birds | Pachycephala_rufiventris | Pachycephala_rufiventris | 0 | 22.5 | 2.4 | 7227395.75 |
| birds | Pachycephala_rufogularis | Pachycephala_rufogularis | 1 | 36.6 | 2.4 | 133566.32 |
| birds | Pachycephala_schlegelii | Pachycephala_schlegelii | 0 | 22.7 | 2 | 132622.24 |
| birds | Pachycephala_simplex | Pachycephala_simplex | 0 | 17.4 | 2 | 93200.99 |
| birds | Pachycephala_soror | Pachycephala_soror | 0 | 24.8 | 1.4 | 221930.33 |
| birds | Pachycephalopsis_poliosoma | Pachycephalopsis_poliosoma | 0 | 38 | 1 | 184655.43 |
| birds | Pachyramphus_aglaiae | Pachyramphus_aglaiae | 0 | 29.7 | 3.5 | 1345543.81 |
| birds | Pachyramphus_cinnamomeus | Pachyramphus_cinnamomeus | 0 | 20.3 | 3.5 | 701563.57 |
| birds | Pachyramphus_minor | Pachyramphus_minor | 0 | 36.6 | 4.9 | 6895984.62 |
| birds | Pachyramphus_niger | Pachyramphus_niger | 0 | 40.2 | 3 | 11021.79 |
| birds | Pachyramphus_polychopterus | Pachyramphus_polychopterus | 0 | 20.8 | 2.8 | 13935213.9 |
| birds | Pachyramphus_rufus | Pachyramphus_rufus | 0 | 18.8 | 3.2 | 3994359.03 |
| birds | Pachyramphus_validus | Pachyramphus_validus | 0 | 43 | 4 | 6063334.02 |
| birds | Pachyramphus_versicolor | Pachyramphus_versicolor | 0 | 15.5 | 2.4 | 302157.64 |
| birds | Pachyramphus_viridis | Pachyramphus_viridis | 0 | 21 | 2.8 | 4959688.09 |
| birds | Padda_fuscata | Padda_fuscata | 0 | 20.7 | 4.2 | 24603.86 |
| birds | Padda_oryzivora | Padda_oryzivora | 1 | 24.8 | 4.9 | 960325.86 |
| birds | Palaeornis_eupatria | Psittacula_eupatria | 0 | 214 | 3.5 | 3231032.2 |
| birds | Pampa_rufa | Campylopterus_rufus | 0 | 7.1 | 2 | 31219.99 |
| birds | Pampusana_beccarii | Gallicolumba_beccarii | 0 | 78.7 | 1 | 167246.95 |
| birds | Pampusana_jobiensis | Gallicolumba_jobiensis | 0 | 135 | 2 | 647409.7 |
| birds | Pampusana_stairi | Gallicolumba_stairi | 1 | 171 | 1.4 | 21209.63 |
| birds | Pandion_haliaetus | Pandion_haliaetus | 0 | 1483.2 | 3 | 91738111.45 |
| birds | Panterpe_insignis | Panterpe_insignis | 0 | 5.7 | 2 | 8271.57 |
| birds | Panurus_biarmicus | Panurus_biarmicus | 0 | 13.8 | 5.7 | 12154631.85 |

| | | | | | | |
|---|---|---|---|---|---|---|
| birds | Parabuteo_leucorrhous | Buteo_leucorrhous | 0 | 290 | 2.4 | 1189248.19 |
| birds | Parabuteo_unicinctus | Parabuteo_unicinctus | 0 | 850.3 | 2.8 | 9015595.35 |
| birds | Paraclaravis_geoffroyi | Claravis_godefrida | 1 | 43.5 | 2 | 612044.34 |
| birds | Paragallinula_angulata | Gallinula_angulata | 0 | 130 | 5.2 | 11738770.12 |
| birds | Pardaliparus_venustulus | Parus_venustulus | 0 | 10.6 | 5.9 | 1942840.09 |
| birds | Pardalotus_punctatus | Pardalotus_punctatus | 0 | 8.7 | 4.2 | 2170812.26 |
| birds | Pardalotus_quadragintus | Pardalotus_quadragintus | 1 | 10.7 | 3.9 | 1433.3 |
| birds | Pardalotus_rubricatus | Pardalotus_rubricatus | 0 | 10.9 | 3.5 | 5191015.83 |
| birds | Pardipicus_caroli | Campethera_caroli | 0 | 44.2 | 2.4 | 3849726.47 |
| birds | Pardipicus_nivosus | Campethera_nivosa | 0 | 37.8 | 2 | 3825842.41 |
| birds | Pardirallus_maculatus | Pardirallus_maculatus | 0 | 171 | 3.7 | 8411430.24 |
| birds | Pardirallus_nigricans | Pardirallus_nigricans | 0 | 217 | 3 | 3254810.23 |
| birds | Pardirallus_sanguinolentus | Pardirallus_sanguinolentus | 0 | 232 | 4.9 | 4579906.01 |
| birds | Parkesia_motacilla | Seiurus_motacilla | 0 | 19.9 | 5 | 4813246.08 |
| birds | Parkesia_noveboracensis | Seiurus_noveboracensis | 0 | 16.3 | 4.5 | 17656885.93 |
| birds | Parmoptila_woodhousei | Parmoptila_woodhousei | 0 | 9.4 | 3.5 | 716141.13 |
| birds | Parus_major | Parus_major | 0 | 16.3 | 7.7 | 32577984.28 |
| birds | Parus_monticolus | Parus_monticolus | 0 | 14 | 5.7 | 1684621.77 |
| birds | Passer_cinnamomeus | Passer_rutilans | 0 | 18.5 | 4.5 | 4964243.4 |
| birds | Passer_domesticus | Passer_domesticus | 0 | 26.5 | 3.7 | 64177419.85 |
| birds | Passer_eminibey | Passer_eminibey | 0 | 13.4 | 3.5 | 549271.11 |
| birds | Passer_flaveolus | Passer_flaveolus | 0 | 19.4 | 2.8 | 1319875.05 |
| birds | Passer_griseus | Passer_griseus | 0 | 27.8 | 3.2 | 10513534.53 |
| birds | Passer_hispaniolensis | Passer_hispaniolensis | 0 | 24.2 | 4.9 | 8060752.71 |
| birds | Passer_luteus | Passer_luteus | 0 | 11.2 | 3.9 | 2874074.9 |
| birds | Passer_melanurus | Passer_melanurus | 0 | 29.5 | 3.5 | 1765360.17 |
| birds | Passer_moabiticus | Passer_moabiticus | 0 | 16.5 | 4.5 | 565850.92 |
| birds | Passer_montanus | Passer_montanus | 0 | 21.4 | 3.7 | 39493440.33 |
| birds | Passer_motitensis | Passer_motitensis | 0 | 30.2 | 3.2 | 1648538.38 |
| birds | Passer_simplex | Passer_simplex | 0 | 19.5 | 3.2 | 660184.35 |
| birds | Passerculus_bairdii | Ammodramus_bairdii | 0 | 18.1 | 4.2 | 2235509.58 |
| birds | Passerculus_henslowii | Ammodramus_henslowii | 0 | 12.8 | 3.9 | 2912755.7 |
| birds | Passerculus_sandwichensis | Passerculus_sandwichensis | 0 | 20 | 4.5 | 19177797.53 |
| birds | Passerella_arborea | Spizella_arborea | 0 | 17.8 | 5 | 14358243.45 |

| | | | | | | |
|---|---|---|---|---|---|---|
| birds | Passerella_iliaca | Passerella_iliaca | 0 | 33.3 | 3.9 | 12314409.99 |
| birds | Passerina_amoena | Passerina_amoena | 0 | 15.5 | 3.9 | 4666546.61 |
| birds | Passerina_caerulea | Passerina_caerulea | 0 | 27.4 | 3.5 | 6742464.86 |
| birds | Passerina_ciris | Passerina_ciris | 0 | 15.5 | 3.5 | 3706093.07 |
| birds | Passerina_cyanea | Passerina_cyanea | 0 | 14.7 | 3.5 | 7436442.54 |
| birds | Passerina_versicolor | Passerina_versicolor | 0 | 12.9 | 3.5 | 1540631.55 |
| birds | Pastor_roseus | Sturnus_roseus | 0 | 73 | 4.2 | 7512915.8 |
| birds | Patagioenas_cayennensis | Patagioenas_cayennensis | 0 | 229 | 1 | 13847694.77 |
| birds | Patagioenas_corensis | Patagioenas_corensis | 0 | 272.3 | 1 | 129079.86 |
| birds | Patagioenas_fasciata | Patagioenas_fasciata | 0 | 366.3 | 1.4 | 2745784.92 |
| birds | Patagioenas_flavirostris | Patagioenas_flavirostris | 0 | 270.8 | 1 | 928061.87 |
| birds | Patagioenas_inornata | Patagioenas_inornata | 0 | 365 | 1.4 | 205258.36 |
| birds | Patagioenas_leucocephala | Patagioenas_leucocephala | 0 | 240.1 | 2 | 243969.1 |
| birds | Patagioenas_maculosa | Patagioenas_maculosa | 0 | 347 | 1.4 | 3659053.31 |
| birds | Patagioenas_nigrirostris | Patagioenas_nigrirostris | 0 | 176 | 1 | 352468.53 |
| birds | Patagioenas_picazuro | Patagioenas_picazuro | 0 | 279 | 1 | 6181623.95 |
| birds | Patagioenas_speciosa | Patagioenas_speciosa | 0 | 258.5 | 2 | 9715779.78 |
| birds | Patagioenas_squamosa | Patagioenas_squamosa | 0 | 312 | 1.4 | 202802.2 |
| birds | Patagioenas_subvinacea | Patagioenas_subvinacea | 0 | 162.5 | 1 | 8499137.27 |
| birds | Patagona_gigas | Patagona_gigas | 0 | 20.2 | 2 | 1270749.2 |
| birds | Pedionomus_torquatus | Pedionomus_torquatus | 1 | 62.5 | 4 | 2237290.61 |
| birds | Pelargopsis_amauroptera | Pelargopsis_amauroptera | 0 | 162 | 3.5 | 41842.36 |
| birds | Pelargopsis_capensis | Pelargopsis_capensis | 0 | 178.7 | 3.2 | 5091973.15 |
| birds | Pelecanus_conspicillatus | Pelecanus_conspicillatus | 0 | 5505 | 2 | 6565542.05 |
| birds | Pelecanus_crispus | Pelecanus_crispus | 0 | 9512.1 | 2 | 12289083.63 |
| birds | Pelecanus_erythrorhynchos | Pelecanus_erythrorhynchos | 0 | 5608.5 | 2 | 8104341.01 |
| birds | Pelecanus_occidentalis | Pelecanus_occidentalis | 0 | 3427.8 | 2.4 | 6500402.25 |
| birds | Pelecanus_onocrotalus | Pelecanus_onocrotalus | 0 | 9322.3 | 1.7 | 16249898.89 |
| birds | Pelecanus_philippensis | Pelecanus_philippensis | 0 | 5011 | 3.5 | 3052519.6 |
| birds | Pelecanus_rufescens | Pelecanus_rufescens | 0 | 5419.6 | 1.7 | 14141427.8 |
| birds | Peliperdix_lathami | Francolinus_lathami | 0 | 269 | 2 | 2655327.38 |
| birds | Pellorneum_bicolor | Trichastoma_bicolor | 0 | 27.2 | 2 | 785296.58 |
| birds | Pellorneum_malaccense | Malacocincla_malaccensis | 0 | 21.4 | 2.4 | 1073369.64 |
| birds | Pellorneum_rostratum | Trichastoma_rostratum | 0 | 21 | 2 | 789435.46 |

| | | | | | | |
|---|---|---|---|---|---|---|
| birds | Peltohyas_australis | Charadrius_australis | 0 | 83.5 | 3 | 2556918.74 |
| birds | Penelopides_manillae | Penelopides_manillae | 0 | 460.9 | 4.2 | 104660.9 |
| birds | Penelopides_panini | Penelopides_panini | 1 | 470.5 | 2.4 | 27587.92 |
| birds | Peneoenanthe_pulverulenta | Eopsaltria_pulverulenta | 0 | 19.2 | 2 | 435422.99 |
| birds | Percnostola_rufifrons | Percnostola_rufifrons | 0 | 23.9 | 1.4 | 1139425.12 |
| birds | Perdicula_argoondah | Perdicula_argoondah | 0 | 60.5 | 5.5 | 1182863.73 |
| birds | Perdicula_asiatica | Perdicula_asiatica | 0 | 68.4 | 5.5 | 2187881.27 |
| birds | Perdicula_erythrorhyncha | Perdicula_erythrorhyncha | 0 | 67.6 | 5.3 | 502809.33 |
| birds | Perdicula_manipurensis | Perdicula_manipurensis | 1 | 70.7 | 4 | 137525.14 |
| birds | Perdix_dauurica | Perdix_dauurica | 0 | 263.4 | 19 | 7725915.52 |
| birds | Perdix_hodgsoniae | Perdix_hodgsoniae | 0 | 363.7 | 8.9 | 1000982.76 |
| birds | Perdix_perdix | Perdix_perdix | 0 | 405.3 | 16 | 12799976.79 |
| birds | Pericrocotus_cinnamomeus | Pericrocotus_cinnamomeus | 0 | 8.6 | 3 | 4793425.28 |
| birds | Pericrocotus_erythropygius | Pericrocotus_erythropygius | 0 | 10.8 | 3 | 1571440.41 |
| birds | Pericrocotus_ethologus | Pericrocotus_ethologus | 0 | 19 | 3.5 | 6237997.8 |
| birds | Pericrocotus_flammeus | Pericrocotus_flammeus | 0 | 23.3 | 2.4 | 6801040.97 |
| birds | Pericrocotus_igneus | Pericrocotus_igneus | 0 | 15 | 2 | 1266299.03 |
| birds | Pericrocotus_roseus | Pericrocotus_roseus | 0 | 19.2 | 3.5 | 3379728.88 |
| birds | Pericrocotus_solaris | Pericrocotus_solaris | 0 | 14.5 | 3 | 2947693.85 |
| birds | Periparus_ater | Parus_ater | 0 | 9.2 | 9.4 | 22198675.93 |
| birds | Periparus_rubidiventris | Parus_rubidiventris | 0 | 11.8 | 2.4 | 949945.6 |
| birds | Periparus_rufonuchalis | Parus_rufonuchalis | 0 | 12.6 | 4.9 | 530583.68 |
| birds | Perisoreus_canadensis | Perisoreus_canadensis | 0 | 71.6 | 4 | 7301159.58 |
| birds | Perisoreus_infaustus | Perisoreus_infaustus | 0 | 84.4 | 3.9 | 11840129.85 |
| birds | Perissocephalus_tricolor | Perissocephalus_tricolor | 0 | 338.9 | 1 | 1655180.77 |
| birds | Pernis_apivorus | Pernis_apivorus | 0 | 754.4 | 2 | 32679909.87 |
| birds | Pernis_celebensis | Pernis_celebensis | 0 | 722 | 2 | 155679.51 |
| birds | Pernis_ptilorhynchus | Pernis_ptilorhyncus | 0 | 1141.1 | 2 | 16484980.32 |
| birds | Petrochelidon_ariel | Hirundo_ariel | 0 | 10.8 | 4.5 | 5371208.21 |
| birds | Petrochelidon_fluvicola | Hirundo_fluvicola | 0 | 9.7 | 3.5 | 2700681.6 |
| birds | Petrochelidon_fulva | Petrochelidon_fulva | 0 | 19 | 2.4 | 1053297.21 |
| birds | Petrochelidon_nigricans | Hirundo_nigricans | 0 | 15.4 | 3.9 | 8047367 |
| birds | Petrochelidon_preussi | Hirundo_preussi | 0 | 13.3 | 2.4 | 2226117.98 |
| birds | Petrochelidon_pyrrhonota | Petrochelidon_pyrrhonota | 0 | 21.6 | 2.4 | 26655865.33 |

| | | | | | | |
|---|---|---|---|---|---|---|
| birds | Petrochelidon_rufigula | Hirundo_rufigula | 0 | 15.9 | 2 | 1298041.09 |
| birds | Petrochelidon_spilodera | Hirundo_spilodera | 0 | 20.6 | 3 | 3660516.72 |
| birds | Petroica_australis | Petroica_australis | 0 | 31.3 | 2.8 | 43012.79 |
| birds | Petroica_goodenovii | Petroica_goodenovii | 0 | 8.9 | 2.4 | 6551523.57 |
| birds | Petroica_macrocephala | Petroica_macrocephala | 0 | 11.8 | 3.9 | 204209.11 |
| birds | Petroica_multicolor | Petroica_multicolor | 1 | 11.4 | 3.5 | 37.92 |
| birds | Petroica_phoenicea | Petroica_phoenicea | 0 | 13 | 3.5 | 667819.8 |
| birds | Petroica_rodinogaster | Petroica_rodinogaster | 0 | 10.2 | 3.5 | 273895.48 |
| birds | Petroica_rosea | Petroica_rosea | 0 | 8.2 | 2.4 | 694633.38 |
| birds | Petronia_petronia | Petronia_petronia | 0 | 30.2 | 5.3 | 8115554.53 |
| birds | Petrophassa_albipennis | Petrophassa_albipennis | 0 | 124.3 | 2 | 235719.79 |
| birds | Petrophassa_rufipennis | Petrophassa_rufipennis | 0 | 159 | 2 | 55159.82 |
| birds | Peucaea_aestivalis | Aimophila_aestivalis | 0 | 19.6 | 3.9 | 761850.7 |
| birds | Peucaea_botterii | Aimophila_botterii | 0 | 19.9 | 3.2 | 998316.54 |
| birds | Peucaea_carpalis | Aimophila_carpalis | 0 | 15.3 | 4 | 175983.96 |
| birds | Peucaea_cassinii | Aimophila_cassinii | 0 | 18.9 | 4 | 2273142.06 |
| birds | Peucaea_humeralis | Aimophila_humeralis | 0 | 23.9 | 3.5 | 83912.61 |
| birds | Peucaea_mystacalis | Aimophila_mystacalis | 0 | 22.2 | 3.5 | 52598.3 |
| birds | Peucaea_ruficauda | Aimophila_ruficauda | 0 | 28.2 | 2.8 | 263649.93 |
| birds | Peucedramus_taeniatus | Peucedramus_taeniatus | 0 | 11 | 3.5 | 692877.18 |
| birds | Pezoporus_occidentalis | Pezoporus_occidentalis | 1 | 130 | 4 | 2740521.52 |
| birds | Pezoporus_wallicus | Pezoporus_wallicus | 0 | 71.2 | 3.5 | 195370.72 |
| birds | Phaenicophaeus_curvirostris | Phaenicophaeus_curvirostris | 0 | 154 | 2.4 | 1217500.23 |
| birds | Phaenicophaeus_sumatranus | Phaenicophaeus_sumatranus | 0 | 91.9 | 2 | 1280940.39 |
| birds | Phaenicophaeus_tristis | Phaenicophaeus_tristis | 0 | 117 | 3 | 4373319.66 |
| birds | Phaenicophaeus_viridirostris | Phaenicophaeus_viridirostris | 0 | 67 | 2 | 785946.33 |
| birds | Phaenicophilus_palmarum | Phaenicophilus_palmarum | 0 | 30.5 | 2.4 | 66937.13 |
| birds | Phaenicophilus_poliocephalus | Phaenicophilus_poliocephalus | 0 | 31 | 2.8 | 9268.82 |
| birds | Phaenostictus_mcleannani | Phaenostictus_mcleannani | 0 | 51.1 | 1.4 | 254363.89 |
| birds | Phaeochroa_cuvierii | Phaeochroa_cuvierii | 0 | 8.9 | 2 | 238479.54 |
| birds | Phaeoptila_sordida | Cynanthus_sordidus | 0 | 4 | 2 | 137024.78 |
| birds | Phaethornis_anthophilus | Phaethornis_anthophilus | 0 | 5.9 | 2 | 426150.71 |
| birds | Phaethornis_augusti | Phaethornis_augusti | 0 | 6 | 2 | 557848.62 |
| birds | Phaethornis_bourcieri | Phaethornis_bourcieri | 0 | 4.3 | 2 | 2982755.21 |

| | | | | | | |
|---|---|---|---|---|---|---|
| birds | Phaethornis_eurynome | Phaethornis_eurynome | 0 | 5.3 | 2 | 934246.25 |
| birds | Phaethornis_guy | Phaethornis_guy | 0 | 6.3 | 2 | 555160.16 |
| birds | Phaethornis_hispidus | Phaethornis_hispidus | 0 | 5 | 2 | 4008483.06 |
| birds | Phaethornis_idaliae | Phaethornis_idaliae | 0 | 2.6 | 2 | 154965.58 |
| birds | Phaethornis_koepckeae | Phaethornis_koepckeae | 0 | 5 | 2 | 96811.65 |
| birds | Phaethornis_longuemareus | Phaethornis_longuemareus | 0 | 3 | 2 | 281728 |
| birds | Phaethornis_malaris | Phaethornis_malaris | 0 | 5.9 | 2 | 3801993.44 |
| birds | Phaethornis_pretrei | Phaethornis_pretrei | 0 | 5.6 | 2 | 3736605.66 |
| birds | Phaethornis_ruber | Phaethornis_ruber | 0 | 2.4 | 2 | 9729125.66 |
| birds | Phaethornis_squalidus | Phaethornis_squalidus | 0 | 3.4 | 2 | 272378.96 |
| birds | Phaethornis_striigularis | Phaethornis_striigularis | 0 | 3 | 2 | 764879.78 |
| birds | Phaethornis_superciliosus | Phaethornis_superciliosus | 0 | 6.3 | 2 | 2562623.83 |
| birds | Phaethornis_syrmatophorus | Phaethornis_syrmatophorus | 0 | 5.8 | 2 | 159638.35 |
| birds | Phaethornis_yaruqui | Phaethornis_yaruqui | 0 | 5.4 | 2 | 119931.58 |
| birds | Phaetusa_simplex | Phaetusa_simplex | 0 | 235 | 2.4 | 13316076.04 |
| birds | Phainopepla_nitens | Phainopepla_nitens | 0 | 22.1 | 2.8 | 1423921.86 |
| birds | Phainoptila_melanoxantha | Phainoptila_melanoxantha | 0 | 56 | 2 | 6565.63 |
| birds | Phalacrocorax_carbo | Phalacrocorax_carbo | 0 | 2529 | 3.5 | 32279804.49 |
| birds | Phalacrocorax_fuscicollis | Phalacrocorax_fuscicollis | 0 | 706 | 4.2 | 1187570.01 |
| birds | Phalacrocorax_sulcirostris | Phalacrocorax_sulcirostris | 0 | 995 | 3.5 | 4510039.58 |
| birds | Phalacrocorax_varius | Phalacrocorax_varius | 0 | 1940.7 | 2.2 | 4379236.12 |
| birds | Phalaenoptilus_nuttallii | Phalaenoptilus_nuttallii | 0 | 48.3 | 2 | 4441424.23 |
| birds | Phalaropus_fulicarius | Phalaropus_fulicarius | 0 | 55.6 | 4 | 15687867.41 |
| birds | Phalaropus_lobatus | Phalaropus_lobatus | 0 | 36.7 | 4 | 18813928.96 |
| birds | Phaps_chalcoptera | Phaps_chalcoptera | 0 | 330.7 | 2 | 5889502.17 |
| birds | Phaps_elegans | Phaps_elegans | 0 | 211.4 | 2 | 694565.11 |
| birds | Phaps_histrionica | Phaps_histrionica | 0 | 289 | 2 | 3734100.7 |
| birds | Phasianus_colchicus | Phasianus_colchicus | 0 | 1120.3 | 9.9 | 16757729.73 |
| birds | Phedina_borbonica | Phedina_borbonica | 0 | 22.2 | 3.5 | 624323.08 |
| birds | Phedinopsis_brazzae | Phedina_brazzae | 0 | 13 | 3 | 403109.29 |
| birds | Phelpsia_inornata | Phelpsia_inornatus | 0 | 29.4 | 2 | 278027.72 |
| birds | Pheucticus_ludovicianus | Pheucticus_ludovicianus | 0 | 42 | 3.9 | 9302746.81 |
| birds | Pheucticus_melanocephalus | Pheucticus_melanocephalus | 0 | 47.1 | 3.5 | 5971695.75 |
| birds | Pheucticus_tibialis | Pheucticus_tibialis | 0 | 62.3 | 2 | 19605.63 |

| | | | | | | |
|---|---|---|---|---|---|---|
| birds | Pheugopedius_coraya | Thryothorus_coraya | 0 | 17.2 | 2 | 3811040.45 |
| birds | Pheugopedius_felix | Thryothorus_felix | 0 | 12.8 | 5 | 364236.12 |
| birds | Pheugopedius_maculipectus | Thryothorus_maculipectus | 0 | 15 | 3.5 | 604088.84 |
| birds | Pheugopedius_rutilus | Thryothorus_rutilus | 0 | 16.5 | 2.8 | 206155.13 |
| birds | Pheugopedius_spadix | Thryothorus_spadix | 0 | 19.1 | 2 | 51078.73 |
| birds | Phibalura_flavirostris | Phibalura_flavirostris | 0 | 46.5 | 2.4 | 1154826.62 |
| birds | Phigys_solitarius | Phigys_solitarius | 0 | 83.3 | 2 | 18131.76 |
| birds | Philemon_argenticeps | Philemon_argenticeps | 0 | 80.4 | 2.4 | 885758.87 |
| birds | Philemon_buceroides | Philemon_buceroides | 0 | 108.6 | 3.5 | 1092195.2 |
| birds | Philemon_citreogularis | Philemon_citreogularis | 0 | 60.6 | 2.4 | 3161053.22 |
| birds | Philemon_corniculatus | Philemon_corniculatus | 0 | 100.7 | 2.8 | 1758116.37 |
| birds | Philentoma_pyrhoptera | Philentoma_pyrhoptera | 0 | 16.9 | 2 | 1224085.41 |
| birds | Philentoma_velata | Philentoma_velata | 0 | 26.1 | 2 | 1151168.58 |
| birds | Philetairus_socius | Philetairus_socius | 0 | 27.4 | 3.5 | 658355.72 |
| birds | Philodice_mitchellii | Calliphlox_mitchellii | 0 | 3.1 | 2 | 59801.22 |
| birds | Philohydor_lictor | Pitangus_lictor | 0 | 25.5 | 2.4 | 9185994.71 |
| birds | Philortyx_fasciatus | Philortyx_fasciatus | 0 | 130 | 5.5 | 133413.97 |
| birds | Philydor_erythrocercum | Philydor_erythrocercum | 0 | 25.3 | 2 | 5418434.13 |
| birds | Phimosus_infuscatus | Phimosus_infuscatus | 0 | 559 | 3.5 | 6585514.18 |
| birds | Phlegopsis_nigromaculata | Phlegopsis_nigromaculata | 0 | 45.2 | 2 | 3887704.81 |
| birds | Phleocryptes_melanops | Phleocryptes_melanops | 0 | 14.6 | 2.8 | 3823377.46 |
| birds | Phlogophilus_hemileucurus | Phlogophilus_hemileucurus | 0 | 2.8 | 2 | 93348.29 |
| birds | Phoenicoparrus_andinus | Phoenicoparrus_andinus | 1 | 4900 | 1 | 354440.55 |
| birds | Phoenicoparrus_jamesi | Phoenicoparrus_jamesi | 0 | 2000 | 1 | 345049.4 |
| birds | Phoenicopterus_chilensis | Phoenicopterus_chilensis | 0 | 2277 | 1 | 3679115.35 |
| birds | Phoenicopterus_ruber | Phoenicopterus_ruber | 0 | 3031.6 | 1 | 490729.15 |
| birds | Phoeniculus_bollei | Phoeniculus_bollei | 0 | 54.6 | 2 | 1297397.49 |
| birds | Phoeniculus_damarensis | Phoeniculus_damarensis | 0 | 81.8 | 3 | 828686.31 |
| birds | Phoeniculus_purpureus | Phoeniculus_purpureus | 0 | 73.2 | 3.2 | 12280558.03 |
| birds | Phoeniculus_somaliensis | Phoeniculus_somaliensis | 0 | 26.5 | 3.9 | 1065209.4 |
| birds | Phoenicurus_auroreus | Phoenicurus_auroreus | 0 | 16.2 | 3.5 | 8472998.97 |
| birds | Phoenicurus_bicolor | Rhyacornis_bicolor | 0 | 23 | 2.8 | 31763.67 |
| birds | Phoenicurus_erythrogastrus | Phoenicurus_erythrogastrus | 0 | 24.5 | 4 | 4461790.55 |
| birds | Phoenicurus_frontalis | Phoenicurus_frontalis | 0 | 15.8 | 3.5 | 3065153.65 |

| | | | | | | |
|---|---|---|---|---|---|---|
| birds | Phoenicurus_fuliginosus | Rhyacornis_fuliginosa | 0 | 20.3 | 3.9 | 5292160.32 |
| birds | Phoenicurus_leucocephalus | Chaimarrornis_leucocephalus | 0 | 30 | 3.9 | 4372940.27 |
| birds | Phoenicurus_moussieri | Phoenicurus_moussieri | 0 | 14.9 | 4.5 | 1093552.5 |
| birds | Phoenicurus_ochruros | Phoenicurus_ochruros | 0 | 16.5 | 4.9 | 29007225.13 |
| birds | Phoenicurus_phoenicurus | Phoenicurus_phoenicurus | 0 | 14.6 | 5.9 | 42756527.38 |
| birds | Phoenicurus_schisticeps | Phoenicurus_schisticeps | 0 | 18.1 | 3 | 754510.63 |
| birds | Pholia_sharpii | Cinnyricinclus_sharpii | 0 | 41 | 3 | 195406.84 |
| birds | Phragmacia_substriata | Phragmacia_substriata | 0 | 10.6 | 3 | 466617.53 |
| birds | Phylidonyris_niger | Phylidonyris_niger | 0 | 18.2 | 2 | 1047225.93 |
| birds | Phylidonyris_novaehollandiae | Phylidonyris_novaehollandiae | 0 | 20.2 | 2.4 | 851798.64 |
| birds | Phylidonyris_pyrrhopterus | Phylidonyris_pyrrhopterus | 0 | 16.3 | 2 | 357930.02 |
| birds | Phyllastrephus_albigularis | Phyllastrephus_albigularis | 0 | 24.5 | 2 | 2136746.72 |
| birds | Phyllastrephus_baumanni | Phyllastrephus_baumanni | 0 | 27.5 | 2 | 446395.6 |
| birds | Phyllastrephus_cabanisi | Phyllastrephus_cabanisi | 0 | 34.4 | 2 | 1995009.62 |
| birds | Phyllastrephus_cerviniventris | Phyllastrephus_cerviniventris | 0 | 27 | 2 | 1170058.22 |
| birds | Phyllastrephus_debilis | Phyllastrephus_debilis | 0 | 14.3 | 2 | 199654.82 |
| birds | Phyllastrephus_fischeri | Phyllastrephus_fischeri | 0 | 27 | 2 | 270855.04 |
| birds | Phyllastrephus_flavostriatus | Phyllastrephus_flavostriatus | 0 | 28.3 | 2 | 444379.94 |
| birds | Phyllastrephus_fulviventris | Phyllastrephus_fulviventris | 0 | 33.3 | 2 | 152087.25 |
| birds | Phyllastrephus_icterinus | Phyllastrephus_icterinus | 0 | 18.9 | 2 | 2473054.92 |
| birds | Phyllastrephus_poensis | Phyllastrephus_poensis | 0 | 28.9 | 2 | 44370.69 |
| birds | Phyllastrephus_terrestris | Phyllastrephus_terrestris | 0 | 31.1 | 2 | 2434586.59 |
| birds | Phyllastrephus_xavieri | Phyllastrephus_xavieri | 0 | 22.6 | 2 | 968575.23 |
| birds | Phyllergates_cucullatus | Orthotomus_cuculatus | 0 | 5.9 | 3.5 | 1549112.24 |
| birds | Phyllolais_pulchella | Phyllolais_pulchella | 0 | 6 | 2.4 | 2078702.06 |
| birds | Phyllomyias_fasciatus | Phyllomyias_fasciatus | 0 | 10.3 | 2 | 3909215.38 |
| birds | Phyllomyias_virescens | Phyllomyias_virescens | 0 | 8.2 | 2 | 1029199.71 |
| birds | Phylloscopus_affinis | Phylloscopus_affinis | 0 | 7 | 3.9 | 6032455.64 |
| birds | Phylloscopus_armandii | Phylloscopus_armandii | 0 | 9.4 | 4.5 | 4630323.06 |
| birds | Phylloscopus_bonelli | Phylloscopus_bonelli | 0 | 8.7 | 3.9 | 10369392.47 |
| birds | Phylloscopus_borealis | Phylloscopus_borealis | 0 | 10.9 | 5.9 | 15296466.25 |
| birds | Phylloscopus_borealoides | Phylloscopus_borealoides | 0 | 9.8 | 4.9 | 2985602.65 |
| birds | Phylloscopus_budongoensis | Phylloscopus_budongoensis | 0 | 8.2 | 3 | 176695.47 |
| birds | Phylloscopus_cantator | Phylloscopus_cantator | 0 | 6 | 3.5 | 788069.43 |

| | | | | | | |
|---|---|---|---|---|---|---|
| birds | Phylloscopus_castaniceps | Seicercus_castaniceps | 0 | 5.3 | 3.9 | 3275730.69 |
| birds | Phylloscopus_chloronotus | Phylloscopus_chloronotus | 0 | 4.9 | 3.9 | 452141.09 |
| birds | Phylloscopus_collybita | Phylloscopus_collybita | 0 | 8.3 | 5.5 | 27400112.71 |
| birds | Phylloscopus_coronatus | Phylloscopus_coronatus | 0 | 10.1 | 5.3 | 6469893.74 |
| birds | Phylloscopus_fuscatus | Phylloscopus_fuscatus | 0 | 8.7 | 4.9 | 20202368.26 |
| birds | Phylloscopus_griseolus | Phylloscopus_griseolus | 0 | 7.6 | 4.5 | 7386291.38 |
| birds | Phylloscopus_herberti | Phylloscopus_herberti | 0 | 8.9 | 3 | 50664.2 |
| birds | Phylloscopus_humei | Phylloscopus_humei | 0 | 6 | 4.5 | 6707584.55 |
| birds | Phylloscopus_inornatus | Phylloscopus_inornatus | 0 | 6.6 | 5.3 | 25153244.43 |
| birds | Phylloscopus_intermedius | Seicercus_affinis | 0 | 6.9 | 4.5 | 658460.77 |
| birds | Phylloscopus_laetus | Phylloscopus_laetus | 0 | 9.5 | 2 | 81632.11 |
| birds | Phylloscopus_magnirostris | Phylloscopus_magnirostris | 0 | 11.6 | 3.9 | 2364522.29 |
| birds | Phylloscopus_montis | Seicercus_montis | 0 | 6.4 | 2 | 369636.97 |
| birds | Phylloscopus_neglectus | Phylloscopus_neglectus | 0 | 5.3 | 3.9 | 3222380.78 |
| birds | Phylloscopus_nitidus | Phylloscopus_nitidus | 0 | 7.8 | 4.5 | 4625690.05 |
| birds | Phylloscopus_occipitalis | Phylloscopus_occipitalis | 0 | 9 | 3.5 | 1246331.27 |
| birds | Phylloscopus_orientalis | Phylloscopus_orientalis | 0 | 8.5 | 4.9 | 3917108.95 |
| birds | Phylloscopus_poliogenys | Seicercus_poliogenys | 0 | 6.3 | 4 | 1049263.64 |
| birds | Phylloscopus_proregulus | Phylloscopus_proregulus | 0 | 6 | 4.9 | 9985414.53 |
| birds | Phylloscopus_pulcher | Phylloscopus_pulcher | 0 | 6.8 | 3.5 | 1521950.82 |
| birds | Phylloscopus_reguloides | Phylloscopus_reguloides | 0 | 7.6 | 4.5 | 3194582.2 |
| birds | Phylloscopus_ruficapilla | Phylloscopus_ruficapilla | 0 | 7.7 | 2.8 | 423407.96 |
| birds | Phylloscopus_sibilatrix | Phylloscopus_sibilatrix | 0 | 9.2 | 5.9 | 23976906.19 |
| birds | Phylloscopus_sindianus | Phylloscopus_sindianus | 0 | 7.5 | 3.2 | 4635151.23 |
| birds | Phylloscopus_subaffinis | Phylloscopus_subaffinis | 0 | 6.2 | 4 | 3400992.32 |
| birds | Phylloscopus_subviridis | Phylloscopus_subviridis | 0 | 5.5 | 3.5 | 523915.03 |
| birds | Phylloscopus_tenellipes | Phylloscopus_tenellipes | 0 | 11.8 | 5.5 | 4607603.52 |
| birds | Phylloscopus_trivirgatus | Phylloscopus_trivirgatus | 0 | 9 | 2 | 539476.75 |
| birds | Phylloscopus_trochiloides | Phylloscopus_trochiloides | 0 | 8.1 | 4.6 | 22292683.08 |
| birds | Phylloscopus_trochilus | Phylloscopus_trochilus | 0 | 8.7 | 5.7 | 59842121.38 |
| birds | Phylloscopus_tytleri | Phylloscopus_tytleri | 0 | 7.2 | 4.5 | 1418121.01 |
| birds | Phylloscopus_umbrovirens | Phylloscopus_umbrovirens | 0 | 8.6 | 2.4 | 611974.74 |
| birds | Phylloscopus_xanthoschistos | Phylloscopus_xanthoschistos | 0 | 6.9 | 3.9 | 269573.71 |
| birds | Phylloscopus_yunnanensis | Phylloscopus_yunnanensis | 0 | 6 | 3.9 | 467272.95 |

| | | | | | | |
|---|---|---|---|---|---|---|
| birds | Phytotoma_raimondii | Phytotoma_raimondii | 1 | 39.8 | 2.8 | 58073.47 |
| birds | Phytotoma_rara | Phytotoma_rara | 0 | 47 | 2.8 | 478137.96 |
| birds | Phytotoma_rutila | Phytotoma_rutila | 0 | 40.5 | 2.8 | 2362046.18 |
| birds | Piaya_cayana | Piaya_cayana | 0 | 102 | 2.4 | 13448425.2 |
| birds | Pica_nutalli | Pica_nuttalli | 1 | 158.3 | 6.3 | 95191.28 |
| birds | Pica_pica | Pica_pica | 0 | 217.5 | 5.3 | 31870223.47 |
| birds | Picoides_arcticus | Picoides_arcticus | 0 | 69.2 | 3.5 | 6771211.07 |
| birds | Picoides_canicapillus | Dendrocopos_canicapillus | 0 | 25.8 | 4.9 | 8010810.88 |
| birds | Picoides_kizuki | Dendrocopos_kizuki | 0 | 19.2 | 5.9 | 1603020.85 |
| birds | Picoides_moluccensis | Dendrocopos_moluccensis | 0 | 16.2 | 2.8 | 742649.6 |
| birds | Picoides_tridactylus | Picoides_tridactylus | 0 | 65.5 | 4.2 | 23486930.71 |
| birds | Piculus_simplex | Piculus_simplex | 0 | 55 | 2.8 | 92589.28 |
| birds | Picumnus_cirratus | Picumnus_cirratus | 0 | 10.2 | 2.8 | 2348460.61 |
| birds | Picumnus_innominatus | Picumnus_innominatus | 0 | 10.2 | 2.8 | 4531626.73 |
| birds | Picumnus_lafresnayi | Picumnus_lafresnayi | 0 | 9.5 | 2 | 779374.9 |
| birds | Picumnus_minutissimus | Picumnus_minutissimus | 0 | 13.3 | 2.4 | 36687.97 |
| birds | Picumnus_olivaceus | Picumnus_olivaceus | 0 | 10.6 | 2.4 | 493974.89 |
| birds | Picus_awokera | Picus_awokera | 0 | 143 | 7.5 | 281098.75 |
| birds | Picus_canus | Picus_canus | 0 | 137 | 6 | 11476968.7 |
| birds | Picus_chlorolophus | Picus_chlorolophus | 0 | 65.4 | 3.5 | 3444974.54 |
| birds | Picus_erythropygius | Picus_erythropygius | 0 | 116.2 | 3.5 | 465293.36 |
| birds | Picus_puniceus | Picus_puniceus | 0 | 79.1 | 2.4 | 1380604.21 |
| birds | Picus_squamatus | Picus_squamatus | 0 | 170 | 4.9 | 781014.45 |
| birds | Picus_viridis | Picus_viridis | 0 | 176 | 6.3 | 5981746.05 |
| birds | Picus_vittatus | Picus_vittatus | 0 | 108.1 | 3.5 | 1352503 |
| birds | Picus_xanthopygaeus | Picus_xanthopygaeus | 0 | 99.7 | 3.9 | 2078020.45 |
| birds | Pinarochroa_sordida | Cercomela_sordida | 0 | 21 | 3 | 188372.16 |
| birds | Pinarocorys_erythropygia | Pinarocorys_erythropygia | 0 | 30.1 | 2 | 5722826.26 |
| birds | Pinarocorys_nigricans | Pinarocorys_nigricans | 0 | 38.1 | 2 | 3553770.13 |
| birds | Pinarornis_plumosus | Pinarornis_plumosus | 0 | 65.8 | 2.4 | 213455.19 |
| birds | Pinicola_enucleator | Pinicola_enucleator | 0 | 56.4 | 3.5 | 24288557.5 |
| birds | Pionites_leucogaster | Pionites_leucogaster | 1 | 155 | 2 | 1982282.57 |
| birds | Pionites_melanocephalus | Pionites_melanocephalus | 0 | 148.8 | 2 | 2920163.54 |
| birds | Pionopsitta_pileata | Pionopsitta_pileata | 0 | 119 | 2 | 728878.23 |

| | | | | | | |
|---|---|---|---|---|---|---|
| birds | Pipile_cumanensis | Pipile_cumanensis | 0 | 1239.2 | 3 | 3138935.13 |
| birds | Pipile_jacutinga | Pipile_jacutinga | 1 | 1241 | 2.4 | 1063311.16 |
| birds | Pipile_pipile | Pipile_pipile | 1 | 1816.6 | 2 | 3403.88 |
| birds | Pipilo_chlorurus | Pipilo_chlorurus | 0 | 29.4 | 4 | 4121928.59 |
| birds | Pipilo_erythrophthalmus | Pipilo_erythrophthalmus | 0 | 40 | 3.5 | 3619655.58 |
| birds | Pipilo_maculatus | Pipilo_maculatus | 0 | 39.3 | 3.5 | 5605136.85 |
| birds | Pipilo_ocai | Pipilo_ocai | 0 | 61 | 4 | 114057.78 |
| birds | Pipra_aureola | Pipra_aureola | 0 | 16.1 | 2 | 671432.52 |
| birds | Pipra_fasciicauda | Pipra_fasciicauda | 0 | 15.9 | 2 | 4779358.8 |
| birds | Pipra_filicauda | Pipra_filicauda | 0 | 15.4 | 2 | 2614150.07 |
| birds | Pipreola_riefferii | Pipreola_riefferii | 0 | 49.3 | 2 | 257255.05 |
| birds | Piranga_bidentata | Piranga_bidentata | 0 | 36.1 | 2.4 | 417994.02 |
| birds | Piranga_flava | Piranga_flava | 0 | 37.7 | 2.8 | 5776535.17 |
| birds | Piranga_ludoviciana | Piranga_ludoviciana | 0 | 28.1 | 3.9 | 7784975.2 |
| birds | Piranga_olivacea | Piranga_olivacea | 0 | 28.2 | 3.5 | 6581671.34 |
| birds | Piranga_rubra | Piranga_rubra | 0 | 29.1 | 3.5 | 8740188.44 |
| birds | Pitangus_sulphuratus | Pitangus_sulphuratus | 0 | 62.9 | 3.9 | 16134345.06 |
| birds | Pithys_albifrons | Pithys_albifrons | 0 | 19.5 | 2 | 3102848.35 |
| birds | Pitta_angolensis | Pitta_angolensis | 0 | 62.7 | 3 | 3968093.86 |
| birds | Pitta_brachyura | Pitta_brachyura | 0 | 55.5 | 4.9 | 2625350.31 |
| birds | Pitta_elegans | Pitta_elegans | 0 | 56.6 | 2.4 | 92181.31 |
| birds | Pitta_iris | Pitta_iris | 0 | 62.2 | 3.9 | 258796.97 |
| birds | Pitta_maxima | Pitta_maxima | 0 | 190 | 1 | 23409.65 |
| birds | Pitta_megarhyncha | Pitta_megarhyncha | 0 | 106 | 2.8 | 56694.66 |
| birds | Pitta_moluccensis | Pitta_moluccensis | 0 | 76 | 4.6 | 2249725.2 |
| birds | Pitta_nympha | Pitta_nympha | 1 | 88.3 | 4.9 | 1374640.89 |
| birds | Pitta_sordida | Pitta_sordida | 0 | 64.5 | 3.2 | 3188235.87 |
| birds | Pitta_versicolor | Pitta_versicolor | 0 | 88.9 | 3.9 | 565329.43 |
| birds | Pittasoma_michleri | Pittasoma_michleri | 0 | 109 | 2 | 49173.73 |
| birds | Platalea_ajaja | Platalea_ajaja | 0 | 1490 | 2.4 | 8415237.44 |
| birds | Platalea_alba | Platalea_alba | 0 | 1521 | 2.4 | 14157612.5 |
| birds | Platalea_flavipes | Platalea_flavipes | 0 | 1741.3 | 3 | 4698362.97 |
| birds | Platalea_leucorodia | Platalea_leucorodia | 0 | 1868 | 3.5 | 9201391.34 |
| birds | Platalea_minor | Platalea_minor | 1 | 1228 | 4.9 | 210695.09 |

| | | | | | | |
|---|---|---|---|---|---|---|
| birds | Platalea_regia | Platalea_regia | 0 | 1723.5 | 3 | 6262402.87 |
| birds | Platycercus_adscitus | Platycercus_adscitus | 0 | 111 | 3.9 | 1465853.97 |
| birds | Platycercus_caledonicus | Platycercus_caledonicus | 0 | 132.6 | 4.9 | 68050.36 |
| birds | Platycercus_elegans | Platycercus_elegans | 0 | 123.9 | 5.3 | 743673.24 |
| birds | Platycercus_eximius | Platycercus_eximius | 0 | 104 | 6 | 1189978.19 |
| birds | Platycercus_icterotis | Platycercus_icterotis | 0 | 63.3 | 4.6 | 214002.52 |
| birds | Platycercus_venustus | Platycercus_venustus | 0 | 89.4 | 2.8 | 785415.83 |
| birds | Platylophus_galericulatus | Platylophus_galericulatus | 0 | 77.9 | 1.4 | 1559387.65 |
| birds | Platyrinchus_cancrominus | Platyrinchus_cancrominus | 0 | 9.2 | 2 | 443828.03 |
| birds | Platyrinchus_coronatus | Platyrinchus_coronatus | 0 | 9.1 | 2 | 5183010.82 |
| birds | Platyrinchus_leucoryphus | Platyrinchus_leucoryphus | 1 | 16 | 2 | 481741.4 |
| birds | Platyrinchus_mystaceus | Platyrinchus_mystaceus | 0 | 9.7 | 2 | 5068239.9 |
| birds | Platysmurus_leucopterus | Platysmurus_leucopterus | 0 | 180 | 2.4 | 624071.23 |
| birds | Platysteira_cyanea | Platysteira_cyanea | 0 | 14.6 | 2 | 5598291.68 |
| birds | Platysteira_peltata | Platysteira_peltata | 0 | 13.7 | 2 | 3620454.8 |
| birds | Plectorhyncha_lanceolata | Plectorhyncha_lanceolata | 0 | 40.6 | 3 | 2020455.09 |
| birds | Plectrophenax_hyperboreus | Plectrophenax_hyperboreus | 0 | 54.5 | 4.9 | 77792.43 |
| birds | Plectrophenax_nivalis | Plectrophenax_nivalis | 0 | 42.2 | 4.9 | 29206717.75 |
| birds | Plegadis_chihi | Plegadis_chihi | 0 | 616.9 | 3 | 9579778.67 |
| birds | Plegadis_falcinellus | Plegadis_falcinellus | 0 | 626.6 | 3.5 | 28264905.8 |
| birds | Plegadis_ridgwayi | Plegadis_ridgwayi | 0 | 579.4 | 2 | 495452.93 |
| birds | Plocepasser_mahali | Plocepasser_mahali | 0 | 41.5 | 1.7 | 3098861.26 |
| birds | Plocepasser_superciliosus | Plocepasser_superciliosus | 0 | 37.1 | 2 | 2062715.24 |
| birds | Ploceus_angolensis | Ploceus_angolensis | 0 | 26.6 | 2 | 260942.79 |
| birds | Ploceus_aurantius | Ploceus_aurantius | 0 | 23 | 2 | 441030.16 |
| birds | Ploceus_badius | Ploceus_badius | 0 | 22 | 2.4 | 365559.99 |
| birds | Ploceus_baglafecht | Ploceus_baglafecht | 0 | 31.6 | 2.4 | 1651063.49 |
| birds | Ploceus_benghalensis | Ploceus_benghalensis | 0 | 19.8 | 3.2 | 1001878.12 |
| birds | Ploceus_bicolor | Ploceus_bicolor | 0 | 34.5 | 2.8 | 1377098.31 |
| birds | Ploceus_capensis | Ploceus_capensis | 0 | 44 | 2.8 | 666169.71 |
| birds | Ploceus_castaneiceps | Ploceus_castaneiceps | 0 | 18.5 | 2.4 | 45796.6 |
| birds | Ploceus_cucullatus | Ploceus_cucullatus | 0 | 36.1 | 2.8 | 13754902.58 |
| birds | Ploceus_heuglini | Ploceus_heuglini | 0 | 22.8 | 2.4 | 2515613.14 |
| birds | Ploceus_hypoxanthus | Ploceus_hypoxanthus | 0 | 18.6 | 2.8 | 861169.04 |

| | | | | | | |
|---|---|---|---|---|---|---|
| birds | Ploceus_intermedius | Ploceus_intermedius | 0 | 20.6 | 2.8 | 3847715.31 |
| birds | Ploceus_jacksoni | Ploceus_jacksoni | 0 | 23.4 | 2.4 | 525034.95 |
| birds | Ploceus_luteolus | Ploceus_luteolus | 0 | 15.1 | 2.4 | 3085847.31 |
| birds | Ploceus_manyar | Ploceus_manyar | 0 | 17.4 | 2.4 | 3778604.76 |
| birds | Ploceus_megarhynchus | Ploceus_megarhynchus | 1 | 34.3 | 2.8 | 125839.04 |
| birds | Ploceus_melanocephalus | Ploceus_melanocephalus | 0 | 23 | 2.4 | 1189276.86 |
| birds | Ploceus_nelicourvi | Ploceus_nelicourvi | 0 | 23.8 | 3 | 139470.31 |
| birds | Ploceus_nigerrimus | Ploceus_nigerrimus | 0 | 34.3 | 2 | 1840056.42 |
| birds | Ploceus_nigricollis | Ploceus_nigricollis | 0 | 27.1 | 1.4 | 2474612.53 |
| birds | Ploceus_ocularis | Ploceus_ocularis | 0 | 25.2 | 2.8 | 2998767.22 |
| birds | Ploceus_olivaceiceps | Ploceus_olivaceiceps | 0 | 20.8 | 2.4 | 233911.57 |
| birds | Ploceus_pelzelni | Ploceus_pelzelni | 0 | 12.7 | 2.4 | 559286.21 |
| birds | Ploceus_philippinus | Ploceus_philippinus | 0 | 28.2 | 3.2 | 6341236.33 |
| birds | Ploceus_rubiginosus | Ploceus_rubiginosus | 0 | 30 | 3.5 | 1228691.95 |
| birds | Ploceus_sakalava | Ploceus_sakalava | 0 | 23.8 | 3.5 | 268618.92 |
| birds | Ploceus_spekei | Ploceus_spekei | 0 | 35.1 | 1.7 | 369189 |
| birds | Ploceus_subaureus | Ploceus_subaureus | 0 | 30.4 | 2.8 | 493658.92 |
| birds | Ploceus_temporalis | Ploceus_temporalis | 0 | 35.8 | 2.8 | 77920.1 |
| birds | Ploceus_velatus | Ploceus_velatus | 0 | 30.9 | 2.8 | 3539457.5 |
| birds | Ploceus_xanthops | Ploceus_xanthops | 0 | 40.6 | 2.4 | 2485649.7 |
| birds | Ploceus_xanthopterus | Ploceus_xanthopterus | 0 | 22 | 2.8 | 299643.55 |
| birds | Pluvialis_apricaria | Pluvialis_apricaria | 0 | 214 | 4 | 4417307.89 |
| birds | Pluvialis_dominica | Pluvialis_dominica | 0 | 151.5 | 4 | 28684530.21 |
| birds | Pluvialis_fulva | Pluvialis_fulva | 0 | 134.9 | 4 | 8070887.15 |
| birds | Pluvialis_squatarola | Pluvialis_squatarola | 0 | 250 | 4 | 15201972.98 |
| birds | Pluvianellus_socialis | Pluvianellus_socialis | 1 | 84.1 | 2 | 172032.99 |
| birds | Pluvianus_aegyptius | Pluvianus_aegyptius | 0 | 82 | 2.4 | 5977374.57 |
| birds | Podargus_ocellatus | Podargus_ocellatus | 0 | 175.4 | 1 | 761242.93 |
| birds | Podargus_papuensis | Podargus_papuensis | 0 | 375.7 | 1 | 958302.87 |
| birds | Podargus_strigoides | Podargus_strigoides | 0 | 308 | 1.7 | 7674636.86 |
| birds | Podica_senegalensis | Podica_senegalensis | 0 | 599 | 2.4 | 11384786.12 |
| birds | Podiceps_auritus | Podiceps_auritus | 1 | 453 | 4.5 | 23714565.82 |
| birds | Podiceps_cristatus | Podiceps_cristatus | 0 | 731 | 3.9 | 35889933.13 |
| birds | Podiceps_gallardoi | Podiceps_gallardoi | 1 | 575 | 2 | 106791.02 |

| | | | | | | |
|---|---|---|---|---|---|---|
| birds | Podiceps_grisegena | Podiceps_grisegena | 0 | 1023 | 4.5 | 23965746.92 |
| birds | Podiceps_major | Podiceps_major | 0 | 1646 | 3.9 | 2909991.5 |
| birds | Podiceps_nigricollis | Podiceps_nigricollis | 0 | 412.4 | 3.5 | 23548498.59 |
| birds | Podiceps_occipitalis | Podiceps_occipitalis | 0 | 320.7 | 2 | 2692417.52 |
| birds | Podiceps_taczanowskii | Podiceps_taczanowskii | 1 | 390.2 | 1.7 | 145.94 |
| birds | Podilymbus_podiceps | Podilymbus_podiceps | 0 | 411.9 | 5.3 | 21861728.49 |
| birds | Poecile_atricapillus | Parus_atricapillus | 0 | 10.8 | 6.9 | 9078514.18 |
| birds | Poecile_carolinensis | Parus_carolinensis | 0 | 10 | 5.5 | 2312694.52 |
| birds | Poecile_cinctus | Parus_cinctus | 0 | 11.4 | 7.7 | 10800997.61 |
| birds | Poecile_gambeli | Parus_gambeli | 0 | 11.2 | 6.7 | 2711124.48 |
| birds | Poecile_hudsonicus | Parus_hudsonicus | 0 | 9.8 | 6.5 | 7102487.58 |
| birds | Poecile_lugubris | Parus_lugubris | 0 | 15.8 | 5.9 | 1281185.72 |
| birds | Poecile_montanus | Parus_montanus | 0 | 11.1 | 7.3 | 21263364.57 |
| birds | Poecile_palustris | Parus_palustris | 0 | 11.1 | 7.1 | 9072135.09 |
| birds | Poecile_rufescens | Parus_rufescens | 0 | 9.4 | 6.9 | 958071.95 |
| birds | Poecile_sclateri | Parus_sclateri | 0 | 10.1 | 6.7 | 201847.75 |
| birds | Poecile_superciliosus | Parus_superciliosus | 0 | 10.7 | 3.5 | 546757.41 |
| birds | Poecilodryas_superciliosa | Poecilodryas_superciliosa | 0 | 19.8 | 2 | 238285.19 |
| birds | Poecilotriccus_plumbeiceps | Poecilotriccus_plumbeiceps | 0 | 5.7 | 2.4 | 1475066.74 |
| birds | Poecilotriccus_sylvia | Poecilotriccus_sylvia | 0 | 7.1 | 2 | 1909899.71 |
| birds | Poephila_acuticauda | Poephila_acuticauda | 0 | 14 | 4.5 | 844828.16 |
| birds | Poephila_cincta | Poephila_cincta | 0 | 14.9 | 4.5 | 826709.36 |
| birds | Poephila_personata | Poephila_personata | 0 | 13 | 4.9 | 972681.32 |
| birds | Pogoniulus_atroflavus | Pogoniulus_atroflavus | 0 | 18.2 | 2.4 | 3231917.57 |
| birds | Pogoniulus_bilineatus | Pogoniulus_bilineatus | 0 | 13.1 | 3 | 7153853.6 |
| birds | Pogoniulus_chrysoconus | Pogoniulus_chrysoconus | 0 | 10.8 | 2.4 | 10202745.34 |
| birds | Pogoniulus_coryphaea | Pogoniulus_coryphaeus | 0 | 10.7 | 3 | 307301.79 |
| birds | Pogoniulus_leucomystax | Pogoniulus_leucomystax | 0 | 11.8 | 2.4 | 552561.75 |
| birds | Pogoniulus_pusillus | Pogoniulus_pusillus | 0 | 9.6 | 3 | 2446279.62 |
| birds | Pogoniulus_scolopaceus | Pogoniulus_scolopaceus | 0 | 14.1 | 2.8 | 3952077.92 |
| birds | Pogoniulus_simplex | Pogoniulus_simplex | 0 | 8.8 | 2.4 | 519676.86 |
| birds | Pogoniulus_subsulphureus | Pogoniulus_subsulphureus | 0 | 10.5 | 2.4 | 3512045.74 |
| birds | Pogonocichla_stellata | Pogonocichla_stellata | 0 | 18.6 | 2.4 | 1563462.55 |
| birds | Pogonornis_bidentatus | Lybius_bidentatus | 0 | 79.3 | 2.8 | 2585581.98 |

| | | | | | | |
|---|---|---|---|---|---|---|
| birds | Pogonornis_dubius | Lybius_dubius | 0 | 91 | 2 | 2333237.28 |
| birds | Pogonornis_melanopterus | Lybius_melanopterus | 0 | 52.4 | 3.5 | 809569.53 |
| birds | Pogonornis_minor | Lybius_minor | 0 | 46.7 | 3.5 | 750053.05 |
| birds | Pogonornis_rolleti | Lybius_rolleti | 0 | 100.4 | 1.4 | 1106455.93 |
| birds | Poicephalus_cryptoxanthus | Poicephalus_cryptoxanthus | 0 | 140 | 2.4 | 1182606.67 |
| birds | Poicephalus_gulielmi | Poicephalus_gulielmi | 0 | 274.6 | 2.8 | 1483974.66 |
| birds | Poicephalus_meyeri | Poicephalus_meyeri | 0 | 117.5 | 2.8 | 5835819.27 |
| birds | Poicephalus_robustus | Poicephalus_robustus | 1 | 343 | 4 | 95197.08 |
| birds | Poicephalus_rueppellii | Poicephalus_rueppellii | 0 | 116 | 3.9 | 597941.08 |
| birds | Poicephalus_senegalus | Poicephalus_senegalus | 0 | 147 | 2.8 | 2714553.58 |
| birds | Polemaetus_bellicosus | Polemaetus_bellicosus | 1 | 3964.5 | 1 | 14888019.22 |
| birds | Polihierax_semitorquatus | Polihierax_semitorquatus | 0 | 61.5 | 3 | 3103663.96 |
| birds | Poliocephalus_poliocephalus | Poliocephalus_poliocephalus | 0 | 239.9 | 3.9 | 7046975.37 |
| birds | Poliocephalus_rufopectus | Poliocephalus_rufopectus | 0 | 249 | 2.4 | 173213.59 |
| birds | Poliocrania_exsul | Myrmeciza_exsul | 0 | 26.5 | 2 | 180127.53 |
| birds | Poliolais_lopezi | Poliolais_lopezi | 0 | 12 | 1.4 | 23329.93 |
| birds | Poliolophus_urostictus | Pycnonotus_urostictus | 0 | 24.8 | 3 | 207679.01 |
| birds | Polioptila_albiloris | Polioptila_albiloris | 0 | 6.3 | 2.4 | 221828.65 |
| birds | Polioptila_caerulea | Polioptila_caerulea | 0 | 5.8 | 3 | 7262670.83 |
| birds | Polioptila_californica | Polioptila_californica | 0 | 5.8 | 3.9 | 135910.75 |
| birds | Polioptila_dumicola | Polioptila_dumicola | 0 | 7 | 3.9 | 4483982.87 |
| birds | Polioptila_lembeyei | Polioptila_lembeyei | 0 | 5 | 3.9 | 13553.51 |
| birds | Polioptila_melanura | Polioptila_melanura | 0 | 5.2 | 3.9 | 1143933.05 |
| birds | Polioptila_plumbea | Polioptila_plumbea | 0 | 6.4 | 2.4 | 7562341.5 |
| birds | Polyboroides_typus | Polyboroides_typus | 0 | 633.2 | 2 | 14893143.16 |
| birds | Polyerata_amabilis | Amazilia_amabilis | 0 | 4.3 | 2 | 298612.12 |
| birds | Polyerata_rosenbergi | Amazilia_rosenbergi | 0 | 3.7 | 2 | 76061.18 |
| birds | Polyplectron_bicalcaratum | Polyplectron_bicalcaratum | 0 | 467.8 | 2 | 1095333.73 |
| birds | Polyplectron_chalcurum | Polyplectron_chalcurum | 0 | 508 | 2 | 83208.6 |
| birds | Polyplectron_germaini | Polyplectron_germaini | 1 | 450 | 1.4 | 117609.35 |
| birds | Polyplectron_inopinatum | Polyplectron_inopinatum | 0 | 497.5 | 2 | 10661.17 |
| birds | Polyplectron_malacense | Polyplectron_malacense | 1 | 608 | 1 | 47675.75 |
| birds | Polyplectron_napoleonis | Polyplectron_napoleonis | 1 | 374.7 | 2 | 11490.41 |
| birds | Polysticta_stelleri | Polysticta_stelleri | 1 | 806.8 | 6.9 | 3544014.16 |

| | | | | | | |
|---|---|---|---|---|---|---|
| birds | Polystictus_pectoralis | Polystictus_pectoralis | 0 | 6.2 | 3 | 4115700.93 |
| birds | Polystictus_superciliaris | Polystictus_superciliaris | 0 | 6 | 1.4 | 313983.58 |
| birds | Polytelis_alexandrae | Polytelis_alexandrae | 0 | 103.7 | 4.9 | 1852628.36 |
| birds | Polytelis_anthopeplus | Polytelis_anthopeplus | 0 | 170 | 4.9 | 387356.37 |
| birds | Polytelis_swainsonii | Polytelis_swainsonii | 0 | 153 | 4.9 | 229296.89 |
| birds | Polytmus_guainumbi | Polytmus_guainumbi | 0 | 5.2 | 2 | 4835885.75 |
| birds | Polytmus_milleri | Polytmus_milleri | 0 | 4.1 | 2 | 74967.65 |
| birds | Polytmus_theresiae | Polytmus_theresiae | 0 | 3.8 | 2 | 1942723.67 |
| birds | Pomatostomus_halli | Pomatostomus_halli | 0 | 41.8 | 1.4 | 487442.04 |
| birds | Pomatostomus_ruficeps | Pomatostomus_ruficeps | 0 | 57.8 | 3.9 | 1074854.31 |
| birds | Pomatostomus_superciliosus | Pomatostomus_superciliosus | 0 | 41.5 | 2.4 | 4621673.34 |
| birds | Pomatostomus_temporalis | Pomatostomus_temporalis | 0 | 75 | 2.4 | 3774038.59 |
| birds | Poodytes_albolimbatus | Megalurus_albolimbatus | 1 | 14 | 3 | 2389.39 |
| birds | Poodytes_carteri | Eremiornis_carteri | 0 | 12.2 | 2 | 1881674.61 |
| birds | Poodytes_gramineus | Megalurus_gramineus | 0 | 14.1 | 3.9 | 3277909.05 |
| birds | Poodytes_punctatus | Bowdleria_punctata | 0 | 25.2 | 1.7 | 196470.83 |
| birds | Pooecetes_gramineus | Pooecetes_gramineus | 0 | 25.7 | 3.9 | 11093552.21 |
| birds | Porphyrio_alleni | Porphyrio_alleni | 0 | 138.7 | 4.9 | 16360428.26 |
| birds | Porphyrio_flavirostris | Porphyrio_flavirostris | 0 | 92.6 | 4.5 | 5270055.29 |
| birds | Porphyrio_martinicus | Porphyrio_martinica | 0 | 235.1 | 6 | 16405883.33 |
| birds | Porphyrio_porphyrio | Porphyrio_porphyrio | 0 | 773.9 | 3.7 | 18686249.25 |
| birds | Porphyriops_melanops | Gallinula_melanops | 0 | 191 | 5.7 | 4136401.21 |
| birds | Porzana_albicollis | Porzana_albicollis | 0 | 100 | 2.4 | 6152059.49 |
| birds | Porzana_carolina | Porzana_carolina | 0 | 74.8 | 11 | 16460975.59 |
| birds | Porzana_fluminea | Porzana_fluminea | 0 | 60.8 | 4.6 | 1975889.79 |
| birds | Porzana_porzana | Porzana_porzana | 0 | 87.1 | 9.8 | 22860334.58 |
| birds | Premnoplex_brunnescens | Premnoplex_brunnescens | 0 | 16.3 | 2 | 406258.31 |
| birds | Primolius_auricollis | Primolius_auricollis | 0 | 245 | 3.5 | 838451.74 |
| birds | Primolius_maracana | Primolius_maracana | 0 | 256 | 2 | 5423813.07 |
| birds | Prinia_atrogularis | Prinia_atrogularis | 0 | 11.8 | 3.9 | 90149.06 |
| birds | Prinia_bairdii | Prinia_bairdii | 0 | 13.4 | 2.4 | 1110524.41 |
| birds | Prinia_buchanani | Prinia_buchanani | 0 | 6.7 | 4.5 | 1899856.57 |
| birds | Prinia_crinigera | Prinia_crinigera | 0 | 13.7 | 4.6 | 2843218.33 |
| birds | Prinia_erythroptera | Heliolais_erythropterus | 0 | 11.5 | 2.4 | 3486114.04 |

| | | | | | | |
|---|---|---|---|---|---|---|
| birds | Prinia_familiaris | Prinia_familiaris | 0 | 9 | 2.8 | 497167.74 |
| birds | Prinia_flavicans | Prinia_flavicans | 0 | 8.7 | 3.5 | 2717287.69 |
| birds | Prinia_flaviventris | Prinia_flaviventris | 0 | 7 | 3.5 | 4037497.01 |
| birds | Prinia_fluviatilis | Prinia_fluviatilis | 0 | 8 | 4 | 98560.02 |
| birds | Prinia_gracilis | Prinia_gracilis | 0 | 7.1 | 3.9 | 4034571.26 |
| birds | Prinia_hodgsonii | Prinia_hodgsonii | 0 | 6.4 | 3.5 | 4680228.23 |
| birds | Prinia_inornata | Prinia_inornata | 0 | 7 | 4.2 | 7403210.36 |
| birds | Prinia_maculosa | Prinia_maculosa | 0 | 9 | 3 | 581622.97 |
| birds | Prinia_polychroa | Prinia_polychroa | 0 | 11 | 4 | 734233.98 |
| birds | Prinia_rufescens | Prinia_rufescens | 0 | 6.5 | 3.5 | 2213362.62 |
| birds | Prinia_rufifrons | Spiloptila_rufifrons | 0 | 6.5 | 3.5 | 1242957.36 |
| birds | Prinia_socialis | Prinia_socialis | 0 | 8 | 3.9 | 2758579.48 |
| birds | Prinia_somalica | Prinia_somalica | 0 | 7.5 | 4 | 1273080.06 |
| birds | Prinia_subflava | Prinia_subflava | 0 | 8.6 | 3.5 | 14611740.02 |
| birds | Prinia_sylvatica | Prinia_sylvatica | 0 | 16.1 | 3.9 | 2267095.6 |
| birds | Prioniturus_discurus | Prioniturus_discurus | 0 | 148.1 | 3 | 171737.97 |
| birds | Prionochilus_maculatus | Prionochilus_maculatus | 0 | 8 | 2 | 1401405.5 |
| birds | Prionochilus_percussus | Prionochilus_percussus | 0 | 8.5 | 1 | 1530525.15 |
| birds | Prionodura_newtoniana | Prionodura_newtoniana | 0 | 78.3 | 2 | 5474.01 |
| birds | Prionops_plumatus | Prionops_plumatus | 0 | 33.8 | 4 | 10670489.61 |
| birds | Prionops_poliolophus | Prionops_poliolophus | 0 | 49 | 1 | 55433.14 |
| birds | Prionops_retzii | Prionops_retzii | 0 | 50.1 | 3.5 | 4451615.16 |
| birds | Prionops_scopifrons | Prionops_scopifrons | 0 | 29.4 | 3 | 827650.54 |
| birds | Probosciger_aterrimus | Probosciger_aterrimus | 0 | 841 | 1 | 715363.09 |
| birds | Procnias_averano | Procnias_averano | 0 | 146.1 | 1 | 1123871.61 |
| birds | Procnias_nudicollis | Procnias_nudicollis | 0 | 172 | 1 | 1131224.33 |
| birds | Prodotiscus_insignis | Prodotiscus_insignis | 0 | 10.4 | 2 | 3022105.25 |
| birds | Prodotiscus_regulus | Prodotiscus_regulus | 0 | 14.1 | 5.5 | 3186750.81 |
| birds | Prodotiscus_zambesiae | Prodotiscus_zambesiae | 0 | 10.2 | 2 | 3997564.3 |
| birds | Progne_chalybea | Progne_chalybea | 0 | 50 | 3.2 | 15629572.14 |
| birds | Progne_cryptoleuca | Progne_cryptoleuca | 0 | 46 | 4.9 | 109839.23 |
| birds | Progne_dominicensis | Progne_dominicensis | 0 | 39.6 | 4.9 | 102598.2 |
| birds | Progne_subis | Progne_subis | 0 | 53.8 | 4.9 | 21193891.3 |
| birds | Progne_tapera | Progne_tapera | 0 | 32 | 4 | 14549505.2 |

| | | | | | | |
|---|---|---|---|---|---|---|
| birds | Promerops_cafer | Promerops_cafer | 0 | 34.7 | 2.4 | 114533.35 |
| birds | Promerops_gurneyi | Promerops_gurneyi | 0 | 34.9 | 1.4 | 103702.82 |
| birds | Protonotaria_citrea | Protonotaria_citrea | 0 | 14.3 | 4.9 | 4050752.01 |
| birds | Prunella_atrogularis | Prunella_atrogularis | 0 | 18.5 | 3.5 | 2939605.66 |
| birds | Prunella_collaris | Prunella_collaris | 0 | 38.6 | 3.5 | 11553059.39 |
| birds | Prunella_immaculata | Prunella_immaculata | 0 | 20.7 | 3.9 | 859718.29 |
| birds | Prunella_modularis | Prunella_modularis | 0 | 20.2 | 4.9 | 9380149.33 |
| birds | Prunella_montanella | Prunella_montanella | 0 | 17 | 4.9 | 14996026.23 |
| birds | Prunella_ocularis | Prunella_ocularis | 0 | 22.5 | 3.5 | 753696.61 |
| birds | Prunella_rubida | Prunella_rubida | 0 | 16.7 | 3.5 | 395971.4 |
| birds | Prunella_strophiata | Prunella_strophiata | 0 | 18.1 | 3.9 | 1390796.42 |
| birds | Psalidoprocne_albiceps | Psalidoprocne_albiceps | 0 | 11.1 | 2.8 | 1147033.42 |
| birds | Psalidoprocne_fuliginosa | Psalidoprocne_fuliginosa | 0 | 12.2 | 2 | 22148.51 |
| birds | Psalidoprocne_nitens | Psalidoprocne_nitens | 0 | 9.8 | 2 | 2890566.41 |
| birds | Psalidoprocne_obscura | Psalidoprocne_obscura | 0 | 9.4 | 2 | 1468905.41 |
| birds | Psalidoprocne_pristoptera | Psalidoprocne_pristoptera | 0 | 10.3 | 2 | 8277210.72 |
| birds | Psaltria_exilis | Psaltria_exilis | 0 | 4 | 2.4 | 11281.75 |
| birds | Psaltriparus_minimus | Psaltriparus_minimus | 0 | 5.3 | 5.9 | 2692275.52 |
| birds | Psarisomus_dalhousiae | Psarisomus_dalhousiae | 0 | 67 | 5.7 | 1318583 |
| birds | Psephotellus_chrysopterygius | Psephotus_chrysopterygius | 1 | 46.8 | 4.2 | 155156.23 |
| birds | Psephotellus_dissimilis | Psephotus_dissimilis | 0 | 45.8 | 4 | 212031.58 |
| birds | Psephotellus_varius | Psephotus_varius | 0 | 61.5 | 5.3 | 3329382.06 |
| birds | Psephotus_haematonotus | Psephotus_haematonotus | 0 | 61.4 | 5.7 | 1548738.74 |
| birds | Pseudasthenes_humicola | Asthenes_humicola | 0 | 20.8 | 3.5 | 118680.46 |
| birds | Pseudasthenes_patagonica | Asthenes_patagonica | 0 | 16.4 | 3.5 | 585684 |
| birds | Pseudasthenes_steinbachi | Asthenes_steinbachi | 0 | 19.5 | 2.4 | 273597.84 |
| birds | Pseudelaenia_leucospodia | Pseudelaenia_leucospodia | 0 | 11.9 | 2.4 | 51916.21 |
| birds | Pseudeos_fuscata | Pseudeos_fuscata | 0 | 149 | 2 | 766350.36 |
| birds | Pseudibis_davisoni | Pseudibis_davisoni | 1 | 1588 | 2.8 | 61742.4 |
| birds | Pseudibis_papillosa | Pseudibis_papillosa | 0 | 823 | 2.8 | 1740563.75 |
| birds | Pseudocolaptes_lawrencii | Pseudocolaptes_lawrencii | 0 | 51 | 1 | 6603.07 |
| birds | Pseudocolopteryx_acutipennis | Pseudocolopteryx_acutipennis | 0 | 6.9 | 3 | 1594435.28 |
| birds | Pseudocolopteryx_flaviventris | Pseudocolopteryx_flaviventris | 0 | 7.5 | 2.8 | 1685669.25 |
| birds | Pseudocolopteryx_sclateri | Pseudocolopteryx_sclateri | 0 | 8 | 2 | 816154.8 |

| | | | | | | |
|---|---|---|---|---|---|---|
| birds | Pseudonigrita_arnaudi | Pseudonigrita_arnaudi | 0 | 20 | 3.9 | 462385.54 |
| birds | Pseudonigrita_cabanisi | Pseudonigrita_cabanisi | 0 | 20.6 | 2.8 | 266415.08 |
| birds | Pseudopipra_pipra | Pipra_pipra | 0 | 11.1 | 2 | 4953783.66 |
| birds | Pseudoseisura_gutturalis | Pseudoseisura_gutturalis | 0 | 70.4 | 3.9 | 779615.04 |
| birds | Pseudoseisura_lophotes | Pseudoseisura_lophotes | 0 | 72.2 | 2.8 | 1313911 |
| birds | Psilopogon_armillaris | Megalaima_armillaris | 0 | 69.4 | 2 | 136023.9 |
| birds | Psilopogon_asiaticus | Megalaima_asiatica | 0 | 90.5 | 3.5 | 1991863.6 |
| birds | Psilopogon_australis | Megalaima_australis | 0 | 33.3 | 2.8 | 78190.52 |
| birds | Psilopogon_chrysopogon | Megalaima_chrysopogon | 0 | 151 | 2 | 924419.88 |
| birds | Psilopogon_corvinus | Megalaima_corvina | 0 | 130 | 2 | 6748.31 |
| birds | Psilopogon_faiostrictus | Megalaima_faiostricta | 0 | 93 | 2 | 709761.14 |
| birds | Psilopogon_flavifrons | Megalaima_flavifrons | 0 | 58.5 | 2.4 | 39998.46 |
| birds | Psilopogon_franklinii | Megalaima_franklinii | 0 | 63.5 | 3.2 | 747848.45 |
| birds | Psilopogon_haemacephalus | Megalaima_haemacephala | 0 | 44.5 | 2.8 | 5975609.97 |
| birds | Psilopogon_javensis | Megalaima_javensis | 0 | 115 | 2 | 10739.02 |
| birds | Psilopogon_lineatus | Megalaima_lineata | 0 | 149 | 2.8 | 1966842.26 |
| birds | Psilopogon_mystacophanos | Megalaima_mystacophanos | 0 | 77.4 | 2.8 | 1239604.69 |
| birds | Psilopogon_pyrolophus | Psilopogon_pyrolophus | 0 | 129 | 2 | 474947.38 |
| birds | Psilopogon_rafflesii | Megalaima_rafflesii | 0 | 118 | 2 | 1091543.7 |
| birds | Psilopogon_virens | Megalaima_virens | 0 | 202 | 3.5 | 3256344.79 |
| birds | Psilopogon_viridis | Megalaima_viridis | 0 | 80.5 | 2.8 | 263685.81 |
| birds | Psilopogon_zeylanicus | Megalaima_zeylanica | 0 | 119 | 2.8 | 2194453.73 |
| birds | Psilopsiagon_aurifrons | Psilopsiagon_aurifrons | 0 | 33.6 | 4.2 | 947209.8 |
| birds | Psilopsiagon_aymara | Psilopsiagon_aymara | 0 | 30 | 4.9 | 382901.13 |
| birds | Psiloscops_flammeolus | Otus_flammeolus | 0 | 59.9 | 2.8 | 1654759 |
| birds | Psittacara_acuticaudatus | Aratinga_acuticaudata | 0 | 171 | 2 | 3363879.68 |
| birds | Psittacara_erythrogenys | Aratinga_erythrogenys | 0 | 151 | 2.8 | 164717.14 |
| birds | Psittacara_euops | Aratinga_euops | 1 | 87 | 3.2 | 108504.92 |
| birds | Psittacara_finschi | Aratinga_finschi | 0 | 150 | 2.8 | 92560.88 |
| birds | Psittacara_holochlorus | Aratinga_holochlora | 0 | 138 | 4 | 299273.47 |
| birds | Psittacara_leucophthalmus | Aratinga_leucophthalma | 0 | 158 | 3.5 | 8731069.11 |
| birds | Psittacara_mitratus | Aratinga_mitrata | 0 | 249 | 2.4 | 291306.04 |
| birds | Psittacula_alexandri | Psittacula_alexandri | 0 | 147 | 3.5 | 2602483.04 |
| birds | Psittacus_erithacus | Psittacus_erithacus | 1 | 333 | 2.4 | 2775743.78 |

| | | | | | | |
|---|---|---|---|---|---|---|
| birds | Psitteuteles_goldiei | Psitteuteles_goldiei | 0 | 58.7 | 2 | 307384.24 |
| birds | Psitteuteles_versicolor | Psitteuteles_versicolor | 0 | 53.5 | 3.2 | 1214241.53 |
| birds | Psittinus_cyanurus | Psittinus_cyanurus | 0 | 60.5 | 3 | 1398599.34 |
| birds | Psophia_crepitans | Psophia_crepitans | 0 | 1026 | 3 | 2708717.4 |
| birds | Psophia_leucoptera | Psophia_leucoptera | 0 | 1315.6 | 3 | 1556201.47 |
| birds | Psophocichla_litsitsirupa | Psophocichla_litsitsirupa | 0 | 74 | 2.4 | 3146030.54 |
| birds | Psophodes_cristatus | Psophodes_cristatus | 0 | 40.7 | 2.4 | 990722.54 |
| birds | Psophodes_nigrogularis | Psophodes_nigrogularis | 0 | 44.5 | 2 | 209988.36 |
| birds | Psophodes_occidentalis | Psophodes_occidentalis | 0 | 39.6 | 2.4 | 2326926.65 |
| birds | Psophodes_olivaceus | Psophodes_olivaceus | 0 | 62.4 | 2 | 428340.7 |
| birds | Pternistis_adspersus | Francolinus_adspersus | 0 | 428 | 6.3 | 1150093.13 |
| birds | Pternistis_afer | Francolinus_afer | 0 | 454.8 | 5.2 | 4561470.06 |
| birds | Pternistis_ahantensis | Francolinus_ahantensis | 0 | 544.1 | 4.9 | 683993.47 |
| birds | Pternistis_bicalcaratus | Francolinus_bicalcaratus | 0 | 439.5 | 6 | 3597861.3 |
| birds | Pternistis_capensis | Francolinus_capensis | 0 | 629.8 | 6.9 | 150806.37 |
| birds | Pternistis_castaneicollis | Francolinus_castaneicollis | 0 | 791.5 | 5.5 | 264511.06 |
| birds | Pternistis_erckelii | Francolinus_erckelii | 0 | 1197.8 | 6.3 | 162711.96 |
| birds | Pternistis_hartlaubi | Francolinus_hartlaubi | 0 | 241.5 | 5.7 | 353895.58 |
| birds | Pternistis_hildebrandti | Francolinus_hildebrandti | 0 | 532 | 6 | 900428.97 |
| birds | Pternistis_icterorhynchus | Francolinus_icterorhynchus | 0 | 571 | 6.9 | 1444290.43 |
| birds | Pternistis_jacksoni | Francolinus_jacksoni | 0 | 1097 | 3 | 16544.06 |
| birds | Pternistis_leucoscepus | Francolinus_leucoscepus | 0 | 640.6 | 5 | 2071108.37 |
| birds | Pternistis_natalensis | Francolinus_natalensis | 0 | 508.1 | 5 | 1136104.66 |
| birds | Pternistis_rufopictus | Francolinus_rufopictus | 0 | 706.1 | 4.5 | 77700.06 |
| birds | Pternistis_squamatus | Francolinus_squamatus | 0 | 449.5 | 6 | 2593426.1 |
| birds | Pternistis_swainsonii | Francolinus_swainsonii | 0 | 597.1 | 5.7 | 1549732.16 |
| birds | Pterocles_alchata | Pterocles_alchata | 0 | 275.8 | 3 | 5331411.15 |
| birds | Pterocles_bicinctus | Pterocles_bicinctus | 0 | 237 | 2.4 | 1654904.57 |
| birds | Pterocles_burchelli | Pterocles_burchelli | 0 | 180.7 | 3 | 1218509.01 |
| birds | Pterocles_coronatus | Pterocles_coronatus | 0 | 300 | 3 | 4815595.16 |
| birds | Pterocles_decoratus | Pterocles_decoratus | 0 | 181.3 | 3 | 1363573.06 |
| birds | Pterocles_exustus | Pterocles_exustus | 0 | 202.8 | 3 | 9277824.09 |
| birds | Pterocles_gutturalis | Pterocles_gutturalis | 0 | 338 | 3 | 810480.4 |
| birds | Pterocles_indicus | Pterocles_indicus | 0 | 178.2 | 2.4 | 1916026.14 |

| | | | | | | |
|---|---|---|---|---|---|---|
| birds | Pterocles_lichtensteinii | Pterocles_lichtensteinii | 0 | 209.1 | 2.4 | 3826174.11 |
| birds | Pterocles_namaqua | Pterocles_namaqua | 0 | 177 | 3 | 2037622.26 |
| birds | Pterocles_orientalis | Pterocles_orientalis | 0 | 404.9 | 3 | 7054392.13 |
| birds | Pterocles_quadricinctus | Pterocles_quadricinctus | 0 | 178.9 | 2.4 | 3917295.51 |
| birds | Pterocles_senegallus | Pterocles_senegallus | 0 | 263.4 | 3 | 9846169 |
| birds | Pteroglossus_aracari | Pteroglossus_aracari | 0 | 250.2 | 2.8 | 3319977.82 |
| birds | Pteroglossus_azara | Pteroglossus_azara | 0 | 144.5 | 2.8 | 3418746.76 |
| birds | Pteroglossus_castanotis | Pteroglossus_castanotis | 0 | 258.1 | 2 | 5071451.43 |
| birds | Pteroglossus_frantzii | Pteroglossus_frantzii | 0 | 253 | 2 | 19662.61 |
| birds | Pteroglossus_torquatus | Pteroglossus_torquatus | 0 | 219.1 | 3.2 | 1018314.61 |
| birds | Pteroglossus_viridis | Pteroglossus_viridis | 0 | 135 | 2.8 | 1558794.85 |
| birds | Pteronetta_hartlaubii | Pteronetta_hartlaubii | 0 | 790 | 8.8 | 4283729.6 |
| birds | Pterophanes_cyanopterus | Pterophanes_cyanopterus | 0 | 9.3 | 2 | 232571.39 |
| birds | Pteroptochos_castaneus | Pteroptochos_castaneus | 0 | 160.6 | 2 | 53074.04 |
| birds | Pteroptochos_megapodius | Pteroptochos_megapodius | 0 | 114 | 2 | 132289.26 |
| birds | Pteroptochos_tarnii | Pteroptochos_tarnii | 0 | 147 | 2 | 275064.61 |
| birds | Pterorhinus_caerulatus | Garrulax_caerulatus | 0 | 85.7 | 3 | 219784.76 |
| birds | Pterorhinus_chinensis | Garrulax_chinensis | 0 | 88.5 | 3.9 | 1498163.61 |
| birds | Pterorhinus_delesserti | Garrulax_delesserti | 0 | 78 | 3 | 125058.65 |
| birds | Pterorhinus_gularis | Garrulax_gularis | 0 | 92 | 3 | 307618.65 |
| birds | Pterorhinus_nuchalis | Garrulax_nuchalis | 0 | 78 | 3 | 60702.13 |
| birds | Pterorhinus_pectoralis | Garrulax_pectoralis | 0 | 145.1 | 3.9 | 2170571.37 |
| birds | Pterorhinus_perspicillatus | Garrulax_perspicillatus | 0 | 117.8 | 3.2 | 1367037.22 |
| birds | Pterorhinus_ruficollis | Garrulax_ruficollis | 0 | 66.2 | 3.5 | 296349.2 |
| birds | Pterorhinus_sannio | Garrulax_sannio | 0 | 67.8 | 3.5 | 2031254.05 |
| birds | Ptilinopus_aurantiifrons | Ptilinopus_aurantiifrons | 0 | 152 | 1 | 479657 |
| birds | Ptilinopus_coronulatus | Ptilinopus_coronulatus | 0 | 76.6 | 1 | 670308.78 |
| birds | Ptilinopus_greyi | Ptilinopus_greyii | 0 | 82.7 | 1 | 32025.73 |
| birds | Ptilinopus_insolitus | Ptilinopus_insolitus | 0 | 136.5 | 1 | 42750.02 |
| birds | Ptilinopus_iozonus | Ptilinopus_iozonus | 0 | 114 | 1 | 674305.79 |
| birds | Ptilinopus_melanospilus | Ptilinopus_melanospilus | 0 | 92 | 1 | 498198.97 |
| birds | Ptilinopus_perlatus | Ptilinopus_perlatus | 0 | 210 | 1 | 673225.36 |
| birds | Ptilinopus_perousii | Ptilinopus_perousii | 0 | 93.5 | 1.4 | 22003.57 |
| birds | Ptilinopus_pulchellus | Ptilinopus_pulchellus | 0 | 70 | 1 | 339387.21 |

| | | | | | | |
|---|---|---|---|---|---|---|
| birds | Ptilinopus_regina | Ptilinopus_regina | 0 | 101.6 | 1 | 553574.47 |
| birds | Ptilinopus_rivoli | Ptilinopus_rivoli | 0 | 149 | 2 | 294245.1 |
| birds | Ptilinopus_solomonensis | Ptilinopus_solomonensis | 0 | 108.5 | 1 | 63045.63 |
| birds | Ptilinopus_superbus | Ptilinopus_superbus | 0 | 118 | 1 | 1524991.57 |
| birds | Ptilinopus_tannensis | Ptilinopus_tannensis | 0 | 208 | 1 | 12046.29 |
| birds | Ptilinopus_viridis | Ptilinopus_viridis | 0 | 121 | 1 | 95441.03 |
| birds | Ptiliogonys_caudatus | Ptilogonys_caudatus | 0 | 36.2 | 2 | 9544.17 |
| birds | Ptiliogonys_cinereus | Ptilogonys_cinereus | 0 | 33.6 | 2 | 524061.64 |
| birds | Ptilonorhynchus_violaceus | Ptilonorhynchus_violaceus | 0 | 218.9 | 1.7 | 523221.88 |
| birds | Ptilopachus_nahani | Francolinus_nahani | 1 | 310 | 4 | 99521.54 |
| birds | Ptilopachus_petrosus | Ptilopachus_petrosus | 0 | 193 | 4.9 | 4691145.79 |
| birds | Ptiloprora_perstriata | Ptiloprora_perstriata | 0 | 26.6 | 1 | 102361.91 |
| birds | Ptilopsis_leucotis | Otus_leucotis | 0 | 204 | 2.4 | 8309644.17 |
| birds | Ptilostomus_afer | Ptilostomus_afer | 0 | 128 | 4.6 | 3779257.09 |
| birds | Ptilotula_flavescens | Lichenostomus_flavescens | 0 | 12.4 | 1.4 | 1038738.89 |
| birds | Ptilotula_fusca | Lichenostomus_fuscus | 0 | 17.7 | 1.7 | 949120.71 |
| birds | Ptilotula_keartlandi | Lichenostomus_keartlandi | 0 | 15.2 | 2 | 2753396.76 |
| birds | Ptilotula_ornata | Lichenostomus_ornatus | 0 | 17.6 | 2 | 1105315.89 |
| birds | Ptilotula_penicillata | Lichenostomus_penicillatus | 0 | 19.4 | 2.4 | 4922061.09 |
| birds | Ptilotula_plumula | Lichenostomus_plumulus | 0 | 16.5 | 2 | 5368167.7 |
| birds | Ptyonoprogne_concolor | Hirundo_concolor | 0 | 13 | 2.8 | 2727893.22 |
| birds | Ptyonoprogne_fuligula | Hirundo_fuligula | 0 | 21.1 | 3.9 | 2657867.37 |
| birds | Ptyonoprogne_obsoleta | Hirundo_obsoleta | 0 | 10.2 | 2.4 | 7338605.45 |
| birds | Ptyonoprogne_rupestris | Hirundo_rupestris | 0 | 20.4 | 3.2 | 12884287.97 |
| birds | Purnella_albifrons | Phylidonyris_albifrons | 0 | 17.4 | 2 | 4027281.71 |
| birds | Purpureicephalus_spurius | Purpureicephalus_spurius | 0 | 116 | 5.3 | 126268.75 |
| birds | Pycnonotus_aurigaster | Pycnonotus_aurigaster | 0 | 44.3 | 3.5 | 3748833.48 |
| birds | Pycnonotus_barbatus | Pycnonotus_barbatus | 0 | 32.1 | 2.4 | 19572184.19 |
| birds | Pycnonotus_blanfordi | Pycnonotus_blanfordi | 0 | 33.4 | 2.4 | 1240819.12 |
| birds | Pycnonotus_brunneus | Pycnonotus_brunneus | 0 | 28.8 | 2 | 1138188.08 |
| birds | Pycnonotus_cafer | Pycnonotus_cafer | 0 | 42.9 | 2.4 | 4241761.35 |
| birds | Pycnonotus_capensis | Pycnonotus_capensis | 0 | 38.6 | 2.4 | 151666.39 |
| birds | Pycnonotus_finlaysoni | Pycnonotus_finlaysoni | 0 | 26.3 | 2.8 | 1253924.24 |
| birds | Pycnonotus_flavescens | Pycnonotus_flavescens | 0 | 28.9 | 2.4 | 917016.17 |

| | | | | | | |
|---|---|---|---|---|---|---|
| birds | Pycnonotus_goiavier | Pycnonotus_goiavier | 0 | 27.8 | 2.8 | 2142578.36 |
| birds | Pycnonotus_jocosus | Pycnonotus_jocosus | 0 | 29.5 | 3 | 5257404.41 |
| birds | Pycnonotus_leucogenys | Pycnonotus_leucogenys | 0 | 35.9 | 3 | 378023.58 |
| birds | Pycnonotus_luteolus | Pycnonotus_luteolus | 0 | 34.7 | 2 | 1386882.2 |
| birds | Pycnonotus_nigricans | Pycnonotus_nigricans | 0 | 30.8 | 3 | 2193980.34 |
| birds | Pycnonotus_penicillatus | Pycnonotus_penicillatus | 0 | 36.5 | 2 | 4919.71 |
| birds | Pycnonotus_plumosus | Pycnonotus_plumosus | 0 | 34.9 | 2 | 1506344.75 |
| birds | Pycnonotus_simplex | Pycnonotus_simplex | 0 | 25.1 | 2 | 1340820.45 |
| birds | Pycnonotus_sinensis | Pycnonotus_sinensis | 0 | 34.2 | 3.5 | 3460936.24 |
| birds | Pycnonotus_xantholaemus | Pycnonotus_xantholaemus | 1 | 28.1 | 2.4 | 362753.3 |
| birds | Pycnonotus_xanthopygos | Pycnonotus_xanthopygos | 0 | 37.6 | 3 | 1313216.5 |
| birds | Pycnonotus_xanthorrhous | Pycnonotus_xanthorrhous | 0 | 26.9 | 2.4 | 1741022.34 |
| birds | Pycnoptilus_floccosus | Pycnoptilus_floccosus | 1 | 28.9 | 1.4 | 79638.34 |
| birds | Pygarrhichas_albogularis | Pygarrhichas_albogularis | 0 | 24 | 2.4 | 470695.91 |
| birds | Pygochelidon_cyanoleuca | Pygochelidon_cyanoleuca | 0 | 9.7 | 2.8 | 10370376.43 |
| birds | Pygochelidon_melanoleuca | Atticora_melanoleuca | 0 | 10.8 | 3 | 2130687.81 |
| birds | Pyrenestes_ostrinus | Pyrenestes_ostrinus | 0 | 22.4 | 3.9 | 4722415.43 |
| birds | Pyrenestes_sanguineus | Pyrenestes_sanguineus | 0 | 19.2 | 3.5 | 420440.4 |
| birds | Pyriglena_leuconota | Pyriglena_leuconota | 0 | 32.3 | 1.4 | 1982603.49 |
| birds | Pyriglena_leucoptera | Pyriglena_leucoptera | 0 | 28.8 | 2 | 1228571.5 |
| birds | Pyrocephalus_rubinus | Pyrocephalus_rubinus | 0 | 14.4 | 2.4 | 17320256.16 |
| birds | Pyroderus_scutatus | Pyroderus_scutatus | 0 | 357 | 1 | 2116677.8 |
| birds | Pyrrhocorax_graculus | Pyrrhocorax_graculus | 0 | 211 | 4.2 | 3556954.7 |
| birds | Pyrrhocorax_pyrrhocorax | Pyrrhocorax_pyrrhocorax | 0 | 276.7 | 3.9 | 12981520.53 |
| birds | Pyrrholaemus_brunneus | Pyrrholaemus_brunneus | 0 | 12.5 | 3.5 | 3285337.24 |
| birds | Pyrrholaemus_sagittatus | Chthonicola_sagittatus | 0 | 13.3 | 3.5 | 1269134.49 |
| birds | Pyrrhomyias_cinnamomeus | Pyrrhomyias_cinnamomeus | 0 | 9.9 | 2 | 539963.44 |
| birds | Pyrrhula_nipalensis | Pyrrhula_nipalensis | 0 | 24.6 | 2 | 2202759.21 |
| birds | Pyrrhula_pyrrhula | Pyrrhula_pyrrhula | 0 | 24.3 | 4.5 | 24467876.15 |
| birds | Pyrrhura_cruentata | Pyrrhura_cruentata | 1 | 75.9 | 2.8 | 233537.34 |
| birds | Pyrrhura_frontalis | Pyrrhura_frontalis | 0 | 72 | 5 | 1704534.47 |
| birds | Pyrrhura_hoffmanni | Pyrrhura_hoffmanni | 0 | 82.2 | 6 | 8001.11 |
| birds | Pyrrhura_leucotis | Pyrrhura_leucotis | 1 | 75.9 | 6.7 | 273322.17 |
| birds | Pyrrhura_melanura | Pyrrhura_melanura | 0 | 70.5 | 4 | 1490802.92 |

| | | | | | | |
|---|---|---|---|---|---|---|
| birds | Pyrrhura_molinae | Pyrrhura_molinae | 0 | 77.1 | 3 | 675243.93 |
| birds | Pyrrhura_perlata | Pyrrhura_perlata | 0 | 75 | 4.2 | 1396027.27 |
| birds | Pyrrhura_picta | Pyrrhura_picta | 0 | 62.1 | 4.5 | 1473164.35 |
| birds | Pyrrhura_rupicola | Pyrrhura_rupicola | 0 | 75 | 7 | 545590.25 |
| birds | Pyrrhurus_scandens | Pyrrhurus_scandens | 0 | 43.8 | 2.4 | 3269576.15 |
| birds | Pytilia_afra | Pytilia_afra | 0 | 16.6 | 3.5 | 3386528.18 |
| birds | Pytilia_hypogrammica | Pytilia_hypogrammica | 0 | 14.8 | 3.5 | 818060.77 |
| birds | Pytilia_melba | Pytilia_melba | 0 | 15.4 | 4.2 | 6033515.56 |
| birds | Pytilia_phoenicoptera | Pytilia_phoenicoptera | 0 | 14.5 | 3.5 | 1507011.59 |
| birds | Quelea_cardinalis | Quelea_cardinalis | 0 | 12.7 | 2.4 | 675544.27 |
| birds | Quelea_erythrops | Quelea_erythrops | 0 | 19.3 | 2 | 2572026.92 |
| birds | Quelea_quelea | Quelea_quelea | 0 | 18.6 | 2.2 | 10621873.64 |
| birds | Querula_purpurata | Querula_purpurata | 0 | 107.4 | 1 | 5636816.57 |
| birds | Quiscalus_lugubris | Quiscalus_lugubris | 0 | 63 | 2.8 | 918524.95 |
| birds | Quiscalus_major | Quiscalus_major | 0 | 151.2 | 3.5 | 207614.15 |
| birds | Quiscalus_mexicanus | Quiscalus_mexicanus | 0 | 160.5 | 3.5 | 5121812.21 |
| birds | Quiscalus_nicaraguensis | Quiscalus_nicaraguensis | 0 | 63.4 | 2.4 | 7099.2 |
| birds | Quiscalus_niger | Quiscalus_niger | 0 | 72.2 | 3.5 | 207006.01 |
| birds | Quiscalus_quiscula | Quiscalus_quiscula | 0 | 105.2 | 4.2 | 8475642.15 |
| birds | Quoyornis_georgianus | Eopsaltria_georgiana | 0 | 19.1 | 2 | 45266.45 |
| birds | Radjah_radjah | Tadorna_radjah | 0 | 885.2 | 8.5 | 2173785.4 |
| birds | Rallicula_rubra | Rallina_rubra | 0 | 81 | 1 | 114047.12 |
| birds | Rallina_eurizonoides | Rallina_eurizonoides | 0 | 109 | 5.7 | 3016592.95 |
| birds | Rallina_fasciata | Rallina_fasciata | 0 | 100 | 4.2 | 1965465.7 |
| birds | Rallina_tricolor | Rallina_tricolor | 0 | 178 | 4.6 | 727507.1 |
| birds | Rallus_antarcticus | Rallus_antarcticus | 1 | 60.4 | 8 | 1079988.15 |
| birds | Rallus_aquaticus | Rallus_aquaticus | 0 | 110.7 | 8.1 | 13788813.36 |
| birds | Rallus_caerulescens | Rallus_caerulescens | 0 | 162.1 | 3.5 | 5673805.66 |
| birds | Rallus_elegans | Rallus_elegans | 0 | 313.3 | 11 | 3180995.76 |
| birds | Rallus_limicola | Rallus_limicola | 0 | 84.1 | 7.2 | 10924844.28 |
| birds | Rallus_longirostris | Rallus_longirostris | 0 | 259.4 | 6.5 | 359148.15 |
| birds | Ramphastos_ambiguus | Ramphastos_ambiguus | 0 | 651.7 | 2.4 | 684339.23 |
| birds | Ramphastos_dicolorus | Ramphastos_dicolorus | 0 | 331 | 2.8 | 1394133.06 |
| birds | Ramphastos_sulfuratus | Ramphastos_sulfuratus | 0 | 409.7 | 2 | 717685.39 |

| | | | | | | |
|---|---|---|---|---|---|---|
| birds | Ramphastos_toco | Ramphastos_toco | 0 | 618 | 2.8 | 4573489.81 |
| birds | Ramphastos_tucanus | Ramphastos_tucanus | 0 | 659.6 | 2.4 | 2365965.2 |
| birds | Ramphastos_vitellinus | Ramphastos_vitellinus | 0 | 360.4 | 2.8 | 1748066.75 |
| birds | Ramphiculus_fischeri | Ptilinopus_fischeri | 0 | 177 | 1 | 41322.27 |
| birds | Ramphiculus_jambu | Ptilinopus_jambu | 0 | 135 | 1 | 1433397.98 |
| birds | Ramphiculus_leclancheri | Ptilinopus_leclancheri | 0 | 164 | 1 | 288493.56 |
| birds | Ramphocaenus_melanurus | Ramphocaenus_melanurus | 0 | 9.7 | 2 | 9179813.28 |
| birds | Ramphocoris_clotbey | Rhamphocoris_clotbey | 0 | 49.1 | 3.9 | 934337.78 |
| birds | Ramphomicron_microrhynchum | Ramphomicron_microrhynchum | 0 | 3.6 | 2 | 203534.21 |
| birds | Ramphotrigon_fuscicauda | Ramphotrigon_fuscicauda | 0 | 18.6 | 2 | 472095.3 |
| birds | Ramphotrigon_megacephalum | Ramphotrigon_megacephalum | 0 | 14.2 | 2 | 2269306.34 |
| birds | Ramphotrigon_ruficauda | Ramphotrigon_ruficauda | 0 | 18.4 | 3 | 6736020.91 |
| birds | Ramsayornis_fasciatus | Ramsayornis_fasciatus | 0 | 12.7 | 1.7 | 931104.02 |
| birds | Ramsayornis_modestus | Ramsayornis_modestus | 0 | 12.1 | 2 | 290171.25 |
| birds | Recurvirostra_americana | Recurvirostra_americana | 0 | 304.5 | 4 | 6045989.84 |
| birds | Recurvirostra_andina | Recurvirostra_andina | 0 | 361 | 4 | 708582.58 |
| birds | Recurvirostra_avosetta | Recurvirostra_avosetta | 0 | 304 | 3.5 | 33474924.46 |
| birds | Recurvirostra_novaehollandiae | Recurvirostra_novaehollandiae | 0 | 325 | 4 | 3538477.51 |
| birds | Regulus_ignicapilla | Regulus_ignicapilla | 0 | 5.6 | 6.9 | 3980020.75 |
| birds | Regulus_regulus | Regulus_regulus | 0 | 5.5 | 9.9 | 16994335.78 |
| birds | Regulus_satrapa | Regulus_satrapa | 0 | 6.2 | 8.5 | 11944577.15 |
| birds | Remiz_consobrinus | Remiz_consobrinus | 0 | 9.1 | 7.1 | 1693475.35 |
| birds | Remiz_coronatus | Remiz_coronatus | 0 | 6.9 | 5.2 | 3914348.34 |
| birds | Remiz_pendulinus | Remiz_pendulinus | 0 | 9.3 | 3.7 | 12487369.78 |
| birds | Rhabdotorrhinus_corrugatus | Aceros_corrugatus | 1 | 1590 | 2.4 | 892566.53 |
| birds | Rhabdotorrhinus_exarhatus | Penelopides_exarhatus | 1 | 370 | 2.8 | 119932.78 |
| birds | Rhaphidura_sabini | Rhaphidura_sabini | 0 | 16 | 2.4 | 1778431.25 |
| birds | Rhea_americana | Rhea_americana | 0 | 23000 | 19.7 | 6949731.03 |
| birds | Rhea_pennata | Rhea_pennata | 0 | 23900 | 17.3 | 1309452.5 |
| birds | Rhegmatorhina_melanosticta | Rhegmatorhina_melanosticta | 0 | 31 | 2 | 1909667.49 |
| birds | Rhinocrypta_lanceolata | Rhinocrypta_lanceolata | 0 | 61.9 | 1.7 | 1006274.62 |
| birds | Rhinopomastus_cyanomelas | Rhinopomastus_cyanomelas | 0 | 27.1 | 2.8 | 6396418.32 |
| birds | Rhinopomastus_minor | Rhinopomastus_minor | 0 | 21.7 | 2.4 | 2064431.67 |
| birds | Rhinoptilus_chalcopterus | Rhinoptilus_chalcopterus | 0 | 150 | 2.4 | 10165054.18 |

| | | | | | | |
|---|---|---|---|---|---|---|
| birds | Rhinoptilus_cinctus | Rhinoptilus_cinctus | 0 | 105 | 2 | 1957707.6 |
| birds | Rhinortha_chlorophaea | Phaenicophaeus_chlorophaeus | 0 | 51.2 | 2.4 | 1422260.56 |
| birds | Rhipidura_albicollis | Rhipidura_albicollis | 0 | 12.9 | 3 | 4734174.12 |
| birds | Rhipidura_atra | Rhipidura_atra | 0 | 12.2 | 1 | 179053.16 |
| birds | Rhipidura_aureola | Rhipidura_aureola | 0 | 11.2 | 3 | 4306980.92 |
| birds | Rhipidura_cyaniceps | Rhipidura_cyaniceps | 0 | 13.6 | 2 | 83397.44 |
| birds | Rhipidura_fuliginosa | Rhipidura_fuliginosa | 0 | 7.5 | 2.2 | 267492.07 |
| birds | Rhipidura_hyperythra | Rhipidura_hyperythra | 0 | 11.2 | 2 | 465792.13 |
| birds | Rhipidura_javanica | Rhipidura_javanica | 0 | 12.5 | 2 | 1594085.24 |
| birds | Rhipidura_leucophrys | Rhipidura_leucophrys | 0 | 27.4 | 2.8 | 8400289.57 |
| birds | Rhipidura_leucothorax | Rhipidura_leucothorax | 0 | 17.6 | 2 | 565343.93 |
| birds | Rhipidura_perlata | Rhipidura_perlata | 0 | 13.4 | 2 | 1287003.58 |
| birds | Rhipidura_phasiana | Rhipidura_phasiana | 0 | 6.9 | 2.4 | 319575.43 |
| birds | Rhipidura_rufidorsa | Rhipidura_rufidorsa | 0 | 9.8 | 2 | 489139.15 |
| birds | Rhipidura_rufifrons | Rhipidura_rufifrons | 0 | 10.1 | 2.4 | 918089.74 |
| birds | Rhipidura_rufiventris | Rhipidura_rufiventris | 0 | 12 | 2.4 | 31219.37 |
| birds | Rhipidura_spilodera | Rhipidura_verreauxi | 0 | 12.9 | 2 | 9510.79 |
| birds | Rhipidura_superflua | Rhipidura_superflua | 0 | 15 | 2.4 | 2601.62 |
| birds | Rhizothera_longirostris | Rhizothera_longirostris | 0 | 746.7 | 3.2 | 1292604.79 |
| birds | Rhodopechys_sanguineus | Rhodopechys_sanguineus | 0 | 38.1 | 4.5 | 1852767.15 |
| birds | Rhodophoneus_cruentus | Rhodophoneus_cruentus | 0 | 53.3 | 2.4 | 1845043.08 |
| birds | Rhodopis_vesper | Rhodopis_vesper | 0 | 3.8 | 2 | 305602.24 |
| birds | Rhodospiza_obsoleta | Rhodopechys_obsoletus | 0 | 25.5 | 4.9 | 3747421.27 |
| birds | Rhynchocyclus_brevirostris | Rhynchocyclus_brevirostris | 0 | 24.3 | 2.4 | 579326.26 |
| birds | Rhynchocyclus_fulvipectus | Rhynchocyclus_fulvipectus | 0 | 23.1 | 2.4 | 196891.42 |
| birds | Rhynchocyclus_olivaceus | Rhynchocyclus_olivaceus | 0 | 21.3 | 2.4 | 2806251.13 |
| birds | Rhynchophanes_mccownii | Calcarius_mccownii | 0 | 25.7 | 3.5 | 2569799.1 |
| birds | Rhynchopsitta_pachyrhyncha | Rhynchopsitta_pachyrhyncha | 1 | 324.8 | 2 | 173654.15 |
| birds | Rhynchospiza_stolzmanni | Aimophila_stolzmanni | 0 | 25.6 | 4 | 48995.91 |
| birds | Rhynchospiza_strigiceps | Aimophila_strigiceps | 0 | 22.2 | 3.5 | 666629.13 |
| birds | Rhynchotus_rufescens | Rhynchotus_rufescens | 0 | 843.5 | 2.2 | 5674200.17 |
| birds | Rhyticeros_cassidix | Aceros_cassidix | 1 | 2360 | 2.4 | 164525.74 |
| birds | Rhyticeros_plicatus | Aceros_plicatus | 0 | 1703.8 | 1.4 | 754091.95 |
| birds | Rhyticeros_subruficollis | Aceros_subruficollis | 1 | 2042 | 1.7 | 72279.84 |

| | | | | | | |
|---|---|---|---|---|---|---|
| birds | Rhyticeros_undulatus | Aceros_undulatus | 1 | 2214.6 | 1.7 | 3225027.08 |
| birds | Riccordia_maugaeus | Chlorostilbon_maugaeus | 0 | 3.1 | 2 | 8728.54 |
| birds | Riccordia_ricordii | Chlorostilbon_ricordii | 0 | 3.2 | 2 | 119014.47 |
| birds | Riccordia_swainsonii | Chlorostilbon_swainsonii | 0 | 4.3 | 2 | 74117.21 |
| birds | Ridgwayia_pinicola | Zoothera_pinicola | 0 | 88 | 2 | 379299.81 |
| birds | Riparia_paludicola | Riparia_paludicola | 0 | 13.5 | 3.2 | 7092614.09 |
| birds | Riparia_riparia | Riparia_riparia | 0 | 12.7 | 4.2 | 71794265.12 |
| birds | Rollandia_microptera | Rollandia_microptera | 1 | 706 | 2 | 48882.25 |
| birds | Rollandia_rolland | Rollandia_rolland | 0 | 324.3 | 1.7 | 4837890.42 |
| birds | Rollulus_rouloul | Rollulus_rouloul | 1 | 216.5 | 5.5 | 1418421.49 |
| birds | Rostratula_benghalensis | Rostratula_benghalensis | 0 | 121 | 3.2 | 24950994.82 |
| birds | Rostrhamus_sociabilis | Rostrhamus_sociabilis | 0 | 366.9 | 2.4 | 11603405.03 |
| birds | Rougetius_rougetii | Rougetius_rougetii | 0 | 193.4 | 4.5 | 344840.95 |
| birds | Rubigula_melanictera | Pycnonotus_melanicterus | 0 | 29.3 | 2.8 | 62208.81 |
| birds | Rufirallus_viridis | Anurolimnas_viridis | 0 | 64.3 | 1.7 | 6139478.63 |
| birds | Rupornis_magnirostris | Buteo_magnirostris | 0 | 269 | 1.4 | 15043360.16 |
| birds | Rynchops_flavirostris | Rynchops_flavirostris | 0 | 164 | 2.4 | 11670589.88 |
| birds | Rynchops_niger | Rynchops_niger | 0 | 297.7 | 3.5 | 14191827.26 |
| birds | Sagittarius_serpentarius | Sagittarius_serpentarius | 1 | 4017 | 2 | 14023594.92 |
| birds | Sakesphorus_canadensis | Sakesphorus_canadensis | 0 | 24.4 | 2.4 | 1626726.49 |
| birds | Salpinctes_obsoletus | Salpinctes_obsoletus | 0 | 16.4 | 4.9 | 5577879.41 |
| birds | Sappho_sparganurus | Sappho_sparganura | 0 | 5.3 | 2 | 409731.95 |
| birds | Sarcophanops_steerii | Eurylaimus_steerii | 1 | 38.7 | 1 | 96675.6 |
| birds | Sarkidiornis_melanotos | Sarkidiornis_melanotos | 0 | 2610 | 11 | 18695511.28 |
| birds | Sarothrura_affinis | Sarothrura_affinis | 0 | 27.8 | 4 | 524225.88 |
| birds | Sarothrura_ayresi | Sarothrura_ayresi | 1 | 31.8 | 3 | 344822.65 |
| birds | Sarothrura_boehmi | Sarothrura_boehmi | 0 | 35 | 3.9 | 1772998.93 |
| birds | Sarothrura_elegans | Sarothrura_elegans | 0 | 46 | 3.9 | 7047799.89 |
| birds | Sarothrura_insularis | Sarothrura_insularis | 0 | 26.8 | 3.5 | 289629.5 |
| birds | Sarothrura_pulchra | Sarothrura_pulchra | 0 | 45.2 | 2 | 4635423.45 |
| birds | Sarothrura_rufa | Sarothrura_rufa | 0 | 37.7 | 2.4 | 4402555 |
| birds | Sasia_abnormis | Sasia_abnormis | 0 | 9.2 | 2.4 | 1159649.15 |
| birds | Sasia_ochracea | Sasia_ochracea | 0 | 9.7 | 2.8 | 1515543.88 |
| birds | Satrapa_icterophrys | Satrapa_icterophrys | 0 | 21.5 | 2.8 | 8028682.88 |

| | | | | | | |
|---|---|---|---|---|---|---|
| birds | Saucerottia_beryllina | Amazilia_beryllina | 0 | 4.5 | 2 | 683715.08 |
| birds | Saucerottia_castaneiventris | Amazilia_castaneiventris | 0 | 5 | 2 | 4916.59 |
| birds | Saucerottia_cyanocephala | Amazilia_cyanocephala | 0 | 5.1 | 2 | 304040.65 |
| birds | Saucerottia_edward | Amazilia_edward | 0 | 4.7 | 2 | 42606.22 |
| birds | Saucerottia_saucerottei | Amazilia_saucerrottei | 0 | 4.5 | 2 | 245059.3 |
| birds | Saucerottia_tobaci | Amazilia_tobaci | 0 | 4.7 | 2 | 434771.48 |
| birds | Saundersilarus_saundersi | Larus_saundersi | 1 | 198 | 3 | 276484.04 |
| birds | Saxicola_caprata | Saxicola_caprata | 0 | 15.2 | 3.9 | 9769943 |
| birds | Saxicola_ferreus | Saxicola_ferreus | 0 | 14.7 | 4.9 | 4791183.22 |
| birds | Saxicola_insignis | Saxicola_insignis | 1 | 15.3 | 4.5 | 377390.34 |
| birds | Saxicola_jerdoni | Saxicola_jerdoni | 0 | 15.3 | 3.5 | 398091.36 |
| birds | Saxicola_rubetra | Saxicola_rubetra | 0 | 16.6 | 5.3 | 18264973.04 |
| birds | Saxicola_torquatus | Saxicola_torquatus | 0 | 14.1 | 4.9 | 54335876.95 |
| birds | Sayornis_nigricans | Sayornis_nigricans | 0 | 18.6 | 2.4 | 3793824.66 |
| birds | Sayornis_phoebe | Sayornis_phoebe | 0 | 19.7 | 4.5 | 9059279.46 |
| birds | Sayornis_saya | Sayornis_saya | 0 | 20.9 | 4.5 | 10053515.71 |
| birds | Scelorchilus_albicollis | Scelorchilus_albicollis | 0 | 47.3 | 2.4 | 112769.27 |
| birds | Scelorchilus_rubecula | Scelorchilus_rubecula | 0 | 56.7 | 2.4 | 271566.99 |
| birds | Scenopoeetes_dentirostris | Scenopoeetes_dentirostris | 0 | 163.4 | 2 | 4997.73 |
| birds | Schiffornis_turdina | Schiffornis_turdina | 0 | 31.7 | 2 | 5587673.67 |
| birds | Schiffornis_virescens | Schiffornis_virescens | 0 | 25.6 | 2 | 1662556.58 |
| birds | Schistes_geoffroyi | Schistes_geoffroyi | 0 | 3.6 | 2 | 546183.79 |
| birds | Schistolais_leucopogon | Prinia_leucopogon | 0 | 11.7 | 2 | 2377144.23 |
| birds | Schoenicola_platyurus | Schoenicola_platyurus | 1 | 20 | 2.4 | 39801.45 |
| birds | Schoenicola_striatus | Chaetornis_striata | 1 | 31.5 | 3.9 | 1189979.43 |
| birds | Schoeniophylax_phryganophilus | Schoeniophylax_phryganophilus | 0 | 18.6 | 4.5 | 2565832.93 |
| birds | Sciaphylax_hemimelaena | Myrmeciza_hemimelaena | 0 | 16 | 2 | 2503854.83 |
| birds | Scleroptila_afra | Francolinus_africanus | 0 | 389.7 | 4.9 | 437877.43 |
| birds | Scleroptila_finschi | Francolinus_finschi | 0 | 569.2 | 5 | 304435.85 |
| birds | Scleroptila_gutturalis | Francolinus_levaillantoides | 0 | 376.7 | 6.3 | 1483139.56 |
| birds | Scleroptila_levaillantii | Francolinus_levaillantii | 0 | 430.9 | 6 | 832216.06 |
| birds | Scleroptila_shelleyi | Francolinus_shelleyi | 0 | 488 | 3.9 | 1271472.68 |
| birds | Scleroptila_streptophora | Francolinus_streptophorus | 0 | 385 | 4.5 | 216468.87 |
| birds | Sclerurus_albigularis | Sclerurus_albigularis | 0 | 34.8 | 2 | 655407.3 |

| | | | | | | |
|---|---|---|---|---|---|---|
| birds | Sclerurus_caudacutus | Sclerurus_caudacutus | 0 | 36.2 | 2 | 5508699.65 |
| birds | Sclerurus_guatemalensis | Sclerurus_guatemalensis | 0 | 34.7 | 2 | 417195.64 |
| birds | Sclerurus_mexicanus | Sclerurus_mexicanus | 0 | 25.1 | 2 | 6362030.4 |
| birds | Sclerurus_scansor | Sclerurus_scansor | 0 | 36.9 | 2.4 | 2117654.84 |
| birds | Scolopax_minor | Scolopax_minor | 0 | 196.3 | 4 | 4661375.09 |
| birds | Scolopax_rusticola | Scolopax_rusticola | 0 | 308.3 | 4 | 20716119.64 |
| birds | Scolopax_saturata | Scolopax_saturata | 0 | 205 | 2 | 19141.4 |
| birds | Scopus_umbretta | Scopus_umbretta | 0 | 472 | 4.2 | 19881333.45 |
| birds | Scotocerca_inquieta | Scotocerca_inquieta | 0 | 8.2 | 3.5 | 4724610.25 |
| birds | Scotopelia_bouvieri | Scotopelia_bouvieri | 0 | 637 | 1.4 | 3377961.54 |
| birds | Scotopelia_peli | Scotopelia_peli | 0 | 1942.8 | 1.4 | 4649310.21 |
| birds | Scytalopus_acutirostris | Scytalopus_acutirostris | 0 | 18.1 | 2 | 21326.44 |
| birds | Scytalopus_affinis | Scytalopus_affinis | 0 | 13.1 | 2.4 | 15372.72 |
| birds | Scytalopus_altirostris | Scytalopus_altirostris | 0 | 19.5 | 2.4 | 24016.16 |
| birds | Scytalopus_argentifrons | Scytalopus_argentifrons | 0 | 17 | 2 | 15549.46 |
| birds | Scytalopus_atratus | Scytalopus_atratus | 0 | 25.7 | 2 | 187476.08 |
| birds | Scytalopus_bolivianus | Scytalopus_bolivianus | 0 | 20.8 | 2 | 64274.24 |
| birds | Scytalopus_canus | Scytalopus_canus | 0 | 18.2 | 2 | 3607.18 |
| birds | Scytalopus_caracae | Scytalopus_caracae | 0 | 24 | 2 | 5235.88 |
| birds | Scytalopus_chocoensis | Scytalopus_chocoensis | 0 | 20 | 2 | 25911.28 |
| birds | Scytalopus_femoralis | Scytalopus_femoralis | 0 | 23.2 | 2 | 83386.49 |
| birds | Scytalopus_griseicollis | Scytalopus_griseicollis | 0 | 18.1 | 2 | 61348.22 |
| birds | Scytalopus_macropus | Scytalopus_macropus | 0 | 37.1 | 2 | 40202.47 |
| birds | Scytalopus_magellanicus | Scytalopus_magellanicus | 0 | 13.7 | 2.4 | 454183.19 |
| birds | Scytalopus_meridanus | Scytalopus_meridanus | 0 | 20 | 2 | 20695.73 |
| birds | Scytalopus_novacapitalis | Scytalopus_novacapitalis | 1 | 18.9 | 2 | 118540.07 |
| birds | Scytalopus_parvirostris | Scytalopus_parvirostris | 0 | 17.8 | 2 | 124858.93 |
| birds | Scytalopus_schulenbergi | Scytalopus_schulenbergi | 0 | 16.1 | 2 | 38231.31 |
| birds | Scytalopus_simonsi | Scytalopus_simonsi | 0 | 16.6 | 2 | 29912.19 |
| birds | Scytalopus_speluncae | Scytalopus_speluncae | 0 | 13 | 2 | 249767.52 |
| birds | Scytalopus_spillmanni | Scytalopus_spillmanni | 0 | 23.5 | 2 | 132293.77 |
| birds | Scytalopus_superciliaris | Scytalopus_superciliaris | 0 | 17.8 | 2 | 53108.46 |
| birds | Scytalopus_unicolor | Scytalopus_unicolor | 0 | 17.6 | 2 | 8500.65 |
| birds | Scytalopus_urubambae | Scytalopus_urubambae | 0 | 13 | 2.4 | 3465.39 |

| | | | | | | |
|---|---|---|---|---|---|---|
| birds | Scytalopus_vicinior | Scytalopus_vicinior | 0 | 23 | 2 | 16812.48 |
| birds | Scytalopus_zimmeri | Scytalopus_zimmeri | 0 | 15.9 | 2.4 | 53350.14 |
| birds | Seiurus_aurocapilla | Seiurus_aurocapilla | 0 | 18.8 | 4.5 | 8879216.96 |
| birds | Selasphorus_calliope | Stellula_calliope | 0 | 2.6 | 2 | 3088048.77 |
| birds | Selasphorus_platycercus | Selasphorus_platycercus | 0 | 3.5 | 2 | 976207.06 |
| birds | Selasphorus_rufus | Selasphorus_rufus | 0 | 3.5 | 2 | 2817202.82 |
| birds | Selasphorus_sasin | Selasphorus_sasin | 0 | 3.4 | 2 | 1128506.96 |
| birds | Selenidera_maculirostris | Selenidera_maculirostris | 0 | 164 | 2.4 | 930797.49 |
| birds | Selenidera_piperivora | Selenidera_piperivora | 0 | 147 | 2.4 | 1090580.68 |
| birds | Selenidera_spectabilis | Selenidera_spectabilis | 0 | 219 | 2.8 | 153224.72 |
| birds | Seleucidis_melanoleucus | Seleucidis_melanoleucus | 0 | 174.4 | 1 | 334716.18 |
| birds | Semnornis_frantzii | Semnornis_frantzii | 0 | 57.3 | 4.5 | 15874.29 |
| birds | Semnornis_ramphastinus | Semnornis_ramphastinus | 0 | 97.7 | 2.4 | 36256.06 |
| birds | Sephanoides_sephaniodes | Sephanoides_sephaniodes | 0 | 5.2 | 2 | 1104451.07 |
| birds | Sericornis_beccarii | Sericornis_beccarii | 0 | 11.9 | 2.8 | 246723.36 |
| birds | Sericornis_frontalis | Sericornis_frontalis | 0 | 13.2 | 2.4 | 1682487.01 |
| birds | Sericornis_humilis | Sericornis_humilis | 0 | 18.3 | 2.4 | 41698.31 |
| birds | Sericornis_keri | Sericornis_keri | 1 | 11.4 | 2 | 24710.75 |
| birds | Sericornis_magnirostra | Sericornis_magnirostra | 0 | 9.8 | 3.5 | 270681.41 |
| birds | Sericulus_chrysocephalus | Sericulus_chrysocephalus | 0 | 99.5 | 2 | 89618.77 |
| birds | Serilophus_lunatus | Serilophus_lunatus | 0 | 34 | 3.7 | 1235508.38 |
| birds | Serinus_alario | Serinus_alario | 0 | 12.2 | 2.8 | 508153.43 |
| birds | Serinus_canaria | Serinus_canaria | 0 | 24.3 | 3.5 | 19528.29 |
| birds | Serinus_canicollis | Serinus_canicollis | 0 | 15.2 | 3.2 | 618013.33 |
| birds | Serinus_pusillus | Serinus_pusillus | 0 | 11.5 | 3.9 | 2537686.87 |
| birds | Serinus_serinus | Serinus_serinus | 0 | 11.2 | 3.5 | 5141384.69 |
| birds | Serinus_syriacus | Serinus_syriacus | 1 | 11.7 | 4 | 78766.52 |
| birds | Serpophaga_cinerea | Serpophaga_cinerea | 0 | 8.3 | 2 | 537348.41 |
| birds | Serpophaga_subcristata | Serpophaga_subcristata | 0 | 6.6 | 2.4 | 5786710.08 |
| birds | Setopagis_parvula | Caprimulgus_parvulus | 0 | 37 | 2 | 6789301.91 |
| birds | Setophaga_adelaidae | Dendroica_adelaidae | 0 | 6.9 | 2.4 | 8923.08 |
| birds | Setophaga_americana | Parula_americana | 0 | 7.8 | 4.5 | 5430698.01 |
| birds | Setophaga_angelae | Dendroica_angelae | 1 | 7.7 | 2 | 163.58 |
| birds | Setophaga_caerulescens | Dendroica_caerulescens | 0 | 10.1 | 4 | 2978266.68 |

| | | | | | | |
|---|---|---|---|---|---|---|
| birds | Setophaga_castanea | Dendroica_castanea | 0 | 11.8 | 4.5 | 8432233.61 |
| birds | Setophaga_cerulea | Dendroica_cerulea | 0 | 9 | 4 | 4658662.6 |
| birds | Setophaga_chrysoparia | Dendroica_chrysoparia | 1 | 9.9 | 4 | 434557.34 |
| birds | Setophaga_citrina | Wilsonia_citrina | 0 | 10.5 | 3.9 | 3115320.7 |
| birds | Setophaga_coronata | Dendroica_coronata | 0 | 11.9 | 4 | 15842705.3 |
| birds | Setophaga_discolor | Dendroica_discolor | 0 | 7.6 | 4 | 2180545.91 |
| birds | Setophaga_dominica | Dendroica_dominica | 0 | 9.7 | 4 | 3207549.9 |
| birds | Setophaga_fusca | Dendroica_fusca | 0 | 9.7 | 4 | 8748176.98 |
| birds | Setophaga_graciae | Dendroica_graciae | 0 | 8.1 | 3 | 1217041.46 |
| birds | Setophaga_kirtlandii | Dendroica_kirtlandii | 0 | 13.9 | 4.5 | 53047.64 |
| birds | Setophaga_magnolia | Dendroica_magnolia | 0 | 8.1 | 4 | 8830411.55 |
| birds | Setophaga_nigrescens | Dendroica_nigrescens | 0 | 8.7 | 4 | 3560024.47 |
| birds | Setophaga_occidentalis | Dendroica_occidentalis | 0 | 10.4 | 4 | 2474138.36 |
| birds | Setophaga_palmarum | Dendroica_palmarum | 0 | 10.3 | 4.5 | 7890903.87 |
| birds | Setophaga_pensylvanica | Dendroica_pensylvanica | 0 | 9.3 | 4 | 7337980.57 |
| birds | Setophaga_petechia | Dendroica_petechia | 0 | 10.2 | 4.5 | 21086713.84 |
| birds | Setophaga_pharetra | Dendroica_pharetra | 0 | 9.9 | 2.8 | 11021.79 |
| birds | Setophaga_pinus | Dendroica_pinus | 0 | 11.8 | 4 | 3767948.83 |
| birds | Setophaga_pitiayumi | Parula_pitiayumi | 0 | 6.8 | 2 | 8364955.27 |
| birds | Setophaga_ruticilla | Setophaga_ruticilla | 0 | 8.2 | 4 | 15136310.84 |
| birds | Setophaga_striata | Dendroica_striata | 0 | 11.8 | 4.5 | 16034144.17 |
| birds | Setophaga_tigrina | Dendroica_tigrina | 0 | 10 | 6.5 | 5707680.08 |
| birds | Setophaga_townsendi | Dendroica_townsendi | 0 | 8.8 | 3.9 | 5348431.12 |
| birds | Setophaga_virens | Dendroica_virens | 0 | 8.7 | 4 | 8653032.87 |
| birds | Sheppardia_aequatorialis | Sheppardia_aequatorialis | 0 | 16.8 | 2 | 138552.17 |
| birds | Sheppardia_bocagei | Sheppardia_bocagei | 0 | 18.3 | 2 | 589392.08 |
| birds | Sheppardia_cyornithopsis | Sheppardia_cyornithopsis | 0 | 18.4 | 2 | 1077673.35 |
| birds | Sheppardia_sharpei | Sheppardia_sharpei | 0 | 14.7 | 2 | 92949.69 |
| birds | Sholicola_major | Myiomela_major | 1 | 38.5 | 2 | 1160.85 |
| birds | Sialia_currucoides | Sialia_currucoides | 0 | 29.6 | 5.5 | 7775403.65 |
| birds | Sialia_mexicana | Sialia_mexicana | 0 | 26.4 | 4.9 | 2595315.96 |
| birds | Sialia_sialis | Sialia_sialis | 0 | 27.5 | 4.5 | 6065720.04 |
| birds | Sibirionetta_formosa | Anas_formosa | 0 | 434 | 7.3 | 13029180.96 |
| birds | Sigelus_silens | Sigelus_silens | 0 | 25.7 | 3 | 982178.67 |

| | | | | | | |
|---|---|---|---|---|---|---|
| birds | Silvicultrix_diadema | Ochthoeca_diadema | 0 | 11.5 | 4 | 130712.81 |
| birds | Sitta_azurea | Sitta_azurea | 0 | 16.3 | 3.5 | 361034.34 |
| birds | Sitta_canadensis | Sitta_canadensis | 0 | 9.8 | 5.5 | 12270262.34 |
| birds | Sitta_carolinensis | Sitta_carolinensis | 0 | 21 | 6.7 | 9154405.08 |
| birds | Sitta_cashmirensis | Sitta_cashmirensis | 0 | 19 | 5.9 | 140130.44 |
| birds | Sitta_castanea | Sitta_castanea | 0 | 19.6 | 4.9 | 1252045.84 |
| birds | Sitta_europaea | Sitta_europaea | 0 | 20.4 | 7.3 | 21575986.15 |
| birds | Sitta_frontalis | Sitta_frontalis | 0 | 16.5 | 4.5 | 5214819.55 |
| birds | Sitta_himalayensis | Sitta_himalayensis | 0 | 14.3 | 5.3 | 421140.6 |
| birds | Sitta_krueperi | Sitta_krueperi | 0 | 12.4 | 5.9 | 282938.85 |
| birds | Sitta_ledanti | Sitta_ledanti | 1 | 17.2 | 2.8 | 324.46 |
| birds | Sitta_leucopsis | Sitta_leucopsis | 0 | 14.8 | 6.9 | 258577.79 |
| birds | Sitta_magna | Sitta_magna | 1 | 41.1 | 3 | 500173.51 |
| birds | Sitta_nagaensis | Sitta_nagaensis | 0 | 14.7 | 3.2 | 1053108.56 |
| birds | Sitta_neumayer | Sitta_neumayer | 0 | 26.5 | 6.3 | 1244088.59 |
| birds | Sitta_pusilla | Sitta_pusilla | 0 | 10.2 | 4.9 | 891316.4 |
| birds | Sitta_pygmaea | Sitta_pygmaea | 0 | 10.6 | 7.3 | 1953869.52 |
| birds | Sitta_tephronota | Sitta_tephronota | 0 | 35.2 | 5.9 | 2326077.82 |
| birds | Sitta_villosa | Sitta_villosa | 0 | 9.4 | 5.5 | 1824265.59 |
| birds | Sitta_whiteheadi | Sitta_whiteheadi | 1 | 11.8 | 5.5 | 2318.9 |
| birds | Sittasomus_griseicapillus | Sittasomus_griseicapillus | 0 | 13.1 | 3 | 10911731.74 |
| birds | Sittiparus_varius | Parus_varius | 0 | 17 | 6.9 | 692847.17 |
| birds | Smicrornis_brevirostris | Smicrornis_brevirostris | 0 | 5.3 | 2.4 | 7230924.88 |
| birds | Smithornis_capensis | Smithornis_capensis | 0 | 23.2 | 2.4 | 3751914 |
| birds | Smithornis_rufolateralis | Smithornis_rufolateralis | 0 | 19 | 2 | 1880114.42 |
| birds | Smithornis_sharpei | Smithornis_sharpei | 0 | 37.5 | 2 | 498936.69 |
| birds | Smutsornis_africanus | Rhinoptilus_africanus | 0 | 87.5 | 1 | 2999057.71 |
| birds | Somateria_fischeri | Somateria_fischeri | 0 | 1557.2 | 4.5 | 969744.29 |
| birds | Somateria_mollissima | Somateria_mollissima | 0 | 2060.9 | 4.9 | 16485948.2 |
| birds | Somateria_spectabilis | Somateria_spectabilis | 0 | 1616.7 | 4.5 | 7316509.6 |
| birds | Spartonoica_maluroides | Spartonoica_maluroides | 0 | 11 | 3.5 | 527135.99 |
| birds | Spatula_clypeata | Anas_clypeata | 0 | 612.6 | 9.9 | 50160551.83 |
| birds | Spatula_cyanoptera | Anas_cyanoptera | 0 | 377.5 | 10.4 | 8842984.67 |
| birds | Spatula_discors | Anas_discors | 0 | 359.4 | 9.4 | 15868657.31 |

| | | | | | | |
|---|---|---|---|---|---|---|
| birds | Spatula_hottentota | Anas_hottentota | 0 | 276.4 | 7.3 | 7754871.01 |
| birds | Spatula_platalea | Anas_platalea | 0 | 465.4 | 6.3 | 2947935.33 |
| birds | Spatula_querquedula | Anas_querquedula | 0 | 325.6 | 8.5 | 34165137.38 |
| birds | Spatula_rhynchotis | Anas_rhynchotis | 0 | 639 | 9.9 | 4454068.38 |
| birds | Spatula_smithii | Anas_smithii | 0 | 591.5 | 7.7 | 2430691.95 |
| birds | Spatula_versicolor | Anas_versicolor | 0 | 595.2 | 7.7 | 3843988.9 |
| birds | Speculanas_specularis | Speculanas_specularis | 0 | 1496.1 | 4.9 | 1108774.14 |
| birds | Spermophaga_haematina | Spermophaga_haematina | 0 | 23 | 4.2 | 1910654.25 |
| birds | Spermophaga_ruficapilla | Spermophaga_ruficapilla | 0 | 23.6 | 3 | 562817.84 |
| birds | Sphenoeacus_afer | Sphenoeacus_afer | 0 | 30.4 | 2.4 | 430218.37 |
| birds | Sphyrapicus_nuchalis | Sphyrapicus_nuchalis | 0 | 48.5 | 4.9 | 3614392.64 |
| birds | Sphyrapicus_ruber | Sphyrapicus_ruber | 0 | 54.8 | 5.3 | 1189115.64 |
| birds | Sphyrapicus_thyroideus | Sphyrapicus_thyroideus | 0 | 47.6 | 4.9 | 1248737.49 |
| birds | Sphyrapicus_varius | Sphyrapicus_varius | 0 | 50.3 | 5.3 | 9946724.6 |
| birds | Spilopelia_chinensis | Stigmatopelia_chinensis | 0 | 159 | 2.4 | 8414112.05 |
| birds | Spilopelia_senegalensis | Stigmatopelia_senegalensis | 0 | 82.1 | 2 | 29236937.74 |
| birds | Spiloptila_clamans | Spiloptila_clamans | 0 | 7 | 3.5 | 2719529.35 |
| birds | Spinus_dominicensis | Carduelis_dominicensis | 0 | 12.6 | 2.4 | 9702.06 |
| birds | Spinus_lawrencei | Carduelis_lawrencei | 0 | 10.9 | 4.5 | 556776.95 |
| birds | Spinus_notatus | Carduelis_notata | 0 | 10.9 | 2 | 488138.48 |
| birds | Spinus_pinus | Carduelis_pinus | 0 | 12.7 | 3.5 | 14061824.42 |
| birds | Spinus_psaltria | Carduelis_psaltria | 0 | 8.8 | 3.5 | 4186163.77 |
| birds | Spinus_spinus | Carduelis_spinus | 0 | 13.2 | 3.9 | 15585639.54 |
| birds | Spinus_tristis | Carduelis_tristis | 0 | 12.8 | 4.9 | 10571095.98 |
| birds | Spinus_xanthogastrus | Carduelis_xanthogastra | 0 | 12.7 | 2.4 | 161327.43 |
| birds | Spiza_americana | Spiza_americana | 0 | 26.2 | 3.5 | 6183307.84 |
| birds | Spizella_atrogularis | Spizella_atrogularis | 0 | 11.3 | 2.8 | 1397656.8 |
| birds | Spizella_breweri | Spizella_breweri | 0 | 10.9 | 3.5 | 5119315.92 |
| birds | Spizella_pallida | Spizella_pallida | 0 | 11.2 | 3.9 | 7311087.96 |
| birds | Spizella_passerina | Spizella_passerina | 0 | 12.2 | 4 | 14987835.95 |
| birds | Spizella_pusilla | Spizella_pusilla | 0 | 12.5 | 4.2 | 4818402.43 |
| birds | Spizella_wortheni | Spizella_wortheni | 1 | 12.4 | 3.5 | 60094.13 |
| birds | Spizixos_canifrons | Spizixos_canifrons | 0 | 44 | 2.4 | 1162572.35 |
| birds | Spizixos_semitorques | Spizixos_semitorques | 0 | 39 | 3.5 | 2328094.86 |

| | | | | | | |
|---|---|---|---|---|---|---|
| birds | Spizocorys_conirostris | Spizocorys_conirostris | 0 | 14.3 | 2.4 | 1128198.56 |
| birds | Spizocorys_fringillaris | Spizocorys_fringillaris | 1 | 19.2 | 2.4 | 3270.79 |
| birds | Spizocorys_sclateri | Spizocorys_sclateri | 0 | 19 | 1 | 226405.69 |
| birds | Spizocorys_starki | Eremalauda_starki | 0 | 18.5 | 2.4 | 2006101.39 |
| birds | Spodiopsar_cineraceus | Sturnus_cineraceus | 0 | 83 | 3.2 | 7910712.35 |
| birds | Sporophila_minuta | Sporophila_minuta | 0 | 7.8 | 2.4 | 2243040.21 |
| birds | Sporophila_nigricollis | Sporophila_nigricollis | 0 | 9.6 | 2.4 | 7194514.7 |
| birds | Sporophila_torqueola | Sporophila_torqueola | 0 | 8.7 | 2.4 | 543612.61 |
| birds | Sporopipes_frontalis | Sporopipes_frontalis | 0 | 17.3 | 3.7 | 3005541.44 |
| birds | Sporopipes_squamifrons | Sporopipes_squamifrons | 0 | 12.3 | 3.7 | 1945507.15 |
| birds | Stactolaema_anchietae | Stactolaema_anchietae | 0 | 47.3 | 3.2 | 1482309.28 |
| birds | Stactolaema_leucotis | Stactolaema_leucotis | 0 | 54.9 | 3.5 | 862758.91 |
| birds | Stactolaema_whytii | Stactolaema_whytii | 0 | 55.6 | 3.2 | 456236.86 |
| birds | Stagonopleura_bella | Stagonopleura_bella | 0 | 14 | 4.5 | 276741.26 |
| birds | Stagonopleura_guttata | Stagonopleura_guttata | 1 | 17.6 | 5.3 | 1192427.37 |
| birds | Stagonopleura_oculata | Stagonopleura_oculata | 0 | 12.9 | 4.9 | 82318.02 |
| birds | Steatornis_caripensis | Steatornis_caripensis | 0 | 408 | 2.8 | 1061013.51 |
| birds | Steganopus_tricolor | Steganopus_tricolor | 0 | 59.4 | 4 | 15117456.65 |
| birds | Stelgidopteryx_ruficollis | Stelgidopteryx_ruficollis | 0 | 16.1 | 4.9 | 13831495.26 |
| birds | Stelgidopteryx_serripennis | Stelgidopteryx_serripennis | 0 | 15.7 | 5.7 | 11481237.67 |
| birds | Stenostira_scita | Stenostira_scita | 0 | 5.9 | 2.4 | 1116923.25 |
| birds | Stephanoaetus_coronatus | Stephanoaetus_coronatus | 0 | 3640 | 2 | 6608300.51 |
| birds | Stephanophorus_diadematus | Stephanophorus_diadematus | 0 | 35.4 | 4 | 1088996.67 |
| birds | Stephanoxis_lalandi | Stephanoxis_lalandi | 0 | 4 | 2 | 300448.07 |
| birds | Stercorarius_longicaudus | Stercorarius_longicaudus | 0 | 287.9 | 2 | 52268431.47 |
| birds | Sterna_forsteri | Sterna_forsteri | 0 | 149 | 2.4 | 10875479.25 |
| birds | Sterna_hirundo | Sterna_hirundo | 0 | 129.2 | 2 | 51361894.71 |
| birds | Sterna_paradisaea | Sterna_paradisaea | 0 | 110 | 2.4 | 61272189.47 |
| birds | Sterna_trudeaui | Sterna_trudeaui | 0 | 153 | 3 | 539946.35 |
| birds | Sternula_superciliaris | Sterna_superciliaris | 0 | 46.4 | 2.4 | 11385573.19 |
| birds | Stictonetta_naevosa | Stictonetta_naevosa | 0 | 903.3 | 7.1 | 5897864.38 |
| birds | Stigmatura_budytoides | Stigmatura_budytoides | 0 | 11.1 | 2 | 1866889.93 |
| birds | Stiltia_isabella | Stiltia_isabella | 0 | 65.5 | 2 | 5216126.15 |
| birds | Stiphrornis_erythrothorax | Stiphrornis_erythrothorax | 0 | 15.6 | 2 | 812936.31 |

| | | | | | | |
|---|---|---|---|---|---|---|
| birds | Stipiturus_malachurus | Stipiturus_malachurus | 0 | 7.3 | 3 | 687011.19 |
| birds | Stipiturus_mallee | Stipiturus_mallee | 1 | 5.5 | 3 | 30237.26 |
| birds | Stipiturus_ruficeps | Stipiturus_ruficeps | 0 | 5 | 2.4 | 568075.87 |
| birds | Stomiopera_flava | Lichenostomus_flavus | 0 | 21.4 | 1.4 | 415063.54 |
| birds | Stomiopera_unicolor | Lichenostomus_unicolor | 0 | 30.7 | 2 | 1189196.61 |
| birds | Strepera_fuliginosa | Strepera_fuliginosa | 0 | 375.5 | 2.8 | 68050.36 |
| birds | Strepera_graculina | Strepera_graculina | 0 | 298.9 | 3 | 1580095.1 |
| birds | Strepera_versicolor | Strepera_versicolor | 0 | 395.5 | 2.4 | 2079178.27 |
| birds | Streptopelia_bitorquata | Streptopelia_bitorquata | 0 | 153 | 2 | 186292.82 |
| birds | Streptopelia_capicola | Streptopelia_capicola | 0 | 142 | 2 | 9534302.21 |
| birds | Streptopelia_decaocto | Streptopelia_decaocto | 0 | 149 | 2 | 19314230.8 |
| birds | Streptopelia_decipiens | Streptopelia_decipiens | 0 | 134 | 2 | 7210175.11 |
| birds | Streptopelia_hypopyrrha | Streptopelia_hypopyrrha | 0 | 162 | 2 | 529885.84 |
| birds | Streptopelia_lugens | Streptopelia_lugens | 0 | 155 | 2 | 945579.48 |
| birds | Streptopelia_orientalis | Streptopelia_orientalis | 0 | 232.9 | 2 | 20050458.53 |
| birds | Streptopelia_reichenowi | Streptopelia_reichenowi | 0 | 119 | 2 | 153525.54 |
| birds | Streptopelia_roseogrisea | Streptopelia_roseogrisea | 0 | 155 | 2 | 6669492.64 |
| birds | Streptopelia_semitorquata | Streptopelia_semitorquata | 0 | 176 | 2 | 13753343.72 |
| birds | Streptopelia_tranquebarica | Streptopelia_tranquebarica | 0 | 103 | 2 | 9537403.56 |
| birds | Streptopelia_turtur | Streptopelia_turtur | 1 | 132 | 2 | 22086231.08 |
| birds | Streptopelia_vinacea | Streptopelia_vinacea | 0 | 110 | 2 | 6146574.47 |
| birds | Streptoprocne_rutila | Streptoprocne_rutila | 0 | 20.2 | 2 | 1392664.2 |
| birds | Streptoprocne_semicollaris | Streptoprocne_semicollaris | 0 | 175 | 2 | 324675.19 |
| birds | Streptoprocne_zonaris | Streptoprocne_zonaris | 0 | 86.8 | 2 | 8315596.31 |
| birds | Strix_aluco | Strix_aluco | 0 | 472.5 | 3.2 | 9594068.11 |
| birds | Strix_butleri | Strix_butleri | 0 | 219 | 2.2 | 5226.95 |
| birds | Strix_hylophila | Strix_hylophila | 0 | 345.4 | 2.4 | 851827.98 |
| birds | Strix_leptogrammica | Strix_leptogrammica | 0 | 970 | 1.4 | 5428569.66 |
| birds | Strix_nebulosa | Strix_nebulosa | 0 | 1061.9 | 4.2 | 19866544.63 |
| birds | Strix_occidentalis | Strix_occidentalis | 0 | 586.4 | 2 | 1222682.34 |
| birds | Strix_rufipes | Strix_rufipes | 0 | 380 | 2.4 | 639116.13 |
| birds | Strix_seloputo | Strix_seloputo | 0 | 1011 | 2.4 | 610935.05 |
| birds | Strix_uralensis | Strix_uralensis | 0 | 780.6 | 2.8 | 12224942.84 |
| birds | Strix_varia | Strix_varia | 0 | 711.5 | 3.2 | 6628563.52 |

| | | | | | | |
|---|---|---|---|---|---|---|
| birds | Strix_woodfordii | Strix_woodfordii | 0 | 350 | 1.7 | 9861983.74 |
| birds | Struthidea_cinerea | Struthidea_cinerea | 0 | 132 | 3.2 | 2500913.56 |
| birds | Struthio_camelus | Struthio_camelus | 0 | 111000 | 7.4 | 9266924.36 |
| birds | Sturnella_magna | Sturnella_magna | 0 | 91.8 | 5 | 7869636.91 |
| birds | Sturnella_neglecta | Sturnella_neglecta | 0 | 100.1 | 4.2 | 8668509.02 |
| birds | Sturnia_malabarica | Sturnus_malabaricus | 0 | 39.6 | 3.9 | 5278207.02 |
| birds | Sturnia_pagodarum | Sturnus_pagodarum | 0 | 48.9 | 3.9 | 3846594.05 |
| birds | Sturnia_sinensis | Sturnus_sinensis | 0 | 61 | 4 | 1582507.91 |
| birds | Sturnus_unicolor | Sturnus_unicolor | 0 | 83.7 | 3.2 | 826392.84 |
| birds | Sturnus_vulgaris | Sturnus_vulgaris | 0 | 77.1 | 5.9 | 40394100.72 |
| birds | Sublegatus_arenarum | Sublegatus_arenarum | 0 | 12.3 | 2 | 940862.17 |
| birds | Sublegatus_modestus | Sublegatus_modestus | 0 | 14 | 2 | 6470238.68 |
| birds | Sugomel_nigrum | Certhionyx_niger | 0 | 9.9 | 2 | 5440427.41 |
| birds | Surnia_ulula | Surnia_ulula | 0 | 319.9 | 7.7 | 21292793.55 |
| birds | Swynnertonia_swynnertoni | Swynnertonia_swynnertoni | 1 | 16 | 2 | 7065.43 |
| birds | Sylvia_atricapilla | Sylvia_atricapilla | 0 | 16.7 | 4.5 | 21049205.58 |
| birds | Sylvia_borin | Sylvia_borin | 0 | 18.2 | 4.5 | 48022839.56 |
| birds | Sylvia_nigricapillus | Lioptilus_nigricapillus | 1 | 29.6 | 2 | 170916.76 |
| birds | Sylvietta_brachyura | Sylvietta_brachyura | 0 | 8.1 | 2 | 5897093.52 |
| birds | Sylvietta_denti | Sylvietta_denti | 0 | 8.4 | 1 | 1130212.97 |
| birds | Sylvietta_isabellina | Sylvietta_isabellina | 0 | 10 | 2 | 901911.55 |
| birds | Sylvietta_leucophrys | Sylvietta_leucophrys | 0 | 11.3 | 2 | 156344.04 |
| birds | Sylvietta_rufescens | Sylvietta_rufescens | 0 | 11.6 | 2 | 4186905.87 |
| birds | Sylvietta_ruficapilla | Sylvietta_ruficapilla | 0 | 11.1 | 2 | 2557018.33 |
| birds | Sylvietta_virens | Sylvietta_virens | 0 | 8.4 | 2 | 3738336.18 |
| birds | Sylvietta_whytii | Sylvietta_whytii | 0 | 9.9 | 2 | 1855770.99 |
| birds | Sylviorthorhynchus_desmurii | Sylviorthorhynchus_desmursii | 0 | 10.9 | 3.5 | 386264.46 |
| birds | Symposiachrus_axillaris | Monarcha_axillaris | 0 | 15.6 | 1 | 118942.75 |
| birds | Symposiachrus_barbatus | Monarcha_barbatus | 0 | 21 | 2 | 21960.62 |
| birds | Symposiachrus_guttula | Monarcha_guttula | 0 | 16.4 | 2 | 657660.08 |
| birds | Symposiachrus_manadensis | Monarcha_manadensis | 0 | 22.5 | 2 | 445281.04 |
| birds | Symposiachrus_trivirgatus | Monarcha_trivirgatus | 0 | 12.7 | 2 | 612420.23 |
| birds | Symposiachrus_verticalis | Monarcha_verticalis | 0 | 21.7 | 2 | 42001.39 |
| birds | Synallaxis_albescens | Synallaxis_albescens | 0 | 11.2 | 2.8 | 9891841.77 |

| | | | | | | |
|---|---|---|---|---|---|---|
| birds | Synallaxis_azarae | Synallaxis_azarae | 0 | 16.9 | 2.8 | 477890.57 |
| birds | Synallaxis_brachyura | Synallaxis_brachyura | 0 | 18.3 | 2.4 | 403853.13 |
| birds | Synallaxis_candei | Synallaxis_candei | 0 | 15.3 | 3.5 | 156033.91 |
| birds | Synallaxis_cinerascens | Synallaxis_cinerascens | 0 | 13.1 | 3 | 1223916.17 |
| birds | Synallaxis_cinnamomea | Synallaxis_cinnamomea | 0 | 18.4 | 3 | 85347.41 |
| birds | Synallaxis_erythrothorax | Synallaxis_erythrothorax | 0 | 17.1 | 2.8 | 355771.38 |
| birds | Synallaxis_frontalis | Synallaxis_frontalis | 0 | 14 | 3.5 | 5745641.87 |
| birds | Synallaxis_gujanensis | Synallaxis_gujanensis | 0 | 18.1 | 2.4 | 4462466.66 |
| birds | Synallaxis_infuscata | Synallaxis_infuscata | 1 | 18.2 | 2 | 25054.16 |
| birds | Synallaxis_ruficapilla | Synallaxis_ruficapilla | 0 | 13.8 | 2.4 | 1174516.01 |
| birds | Synallaxis_rutilans | Synallaxis_rutilans | 0 | 16.7 | 3.5 | 5856806.86 |
| birds | Synallaxis_scutata | Synallaxis_scutata | 0 | 15.4 | 2.4 | 3789577.7 |
| birds | Synallaxis_spixi | Synallaxis_spixi | 0 | 12.6 | 3.9 | 1343787.35 |
| birds | Synallaxis_zimmeri | Synallaxis_zimmeri | 1 | 13 | 2 | 4318.41 |
| birds | Syndactyla_rufosuperciliata | Syndactyla_rufosuperciliata | 0 | 25.6 | 2.8 | 1726834 |
| birds | Synoicus_chinensis | Coturnix_chinensis | 0 | 40.4 | 5.3 | 8165093.61 |
| birds | Synoicus_ypsilophorus | Coturnix_ypsilophora | 0 | 107.7 | 8.8 | 5840602.83 |
| birds | Sypheotides_indicus | Sypheotides_indicus | 1 | 614.3 | 3.9 | 2471338.06 |
| birds | Syrigma_sibilatrix | Syrigma_sibilatrix | 0 | 463 | 3.5 | 4235220.19 |
| birds | Syrrhaptes_paradoxus | Syrrhaptes_paradoxus | 0 | 256.4 | 3 | 8706694.08 |
| birds | Syrrhaptes_tibetanus | Syrrhaptes_tibetanus | 0 | 343.7 | 3 | 1238571.56 |
| birds | Systellura_longirostris | Caprimulgus_longirostris | 0 | 45.6 | 1.4 | 5080733.59 |
| birds | Taccocua_leschenaultii | Phaenicophaeus_leschenaultii | 0 | 174.7 | 2.4 | 2673797.89 |
| birds | Tachuris_rubrigastra | Tachuris_rubrigastra | 0 | 7.8 | 3.5 | 2891313.77 |
| birds | Tachybaptus_dominicus | Tachybaptus_dominicus | 0 | 132.6 | 4.9 | 13082260.67 |
| birds | Tachybaptus_novaehollandiae | Tachybaptus_novaehollandiae | 0 | 219 | 4.5 | 8308868.82 |
| birds | Tachybaptus_pelzelnii | Tachybaptus_pelzelnii | 1 | 184 | 3.5 | 590930.76 |
| birds | Tachybaptus_ruficollis | Tachybaptus_ruficollis | 0 | 169.4 | 3.7 | 37429277.13 |
| birds | Tachycineta_albilinea | Tachycineta_albilinea | 0 | 16.7 | 3.9 | 755005.82 |
| birds | Tachycineta_albiventer | Tachycineta_albiventer | 0 | 17.7 | 4.2 | 11742614.01 |
| birds | Tachycineta_bicolor | Tachycineta_bicolor | 0 | 21.2 | 5.5 | 17478048.63 |
| birds | Tachycineta_cyaneoviridis | Tachycineta_cyaneoviridis | 1 | 17.5 | 3 | 14730.66 |
| birds | Tachycineta_euchrysea | Tachycineta_euchrysea | 1 | 17 | 3 | 14021.59 |
| birds | Tachycineta_leucorrhoa | Tachycineta_leucorrhoa | 0 | 15.3 | 5.3 | 6386014.55 |

| | | | | | | |
|---|---|---|---|---|---|---|
| birds | Tachycineta_meyeni | Tachycineta_meyeni | 0 | 14.9 | 4.2 | 4066377.87 |
| birds | Tachycineta_thalassina | Tachycineta_thalassina | 0 | 14.1 | 4.9 | 6579514.33 |
| birds | Tachymarptis_aequatorialis | Tachymarptis_aequatorialis | 0 | 92.8 | 1.4 | 1399846.87 |
| birds | Tachymarptis_melba | Tachymarptis_melba | 0 | 100.4 | 2 | 12945294.11 |
| birds | Tadorna_cana | Tadorna_cana | 0 | 1182 | 10 | 1631456.11 |
| birds | Tadorna_ferruginea | Tadorna_ferruginea | 0 | 1235 | 8.5 | 24406375.91 |
| birds | Tadorna_tadorna | Tadorna_tadorna | 0 | 1146.8 | 8.9 | 11207269.88 |
| birds | Tadorna_tadornoides | Tadorna_tadornoides | 0 | 1418.7 | 8.9 | 3300063.22 |
| birds | Tadorna_variegata | Tadorna_variegata | 0 | 1541 | 8.9 | 268823.5 |
| birds | Taeniopygia_bichenovii | Taeniopygia_bichenovii | 0 | 9.6 | 4.5 | 3108066.1 |
| birds | Taeniopygia_guttata | Taeniopygia_guttata | 0 | 12 | 4.5 | 83399.73 |
| birds | Talaphorus_chlorocercus | Leucippus_chlorocercus | 0 | 4.6 | 2 | 221320.45 |
| birds | Tangara_arthus | Tangara_arthus | 0 | 22 | 1.4 | 38919.72 |
| birds | Tangara_cayana | Tangara_cayana | 0 | 18 | 2 | 1467704.42 |
| birds | Tangara_chilensis | Tangara_chilensis | 0 | 19.8 | 2 | 4594777.46 |
| birds | Tangara_cyanicollis | Tangara_cyanicollis | 0 | 17 | 2 | 1248379.88 |
| birds | Tangara_cyanocephala | Tangara_cyanocephala | 0 | 18 | 3 | 444844.92 |
| birds | Tangara_desmaresti | Tangara_desmaresti | 0 | 20.4 | 3 | 260465.02 |
| birds | Tangara_dowii | Tangara_dowii | 0 | 15 | 2 | 12083.47 |
| birds | Tangara_florida | Tangara_florida | 0 | 19.3 | 2 | 37066.94 |
| birds | Tangara_guttata | Tangara_guttata | 0 | 18.4 | 2 | 376030.94 |
| birds | Tangara_gyrola | Tangara_gyrola | 0 | 21 | 2 | 4181131.62 |
| birds | Tangara_icterocephala | Tangara_icterocephala | 0 | 22 | 2 | 75164 |
| birds | Tangara_inornata | Tangara_inornata | 0 | 18 | 2 | 180601.4 |
| birds | Tangara_larvata | Tangara_larvata | 0 | 20 | 2 | 480330.12 |
| birds | Tangara_lavinia | Tangara_lavinia | 0 | 24 | 3 | 135531.53 |
| birds | Tangara_mexicana | Tangara_mexicana | 0 | 20.5 | 2.4 | 7040026.36 |
| birds | Tangara_nigrocincta | Tangara_nigrocincta | 0 | 17 | 2 | 3416902.14 |
| birds | Tangara_ruficervix | Tangara_ruficervix | 0 | 19 | 2 | 134578.1 |
| birds | Tangara_schrankii | Tangara_schrankii | 0 | 19 | 2 | 2751315.13 |
| birds | Tangara_seledon | Tangara_seledon | 0 | 18.7 | 3 | 672563.84 |
| birds | Tangara_vitriolina | Tangara_vitriolina | 0 | 23 | 2 | 123913.11 |
| birds | Tangara_xanthocephala | Tangara_xanthocephala | 0 | 19 | 2 | 356998.99 |
| birds | Taphrospilus_hypostictus | Taphrospilus_hypostictus | 0 | 7 | 2 | 110112.14 |

| | | | | | | |
|---|---|---|---|---|---|---|
| birds | Taraba_major | Taraba_major | 0 | 59.2 | 2.4 | 12899627.84 |
| birds | Tarphonomus_certhioides | Tarphonomus_certhioides | 0 | 22.1 | 3.5 | 1267589.78 |
| birds | Tarsiger_chrysaeus | Tarsiger_chrysaeus | 0 | 13.8 | 3.5 | 1061370 |
| birds | Tarsiger_cyanurus | Tarsiger_cyanurus | 0 | 13.4 | 5.9 | 13872524.62 |
| birds | Tarsiger_indicus | Tarsiger_indicus | 0 | 14.6 | 3.5 | 533602.16 |
| birds | Tauraco_bannermani | Tauraco_bannermani | 1 | 224 | 2 | 1978.6 |
| birds | Tauraco_corythaix | Tauraco_corythaix | 0 | 308 | 2.4 | 124836.47 |
| birds | Tauraco_fischeri | Tauraco_fischeri | 0 | 250 | 2 | 48655.56 |
| birds | Tauraco_hartlaubi | Tauraco_hartlaubi | 0 | 224 | 2 | 264805.15 |
| birds | Tauraco_leucolophus | Tauraco_leucolophus | 0 | 219 | 2 | 1433202.57 |
| birds | Tauraco_leucotis | Tauraco_leucotis | 0 | 265 | 2 | 807016.24 |
| birds | Tauraco_livingstonii | Tauraco_livingstonii | 0 | 249 | 2 | 974104.7 |
| birds | Tauraco_macrorhynchus | Tauraco_macrorhynchus | 0 | 243.7 | 2 | 1273570.69 |
| birds | Tauraco_persa | Tauraco_persa | 0 | 306 | 2 | 3260169.31 |
| birds | Tauraco_schalowi | Tauraco_schalowi | 0 | 206.4 | 2 | 1899419.96 |
| birds | Tauraco_schuettii | Tauraco_schuetti | 0 | 235 | 2 | 2085207.97 |
| birds | Tchagra_australis | Tchagra_australis | 0 | 32.3 | 2.4 | 8305150.13 |
| birds | Tchagra_jamesi | Tchagra_jamesi | 0 | 27.4 | 2.4 | 1196428.5 |
| birds | Tchagra_senegalus | Tchagra_senegalus | 0 | 53.5 | 2.4 | 14953687.11 |
| birds | Tchagra_tchagra | Tchagra_tchagra | 0 | 50.3 | 2 | 251680.9 |
| birds | Telacanthura_ussheri | Telacanthura_ussheri | 0 | 32.6 | 4 | 5188546 |
| birds | Teledromas_fuscus | Teledromas_fuscus | 0 | 37.9 | 2 | 521735.99 |
| birds | Telophorus_dohertyi | Telophorus_dohertyi | 0 | 35 | 3 | 113100.74 |
| birds | Telophorus_viridis | Telophorus_viridis | 0 | 37.1 | 2 | 1025110.51 |
| birds | Telophorus_zeylonus | Telophorus_zeylonus | 0 | 65 | 3 | 1314512.43 |
| birds | Temnurus_temnurus | Temnurus_temnurus | 0 | 138 | 2 | 266868.95 |
| birds | Tephrodornis_pondicerianus | Tephrodornis_pondicerianus | 0 | 20.2 | 3 | 4411317.05 |
| birds | Terathopius_ecaudatus | Terathopius_ecaudatus | 1 | 2200 | 1 | 14023477.79 |
| birds | Terenotriccus_erythrurus | Terenotriccus_erythrurus | 0 | 7.4 | 2 | 7494686.86 |
| birds | Teretistris_fernandinae | Teretistris_fernandinae | 0 | 9.9 | 2.4 | 25700.55 |
| birds | Teretistris_fornsi | Teretistris_fornsi | 0 | 9.7 | 2.4 | 38336.73 |
| birds | Terpsiphone_atrocaudata | Terpsiphone_atrocaudata | 0 | 18.7 | 3.9 | 2897084.96 |
| birds | Terpsiphone_bedfordi | Terpsiphone_bedfordi | 0 | 15.4 | 2 | 42552.86 |
| birds | Terpsiphone_bourbonnensis | Terpsiphone_bourbonnensis | 0 | 10.8 | 2.4 | 3297.84 |

| | | | | | | |
|---|---|---|---|---|---|---|
| birds | Terpsiphone_cinnamomea | Terpsiphone_cinnamomea | 0 | 20.4 | 3 | 109426.27 |
| birds | Terpsiphone_mutata | Terpsiphone_mutata | 0 | 13.2 | 3 | 592893.86 |
| birds | Terpsiphone_paradisi | Terpsiphone_paradisi | 0 | 19.2 | 3.5 | 4205670.73 |
| birds | Terpsiphone_rufiventer | Terpsiphone_rufiventer | 0 | 15.1 | 2.4 | 4275150.97 |
| birds | Terpsiphone_rufocinerea | Terpsiphone_rufocinerea | 0 | 15.4 | 1.4 | 444393.52 |
| birds | Terpsiphone_viridis | Terpsiphone_viridis | 0 | 13.2 | 2.2 | 14483913.96 |
| birds | Tesia_cyaniventer | Tesia_cyaniventer | 0 | 9.7 | 3.9 | 2098380.83 |
| birds | Tetrao_urogalloides | Tetrao_parvirostris | 0 | 2752.3 | 6.5 | 7003788.43 |
| birds | Tetrao_urogallus | Tetrao_urogallus | 0 | 2716.6 | 7.3 | 11330230.84 |
| birds | Tetrastes_bonasia | Bonasa_bonasia | 0 | 429 | 8.8 | 15669355.23 |
| birds | Tetrastes_sewerzowi | Bonasa_sewerzowi | 0 | 267.3 | 6.3 | 394309.68 |
| birds | Tetrax_tetrax | Tetrax_tetrax | 0 | 833.5 | 3.5 | 10243202.49 |
| birds | Thalurania_colombica | Thalurania_colombica | 0 | 4.5 | 2 | 705818.09 |
| birds | Thalurania_glaucopis | Thalurania_glaucopis | 0 | 4.8 | 2 | 1415794.42 |
| birds | Thalurania_watertonii | Thalurania_watertonii | 1 | 4.6 | 2 | 44734.57 |
| birds | Thamnistes_anabatinus | Thamnistes_anabatinus | 0 | 20.7 | 2 | 398111.82 |
| birds | Thamnolaea_cinnamomeiventris | Myrmecocichla_cinnamomeiventris | 0 | 39.8 | 2.8 | 2978092.64 |
| birds | Thamnomanes_ardesiacus | Thamnomanes_ardesiacus | 0 | 17.7 | 2 | 3380827.14 |
| birds | Thamnomanes_caesius | Thamnomanes_caesius | 0 | 15.7 | 2 | 6049355.84 |
| birds | Thamnomanes_schistogynus | Thamnomanes_schistogynus | 0 | 17.2 | 2 | 1226196.42 |
| birds | Thamnophilus_aethiops | Thamnophilus_aethiops | 0 | 25.7 | 2 | 4946417.18 |
| birds | Thamnophilus_amazonicus | Thamnophilus_amazonicus | 0 | 18.7 | 2 | 6192801.73 |
| birds | Thamnophilus_atrinucha | Thamnophilus_atrinucha | 0 | 23.6 | 2 | 583507.07 |
| birds | Thamnophilus_bernardi | Sakesphorus_bernardi | 0 | 34.4 | 2.4 | 102383.98 |
| birds | Thamnophilus_bridgesi | Thamnophilus_bridgesi | 0 | 27 | 2.4 | 28247.58 |
| birds | Thamnophilus_caerulescens | Thamnophilus_caerulescens | 0 | 21.1 | 2.4 | 3969673.89 |
| birds | Thamnophilus_doliatus | Thamnophilus_doliatus | 0 | 27 | 1.7 | 10725968.4 |
| birds | Thamnophilus_melanonotus | Sakesphorus_melanonotus | 0 | 25.5 | 2 | 113947.78 |
| birds | Thamnophilus_murinus | Thamnophilus_murinus | 0 | 19.3 | 2 | 4198741.89 |
| birds | Thamnophilus_nigrocinereus | Thamnophilus_nigrocinereus | 0 | 30 | 2 | 1299292.69 |
| birds | Thamnophilus_palliatus | Thamnophilus_palliatus | 0 | 27 | 2 | 2542297.14 |
| birds | Thamnophilus_punctatus | Thamnophilus_punctatus | 0 | 19.4 | 2 | 1569761.66 |
| birds | Thamnophilus_ruficapillus | Thamnophilus_ruficapillus | 0 | 20.4 | 2.4 | 1391729.07 |
| birds | Thamnophilus_schistaceus | Thamnophilus_schistaceus | 0 | 20.3 | 2 | 3991785.91 |

| | | | | | | |
|---|---|---|---|---|---|---|
| birds | Thamnophilus_tenuepunctatus | Thamnophilus_tenuepunctatus | 0 | 23.3 | 2 | 136115.51 |
| birds | Thamnophilus_torquatus | Thamnophilus_torquatus | 0 | 21.1 | 2.4 | 3732484.24 |
| birds | Thamnornis_chloropetoides | Thamnornis_chloropetoides | 0 | 14.3 | 3 | 69624.44 |
| birds | Thaumastura_cora | Thaumastura_cora | 0 | 1.9 | 2 | 203316.51 |
| birds | Thaumatibis_gigantea | Thaumatibis_gigantea | 1 | 3515 | 2.4 | 50313.06 |
| birds | Theristicus_caerulescens | Theristicus_caerulescens | 0 | 1500 | 2.4 | 1749653.63 |
| birds | Theristicus_caudatus | Theristicus_caudatus | 0 | 1726 | 2.8 | 5146136.12 |
| birds | Theristicus_melanopis | Theristicus_melanopis | 0 | 1250 | 2.4 | 2020059.98 |
| birds | Thinocorus_orbignyianus | Thinocorus_orbignyianus | 0 | 115 | 4 | 1770605.52 |
| birds | Thinocorus_rumicivorus | Thinocorus_rumicivorus | 0 | 53.1 | 4 | 2241860.15 |
| birds | Threnetes_niger | Threnetes_niger | 0 | 5.6 | 2 | 313115.48 |
| birds | Threnetes_ruckeri | Threnetes_ruckeri | 0 | 5.2 | 2 | 482644.97 |
| birds | Threskiornis_aethiopicus | Threskiornis_aethiopicus | 0 | 1493.2 | 2.4 | 17731713.86 |
| birds | Threskiornis_melanocephalus | Threskiornis_melanocephalus | 0 | 1303 | 2.8 | 2840531.68 |
| birds | Threskiornis_moluccus | Threskiornis_molucca | 0 | 1794.9 | 2 | 5697443.52 |
| birds | Threskiornis_spinicollis | Threskiornis_spinicollis | 0 | 1346.7 | 3.2 | 6556820.35 |
| birds | Thripadectes_melanorhynchus | Thripadectes_melanorhynchus | 0 | 44.3 | 3 | 121002.94 |
| birds | Thripadectes_rufobrunneus | Thripadectes_rufobrunneus | 0 | 56.2 | 2 | 17593.39 |
| birds | Thripadectes_virgaticeps | Thripadectes_virgaticeps | 0 | 65.8 | 2 | 62554.86 |
| birds | Thryomanes_bewickii | Thryomanes_bewickii | 0 | 9.9 | 5.5 | 3814727.24 |
| birds | Thryophilus_pleurostictus | Thryothorus_pleurostictus | 0 | 17.7 | 3.2 | 185486.45 |
| birds | Thryophilus_rufalbus | Thryothorus_rufalbus | 0 | 24.9 | 2.8 | 821636.36 |
| birds | Thryophilus_sinaloa | Thryothorus_sinaloa | 0 | 15.1 | 5 | 304927.47 |
| birds | Thryorchilus_browni | Thryorchilus_browni | 0 | 14 | 2 | 1835.35 |
| birds | Thryothorus_ludovicianus | Thryothorus_ludovicianus | 0 | 19 | 4.2 | 3040027.22 |
| birds | Tichodroma_muraria | Tichodroma_muraria | 0 | 17.6 | 4.5 | 9663228.8 |
| birds | Tigriornis_leucolopha | Tigriornis_leucolopha | 0 | 846 | 1 | 3869151.95 |
| birds | Tigrisoma_lineatum | Tigrisoma_lineatum | 0 | 813 | 3 | 13239678.28 |
| birds | Tigrisoma_mexicanum | Tigrisoma_mexicanum | 0 | 1154.4 | 1.7 | 928353.01 |
| birds | Timeliopsis_fulvigula | Timeliopsis_fulvigula | 0 | 18.3 | 1 | 137140.89 |
| birds | Tinamotis_ingoufi | Tinamotis_ingoufi | 0 | 950 | 11 | 403733.45 |
| birds | Tinamotis_pentlandii | Tinamotis_pentlandii | 0 | 650 | 5.3 | 632710.16 |
| birds | Tinamus_guttatus | Tinamus_guttatus | 0 | 686.2 | 5.5 | 3971186.8 |
| birds | Tinamus_major | Tinamus_major | 0 | 1026.2 | 4.2 | 8123168.48 |

| | | | | | | |
|---|---|---|---|---|---|---|
| birds | Tinamus_solitarius | Tinamus_solitarius | 0 | 1386.4 | 9.2 | 1069608.94 |
| birds | Tinamus_tao | Tinamus_tao | 1 | 1600.1 | 4.2 | 3810484.55 |
| birds | Tityra_cayana | Tityra_cayana | 0 | 68.1 | 3 | 5115788.26 |
| birds | Tityra_inquisitor | Tityra_inquisitor | 0 | 43.1 | 3 | 11190798.84 |
| birds | Tityra_semifasciata | Tityra_semifasciata | 0 | 79.3 | 2.4 | 6821138.58 |
| birds | Tmetothylacus_tenellus | Tmetothylacus_tenellus | 0 | 20.1 | 2.8 | 1285795.15 |
| birds | Tockus_deckeni | Tockus_deckeni | 0 | 167.7 | 2.8 | 1628868.18 |
| birds | Tockus_erythrorhynchus | Tockus_erythrorhynchus | 0 | 138.6 | 3.7 | 9150146.92 |
| birds | Tockus_flavirostris | Tockus_flavirostris | 0 | 216.7 | 2.4 | 1449740.42 |
| birds | Tockus_leucomelas | Tockus_leucomelas | 0 | 188.3 | 3.5 | 1820119.32 |
| birds | Tockus_monteiri | Tockus_monteiri | 0 | 289.8 | 4 | 399497.56 |
| birds | Todiramphus_albonotatus | Todiramphus_albonotatus | 0 | 32 | 2.4 | 30015.58 |
| birds | Todiramphus_chloris | Todiramphus_chloris | 0 | 66.1 | 3.2 | 3828153.1 |
| birds | Todiramphus_farquhari | Todiramphus_farquhari | 0 | 38.8 | 3 | 5637.52 |
| birds | Todiramphus_leucopygius | Todiramphus_leucopygius | 0 | 47.9 | 3 | 23194.38 |
| birds | Todiramphus_macleayii | Todiramphus_macleayii | 0 | 37.3 | 4.2 | 1316178.4 |
| birds | Todiramphus_pyrrhopygius | Todiramphus_pyrrhopygius | 0 | 51.7 | 4.5 | 7435153.36 |
| birds | Todiramphus_sanctus | Todiramphus_sanctus | 0 | 53 | 4.5 | 8193818.92 |
| birds | Todiramphus_saurophagus | Todiramphus_saurophaga | 0 | 126 | 2.8 | 134276.08 |
| birds | Todirostrum_chrysocrotaphum | Todirostrum_chrysocrotaphum | 0 | 7 | 2 | 4205502.11 |
| birds | Todirostrum_cinereum | Todirostrum_cinereum | 0 | 6.3 | 2.4 | 8243668.88 |
| birds | Todirostrum_maculatum | Todirostrum_maculatum | 0 | 7.3 | 1.4 | 4363496.68 |
| birds | Todirostrum_nigriceps | Todirostrum_nigriceps | 0 | 6.5 | 1 | 423081.58 |
| birds | Tolmomyias_flaviventris | Tolmomyias_flaviventris | 0 | 12.2 | 2.4 | 7836566.05 |
| birds | Tolmomyias_sulphurescens | Tolmomyias_sulphurescens | 0 | 14.3 | 2.4 | 12413249.74 |
| birds | Topaza_pella | Topaza_pella | 0 | 12.1 | 2 | 1678442.91 |
| birds | Torgos_tracheliotos | Torgos_tracheliotos | 1 | 6969 | 1 | 14737739.36 |
| birds | Torreornis_inexpectata | Torreornis_inexpectata | 0 | 25.5 | 1.4 | 4568.06 |
| birds | Touit_batavicus | Touit_batavicus | 0 | 55 | 5.5 | 333487.33 |
| birds | Touit_costaricensis | Touit_costaricensis | 0 | 80 | 2.4 | 19135.59 |
| birds | Touit_purpuratus | Touit_purpuratus | 0 | 59.7 | 3.9 | 3821297.27 |
| birds | Toxorhamphus_poliopterus | Toxorhamphus_poliopterus | 0 | 11.4 | 1 | 179355.79 |
| birds | Toxostoma_bendirei | Toxostoma_bendirei | 1 | 62.2 | 3.5 | 822797.04 |
| birds | Toxostoma_cinereum | Toxostoma_cinereum | 0 | 59 | 2.4 | 99643.4 |

| | | | | | | |
|---|---|---|---|---|---|---|
| birds | Toxostoma_crissale | Toxostoma_crissale | 0 | 62.7 | 2.4 | 1010689.86 |
| birds | Toxostoma_curvirostre | Toxostoma_curvirostre | 0 | 80.5 | 3.9 | 2044910.56 |
| birds | Toxostoma_lecontei | Toxostoma_lecontei | 0 | 61.9 | 3.2 | 251262.3 |
| birds | Toxostoma_longirostre | Toxostoma_longirostre | 0 | 67.6 | 3.9 | 300961.71 |
| birds | Toxostoma_ocellatum | Toxostoma_ocellatum | 0 | 84.3 | 2 | 88725.79 |
| birds | Toxostoma_redivivum | Toxostoma_redivivum | 0 | 84.4 | 3.5 | 167320.21 |
| birds | Toxostoma_rufum | Toxostoma_rufum | 0 | 68.8 | 3.9 | 5859124.47 |
| birds | Trachylaemus_purpuratus | Trachyphonus_purpuratus | 0 | 76.1 | 2.8 | 3484026.33 |
| birds | Trachyphonus_darnaudii | Trachyphonus_darnaudii | 0 | 35.8 | 2.8 | 1278810.54 |
| birds | Trachyphonus_erythrocephalus | Trachyphonus_erythrocephalus | 0 | 63.5 | 3.5 | 1574108.27 |
| birds | Trachyphonus_margaritatus | Trachyphonus_margaritatus | 0 | 64 | 4.2 | 2214289.93 |
| birds | Trachyphonus_vaillantii | Trachyphonus_vaillantii | 0 | 69.4 | 3.2 | 3120428.04 |
| birds | Tragopan_blythii | Tragopan_blythii | 1 | 1930 | 3.2 | 120944.17 |
| birds | Tragopan_caboti | Tragopan_caboti | 1 | 1122.5 | 3.9 | 396418.35 |
| birds | Tragopan_melanocephalus | Tragopan_melanocephalus | 1 | 1613.2 | 4.2 | 21593.78 |
| birds | Tragopan_satyra | Tragopan_satyra | 0 | 1417 | 2.4 | 99176.64 |
| birds | Tragopan_temminckii | Tragopan_temminckii | 0 | 1163.4 | 3.9 | 1261852.14 |
| birds | Tregellasia_capito | Tregellasia_capito | 0 | 13.9 | 2 | 155466.5 |
| birds | Tregellasia_leucops | Tregellasia_leucops | 0 | 15.6 | 1.4 | 201214.41 |
| birds | Treron_apicauda | Treron_apicauda | 0 | 217.2 | 2 | 1636420.93 |
| birds | Treron_australis | Treron_australis | 0 | 229 | 2 | 505554.32 |
| birds | Treron_calvus | Treron_calvus | 0 | 197.5 | 2 | 10739326.33 |
| birds | Treron_formosae | Treron_formosae | 0 | 140 | 2 | 36703.57 |
| birds | Treron_fulvicollis | Treron_fulvicollis | 1 | 167 | 2 | 932712.23 |
| birds | Treron_griseicauda | Treron_griseicauda | 0 | 205.7 | 2 | 245187.84 |
| birds | Treron_phoenicopterus | Treron_phoenicopterus | 0 | 235 | 2 | 4843995.11 |
| birds | Treron_sieboldii | Treron_sieboldii | 0 | 245 | 2 | 2159328.24 |
| birds | Treron_sphenurus | Treron_sphenurus | 0 | 210 | 1.4 | 1739910.38 |
| birds | Treron_waalia | Treron_waalia | 0 | 254 | 1.4 | 5454383.59 |
| birds | Tribonyx_mortierii | Gallinula_mortierii | 0 | 1291.8 | 6.3 | 42670.17 |
| birds | Tribonyx_ventralis | Gallinula_ventralis | 0 | 386.3 | 5.9 | 4599237.18 |
| birds | Trichoglossus_chlorolepidotus | Trichoglossus_chlorolepidotus | 0 | 87.1 | 2 | 1109928.47 |
| birds | Trichoglossus_euteles | Trichoglossus_euteles | 0 | 73.7 | 3 | 37893.11 |
| birds | Trichoglossus_flavoviridis | Trichoglossus_flavoviridis | 0 | 90.5 | 2 | 4102.71 |

| | | | | | | |
|---|---|---|---|---|---|---|
| birds | Trichoglossus_haematodus | Trichoglossus_haematodus | 0 | 113.3 | 1.7 | 876908.04 |
| birds | Trichoglossus_johnstoniae | Trichoglossus_johnstoniae | 0 | 55.2 | 2 | 8694.55 |
| birds | Trichoglossus_ornatus | Trichoglossus_ornatus | 0 | 120 | 2 | 172993.28 |
| birds | Tricholaema_diademata | Tricholaema_diademata | 0 | 30.6 | 2.8 | 578757.82 |
| birds | Tricholaema_frontata | Tricholaema_frontata | 0 | 26.8 | 2.4 | 1375938.26 |
| birds | Tricholaema_lacrymosa | Tricholaema_lacrymosa | 0 | 22.9 | 2.8 | 1269448.61 |
| birds | Tricholaema_leucomelas | Tricholaema_leucomelas | 0 | 32.2 | 2 | 3123368.53 |
| birds | Tricholaema_melanocephala | Tricholaema_melanocephala | 0 | 20.5 | 2.4 | 1969018.58 |
| birds | Trichothraupis_melanops | Trichothraupis_melanops | 0 | 22.6 | 3 | 2473090.35 |
| birds | Triclaria_malachitacea | Triclaria_malachitacea | 0 | 90 | 2.8 | 99479.82 |
| birds | Trigonoceps_occipitalis | Trigonoceps_occipitalis | 1 | 3016 | 1 | 11472482.17 |
| birds | Tringa_brevipes | Heteroscelus_brevipes | 0 | 127 | 4 | 4918509.31 |
| birds | Tringa_erythropus | Tringa_erythropus | 0 | 158 | 4 | 17465707.92 |
| birds | Tringa_flavipes | Tringa_flavipes | 0 | 77.5 | 4 | 35216181.96 |
| birds | Tringa_glareola | Tringa_glareola | 0 | 62.1 | 4 | 47087120.54 |
| birds | Tringa_guttifer | Tringa_guttifer | 1 | 158 | 4 | 413621.64 |
| birds | Tringa_incana | Heteroscelus_incanus | 0 | 108.2 | 4 | 2865961.57 |
| birds | Tringa_melanoleuca | Tringa_melanoleuca | 0 | 161.7 | 4 | 34094264.54 |
| birds | Tringa_nebularia | Tringa_nebularia | 0 | 187 | 4 | 46520227.98 |
| birds | Tringa_ochropus | Tringa_ochropus | 0 | 71.4 | 4 | 43983761.47 |
| birds | Tringa_solitaria | Tringa_solitaria | 0 | 48.4 | 4 | 29415391.2 |
| birds | Tringa_stagnatilis | Tringa_stagnatilis | 0 | 77.5 | 4 | 33164442.82 |
| birds | Tringa_totanus | Tringa_totanus | 0 | 129 | 4 | 46612790.94 |
| birds | Trochalopteron_affine | Garrulax_affinis | 0 | 72.7 | 2.4 | 740183.99 |
| birds | Trochalopteron_austeni | Garrulax_austeni | 0 | 59 | 3.5 | 54762.06 |
| birds | Trochalopteron_erythrocephalum | Garrulax_erythrocephalus | 0 | 71.7 | 2.4 | 334402.42 |
| birds | Trochalopteron_henrici | Garrulax_henrici | 0 | 65 | 2.4 | 151106.19 |
| birds | Trochalopteron_lineatum | Garrulax_lineatus | 0 | 40.7 | 3 | 395193.54 |
| birds | Trochalopteron_milnei | Garrulax_milnei | 0 | 79.5 | 2.4 | 1489338.52 |
| birds | Trochalopteron_squamatum | Garrulax_squamatus | 0 | 84 | 2.4 | 639106.38 |
| birds | Trochalopteron_subunicolor | Garrulax_subunicolor | 0 | 65.9 | 3.5 | 340826.36 |
| birds | Trochalopteron_variegatum | Garrulax_variegatus | 0 | 64.5 | 3 | 281662.13 |
| birds | Trochalopteron_virgatum | Garrulax_virgatus | 0 | 48 | 2.4 | 112099.79 |
| birds | Trochilus_polytmus | Trochilus_polytmus | 0 | 4.8 | 2 | 10224.99 |

| | | | | | | |
|---|---|---|---|---|---|---|
| birds | Trochocercus_cyanomelas | Trochocercus_cyanomelas | 0 | 10.2 | 2.4 | 179874.79 |
| birds | Trochocercus_nitens | Trochocercus_nitens | 0 | 11.8 | 2 | 3680877.59 |
| birds | Troglodytes_aedon | Troglodytes_aedon | 0 | 10.9 | 5.7 | 28997428.72 |
| birds | Troglodytes_cobbi | Troglodytes_cobbi | 0 | 19.1 | 3.5 | 283.46 |
| birds | Troglodytes_ochraceus | Troglodytes_ochraceus | 0 | 9.4 | 2 | 17334.67 |
| birds | Troglodytes_rufociliatus | Troglodytes_rufociliatus | 0 | 10 | 3 | 135313.92 |
| birds | Troglodytes_solstitialis | Troglodytes_solstitialis | 0 | 11.4 | 2 | 553583.66 |
| birds | Troglodytes_troglodytes | Troglodytes_troglodytes | 0 | 9.7 | 6.3 | 18137186.27 |
| birds | Trogon_bairdii | Trogon_bairdii | 0 | 94.4 | 2.4 | 12068.83 |
| birds | Trogon_citreolus | Trogon_citreolus | 0 | 79 | 2.8 | 124015.58 |
| birds | Trogon_collaris | Trogon_collaris | 0 | 65.4 | 2.4 | 6799060.77 |
| birds | Trogon_comptus | Trogon_comptus | 0 | 114 | 3.5 | 79435.54 |
| birds | Trogon_curucui | Trogon_curucui | 0 | 54 | 2.4 | 7396318.69 |
| birds | Trogon_elegans | Trogon_elegans | 0 | 70.9 | 2.4 | 205850.35 |
| birds | Trogon_massena | Trogon_massena | 0 | 141 | 3 | 493358.7 |
| birds | Trogon_melanocephalus | Trogon_melanocephalus | 0 | 85.1 | 3 | 443923.31 |
| birds | Trogon_melanurus | Trogon_melanurus | 0 | 114 | 2.4 | 6755571.08 |
| birds | Trogon_mexicanus | Trogon_mexicanus | 0 | 71 | 2.4 | 375078.57 |
| birds | Trogon_personatus | Trogon_personatus | 0 | 63.4 | 2 | 745904.62 |
| birds | Trogon_rufus | Trogon_rufus | 0 | 53.8 | 2 | 7239946.91 |
| birds | Trogon_violaceus | Trogon_violaceus | 0 | 46.5 | 2.4 | 7284998.29 |
| birds | Trogon_viridis | Trogon_viridis | 0 | 89.7 | 2.4 | 7907953.76 |
| birds | Trugon_terrestris | Trugon_terrestris | 0 | 400 | 1 | 651761.91 |
| birds | Turdoides_bicolor | Turdoides_bicolor | 0 | 78.3 | 3.2 | 1132138.95 |
| birds | Turdoides_gymnogenys | Turdoides_gymnogenys | 0 | 77 | 2 | 411624.93 |
| birds | Turdoides_hartlaubii | Turdoides_hartlaubii | 0 | 86.2 | 3.2 | 1716629.63 |
| birds | Turdoides_hindei | Turdoides_hindei | 1 | 67.6 | 2.4 | 24091.46 |
| birds | Turdoides_hypoleuca | Turdoides_hypoleuca | 0 | 74 | 3 | 89042.46 |
| birds | Turdoides_jardineii | Turdoides_jardineii | 0 | 78.2 | 3.2 | 5822352.5 |
| birds | Turdoides_leucopygia | Turdoides_leucopygia | 0 | 82.4 | 3.2 | 693637.92 |
| birds | Turdoides_melanops | Turdoides_melanops | 0 | 77 | 3.2 | 224027.26 |
| birds | Turdoides_plebejus | Turdoides_plebejus | 0 | 62.5 | 2.8 | 5173060.81 |
| birds | Turdoides_reinwardtii | Turdoides_reinwardii | 0 | 78.8 | 2.4 | 2152069.64 |
| birds | Turdoides_sharpei | Turdoides_sharpei | 0 | 79.3 | 3.2 | 532508.01 |

| | | | | | | |
|---|---|---|---|---|---|---|
| birds | Turdoides_tenebrosa | Turdoides_tenebrosa | 0 | 76 | 2.4 | 294743.09 |
| birds | Turdus_albicollis | Turdus_albicollis | 0 | 54 | 2.4 | 8533196.81 |
| birds | Turdus_albocinctus | Turdus_albocinctus | 0 | 99.4 | 3.5 | 438247.5 |
| birds | Turdus_amaurochalinus | Turdus_amaurochalinus | 0 | 57.9 | 3 | 8248771.08 |
| birds | Turdus_assimilis | Turdus_assimilis | 0 | 70.2 | 2.4 | 548279.87 |
| birds | Turdus_aurantius | Turdus_aurantius | 0 | 81.9 | 2.8 | 11021.79 |
| birds | Turdus_boulboul | Turdus_boulboul | 0 | 97.1 | 3.5 | 1054926.29 |
| birds | Turdus_cardis | Turdus_cardis | 0 | 58.9 | 4 | 1354131.51 |
| birds | Turdus_chrysolaus | Turdus_chrysolaus | 0 | 65.4 | 3.2 | 824574.83 |
| birds | Turdus_dissimilis | Turdus_dissimilis | 0 | 75 | 3.5 | 820644.32 |
| birds | Turdus_falcklandii | Turdus_falcklandii | 0 | 93.9 | 2.4 | 987597.35 |
| birds | Turdus_feae | Turdus_feae | 1 | 71 | 4.5 | 2133337.98 |
| birds | Turdus_fumigatus | Turdus_fumigatus | 0 | 75.7 | 2.4 | 4070825.39 |
| birds | Turdus_fuscater | Turdus_fuscater | 0 | 143 | 2 | 411278.64 |
| birds | Turdus_grayi | Turdus_grayi | 0 | 79.5 | 2.4 | 1125160.83 |
| birds | Turdus_helleri | Turdus_helleri | 1 | 68.7 | 2.4 | 5.37 |
| birds | Turdus_hortulorum | Turdus_hortulorum | 0 | 66.7 | 4.5 | 4088520.4 |
| birds | Turdus_ignobilis | Turdus_ignobilis | 0 | 63.8 | 2 | 272969.91 |
| birds | Turdus_iliacus | Turdus_iliacus | 0 | 61.2 | 4.9 | 23032956.13 |
| birds | Turdus_jamaicensis | Turdus_jamaicensis | 0 | 59.1 | 2.4 | 11021.79 |
| birds | Turdus_leucomelas | Turdus_leucomelas | 0 | 69.1 | 2.4 | 7690463.38 |
| birds | Turdus_libonyana | Turdus_libonyanus | 0 | 60.6 | 2.4 | 4390032.16 |
| birds | Turdus_ludoviciae | Turdus_ludoviciae | 0 | 62 | 2.4 | 12100.21 |
| birds | Turdus_maculirostris | Turdus_maculirostris | 0 | 69.6 | 3 | 88017.54 |
| birds | Turdus_menachensis | Turdus_menachensis | 0 | 75.4 | 2 | 66735.6 |
| birds | Turdus_merula | Turdus_merula | 0 | 102.7 | 3.9 | 14326330.94 |
| birds | Turdus_migratorius | Turdus_migratorius | 0 | 78.5 | 3.5 | 17922995.71 |
| birds | Turdus_naumanni | Turdus_naumanni | 0 | 73.2 | 3.9 | 4484419.71 |
| birds | Turdus_nigrescens | Turdus_nigrescens | 0 | 96 | 2 | 2072.87 |
| birds | Turdus_nigriceps | Turdus_nigriceps | 0 | 52.7 | 3 | 655026.18 |
| birds | Turdus_nudigenis | Turdus_nudigenis | 0 | 63.9 | 2.4 | 1554169.64 |
| birds | Turdus_obscurus | Turdus_obscurus | 0 | 62.6 | 4.9 | 13237348.7 |
| birds | Turdus_obsoletus | Turdus_obsoletus | 0 | 74.8 | 2.4 | 247412.36 |
| birds | Turdus_olivaceus | Turdus_olivaceus | 0 | 67.3 | 2.4 | 399268.73 |

| | | | | | | |
|---|---|---|---|---|---|---|
| birds | Turdus_olivater | Turdus_olivater | 0 | 86.6 | 2.4 | 227974.73 |
| birds | Turdus_pallidus | Turdus_pallidus | 0 | 72.1 | 4.5 | 3500844.87 |
| birds | Turdus_pelios | Turdus_pelios | 0 | 65.4 | 2.4 | 8376209.44 |
| birds | Turdus_philomelos | Turdus_philomelos | 0 | 67.7 | 3.9 | 17749014.34 |
| birds | Turdus_pilaris | Turdus_pilaris | 0 | 106 | 5.5 | 20398397.01 |
| birds | Turdus_plebejus | Turdus_plebejus | 0 | 86.3 | 2.4 | 160186.83 |
| birds | Turdus_plumbeus | Turdus_plumbeus | 0 | 75.3 | 3.9 | 7073.93 |
| birds | Turdus_poliocephalus | Turdus_poliocephalus | 0 | 61.6 | 1.7 | 246956.22 |
| birds | Turdus_rubrocanus | Turdus_rubrocanus | 0 | 92.2 | 2.8 | 1637844.19 |
| birds | Turdus_ruficollis | Turdus_ruficollis | 0 | 80.6 | 4.5 | 7170455.49 |
| birds | Turdus_rufitorques | Turdus_rufitorques | 0 | 72 | 2.4 | 76537.37 |
| birds | Turdus_rufiventris | Turdus_rufiventris | 0 | 69.4 | 2.4 | 5028683.78 |
| birds | Turdus_serranus | Turdus_serranus | 0 | 84.9 | 2 | 408915.41 |
| birds | Turdus_swalesi | Turdus_swalesi | 1 | 75 | 2.4 | 4953.98 |
| birds | Turdus_tephronotus | Turdus_tephronotus | 0 | 49.4 | 2.4 | 998012.21 |
| birds | Turdus_torquatus | Turdus_torquatus | 0 | 109 | 4.5 | 7426107.99 |
| birds | Turdus_unicolor | Turdus_unicolor | 0 | 64.2 | 3.5 | 1400167.39 |
| birds | Turdus_viscivorus | Turdus_viscivorus | 0 | 117.4 | 3.9 | 15469229.78 |
| birds | Turnix_castanotus | Turnix_castanotus | 0 | 82.7 | 4 | 432722.67 |
| birds | Turnix_hottentottus | Turnix_hottentottus | 0 | 50.6 | 3 | 58126.33 |
| birds | Turnix_maculosus | Turnix_maculosus | 0 | 39.1 | 4 | 1110140.42 |
| birds | Turnix_melanogaster | Turnix_melanogaster | 1 | 87.1 | 3.5 | 109323.75 |
| birds | Turnix_nigricollis | Turnix_nigricollis | 0 | 61.3 | 3.9 | 595286.92 |
| birds | Turnix_olivii | Turnix_olivii | 1 | 56.4 | 3.5 | 2066.24 |
| birds | Turnix_pyrrhothorax | Turnix_pyrrhothorax | 0 | 45.7 | 4 | 4287341.81 |
| birds | Turnix_suscitator | Turnix_suscitator | 0 | 49.1 | 4.2 | 5511938.06 |
| birds | Turnix_sylvaticus | Turnix_sylvaticus | 0 | 52.7 | 3.7 | 17314162.23 |
| birds | Turnix_tanki | Turnix_tanki | 0 | 52.8 | 4 | 9318008.55 |
| birds | Turnix_varius | Turnix_varius | 0 | 90.2 | 3.5 | 2276694.74 |
| birds | Turnix_velox | Turnix_velox | 0 | 44.3 | 3.5 | 6979116.25 |
| birds | Turtur_abyssinicus | Turtur_abyssinicus | 0 | 62.7 | 2 | 4644147.82 |
| birds | Turtur_afer | Turtur_afer | 0 | 65.6 | 2 | 10011361.74 |
| birds | Turtur_brehmeri | Turtur_brehmeri | 0 | 116 | 1.4 | 2760838.43 |
| birds | Turtur_chalcospilos | Turtur_chalcospilos | 0 | 60.6 | 2 | 5888722.68 |

| | | | | | | |
|---|---|---|---|---|---|---|
| birds | Turtur_tympanistria | Turtur_tympanistria | 0 | 70.6 | 2 | 8134665.65 |
| birds | Tychaedon_barbata | Erythropygia_barbata | 0 | 25.6 | 2.4 | 1773141.4 |
| birds | Tychaedon_coryphoeus | Erythropygia_coryphaeus | 0 | 20.4 | 3 | 875738.69 |
| birds | Tychaedon_quadrivirgata | Erythropygia_quadrivirgata | 0 | 26.1 | 2.4 | 1616296.2 |
| birds | Tychaedon_signata | Erythropygia_signata | 0 | 38.2 | 2.4 | 167351 |
| birds | Tylas_eduardi | Tylas_eduardi | 0 | 46.8 | 2 | 231862.91 |
| birds | Tympanuchus_cupido | Tympanuchus_cupido | 0 | 870.1 | 10.2 | 3289884.57 |
| birds | Tympanuchus_pallidicinctus | Tympanuchus_pallidicinctus | 1 | 745.8 | 11 | 453431.18 |
| birds | Tympanuchus_phasianellus | Tympanuchus_phasianellus | 0 | 882.4 | 12 | 6080574.6 |
| birds | Tyrannopsis_sulphurea | Tyrannopsis_sulphurea | 0 | 53.6 | 2 | 6496396.06 |
| birds | Tyrannulus_elatus | Tyrannulus_elatus | 0 | 7 | 1.4 | 6406376.29 |
| birds | Tyrannus_caudifasciatus | Tyrannus_caudifasciatus | 0 | 42.4 | 2.8 | 211089.91 |
| birds | Tyrannus_couchii | Tyrannus_couchii | 0 | 39 | 3.5 | 551401.87 |
| birds | Tyrannus_crassirostris | Tyrannus_crassirostris | 0 | 55.9 | 3.5 | 689580.18 |
| birds | Tyrannus_cubensis | Tyrannus_cubensis | 1 | 93.6 | 2.4 | 110766.9 |
| birds | Tyrannus_dominicensis | Tyrannus_dominicensis | 0 | 46.5 | 2.8 | 1743599.43 |
| birds | Tyrannus_forficatus | Tyrannus_forficatus | 0 | 39.3 | 4.2 | 2047593.81 |
| birds | Tyrannus_melancholicus | Tyrannus_melancholicus | 0 | 37.4 | 3.2 | 16507028.16 |
| birds | Tyrannus_niveigularis | Tyrannus_niveigularis | 0 | 34.4 | 2.8 | 113185.59 |
| birds | Tyrannus_savana | Tyrannus_savana | 0 | 31.9 | 2.4 | 13572472.85 |
| birds | Tyrannus_tyrannus | Tyrannus_tyrannus | 0 | 39.9 | 3.9 | 16650317.21 |
| birds | Tyrannus_verticalis | Tyrannus_verticalis | 0 | 39.6 | 3.5 | 6509951.48 |
| birds | Tyrannus_vociferans | Tyrannus_vociferans | 0 | 45.6 | 3.5 | 2291219.9 |
| birds | Tyto_alba | Tyto_alba | 0 | 403.3 | 5.3 | 63140770.91 |
| birds | Tyto_capensis | Tyto_capensis | 0 | 364.7 | 3.2 | 2692718.84 |
| birds | Tyto_glaucops | Tyto_glaucops | 0 | 397.5 | 4.6 | 75807.64 |
| birds | Tyto_longimembris | Tyto_longimembris | 0 | 423.5 | 4.9 | 4282168.16 |
| birds | Tyto_novaehollandiae | Tyto_novaehollandiae | 0 | 631.9 | 2.8 | 3178834.97 |
| birds | Tyto_tenebricosa | Tyto_tenebricosa | 0 | 660.6 | 1.4 | 1125489.71 |
| birds | Upucerthia_dumetaria | Upucerthia_dumetaria | 0 | 48.7 | 2.8 | 2678611.87 |
| birds | Upucerthia_validirostris | Upucerthia_validirostris | 0 | 33.6 | 2 | 731195.97 |
| birds | Upupa_epops | Upupa_epops | 0 | 66.9 | 5.7 | 61808940.04 |
| birds | Uraeginthus_angolensis | Uraeginthus_angolensis | 0 | 9.9 | 4.2 | 3460240.46 |
| birds | Uraeginthus_bengalus | Uraeginthus_bengalus | 0 | 9.9 | 4.2 | 7659842.25 |

| | | | | | | |
|---|---|---|---|---|---|---|
| birds | Uraeginthus_cyanocephalus | Uraeginthus_cyanocephalus | 0 | 9.8 | 4.9 | 756367.53 |
| birds | Urochroa_bougueri | Urochroa_bougueri | 0 | 11.4 | 2 | 40202.28 |
| birds | Urocolius_indicus | Urocolius_indicus | 0 | 52.7 | 3.5 | 3462430.5 |
| birds | Urocolius_macrourus | Urocolius_macrourus | 0 | 45.5 | 2.4 | 5779732.9 |
| birds | Uropsila_leucogastra | Uropsila_leucogastra | 0 | 9 | 4 | 409367.84 |
| birds | Urosticte_benjamini | Urosticte_benjamini | 0 | 3.9 | 2 | 17377.94 |
| birds | Urosticte_ruficrissa | Urosticte_ruficrissa | 0 | 4.3 | 2 | 31247.82 |
| birds | Vanellus_albiceps | Vanellus_albiceps | 0 | 192 | 2.8 | 7550710.29 |
| birds | Vanellus_armatus | Vanellus_armatus | 0 | 156 | 3 | 5662037.2 |
| birds | Vanellus_chilensis | Vanellus_chilensis | 0 | 327 | 4 | 13321164.29 |
| birds | Vanellus_cinereus | Vanellus_cinereus | 0 | 270 | 4 | 5700503.93 |
| birds | Vanellus_coronatus | Vanellus_coronatus | 0 | 165 | 2.4 | 6790147.37 |
| birds | Vanellus_crassirostris | Vanellus_crassirostris | 0 | 170 | 2.8 | 3461973.99 |
| birds | Vanellus_duvaucelii | Vanellus_duvaucelii | 0 | 143 | 4 | 1666558.67 |
| birds | Vanellus_gregarius | Vanellus_gregarius | 1 | 224.5 | 3.2 | 11070005.97 |
| birds | Vanellus_indicus | Vanellus_indicus | 0 | 181 | 3.5 | 7325570.07 |
| birds | Vanellus_leucurus | Vanellus_leucurus | 0 | 132 | 4 | 6537401.05 |
| birds | Vanellus_lugubris | Vanellus_lugubris | 0 | 122 | 3 | 3561388.31 |
| birds | Vanellus_malabaricus | Vanellus_malarbaricus | 0 | 140 | 4 | 2548296.23 |
| birds | Vanellus_melanocephalus | Vanellus_melanocephalus | 0 | 214 | 4 | 179801.06 |
| birds | Vanellus_melanopterus | Vanellus_melanopterus | 0 | 184.9 | 3 | 959143.34 |
| birds | Vanellus_miles | Vanellus_miles | 0 | 387 | 4 | 3058008.89 |
| birds | Vanellus_senegallus | Vanellus_senegallus | 0 | 248 | 2.8 | 11259086.33 |
| birds | Vanellus_spinosus | Vanellus_spinosus | 0 | 188 | 2.8 | 8392007.79 |
| birds | Vanellus_superciliosus | Vanellus_superciliosus | 0 | 150 | 2.8 | 3364640.18 |
| birds | Vanellus_tectus | Vanellus_tectus | 0 | 100 | 2.4 | 5265920.67 |
| birds | Vanellus_vanellus | Vanellus_vanellus | 0 | 218.4 | 4 | 24038094.92 |
| birds | Vauriella_albigularis | Rhinomyias_albigularis | 1 | 35.7 | 2.4 | 23988.06 |
| birds | Vauriella_goodfellowi | Rhinomyias_goodfellowi | 0 | 31.7 | 2.4 | 7732.35 |
| birds | Vauriella_gularis | Rhinomyias_gularis | 0 | 25.1 | 2.4 | 34379.01 |
| birds | Vauriella_insignis | Rhinomyias_insignis | 1 | 28.7 | 2.4 | 9756.91 |
| birds | Veniliornis_kirkii | Veniliornis_kirkii | 0 | 28.8 | 2.4 | 846131.89 |
| birds | Veniliornis_lignarius | Veniliornis_lignarius | 0 | 34 | 3.9 | 597481.81 |
| birds | Veniliornis_mixtus | Veniliornis_mixtus | 0 | 28.7 | 4 | 3656621.36 |

| | | | | | | |
|---|---|---|---|---|---|---|
| birds | Veniliornis_sanguineus | Veniliornis_sanguineus | 0 | 26.5 | 1.4 | 36223.6 |
| birds | Vermivora_chrysoptera | Vermivora_chrysoptera | 0 | 8.7 | 4.5 | 4556072.36 |
| birds | Vermivora_cyanoptera | Vermivora_pinus | 0 | 8.9 | 5.5 | 3546336.02 |
| birds | Verreauxia_africana | Sasia_africana | 0 | 9.4 | 2 | 2527784.99 |
| birds | Vireo_altiloquus | Vireo_altiloquus | 0 | 19 | 2.4 | 4891227.03 |
| birds | Vireo_atricapilla | Vireo_atricapilla | 0 | 9 | 3.9 | 1513757.31 |
| birds | Vireo_bellii | Vireo_bellii | 0 | 8.5 | 3.9 | 3853184.17 |
| birds | Vireo_carmioli | Vireo_carmioli | 0 | 13.1 | 2 | 4187.03 |
| birds | Vireo_cassinii | Vireo_cassinii | 0 | 14.7 | 3.9 | 4111688.48 |
| birds | Vireo_crassirostris | Vireo_crassirostris | 0 | 13 | 2.4 | 19800.8 |
| birds | Vireo_flavifrons | Vireo_flavifrons | 0 | 18 | 3.9 | 5029981.24 |
| birds | Vireo_flavoviridis | Vireo_flavoviridis | 0 | 17.6 | 2.4 | 3010055.81 |
| birds | Vireo_gilvus | Vireo_gilvus | 0 | 12.7 | 3.9 | 11735657.6 |
| birds | Vireo_griseus | Vireo_griseus | 0 | 11.4 | 3.9 | 3615420.04 |
| birds | Vireo_gundlachii | Vireo_gundlachii | 0 | 12.9 | 3 | 109844.77 |
| birds | Vireo_huttoni | Vireo_huttoni | 0 | 11.3 | 3.9 | 1214843.43 |
| birds | Vireo_latimeri | Vireo_latimeri | 0 | 12 | 3 | 6995.48 |
| birds | Vireo_magister | Vireo_magister | 0 | 15.4 | 2 | 8592.85 |
| birds | Vireo_modestus | Vireo_modestus | 0 | 9.6 | 2.4 | 11021.79 |
| birds | Vireo_nanus | Vireo_nanus | 0 | 10.5 | 2 | 13196.28 |
| birds | Vireo_olivaceus | Vireo_olivaceus | 0 | 16.1 | 3.9 | 21662377.65 |
| birds | Vireo_pallens | Vireo_pallens | 0 | 11 | 2.4 | 264054.35 |
| birds | Vireo_philadelphicus | Vireo_philadelphicus | 0 | 11.5 | 3.9 | 6279191.56 |
| birds | Vireo_plumbeus | Vireo_plumbeus | 0 | 16.4 | 3.9 | 2446894.17 |
| birds | Vireo_solitarius | Vireo_solitarius | 0 | 15.3 | 3.9 | 8397663.45 |
| birds | Vireo_vicinior | Vireo_vicinior | 0 | 12.8 | 3.9 | 966622.87 |
| birds | Xanthocephalus_xanthocephalus | Xanthocephalus_xanthocephalus | 0 | 62.7 | 3.5 | 8088084.37 |
| birds | Xanthomixis_cinereiceps | Bernieria_cinereiceps | 0 | 18.2 | 3 | 163803.27 |
| birds | Xanthomixis_zosterops | Bernieria_zosterops | 0 | 10.9 | 3.5 | 210073.52 |
| birds | Xanthopsar_flavus | Xanthopsar_flavus | 1 | 43 | 4 | 732681.65 |
| birds | Xanthotis_flaviventer | Xanthotis_flaviventer | 0 | 40 | 2 | 759089.47 |
| birds | Xanthotis_macleayanus | Xanthotis_macleayanus | 0 | 31 | 2 | 27328.2 |
| birds | Xenops_minutus | Xenops_minutus | 0 | 10.6 | 2 | 945529.29 |
| birds | Xenops_rutilus | Xenops_rutilans | 0 | 11.2 | 2.4 | 6929954.15 |

| | | | | | | |
|---|---|---|---|---|---|---|
| birds | Xenopsaris_albinucha | Xenopsaris_albinucha | 0 | 10 | 3 | 4453736.32 |
| birds | Xenornis_setifrons | Xenornis_setifrons | 0 | 24.6 | 2 | 17510.59 |
| birds | Xenotriccus_callizonus | Xenotriccus_callizonus | 0 | 11.2 | 3 | 53967.15 |
| birds | Xenotriccus_mexicanus | Xenotriccus_mexicanus | 0 | 13.8 | 2.4 | 112766.33 |
| birds | Xenus_cinereus | Xenus_cinereus | 0 | 78.8 | 4 | 14264708.24 |
| birds | Xiphidiopicus_percussus | Xiphidiopicus_percussus | 0 | 60.9 | 3.5 | 110277.34 |
| birds | Xiphocolaptes_albicollis | Xiphocolaptes_albicollis | 0 | 118 | 2 | 2097683.15 |
| birds | Xiphocolaptes_major | Xiphocolaptes_major | 0 | 156 | 2.4 | 1461863.85 |
| birds | Xipholena_atropurpurea | Xipholena_atropurpurea | 1 | 61.3 | 1 | 214393.1 |
| birds | Xipholena_lamellipennis | Xipholena_lamellipennis | 0 | 53.8 | 1 | 1084022.35 |
| birds | Xipholena_punicea | Xipholena_punicea | 0 | 68.1 | 1 | 3624635.86 |
| birds | Xiphorhynchus_erythropygius | Xiphorhynchus_erythropygius | 0 | 46.8 | 2 | 175302.58 |
| birds | Xiphorhynchus_flavigaster | Xiphorhynchus_flavigaster | 0 | 46.2 | 2.4 | 767413.14 |
| birds | Xiphorhynchus_fuscus | Xiphorhynchus_fuscus | 0 | 21.8 | 2.4 | 1948346.79 |
| birds | Xiphorhynchus_guttatus | Xiphorhynchus_guttatus | 0 | 59.7 | 1.7 | 2005419.73 |
| birds | Xiphorhynchus_lachrymosus | Xiphorhynchus_lachrymosus | 0 | 56.4 | 2 | 271654.21 |
| birds | Xiphorhynchus_obsoletus | Xiphorhynchus_obsoletus | 0 | 32.6 | 2 | 6665026.21 |
| birds | Xiphorhynchus_pardalotus | Xiphorhynchus_pardalotus | 0 | 34.6 | 1 | 1710329.56 |
| birds | Xiphorhynchus_spixii | Xiphorhynchus_spixii | 0 | 31.2 | 2 | 993444.47 |
| birds | Xiphorhynchus_susurrans | Xiphorhynchus_susurrans | 0 | 46.7 | 2 | 640926.31 |
| birds | Xolmis_cinereus | Xolmis_cinereus | 0 | 57.1 | 2.4 | 6894893.41 |
| birds | Xolmis_coronatus | Xolmis_coronatus | 0 | 46.8 | 2.4 | 2279881.38 |
| birds | Xolmis_dominicanus | Xolmis_dominicanus | 1 | 42.8 | 3.5 | 344144.45 |
| birds | Xolmis_irupero | Xolmis_irupero | 0 | 28.7 | 3.5 | 3570599.32 |
| birds | Xolmis_pyrope | Xolmis_pyrope | 0 | 35.3 | 2.4 | 747754.31 |
| birds | Xolmis_rubetra | Xolmis_rubetra | 0 | 36.3 | 2 | 1640531.88 |
| birds | Zanda_baudinii | Calyptorhynchus_baudinii | 1 | 620 | 2 | 56186.02 |
| birds | Zanda_funerea | Calyptorhynchus_funereus | 0 | 674.7 | 2 | 1045925.98 |
| birds | Zanda_latirostris | Calyptorhynchus_latirostris | 1 | 612 | 1.4 | 235747.48 |
| birds | Zapornia_akool | Amaurornis_akool | 0 | 131.4 | 5.5 | 3895470.77 |
| birds | Zapornia_flavirostra | Amaurornis_flavirostra | 0 | 86.2 | 3 | 16710583.49 |
| birds | Zapornia_fusca | Porzana_fusca | 0 | 57.3 | 5.2 | 6843713.64 |
| birds | Zapornia_parva | Porzana_parva | 0 | 49.7 | 7.9 | 12207181.68 |
| birds | Zapornia_paykullii | Porzana_paykullii | 0 | 111 | 6.7 | 2630845.25 |

| | | | | | | |
|---|---|---|---|---|---|---|
| birds | Zapornia_pusilla | Porzana_pusilla | 0 | 35.4 | 6.6 | 41547271.71 |
| birds | Zapornia_tabuensis | Porzana_tabuensis | 0 | 42.2 | 3.5 | 2288129.04 |
| birds | Zaratornis_stresemanni | Zaratornis_stresemanni | 0 | 52.1 | 3 | 61015.41 |
| birds | Zenaida_asiatica | Zenaida_asiatica | 0 | 153 | 2 | 3221953.54 |
| birds | Zenaida_auriculata | Zenaida_auriculata | 0 | 110.2 | 2 | 11530888.21 |
| birds | Zenaida_aurita | Zenaida_aurita | 0 | 156 | 2 | 230814.88 |
| birds | Zenaida_galapagoensis | Zenaida_galapagoensis | 0 | 111 | 2 | 8331.05 |
| birds | Zenaida_macroura | Zenaida_macroura | 0 | 118.9 | 2 | 12173012.56 |
| birds | Zentrygon_albifacies | Geotrygon_albifacies | 0 | 238 | 1.4 | 192106.95 |
| birds | Zentrygon_carrikeri | Geotrygon_carrikeri | 1 | 267 | 1 | 127.54 |
| birds | Zentrygon_chiriquensis | Geotrygon_chiriquensis | 0 | 308 | 2 | 16384.72 |
| birds | Zentrygon_frenata | Geotrygon_frenata | 0 | 307 | 1 | 442995.43 |
| birds | Zentrygon_lawrencii | Geotrygon_lawrencii | 0 | 220 | 1 | 7970.03 |
| birds | Zentrygon_linearis | Geotrygon_linearis | 0 | 261 | 2 | 162092.83 |
| birds | Zimmerius_vilissimus | Zimmerius_vilissimus | 0 | 9.7 | 2 | 79107.68 |
| birds | Zonotrichia_albicollis | Zonotrichia_albicollis | 0 | 24.4 | 3.9 | 11733130.06 |
| birds | Zonotrichia_atricapilla | Zonotrichia_atricapilla | 0 | 32 | 3.9 | 2344274.02 |
| birds | Zonotrichia_capensis | Zonotrichia_capensis | 0 | 20.3 | 2.8 | 12703362.39 |
| birds | Zonotrichia_leucophrys | Zonotrichia_leucophrys | 0 | 28 | 3.9 | 17967442.74 |
| birds | Zonotrichia_querula | Zonotrichia_querula | 0 | 35.5 | 3.9 | 6150624.35 |
| birds | Zoothera_andromedae | Zoothera_andromedae | 0 | 90.5 | 2.4 | 132967.79 |
| birds | Zoothera_dauma | Zoothera_dauma | 0 | 134.1 | 3.5 | 3762059.45 |
| birds | Zoothera_dixoni | Zoothera_dixoni | 0 | 90 | 3 | 1131286.29 |
| birds | Zoothera_heinei | Zoothera_heinei | 0 | 82.8 | 2.4 | 469278.97 |
| birds | Zoothera_lunulata | Zoothera_lunulata | 0 | 130 | 2.4 | 578456.61 |
| birds | Zoothera_marginata | Zoothera_marginata | 0 | 81 | 3.5 | 939864.56 |
| birds | Zoothera_mollissima | Zoothera_mollissima | 0 | 98.2 | 4 | 422988.16 |
| birds | Zoothera_monticola | Zoothera_monticola | 0 | 126 | 2.8 | 323451.38 |
| birds | Zosterops_abyssinicus | Zosterops_abyssinicus | 0 | 9.4 | 2.8 | 665331.72 |
| birds | Zosterops_citrinella | Zosterops_citrinella | 0 | 11.5 | 2.8 | 70904.8 |
| birds | Zosterops_griseotinctus | Zosterops_griseotinctus | 0 | 14 | 2 | 1046.45 |
| birds | Zosterops_hypoxanthus | Zosterops_hypoxanthus | 0 | 12.5 | 2.4 | 45547.01 |
| birds | Zosterops_japonicus | Zosterops_japonicus | 0 | 11.3 | 2.8 | 621194.12 |
| birds | Zosterops_lateralis | Zosterops_lateralis | 0 | 12.7 | 2.8 | 2702683.6 |

| | | | | | | |
|---|---|---|---|---|---|---|
| birds | Zosterops_luteus | Zosterops_luteus | 0 | 9.7 | 3.5 | 672778.06 |
| birds | Zosterops_maderaspatanus | Zosterops_maderaspatanus | 0 | 8.9 | 2.4 | 592006.09 |
| birds | Zosterops_novaeguineae | Zosterops_novaeguineae | 0 | 12.5 | 2 | 103133.53 |
| birds | Zosterops_pallidus | Zosterops_pallidus | 0 | 11.1 | 2.8 | 420000.76 |
| birds | Zosterops_palpebrosus | Zosterops_palpebrosus | 0 | 8.6 | 2.8 | 6382482.99 |
| birds | Zosterops_poliogastrus | Zosterops_poliogastrus | 0 | 11 | 2.4 | 324865.26 |
| birds | Zosterops_senegalensis | Zosterops_senegalensis | 0 | 10.9 | 2 | 8737492.27 |
