## Supplementary Methods for "Occam’s bias undermines inferences from phylogenetic linear models"

To predict the impact of Occam's bias using model structure Trait 1 ~ Trait 2 + Trait 3, we defined the bivariate correlation between Trait 1 and Trait 2 as zero, and then determined the conditions that produce falsely significant Trait 2 coefficients using an inequality statement (Equation 4) based on the partial correlation formula, the standard error of the partial correlation formula, and the significance threshold (Equations 1-3).

***Equation 1. Partial correlation***

$$r_{12.3} = (r_{12} - r_{13} \cdot r_{23}) / \sqrt{[(1 - r_{13}^2)(1 - r_{23}^2)]}$$

where $r_{12.3}$ is the correlation coefficient of Trait 1 and Trait 2 given Trait 3

***Equation 2. Standard error of a partial correlation***

$$SE(r_{12.3}) = \sqrt{[(1 - r_{12.3}^2) / (n - k - 1)]}$$

where n is sample size and k is number of predictors. For null hypothesis H0: $r_{12.3} = 0$, we substitute 0 into the formula, giving

$$SE(r_{12.3}) = 1/\sqrt{(n - k - 1)}$$

***Equation 3. Significance threshold***

$$| t\_statistic | = r_{12.3} / SE(r_{12.3})$$

which was obtained from t-statistic formula of linear regression. To reject the null hypothesis H0: $r_{12.3} = 0$,

$$| t\_statistic | > t\_critical$$

where t_critical is obtained using the inverse cumulative density function, alpha (0.05) and degrees of freedom (df = n - k -1). Therefore,

$$| r_{12.3} | / SE(r_{12.3}) > t\_critical$$

which rearranges to

$$| r_{12.3} | > t\_critical \cdot SE(r_{12.3})$$

Substituting the SE formula gives,

$$| r_{12.3} | > t\_critical \cdot 1/\sqrt{(n - k - 1)}$$

The right-hand side is the significance threshold which any $r_{12.3}$ must be greater than to be significant. Because t_critical is dimensionless, relevant units are inherited from $SE(r_{12.3})$, which are in this case correlations.

***Equation 4: Occam's bias inequality***

We used the partial correlation formula (Equation 1) with significance threshold (Equation 3) to obtain Occam's bias inequality, which predicts false significance in $r_{12.3}$. The coefficient $r_{12.3}$ would be falsely significant if

$$|r_{12.3}| > \text{significance threshold}$$

Substituting the significance threshold with its formula gives

$$|r_{12.3}| > t\_critical \cdot 1/\sqrt{(n - k - 1)}$$

Substituting $r_{12.3}$ with the partial correlation formula gives

$$|r_{12} - r_{13} \cdot r_{23} / \sqrt{[(1 - r_{13}^2)(1 - r_{23}^2)]}| > t\_critical \cdot 1/\sqrt{(n - k - 1)}$$

In these simulations, $r_{12}$ is always 0, therefore

$$|r_{13} \cdot r_{23} / \sqrt{[(1 - r_{13}^2)(1 - r_{23}^2)]}| > t\_critical \cdot 1/\sqrt{(n - k - 1)}$$

Which rearranges to the Occam's bias inequality

$$|r_{13} \cdot r_{23}| > t\_critical \cdot 1/\sqrt{(n - k - 1)} \cdot \sqrt{[(1 - r_{13}^2)(1 - r_{23}^2)]}$$

As we defined the bivariate correlations, we can solve this inequality. If the inequality is true, then Occam's bias will cause a false significance in Trait 2's effect on Trait 1.
