## Supplementary material for "Occam’s bias undermines inferences from phylogenetic linear models": Table S1

**Table S1**. Outputs from Bayesian phylogenetic multivariate response (MR) models, performed using the R package *MCMCglmm* to test the prediction that life history traits (body mass, fecundity and geographic range size; all log-transformed then *z*-transformed) are associated with extinction risk (binary, the probability of being threatened) among global tetrapods (amphibians, lizards, mammals and birds). Fixed effects were restricted to fixed intercepts for each response variable. For random effects, we utilised an unstructured, multidimensional variance-covariance matrix, which allows for the estimation of the phylogenetic and residual variances of each response variable as well as the phylogenetic and residual covariances of each pairwise combination of response variables. Consequently, evidence for our hypothesis was derived from observing significant phylogenetic or residual covariances, with the former indicating a significant relationship with clustering by clades while the latter indicates a significant relationship without clustering by clades. Extinction risk was modelled using the threshold likelihood function, therefore we fixed its residual variances 1 for identifiability. Continuous response variables were modelled using a Gaussian likelihood function. One MR model ran for each taxon. Table shows the model name with sample size (*n*), parameter type, parameter name, posterior mean, 95% credible interval (95% CI), and effective sample size (*ESS*). Phylogenetic and residual covariances are considered significant when the 95% credible interval excludes zero. Significance is indicated in the table by the bold-face text. Convergence of the MCMC chain was determined by examination of trace plots and we ensured *ESS* > 1,000.

| Model | Parameter Type | Parameter | Posterior Mean | 95% CI | *ESS* |
|---|---|---|---|---|---|
| **Amphibians (n = 2,105)** | fixed intercept | threatened_1_not_0 | -0.986 | -2.406, 0.393 | 49299 |
| | fixed intercept | body_mass | 0.558 | -0.458, 1.615 | 50000 |
| | fixed intercept | fecundity | -0.383 | -1.27, 0.495 | 50000 |
| | fixed intercept | geographic_range_size | -0.042 | -0.967, 0.845 | 50000 |
| | phylogenetic variance | threatened_1_not_0:threatened_1_not_0 | 2.953 | 1.766, 4.231 | 5917 |
| | phylogenetic variance | body_mass:body_mass | 1.686 | 1.48, 1.899 | 35693 |
| | phylogenetic variance | fecundity:fecundity | 1.213 | 1.058, 1.373 | 36597 |
| | phylogenetic variance | geographic_range_size:geographic_range_size | 1.276 | 1.011, 1.546 | 17476 |
| | phylogenetic covariance | threatened_1_not_0:body_mass | **-0.338** | **-0.615, -0.063** | 12870 |
| | phylogenetic covariance | threatened_1_not_0:fecundity | **-0.675** | **-0.937, -0.41** | 11278 |
| | phylogenetic covariance | threatened_1_not_0:geographic_range_size | **-1.374** | **-1.857, -0.924** | 6703 |
| | phylogenetic covariance | body_mass:fecundity | **0.902** | **0.759, 1.047** | 27894 |
| | phylogenetic covariance | body_mass:geographic_range_size | **0.502** | **0.344, 0.667** | 18916 |
| | phylogenetic covariance | fecundity:geographic_range_size | **0.661** | **0.516, 0.811** | 16909 |
| | residual variance | threatened_1_not_0:threatened_1_not_0 | 1 | 1, 1 | 0 |
| | residual variance | body_mass:body_mass | 0.255 | 0.232, 0.278 | 45551 |
| | residual variance | fecundity:fecundity | 0.241 | 0.221, 0.261 | 45257 |
| | residual variance | geographic_range_size:geographic_range_size | 0.552 | 0.507, 0.599 | 21322 |

| | | | | | |
|---|---|---|---|---|---|
| | residual covariance | threatened_1_not_0:body_mass | **-0.048** | **-0.083, -0.01** | 35568 |
| | residual covariance | threatened_1_not_0:fecundity | **-0.054** | **-0.088, -0.02** | 32634 |
| | residual covariance | threatened_1_not_0:geographic_range_size | **-0.444** | **-0.488, -0.399** | 18383 |
| | residual covariance | body_mass:fecundity | **0.076** | **0.06, 0.092** | 43149 |
| | residual covariance | body_mass:geographic_range_size | **0.054** | **0.031, 0.077** | 33807 |
| | residual covariance | fecundity:geographic_range_size | **0.063** | **0.041, 0.085** | 32229 |
| **Lizards (n = 3,272)** | fixed intercept | threatened_1_not_0 | -1.531 | -3.212, 0.149 | 40771 |
| | fixed intercept | body_mass | 0.217 | -0.511, 0.944 | 50000 |
| | fixed intercept | fecundity | -0.113 | -0.729, 0.511 | 50000 |
| | fixed intercept | geographic_range_size | -0.008 | -0.863, 0.801 | 50000 |
| | phylogenetic variance | threatened_1_not_0:threatened_1_not_0 | 5.523 | 3.176, 8.304 | 1738 |
| | phylogenetic variance | body_mass:body_mass | 1.033 | 0.94, 1.126 | 28179 |
| | phylogenetic variance | fecundity:fecundity | 0.731 | 0.653, 0.812 | 26216 |
| | phylogenetic variance | geographic_range_size:geographic_range_size | 1.359 | 1.14, 1.579 | 15688 |
| | phylogenetic covariance | threatened_1_not_0:body_mass | **-0.281** | **-0.503, -0.063** | 7938 |
| | phylogenetic covariance | threatened_1_not_0:fecundity | **-0.271** | **-0.472, -0.073** | 7757 |
| | phylogenetic covariance | threatened_1_not_0:geographic_range_size | **-1.631** | **-2.192, -1.102** | 2396 |
| | phylogenetic covariance | body_mass:fecundity | **0.439** | **0.374, 0.503** | 26695 |
| | phylogenetic covariance | body_mass:geographic_range_size | **0.208** | **0.116, 0.304** | 20479 |
| | phylogenetic covariance | fecundity:geographic_range_size | **0.268** | **0.18, 0.355** | 17918 |
| | residual variance | threatened_1_not_0:threatened_1_not_0 | 1 | 1, 1 | 0 |
| | residual variance | body_mass:body_mass | 0.175 | 0.161, 0.189 | 44175 |
| | residual variance | fecundity:fecundity | 0.196 | 0.182, 0.211 | 45078 |
| | residual variance | geographic_range_size:geographic_range_size | 0.497 | 0.457, 0.539 | 16624 |
| | residual covariance | threatened_1_not_0:body_mass | -0.025 | -0.054, 0.004 | 32209 |
| | residual covariance | threatened_1_not_0:fecundity | -0.025 | -0.055, 0.005 | 31663 |
| | residual covariance | threatened_1_not_0:geographic_range_size | **-0.28** | **-0.332, -0.228** | 9900 |
| | residual covariance | body_mass:fecundity | **0.041** | **0.031, 0.052** | 41198 |
| | residual covariance | body_mass:geographic_range_size | **0.031** | **0.015, 0.048** | 33596 |
| | residual covariance | fecundity:geographic_range_size | **0.025** | **0.009, 0.042** | 31947 |
| **Mammals (n = 3,462)** | fixed intercept | threatened_1_not_0 | -0.71 | -2.406, 0.906 | 50000 |
| | fixed intercept | body_mass | 0.529 | -0.337, 1.403 | 52061 |
| | fixed intercept | fecundity | -0.119 | -1.231, 0.987 | 50000 |
| | fixed intercept | geographic_range_size | 0.033 | -1.06, 1.104 | 48444 |
| | phylogenetic variance | threatened_1_not_0:threatened_1_not_0 | 2.705 | 1.697, 3.822 | 3926 |
| | phylogenetic variance | body_mass:body_mass | 0.744 | 0.68, 0.809 | 9378 |
| | phylogenetic variance | fecundity:fecundity | 1.247 | 1.119, 1.381 | 12253 |

| | | | | | |
|---|---|---|---|---|---|
| | phylogenetic variance | geographic_range_size:geographic_range_size | 1.122 | 0.851, 1.416 | 14761 |
| | phylogenetic covariance | threatened_1_not_0:body_mass | **0.387** | **0.251, 0.528** | 14447 |
| | phylogenetic covariance | threatened_1_not_0:fecundity | **-0.316** | **-0.519, -0.112** | 10449 |
| | phylogenetic covariance | threatened_1_not_0:geographic_range_size | **-0.697** | **-1.039, -0.377** | 4659 |
| | phylogenetic covariance | body_mass:fecundity | **-0.188** | **-0.248, -0.126** | 29828 |
| | phylogenetic covariance | body_mass:geographic_range_size | 0.028 | -0.051, 0.107 | 17975 |
| | phylogenetic covariance | fecundity:geographic_range_size | **0.263** | **0.138, 0.388** | 13710 |
| | residual variance | threatened_1_not_0:threatened_1_not_0 | 1 | 1, 1 | 0 |
| | residual variance | body_mass:body_mass | 0.075 | 0.07, 0.079 | 50000 |
| | residual variance | fecundity:fecundity | 0.129 | 0.121, 0.137 | 43302 |
| | residual variance | geographic_range_size:geographic_range_size | 0.692 | 0.654, 0.733 | 29690 |
| | residual covariance | threatened_1_not_0:body_mass | 0 | -0.012, 0.013 | 44331 |
| | residual covariance | threatened_1_not_0:fecundity | **-0.025** | **-0.044, -0.006** | 37365 |
| | residual covariance | threatened_1_not_0:geographic_range_size | **-0.509** | **-0.545, -0.473** | 26284 |
| | residual covariance | body_mass:fecundity | 0.002 | -0.002, 0.007 | 48010 |
| | residual covariance | body_mass:geographic_range_size | 0.008 | -0.002, 0.017 | 43556 |
| | residual covariance | fecundity:geographic_range_size | **0.032** | **0.019, 0.045** | 35422 |
| **Birds (n = 5,107)** | fixed intercept | threatened_1_not_0 | -1.377 | -1.978, -0.79 | 37747 |
| | fixed intercept | body_mass | 1.496 | 1.067, 1.935 | 49437 |
| | fixed intercept | fecundity | 0.686 | -0.036, 1.395 | 49127 |
| | fixed intercept | geographic_range_size | 0.172 | -0.357, 0.685 | 48930 |
| | phylogenetic variance | threatened_1_not_0:threatened_1_not_0 | 0.591 | 0.316, 0.898 | 4039 |
| | phylogenetic variance | body_mass:body_mass | 0.379 | 0.356, 0.403 | 6548 |
| | phylogenetic variance | fecundity:fecundity | 1.071 | 0.976, 1.165 | 23205 |
| | phylogenetic variance | geographic_range_size:geographic_range_size | 0.524 | 0.411, 0.638 | 19900 |
| | phylogenetic covariance | threatened_1_not_0:body_mass | **0.147** | **0.101, 0.195** | 9500 |
| | phylogenetic covariance | threatened_1_not_0:fecundity | -0.088 | -0.186, 0.01 | 5123 |
| | phylogenetic covariance | threatened_1_not_0:geographic_range_size | -0.08 | -0.171, 0.007 | 3869 |
| | phylogenetic covariance | body_mass:fecundity | **-0.081** | **-0.111, -0.049** | 33854 |
| | phylogenetic covariance | body_mass:geographic_range_size | -0.001 | -0.032, 0.029 | 18740 |
| | phylogenetic covariance | fecundity:geographic_range_size | **0.246** | **0.174, 0.318** | 12121 |
| | residual variance | threatened_1_not_0:threatened_1_not_0 | 1 | 1, 1 | 0 |
| | residual variance | body_mass:body_mass | 0.063 | 0.06, 0.067 | 50000 |
| | residual variance | fecundity:fecundity | 0.215 | 0.201, 0.229 | 37090 |
| | residual variance | geographic_range_size:geographic_range_size | 0.718 | 0.683, 0.753 | 35116 |
| | residual covariance | threatened_1_not_0:body_mass | 0.006 | -0.005, 0.017 | 44664 |
| | residual covariance | threatened_1_not_0:fecundity | **-0.046** | **-0.074, -0.02** | 26521 |

| | | | | | |
|---|---|---|---|---|---|
| | residual covariance | threatened_1_not_0:geographic_range_size | **-0.373** | **-0.413, -0.333** | 22145 |
| | residual covariance | body_mass:fecundity | -0.001 | -0.005, 0.004 | 47269 |
| | residual covariance | body_mass:geographic_range_size | -0.005 | -0.013, 0.003 | 41566 |
| | residual covariance | fecundity:geographic_range_size | **0.104** | **0.087, 0.12** | 28491 |

11
