## Supplementary material for "Occam’s bias undermines inferences from phylogenetic linear models": Table S2

**Table S2**. Outputs from Bayesian phylogenetic single-response (SR) models, performed using the R package *MCMCglmm* to test the prediction that life history traits (body mass, fecundity and geographic range size; each log-transformed then *z*-transformed) are associated with extinction risk (binary, the probability of being threatened as a binary variable; modelled using a threshold liability likelihood function) among global tetrapods (amphibians, lizards, mammals and birds). For each tetrapod group, we ran four models. The response variable (extinction risk) is the same across all models, while predictors differ. The first model has body mass as the predictor. The second model has body mass and fecundity as predictors. The third model has body mass and geographic range size as predictors. The fourth model has body mass, fecundity and geographic range size as predictors. We estimated the phylogenetic variance as a random effect, and fixed the residual variance to 1 for identifiability. Table shows the taxon with sample size (*n*), model number, deviance information criterion (*DIC*), parameter, posterior mean, 95% credible interval (95% CI), and effective sample size (*ESS*). Predictors are considered significant when the 95% credible interval excludes include zero. Significance is indicated in the table by the bold-face text. Convergence of the MCMC chain was determined by examination of trace plots and we ensured *ESS* > 1,000.

| **Taxon** | **Model** | ***DIC*** | **Parameter** | **Posterior Mean** | **95% CI** | ***ESS*** |
|---|---|---|---|---|---|---|
| **Amphibians (*n* = 2,105)** | 1 | 1993.799 | fixed_intercept | -1.103 | -2.956, 0.698 | 24066 |
| | | | phylogenetic_variance | 5.251 | 2.856, 7.912 | 2227 |
| | | | residual_variance | 1 | 1, 1 | 0 |
| | | | body_mass | **-0.281** | **-0.406, -0.148** | 11458 |
| | 2 | 1993.198 | fixed_intercept | -1.34 | -2.922, 0.231 | 23452 |
| | | | phylogenetic_variance | 3.779 | 2.009, 5.75 | 2387 |
| | | | residual_variance | 1 | 1, 1 | 0 |
| | | | body_mass | 0.001 | -0.139, 0.136 | 19662 |
| | | | fecundity | **-0.527** | **-0.675, -0.373** | 20448 |
| | 3 | 1157.683 | fixed_intercept | -1.739 | -3.196, -0.271 | 16694 |
| | | | phylogenetic_variance | 3.02 | 1.182, 5.144 | 1005 |
| | | | residual_variance | 1 | 1, 1 | 0 |
| | | | body_mass | **0.251** | **0.11, 0.397** | 17762 |
| | | | geographic_range_size | **-1.938** | **-2.222, -1.664** | 1111 |
| | 4 | 1158.281 | fixed_intercept | -1.731 | -3.21, -0.283 | 18281 |
| | | | phylogenetic_variance | 3.063 | 1.288, 5.266 | 1014 |
| | | | residual_variance | 1 | 1, 1 | 0 |
| | | | body_mass | **0.246** | **0.078, 0.416** | 18614 |
| | | | fecundity | 0.012 | -0.185, 0.199 | 19069 |

| | | | | | | |
|---|---|---|---|---|---|---|
| | | | geographic_range_size | **-1.945** | **-2.236, -1.67** | 1108 |
| **Lizards (*n* = 3,272)** | 1 | 2333.175 | fixed_intercept | -1.735 | -3.661, 0.203 | 50659 |
| | | | phylogenetic_variance | 7.365 | 3.839, 11.48 | 1761 |
| | | | residual_variance | 1 | 1, 1 | 0 |
| | | | body_mass | **-0.264** | **-0.411, -0.12** | 17545 |
| | 2 | 2314.493 | fixed_intercept | -1.796 | -3.778, 0.162 | 39398 |
| | | | phylogenetic_variance | 7.716 | 3.848, 12.167 | 1494 |
| | | | residual_variance | 1 | 1, 1 | 0 |
| | | | body_mass | **-0.163** | **-0.325, -0.002** | 24386 |
| | | | fecundity | **-0.27** | **-0.443, -0.094** | 18373 |
| | 3 | 1961.242 | fixed_intercept | -1.744 | -3.33, -0.239 | 41667 |
| | | | phylogenetic_variance | 4.412 | 2.165, 7.138 | 1981 |
| | | | residual_variance | 1 | 1, 1 | 0 |
| | | | body_mass | -0.065 | -0.204, 0.072 | 39494 |
| | | | geographic_range_size | **-1.068** | **-1.246, -0.911** | 2581 |
| | 4 | 1958.084 | fixed_intercept | -1.756 | -3.31, -0.186 | 46088 |
| | | | phylogenetic_variance | 4.53 | 2.165, 7.174 | 2100 |
| | | | residual_variance | 1 | 1, 1 | 0 |
| | | | body_mass | -0.066 | -0.216, 0.082 | 53116 |
| | | | fecundity | 0.002 | -0.164, 0.163 | 56999 |
| | | | geographic_range_size | **-1.075** | **-1.248, -0.915** | 2896 |
| **Mammals (*n* = 3,462)** | 1 | 2867.234 | fixed_intercept | -1.075 | -3.079, 0.814 | 25000 |
| | | | phylogenetic_variance | 3.948 | 2.198, 5.853 | 2217 |
| | | | residual_variance | 1 | 1, 1 | 0 |
| | | | body_mass | **0.485** | **0.322, 0.648** | 22586 |
| | 2 | 2854.917 | fixed_intercept | -1.084 | -2.984, 0.906 | 25000 |
| | | | phylogenetic_variance | 3.876 | 2.154, 5.952 | 2144 |
| | | | residual_variance | 1 | 1, 1 | 0 |
| | | | body_mass | **0.431** | **0.266, 0.595** | 25000 |
| | | | fecundity | **-0.276** | **-0.406, -0.143** | 19350 |
| | 3 | 1935.433 | fixed_intercept | -1.373 | -3.31, 0.554 | 25000 |
| | | | phylogenetic_variance | 3.793 | 1.87, 5.961 | 1285 |
| | | | residual_variance | 1 | 1, 1 | 0 |
| | | | body_mass | **0.829** | **0.646, 1.029** | 10513 |
| | | | geographic_range_size | **-1.241** | **-1.369, -1.119** | 2268 |
| | 4 | 1935.935 | fixed_intercept | -1.377 | -3.328, 0.549 | 24569 |
| | | | phylogenetic_variance | 3.83 | 1.856, 5.817 | 1397 |

| | | | | | | |
|---|---|---|---|---|---|---|
| | | | residual_variance | 1 | 1, 1 | 0 |
| | | | body_mass | **0.832** | **0.641, 1.03** | 10594 |
| | | | fecundity | 0.002 | -0.153, 0.155 | 21222 |
| | | | geographic_range_size | **-1.244** | **-1.371, -1.119** | 2319 |
| **Birds (*n* = 5,107)** | 1 | 2340.179 | fixed_intercept | -1.985 | -2.573, -1.429 | 12967 |
| | | | phylogenetic_variance | 0.48 | 0.186, 0.834 | 1945 |
| | | | residual_variance | 1 | 1, 1 | 0 |
| | | | body_mass | **0.411** | **0.301, 0.516** | 19460 |
| | 2 | 2335.286 | fixed_intercept | -1.878 | -2.437, -1.35 | 14380 |
| | | | phylogenetic_variance | 0.43 | 0.174, 0.75 | 2347 |
| | | | residual_variance | 1 | 1, 1 | 0 |
| | | | body_mass | **0.386** | **0.28, 0.492** | 20844 |
| | | | fecundity | **-0.123** | **-0.196, -0.045** | 22161 |
| | 3 | 1969.873 | fixed_intercept | -2.184 | -2.799, -1.591 | 12237 |
| | | | phylogenetic_variance | 0.534 | 0.196, 0.927 | 1792 |
| | | | residual_variance | 1 | 1, 1 | 0 |
| | | | body_mass | **0.521** | **0.405, 0.641** | 18340 |
| | | | geographic_range_size | **-0.591** | **-0.666, -0.524** | 4537 |
| | 4 | 1965.025 | fixed_intercept | -2.271 | -2.955, -1.678 | 10336 |
| | | | phylogenetic_variance | 0.581 | 0.201, 0.991 | 1573 |
| | | | residual_variance | 1 | 1, 1 | 0 |
| | | | body_mass | **0.542** | **0.414, 0.661** | 16192 |
| | | | fecundity | 0.079 | -0.006, 0.168 | 12845 |
| | | | geographic_range_size | **-0.609** | **-0.684, -0.535** | 3821 |
