## Supplementary material for "Occam’s bias undermines inferences from phylogenetic linear models": Table S3

**Table S3**. Priors specified for each *MCMCglmm* model. The table gives dataset, model number, variables included, statistical approach, iterations, burn-in percentage, thinning interval, random effects covariance structure, as well as prior specifications for the phylogenetic, residual and fixed effects. Variables abbreviated in the table are simulated variables of Trait 1 (T1), Trait 2 (T2), Trait 3 (T3), as well as the empirical variables of extinction risk (E), body mass (B), fecundity (F) and geographic range size (G). Statistical approaches are either single-response mixed effects models (SR) or multiple-response mixed effects models (MR). For the random effects covariance structure (applies to MR approaches only), we implemented an unstructured multidimensional covariance matrix ('us') that estimates variances of each response variable as well as the covariances of all pairwise combinations of response variables. Random (phylogenetic and residual) effects prior is an inverse-Wishart distribution, with scale *g_v/r_v* and degrees of freedom *g_nu/r_nu* for phylogenetic/residual effects respectively. In the table, *k* equals the number of response variables in the model. For MR models, we use a *k**k* identity matrix as the scale matrix (indicated by *diag(k)*) and *k* + 100 degrees of freedom, which is is moderately regularising in a multiple-response context. For SR models, we used 1-dimensional matrix (i.e., 1 x 1 matrix) as the scale matrix and 1 degree of freedom for simulated analyses and 0.02 degrees of freedom for empirical analyses, which creates a moderately and weakly informative priors (respectively) that has little influence on the posterior distribution. For both MR and SR models on empirical analyses, we implemented parameter expansion algorithms to aid in convergence of phylogenetic effects of binary response variables, specified by *alpha_mu* and *alpha_v*. For identifiability, the residual variance of extinction risk is fixed to 1 in both SR and MR models. Fixed effects priors are normal distributions with standard deviation equal to 3 (variance of 9). Continuous variables were centred on 0 while extinction risk was centred on the base rate obtained from the IUCN (i.e., the proportion of species threatened with extinction obtained from the entire IUCN Red List) transformed to the liability scale.

| Dataset | Model | Variables | Approach | Iterations | Burn-in (%) | Thin | Covariance Structure | g_v | g_nu | alpha_mu | alpha_v | r_v | r_nu | b_mus | b_variances |
|---|---|---|---|---|---|---|---|---|---|---|---|---|---|---|---|
| **Simulations (*n* = 100)** | 1 | T1,T2,T3 | SR | 100,000 | 50 | 10 | NA | 1 | 1 | NA | NA | 1 | 1 | 0 | 3^2 |
| | 2 | T1,T2,T3 | MR | 100,000 | 50 | 10 | us | diag(k) | k | NA | NA | diag(k) | k | 0 | 3^2 |
| **Simulations (*n* = 1,500)** | 3 | T1,T2,T3 | SR | 300,000 | 50 | 10 | NA | 1 | 1 | NA | NA | 1 | 1 | 0 | 3^2 |
| | 4 | T1,T2,T3 | MR | 300,000 | 50 | 10 | us | diag(k) | k | NA | NA | diag(k) | k | 0 | 3^2 |
| **Simulations (*n* = 5,000)** | 5 | T1,T2,T3 | SR | 500,000 | 50 | 10 | NA | 1 | 1 | NA | NA | 1 | 1 | 0 | 3^2 |
| | 6 | T1,T2,T3 | MR | 500,000 | 50 | 10 | us | diag(k) | k | NA | NA | diag(k) | k | 0 | 3^2 |
| **Amphibians** | 1 | E,B | SR | 500,000 | 50 | 10 | NA | 1 | 0.02 | 0 | 1000 | NA | NA | 0/base_rate_liability | 3^2 |
| | 2 | E,B,F | SR | 500,000 | 50 | 10 | NA | 1 | 0.02 | 0 | 1000 | NA | NA | 0/base_rate_liability | 3^2 |
| | 3 | E,B,G | SR | 500,000 | 50 | 10 | NA | 1 | 0.02 | 0 | 1000 | NA | NA | 0/base_rate_liability | 3^2 |
| | 4 | E,B,F,G | SR | 500,000 | 50 | 10 | NA | 1 | 0.02 | 0 | 1000 | NA | NA | 0/base_rate_liability | 3^2 |

| | | | | | | | | | | | | | | | |
|---|---|---|---|---|---|---|---|---|---|---|---|---|---|---|---|
| | 5 | E,B,F,G | MR | 500,000 | 50 | 10 | us | diag(k) | k+100 | 0 | diag(k)*1000 | diag(k) | k+100 | 0/base_rate_liability | 3^2 |
| **Lizards** | 1 | E,B | SR | 1,500,000 | 50 | 10 | NA | 1 | 0.02 | 0 | 1000 | NA | NA | 0/base_rate_liability | 3^2 |
| | 2 | E,B,F | SR | 1,500,000 | 50 | 10 | NA | 1 | 0.02 | 0 | 1000 | NA | NA | 0/base_rate_liability | 3^2 |
| | 3 | E,B,G | SR | 1,500,000 | 50 | 10 | NA | 1 | 0.02 | 0 | 1000 | NA | NA | 0/base_rate_liability | 3^2 |
| | 4 | E,B,F,G | SR | 1,500,000 | 50 | 10 | NA | 1 | 0.02 | 0 | 1000 | NA | NA | 0/base_rate_liability | 3^2 |
| | 5 | E,B,F,G | MR | 500,000 | 50 | 10 | us | diag(k) | k+100 | 0 | diag(k)*1000 | diag(k) | k+100 | 0/base_rate_liability | 3^2 |
| **Mammals** | 1 | E,B | SR | 500,000 | 50 | 10 | NA | 1 | 0.02 | 0 | 1000 | NA | NA | 0/base_rate_liability | 3^2 |
| | 2 | E,B,F | SR | 500,000 | 50 | 10 | NA | 1 | 0.02 | 0 | 1000 | NA | NA | 0/base_rate_liability | 3^2 |
| | 3 | E,B,G | SR | 500,000 | 50 | 10 | NA | 1 | 0.02 | 0 | 1000 | NA | NA | 0/base_rate_liability | 3^2 |
| | 4 | E,B,F,G | SR | 500,000 | 50 | 10 | NA | 1 | 0.02 | 0 | 1000 | NA | NA | 0/base_rate_liability | 3^2 |
| | 5 | E,B,F,G | MR | 500,000 | 50 | 10 | us | diag(k) | k+100 | 0 | diag(k)*1000 | diag(k) | k+100 | 0/base_rate_liability | 3^2 |
| **Birds** | 1 | E,B | SR | 500,000 | 50 | 10 | NA | 1 | 0.02 | 0 | 1000 | NA | NA | 0/base_rate_liability | 3^2 |
| | 2 | E,B,F | SR | 500,000 | 50 | 10 | NA | 1 | 0.02 | 0 | 1000 | NA | NA | 0/base_rate_liability | 3^2 |
| | 3 | E,B,G | SR | 500,000 | 50 | 10 | NA | 1 | 0.02 | 0 | 1000 | NA | NA | 0/base_rate_liability | 3^2 |
| | 4 | E,B,F,G | SR | 500,000 | 50 | 10 | NA | 1 | 0.02 | 0 | 1000 | NA | NA | 0/base_rate_liability | 3^2 |
| | 5 | E,B,F,G | MR | 500,000 | 50 | 10 | us | diag(k) | k+100 | 0 | diag(k)*1000 | diag(k) | k+100 | 0/base_rate_liability | 3^2 |
